# Normalization for sampled count data

**DOI:** 10.1101/2022.05.06.490859

**Authors:** A. Sina Booeshaghi, Ingileif B. Hallgrímsdóttir, Ángel Gálvez-Merchán, Lior Pachter

## Abstract

Genomics data analysis requires normalization of feature counts that stabilizes technical variance, accounts for variable cell sequencing depth, and preserves monotonicity of within-cell feature abundances. We show that normalization via an optimal variance stabilizing transform for negative binomial count data followed by a proportional fitting step (PFlog) is the only feature-relabeling-equivariant method satisfying the three desiderata. We demonstrate superior performance of this method, which is equivalent to a shifted centered-log ratio transform, in comparison to other normalizations on numerous benchmarks across hundreds of single-cell RNA-seq datasets. We further show that both the shifted-log scale and centered-log ratio geometry are important for preserving PCA and *k*-NN structure.

## 1. Introduction

A central theme in genomics count normalization is the importance of achieving depth normalization alongside stabilization of technical variance (Vallejos et al., 2017; Evans et al., 2017; Robinson and Oshlack, 2010). While variance-stabilizing transformations have been studied for over 90 years (Bartlett, 1936), the question of how to achieve both technical variance stabilization and depth normalization is a matter of ongoing debate (Booeshaghi and Pachter, 2021; Lun et al., 2016; Lun, 2018; Ahlmann-Eltze and Huber, 2023; Cuevas-Diaz Duran et al., 2024; Lu et al., 2025; Huang et al., 2025).

While many methods have been proposed for genomics conut data normalization (Bacher et al., 2017; Cole et al., 2019; Tian et al., 2019; You et al., 2021; Lytal et al., 2020; Borella et al., 2021; Breda et al., 2021; Ge et al., 2025; Park and Hauschild, 2024), the approach of equalizing depth for all cells via a proportional fitting step (PF), often to a “size factor” such as ten thousand (CP10k) or one million (CPM), followed by the application of a technical variance-stabilizing transform such as log plus one (log1p) is most popular and remains common in best-practice workflows (Luecken and Theis, 2019; Heumos et al., 2023). Depth normalization is performed because in most genomic studies cells are not sequenced to saturation. Alternative size-factor strategies, including pooling-based deconvolution, were developed specifically to address sparsity in single-cell count matrices (Lun et al., 2016). The log1pPF workflow is implemented in the widely used Seurat (Hao et al., 2023) and Scanpy (Wolf et al., 2018) programs (Rich et al., 2026), but it does not explicitly model cell depth as a covariate. The sctransform method (Hafemeister and Satija, 2019) aims to address the challenge of technical variance stabilization and depth normalization by transforming data to Pearson residuals derived from a regularized negative binomial regression. This regression-based method incorporates sequencing depth as a covariate in a model, rather than utilizing a size factor (Anders and Huber, 2010). However, Figure 1 of (Hafemeister and Satija, 2019) belies claims that sctransform effectively controls for the effects of variable depth, an observation that has been corroborated in (Crowell et al., 2020); sctransform v2 (Choudhary and Satija, 2022) updates the approach by fixing the coefficient of log sequencing depth in the negative binomial model and regularizing gene-level residual variance.

**Figure 1.**
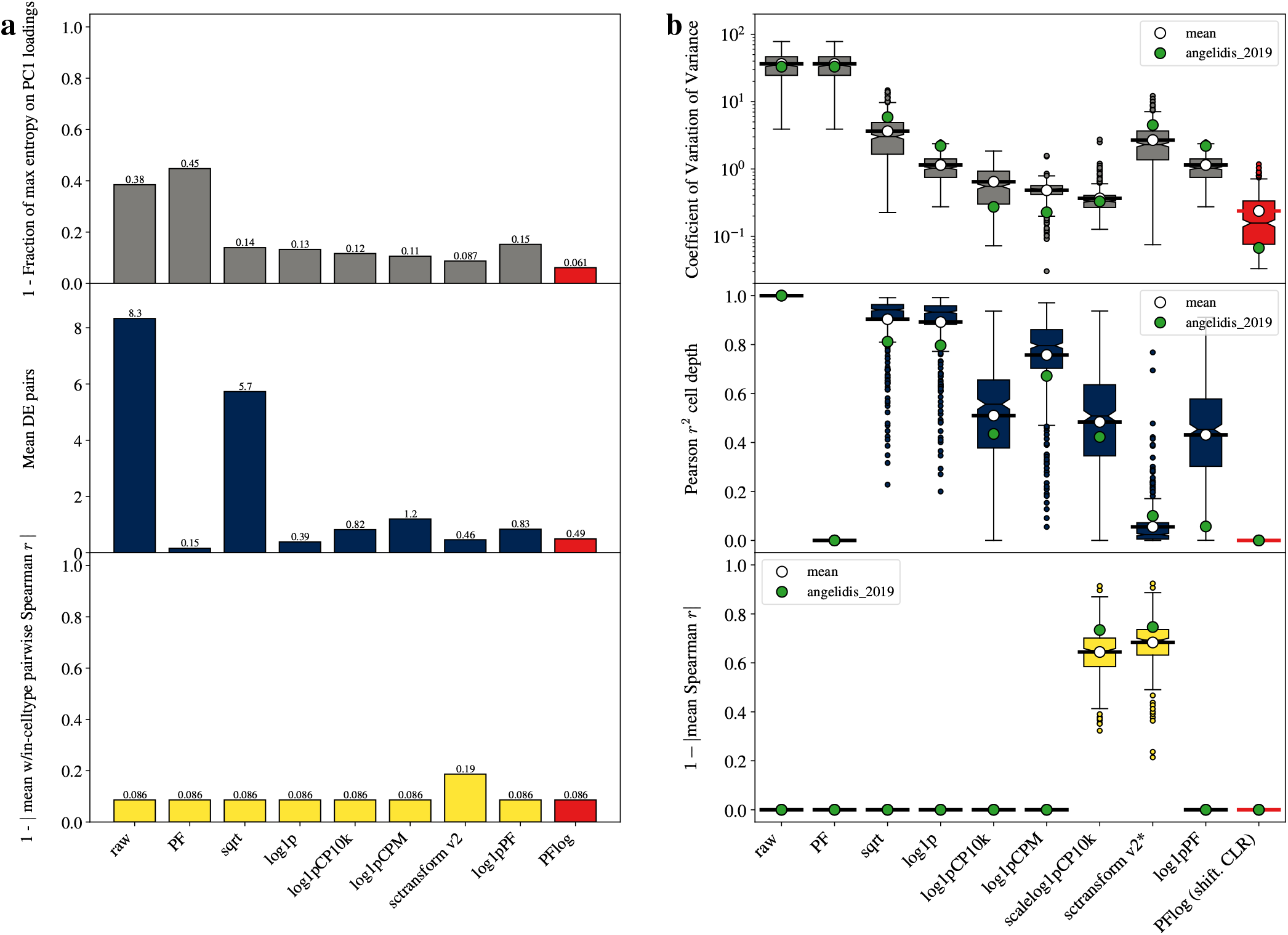
Desired properties for normalization methods. (a) Measurement of the impact of technical variance stabilization, depth normalization, and monotonicity measured via negative controls on the Angelidis2019 dataset (Angelidis et al., 2019) with PFlog shown in red (lower is better in all assessments). The top panel reports the fraction of the max entropy on the PC1 loadings for all genes. The middle panel reports the mean number of significant abundance-matched log-ratio gene pairs across randomized pairings. The bottom panel reports the median within cell type pairwise Spearman correlation. (b) Technical variance stabilization, depth normalization, and monotonicity measured on 526 datasets, with 437 passing quality control (lower is better in all assessments). The Angelidis2019 data results are shown in green. sctransform* is measured by empirical residual-variance CV rather than CV(*q*).

In a comprehensive benchmark (Ahlmann-Eltze and Huber, 2023), 22 normalization methods were compared specifically with respect to their efficacy when coupled to principal component analysis (PCA). That study found that proportional fitting followed by log transformation performed well relative to many alternatives. However, the benchmark also made clear that good relative performance does not translate to complete removal of depth effects. In particular, the depth-associated structure remained visible after normalization, and a subsampling-based nearest-neighbor concordance analysis suggested that even the best performing methods still had substantial room for improvement in absolute terms. Model-based work has also emphasized that UMI count vectors can be treated as multinomial samples conditional on cell depth, so normalization and feature selection directly shape dimension reduction (Townes et al., 2019). Related work highlights how strongly downstream structure depends on the normalization and transformation step itself (Lingen et al., 2024).

The PF step in log1pPF appears to remove depth dependence at the count level, but the form of the transform is just an illusion; in practice, the PF step merely serves to set a pseudocount for the logarithm of the counts. It is therefore natural to ask whether the pseudocount matters in practice (beyond just preventing the pathology of evaluating the logarithm of zero counts), how to set it if it does, and whether a second normalization step after the logarithm could achieve depth equalization while retaining the benefits of technical variance stabilization. Because the logarithm has already been applied, such a second proportional fitting step should be multiplicative in the raw counts rather than additive. We refer to this family of transforms as PFlog. This construction separates two practical issues that are often conflated in log-normalized workflows: choosing the pseudocount and then applying the centering that gives the transformed data its scale-invariant geometry.

## 2. Results

### 2.1. Desired properties

In considering what one would want from a normalization method, we focused on downstream applications and their assumptions. Three properties are especially important: technical variance stabilization, depth normalization, and monotonicity. These can be measured using the coefficient of variation (CV) of transformed technical variances, negative-control tests for residual depth-associated signal, and rank-based correlation metrics that quantify within-cell ordering changes after transformation, respectively. Figure 1a shows the results of measuring these properties on a lung single-cell RNA-seq dataset (Angelidis et al., 2019), which was used in the benchmark of (Ahlmann-Eltze and Huber, 2023).

Technical variance stabilization matters because PCA and related downstream methods assume that high technical variance alone should not cause high-mean genes to dominate principal components (Nguyen and Holmes, 2019). On the Angelidis2019 dataset (Angelidis et al., 2019), methods without explicit technical variance stabilization show low entropy in PC loadings, while transformations such as log1p and sqrt flatten the technical mean-variance trend substantially (Figure 1a, top). This distortion can also alter interpretable signal structure, which is why some workflows now combine log normalization with cell-vector length equalization strategies (Kim et al., 2024).

Depth normalization is equally important. In the absence of depth normalization, transformed matrices can retain sequencing depth bias that inflates false-positive differential signal and perturbs cell-cell distances used for graph construction and clustering. For example, in Type 2 pneumocytes in the Angelidis2019 dataset, depth-biased transforms produce many more depth-associated log-ratio gene-pair differences than transforms that remove depth effects (Figure 1a, middle). In other words, scale invariance is important.

Monotonicity of a transform is important because marker analysis and heatmap interpretation depend on preserving within-cell gene ordering. Non-monotonic transforms alter perceived marker specificity (Figure 1a, bottom; Supplementary Note Fig. 1).

### 2.2. PFlog (shifted CLR)

Figure 1a shows that no existing method uniformly satisfies all three criteria in the Angelidis2019 dataset. Some methods stabilize technical variance but retain depth effects, while others improve depth behavior but don’t preserve rank. This gap motivates studying a transform that jointly targets technical variance stabilization, depth normalization, and monotonicity. In Supplementary Note Section 1, we show that the shifted centered log-ratio is the only count-data transform that is compatible with the desired properties of technical variance stabilization, scale (sequencing depth) invariance, and monotonicity, while satisfying perturbation additivity (so that it is compatible with PCA) and remaining invariant to the order of the input (Supplementary Note Theorem 1.1, Propositions 1.2 and 1.3).

The centered log-ratio transform (CLR) was first proposed in (Aitchison, 1982), and the shifted centered log-ratio transform can be understood as an extra proportional fitting step after a shifted logarithm. Specifically, for a cell *c* with counts 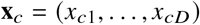, let 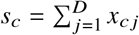 and *u*_*ci*_ = *x*_*ci*_/*s*_*c*_. The PFlog family applies a shifted log and then centers in log space:

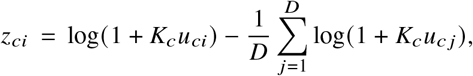

where *K*_*c*_ *>* 0 sets the shifted-log scale for cell *c*. Equivalently, this is the shifted-CLR form with normalized-composition pseudocount 1/*K*_*c*_. On the original count scale,

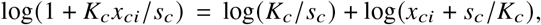

so depth-normalizing before the logarithm has a simple interpretation: it changes the effective raw-count pseudocount to *s*_*c*_ */K*_*c*_. The extra term log (*K*_*c*_ */s*_*c*_) is a scalar shift applied to every gene in the cell, and in PFlog it cancels exactly under the subsequent within-cell centering. Thus the substantive choice made by *K*_*c*_ dictates the count-scale pseudocount. The exact Anscombe-derived count-scale pseudocount is 1/ (4*α*), which corresponds to the cell-specific scale 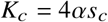 (Anscombe, 1948) (Supplementary Note Sections 1 and 3).

This expression is exactly a centered log-ratio transform on the shifted composition (*u*_*c*1_ + 1 /*K*_*c*_, …, *u*_*cD*_ + 1 /*K*_*c*_) up to an additive constant that cancels under centering (see Supplementary Note Section 2). The shifted CLR transform effectively adds pseudocounts in the right way to make the CLR suitable for count data with zeros: under the Anscombe calibration, the shift is the raw-count operation *x*_*ci*_ → *x*_*ci*_ 1/(4*α)*. The shift can therefore be chosen to optimize technical variance stabilization (Supplementary Note Fig. 2). Moreover, PFlog can be computed efficiently and easily (Supplementary Note Section 5, Supplementary Note Fig. 4).

### 2.3. Benchmarks of 526 datasets

The practice of assessing the performance of normalization methods on only a single dataset such as Angelidis2019, or even a handful of examples, can be misleading, and conclusions may not generalize. For this reason, we performed a benchmark of 526 datasets, of which 437 passed quality control (Fig. 1b, Supp. Fig. 1.1–1.526; Methods). We compared eight normalization methods, including sctransform and widely used PF/log-based workflows from Seurat (Hao et al., 2023), Scanpy (Wolf et al., 2018), Monocle (Trapnell et al., 2014), and scprep (Gigante et al., 2020). These results are consistent with recent multi-cohort evaluations that show method ranking is highly context-dependent (Ge et al., 2025; Cuevas-Diaz Duran et al., 2024; Saxena, 2026).

Across datasets, technical variance stabilization showed high method- and dataset-specific variability, reinforcing that relative performance on one dataset can be misleading. Depth effects also persisted for methods that nominally correct for depth variability; importantly, the log1pPF transform and some sctransform outputs retained substantial depth correlation in many datasets. For monotonicity, transformations based on residuals and scaling showed within-cell rank changes.

Our benchmark also highlighted a software constraint that affects method choice: some transforms operate on dense matrices that exceed memory limits on standard hardware (Supplementary Note Fig. 4). For example, the sctransform matrices are much larger than sparse matrices (Supplementary Note Fig. 4b), creating practical limits for large-scale analysis.

Most importantly, PFlog provided the best overall performance across criteria in this large benchmark. The additional proportional fitting step after log completely eliminated the residual depth structure while fully preserving monotonicity and exhibiting strong technical variance stabilization, equal to that of log1pPF. While scaling can provide perfect empirical gene-variance equalization, it comes at a cost of scrambling gene order within cells and not removing depth dependence. The sctransform procedure rescues depth but scrambles within-cell gene order while operating on a dense matrix. These results are corroborated by adding a PFlog row to the previous Ahlmann-Eltze–Huber benchmark, where the added transform performs favorably on the retained simulation and downsampling comparisons even under conservative holdout scoring (Supplementary Note Fig. 6).

### 2.4. Implications for cell geometry

Normalization determines the Euclidean geometry seen by PCA and by the *k*-NN graphs constructed from PC coordinates. Two choices are especially important: the pseudocount, which controls technical variance stabilization, and the CLR centering step, which removes cell-wise scale. If either is poorly chosen, PCA can distort the axes, loadings, and local neighborhoods used downstream.

To understand the impact of these choices, we first analyzed mouse-level pseudobulk counts from the Angelidis2019 lung dataset (Angelidis et al., 2019). This example isolates both parts of the PFlog construction: the choice of count-scale pseudocount and the subsequent CLR centering step. In a PCA of young and old samples, log1p(CP10K), the default normalization scale used by Seurat and Scanpy, and PFlog both separate the age groups visually (Fig. 2a,b). However, CP10K fixes the shifted-log scale at *K* = 10,000, whereas the Anscombe-calibrated PFlog estimates *α*, uses the cell-specific scale 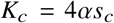, and then applies CLR centering. Equivalently, PFlog uses the count-scale pseudocount 1 /(4*α)*, while CP10K fixes a different pseudocount *s*_*c*_ */*10,000. The pseudobulk sample depths vary by more than two-fold (Fig. 2e); expressed at a representative depth *s*_*_, the estimated scale is *K*_*_ = 4*αs*_*_ *≈* 1.3 × 10^6^, so the default is about two orders of magnitude off from the optimal representative-depth value.

**Figure 2.**
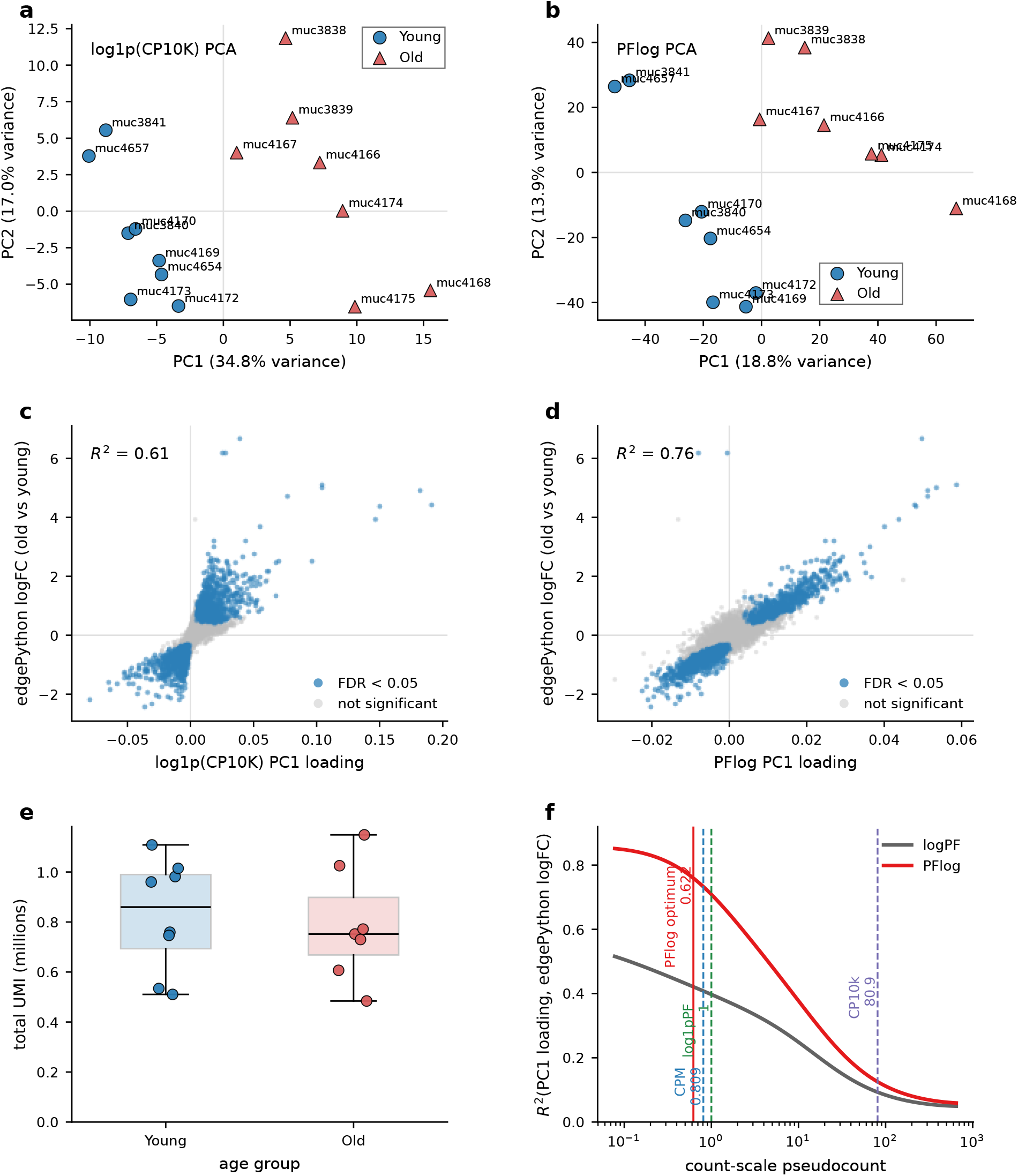
PFlog, which uses an estimated shifted-log scale and CLR centering, improves the correspondence between unsupervised PCA loadings and supervised old-vs-young differential expression in Angelidis2019 lung pseudobulk samples. **(a)** PCA after log1p(CP10K) normalization. **(b)** PCA after PFlog normalization. **(c)** log1p(CP10K) PC1 gene loadings compared with edgePython old-vs-young log fold changes. **(d)** PFlog PC1 gene loadings compared with the same supervised log fold changes. Blue points indicate genes with edgePython FDR < 0.05. **(e)** Total pseudobulk UMI depth for each mouse sample. **(f)** Effect of the pseudocount and centering on the agreement between PC1 loadings and supervised log fold changes.

The consequence is visible in the PC1 loadings. PC1 explains much more variance after log1p(CP10K) than after PFlog (34.8% versus 18.8%), but this additional variance is not necessarily biological. The largest log1p(CP10K) PC1 loadings include highly expressed immune-abundance genes such as *Igkc, Ighm, Igj, Igha, Ccl5*, and *Lyz1*, which are not the old-vs-young expression program. After PFlog, the unsupervised PC1 loadings align much more closely with supervised old-vs-young differential expression: the coefficient of determination between PC1 loading and supervised log fold change increases from *R*^2^ = 0.61 for log1p(CP10K) to *R*^2^ = 0.76 for PFlog (Fig. 2c,d). Sweeping the count-scale pseudocount separates the effect of the pseudocount and centering (Fig. 2f). First, along either curve, the agreement is highest near the estimated Anscombe pseudocount and deteriorates at the much larger CP10K pseudocount, showing that the shifted-log scale is consequential. Second, at the same pseudocount, the CLR-centered PFlog curve lies above the uncentered shifted-log curve, showing that centering is not just a cosmetic rescaling but improves the biological alignment of the leading loading vector. Thus, both choosing the pseudocount correctly and applying CLR centering can change PC1 from a high-abundance axis into an interpretable biological contrast. This also shows that PFlog is useful beyond single-cell RNA-seq, because the example is based on pseudobulk RNA-seq profiles.

Second, we tested whether the same PC loading geometry is stable across large depth differences in single-cell data. We compared lung cells from the integrated Human Lung Cell Atlas (HLCA)(Sikkema et al., 2023) measured with Seq-Well and 10x 3’ v3, whose mean depths differ by approximately 12-fold. With log1p(CP10K), the sign-aligned PC1 loading *R*^2^ between the two assays is only 0.06 (Fig. 3a). Estimating the pseudocount while omitting CLR centering improves this value to 0.15 (Fig. 3b), showing that the shifted-log scale matters. Applying the full PFlog transform raises *R*^2^ to 0.57 (Fig. 3c), showing that CLR centering matters as well for data integration. The HLCA analysis did not use default log1p(CP10K) normalization for integration, but instead used scANVI (Xu et al., 2021; Sikkema et al., 2023); this approach to data integration does not by itself solve this geometry problem, since in a muscat 12-fold downsampling control the downsampled cells recovered only 17.2% of their full-depth *k* = 10 neighbors in the scANVI latent space, compared with 31.0% after PFlog PCA.

**Figure 3.**
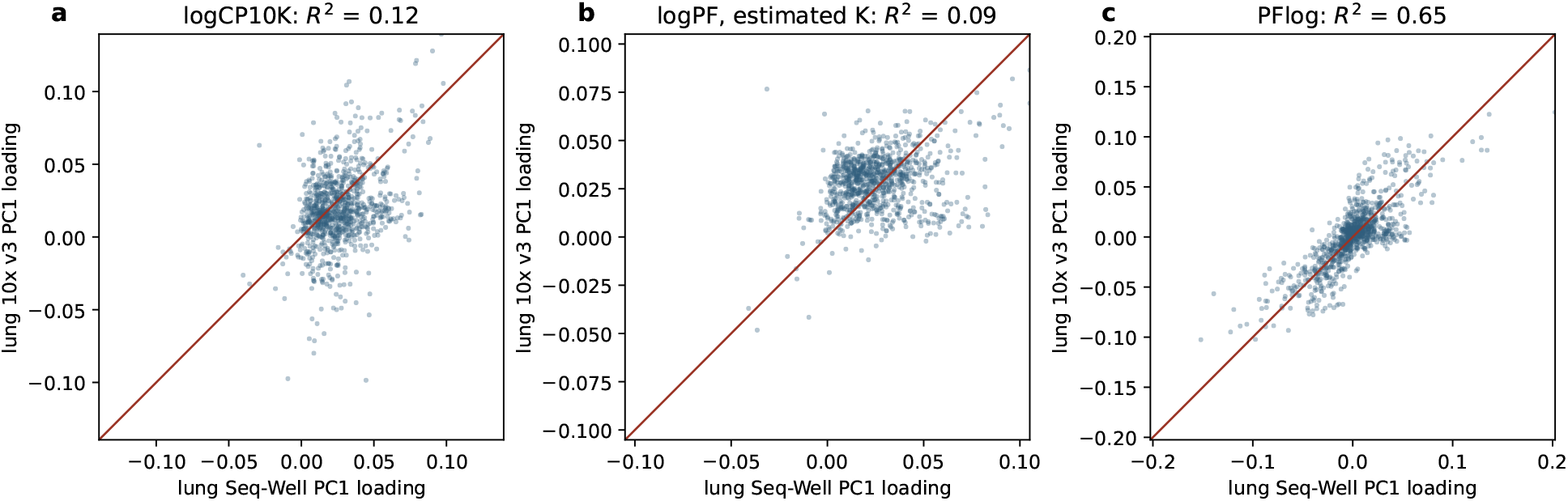
PFlog stabilizes PC1 loading geometry across single-cell lung datasets with different sequencing depths. PC1 loadings were compared between Seq-Well and 10x 3’ v3 lung cells from the integrated Human Lung Cell Atlas, using the same selected genes and matched cell-type composition. **(a)** log1p(CP10K) gives weak loading agreement across assays. **(b)** logPF with estimated *K*, but without CLR centering, improves agreement. **(c)** PFlog, with estimated *K* and CLR centering, gives substantially stronger agreement.

The effect also propagates from PC loadings to local neighborhoods and clustering. In the 10x Genomics mouse brain `neuron_1k_v3` dataset, using the CP10K pseudocount recovered only 38.0 of 50 PFlog neighbors from the graph obtained at the estimated Anscombe pseudocount, while log1pPF recovered 37.7 (Supplementary Note Fig. 3). With 50 PCs and Leiden clustering using *n*_neighbors_ = 30, PFlog produced 13 clusters whereas CP10K produced 15. One CP10K cluster appeared to over-split a PFlog low-depth vascular/immune cluster. It showed no significant targeted marker differences relative to the full PFlog cluster and primarily separated cells by depth. Thus, the effect of the count-scale pseudocount is not confined to impacting PC loadings; it can also change the neighborhood graph and induce depth-associated over-splitting of marker-coherent cells (Supplementary Note Section 1).

Together, Figs. 2 and 3 show that PFlog improves cell and sample geometry for two complementary reasons. A data-calibrated pseudocount appropriately balances the influence of low- and high-count features by setting the transition between the linear and logarithmic regimes of the shifted log, while CLR centering removes residual depth effects so that PCA compares compositions rather than library sizes. Without both steps, PCA axes and the *k*-NN graphs derived from them can be distorted even when low-dimensional visualizations appear superficially reasonable.

## 3. Discussion

Count normalization is a crucial first step in all scRNA-seq analysis that, in principle, comprises a single step in a standard workflow. However, in practice normalization is a collection of techniques, data representations, analysis types, and visualizations that interact with each other in non-obvious and frequently undocumented ways. In Seurat and Scanpy, the analysis software used for the majority of scRNA-seq analysis, some normalization implementations can also limit users by requiring large amounts of memory. Thus, while users frequently think of normalization as a single data-transformation step in analyses, it is often not; the software engineering choices made by developers of the tools used can affect analyses in unpredictable, and sometimes unintended, ways.

A concrete example is Seurat’s function named “CLR”, which although labeled as the centered log-ratio transform, does not function as such (Scanpy does not implement CLR normalization at all). The relevant Seurat normalization step appears early in the workflow (Supplementary Note Fig. 8). As shown in Supplementary Note Section 4, the implementation path computes a zero-sensitive shifted log heuristic based on a geometric-mean-like factor from the positive entries, excludes zeros from that factor, and does not enforce CLR centering (Σ_*i*_ *z*_*i*_ = 0). This is a categorical mismatch with CLR, not a small stylistic difference.

This mismatch is consequential. The transform becomes dependent on zero patterns, and therefore depth-sensitive in exactly the regime tested by downsampling (Supplementary Note Section 4; Supplementary Note Fig. 5). In practical terms, selecting “CLR” in Seurat does not execute CLR in the Aitchison sense; it is a different transform no longer satisfying desirable properties, and benchmark results are consistent with that behavior (Borella et al., 2021).

Despite the complexity of normalization in practice, much work on scRNA-seq has focused on statistical details that, while important, are not necessarily the primary determinants of results. For example, the debate over whether gene-specific over-dispersion parameters should be used when computing Pearson residuals (Hafemeister and Satija, 2019; Lause et al., 2021; Hafemeister and Satija, 2020; Choudhary and Satija, 2022) ignores the fact that Pearson residuals are not the result of a monotonic transformation and are typically stored as dense matrices. Such dense or memory-intensive representations can lead to significant analysis limitations in large scRNA-seq workflows (Borella et al., 2021), and can slow downstream computations enough to make million- or tens-of-millions-cell analyses problematic or intractable (Dicks et al., 2026). These problems have significant implications for common tasks, such as finding marker genes, as discussed above.

The shifted-log scale *K* is also not a minor implementation detail. Figure 2 shows a case where the Seurat and Scanpy default *K* = 10,000 is off by about two orders of magnitude: the Angelidis2019 pseudobulk analysis gives *K*_*_ ≈ 1.3 × 10^6^ at representative depth. Since a fixed *K* determines the cell-specific count-scale pseudocount *s*_*c*_ */K*, this is equivalently an error in the amount added to each count before the logarithm. This is not an isolated example, since Fig. 1b shows that across many datasets a scale near one million is usually better matched to the data than ten thousand. If a single fixed default had to be chosen, one million would therefore be more defensible than ten thousand, but the better solution is to estimate overdispersion and use the corresponding pseudocount 1 /(4*α)*. The practical consequence is not limited to the value of a technical-variance diagnostic: changing the pseudocount can change PC loadings, the *k*-NN graph built from the PCs, and ultimately Leiden clusters (Fig. 2f and Supplementary Note Fig. 3). These finding show the while it may be true that for (Ahlmann-Eltze and Huber, 2023) “benchmarks showed limited performance benefits for [an adaptive pseudocount”, this does not mean that it is not important to set the pseudocount optimally according to the theory of (Anscombe, 1948); unfortunately, the benchmarks of (Ahlmann-Eltze and Huber, 2023) were inadequate.

While locally, within one dataset, the pseudocount can dominate the effect of the subsequent CLR centering; globally, when combining many datasets sequenced to different depths, the scale-invariance provided by CLR centering becomes increasingly important. Figure 3 illustrates this second point directly: estimating the pseudocount improves cross-depth PC loading agreement, but the full shifted CLR is needed to recover stable loading geometry. The same lesson applies to learned integration methods. Models such as scANVI can be useful for annotation and integration, but they do not remove the need to understand how normalization and latent geometry affect *k*-NN graphs. When the same muscat cells were downsampled 12-fold, PFlog PCA preserved substantially more of the full-depth local neighborhood at *k* = 10 than the scANVI latent space. Thus, model-based integration should be evaluated against the geometry required by the downstream task, rather than treated as a substitute for normalization.

PFlog also illustrates that the mathematical form of a normalization and its data representation should be considered together. Naively applying the final CLR-centering step materializes a dense matrix, because structural zeros are shifted by a cell-specific centering constant. This apparent obstacle can be avoided by storing the sparse log-transformed matrix together with its centering vector, and by carrying out downstream linear algebra through the corresponding sparse-plus-low-rank operator (Supplementary Note Section 5). We provide efficient implementations for PFlog via scclr `(Python)` and `scclrR` (R), making the normalization practical for large single-cell matrices without requiring dense storage.

Although this paper focuses on single-cell RNA-seq, the same normalization issues arise in other genomics count assays. Figure 2 is based on mouse-level pseudobulk RNA-seq profiles rather than individual cells, and it shows that PFlog can improve the interpretation of sample-level PCA by reducing the influence of high-count baseline effects and aligning the leading unsupervised axis with supervised differential expression. This shows that PFlog may be useful for bulk RNA-seq and related count-based genomics assays where library depth and highly expressed features can dominate geometry.

On a cautionary note, the goal of eliminating residual depth after transformation presumes that all depth variation is technical, which is known not to be the case (Lu et al., 2025; Huang et al., 2025; Gorin, Vastola, et al., 2023; Gorin, Chari, et al., 2025). Normalization should ideally separate biological from technical effects, and doing so requires modeling transcriptional dynamics to be able to evaluate the contribution of technical noise to count data (Gorin, Vastola, et al., 2023; Gorin, Chari, et al., 2025).

Regardless of the normalization transformation that is applied, our work shows that assessment of data quality and normalization effectiveness is crucial in practice. No normalization will rescue poor quality data. Measures such as overdispersion, coefficient of variation of transformed technical variances, and coefficient of determination between raw and transformed cell depth ought to be collected as part of standard quality control of experiments. It is also crucial that practitioners understand the assumptions implicit in the normalizations applied, and the implications for interpretation of results, such as whether variation is technical or biological. Such assessments and considerations are now essential as multiple datasets, frequently normalized via different approaches, are used for training foundation models. Due to the absence of these critical assessments, dozens of normalization methods that perform poorly in absolute terms (Ahlmann-Eltze and Huber, 2023) have been used in hundreds of thousands of studies, despite the existence of an optimal approach (Supplementary Note Section 1) developed more than 40 years ago (Aitchison, 1982).

## Data and code availability

All data and code to reproduce the figures and results in the paper are available at https://github.com/pachterlab/BHGP_2022. Code for performing sparse PFlog (shifted CLR) normalization is available at https://github.com/cleartools/scclr.

## Acknowledgments

This project started with an investigation of normalization of orthogonal barcoding tags. We thank John Thompson and Linda Hsieh-Wilson for helpful initial discussions related to that problem. We thank Tara Chari for helpful insights on normalization and clustering. Thanks to Meichen Fang and Gennady Gorin for discussions on the role of normalization in separating technical noise from biological signal. Lambda Moses and Joseph Rich helped review the Seurat source code. Code was written with the assistance of GPT 5.5 and Claude 4.8.

## Author contributions

A.S.B. and L.P. developed the project idea. A.G.M. pre-processed the datasets. A.S.B. and L.P. performed the analysis. I.B.H. performed an overdispersion simulation. A.S.B. and A.G.M. compiled the 526-dataset supplement. A.S.B. drafted an initial version of this paper. A.S.B., I.B.H., A.G.M., and L.P. wrote, reviewed, and edited the paper.

## Competing interests

The authors declare no competing interests.

## 4. Methods

### 4.1. Preprocessing

The raw matrices were filtered by removing cells below a selected knee-plot threshold (see the dataset folders). Datasets for which the average count per cell was less than 818.46 (the average count per cell in (Angelidis et al., 2019)) were not used in Fig. 1.

### 4.2. Collecting metadata

Dataset metadata was collected with the ffq program version 0.2.1 (Gálvez-Merchán et al., 2023) by running `ffq -l 2 -o DATASETID_metadata.json DATASETID`. Eighteen out of the 526 datasets processed did not have metadata associated with their dataset ID.

### 4.3. Normalizing matrices

We applied seven non-sctransform normalization methods to each cell-filtered matrix: PF, sqrt, log1p, log1pCP10k, log1pPF, log1pCPM, and PFlog. The PF, sqrt, log1p, log1pCP10k, log1pPF, and log1pCPM transformations were computed by running the `norm_sparse.sh` script. The reported PFlog transformation is the shifted CLR transform defined in the main text and Supplementary Note: for each analyzed matrix, the overdispersion *α* was estimated, the count-scale pseudocount was set to 1 /(4*α)*, equivalently the cell-specific normalized shifted-log scale was *K*_*c*_ = 4*αs*_*c*_, counts were divided by each cell’s depth to obtain *u*_*ci*_, log (1 + *K*_*c*_*u*_*ci*_ ) was computed for each gene, and the within-cell mean of those log values was subtracted. Together with sctransform, this gives the eight normalization in Fig. 1b. We then ran `norm_sctransform.sh` on the original cell-filtered matrix to generate the sc-transform matrix. The sctransform function was called with `var_features_n=number_of_genes_in_dataset, vst_flavor=“v2”,` and default parameters. In order to perform a uniform comparison with sctransform, we additionally filtered the original cell-filtered matrix to the set of genes returned by sctransform, since sctransform has a built-in gene filtering step. Finally, we repeated these normalizations on the sctransform-selected gene subset, and ran `norm_cp10k_log_scale.sh.`

### 4.4. Angelidis pseudobulk PCA

Figure 2 was generated from Angelidis2019 lung counts aggregated to mouse-level pseudobulk profiles by summing raw counts over cells from each mouse sample. The pseudobulk matrix contained 15 samples, 21,969 genes, and 14,813 cells in aggregate, with 8 young (3 month) and 7 old (24 month) samples. We compared log1p(CP10K) with PFlog. For log1p(CP10K), each pseudobulk count vector was divided by its total count, multiplied by 10,000, and transformed with log (1 + *x)*. For PFlog, we used `scclr` with `target=“auto”` to estimate *α*, use the corresponding cell-specific scale *K*_*c*_ = 4*αs*_*c*_, apply the shifted log, and perform CLR centering. PCA was run with five components; PC1 was oriented so that old samples had larger mean PC1 scores than young samples. Sample depths in panel (e) are total pseudobulk UMI counts. For panel (f), we swept count-scale pseudocounts *y*_0_. For each *y*_0_, raw pseudobulk counts were transformed either by the shifted log alone, log (*y + y*_0_), or by shifted log followed by CLR centering, equivalently by running `scclr` with *α* = 1 /(4*y*_0_). The plotted value is the coefficient of determination between unsupervised PC1 gene loadings and supervised edgePython old-vs-young log fold changes.

Old-vs-young differential expression was computed from the raw pseudobulk counts with edgePython. Counts were supplied as genes by samples, the age indicator was used as the group variable, genes were filtered with `filter_by_expr,` normalization factors were computed, and a quasi-likelihood negative-binomial GLM was fit with an intercept and age coefficient. PC1 gene loadings from each normalization were merged with edgePython log fold changes by gene, and the reported *R*^2^ values were squared Pearson correlations between PC1 loadings and old-vs-young log fold changes over genes retained in the differential-expression analysis.

### 4.5. HLCA cross-depth PC loading analysis

Figure 3 was generated from a raw-count subset of the integrated Human Lung Cell Atlas containing Seq-Well and 10x 3’ v3 lung cells. The subset contained 560 cells from each assay, balanced across eight cell types with 70 cells per assay and cell type. The mean raw depths were 726 counts per cell for Seq-Well and 8,509 counts per cell for 10x 3’ v3. Genes were retained if detected in more than 1% of cells in both assay groups; among those genes, the 3,000 genes with largest summed mean abundance across the two groups were used.

For each assay group separately, we computed PC1 gene loadings after three transformations: log1p(CP10K), logPF with estimated *K* but without CLR centering, and PFlog. The log1p(CP10K) transform divided each cell by its depth, multiplied by 10,000, and applied log (1 + *x)*. For logPF with estimated *K, K* was obtained from the corresponding PFlog fit for that assay group and then used in log (1 + *Kx /s*) without the final centering step. PFlog was computed with `scclr.normalize_pca` using `target=“auto”` and one principal component. PCA was run independently within the Seq-Well and 10x 3’ v3 groups. For each method, the 10x 3’ v3 PC1 loading vector was sign-aligned to the Seq-Well loading vector, and the plotted *R*^2^ is the squared Pearson correlation between the two matched loading vectors.

For the scANVI downsampling comparison quoted in Section 2.4, we used the stored muscat seed 1 simulated count matrix, created a paired low-depth copy by binomially down-sampling each count with probability 1 12, and embedded the combined full-depth and downsampled matrices with PFlog PCA or scVI followed by scANVI. For each original cell, the reference neighborhood was its *k* = 10 nearest neighbors among full-depth cells; the recovery score was the fraction of those neighbors recovered by the corresponding downsampled cell when queried against the full-depth cells, excluding the exact full-depth twin.

### 4.6. Adapted Ahlmann-Eltze–Huber benchmark

The adapted Ahlmann-Eltze–Huber benchmark figure in the Supplementary Note was generated from the checked-in benchmark tables of Ahlmann-Eltze and Huber (2023), with an added row labeled CLR corresponding to PFlog/shifted CLR recomputed with `scclr` using `target=“auto”`. This CLR row was not present in the original benchmark. The consistency panel used the same gene-subset *k*-NN overlap metric and parameter choices as the original benchmark. For the simulation panel, the original benchmark considered muscat, dyngen, linear walk, random walk, and scDesign2. The adapted figure retains muscat, dyngen, and scDesign2, but excludes the linear-walk and random-walk simulations because those frameworks multiply the simulated expression state by real cell-specific depths; depth is therefore part of the benchmark ground-truth geometry, leading to an unrealistic simulation (Supplementary Note Fig. 7). For the retained simulations, PFlog was scored by comparing *k* = 50 neighborhoods after PFlog PCA to the simulator ground-truth *k* = 50 neighborhoods, using the same simulator-specific PCA dimensions and seeds as the plotted original methods.

For the downsampling panel, we followed the consensus-neighborhood structure of Ahlmann-Eltze and Huber (2023) rather than using PFlog full-depth-vs-downsampled self-overlap. The full-depth target neighborhood for each cell was defined as the intersection of the full-depth *k* = 50 neighborhoods from the log-family methods highlighted by Ahlmann-Eltze and Huber (2023): log (*y/ s* +1), log(*y/ s* +1) ⟶ HVG, log (*y /s* + 1) ⟶Z, log (*y/ s* +1) ⟶HVG ⟶Z, log (*y/ s* + 1)/ *u*, and logCPM. PFlog was omitted from the construction of this consensus, so that the addition of PFlog did not change the target against which it was scored. PFlog neighborhoods were computed on the downsampled matrices with scclr and then scored against this fixed log-method consensus by averaging, over cells passing the same consensus-size filter as in the original downsampling code, the number of PFlog downsampled neighbors contained in the full-depth consensus. This is a holdout comparison against the methods that Ahlmann-Eltze–Huber recommended as strong log-based baselines, and it is conservative in the sense that PFlog cannot contribute to the consensus target.

For the walk-simulation diagnostic in Supplementary Note Fig. 7, we regenerated or loaded the cached linear-walk and random-walk count matrices and simulator ground-truth *k* = 50 neighborhoods for seeds 1–5 from the Ahlmann-Eltze and Huber (2023) benchmark. For each seed, total raw depth was computed per cell. We plotted the raw depth across simulator cell order for seed 1 and compared the mean absolute depth difference between each cell and its ground-truth neighbors with the corresponding quantity for random non-self neighbors sampled with the same *k*. Ratios below one indicate that the simulator ground-truth neighborhood is itself depth-similar.

### 4.7. Mouse brain pseudocount and clustering analysis

The mouse-brain analysis in Supplementary Note Fig. 3used the 10x Genomics `neuron_1k_v3` filtered feature-barcode matrix. The analysis retained 1,301 cells and the 3,000 genes with the largest total counts. The estimated overdispersion was *α* = 4.57, giving the Anscombe count-scale pseudocount 1 /(4*α)*= 0.0547; at the filtered mean depth of 9,348 counts, the CP10K and log1pPF count-scale pseudocounts were 0.935 and 1.0, respectively. For the pseudocount sweep, counts were transformed by the centered shifted log over a grid of pseudocounts, PCA was run, a *k* = 50 nearest-neighbor graph was constructed, and neighbor overlap was measured against the graph at the estimated Anscombe pseudocount. For the clustering comparison, PFlog and CP10K embeddings were computed with 50 PCs and clustered with Leiden using *n*_neighbors_ = 30; cluster concordance was summarized with adjusted Rand index and normalized mutual information. The putative CP10K over-split was assessed by comparing its cells with the matched PFlog cluster using raw depth distributions and a targeted panel of canonical brain cell-type marker genes, with multiple-testing adjustment across the targeted tests.

### 4.8. Seurat “CLR” analysis

For the Seurat workflow diagram in Supplementary Note Fig. 8, we inspected the Seurat source code path through `NormalizeData` and summarized where normalization occurs in the standard workflow. For the Seurat “CLR” downsampling experiment in Supplementary Note Fig. 5, we used the Smart-seq3 fibroblast UMI matrix from the Ahlmann-Eltze and Huber (2023) downsampling benchmark after removing zero-count genes and cells. For each of 20 seeds, each full-depth cell was paired with a multinomially downsampled twin at 10% of its original depth. The joint full-depth/downsampled matrix was transformed with raw counts, Seurat “CLR” with `margin=2`, log1pPF, the Ahlmann-Eltze–Huber shifted log log(*y /s* + 1 /(4*α*)), or PFlog. PCA was run with 10 components. We computed a per-cell twin-recovery score from cross-depth distances between full-depth cells and downsampled twins, and computed the coefficient of determination from regressing PC1 on the full-depth/downsampled group label.

### 4.9. Dataset-level benchmark metrics

The pseudocount simulation in Supplementary Note Fig. 2used eight overdispersion/depth scenarios chosen from representative datasets. For each scenario, 20 replicates of 2,000 genes and 500 cells were simulated by drawing gene weights on a log scale, normalizing them to gene proportions, and drawing cell depths from a log-normal distribution around the scenario mean depth. For each value of *K*, we evaluated the delta-method technical variance *q*_*gc*_ (*K*), averaged it over cells to obtain gene-level *q*_*g*_ (*K)*, and plotted the coefficient of variation of *q*_*g*_ *(K)* over genes. The grid included log-spaced *K* values, CP10K, CPM, and the representative-depth Anscombe value 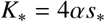, corresponding to the cell-specific rule 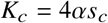 at *s*_*c*_ = *s*_*_.

For Fig. 1b we computed three dataset-level metrics: the coefficient of variation of the model-based transformed technical variance *q*_*g*_, the coefficient of determination between transformed cell depth and raw cell depth, and the average Spearman *r* between the transformed and raw within-cell gene ranks. For the technical variance-stabilization panel, let *Y*_*g*_ denote a count for gene *g* with mean *μ*_*g*_, and model the technical count variance as

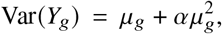

where *α* is the empirical overdispersion estimated for the matrix. For a scalar transformation *h*, the delta method gives

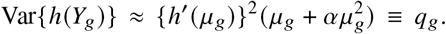

For shifted-log normalizations with representative depth *s*\* and scale *K*,

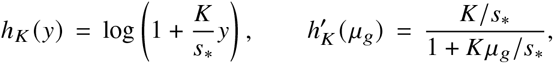

so the quantity used in the top panel was

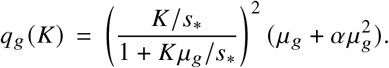

Equivalently, *h*_*K*_ (*y*) = *log*(*y*+*s*_*_/*K*) up to an additive constant, so the parameter being varied is the representative count-scale pseudocount *s*_*_ /*K*. The Anscombe-derived pseudocount is 1/(4*α)*, corresponding exactly to 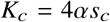 cell by cell, or to 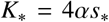 when plotted at representative depth *s*_*_. The reported score was the coefficient of variation across retained genes,

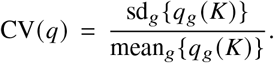

For PFlog in this representative-depth diagnostic, the plotted scale was 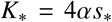 for log1pCP10k and log1pCPM, *K* was fixed at 10,000 and 10^6^, respectively. Because sc-transform returns regularized Pearson residuals from a fitted negative-binomial regression rather than applying a shared scalar function *h*, it does not have an analytic *q*_*g*_ *(K)* comparable to the fixed transforms. For the sctransform (labeled sctransform* in Fig. 1b), we therefore computed an empirical residual-variance analogue: if *R*_*cg*_ is the sctransform residual for cell *c* and gene *g*, we computed

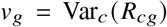

and reported

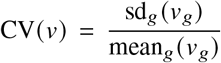

across the genes returned by sctransform. This is not an analytic CV *q* ; the model-based Pearson residual variance is standardized by construction under the fitted model, whereas the realized residual matrix can have nonzero empirical variance heterogeneity because of regularization, clipping, gene filtering, finite-sample fitting variability, and model misspecification. The coefficient of determination was computed by summing each transformed cell vector, regressing transformed cell depth on raw cell depth with `sklearn.linear_model.LinearRegression().fit()`, and calling `score().` The average Spearman *r* was computed by applying `stats.spearmanr` to each transformed-raw cell-vector pair and then taking the mean across cells.

### 4.10. Angelidis cell-type metrics

Cell-type metrics were computed on Type 2 pneumocytes in the Angelidis2019 dataset (Angelidis et al., 2019). For each normalization method, `sklearn.decomposition.PCA()` was run with `n_components=1 and svd_solver=“full”` and the absolute value of the loadings was *l*_1_-normalized. The entropy was computed with `scipy.stats.entropy()` and the max entropy was computed with `np.log(ngenes).` Additionally, the coefficient of determination was computed by regressing PC1, derived from PCA on the normalized matrix, on raw cell depth. To compute the depth negative-control metric in Fig. 1a, we restricted to 3-month-old Type 2 pneumocytes and, within each of eight mice, selected the 50 cells with the largest raw depth and the immediately next 50 cells by raw depth. This produced two 400-cell depth strata with identical mouse composition. Genes detected in more than 25% of cells in both strata were sorted by mean raw count and paired within 12 abundance bins to form 200 randomized sets of abundance-matched gene pairs. For each normalization method, including sctransform v2 residuals, and each pairing, we tested log-ratio balances 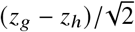 for a depth-stratum effect after subtracting mouse means. The plotted value is the mean number of balances with Bonferroni-corrected *p <* 0.01 across randomized pairings. To compute the average pairwise-Spearman gene-rank correlation, we first found the smallest non-zero difference in counts between entries in each normalization matrix. We added a random number between zero and one-fourth of this minimum to each gene vector to break ties. After adjusting the matrix counts, pairwise-Spearman correlations were calculated on all cells and the average was computed. To compute the association between pairwise differences in cell depth and pairwise *l*_1_ distances between transformed expression profiles, for each matrix we subsampled to 1,000 cells. We computed all pairwise absolute differences in cell depth `withsklearn.metrics.pairwise_distances` with metric=“l1” on the cell sums. We then computed pairwise *l*_1_ distances between the corresponding transformed gene vectors. Finally, `sklearn.linear_model.LinearRegression().fit()` and `score()` were used to compute the coefficient of determination. The monotonicity heatmap in Supplementary Note Fig. 1 was generated from the Angelidis2019 cell-filtered matrix and cell-type metadata. Cells without cell-type labels were removed. For each normalization shown, gene expression was averaged within each cell type. For each cell type, the 100 highest-mean genes in the raw counts were selected as the reference set; the corresponding genes were ranked within that cell type for raw counts, log1pPF, PFlog, and sctransform v2 using ordinal ranks, and the rank matrices were plotted as heatmaps.

### 4.11. Matrix size and summary metrics

The following matrix-level metrics were computed for each matrix, on both all genes and those subset by sctransform: the number of cells (ncells), the number of genes (ngenes), the number of non-zero entries in the matrix (nvals), the fraction of non-zero entries (density), the average depth per cell (avg_per_cell), the average depth per gene (avg_per_gene), the minimum depth per cell (min_cell), the maximum depth per cell (max_cell), the total number of counts in the matrix (total_count), and the empirical overdispersion (overdispersion). These metrics were computed with `metrics_matrix.sh.` Supplementary Note Fig. 4combines two memory/runtime summaries. Panel (a) reproduces the published sctransform v2 runtime curve from Choudhary and Satija (2022). Panel (b) was generated from benchmark size tables recording, for each dataset and normalization, the number of cells and in-memory matrix size in bytes; sparse raw-matrix sizes and dense sctransform-matrix sizes were converted to gigabytes and plotted against cell number.

### 4.12. Creating multi-panel normalization figures

For each dataset and normalization, the following plots were made: 1. a scatterplot of the transformed gene variance vs raw gene mean, 2. a scatterplot of the transformed cell depth vs raw cell depth, and 3. a histogram of the distribution of transformed-to-raw cell Spearman rank correlations. A minmax procedure was performed to scale the *x* and *y* axes of plot 2 where the minimum cell depth was subtracted from each cell and the result was divided by the maximum cell depth. With the exception of sctransform (which chooses a subset of genes), the figures were made with all genes.

### 4.13. Lean formalization

The proofs in the Supplementary Note were formalized with Lean (Moura and Ullrich, 2021), the span tool for relating the Lean declarations to the LATEX (Pachter, 2026), and the assistance of GPT 5.5.

## Notes

https://github.com/pachterlab/BHGP_2022

### Competing Interest Statement

The authors have declared no competing interest.

### Summary of Updates

In this updated version we expanded the benchmarking to address the impact that both the pseudocount and CLR centering has on various metrics, including the corrected AEH benchmarks. We've performed extensive tests to understand the impact that normalization has on single-cell geometry (via PCA concordance with differential expression tests) and cross-study concordance of different normalization strategies.

## References

Ahlmann-Eltze, C. and W. Huber (2023). “Comparison of transformations for single-cell RNA-seq data”. In: Nature Methods 20.5, pp. 665–672. doi: 10.1038/s41592-023-01814-1.

Aitchison, J. (Jan. 1982). “The statistical analysis of compositional data”. In: Journal of the Royal Statistical Society Series B: Statistical Methodology 44.2, pp. 139–160. doi: 10.1111/j.2517-6161.1982.tb01195.x.

Anders, S. and W. Huber (2010). “Differential expression analysis for sequence count data”. In: Genome Biology 11.10. doi: 10.1186/gb-2010-11-10-r106.

Angelidis, I., L. M. Simon, I. E. Fernandez, M. Strunz, C. H. Mayr, F. R. Greiffo, G. Tsitsiridis, M. Ansari, E. Graf, T.-M. Strom, et al. (2019). “An atlas of the aging lung mapped by single cell transcriptomics and deep tissue proteomics”. In: Nature Communications 10.1, p. 963. doi: 10.1038/s41467-019-08831-9.

Anscombe, F. J. (1948). “The Transformation of Poisson, Binomial and Negative-Binomial Data”. In: Biometrika 35.3/4, pp. 246–254.

Bacher, R., L.-F. Chu, N. Leng, A. P. Gasch, J. A. Thomson, R. M. Stewart, M. Newton, and C. Kendziorski (2017). “SCnorm: robust normalization of single-cell RNA-seq data”. In: Nature Methods 14.6, pp. 584–586. doi: 10.1038/nmeth.4263.

Bartlett, M. S. (1936). “The square root transformation in analysis of variance”. In: Journal of the Royal Statistical Society Series B: Statistical Methodology 3.1, pp. 68–78. doi: 10.2307/2983678.

Booeshaghi, A. S. and L. Pachter (2021). “Normalization of single-cell RNA-seq counts by log(x + 1) or log(1 + x)”. In: Bioinformatics 37.15, pp. 2223–2224. doi: 10.1093/bioinformatics/btab085.

Borella, M., G. Martello, D. Risso, and C. Romualdi (2021). “PsiNorm: a scalable normalization for single-cell RNA-seq data”. In: Bioinformatics 38.1, pp. 164–172. doi: 10.1093/bioinformatics/btab641.

Breda, J., M. Zavolan, and E. van Nimwegen (2021). “Bayesian inference of gene expression states from single-cell RNA-seq data”. In: Nature Biotechnology 39.8, pp. 1008–1016. doi: 10.1038/s41587-021-00875-x.

Choudhary, S. and R. Satija (2022). “Comparison and evaluation of statistical error models for scRNA-seq”. In: Genome Biology 23.1, p. 27. doi: 10.1186/s13059-021-02584-9.

Cole, M. B., D. Risso, A. Wagner, D. DeTomaso, J. Ngai, E. Purdom, S. Dudoit, and N. Yosef (2019). “Performance assessment and selection of normalization procedures for single-cell RNA-seq”. In: Cell Systems 8.4, 315–328.e8. doi: 10.1016/j.cels.2019.03.010.

Crowell, H. L., C. Soneson, P.-L. Germain, D. Calini, L. Collin, C. Raposo, D. Malhotra, and M. D. Robinson (2020). “muscat detects subpopulation-specific state transitions from multi-sample multi-condition single-cell transcriptomics data”. In: Nature Communications 11.1, p. 6077. doi: 10.1038/s41467-020-19894-4.

Cuevas-Diaz Duran, R., H. Wei, and J. Wu (2024). “Data normalization for addressing the challenges in the analysis of single-cell transcriptomic datasets”. In: BMC Genomics 25.1, p. 444. doi: 10.1186/s12864-024-10364-5.

Dicks, S., L. Heumos, L. May, S. Jimenez, P. Angerer, I. Gold, I. Virshup, F. Fischer, M. Gill, M. Boerries, et al. (2026). GPU-accelerated single-cell analysis at scale with rapids-singlecell. doi: 10.48550/arXiv.2603.02402.

Evans, C., J. Hardin, and D. M. Stoebel (2017). “Selecting between-sample RNA-seq normalization methods from the perspective of their assumptions”. In: Briefings in Bioinformatics 19.5, pp. 776–792. doi: 10.1093/bib/bbx008.

Gálvez-Merchán, Á., K. H. J. Min, L. Pachter, and A. S. Booeshaghi (Jan. 2023). “Metadata retrieval from sequence databases with ffq”. In: Bioinformatics 39.1. doi: 10.1093/bioinformatics/btac667.

Ge, Q., Y. Sheng, J. Lu, Y. Yang, and M. Pan (2025). “Single-cell RNA-seq data normalization: a benchmarking study”. In: PLOS One 20.12, e0335102. doi: 10.1371/journal.pone.0335102.

Gigante, S., D. B. Burkhardt, D. Dager, J. Stanley, and A. Tong (2020). scprep.

Gorin, G., T. Chari, M. Carilli, J. J. Vastola, and L. Pachter (2025). “Monod: model-based discovery and integration through fitting stochastic transcriptional dynamics to single-cell sequencing data”. In: Nature Methods 22.11, pp. 2286–2300. doi: 10.1038/s41592-025-02832-x.

Gorin, G., J. J. Vastola, and L. Pachter (2023). “Studying stochastic systems biology of the cell with single-cell genomics data”. In: Cell Systems 14.10, 822–843.e22. doi: 10.1016/j.cels.2023.08.004.

Hafemeister, C. and R. Satija (2019). “Normalization and variance stabilization of single-cell RNA-seq data using regularized negative binomial regression”. In: Genome Biology 20.1, p. 296. doi: 10.1186/s13059-019-1874-1.

Hafemeister, C. and R. Satija (2020). Analyzing scRNA-seq data with the sctransform and offset models.

Hao, Y., T. Stuart, M. H. Kowalski, S. Choudhary, P. Hoffman, A. Hartman, A. Srivastava, G. Molla, S. Madad, C. Fernandez-Granda, et al. (2023). “Dictionary learning for integrative, multimodal and scalable single-cell analysis”. In: Nature Biotechnology 42.2, pp. 293–304. doi: 10.1038/s41587-023-01767-y.

Heumos, L., A. C. Schaar, C. Lance, A. Litinetskaya, F. Drost, L. Zappia, M. D. Luecken, D. C. Strobl, J. Henao, F. Curion, et al. (2023). “Best practices for single-cell analysis across modalities”. In: Nature Reviews Genetics 24.8, pp. 550–572. doi: 10.1038/s41576-023-00586-w.

Huang, J., S. C. P. Yam, K. S. Leung, M. Deng, and N. L. S. Tang (2025). “Compositional data modeling of high-dimensional single cell RNA-seq (CoDA-hd): its advantages over commonly used normalization approaches”. In: Journal of Translational Medicine 23.1, p. 1143. doi: 10.1186/s12967-025-07157-z.

Kim, H., W. Chang, S. J. Chae, J.-E. Park, M. Seo, and J. K. Kim (2024). “scLENS: data-driven signal detection for unbiased scRNA-seq data analysis”. In: Nature Communications 15.1, p. 3575. doi: 10.1038/s41467-024-47884-3.

Lause, J., P. Berens, and D. Kobak (2021). “Analytic Pearson residuals for normalization of single-cell RNA-seq UMI data”. In: Genome Biology 22.1, p. 258. doi: 10.1186/s13059-021-02451-7.

Lingen, H. J. van, M. Suarez-Diez, and E. Saccenti (2024). “Normalization of gene counts affects principal components-based exploratory analysis of RNA-sequencing data”. In: Biochimica et Biophysica Acta (BBA) - Gene Regulatory Mechanisms 1867.4, p. 195058. doi: 10.1016/j.bbagrm.2024.195058.

Lu, S., J. Yang, L. Yan, J. Liu, J. J. Wang, R. Jain, and J. Yu (2025). “Transcriptome size matters for single-cell RNA-seq normalization and bulk deconvolution”. In: Nature Communications 16.1, p. 1246. doi: 10.1038/s41467-025-56623-1.

Luecken, M. D. and F. J. Theis (2019). “Current best practices in single-cell RNA-seq analysis: a tutorial”. In: Molecular Systems Biology 15.6. doi: 10.15252/msb.20188746.

Lun, A. (Aug. 2018). “Overcoming systematic errors caused by log-transformation of normalized single-cell RNA sequencing data”. In: bioRxiv. doi: 10.1101/404962.

Lun, A., K. Bach, and J. C. Marioni (2016). “Pooling across cells to normalize single-cell RNA sequencing data with many zero counts”. In: Genome Biology 17.1. doi: 10.1186/s13059-016-0947-7.

Lytal, N., D. Ran, and L. An (2020). “Normalization methods on single-cell RNA-seq data: an empirical survey”. In: Frontiers in Genetics 11, p. 41. doi: 10.3389/fgene.2020.00041.

Moura, L. de and S. Ullrich (2021). “The Lean 4 Theorem Prover and Programming Language”. In: Automated De-duction – CADE 28. Springer-Verlag, pp. 625–635. doi: 10.1007/978-3-030-79876-5_37.

Nguyen, L. H. and S. Holmes (2019). “Ten quick tips for effective dimensionality reduction”. In: PLOS Computational Biology 15.6, e1006907. doi: 10.1371/journal.pcbi.1006907.

Pachter, L. (2026). Optimal pebbling of the hypercube. doi: 10.48550/arXiv.2606.01685.

Park, Y. and A.-C. Hauschild (2024). “The effect of data transformation on low-dimensional integration of single-cell RNA-seq”. In: BMC Bioinformatics 25.1, p. 171. doi: 10.1186/s12859-024-05788-5.

Rich, J. M., L. Moses, P. H. Einarsson, K. Jackson, L. Luebbert, A. S. Booeshaghi, S. Antonsson, D. K. Sullivan, N. Bray, P. Melsted, et al. (2026). “The impact of package selection and versioning on single-cell RNA-seq analysis”. In: Cell Systems 17.4, p. 101560. doi: 10.1016/j.cels.2026.101560.

Robinson, M. D. and A. Oshlack (2010). “A scaling normalization method for differential expression analysis of RNA-seq data”. In: Genome Biology 11.3, R25. doi: 10.1186/gb-2010-11-3-r25.

Saxena, N. (2026). “Normalization choice drives biological interpretation in single-cell RNA-seq cancer studies: a systematic benchmarking of 465 computational pipelines”. In: Computational Biology and Chemistry 124.1, p. 109100. doi: 10.1016/j.compbiolchem.2026.109100.

Sikkema, L., C. Ramírez-Suástegui, D. C. Strobl, T. E. Gillett, L. Zappia, E. Madissoon, N. S. Markov, L.-E. Zaragosi, Y. Ji, M. Ansari, et al. (2023). “An integrated cell atlas of the lung in health and disease”. In: Nature Medicine 29, pp. 1563–1577. doi: 10.1038/s41591-023-02327-2.

Tian, L., X. Dong, S. Freytag, K.-A. Lê Cao, S. Su, A. Jalal-Abadi, D. Amann-Zalcenstein, T. S. Weber, A. Seidi, J. S. Jabbari, et al. (2019). “Benchmarking single cell RNA-sequencing analysis pipelines using mixture control experiments”. In: Nature Methods 16.6, pp. 479–487. doi: 10.1038/s41592-019-0425-8.

Townes, F. W., S. C. Hicks, M. J. Aryee, and R. A. Irizarry (2019). “Feature selection and dimension reduction for single-cell RNA-Seq based on a multinomial model”. In: Genome Biology 20.1. doi: 10.1186/s13059-019-1861-6.

Trapnell, C., D. Cacchiarelli, J. Grimsby, P. Pokharel, S. Li, M. Morse, N. J. Lennon, K. J. Livak, T. S. Mikkelsen, and J. L. Rinn (2014). “The dynamics and regulators of cell fate decisions are revealed by pseudotemporal ordering of single cells”. In: Nature Biotechnology 32.4, pp. 381–386. doi: 10.1038/nbt.2859.

Vallejos, C. A., D. Risso, A. Scialdone, S. Dudoit, and J. C. Marioni (2017). “Normalizing single-cell RNA sequencing data: challenges and opportunities”. In: Nature Methods 14.6, pp. 565–571. doi: 10.1038/nmeth.4292.

Wolf, F. A., P. Angerer, and F. J. Theis (2018). “SCANPY: large-scale single-cell gene expression data analysis”. In: Genome Biology 19.1, p. 15. doi: 10.1186/s13059-017-1382-0.

Xu, C., R. Lopez, E. Mehlman, J. Regier, M. I. Jordan, and N. Yosef (2021). “Probabilistic harmonization and annotation of single-cell transcriptomics data with deep generative models”. In: Molecular Systems Biology 17.1, e9620. doi: 10.15252/msb.20209620.

You, Y., L. Tian, S. Su, X. Dong, J. S. Jabbari, P. F. Hickey, and M. E. Ritchie (2021). “Benchmarking UMI-based single-cell RNA-seq preprocessing workflows”. In: Genome Biology 22.1, p. 339. doi: 10.1186/s13059-021-02552-3.

