## Supplementary Note for "Normalization for sampled count data"

#### 1 The CLR transform

Many transformations have been applied to biological data, and while it is useful to establish, for different transformations, how they perform on benchmarks, we posit that a principled approach to selecting a transformation is also of value. Specifically, rather than asking what properties different transformations have, we can ask which transformations exhibit properties that are important for biological analysis. Below, we show that asking this question leads to a result: the properties that form desiderata for single-cell genomics count data normalization can be satisfied, and only by a single transformation.

Consider a transformation

$$T_D : \mathbb{R}^D_{>0} \to \mathbb{R}^D,$$

which we wish to apply to compositional biological data. Here $\mathbb{R}^D_{>0}$ refers to a vector of length $D$ of positive gene counts. To facilitate analysis, we may wish the transformation $T_D$ to satisfy the following properties:

1. *Rank monotonicity.* Within a cell, the transformation should preserve the ordering of parts (see Supplementary Note Figure 1 for why this is desirable). If one part is more abundant than another, the transformed value of the more abundant part should be larger. Formally, for all $\mathbf{x} \in \mathbb{R}^D_{>0}$ and all $i, j$,

   $$x_i > x_j \iff T_D(\mathbf{x})_i > T_D(\mathbf{x})_j.$$

   Equivalently, the transform preserves within-composition ranks.

2. *Perturbation additivity.* In order to utilize linear algebra on the transformed data, in particular principal component analysis (PCA), we require additivity (Aitchison, 1982; Pawlowsky-Glahn and Egozcue, 2001). Let $\odot$ denote componentwise multiplication. For all $\mathbf{x}, \mathbf{y} \in \mathbb{R}^D_{>0}$,

   $$T_D(\mathbf{x} \odot \mathbf{y}) = T_D(\mathbf{x}) + T_D(\mathbf{y}).$$

   Thus, multiplicative perturbations of the original composition are represented additively after transformation. Intuitively, fold-change effects should add on the transformed scale.

3. *Relabeling equivariance.* The transformation should not depend on the order of the input. This requirement translates to asking, for every permutation matrix $P$, that

$$T_D(P\mathbf{x}) \;=\; PT_D(\mathbf{x}).$$

Thus, relabeling the parts only relabels the transformed coordinates.

4. *Scale (depth) invariance.* The transformation should facilitate compositional analysis (Aitchison, 1982; Egozcue and Pawlowsky-Glahn, 2019; Huang et al., 2025). That is, rescaling the input, for instance by changing sequencing depth, should not affect the result of the transformation. This translates to asking that for every $a > 0$,

$$T_D(a\mathbf{x}) \;=\; T_D(\mathbf{x}).$$

5. *Natural-log calibration.* Finally, for consistency, we ask for a calibration that allows for comparison of different datasets. The size of a one unit log-fold change is fixed by

$$T_D(e, 1, \ldots, 1)_1 \;=\; \frac{D-1}{D}.$$

Equivalently, after passing to log-coordinates, the first standard basis vector has first transformed coordinate $(D-1)/D$.

The centered log-ratio transform (Aitchison, 1982) can be understood as the unique transformation satisfying a set of desired invariance properties, which we now make precise.

**Theorem 1.1** (Uniqueness of CLR). *Let $D \geq 2$. A transformation $T_D : \mathbb{R}^D_{>0} \to \mathbb{R}^D$ satisfies rank monotonicity, perturbation additivity, relabeling equivariance, scale invariance, and natural-log calibration if, and only if,*

$$T_D(\mathbf{x})_i \;=\; \log x_i - \frac{1}{D}\sum_{j=1}^{D} \log x_j \;=\; \mathrm{clr}(\mathbf{x})_i.$$

*If the calibration axiom is omitted, the only remaining freedom is multiplication by a positive scalar.*

*Proof.* The lower bound $D \geq 2$ is needed only so that we can distinguish the calibrated coordinate from at least one off-diagonal coordinate. To derive the form of $T_D$, write

$$\ell(\mathbf{x}) \;=\; (\log x_1, \ldots, \log x_D),$$

which is a continuous bijection from $\mathbb{R}^D_{>0}$ onto $\mathbb{R}^D$ with continuous inverse $\mathbf{t} \mapsto (e^{t_1}, \ldots, e^{t_D})$. Define $F : \mathbb{R}^D \to \mathbb{R}^D$ by

$$F(\mathbf{t}) \;=\; T_D(e^{t_1}, \ldots, e^{t_D}), \qquad \text{so that} \qquad T_D(\mathbf{x}) \;=\; F(\ell(\mathbf{x})).$$

From this point on, the proof is a statement about the log-coordinate map $F : \mathbb{R}^D \to \mathbb{R}^D$. In these coordinates, rank monotonicity says that $t_i > t_j$ if and only if $F(\mathbf{t})_i > F(\mathbf{t})_j$, perturbation additivity is ordinary additivity, relabeling equivariance is $F(P\mathbf{t}) = PF(\mathbf{t})$, scale invariance is invariance under adding a constant vector, and the calibration is

$$F(e_1)_1 \;=\; \frac{D-1}{D},$$

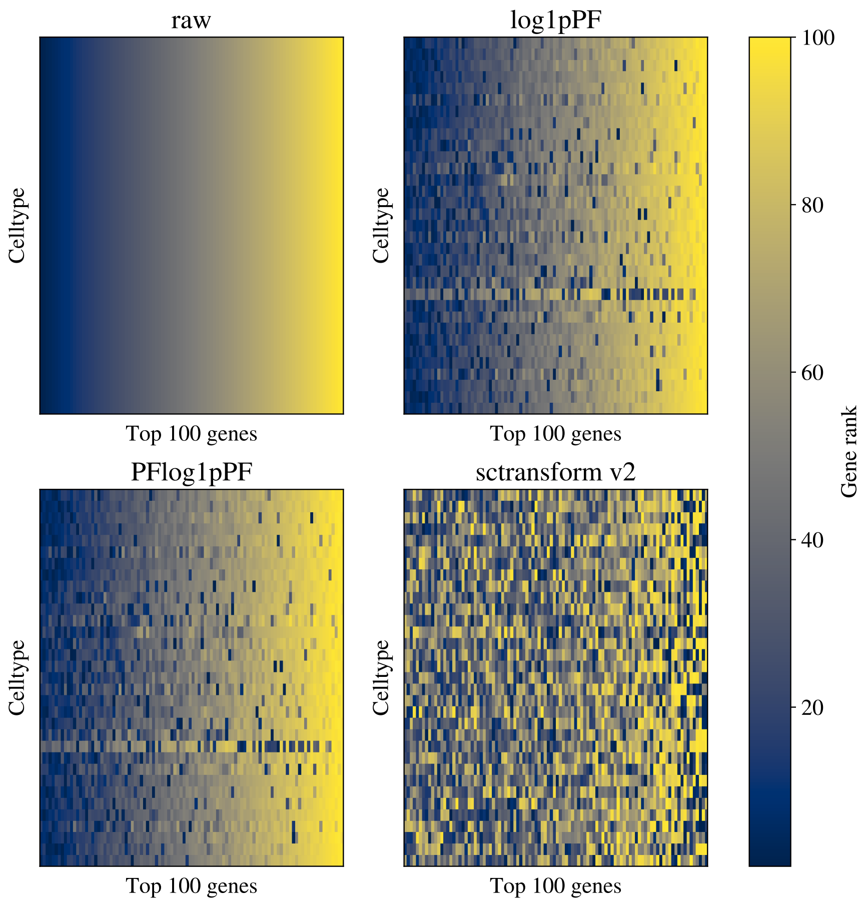


Supplementary Note Figure 1: Gene-rank scramble. For each cell type in the Angelidis et al. (2019) dataset, the 100 most highly expressed genes—ranked by mean expression in the raw counts—are selected, and within each cell type those genes are ranked from lowest to highest expressed; the resulting matrix of within-cell-type ranks is shown as a heatmap (rows are cell types, columns are rank position) for raw counts (the reference) and three normalizations (log1pPF, PFlog, sctransform v2). Because expression is averaged across the cells within each cell type before ranking, even strictly monotonic per-cell transformations (log1pPF, PFlog) shift a few genes slightly out of order — the mean of a nonlinear transform is not the transform of the mean — whereas sctransform v2, which is not monotonic, scrambles the gene order.

where $e_1 = (1, 0, \ldots, 0)$. Since $\ell(\mathbf{x} \odot \mathbf{y}) = \ell(\mathbf{x}) + \ell(\mathbf{y})$, perturbation additivity becomes Cauchy's functional equation

$$F(\mathbf{t} + \mathbf{s}) \;=\; F(\mathbf{t}) + F(\mathbf{s}).$$

Additivity gives $F(\mathbf{0}) = \mathbf{0}$ and $F(-\mathbf{t}) = -F(\mathbf{t})$. Relabeling equivariance implies that, for each $r \in \mathbb{R}$, the vector $F(re_1)$ has one value in the first coordinate and a common value in all other coordinates. Write

$$F(re_1) \;=\; \big(f(r), g(r), \ldots, g(r)\big).$$

By additivity of $F$, both $f$ and $g$ are additive functions $\mathbb{R} \to \mathbb{R}$. For $r > 0$, the first coordinate of $re_1$ is larger than the second, so rank monotonicity gives $f(r) > g(r)$. Thus, $h(r) = f(r) - g(r)$ is an additive function with $h(r) > 0$ for every $r > 0$. An additive function on $\mathbb{R}$ that is positive on the positive half-line is monotone, and hence linear. Therefore $h(r) = \alpha r$ for some $\alpha > 0$.

*Scale invariance forces centering.* Because $\ell(a\mathbf{x}) = \ell(\mathbf{x}) + (\log a)\mathbf{1}$, scale invariance is equivalent in log-coordinates to

$$F(\mathbf{t} + c\mathbf{1}) \;=\; F(\mathbf{t}) \qquad \text{for all } \mathbf{t} \in \mathbb{R}^D,\ c \in \mathbb{R}.$$

Using additivity, this is the same as $F(c\mathbf{1}) = \mathbf{0}$ for all $c \in \mathbb{R}$. In particular,

$$\mathbf{0} \;=\; F(c\mathbf{1}) \;=\; \sum_{j=1}^{D} F(ce_j),$$

so the first coordinate gives

$$f(c) + (D-1)g(c) \;=\; 0 \qquad \text{for all } c \in \mathbb{R}.$$

Together with $f - g = h = \alpha\,\mathrm{id}$, this yields

$$f(c) \;=\; \alpha\frac{D-1}{D}c, \qquad g(c) \;=\; -\alpha\frac{1}{D}c.$$

By additivity and equivariance, for any $\mathbf{t} \in \mathbb{R}^D$,

$$F(\mathbf{t})_i \;=\; \alpha\left(t_i - \frac{1}{D}\sum_{j=1}^{D} t_j\right).$$

Equivalently, there is a matrix

$$A \;=\; \alpha\left(I - \frac{1}{D}\mathbf{1}\mathbf{1}^\top\right)$$

such that $F(\mathbf{t}) = A\mathbf{t}$, and $T_D(\mathbf{x}) = A\,\ell(\mathbf{x})$. Therefore, for *every* $\mathbf{x}$,

$$\sum_{i=1}^{D} T_D(\mathbf{x})_i \;=\; \mathbf{1}^\top A\,\ell(\mathbf{x}) \;=\; 0,$$

so $T_D$ takes values in the zero-sum hyperplane

$$H_D \;=\; \left\{ z \in \mathbb{R}^D : \sum_{i=1}^{D} z_i \;=\; 0 \right\}.$$

Centering is thus derived and not assumed.

*Calibration fixes the scale.* In log-coordinates, the calibration is $F(e_1)_1 = (D-1)/D$. Since $F(e_1)$ is the first column of $A$, its first entry is $\alpha\big(1 - \frac{1}{D}\big)$. The calibration therefore forces $\alpha = 1$. Equivalently, this is the original-scale statement $T_D(e, 1, \ldots, 1)_1 = (D-1)/D$. Therefore

$$T_D(\mathbf{x})_i \;=\; \log x_i - \frac{1}{D}\sum_{j=1}^{D}\log x_j \;=\; \mathrm{clr}(\mathbf{x})_i.$$

Conversely, clr satisfies all five axioms, so it is the unique transform satisfying them. If the calibration is omitted, the only remaining freedom is the positive scalar $\alpha$, multiplication by a positive constant. □

**Proposition 1.2** (Necessity of the axioms)**.** *For $D \geq 3$, none of the five axioms in Theorem 1.1 is redundant: dropping any single one of them admits a transform that satisfies the remaining four but differs from* clr*.*

*Proof.*

- *Without rank monotonicity (Axiom 1).* Let $\phi : \mathbb{R} \to \mathbb{R}$ be additive with $\phi(1) = 1$ but $\phi \neq \mathrm{id}$; such maps exist as Hamel-basis solutions of Cauchy's equation and are necessarily non-monotone. Then

$$T_D(\mathbf{x}) = \left(I - \tfrac{1}{D}\mathbf{1}\mathbf{1}^\top\right)\left(\phi(\log x_1), \ldots, \phi(\log x_D)\right)$$

  is additive, equivariant ($\phi$ acts coordinatewise), scale invariant (the projection kills the common shift $\phi(\log a)\mathbf{1}$), and calibrated ($\phi(1) = 1$), yet it is not rank-monotone and $T_D \neq \mathrm{clr}$.

- *Without perturbation additivity (Axiom 2).* For $D \geq 3$, median centering

$$T_D(\mathbf{x})_i = \frac{D-1}{D}\Big(\log x_i - \mathrm{median}_j \log x_j\Big)$$

  is rank-monotone, equivariant, scale invariant, and calibrated, but the median is not additive, so $T_D(\mathbf{x} \odot \mathbf{y}) \neq T_D(\mathbf{x}) + T_D(\mathbf{y})$ in general and $T_D \neq \mathrm{clr}$.

- *Without relabeling equivariance (Axiom 3).* For $D \geq 3$, a weighted centering

$$T_D(\mathbf{x})_i = \log x_i - \sum_{j=1}^{D} w_j \log x_j, \qquad \sum_{j=1}^{D} w_j = 1, \quad w_1 = \tfrac{1}{D},$$

  with the remaining weights non-uniform, is rank-monotone, additive, scale invariant (the weights sum to one), and meets the calibration, yet it weights the parts unequally and is not clr.

- *Without scale invariance (Axiom 4).* The scaled logarithm

$$T_D(\mathbf{x})_i = \frac{D-1}{D}\log x_i$$

  is rank-monotone, additive, equivariant, and calibrated, but it is not centered and obeys $T_D(a\mathbf{x}) = T_D(\mathbf{x}) + \frac{D-1}{D}(\log a)\mathbf{1}$, so it still carries the overall depth.

- *Without natural-log calibration (Axiom 5).* Every map $\alpha\,\mathrm{clr}$ with $\alpha > 0$ satisfies Axioms 1–4. The calibration is exactly what removes this one-parameter scale freedom; without it the transform is pinned down only up to a positive scalar. Positive choices of $\alpha$ correspond to changes of logarithm base or unit (for instance $\alpha = 1/\ln 10$ gives the base-10 log-ratio).

□

Table 1 summarizes how the benchmarked transformations compare against the structural axioms used above. Empirically, of the three popular transforms for single-cell genomics data, sctransform fails monotonicity, log1pPF fails to remove depth dependency, and PF fails to stabilize technical variance (main Fig. 1).

For count data with zeros, the ordinary CLR is not defined on the boundary of the positive orthant. Count vectors are finite-depth measurements of an underlying relative abundance vector. The rank-monotonicity axiom has a direct count-data interpretation: if one feature is more abundant than another in a cell, the normalized values should preserve that ordering. This is the formal version

| Method | Monot. | Add. | Equiv. | Scale inv. |
|---|---|---|---|---|
| PFlog (CLR of $u + \tau$) | ✓ | ✓ | ✓ | ✓ |
| $\log(y/s + 1)$ | ✓ | × | ✓ | ✓ |
| $\operatorname{acosh}(2\alpha y/s + 1)$ | ✓ | × | ✓ | ✓ |
| $\log(y/s + 1/(4\alpha))$ | ✓ | × | ✓ | ✓ |
| $\log(\mathrm{CPM} + 1)$ | ✓ | × | ✓ | ✓ |
| $\log(y/s + 1)/u$ | ✓ | × | ✓ | ✓ |
| $\log(y/s + 1) \to$ HVG | – | – | – | – |
| $\log(y/s + 1) \to Z$ | × | × | ✓ | ✓ |
| $\log(y/s + 1) \to$ HVG $\to Z$ | – | – | – | – |
| Pearson | × | × | ✓ | × |
| Pearson (no clip) | × | × | ✓ | × |
| Analytic Pearson | × | × | ✓ | × |
| sctransform | – | – | – | – |
| Random quantile | – | – | – | – |
| Pearson$\to$ HVG | – | – | – | – |
| Pearson$\to Z$ | × | × | ✓ | × |
| Pearson$\to$ HVG $\to Z$ | – | – | – | – |
| Sanity MAP | – | – | – | – |
| Sanity distance | – | – | – | – |
| Dino | – | – | – | – |
| Normalisr | – | – | – | – |
| GLM PCA | – | – | – | – |
| NewWave | – | – | – | – |

Table 1: Axiom check for PFlog and the 22 transformations benchmarked in (Ahlmann-Eltze and Huber, 2023), excluding the natural-log calibration axiom. The columns correspond to rank monotonicity, perturbation additivity, relabeling equivariance, and scale invariance. For PFlog, additivity is the ordinary CLR perturbation additivity of the positive shifted composition $u + \tau\mathbf{1}$ after the fixed normalized shift $\tau$ has been chosen; PFlog is not additive for ordinary perturbations of the unshifted composition $u$. The table assesses the transformations as benchmarked. A dash indicates that the method is not an ordinary map $\mathbb{R}^D_{>0} \to \mathbb{R}^D$ to which the axioms directly apply, for example because it includes feature selection, randomization, distances, or low-dimensional model fitting.

of the within-cell rank preservation measured empirically by the Spearman correlations in the main text. The fixed pseudocount changes the algebra: PFlog is not additive for ordinary perturbations of the unshifted composition $u$. Instead, for a fixed normalized shift $\tau$, the count-domain transform should be compatible with a rank-monotone compositional extension of the shifted positive vector on the positive orthant satisfying the same structural axioms. This excludes transformations that depend on the accidental integer representation of the counts rather than on the underlying composition. Once a fixed shift has been added on the normalized-composition scale, the shifted vector

$$\mathbf{u}^{(\tau)} = (u_1 + \tau, \ldots, u_D + \tau)$$

lies in $\mathbb{R}^D_{>0}$. Applying the same invariance characterization to the rank-monotone extension on $\mathbf{u}^{(\tau)}$ forces the transformed coordinates to be

$$\operatorname{clr}\big(\mathbf{u}^{(\tau)}\big)_i = \log(u_i + \tau) - \frac{1}{D}\sum_{j=1}^{D} \log(u_j + \tau),$$

which is the PFlog transform derived below.

**Proposition 1.3** (Shifted CLR with fixed normalized shift)**.** *Let $D \geq 2$, $\tau > 0$, and*

$$\mathcal{E}_{D,\tau} = \left\{ \left( \frac{x_1}{s} + \tau, \dots, \frac{x_D}{s} + \tau \right) : x \in \mathbb{N}_{\geq 0}^D,\ s = \sum_{j=1}^{D} x_j > 0 \right\}$$

*be the empirical shifted compositions. Suppose a count-data transform*

$$S : \mathcal{E}_{D,\tau} \to \mathbb{R}^D$$

*is the restriction of a map $\widetilde{T} : \mathbb{R}_{>0}^D \to \mathbb{R}^D$ satisfying rank monotonicity, perturbation additivity, relabeling equivariance, scale invariance, and the natural-log calibration above. Then, for every $\mathbf{u}^{(\tau)} \in \mathcal{E}_{D,\tau}$,*

$$S(\mathbf{u}^{(\tau)})_i = \log(u_i + \tau) - \frac{1}{D} \sum_{j=1}^{D} \log(u_j + \tau).$$

*Thus, shifted CLR is the unique count-data transform, for the fixed normalized shift $\tau$, that admits such a rank-monotone compositional extension. If the calibration axiom is omitted, the same conclusion holds up to multiplication by a positive scalar.*

*Proof.* Since $\widetilde{T}$ satisfies rank monotonicity and the four structural axioms together with the calibration, the uniqueness argument above applies to $\widetilde{T}$. Hence $\widetilde{T} = \mathrm{clr}$ on $\mathbb{R}_{>0}^D$. Restricting this identity to $\mathcal{E}_{D,\tau}$ gives

$$S(\mathbf{u}^{(\tau)})_i = \widetilde{T}(\mathbf{u}^{(\tau)})_i = \mathrm{clr}(\mathbf{u}^{(\tau)})_i = \log(u_i + \tau) - \frac{1}{D} \sum_{j=1}^{D} \log(u_j + \tau).$$

The uncalibrated statement follows from the corresponding uncalibrated uniqueness result above.

The invariance argument fixes the transformation only after the normalized shift $\tau$ has been chosen. That choice is distinct from the Anscombe or delta-method question of how to choose a count-scale pseudocount. Single-cell count data has been empirically observed to fit well with negative-binomial mean-variance models (Grün et al., 2014). Under such models,

$$\mathrm{Var}(X_i) \approx \mu_i + \alpha \mu_i^2,$$

delta-method calculations motivate a shifted logarithm with pseudocount proportional to $1/\alpha$, commonly $y_0 = 1/(4\alpha)$ on the original count scale (Anscombe, 1948; Ahlmann-Eltze and Huber, 2023; Booeshaghi and Pachter, 2021). For a cell $c$, let

$$s_c = \sum_{j=1}^{D} x_{cj}, \qquad u_{ci} = \frac{x_{ci}}{s_c}.$$

If a shifted log is written in the common software form

$$\log(1 + K_c u_{ci}),$$

then matching the count-scale Anscombe pseudocount exactly requires

$$\frac{s_c}{K_c} = \frac{1}{4\alpha}, \qquad \text{or equivalently} \qquad K_c = 4\alpha s_c.$$

With this cell-specific choice, the cell depth cancels inside the shifted log:

$$\log(1 + K_c u_{ci}) \;=\; \log(4\alpha) + \log\left(x_{ci} + \frac{1}{4\alpha}\right).$$

The additive constant $\log(4\alpha)$ is common to all genes in the cell and has no effect on subsequent within-cell centering. Thus the exact Anscombe shifted-log calculation corresponds to applying a shifted logarithm on the original count scale after adding the count-scale pseudocount $1/(4\alpha)$. The PFlog transform then applies the final CLR-centering step,

$$z_{ci} \;=\; \log\left(x_{ci} + \frac{1}{4\alpha}\right) - \frac{1}{D}\sum_{j=1}^{D}\log\left(x_{cj} + \frac{1}{4\alpha}\right),$$

but that centering is the second PF/CLR step; it is not part of the delta-method derivation of the shifted logarithm itself.

By contrast, fixed-$K$ PFlog uses one software scale factor for all cells:

$$T^{\text{PFlog}}_{D,K}(\mathbf{x}_c)_i \;=\; \log(1 + K u_{ci}) - \frac{1}{D}\sum_{j=1}^{D}\log(1 + K u_{cj}).$$

For fixed $K$, the normalized-composition shift is fixed, $\tau = 1/K$, but the effective count-scale pseudocount is cell-specific:

$$y_{0c} \;=\; \frac{s_c}{K}.$$

Choosing $K = 4\alpha s_*$ for a representative depth $s_*$, such as the mean depth of a matrix, is therefore a representative-depth calibration of fixed-$K$ PFlog, not the exact cellwise Anscombe choice unless $s_c = s_*$. □

In other words, the exact Anscombe shifted-log rule fixes the count-scale pseudocount $1/(4\alpha)$ and therefore uses $K_c = 4\alpha s_c$. Fixed-$K$ PFlog instead fixes the normalized-composition shift $1/K$, and consequently assigns different count-scale pseudocounts to cells with different depths. For a representative depth $s_* = 5{,}000$, Seurat's $K = 10{,}000$ corresponds to $y_0 = 0.5$ only at that representative depth, and hence to $\alpha = 0.5$ under $y_0 = 1/(4\alpha)$, as noted by Ahlmann-Eltze and Huber (2023). The degree of technical variance stabilization is sensitive to the pseudocount chosen (Supplementary Note Figure 2), so the distinction between cell-specific Anscombe scaling and fixed-$K$ shifted CLR should be kept explicit.

This sensitivity also appears in the Angelidis2019 pseudobulk analysis used in the main text. Sweeping the count-scale pseudocount shows that the loading–log fold-change agreement is high near the estimated Anscombe value ($R^2 = 0.759$ for PFlog) but deteriorates when the CP10k pseudocount is used ($R^2 = 0.125$; main text Fig. 2f). Thus, the pseudocount selected for variance stabilization can change the biological interpretation of PC loadings.

The same issue propagates from PC loadings to cell geometry. In the 10x Genomics mouse brain `neuron_1k_v3` dataset, we recomputed PFlog with a range of count-scale pseudocounts and compared each resulting PCA $k$-NN graph to the graph obtained at the estimated Anscombe pseudocount (Supplementary Note Figure 3). The estimated pseudocount recovered 50.0 of 50 neighbors by construction, whereas the CP10k pseudocount recovered 38.0 and log1pPF recovered 37.7. This change was large enough to affect clustering: using 50 PCs and Leiden clustering with $n_{\text{neighbors}} = 30$, matching the default number of neighbors used by Seurat for UMAP, PFlog produced 13 clusters and CP10k produced 15 clusters (adjusted Rand index 0.656; normalized mutual information 0.787).

One extra CP10k cluster mapped primarily to a PFlog low-depth vascular/immune cluster, had no significant targeted marker differences from that PFlog cluster as a whole, and separated cells within that PFlog cluster by depth (median depth 2,543 versus 775 counts; two-sample Kolmogorov–Smirnov $p = 9.8 \times 10^{-14}$). Thus, the CP10k pseudocount can induce depth-associated over-splitting of a marker-coherent neighborhood. In summary, the invariance axioms select shifted CLR among transforms with a fixed normalized shift, while the delta-method Anscombe calculation selects a fixed count-scale pseudocount. These criteria coincide only when all cells have the same depth, and when they do not, the choice can affect loadings, neighborhoods, and clusters.

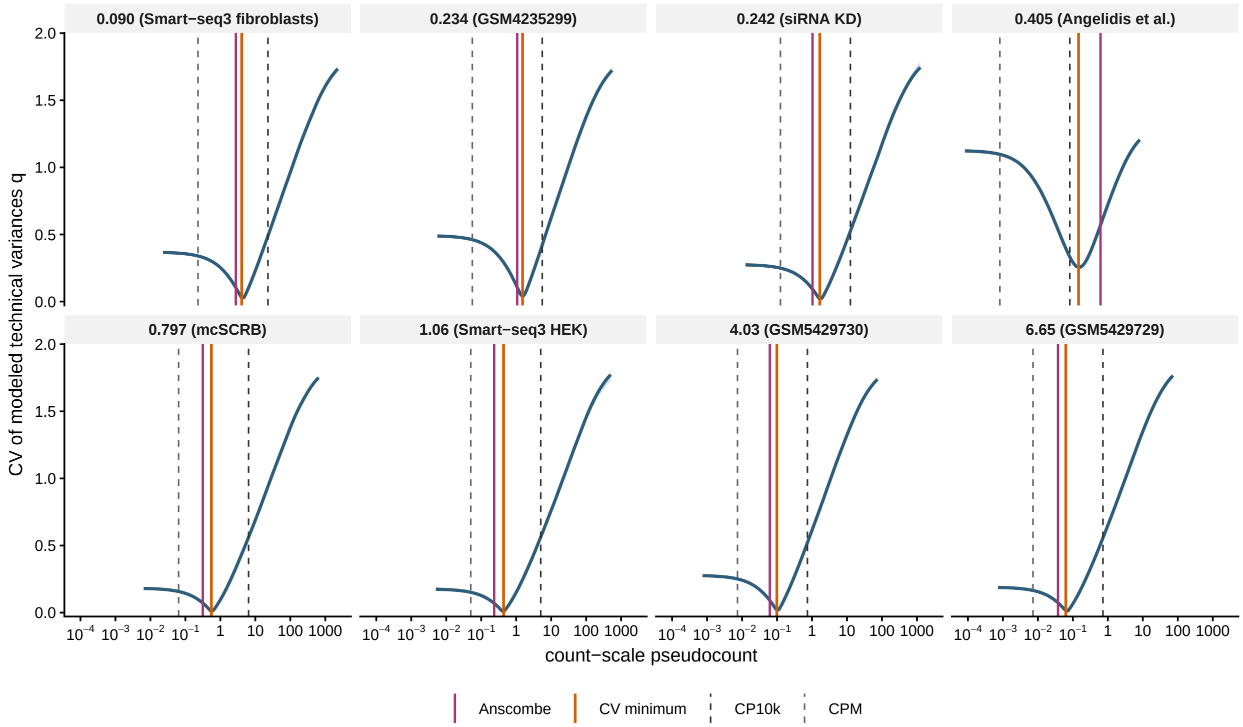


Supplementary Note Figure 2: Pseudocount effect simulation. For each simulated overdispersion/depth scenario, negative-binomial counts were transformed across a grid of shifted-log scales and the coefficient of variation of the delta-method technical gene variances was computed. Lower values indicate better technical variance stabilization. Vertical reference lines mark commonly used fixed scales and the Anscombe-derived count-scale pseudocount $1/(4\alpha)$, plotted at the scenario's representative depth. The curves show that technical variance stabilization is sensitive to the pseudocount and that fixed scales such as CP10k can be far from the Anscombe-calibrated value.

#### 2 Equivalence of PFlog and CLR

Following Aitchison's original logratio formulation (Aitchison, 1982), let

$$\mathbf{x} = (x_1, \ldots, x_D)$$

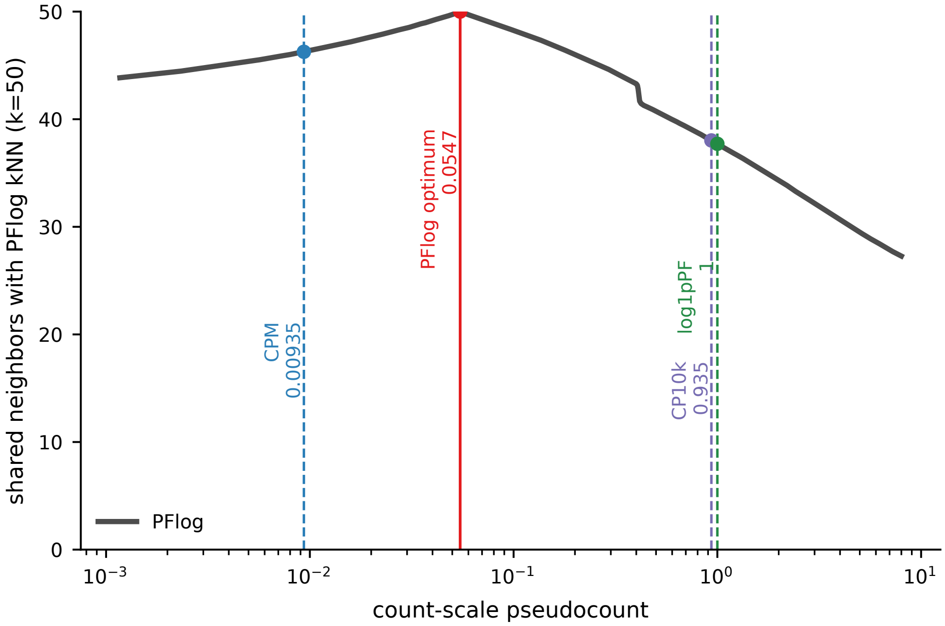


Supplementary Note Figure 3: Effect of the count-scale pseudocount on the 10x Genomics mouse brain PCA neighborhood graph. The `neuron_1k_v3` raw count matrix contained 1,301 cells and was filtered to 3,000 genes. For each pseudocount $y_0$, raw counts were transformed by PFlog, equivalently by running scclr with $\alpha = 1/(4y_0)$, embedded by PCA, and used to construct a $k = 50$ nearest-neighbor graph. The plotted value is the mean number of neighbors shared with the PFlog graph obtained at the estimated Anscombe pseudocount. Vertical lines mark the Anscombe/delta-method pseudocount estimated from the data, and the pseudocounts corresponding to CPM, log1pPF, and CP10k at the representative cell depth.

denote a $D$-part composition. Aitchison defines the centered log-ratio coordinates through the log-ratio of each part to the geometric mean of the composition:

$$z_i = \log\left(\frac{x_i}{g(\mathbf{x})}\right), \qquad g(\mathbf{x}) = \left(\prod_{j=1}^{D} x_j\right)^{1/D}, \qquad i = 1, \ldots, D.$$

Since

$$\log g(\mathbf{x}) = \frac{1}{D}\sum_{j=1}^{D} \log x_j,$$

the same expression can be written as

$$z_i = \log x_i - \frac{1}{D}\sum_{j=1}^{D} \log x_j.$$

Now let

$$\mathbf{x} = (x_1, \ldots, x_D) \in [0, \infty)^D$$

be the count vector for a cell, and let

$$s = \sum_{j=1}^{D} x_j$$

denote its total count. The first PF step divides by cell depth:

$$u_i = \frac{x_i}{s}, \qquad i = 1, \ldots, D.$$

Thus, $\mathbf{u} = (u_1, \ldots, u_D)$ is a composition in Aitchison's notation.

Let $c > 0$ be the normalized-composition shift used before the logarithm. Define the uncentered shifted-log coordinates

$$\ell_c(x)_i = \log(u_i + c).$$

Define the shifted composition

$$\mathbf{u}^{(c)} = (u_1 + c, \ldots, u_D + c).$$

Applying Aitchison's logratio definition to $\mathbf{u}^{(c)}$ gives

$$\mathrm{clr}\big(\mathbf{u}^{(c)}\big)_i = \log\left(\frac{u_i + c}{g(\mathbf{u}^{(c)})}\right),$$

where

$$g(\mathbf{u}^{(c)}) = \left(\prod_{j=1}^{D}(u_j + c)\right)^{1/D}.$$

Using the logarithm of the geometric mean,

$$\log g(\mathbf{u}^{(c)}) = \frac{1}{D}\sum_{j=1}^{D}\log(u_j + c),$$

we obtain

$$\mathrm{clr}\big(\mathbf{u}^{(c)}\big)_i = \log(u_i + c) - \frac{1}{D}\sum_{j=1}^{D}\log(u_j + c).$$

Thus CLR centering the uncentered shifted-log coordinates is exactly shifted CLR:

$$T_{\mathrm{PFlog},c}(x)_i = \ell_c(x)_i - \frac{1}{D}\sum_{j=1}^{D}\ell_c(x)_j = \log(u_i + c) - \frac{1}{D}\sum_{j=1}^{D}\log(u_j + c) = \mathrm{clr}\big(\mathbf{u}^{(c)}\big)_i.$$

So PFlog is the shifted logarithm after the first proportional fitting step together with the final per-cell centering operation. That centering operation is exactly the CLR centering step, and it is what makes PFlog the centered log-ratio transform of the shifted, depth-normalized composition. In particular,

$$\sum_{i=1}^{D} T_{\mathrm{PFlog},c}(x)_i = 0,$$

and, after exponentiation, the transformed coordinates have geometric mean 1.

PFlog is not only easy to compute, it is also efficient to work with: the sparse shifted-log matrix can be stored together with its centering vector, so downstream linear algebra can use the implicit centered matrix without materializing it. This is not the case for sctransform, which has high memory requirements in addition to slow runtimes (Supplementary Note Figure 4).

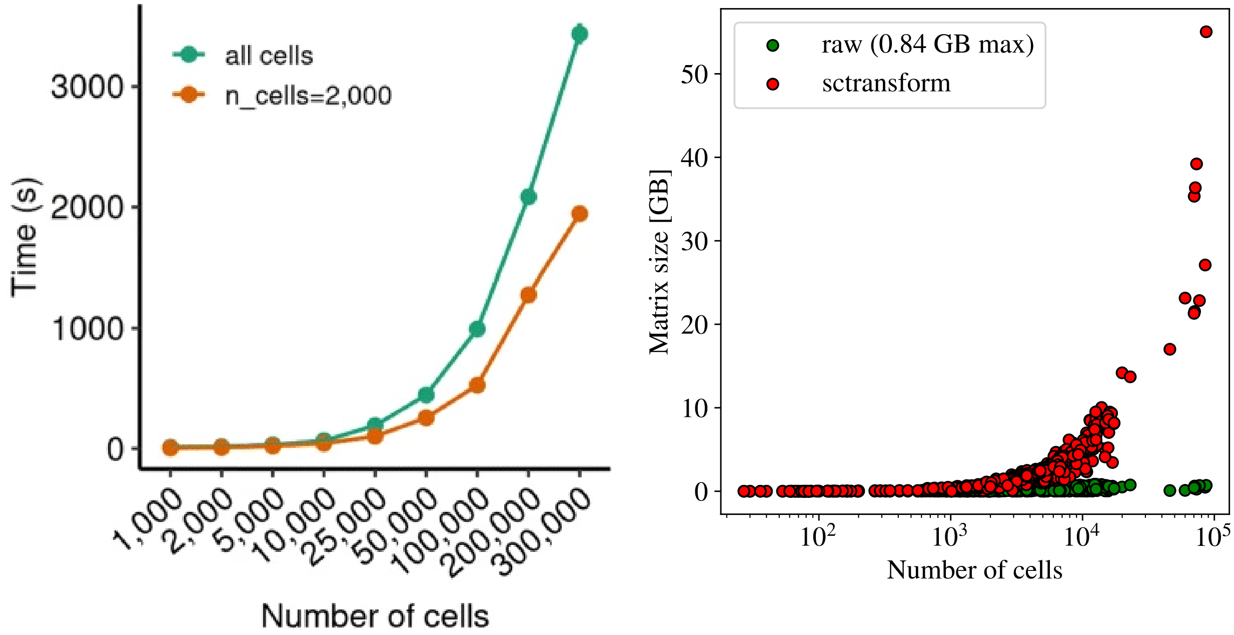


Supplementary Note Figure 4: Speed and memory comparison. (a) Runtime of sctransform v2 vs. the number of cells. The figure is a reproduction, without alteration, of Figure 3c from (Choudhary and Satija, 2022), which is licensed under the Creative Commons Attribution 4.0 International License. (b) In-memory size of dense sctransform matrices compared with sparse raw-count matrices as a function of cell number across the benchmarked datasets. The comparison illustrates why methods that materialize dense normalized matrices can become limiting for large single-cell analyses, whereas PFlog can be represented as a sparse shifted-log matrix plus a centering vector.

#### 3 Ahlmann-Eltze–Huber pseudocount analysis

Ahlmann-Eltze and Huber (2023) derive the negative-binomial Anscombe transform

$$g(y) = \frac{1}{\sqrt{\alpha}} \operatorname{acosh}(2\alpha y + 1),$$

and its shifted-log large-count approximation

$$g(y) \approx \frac{1}{\sqrt{\alpha}} \left[ \log(4\alpha) + \log\left( y + \frac{1}{4\alpha} \right) \right].$$

Ignoring the additive constant and the overall scale, this is the shifted log

$$\log\left( y + \frac{1}{4\alpha} \right),$$

so the Anscombe-derived pseudocount is $1/(4\alpha)$ on the original count scale.

However, the implementation benchmarked by Ahlmann-Eltze and Huber (2023) first divides each count by a cell size factor and then applies the shifted log. Writing $d_c = \sum_i y_{ci}$ for the cell depth, $\bar{d}$ for the mean depth, and $r_c = d_c/\bar{d}$ for the relative size factor used in the benchmark code, the implemented adaptive-pseudocount transform is, up to constants,

$$\log\left( \frac{y_{ci}}{r_c} + \frac{1}{4\alpha} \right).$$

This is not the Anscombe shifted log on the original count scale. Algebraically,

$$\log\left(\frac{y_{ci}}{r_c} + \frac{1}{4\alpha}\right) = \log\left(y_{ci} + \frac{r_c}{4\alpha}\right) - \log r_c.$$

The final term is constant within a cell and cancels under a within-cell centering operation, but the effective count-scale pseudocount is

$$y_{0c}^{\mathrm{AEH}} = \frac{r_c}{4\alpha},$$

not $1/(4\alpha)$. Thus the benchmarked transform is off from the Anscombe approximation by a factor of $r_c$ in its count-scale pseudocount. A cell at the mean depth has $r_c = 1$ and no discrepancy, but a cell at one tenth of the mean depth has a ten-fold too small pseudocount, while a cell at ten times the mean depth has a ten-fold too large pseudocount. In the 10% downsampling setting, if full-depth and downsampled cells are pooled in equal numbers, the relative size factors are approximately $1/0.55 \approx 1.82$ and $0.1/0.55 \approx 0.18$, so the two members of a full-depth/downsampled pair are assigned count-scale pseudocounts differing by a factor of ten.

The issue is therefore not merely notational. The correct shifted-log approximation to the Anscombe transform is

$$\log\left(y_{ci} + \frac{1}{4\alpha}\right),$$

with the additive constant $\log(4\alpha)$ suppressed. Dividing by $r_c$ before applying the same $1/(4\alpha)$ shift changes the pseudocount into $r_c/(4\alpha)$, making the transformation explicitly depth dependent. The same normalization-before-transformation issue also affects the acosh transform when it is applied as $\operatorname{acosh}(2\alpha y_{ci}/r_c + 1)$: it is no longer the raw-count Anscombe transform derived from $\operatorname{Var}(Y) = \mu + \alpha\mu^2$ with the same $\alpha$.

This explains why the Ahlmann-Eltze–Huber benchmark can understate the role of the Anscombe pseudocount. Their discussion says that "our benchmarks showed limited performance benefits for these," referring to the acosh transform and the adaptive pseudocount. But the adaptive pseudocount they benchmarked was not the count-scale Anscombe pseudocount $1/(4\alpha)$; it was a depth-varying pseudocount $r_c/(4\alpha)$. Supplementary Note Figure 2 shows that technical variance stabilization is sensitive to pseudocount choice. The relevant diagnostic comparison is therefore between the raw-count Anscombe approximation $\log(y + 1/(4\alpha))$ and the normalized-count implementation $\log(y/r_c + 1/(4\alpha))$, especially in the downsampling benchmark where $r_c$ is deliberately changed. The practical consequence is evident in the downstream benchmark of Ahlmann-Eltze and Huber (2023): in their Figure 2, reproduced and adapted in Supplementary Note Figure 6, many methods appear nearly indistinguishable, even though the pseudocount analysis above shows that the benchmarked shifted-log and acosh implementations are not the Anscombe transforms they are meant to approximate.

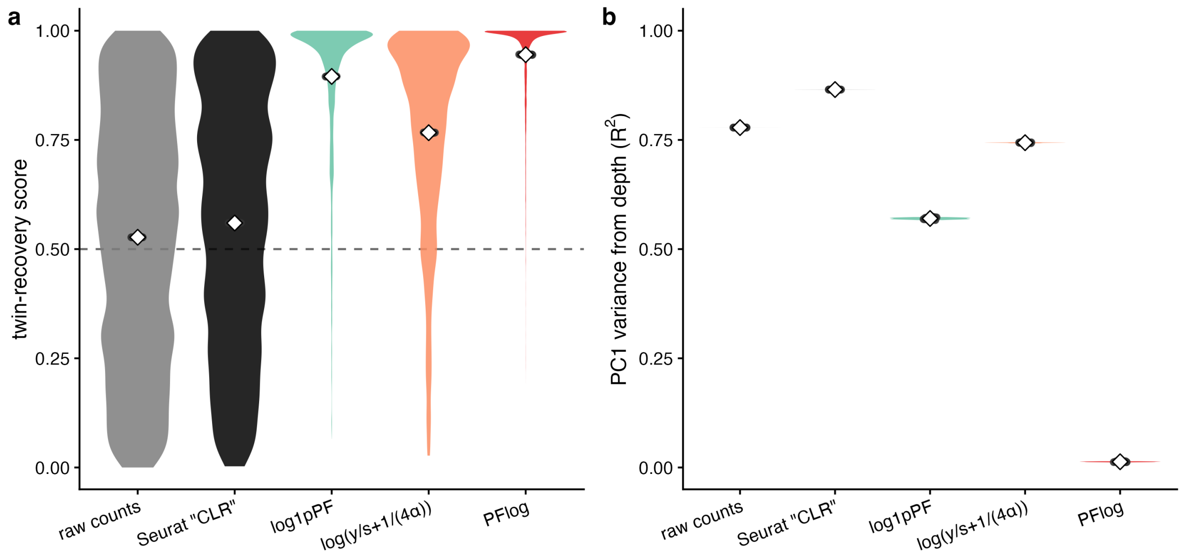


Supplementary Note Figure 5: Depth effects in a full-depth/downsampled twin experiment based on the downsampling setting of (Ahlmann-Eltze and Huber, 2023). Deeply-sequenced Smart-seq3 fibroblasts and their own twins, each downsampled to 10% of its depth, were placed in a single PCA embedding. The only difference between a cell and its twin is sequencing depth, so a depth-invariant transform should mix the two depth groups. **(a)** Per-cell twin-recovery score (1 = a cell's own twin is its nearest cross-depth neighbor, 0.5 = random; white diamonds mark the mean), pooled over cells and 20 downsampling seeds. **(b)** Coefficient of determination for PC1 regressed on depth group ($R^2$; 0 = depth fully removed), one value per seed. The $\log(y/s + 1/(4\alpha))$ label denotes the benchmarked shifted-log transform of Ahlmann-Eltze and Huber (2023), with $\alpha$ estimated from each joint full-depth/downsampled matrix.

There are additional issues with the assessment. In the simulation panel, two of the original simulation frameworks, linear walk and random walk, confound depth with the biological effect: the simulated expression state is multiplied by real cell-specific depths, so a method that removes depth can be penalized for removing variation encoded in the benchmark's ground truth. For that reason, Supplementary Note Figure 6 excludes those two simulations and retains muscat, dyngen, and scDesign2. Even in the retained simulations, the absolute $k$-NN recovery is low for all methods because the methods must infer a latent ground-truth neighborhood from noisy sampled counts. Thus, although PFlog ranks second of 25 methods on muscat and is above the method average on the retained simulations, the absolute improvement in panel (b) is necessarily modest.

The downsampling panel has a different caveat. The original benchmark scored each method against a consensus neighborhood derived from the methods being compared, so adding a new method would itself change the target. To avoid that circularity, PFlog was not used to build the consensus in Supplementary Note Figure 6c. Instead, the full-depth consensus neighborhood was computed from the log-family methods highlighted as strong performers by Ahlmann-Eltze and Huber (2023), and PFlog downsampled neighbors were scored against that independently defined log-method consensus. Under this holdout-consensus metric, PFlog recovers, on average, 11.3 neighbors out of 50, compared to 5.77 for the other methods.

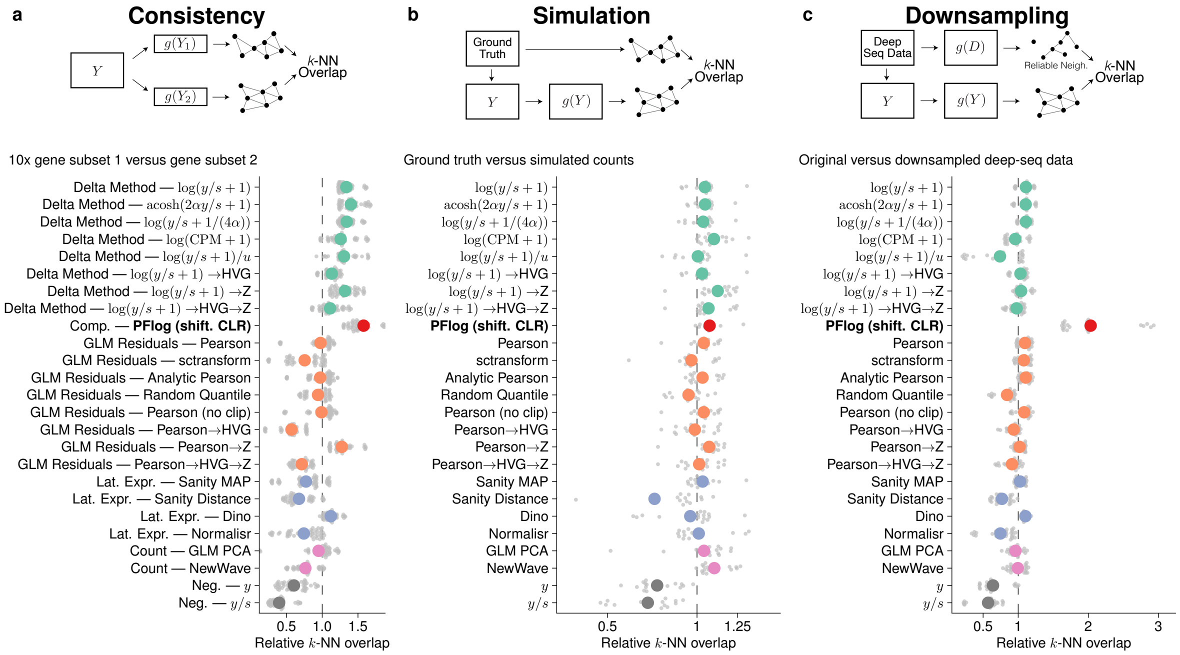


Supplementary Note Figure 6: Adapted reproduction of Ahlmann-Eltze and Huber (2023) Figure 2a–c, with an added CLR row shown in red. This row corresponds to PFlog (shifted CLR) and was not present in the original benchmark. The original figure is licensed under the Creative Commons Attribution 4.0 International License. **(a)** Consistency: overlap between $k$-NN neighborhood graphs from two 10x gene subsets. **(b)** Simulation: overlap between $k$-NN neighborhoods from simulated counts and ground truth, retaining muscat, dyngen, and scDesign2 and excluding the linear-walk and random-walk simulations. **(c)** Downsampling: overlap between $k$-NN neighborhoods from original deep-seq data and downsampled deep-seq data. Vertical dashed lines mark relative overlap of 1.

The excluded walk simulations are problematic not only because they encode depth in the ground-truth geometry, but also because the resulting count matrices have unrealistic depth behavior. Supplementary Note Figure 7 shows this directly for the cached simulations used in the benchmark audit. In the linear-walk simulation, ground-truth neighbors have approximately half the raw-depth gap of random pairs. In the random-walk simulation, the effect is weaker but still present, and the simulation also produces extreme total-count outliers: in seed 1, the median depth is 141 counts and the mean depth is 487 counts, but the maximum depth is 594,005 counts, with four cells above 50,000 counts. This is not a realistic model of technical sequencing depth. It illustrates why benchmarks that define biological ground truth from these walk processes can penalize a normalization method for removing depth-associated variation that the simulator itself has made part of the target.

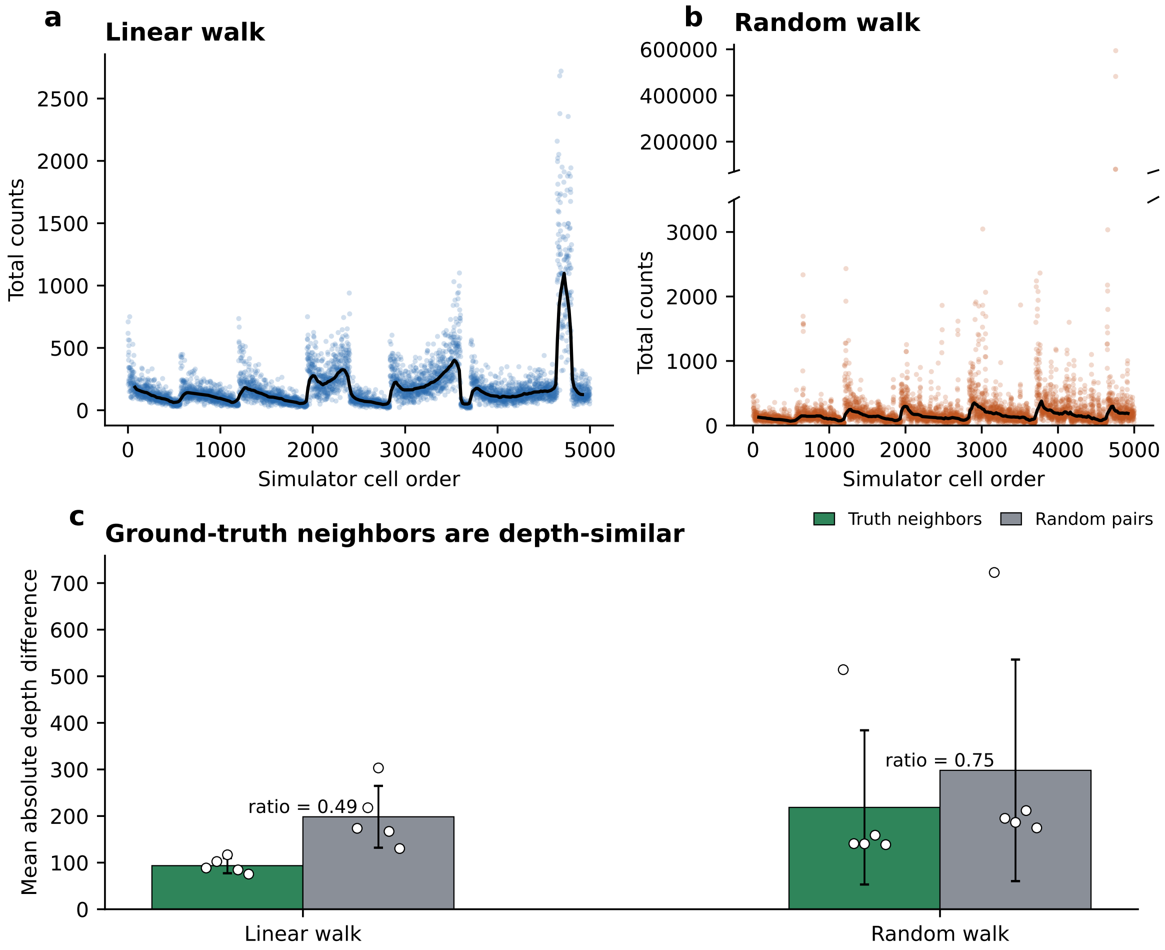


Supplementary Note Figure 7: Depth confounding in the linear-walk and random-walk simulations. **(a,b)** Raw total counts across simulator cell order for seed 1. The random-walk panel uses a broken y-axis to show both the main depth variation and the handful of extreme outliers. **(c)** Mean absolute raw-depth difference between cells and their simulator ground-truth $k = 50$ neighbors, compared with random cell pairs, across five seeds. Values below the random-pair bar indicate that the simulator ground-truth neighborhood structure is depth-similar. Thus, total depth is predictive of the benchmark target in these simulations.

### 4 The Seurat "CLR" transform

Normalization is the first analysis step in the Seurat workflow (see Supplementary Note Figure 8), and the procedure therefore affects all downstream analyses and derived results. The default normalization in Seurat in the notation of (Ahlmann-Eltze and Huber, 2023) is $\log(\gamma x_i/s+1)$, where $s$ is the cell depth and $\gamma = 10{,}000$, i.e. log1pCP10k (Rich et al., 2026).

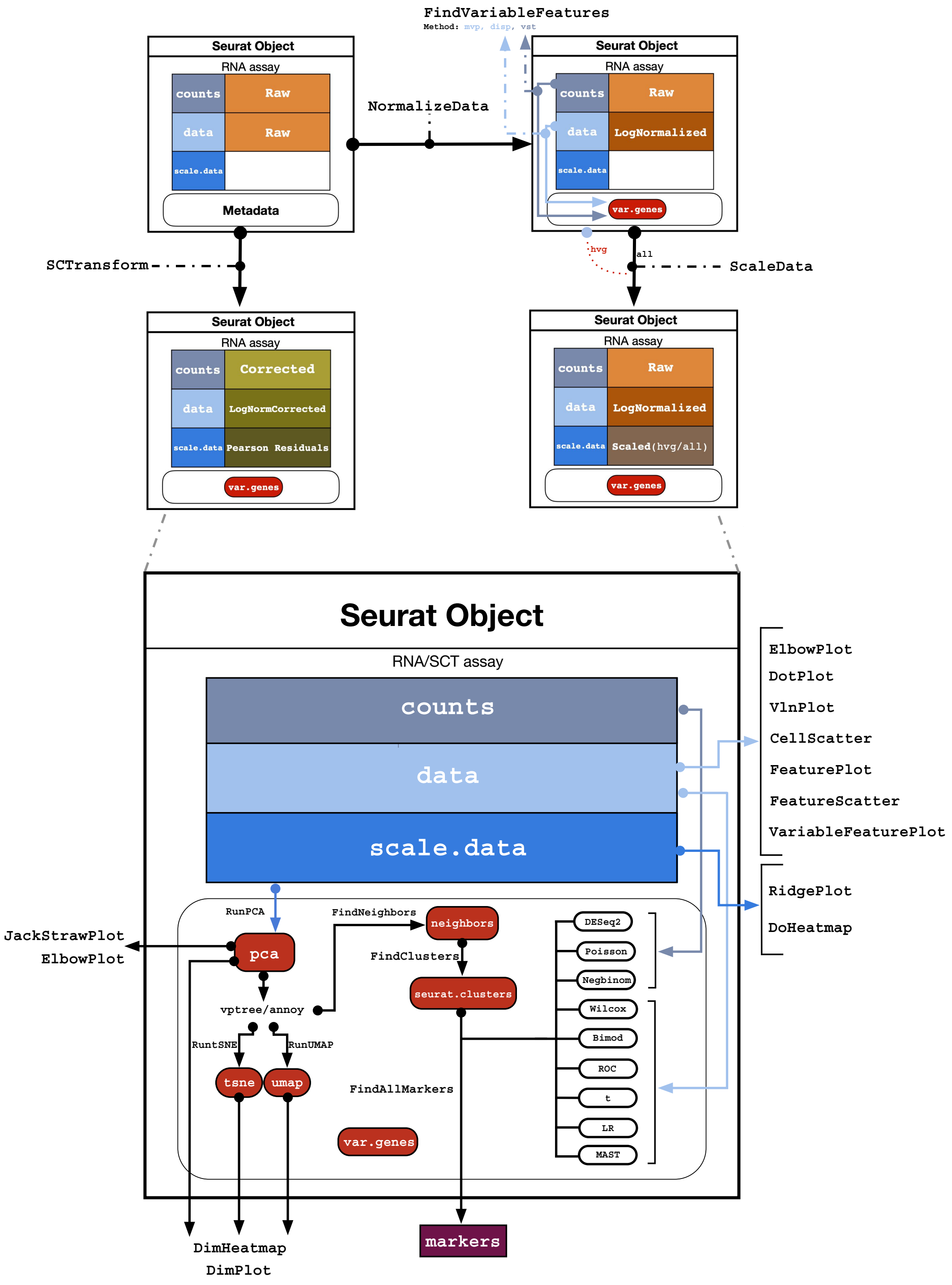


Supplementary Note Figure 8: The Seurat analysis workflow as implemented in the Seurat source code. Normalization is performed early through `NormalizeData`, before variable-feature selection, scaling, PCA, neighborhood graph construction, clustering, and UMAP. Consequently, the mathematical behavior of a selected normalization method, including Seurat's method named "CLR", propagates to the downstream geometry and clustering results.

Seurat also provides a normalization called "CLR"[1], which is recommended, and therefore widely used, for normalizing antibody-derived tags (ADTs) in CITE-seq data (Stoeckius et al., 2017). Below, we show that the Seurat implementation of CLR is neither CLR nor shifted CLR, and can behave poorly as a result.

For a vector $x = (x_1, \ldots, x_p) \in [0, \infty)^p$, let

$$P(x) \;=\; \{j : x_j \;>\; 0\}, \qquad p \;=\; \text{length}(x).$$

Define Seurat's normalization factor by

$$h_+(x) \;=\; \exp\left(\frac{1}{p}\sum_{j\,\in\,P(x)} \log(1+x_j)\right) \;=\; \left(\prod_{j\,\in\,P(x)} (1+x_j)\right)^{1/p}.$$

Then the Seurat transform applied coordinatewise can be written as

$$T_{\text{Seurat}}(x)_i \;=\; \log\left(1 + \frac{x_i}{h_+(x)}\right).$$

This is not the centered log-ratio transform, and the problem is not just the exclusion of zeros. The more basic issue is that the pseudocount is being added after division by a data-dependent normalization factor rather than before the logarithm on the original count scale. In other words, Seurat computes

$$\log\left(1 + \frac{x_i}{h_+(x)}\right)$$

rather than a shifted log of the form

$$\log(x_i + c) - \text{centering term}.$$

These are fundamentally different operations. In the Seurat expression, the effective pseudocount on the original scale is $h_+(x)$, which is data dependent and changes from cell to cell. Thus, the shift is not fixed; it depends on the observed positive entries of the very vector being transformed. A transformation of the form $\log(x_i + c)$ with fixed $c$ preserves the meaning of the pseudocount across cells. A transformation of the form $\log(1 + x_i/h_+(x))$ does not.

There are therefore two distinct departures from CLR. First, the normalization factor is computed from $\log(1 + x_j)$ only over the positive entries, so zeros are excluded from the product while the full vector length remains in the denominator. Second, after forming the ratio to that data-dependent factor, Seurat applies $\log(1 + t)$ rather than $\log t$. As a consequence,

$$T_{\text{Seurat}}(x)_i \;=\; 0 \qquad \text{whenever } x_i \;=\; 0,$$

whereas ordinary CLR is not defined at zero unless a pseudocount is introduced first. This discrepancy was noticed by Seurat users; issue #1268 reports that Seurat's "CLR" output remains positive and

[1]At commit `4608b3aa98d7089c57799e6c751c416e400c67d6`, the relevant code (for the setting margin = 2 recommended for ADT data) in `NormalizeData.default` defines

$$x \;\mapsto\; \log\left(1 + \frac{x}{\exp\left(\frac{1}{p}\sum_{x_j\,>\,0}\log(1+x_j)\right)}\right),$$

implemented as `log1p(x = x / (exp(x = sum(log1p(x = x[x > 0]), na.rm = TRUE) / length(x = x))))`.

broadens the low-signal distribution relative to a manually computed CLR (satijalab/seurat, 2019). For positive entries there is an exact simplification:

$$T_{\text{Seurat}}(x)_i \;=\; \log\bigl(x_i + h_+(x)\bigr) - \log h_+(x), \qquad i \,\in\, P(x).$$

So Seurat's "CLR" is simply a shifted log transform with a shift equal to a data-dependent factor computed from the positive entries, followed by subtraction of the same data-dependent constant. That is not CLR.

The point can already be seen with only two parts. Let $x = (x_1, x_2)$ with both entries positive. Then

$$h_+(x) \;=\; \sqrt{(1 + x_1)(1 + x_2)},$$

and Seurat's transform becomes

$$T_{\text{Seurat}}(x)_1 \;=\; \log\left(1 + \frac{x_1}{\sqrt{(1 + x_1)(1 + x_2)}}\right), \qquad T_{\text{Seurat}}(x)_2 \;=\; \log\left(1 + \frac{x_2}{\sqrt{(1 + x_1)(1 + x_2)}}\right).$$

To compare this to shifted CLR in the PFlog sense, first depth-normalize:

$$u_1 \;=\; \frac{x_1}{x_1 + x_2}, \qquad u_2 \;=\; \frac{x_2}{x_1 + x_2}.$$

With shift $c = 1$, the PFlog instance of the shifted CLR / PFlog transform is

$$T_{\text{PFlog}}(x)_i \;=\; \log(u_i + 1) - \frac{1}{2}\bigl[\log(u_1 + 1) + \log(u_2 + 1)\bigr].$$

Since $u_2 = 1 - u_1$, the two coordinates are exact negatives:

$$T_{\text{PFlog}}(x)_1 \;=\; \frac{1}{2}\log\left(\frac{u_1 + 1}{u_2 + 1}\right), \qquad T_{\text{PFlog}}(x)_2 \;=\; -\frac{1}{2}\log\left(\frac{u_1 + 1}{u_2 + 1}\right).$$

Writing $r = x_1/x_2$, so that

$$u_1 \;=\; \frac{r}{1 + r}, \qquad u_2 \;=\; \frac{1}{1 + r},$$

gives

$$T_{\text{PFlog}}(x)_1 \;=\; \frac{1}{2}\log\left(\frac{1 + 2r}{r + 2}\right), \qquad T_{\text{PFlog}}(x)_2 \;=\; -\frac{1}{2}\log\left(\frac{1 + 2r}{r + 2}\right).$$

So in the two-part case the contrast is very clear:

$$T_{\text{Seurat}}(x) \;=\; \left(\log\left(1 + \frac{x_1}{h_+(x)}\right), \log\left(1 + \frac{x_2}{h_+(x)}\right)\right),$$

whereas

$$T_{\text{PFlog}}(x) \;=\; \left(\frac{1}{2}\log\left(\frac{1 + 2r}{r + 2}\right), -\frac{1}{2}\log\left(\frac{1 + 2r}{r + 2}\right)\right).$$

Both contain a "+1", but the +1 is being used in completely different ways. In PFlog, the +1 is added before the logarithm to each normalized part and the result is then centered, so the two coordinates still form a genuine log-ratio pair. In Seurat, the +1 is added only after division by a data-dependent normalization factor, so the two coordinates are not centered and no longer represent opposite sides of the same log-ratio.

For orientation, the ordinary two-part CLR is

$$\operatorname{clr}(x)_1 = \frac{1}{2}\log\left(\frac{x_1}{x_2}\right), \qquad \operatorname{clr}(x)_2 = \frac{1}{2}\log\left(\frac{x_2}{x_1}\right).$$

These coordinates are exact negatives of one another. Seurat's are not:

$$T_{\text{Seurat}}(x)_1 + T_{\text{Seurat}}(x)_2 = \log\left(1+\frac{x_1}{h_+(x)}\right) + \log\left(1+\frac{x_2}{h_+(x)}\right) > 0.$$

By contrast,

$$T_{\text{PFlog}}(x)_1 + T_{\text{PFlog}}(x)_2 = 0.$$

So even in the simplest possible case, Seurat does not land in the CLR hyperplane, whereas shifted CLR / PFlog does.

The two-part case also shows how far off the Seurat transform can be. Write $r = x_1/x_2$. Then

$$T_{\text{Seurat}}(x)_1 = \log\left(1+\frac{x_1}{\sqrt{(1+x_1)(1+x_2)}}\right), \qquad T_{\text{PFlog}}(x)_1 = \frac{1}{2}\log\left(\frac{1+2r}{r+2}\right).$$

For a fixed small component and $r \gg 1$, the Seurat coordinate grows according to the data-dependent denominator rather than remaining a centered log-ratio contrast, whereas

$$T_{\text{PFlog}}(x)_1 \longrightarrow \frac{1}{2}\log 2 \qquad \text{as } r \to \infty.$$

That limiting value is finite because PFlog is being applied after proportional fitting, so once one component dominates the composition there is a hard upper limit to the shifted log-ratio with shift 1. But the two coordinates still remain perfectly centered:

$$T_{\text{PFlog}}(x)_2 = -T_{\text{PFlog}}(x)_1.$$

For Seurat, the small component behaves very differently:

$$T_{\text{Seurat}}(x)_2 = \log\left(1+\frac{x_2}{h_+(x)}\right),$$

so the small coordinate is forced toward a small nonnegative value rather than becoming the negative counterpart of the large coordinate. For example, if $x = (10^6, 1)$, then

$$T_{\text{Seurat}}(x) \approx (6.56, 0.0007),$$

whereas

$$\frac{x}{x_1 + x_2} \approx (0.999999, 0.000001),$$

and hence

$$T_{\text{PFlog}}(x) \approx \left(\frac{1}{2}\log\frac{1.999999}{1.000001}, -\frac{1}{2}\log\frac{1.999999}{1.000001}\right) \approx (0.347, -0.347).$$

So Seurat produces one large positive coordinate and one nearly zero coordinate, while shifted CLR / PFlog produces a centered positive/negative pair. That is exactly the distortion introduced by putting the +1 in the wrong place.

A shifted-log version shows the same point. If one instead used a fixed pseudocount $c > 0$ before the log, then for two parts

$$\log(x_i + c) - \frac{1}{2}\big[\log(x_1 + c) + \log(x_2 + c)\big]$$

remains centered and still represents a genuine log-ratio of shifted parts. By contrast, Seurat's

$$\log\left(1 + \frac{x_i}{h_+(x)}\right)$$

adds the pseudocount only after rescaling by a data-dependent normalization factor. This is exactly the wrong place to apply log1p, because it turns a fixed shift on the count scale into a variable shift whose size depends on the cell.

Unlike CLR, the Seurat transform is not centered:

$$\sum_{i=1}^{p} T_{\text{Seurat}}(x)_i \neq 0$$

in general. Therefore, it does not map data into the CLR hyperplane and it is not a log-ratio transformation in the Aitchison sense. A concise summary is that Seurat's "CLR" is a zero-preserving shifted log heuristic relative to a data-dependent factor computed from positive entries. It should not be described as the centered log-ratio transform.

The reason this transformation does not remove depth dependence is that there is no final centering or proportional fitting step after the shifted log. If CLR normalization is applied across cells rather than across features, as allowed by Seurat's `margin` argument, then it is not a within-cell normalization at all, so cell depth is not removed by construction (satijalab/seurat, 2023). If it is applied across features within a cell, the formula still depends on the set of positive entries through $P(x)$ and on the data-dependent factor $h_+(x)$. Real changes in sequencing depth are not exact multiplicative rescalings of a cell; they change which low-count features are observed as zero, and therefore change both $P(x)$ and $h_+(x)$. Because Seurat excludes zeros from the normalization factor, depth-dependent detection directly affects the transform.

This interpretation is consistent with the benchmark in the PsiNorm paper, where Borella et al. reported that CLR had the largest average correlation between PCA and depth among the methods they compared (Borella et al., 2021). To confirm that finding, we examined the performance of Seurat "CLR" in the context of the downsampling experiment of (Ahlmann-Eltze and Huber, 2023). In agreement with (Borella et al., 2021), we find that the "CLR" underperforms on removing depth via the normalization. In fact, we find that Seurat "CLR" is worse than not transforming the data at all (see Supplementary Note Figure 5).

Concretely, when full-depth cells and their own downsampled twins are placed in a single PCA embedding, depth accounts for 87% of the variance of the first principal component under Seurat "CLR", compared with 78% for raw counts, 57% for $\log(y/s + 1)$, 74% for the Ahlmann-Eltze–Huber shifted log, and only 1% for PFlog (Supplementary Note Figure 5b). Correspondingly, under Seurat "CLR" a cell embeds next to other cells of similar depth rather than next to its own twin, whereas PFlog very nearly recovers the twin pairing (Supplementary Note Figure 5a). The practical implication is that applying Seurat "CLR" does not merely fail to remove sequencing depth, it amplifies the dependence on depth relative to leaving the counts untransformed, so any downstream dimensionality reduction, clustering, or differential expression built on "CLR"-normalized data inherits a depth artifact. That is exactly what one would expect from a zero-sensitive shifted log heuristic rather than a true centered log-ratio normalization. Related confusion about the use and interpretation of Seurat's "CLR" is also evident in issue #3550 (satijalab/seurat, 2020).

### 5 Sparse shifted CLR

The final centering step in shifted CLR has an apparent computational drawback. Let $A \in \mathbb{R}^{n \times D}$ denote the sparse matrix obtained after proportional fitting and $\log(1+x)$, with cells as rows and genes as columns. Since zero counts remain zero through these first two steps, $A$ can be stored sparsely. The shifted CLR matrix is

$$Z_{ij} \;=\; A_{ij} - m_i, \qquad m_i \;=\; \frac{1}{D}\sum_{j=1}^{D} A_{ij}.$$

If $Z$ is represented naively, every structural zero in $A$ becomes $-m_i$. Except in the degenerate case $m_i = 0$, the centered matrix therefore does not have structural zeros within the row $i$, and the output is dense. This is the source of the usual memory concern for applying CLR-style centering to large single-cell matrices.

However, the dense matrix need not be stored, nor is it required for downstream operations. The same object can be represented exactly by the sparse matrix $A$ together with the centering vector $m$. Algebraically,

$$Z \;=\; A - m\mathbf{1}^T,$$

where $\mathbf{1} \in \mathbb{R}^D$ is the all-ones vector. Thus, the shifted CLR matrix is a sparse matrix plus a rank-one correction. Any computation that only requires products with $Z$ or $Z^T$ can use

$$Zv \;=\; Av - m(\mathbf{1}^T v), \qquad Z^T u \;=\; A^T u - \mathbf{1}(m^T u),$$

with the sparse products $Av$ and $A^T u$ performed on $A$ and the remaining terms computed from dense vectors of length $n$ or $D$. This is an exact representation of the shifted CLR matrix, not an approximation.

This representation is compatible with sparse PCA. The large-scale PCA problem does not require random access to every entry of a dense normalized matrix; it requires only repeated multiplication by the data matrix, its transpose, or the associated covariance operator. Since those products can be evaluated exactly from $A$ and $m$, an iterative sparse PCA routine can compute principal components for shifted CLR data while keeping the normalized matrix in the sparse-plus-centering form. Column centering or scaling, when used, can be folded into the same linear-operator view by applying the corresponding dense diagonal or low-rank corrections during the matrix-vector products.

The `scclr` implementation uses this observation directly. Rather than requiring shifted CLR to be written out as a dense `AnnData` matrix, `scclr` stores the sparse transformed matrix and the row-centering vector in a custom Rust `AnnData` representation. Downstream routines can then treat the object as if it were the dense shifted CLR matrix while the storage remains sparse. In particular, the Python interface can expose a conventional `AnnData`-like workflow, while the native implementation keeps the normalization state in the sparse-plus-centering representation.

The PCA backend used for this purpose is `rupca`. It was written to operate on this shifted sparse representation rather than on a materialized dense matrix. Its eigensolver machinery relies on a Rust port of the relevant ARPACK material for implicitly restarted Krylov subspace iteration, with the matrix operation supplied by the shifted CLR linear operator above. Thus each iteration uses sparse matrix-vector products with $A$, together with the rank-one centering correction, instead of dense products with $Z$.

Together, `scclr` and `rupca` provide a native Rust implementation of sparse shifted CLR normalization and sparse PCA on the resulting data structure. The same functionality is wrapped in Python through `scclr`, so shifted CLR can be used from standard Python single-cell workflows without giving up the memory advantages of sparse storage.

#### References


Ahlmann-Eltze, C. and W. Huber (2023). "Comparison of transformations for single-cell RNA-seq data". In: *Nature Methods* 20.5, pp. 665–672. DOI: 10.1038/s41592-023-01814-1.

Aitchison, J. (Jan. 1982). "The statistical analysis of compositional data". In: *Journal of the Royal Statistical Society Series B: Statistical Methodology* 44.2, pp. 139–160. DOI: 10.1111/j.2517-6161.1982.tb01195.x.

Angelidis, I., L. M. Simon, I. E. Fernandez, M. Strunz, C. H. Mayr, F. R. Greiffo, G. Tsitsiridis, M. Ansari, E. Graf, T.-M. Strom, et al. (2019). "An atlas of the aging lung mapped by single cell transcriptomics and deep tissue proteomics". In: *Nature Communications* 10.1, p. 963. DOI: 10.1038/s41467-019-08831-9.

Anscombe, F. J. (1948). "The transformation of Poisson, binomial and negative-binomial data". In: *Biometrika* 35.3-4, pp. 246–254. DOI: 10.1093/biomet/35.3-4.246.

Booeshaghi, A. S. and L. Pachter (2021). "Normalization of single-cell RNA-seq counts by log(x + 1) or log(1 + x)". In: *Bioinformatics* 37.15, pp. 2223–2224. DOI: 10.1093/bioinformatics/btab085.

Borella, M., G. Martello, D. Risso, and C. Romualdi (2021). "PsiNorm: a scalable normalization for single-cell RNA-seq data". In: *Bioinformatics* 38.1, pp. 164–172. DOI: 10.1093/bioinformatics/btab641.

Choudhary, S. and R. Satija (2022). "Comparison and evaluation of statistical error models for scRNA-seq". In: *Genome Biology* 23.1, p. 27. DOI: 10.1186/s13059-021-02584-9.

Egozcue, J. J. and V. Pawlowsky-Glahn (July 2019). "Compositional data: the sample space and its structure". In: *TEST* 28.3, pp. 599–638. DOI: 10.1007/s11749-019-00670-6.

Grün, D., L. Kester, and A. van Oudenaarden (June 2014). "Validation of noise models for single-cell transcriptomics". In: *Nature Methods* 11.6, pp. 637–640. DOI: 10.1038/nmeth.2930.

Huang, J., S. C. P. Yam, K. S. Leung, M. Deng, and N. L. S. Tang (2025). "Compositional data modeling of high-dimensional single cell RNA-seq (CoDA-hd): its advantages over commonly used normalization approaches". In: *Journal of Translational Medicine* 23.1, p. 1143. DOI: 10.1186/s12967-025-07157-z.

Pawlowsky-Glahn, V. and J. J. Egozcue (Oct. 2001). "Geometric approach to statistical analysis on the simplex". In: *Stochastic Environmental Research and Risk Assessment* 15.5, pp. 384–398. DOI: 10.1007/s004770100077.

Rich, J. M., L. Moses, P. H. Einarsson, K. Jackson, L. Luebbert, A. S. Booeshaghi, S. Antonsson, D. K. Sullivan, N. Bray, P. Melsted, et al. (2026). "The impact of package selection and versioning on single-cell RNA-seq analysis". In: *Cell Systems* 17.4, p. 101560. DOI: 10.1016/j.cels.2026.101560.

satijalab/seurat (2019). *CLR Normalization and Scaling.* https://github.com/satijalab/seurat/issues/1268. GitHub issue #1268.

— (2020). *Rationale for CLR Transformation in ADT Counts.* https://github.com/satijalab/seurat/issues/3550. GitHub issue #3550.

— (2023). *Order of Data Normalization for Layer Integration.* https://github.com/satijalab/seurat/issues/7585. GitHub issue #7585.

Stoeckius, M., C. Hafemeister, W. Stephenson, B. Houck-Loomis, P. K. Chattopadhyay, H. Swerdlow, R. Satija, and P. Smibert (July 2017). "Simultaneous epitope and transcriptome measurement in single cells". In: *Nature Methods* 14.9, pp. 865–868. DOI: 10.1038/nmeth.4380.
