## Supplementary Figures for "Normalization for sampled count data"

### Supplementary Figures: per-dataset method comparisons

Booeshaghi et al.

526 datasets, numbered Supplementary Figure 1.1–1.526. Use the PDF bookmarks (outline) to navigate by accession.

| 10xv3_hek_293t | |
|---|---|
| cells | 2,378 |
| genes | 57,905 |
| nonzero entries | 13,167,099 |
| density | 0.0956 |
| total counts | 79,525,261 |
| mean counts / cell | 33,442 |
| mean counts / gene | 1,373 |
| min counts / cell | 2,083 |
| max counts / cell | 98,397 |
| overdispersion | 0.136 |

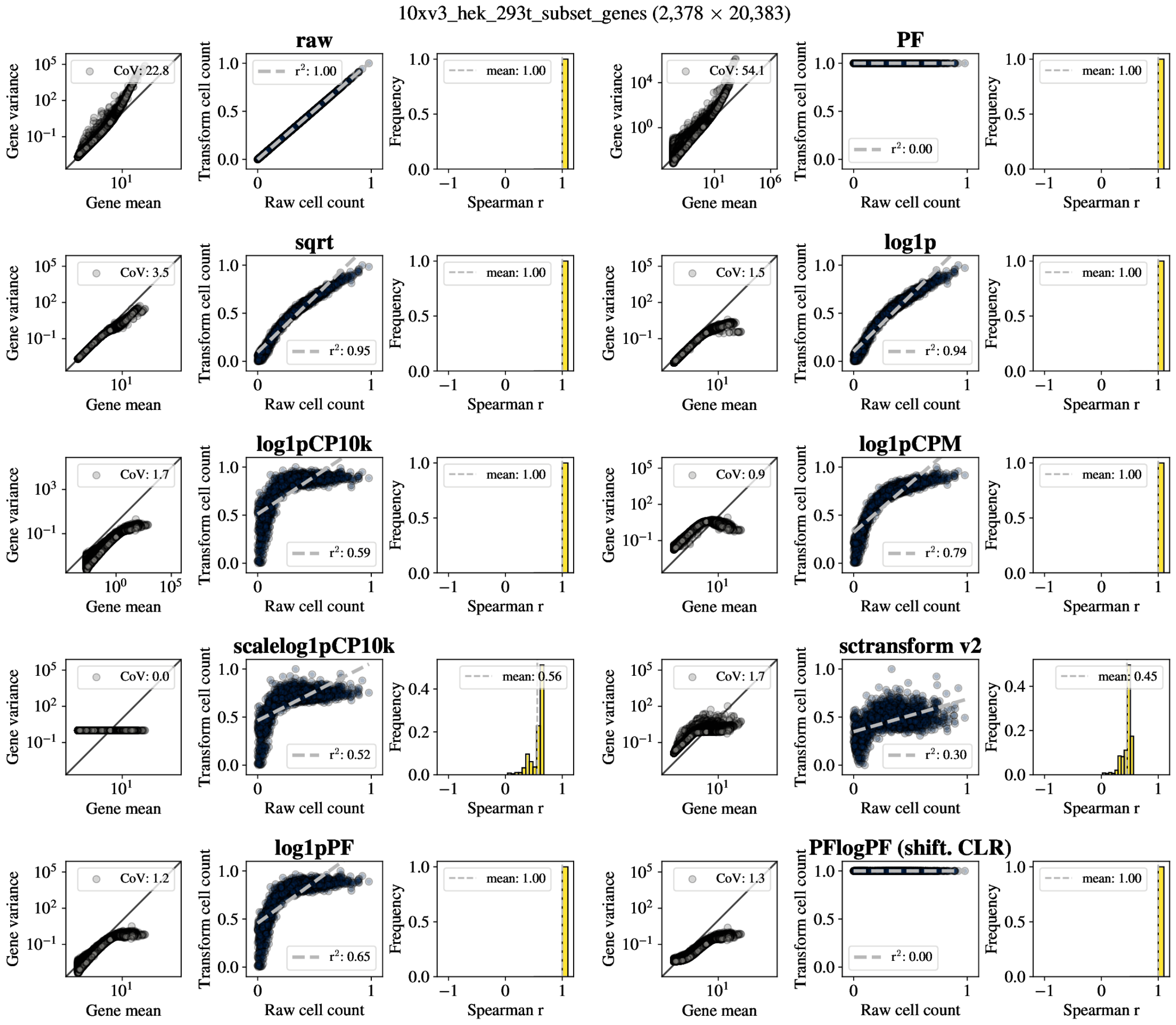


Supplementary Figure 1.1. **10xv3_hek_293t**. Per-method comparison of variance stabilization (left), depth normalization (middle), and monotonicity (right) for each normalization on dataset 10xv3_hek_293t.

| **10xv3_nih_3t3** | |
|---|---|
| cells | 2,458 |
| genes | 54,232 |
| nonzero entries | 9,731,396 |
| density | 0.073 |
| total counts | 39,781,517 |
| mean counts / cell | 16,185 |
| mean counts / gene | 734 |
| min counts / cell | 2,345 |
| max counts / cell | 70,115 |
| overdispersion | 0.24 |

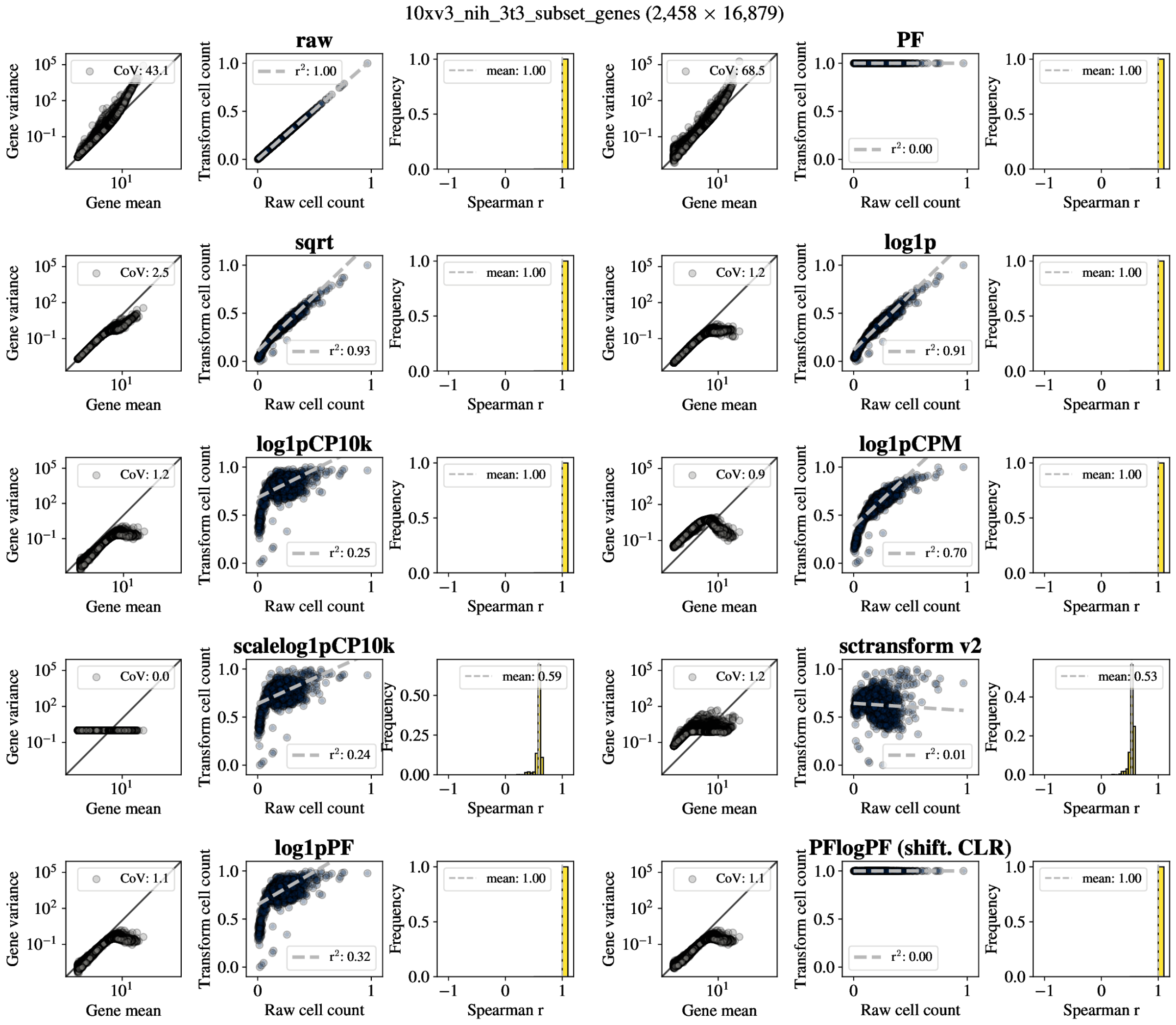


Supplementary Figure 1.2. **10xv3_nih_3t3**. Per-method comparison of variance stabilization (left), depth normalization (middle), and monotonicity (right) for each normalization on dataset 10xv3_nih_3t3.

| **10xv3_pbmc** | |
|---|---|
| cells | 1,222 |
| genes | 33,538 |
| nonzero entries | 2,498,225 |
| density | 0.061 |
| total counts | 9,235,144 |
| mean counts / cell | 7,557 |
| mean counts / gene | 275 |
| min counts / cell | 503 |
| max counts / cell | 56,884 |
| overdispersion | 0.619 |

10xv3_pbmc_subset_genes (1,222 × 14,043)

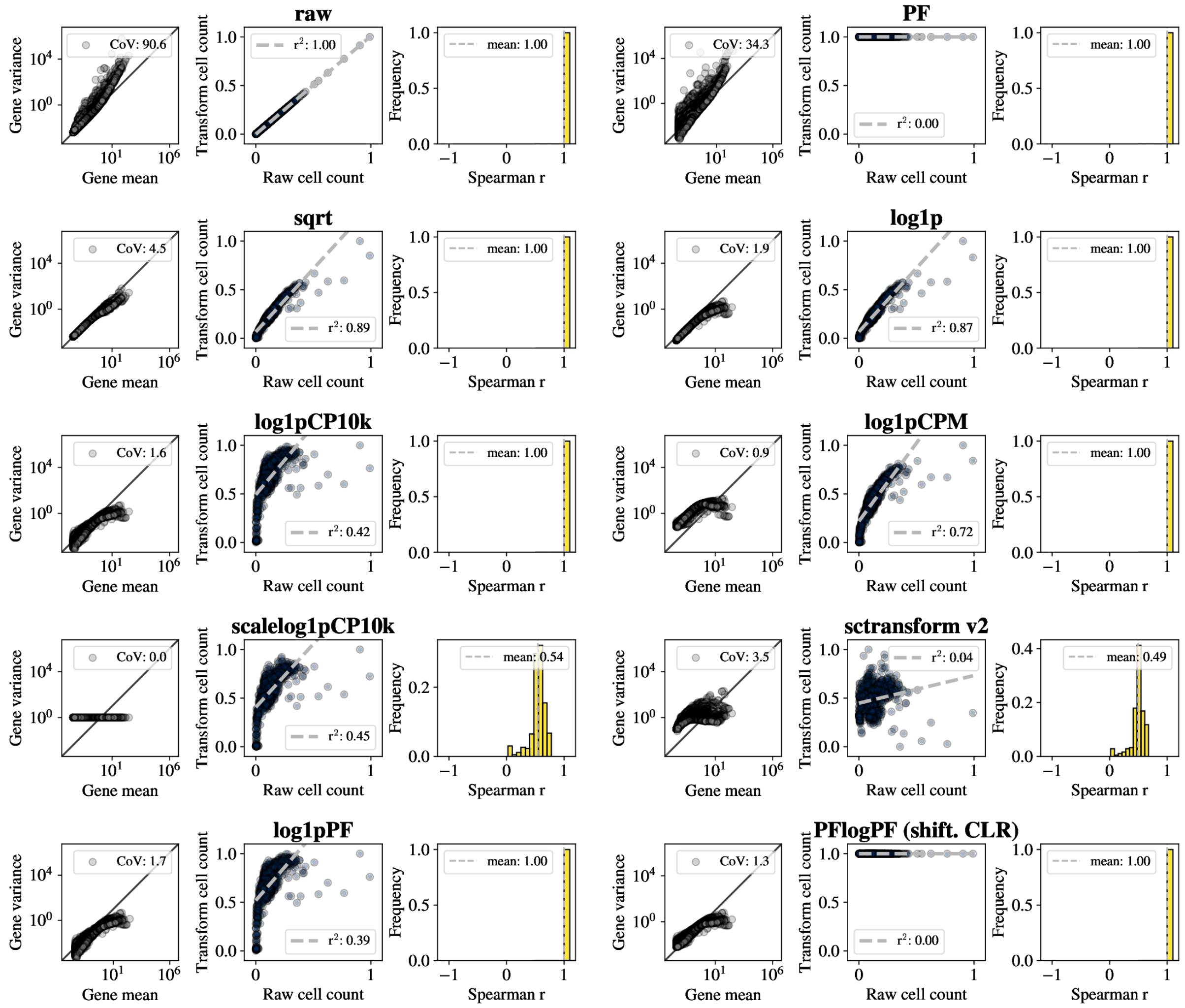


Supplementary Figure 1.3. **10xv3_pbmc**. Per-method comparison of variance stabilization (left), depth normalization (middle), and monotonicity (right) for each normalization on dataset 10xv3_pbmc.

| **angelidis_2019** | |
|---|---|
| cells | 14,813 |
| genes | 21,969 |
| nonzero entries | 6,607,982 |
| density | 0.0203 |
| total counts | 12,132,443 |
| mean counts / cell | 819 |
| mean counts / gene | 552 |
| min counts / cell | 209 |
| max counts / cell | 14,787 |
| overdispersion | 12.5 |

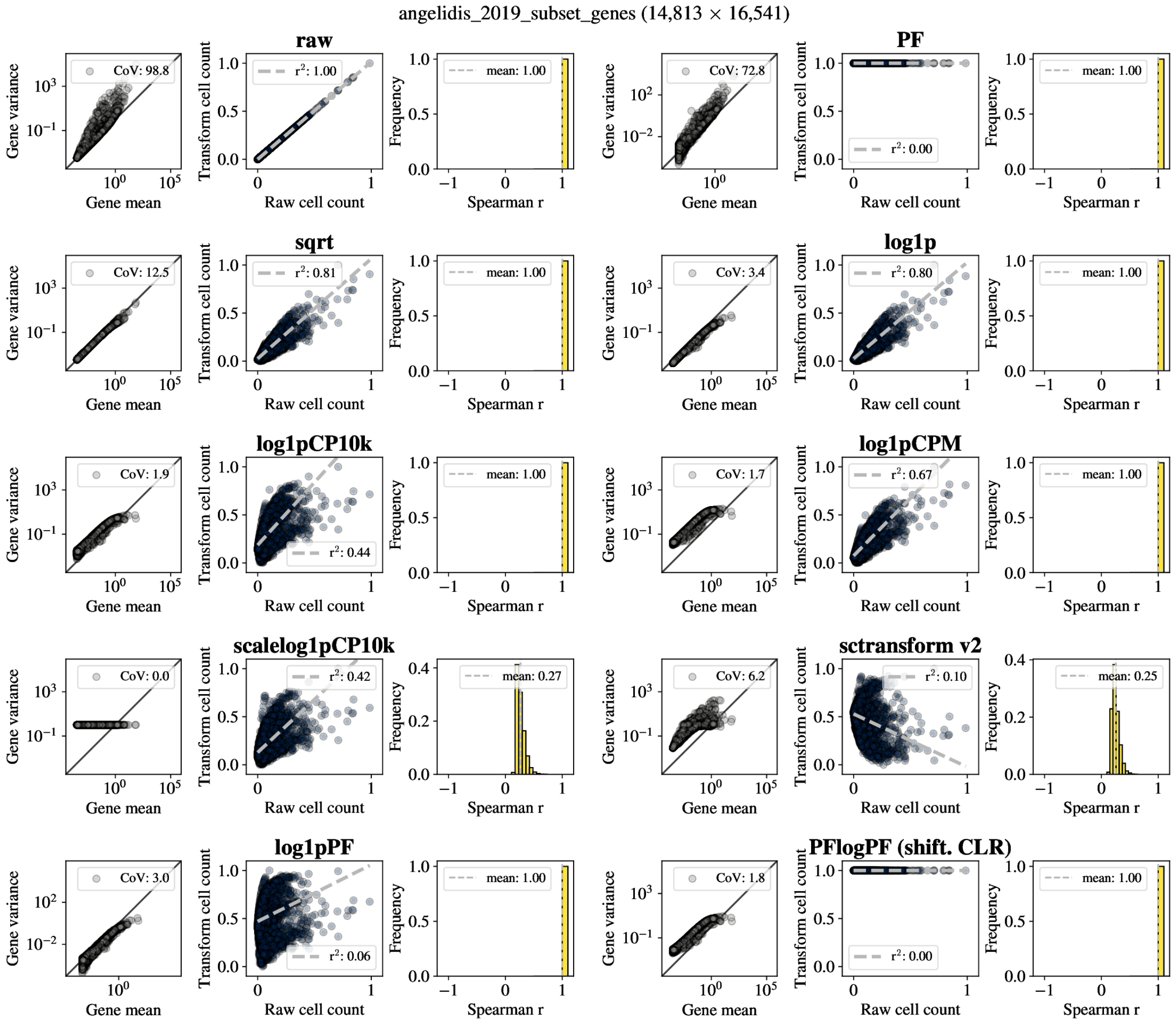


Supplementary Figure 1.4. **angelidis_2019**. Per-method comparison of variance stabilization (left), depth normalization (middle), and monotonicity (right) for each normalization on dataset angelidis_2019.

| **CRX102285** | |
|---|---|
| cells | 8,991 |
| genes | 60,623 |
| nonzero entries | 8,938,174 |
| density | 0.0164 |
| total counts | 24,363,912 |
| mean counts / cell | 2,710 |
| mean counts / gene | 402 |
| min counts / cell | 214 |
| max counts / cell | 41,356 |
| overdispersion | 1.76 |

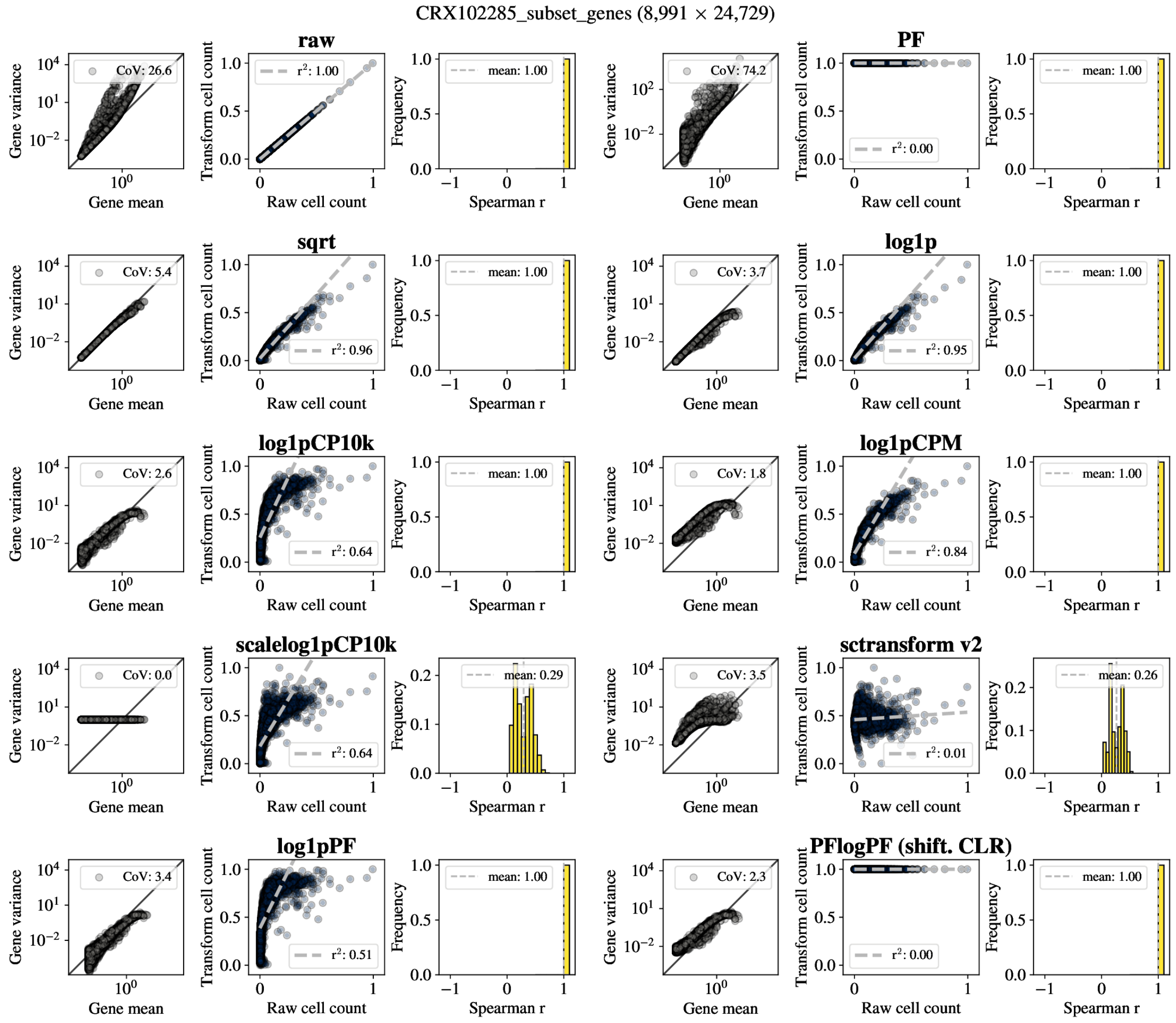


Supplementary Figure 1.5. **CRX102285**. Per-method comparison of variance stabilization (left), depth normalization (middle), and monotonicity (right) for each normalization on dataset CRX102285.

| CRX102286 | |
|---|---|
| cells | 70,820 |
| genes | 60,623 |
| nonzero entries | 18,408,410 |
| density | 0.00429 |
| total counts | 54,319,644 |
| mean counts / cell | 767 |
| mean counts / gene | 896 |
| min counts / cell | 24 |
| max counts / cell | 67,369 |
| overdispersion | 16.5 |

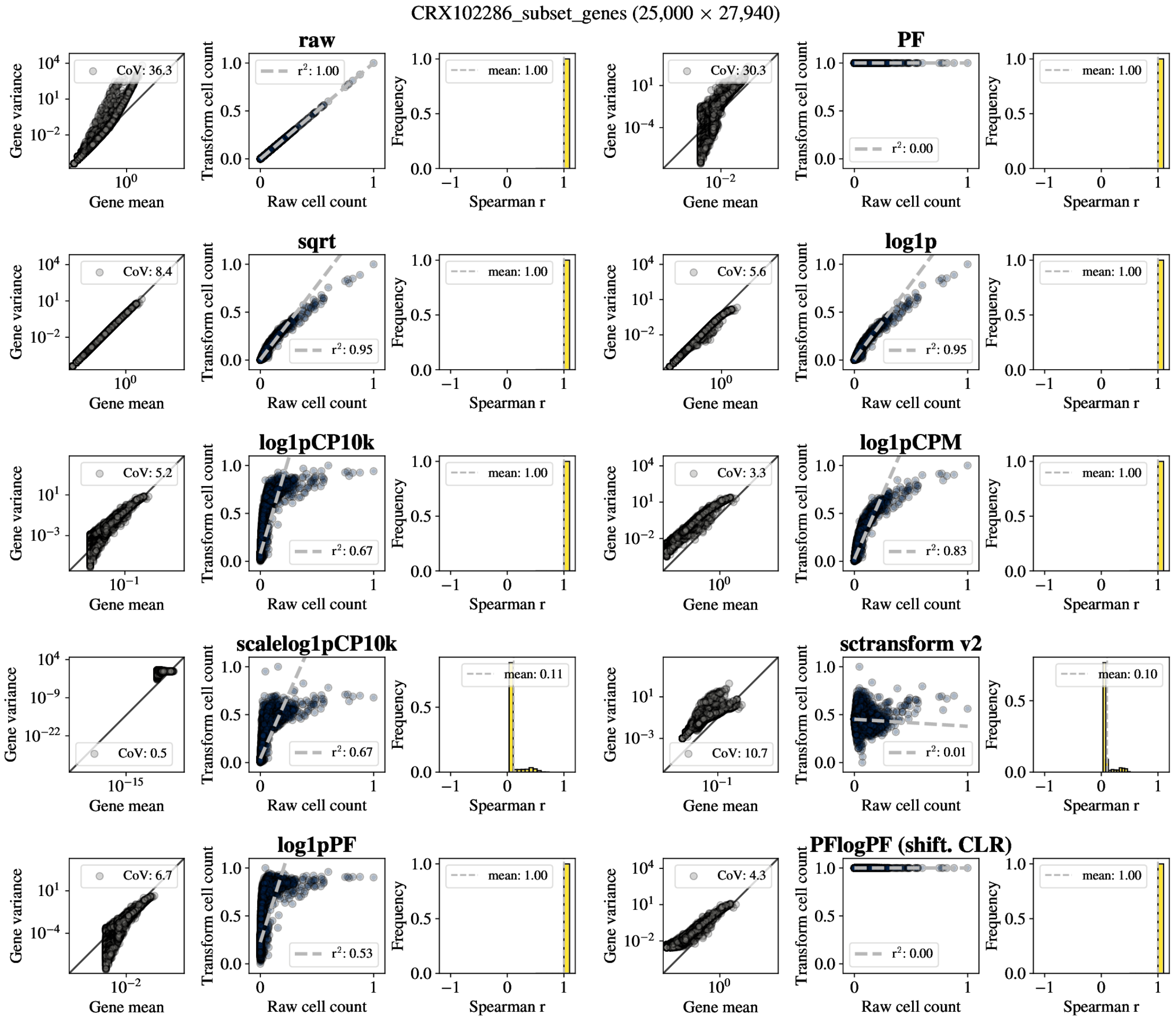


Supplementary Figure 1.6. **CRX102286**. Per-method comparison of variance stabilization (left), depth normalization (middle), and monotonicity (right) for each normalization on dataset CRX102286. Figure panels were computed on a random subsample of 25,000 of 70,820 cells to bound memory; the metrics table above reflects the full dataset.

| **CRX102287** | |
|---|---|
| cells | 10,785 |
| genes | 60,623 |
| nonzero entries | 14,273,310 |
| density | 0.0218 |
| total counts | 44,338,592 |
| mean counts / cell | 4,111 |
| mean counts / gene | 731 |
| min counts / cell | 182 |
| max counts / cell | 62,634 |
| overdispersion | 2.61 |

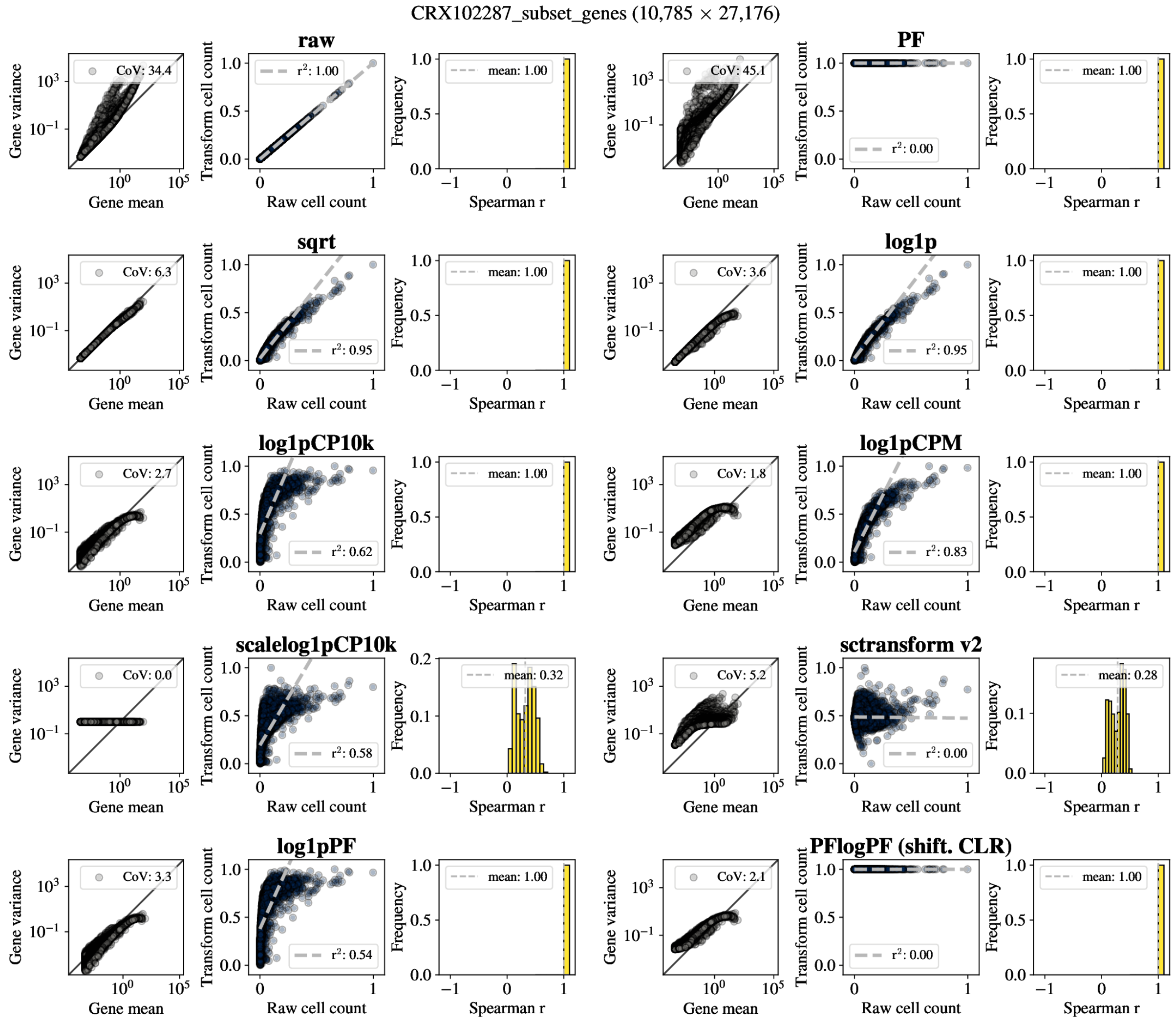


Supplementary Figure 1.7. **CRX102287**. Per-method comparison of variance stabilization (left), depth normalization (middle), and monotonicity (right) for each normalization on dataset CRX102287.

| CRX102288 | |
|---|---|
| cells | 12,531 |
| genes | 60,623 |
| nonzero entries | 19,302,558 |
| density | 0.0254 |
| total counts | 61,801,673 |
| mean counts / cell | 4,932 |
| mean counts / gene | 1,019 |
| min counts / cell | 205 |
| max counts / cell | 50,011 |
| overdispersion | 2.02 |

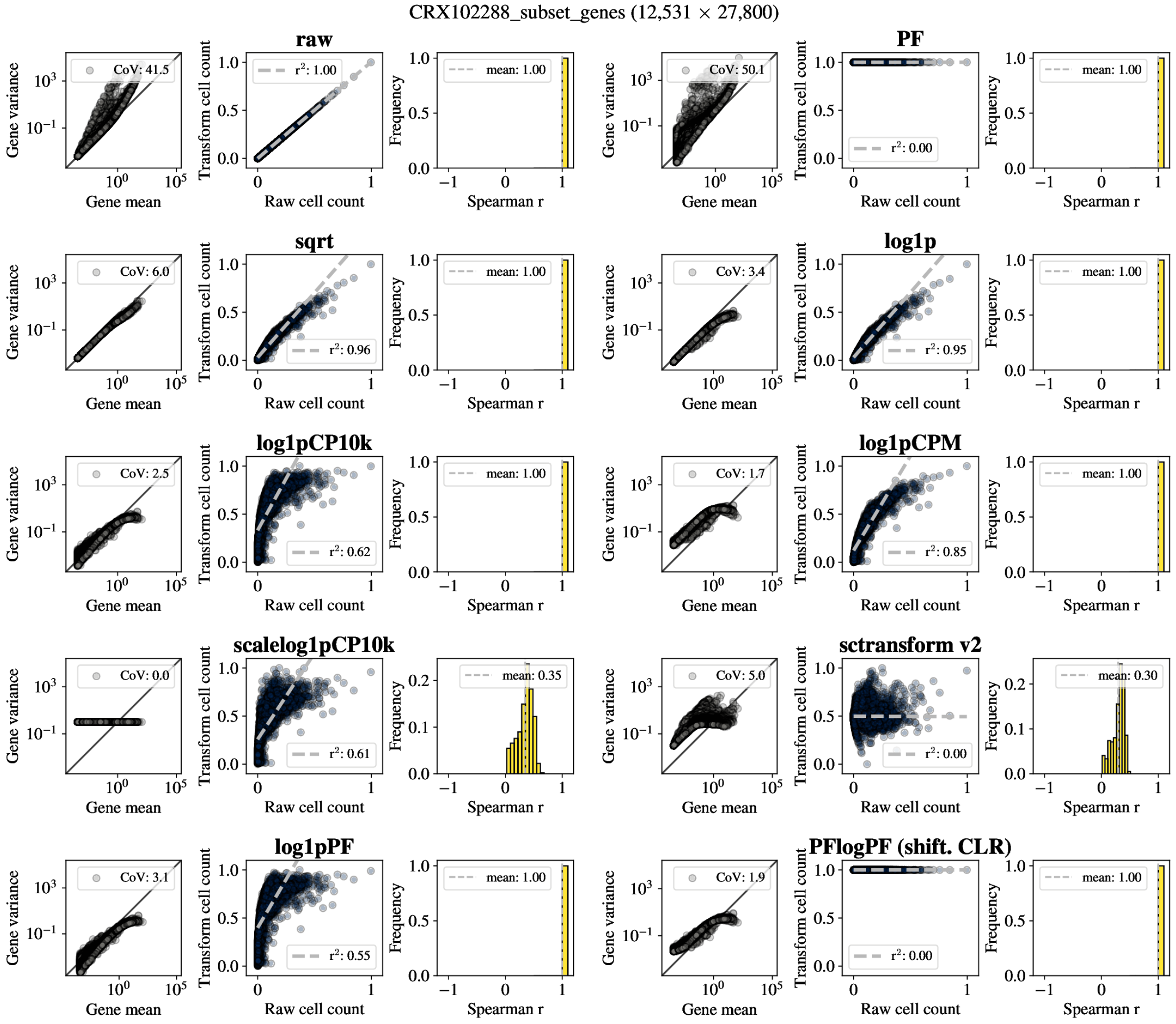


Supplementary Figure 1.8. **CRX102288**. Per-method comparison of variance stabilization (left), depth normalization (middle), and monotonicity (right) for each normalization on dataset CRX102288.

| CRX102289 | |
|---|---|
| cells | 9,200 |
| genes | 60,623 |
| nonzero entries | 11,280,911 |
| density | 0.0202 |
| total counts | 32,344,833 |
| mean counts / cell | 3,516 |
| mean counts / gene | 534 |
| min counts / cell | 154 |
| max counts / cell | 44,737 |
| overdispersion | 2.61 |

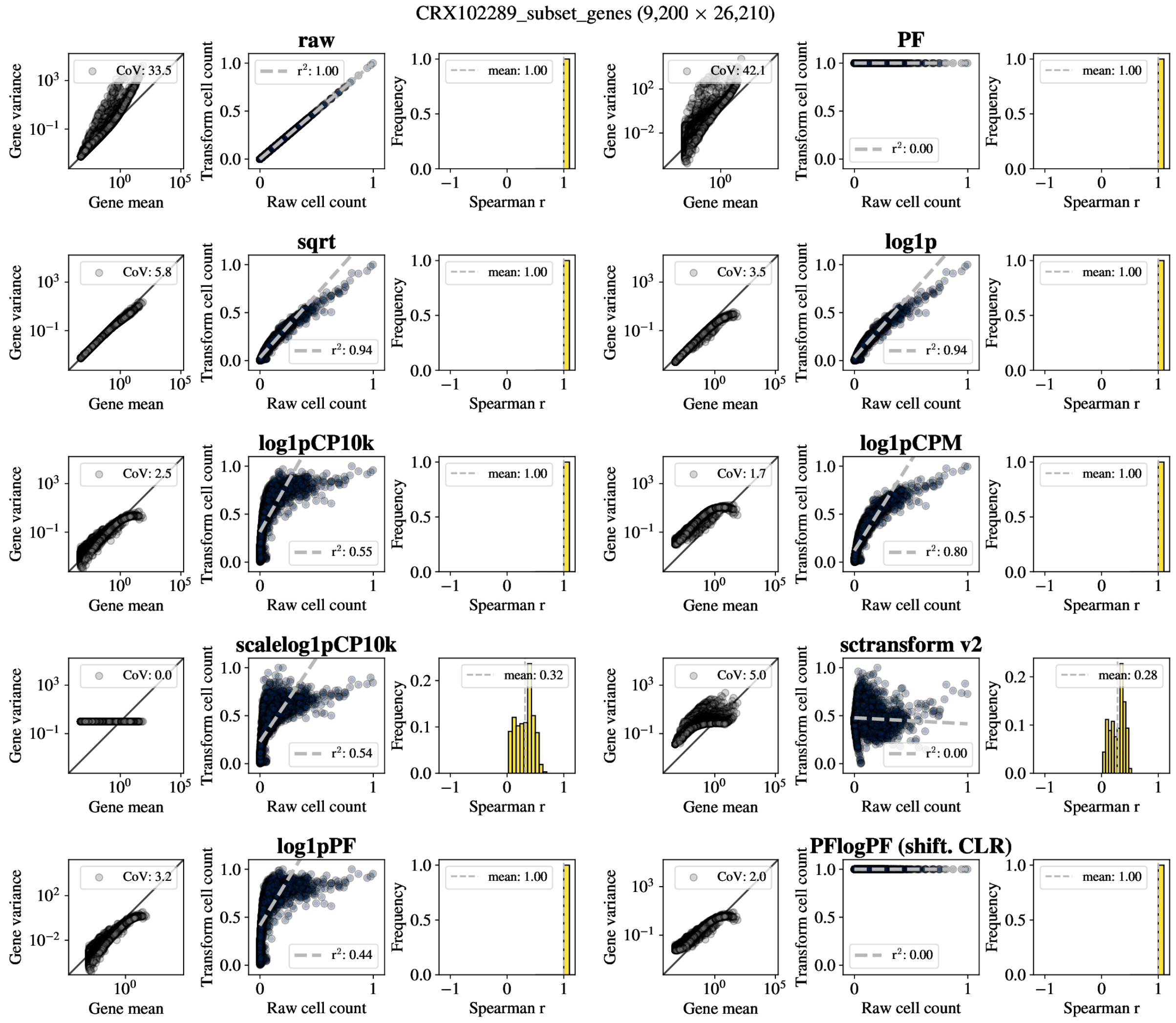


Supplementary Figure 1.9. **CRX102289**. Per-method comparison of variance stabilization (left), depth normalization (middle), and monotonicity (right) for each normalization on dataset CRX102289.

| CRX102290 | |
|---|---|
| cells | 7,632 |
| genes | 60,623 |
| nonzero entries | 5,008,777 |
| density | 0.0108 |
| total counts | 13,404,917 |
| mean counts / cell | 1,756 |
| mean counts / gene | 221 |
| min counts / cell | 117 |
| max counts / cell | 42,486 |
| overdispersion | 1.94 |

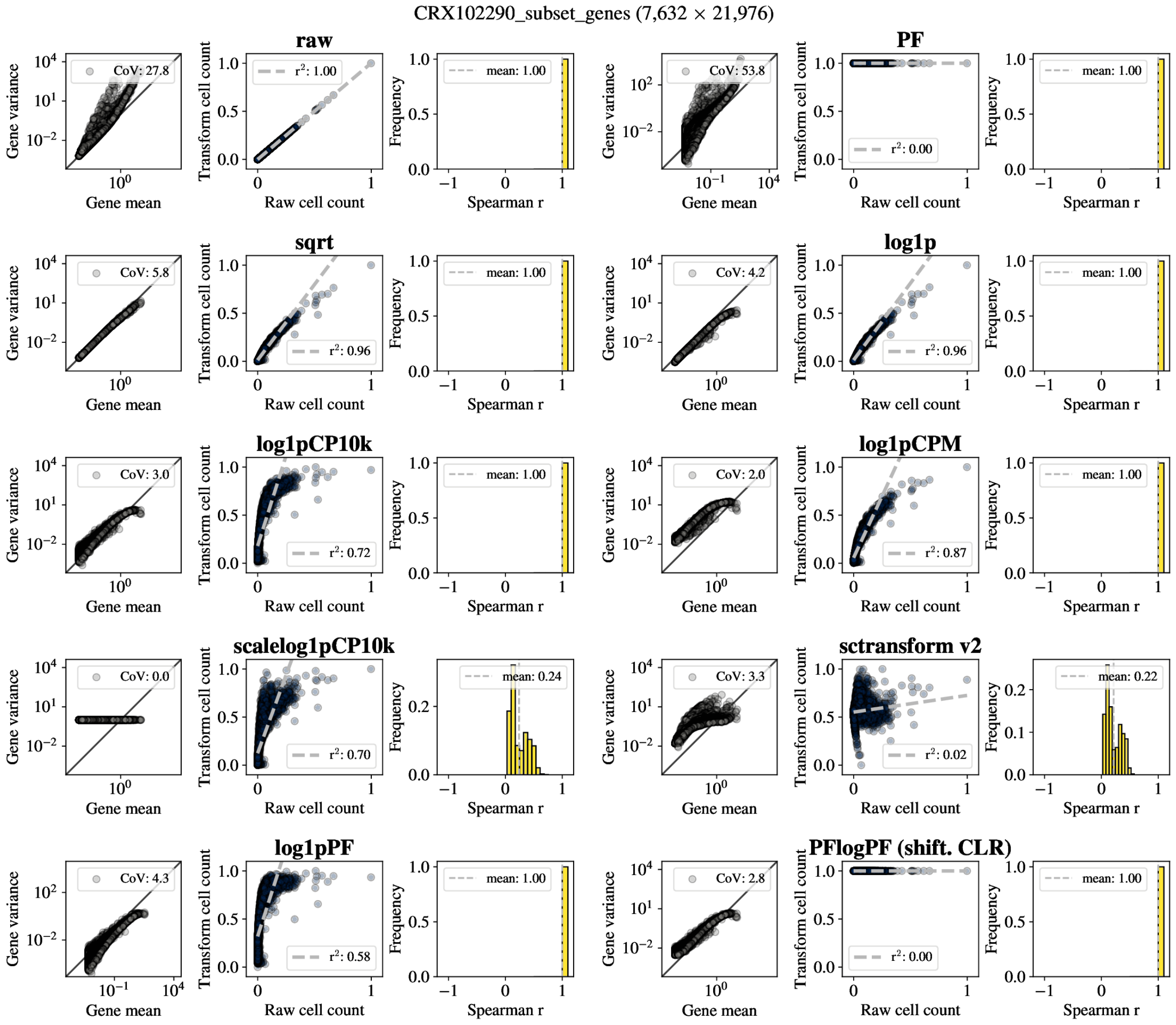


Supplementary Figure 1.10. **CRX102290**. Per-method comparison of variance stabilization (left), depth normalization (middle), and monotonicity (right) for each normalization on dataset CRX102290.

| CRX102291 | |
|---|---|
| cells | 3,963 |
| genes | 28,239 |
| nonzero entries | 7,735,368 |
| density | 0.0691 |
| total counts | 23,579,386 |
| mean counts / cell | 5,950 |
| mean counts / gene | 835 |
| min counts / cell | 3,337 |
| max counts / cell | 57,254 |
| overdispersion | 0.52 |

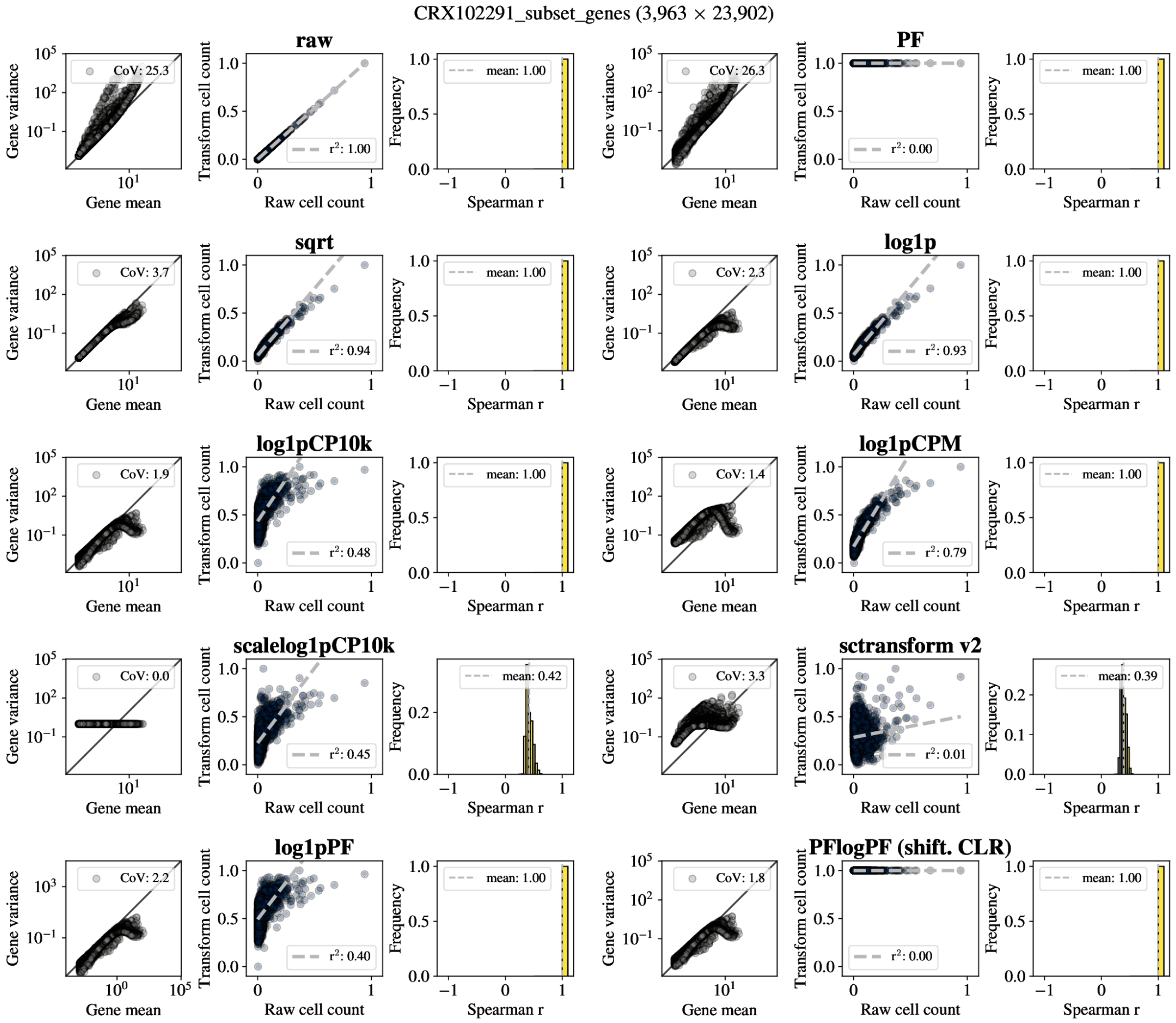


Supplementary Figure 1.11. **CRX102291**. Per-method comparison of variance stabilization (left), depth normalization (middle), and monotonicity (right) for each normalization on dataset CRX102291.

| **CRX102292** | |
|---|---|
| cells | 85,238 |
| genes | 60,623 |
| nonzero entries | 14,227,453 |
| density | 0.00275 |
| total counts | 38,639,910 |
| mean counts / cell | 453 |
| mean counts / gene | 637 |
| min counts / cell | 8 |
| max counts / cell | 57,514 |
| overdispersion | 15.1 |

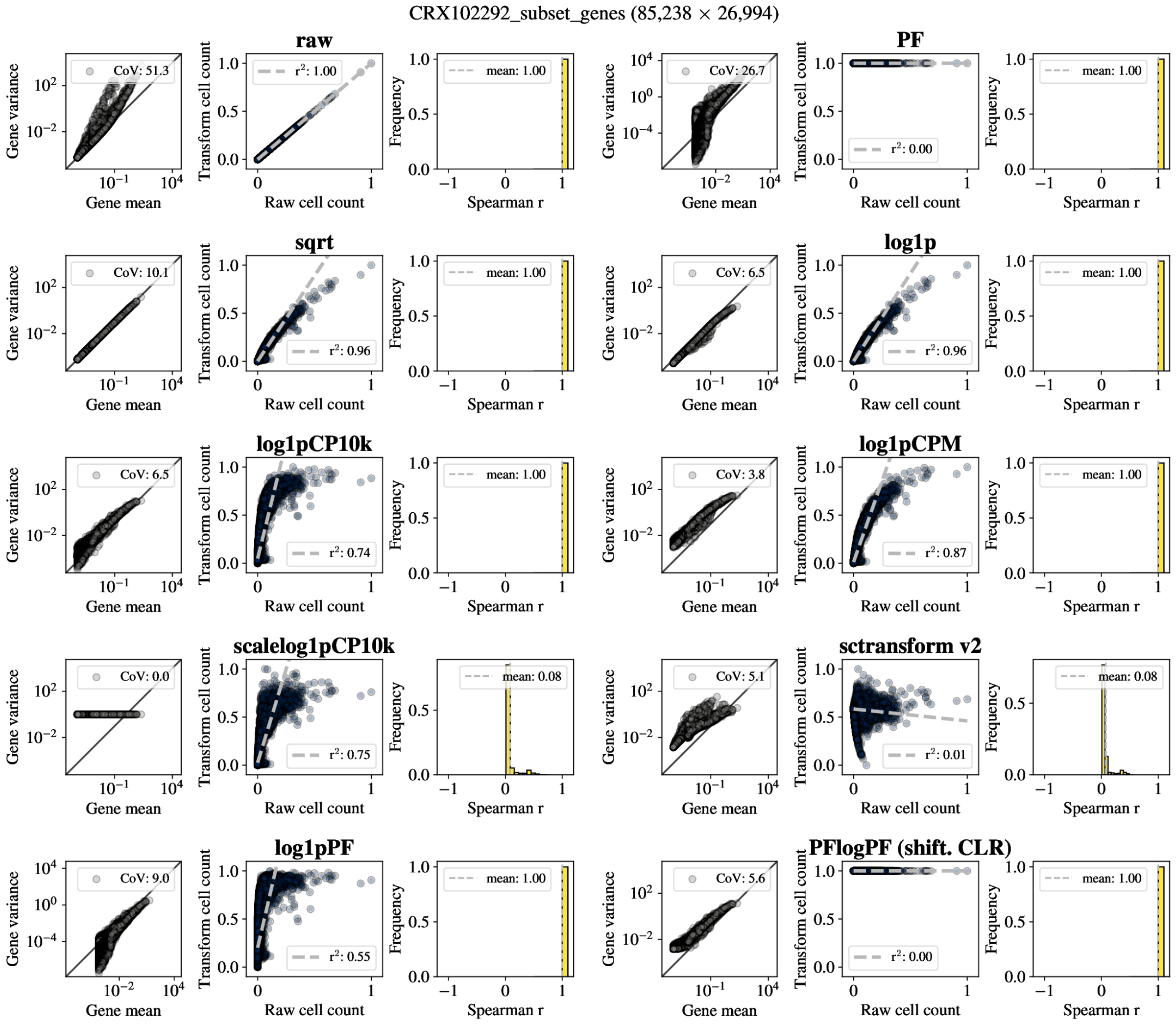


Supplementary Figure 1.12. **CRX102292**. Per-method comparison of variance stabilization (left), depth normalization (middle), and monotonicity (right) for each normalization on dataset CRX102292.

| CRX102293 | |
|---|---|
| cells | 12,626 |
| genes | 60,623 |
| nonzero entries | 12,231,616 |
| density | 0.016 |
| total counts | 32,099,058 |
| mean counts / cell | 2,542 |
| mean counts / gene | 529 |
| min counts / cell | 462 |
| max counts / cell | 42,831 |
| overdispersion | 1.5 |

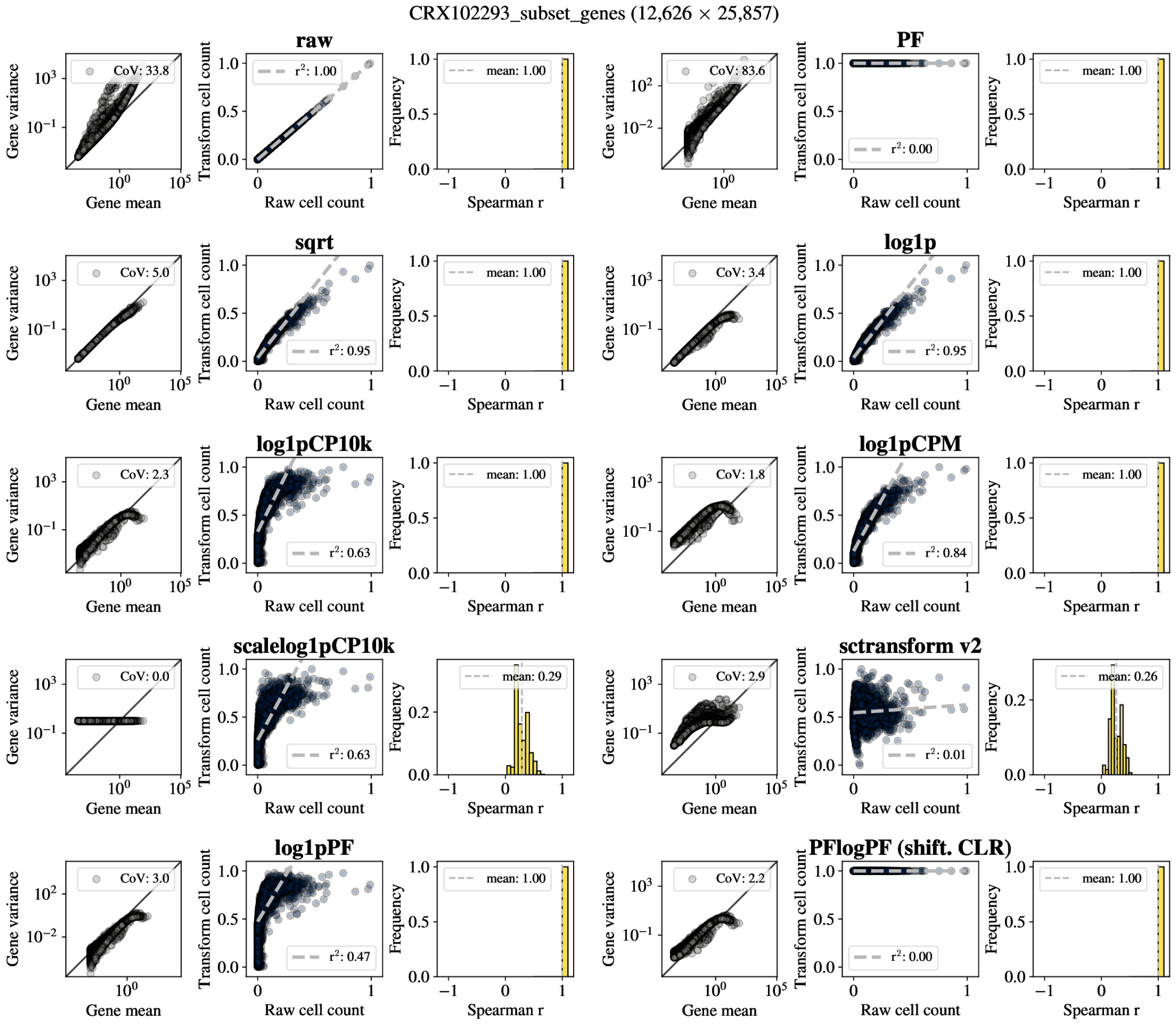


Supplementary Figure 1.13. **CRX102293**. Per-method comparison of variance stabilization (left), depth normalization (middle), and monotonicity (right) for each normalization on dataset CRX102293.

| **CRX102294** | |
|---|---|
| cells | 10,577 |
| genes | 60,623 |
| nonzero entries | 13,971,738 |
| density | 0.0218 |
| total counts | 43,669,559 |
| mean counts / cell | 4,129 |
| mean counts / gene | 720 |
| min counts / cell | 187 |
| max counts / cell | 56,598 |
| overdispersion | 2.45 |

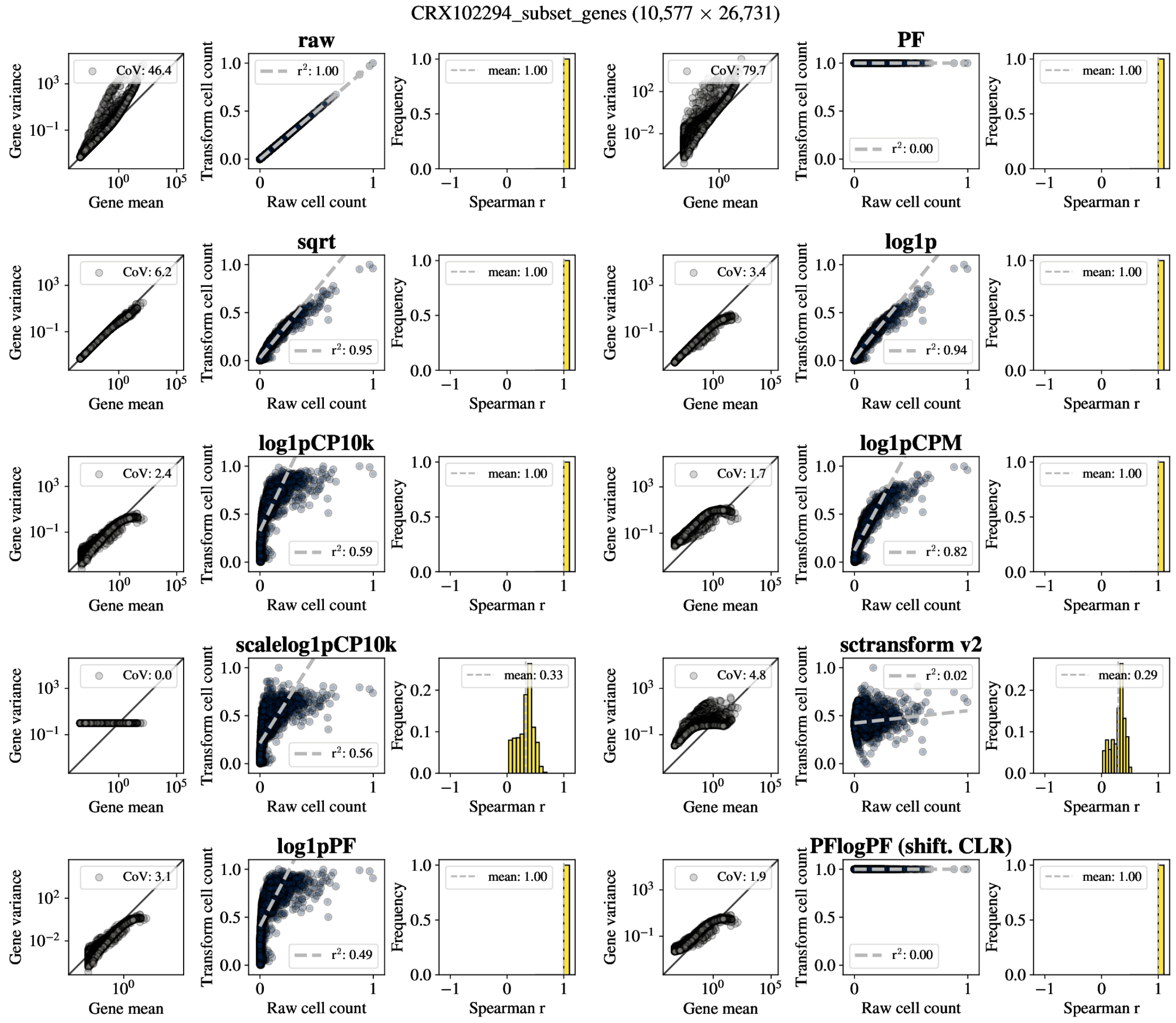


Supplementary Figure 1.14. **CRX102294**. Per-method comparison of variance stabilization (left), depth normalization (middle), and monotonicity (right) for each normalization on dataset CRX102294.

| **CRX118012** | |
|---|---|
| cells | 14,992 |
| genes | 60,623 |
| nonzero entries | 21,216,502 |
| density | 0.0233 |
| total counts | 58,988,833 |
| mean counts / cell | 3,935 |
| mean counts / gene | 973 |
| min counts / cell | 209 |
| max counts / cell | 44,131 |
| overdispersion | 1.65 |

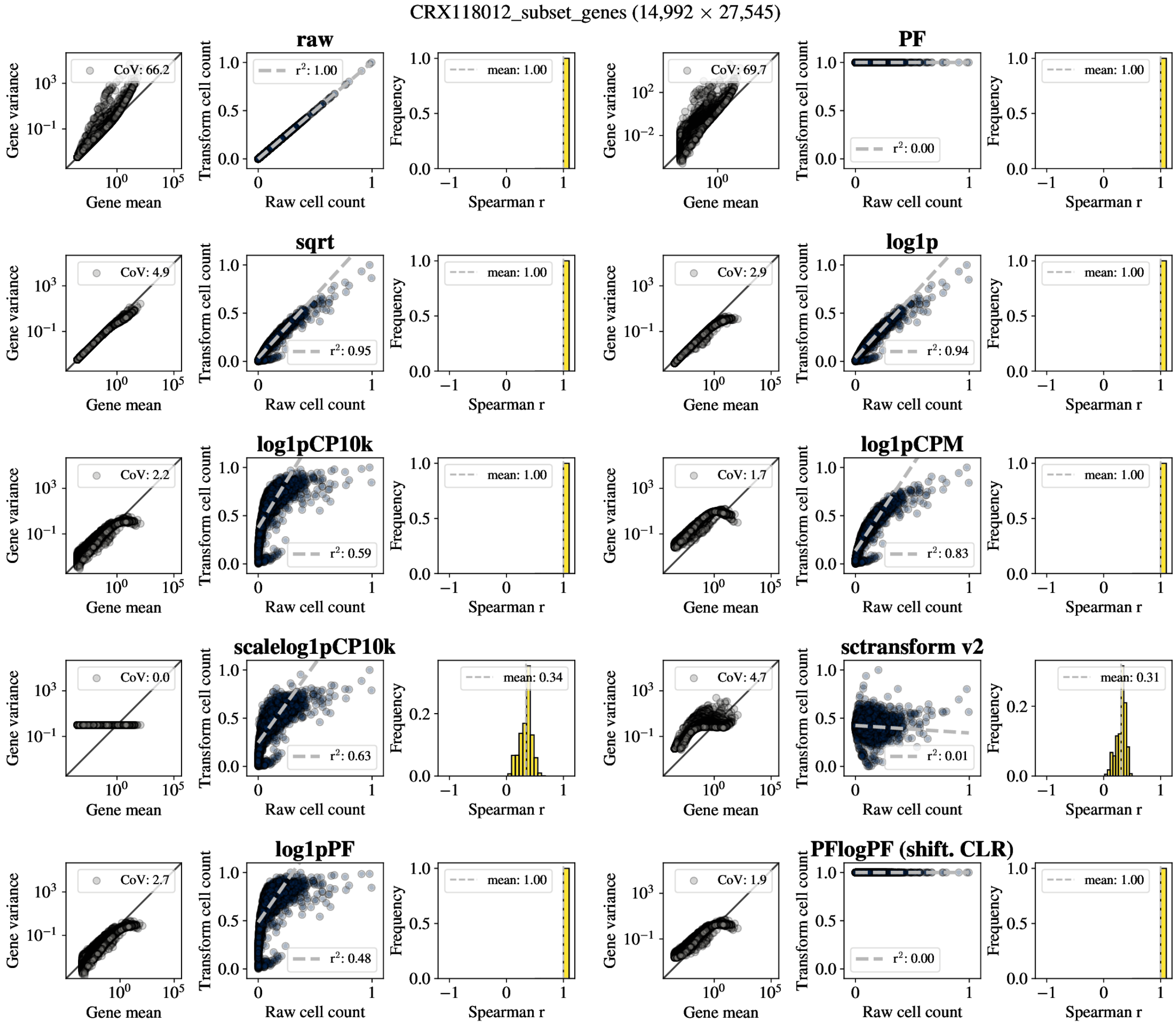


Supplementary Figure 1.15. **CRX118012**. Per-method comparison of variance stabilization (left), depth normalization (middle), and monotonicity (right) for each normalization on dataset CRX118012.

| **CRX118013** | |
|---|---|
| cells | 10,963 |
| genes | 60,623 |
| nonzero entries | 14,759,943 |
| density | 0.0222 |
| total counts | 41,549,803 |
| mean counts / cell | 3,790 |
| mean counts / gene | 685 |
| min counts / cell | 328 |
| max counts / cell | 44,874 |
| overdispersion | 1.66 |

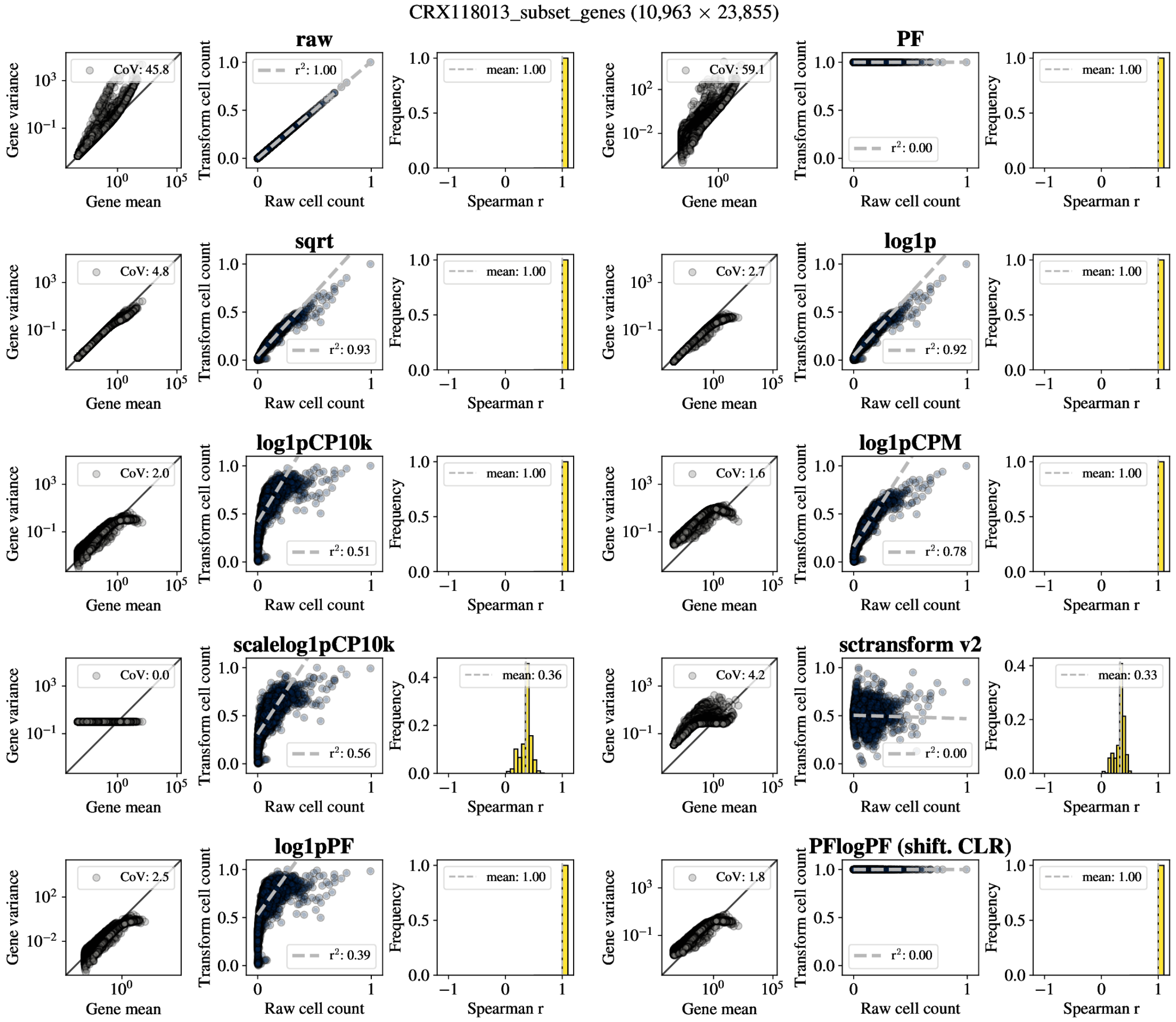


Supplementary Figure 1.16. **CRX118013**. Per-method comparison of variance stabilization (left), depth normalization (middle), and monotonicity (right) for each normalization on dataset CRX118013.

| **CRX118014** | |
|---|---|
| cells | 16,881 |
| genes | 60,623 |
| nonzero entries | 18,287,785 |
| density | 0.0179 |
| total counts | 51,391,489 |
| mean counts / cell | 3,044 |
| mean counts / gene | 848 |
| min counts / cell | 158 |
| max counts / cell | 28,410 |
| overdispersion | 1.83 |

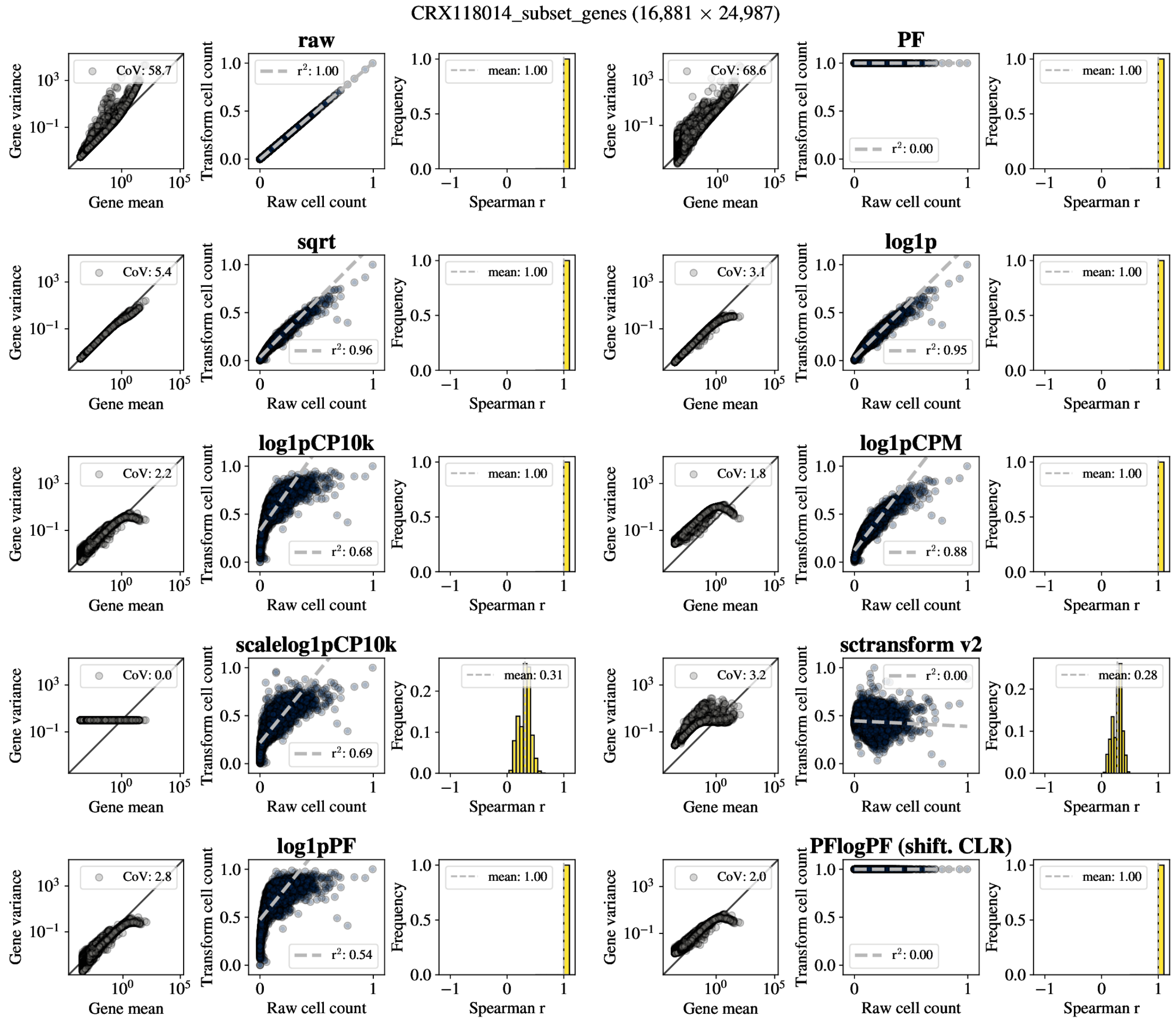


Supplementary Figure 1.17. **CRX118014**. Per-method comparison of variance stabilization (left), depth normalization (middle), and monotonicity (right) for each normalization on dataset CRX118014.

| CRX118015 | |
|---|---|
| cells | 9,789 |
| genes | 60,623 |
| nonzero entries | 12,572,513 |
| density | 0.0212 |
| total counts | 36,679,830 |
| mean counts / cell | 3,747 |
| mean counts / gene | 605 |
| min counts / cell | 249 |
| max counts / cell | 30,797 |
| overdispersion | 1.6 |

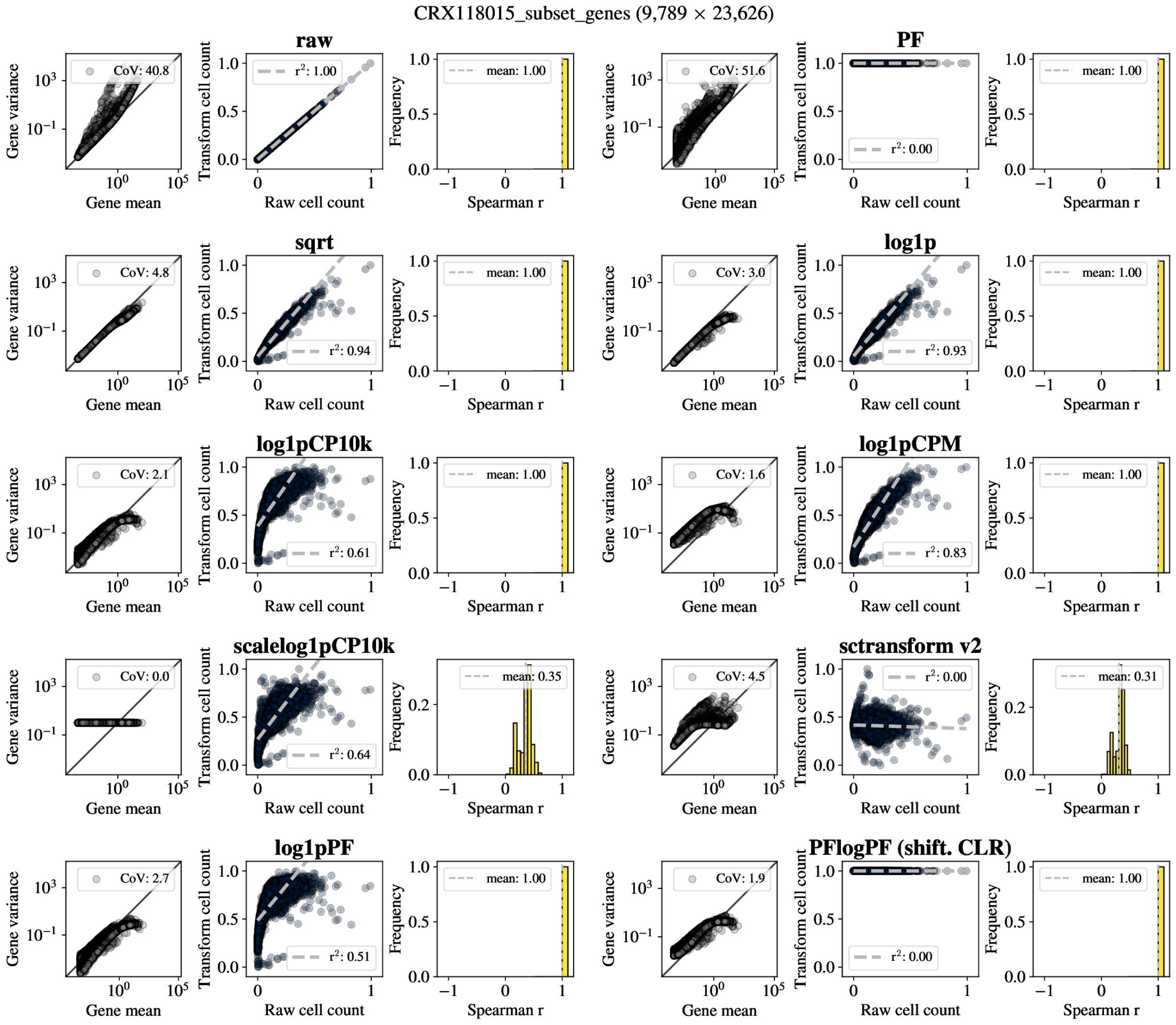


Supplementary Figure 1.18. **CRX118015**. Per-method comparison of variance stabilization (left), depth normalization (middle), and monotonicity (right) for each normalization on dataset CRX118015.

| ERS3861771 | |
|---|---|
| cells | 1,108 |
| genes | 60,623 |
| nonzero entries | 4,619,357 |
| density | 0.0688 |
| total counts | 15,392,592 |
| mean counts / cell | 13,892 |
| mean counts / gene | 254 |
| min counts / cell | 2,892 |
| max counts / cell | 119,815 |
| overdispersion | 0.442 |

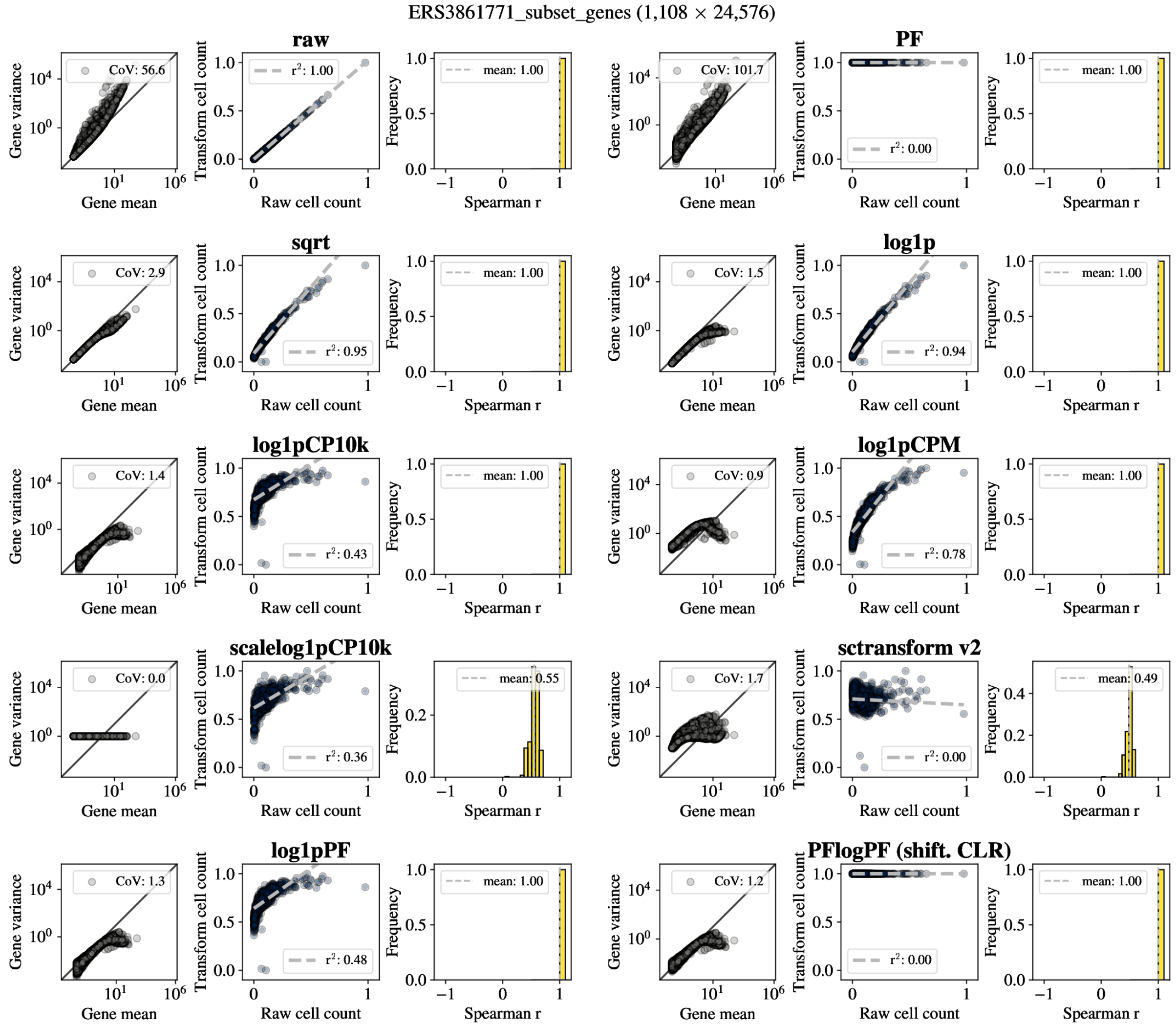


Supplementary Figure 1.19. **ERS3861771**. Per-method comparison of variance stabilization (left), depth normalization (middle), and monotonicity (right) for each normalization on dataset ERS3861771.

| **ERS3861772** | |
|---|---|
| cells | 850 |
| genes | 60,623 |
| nonzero entries | 2,755,351 |
| density | 0.0535 |
| total counts | 9,620,644 |
| mean counts / cell | 11,318 |
| mean counts / gene | 159 |
| min counts / cell | 1,988 |
| max counts / cell | 49,473 |
| overdispersion | 0.573 |

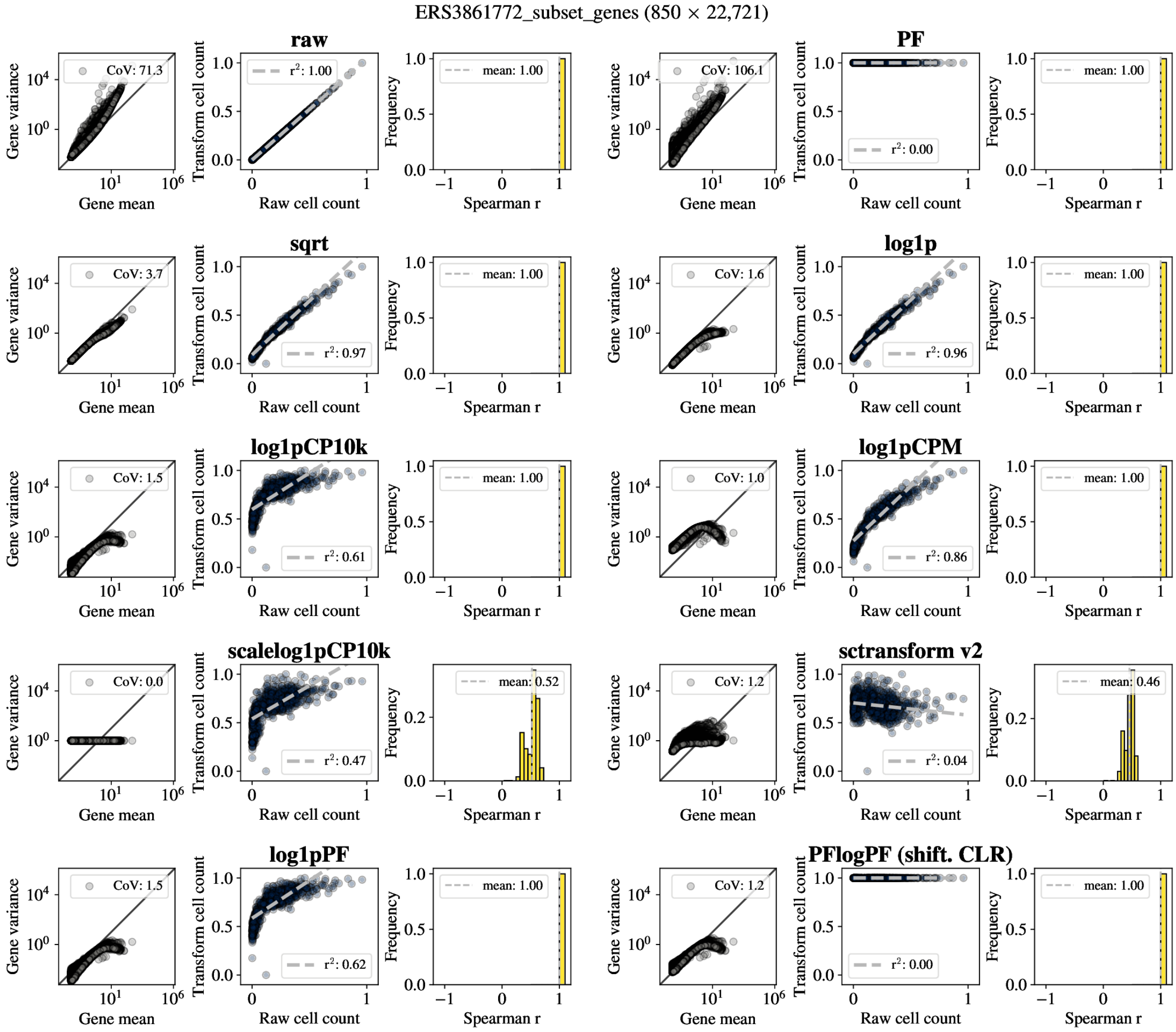


Supplementary Figure 1.20. **ERS3861772**. Per-method comparison of variance stabilization (left), depth normalization (middle), and monotonicity (right) for each normalization on dataset ERS3861772.

| **ERS3861773** | |
|---|---|
| cells | 15,628 |
| genes | 60,623 |
| nonzero entries | 1,204,660 |
| density | 0.00127 |
| total counts | 3,571,748 |
| mean counts / cell | 229 |
| mean counts / gene | 59 |
| min counts / cell | 41 |
| max counts / cell | 58,977 |
| overdispersion | 216 |

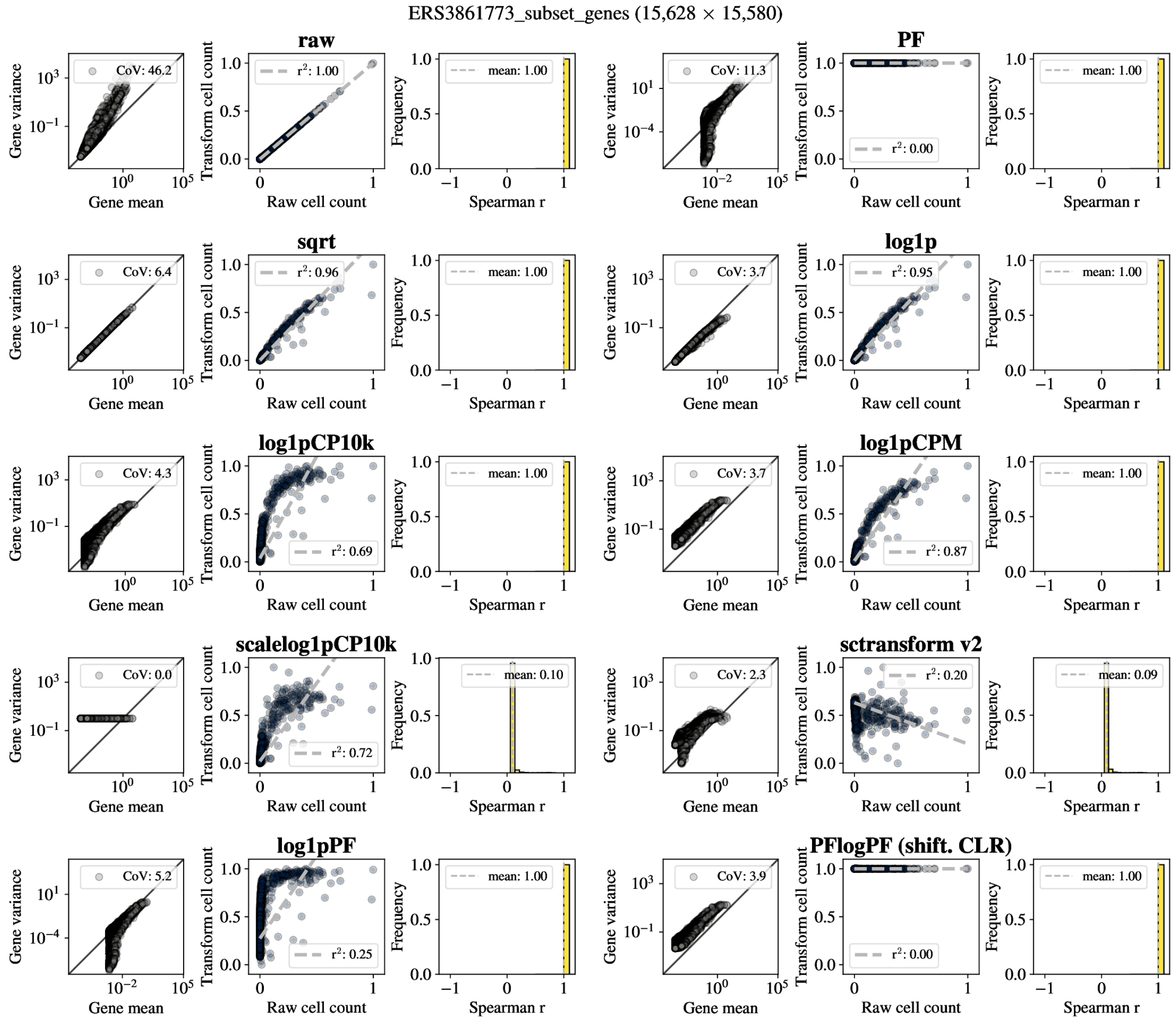


Supplementary Figure 1.21. **ERS3861773**. Per-method comparison of variance stabilization (left), depth normalization (middle), and monotonicity (right) for each normalization on dataset ERS3861773.

| **ERS3861774** | |
|---|---|
| cells | 2,075 |
| genes | 60,623 |
| nonzero entries | 6,287,592 |
| density | 0.05 |
| total counts | 43,692,297 |
| mean counts / cell | 21,057 |
| mean counts / gene | 721 |
| min counts / cell | 613 |
| max counts / cell | 98,825 |
| overdispersion | 2.02 |

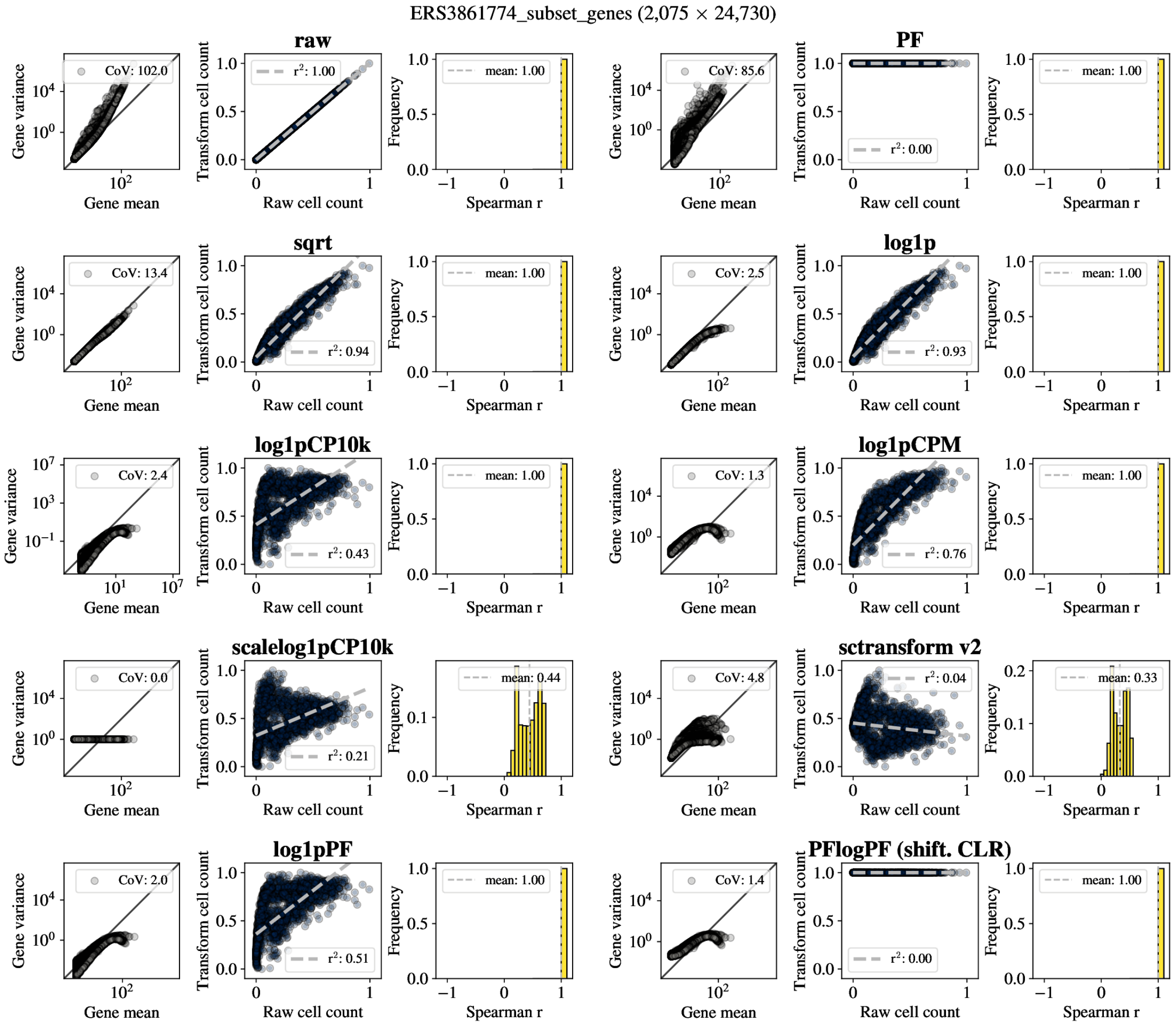


Supplementary Figure 1.22. **ERS3861774**. Per-method comparison of variance stabilization (left), depth normalization (middle), and monotonicity (right) for each normalization on dataset ERS3861774.

| ERS3861775 | |
|---|---|
| cells | 9,683 |
| genes | 60,623 |
| nonzero entries | 2,757,090 |
| density | 0.0047 |
| total counts | 9,752,907 |
| mean counts / cell | 1,007 |
| mean counts / gene | 161 |
| min counts / cell | 78 |
| max counts / cell | 97,238 |
| overdispersion | 23 |

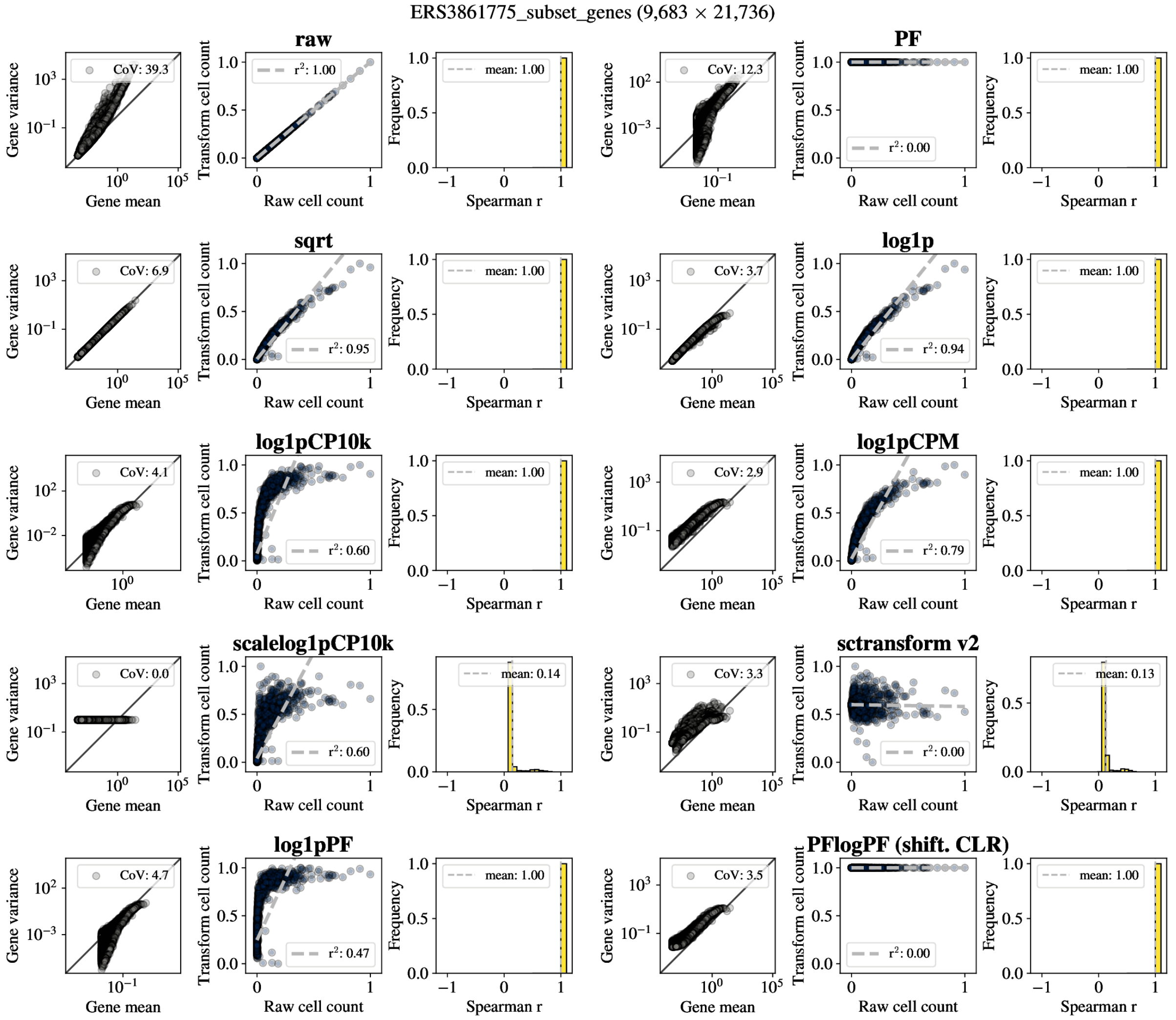


Supplementary Figure 1.23. **ERS3861775**. Per-method comparison of variance stabilization (left), depth normalization (middle), and monotonicity (right) for each normalization on dataset ERS3861775.

| ERS3861776 | |
|---|---|
| cells | 12,126 |
| genes | 60,623 |
| nonzero entries | 2,590,316 |
| density | 0.00352 |
| total counts | 7,851,758 |
| mean counts / cell | 648 |
| mean counts / gene | 130 |
| min counts / cell | 92 |
| max counts / cell | 94,417 |
| overdispersion | 268 |

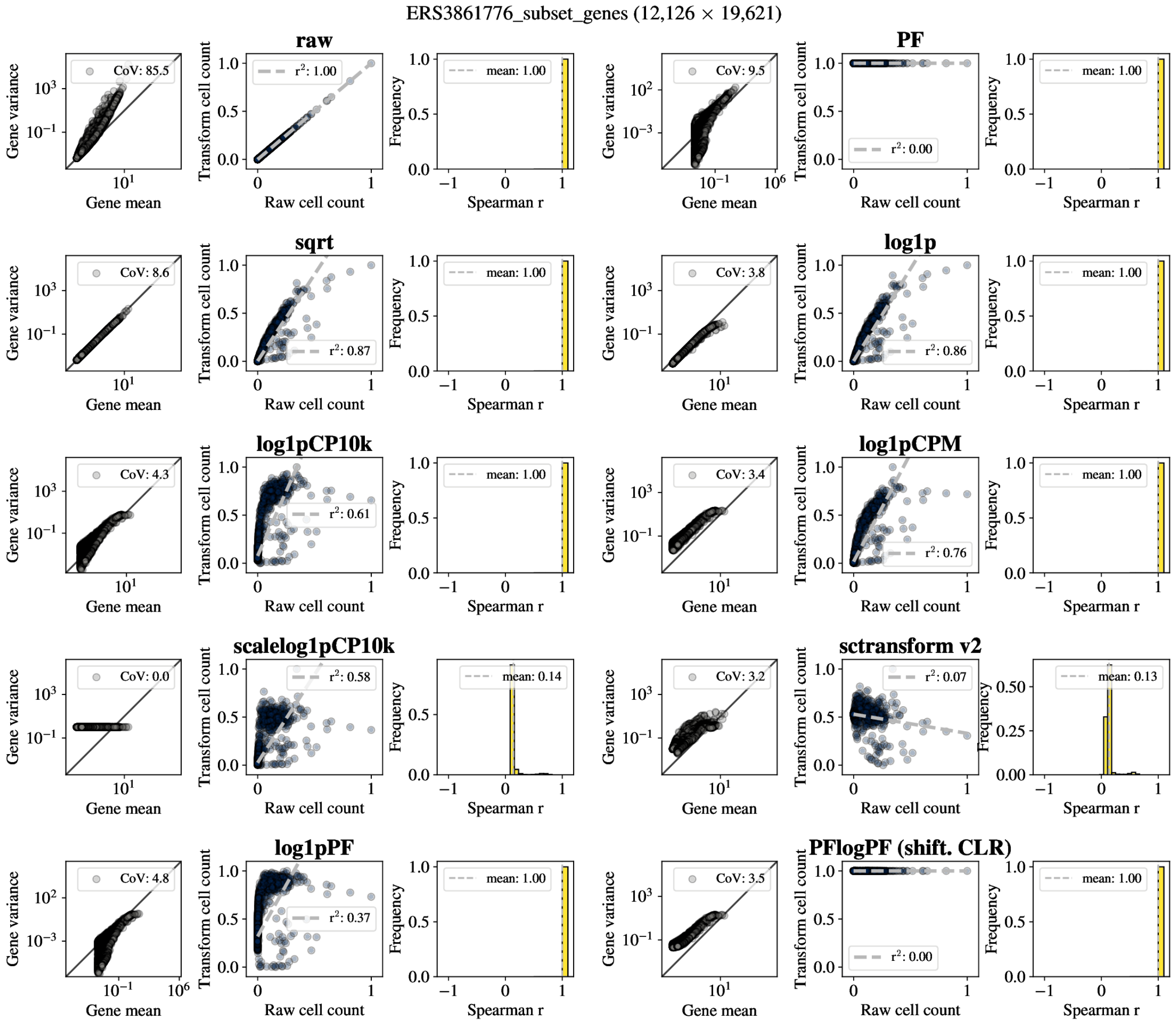


Supplementary Figure 1.24. **ERS3861776**. Per-method comparison of variance stabilization (left), depth normalization (middle), and monotonicity (right) for each normalization on dataset ERS3861776.

| **ERS3861777** | |
|---|---|
| cells | 1,322 |
| genes | 60,623 |
| nonzero entries | 4,547,992 |
| density | 0.0567 |
| total counts | 15,008,328 |
| mean counts / cell | 11,353 |
| mean counts / gene | 248 |
| min counts / cell | 1,420 |
| max counts / cell | 128,348 |
| overdispersion | 0.553 |

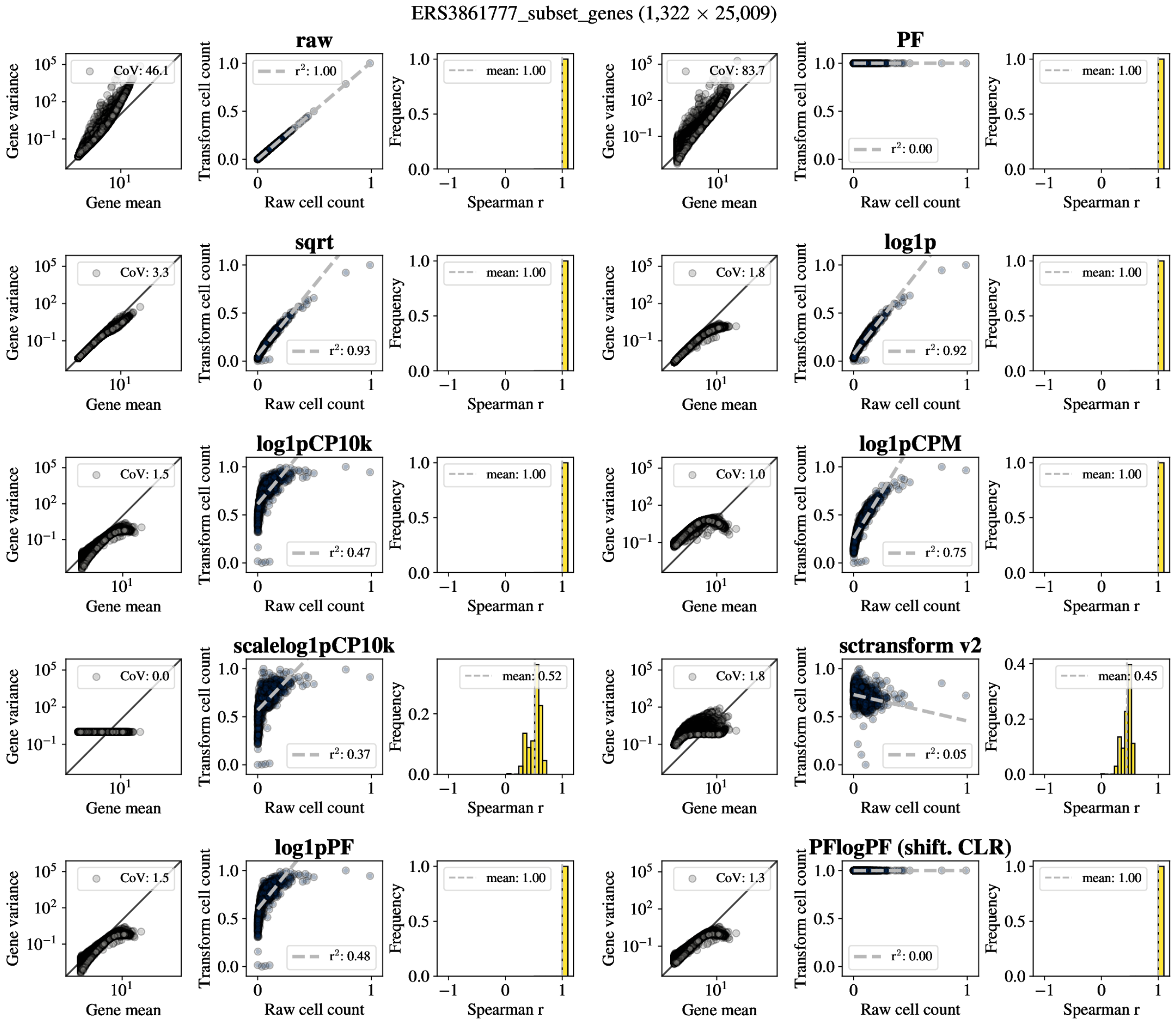


Supplementary Figure 1.25. **ERS3861777**. Per-method comparison of variance stabilization (left), depth normalization (middle), and monotonicity (right) for each normalization on dataset ERS3861777.

| **ERS3861778** | |
|---|---|
| cells | 1,954 |
| genes | 60,623 |
| nonzero entries | 6,377,497 |
| density | 0.0538 |
| total counts | 40,231,990 |
| mean counts / cell | 20,590 |
| mean counts / gene | 664 |
| min counts / cell | 549 |
| max counts / cell | 98,675 |
| overdispersion | 2.11 |

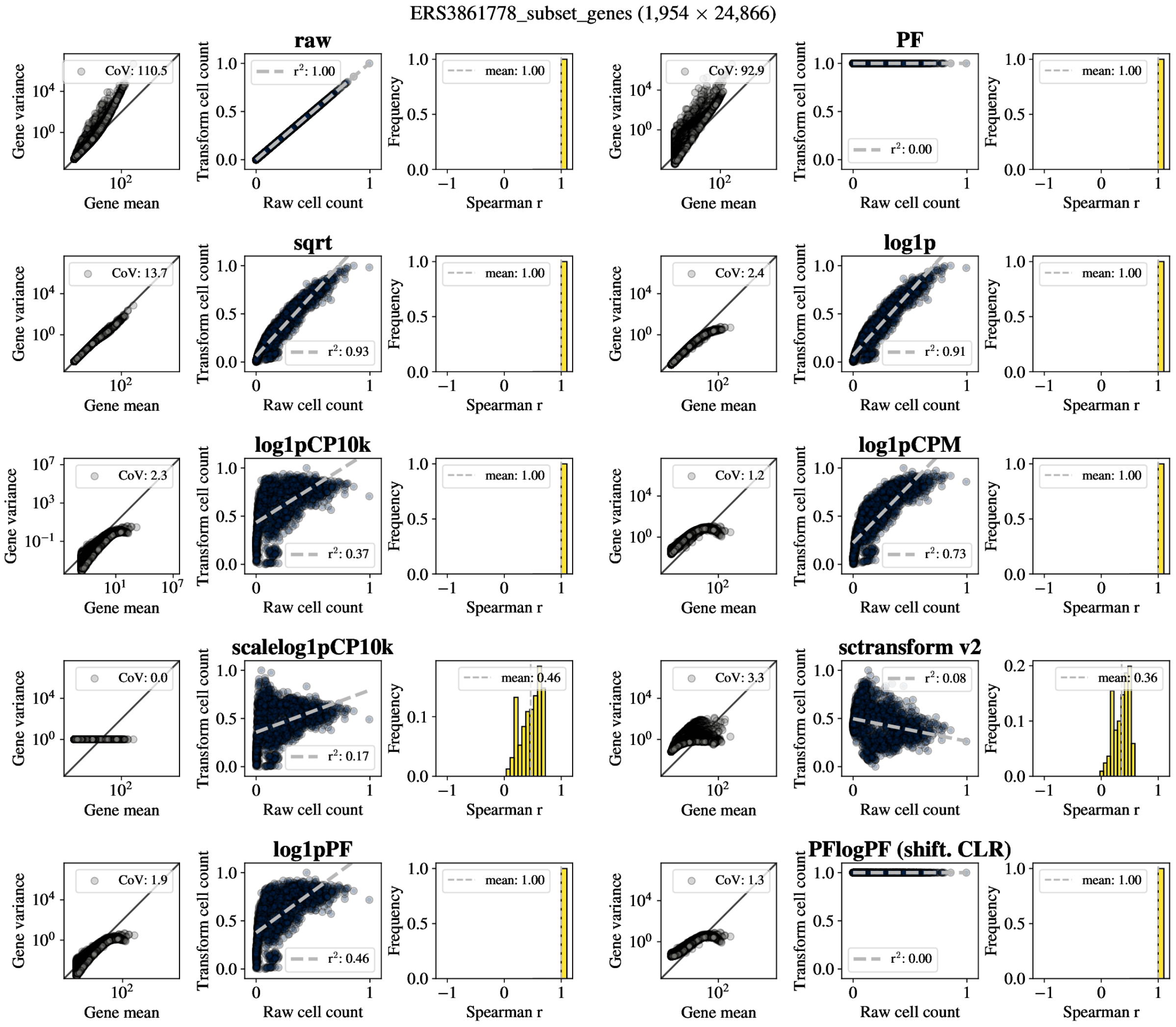


Supplementary Figure 1.26. **ERS3861778**. Per-method comparison of variance stabilization (left), depth normalization (middle), and monotonicity (right) for each normalization on dataset ERS3861778.

| **ERS3861779** | |
|---|---|
| cells | 1,645 |
| genes | 60,623 |
| nonzero entries | 3,948,039 |
| density | 0.0396 |
| total counts | 13,728,949 |
| mean counts / cell | 8,346 |
| mean counts / gene | 226 |
| min counts / cell | 435 |
| max counts / cell | 65,830 |
| overdispersion | 3.54 |

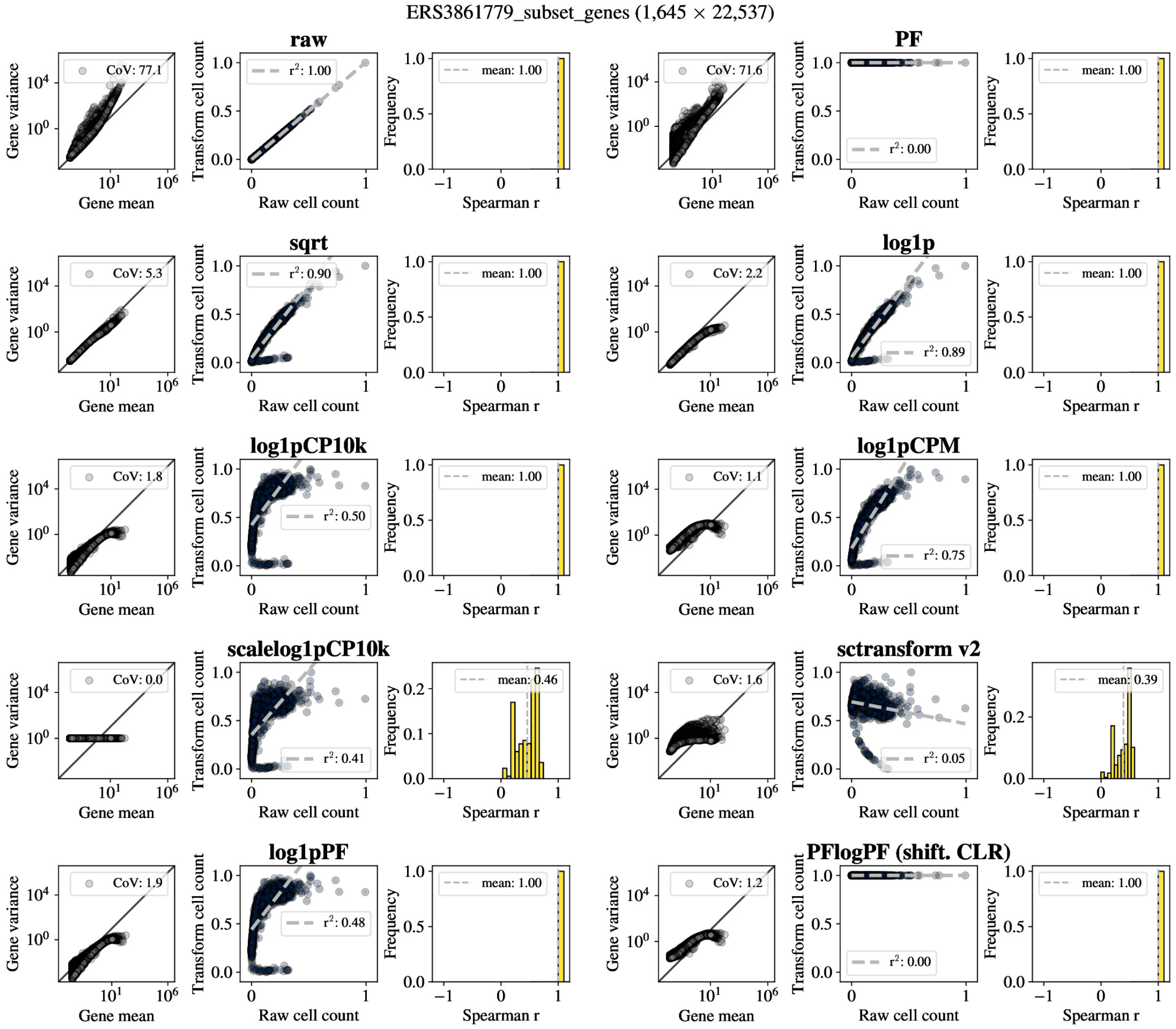


Supplementary Figure 1.27. **ERS3861779**. Per-method comparison of variance stabilization (left), depth normalization (middle), and monotonicity (right) for each normalization on dataset ERS3861779.

| **ERS3861780** | |
|---|---|
| cells | 10,669 |
| genes | 60,623 |
| nonzero entries | 3,062,753 |
| density | 0.00474 |
| total counts | 9,121,936 |
| mean counts / cell | 855 |
| mean counts / gene | 150 |
| min counts / cell | 92 |
| max counts / cell | 88,303 |
| overdispersion | 24.5 |

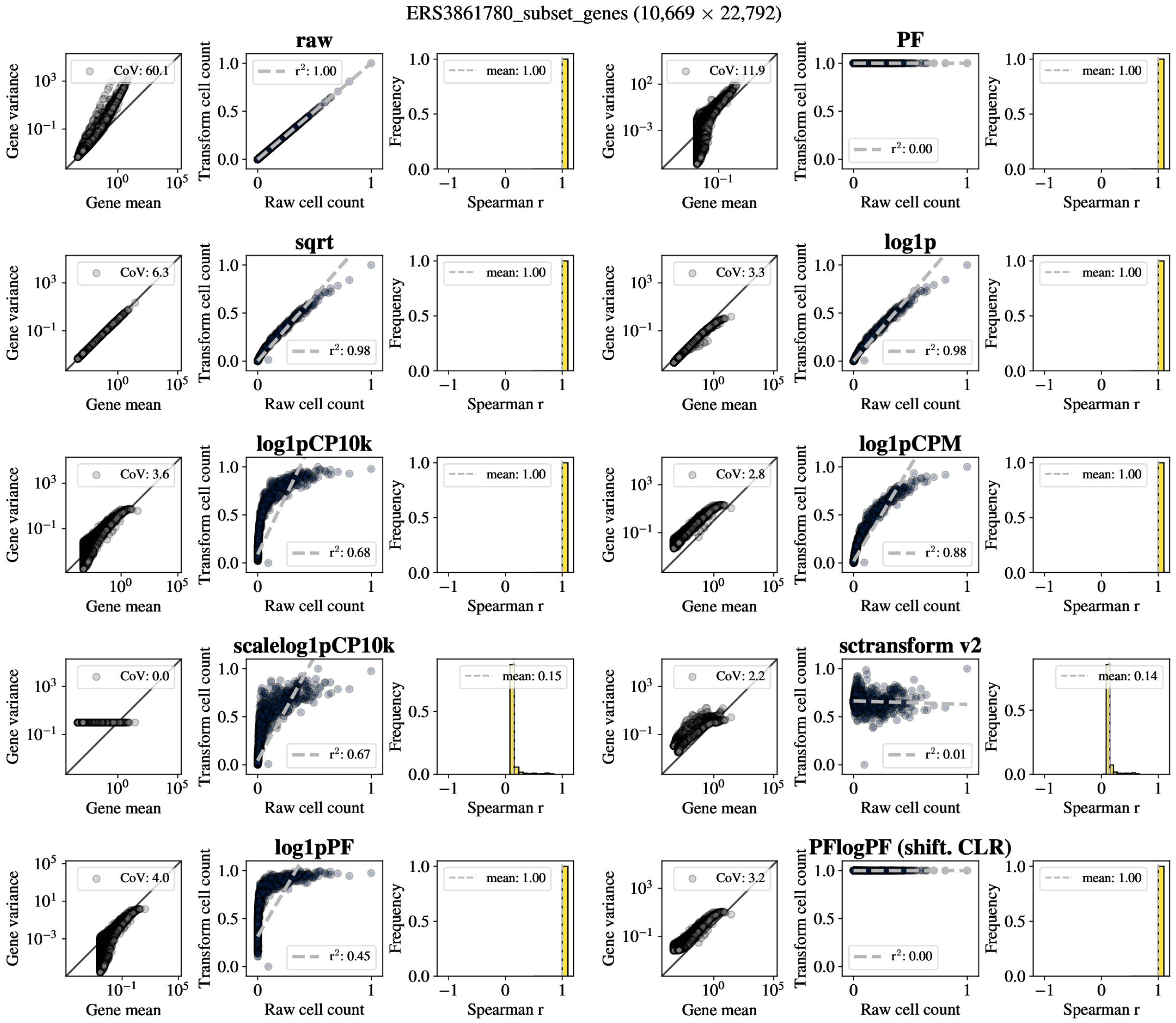


Supplementary Figure 1.28. **ERS3861780**. Per-method comparison of variance stabilization (left), depth normalization (middle), and monotonicity (right) for each normalization on dataset ERS3861780.

| **ERS3861781** | |
|---|---|
| cells | 1,128 |
| genes | 60,623 |
| nonzero entries | 4,644,841 |
| density | 0.0679 |
| total counts | 15,295,797 |
| mean counts / cell | 13,560 |
| mean counts / gene | 252 |
| min counts / cell | 2,784 |
| max counts / cell | 116,266 |
| overdispersion | 0.447 |

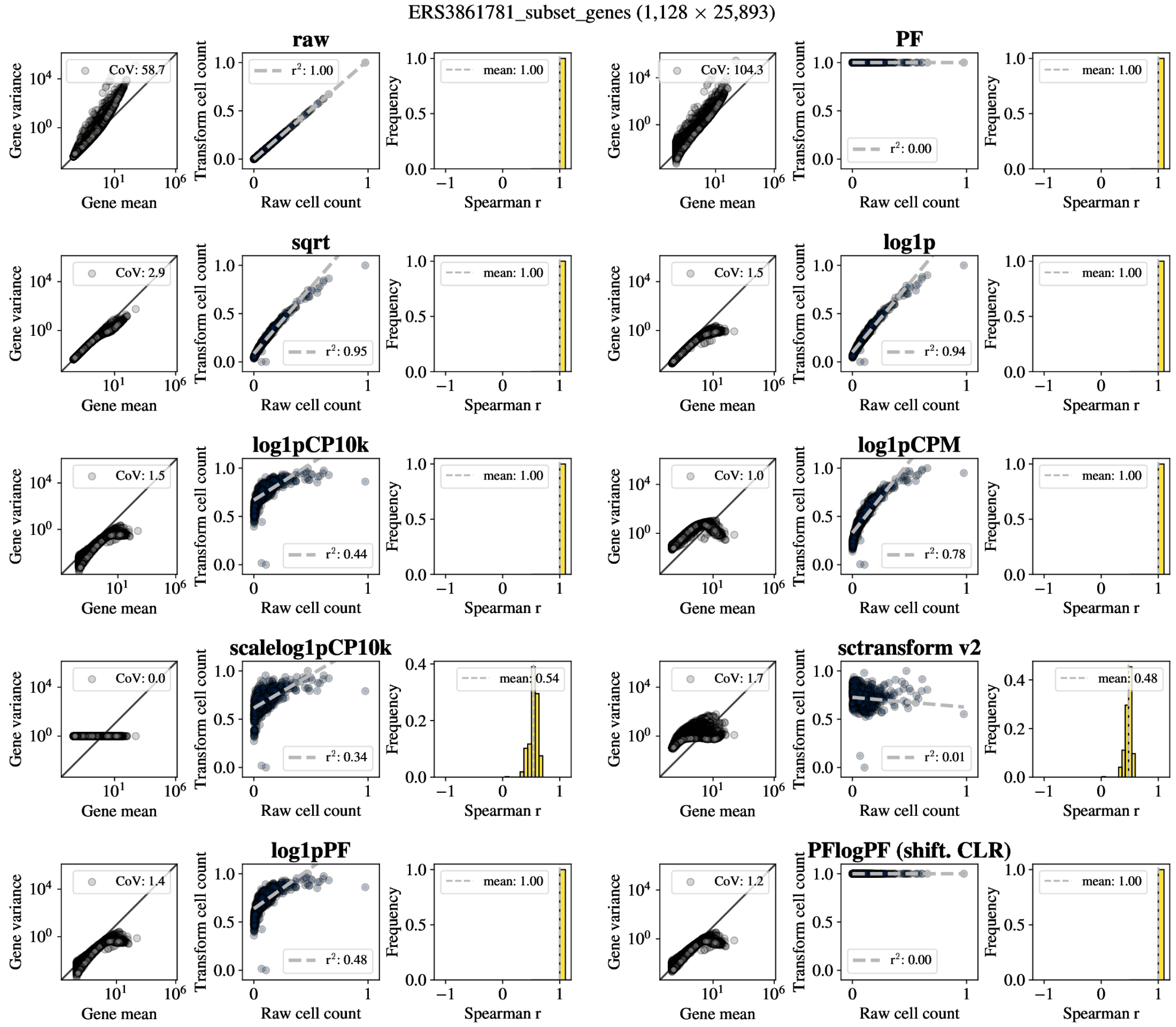


Supplementary Figure 1.29. **ERS3861781**. Per-method comparison of variance stabilization (left), depth normalization (middle), and monotonicity (right) for each normalization on dataset ERS3861781.

| ERS3861782 | |
|---|---|
| cells | 1,683 |
| genes | 60,623 |
| nonzero entries | 933,861 |
| density | 0.00915 |
| total counts | 4,511,798 |
| mean counts / cell | 2,681 |
| mean counts / gene | 74 |
| min counts / cell | 65 |
| max counts / cell | 52,575 |
| overdispersion | 6.82 |

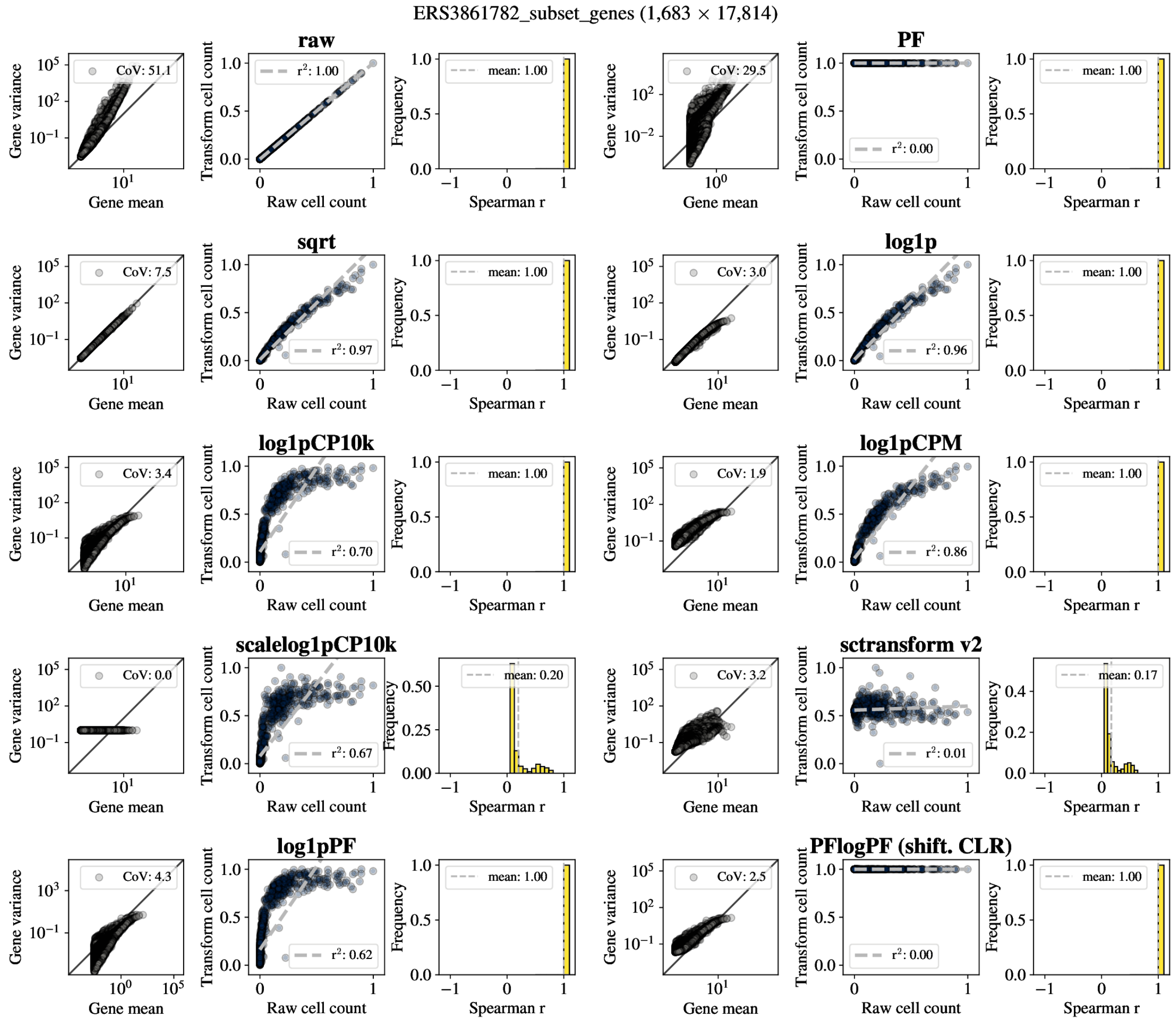


Supplementary Figure 1.30. **ERS3861782**. Per-method comparison of variance stabilization (left), depth normalization (middle), and monotonicity (right) for each normalization on dataset ERS3861782.

| **ERS3861783** | |
|---|---|
| cells | 4,866 |
| genes | 60,623 |
| nonzero entries | 4,312,827 |
| density | 0.0146 |
| total counts | 24,181,819 |
| mean counts / cell | 4,970 |
| mean counts / gene | 399 |
| min counts / cell | 173 |
| max counts / cell | 91,975 |
| overdispersion | 8.77 |

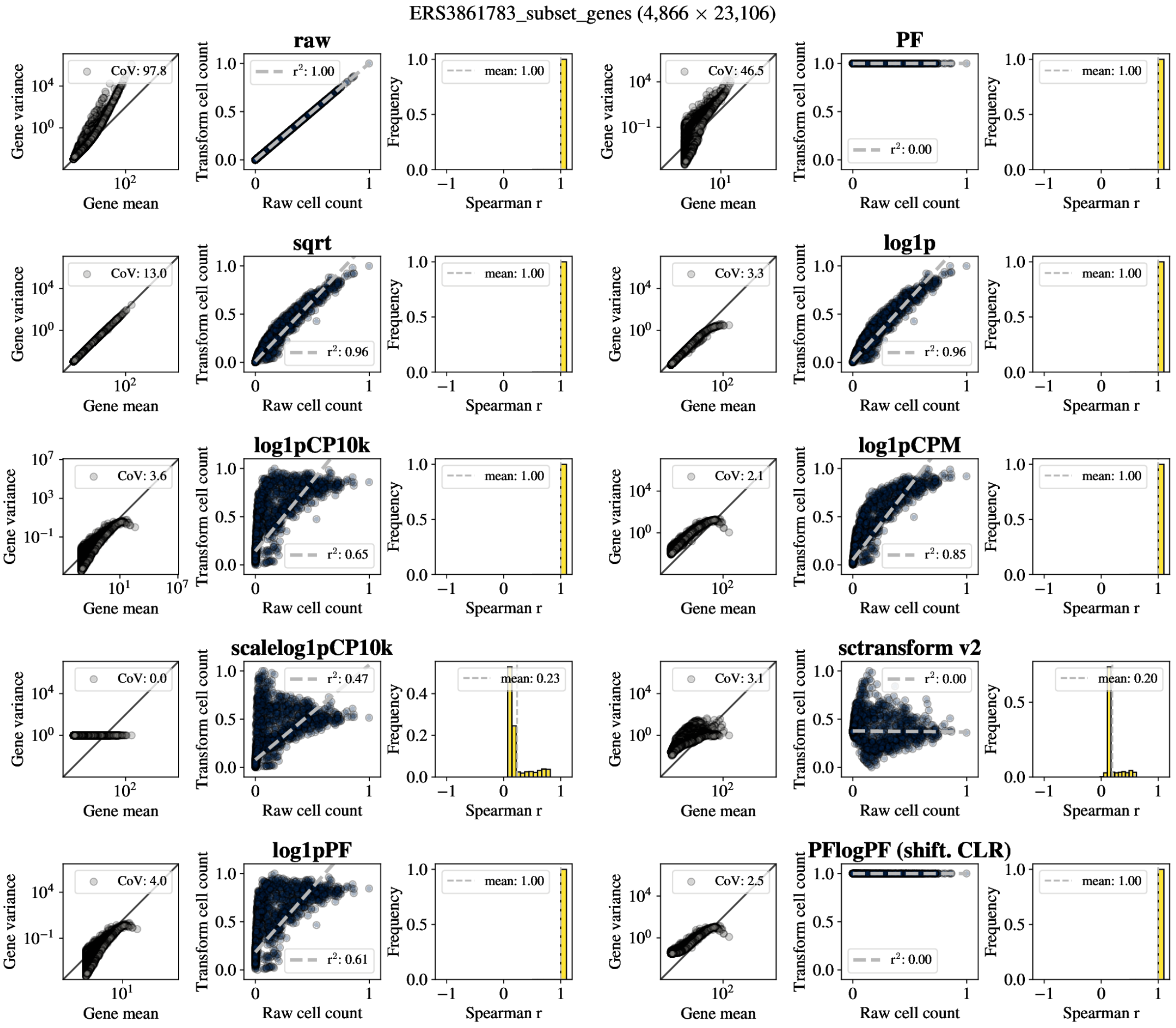


Supplementary Figure 1.31. **ERS3861783**. Per-method comparison of variance stabilization (left), depth normalization (middle), and monotonicity (right) for each normalization on dataset ERS3861783.

| **ERS3861784** | |
|---|---|
| cells | 2,959 |
| genes | 60,623 |
| nonzero entries | 1,110,357 |
| density | 0.00619 |
| total counts | 3,864,533 |
| mean counts / cell | 1,306 |
| mean counts / gene | 64 |
| min counts / cell | 83 |
| max counts / cell | 52,464 |
| overdispersion | 10.6 |

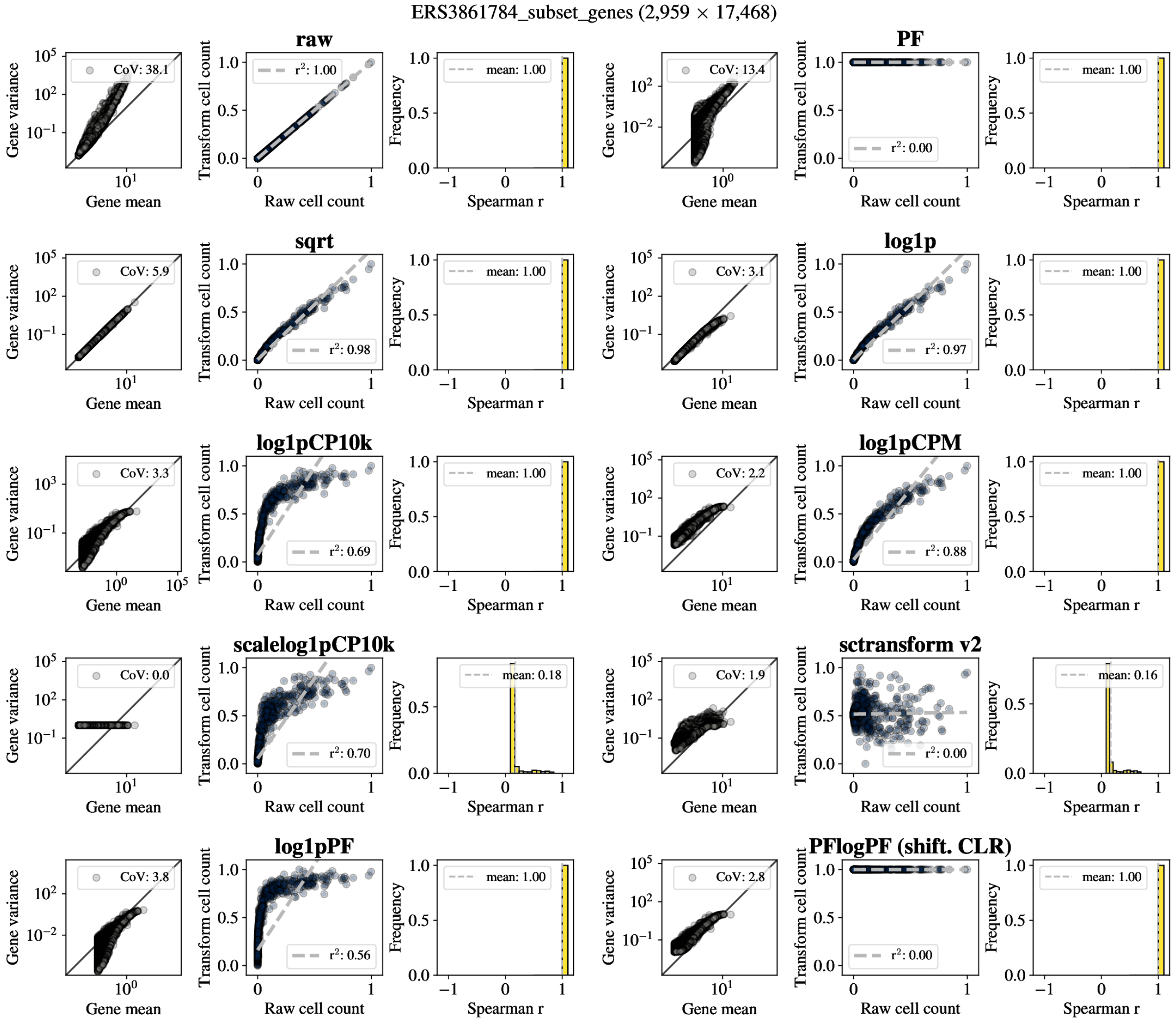


Supplementary Figure 1.32. **ERS3861784**. Per-method comparison of variance stabilization (left), depth normalization (middle), and monotonicity (right) for each normalization on dataset ERS3861784.

| ERS3861785 | |
|---|---|
| cells | 1,396 |
| genes | 60,623 |
| nonzero entries | 2,868,858 |
| density | 0.0339 |
| total counts | 10,516,537 |
| mean counts / cell | 7,533 |
| mean counts / gene | 173 |
| min counts / cell | 351 |
| max counts / cell | 40,880 |
| overdispersion | 1.38 |

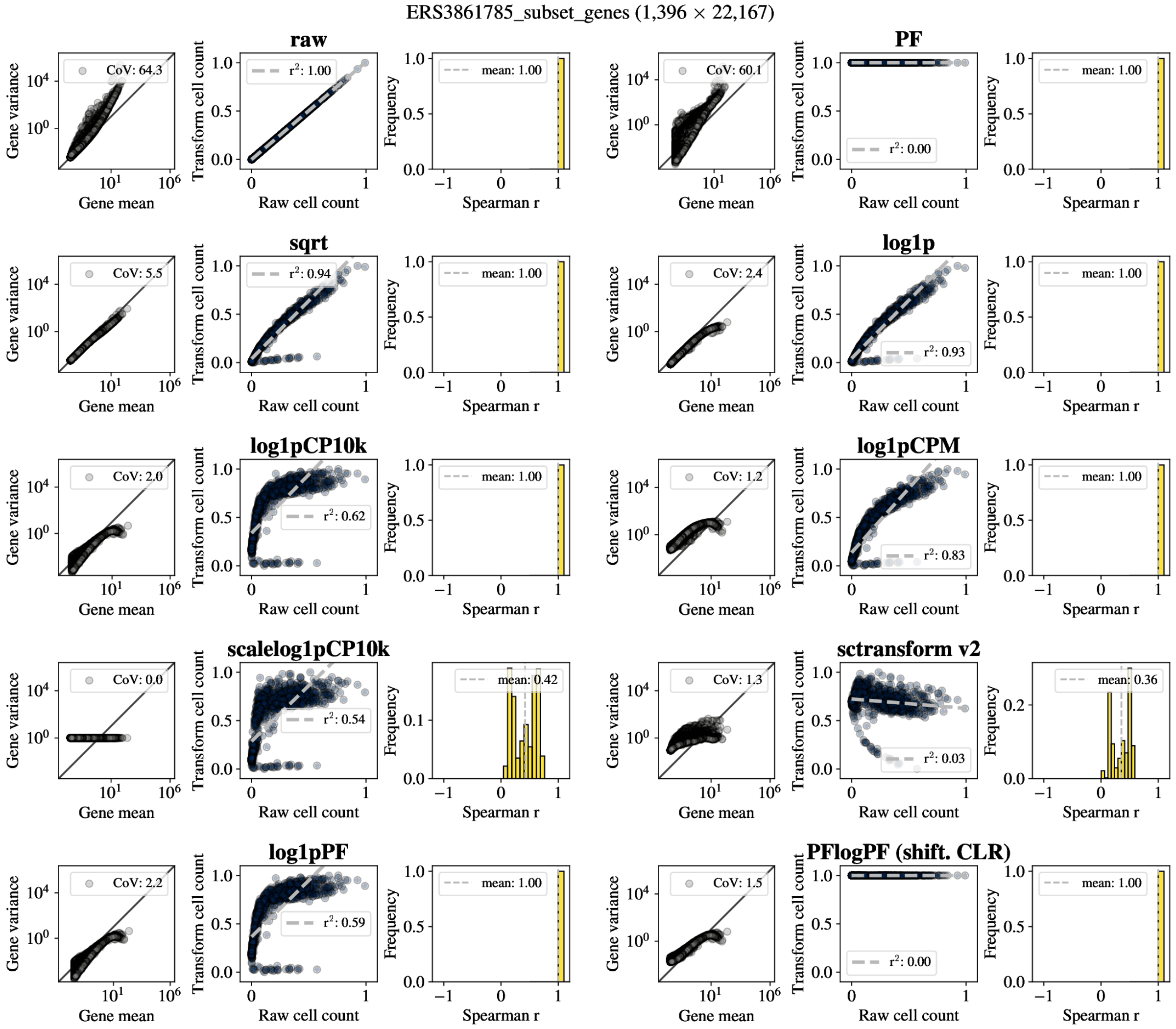


Supplementary Figure 1.33. **ERS3861785**. Per-method comparison of variance stabilization (left), depth normalization (middle), and monotonicity (right) for each normalization on dataset ERS3861785.

| **ERS3861786** | |
|---|---|
| cells | 5,779 |
| genes | 60,623 |
| nonzero entries | 10,371,212 |
| density | 0.0296 |
| total counts | 41,670,515 |
| mean counts / cell | 7,211 |
| mean counts / gene | 687 |
| min counts / cell | 80 |
| max counts / cell | 66,210 |
| overdispersion | 5.55 |

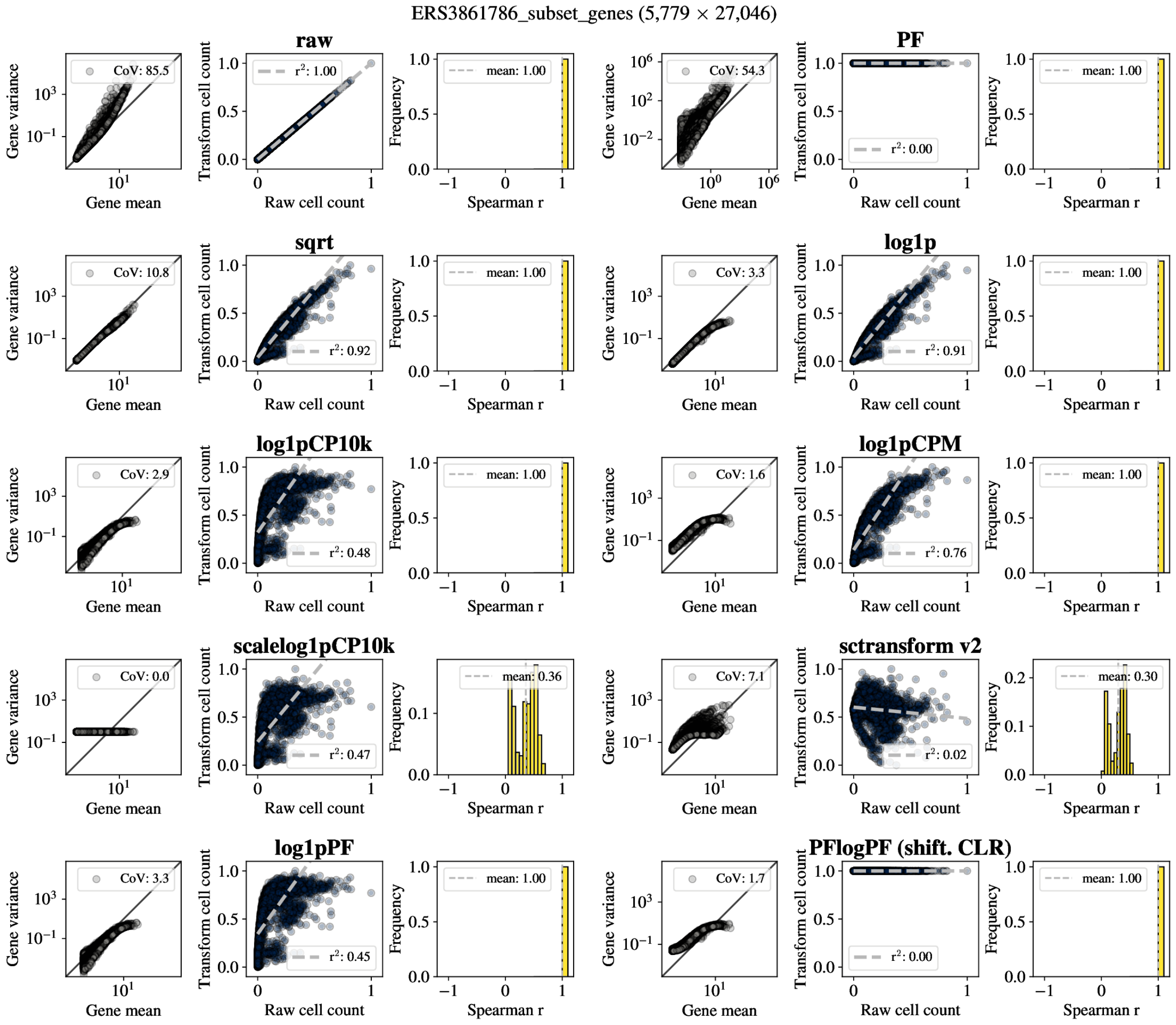


Supplementary Figure 1.34. **ERS3861786**. Per-method comparison of variance stabilization (left), depth normalization (middle), and monotonicity (right) for each normalization on dataset ERS3861786.

| **ERS3861787** | |
|---|---|
| cells | 13,328 |
| genes | 60,623 |
| nonzero entries | 20,374,804 |
| density | 0.0252 |
| total counts | 94,444,603 |
| mean counts / cell | 7,086 |
| mean counts / gene | 1,558 |
| min counts / cell | 155 |
| max counts / cell | 77,315 |
| overdispersion | 4.4 |

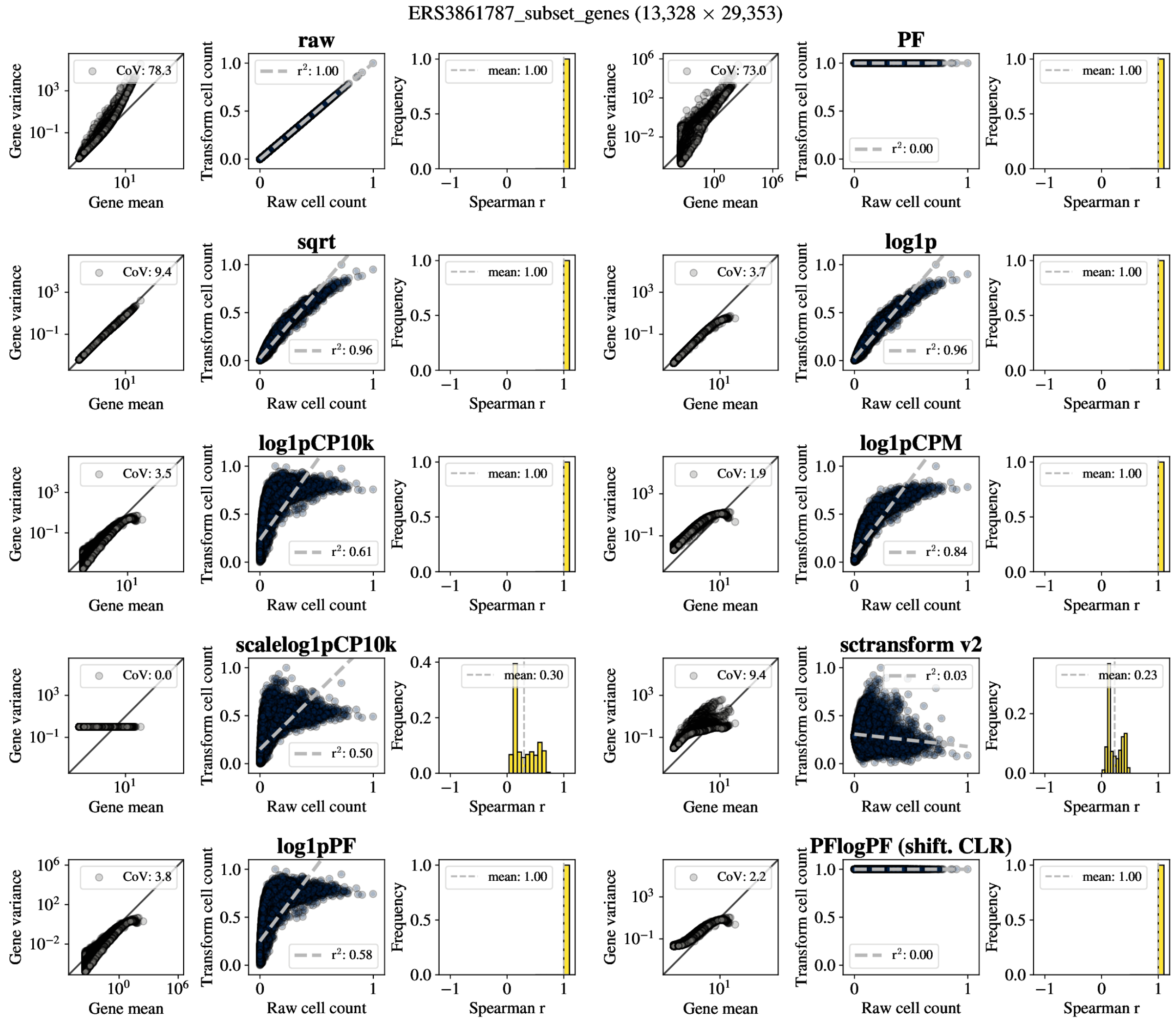


Supplementary Figure 1.35. **ERS3861787**. Per-method comparison of variance stabilization (left), depth normalization (middle), and monotonicity (right) for each normalization on dataset ERS3861787.

| **ERS3861788** | |
|---|---|
| cells | 4,079 |
| genes | 60,623 |
| nonzero entries | 11,612,050 |
| density | 0.047 |
| total counts | 54,024,863 |
| mean counts / cell | 13,245 |
| mean counts / gene | 891 |
| min counts / cell | 4,984 |
| max counts / cell | 76,783 |
| overdispersion | 0.736 |

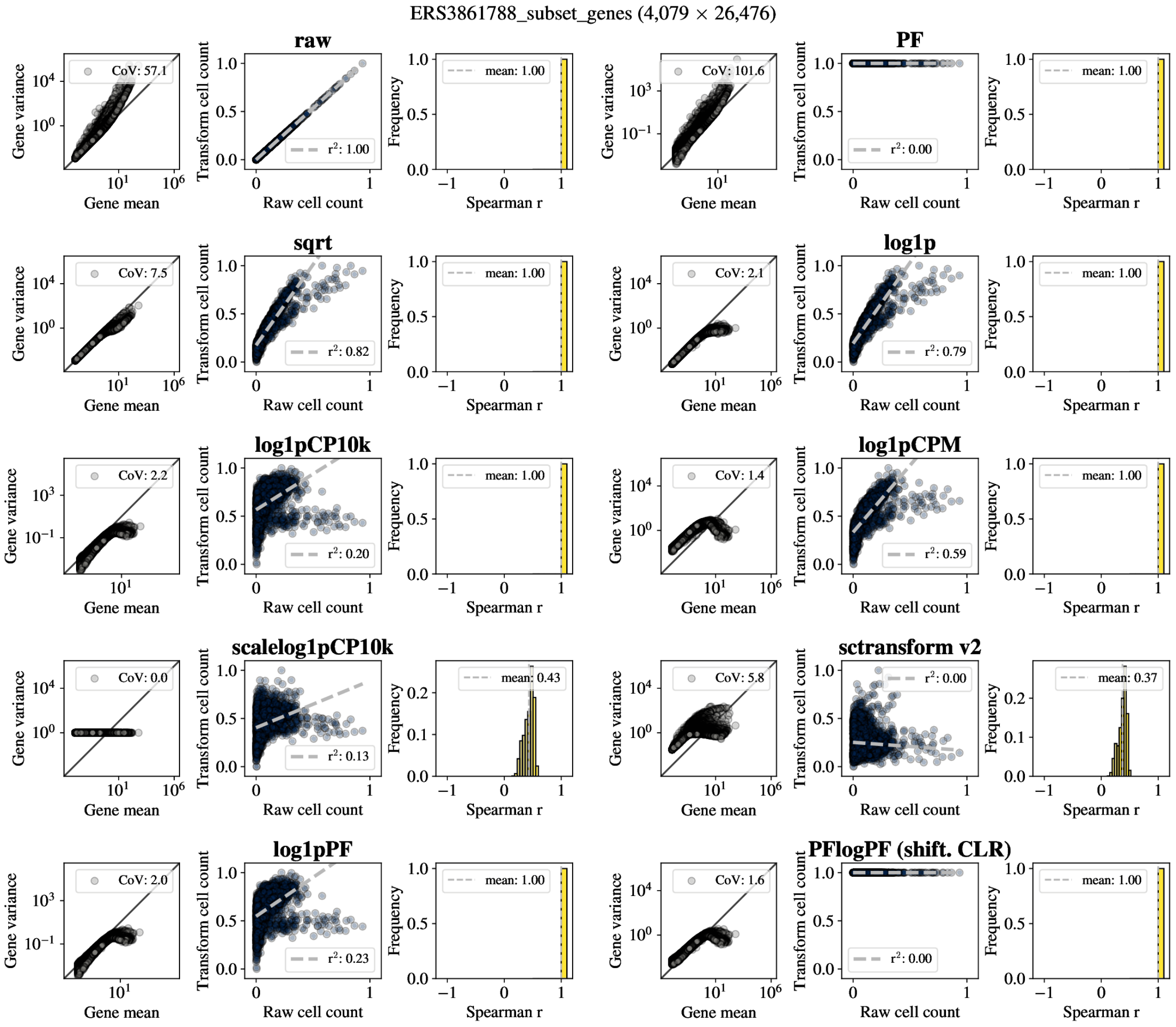


Supplementary Figure 1.36. **ERS3861788**. Per-method comparison of variance stabilization (left), depth normalization (middle), and monotonicity (right) for each normalization on dataset ERS3861788.

| ERS3861789 | |
|---|---|
| cells | 7,037 |
| genes | 60,623 |
| nonzero entries | 10,031,393 |
| density | 0.0235 |
| total counts | 46,068,408 |
| mean counts / cell | 6,547 |
| mean counts / gene | 760 |
| min counts / cell | 77 |
| max counts / cell | 64,054 |
| overdispersion | 3.95 |

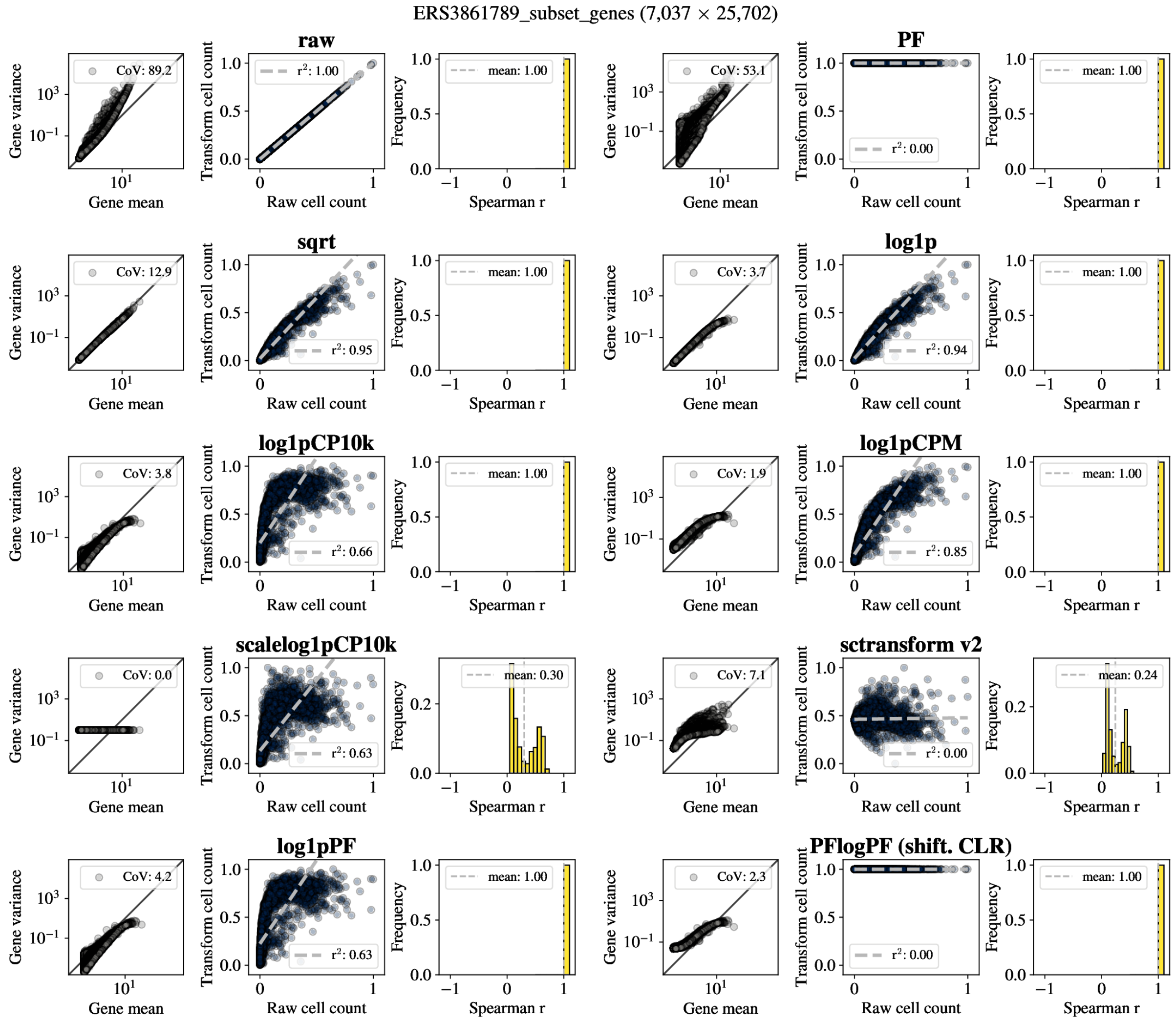


Supplementary Figure 1.37. **ERS3861789**. Per-method comparison of variance stabilization (left), depth normalization (middle), and monotonicity (right) for each normalization on dataset ERS3861789.

| **ERS3861790** | |
|---|---|
| cells | 4,915 |
| genes | 60,623 |
| nonzero entries | 15,633,451 |
| density | 0.0525 |
| total counts | 92,573,088 |
| mean counts / cell | 18,835 |
| mean counts / gene | 1,527 |
| min counts / cell | 2,191 |
| max counts / cell | 95,356 |
| overdispersion | 1.24 |

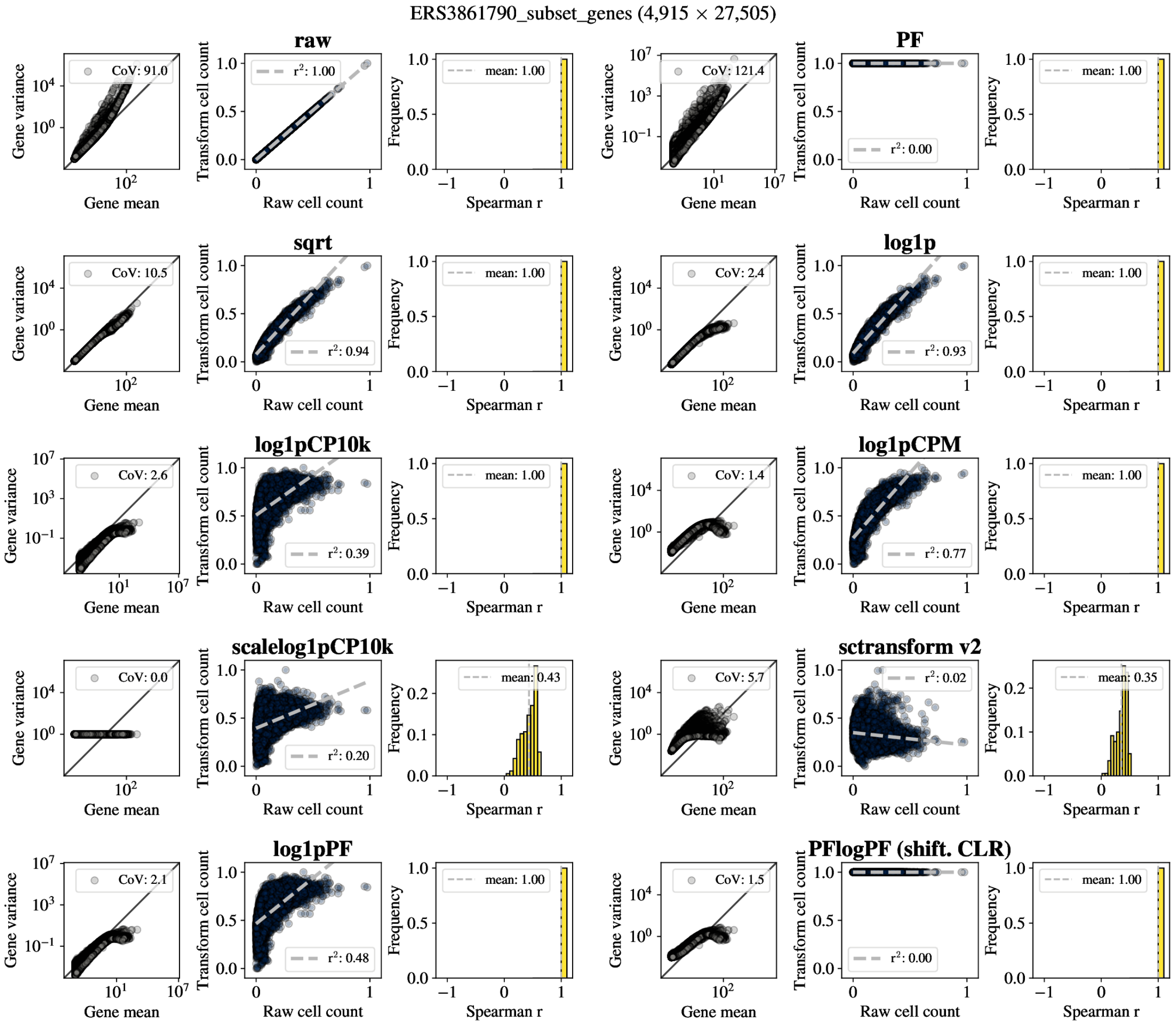


Supplementary Figure 1.38. **ERS3861790**. Per-method comparison of variance stabilization (left), depth normalization (middle), and monotonicity (right) for each normalization on dataset ERS3861790.

| ERS3861791 | |
|---|---|
| cells | 850 |
| genes | 60,623 |
| nonzero entries | 2,755,351 |
| density | 0.0535 |
| total counts | 9,620,644 |
| mean counts / cell | 11,318 |
| mean counts / gene | 159 |
| min counts / cell | 1,988 |
| max counts / cell | 49,473 |
| overdispersion | 0.573 |

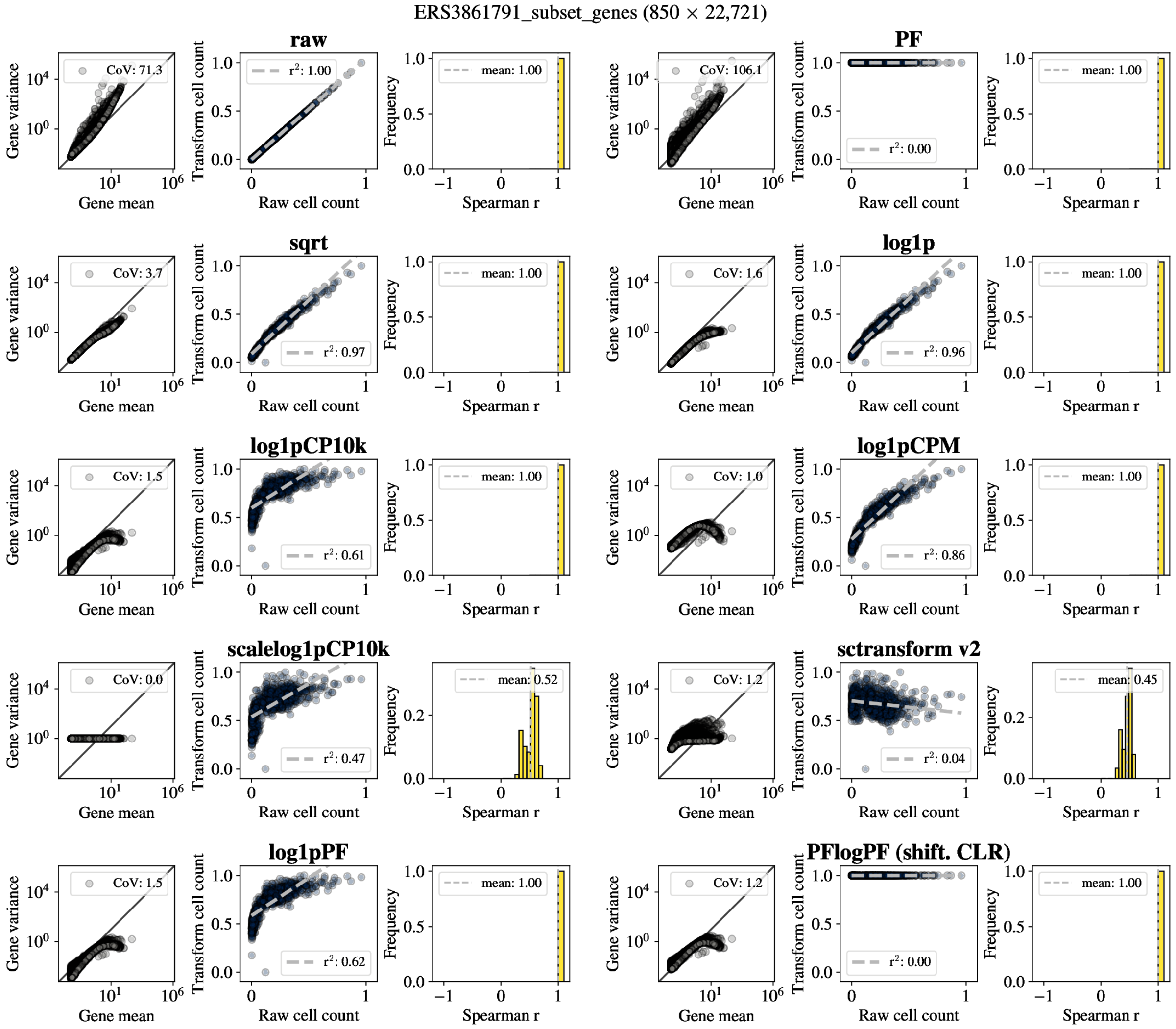


Supplementary Figure 1.39. **ERS3861791**. Per-method comparison of variance stabilization (left), depth normalization (middle), and monotonicity (right) for each normalization on dataset ERS3861791.

| **ERS3861792** | |
|---|---|
| cells | 3,952 |
| genes | 60,623 |
| nonzero entries | 12,942,079 |
| density | 0.054 |
| total counts | 80,271,464 |
| mean counts / cell | 20,312 |
| mean counts / gene | 1,324 |
| min counts / cell | 2,791 |
| max counts / cell | 77,186 |
| overdispersion | 1.07 |

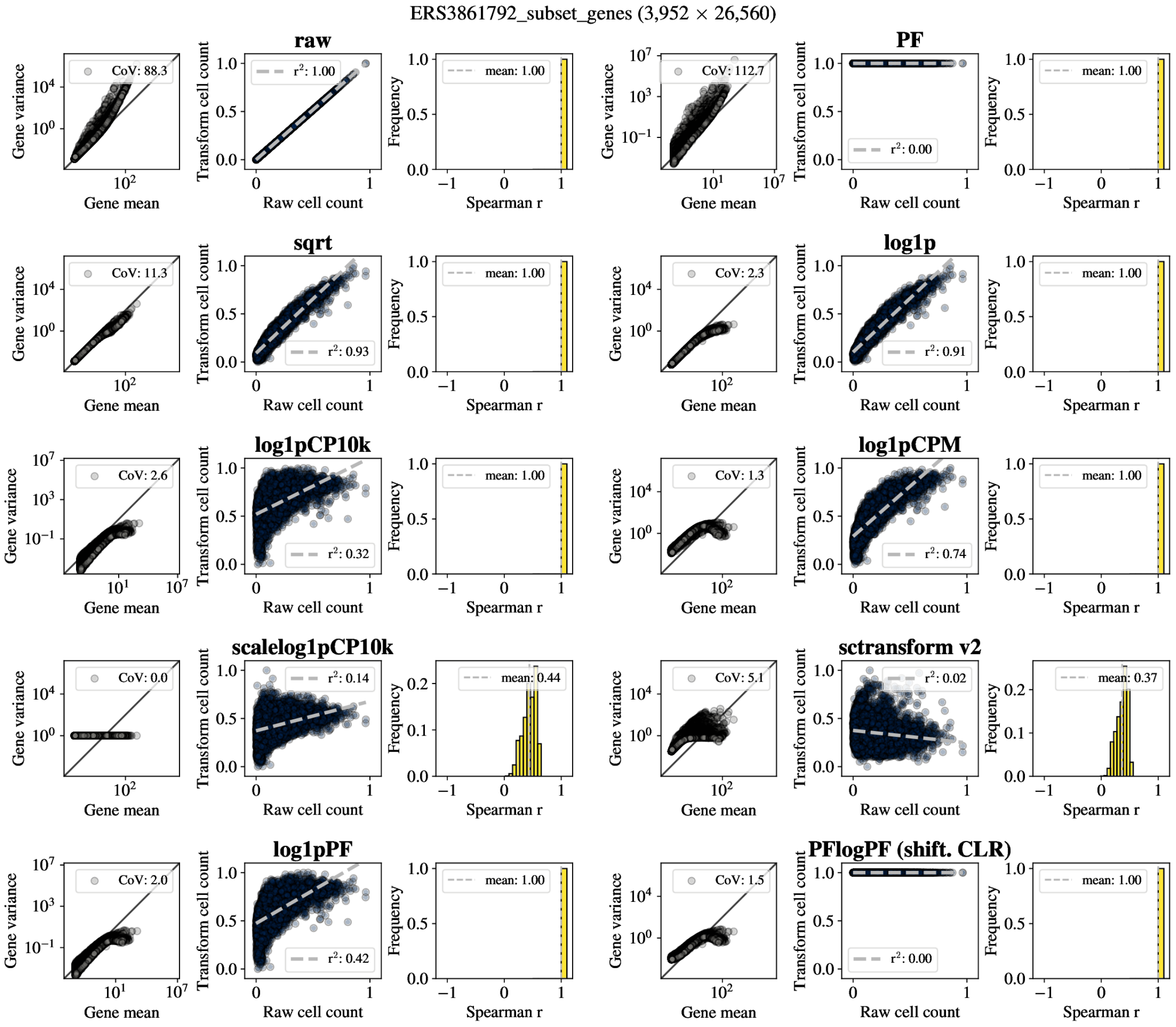


Supplementary Figure 1.40. **ERS3861792**. Per-method comparison of variance stabilization (left), depth normalization (middle), and monotonicity (right) for each normalization on dataset ERS3861792.

| **ERS3861793** | |
|---|---|
| cells | 11,124 |
| genes | 60,623 |
| nonzero entries | 1,975,997 |
| density | 0.00293 |
| total counts | 6,128,011 |
| mean counts / cell | 551 |
| mean counts / gene | 101 |
| min counts / cell | 60 |
| max counts / cell | 59,743 |
| overdispersion | 38.3 |

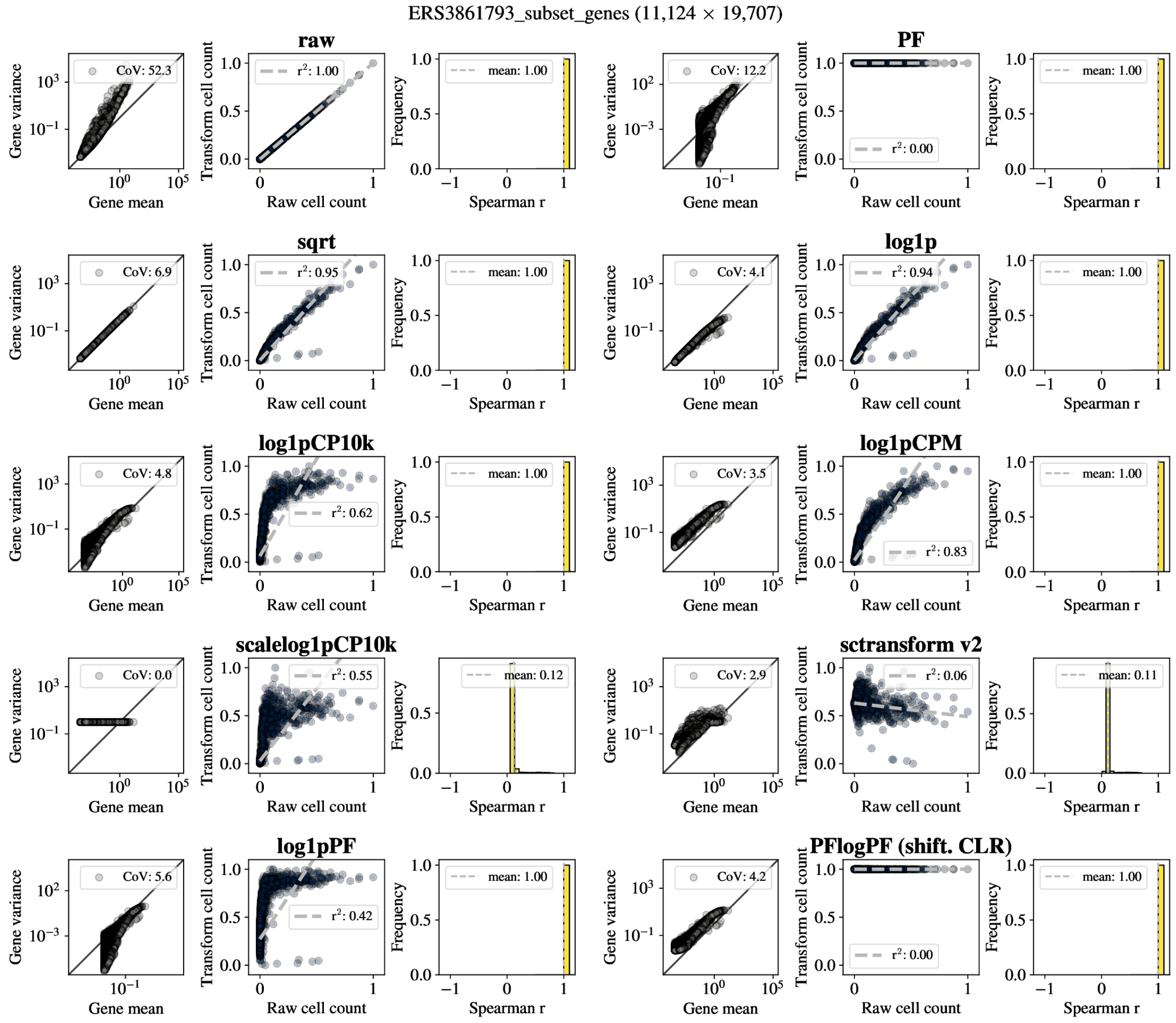


Supplementary Figure 1.41. **ERS3861793**. Per-method comparison of variance stabilization (left), depth normalization (middle), and monotonicity (right) for each normalization on dataset ERS3861793.

| **ERS3861794** | |
|---|---|
| cells | 4,954 |
| genes | 60,623 |
| nonzero entries | 14,966,635 |
| density | 0.0498 |
| total counts | 50,897,401 |
| mean counts / cell | 10,274 |
| mean counts / gene | 840 |
| min counts / cell | 4,243 |
| max counts / cell | 57,856 |
| overdispersion | 0.241 |

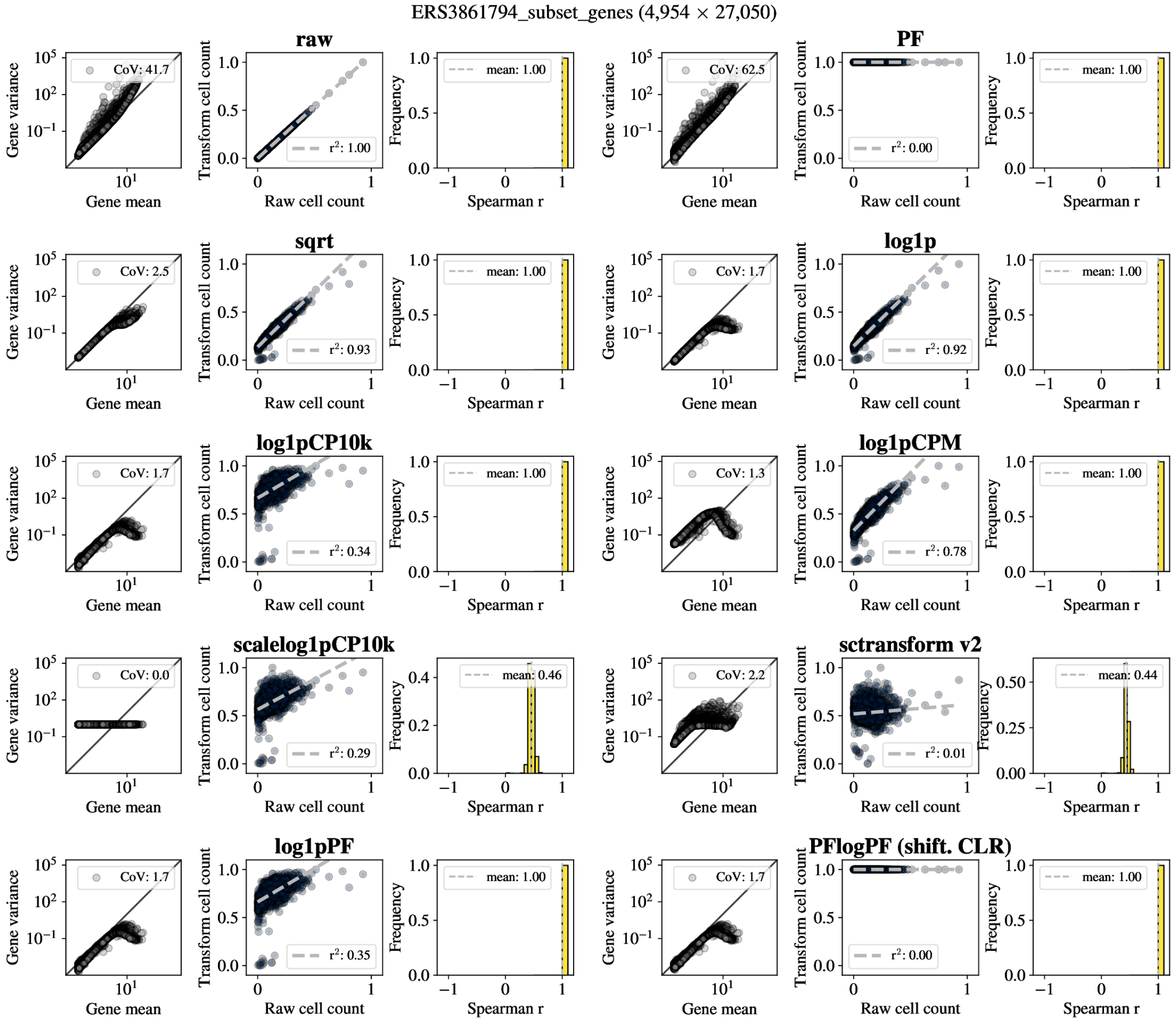


Supplementary Figure 1.42. **ERS3861794**. Per-method comparison of variance stabilization (left), depth normalization (middle), and monotonicity (right) for each normalization on dataset ERS3861794.

| ERS3861795 | |
|---|---|
| cells | 6,203 |
| genes | 60,623 |
| nonzero entries | 10,405,038 |
| density | 0.0277 |
| total counts | 41,287,053 |
| mean counts / cell | 6,656 |
| mean counts / gene | 681 |
| min counts / cell | 95 |
| max counts / cell | 131,982 |
| overdispersion | 4.84 |

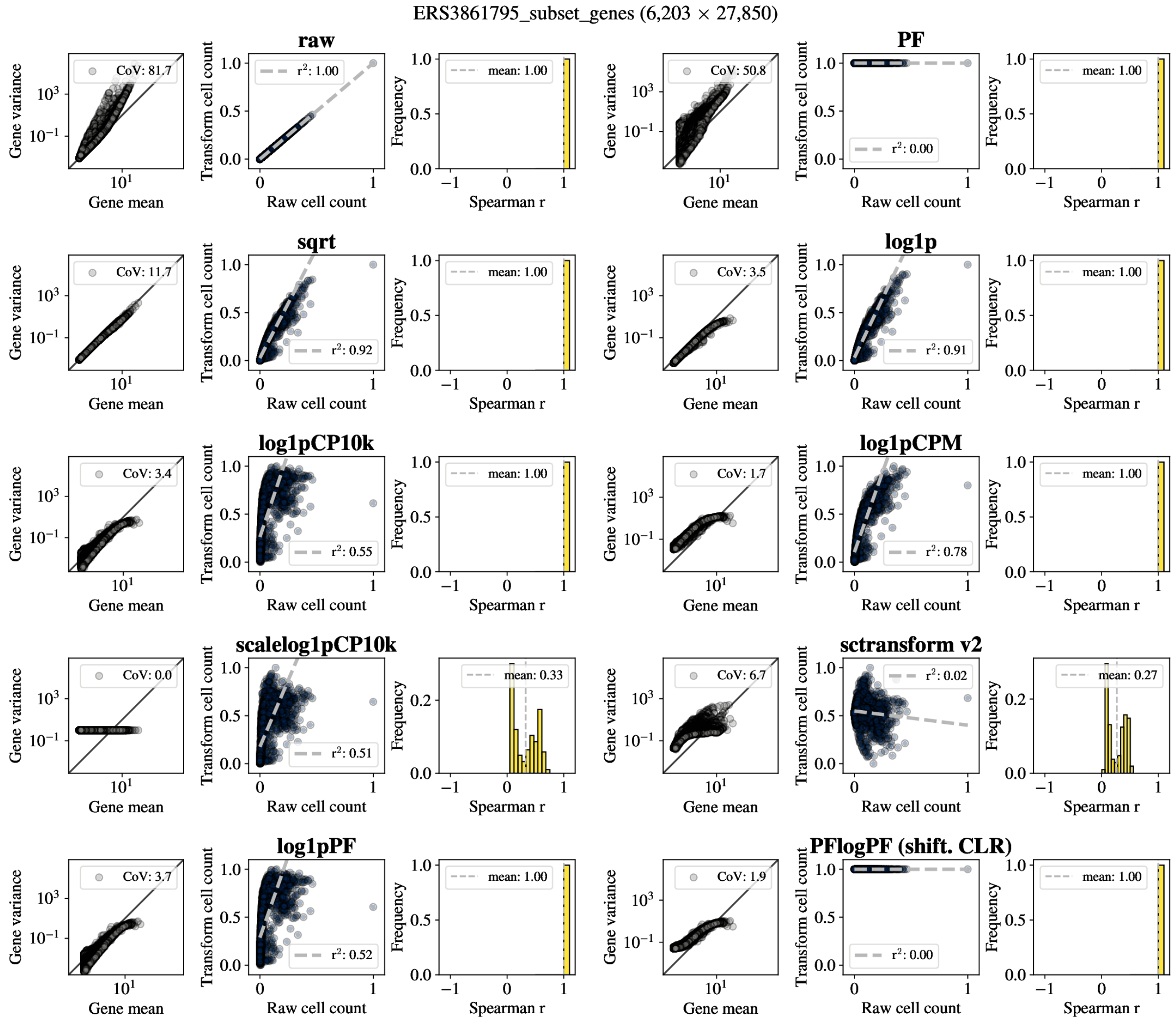


Supplementary Figure 1.43. **ERS3861795**. Per-method comparison of variance stabilization (left), depth normalization (middle), and monotonicity (right) for each normalization on dataset ERS3861795.

| ERS3861796 | |
|---|---|
| cells | 3,107 |
| genes | 60,623 |
| nonzero entries | 14,288,415 |
| density | 0.0759 |
| total counts | 84,141,520 |
| mean counts / cell | 27,081 |
| mean counts / gene | 1,388 |
| min counts / cell | 9,541 |
| max counts / cell | 70,539 |
| overdispersion | 1.76 |

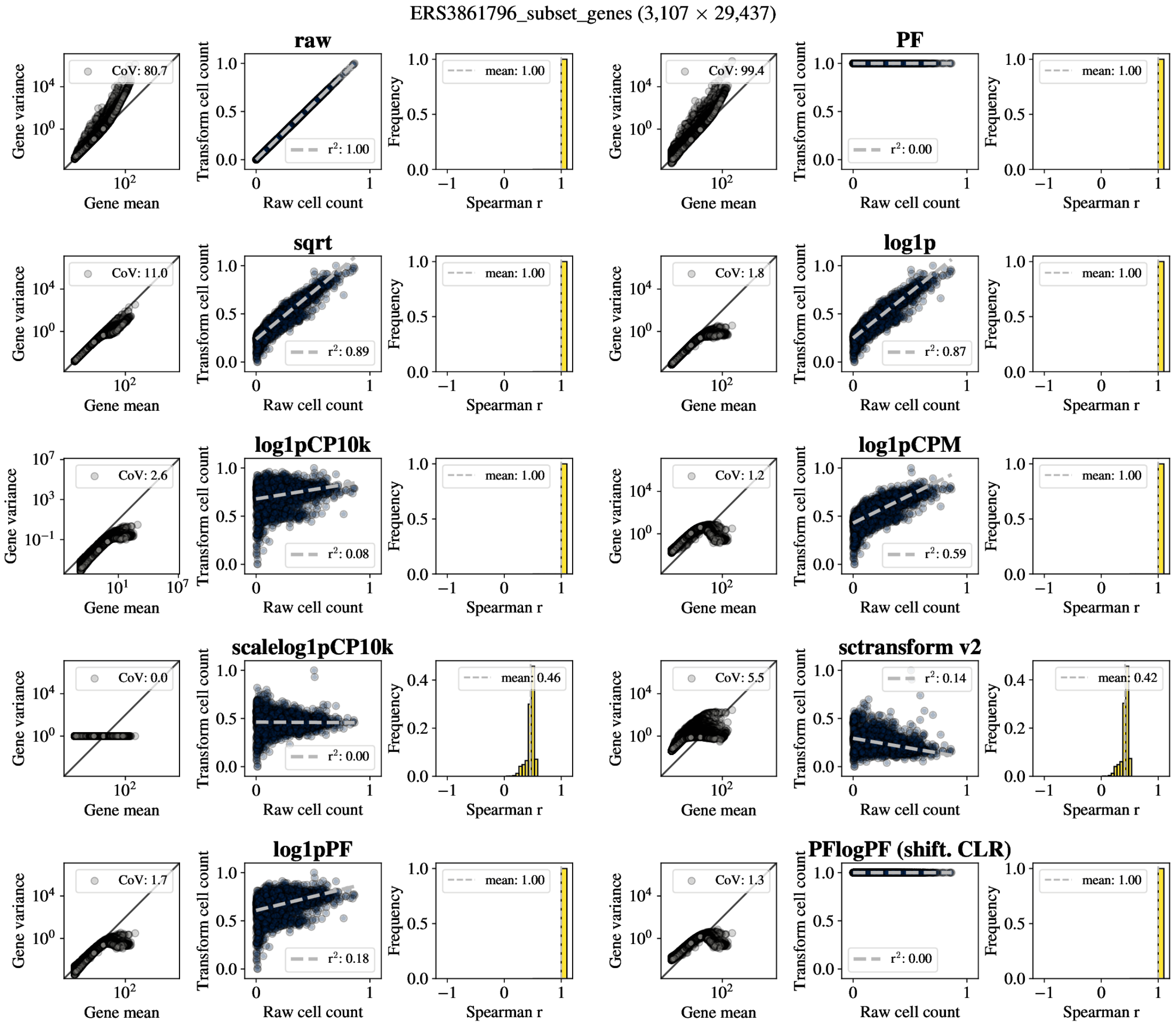


Supplementary Figure 1.44. **ERS3861796**. Per-method comparison of variance stabilization (left), depth normalization (middle), and monotonicity (right) for each normalization on dataset ERS3861796.

| **ERS3861797** | |
|---|---|
| cells | 10,255 |
| genes | 60,623 |
| nonzero entries | 15,270,224 |
| density | 0.0246 |
| total counts | 54,139,613 |
| mean counts / cell | 5,279 |
| mean counts / gene | 893 |
| min counts / cell | 66 |
| max counts / cell | 49,189 |
| overdispersion | 2.8 |

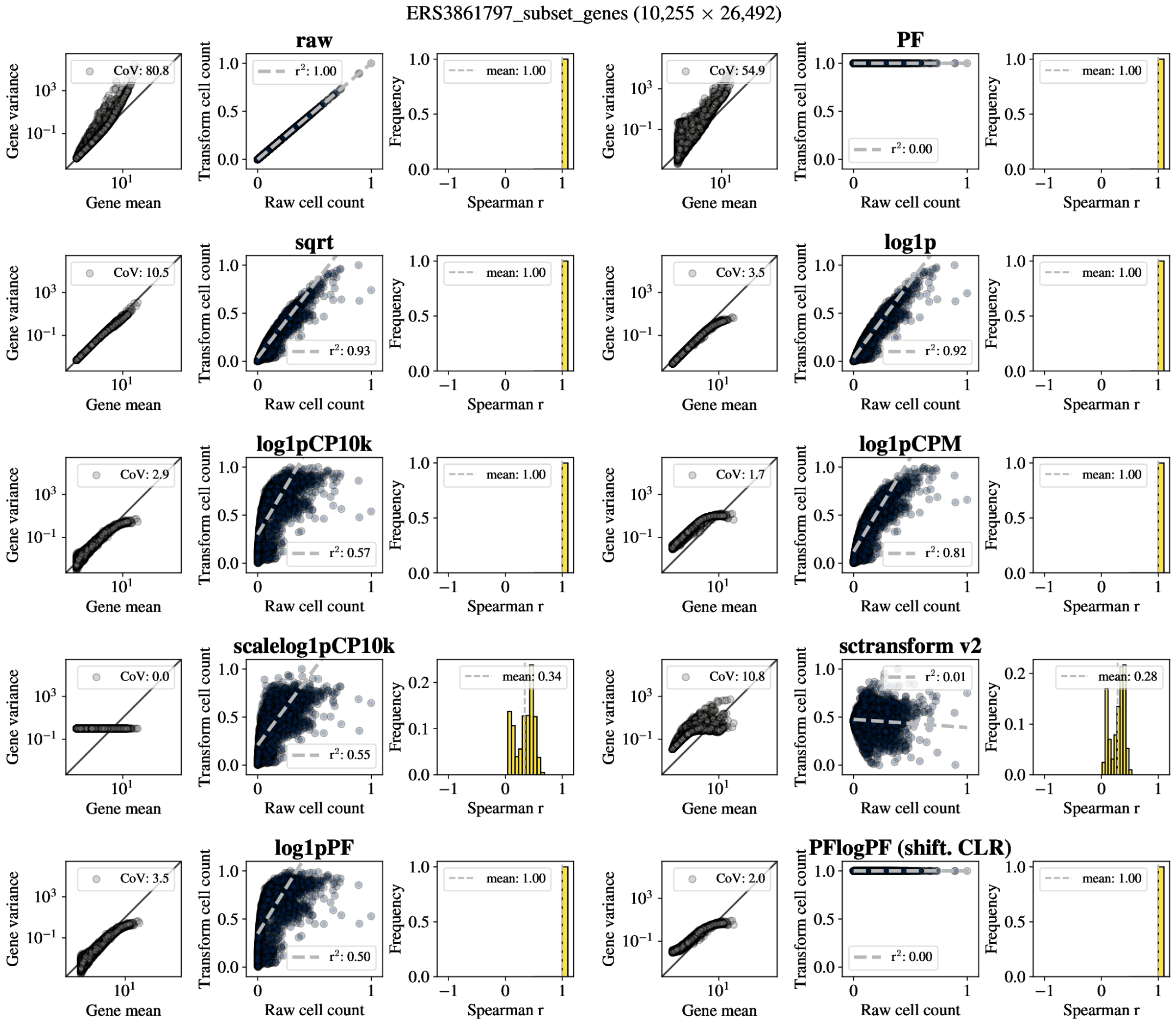


Supplementary Figure 1.45. **ERS3861797**. Per-method comparison of variance stabilization (left), depth normalization (middle), and monotonicity (right) for each normalization on dataset ERS3861797.

| **ERS3861798** | |
|---|---|
| cells | 5,399 |
| genes | 60,623 |
| nonzero entries | 19,741,952 |
| density | 0.0603 |
| total counts | 103,575,256 |
| mean counts / cell | 19,184 |
| mean counts / gene | 1,709 |
| min counts / cell | 3,517 |
| max counts / cell | 67,302 |
| overdispersion | 1.23 |

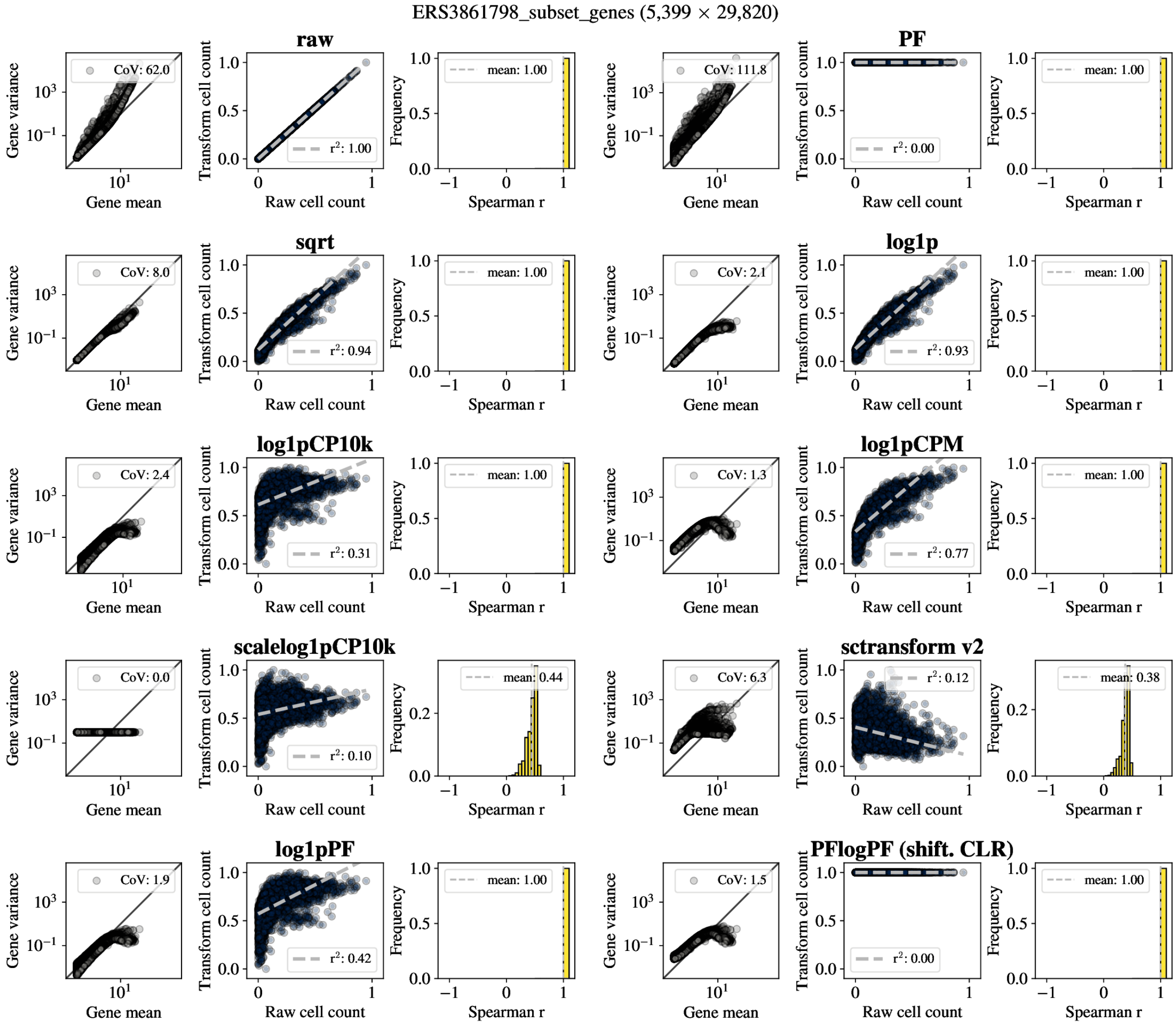


Supplementary Figure 1.46. **ERS3861798**. Per-method comparison of variance stabilization (left), depth normalization (middle), and monotonicity (right) for each normalization on dataset ERS3861798.

| ERS3861799 | |
|---|---|
| cells | 5,764 |
| genes | 60,623 |
| nonzero entries | 9,631,597 |
| density | 0.0276 |
| total counts | 43,792,694 |
| mean counts / cell | 7,598 |
| mean counts / gene | 722 |
| min counts / cell | 51 |
| max counts / cell | 108,527 |
| overdispersion | 4.09 |

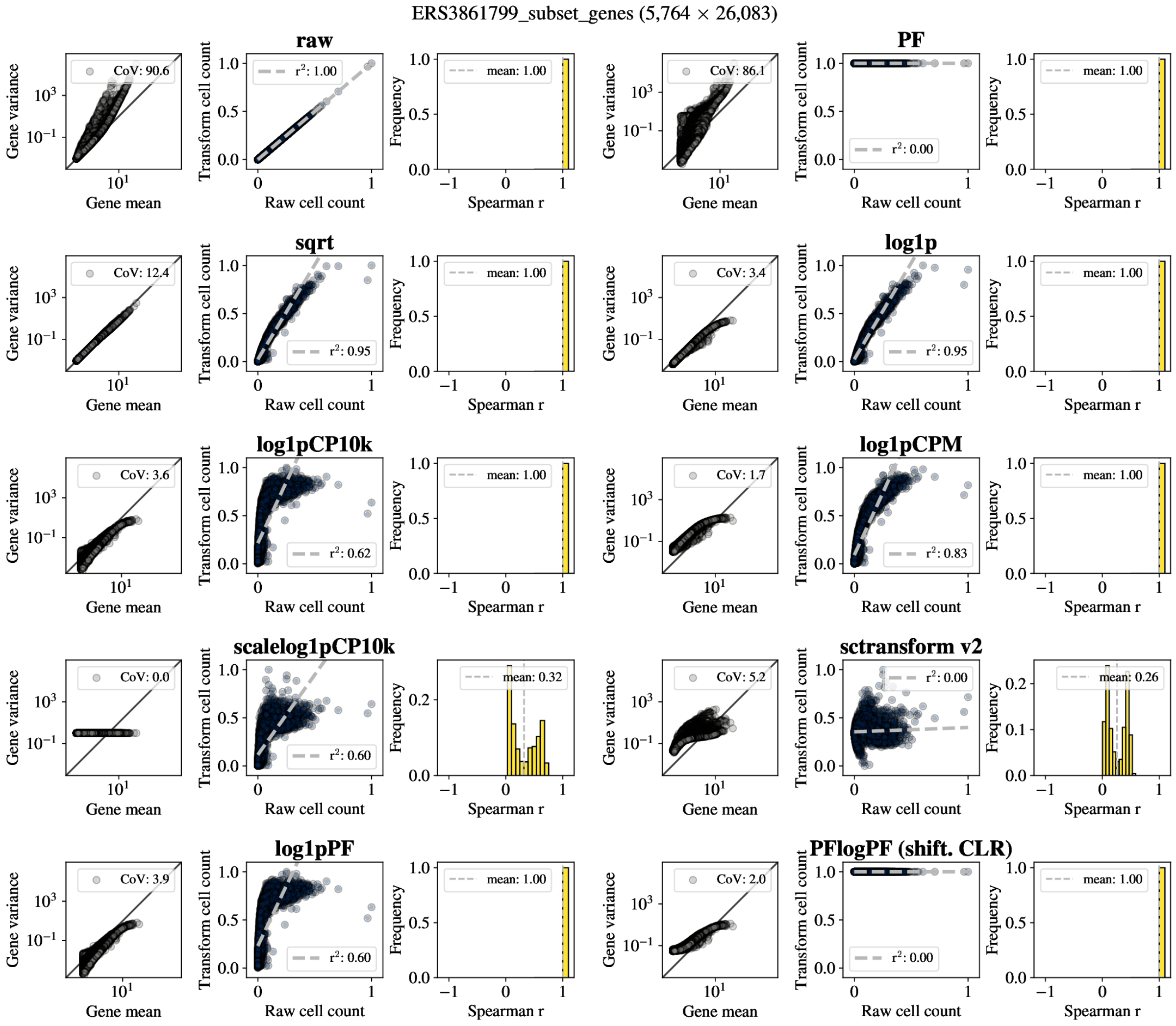


Supplementary Figure 1.47. **ERS3861799**. Per-method comparison of variance stabilization (left), depth normalization (middle), and monotonicity (right) for each normalization on dataset ERS3861799.

| ERS3861800 | |
|---|---|
| cells | 5,105 |
| genes | 60,623 |
| nonzero entries | 14,157,941 |
| density | 0.0457 |
| total counts | 58,168,377 |
| mean counts / cell | 11,394 |
| mean counts / gene | 960 |
| min counts / cell | 4,770 |
| max counts / cell | 50,175 |
| overdispersion | 0.918 |

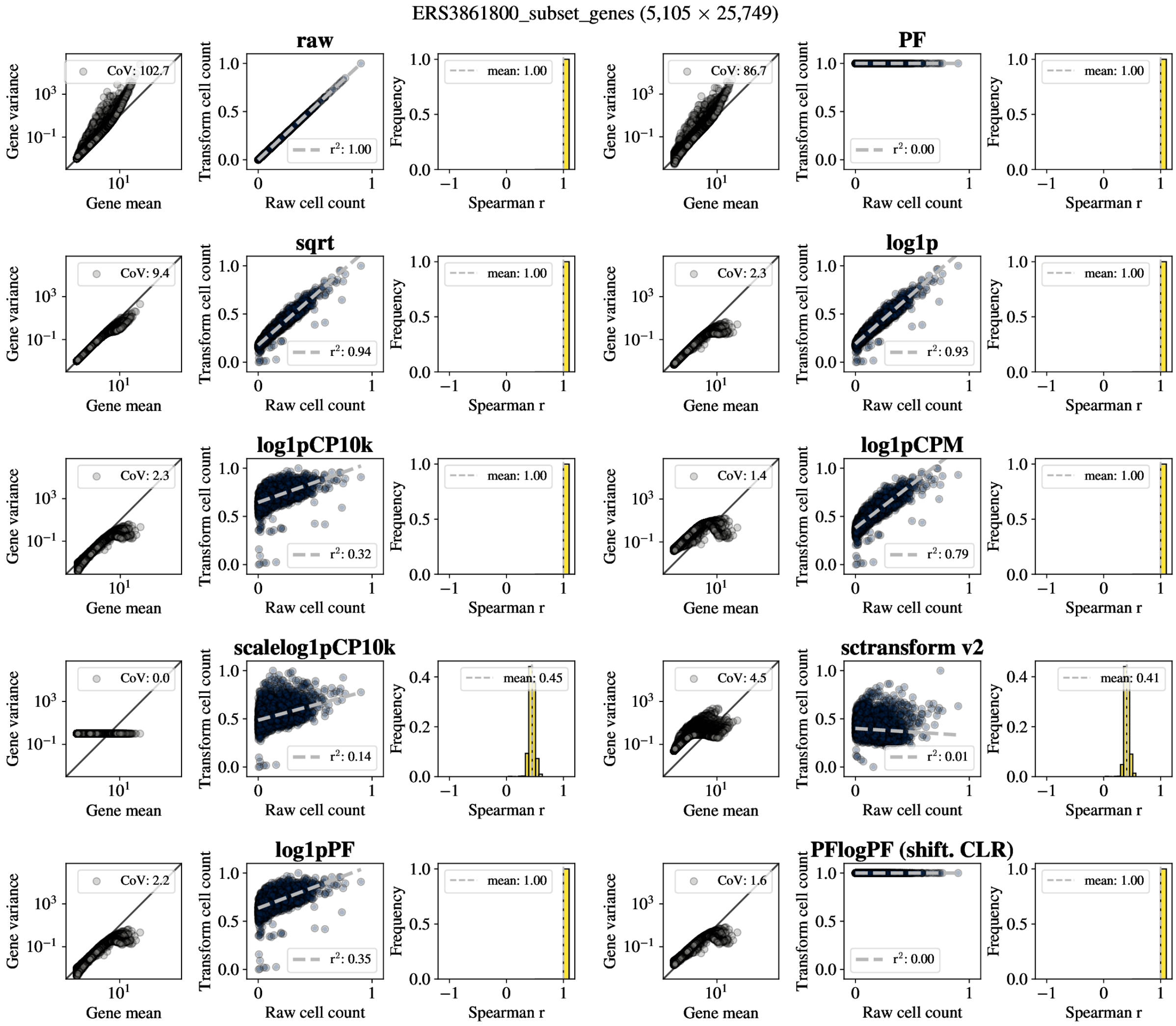


Supplementary Figure 1.48. **ERS3861800**. Per-method comparison of variance stabilization (left), depth normalization (middle), and monotonicity (right) for each normalization on dataset ERS3861800.

| **ERS3861801** | |
|---|---|
| cells | 6,473 |
| genes | 60,623 |
| nonzero entries | 14,110,039 |
| density | 0.036 |
| total counts | 51,327,746 |
| mean counts / cell | 7,930 |
| mean counts / gene | 847 |
| min counts / cell | 176 |
| max counts / cell | 50,135 |
| overdispersion | 0.968 |

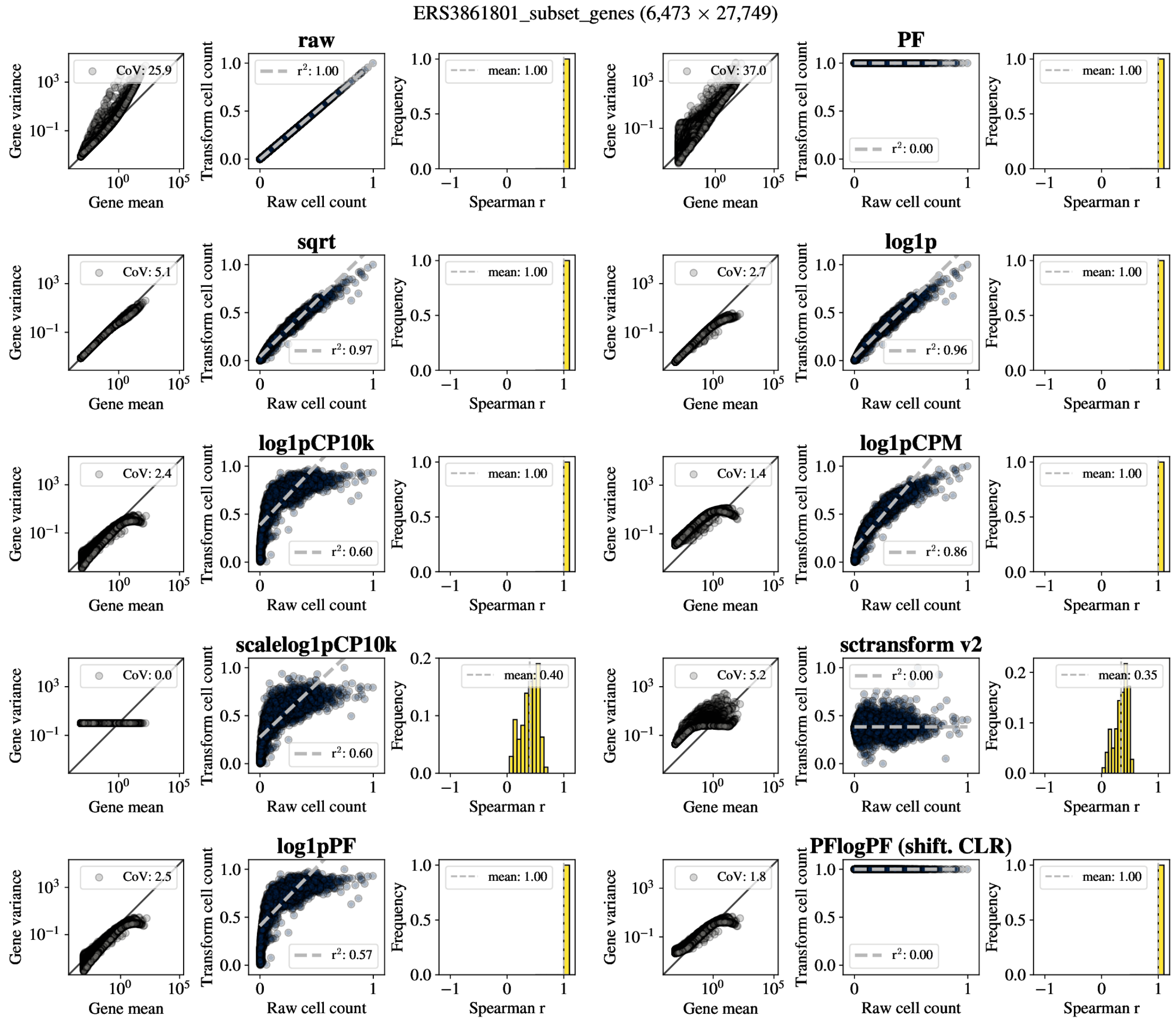


Supplementary Figure 1.49. **ERS3861801**. Per-method comparison of variance stabilization (left), depth normalization (middle), and monotonicity (right) for each normalization on dataset ERS3861801.

| ERS3861802 | |
|---|---|
| cells | 6,286 |
| genes | 60,623 |
| nonzero entries | 13,650,045 |
| density | 0.0358 |
| total counts | 49,468,625 |
| mean counts / cell | 7,870 |
| mean counts / gene | 816 |
| min counts / cell | 158 |
| max counts / cell | 45,161 |
| overdispersion | 0.925 |

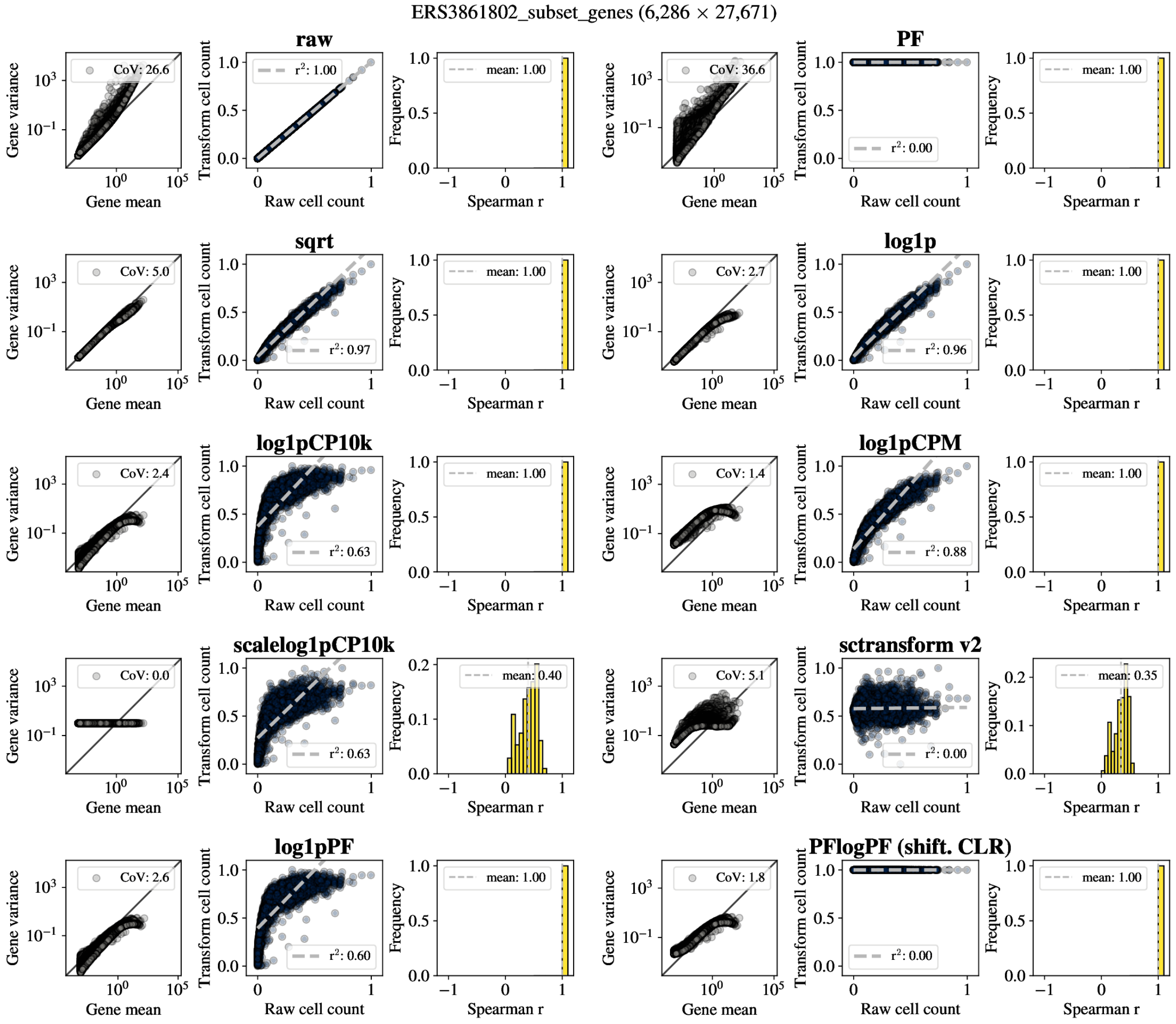


Supplementary Figure 1.50. **ERS3861802**. Per-method comparison of variance stabilization (left), depth normalization (middle), and monotonicity (right) for each normalization on dataset ERS3861802.

| **ERS3861803** | |
|---|---|
| cells | 7,791 |
| genes | 60,623 |
| nonzero entries | 17,897,449 |
| density | 0.0379 |
| total counts | 58,602,015 |
| mean counts / cell | 7,522 |
| mean counts / gene | 967 |
| min counts / cell | 184 |
| max counts / cell | 43,029 |
| overdispersion | 0.677 |

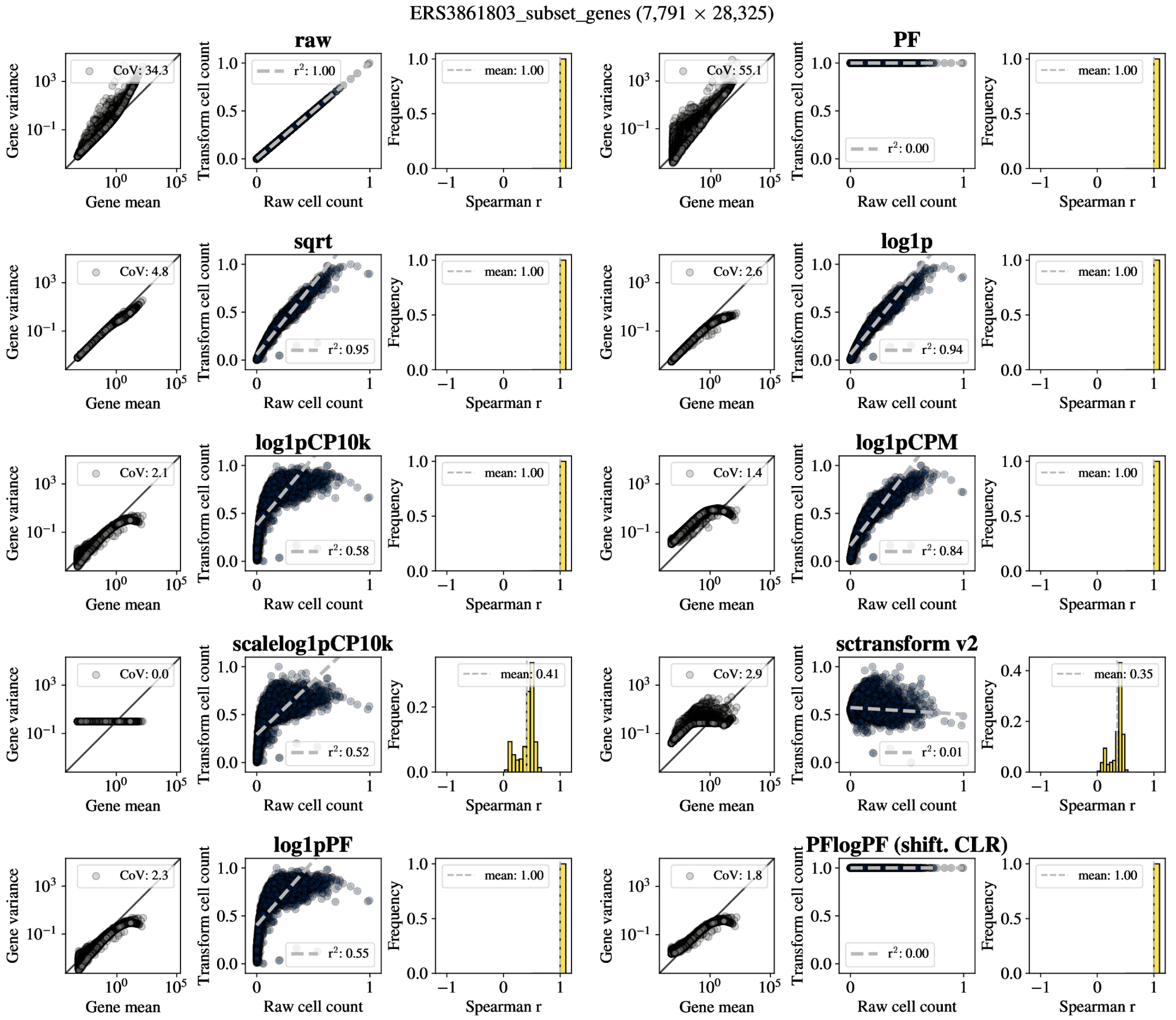


Supplementary Figure 1.51. **ERS3861803**. Per-method comparison of variance stabilization (left), depth normalization (middle), and monotonicity (right) for each normalization on dataset ERS3861803.

| **ERS3861804** | |
|---|---|
| cells | 7,633 |
| genes | 60,623 |
| nonzero entries | 13,800,911 |
| density | 0.0298 |
| total counts | 53,313,220 |
| mean counts / cell | 6,985 |
| mean counts / gene | 879 |
| min counts / cell | 121 |
| max counts / cell | 93,237 |
| overdispersion | 3.33 |

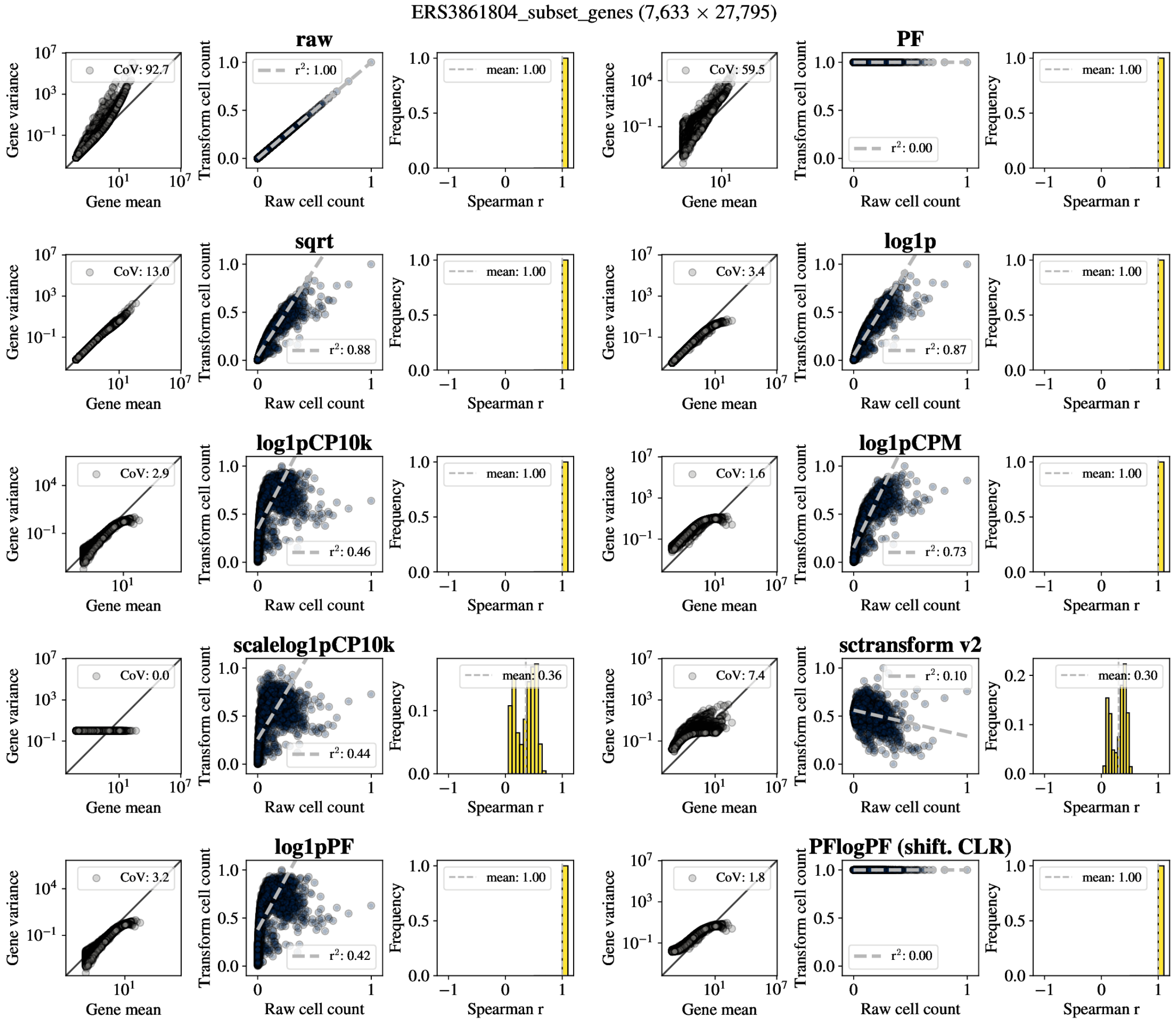


Supplementary Figure 1.52. **ERS3861804**. Per-method comparison of variance stabilization (left), depth normalization (middle), and monotonicity (right) for each normalization on dataset ERS3861804.

| ERS3861805 | |
|---|---|
| cells | 8,094 |
| genes | 60,623 |
| nonzero entries | 14,819,582 |
| density | 0.0302 |
| total counts | 56,420,966 |
| mean counts / cell | 6,971 |
| mean counts / gene | 931 |
| min counts / cell | 134 |
| max counts / cell | 62,110 |
| overdispersion | 2.93 |

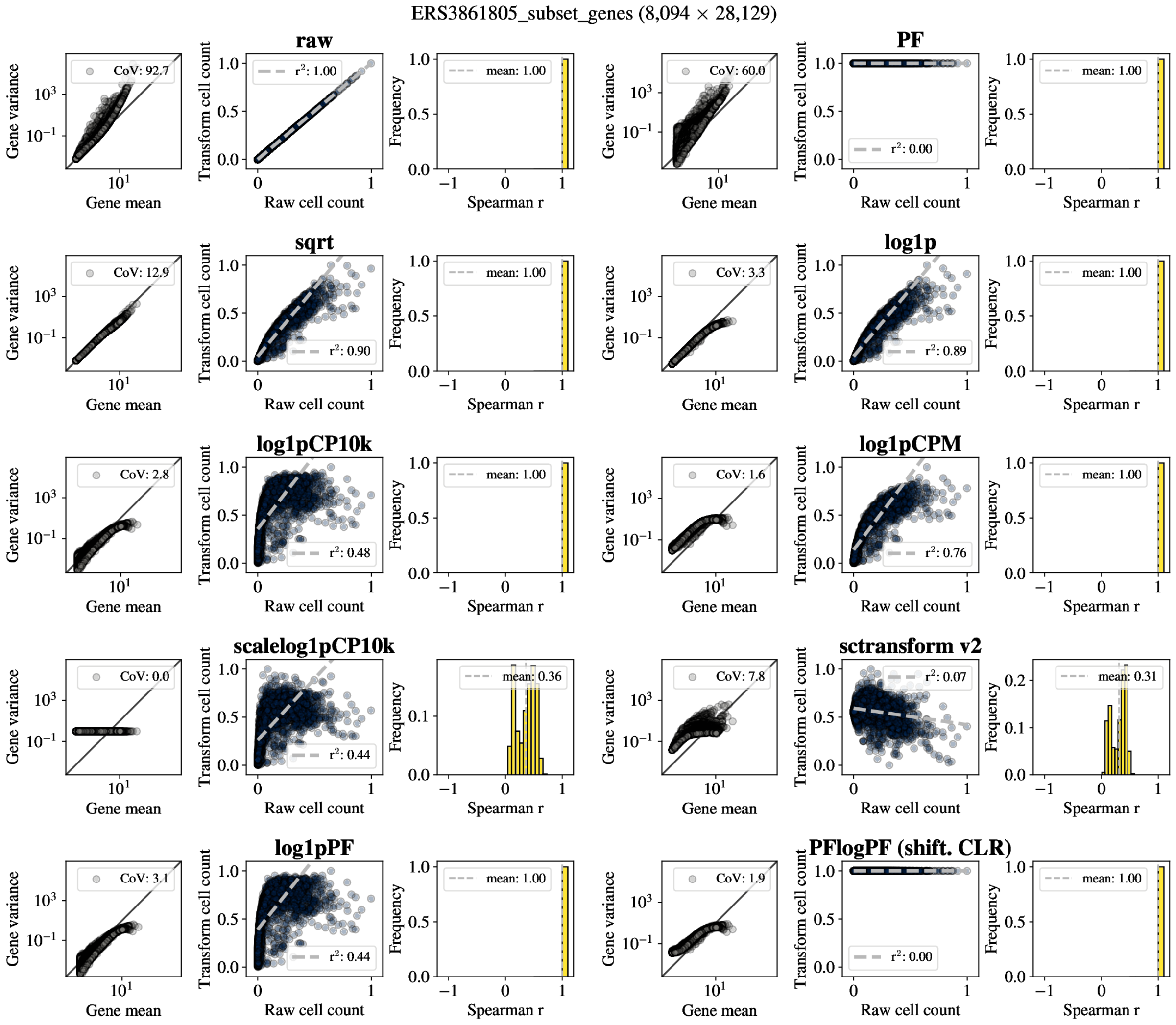


Supplementary Figure 1.53. **ERS3861805**. Per-method comparison of variance stabilization (left), depth normalization (middle), and monotonicity (right) for each normalization on dataset ERS3861805.

| ERS3861806 | |
|---|---|
| cells | 7,970 |
| genes | 60,623 |
| nonzero entries | 14,522,164 |
| density | 0.0301 |
| total counts | 56,456,060 |
| mean counts / cell | 7,084 |
| mean counts / gene | 931 |
| min counts / cell | 136 |
| max counts / cell | 69,435 |
| overdispersion | 2.93 |

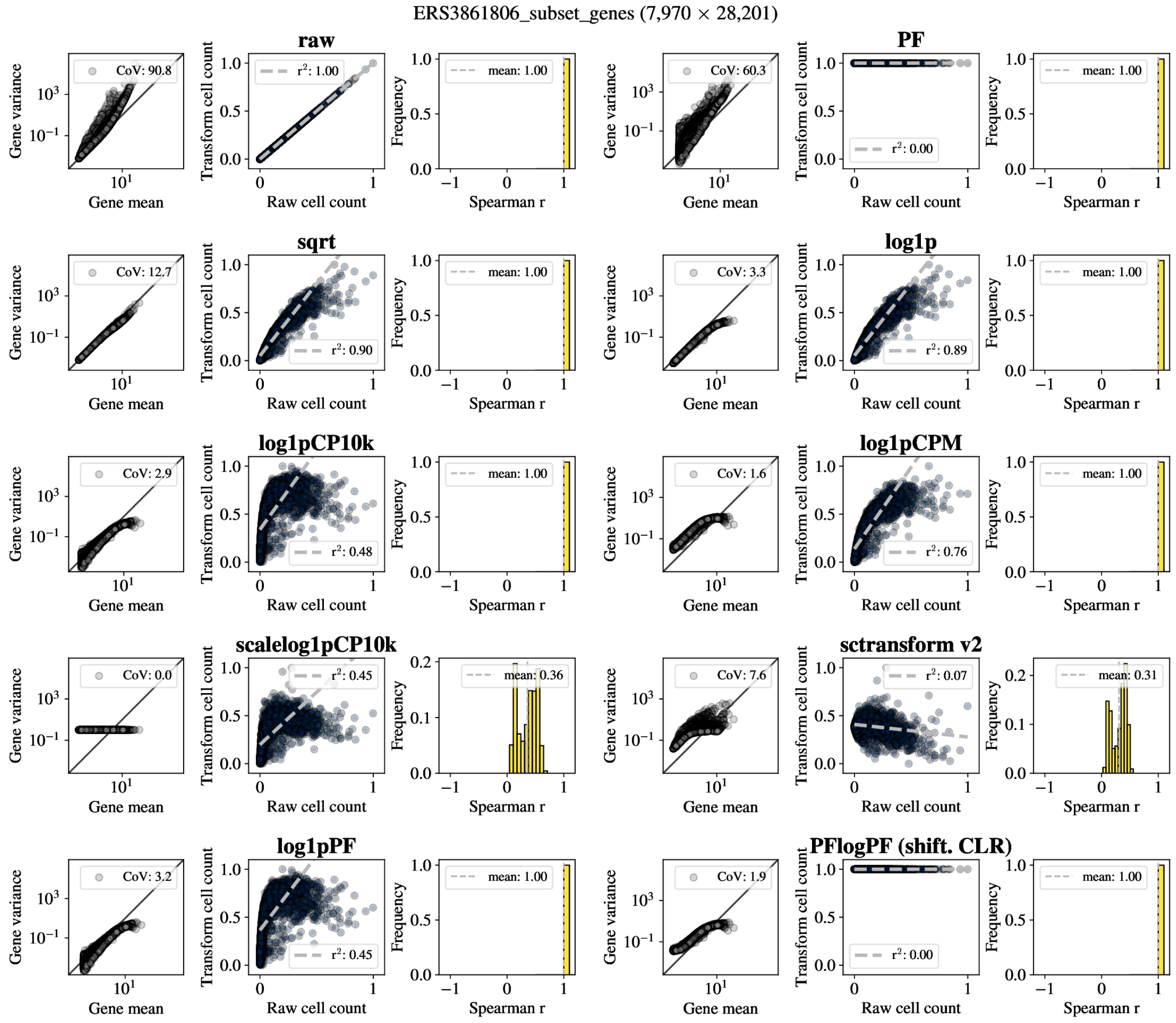


Supplementary Figure 1.54. **ERS3861806**. Per-method comparison of variance stabilization (left), depth normalization (middle), and monotonicity (right) for each normalization on dataset ERS3861806.

| **ERS3861807** | |
|---|---|
| cells | 6,601 |
| genes | 60,623 |
| nonzero entries | 16,511,484 |
| density | 0.0413 |
| total counts | 82,536,919 |
| mean counts / cell | 12,504 |
| mean counts / gene | 1,361 |
| min counts / cell | 381 |
| max counts / cell | 75,368 |
| overdispersion | 3.32 |

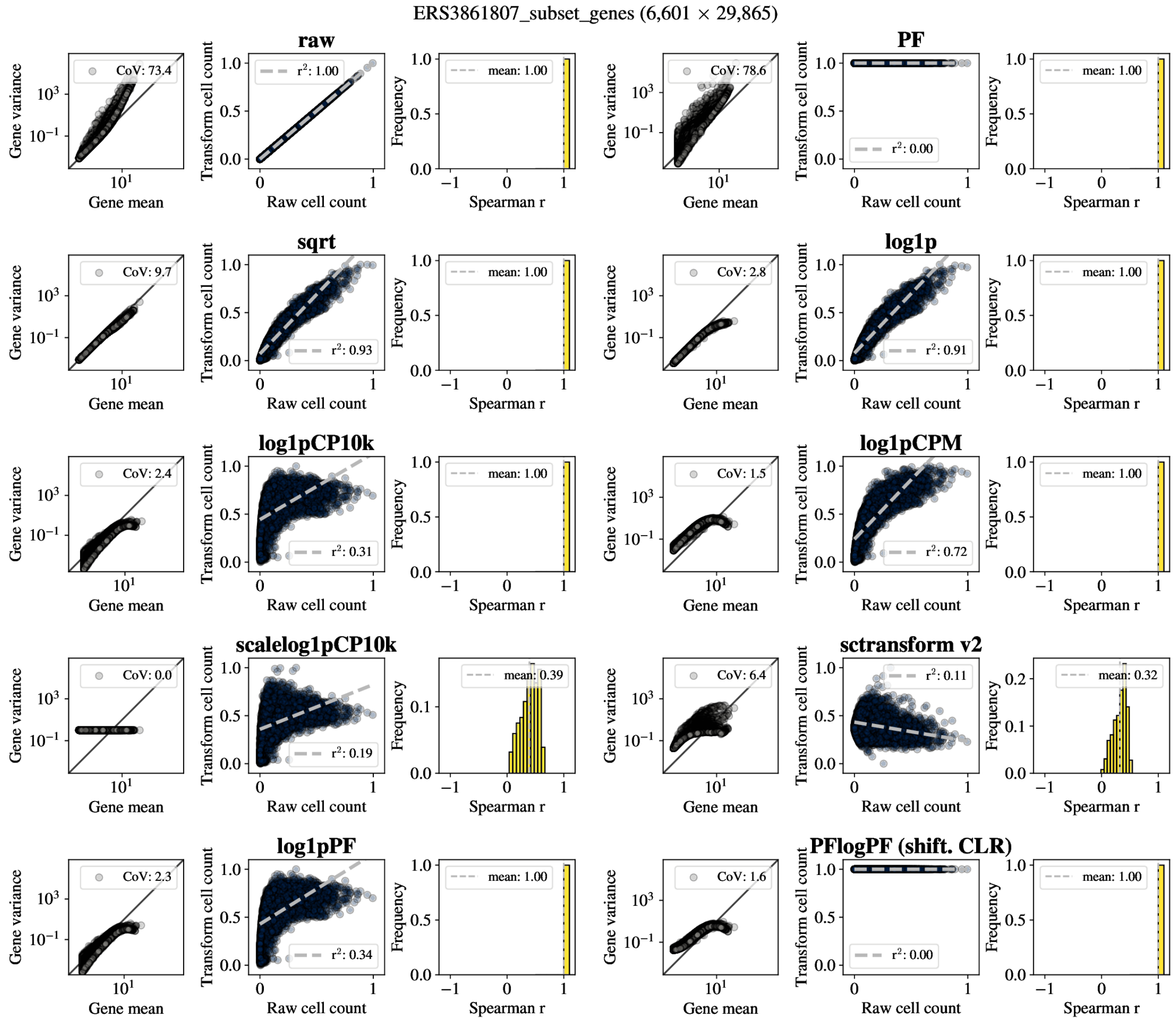


Supplementary Figure 1.55. **ERS3861807**. Per-method comparison of variance stabilization (left), depth normalization (middle), and monotonicity (right) for each normalization on dataset ERS3861807.

| ERS3861808 | |
|---|---|
| cells | 6,684 |
| genes | 60,623 |
| nonzero entries | 16,135,583 |
| density | 0.0398 |
| total counts | 84,162,850 |
| mean counts / cell | 12,592 |
| mean counts / gene | 1,388 |
| min counts / cell | 343 |
| max counts / cell | 89,165 |
| overdispersion | 3.28 |

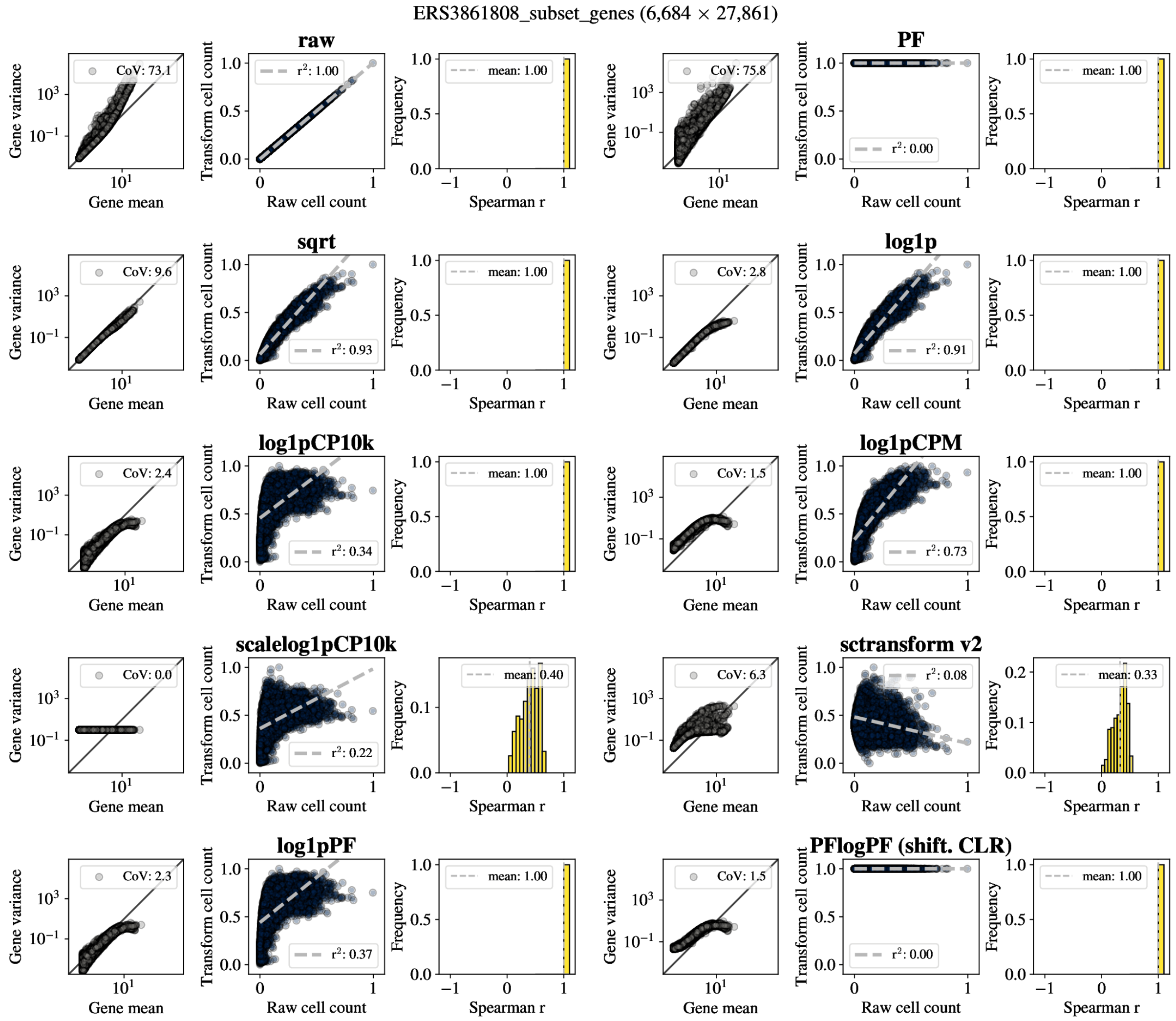


Supplementary Figure 1.56. **ERS3861808**. Per-method comparison of variance stabilization (left), depth normalization (middle), and monotonicity (right) for each normalization on dataset ERS3861808.

| ERS3861809 | |
|---|---|
| cells | 6,935 |
| genes | 60,623 |
| nonzero entries | 14,017,840 |
| density | 0.0333 |
| total counts | 54,591,014 |
| mean counts / cell | 7,872 |
| mean counts / gene | 901 |
| min counts / cell | 101 |
| max counts / cell | 64,807 |
| overdispersion | 2.63 |

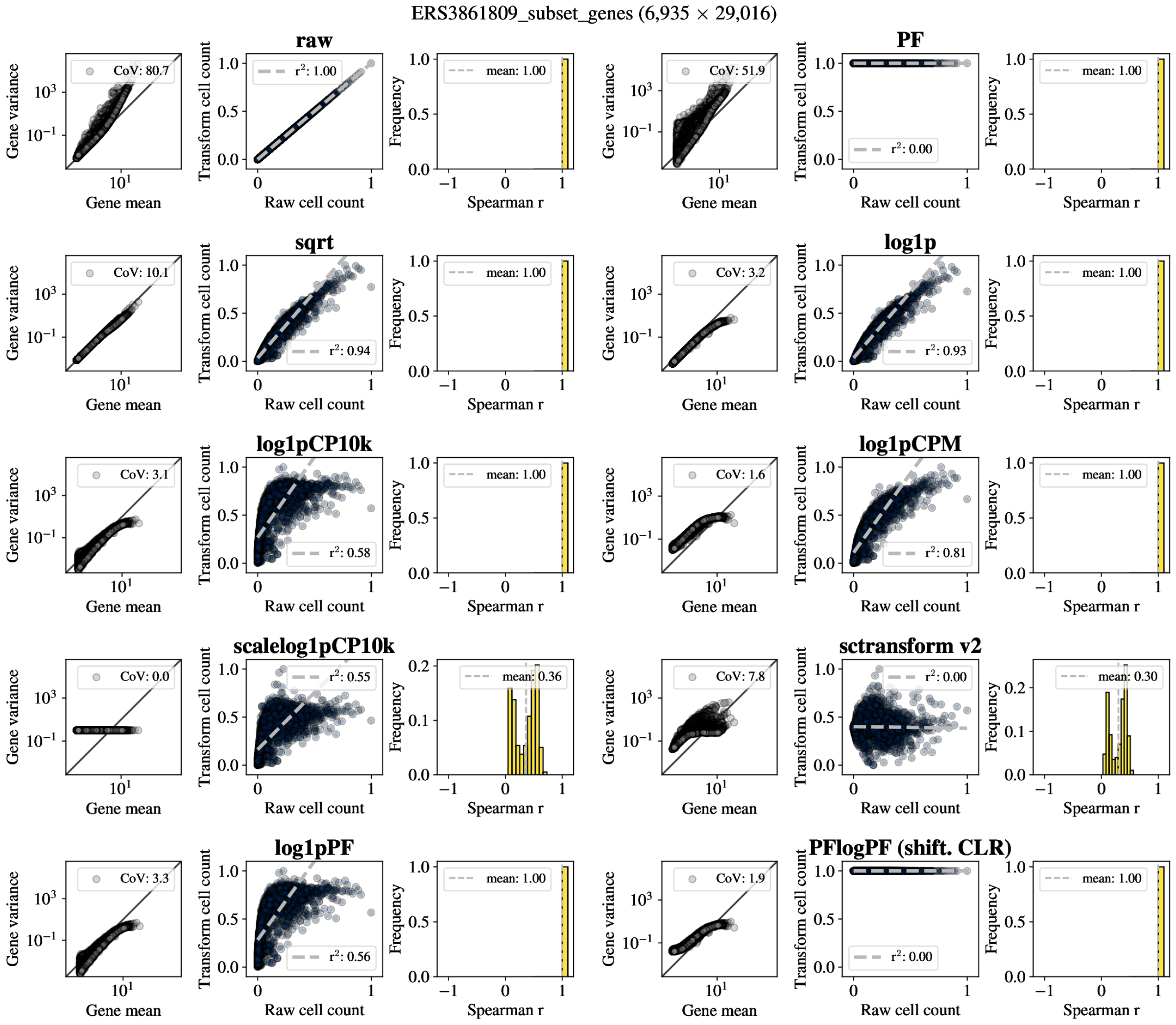


Supplementary Figure 1.57. **ERS3861809**. Per-method comparison of variance stabilization (left), depth normalization (middle), and monotonicity (right) for each normalization on dataset ERS3861809.

| ERS3861810 | |
|---|---|
| cells | 86,769 |
| genes | 60,623 |
| nonzero entries | 21,118,631 |
| density | 0.00401 |
| total counts | 90,942,272 |
| mean counts / cell | 1,048 |
| mean counts / gene | 1,500 |
| min counts / cell | 24 |
| max counts / cell | 116,644 |
| overdispersion | 32.5 |

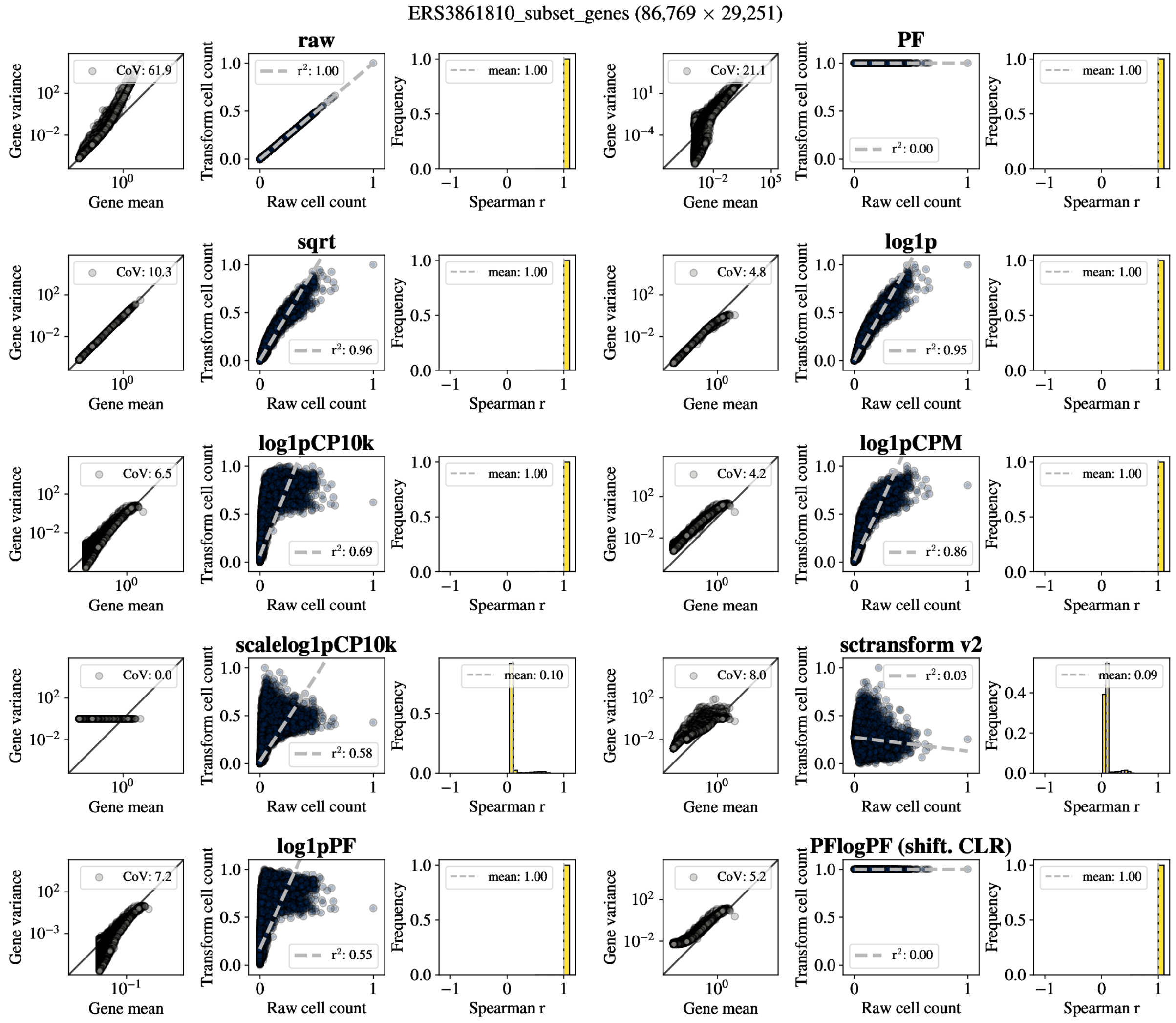


Supplementary Figure 1.58. **ERS3861810**. Per-method comparison of variance stabilization (left), depth normalization (middle), and monotonicity (right) for each normalization on dataset ERS3861810.

| **ERS3861811** | |
|---|---|
| cells | 3,325 |
| genes | 60,623 |
| nonzero entries | 4,661,604 |
| density | 0.0231 |
| total counts | 12,087,316 |
| mean counts / cell | 3,635 |
| mean counts / gene | 199 |
| min counts / cell | 483 |
| max counts / cell | 28,043 |
| overdispersion | 0.585 |

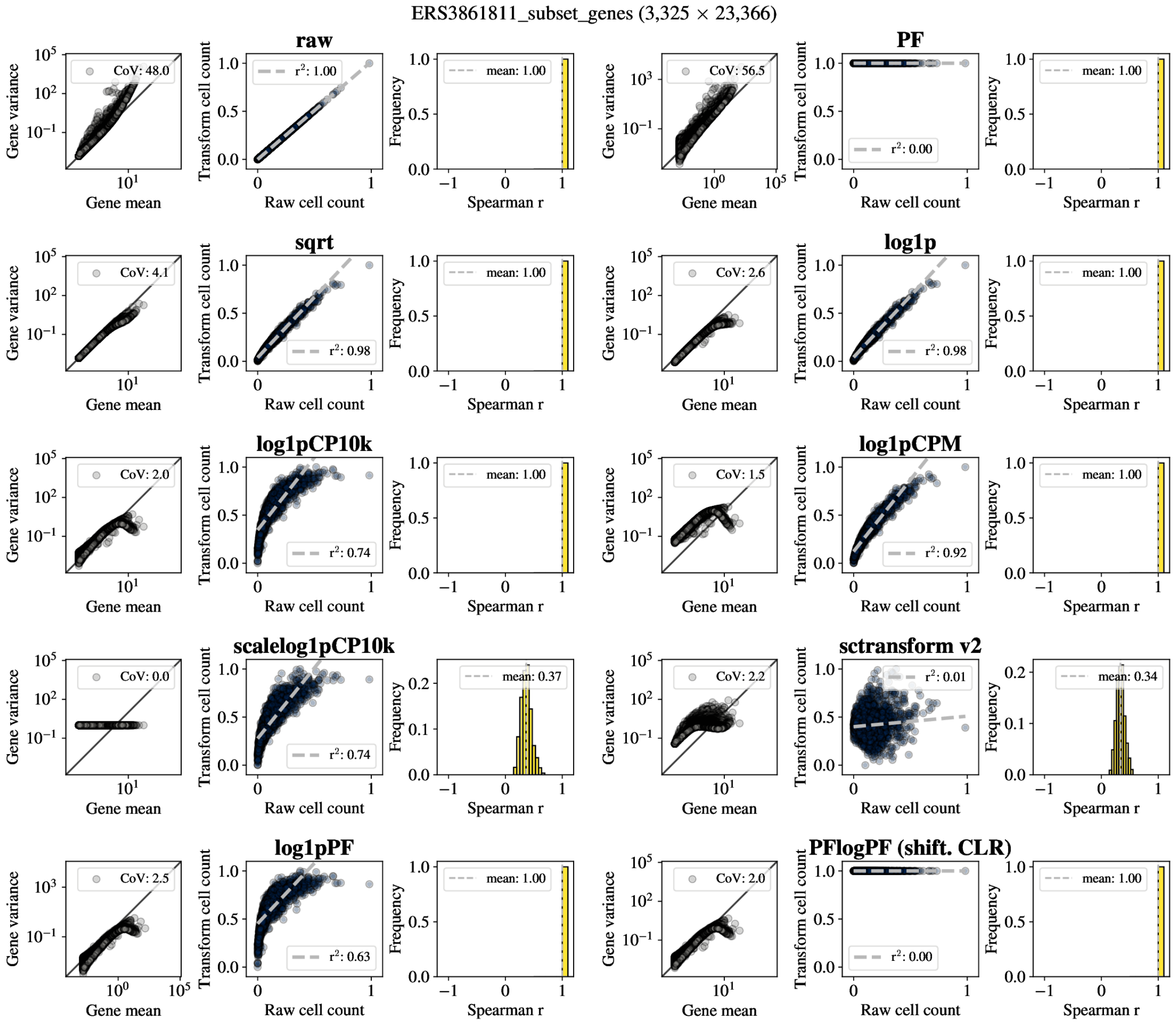


Supplementary Figure 1.59. **ERS3861811**. Per-method comparison of variance stabilization (left), depth normalization (middle), and monotonicity (right) for each normalization on dataset ERS3861811.

| **ERS3861813** | |
|---|---|
| cells | 5,218 |
| genes | 60,623 |
| nonzero entries | 930,754 |
| density | 0.00294 |
| total counts | 6,310,811 |
| mean counts / cell | 1,209 |
| mean counts / gene | 104 |
| min counts / cell | 13 |
| max counts / cell | 14,930 |
| overdispersion | 1.26 |

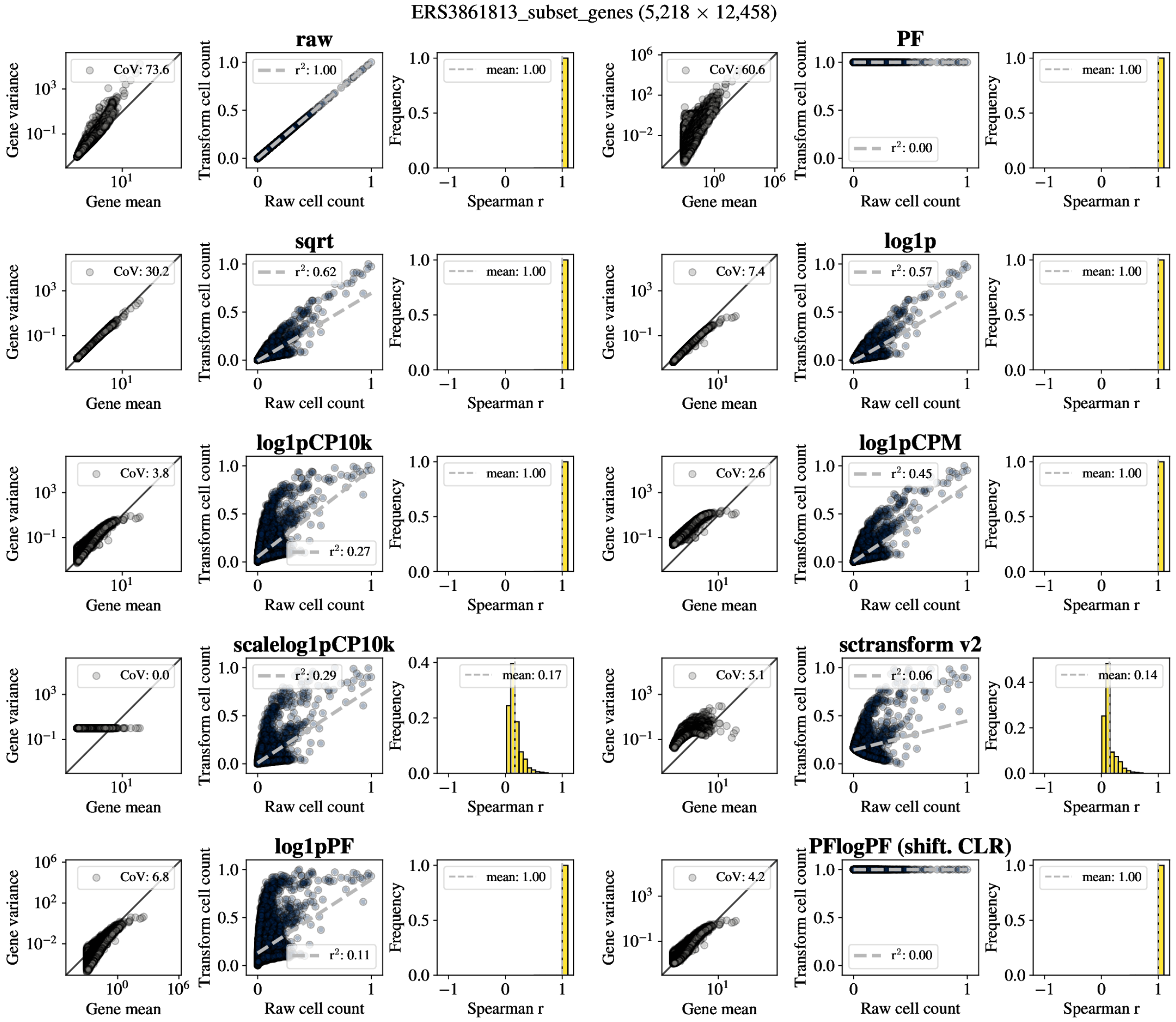


Supplementary Figure 1.60. **ERS3861813**. Per-method comparison of variance stabilization (left), depth normalization (middle), and monotonicity (right) for each normalization on dataset ERS3861813.

| ERS3861814 | |
|---|---|
| cells | 3,805 |
| genes | 60,623 |
| nonzero entries | 2,386,655 |
| density | 0.0103 |
| total counts | 5,038,419 |
| mean counts / cell | 1,324 |
| mean counts / gene | 83 |
| min counts / cell | 372 |
| max counts / cell | 5,702 |
| overdispersion | 2.06 |

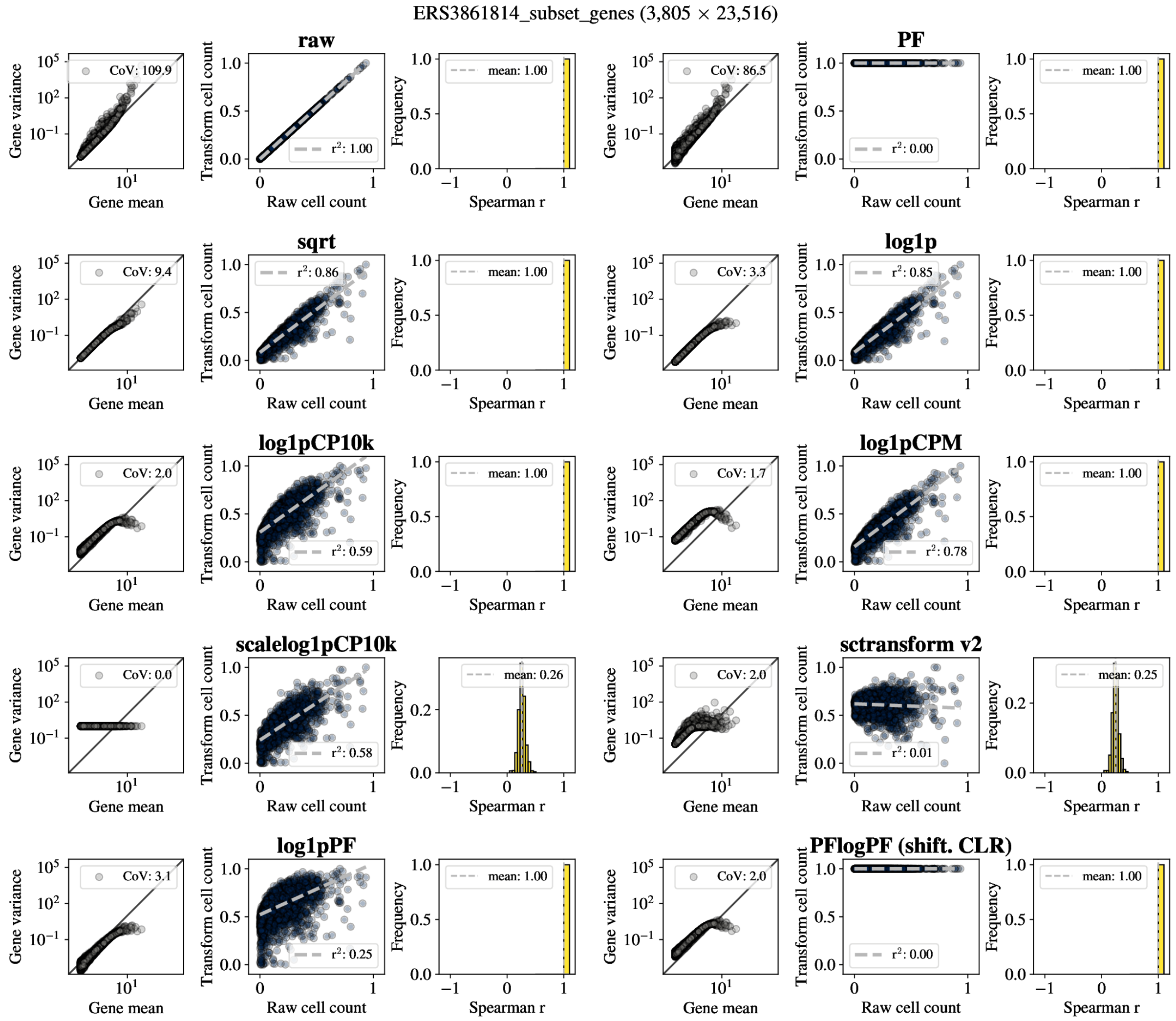


Supplementary Figure 1.61. **ERS3861814**. Per-method comparison of variance stabilization (left), depth normalization (middle), and monotonicity (right) for each normalization on dataset ERS3861814.

| ERS3861815 | |
|---|---|
| cells | 3,318 |
| genes | 60,623 |
| nonzero entries | 6,847,259 |
| density | 0.034 |
| total counts | 23,744,061 |
| mean counts / cell | 7,156 |
| mean counts / gene | 392 |
| min counts / cell | 130 |
| max counts / cell | 81,248 |
| overdispersion | 1.35 |

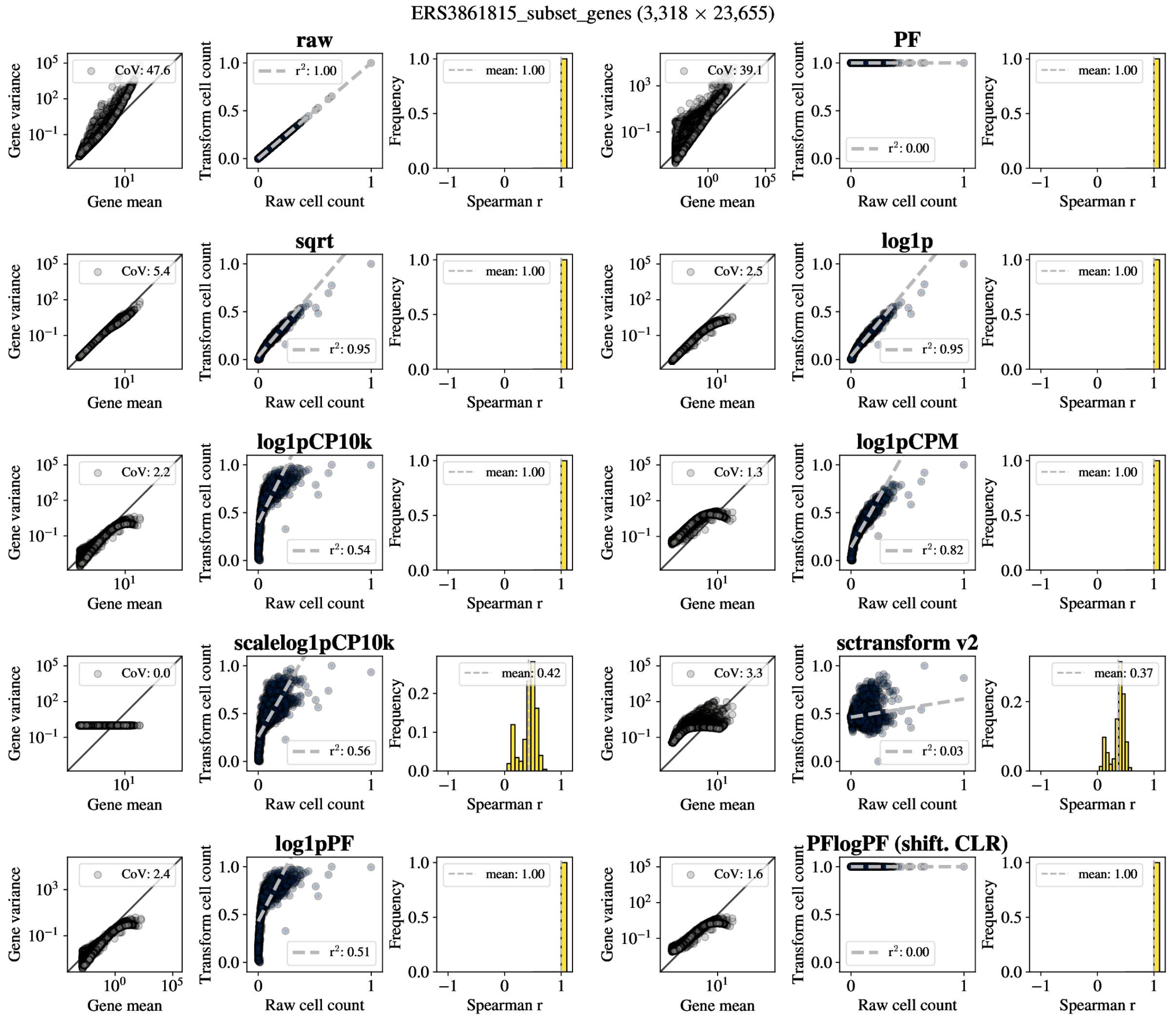


Supplementary Figure 1.62. **ERS3861815**. Per-method comparison of variance stabilization (left), depth normalization (middle), and monotonicity (right) for each normalization on dataset ERS3861815.

| **ERS3861816** | |
|---|---|
| cells | 7,621 |
| genes | 60,623 |
| nonzero entries | 19,047,167 |
| density | 0.0412 |
| total counts | 63,382,163 |
| mean counts / cell | 8,317 |
| mean counts / gene | 1,046 |
| min counts / cell | 228 |
| max counts / cell | 35,218 |
| overdispersion | 0.483 |

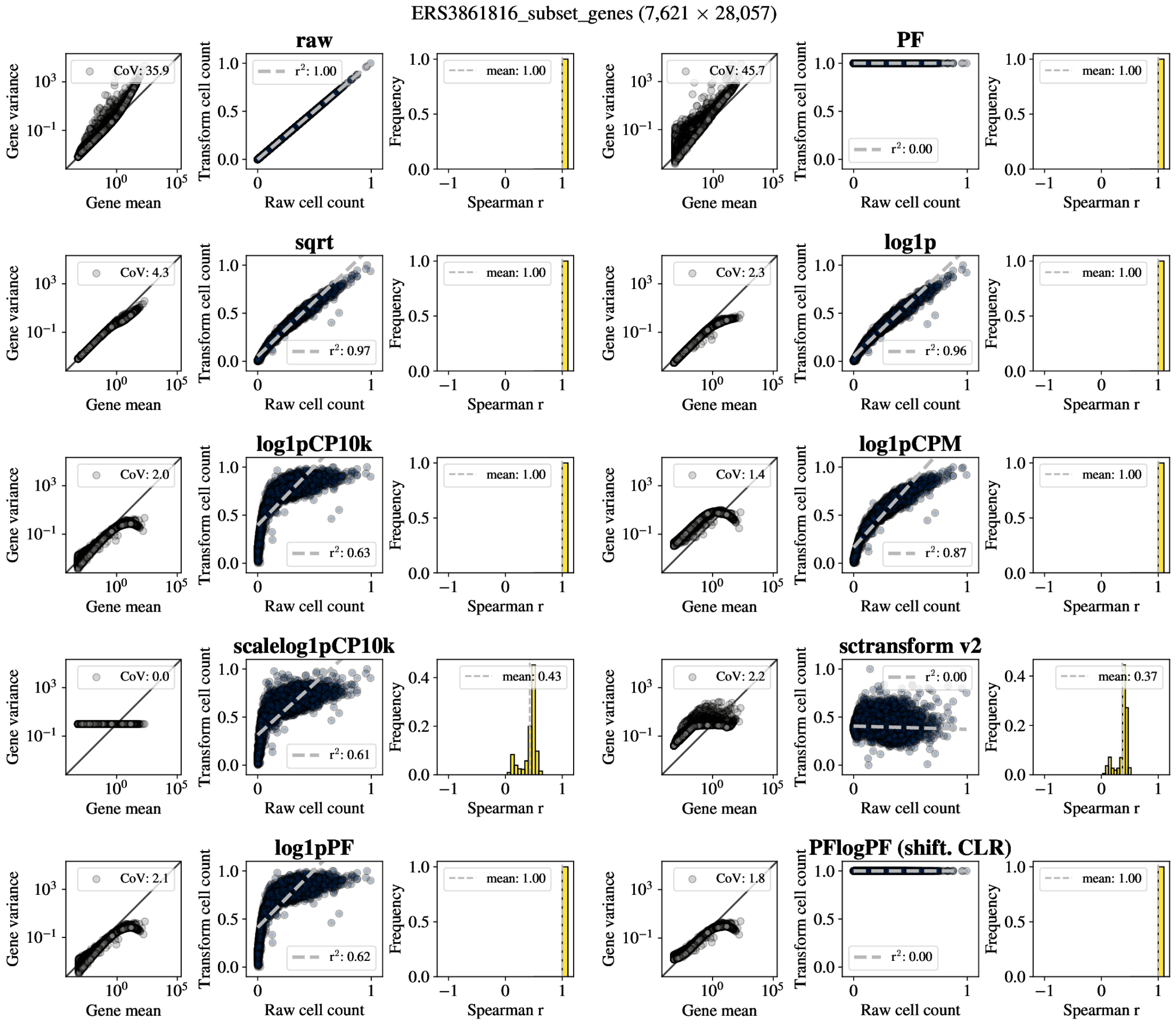


Supplementary Figure 1.63. **ERS3861816**. Per-method comparison of variance stabilization (left), depth normalization (middle), and monotonicity (right) for each normalization on dataset ERS3861816.

| ERS3861817 | |
|---|---|
| cells | 7,681 |
| genes | 60,623 |
| nonzero entries | 20,438,881 |
| density | 0.0439 |
| total counts | 69,235,047 |
| mean counts / cell | 9,014 |
| mean counts / gene | 1,142 |
| min counts / cell | 266 |
| max counts / cell | 40,379 |
| overdispersion | 0.455 |

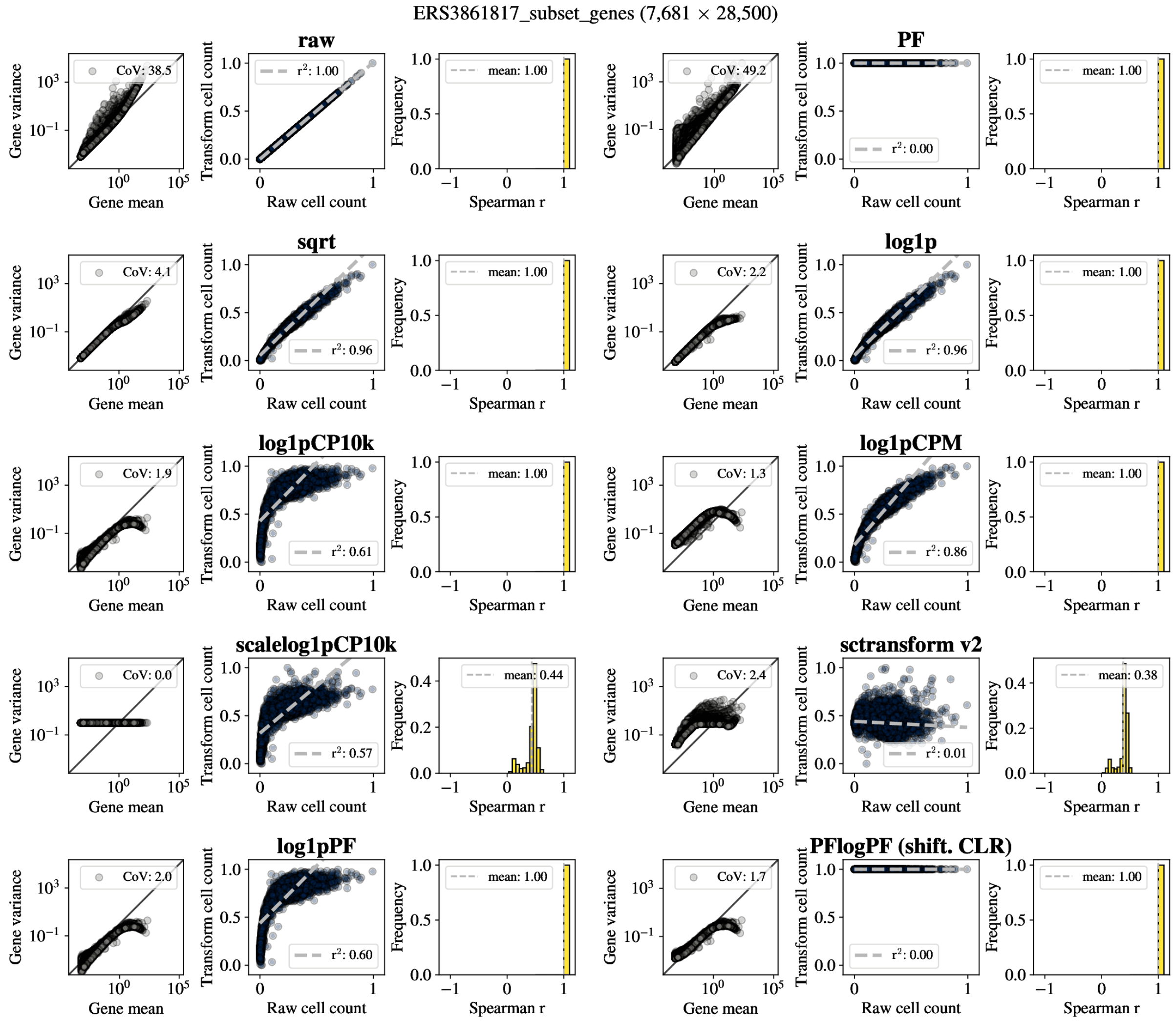


Supplementary Figure 1.64. **ERS3861817**. Per-method comparison of variance stabilization (left), depth normalization (middle), and monotonicity (right) for each normalization on dataset ERS3861817.

| ERS3861818 | |
|---|---|
| cells | 10,515 |
| genes | 60,623 |
| nonzero entries | 16,725,281 |
| density | 0.0262 |
| total counts | 61,665,658 |
| mean counts / cell | 5,865 |
| mean counts / gene | 1,017 |
| min counts / cell | 130 |
| max counts / cell | 133,389 |
| overdispersion | 2.88 |

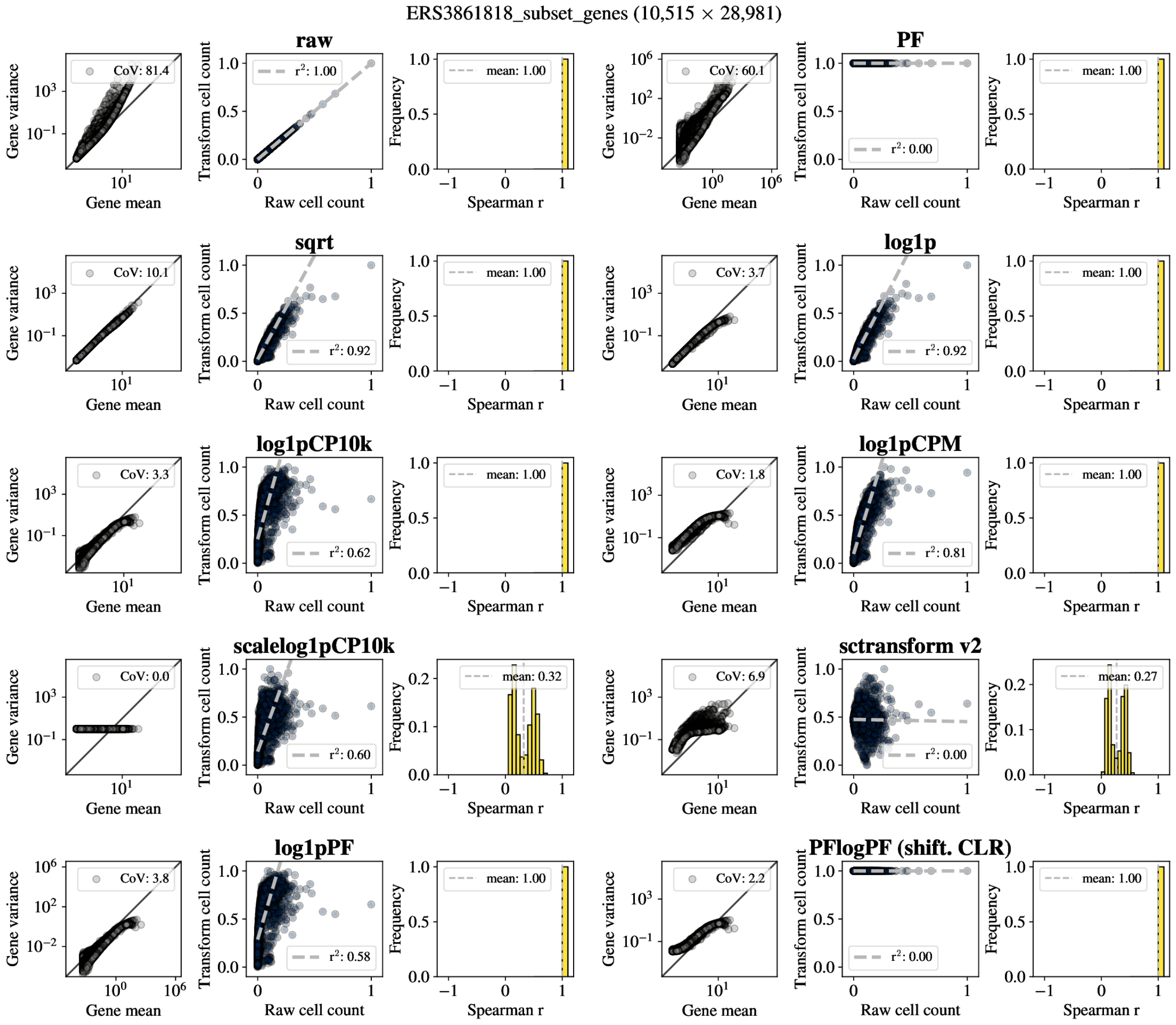


Supplementary Figure 1.65. **ERS3861818**. Per-method comparison of variance stabilization (left), depth normalization (middle), and monotonicity (right) for each normalization on dataset ERS3861818.

| ERS3861819 | |
|---|---|
| cells | 11,207 |
| genes | 60,623 |
| nonzero entries | 18,012,383 |
| density | 0.0265 |
| total counts | 66,098,917 |
| mean counts / cell | 5,898 |
| mean counts / gene | 1,090 |
| min counts / cell | 128 |
| max counts / cell | 83,093 |
| overdispersion | 2.75 |

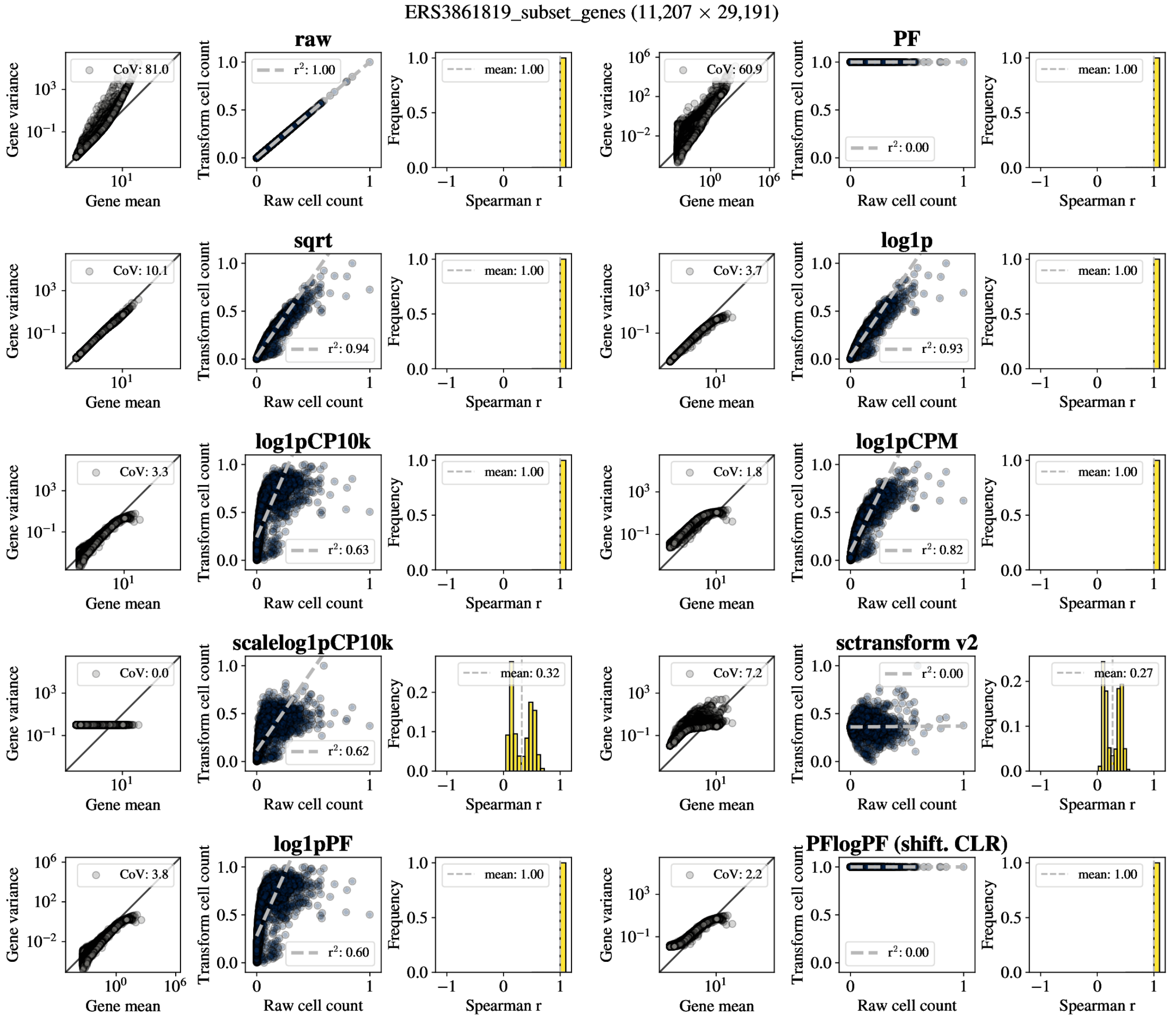


Supplementary Figure 1.66. **ERS3861819**. Per-method comparison of variance stabilization (left), depth normalization (middle), and monotonicity (right) for each normalization on dataset ERS3861819.

| **ERS3861820** | |
|---|---|
| cells | 6,831 |
| genes | 60,623 |
| nonzero entries | 17,312,512 |
| density | 0.0418 |
| total counts | 91,866,183 |
| mean counts / cell | 13,448 |
| mean counts / gene | 1,515 |
| min counts / cell | 262 |
| max counts / cell | 97,466 |
| overdispersion | 3.69 |

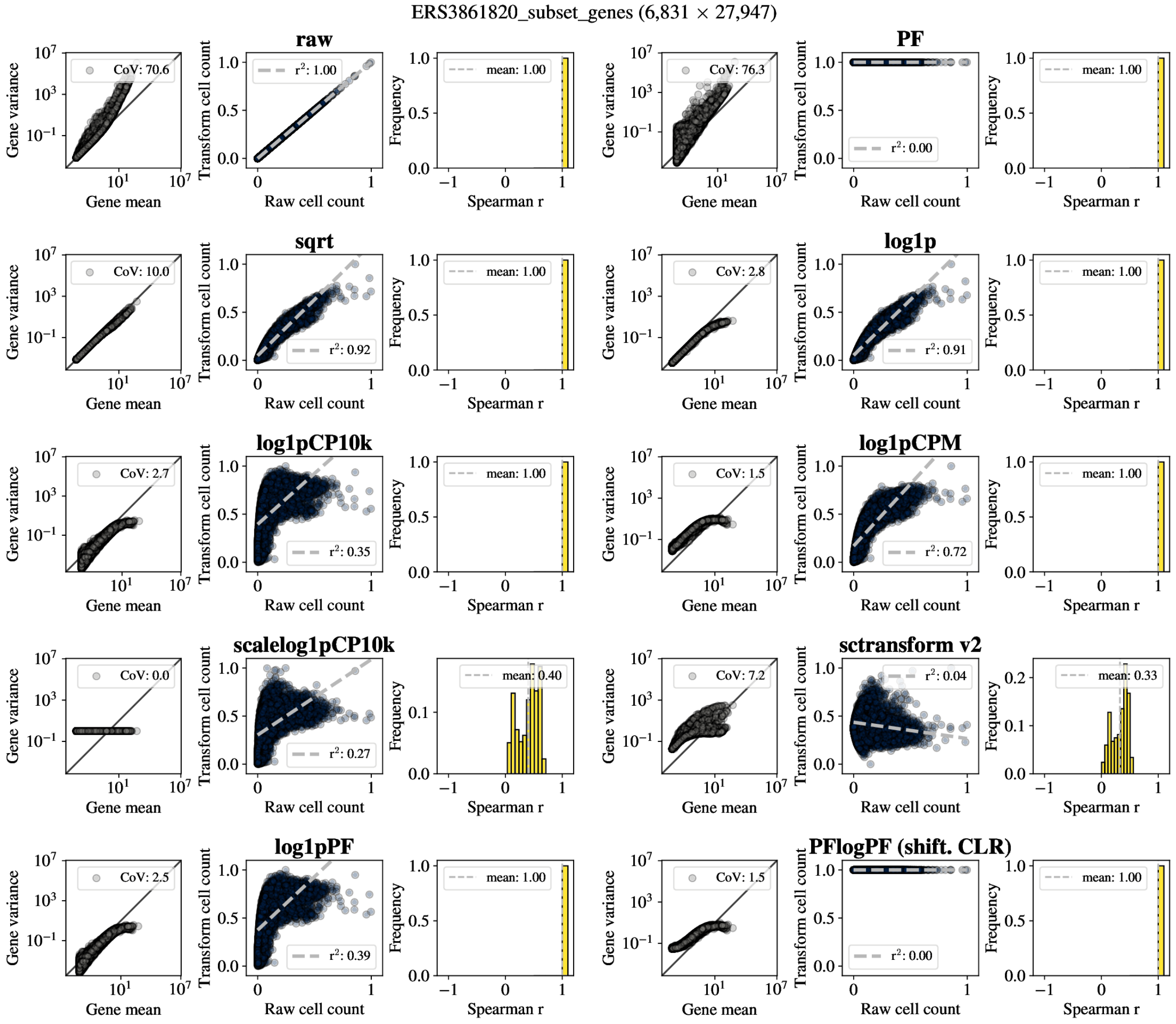


Supplementary Figure 1.67. **ERS3861820**. Per-method comparison of variance stabilization (left), depth normalization (middle), and monotonicity (right) for each normalization on dataset ERS3861820.

| **ERS3861821** | |
|---|---|
| cells | 7,041 |
| genes | 60,623 |
| nonzero entries | 17,543,455 |
| density | 0.0411 |
| total counts | 92,904,881 |
| mean counts / cell | 13,195 |
| mean counts / gene | 1,533 |
| min counts / cell | 282 |
| max counts / cell | 82,435 |
| overdispersion | 3.47 |

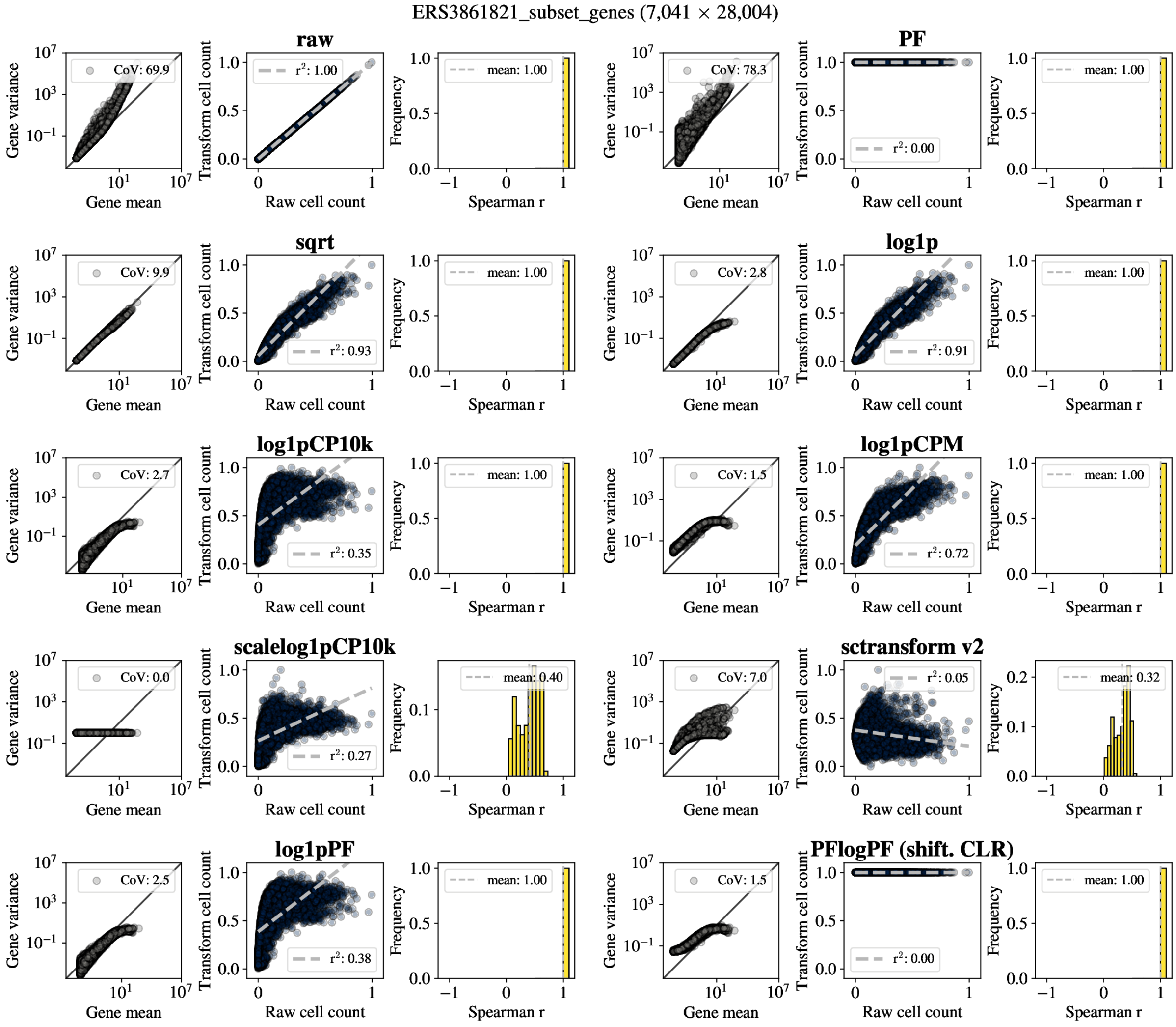


Supplementary Figure 1.68. **ERS3861821**. Per-method comparison of variance stabilization (left), depth normalization (middle), and monotonicity (right) for each normalization on dataset ERS3861821.

| **ERS3861822** | |
|---|---|
| cells | 10,285 |
| genes | 60,623 |
| nonzero entries | 16,461,002 |
| density | 0.0264 |
| total counts | 63,515,172 |
| mean counts / cell | 6,176 |
| mean counts / gene | 1,048 |
| min counts / cell | 113 |
| max counts / cell | 78,183 |
| overdispersion | 3.14 |

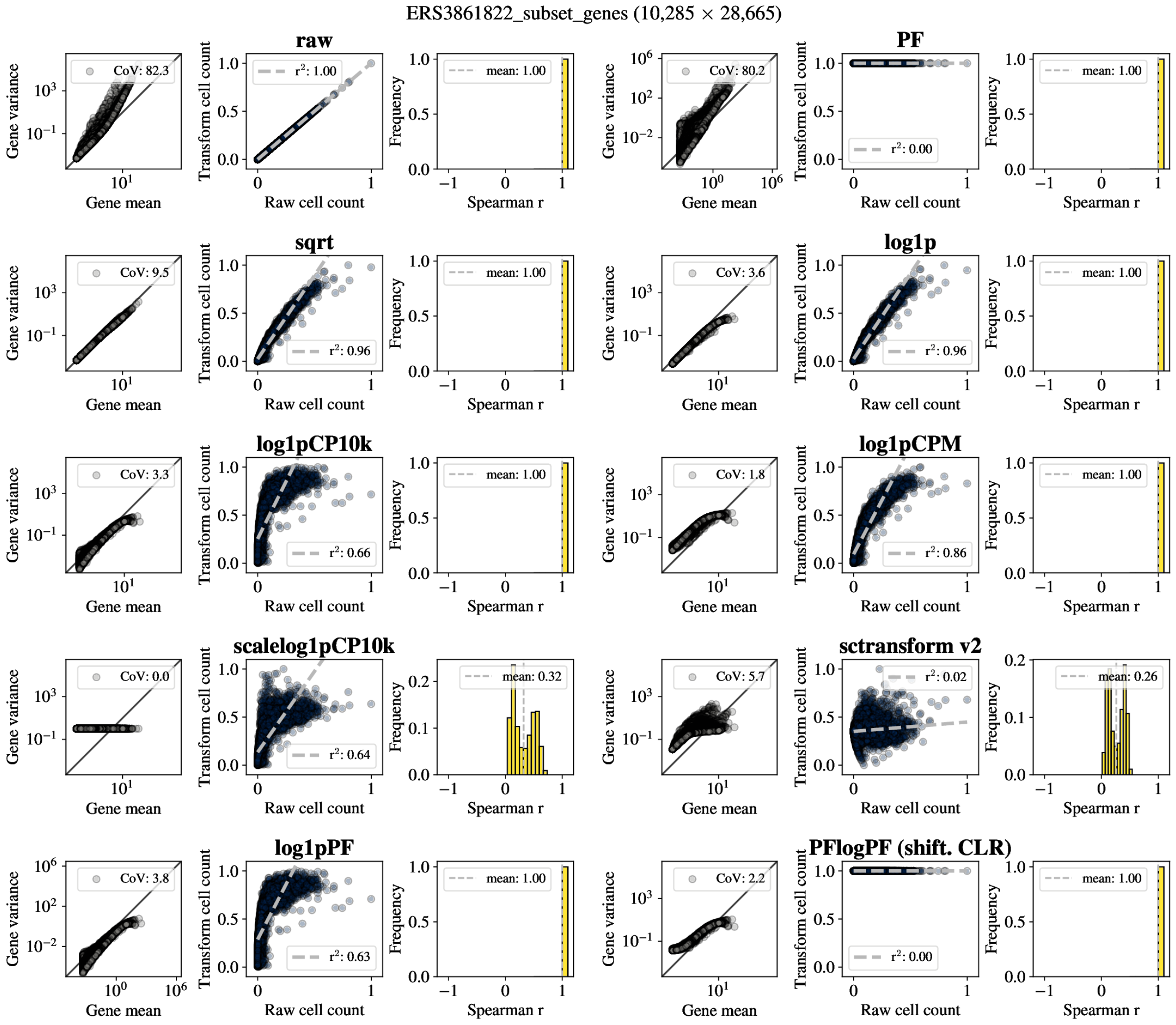


Supplementary Figure 1.69. **ERS3861822**. Per-method comparison of variance stabilization (left), depth normalization (middle), and monotonicity (right) for each normalization on dataset ERS3861822.

| **ERS3861823** | |
|---|---|
| cells | 3,490 |
| genes | 60,623 |
| nonzero entries | 7,311,793 |
| density | 0.0346 |
| total counts | 24,200,905 |
| mean counts / cell | 6,934 |
| mean counts / gene | 399 |
| min counts / cell | 1,706 |
| max counts / cell | 35,400 |
| overdispersion | 2.06 |

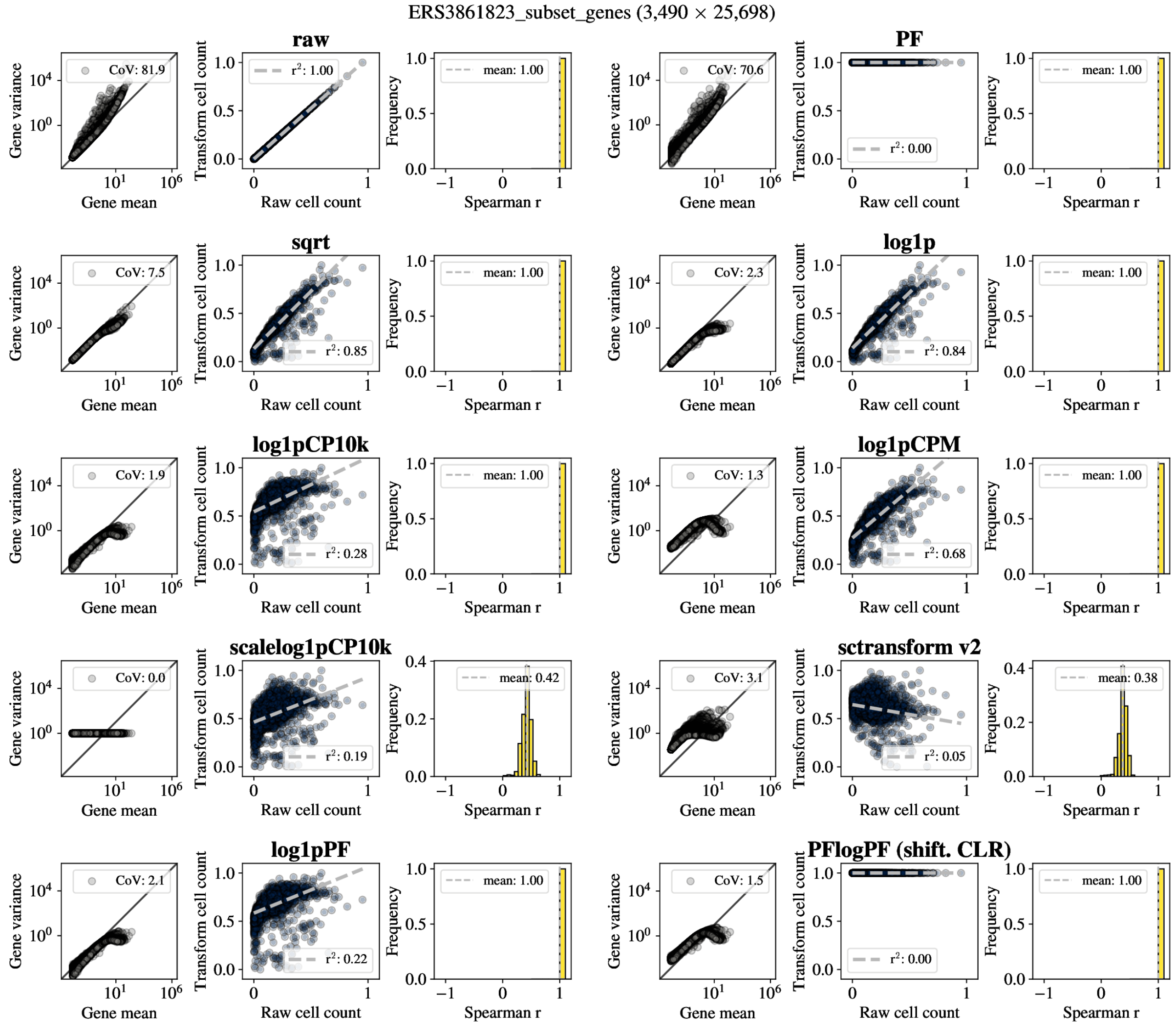


Supplementary Figure 1.70. **ERS3861823**. Per-method comparison of variance stabilization (left), depth normalization (middle), and monotonicity (right) for each normalization on dataset ERS3861823.

| ERS3861824 | |
|---|---|
| cells | 7,337 |
| genes | 60,623 |
| nonzero entries | 7,376,029 |
| density | 0.0166 |
| total counts | 24,793,917 |
| mean counts / cell | 3,379 |
| mean counts / gene | 409 |
| min counts / cell | 53 |
| max counts / cell | 47,730 |
| overdispersion | 4.66 |

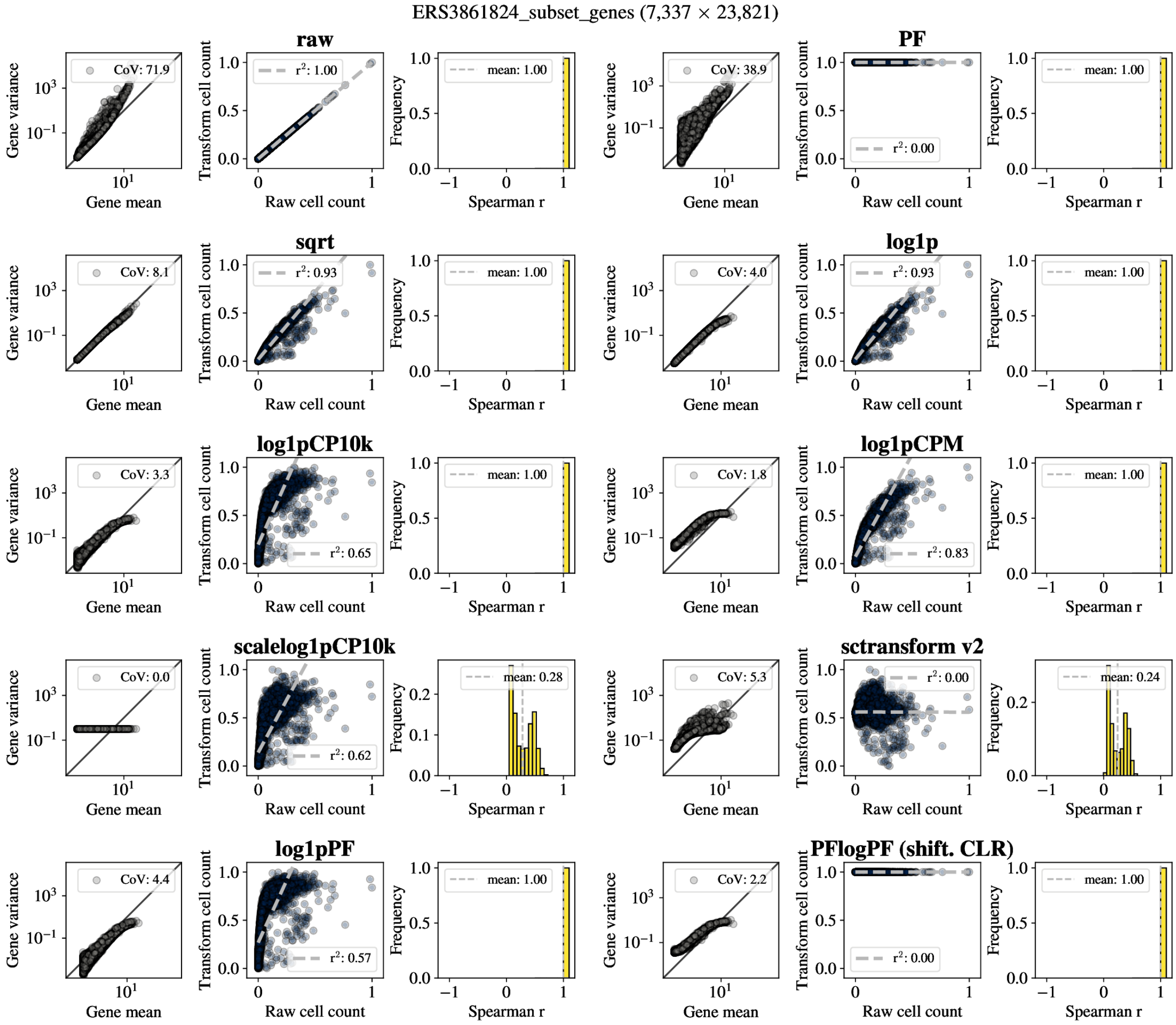


Supplementary Figure 1.71. **ERS3861824**. Per-method comparison of variance stabilization (left), depth normalization (middle), and monotonicity (right) for each normalization on dataset ERS3861824.

| ERS3861825 | |
|---|---|
| cells | 8,050 |
| genes | 60,623 |
| nonzero entries | 13,974,383 |
| density | 0.0286 |
| total counts | 63,231,002 |
| mean counts / cell | 7,855 |
| mean counts / gene | 1,043 |
| min counts / cell | 110 |
| max counts / cell | 69,640 |
| overdispersion | 4.97 |

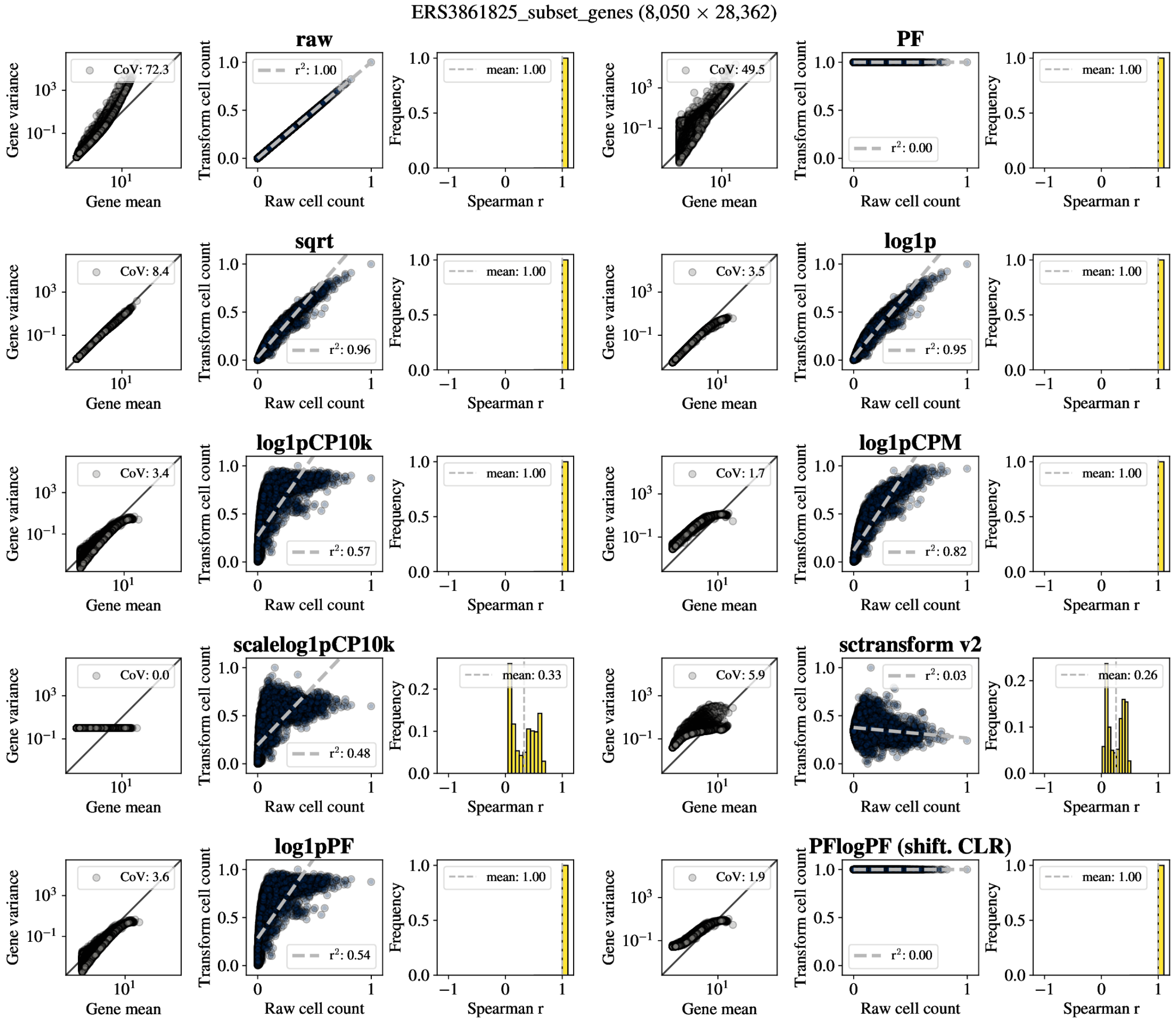


Supplementary Figure 1.72. **ERS3861825**. Per-method comparison of variance stabilization (left), depth normalization (middle), and monotonicity (right) for each normalization on dataset ERS3861825.

| **ERS3861826** | |
|---|---|
| cells | 4,495 |
| genes | 60,623 |
| nonzero entries | 6,045,171 |
| density | 0.0222 |
| total counts | 20,084,713 |
| mean counts / cell | 4,468 |
| mean counts / gene | 331 |
| min counts / cell | 55 |
| max counts / cell | 29,133 |
| overdispersion | 0.995 |

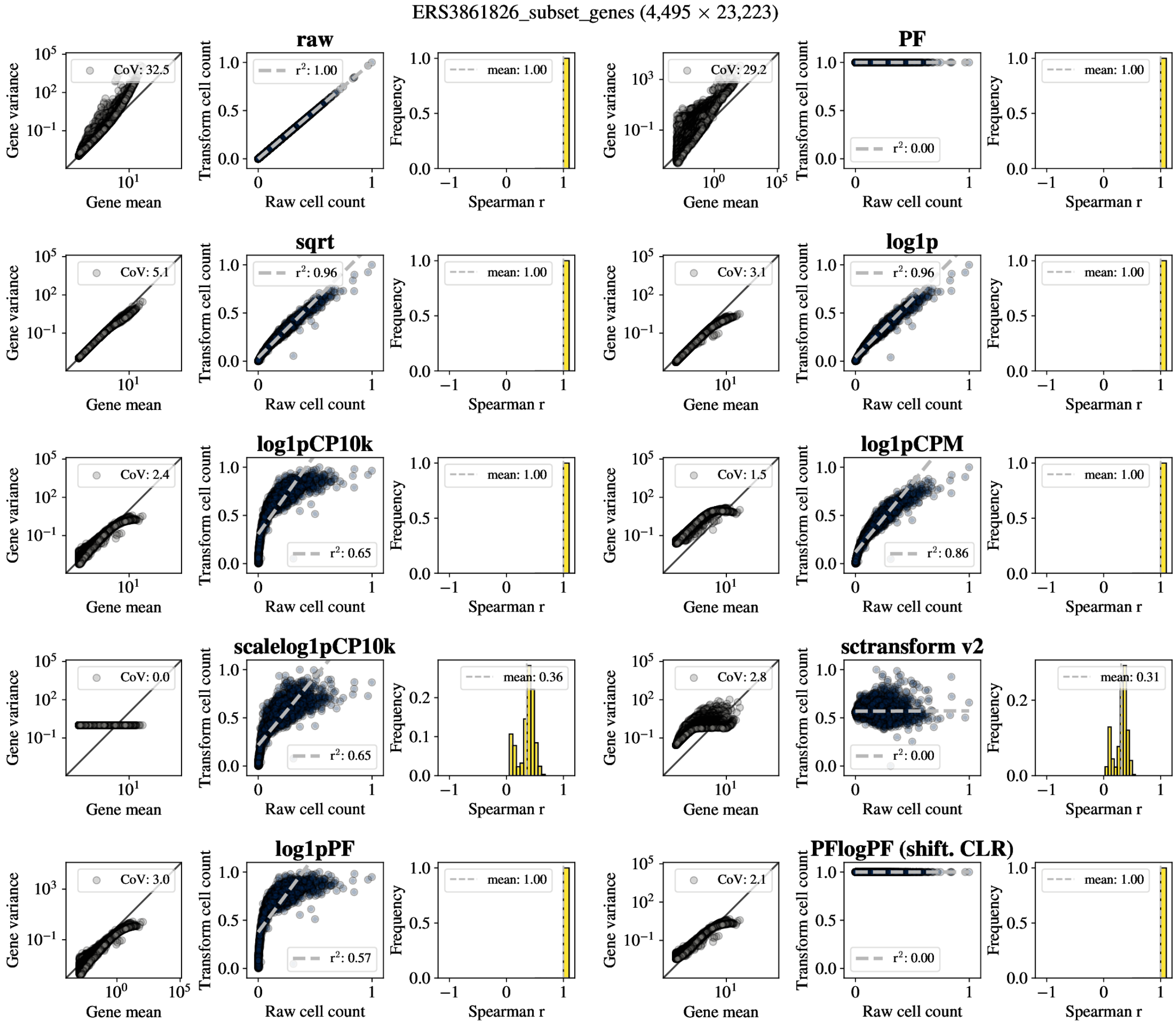


Supplementary Figure 1.73. **ERS3861826**. Per-method comparison of variance stabilization (left), depth normalization (middle), and monotonicity (right) for each normalization on dataset ERS3861826.

| **ERS3861827** | |
|---|---|
| cells | 1,914 |
| genes | 60,623 |
| nonzero entries | 6,020,763 |
| density | 0.0519 |
| total counts | 22,796,334 |
| mean counts / cell | 11,910 |
| mean counts / gene | 376 |
| min counts / cell | 8,844 |
| max counts / cell | 40,599 |
| overdispersion | 0.159 |

ERS3861827_subset_genes (1,914 × 23,664)

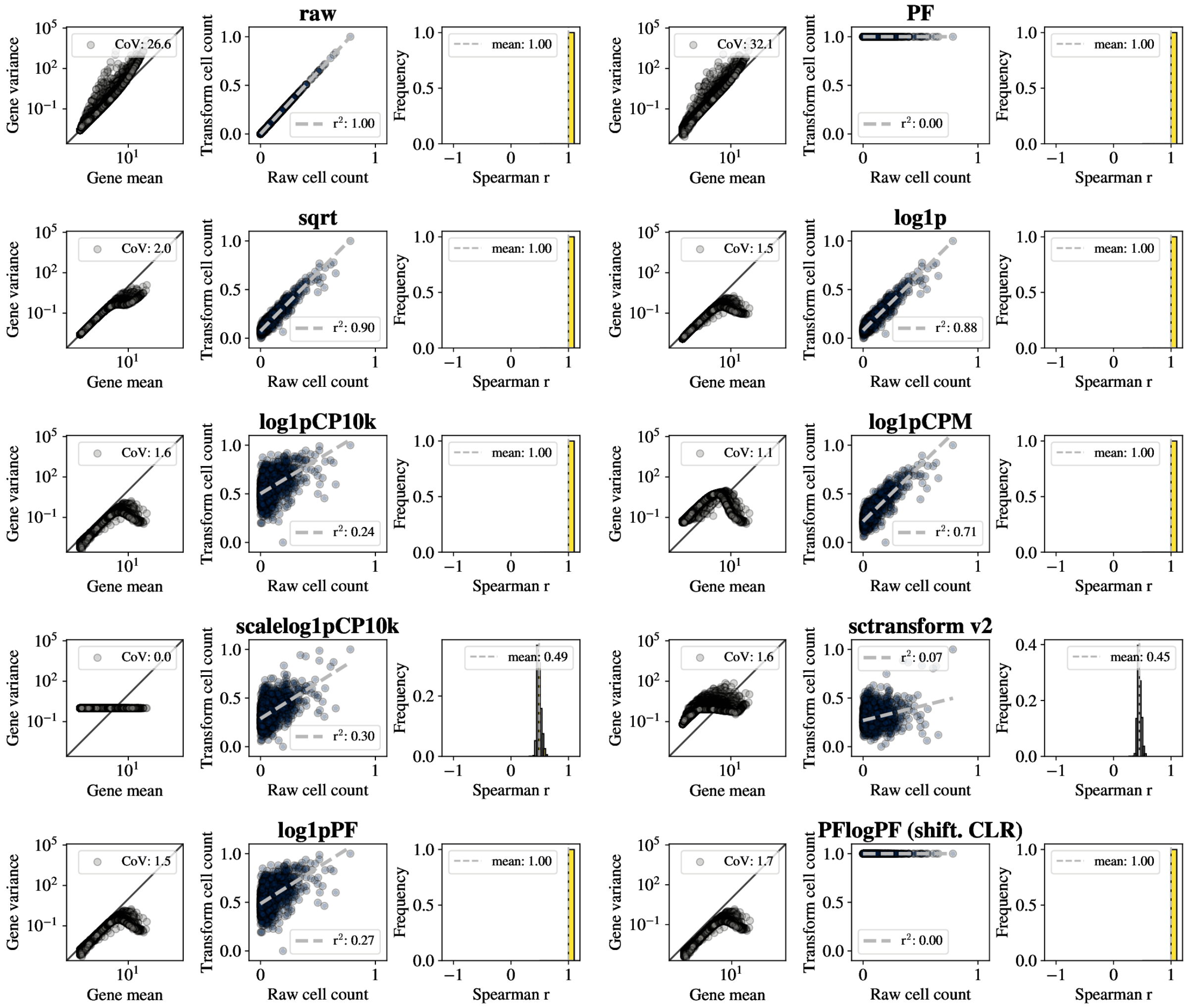


Supplementary Figure 1.74. **ERS3861827**. Per-method comparison of variance stabilization (left), depth normalization (middle), and monotonicity (right) for each normalization on dataset ERS3861827.

| ERS3861828 | |
|---|---|
| cells | 1,662 |
| genes | 60,623 |
| nonzero entries | 2,815,939 |
| density | 0.0279 |
| total counts | 8,572,740 |
| mean counts / cell | 5,158 |
| mean counts / gene | 141 |
| min counts / cell | 2,460 |
| max counts / cell | 25,330 |
| overdispersion | 0.377 |

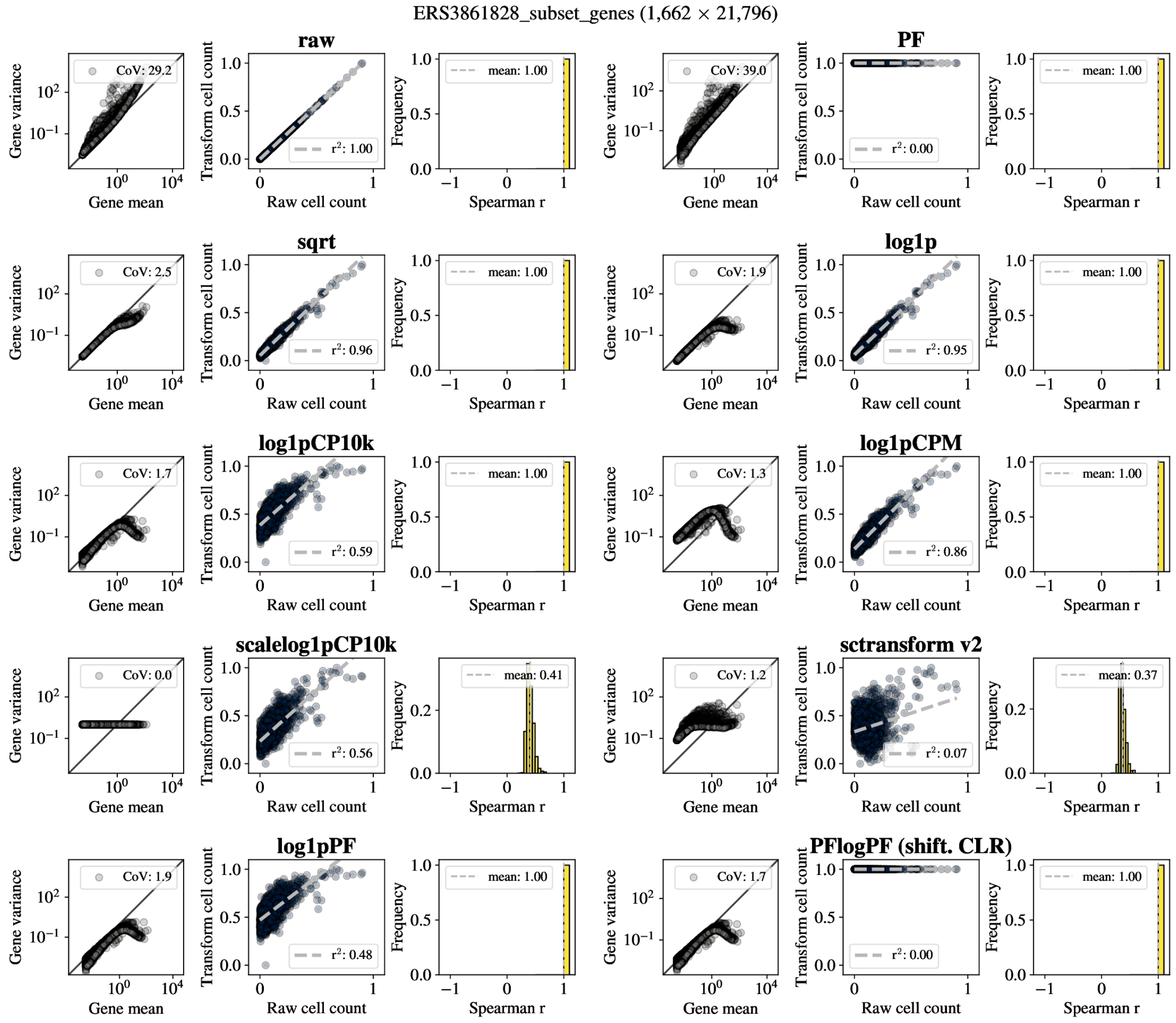


Supplementary Figure 1.75. **ERS3861828**. Per-method comparison of variance stabilization (left), depth normalization (middle), and monotonicity (right) for each normalization on dataset ERS3861828.

| **ERS3861829** | |
|---|---|
| cells | 6,733 |
| genes | 60,623 |
| nonzero entries | 15,166,413 |
| density | 0.0372 |
| total counts | 45,586,190 |
| mean counts / cell | 6,771 |
| mean counts / gene | 752 |
| min counts / cell | 69 |
| max counts / cell | 114,380 |
| overdispersion | 0.928 |

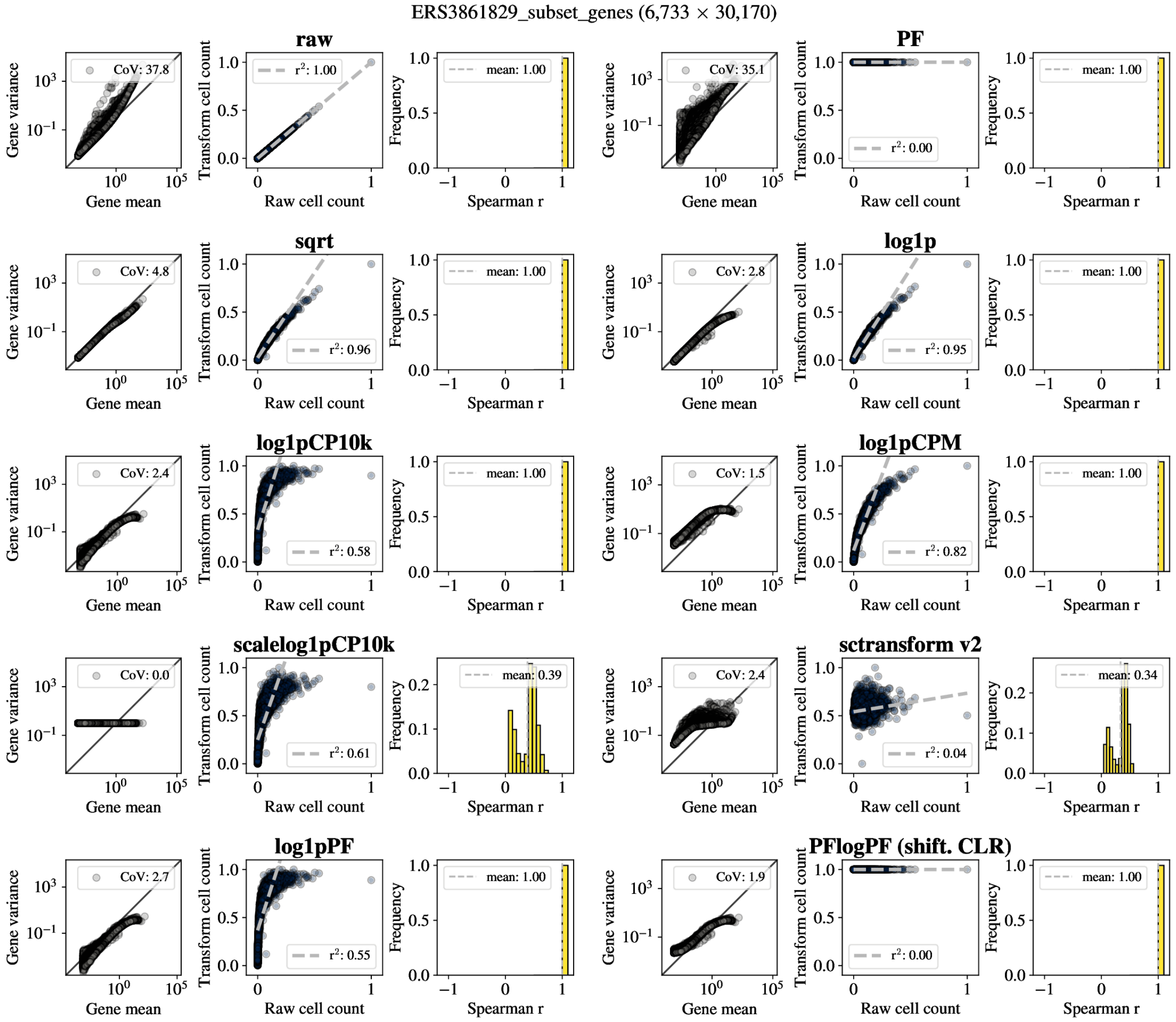


Supplementary Figure 1.76. **ERS3861829**. Per-method comparison of variance stabilization (left), depth normalization (middle), and monotonicity (right) for each normalization on dataset ERS3861829.

| **ERS3861831** | |
|---|---|
| cells | 3,530 |
| genes | 60,623 |
| nonzero entries | 6,852,453 |
| density | 0.032 |
| total counts | 31,785,192 |
| mean counts / cell | 9,004 |
| mean counts / gene | 524 |
| min counts / cell | 424 |
| max counts / cell | 55,171 |
| overdispersion | 4.77 |

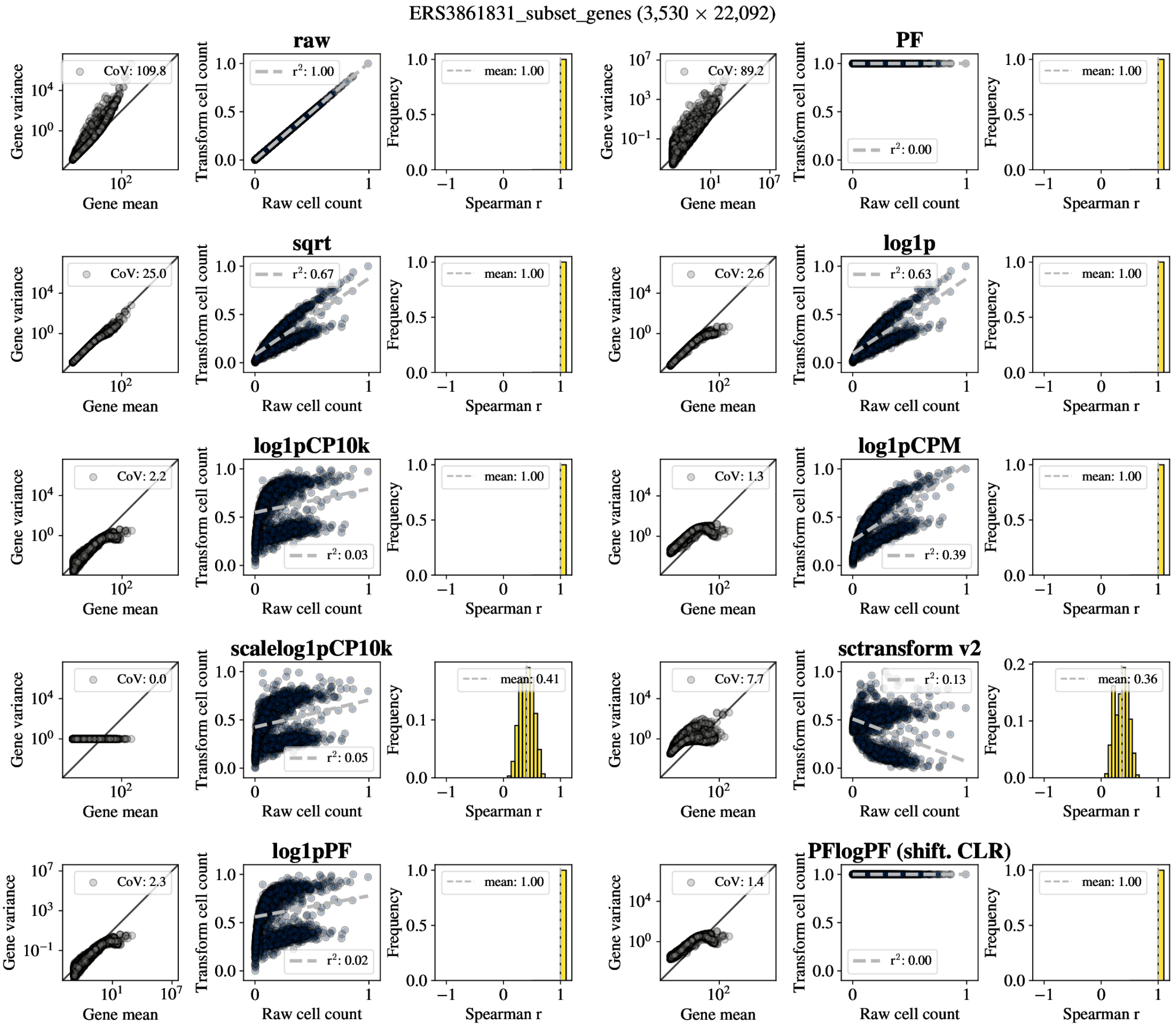


Supplementary Figure 1.77. **ERS3861831**. Per-method comparison of variance stabilization (left), depth normalization (middle), and monotonicity (right) for each normalization on dataset ERS3861831.

| ERS3861832 | |
|---|---|
| cells | 1,638 |
| genes | 60,623 |
| nonzero entries | 4,192,410 |
| density | 0.0422 |
| total counts | 23,228,386 |
| mean counts / cell | 14,181 |
| mean counts / gene | 383 |
| min counts / cell | 6,390 |
| max counts / cell | 56,475 |
| overdispersion | 2.36 |

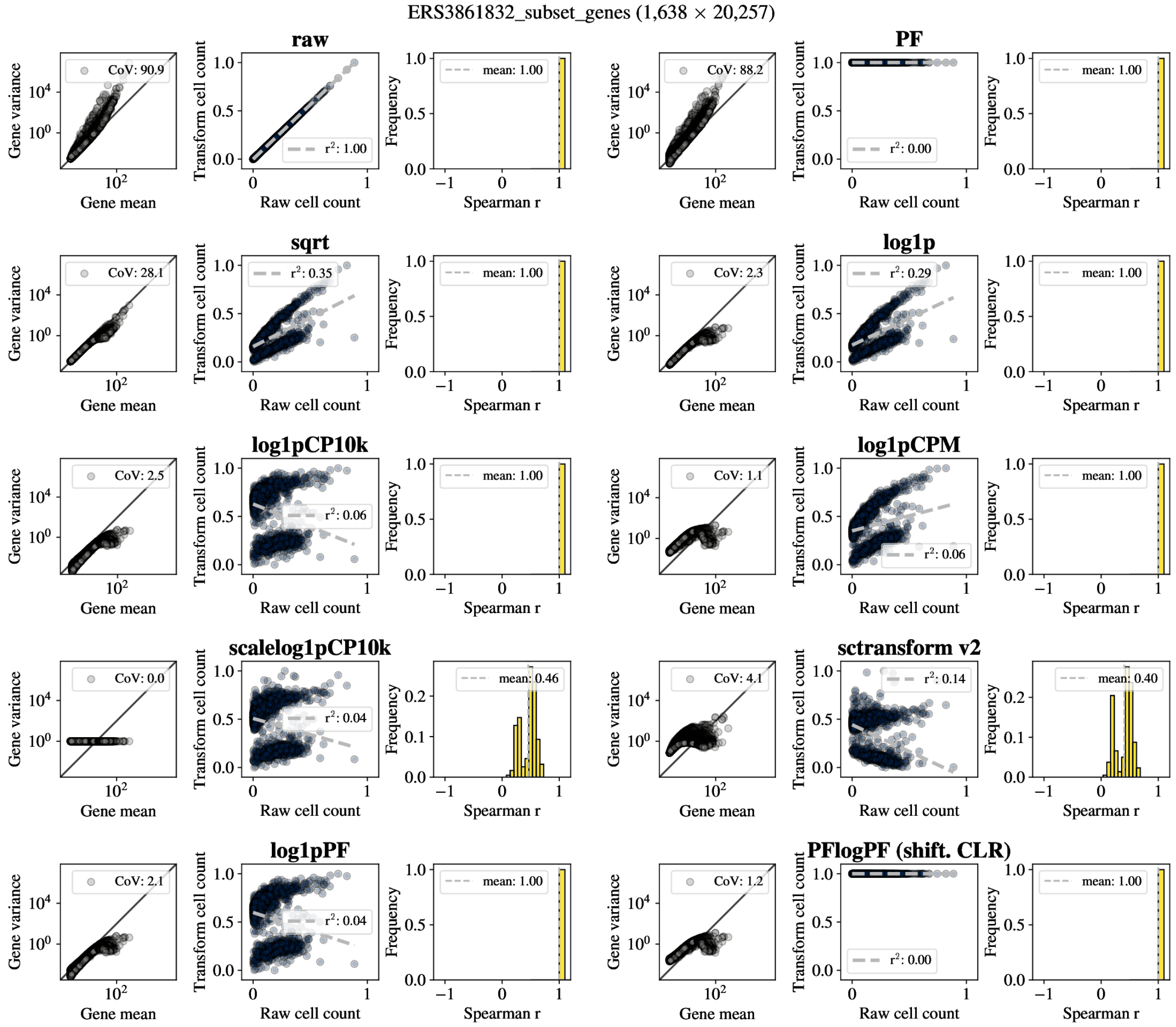


Supplementary Figure 1.78. **ERS3861832**. Per-method comparison of variance stabilization (left), depth normalization (middle), and monotonicity (right) for each normalization on dataset ERS3861832.

| ERS3861833 | |
|---|---|
| cells | 5,943 |
| genes | 60,623 |
| nonzero entries | 8,675,340 |
| density | 0.0241 |
| total counts | 34,731,490 |
| mean counts / cell | 5,844 |
| mean counts / gene | 573 |
| min counts / cell | 56 |
| max counts / cell | 77,505 |
| overdispersion | 12 |

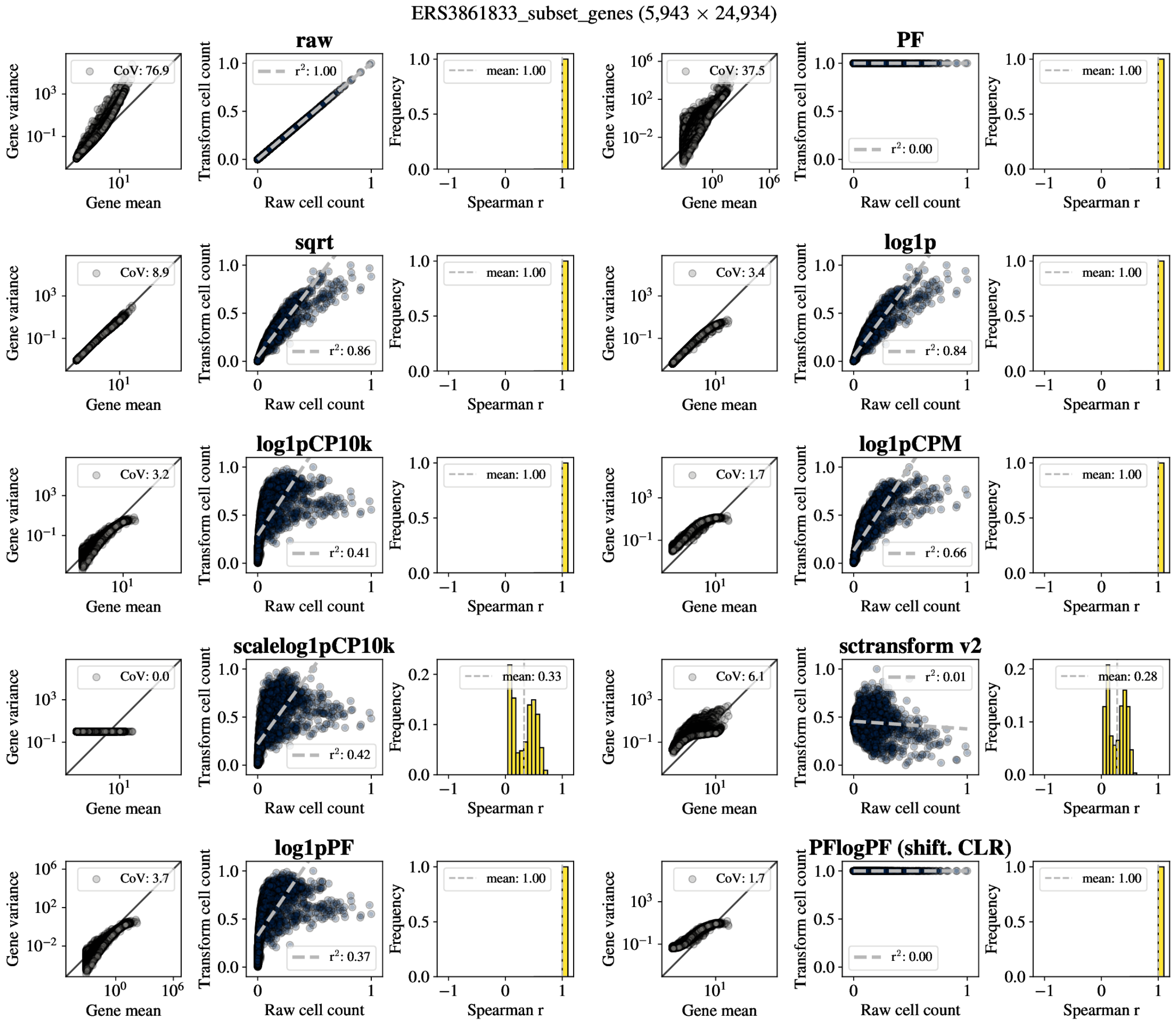


Supplementary Figure 1.79. **ERS3861833**. Per-method comparison of variance stabilization (left), depth normalization (middle), and monotonicity (right) for each normalization on dataset ERS3861833.

| ERS3861834 | |
|---|---|
| cells | 5,193 |
| genes | 60,623 |
| nonzero entries | 8,184,013 |
| density | 0.026 |
| total counts | 32,429,469 |
| mean counts / cell | 6,245 |
| mean counts / gene | 535 |
| min counts / cell | 56 |
| max counts / cell | 82,726 |
| overdispersion | 7.51 |

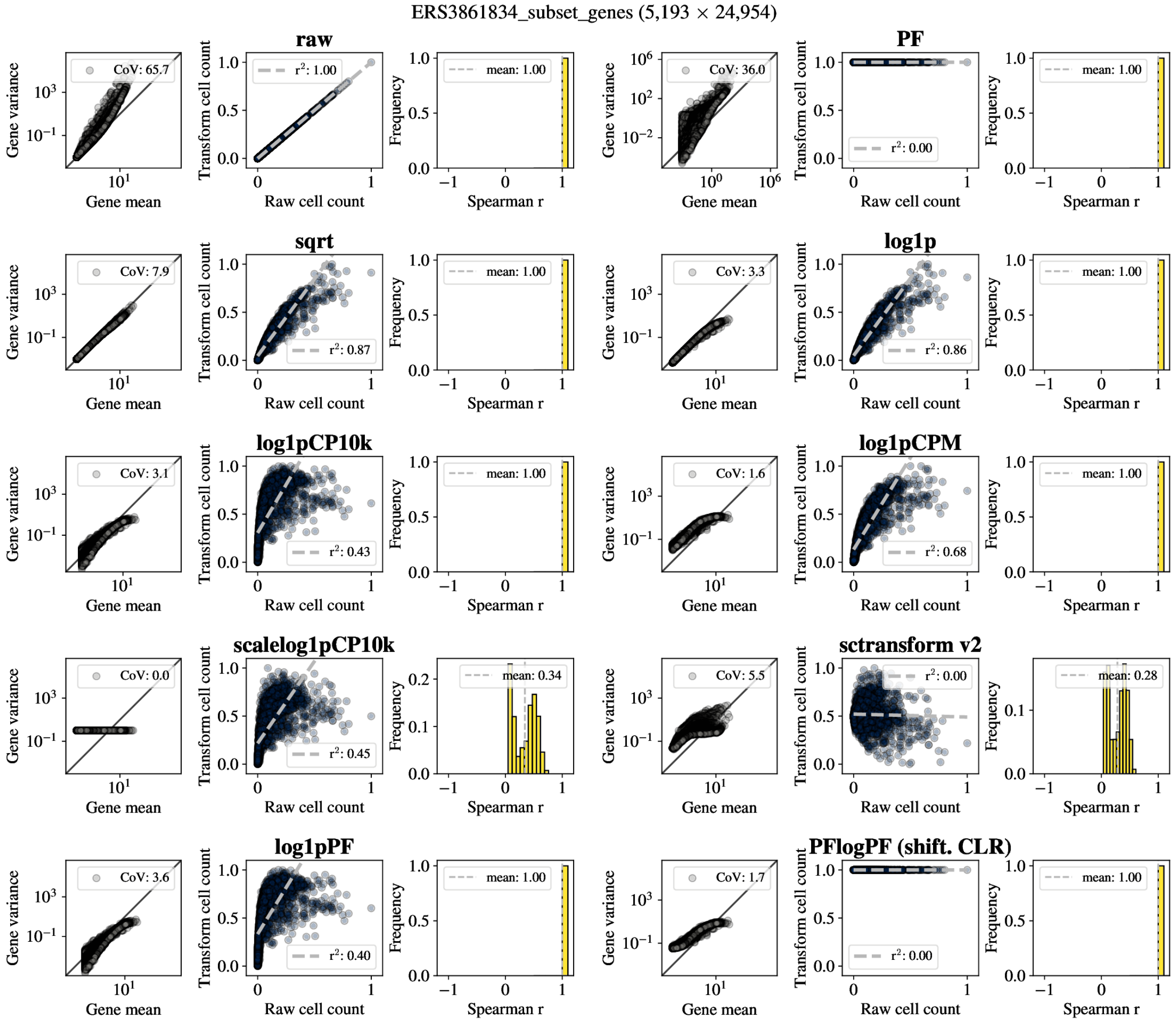


Supplementary Figure 1.80. **ERS3861834**. Per-method comparison of variance stabilization (left), depth normalization (middle), and monotonicity (right) for each normalization on dataset ERS3861834.

| ERS3861835 | |
|---|---|
| cells | 5,360 |
| genes | 60,623 |
| nonzero entries | 7,845,026 |
| density | 0.0241 |
| total counts | 30,575,591 |
| mean counts / cell | 5,704 |
| mean counts / gene | 504 |
| min counts / cell | 49 |
| max counts / cell | 71,534 |
| overdispersion | 7.15 |

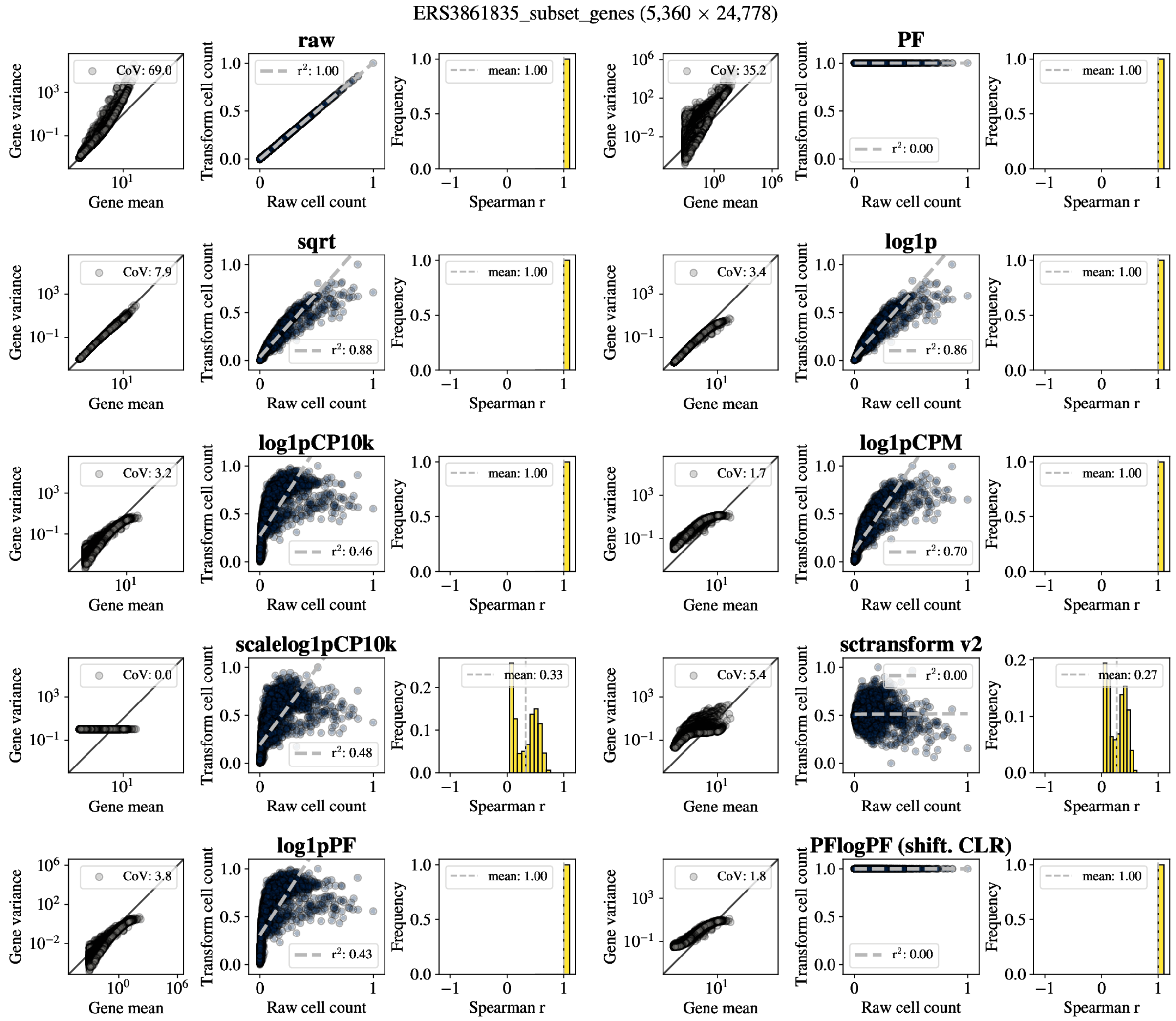


Supplementary Figure 1.81. **ERS3861835**. Per-method comparison of variance stabilization (left), depth normalization (middle), and monotonicity (right) for each normalization on dataset ERS3861835.

| **ERS3861836** | |
|---|---|
| cells | 2,693 |
| genes | 60,623 |
| nonzero entries | 5,981,598 |
| density | 0.0366 |
| total counts | 20,765,795 |
| mean counts / cell | 7,711 |
| mean counts / gene | 343 |
| min counts / cell | 1,837 |
| max counts / cell | 51,488 |
| overdispersion | 0.769 |

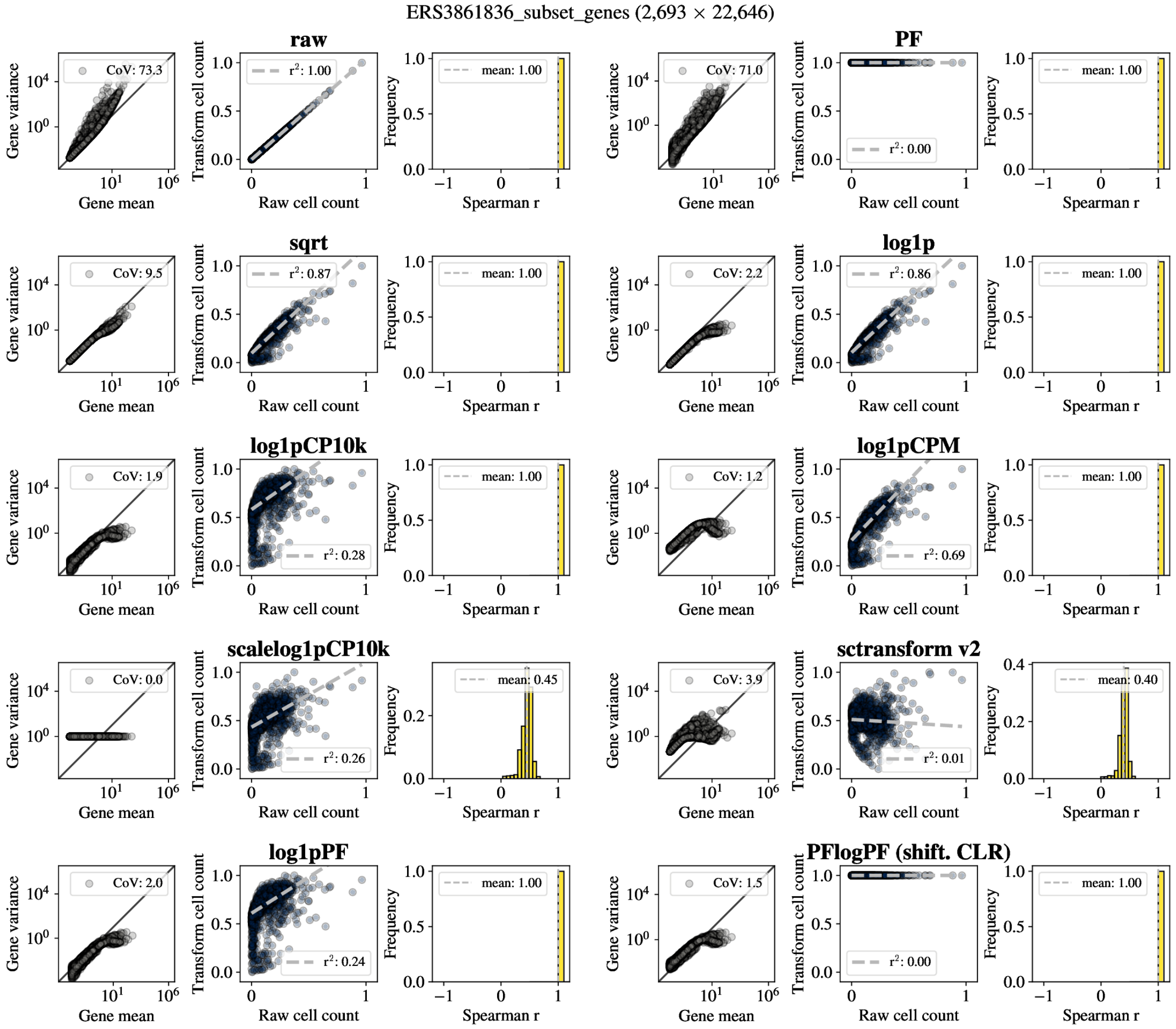


Supplementary Figure 1.82. **ERS3861836**. Per-method comparison of variance stabilization (left), depth normalization (middle), and monotonicity (right) for each normalization on dataset ERS3861836.

| **ERS3861837** | |
|---|---|
| cells | 4,687 |
| genes | 60,623 |
| nonzero entries | 6,118,560 |
| density | 0.0215 |
| total counts | 21,066,623 |
| mean counts / cell | 4,495 |
| mean counts / gene | 348 |
| min counts / cell | 55 |
| max counts / cell | 75,458 |
| overdispersion | 2.93 |

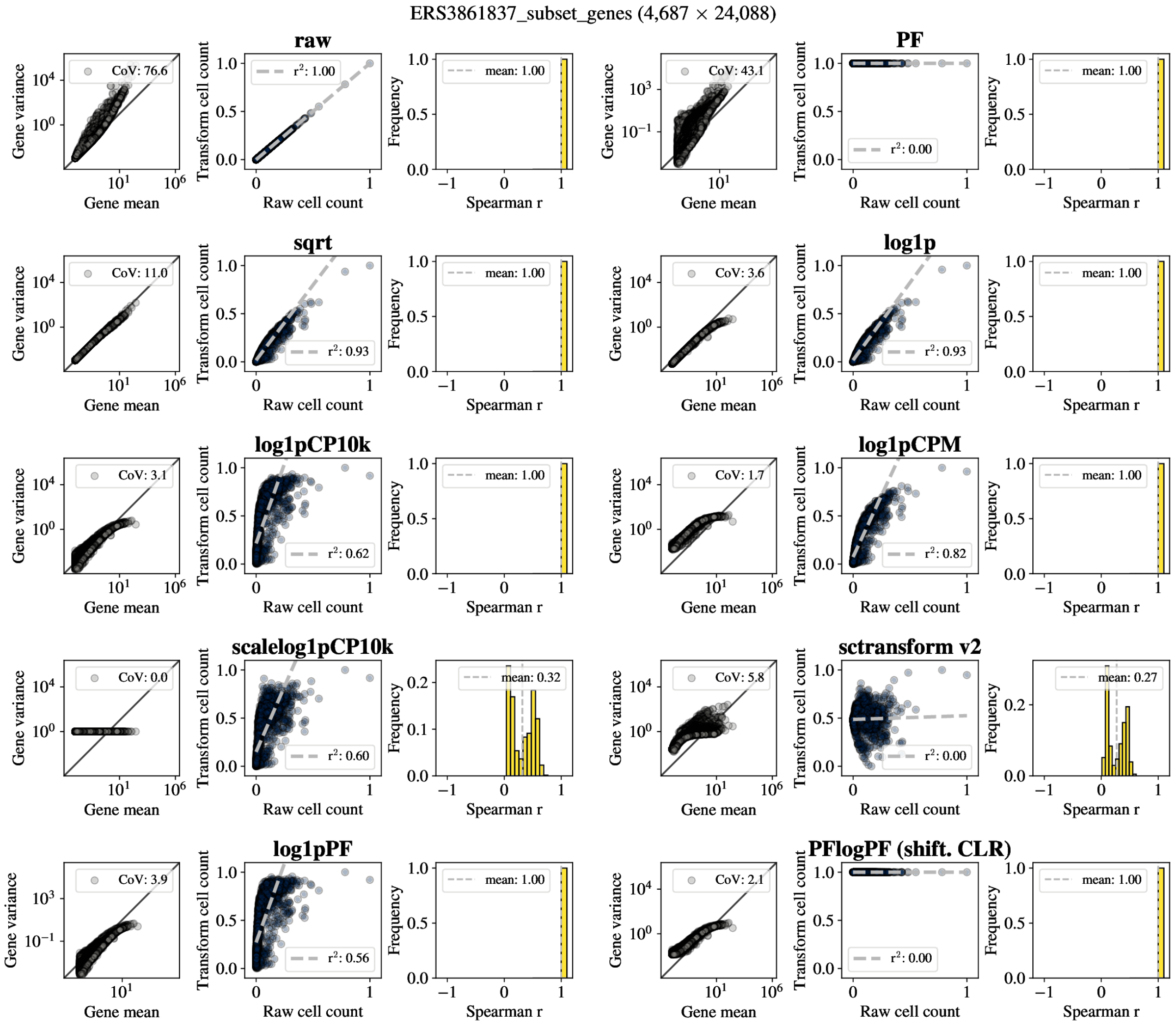


Supplementary Figure 1.83. **ERS3861837**. Per-method comparison of variance stabilization (left), depth normalization (middle), and monotonicity (right) for each normalization on dataset ERS3861837.

| ERS3861838 | |
|---|---|
| cells | 6,095 |
| genes | 60,623 |
| nonzero entries | 11,966,050 |
| density | 0.0324 |
| total counts | 66,948,522 |
| mean counts / cell | 10,984 |
| mean counts / gene | 1,104 |
| min counts / cell | 76 |
| max counts / cell | 66,652 |
| overdispersion | 2.89 |

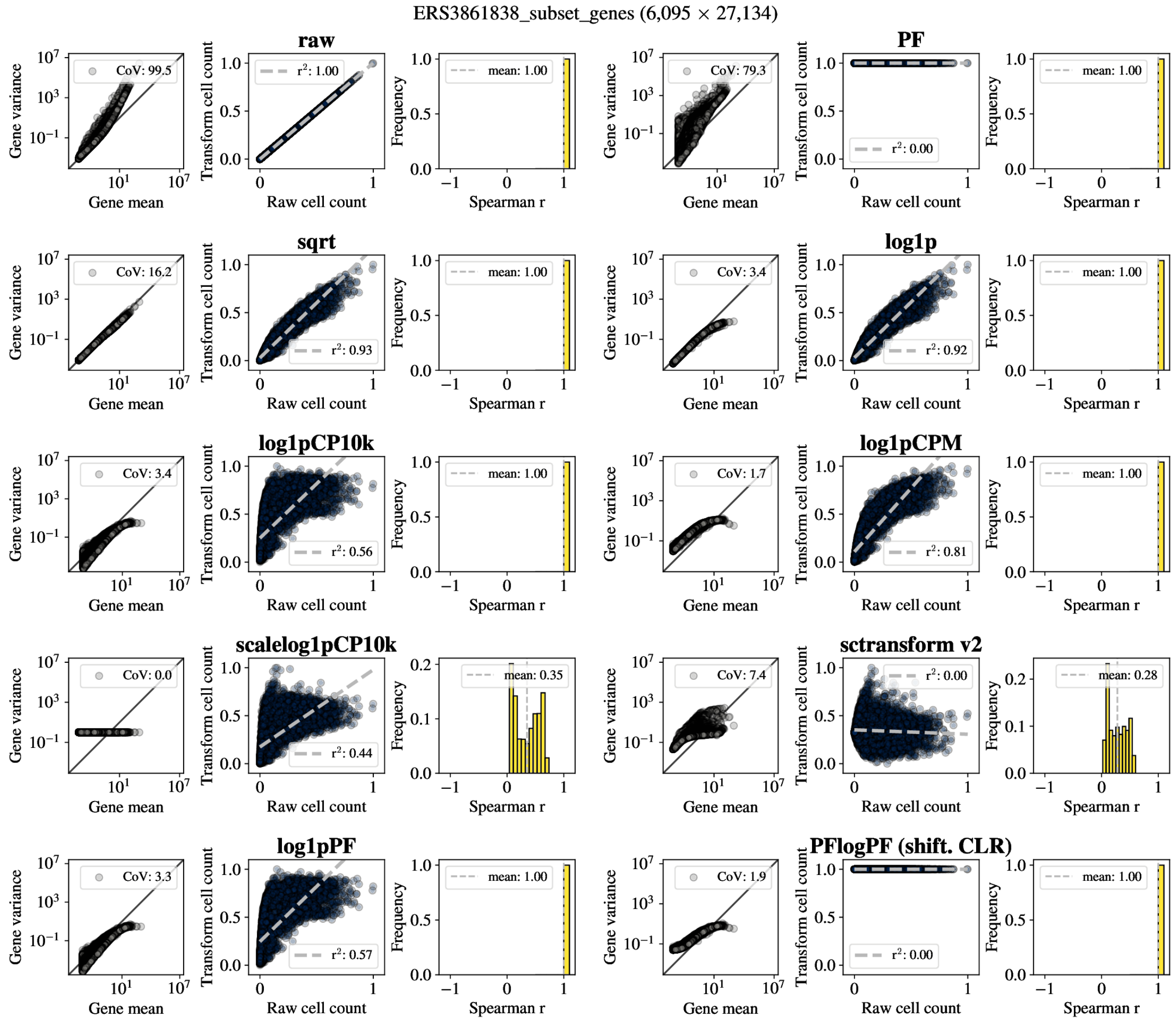


Supplementary Figure 1.84. **ERS3861838**. Per-method comparison of variance stabilization (left), depth normalization (middle), and monotonicity (right) for each normalization on dataset ERS3861838.

| ERS3861839 | |
|---|---|
| cells | 3,245 |
| genes | 60,623 |
| nonzero entries | 10,875,436 |
| density | 0.0553 |
| total counts | 62,491,297 |
| mean counts / cell | 19,258 |
| mean counts / gene | 1,031 |
| min counts / cell | 3,069 |
| max counts / cell | 68,466 |
| overdispersion | 1.25 |

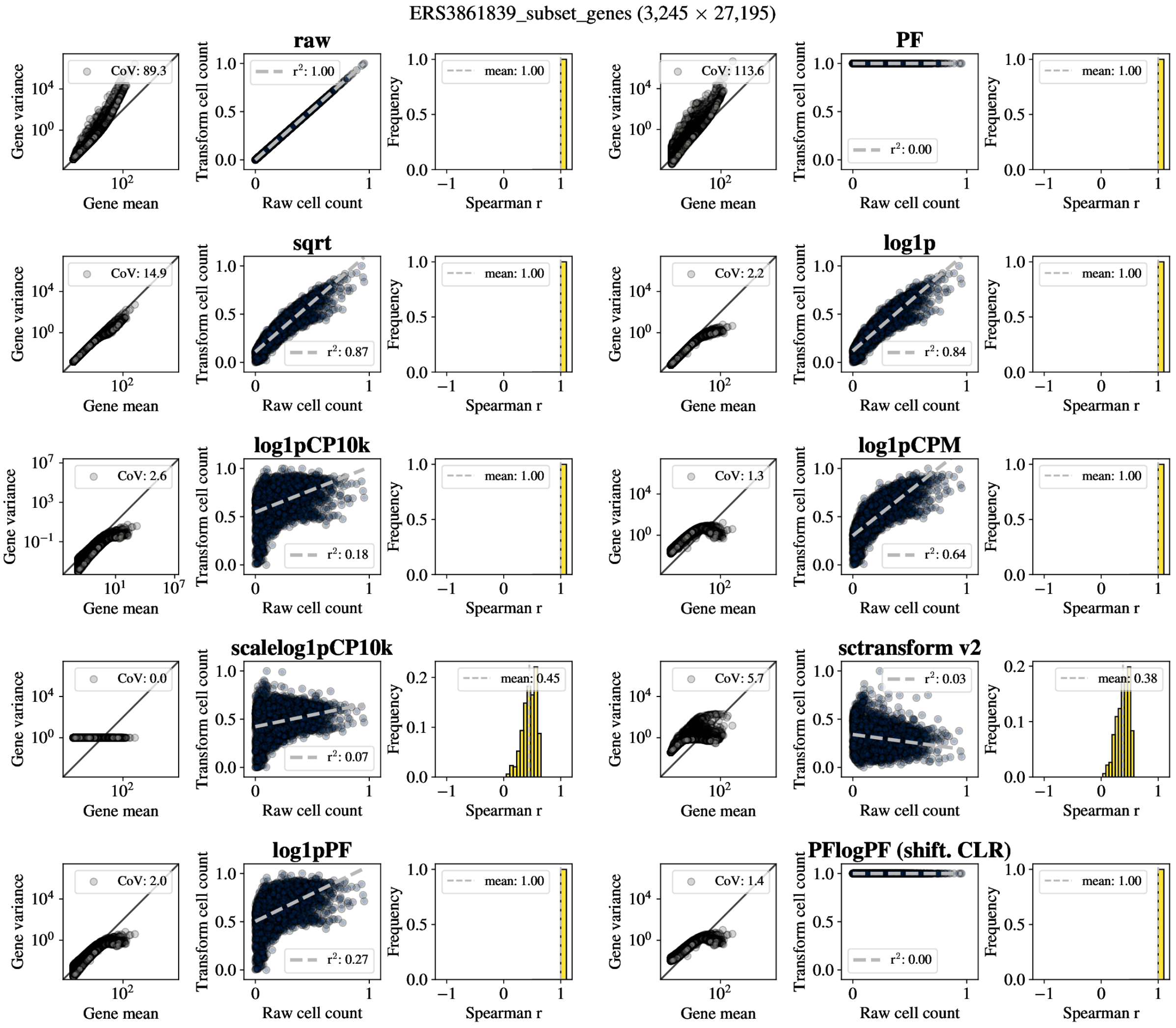


Supplementary Figure 1.85. **ERS3861839**. Per-method comparison of variance stabilization (left), depth normalization (middle), and monotonicity (right) for each normalization on dataset ERS3861839.

| ERS3861840 | |
|---|---|
| cells | 3,247 |
| genes | 60,623 |
| nonzero entries | 10,017,197 |
| density | 0.0509 |
| total counts | 47,439,493 |
| mean counts / cell | 14,610 |
| mean counts / gene | 783 |
| min counts / cell | 2,575 |
| max counts / cell | 70,650 |
| overdispersion | 1.23 |

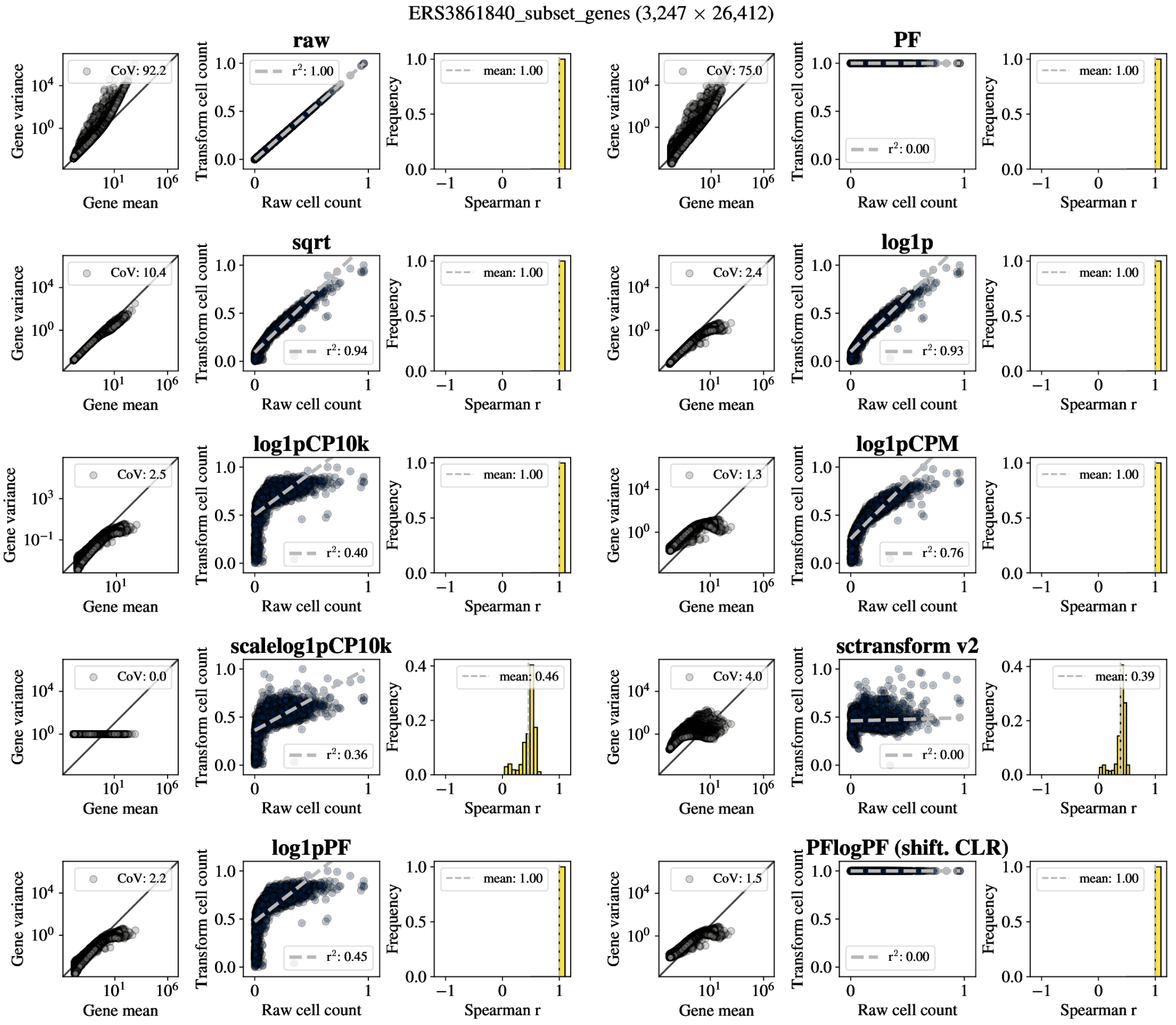


Supplementary Figure 1.86. **ERS3861840**. Per-method comparison of variance stabilization (left), depth normalization (middle), and monotonicity (right) for each normalization on dataset ERS3861840.

| **ERS3861841** | |
|---|---|
| cells | 7,068 |
| genes | 60,623 |
| nonzero entries | 11,823,068 |
| density | 0.0276 |
| total counts | 38,291,784 |
| mean counts / cell | 5,418 |
| mean counts / gene | 632 |
| min counts / cell | 172 |
| max counts / cell | 57,512 |
| overdispersion | 7.39 |

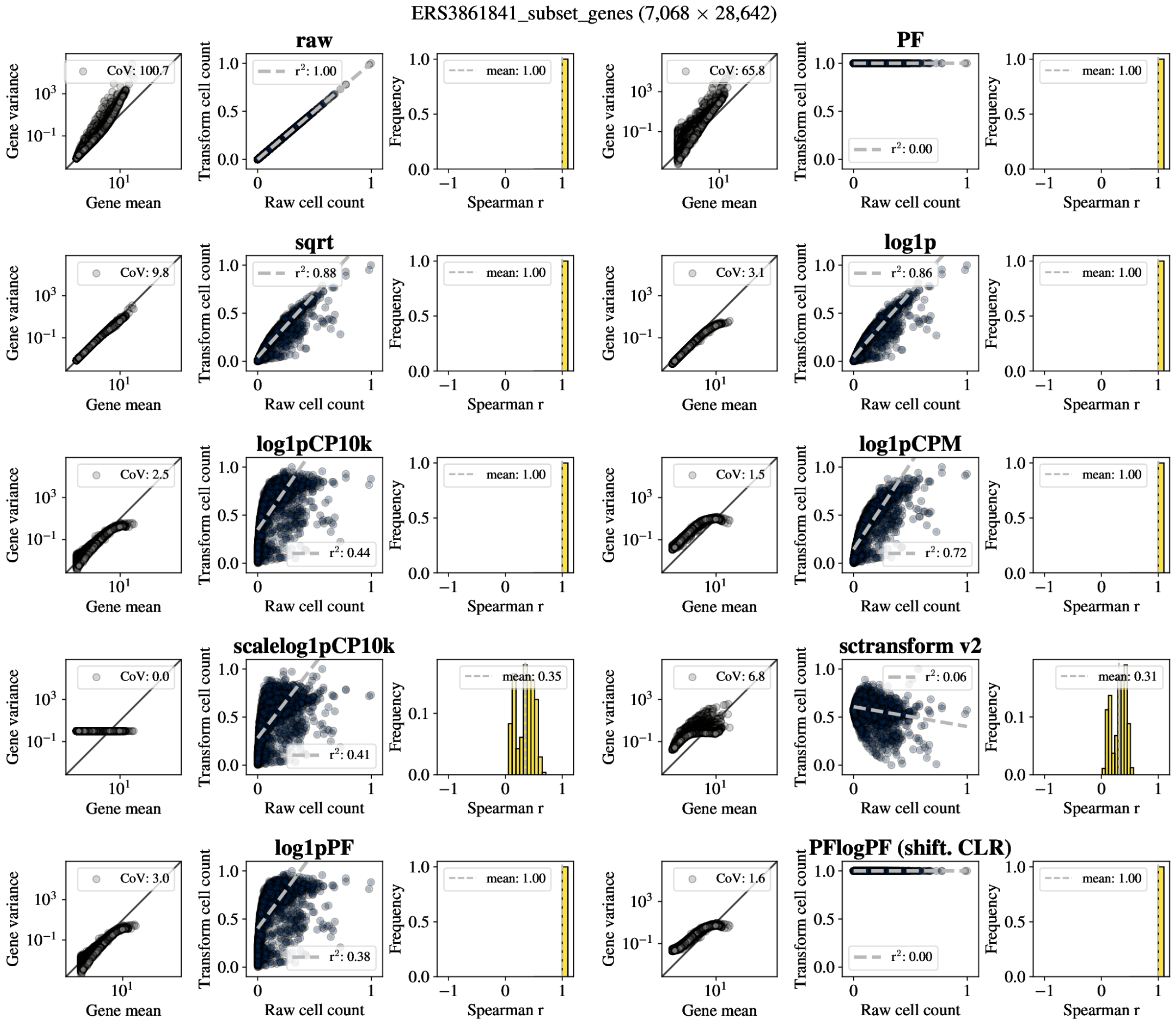


Supplementary Figure 1.87. **ERS3861841**. Per-method comparison of variance stabilization (left), depth normalization (middle), and monotonicity (right) for each normalization on dataset ERS3861841.

| **ERS3861842** | |
|---|---|
| cells | 8,406 |
| genes | 60,623 |
| nonzero entries | 17,420,638 |
| density | 0.0342 |
| total counts | 88,964,954 |
| mean counts / cell | 10,584 |
| mean counts / gene | 1,468 |
| min counts / cell | 186 |
| max counts / cell | 73,791 |
| overdispersion | 1.82 |

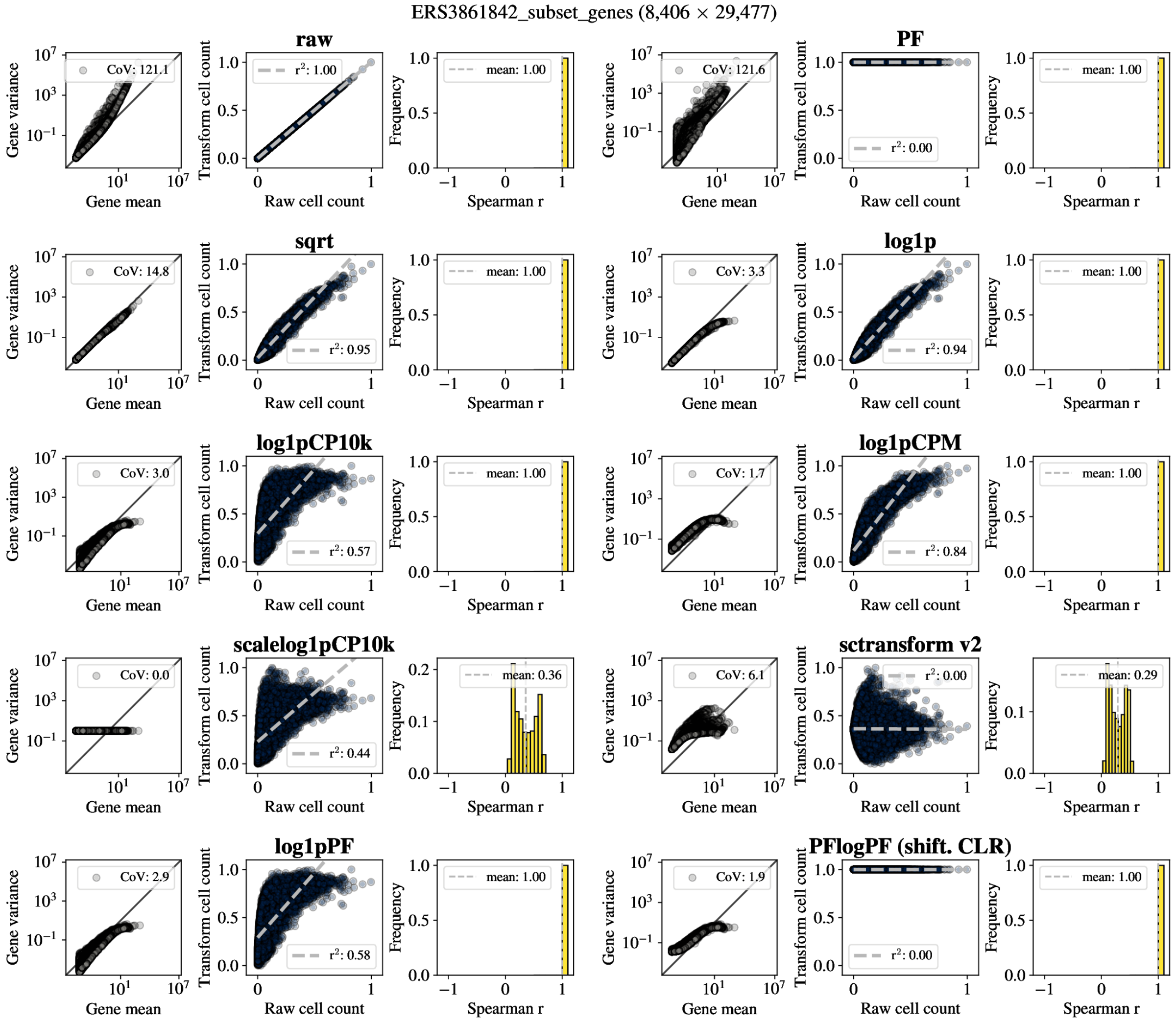


Supplementary Figure 1.88. **ERS3861842**. Per-method comparison of variance stabilization (left), depth normalization (middle), and monotonicity (right) for each normalization on dataset ERS3861842.

| **ERS3861843** | |
|---|---|
| cells | 4,089 |
| genes | 60,623 |
| nonzero entries | 9,485,725 |
| density | 0.0383 |
| total counts | 35,518,397 |
| mean counts / cell | 8,686 |
| mean counts / gene | 586 |
| min counts / cell | 2,861 |
| max counts / cell | 76,402 |
| overdispersion | 1.47 |

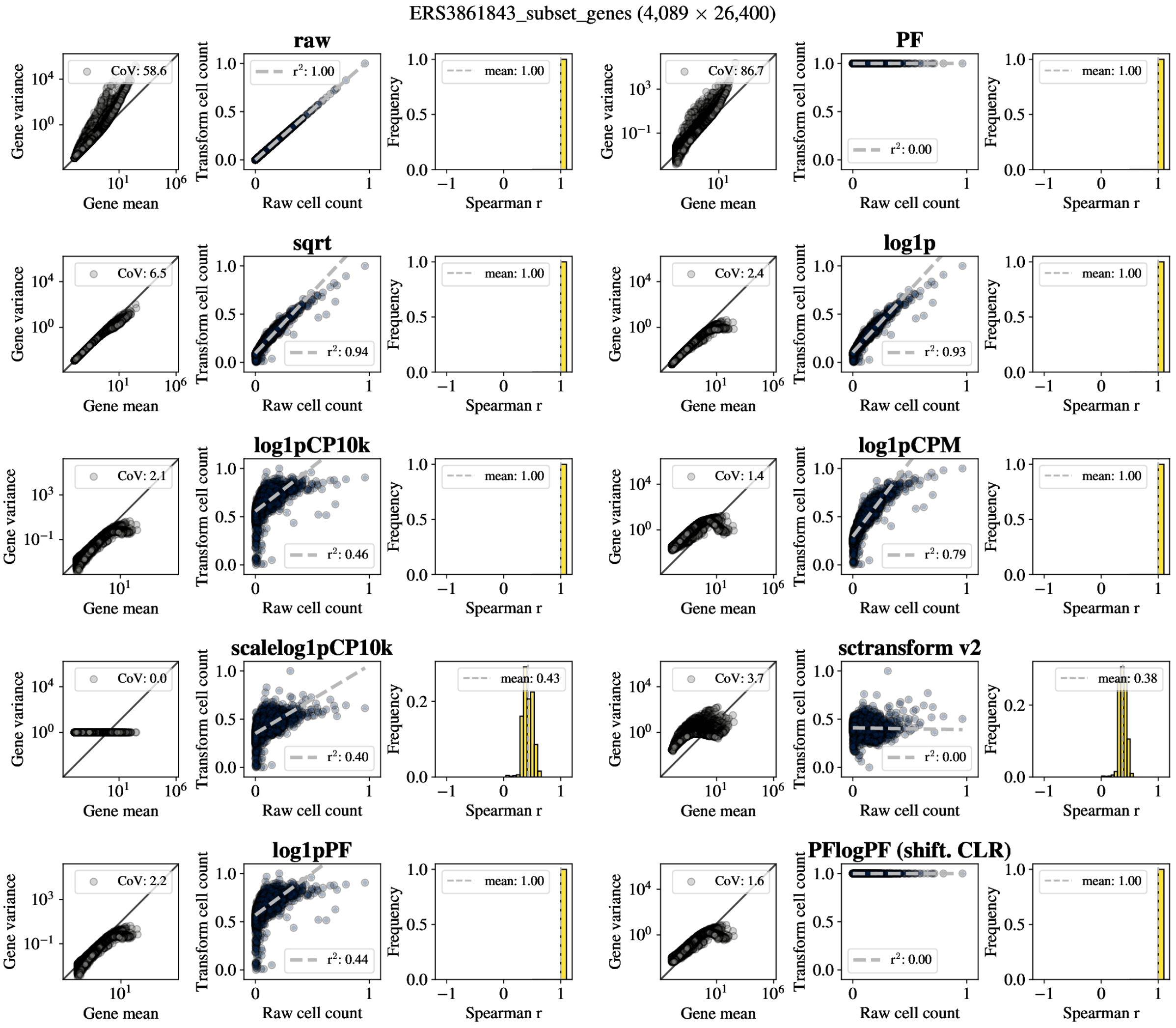


Supplementary Figure 1.89. **ERS3861843**. Per-method comparison of variance stabilization (left), depth normalization (middle), and monotonicity (right) for each normalization on dataset ERS3861843.

| **ERS3861844** | |
|---|---|
| cells | 3,505 |
| genes | 60,623 |
| nonzero entries | 15,041,956 |
| density | 0.0708 |
| total counts | 84,955,098 |
| mean counts / cell | 24,238 |
| mean counts / gene | 1,401 |
| min counts / cell | 7,075 |
| max counts / cell | 82,522 |
| overdispersion | 0.982 |

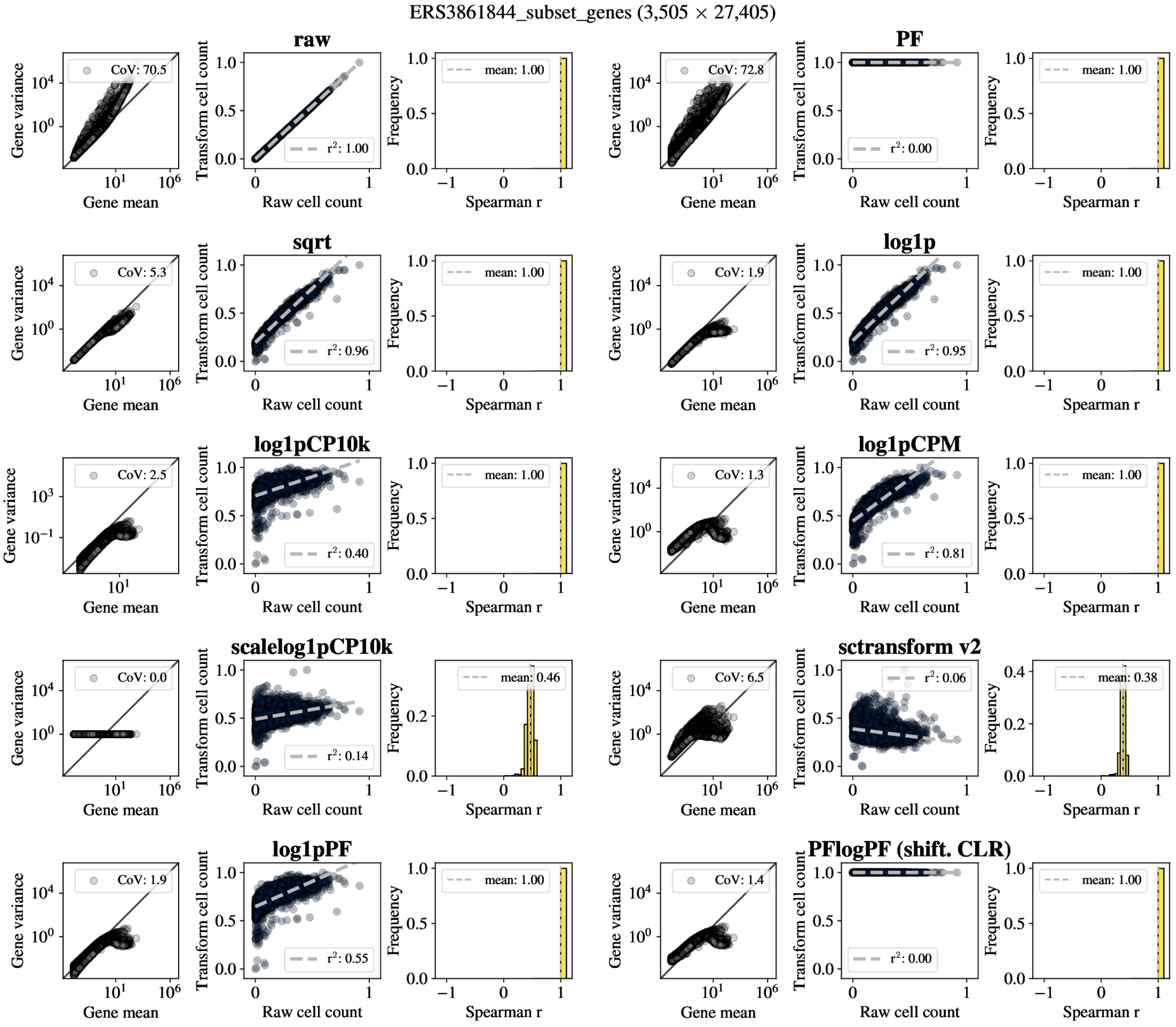


Supplementary Figure 1.90. **ERS3861844**. Per-method comparison of variance stabilization (left), depth normalization (middle), and monotonicity (right) for each normalization on dataset ERS3861844.

| **ERS3861845** | |
|---|---|
| cells | 8,181 |
| genes | 60,623 |
| nonzero entries | 12,951,927 |
| density | 0.0261 |
| total counts | 43,056,607 |
| mean counts / cell | 5,263 |
| mean counts / gene | 710 |
| min counts / cell | 119 |
| max counts / cell | 73,997 |
| overdispersion | 0.984 |

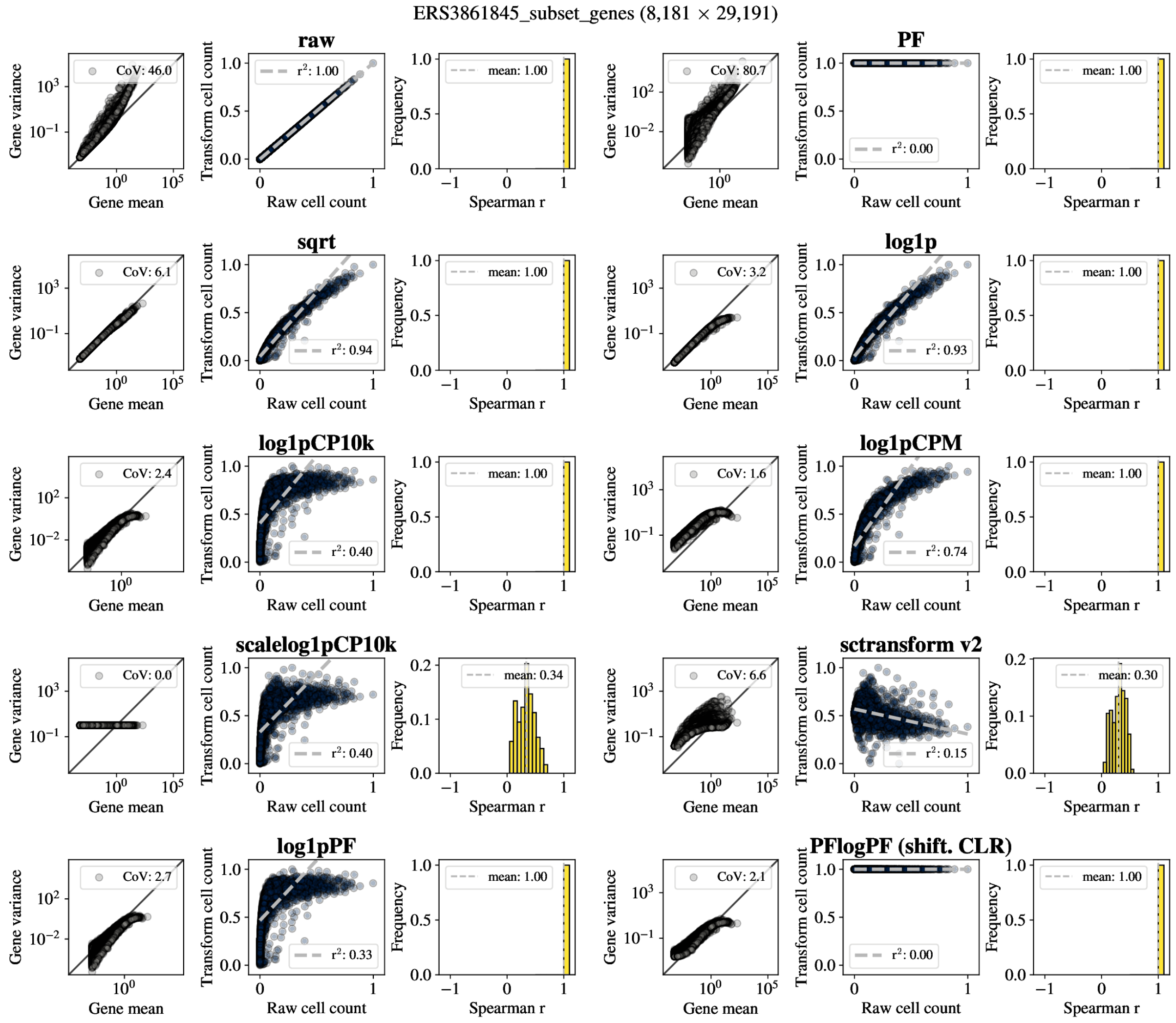


Supplementary Figure 1.91. **ERS3861845**. Per-method comparison of variance stabilization (left), depth normalization (middle), and monotonicity (right) for each normalization on dataset ERS3861845.

| **ERS3861846** | |
|---|---|
| cells | 8,164 |
| genes | 60,623 |
| nonzero entries | 10,853,957 |
| density | 0.0219 |
| total counts | 56,898,872 |
| mean counts / cell | 6,969 |
| mean counts / gene | 939 |
| min counts / cell | 113 |
| max counts / cell | 79,195 |
| overdispersion | 5.53 |

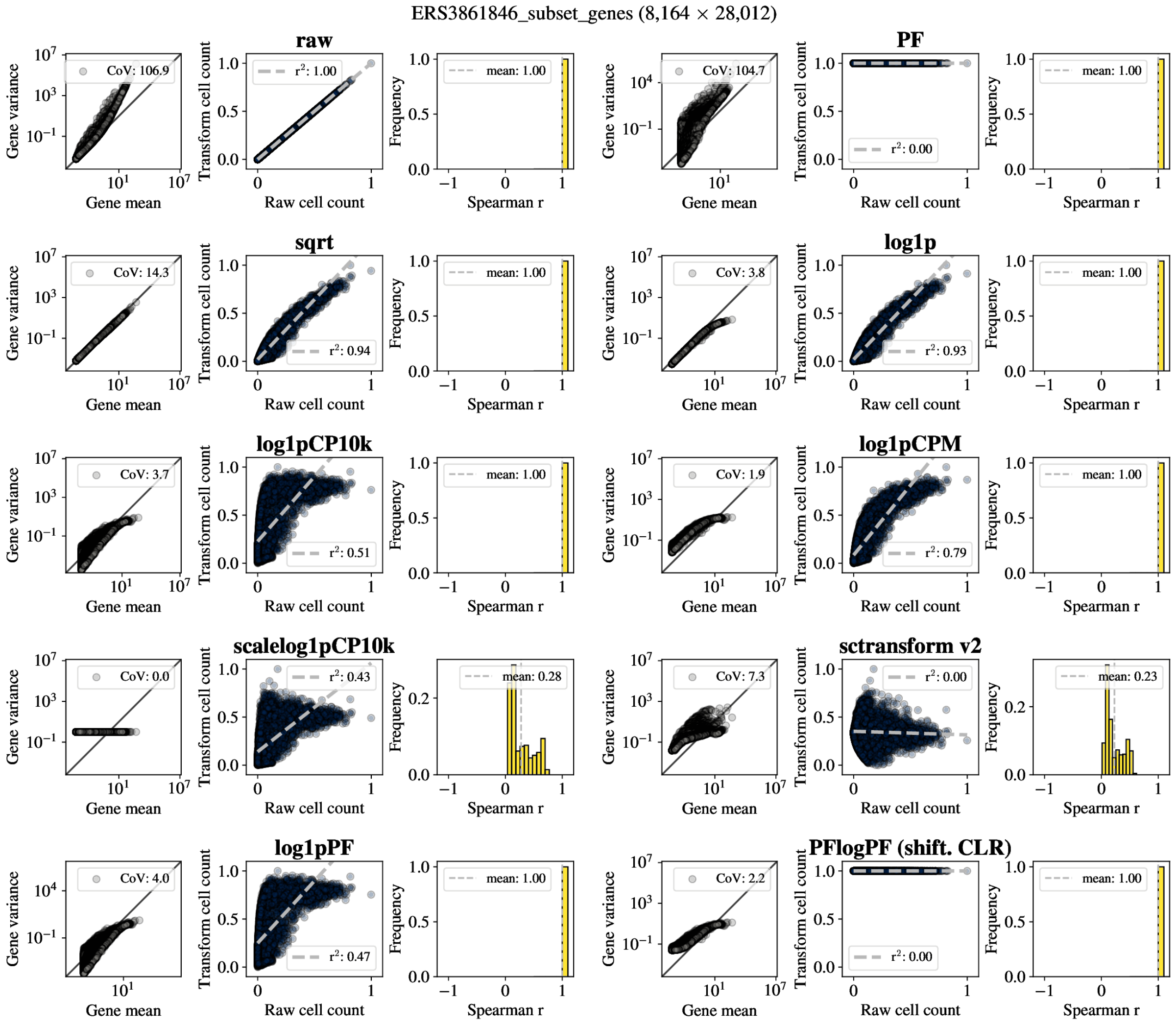


Supplementary Figure 1.92. **ERS3861846**. Per-method comparison of variance stabilization (left), depth normalization (middle), and monotonicity (right) for each normalization on dataset ERS3861846.

| **ERS3861847** | |
|---|---|
| cells | 6,809 |
| genes | 60,623 |
| nonzero entries | 10,469,859 |
| density | 0.0254 |
| total counts | 46,194,190 |
| mean counts / cell | 6,784 |
| mean counts / gene | 762 |
| min counts / cell | 84 |
| max counts / cell | 83,268 |
| overdispersion | 14.8 |

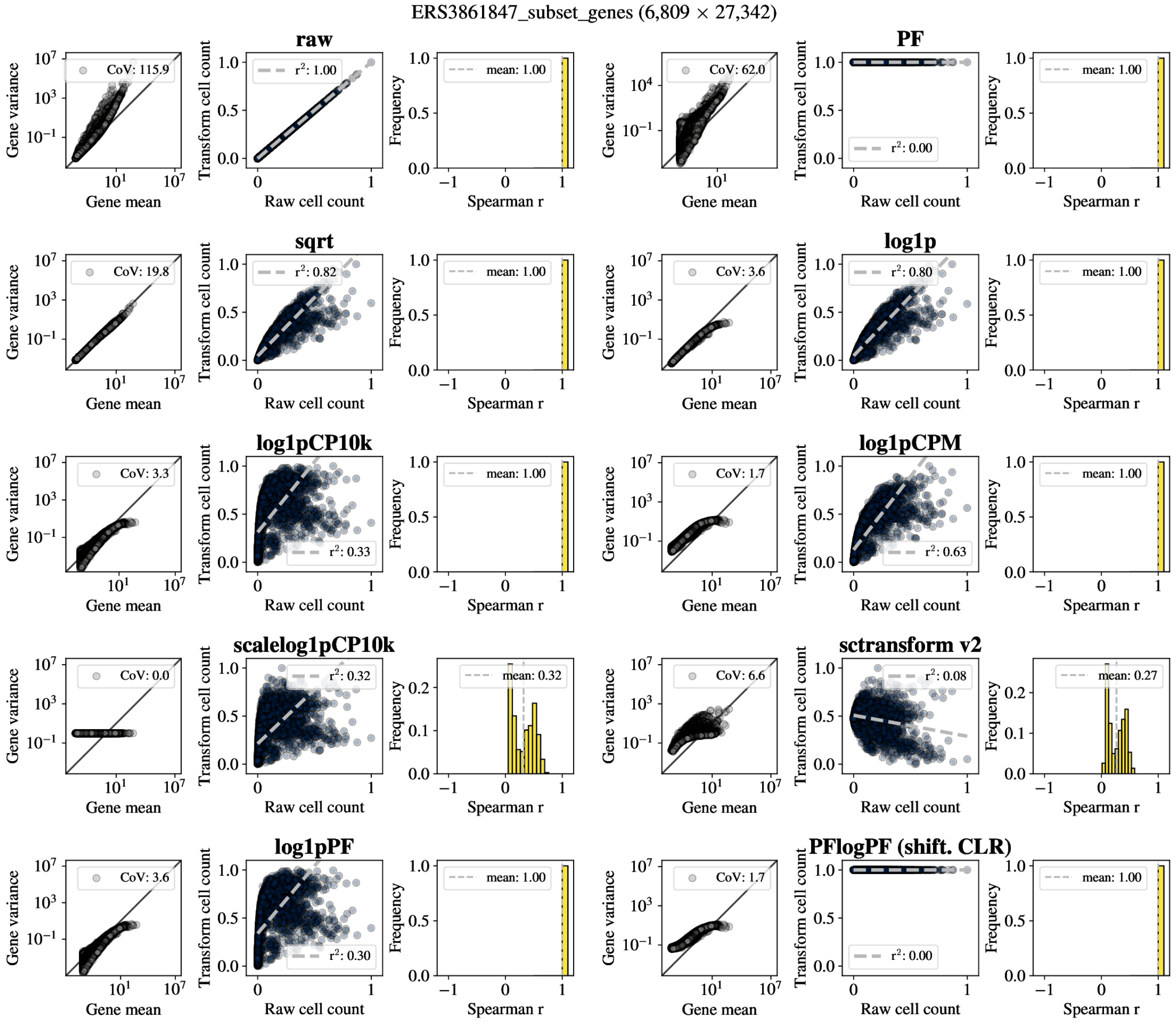


Supplementary Figure 1.93. **ERS3861847**. Per-method comparison of variance stabilization (left), depth normalization (middle), and monotonicity (right) for each normalization on dataset ERS3861847.

| ERS3861848 | |
|---|---|
| cells | 6,100 |
| genes | 60,623 |
| nonzero entries | 11,516,249 |
| density | 0.0311 |
| total counts | 69,474,982 |
| mean counts / cell | 11,389 |
| mean counts / gene | 1,146 |
| min counts / cell | 123 |
| max counts / cell | 80,088 |
| overdispersion | 8.8 |

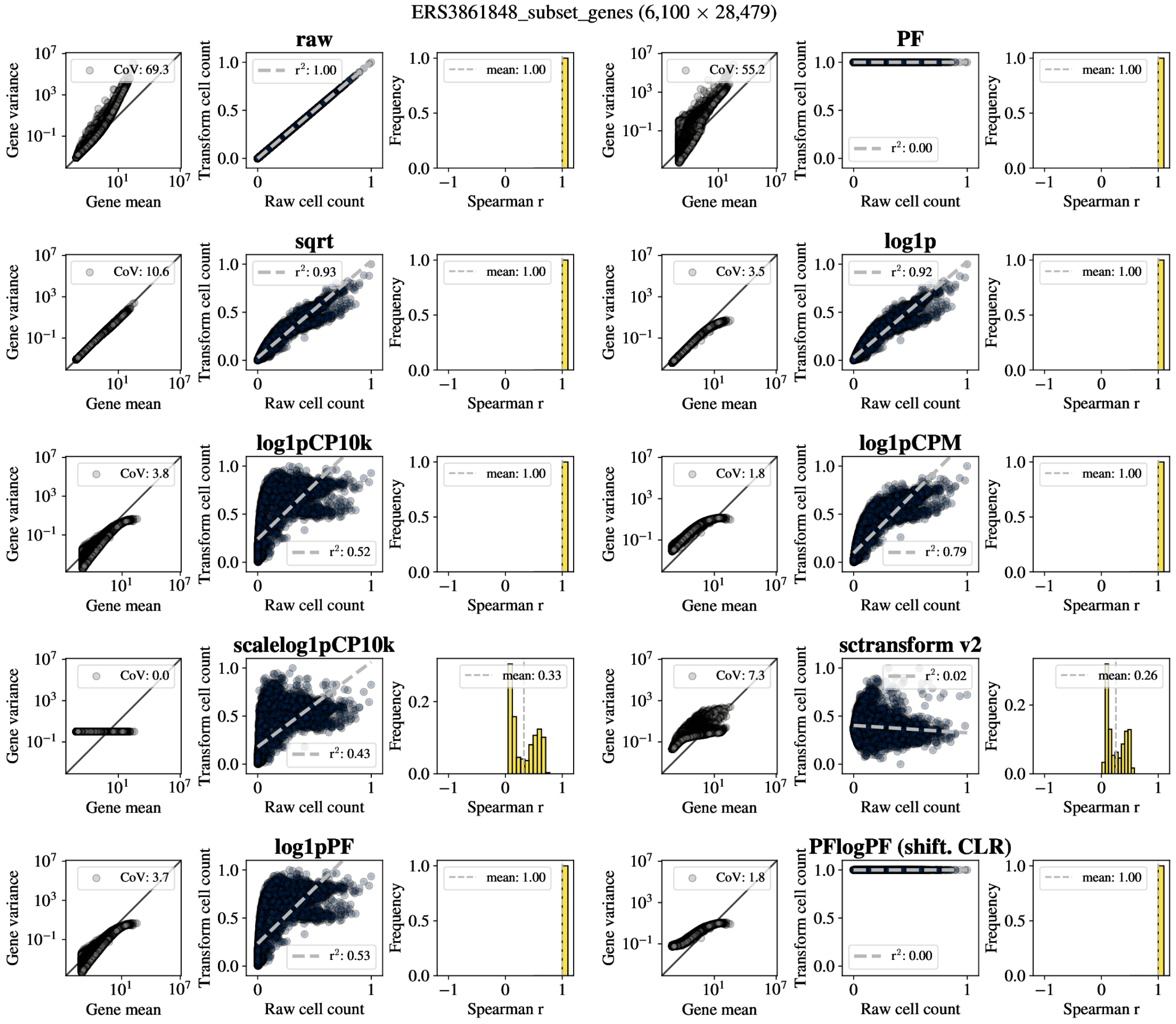


Supplementary Figure 1.94. **ERS3861848**. Per-method comparison of variance stabilization (left), depth normalization (middle), and monotonicity (right) for each normalization on dataset ERS3861848.

| **ERS3861849** | |
|---|---|
| cells | 3,985 |
| genes | 60,623 |
| nonzero entries | 14,334,152 |
| density | 0.0593 |
| total counts | 68,629,164 |
| mean counts / cell | 17,222 |
| mean counts / gene | 1,132 |
| min counts / cell | 7,266 |
| max counts / cell | 72,929 |
| overdispersion | 1.16 |

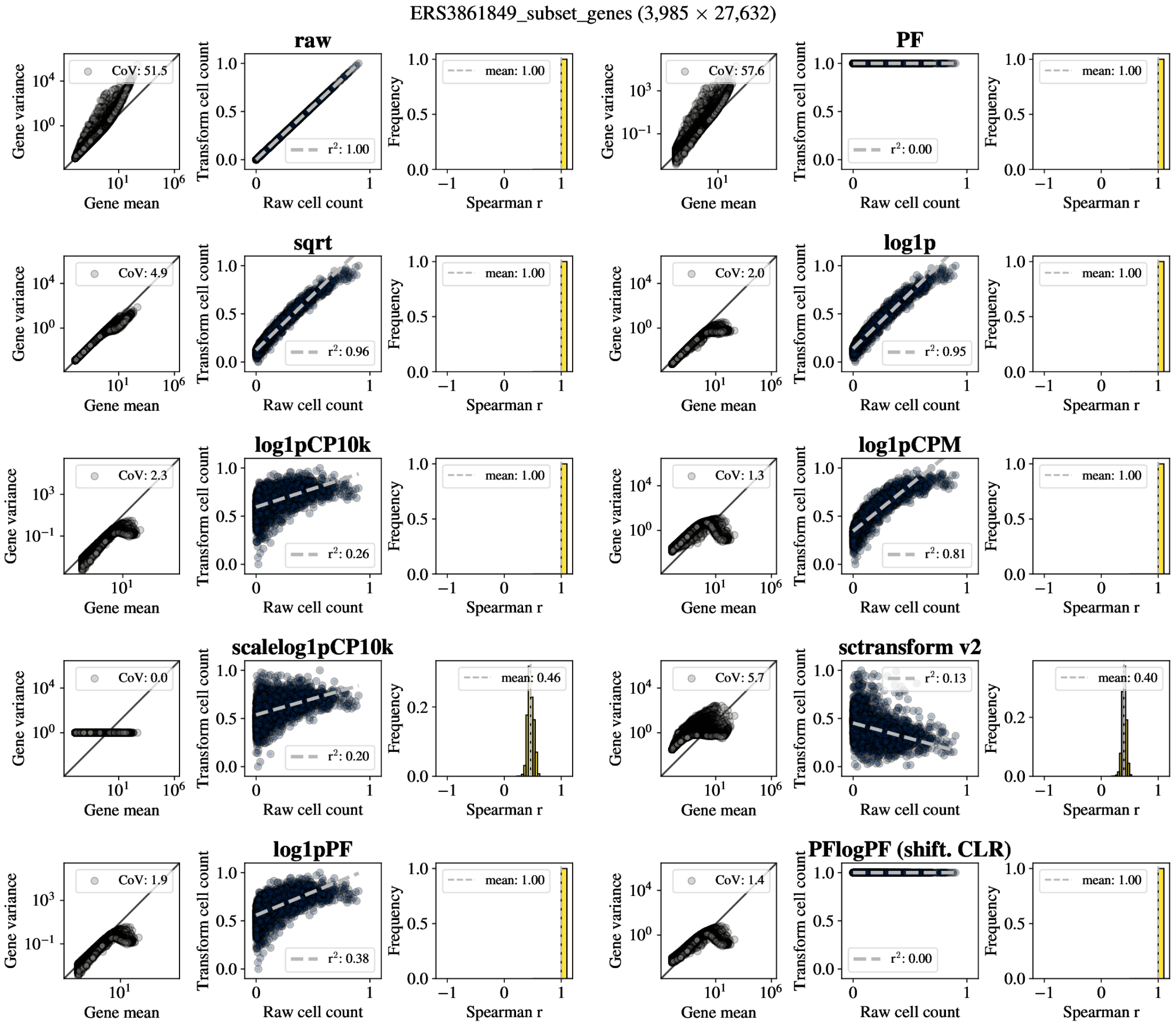


Supplementary Figure 1.95. **ERS3861849**. Per-method comparison of variance stabilization (left), depth normalization (middle), and monotonicity (right) for each normalization on dataset ERS3861849.

| **ERS3861850** | |
|---|---|
| cells | 5,154 |
| genes | 60,623 |
| nonzero entries | 9,832,663 |
| density | 0.0315 |
| total counts | 59,137,544 |
| mean counts / cell | 11,474 |
| mean counts / gene | 975 |
| min counts / cell | 80 |
| max counts / cell | 97,078 |
| overdispersion | 3.91 |

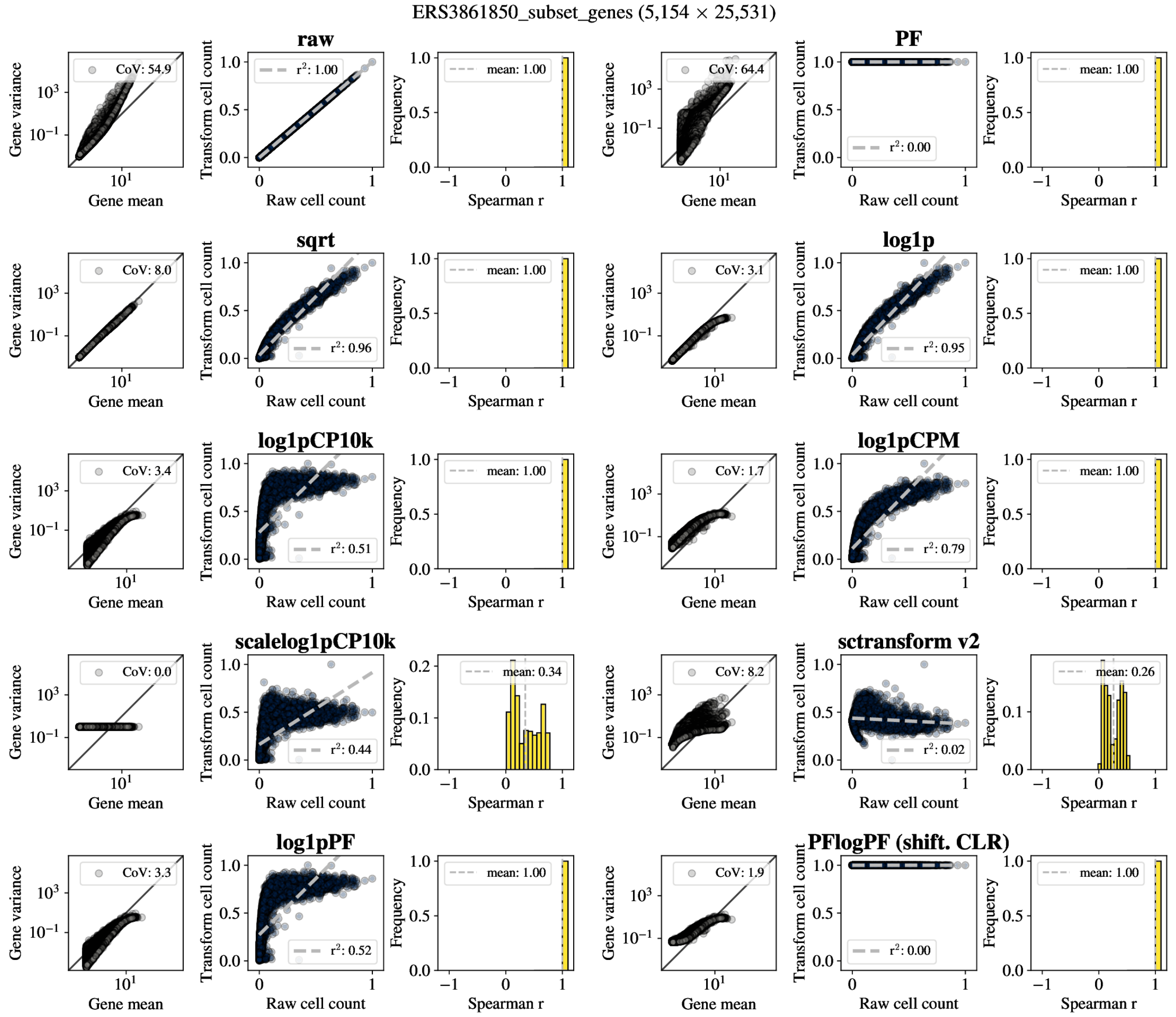


Supplementary Figure 1.96. **ERS3861850**. Per-method comparison of variance stabilization (left), depth normalization (middle), and monotonicity (right) for each normalization on dataset ERS3861850.

| ERS4228628 | |
|---|---|
| cells | 1,000 |
| genes | 60,623 |
| nonzero entries | 1,645,629 |
| density | 0.0271 |
| total counts | 3,797,641 |
| mean counts / cell | 3,798 |
| mean counts / gene | 63 |
| min counts / cell | 1,464 |
| max counts / cell | 28,224 |
| overdispersion | 0.889 |

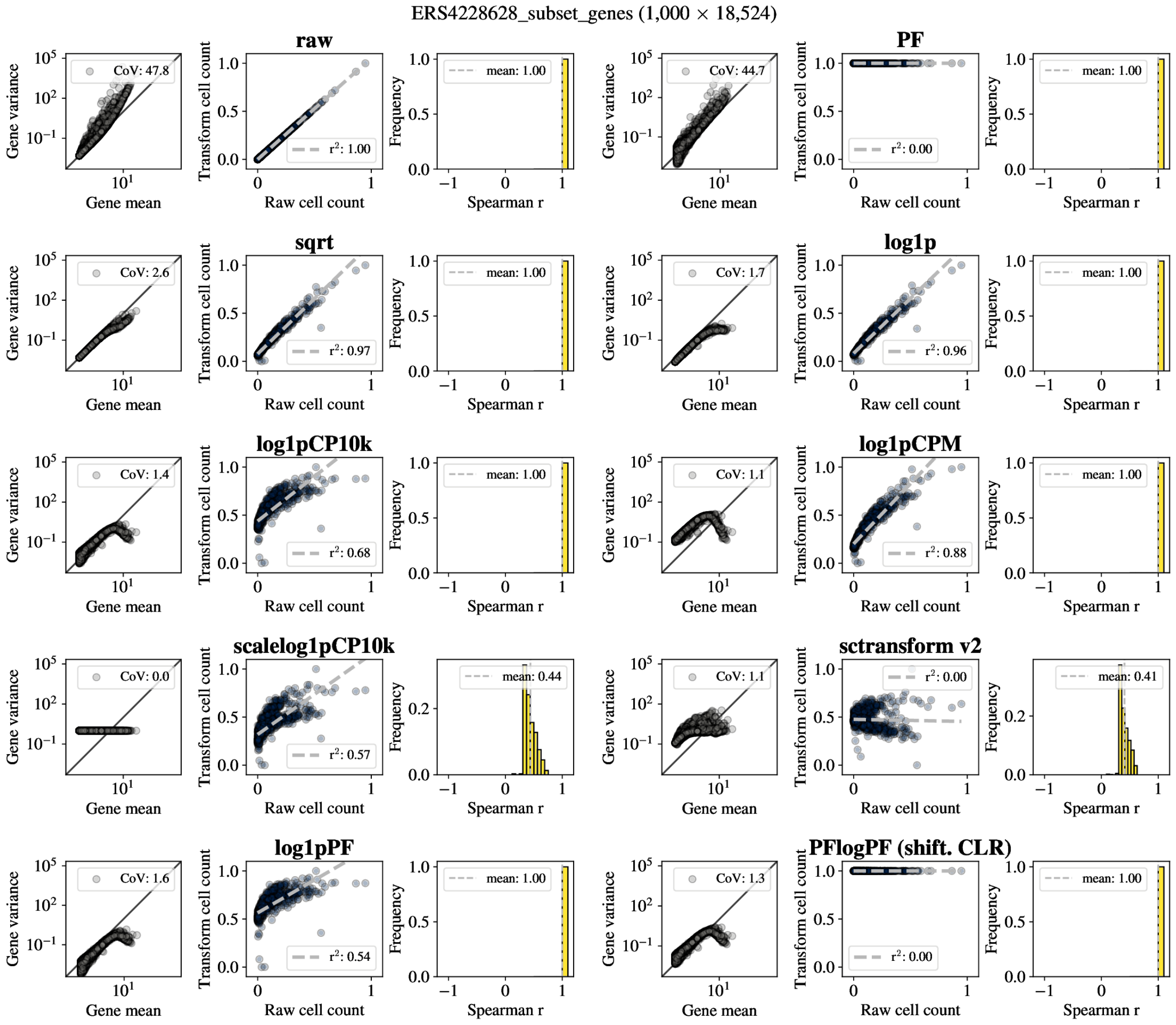


Supplementary Figure 1.97. **ERS4228628**. Per-method comparison of variance stabilization (left), depth normalization (middle), and monotonicity (right) for each normalization on dataset ERS4228628.

| ERS4228634 | |
|---|---|
| cells | 6,496 |
| genes | 60,623 |
| nonzero entries | 11,871,652 |
| density | 0.0301 |
| total counts | 36,395,911 |
| mean counts / cell | 5,603 |
| mean counts / gene | 600 |
| min counts / cell | 250 |
| max counts / cell | 60,247 |
| overdispersion | 1.06 |

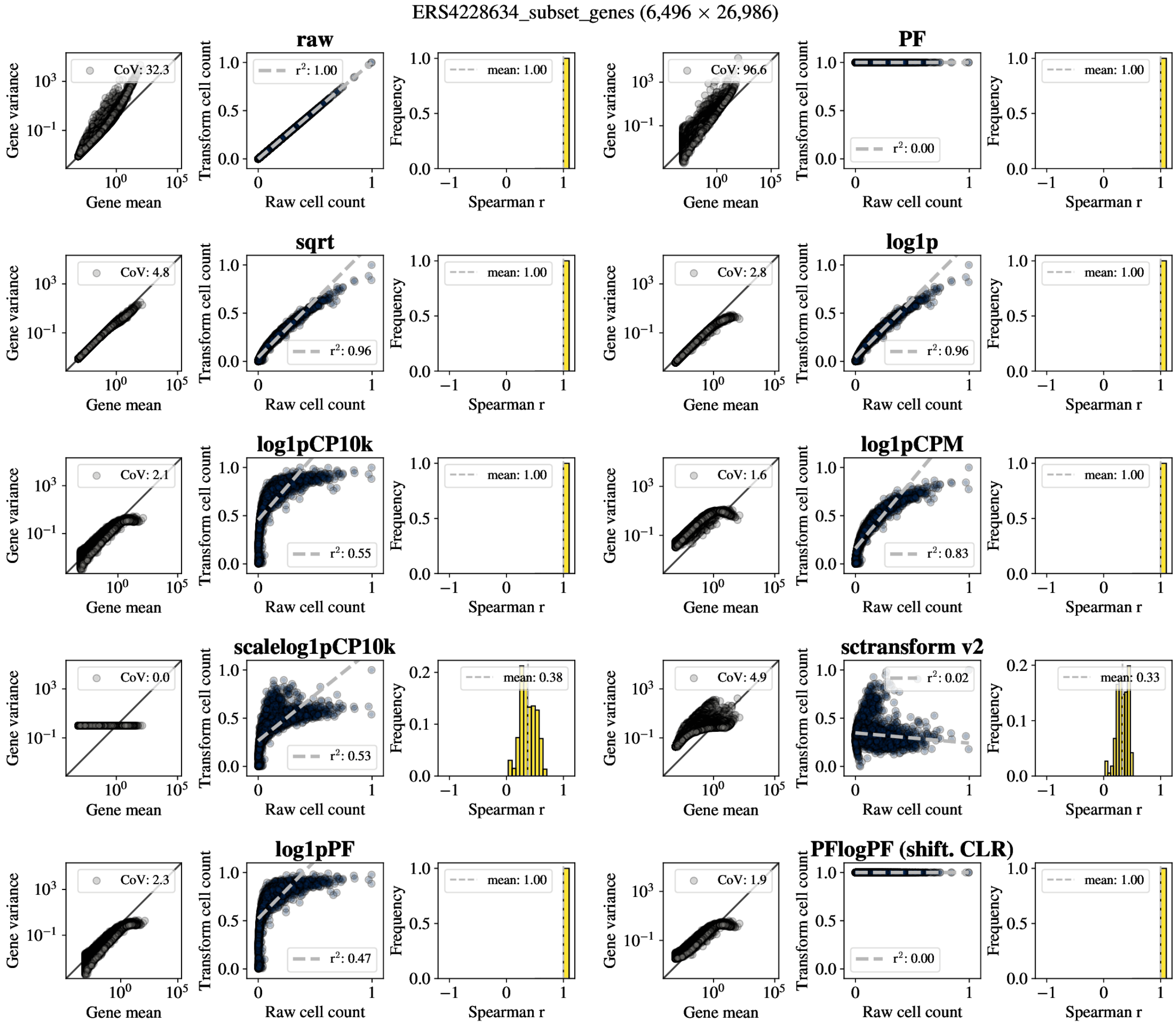


Supplementary Figure 1.98. **ERS4228634**. Per-method comparison of variance stabilization (left), depth normalization (middle), and monotonicity (right) for each normalization on dataset ERS4228634.

| **ERS4228635** | |
|---|---|
| cells | 7,247 |
| genes | 60,623 |
| nonzero entries | 11,713,592 |
| density | 0.0267 |
| total counts | 33,846,055 |
| mean counts / cell | 4,670 |
| mean counts / gene | 558 |
| min counts / cell | 160 |
| max counts / cell | 51,931 |
| overdispersion | 1.07 |

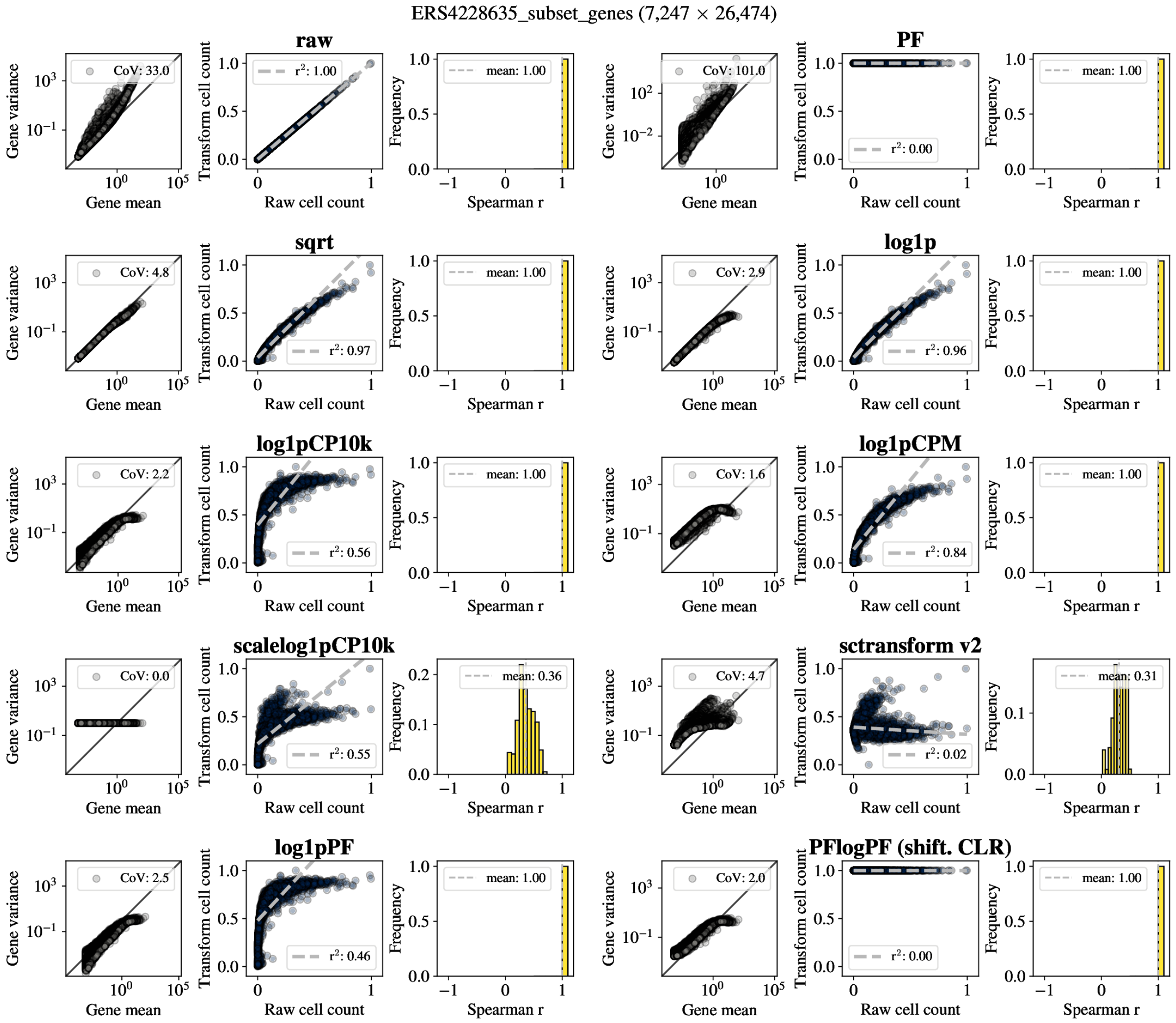


Supplementary Figure 1.99. **ERS4228635**. Per-method comparison of variance stabilization (left), depth normalization (middle), and monotonicity (right) for each normalization on dataset ERS4228635.

| ERS4228636 | |
|---|---|
| cells | 4,000 |
| genes | 60,623 |
| nonzero entries | 8,771,297 |
| density | 0.0362 |
| total counts | 26,889,181 |
| mean counts / cell | 6,722 |
| mean counts / gene | 444 |
| min counts / cell | 1,597 |
| max counts / cell | 53,382 |
| overdispersion | 0.968 |

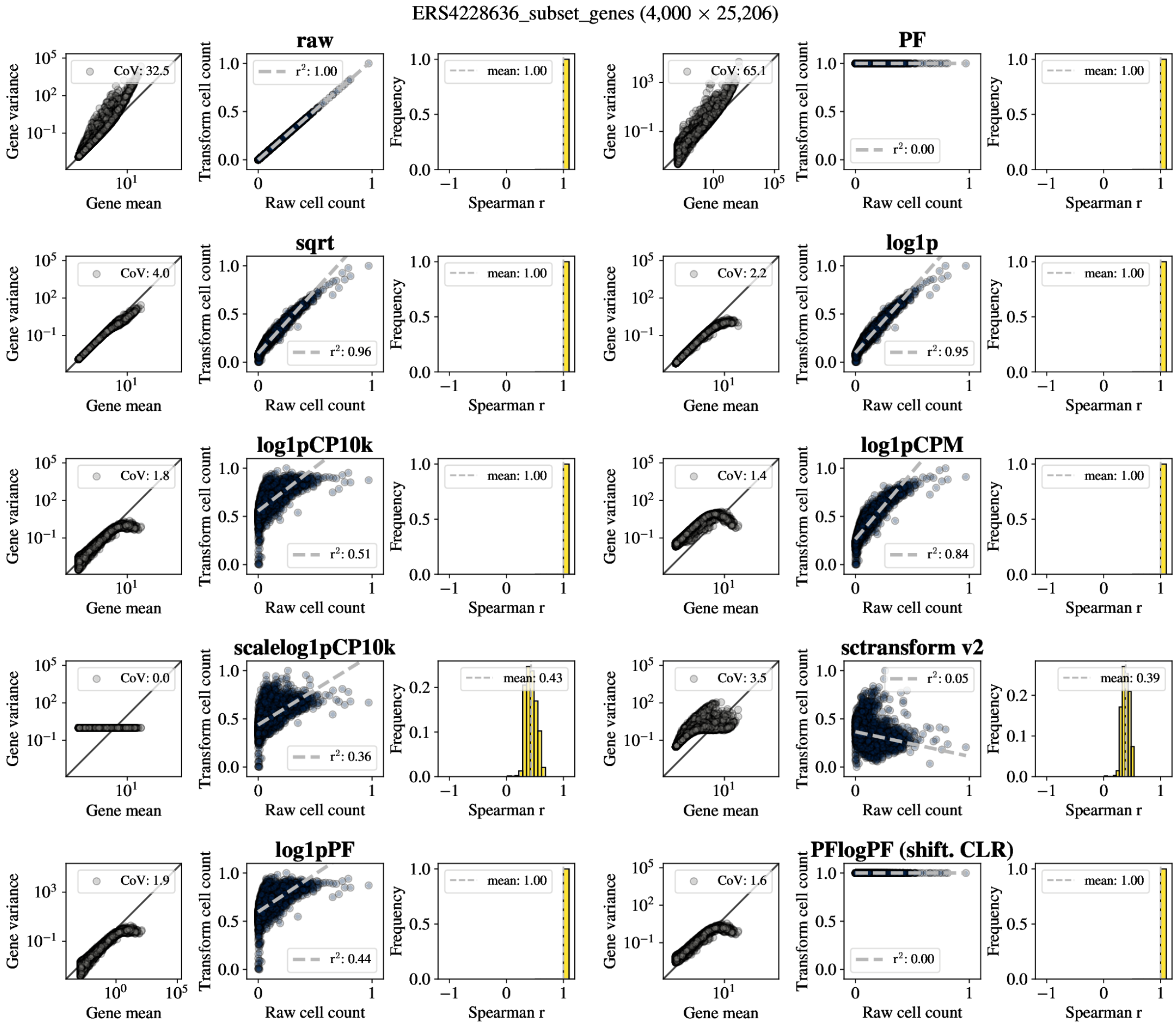


Supplementary Figure 1.100. **ERS4228636**. Per-method comparison of variance stabilization (left), depth normalization (middle), and monotonicity (right) for each normalization on dataset ERS4228636.

| ERS4228637 | |
|---|---|
| cells | 4,822 |
| genes | 60,623 |
| nonzero entries | 8,883,306 |
| density | 0.0304 |
| total counts | 26,983,322 |
| mean counts / cell | 5,596 |
| mean counts / gene | 445 |
| min counts / cell | 623 |
| max counts / cell | 57,735 |
| overdispersion | 1.11 |

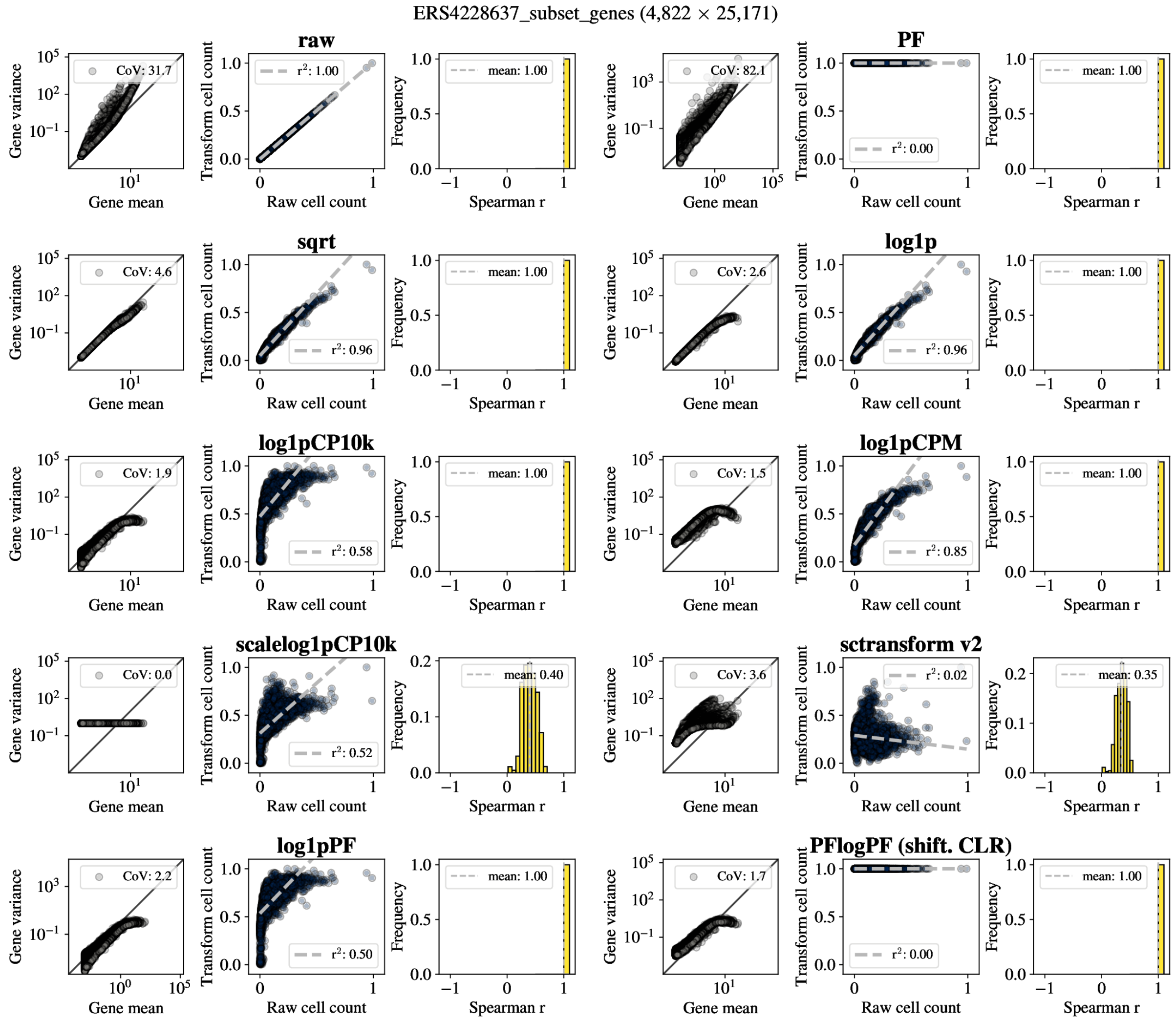


Supplementary Figure 1.101. **ERS4228637**. Per-method comparison of variance stabilization (left), depth normalization (middle), and monotonicity (right) for each normalization on dataset ERS4228637.

| **ERS4228638** | |
|---|---|
| cells | 5,289 |
| genes | 60,623 |
| nonzero entries | 10,626,705 |
| density | 0.0331 |
| total counts | 33,914,538 |
| mean counts / cell | 6,412 |
| mean counts / gene | 559 |
| min counts / cell | 573 |
| max counts / cell | 69,569 |
| overdispersion | 1.23 |

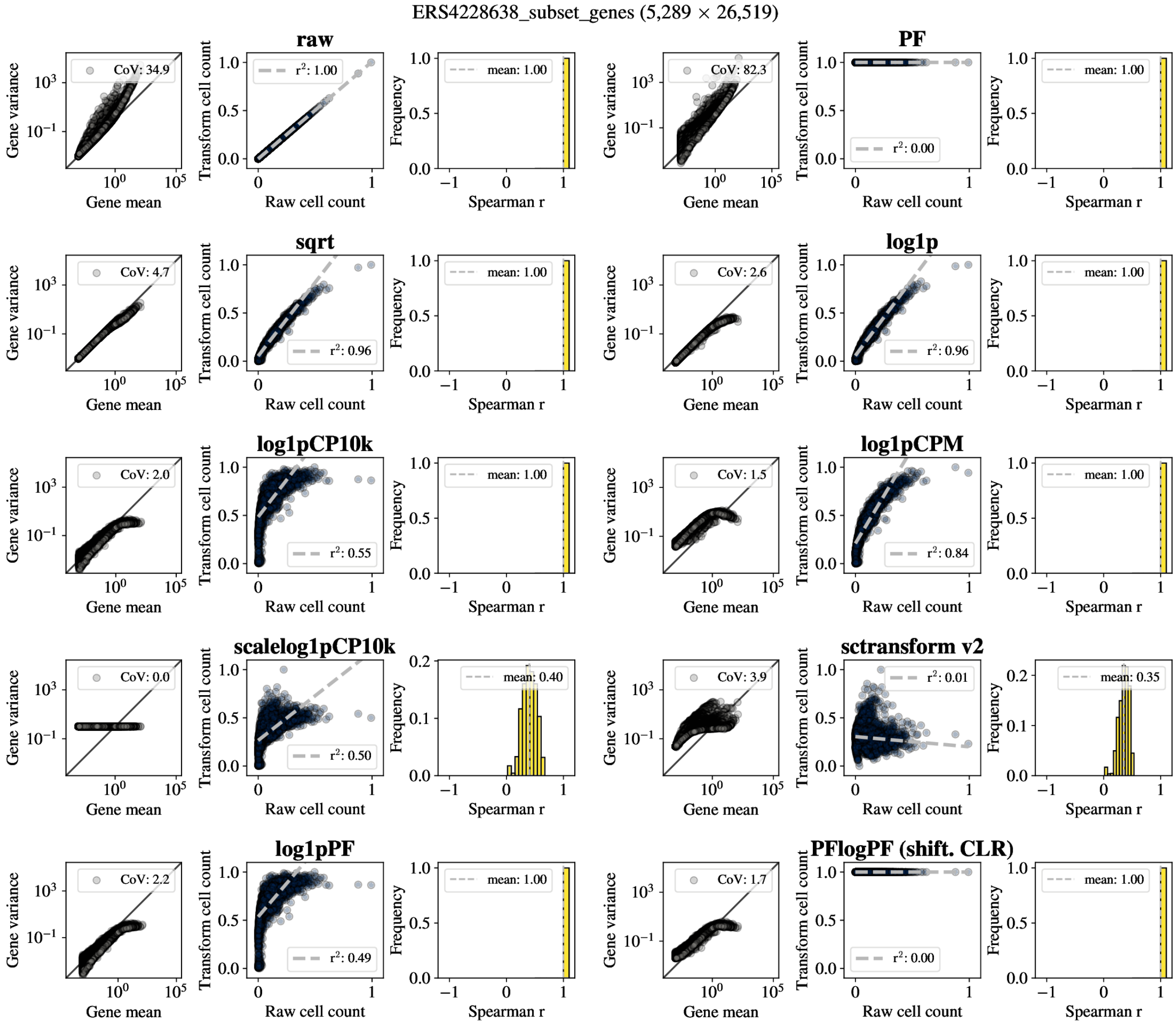


Supplementary Figure 1.102. **ERS4228638**. Per-method comparison of variance stabilization (left), depth normalization (middle), and monotonicity (right) for each normalization on dataset ERS4228638.

| ERS4228639 | |
|---|---|
| cells | 5,695 |
| genes | 60,623 |
| nonzero entries | 7,983,221 |
| density | 0.0231 |
| total counts | 21,787,329 |
| mean counts / cell | 3,826 |
| mean counts / gene | 359 |
| min counts / cell | 316 |
| max counts / cell | 38,972 |
| overdispersion | 1.32 |

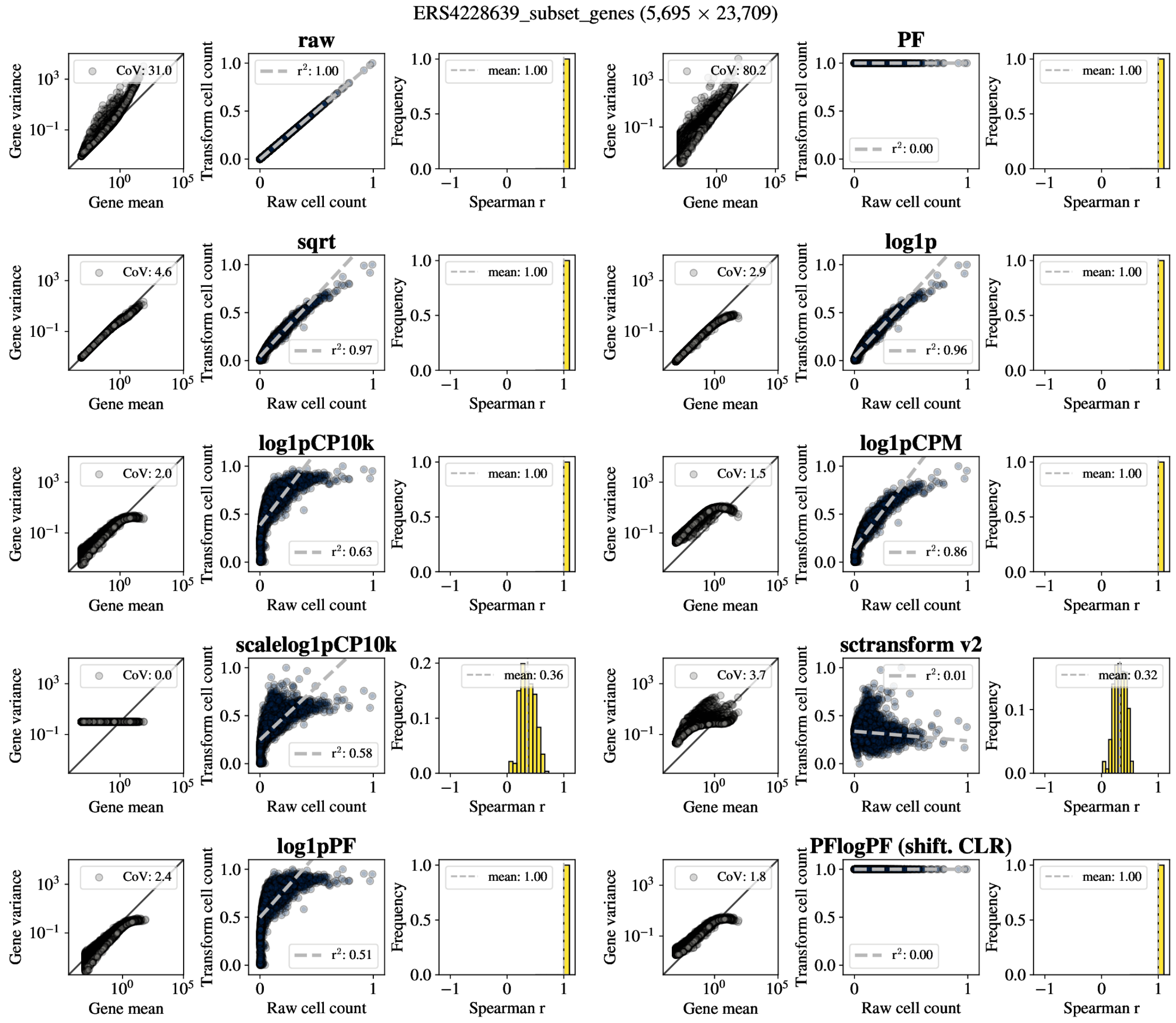


Supplementary Figure 1.103. **ERS4228639**. Per-method comparison of variance stabilization (left), depth normalization (middle), and monotonicity (right) for each normalization on dataset ERS4228639.

| ERS4228662 | |
|---|---|
| cells | 10,774 |
| genes | 60,623 |
| nonzero entries | 530,173 |
| density | 0.000812 |
| total counts | 919,373 |
| mean counts / cell | 85 |
| mean counts / gene | 15 |
| min counts / cell | 10 |
| max counts / cell | 2,485 |
| overdispersion | 4.09 |

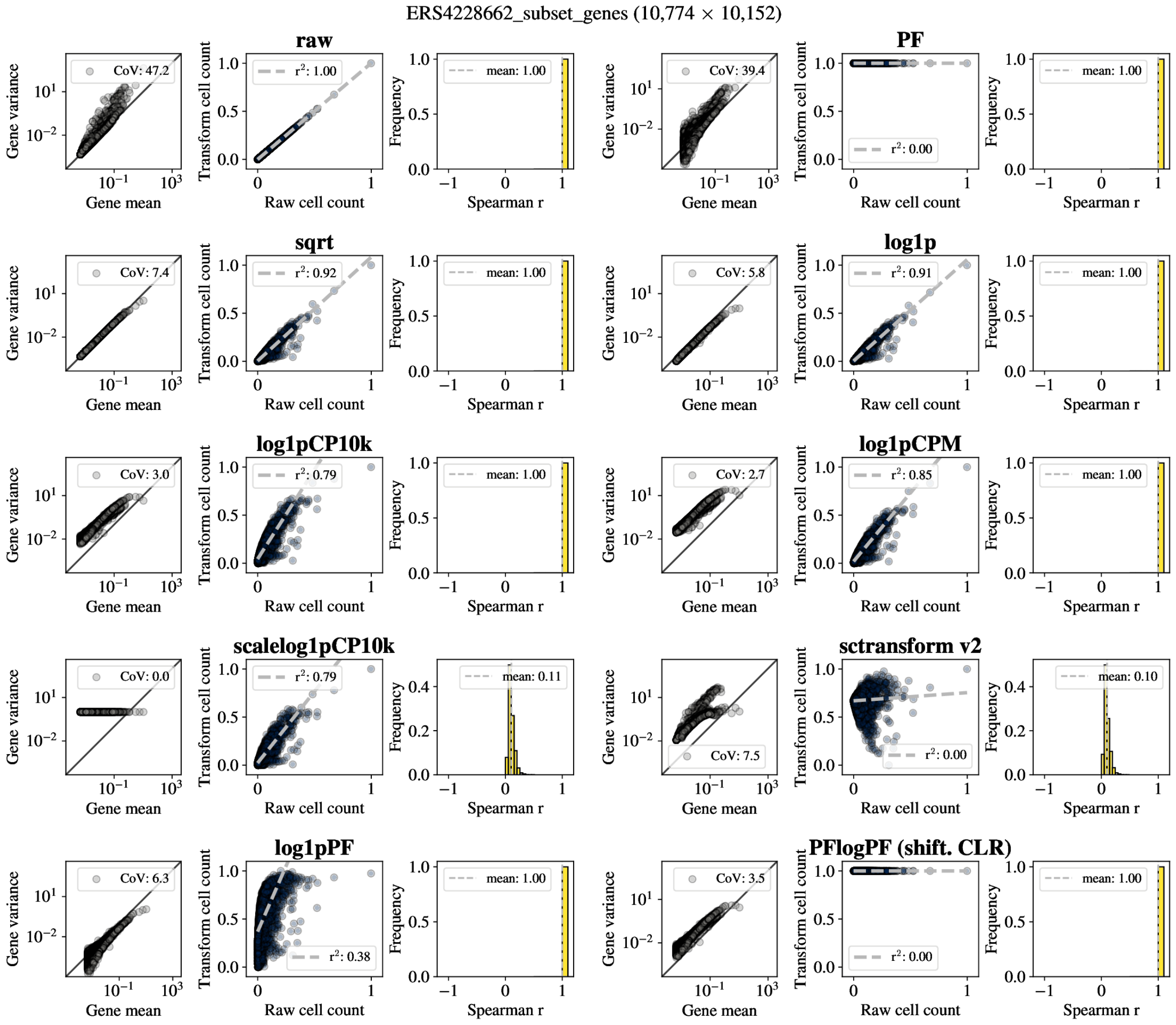


Supplementary Figure 1.104. **ERS4228662**. Per-method comparison of variance stabilization (left), depth normalization (middle), and monotonicity (right) for each normalization on dataset ERS4228662.

| **ERS4228663** | |
|---|---|
| cells | 3,140 |
| genes | 60,623 |
| nonzero entries | 241,579 |
| density | 0.00127 |
| total counts | 402,988 |
| mean counts / cell | 128 |
| mean counts / gene | 7 |
| min counts / cell | 27 |
| max counts / cell | 1,893 |
| overdispersion | 5.27 |

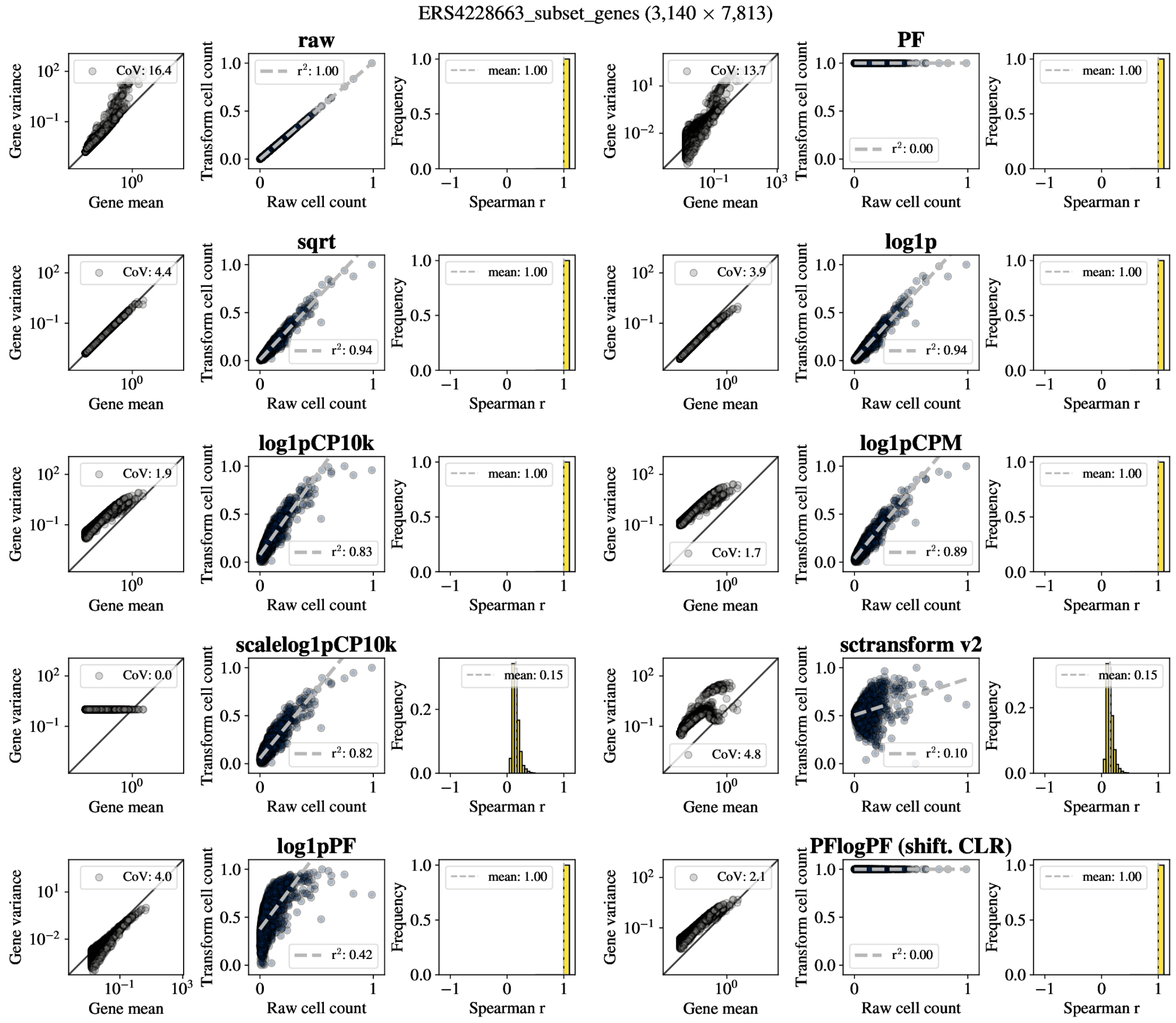


Supplementary Figure 1.105. **ERS4228663**. Per-method comparison of variance stabilization (left), depth normalization (middle), and monotonicity (right) for each normalization on dataset ERS4228663.

| ERS4228664 | |
|---|---|
| cells | 14,824 |
| genes | 60,623 |
| nonzero entries | 1,314,850 |
| density | 0.00146 |
| total counts | 1,981,708 |
| mean counts / cell | 134 |
| mean counts / gene | 33 |
| min counts / cell | 45 |
| max counts / cell | 2,352 |
| overdispersion | 1.27 |

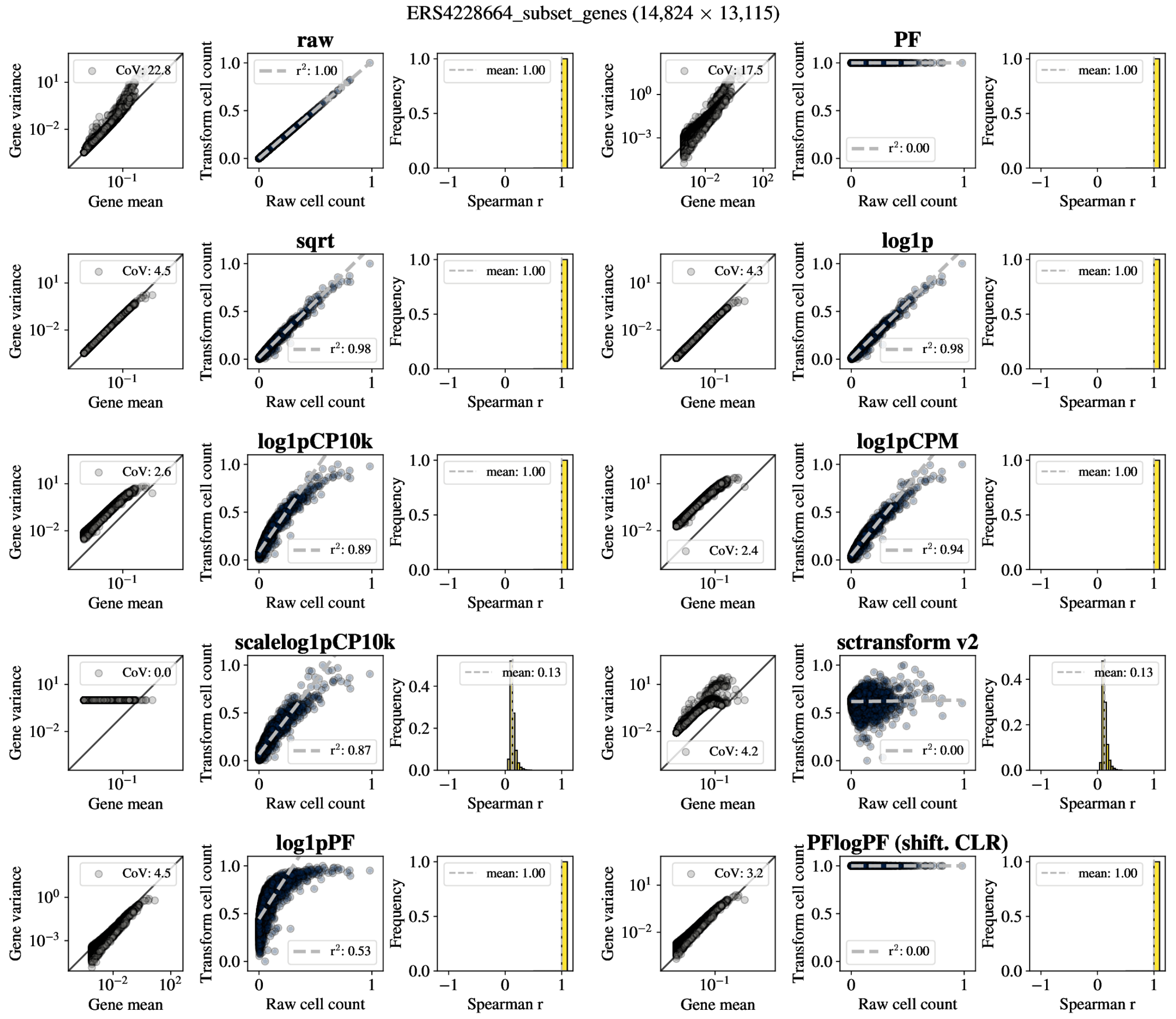


Supplementary Figure 1.106. **ERS4228664**. Per-method comparison of variance stabilization (left), depth normalization (middle), and monotonicity (right) for each normalization on dataset ERS4228664.

| **ERS4228665** | |
|---|---|
| cells | 1,183 |
| genes | 60,623 |
| nonzero entries | 126,698 |
| density | 0.00177 |
| total counts | 224,245 |
| mean counts / cell | 190 |
| mean counts / gene | 4 |
| min counts / cell | 88 |
| max counts / cell | 1,161 |
| overdispersion | 6.59 |

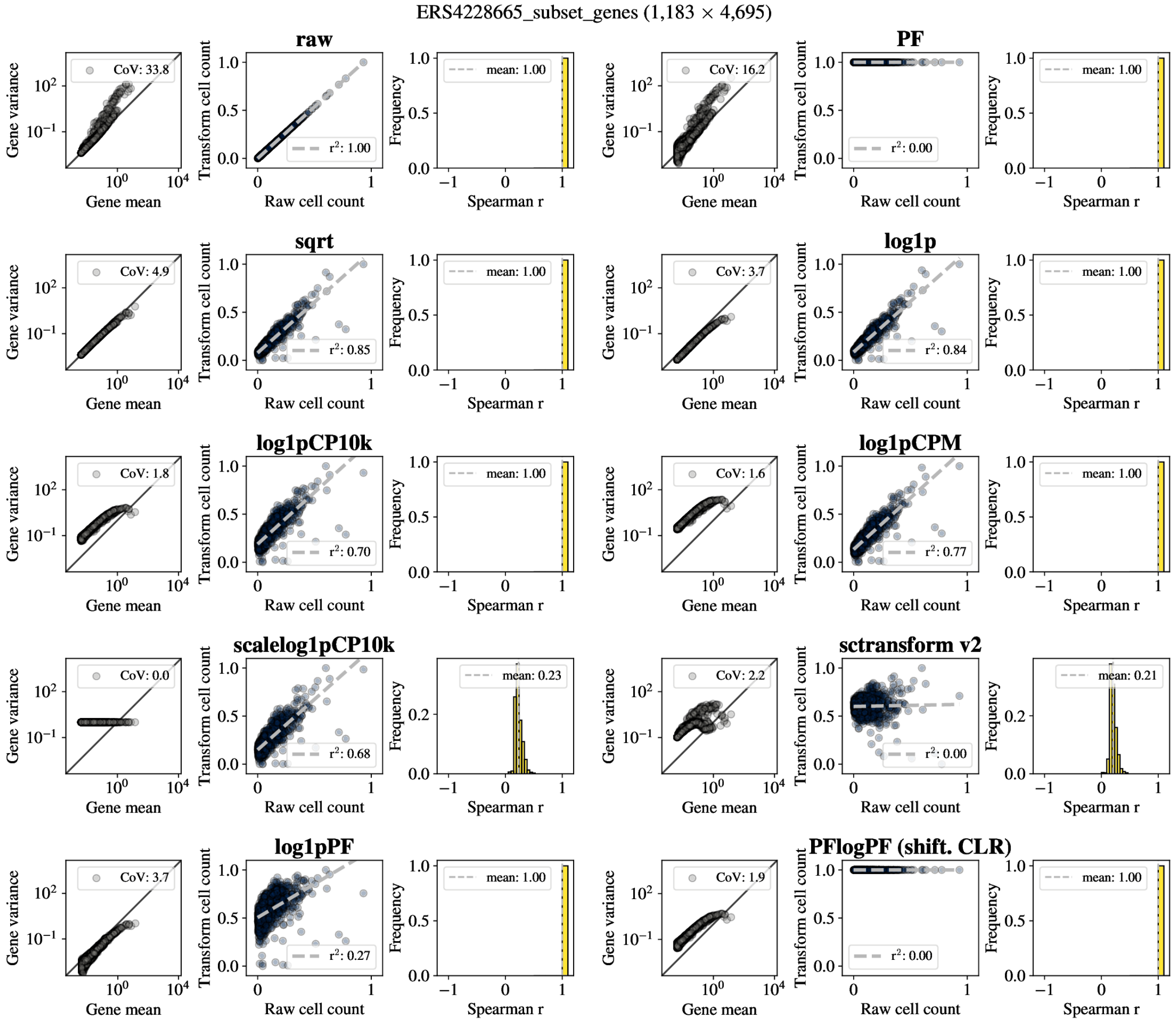


Supplementary Figure 1.107. **ERS4228665**. Per-method comparison of variance stabilization (left), depth normalization (middle), and monotonicity (right) for each normalization on dataset ERS4228665.

| **ERS4228668** | |
|---|---|
| cells | 6,117 |
| genes | 60,623 |
| nonzero entries | 471,203 |
| density | 0.00127 |
| total counts | 751,946 |
| mean counts / cell | 123 |
| mean counts / gene | 12 |
| min counts / cell | 16 |
| max counts / cell | 1,337 |
| overdispersion | 8.06 |

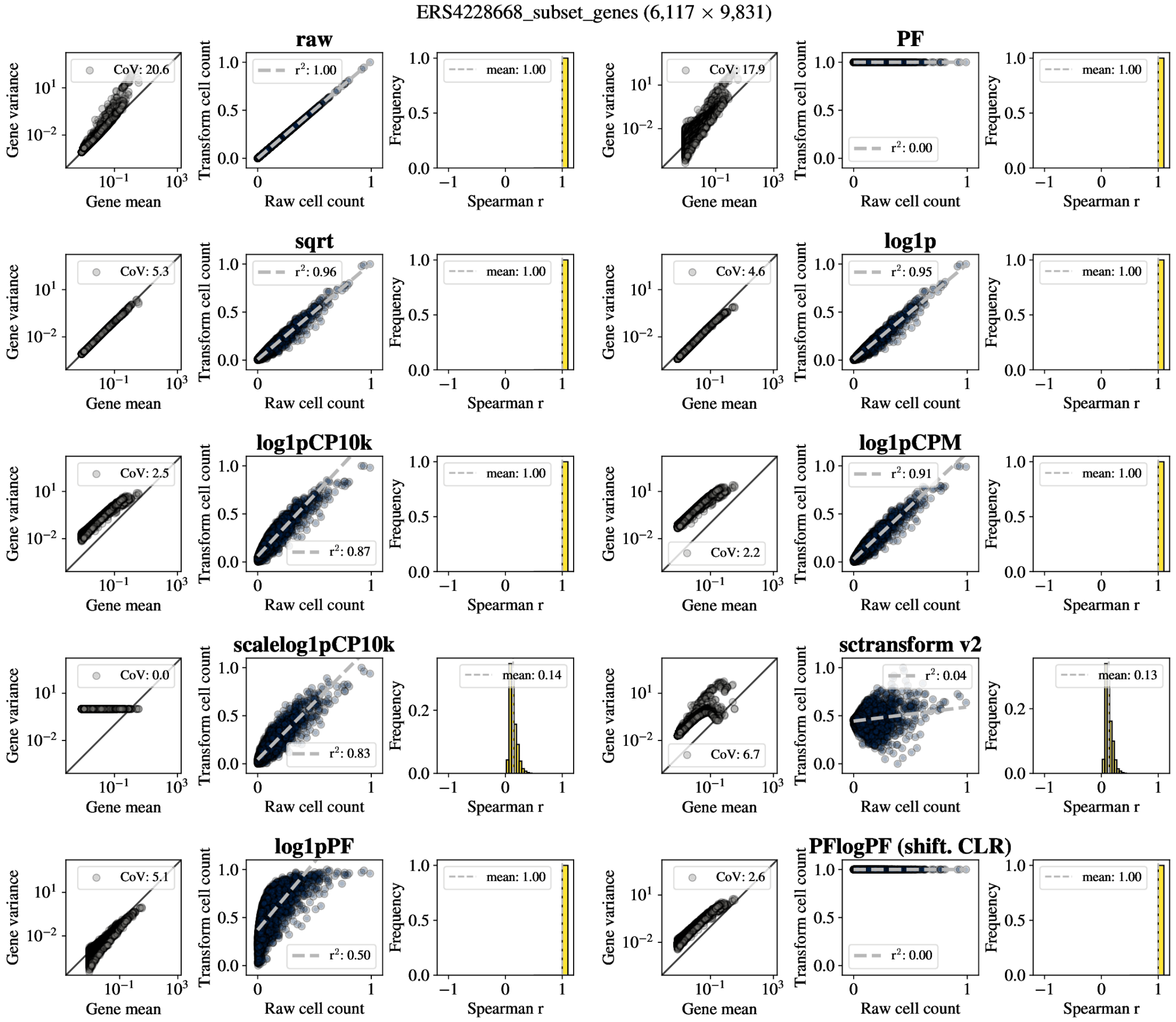


Supplementary Figure 1.108. **ERS4228668**. Per-method comparison of variance stabilization (left), depth normalization (middle), and monotonicity (right) for each normalization on dataset ERS4228668.

| **ERS4228669** | |
|---|---|
| cells | 6,414 |
| genes | 60,623 |
| nonzero entries | 463,174 |
| density | 0.00119 |
| total counts | 722,252 |
| mean counts / cell | 113 |
| mean counts / gene | 12 |
| min counts / cell | 18 |
| max counts / cell | 1,253 |
| overdispersion | 6.63 |

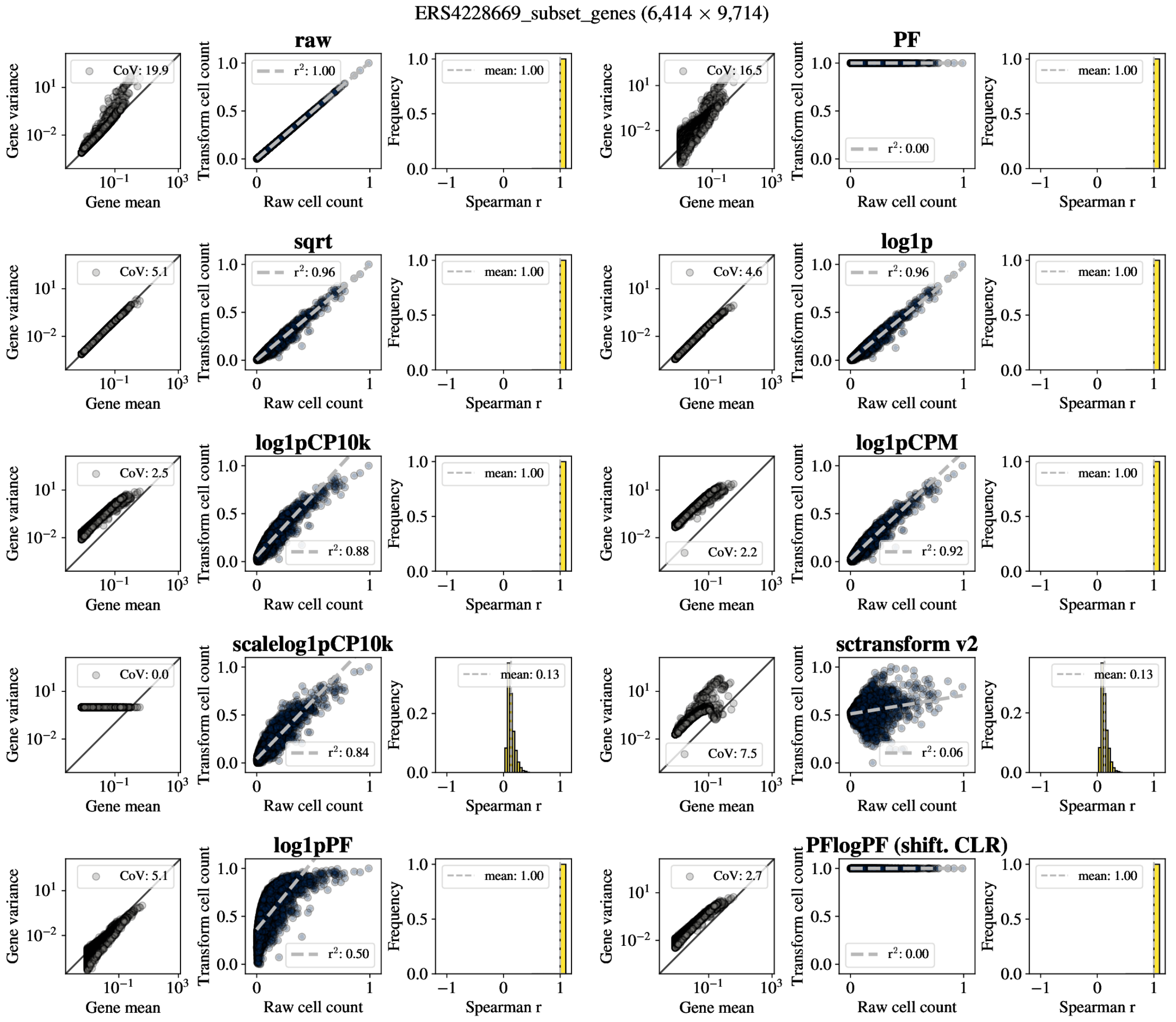


Supplementary Figure 1.109. **ERS4228669**. Per-method comparison of variance stabilization (left), depth normalization (middle), and monotonicity (right) for each normalization on dataset ERS4228669.

| ERS4228670 | |
|---|---|
| cells | 4,856 |
| genes | 60,623 |
| nonzero entries | 404,352 |
| density | 0.00137 |
| total counts | 662,769 |
| mean counts / cell | 136 |
| mean counts / gene | 11 |
| min counts / cell | 21 |
| max counts / cell | 2,211 |
| overdispersion | 4.7 |

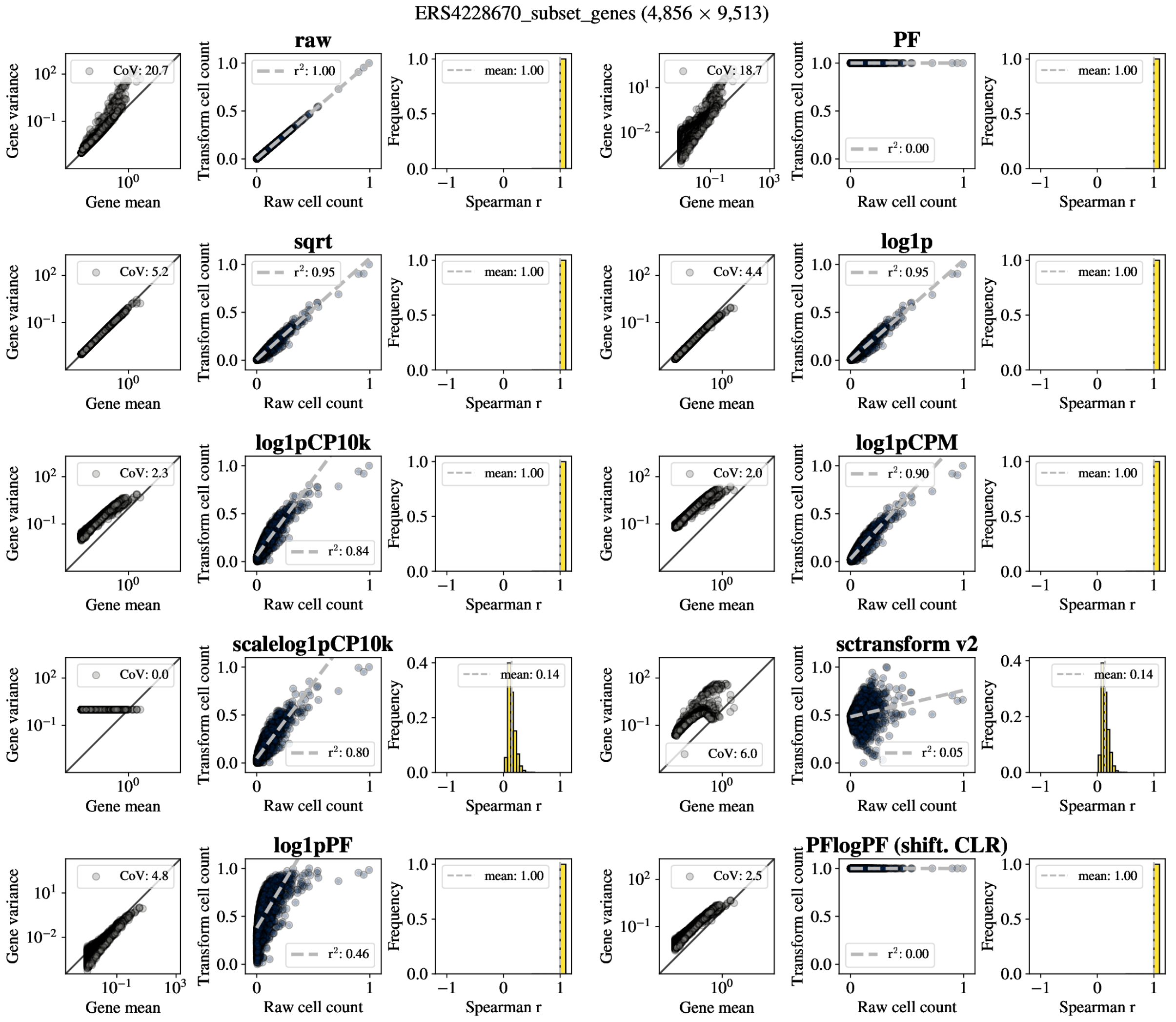


Supplementary Figure 1.110. **ERS4228670**. Per-method comparison of variance stabilization (left), depth normalization (middle), and monotonicity (right) for each normalization on dataset ERS4228670.

| ERS4228671 | |
|---|---|
| cells | 1,987 |
| genes | 60,623 |
| nonzero entries | 271,153 |
| density | 0.00225 |
| total counts | 446,408 |
| mean counts / cell | 225 |
| mean counts / gene | 7 |
| min counts / cell | 104 |
| max counts / cell | 1,823 |
| overdispersion | 3.97 |

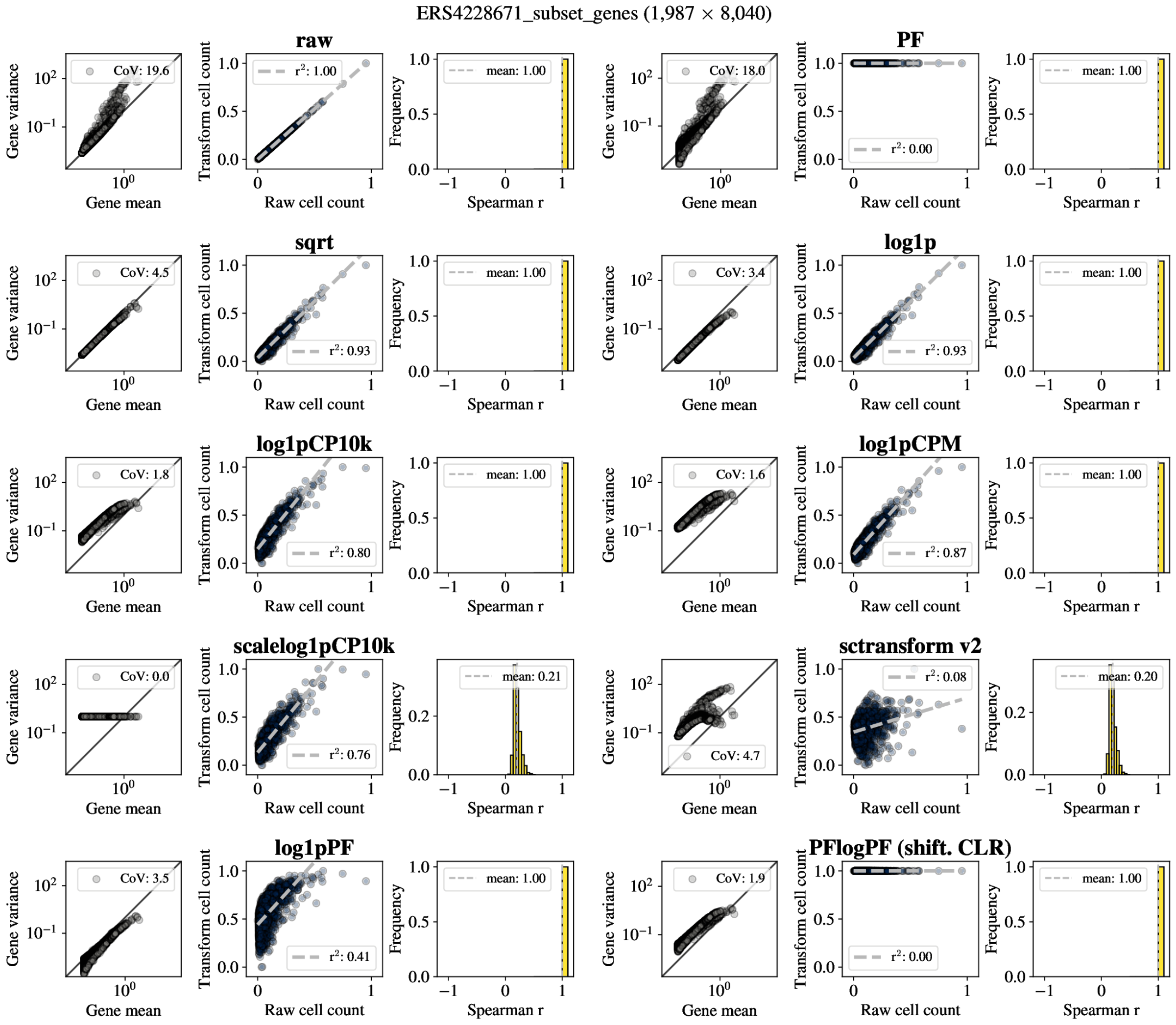


Supplementary Figure 1.111. **ERS4228671**. Per-method comparison of variance stabilization (left), depth normalization (middle), and monotonicity (right) for each normalization on dataset ERS4228671.

| ERS4228672 | |
|---|---|
| cells | 5,029 |
| genes | 60,623 |
| nonzero entries | 445,724 |
| density | 0.00146 |
| total counts | 707,571 |
| mean counts / cell | 141 |
| mean counts / gene | 12 |
| min counts / cell | 20 |
| max counts / cell | 1,737 |
| overdispersion | 3.56 |

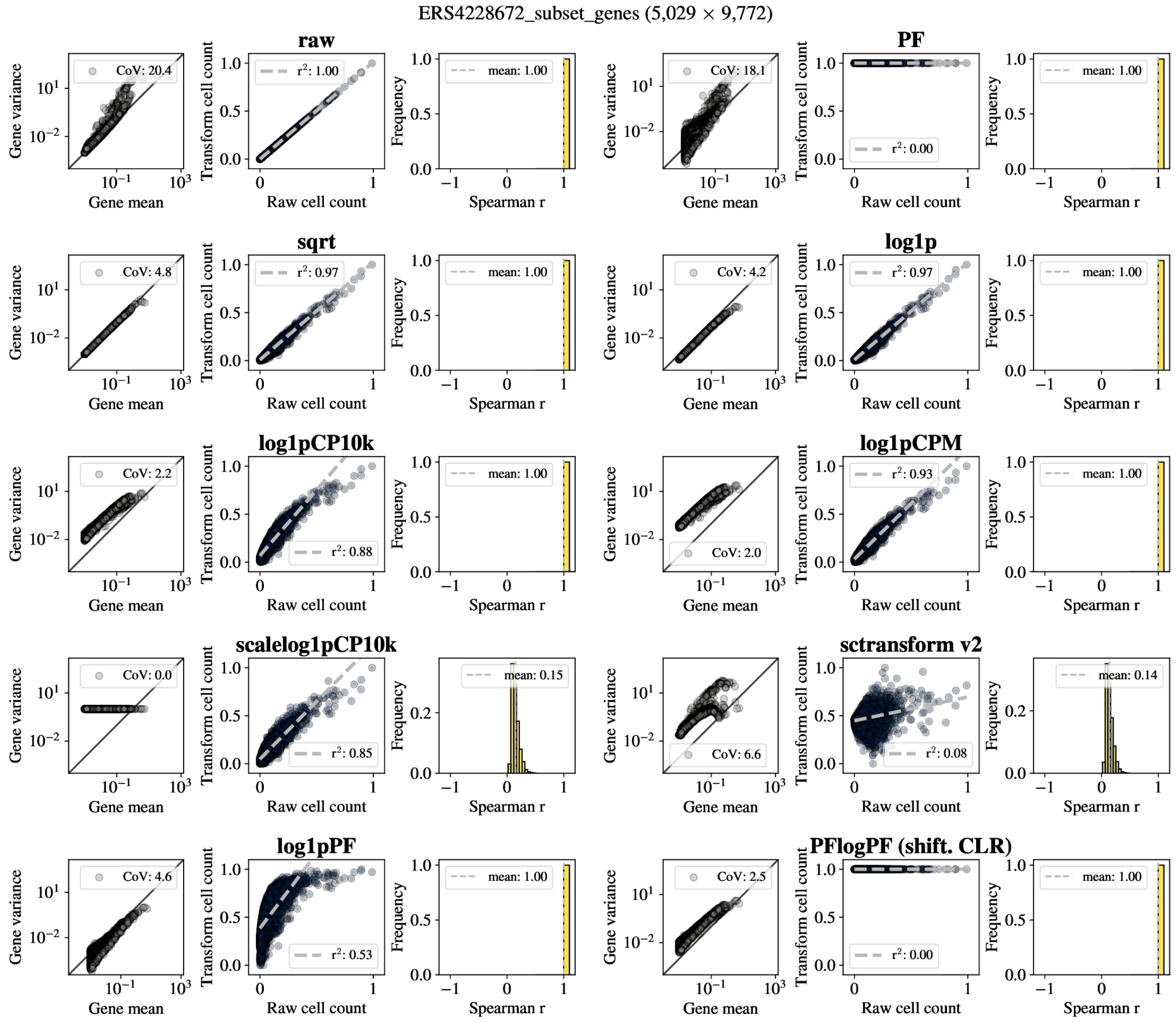


Supplementary Figure 1.112. **ERS4228672**. Per-method comparison of variance stabilization (left), depth normalization (middle), and monotonicity (right) for each normalization on dataset ERS4228672.

| ERS4228673 | |
|---|---|
| cells | 5,134 |
| genes | 60,623 |
| nonzero entries | 369,247 |
| density | 0.00119 |
| total counts | 622,727 |
| mean counts / cell | 121 |
| mean counts / gene | 10 |
| min counts / cell | 21 |
| max counts / cell | 1,301 |
| overdispersion | 6.19 |

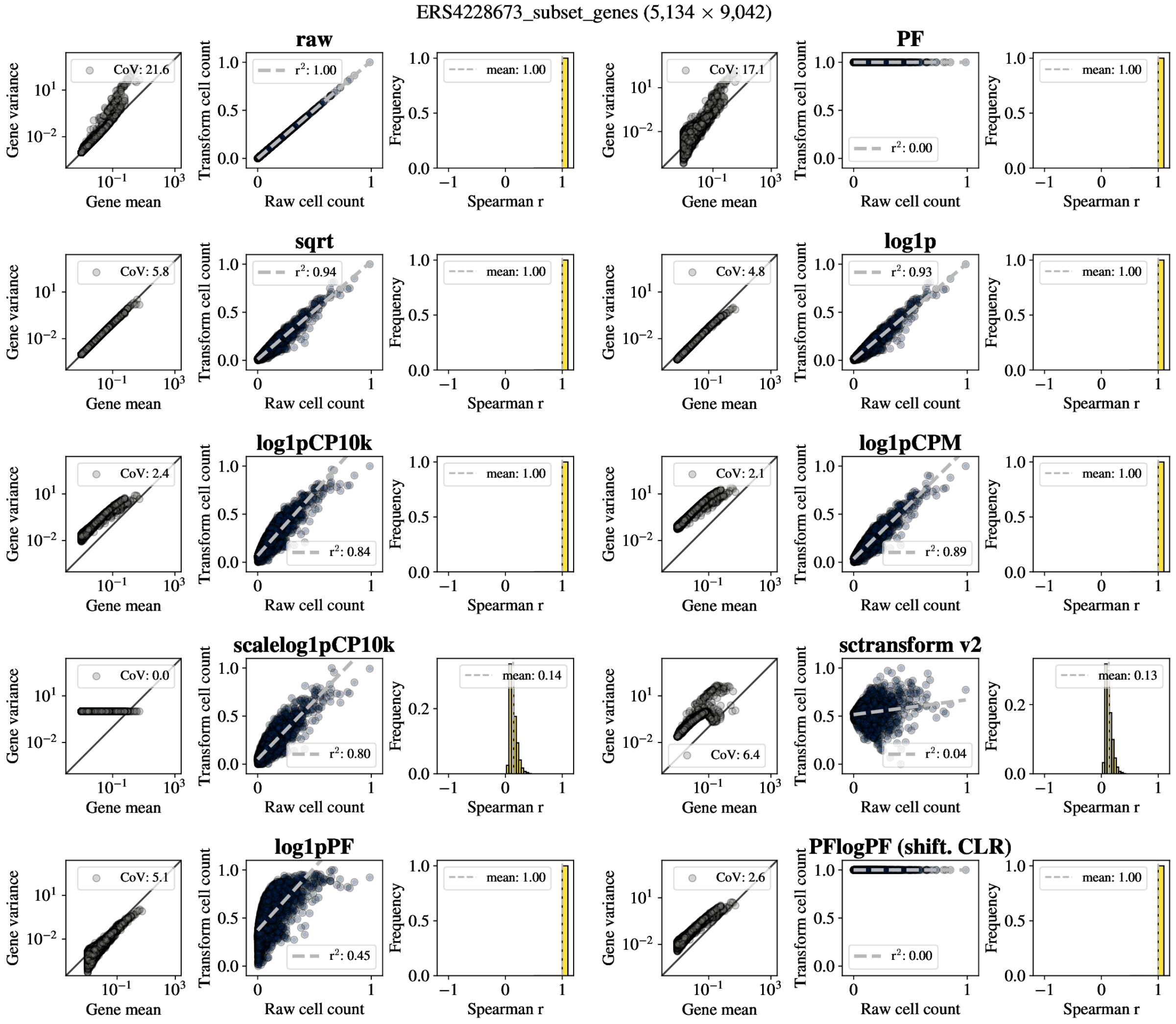


Supplementary Figure 1.113. **ERS4228673**. Per-method comparison of variance stabilization (left), depth normalization (middle), and monotonicity (right) for each normalization on dataset ERS4228673.

| ERX2756708 | |
|---|---|
| cells | 1,289 |
| genes | 60,623 |
| nonzero entries | 721,209 |
| density | 0.00923 |
| total counts | 2,017,149 |
| mean counts / cell | 1,565 |
| mean counts / gene | 33 |
| min counts / cell | 46 |
| max counts / cell | 54,628 |
| overdispersion | 3.6 |

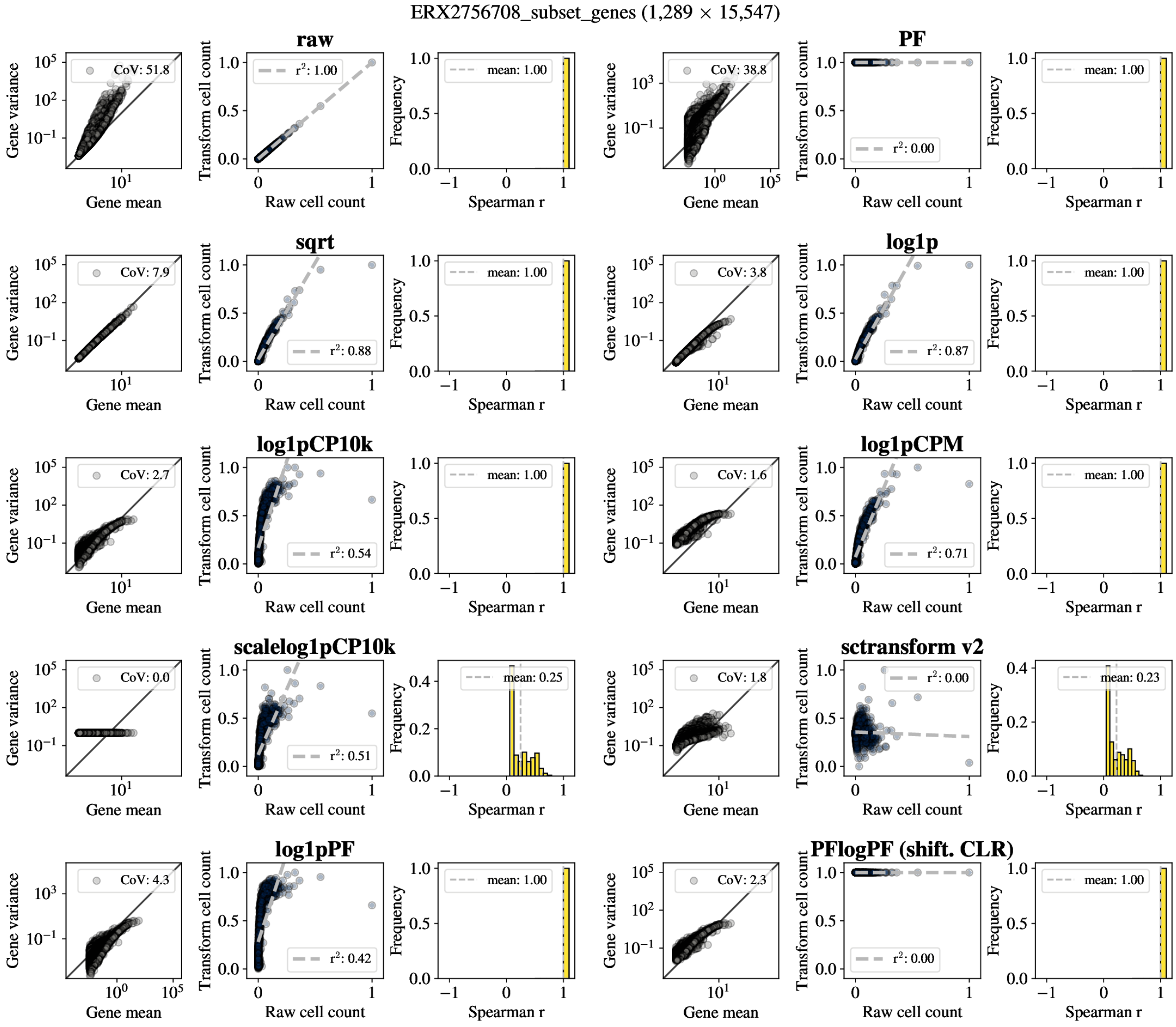


Supplementary Figure 1.114. **ERX2756708**. Per-method comparison of variance stabilization (left), depth normalization (middle), and monotonicity (right) for each normalization on dataset ERX2756708.

| ERX2756710 | |
|---|---|
| cells | 2,055 |
| genes | 60,623 |
| nonzero entries | 1,331,166 |
| density | 0.0107 |
| total counts | 3,918,662 |
| mean counts / cell | 1,907 |
| mean counts / gene | 65 |
| min counts / cell | 63 |
| max counts / cell | 44,025 |
| overdispersion | 4.63 |

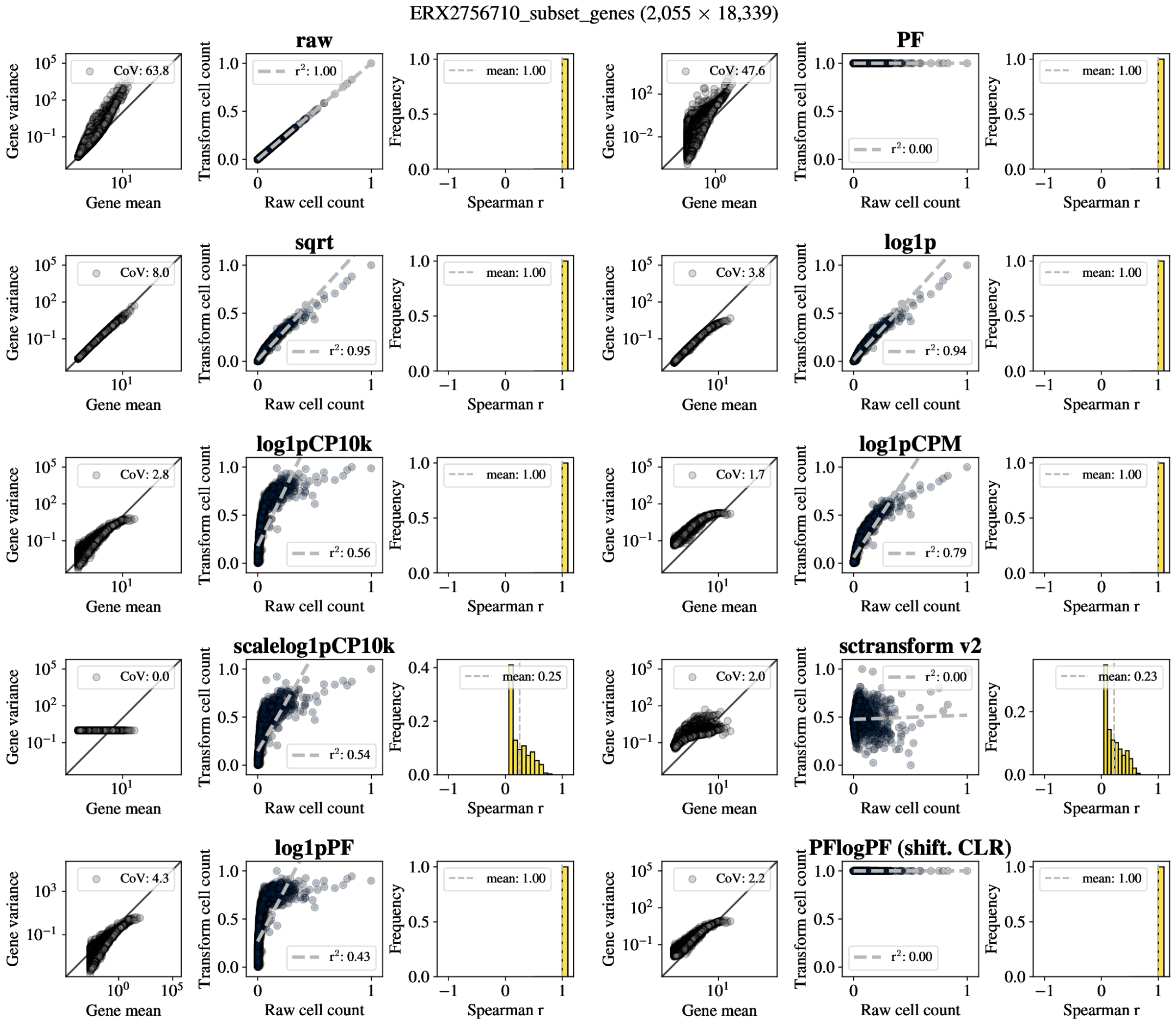


Supplementary Figure 1.115. **ERX2756710**. Per-method comparison of variance stabilization (left), depth normalization (middle), and monotonicity (right) for each normalization on dataset ERX2756710.

| ERX2756711 | |
|---|---|
| cells | 700 |
| genes | 60,623 |
| nonzero entries | 652,674 |
| density | 0.0154 |
| total counts | 1,516,586 |
| mean counts / cell | 2,167 |
| mean counts / gene | 25 |
| min counts / cell | 1,171 |
| max counts / cell | 7,371 |
| overdispersion | 0.312 |

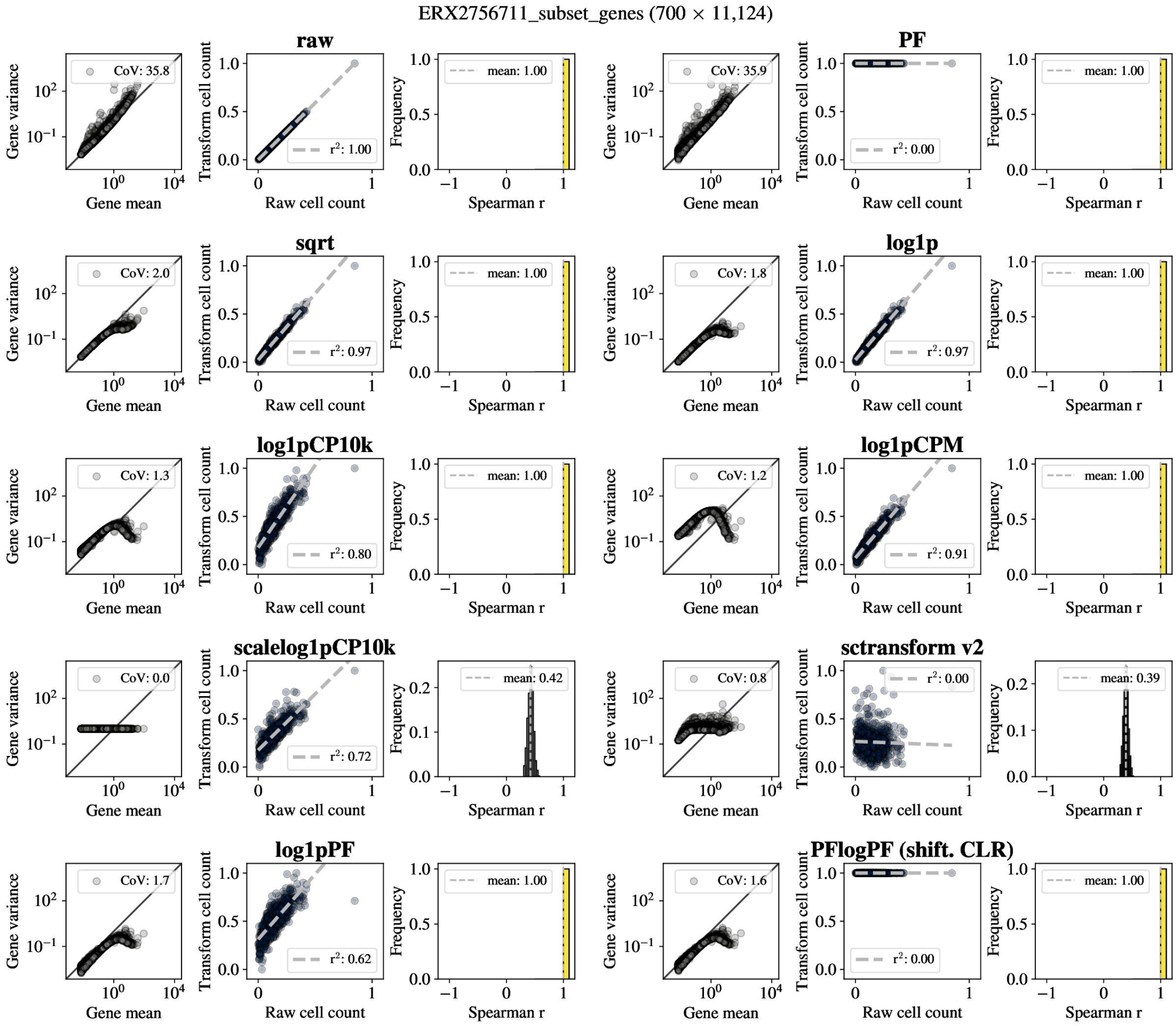


Supplementary Figure 1.116. **ERX2756711**. Per-method comparison of variance stabilization (left), depth normalization (middle), and monotonicity (right) for each normalization on dataset ERX2756711.

| **ERX2756712** | |
|---|---|
| cells | 70,373 |
| genes | 60,623 |
| nonzero entries | 10,623,887 |
| density | 0.00249 |
| total counts | 19,501,000 |
| mean counts / cell | 277 |
| mean counts / gene | 322 |
| min counts / cell | 66 |
| max counts / cell | 47,456 |
| overdispersion | 10.5 |

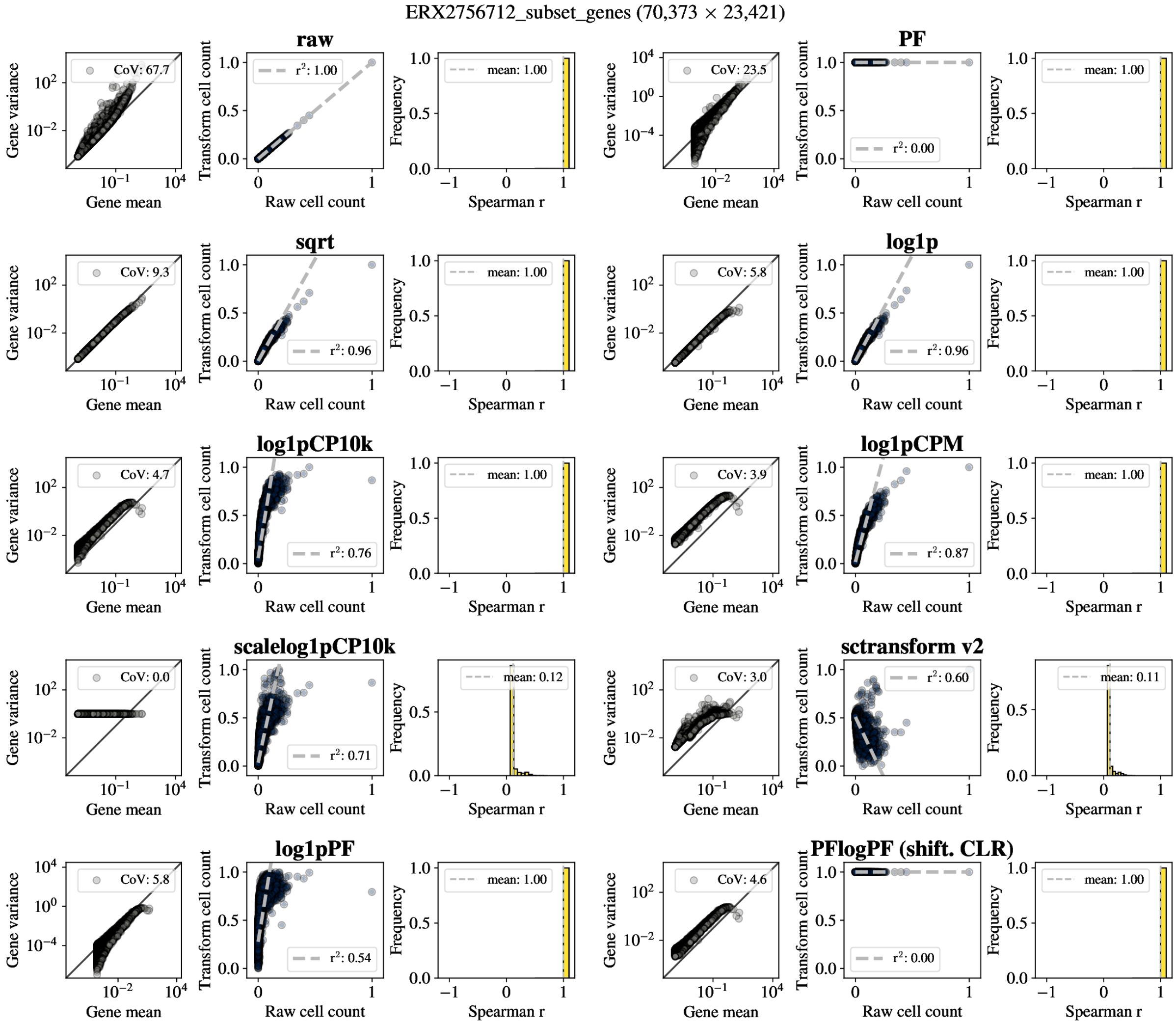


Supplementary Figure 1.117. **ERX2756712**. Per-method comparison of variance stabilization (left), depth normalization (middle), and monotonicity (right) for each normalization on dataset ERX2756712.

| ERX2756713 | |
|---|---|
| cells | 7,362 |
| genes | 60,623 |
| nonzero entries | 6,162,146 |
| density | 0.0138 |
| total counts | 18,538,652 |
| mean counts / cell | 2,518 |
| mean counts / gene | 306 |
| min counts / cell | 201 |
| max counts / cell | 40,666 |
| overdispersion | 4.58 |

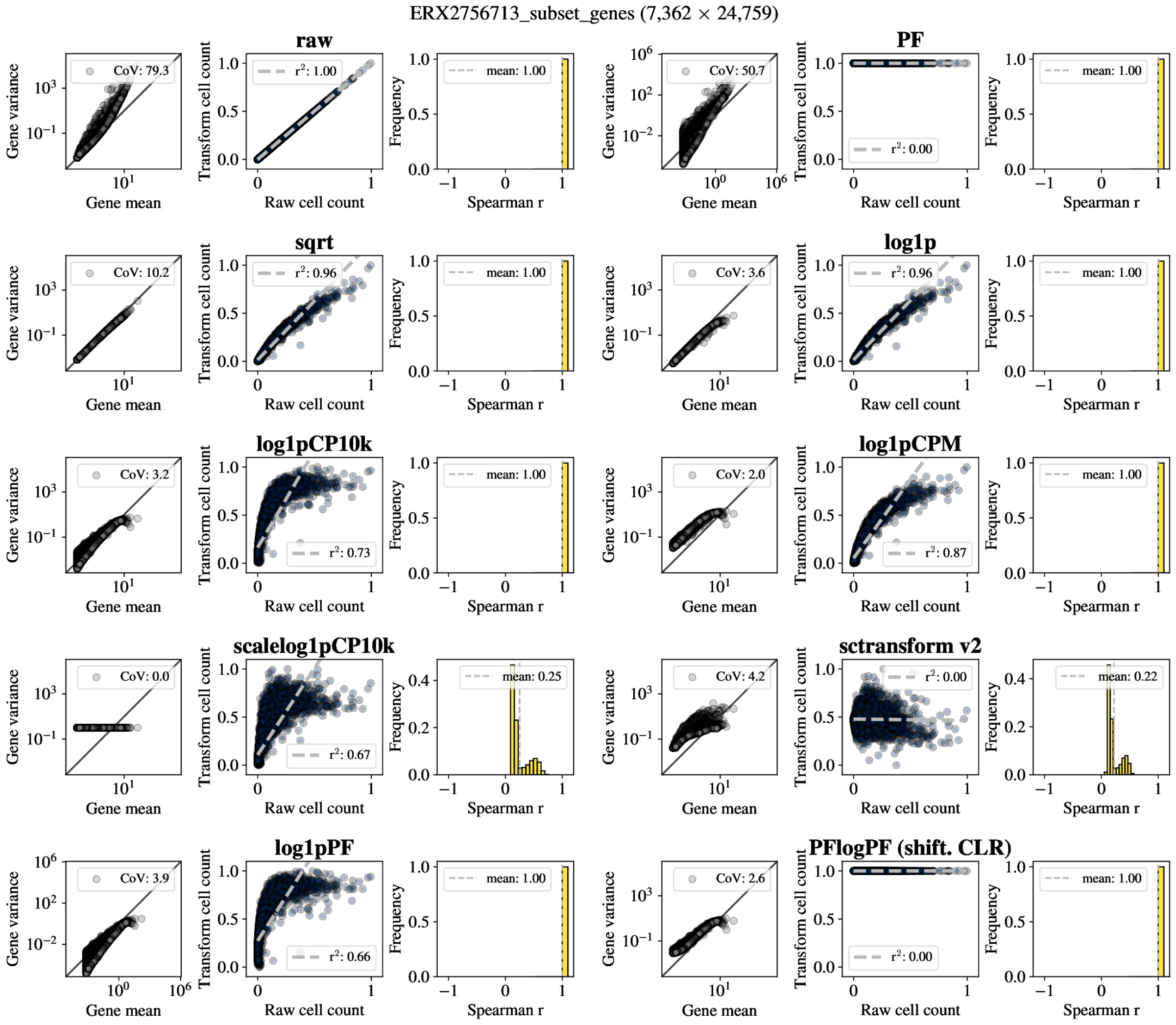


Supplementary Figure 1.118. **ERX2756713**. Per-method comparison of variance stabilization (left), depth normalization (middle), and monotonicity (right) for each normalization on dataset ERX2756713.

| ERX2756714 | |
|---|---|
| cells | 12,365 |
| genes | 60,623 |
| nonzero entries | 17,160,673 |
| density | 0.0229 |
| total counts | 56,689,528 |
| mean counts / cell | 4,585 |
| mean counts / gene | 935 |
| min counts / cell | 543 |
| max counts / cell | 47,055 |
| overdispersion | 1.7 |

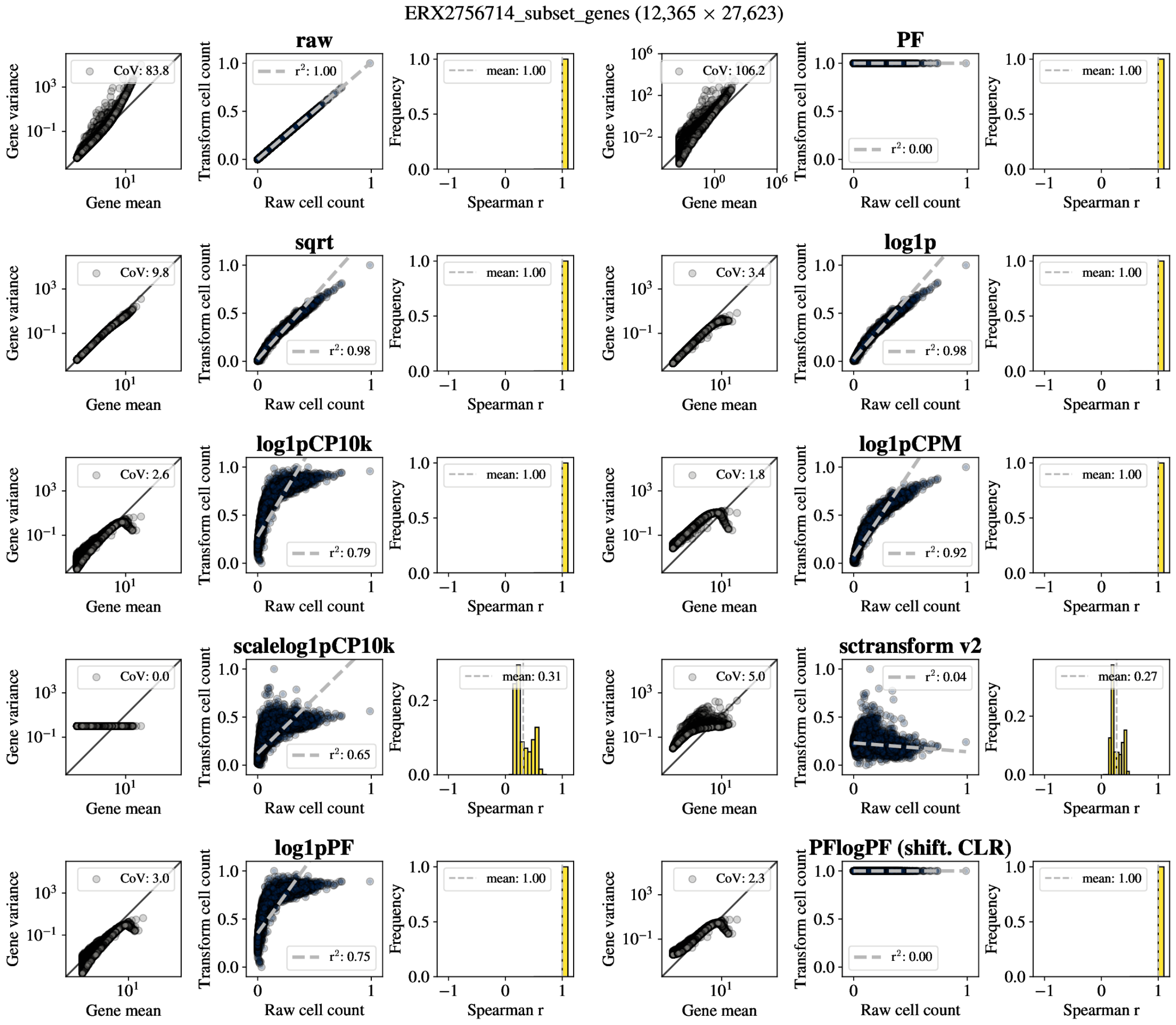


Supplementary Figure 1.119. **ERX2756714**. Per-method comparison of variance stabilization (left), depth normalization (middle), and monotonicity (right) for each normalization on dataset ERX2756714.

| ERX2756718 | |
|---|---|
| cells | 5,224 |
| genes | 60,623 |
| nonzero entries | 922,407 |
| density | 0.00291 |
| total counts | 2,522,399 |
| mean counts / cell | 483 |
| mean counts / gene | 42 |
| min counts / cell | 76 |
| max counts / cell | 40,865 |
| overdispersion | 13.2 |

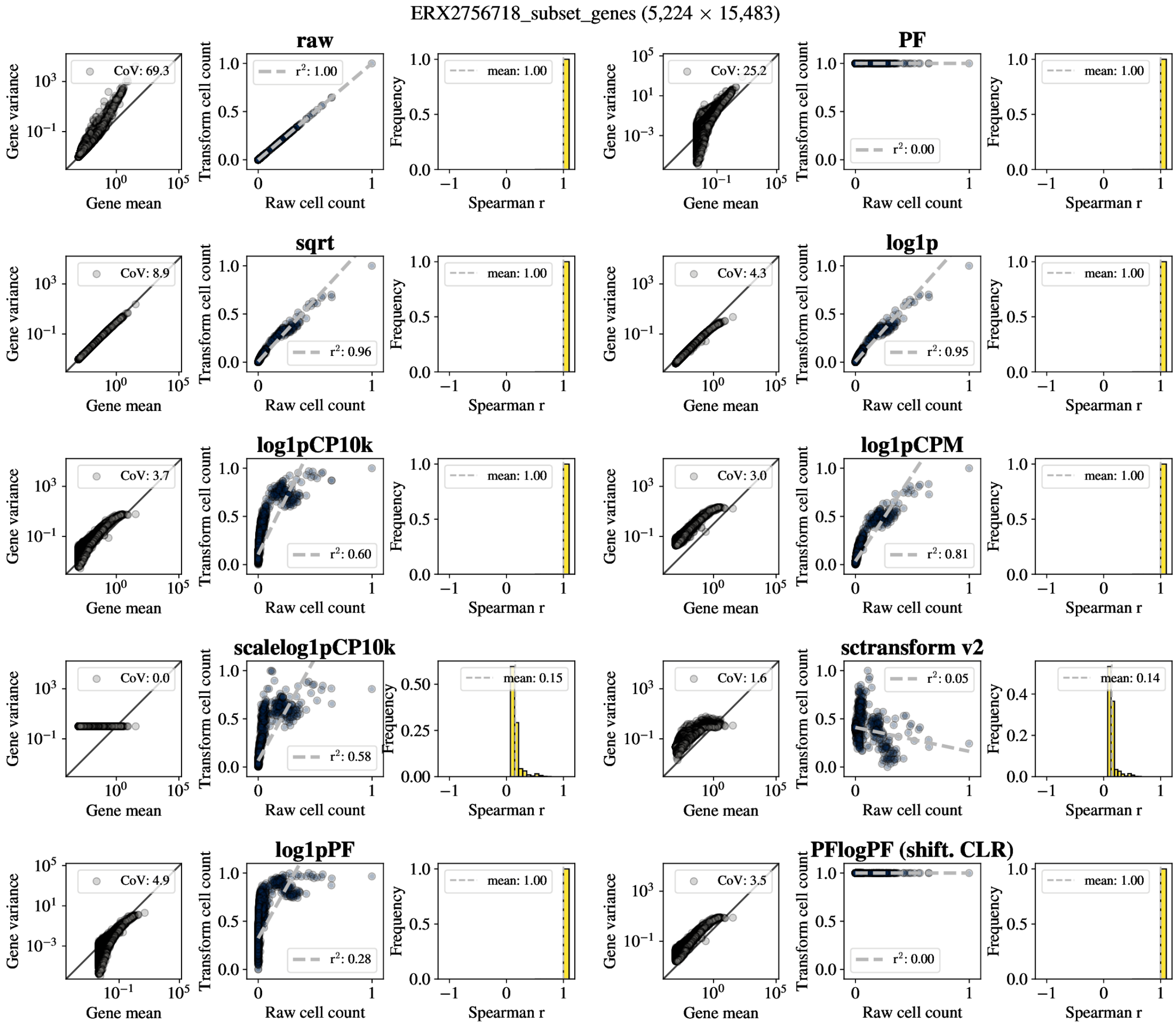


Supplementary Figure 1.120. **ERX2756718**. Per-method comparison of variance stabilization (left), depth normalization (middle), and monotonicity (right) for each normalization on dataset ERX2756718.

| ERX2756720 | |
|---|---|
| cells | 60,000 |
| genes | 60,623 |
| nonzero entries | 3,212,433 |
| density | 0.000883 |
| total counts | 4,939,009 |
| mean counts / cell | 82 |
| mean counts / gene | 81 |
| min counts / cell | 20 |
| max counts / cell | 7,886 |
| overdispersion | 13.9 |

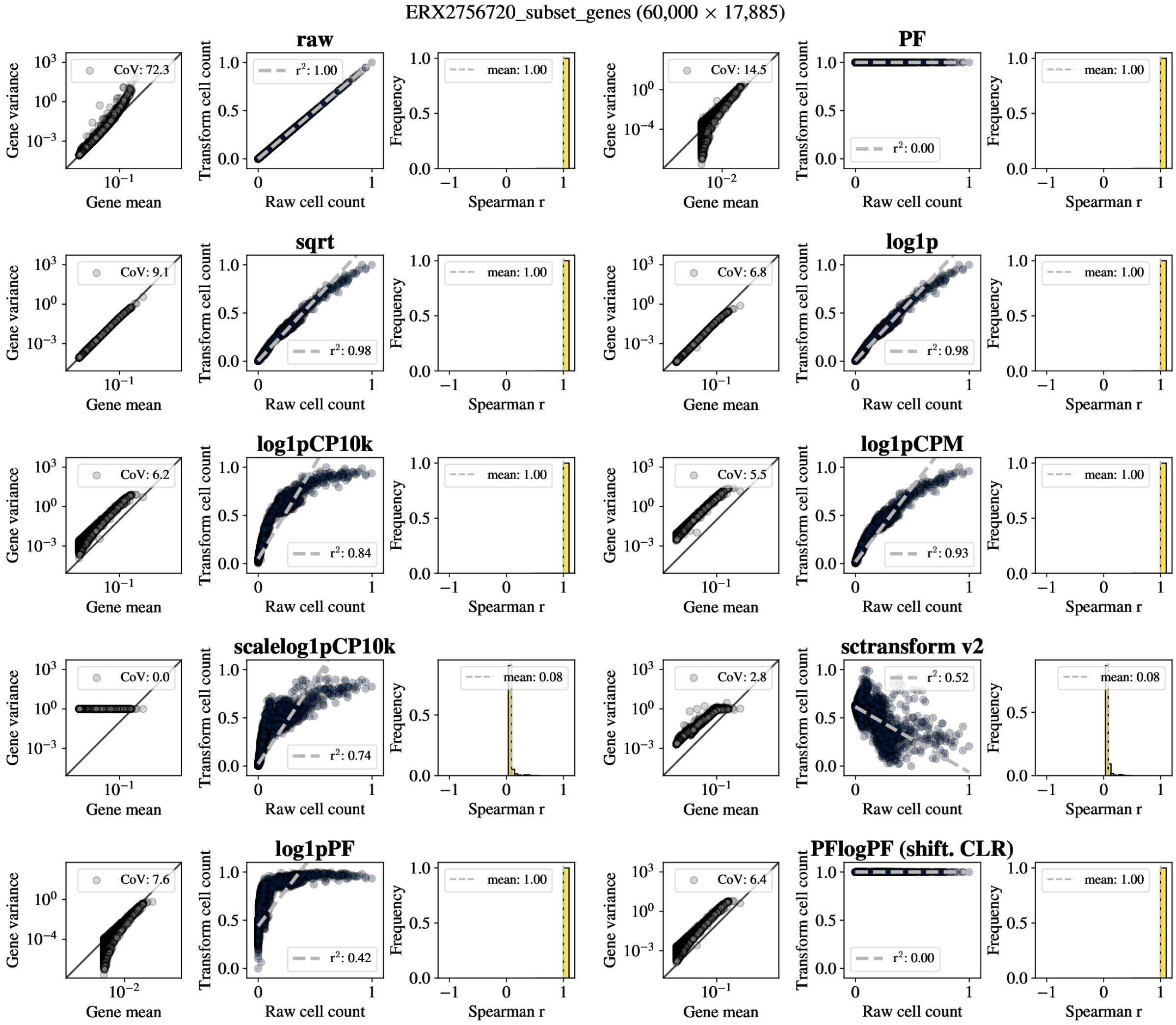


Supplementary Figure 1.121. **ERX2756720**. Per-method comparison of variance stabilization (left), depth normalization (middle), and monotonicity (right) for each normalization on dataset ERX2756720.

| **ERX2756721** | |
|---|---|
| cells | 9,293 |
| genes | 60,623 |
| nonzero entries | 1,636,093 |
| density | 0.0029 |
| total counts | 2,518,054 |
| mean counts / cell | 271 |
| mean counts / gene | 42 |
| min counts / cell | 31 |
| max counts / cell | 5,888 |
| overdispersion | 3.01 |

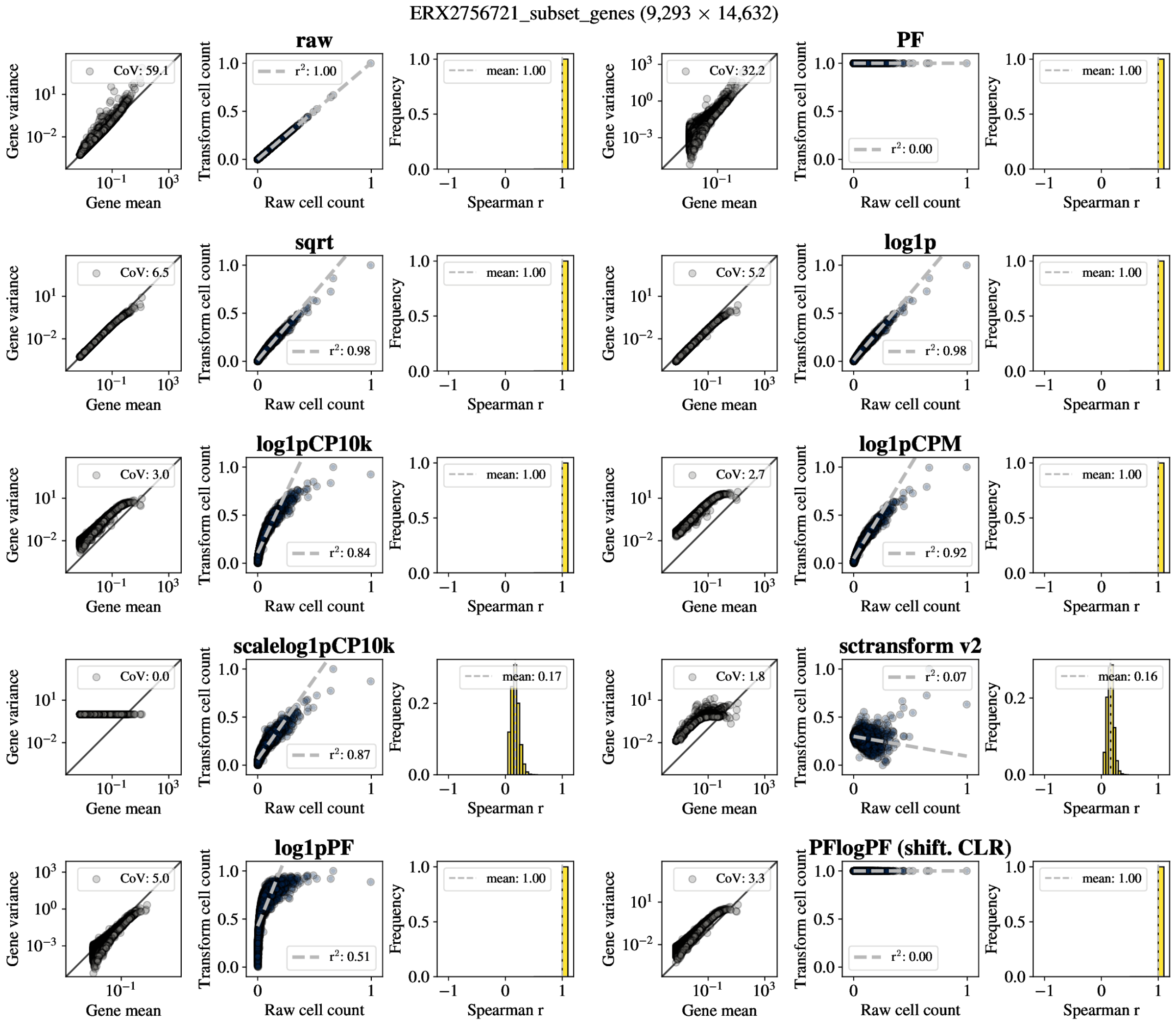


Supplementary Figure 1.122. **ERX2756721**. Per-method comparison of variance stabilization (left), depth normalization (middle), and monotonicity (right) for each normalization on dataset ERX2756721.

| ERX2756722 | |
|---|---|
| cells | 4,916 |
| genes | 60,623 |
| nonzero entries | 1,989,523 |
| density | 0.00668 |
| total counts | 3,831,458 |
| mean counts / cell | 779 |
| mean counts / gene | 63 |
| min counts / cell | 6 |
| max counts / cell | 5,354 |
| overdispersion | 0.847 |

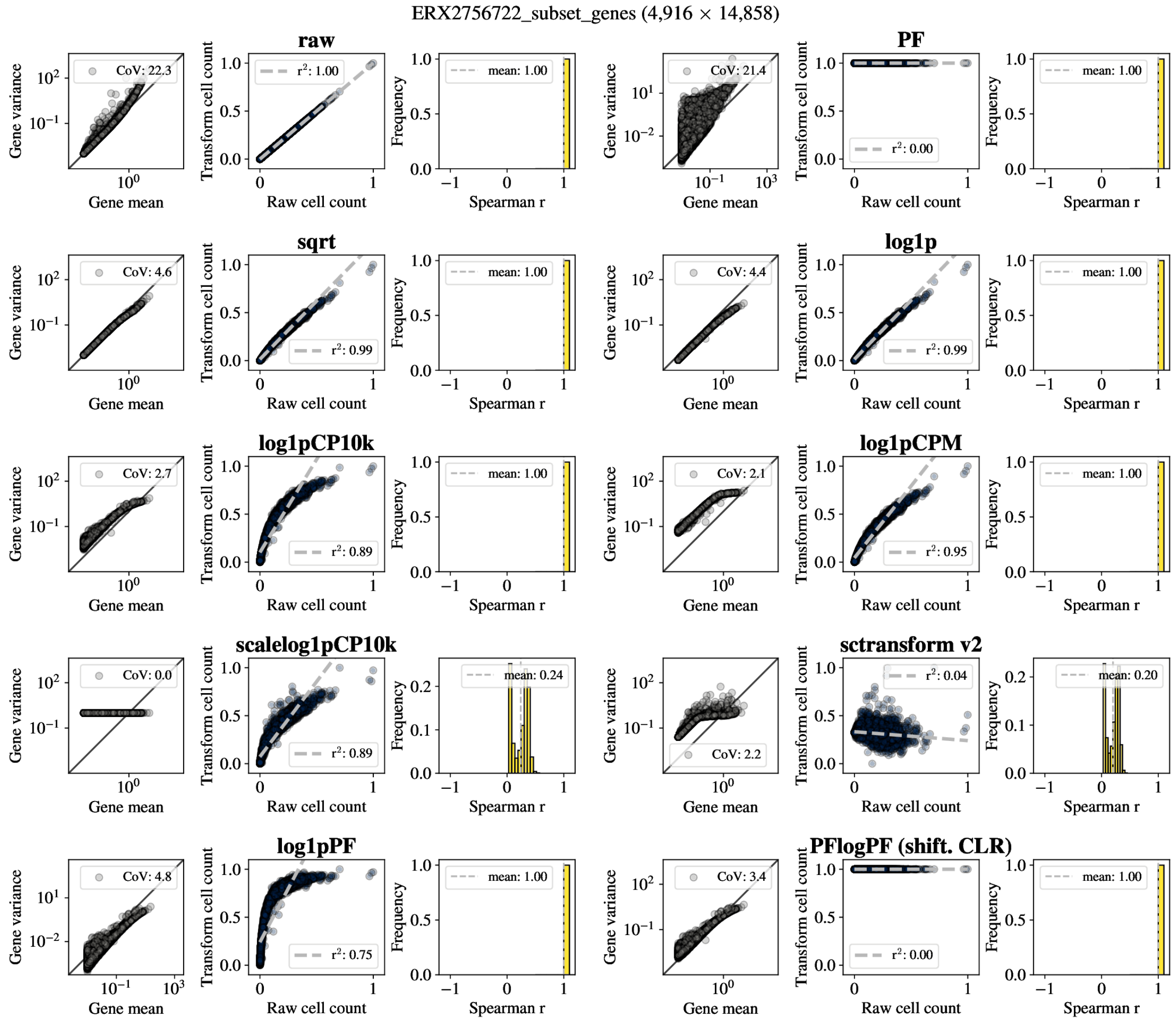


Supplementary Figure 1.123. **ERX2756722**. Per-method comparison of variance stabilization (left), depth normalization (middle), and monotonicity (right) for each normalization on dataset ERX2756722.

| **ERX2756723** | |
|---|---|
| cells | 7,564 |
| genes | 60,623 |
| nonzero entries | 15,956,527 |
| density | 0.0348 |
| total counts | 47,350,861 |
| mean counts / cell | 6,260 |
| mean counts / gene | 781 |
| min counts / cell | 383 |
| max counts / cell | 40,359 |
| overdispersion | 2.29 |

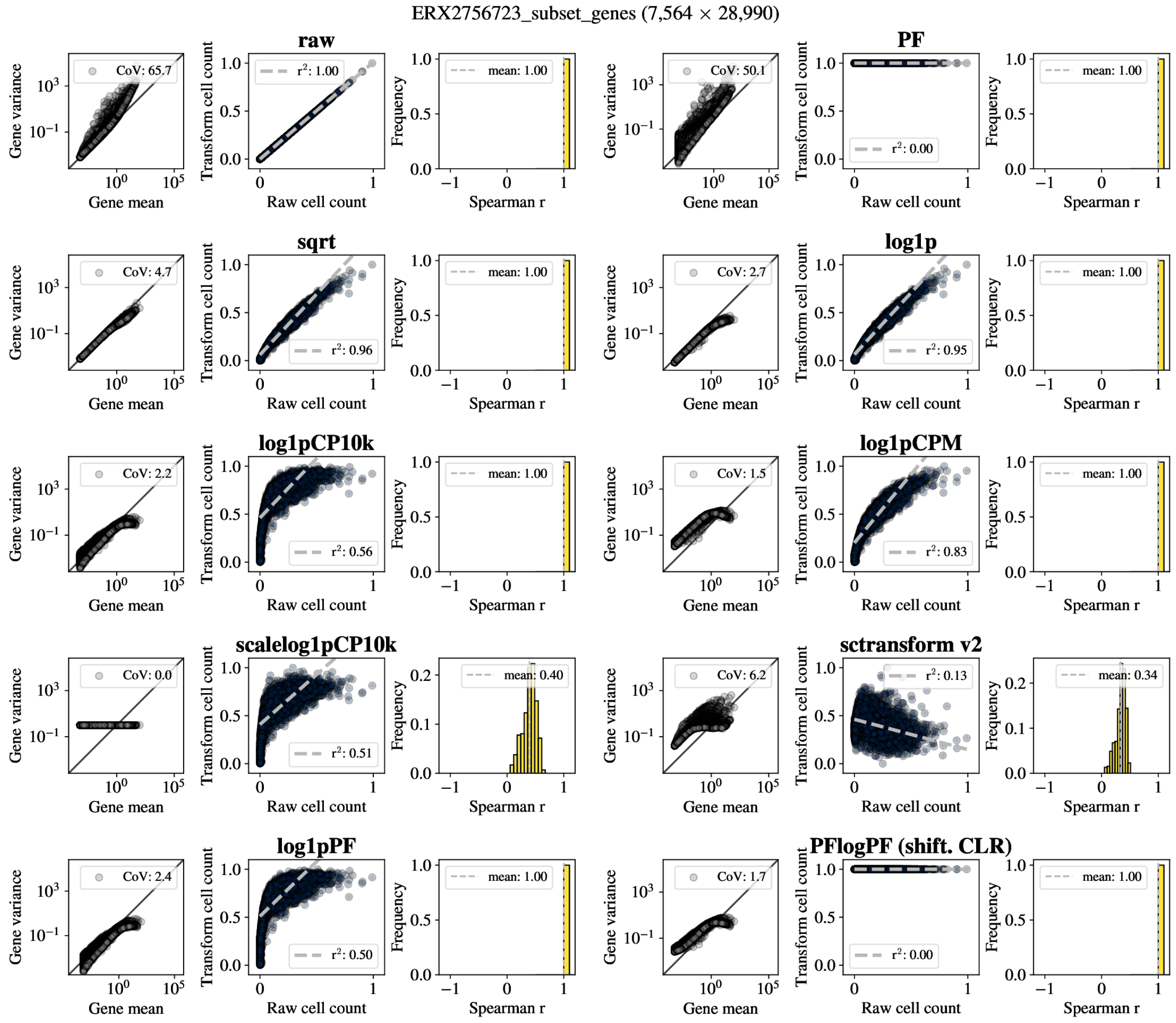


Supplementary Figure 1.124. **ERX2756723**. Per-method comparison of variance stabilization (left), depth normalization (middle), and monotonicity (right) for each normalization on dataset ERX2756723.

| ERX2756724 | |
|---|---|
| cells | 6,415 |
| genes | 60,623 |
| nonzero entries | 975,696 |
| density | 0.00251 |
| total counts | 1,397,798 |
| mean counts / cell | 218 |
| mean counts / gene | 23 |
| min counts / cell | 87 |
| max counts / cell | 6,014 |
| overdispersion | 0.546 |

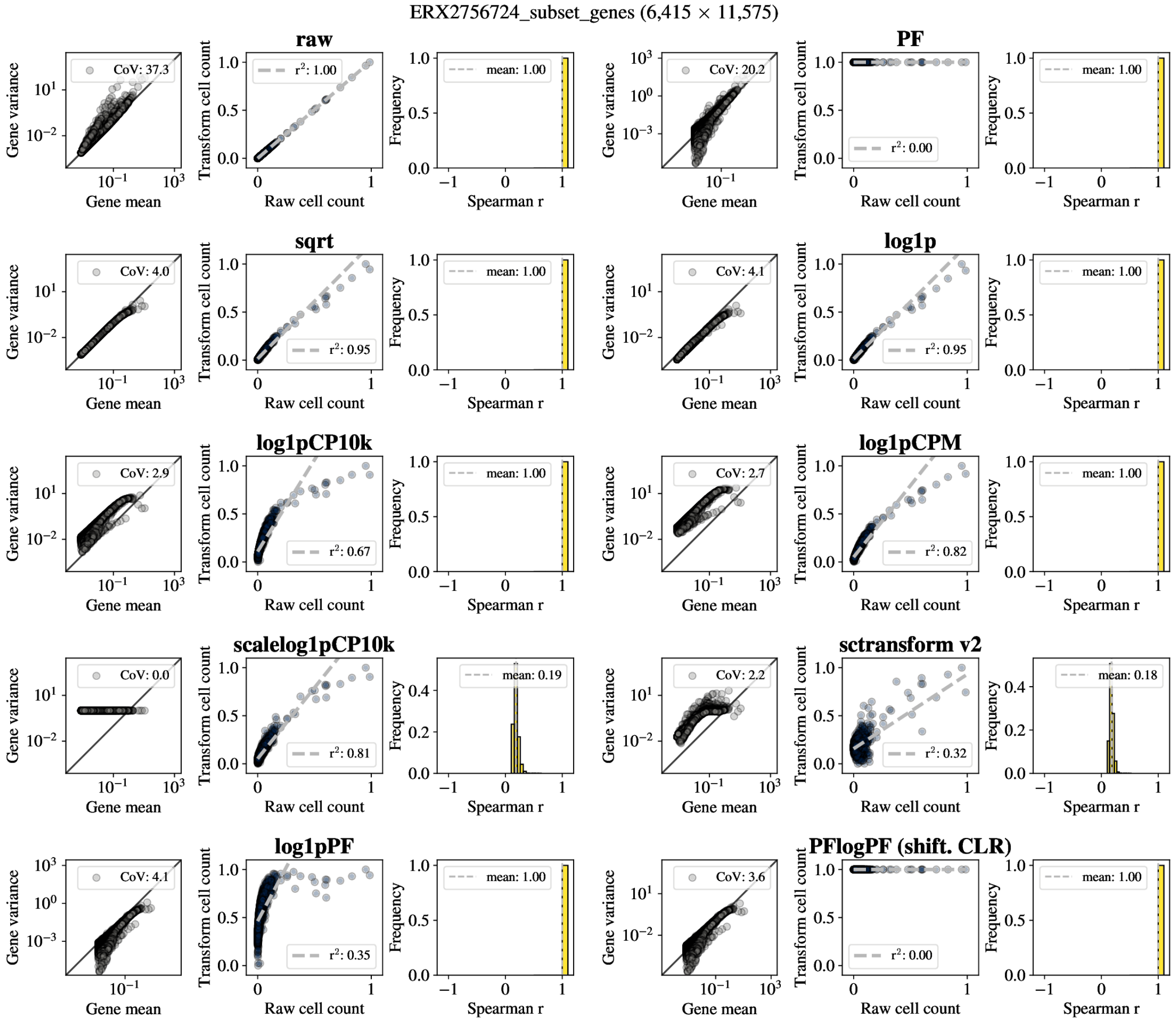


Supplementary Figure 1.125. **ERX2756724**. Per-method comparison of variance stabilization (left), depth normalization (middle), and monotonicity (right) for each normalization on dataset ERX2756724.

| ERX2756725 | |
|---|---|
| cells | 1,754 |
| genes | 60,623 |
| nonzero entries | 66,238 |
| density | 0.000623 |
| total counts | 75,095 |
| mean counts / cell | 43 |
| mean counts / gene | 1 |
| min counts / cell | 22 |
| max counts / cell | 322 |
| overdispersion | 0.396 |

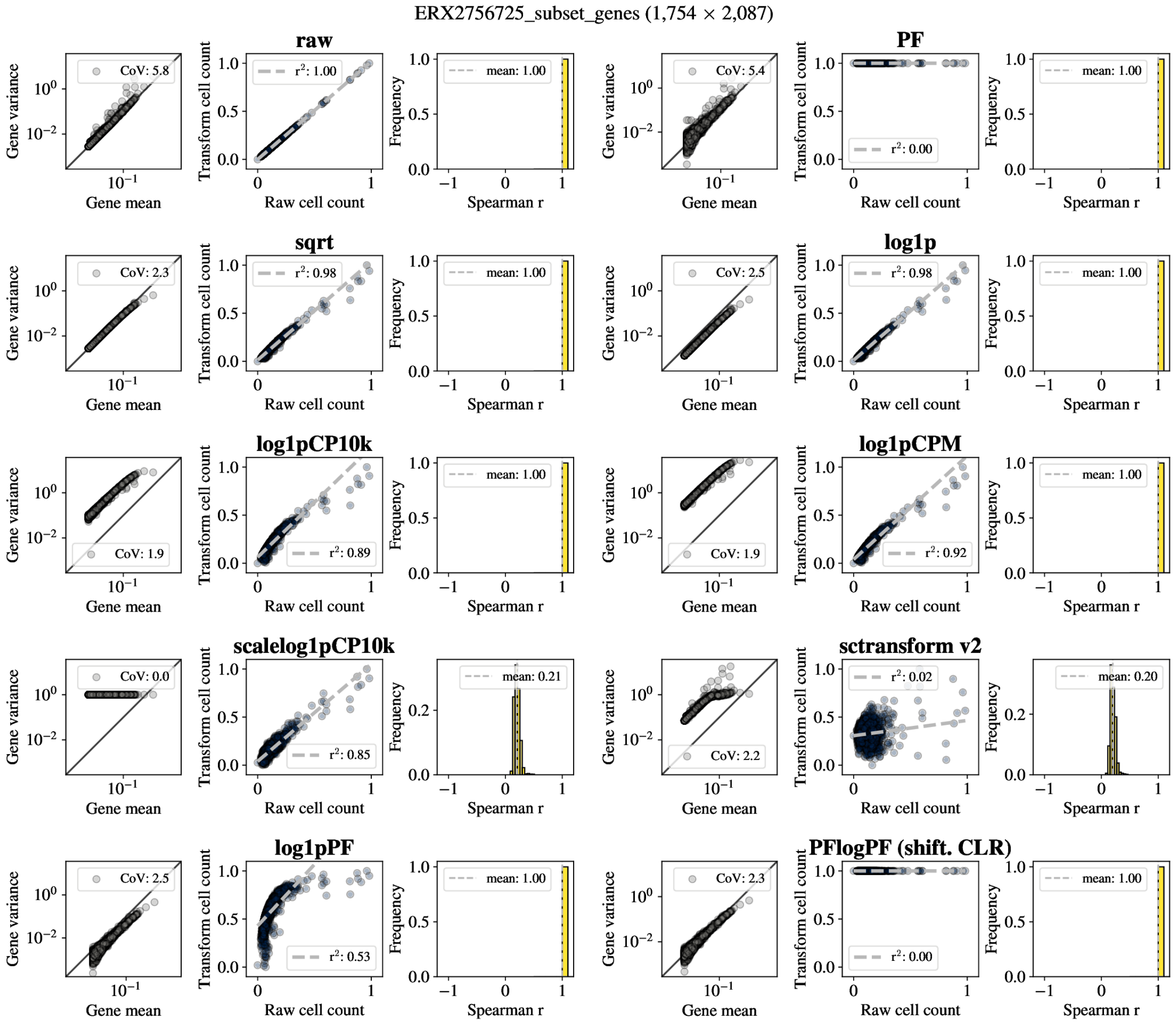


Supplementary Figure 1.126. **ERX2756725**. Per-method comparison of variance stabilization (left), depth normalization (middle), and monotonicity (right) for each normalization on dataset ERX2756725.

| ERX2756726 | |
|---|---|
| cells | 2,036 |
| genes | 60,623 |
| nonzero entries | 3,749,344 |
| density | 0.0304 |
| total counts | 11,849,392 |
| mean counts / cell | 5,820 |
| mean counts / gene | 195 |
| min counts / cell | 1,575 |
| max counts / cell | 25,795 |
| overdispersion | 2.72 |

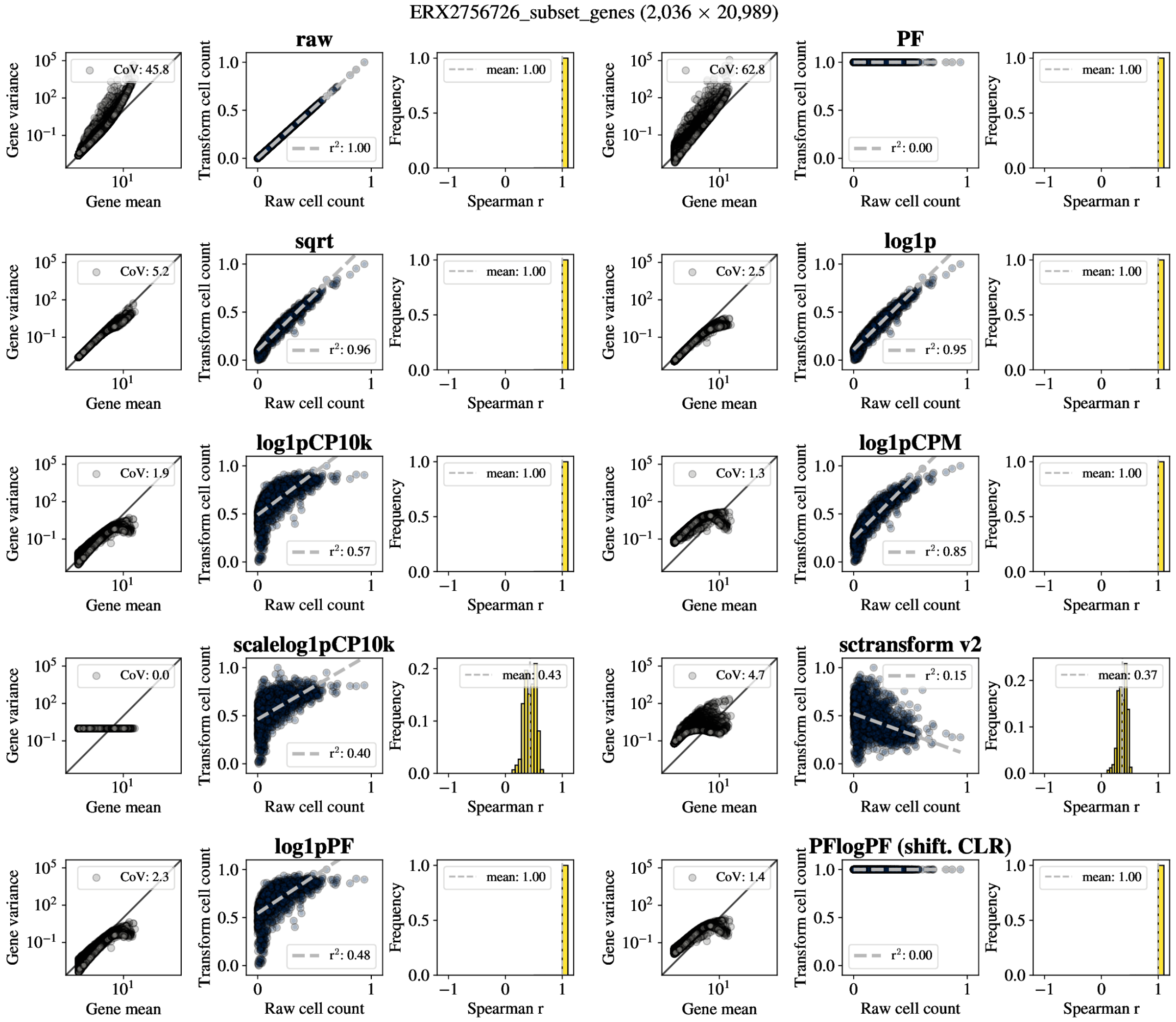


Supplementary Figure 1.127. **ERX2756726**. Per-method comparison of variance stabilization (left), depth normalization (middle), and monotonicity (right) for each normalization on dataset ERX2756726.

| ERX2756727 | |
|---|---|
| cells | 7,125 |
| genes | 60,623 |
| nonzero entries | 10,194,932 |
| density | 0.0236 |
| total counts | 50,617,545 |
| mean counts / cell | 7,104 |
| mean counts / gene | 835 |
| min counts / cell | 60 |
| max counts / cell | 87,686 |
| overdispersion | 9.02 |

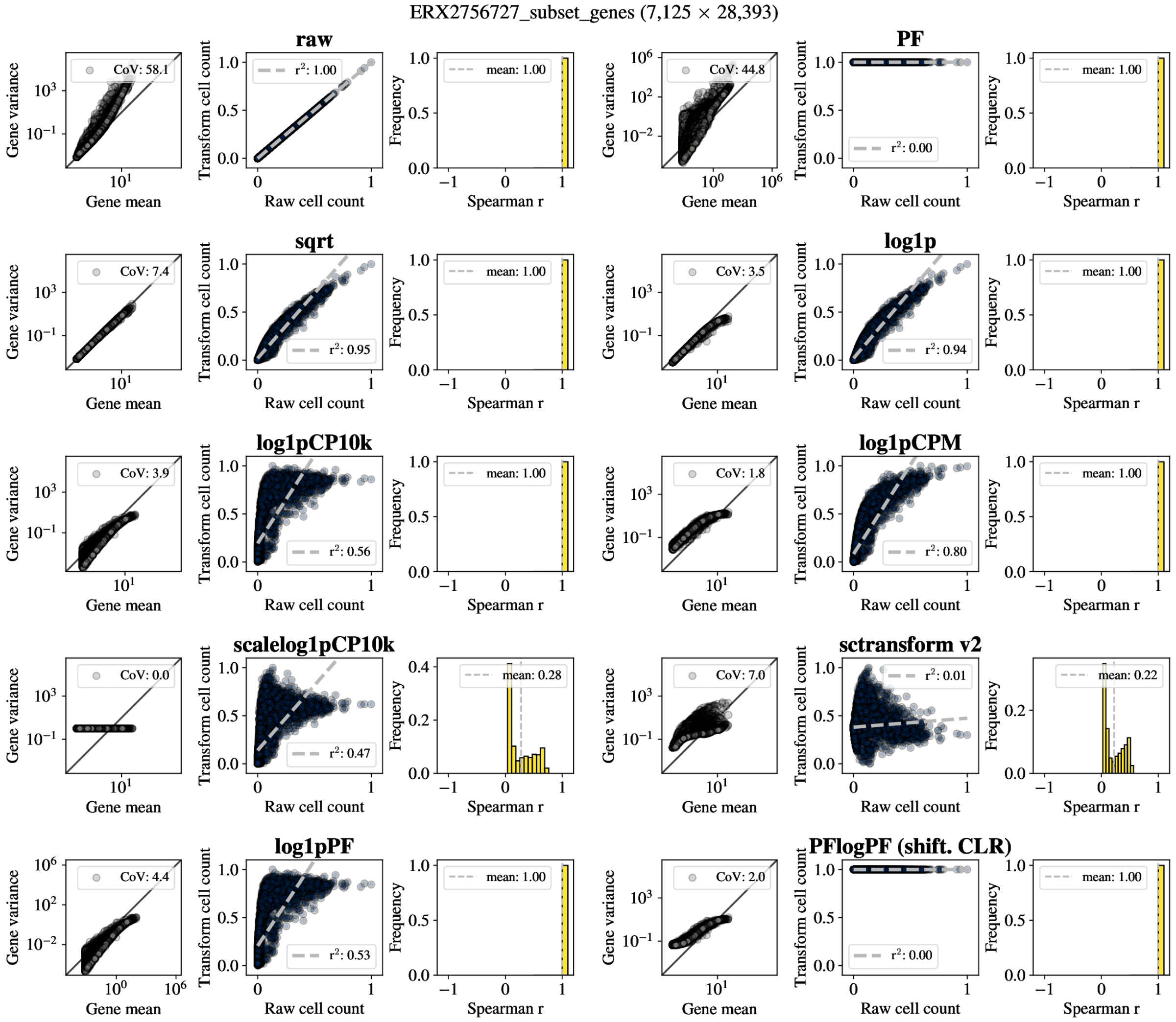


Supplementary Figure 1.128. **ERX2756727**. Per-method comparison of variance stabilization (left), depth normalization (middle), and monotonicity (right) for each normalization on dataset ERX2756727.

| ERX2756728 | |
|---|---|
| cells | 4,529 |
| genes | 60,623 |
| nonzero entries | 4,727,813 |
| density | 0.0172 |
| total counts | 14,649,624 |
| mean counts / cell | 3,235 |
| mean counts / gene | 242 |
| min counts / cell | 1,204 |
| max counts / cell | 23,124 |
| overdispersion | 0.131 |

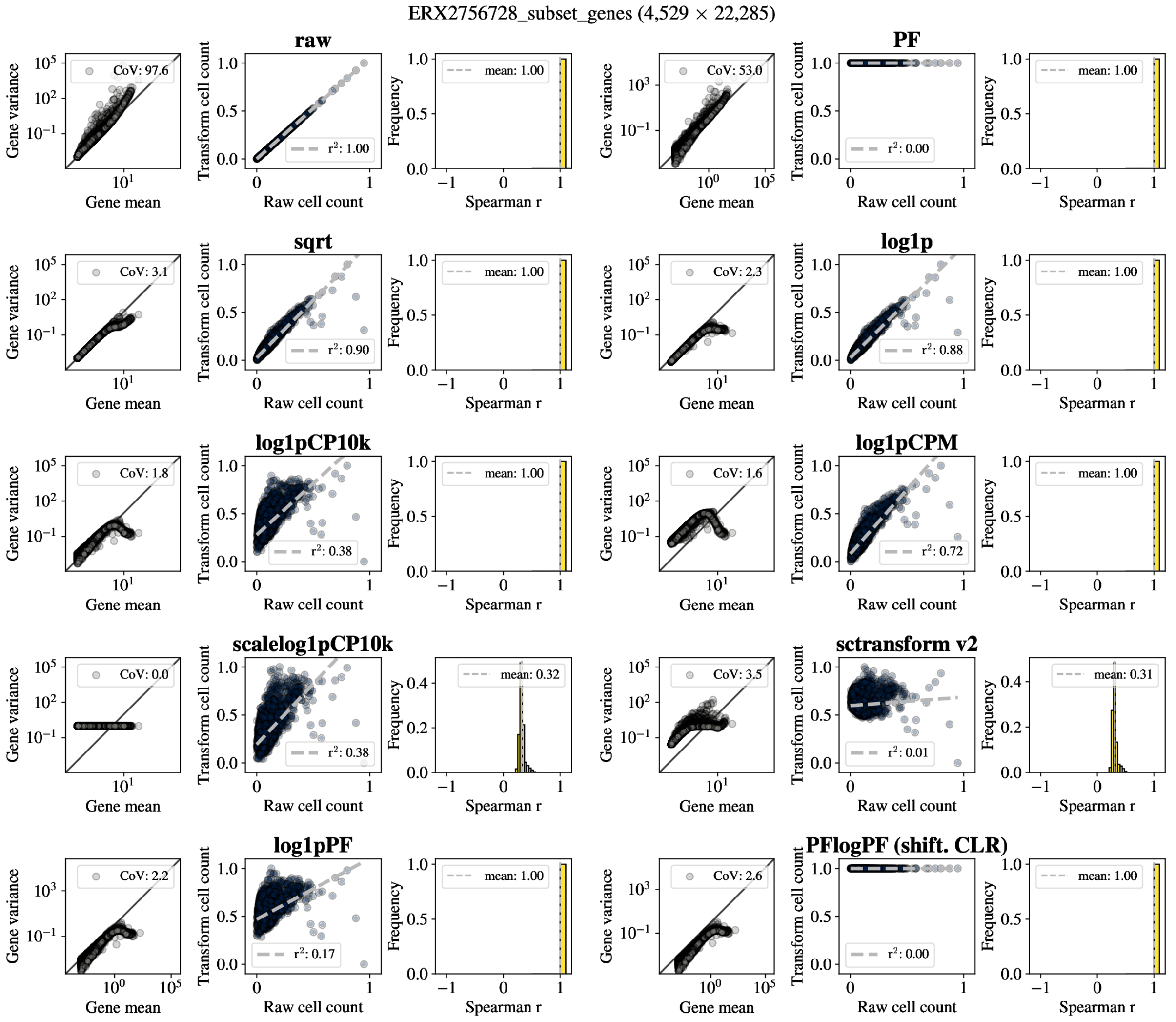


Supplementary Figure 1.129. **ERX2756728**. Per-method comparison of variance stabilization (left), depth normalization (middle), and monotonicity (right) for each normalization on dataset ERX2756728.

| ERX2756729 | |
|---|---|
| cells | 73,299 |
| genes | 60,623 |
| nonzero entries | 7,464,264 |
| density | 0.00168 |
| total counts | 20,091,476 |
| mean counts / cell | 274 |
| mean counts / gene | 331 |
| min counts / cell | 12 |
| max counts / cell | 52,610 |
| overdispersion | 70 |

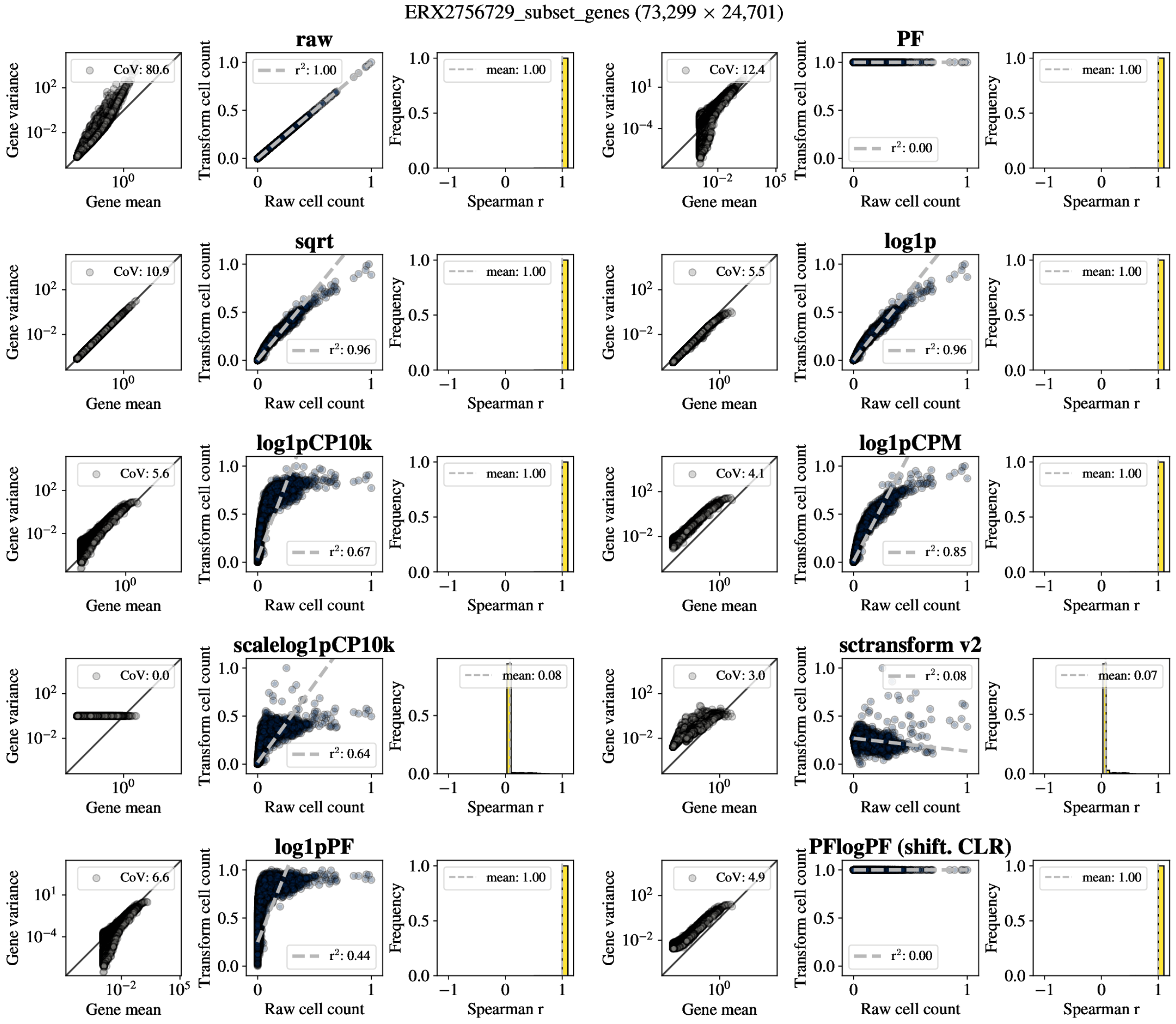


Supplementary Figure 1.130. **ERX2756729**. Per-method comparison of variance stabilization (left), depth normalization (middle), and monotonicity (right) for each normalization on dataset ERX2756729.

| **ERX2756730** | |
|---|---|
| cells | 4,249 |
| genes | 60,623 |
| nonzero entries | 9,929,254 |
| density | 0.0385 |
| total counts | 50,725,460 |
| mean counts / cell | 11,938 |
| mean counts / gene | 837 |
| min counts / cell | 92 |
| max counts / cell | 99,968 |
| overdispersion | 1.06 |

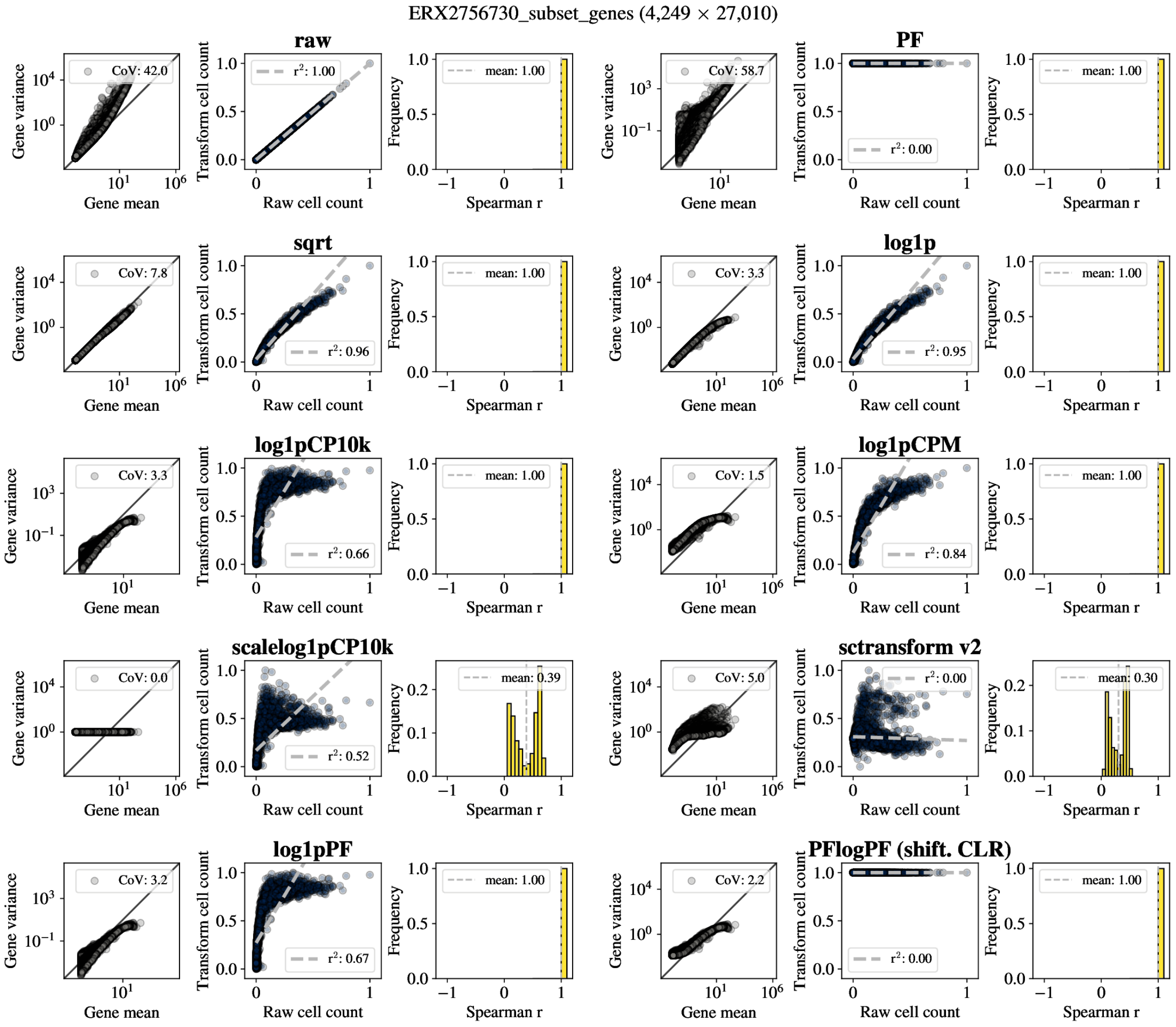


Supplementary Figure 1.131. **ERX2756730**. Per-method comparison of variance stabilization (left), depth normalization (middle), and monotonicity (right) for each normalization on dataset ERX2756730.

| **ERX2756731** | |
|---|---|
| cells | 77,083 |
| genes | 60,623 |
| nonzero entries | 20,464,392 |
| density | 0.00438 |
| total counts | 79,070,789 |
| mean counts / cell | 1,026 |
| mean counts / gene | 1,304 |
| min counts / cell | 11 |
| max counts / cell | 94,061 |
| overdispersion | 29.5 |

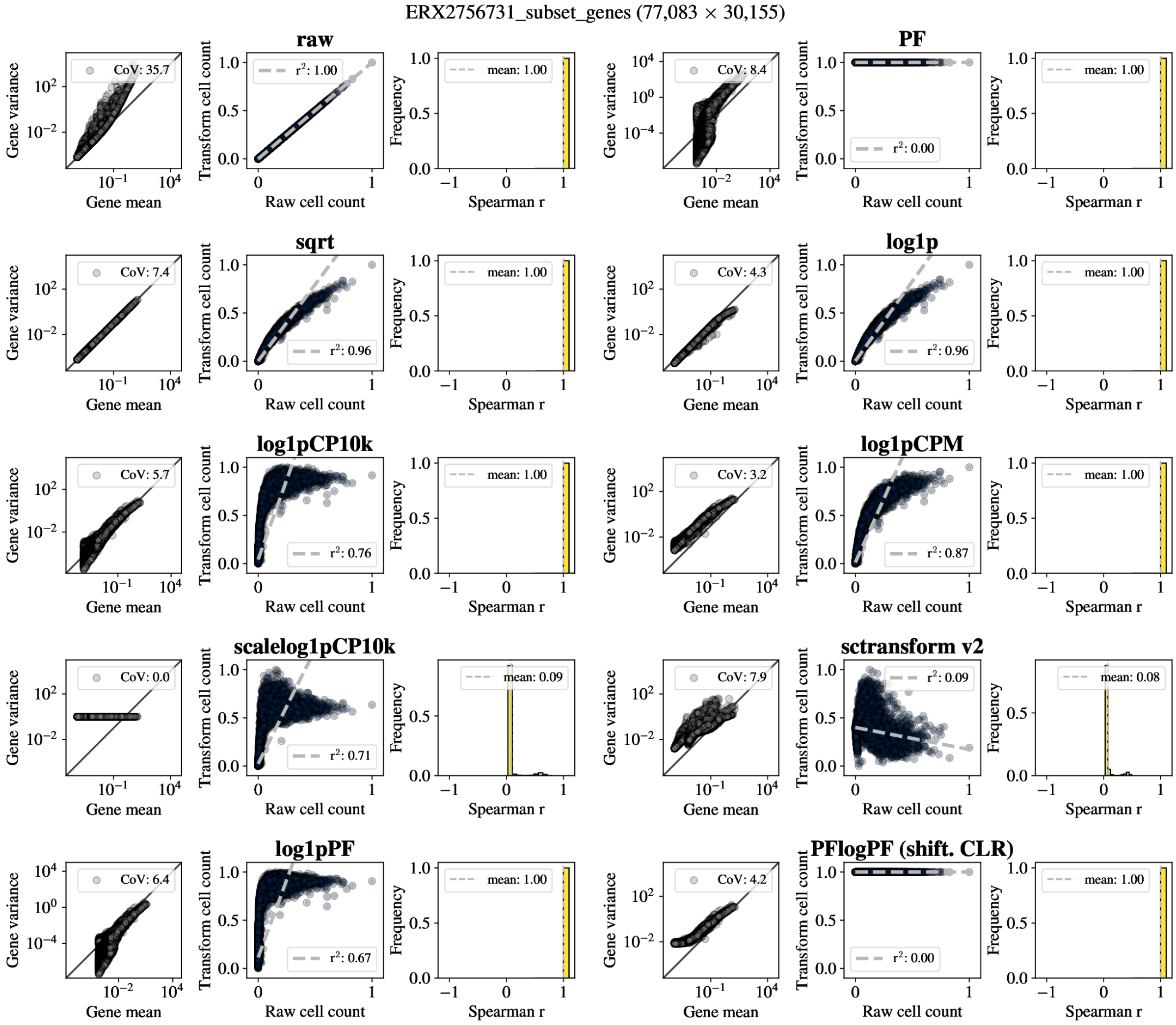


Supplementary Figure 1.132. **ERX2756731**. Per-method comparison of variance stabilization (left), depth normalization (middle), and monotonicity (right) for each normalization on dataset ERX2756731.

| ERX2756732 | |
|---|---|
| cells | 6,045 |
| genes | 60,623 |
| nonzero entries | 17,964,112 |
| density | 0.049 |
| total counts | 76,726,333 |
| mean counts / cell | 12,693 |
| mean counts / gene | 1,266 |
| min counts / cell | 108 |
| max counts / cell | 75,547 |
| overdispersion | 1.68 |

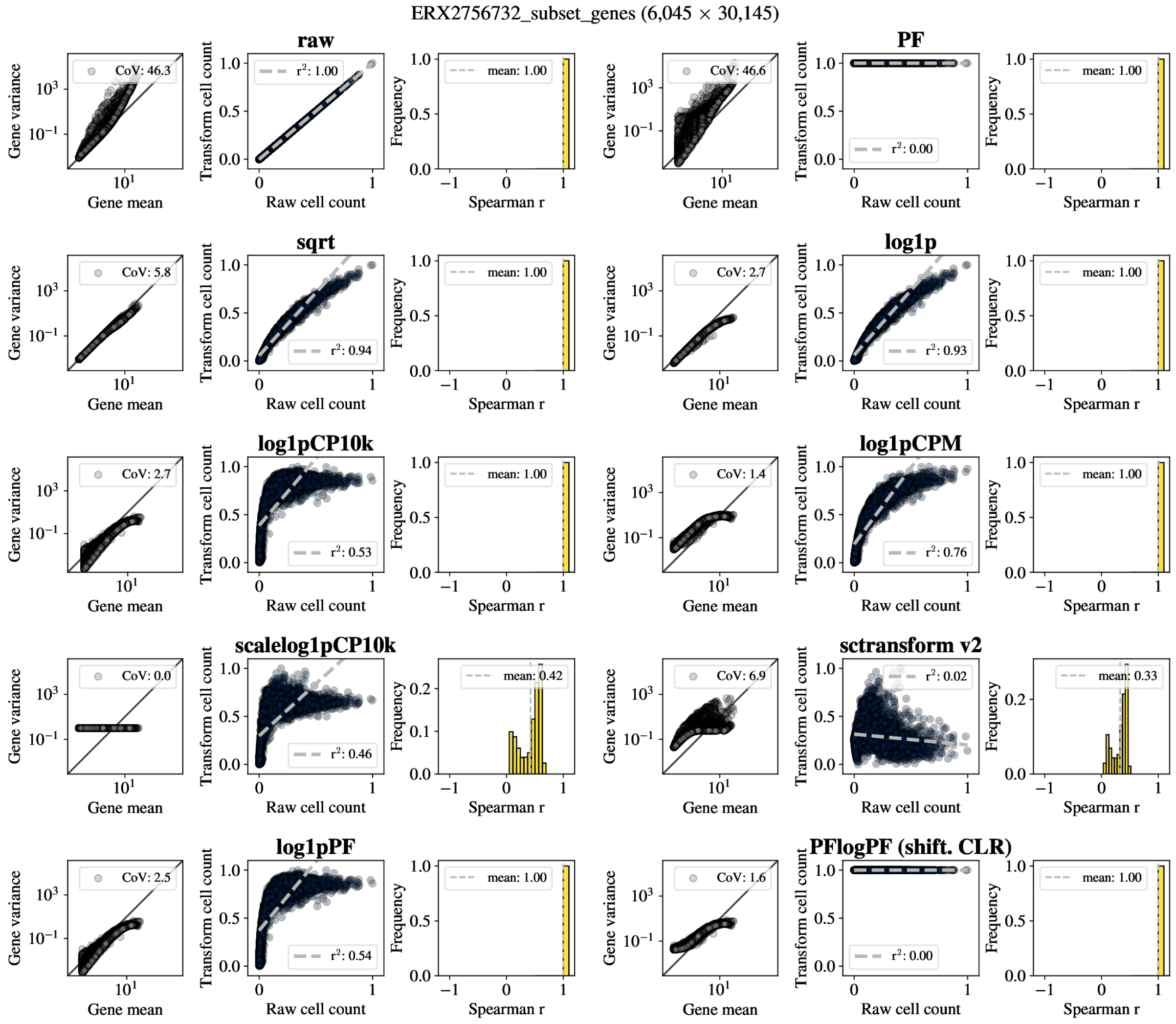


Supplementary Figure 1.133. **ERX2756732**. Per-method comparison of variance stabilization (left), depth normalization (middle), and monotonicity (right) for each normalization on dataset ERX2756732.

| ERX2756733 | |
|---|---|
| cells | 6,259 |
| genes | 60,623 |
| nonzero entries | 20,669,072 |
| density | 0.0545 |
| total counts | 95,176,050 |
| mean counts / cell | 15,206 |
| mean counts / gene | 1,570 |
| min counts / cell | 93 |
| max counts / cell | 84,655 |
| overdispersion | 0.791 |

ERX2756733_subset_genes (6,259 × 29,621)

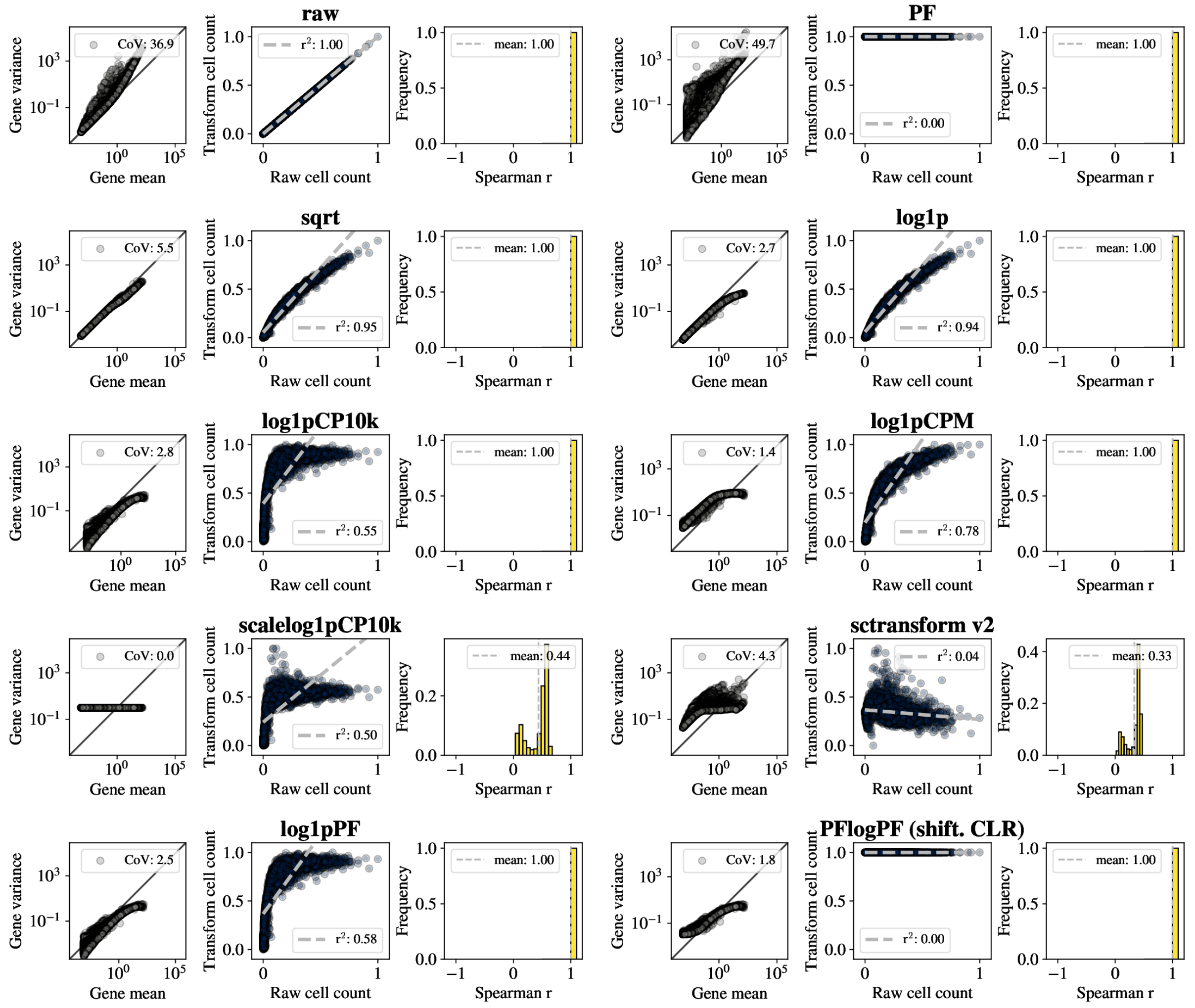


Supplementary Figure 1.134. **ERX2756733**. Per-method comparison of variance stabilization (left), depth normalization (middle), and monotonicity (right) for each normalization on dataset ERX2756733.

| **ERX2756735** | |
|---|---|
| cells | 8,272 |
| genes | 60,623 |
| nonzero entries | 13,906,886 |
| density | 0.0277 |
| total counts | 77,177,978 |
| mean counts / cell | 9,330 |
| mean counts / gene | 1,273 |
| min counts / cell | 113 |
| max counts / cell | 95,532 |
| overdispersion | 3.14 |

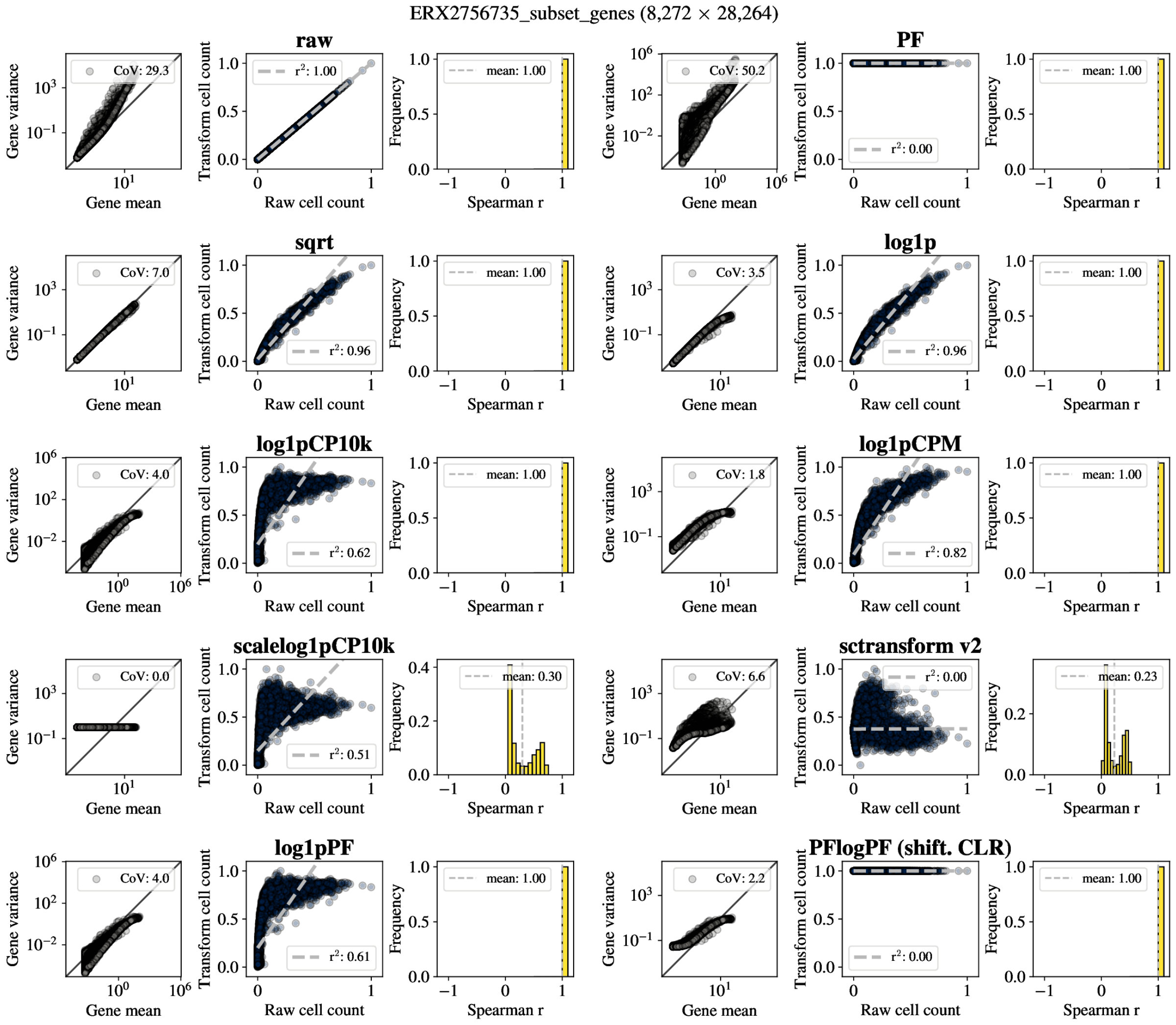


Supplementary Figure 1.135. **ERX2756735**. Per-method comparison of variance stabilization (left), depth normalization (middle), and monotonicity (right) for each normalization on dataset ERX2756735.

| ERX2756736 | |
|---|---|
| cells | 4,996 |
| genes | 60,623 |
| nonzero entries | 1,269,689 |
| density | 0.00419 |
| total counts | 6,945,725 |
| mean counts / cell | 1,390 |
| mean counts / gene | 115 |
| min counts / cell | 477 |
| max counts / cell | 51,274 |
| overdispersion | 0.372 |

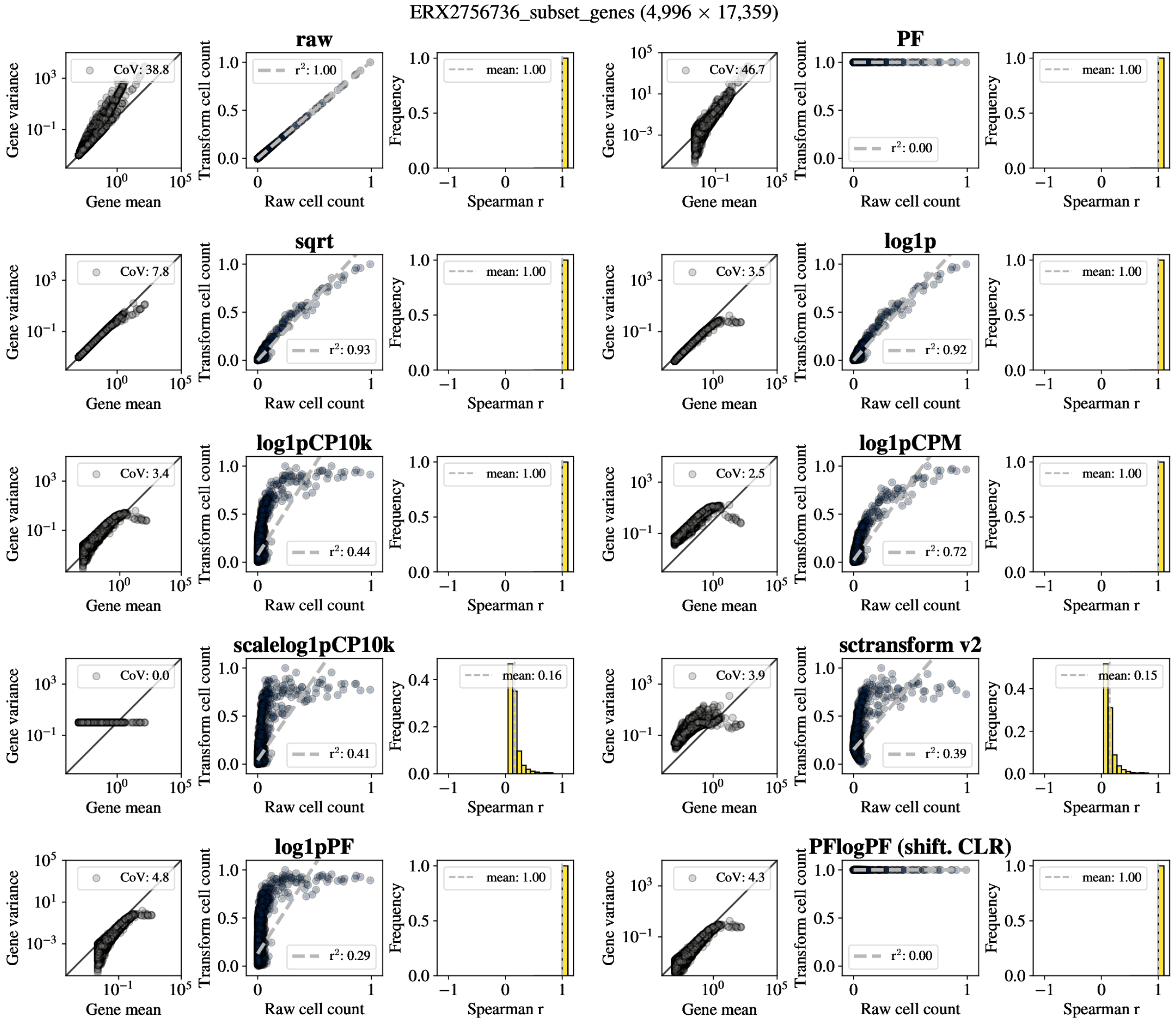


Supplementary Figure 1.136. **ERX2756736**. Per-method comparison of variance stabilization (left), depth normalization (middle), and monotonicity (right) for each normalization on dataset ERX2756736.

| ERX2756737 | |
|---|---|
| cells | 2,996 |
| genes | 60,623 |
| nonzero entries | 1,435,376 |
| density | 0.0079 |
| total counts | 5,260,306 |
| mean counts / cell | 1,756 |
| mean counts / gene | 87 |
| min counts / cell | 149 |
| max counts / cell | 17,264 |
| overdispersion | 2.31 |

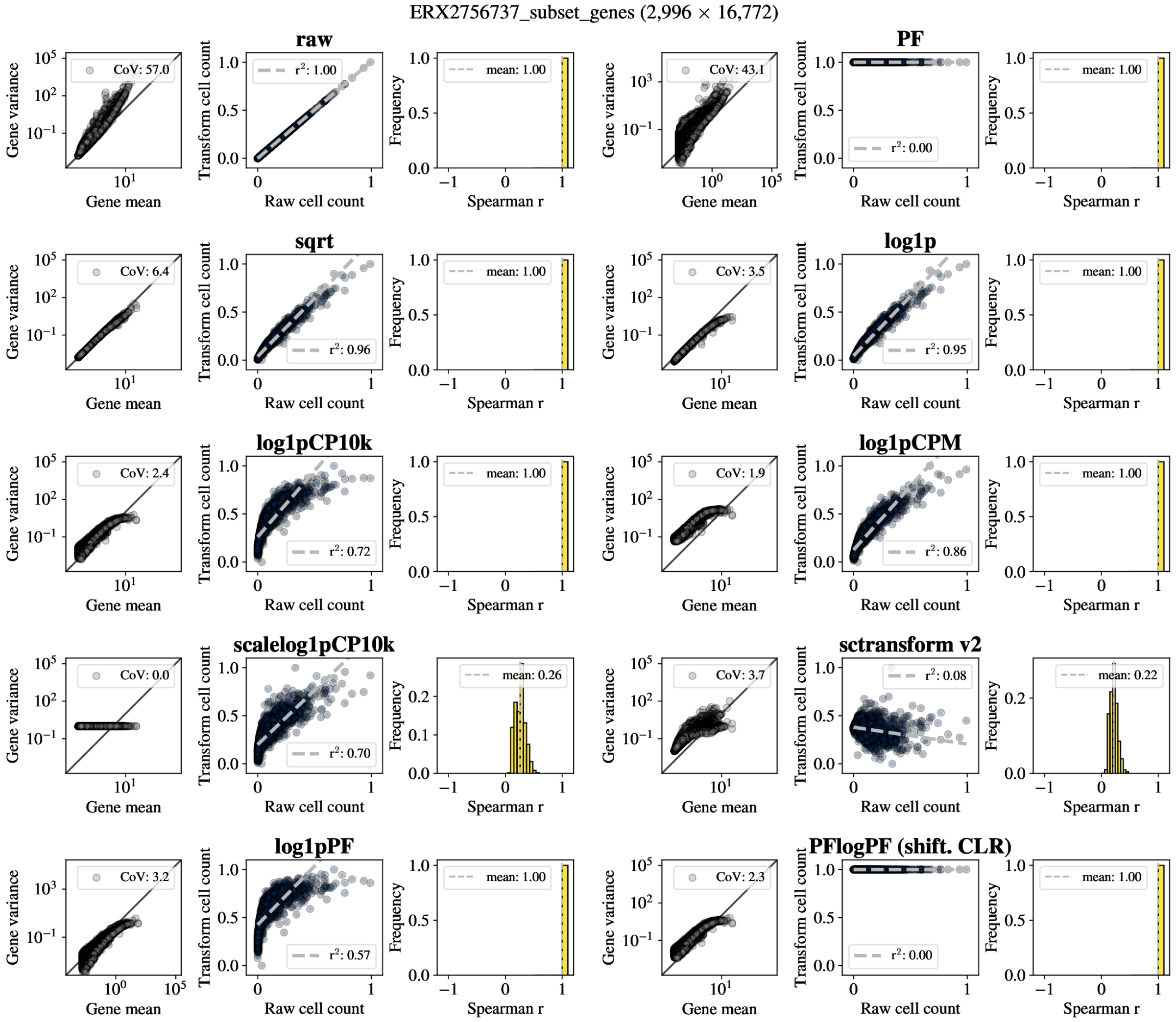


Supplementary Figure 1.137. **ERX2756737**. Per-method comparison of variance stabilization (left), depth normalization (middle), and monotonicity (right) for each normalization on dataset ERX2756737.

| **ERX2756738** | |
|---|---|
| cells | 1,443 |
| genes | 60,623 |
| nonzero entries | 4,715,034 |
| density | 0.0539 |
| total counts | 23,933,943 |
| mean counts / cell | 16,586 |
| mean counts / gene | 395 |
| min counts / cell | 8,684 |
| max counts / cell | 53,784 |
| overdispersion | 0.191 |

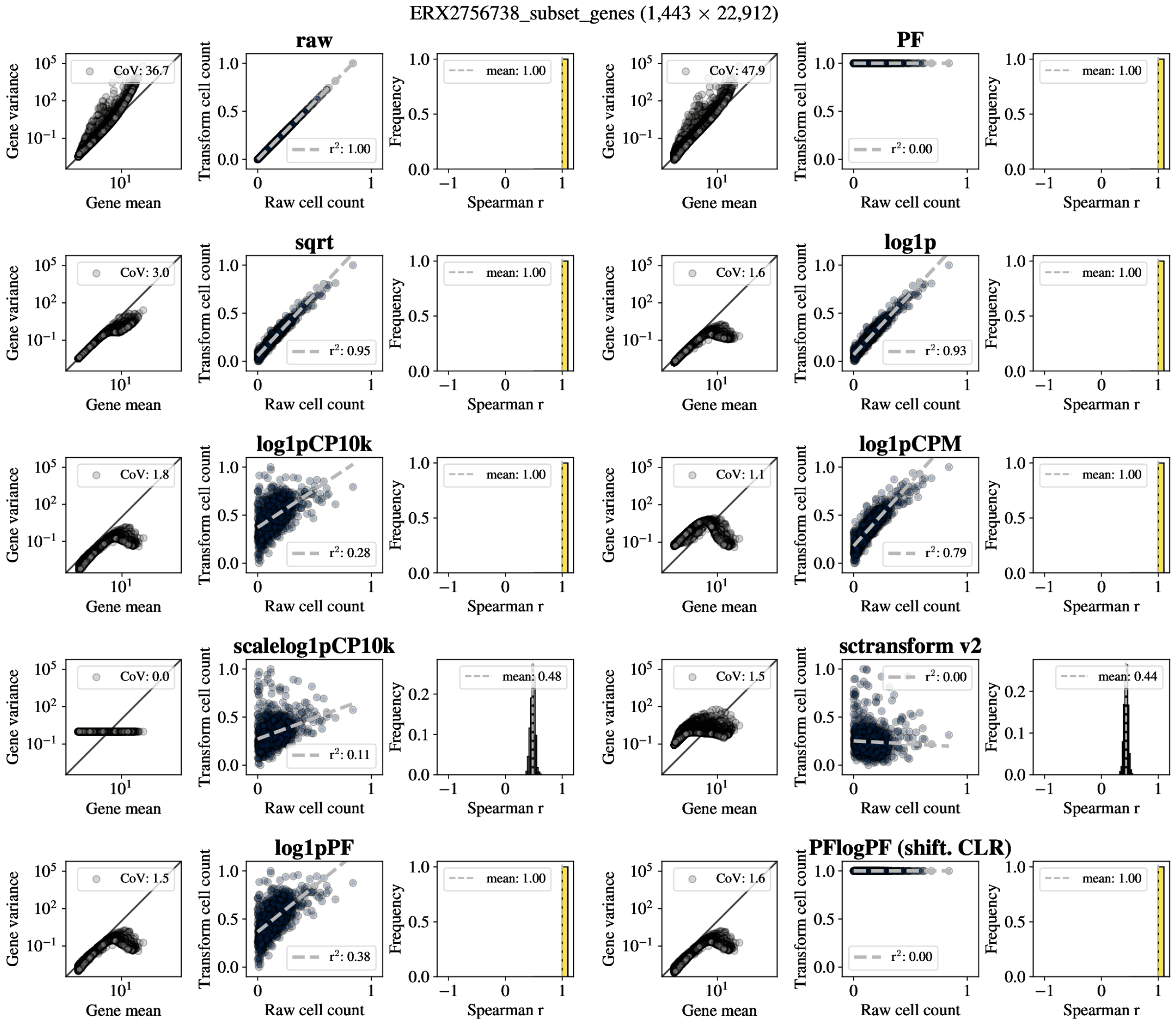


Supplementary Figure 1.138. **ERX2756738**. Per-method comparison of variance stabilization (left), depth normalization (middle), and monotonicity (right) for each normalization on dataset ERX2756738.

| ERX2756739 | |
|---|---|
| cells | 604 |
| genes | 60,623 |
| nonzero entries | 289,936 |
| density | 0.00792 |
| total counts | 1,205,213 |
| mean counts / cell | 1,995 |
| mean counts / gene | 20 |
| min counts / cell | 150 |
| max counts / cell | 11,500 |
| overdispersion | 1.75 |

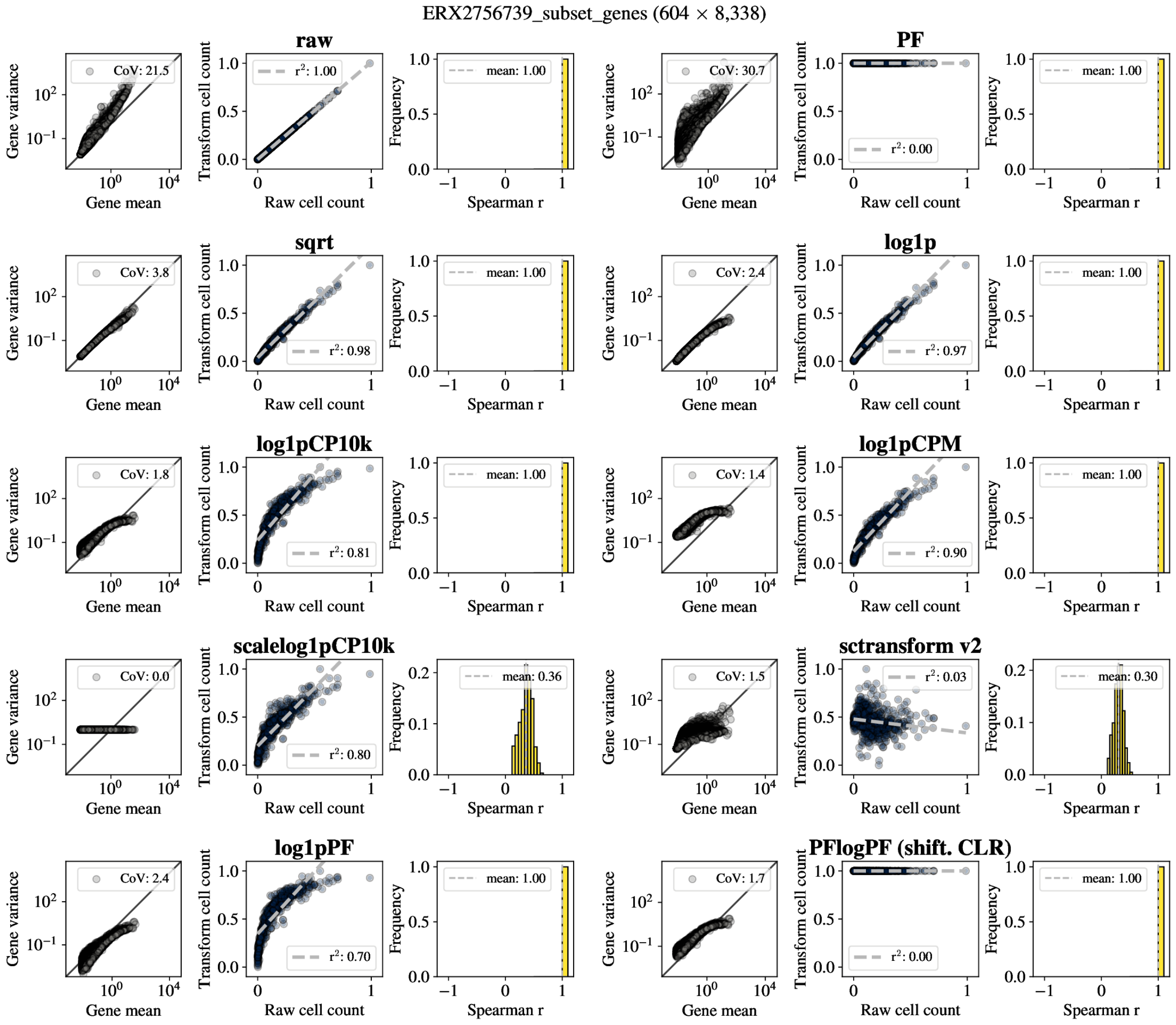


Supplementary Figure 1.139. **ERX2756739**. Per-method comparison of variance stabilization (left), depth normalization (middle), and monotonicity (right) for each normalization on dataset ERX2756739.

| ERX2756740 | |
|---|---|
| cells | 7,265 |
| genes | 60,623 |
| nonzero entries | 12,094,314 |
| density | 0.0275 |
| total counts | 51,481,720 |
| mean counts / cell | 7,086 |
| mean counts / gene | 849 |
| min counts / cell | 68 |
| max counts / cell | 68,161 |
| overdispersion | 3.11 |

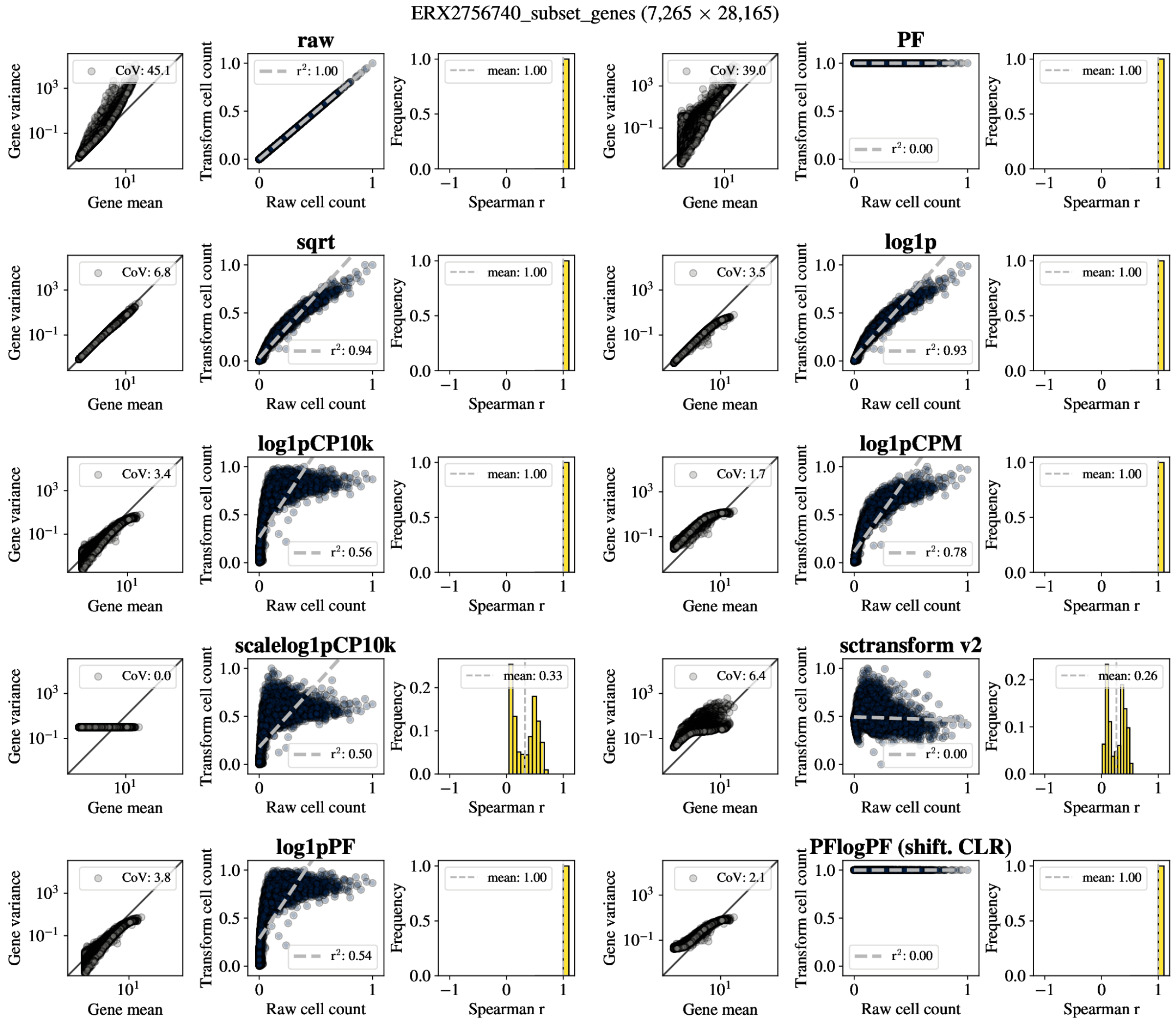


Supplementary Figure 1.140. **ERX2756740**. Per-method comparison of variance stabilization (left), depth normalization (middle), and monotonicity (right) for each normalization on dataset ERX2756740.

| **ERX2756741** | |
|---|---|
| cells | 2,614 |
| genes | 60,623 |
| nonzero entries | 8,993,473 |
| density | 0.0568 |
| total counts | 47,093,775 |
| mean counts / cell | 18,016 |
| mean counts / gene | 777 |
| min counts / cell | 6,801 |
| max counts / cell | 62,584 |
| overdispersion | 1.33 |

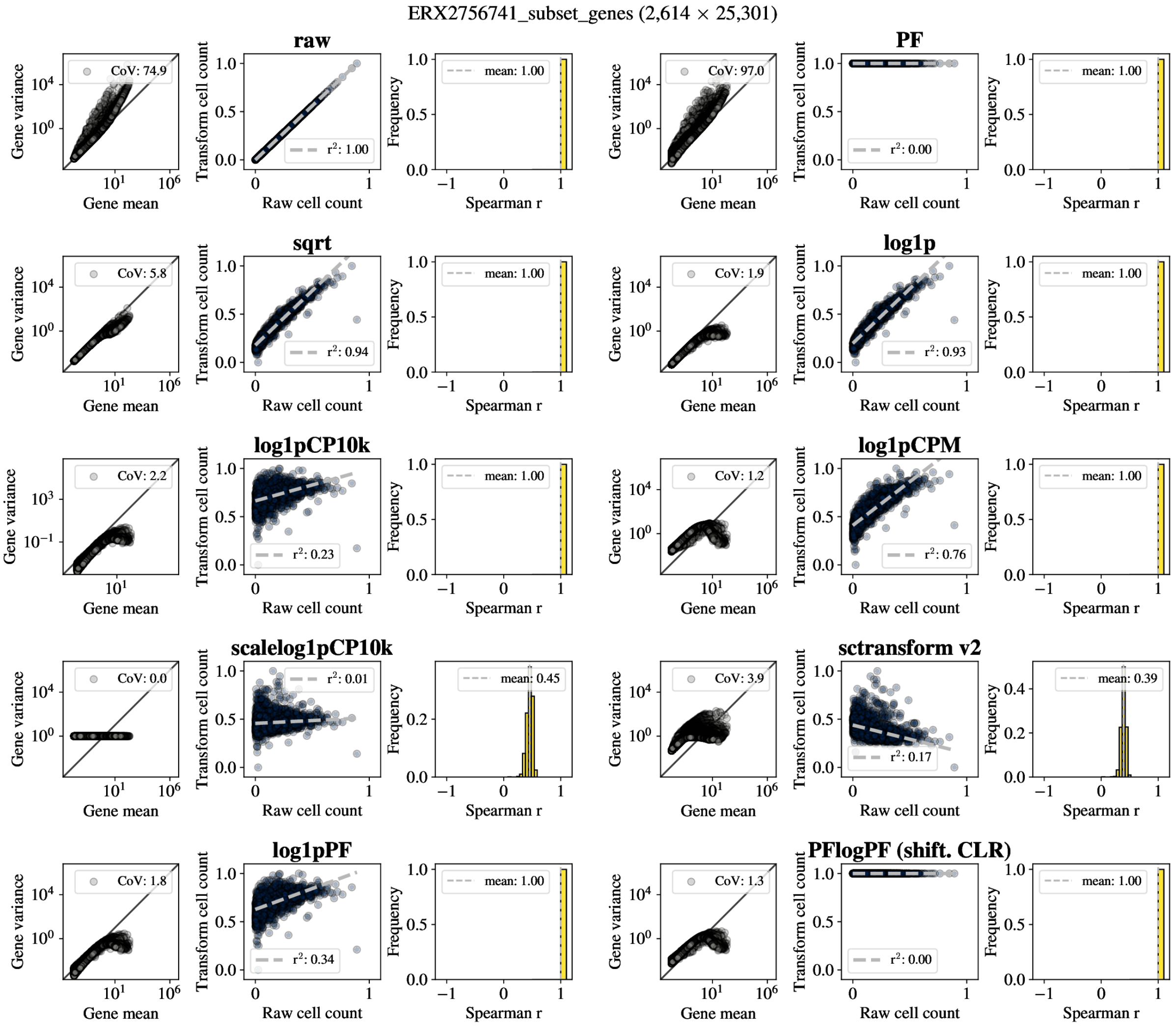


Supplementary Figure 1.141. **ERX2756741**. Per-method comparison of variance stabilization (left), depth normalization (middle), and monotonicity (right) for each normalization on dataset ERX2756741.

| **ERX2756742** | |
|---|---|
| cells | 5,556 |
| genes | 60,623 |
| nonzero entries | 8,739,031 |
| density | 0.0259 |
| total counts | 39,404,510 |
| mean counts / cell | 7,092 |
| mean counts / gene | 650 |
| min counts / cell | 59 |
| max counts / cell | 60,975 |
| overdispersion | 11.6 |

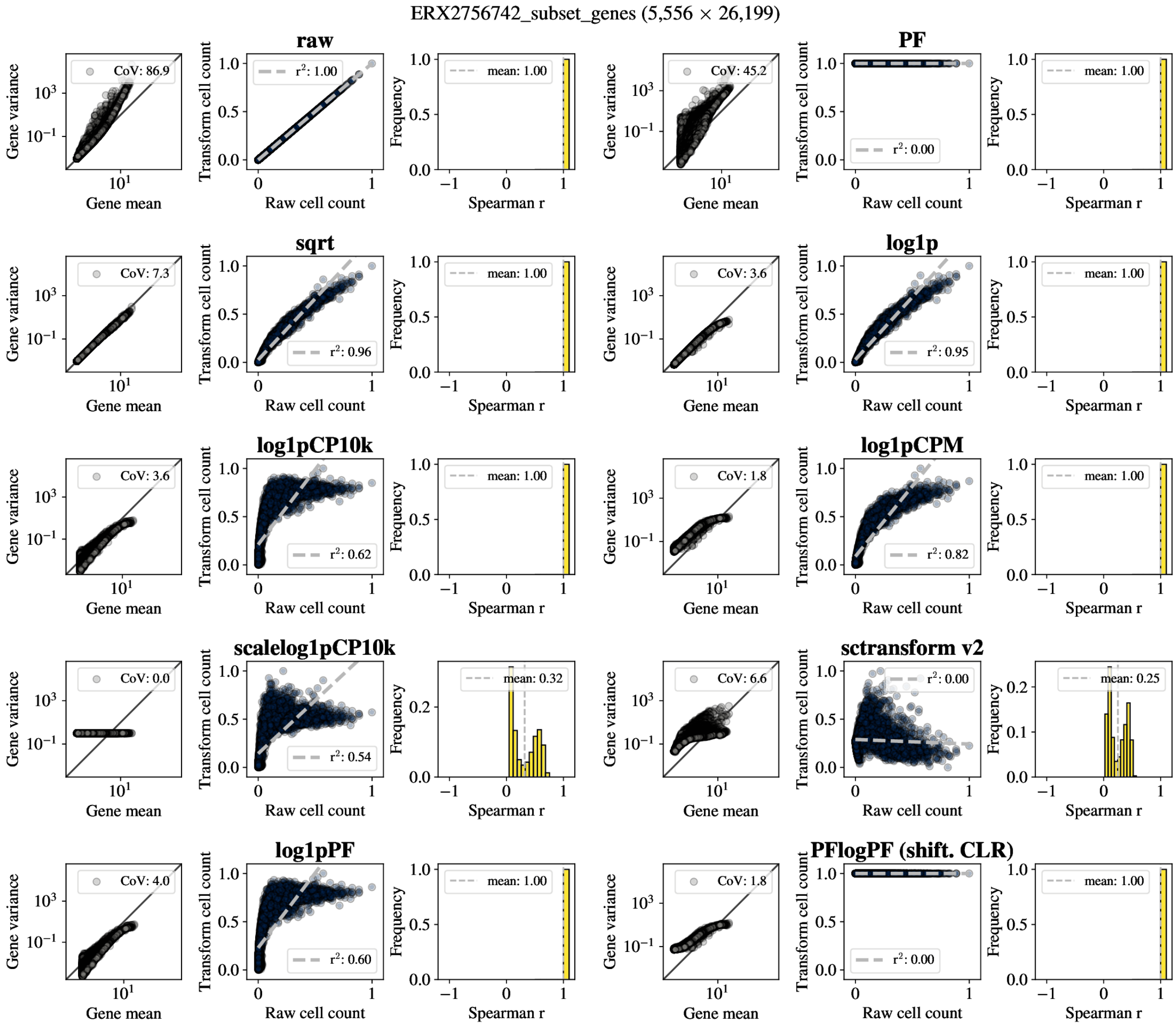


Supplementary Figure 1.142. **ERX2756742**. Per-method comparison of variance stabilization (left), depth normalization (middle), and monotonicity (right) for each normalization on dataset ERX2756742.

| GSM2560245 | |
|---|---|
| cells | 3,679 |
| genes | 60,623 |
| nonzero entries | 1,562,140 |
| density | 0.007 |
| total counts | 5,074,386 |
| mean counts / cell | 1,379 |
| mean counts / gene | 84 |
| min counts / cell | 340 |
| max counts / cell | 9,804 |
| overdispersion | 0.309 |

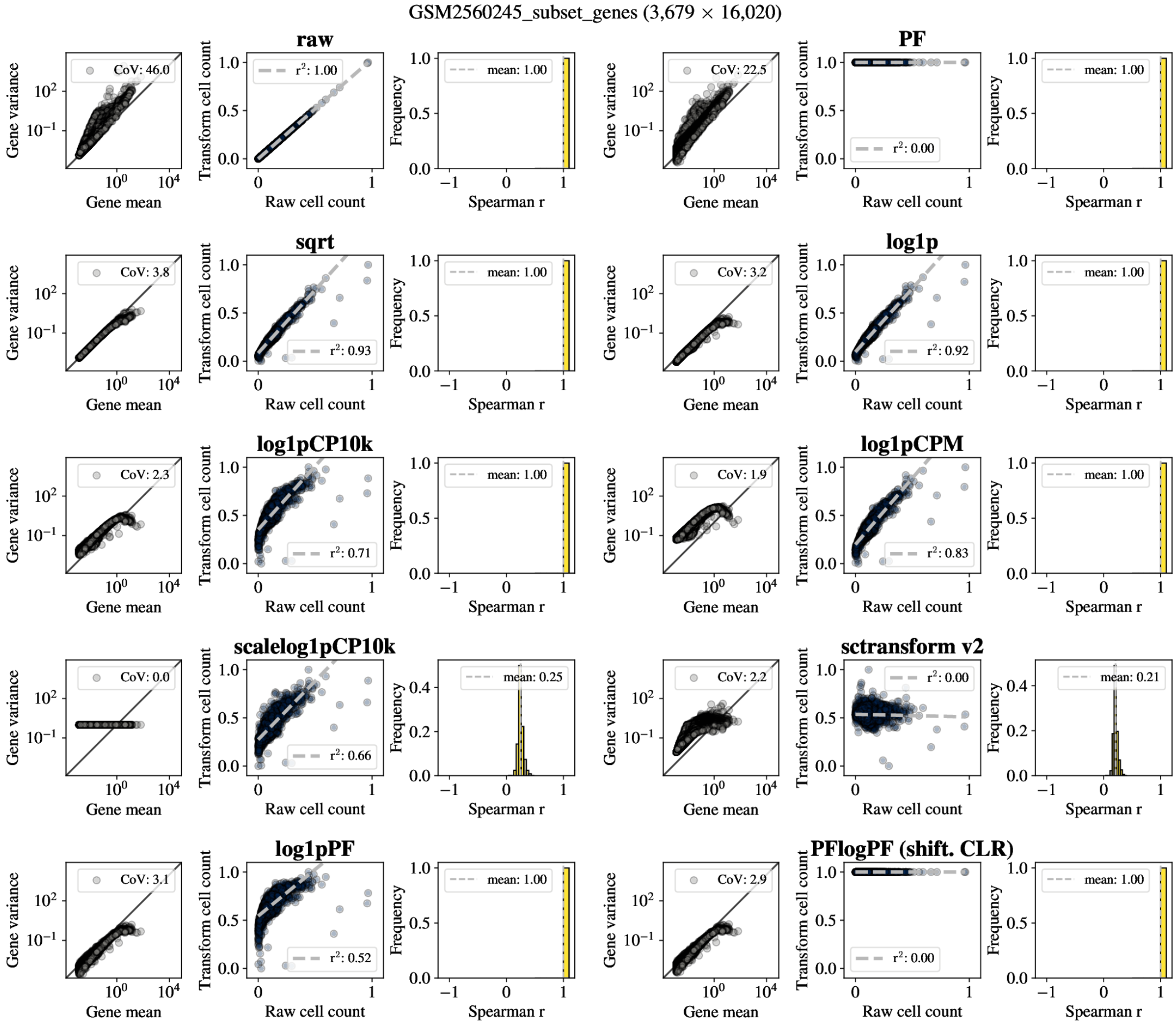


Supplementary Figure 1.143. **GSM2560245**. Per-method comparison of variance stabilization (left), depth normalization (middle), and monotonicity (right) for each normalization on dataset GSM2560245.

| GSM2560248 | |
|---|---|
| cells | 15,707 |
| genes | 60,623 |
| nonzero entries | 9,956,559 |
| density | 0.0105 |
| total counts | 25,288,568 |
| mean counts / cell | 1,610 |
| mean counts / gene | 417 |
| min counts / cell | 244 |
| max counts / cell | 8,713 |
| overdispersion | 0.64 |

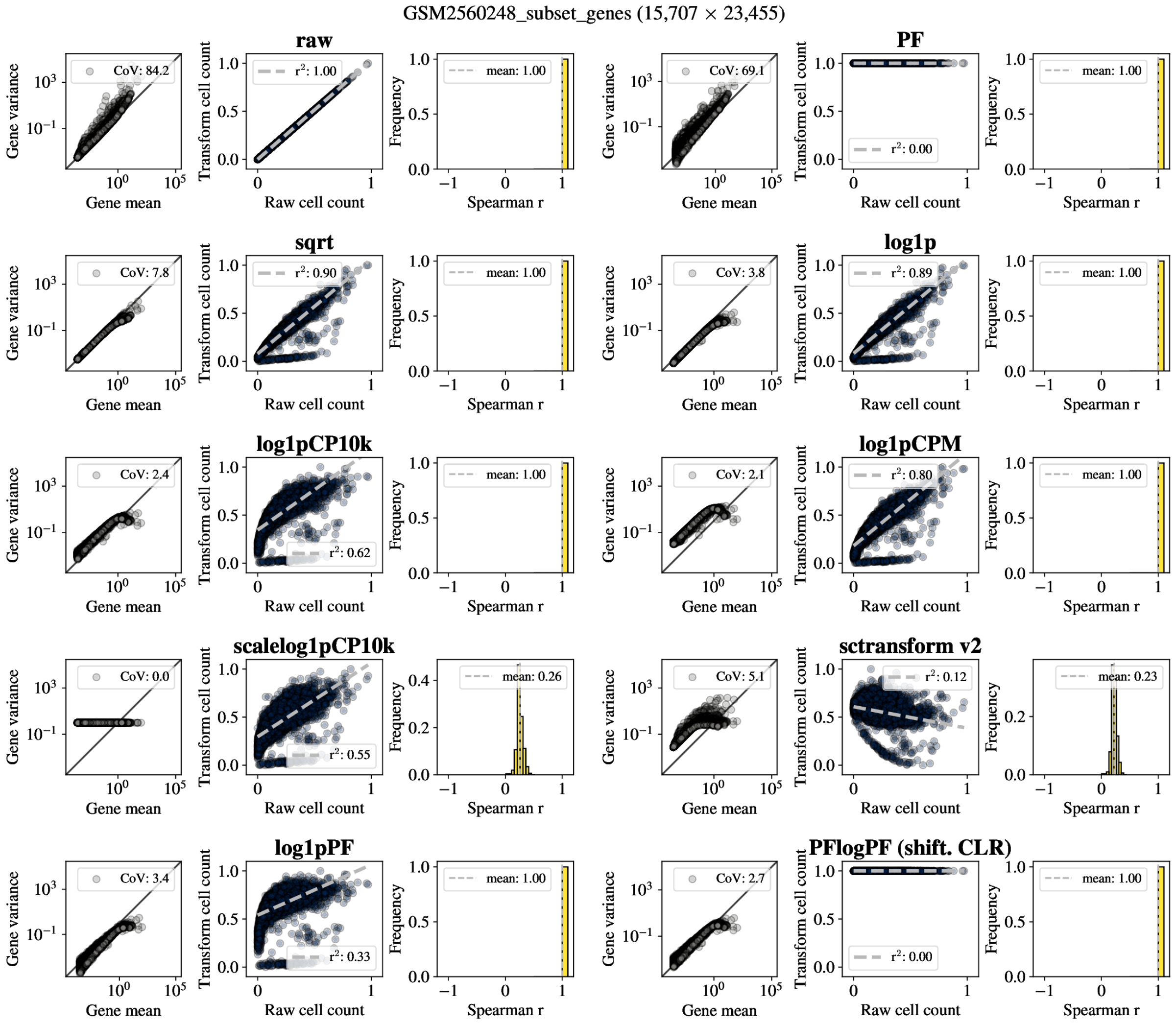


Supplementary Figure 1.144. **GSM2560248**. Per-method comparison of variance stabilization (left), depth normalization (middle), and monotonicity (right) for each normalization on dataset GSM2560248.

| GSM2786157 | |
|---|---|
| cells | 666 |
| genes | 60,623 |
| nonzero entries | 4,056,717 |
| density | 0.1 |
| total counts | 27,585,199 |
| mean counts / cell | 41,419 |
| mean counts / gene | 455 |
| min counts / cell | 13,776 |
| max counts / cell | 131,443 |
| overdispersion | 1.02 |

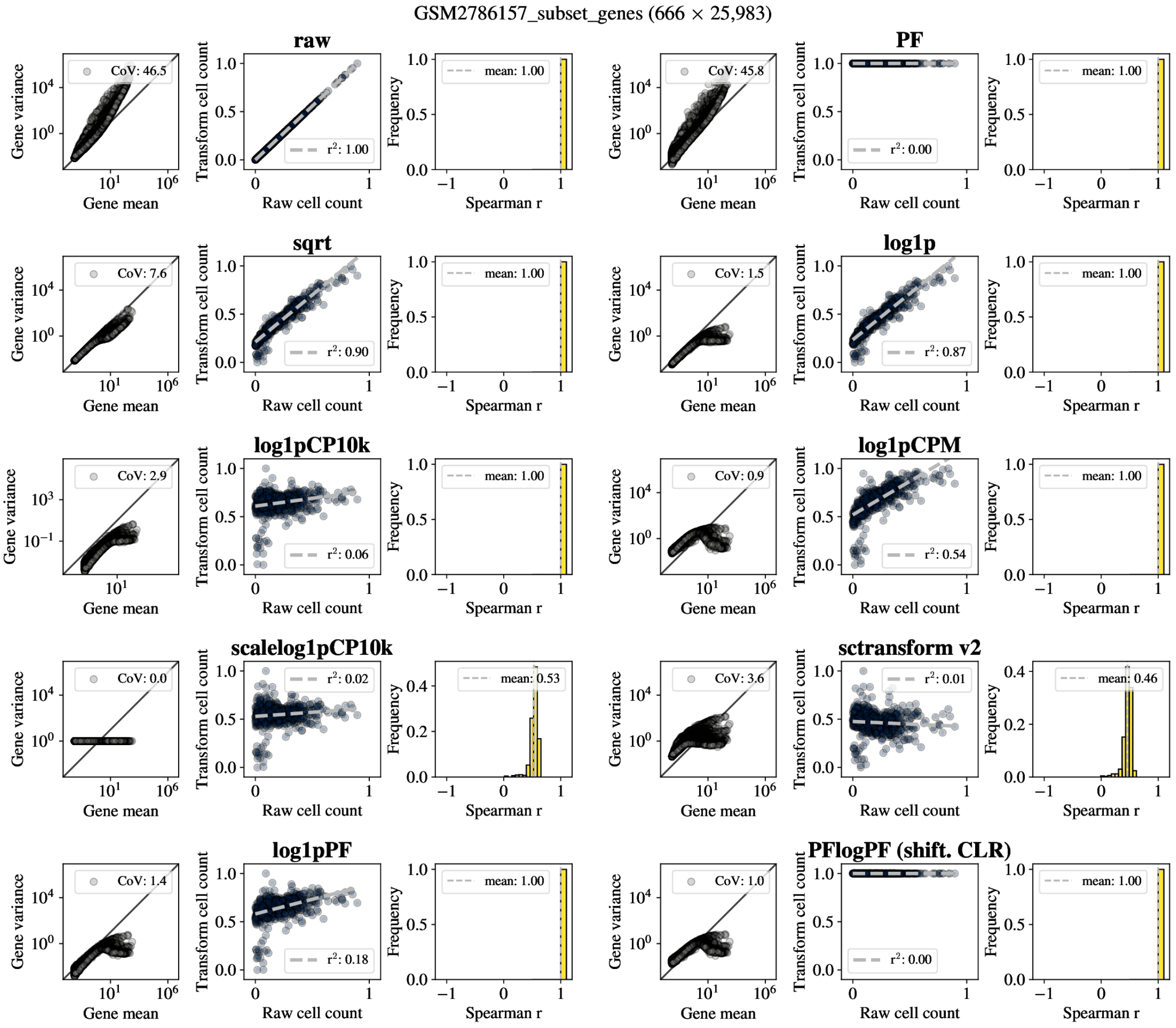


Supplementary Figure 1.145. **GSM2786157**. Per-method comparison of variance stabilization (left), depth normalization (middle), and monotonicity (right) for each normalization on dataset GSM2786157.

| GSM2786158 | |
|---|---|
| cells | 753 |
| genes | 60,623 |
| nonzero entries | 3,640,412 |
| density | 0.0797 |
| total counts | 20,183,218 |
| mean counts / cell | 26,804 |
| mean counts / gene | 333 |
| min counts / cell | 4,526 |
| max counts / cell | 100,990 |
| overdispersion | 0.477 |

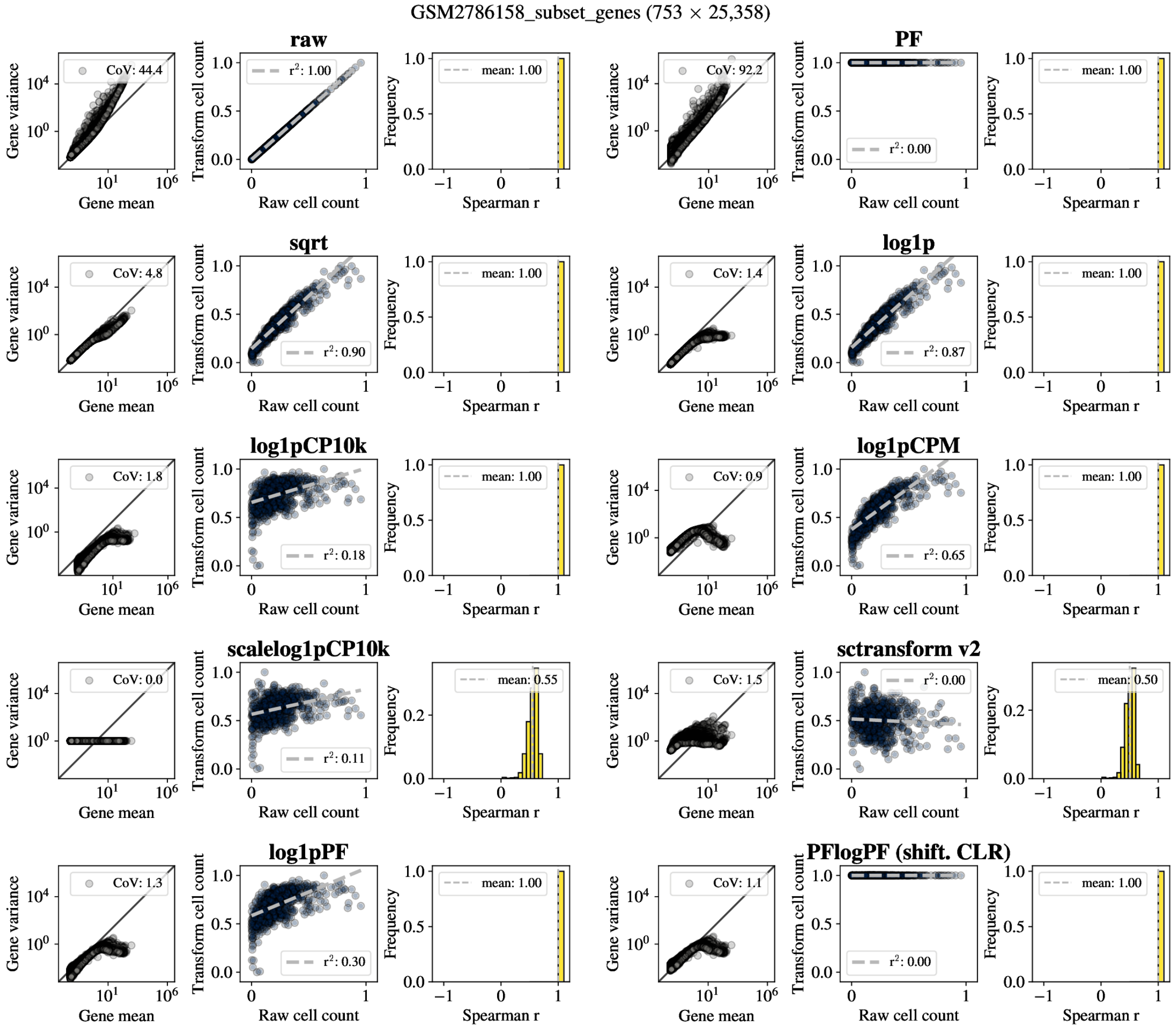


Supplementary Figure 1.146. **GSM2786158**. Per-method comparison of variance stabilization (left), depth normalization (middle), and monotonicity (right) for each normalization on dataset GSM2786158.

| GSM2803967 | |
|---|---|
| cells | 266 |
| genes | 60,623 |
| nonzero entries | 433,171 |
| density | 0.0269 |
| total counts | 2,039,560 |
| mean counts / cell | 7,668 |
| mean counts / gene | 34 |
| min counts / cell | 2,842 |
| max counts / cell | 26,561 |
| overdispersion | 0.267 |

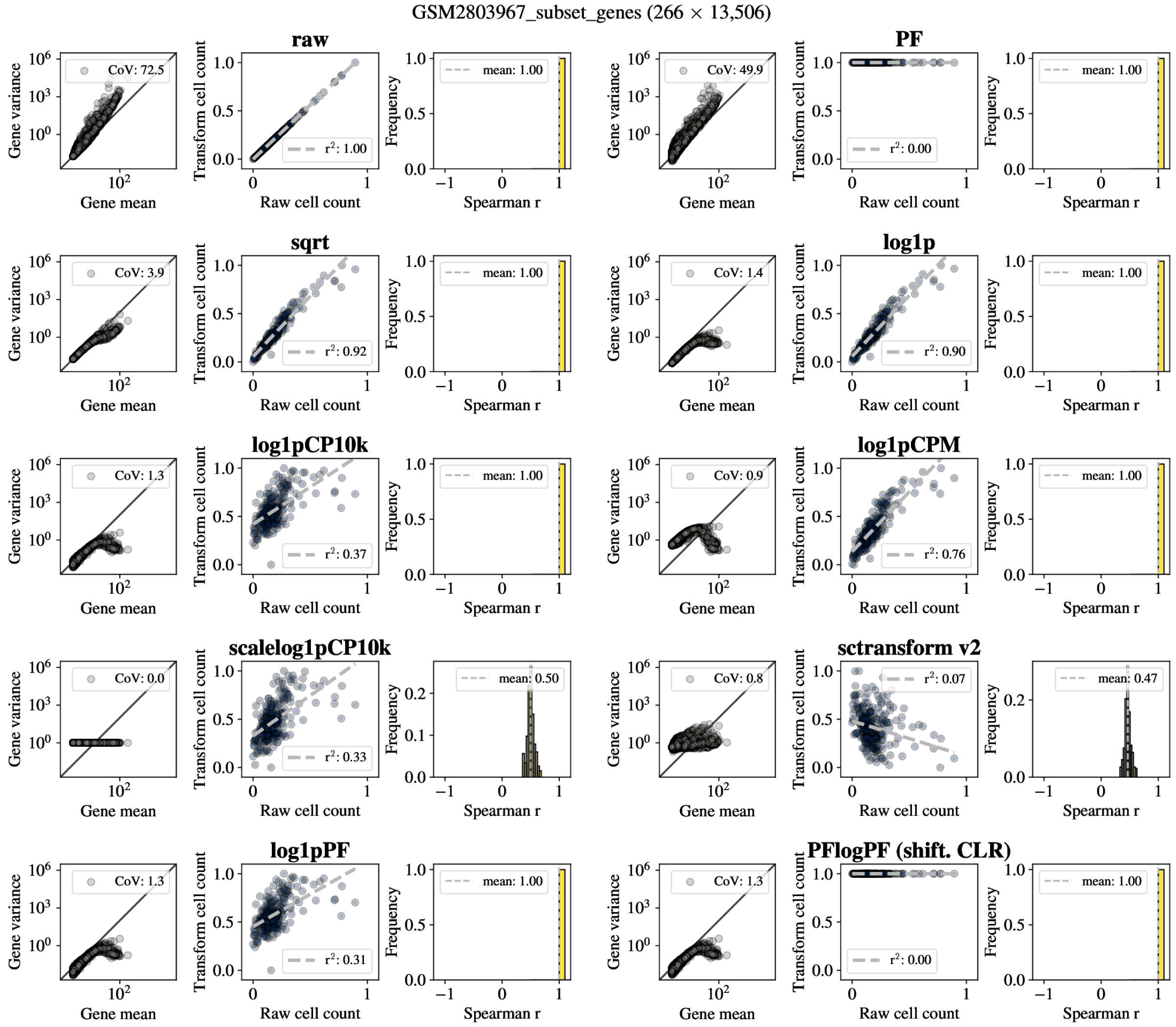


Supplementary Figure 1.147. **GSM2803967**. Per-method comparison of variance stabilization (left), depth normalization (middle), and monotonicity (right) for each normalization on dataset GSM2803967.

| GSM2877959 | |
|---|---|
| cells | 17,075 |
| genes | 60,623 |
| nonzero entries | 10,345,060 |
| density | 0.00999 |
| total counts | 27,517,121 |
| mean counts / cell | 1,612 |
| mean counts / gene | 454 |
| min counts / cell | 87 |
| max counts / cell | 20,858 |
| overdispersion | 5.98 |

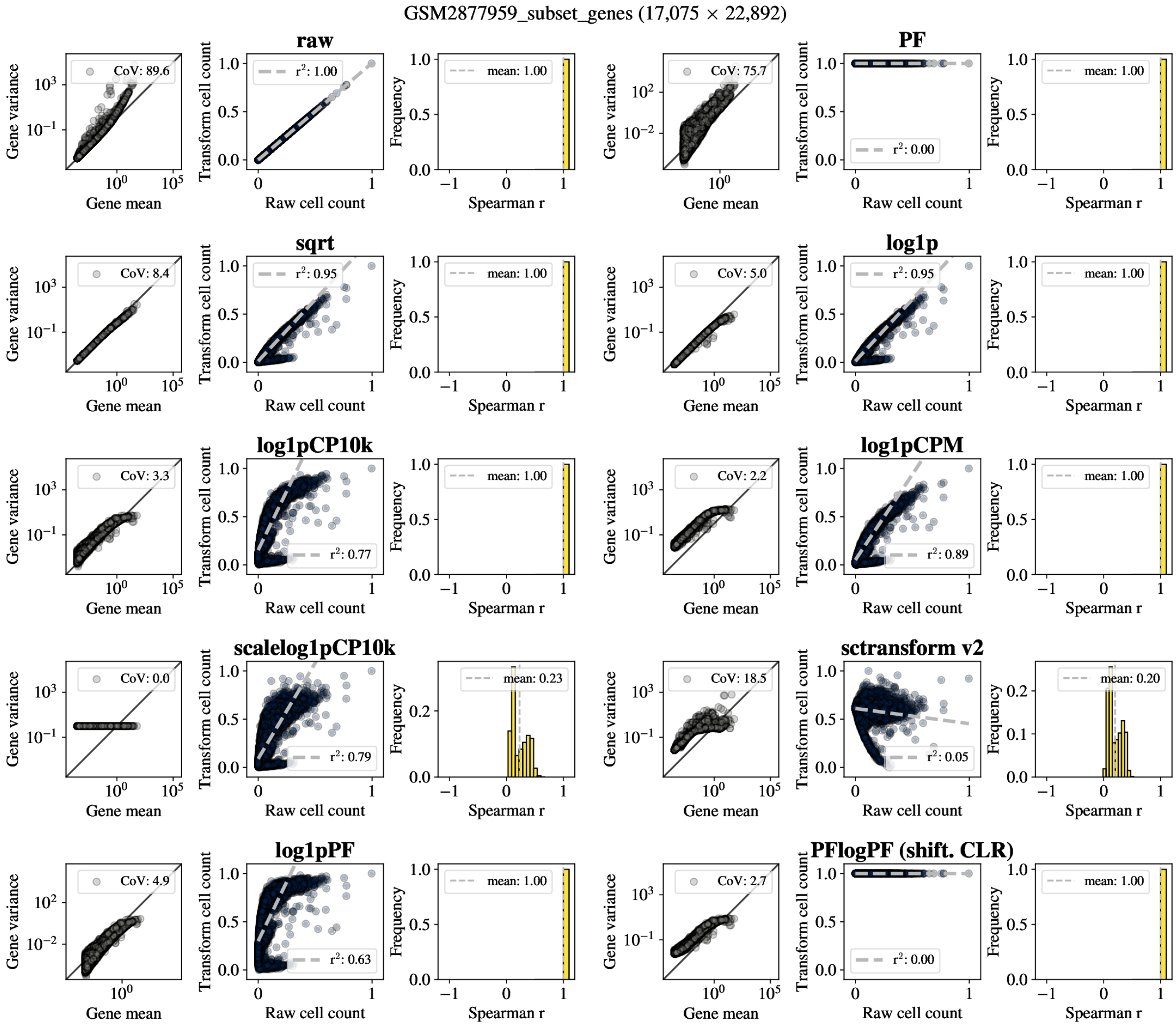


Supplementary Figure 1.148. **GSM2877959**. Per-method comparison of variance stabilization (left), depth normalization (middle), and monotonicity (right) for each normalization on dataset GSM2877959.

| GSM2877960 | |
|---|---|
| cells | 6,277 |
| genes | 60,623 |
| nonzero entries | 4,338,653 |
| density | 0.0114 |
| total counts | 13,721,030 |
| mean counts / cell | 2,186 |
| mean counts / gene | 226 |
| min counts / cell | 364 |
| max counts / cell | 29,870 |
| overdispersion | 6.44 |

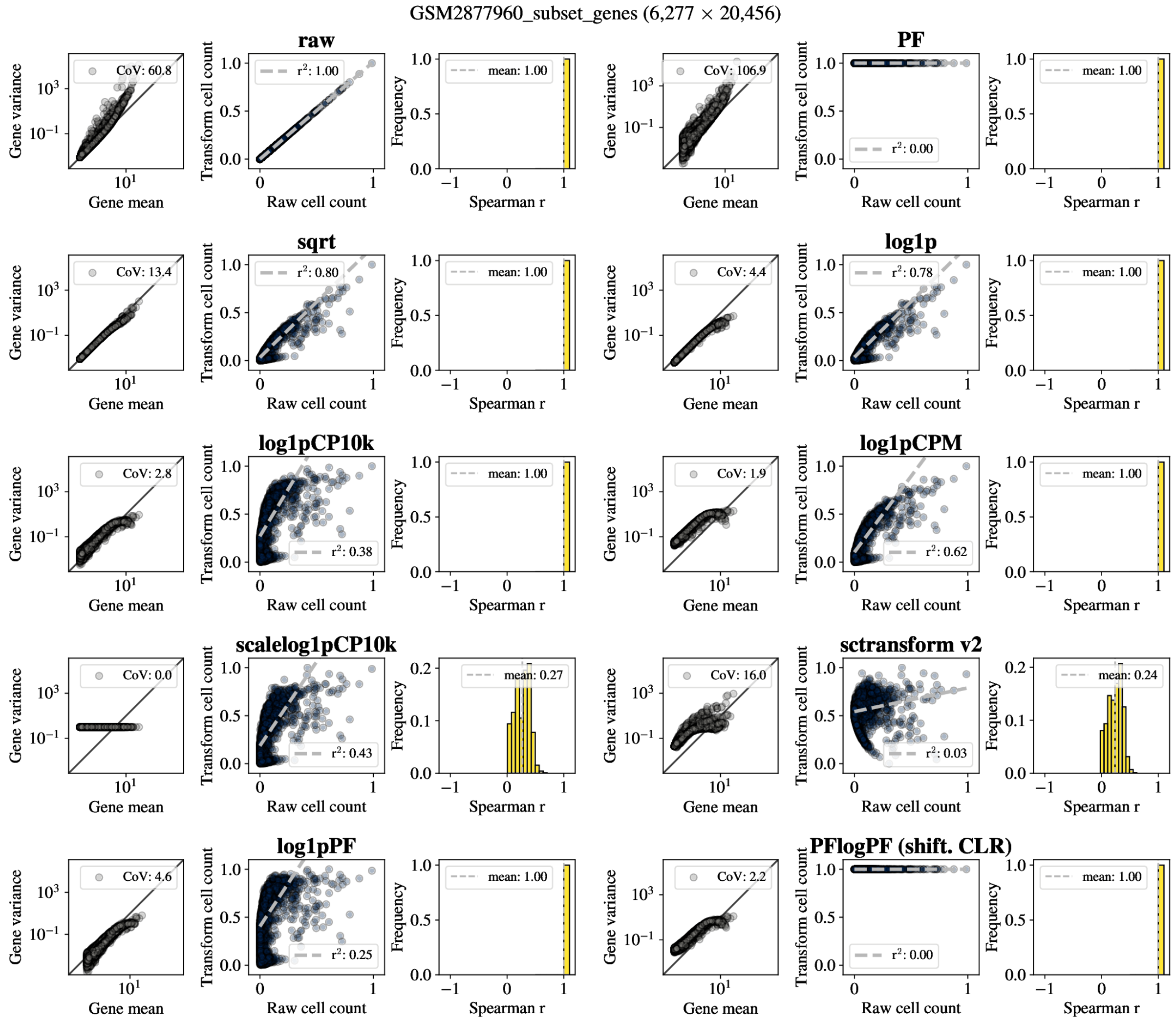


Supplementary Figure 1.149. **GSM2877960**. Per-method comparison of variance stabilization (left), depth normalization (middle), and monotonicity (right) for each normalization on dataset GSM2877960.

| GSM2894834 | |
|---|---|
| cells | 12,728 |
| genes | 60,623 |
| nonzero entries | 25,319,748 |
| density | 0.0328 |
| total counts | 79,420,670 |
| mean counts / cell | 6,240 |
| mean counts / gene | 1,310 |
| min counts / cell | 373 |
| max counts / cell | 37,684 |
| overdispersion | 2.8 |

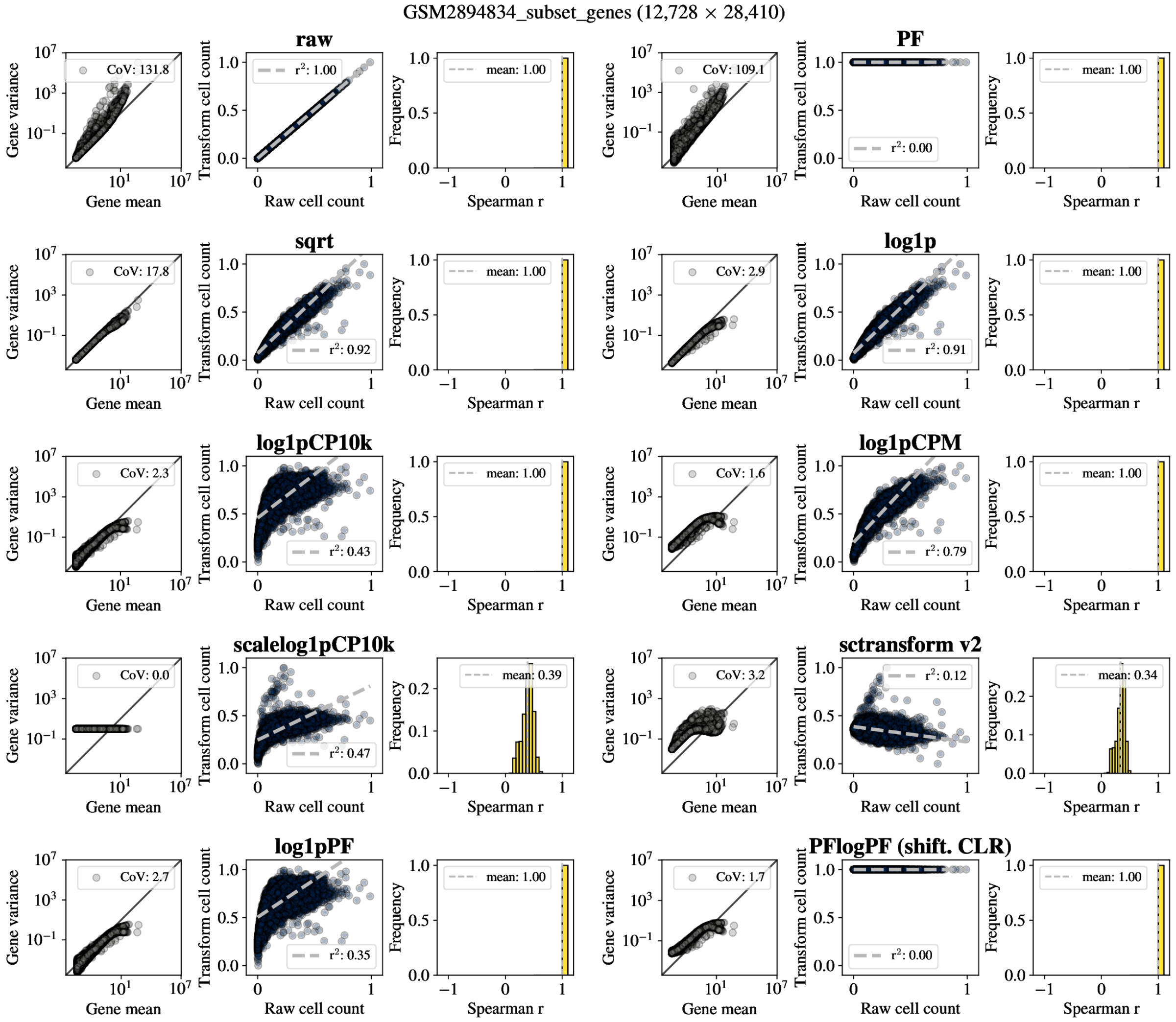


Supplementary Figure 1.150. **GSM2894834**. Per-method comparison of variance stabilization (left), depth normalization (middle), and monotonicity (right) for each normalization on dataset GSM2894834.

| GSM2894835 | |
|---|---|
| cells | 12,259 |
| genes | 60,623 |
| nonzero entries | 25,647,743 |
| density | 0.0345 |
| total counts | 80,242,513 |
| mean counts / cell | 6,546 |
| mean counts / gene | 1,324 |
| min counts / cell | 372 |
| max counts / cell | 35,045 |
| overdispersion | 2.59 |

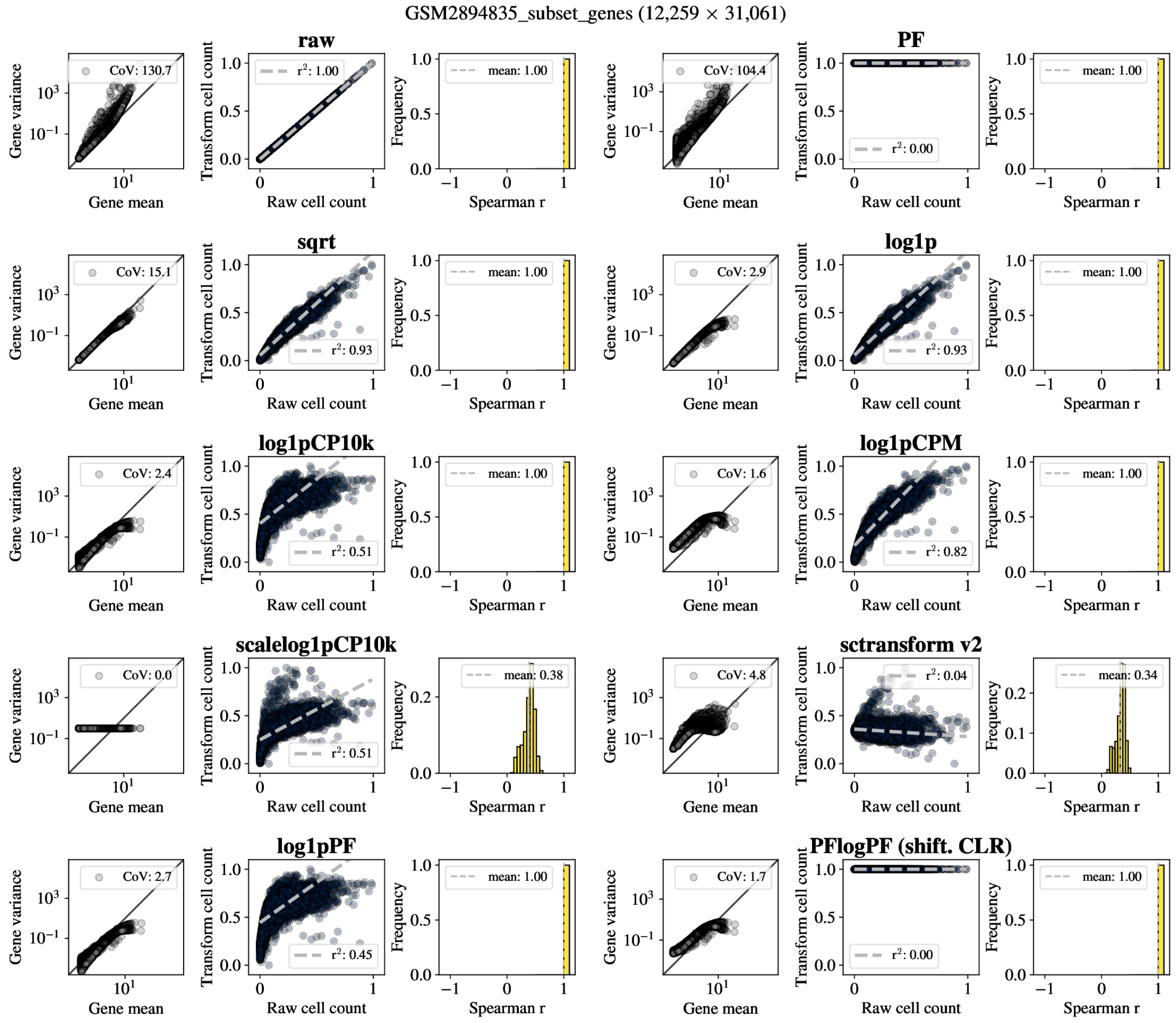


Supplementary Figure 1.151. **GSM2894835**. Per-method comparison of variance stabilization (left), depth normalization (middle), and monotonicity (right) for each normalization on dataset GSM2894835.

| GSM2905290 | |
|---|---|
| cells | 8,319 |
| genes | 60,623 |
| nonzero entries | 6,718,390 |
| density | 0.0133 |
| total counts | 19,515,450 |
| mean counts / cell | 2,346 |
| mean counts / gene | 322 |
| min counts / cell | 110 |
| max counts / cell | 39,720 |
| overdispersion | 5.29 |

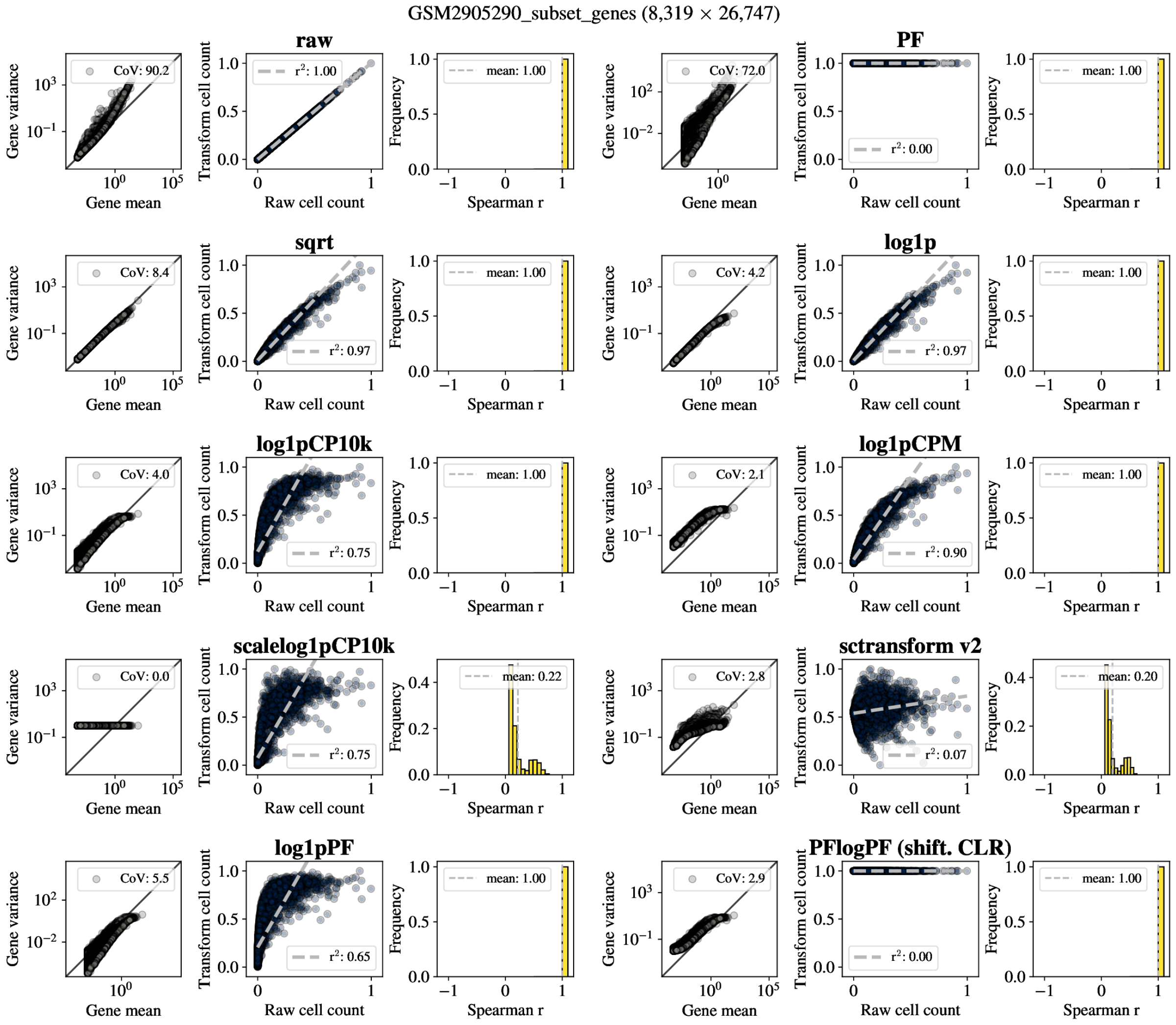


Supplementary Figure 1.152. **GSM2905290**. Per-method comparison of variance stabilization (left), depth normalization (middle), and monotonicity (right) for each normalization on dataset GSM2905290.

| GSM2905291 | |
|---|---|
| cells | 8,296 |
| genes | 60,623 |
| nonzero entries | 25,277,483 |
| density | 0.0503 |
| total counts | 88,151,013 |
| mean counts / cell | 10,626 |
| mean counts / gene | 1,454 |
| min counts / cell | 1,091 |
| max counts / cell | 162,529 |
| overdispersion | 0.788 |

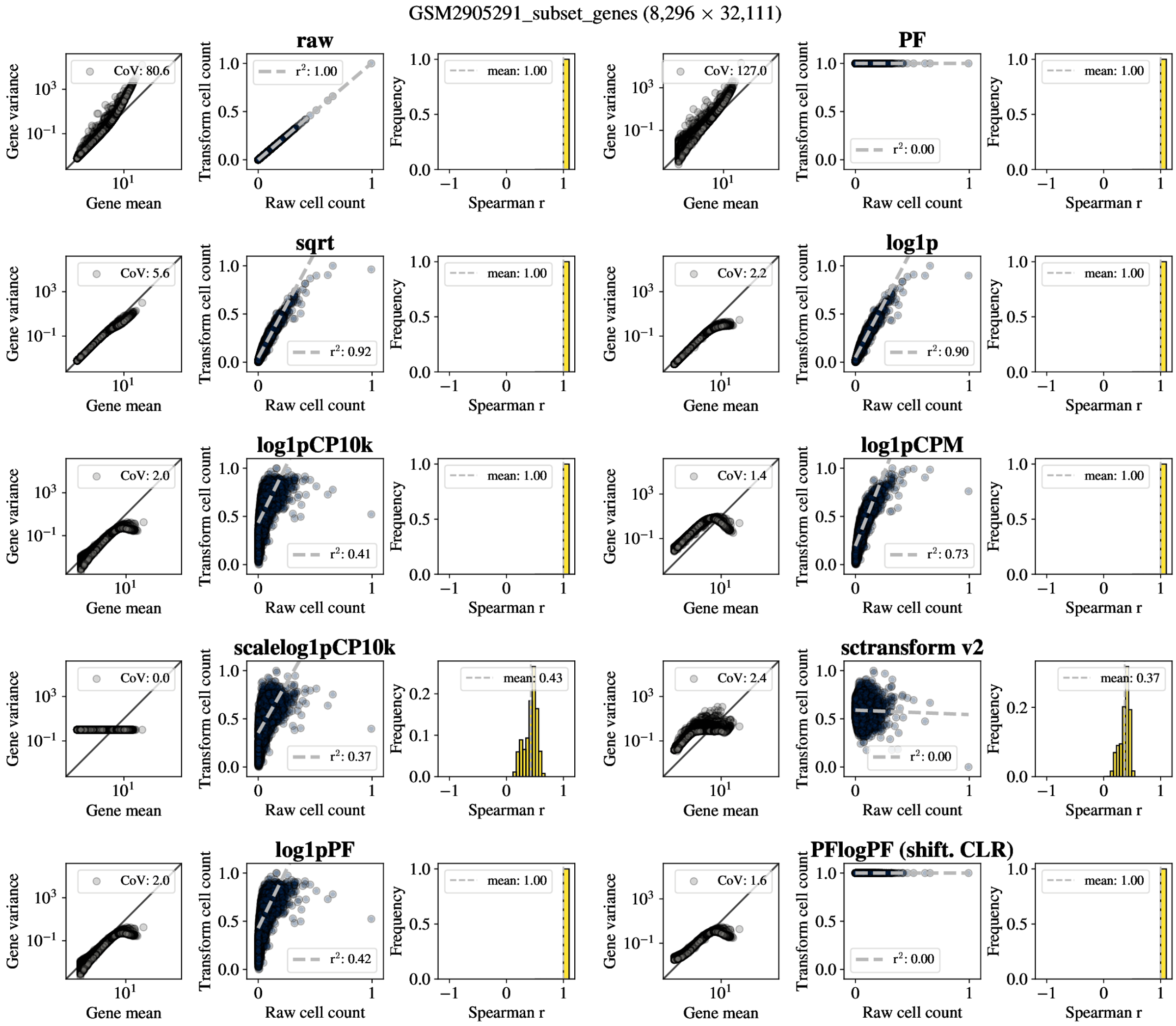


Supplementary Figure 1.153. **GSM2905291**. Per-method comparison of variance stabilization (left), depth normalization (middle), and monotonicity (right) for each normalization on dataset GSM2905291.

| GSM2905292 | |
|---|---|
| cells | 9,939 |
| genes | 60,623 |
| nonzero entries | 5,139,873 |
| density | 0.00853 |
| total counts | 13,385,843 |
| mean counts / cell | 1,347 |
| mean counts / gene | 221 |
| min counts / cell | 82 |
| max counts / cell | 53,510 |
| overdispersion | 8.7 |

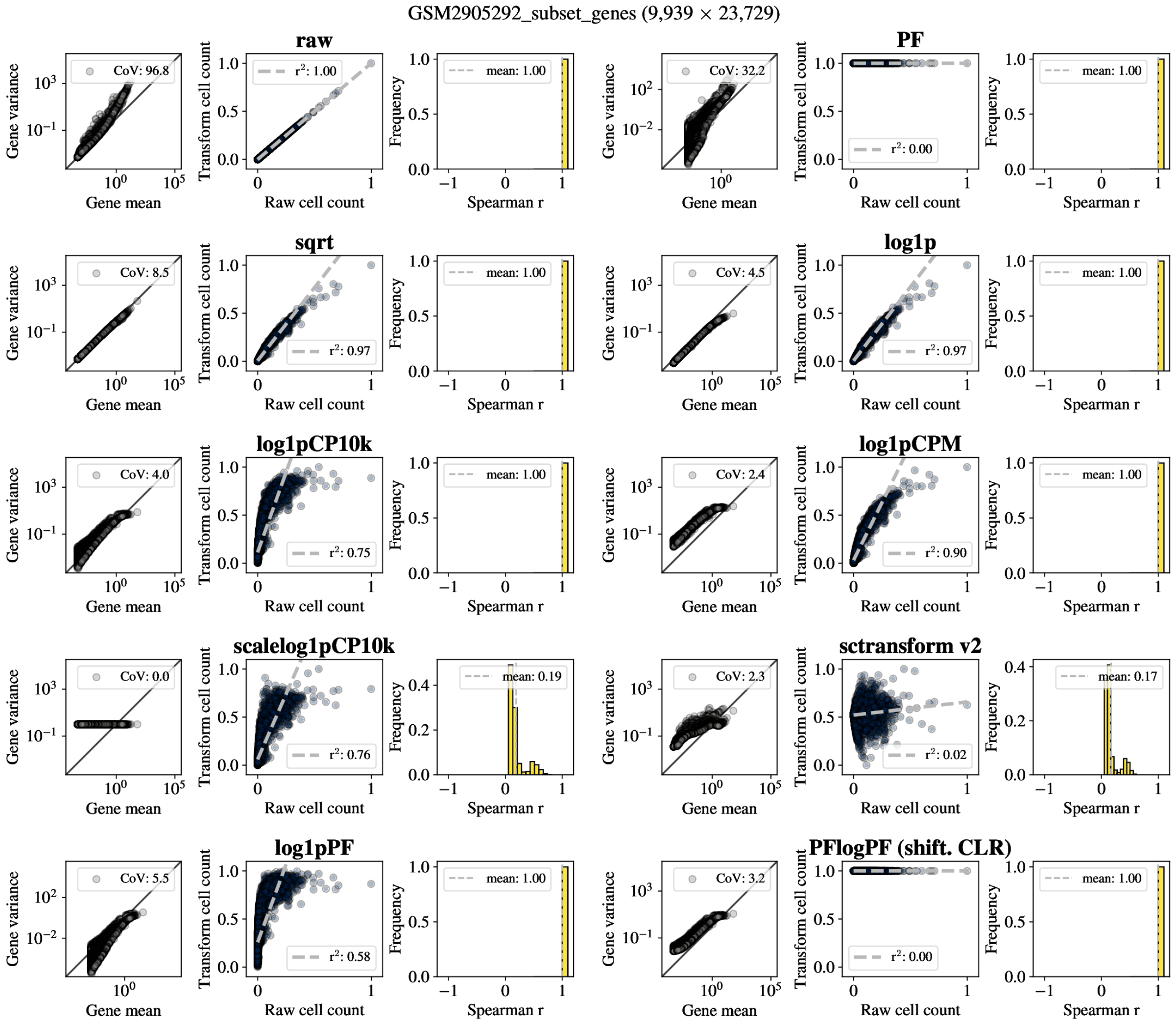


Supplementary Figure 1.154. **GSM2905292**. Per-method comparison of variance stabilization (left), depth normalization (middle), and monotonicity (right) for each normalization on dataset GSM2905292.

| GSM2905293 | |
|---|---|
| cells | 8,509 |
| genes | 60,623 |
| nonzero entries | 24,557,461 |
| density | 0.0476 |
| total counts | 89,876,300 |
| mean counts / cell | 10,562 |
| mean counts / gene | 1,483 |
| min counts / cell | 285 |
| max counts / cell | 160,187 |
| overdispersion | 1.24 |

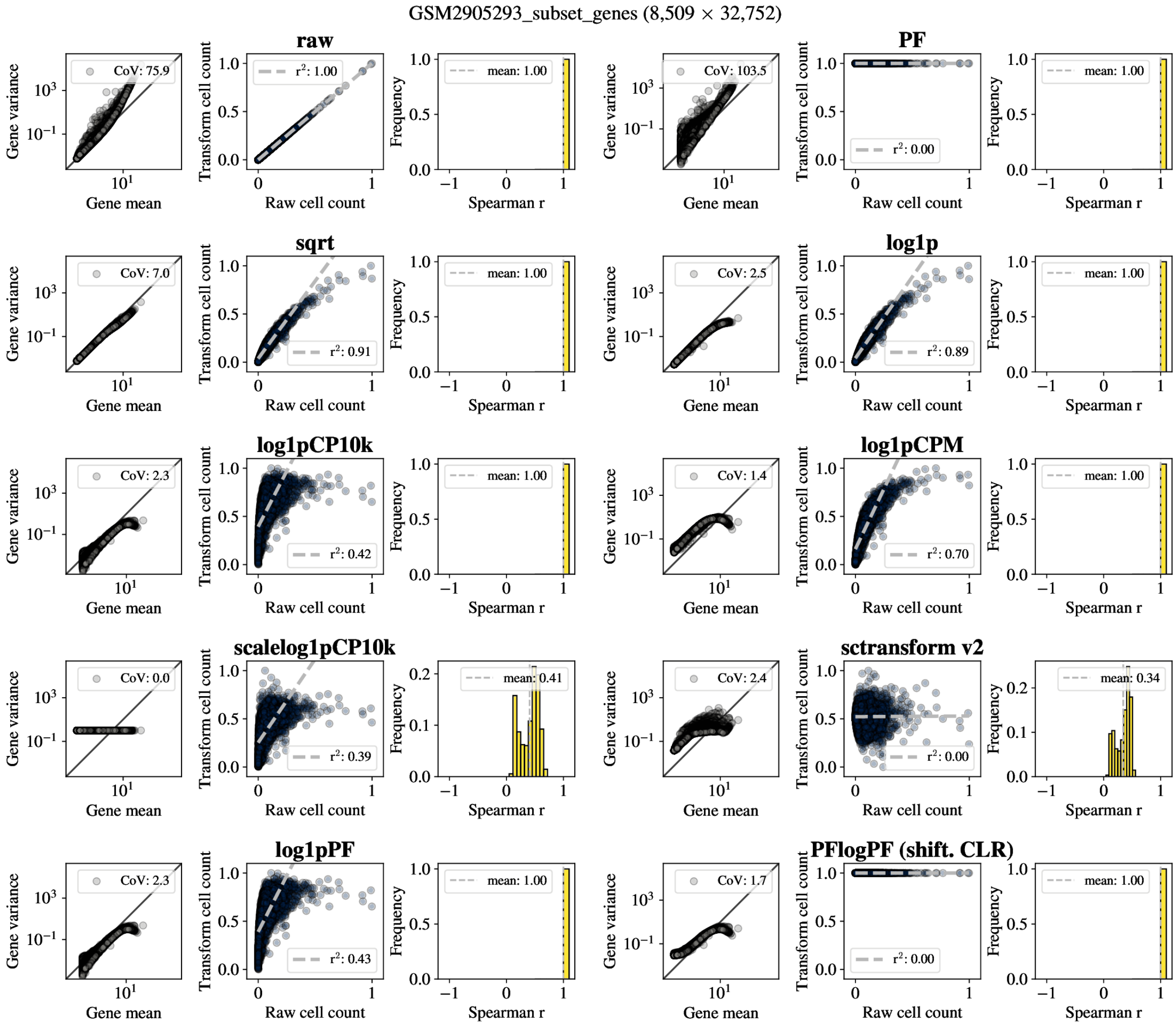


Supplementary Figure 1.155. **GSM2905293**. Per-method comparison of variance stabilization (left), depth normalization (middle), and monotonicity (right) for each normalization on dataset GSM2905293.

| GSM2928377 | |
|---|---|
| cells | 2,893 |
| genes | 60,623 |
| nonzero entries | 13,505,179 |
| density | 0.077 |
| total counts | 34,785,308 |
| mean counts / cell | 12,024 |
| mean counts / gene | 574 |
| min counts / cell | 8,210 |
| max counts / cell | 35,002 |
| overdispersion | 0.293 |

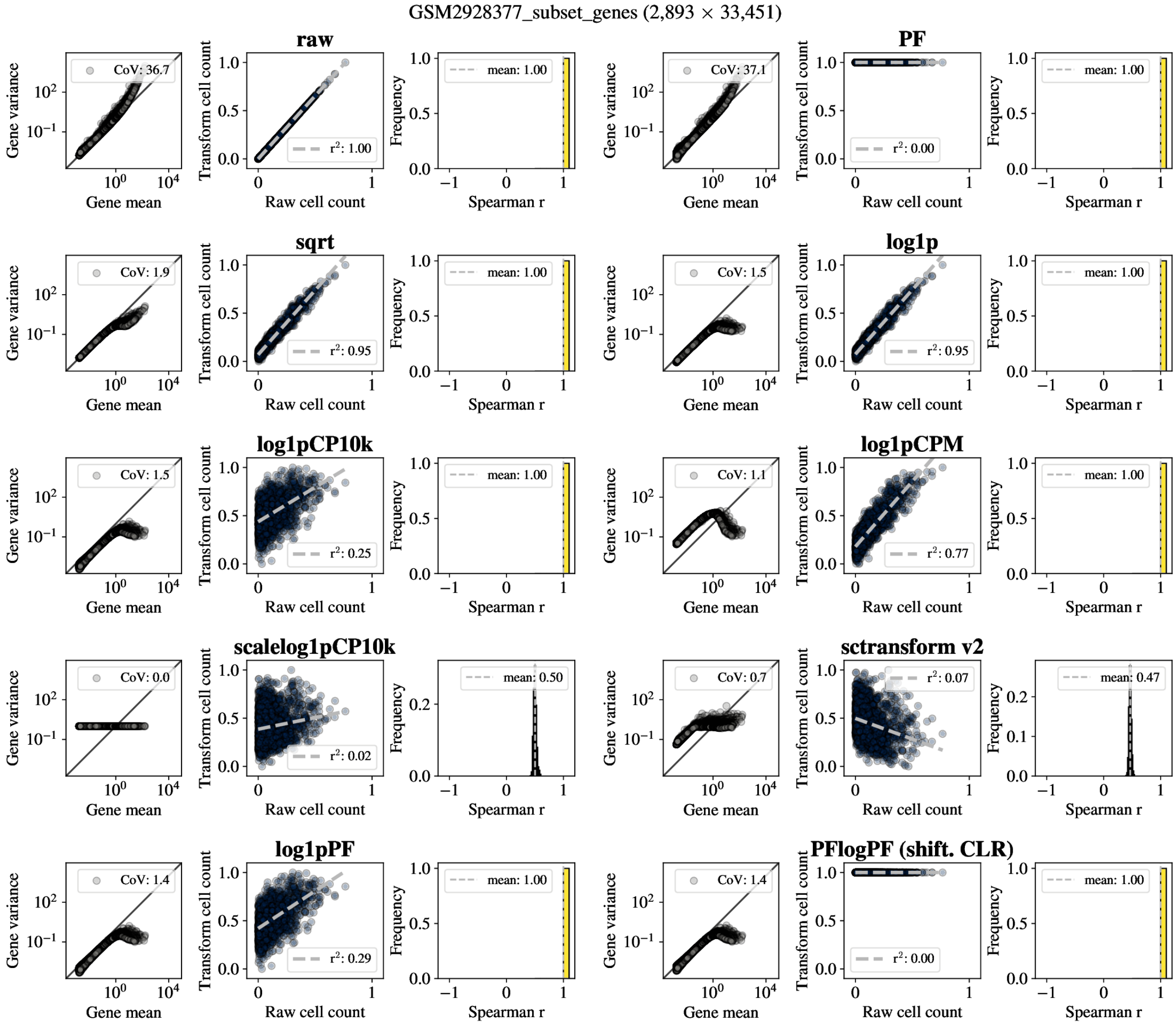


Supplementary Figure 1.156. **GSM2928377**. Per-method comparison of variance stabilization (left), depth normalization (middle), and monotonicity (right) for each normalization on dataset GSM2928377.

| GSM2928378 | |
|---|---|
| cells | 3,462 |
| genes | 60,623 |
| nonzero entries | 12,657,332 |
| density | 0.0603 |
| total counts | 30,790,949 |
| mean counts / cell | 8,894 |
| mean counts / gene | 508 |
| min counts / cell | 2,581 |
| max counts / cell | 29,904 |
| overdispersion | 0.317 |

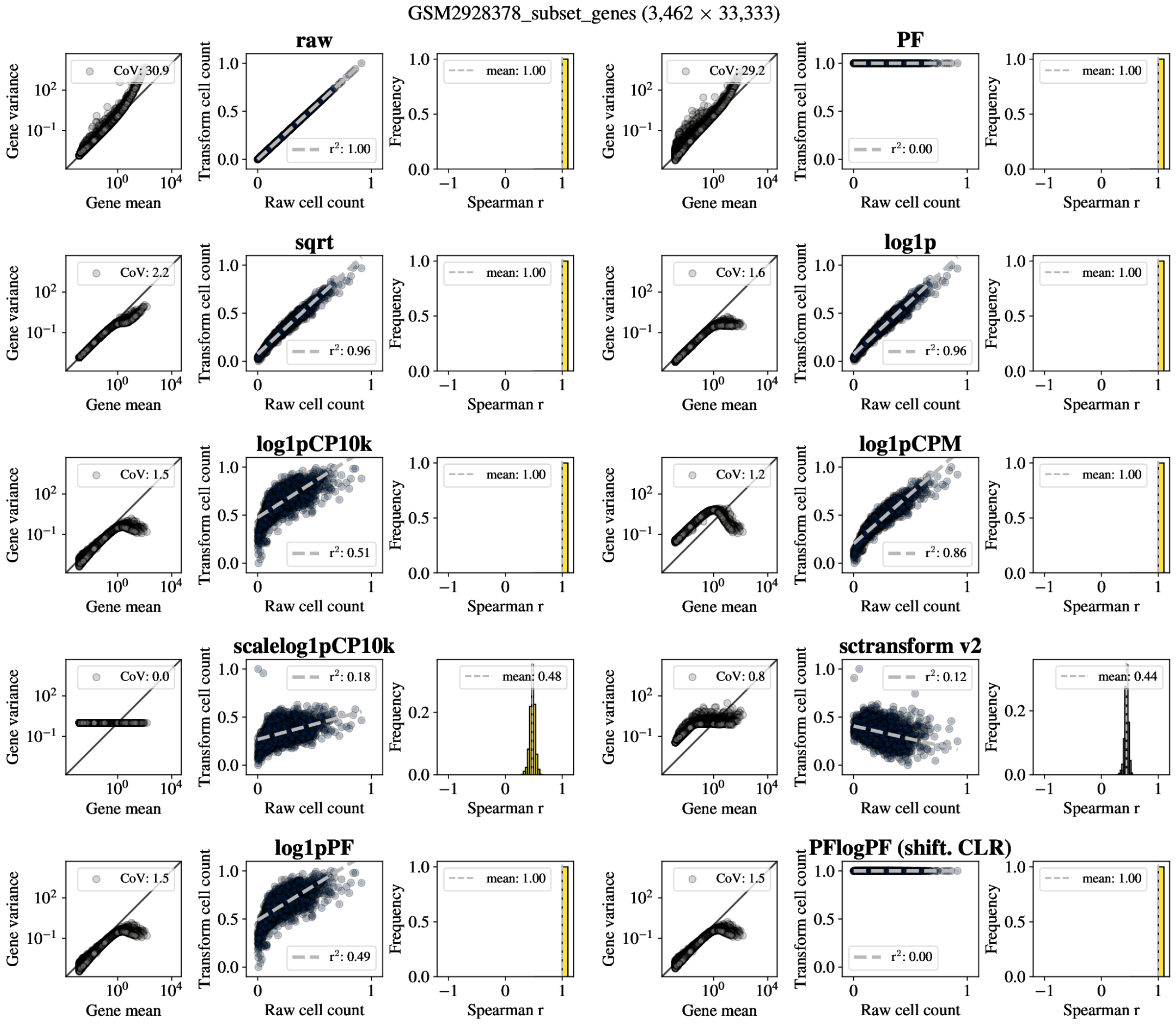


Supplementary Figure 1.157. **GSM2928378**. Per-method comparison of variance stabilization (left), depth normalization (middle), and monotonicity (right) for each normalization on dataset GSM2928378.

| GSM2928379 | |
|---|---|
| cells | 3,494 |
| genes | 60,623 |
| nonzero entries | 13,030,023 |
| density | 0.0615 |
| total counts | 35,872,390 |
| mean counts / cell | 10,267 |
| mean counts / gene | 592 |
| min counts / cell | 2,695 |
| max counts / cell | 36,288 |
| overdispersion | 0.409 |

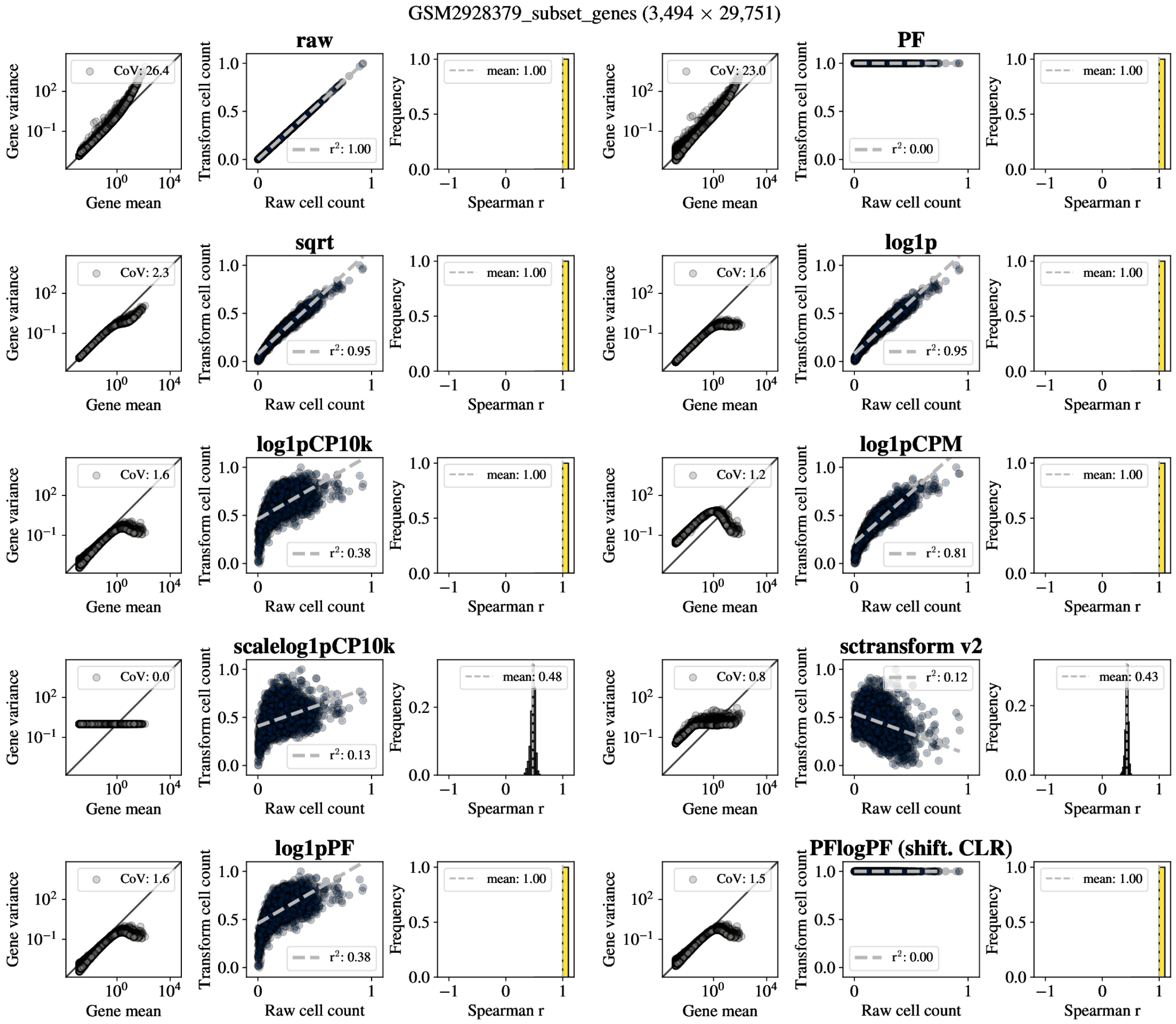


Supplementary Figure 1.158. **GSM2928379**. Per-method comparison of variance stabilization (left), depth normalization (middle), and monotonicity (right) for each normalization on dataset GSM2928379.

| GSM2928380 | |
|---|---|
| cells | 19,962 |
| genes | 60,623 |
| nonzero entries | 15,286,385 |
| density | 0.0126 |
| total counts | 46,049,097 |
| mean counts / cell | 2,307 |
| mean counts / gene | 760 |
| min counts / cell | 206 |
| max counts / cell | 132,763 |
| overdispersion | 17.2 |

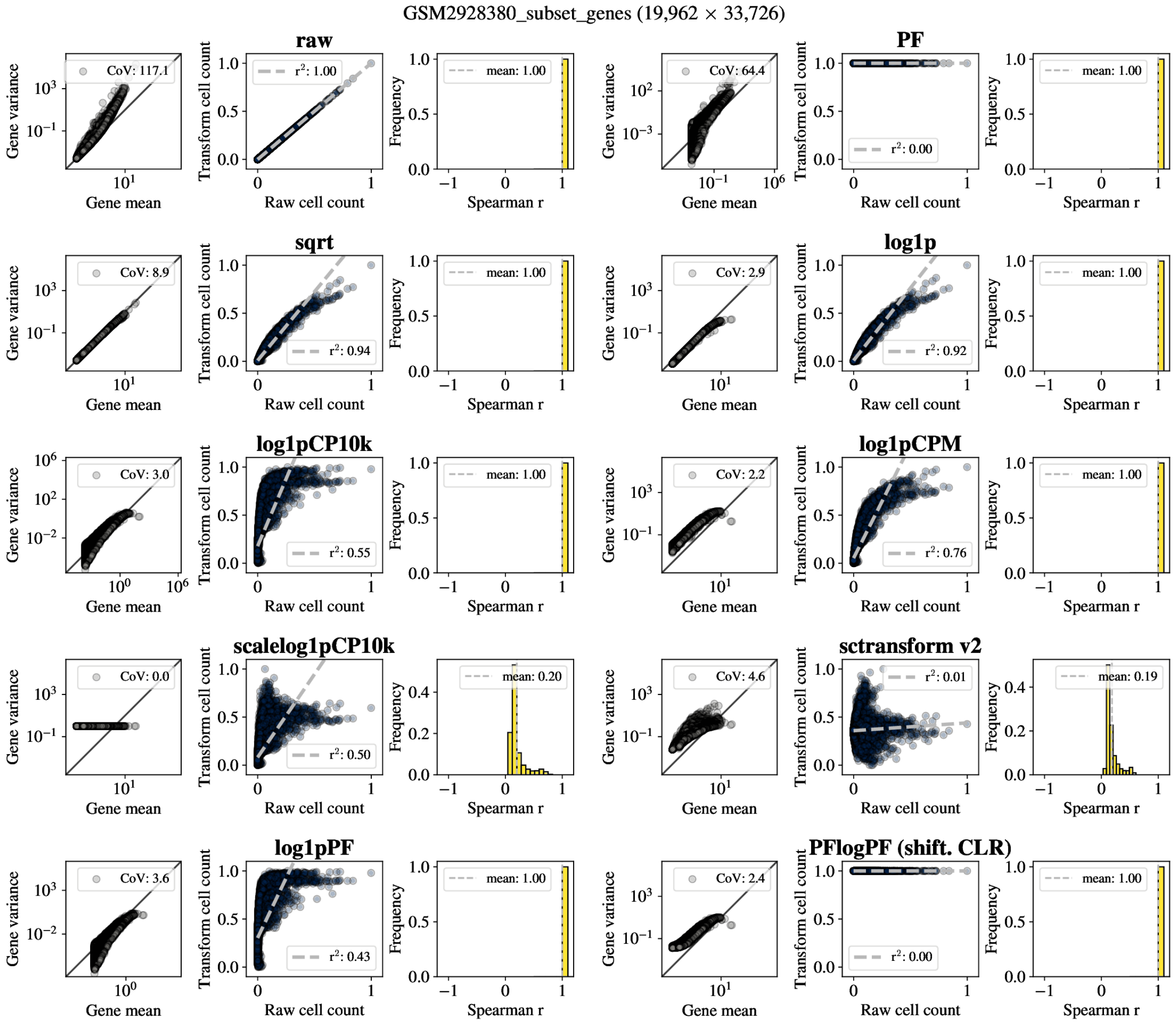


Supplementary Figure 1.159. **GSM2928380**. Per-method comparison of variance stabilization (left), depth normalization (middle), and monotonicity (right) for each normalization on dataset GSM2928380.

| GSM2928381 | |
|---|---|
| cells | 6,280 |
| genes | 60,623 |
| nonzero entries | 17,455,400 |
| density | 0.0458 |
| total counts | 59,548,961 |
| mean counts / cell | 9,482 |
| mean counts / gene | 982 |
| min counts / cell | 734 |
| max counts / cell | 138,104 |
| overdispersion | 12.5 |

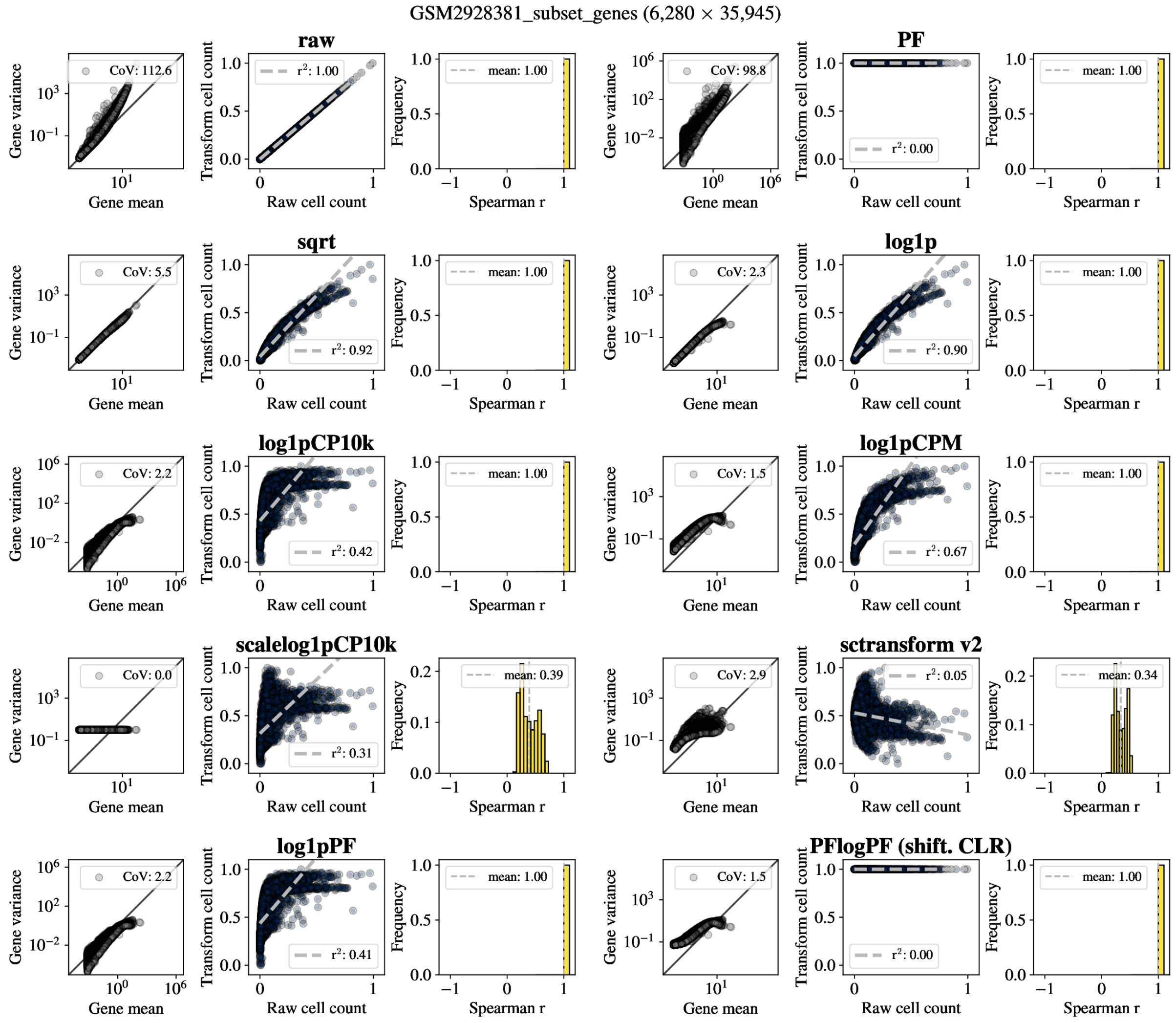


Supplementary Figure 1.160. **GSM2928381**. Per-method comparison of variance stabilization (left), depth normalization (middle), and monotonicity (right) for each normalization on dataset GSM2928381.

| **GSM2928382** | |
|---|---|
| cells | 7,899 |
| genes | 60,623 |
| nonzero entries | 20,010,994 |
| density | 0.0418 |
| total counts | 68,722,931 |
| mean counts / cell | 8,700 |
| mean counts / gene | 1,134 |
| min counts / cell | 367 |
| max counts / cell | 188,299 |
| overdispersion | 18.5 |

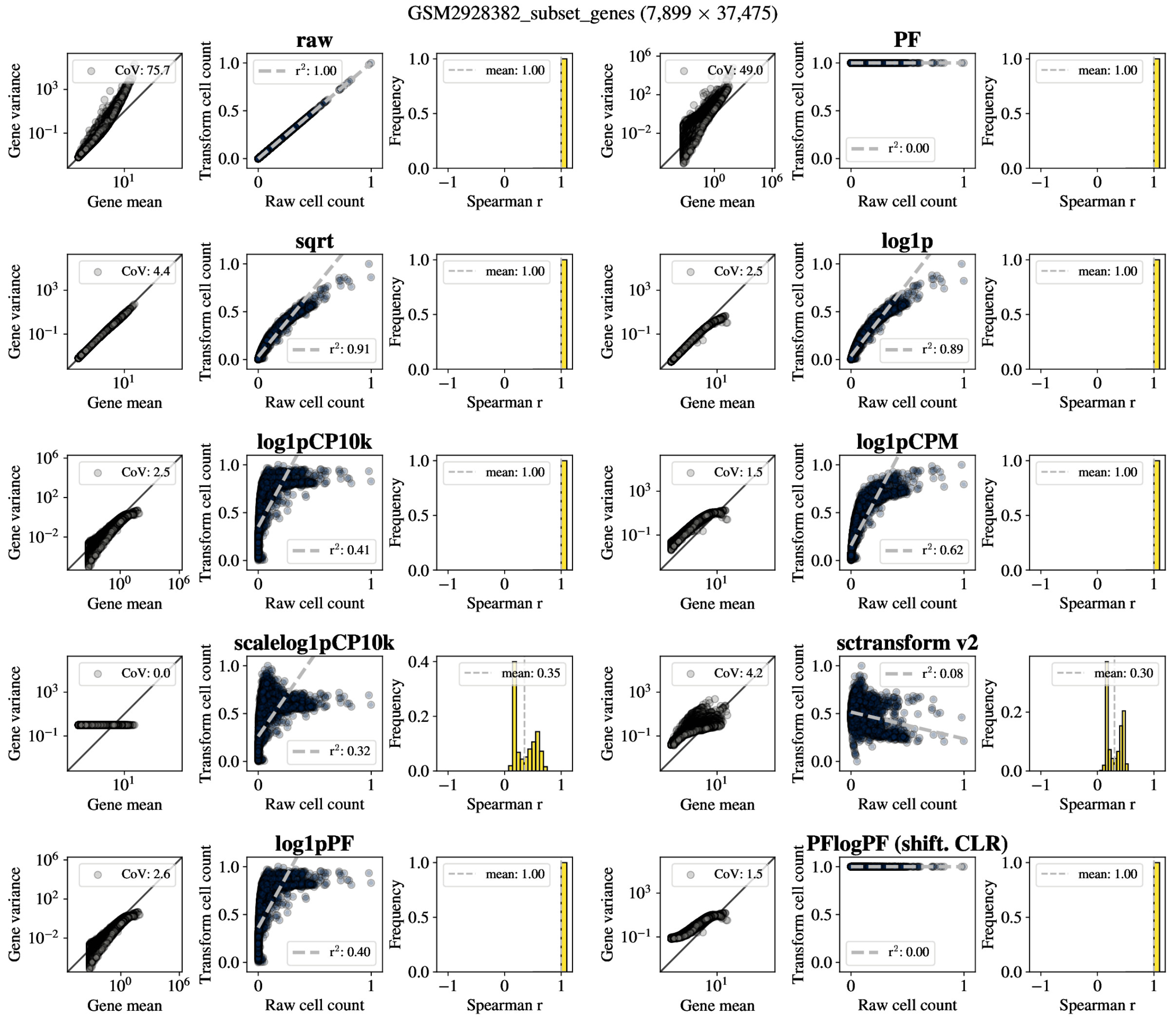


Supplementary Figure 1.161. **GSM2928382**. Per-method comparison of variance stabilization (left), depth normalization (middle), and monotonicity (right) for each normalization on dataset GSM2928382.

| GSM2928383 | |
|---|---|
| cells | 2,212 |
| genes | 60,623 |
| nonzero entries | 15,219,136 |
| density | 0.113 |
| total counts | 82,243,585 |
| mean counts / cell | 37,181 |
| mean counts / gene | 1,357 |
| min counts / cell | 16,288 |
| max counts / cell | 134,720 |
| overdispersion | 1.9 |

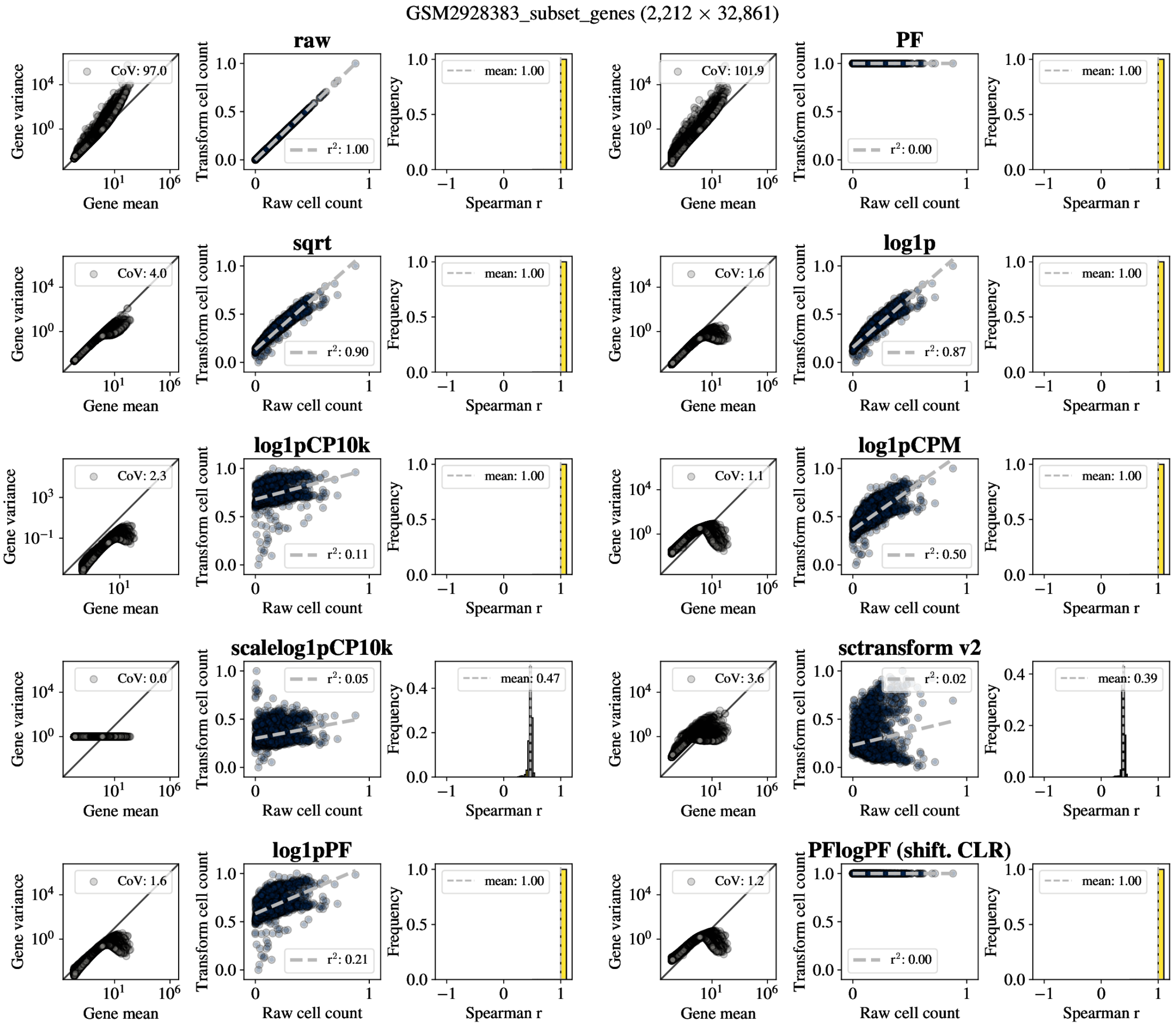


Supplementary Figure 1.162. **GSM2928383**. Per-method comparison of variance stabilization (left), depth normalization (middle), and monotonicity (right) for each normalization on dataset GSM2928383.

| GSM2928384 | |
|---|---|
| cells | 11,360 |
| genes | 60,623 |
| nonzero entries | 22,483,374 |
| density | 0.0326 |
| total counts | 69,479,680 |
| mean counts / cell | 6,116 |
| mean counts / gene | 1,146 |
| min counts / cell | 413 |
| max counts / cell | 88,542 |
| overdispersion | 20.9 |

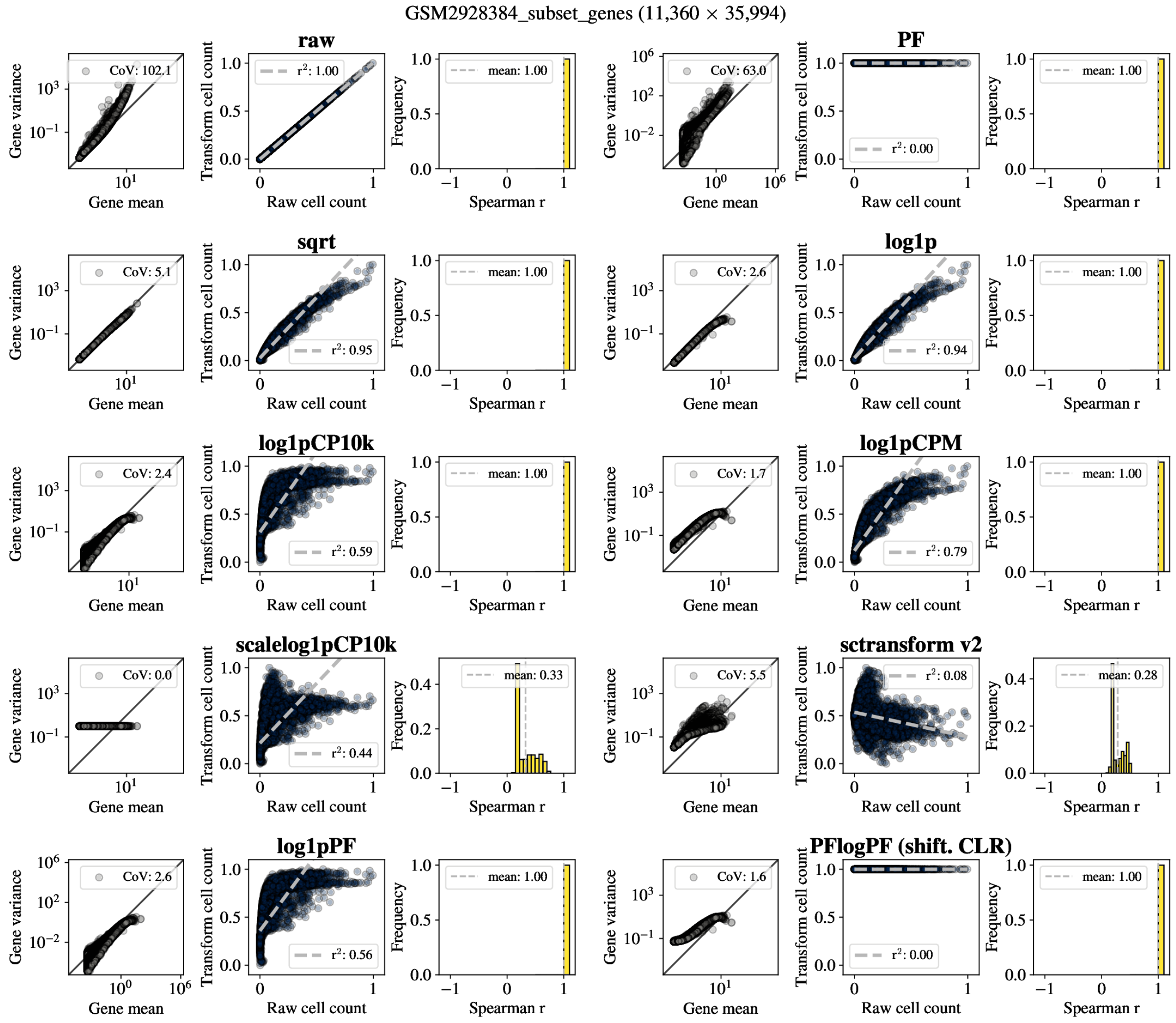


Supplementary Figure 1.163. **GSM2928384**. Per-method comparison of variance stabilization (left), depth normalization (middle), and monotonicity (right) for each normalization on dataset GSM2928384.

| GSM3020393 | |
|---|---|
| cells | 3,017 |
| genes | 60,623 |
| nonzero entries | 3,414,587 |
| density | 0.0187 |
| total counts | 7,217,895 |
| mean counts / cell | 2,392 |
| mean counts / gene | 119 |
| min counts / cell | 385 |
| max counts / cell | 14,547 |
| overdispersion | 0.169 |

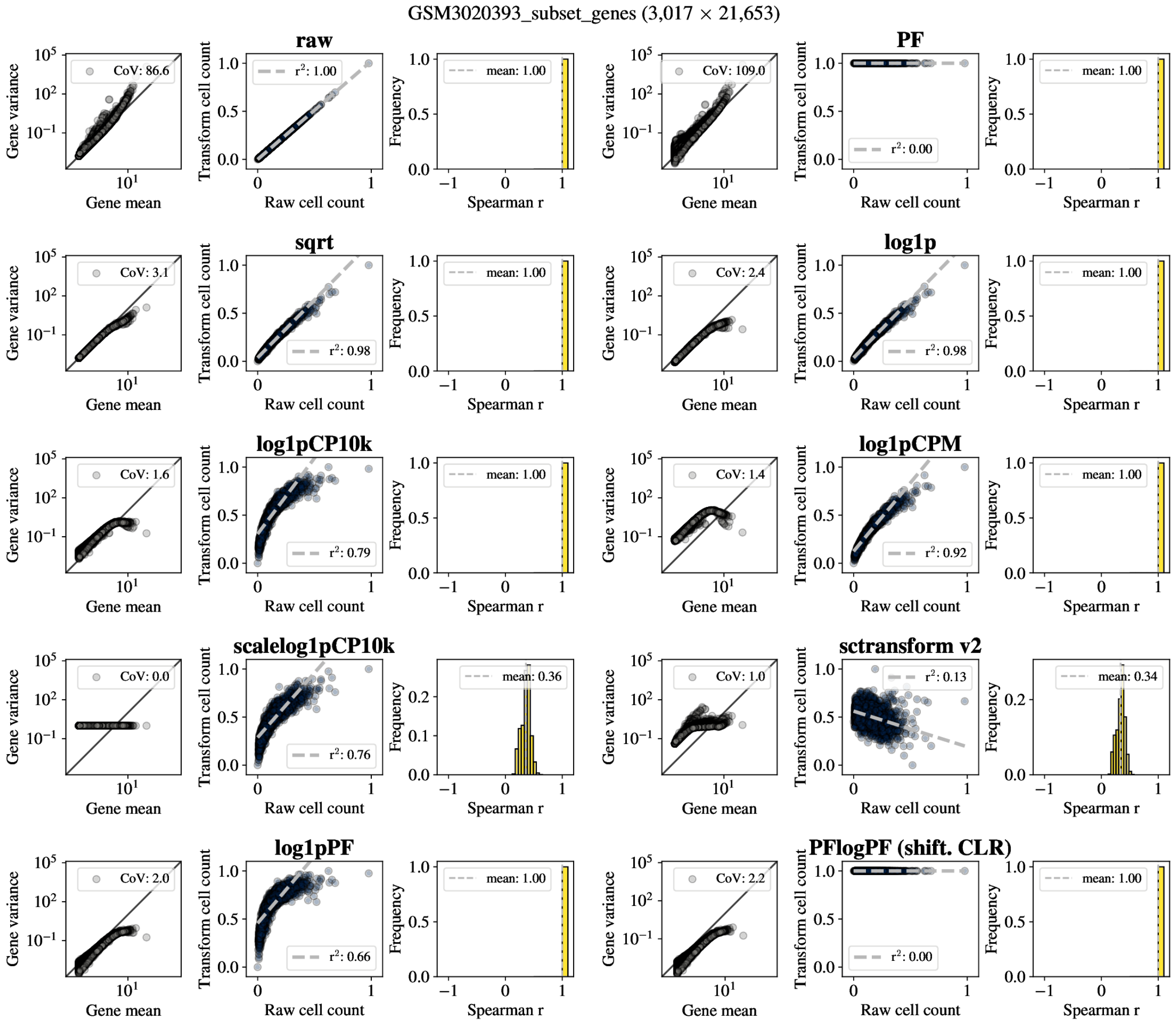


Supplementary Figure 1.164. **GSM3020393**. Per-method comparison of variance stabilization (left), depth normalization (middle), and monotonicity (right) for each normalization on dataset GSM3020393.

| GSM3052917 | |
|---|---|
| cells | 11,420 |
| genes | 60,623 |
| nonzero entries | 22,030,294 |
| density | 0.0318 |
| total counts | 65,478,192 |
| mean counts / cell | 5,734 |
| mean counts / gene | 1,080 |
| min counts / cell | 384 |
| max counts / cell | 83,706 |
| overdispersion | 20.8 |

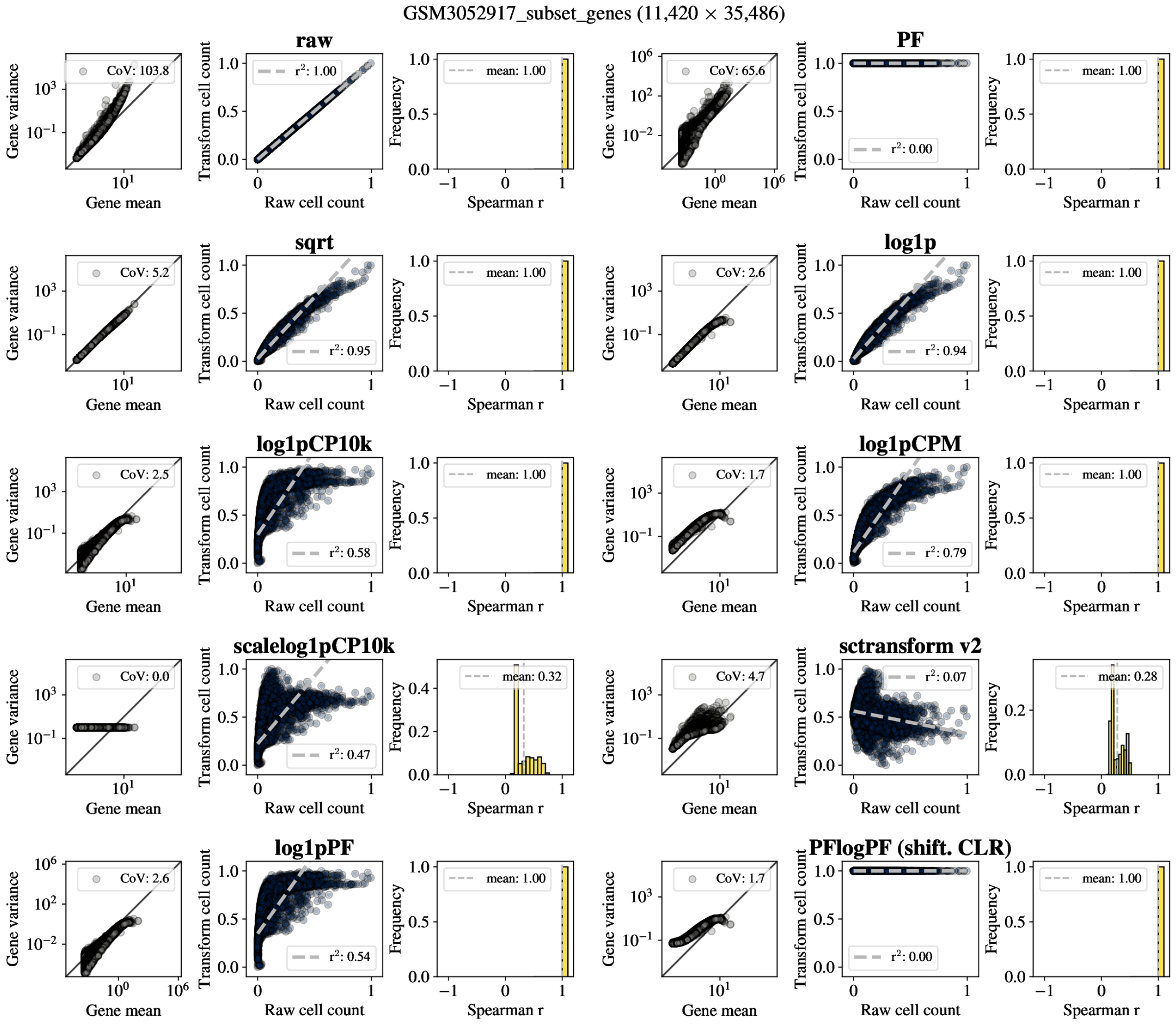


Supplementary Figure 1.165. **GSM3052917**. Per-method comparison of variance stabilization (left), depth normalization (middle), and monotonicity (right) for each normalization on dataset GSM3052917.

| GSM3052919 | |
|---|---|
| cells | 3,800 |
| genes | 60,623 |
| nonzero entries | 6,774,883 |
| density | 0.0294 |
| total counts | 23,977,612 |
| mean counts / cell | 6,310 |
| mean counts / gene | 396 |
| min counts / cell | 614 |
| max counts / cell | 142,012 |
| overdispersion | 8.39 |

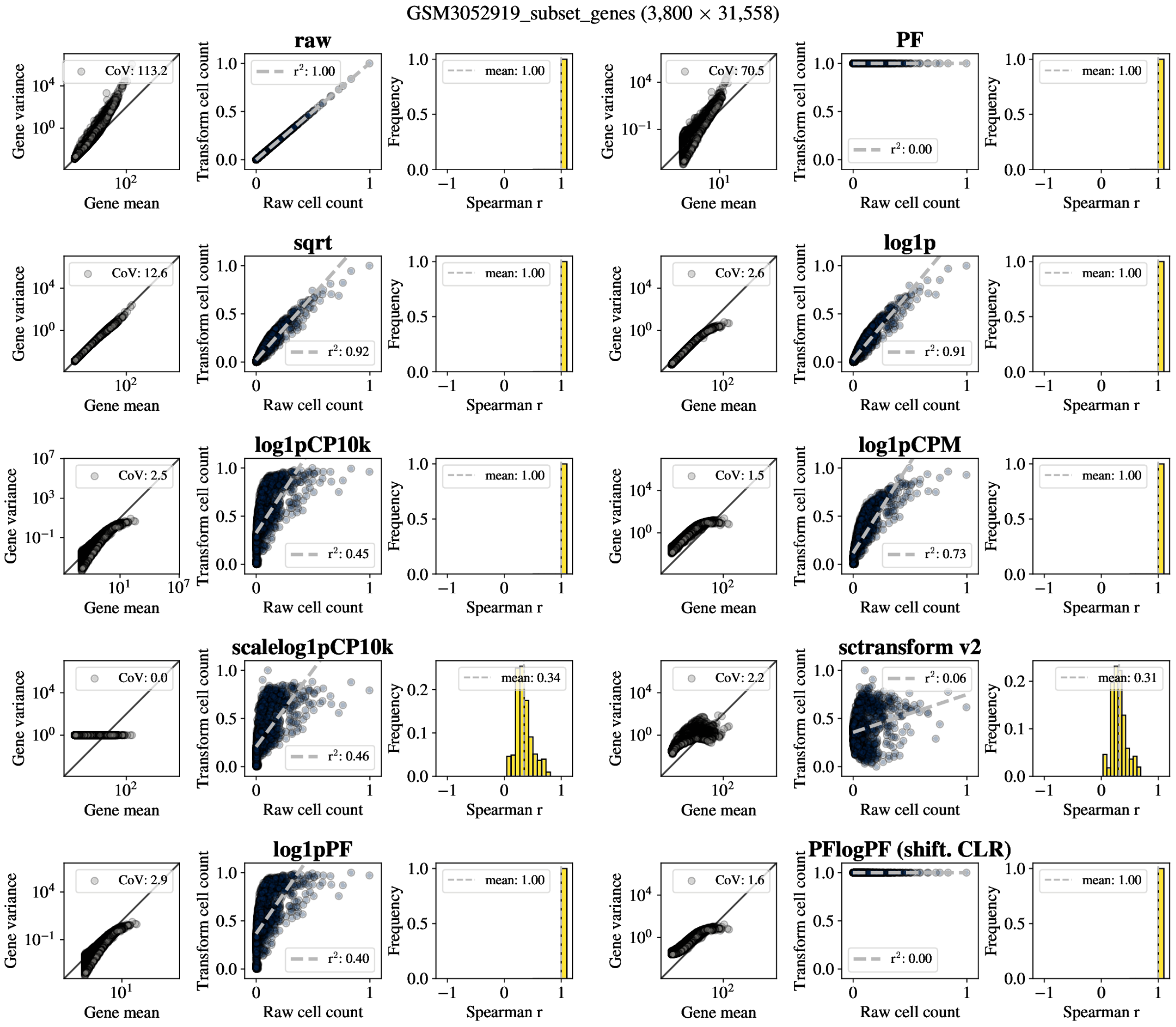


Supplementary Figure 1.166. **GSM3052919**. Per-method comparison of variance stabilization (left), depth normalization (middle), and monotonicity (right) for each normalization on dataset GSM3052919.

| GSM3052921 | |
|---|---|
| cells | 7,247 |
| genes | 60,623 |
| nonzero entries | 7,173,610 |
| density | 0.0163 |
| total counts | 24,554,060 |
| mean counts / cell | 3,388 |
| mean counts / gene | 405 |
| min counts / cell | 381 |
| max counts / cell | 93,541 |
| overdispersion | 10.7 |

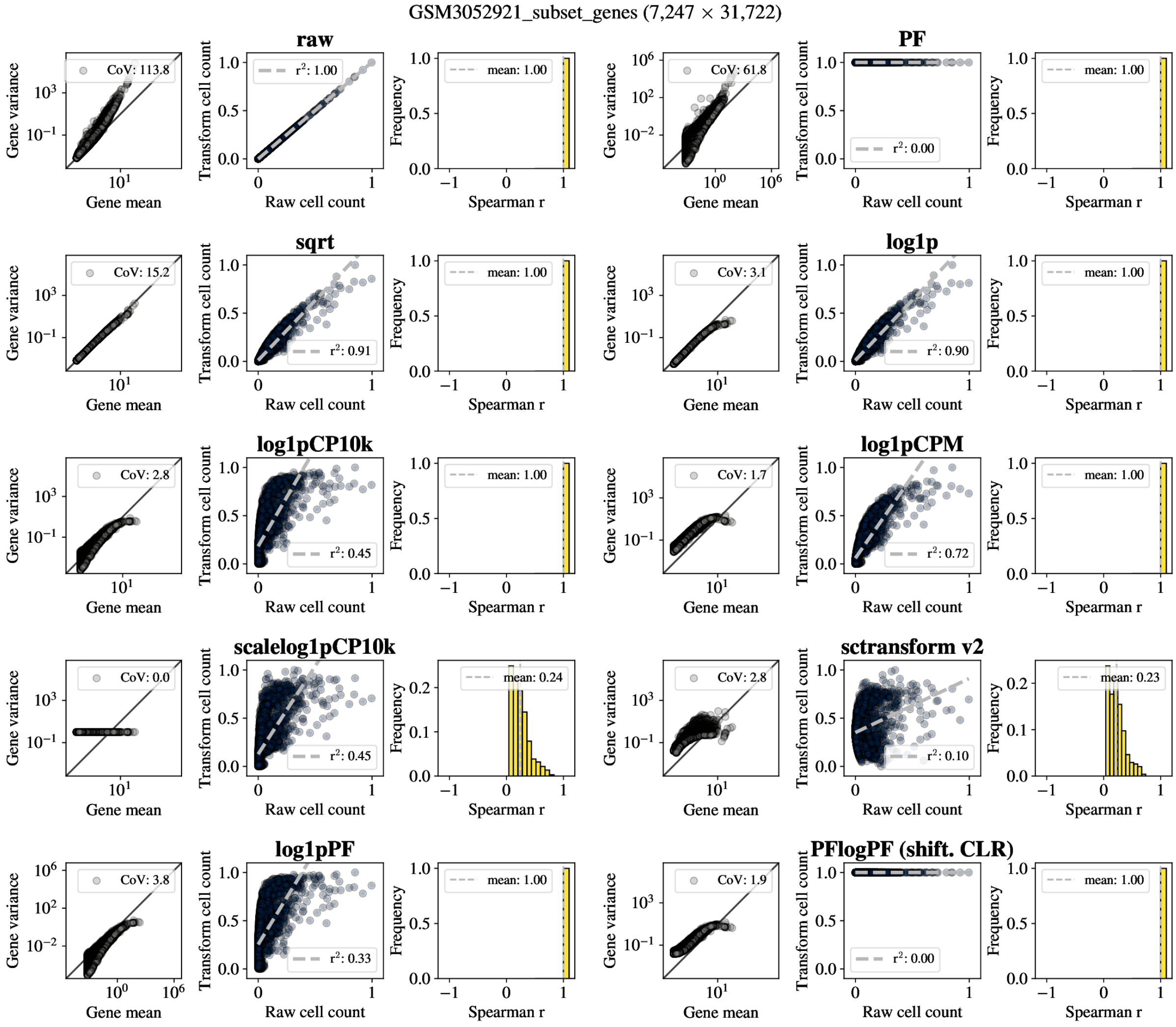


Supplementary Figure 1.167. **GSM3052921**. Per-method comparison of variance stabilization (left), depth normalization (middle), and monotonicity (right) for each normalization on dataset GSM3052921.

| GSM3073088 | |
|---|---|
| cells | 6,799 |
| genes | 60,623 |
| nonzero entries | 10,690,488 |
| density | 0.0259 |
| total counts | 33,955,911 |
| mean counts / cell | 4,994 |
| mean counts / gene | 560 |
| min counts / cell | 61 |
| max counts / cell | 45,736 |
| overdispersion | 0.949 |

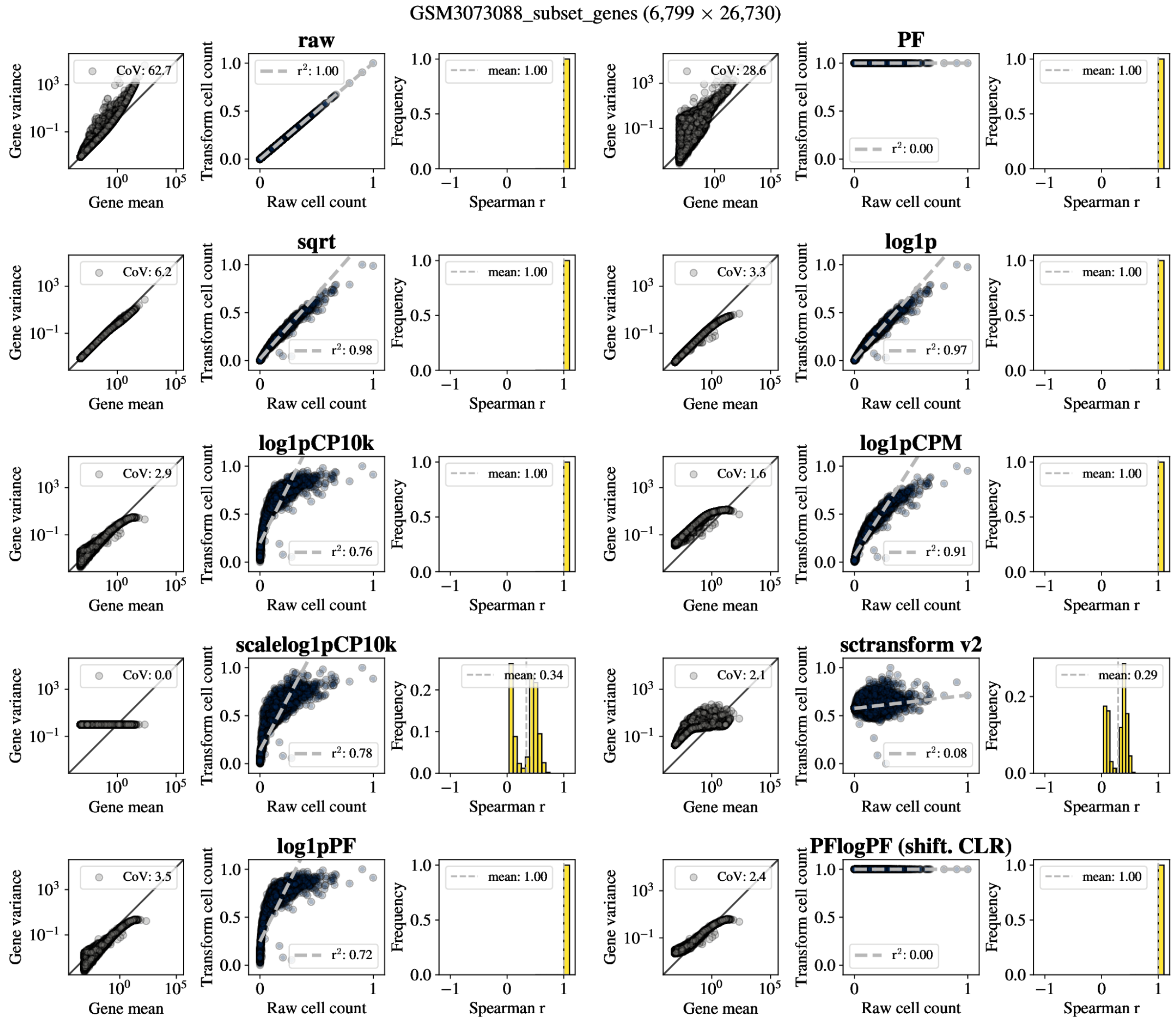


Supplementary Figure 1.168. **GSM3073088**. Per-method comparison of variance stabilization (left), depth normalization (middle), and monotonicity (right) for each normalization on dataset GSM3073088.

| GSM3073089 | |
|---|---|
| cells | 4,552 |
| genes | 60,623 |
| nonzero entries | 9,352,793 |
| density | 0.0339 |
| total counts | 30,139,980 |
| mean counts / cell | 6,621 |
| mean counts / gene | 497 |
| min counts / cell | 81 |
| max counts / cell | 51,543 |
| overdispersion | 0.491 |

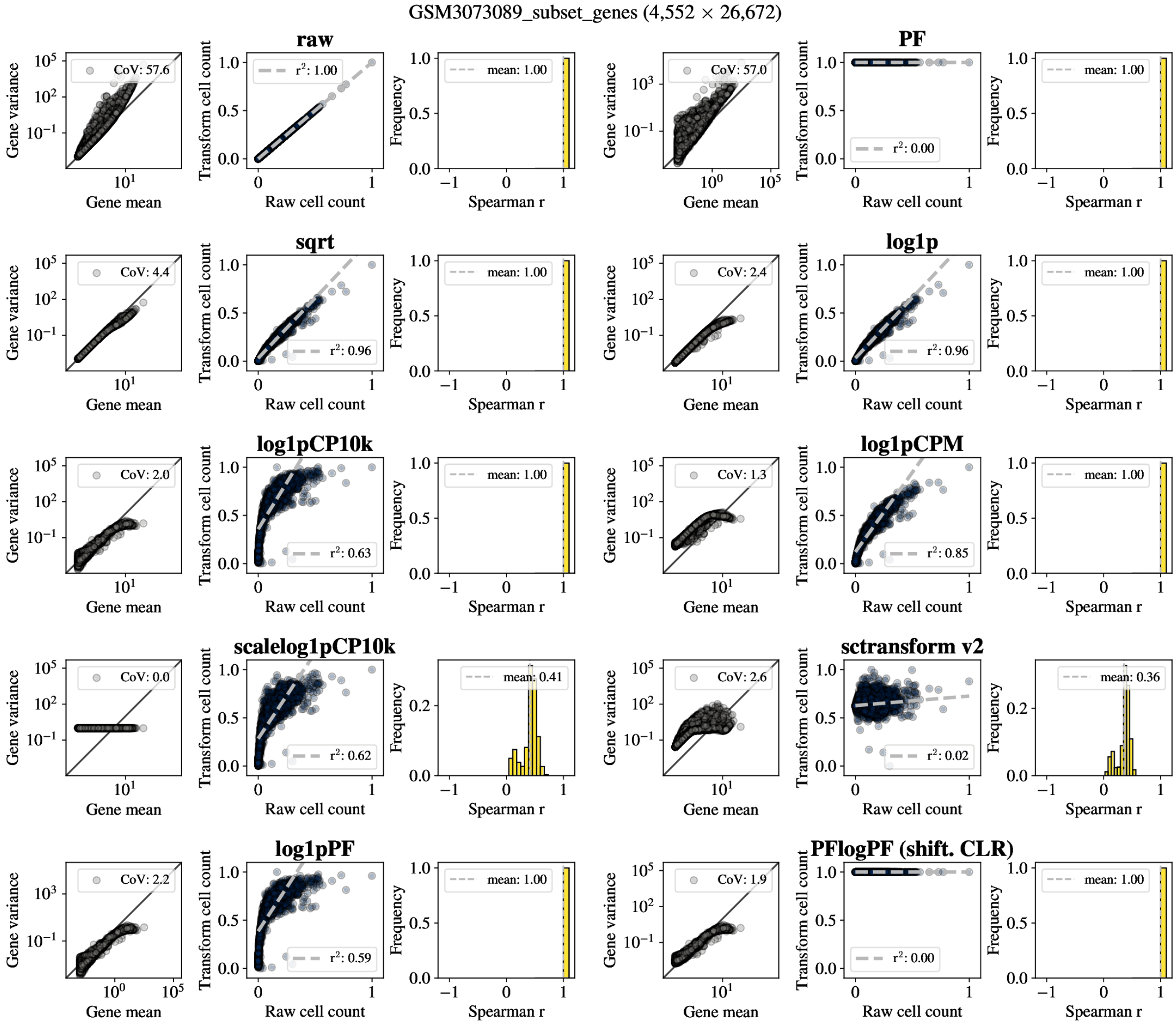


Supplementary Figure 1.169. **GSM3073089**. Per-method comparison of variance stabilization (left), depth normalization (middle), and monotonicity (right) for each normalization on dataset GSM3073089.

| **GSM3143601** | |
|---|---|
| cells | 10,711 |
| genes | 60,623 |
| nonzero entries | 19,046,880 |
| density | 0.0293 |
| total counts | 45,057,512 |
| mean counts / cell | 4,207 |
| mean counts / gene | 743 |
| min counts / cell | 807 |
| max counts / cell | 65,239 |
| overdispersion | 0.596 |

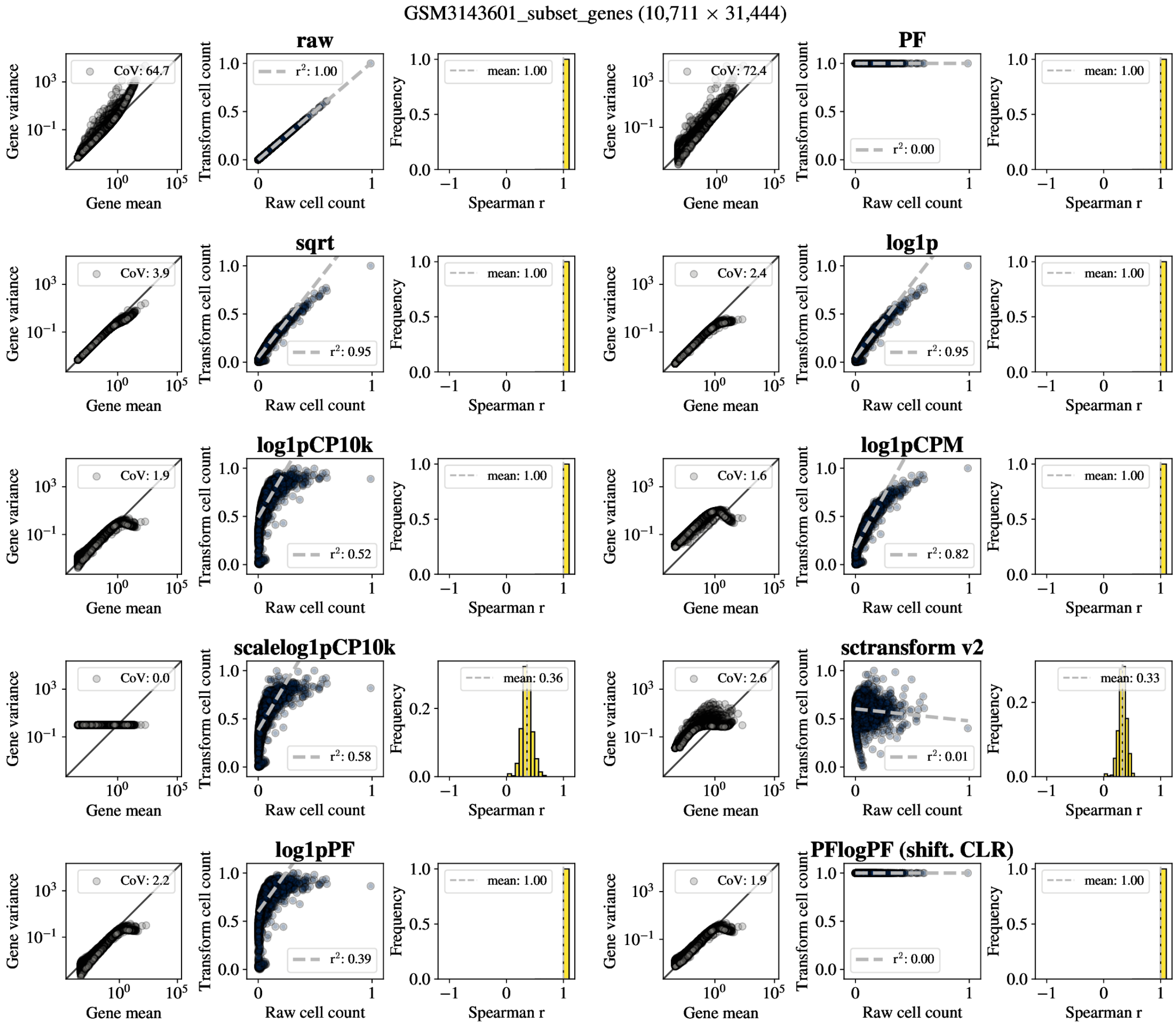


Supplementary Figure 1.170. **GSM3143601**. Per-method comparison of variance stabilization (left), depth normalization (middle), and monotonicity (right) for each normalization on dataset GSM3143601.

| GSM3148575 | |
|---|---|
| cells | 1,673 |
| genes | 60,623 |
| nonzero entries | 515,600 |
| density | 0.00508 |
| total counts | 1,300,981 |
| mean counts / cell | 778 |
| mean counts / gene | 21 |
| min counts / cell | 539 |
| max counts / cell | 3,742 |
| overdispersion | 1.56 |

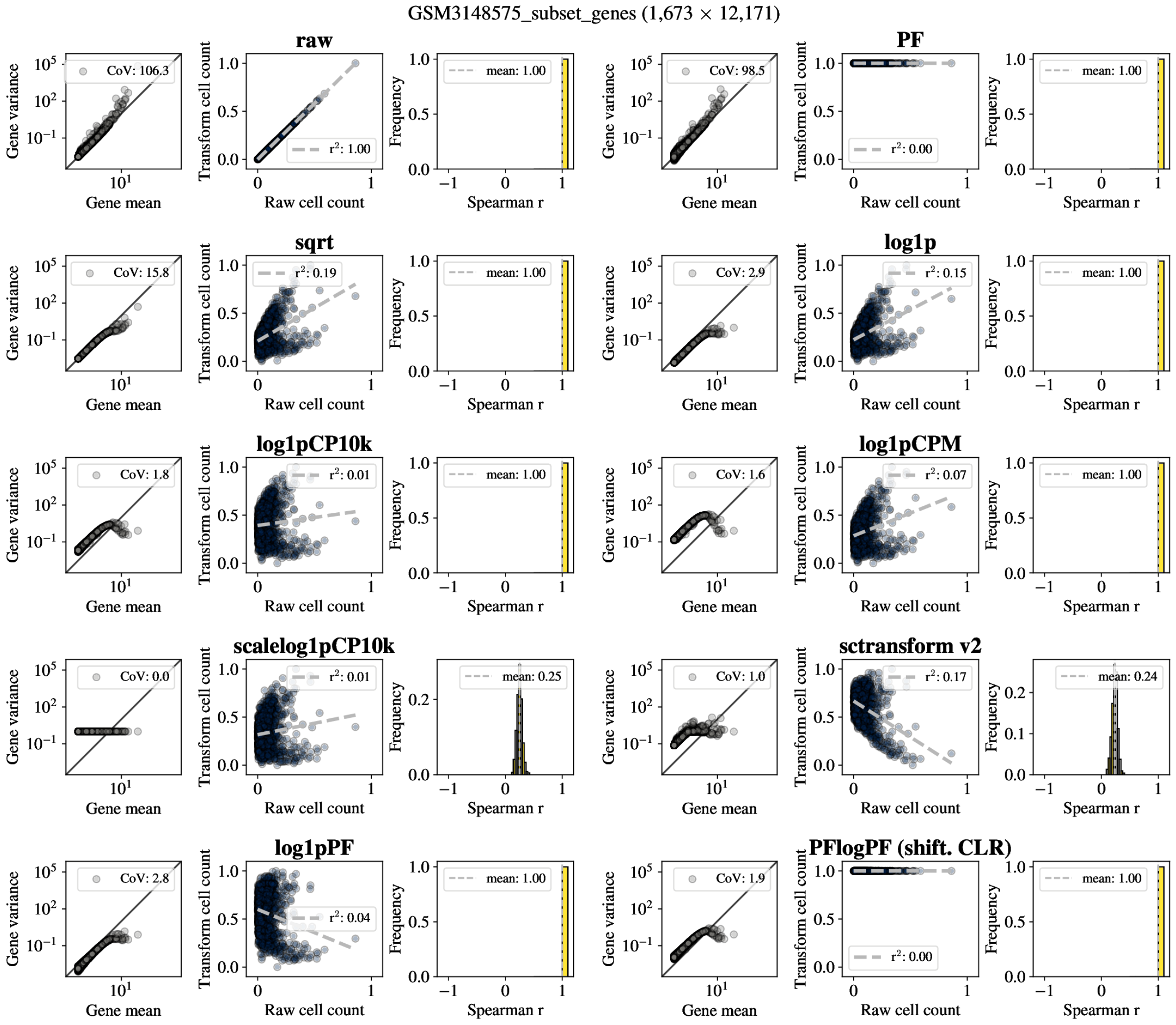


Supplementary Figure 1.171. **GSM3148575**. Per-method comparison of variance stabilization (left), depth normalization (middle), and monotonicity (right) for each normalization on dataset GSM3148575.

| GSM3148576 | |
|---|---|
| cells | 1,517 |
| genes | 60,623 |
| nonzero entries | 505,884 |
| density | 0.0055 |
| total counts | 1,235,306 |
| mean counts / cell | 814 |
| mean counts / gene | 20 |
| min counts / cell | 563 |
| max counts / cell | 2,885 |
| overdispersion | 1.57 |

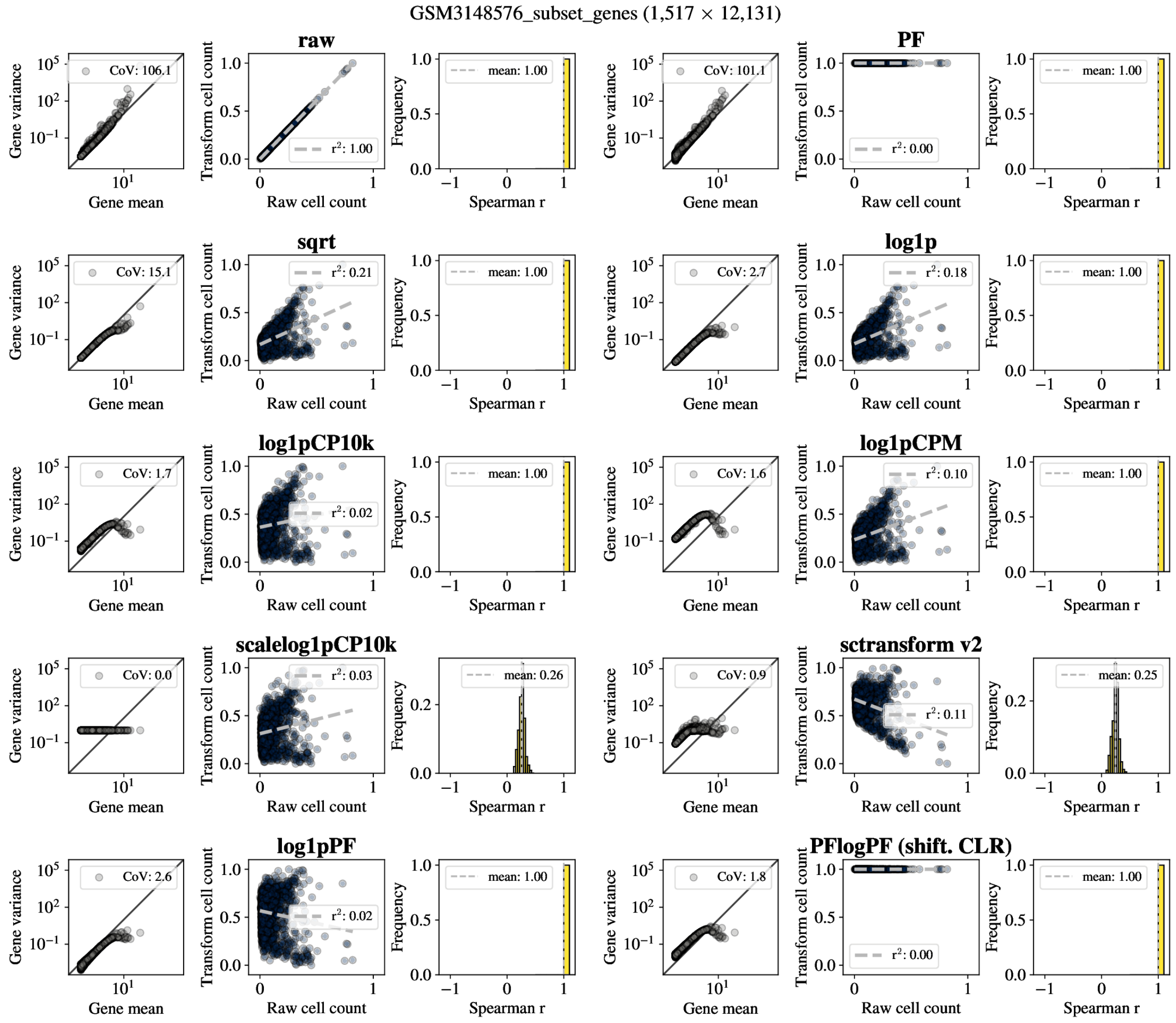


Supplementary Figure 1.172. **GSM3148576**. Per-method comparison of variance stabilization (left), depth normalization (middle), and monotonicity (right) for each normalization on dataset GSM3148576.

| GSM3148577 | |
|---|---|
| cells | 992 |
| genes | 60,623 |
| nonzero entries | 475,847 |
| density | 0.00791 |
| total counts | 1,138,530 |
| mean counts / cell | 1,148 |
| mean counts / gene | 19 |
| min counts / cell | 696 |
| max counts / cell | 6,505 |
| overdispersion | 1.81 |

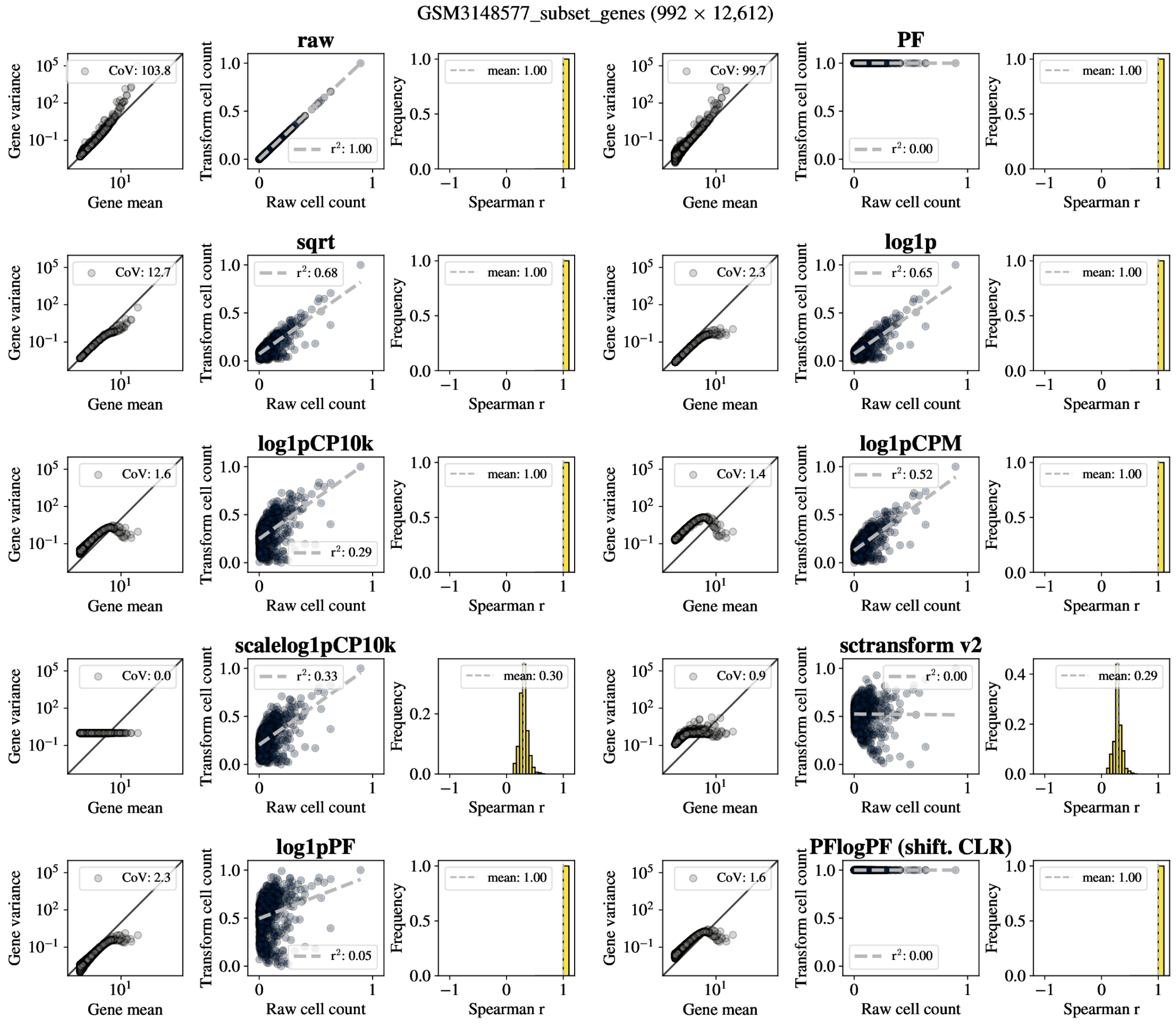


Supplementary Figure 1.173. **GSM3148577**. Per-method comparison of variance stabilization (left), depth normalization (middle), and monotonicity (right) for each normalization on dataset GSM3148577.

| **GSM3148578** | |
|---|---|
| cells | 1,378 |
| genes | 60,623 |
| nonzero entries | 592,378 |
| density | 0.00709 |
| total counts | 1,241,839 |
| mean counts / cell | 901 |
| mean counts / gene | 20 |
| min counts / cell | 596 |
| max counts / cell | 6,268 |
| overdispersion | 2.07 |

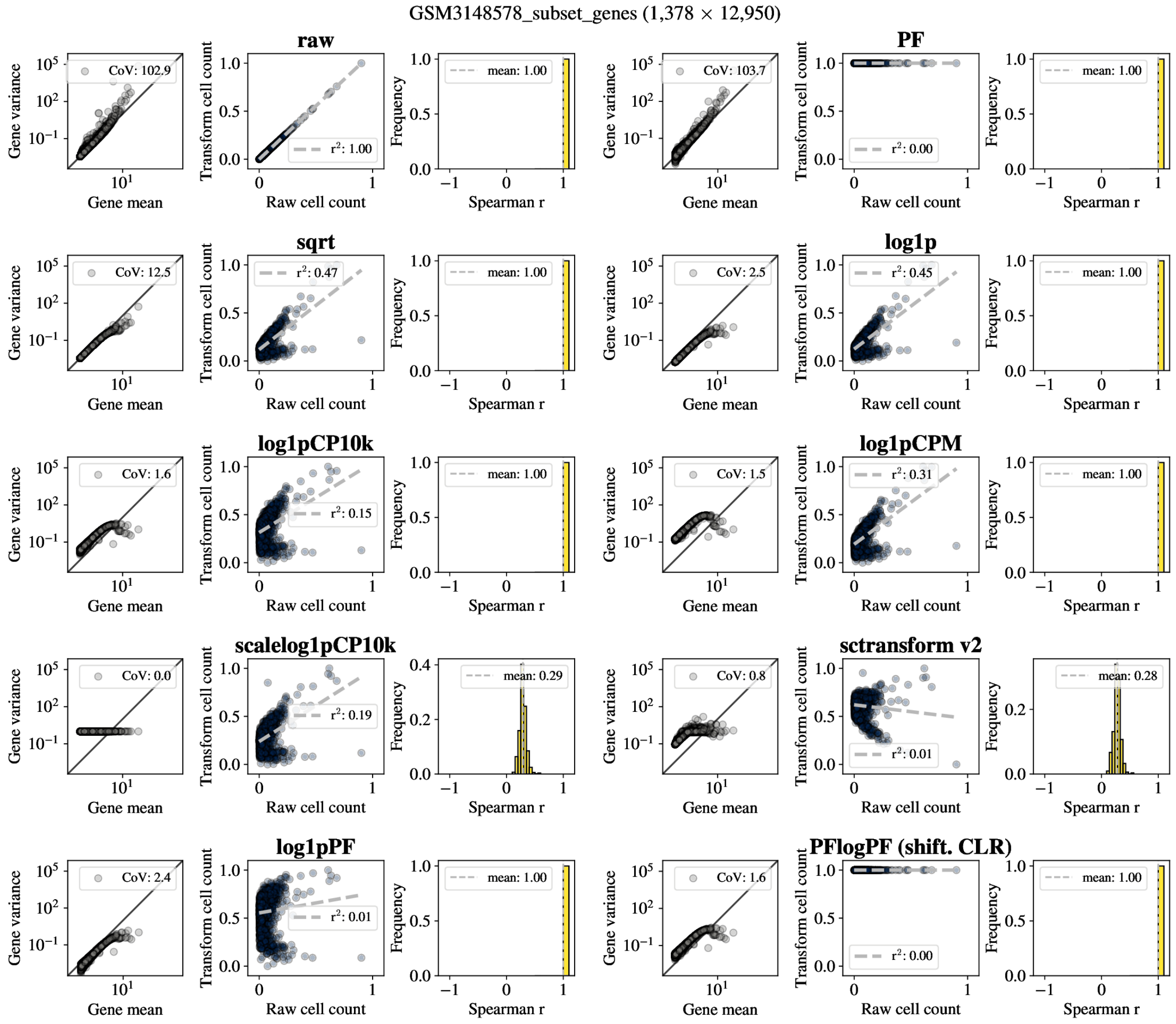


Supplementary Figure 1.174. **GSM3148578**. Per-method comparison of variance stabilization (left), depth normalization (middle), and monotonicity (right) for each normalization on dataset GSM3148578.

| GSM3148579 | |
|---|---|
| cells | 1,290 |
| genes | 60,623 |
| nonzero entries | 569,905 |
| density | 0.00729 |
| total counts | 1,205,273 |
| mean counts / cell | 934 |
| mean counts / gene | 20 |
| min counts / cell | 614 |
| max counts / cell | 4,831 |
| overdispersion | 2.27 |

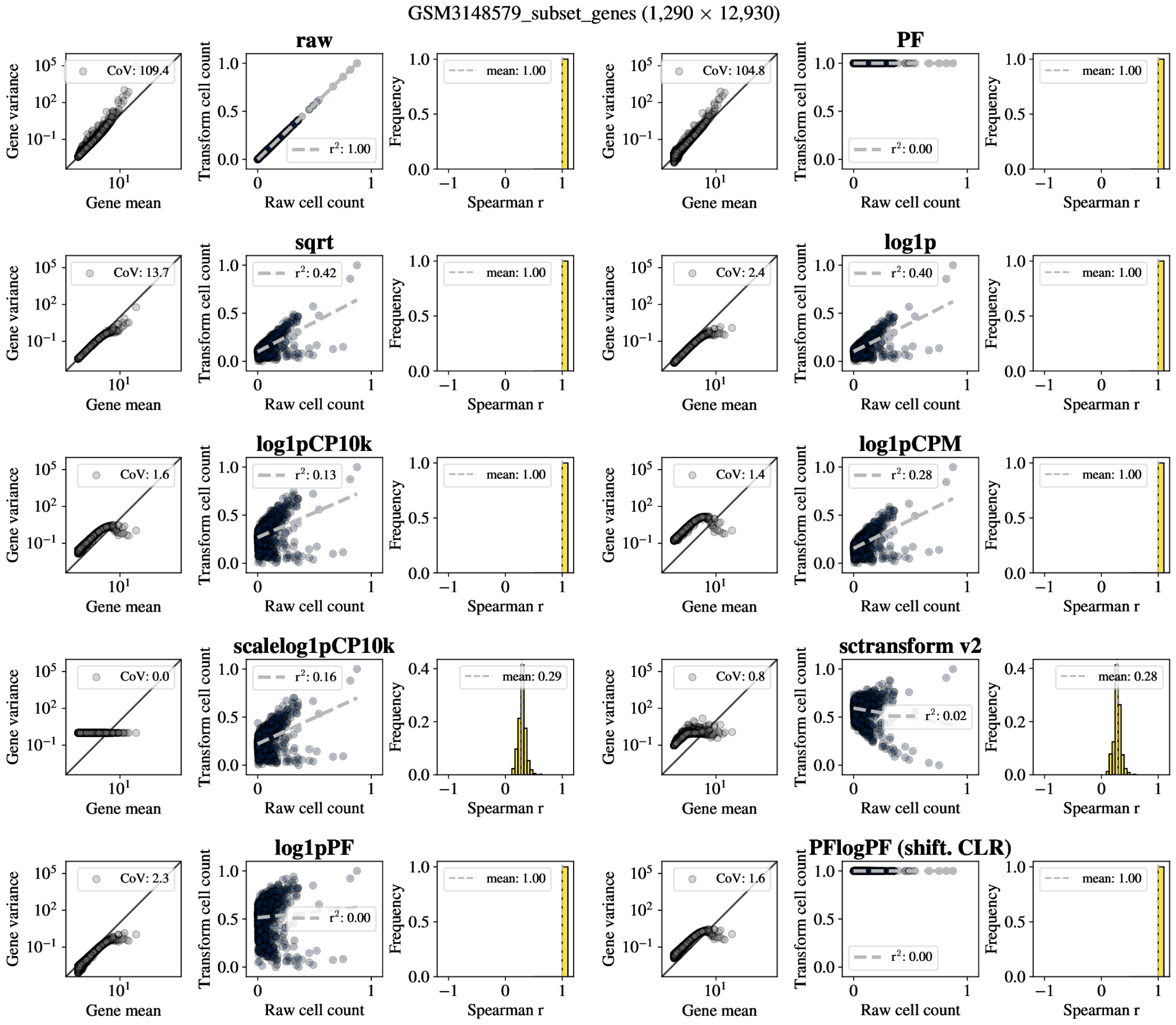


Supplementary Figure 1.175. **GSM3148579**. Per-method comparison of variance stabilization (left), depth normalization (middle), and monotonicity (right) for each normalization on dataset GSM3148579.

| GSM3168248 | |
|---|---|
| cells | 8,333 |
| genes | 60,623 |
| nonzero entries | 10,980,622 |
| density | 0.0217 |
| total counts | 32,106,483 |
| mean counts / cell | 3,853 |
| mean counts / gene | 530 |
| min counts / cell | 645 |
| max counts / cell | 56,428 |
| overdispersion | 1.71 |

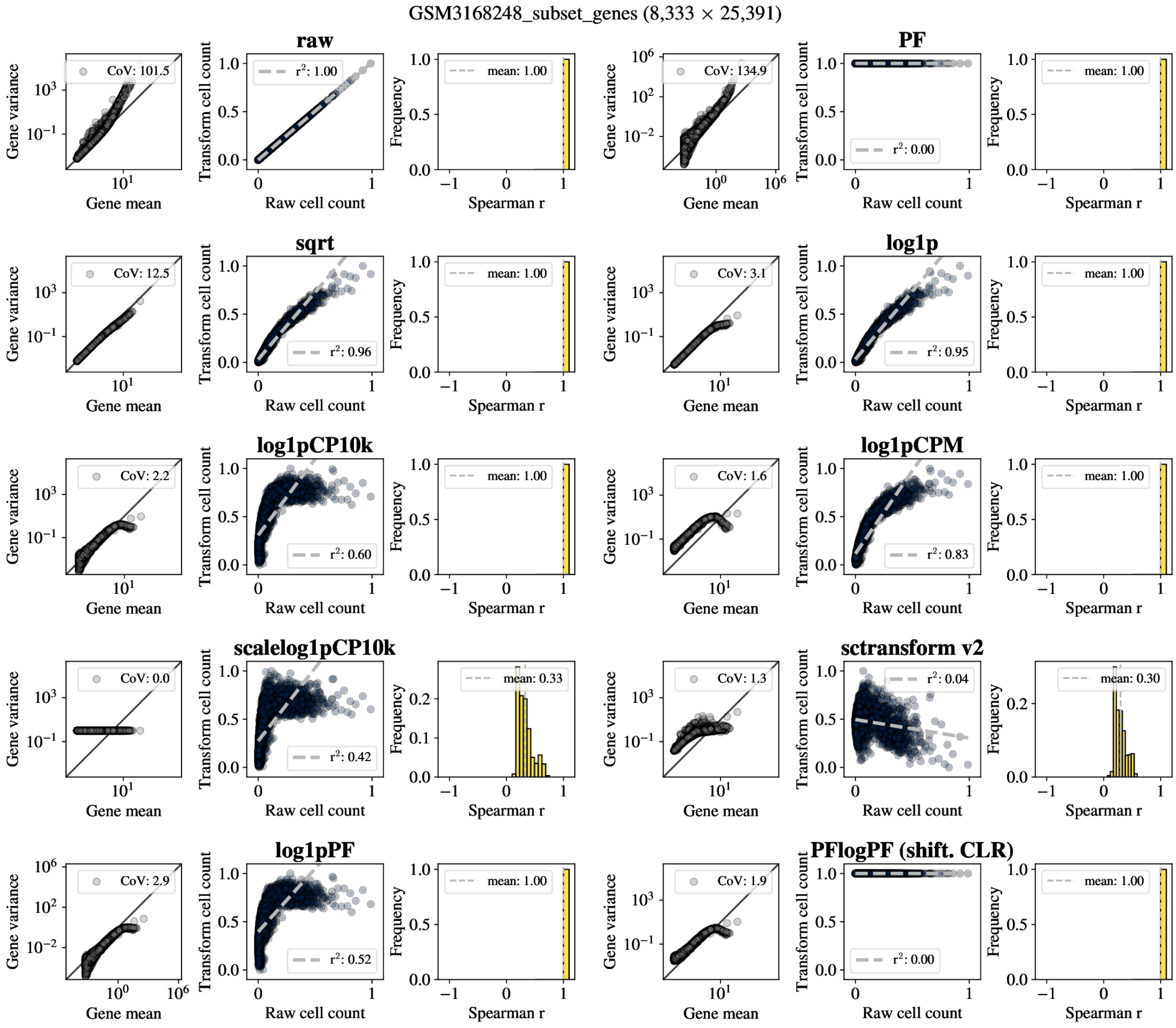


Supplementary Figure 1.176. **GSM3168248**. Per-method comparison of variance stabilization (left), depth normalization (middle), and monotonicity (right) for each normalization on dataset GSM3168248.

| **GSM3169075** | |
|---|---|
| cells | 2,949 |
| genes | 60,623 |
| nonzero entries | 3,741,857 |
| density | 0.0209 |
| total counts | 10,573,288 |
| mean counts / cell | 3,585 |
| mean counts / gene | 174 |
| min counts / cell | 1,877 |
| max counts / cell | 26,263 |
| overdispersion | 0.148 |

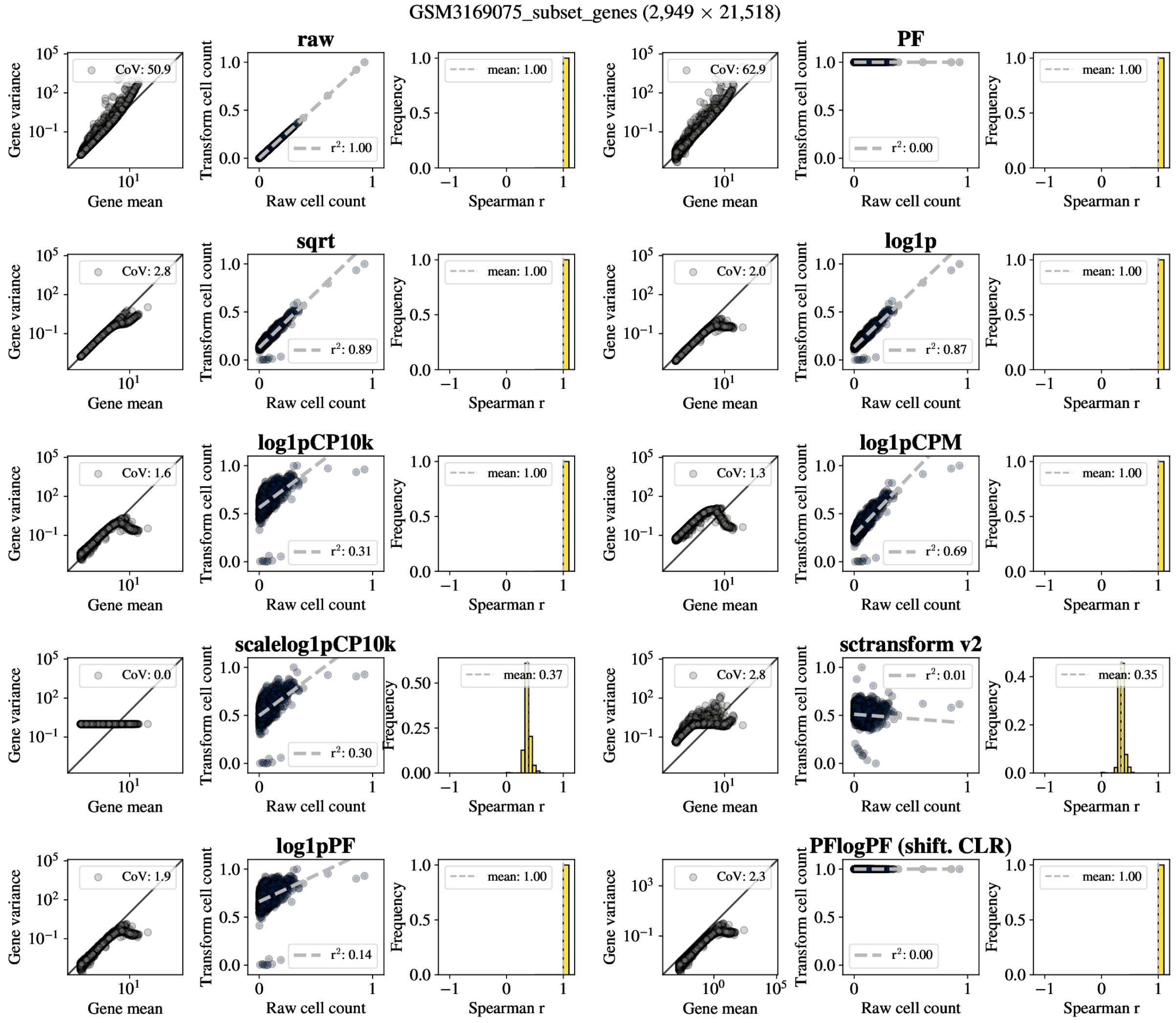


Supplementary Figure 1.177. **GSM3169075**. Per-method comparison of variance stabilization (left), depth normalization (middle), and monotonicity (right) for each normalization on dataset GSM3169075.

| GSM3178782 | |
|---|---|
| cells | 15,841 |
| genes | 60,623 |
| nonzero entries | 3,498,952 |
| density | 0.00364 |
| total counts | 21,539,679 |
| mean counts / cell | 1,360 |
| mean counts / gene | 355 |
| min counts / cell | 192 |
| max counts / cell | 62,248 |
| overdispersion | 3.44 |

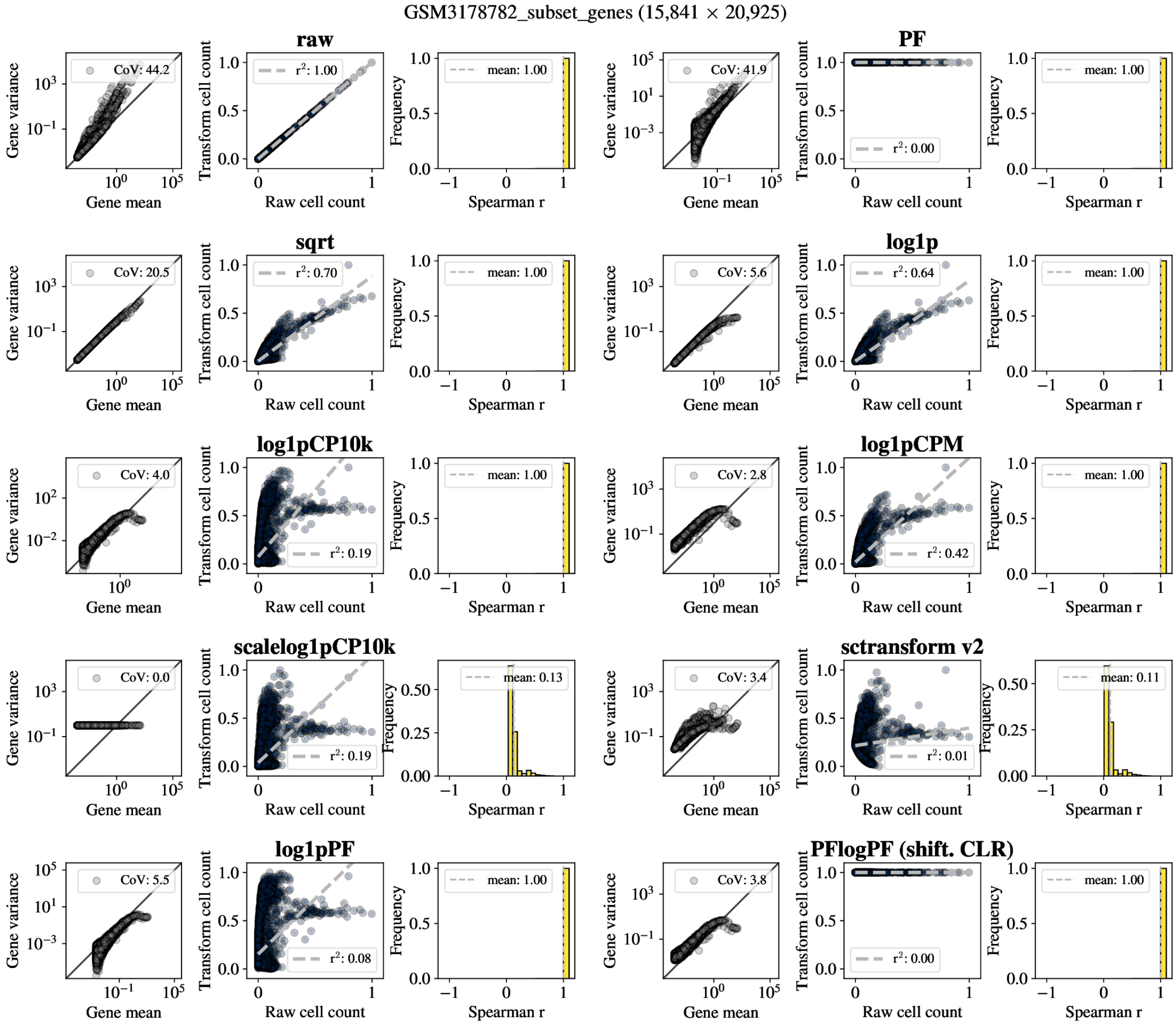


Supplementary Figure 1.178. **GSM3178782**. Per-method comparison of variance stabilization (left), depth normalization (middle), and monotonicity (right) for each normalization on dataset GSM3178782.

| GSM3178783 | |
|---|---|
| cells | 71,720 |
| genes | 60,623 |
| nonzero entries | 14,954,743 |
| density | 0.00344 |
| total counts | 65,293,177 |
| mean counts / cell | 910 |
| mean counts / gene | 1,077 |
| min counts / cell | 332 |
| max counts / cell | 12,414 |
| overdispersion | 0.166 |

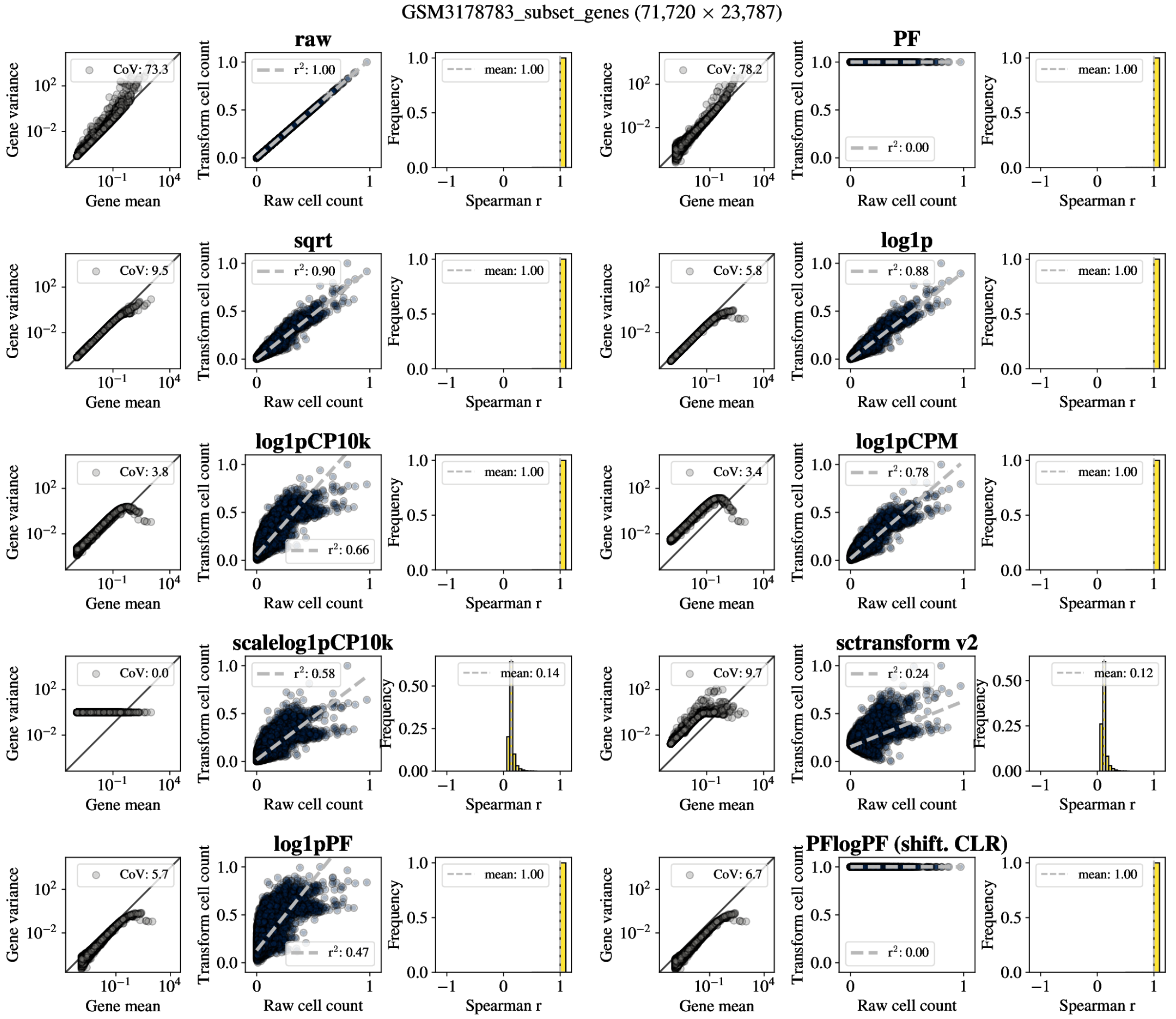


Supplementary Figure 1.179. **GSM3178783**. Per-method comparison of variance stabilization (left), depth normalization (middle), and monotonicity (right) for each normalization on dataset GSM3178783.

| GSM3178784 | |
|---|---|
| cells | 17,262 |
| genes | 60,623 |
| nonzero entries | 7,744,060 |
| density | 0.0074 |
| total counts | 34,731,963 |
| mean counts / cell | 2,012 |
| mean counts / gene | 573 |
| min counts / cell | 389 |
| max counts / cell | 26,520 |
| overdispersion | 0.842 |

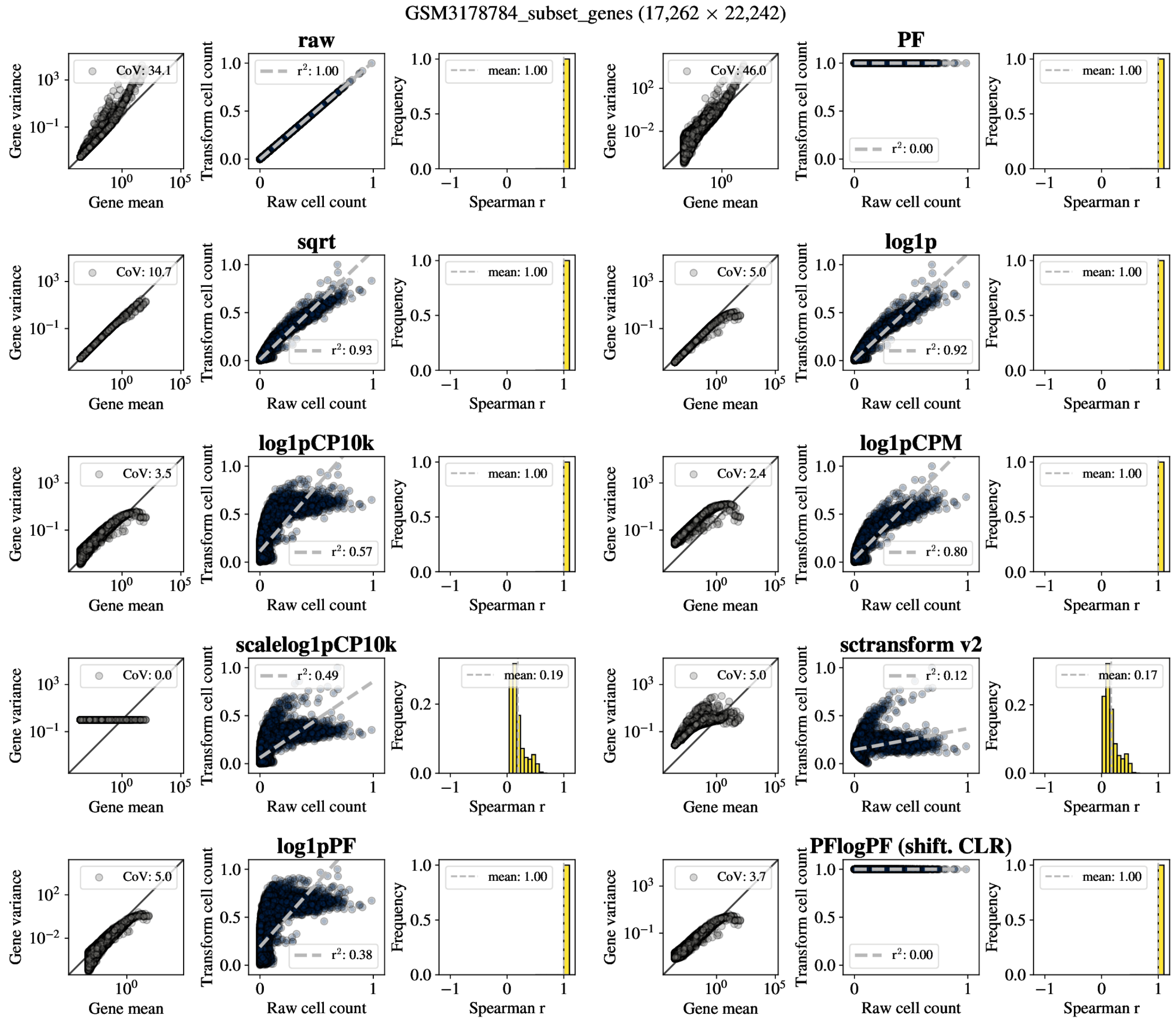


Supplementary Figure 1.180. **GSM3178784**. Per-method comparison of variance stabilization (left), depth normalization (middle), and monotonicity (right) for each normalization on dataset GSM3178784.

| GSM3178785 | |
|---|---|
| cells | 6,319 |
| genes | 60,623 |
| nonzero entries | 2,989,664 |
| density | 0.0078 |
| total counts | 15,530,916 |
| mean counts / cell | 2,458 |
| mean counts / gene | 256 |
| min counts / cell | 1,049 |
| max counts / cell | 63,358 |
| overdispersion | 0.413 |

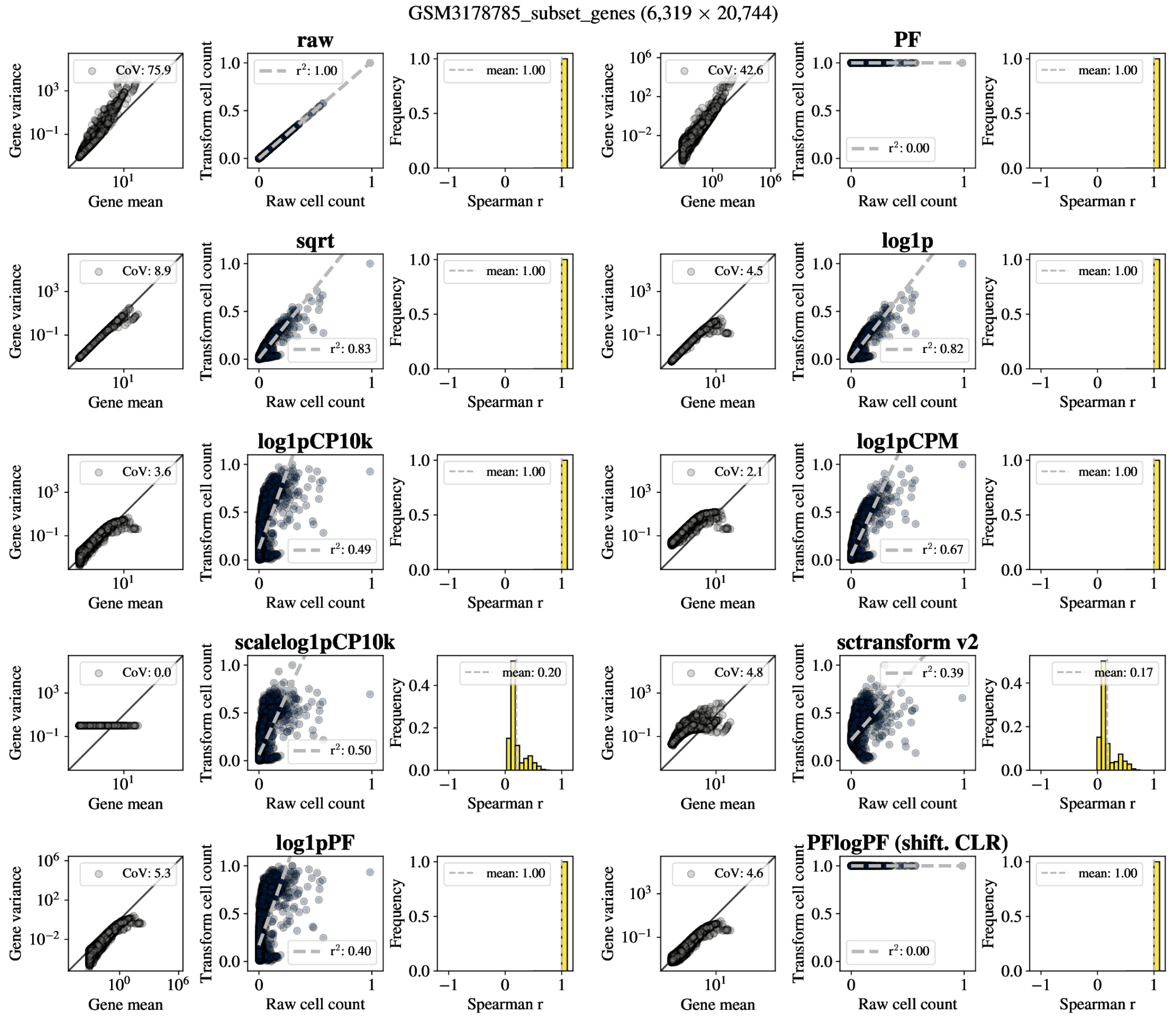


Supplementary Figure 1.181. **GSM3178785**. Per-method comparison of variance stabilization (left), depth normalization (middle), and monotonicity (right) for each normalization on dataset GSM3178785.

| GSM3178786 | |
|---|---|
| cells | 8,609 |
| genes | 60,623 |
| nonzero entries | 3,755,218 |
| density | 0.0072 |
| total counts | 23,635,956 |
| mean counts / cell | 2,745 |
| mean counts / gene | 390 |
| min counts / cell | 627 |
| max counts / cell | 58,068 |
| overdispersion | 0.894 |

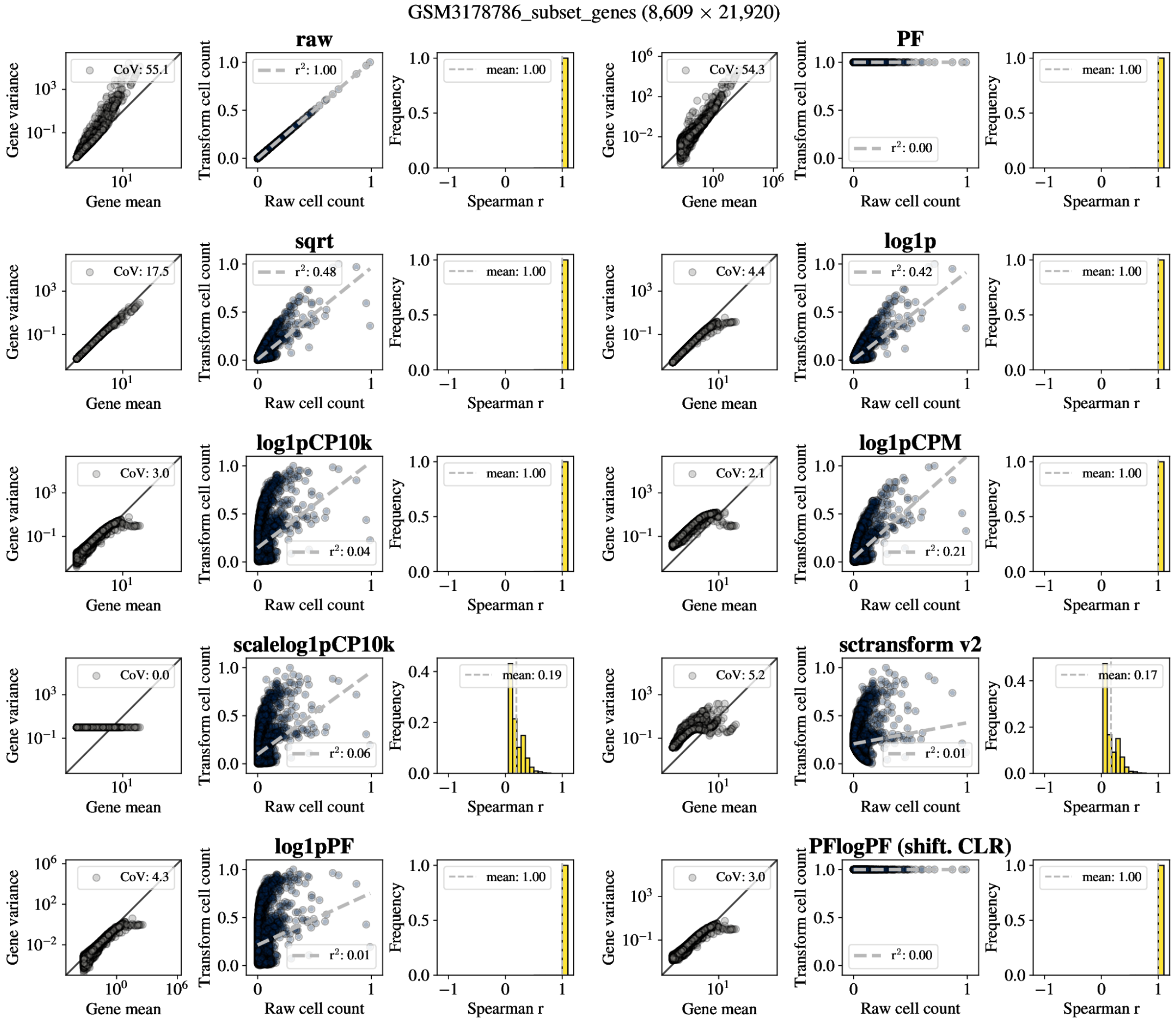


Supplementary Figure 1.182. **GSM3178786**. Per-method comparison of variance stabilization (left), depth normalization (middle), and monotonicity (right) for each normalization on dataset GSM3178786.

| GSM3273570 | |
|---|---|
| cells | 3,244 |
| genes | 60,623 |
| nonzero entries | 12,194,113 |
| density | 0.062 |
| total counts | 52,705,676 |
| mean counts / cell | 16,247 |
| mean counts / gene | 869 |
| min counts / cell | 2,615 |
| max counts / cell | 133,753 |
| overdispersion | 0.776 |

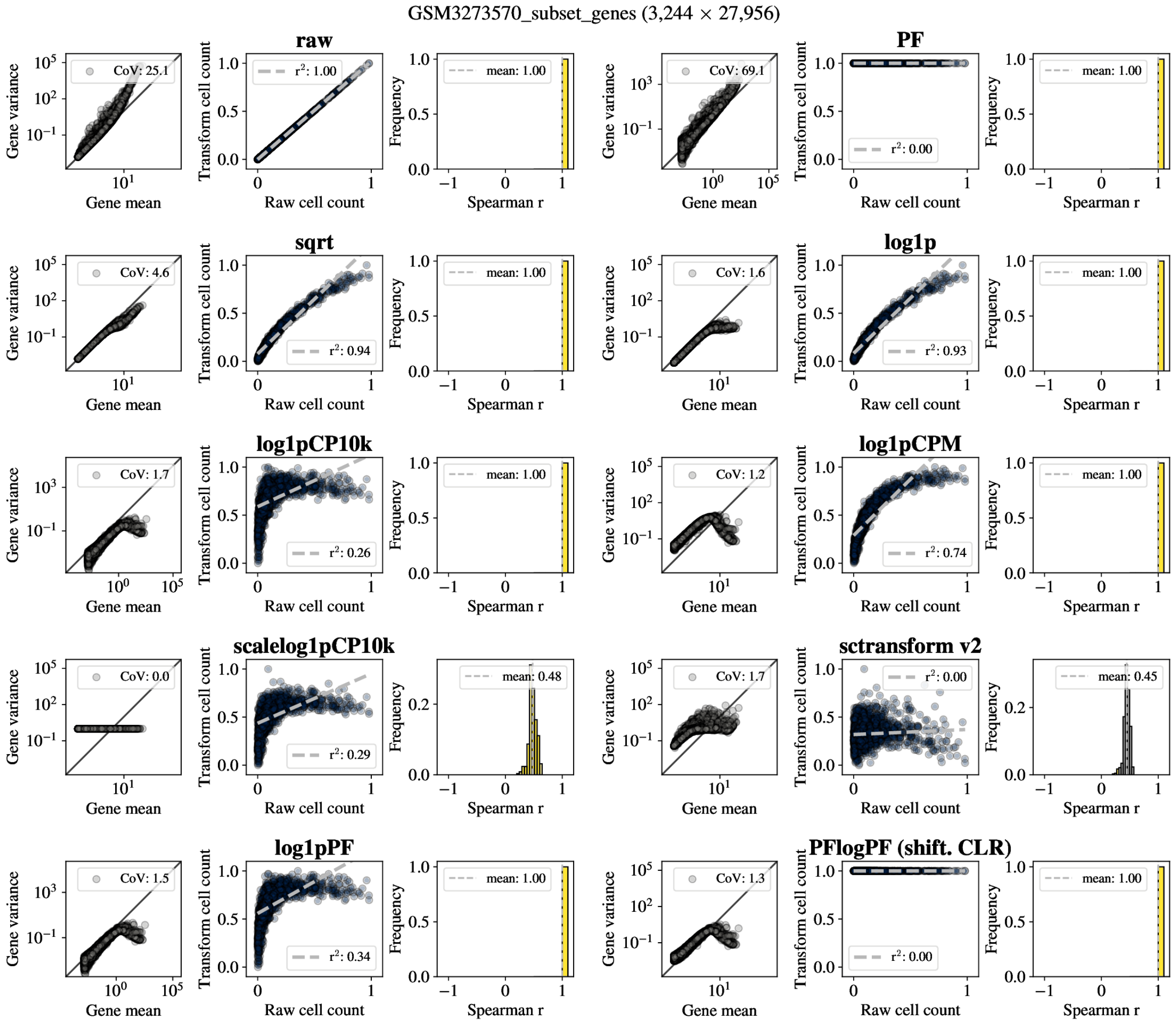


Supplementary Figure 1.183. **GSM3273570**. Per-method comparison of variance stabilization (left), depth normalization (middle), and monotonicity (right) for each normalization on dataset GSM3273570.

| GSM3302524 | |
|---|---|
| cells | 13,945 |
| genes | 60,623 |
| nonzero entries | 16,788,759 |
| density | 0.0199 |
| total counts | 55,088,339 |
| mean counts / cell | 3,950 |
| mean counts / gene | 909 |
| min counts / cell | 192 |
| max counts / cell | 94,332 |
| overdispersion | 67.8 |

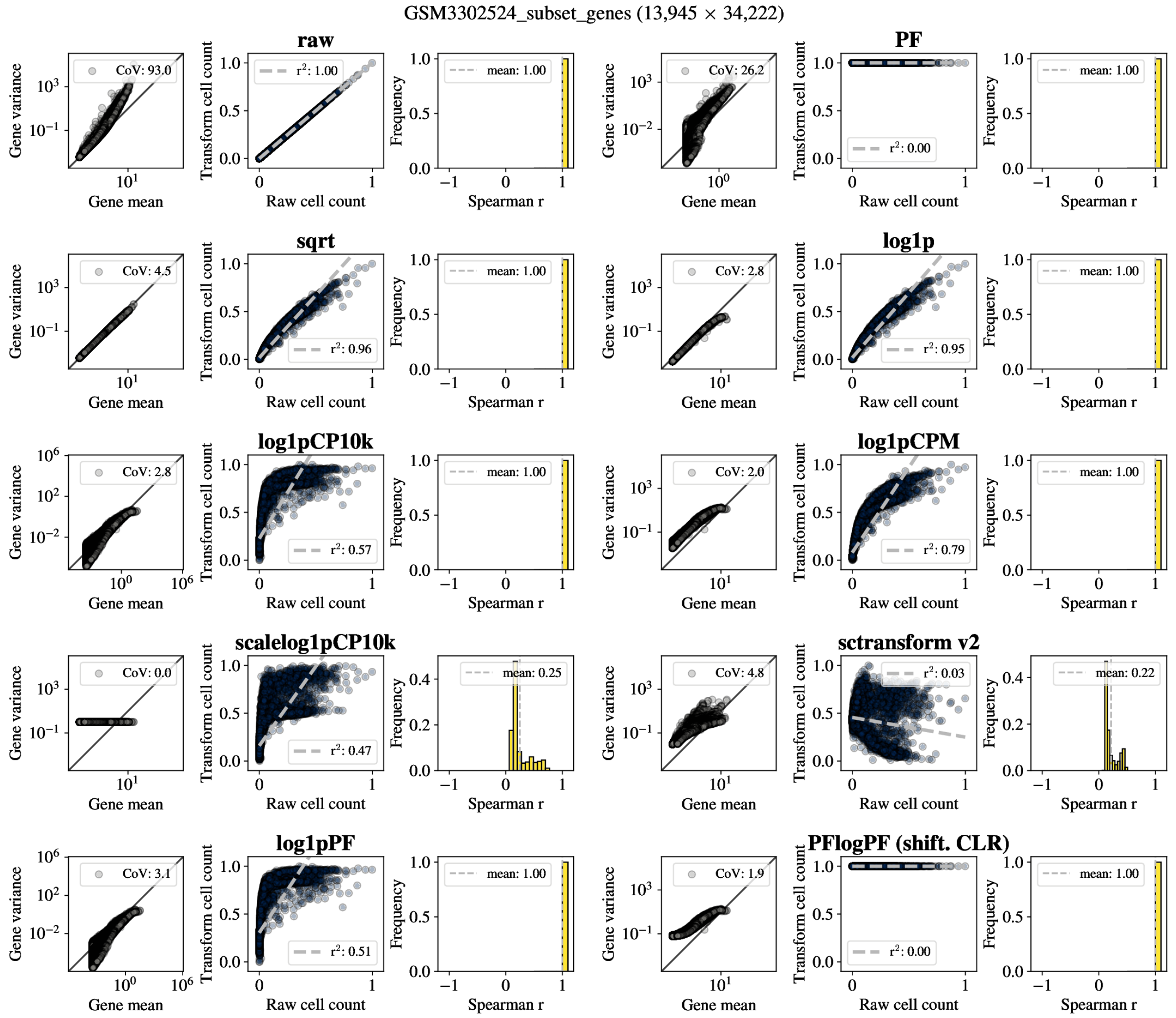


Supplementary Figure 1.184. **GSM3302524**. Per-method comparison of variance stabilization (left), depth normalization (middle), and monotonicity (right) for each normalization on dataset GSM3302524.

| GSM3302525 | |
|---|---|
| cells | 6,066 |
| genes | 60,623 |
| nonzero entries | 18,189,696 |
| density | 0.0495 |
| total counts | 53,113,541 |
| mean counts / cell | 8,756 |
| mean counts / gene | 876 |
| min counts / cell | 291 |
| max counts / cell | 54,589 |
| overdispersion | 5.11 |

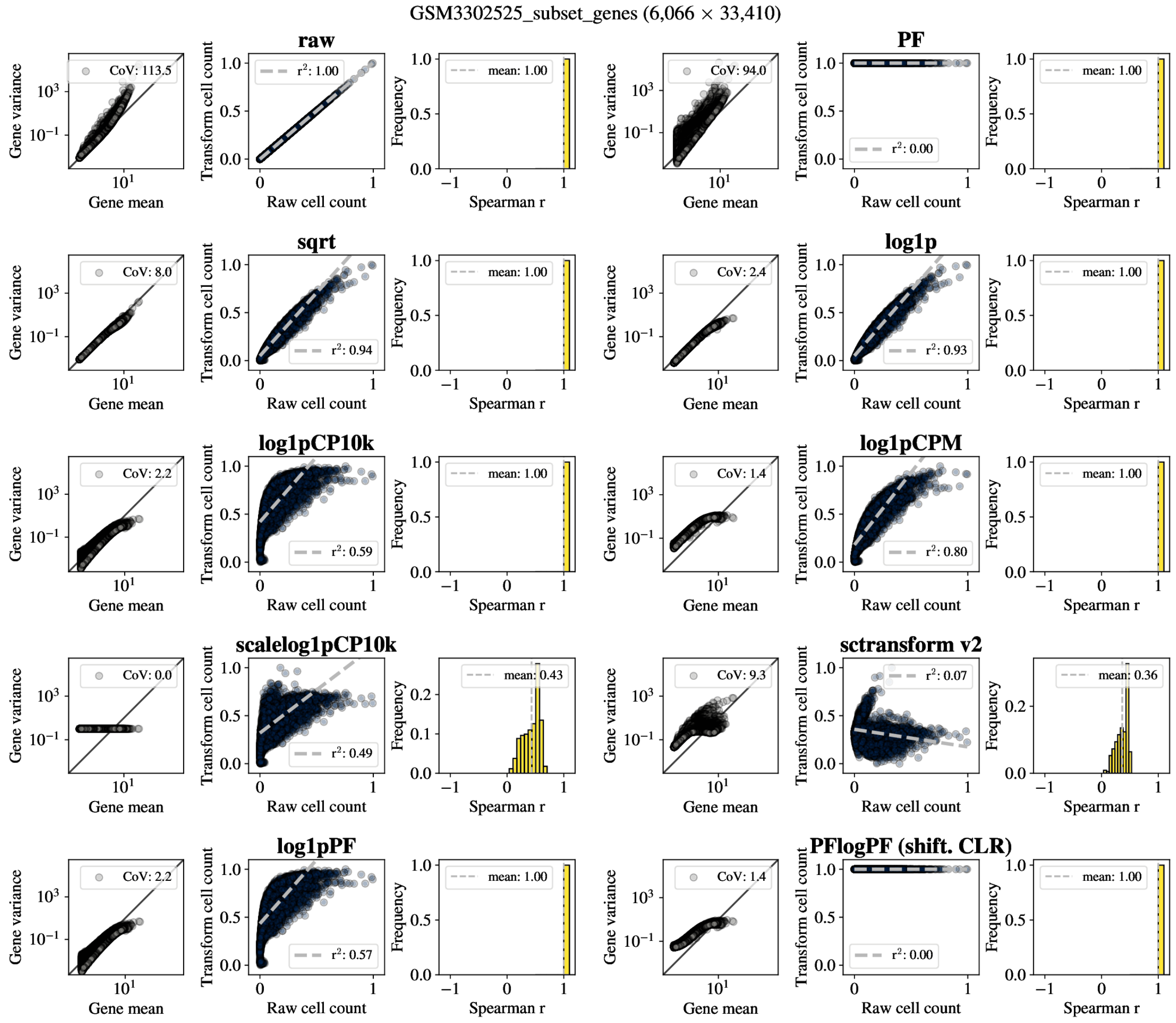


Supplementary Figure 1.185. **GSM3302525**. Per-method comparison of variance stabilization (left), depth normalization (middle), and monotonicity (right) for each normalization on dataset GSM3302525.

| GSM3309832 | |
|---|---|
| cells | 7,236 |
| genes | 60,623 |
| nonzero entries | 11,640,516 |
| density | 0.0265 |
| total counts | 40,836,408 |
| mean counts / cell | 5,644 |
| mean counts / gene | 674 |
| min counts / cell | 1,680 |
| max counts / cell | 28,569 |
| overdispersion | 0.264 |

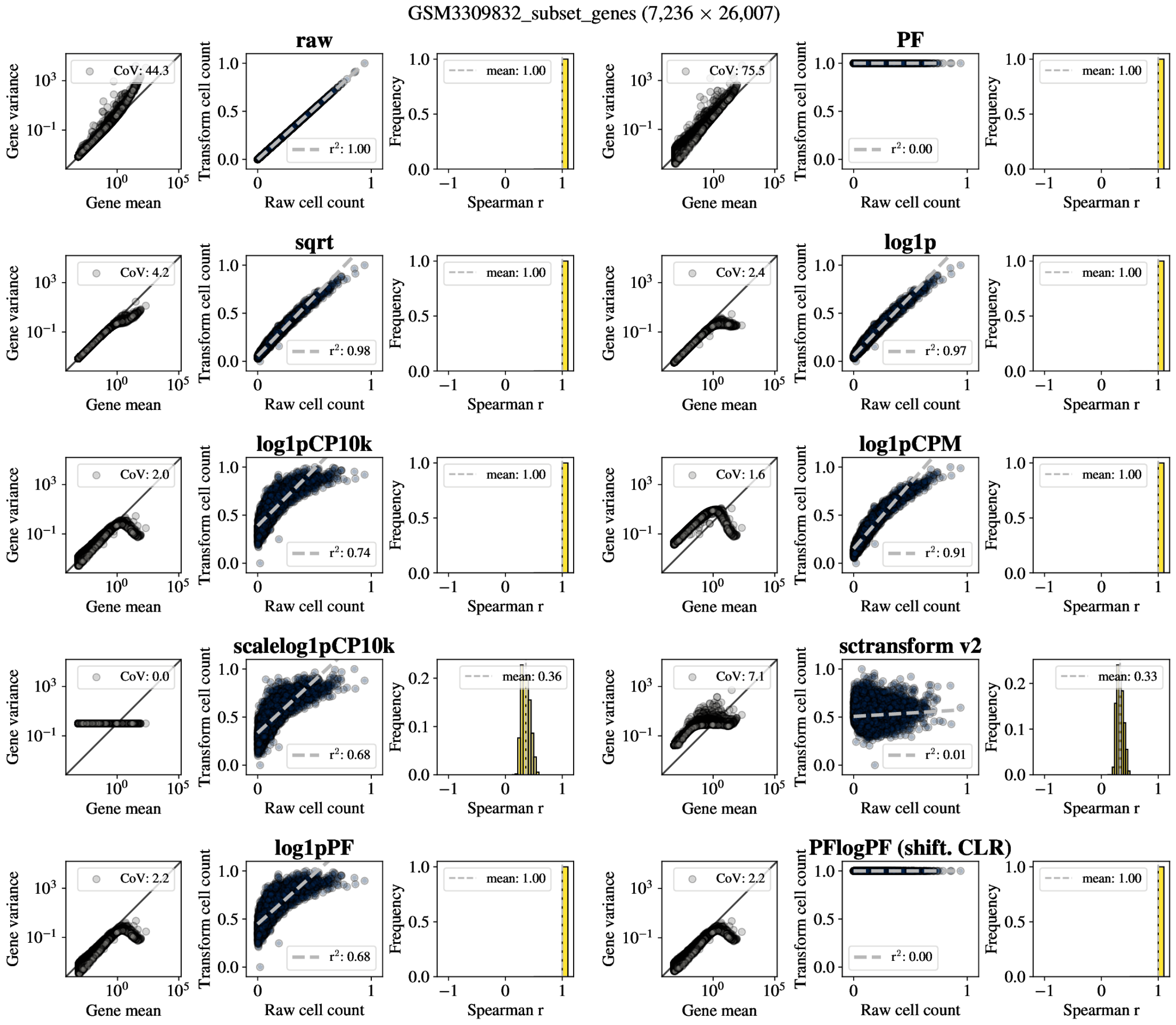


Supplementary Figure 1.186. **GSM3309832**. Per-method comparison of variance stabilization (left), depth normalization (middle), and monotonicity (right) for each normalization on dataset GSM3309832.

| GSM3309833 | |
|---|---|
| cells | 1,760 |
| genes | 60,623 |
| nonzero entries | 4,784,190 |
| density | 0.0448 |
| total counts | 20,615,985 |
| mean counts / cell | 11,714 |
| mean counts / gene | 340 |
| min counts / cell | 708 |
| max counts / cell | 56,022 |
| overdispersion | 0.876 |

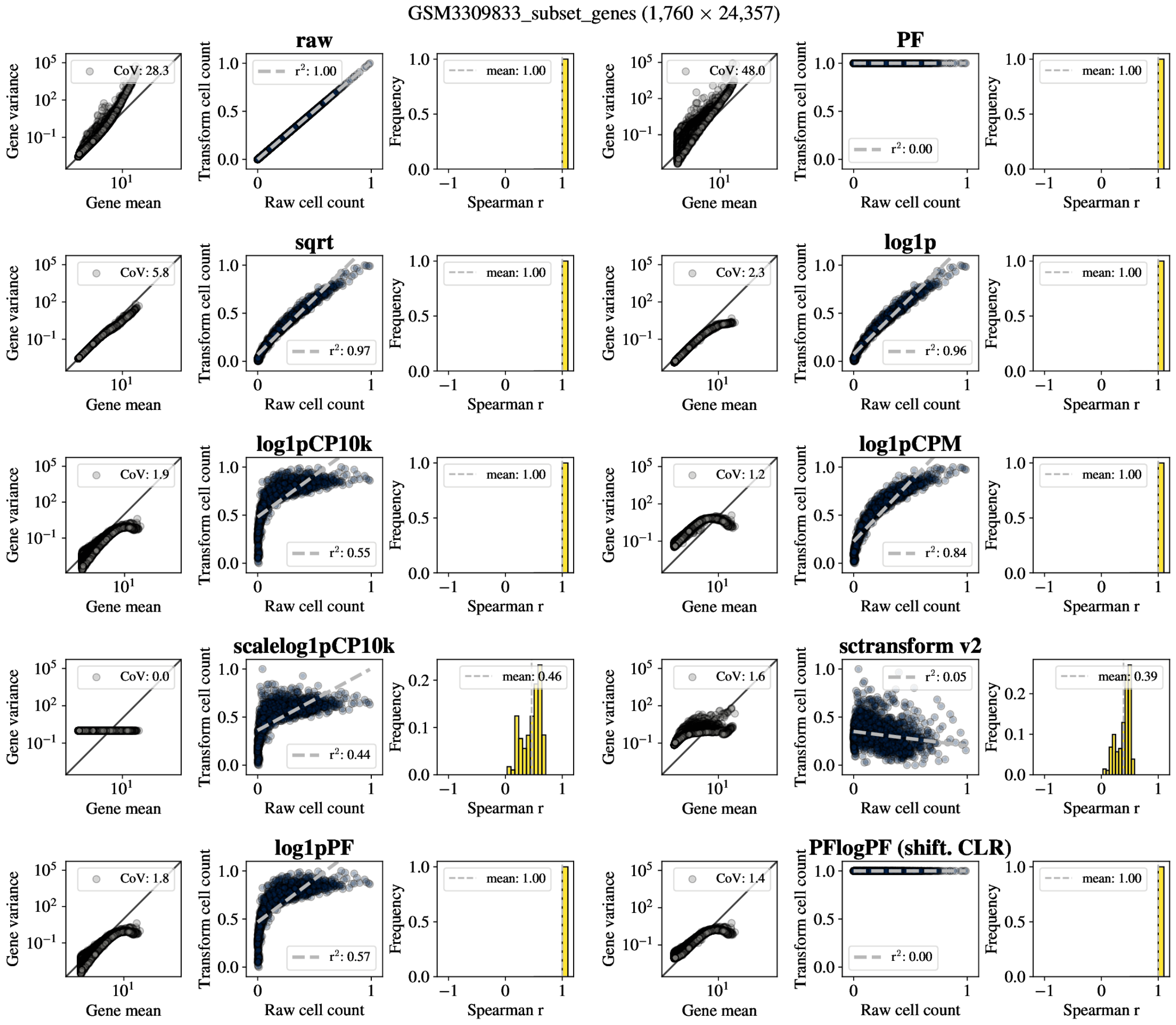


Supplementary Figure 1.187. **GSM3309833**. Per-method comparison of variance stabilization (left), depth normalization (middle), and monotonicity (right) for each normalization on dataset GSM3309833.

| GSM3309834 | |
|---|---|
| cells | 4,805 |
| genes | 60,623 |
| nonzero entries | 11,539,296 |
| density | 0.0396 |
| total counts | 43,688,057 |
| mean counts / cell | 9,092 |
| mean counts / gene | 721 |
| min counts / cell | 148 |
| max counts / cell | 49,262 |
| overdispersion | 0.608 |

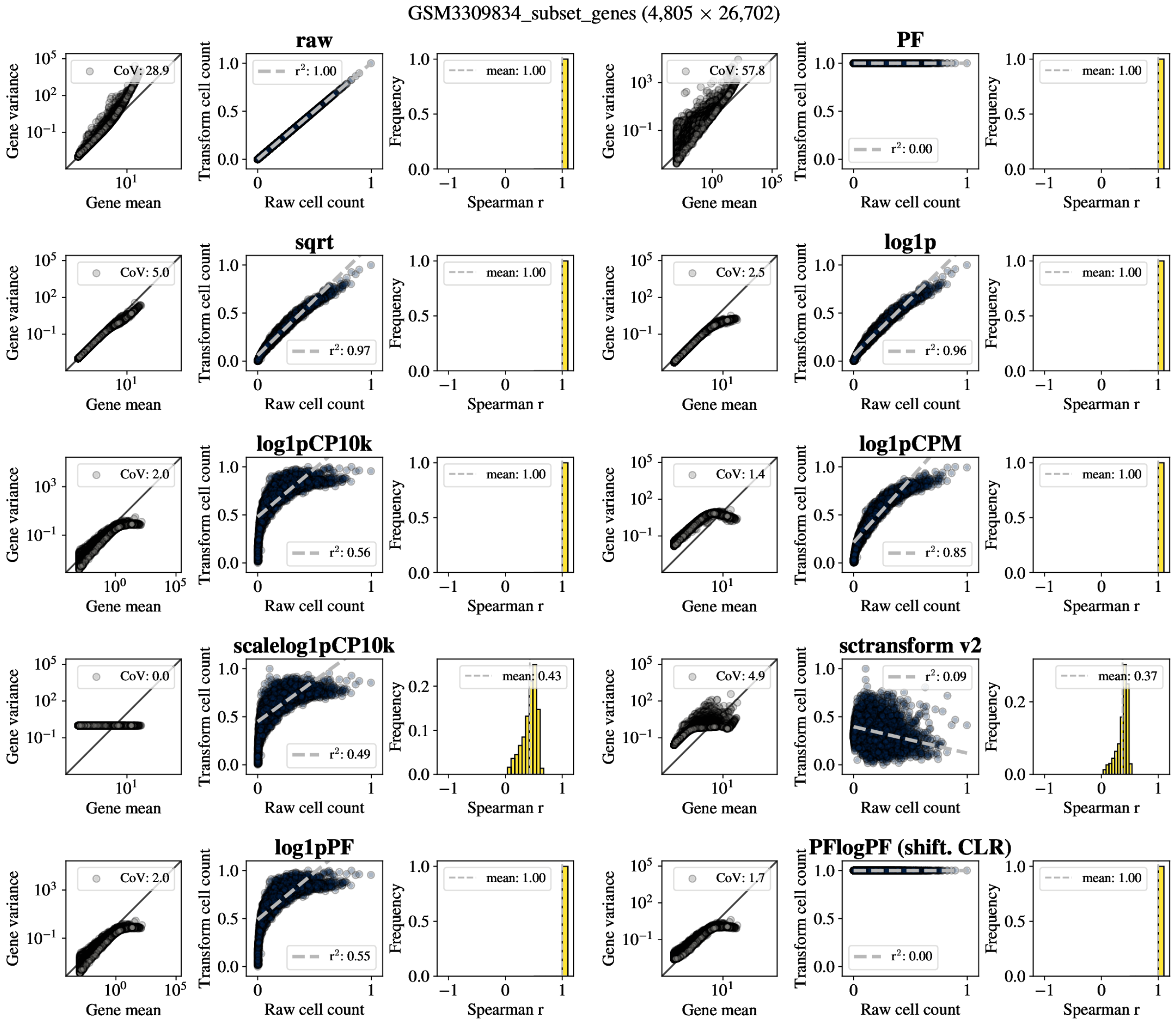


Supplementary Figure 1.188. **GSM3309834**. Per-method comparison of variance stabilization (left), depth normalization (middle), and monotonicity (right) for each normalization on dataset GSM3309834.

| GSM3309836 | |
|---|---|
| cells | 9,514 |
| genes | 60,623 |
| nonzero entries | 23,079,845 |
| density | 0.04 |
| total counts | 65,277,912 |
| mean counts / cell | 6,861 |
| mean counts / gene | 1,077 |
| min counts / cell | 1,176 |
| max counts / cell | 27,791 |
| overdispersion | 0.355 |

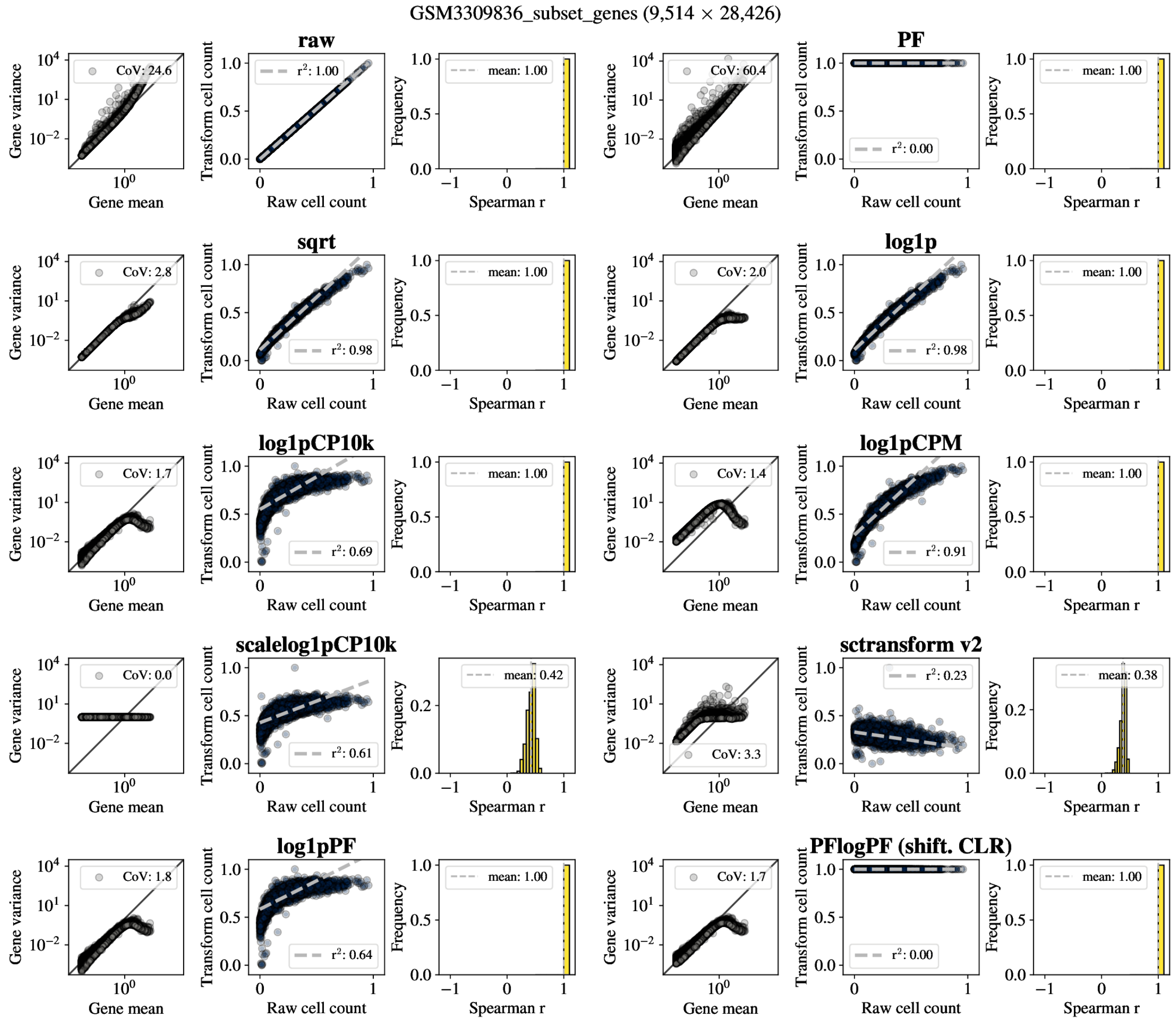


Supplementary Figure 1.189. **GSM3309836**. Per-method comparison of variance stabilization (left), depth normalization (middle), and monotonicity (right) for each normalization on dataset GSM3309836.

| GSM3319032 | |
|---|---|
| cells | 5,089 |
| genes | 60,623 |
| nonzero entries | 4,833,437 |
| density | 0.0157 |
| total counts | 11,282,425 |
| mean counts / cell | 2,217 |
| mean counts / gene | 186 |
| min counts / cell | 918 |
| max counts / cell | 9,151 |
| overdispersion | 0.244 |

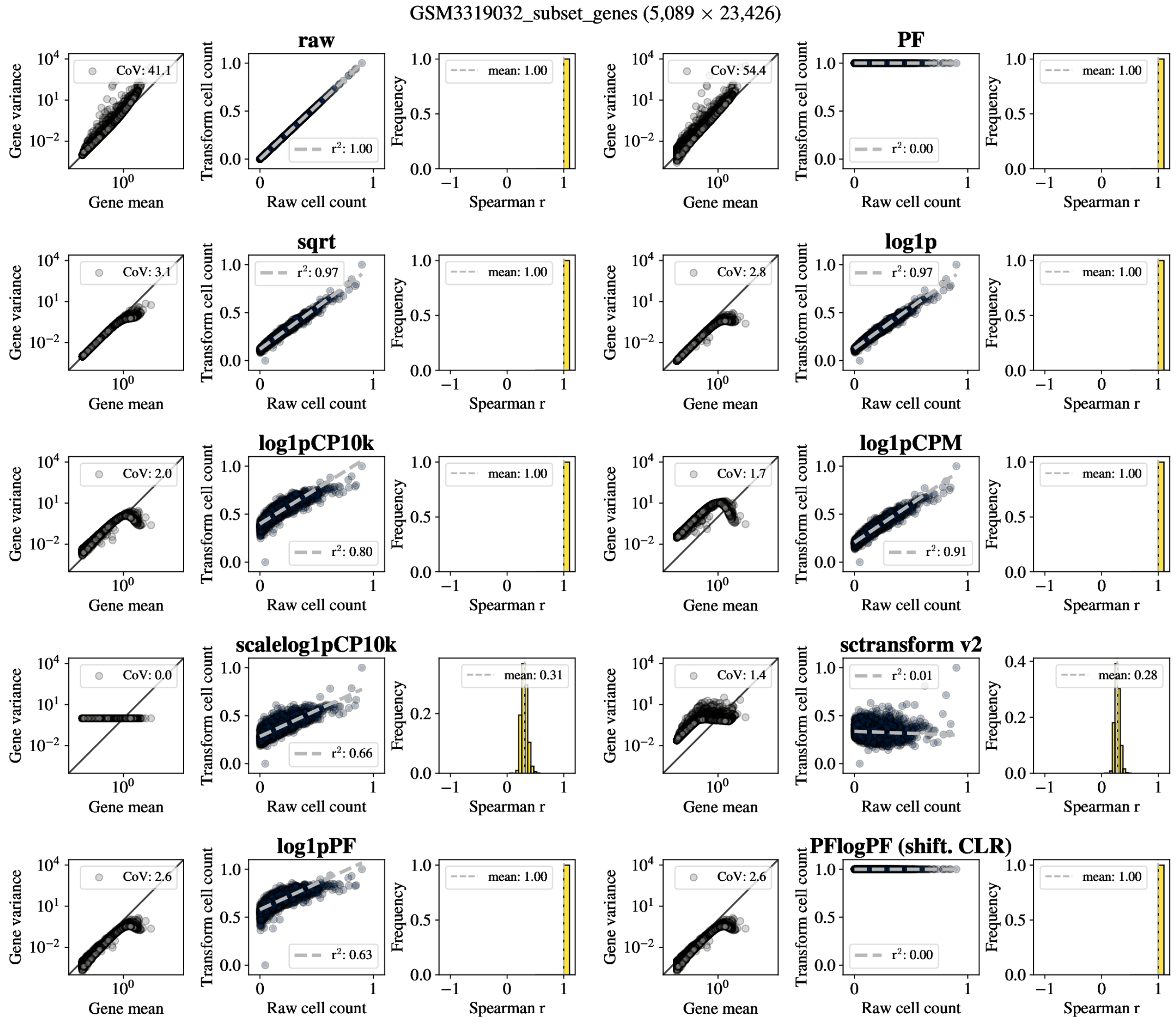


Supplementary Figure 1.190. **GSM3319032**. Per-method comparison of variance stabilization (left), depth normalization (middle), and monotonicity (right) for each normalization on dataset GSM3319032.

| GSM3319033 | |
|---|---|
| cells | 2,249 |
| genes | 60,623 |
| nonzero entries | 2,372,098 |
| density | 0.0174 |
| total counts | 5,453,680 |
| mean counts / cell | 2,425 |
| mean counts / gene | 90 |
| min counts / cell | 224 |
| max counts / cell | 15,749 |
| overdispersion | 0.315 |

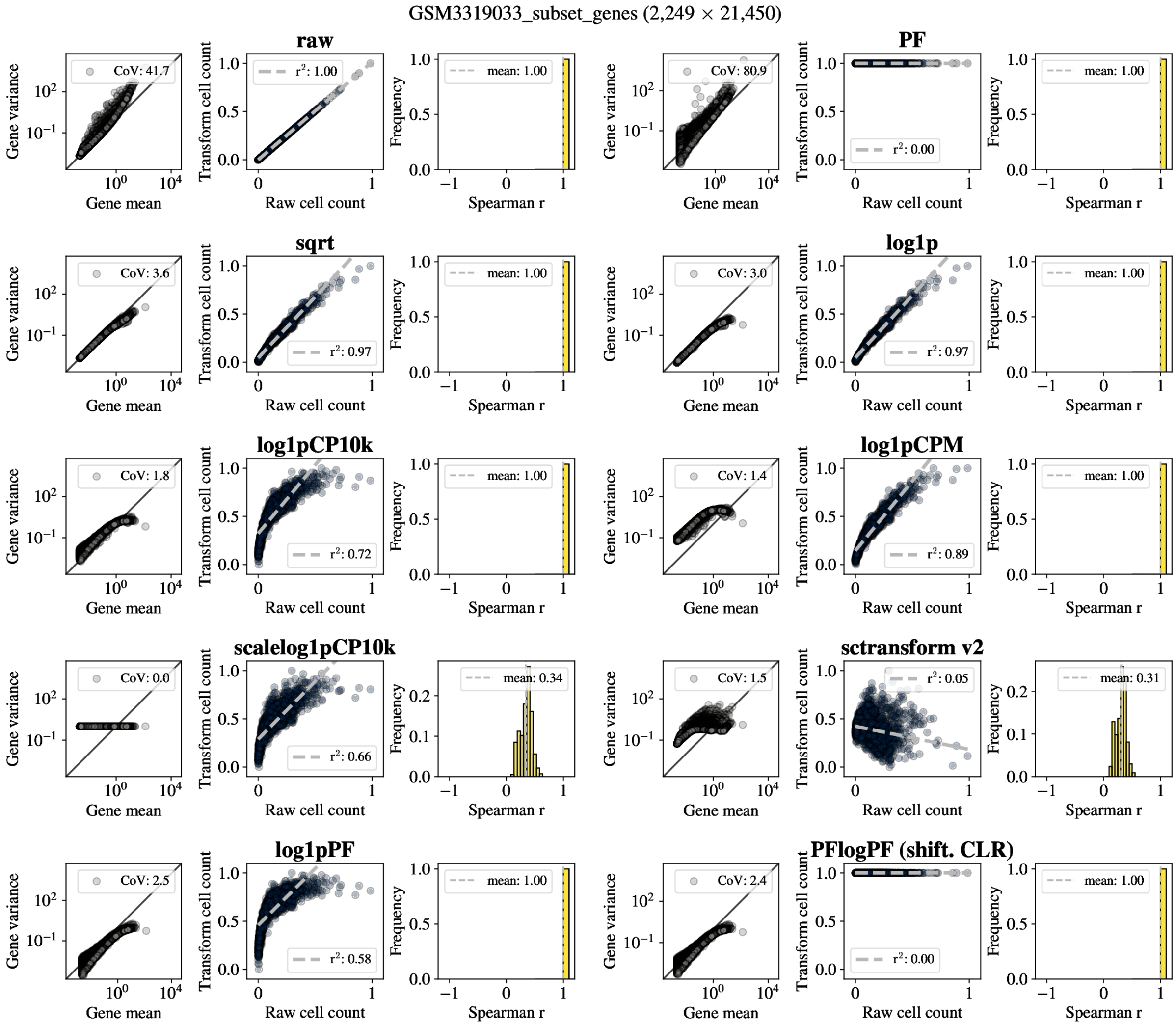


Supplementary Figure 1.191. **GSM3319033**. Per-method comparison of variance stabilization (left), depth normalization (middle), and monotonicity (right) for each normalization on dataset GSM3319033.

| GSM3319034 | |
|---|---|
| cells | 2,848 |
| genes | 60,623 |
| nonzero entries | 3,628,926 |
| density | 0.021 |
| total counts | 8,206,391 |
| mean counts / cell | 2,881 |
| mean counts / gene | 135 |
| min counts / cell | 1,557 |
| max counts / cell | 11,872 |
| overdispersion | 0.163 |

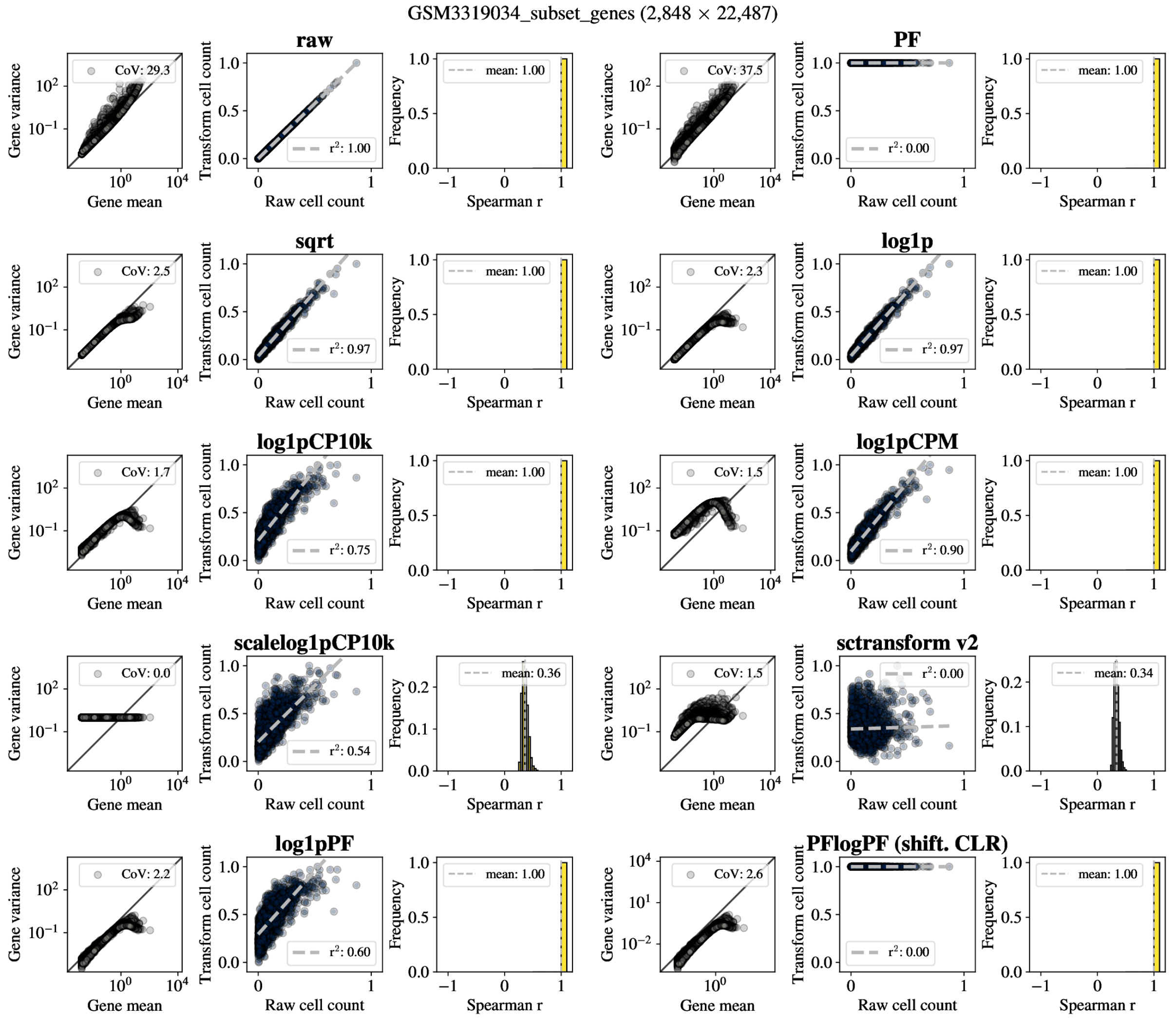


Supplementary Figure 1.192. **GSM3319034**. Per-method comparison of variance stabilization (left), depth normalization (middle), and monotonicity (right) for each normalization on dataset GSM3319034.

| GSM3319035 | |
|---|---|
| cells | 3,311 |
| genes | 60,623 |
| nonzero entries | 3,820,856 |
| density | 0.019 |
| total counts | 9,137,713 |
| mean counts / cell | 2,760 |
| mean counts / gene | 151 |
| min counts / cell | 636 |
| max counts / cell | 17,569 |
| overdispersion | 0.264 |

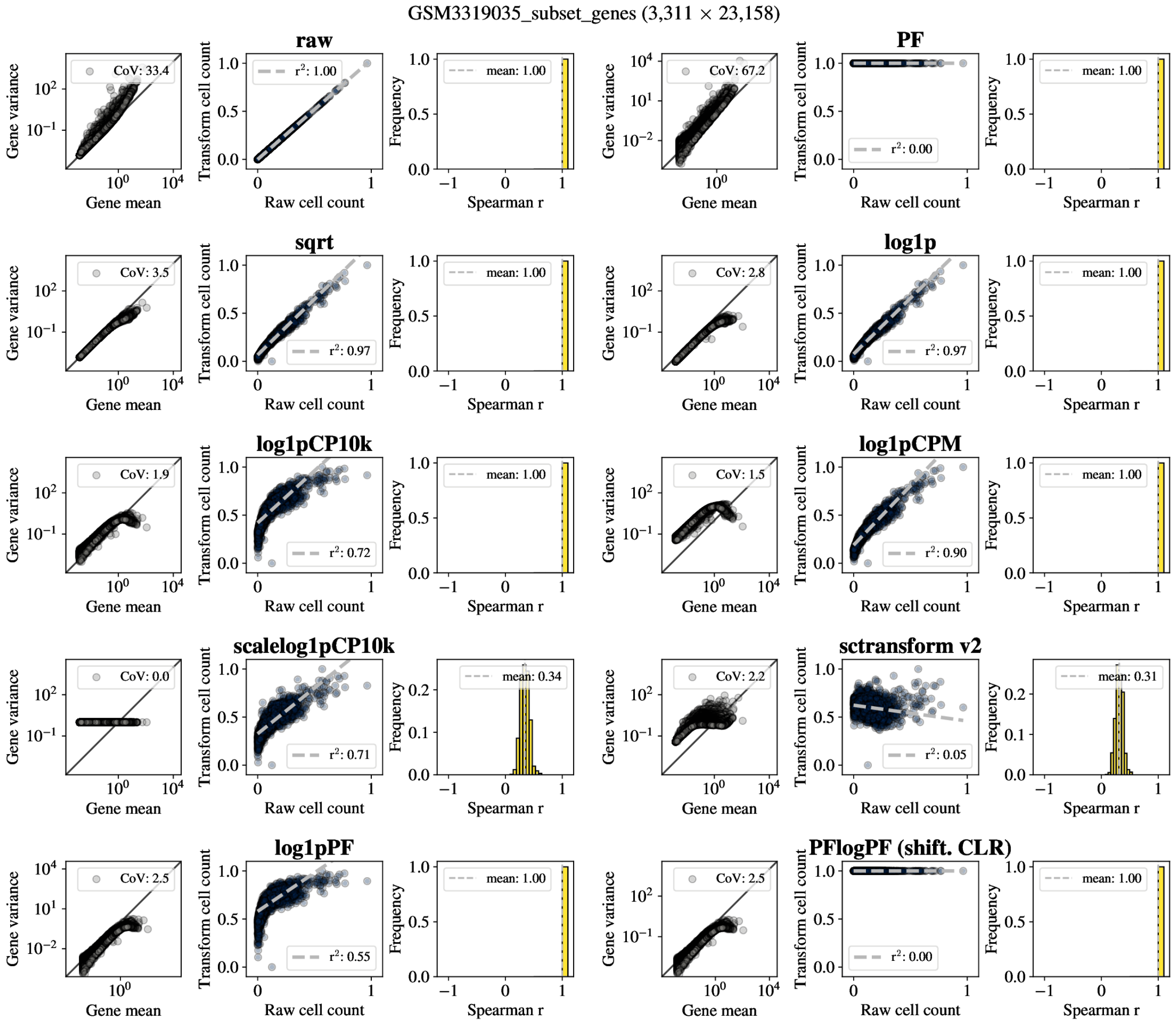


Supplementary Figure 1.193. **GSM3319035**. Per-method comparison of variance stabilization (left), depth normalization (middle), and monotonicity (right) for each normalization on dataset GSM3319035.

| GSM3319036 | |
| --- | --- |
| cells | 3,683 |
| genes | 60,623 |
| nonzero entries | 4,463,600 |
| density | 0.02 |
| total counts | 9,811,726 |
| mean counts / cell | 2,664 |
| mean counts / gene | 162 |
| min counts / cell | 68 |
| max counts / cell | 13,557 |
| overdispersion | 0.471 |

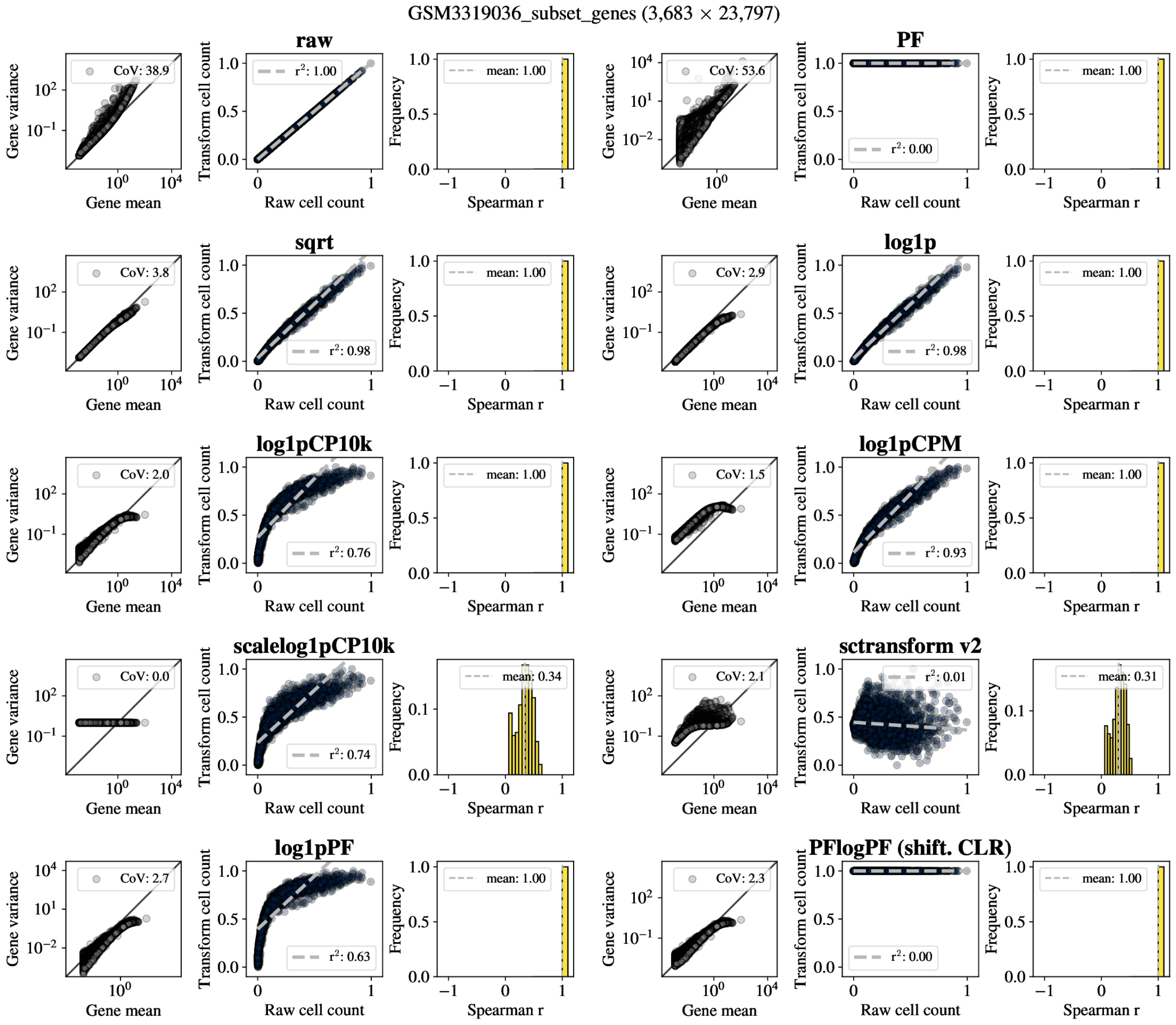


Supplementary Figure 1.194. **GSM3319036**. Per-method comparison of variance stabilization (left), depth normalization (middle), and monotonicity (right) for each normalization on dataset GSM3319036.

| GSM3319037 | |
|---|---|
| cells | 2,032 |
| genes | 60,623 |
| nonzero entries | 2,665,672 |
| density | 0.0216 |
| total counts | 6,824,934 |
| mean counts / cell | 3,359 |
| mean counts / gene | 113 |
| min counts / cell | 1,296 |
| max counts / cell | 13,851 |
| overdispersion | 0.319 |

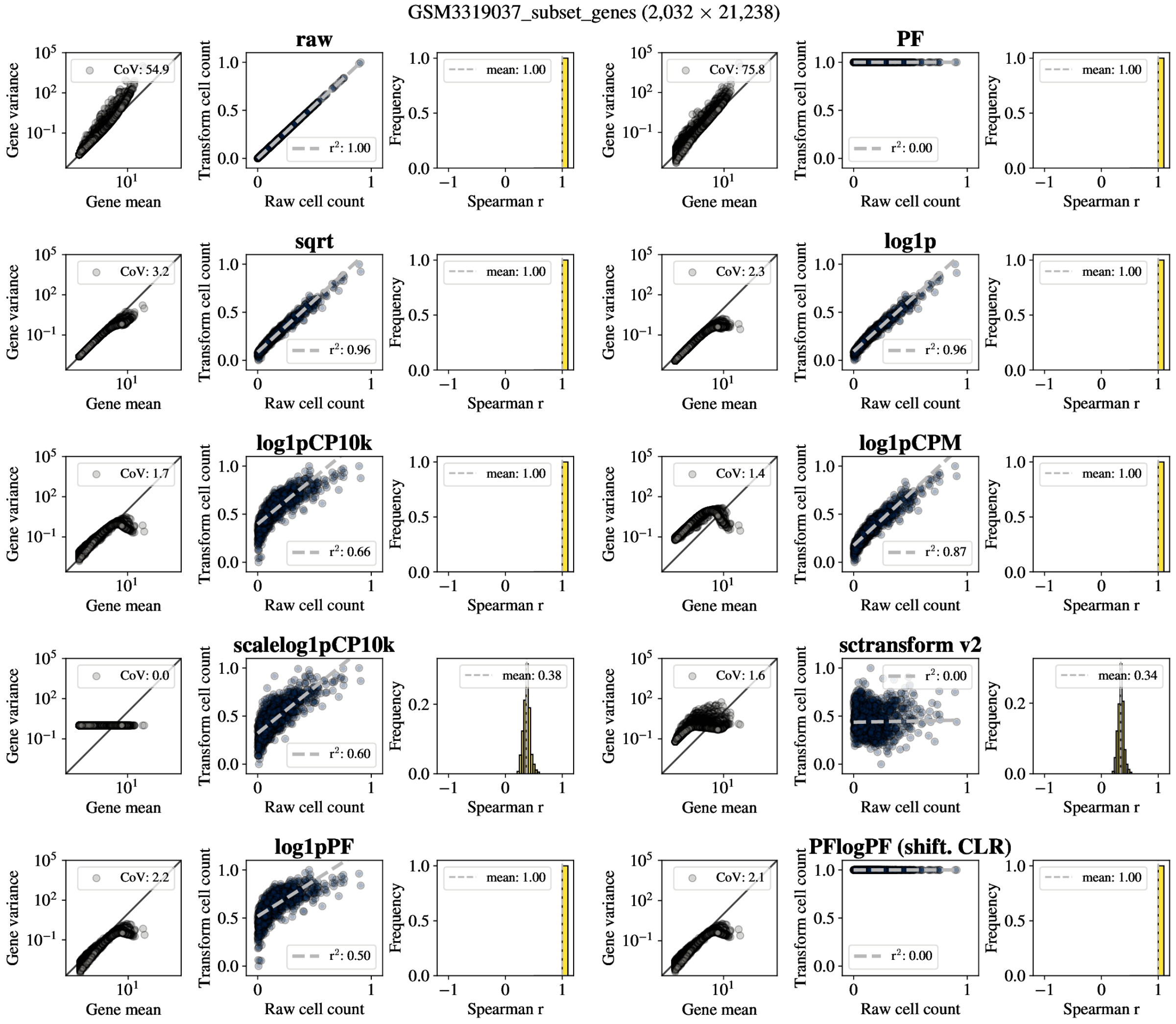


Supplementary Figure 1.195. **GSM3319037**. Per-method comparison of variance stabilization (left), depth normalization (middle), and monotonicity (right) for each normalization on dataset GSM3319037.

| **GSM3319038** | |
|---|---|
| cells | 4,671 |
| genes | 60,623 |
| nonzero entries | 4,728,369 |
| density | 0.0167 |
| total counts | 10,082,900 |
| mean counts / cell | 2,159 |
| mean counts / gene | 166 |
| min counts / cell | 390 |
| max counts / cell | 13,676 |
| overdispersion | 0.285 |

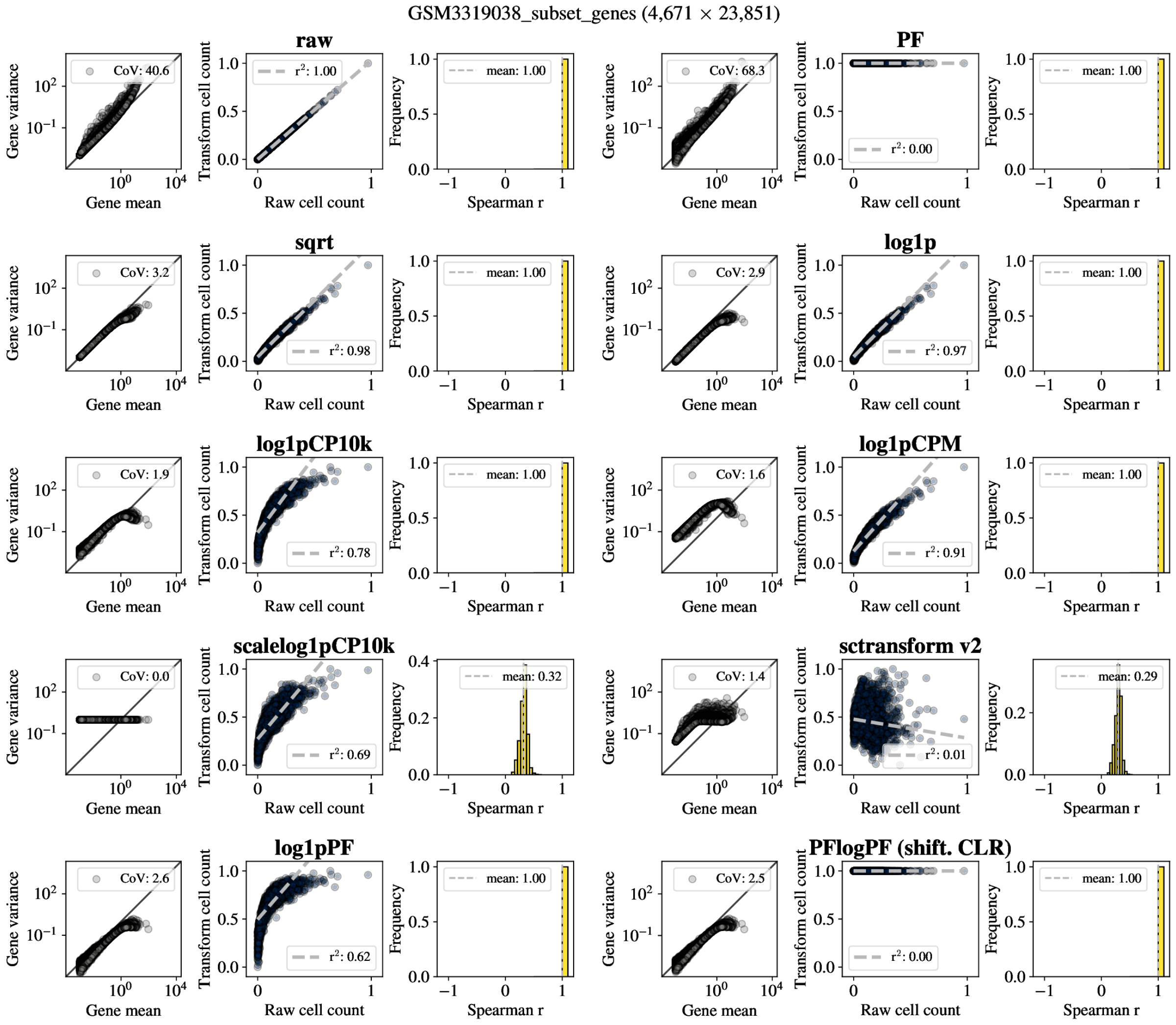


Supplementary Figure 1.196. **GSM3319038**. Per-method comparison of variance stabilization (left), depth normalization (middle), and monotonicity (right) for each normalization on dataset GSM3319038.

| GSM3319039 | |
|---|---|
| cells | 3,829 |
| genes | 60,623 |
| nonzero entries | 3,959,490 |
| density | 0.0171 |
| total counts | 7,957,927 |
| mean counts / cell | 2,078 |
| mean counts / gene | 131 |
| min counts / cell | 116 |
| max counts / cell | 18,141 |
| overdispersion | 0.471 |

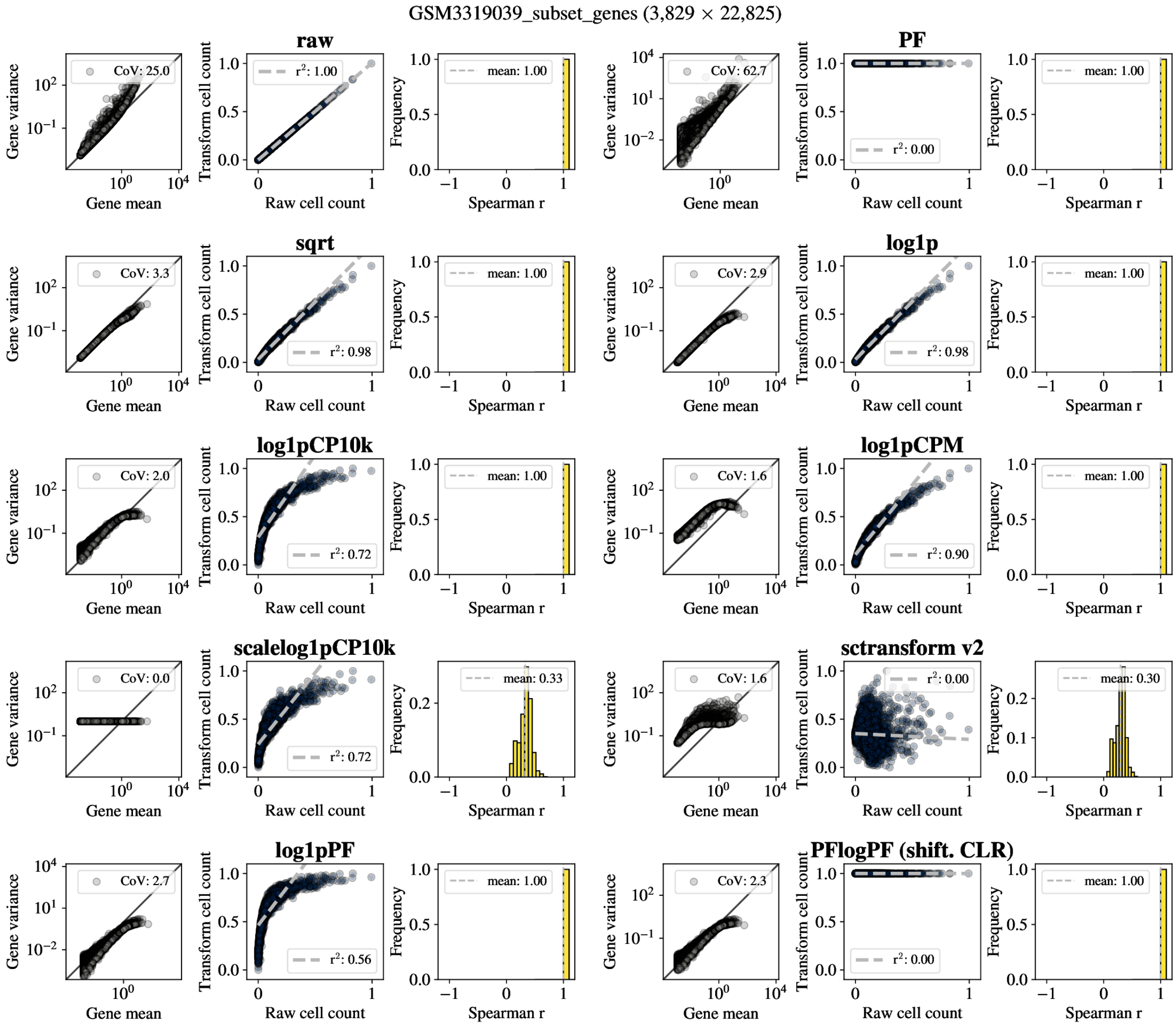


Supplementary Figure 1.197. **GSM3319039**. Per-method comparison of variance stabilization (left), depth normalization (middle), and monotonicity (right) for each normalization on dataset GSM3319039.

| GSM3319040 | |
|---|---|
| cells | 643 |
| genes | 60,623 |
| nonzero entries | 653,412 |
| density | 0.0168 |
| total counts | 1,488,196 |
| mean counts / cell | 2,314 |
| mean counts / gene | 25 |
| min counts / cell | 238 |
| max counts / cell | 10,709 |
| overdispersion | 0.566 |

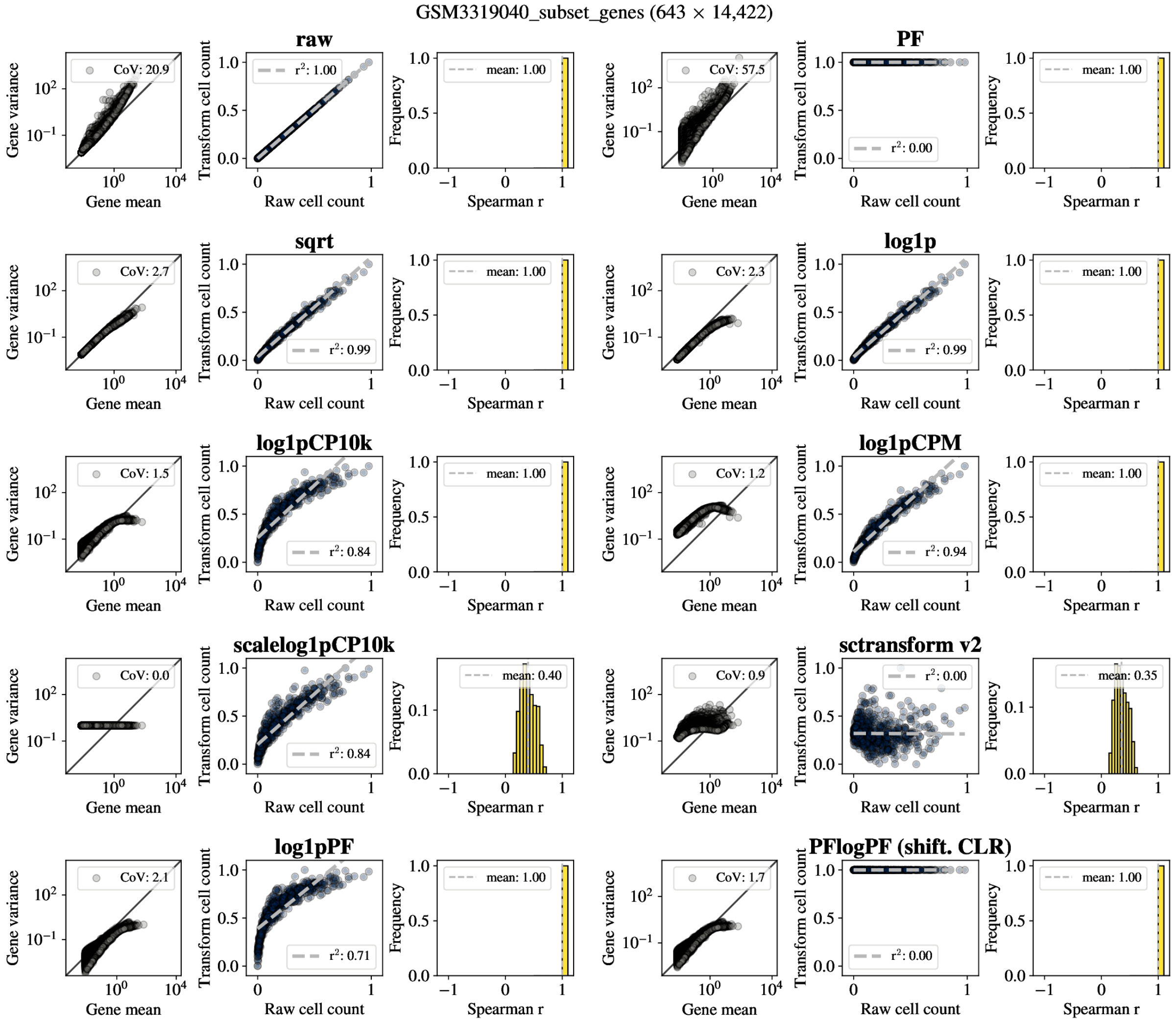


Supplementary Figure 1.198. **GSM3319040**. Per-method comparison of variance stabilization (left), depth normalization (middle), and monotonicity (right) for each normalization on dataset GSM3319040.

| GSM3319041 | |
|---|---|
| cells | 1,270 |
| genes | 60,623 |
| nonzero entries | 1,529,029 |
| density | 0.0199 |
| total counts | 3,349,168 |
| mean counts / cell | 2,637 |
| mean counts / gene | 55 |
| min counts / cell | 741 |
| max counts / cell | 20,020 |
| overdispersion | 0.634 |

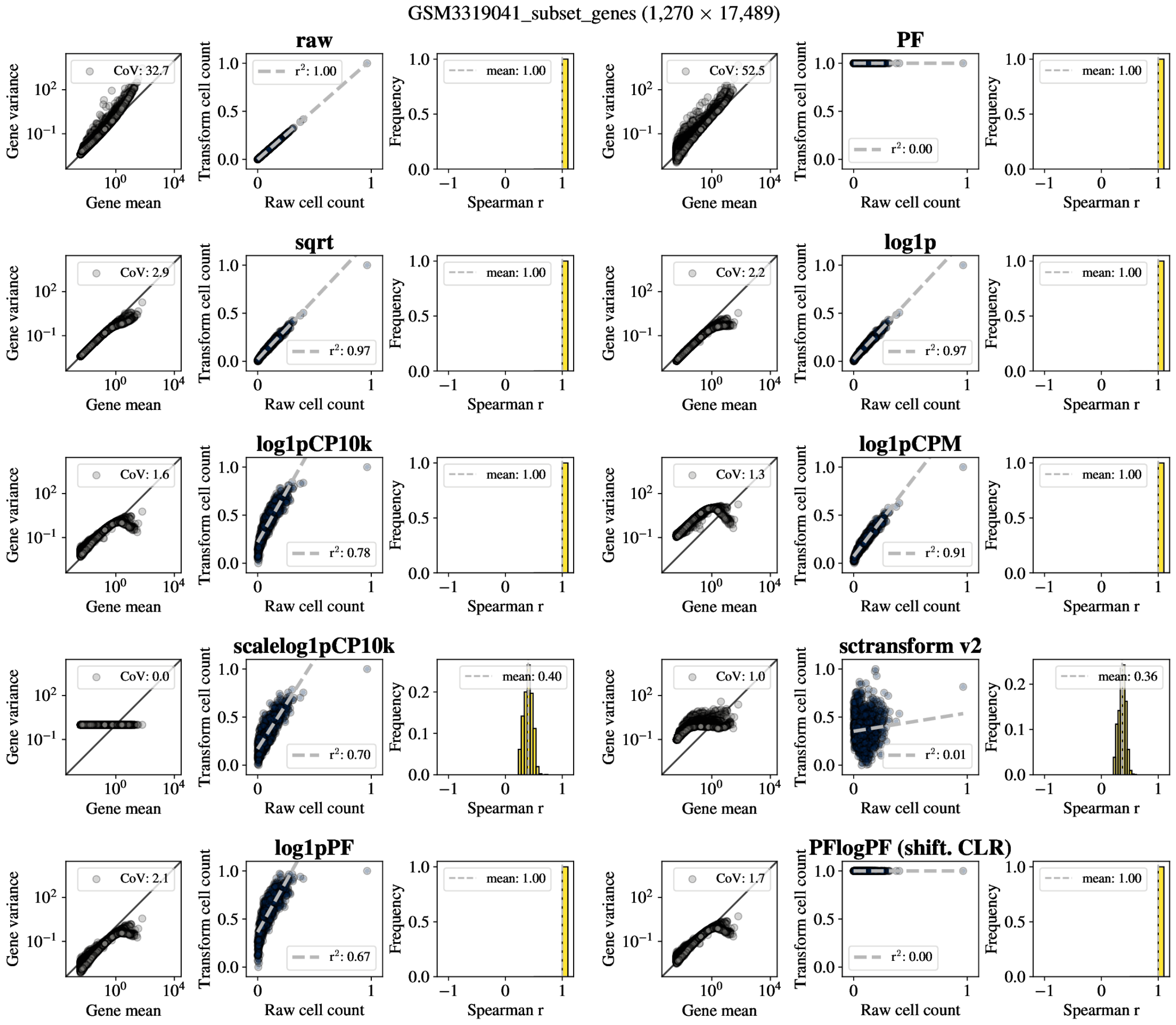


Supplementary Figure 1.199. **GSM3319041**. Per-method comparison of variance stabilization (left), depth normalization (middle), and monotonicity (right) for each normalization on dataset GSM3319041.

| GSM3319042 | |
|---|---|
| cells | 1,181 |
| genes | 60,623 |
| nonzero entries | 694,865 |
| density | 0.00971 |
| total counts | 1,399,649 |
| mean counts / cell | 1,185 |
| mean counts / gene | 23 |
| min counts / cell | 108 |
| max counts / cell | 7,230 |
| overdispersion | 0.648 |

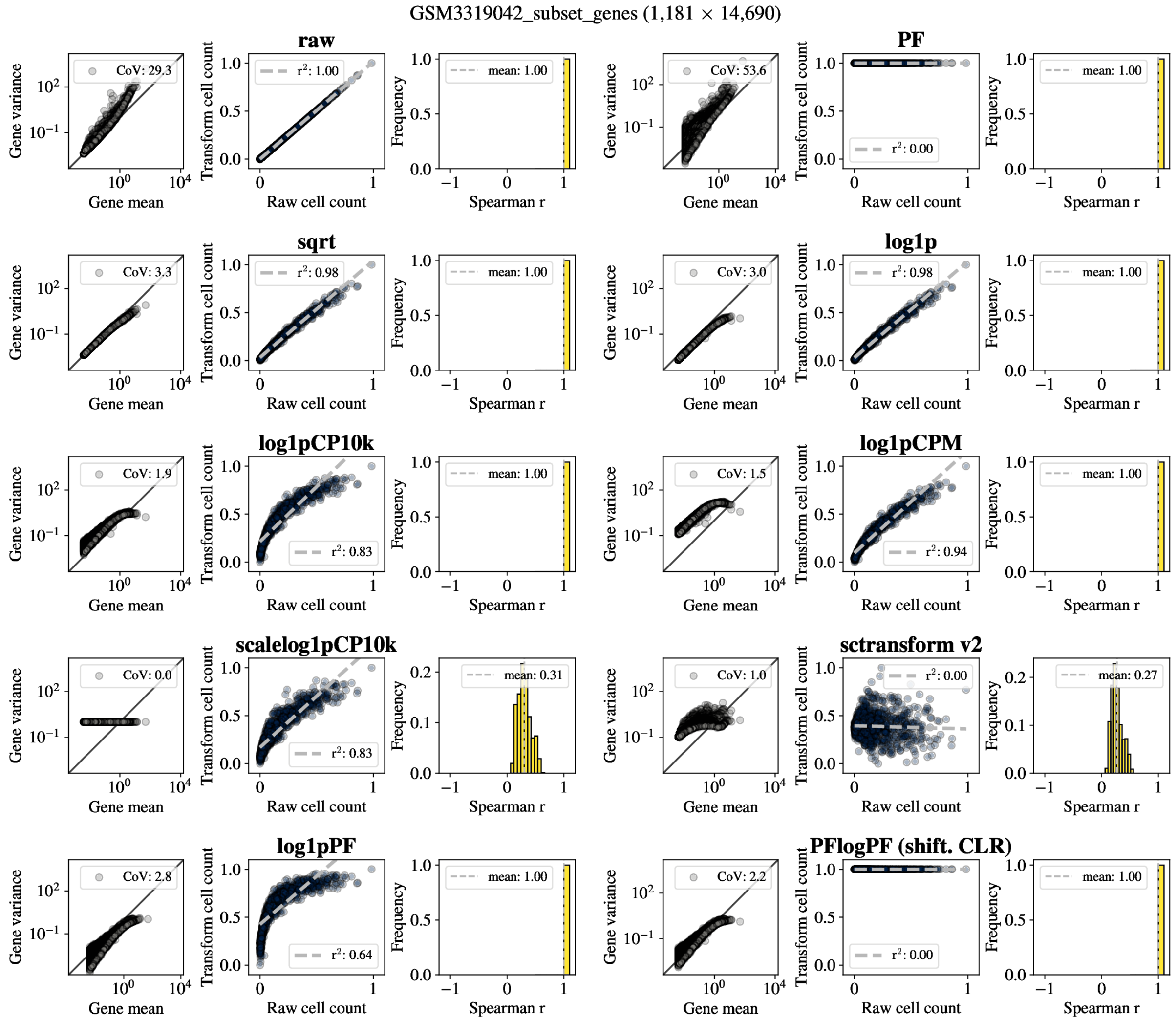


Supplementary Figure 1.200. **GSM3319042**. Per-method comparison of variance stabilization (left), depth normalization (middle), and monotonicity (right) for each normalization on dataset GSM3319042.

| GSM3319043 | |
|---|---|
| cells | 7,286 |
| genes | 60,623 |
| nonzero entries | 3,959,038 |
| density | 0.00896 |
| total counts | 7,412,483 |
| mean counts / cell | 1,017 |
| mean counts / gene | 122 |
| min counts / cell | 73 |
| max counts / cell | 9,710 |
| overdispersion | 0.748 |

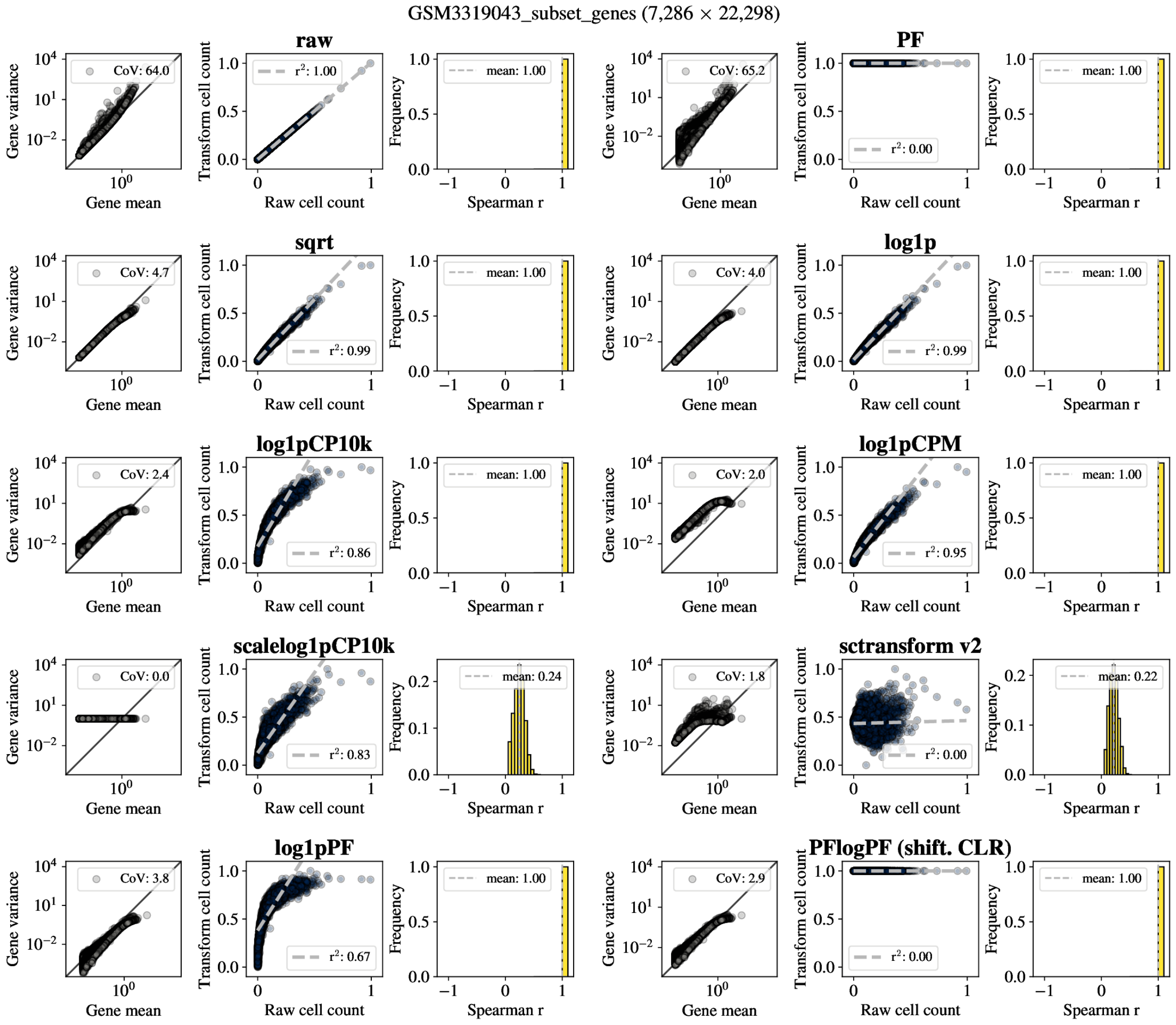


Supplementary Figure 1.201. **GSM3319043**. Per-method comparison of variance stabilization (left), depth normalization (middle), and monotonicity (right) for each normalization on dataset GSM3319043.

| GSM3319044 | |
|---|---|
| cells | 4,274 |
| genes | 60,623 |
| nonzero entries | 2,503,462 |
| density | 0.00966 |
| total counts | 4,747,538 |
| mean counts / cell | 1,111 |
| mean counts / gene | 78 |
| min counts / cell | 422 |
| max counts / cell | 5,493 |
| overdispersion | 0.302 |

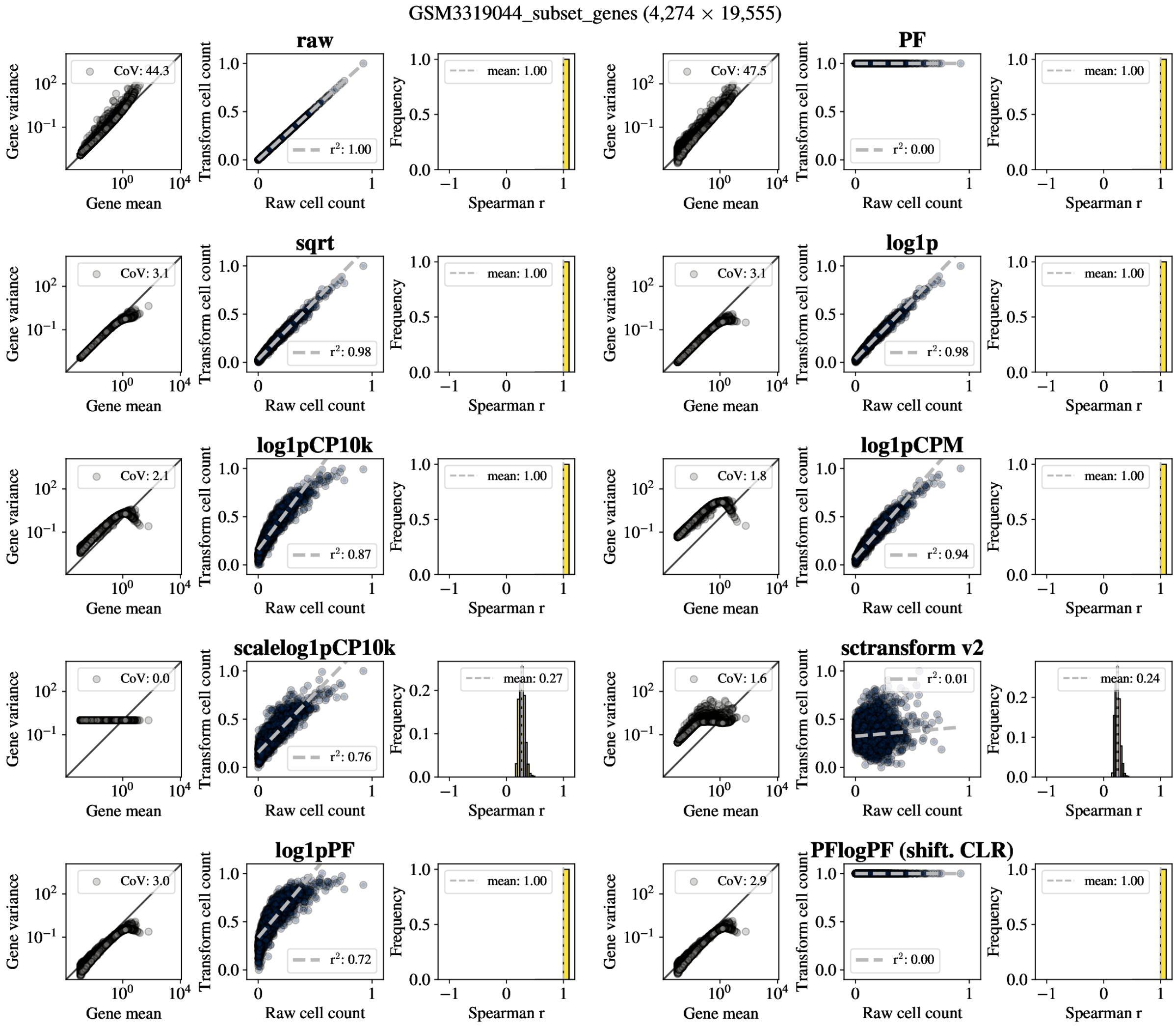


Supplementary Figure 1.202. **GSM3319044**. Per-method comparison of variance stabilization (left), depth normalization (middle), and monotonicity (right) for each normalization on dataset GSM3319044.

| GSM3319045 | |
|---|---|
| cells | 7,040 |
| genes | 60,623 |
| nonzero entries | 2,921,683 |
| density | 0.00685 |
| total counts | 5,591,512 |
| mean counts / cell | 794 |
| mean counts / gene | 92 |
| min counts / cell | 55 |
| max counts / cell | 4,568 |
| overdispersion | 0.31 |

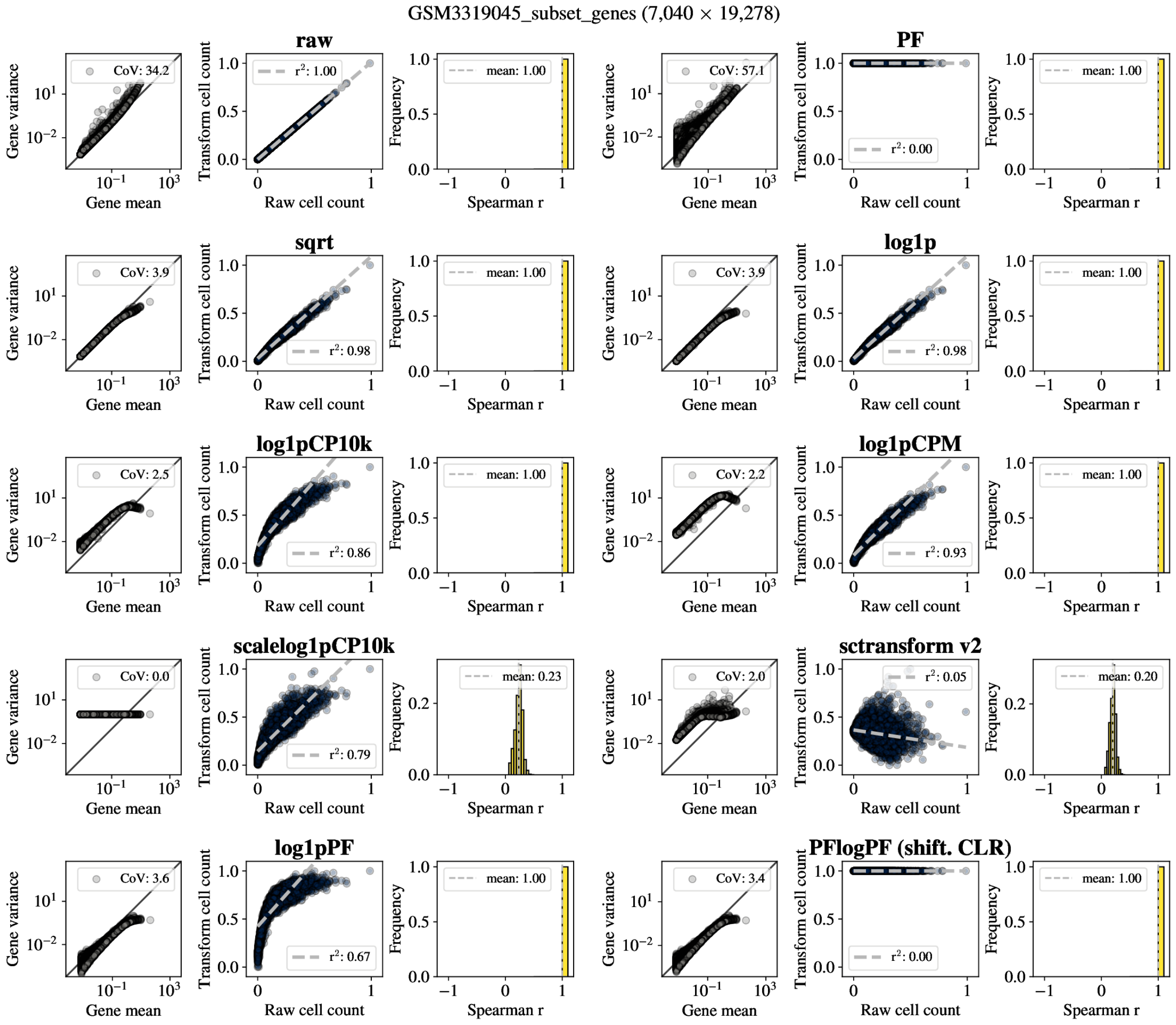


Supplementary Figure 1.203. **GSM3319045**. Per-method comparison of variance stabilization (left), depth normalization (middle), and monotonicity (right) for each normalization on dataset GSM3319045.

| GSM3319046 | |
|---|---|
| cells | 1,675 |
| genes | 60,623 |
| nonzero entries | 443,100 |
| density | 0.00436 |
| total counts | 864,319 |
| mean counts / cell | 516 |
| mean counts / gene | 14 |
| min counts / cell | 21 |
| max counts / cell | 6,941 |
| overdispersion | 2.4 |

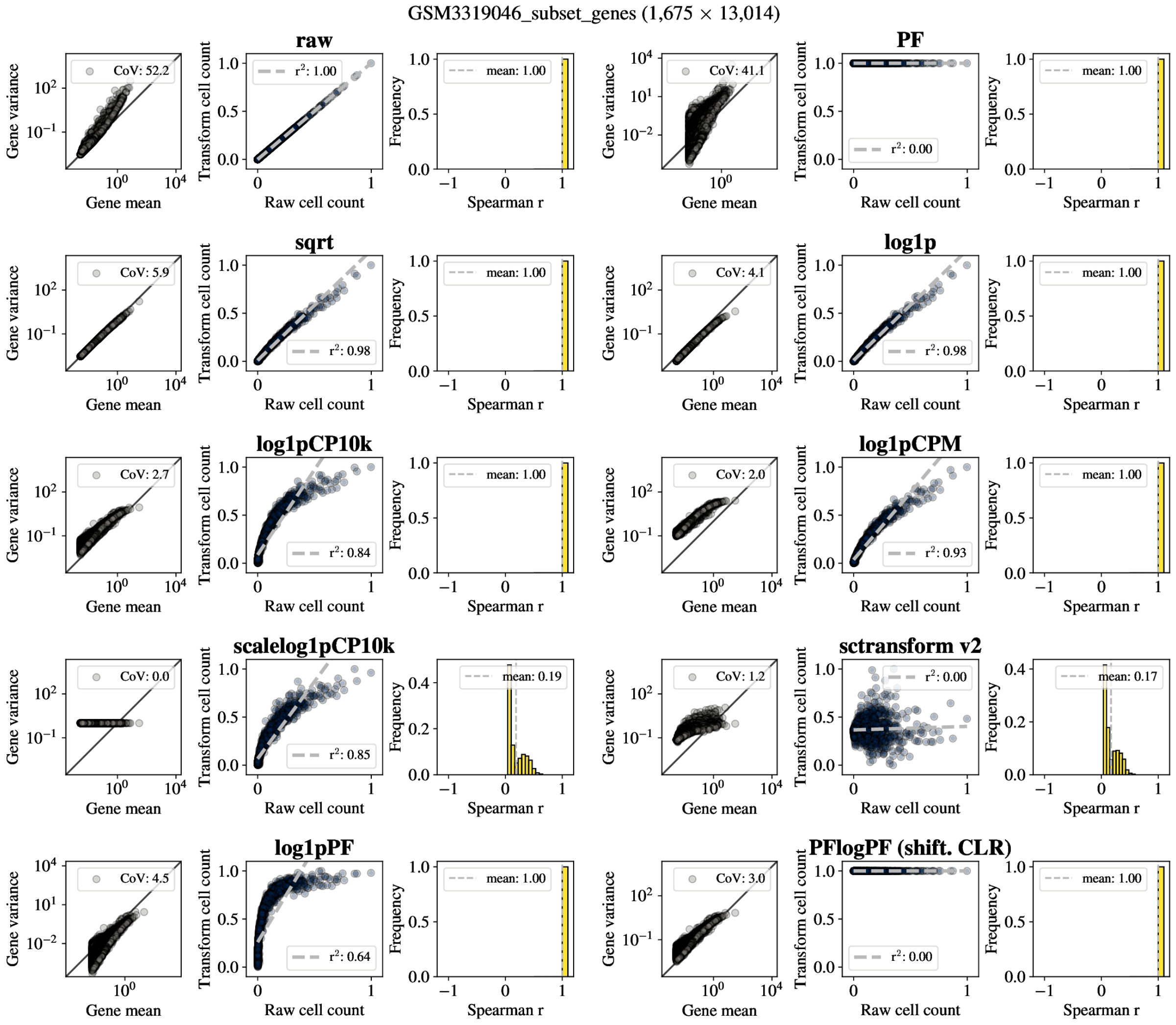


Supplementary Figure 1.204. **GSM3319046**. Per-method comparison of variance stabilization (left), depth normalization (middle), and monotonicity (right) for each normalization on dataset GSM3319046.

| GSM3319047 | |
|---|---|
| cells | 137 |
| genes | 60,623 |
| nonzero entries | 66,143 |
| density | 0.00796 |
| total counts | 148,803 |
| mean counts / cell | 1,086 |
| mean counts / gene | 2 |
| min counts / cell | 191 |
| max counts / cell | 4,557 |
| overdispersion | 0.341 |

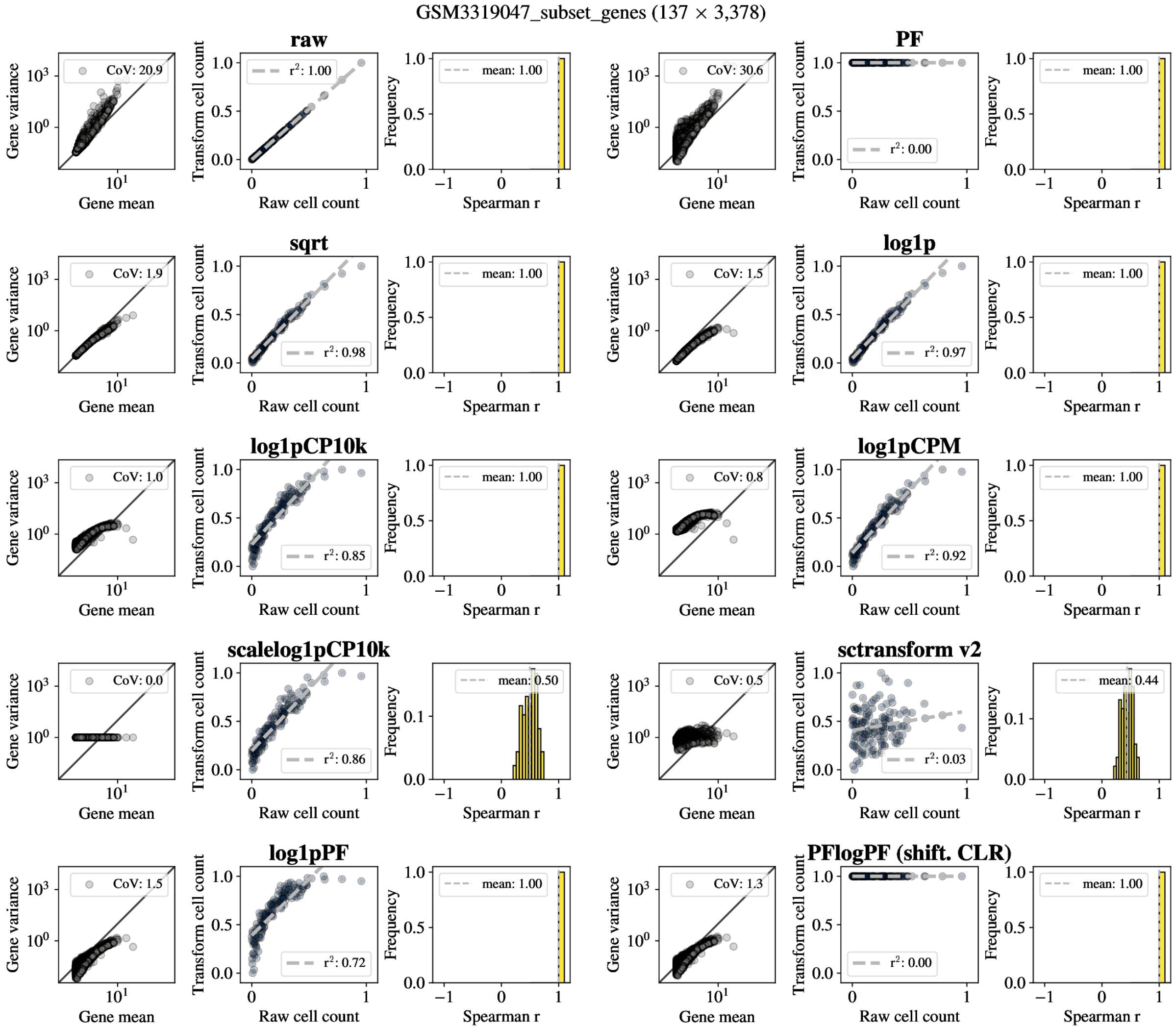


Supplementary Figure 1.205. **GSM3319047**. Per-method comparison of variance stabilization (left), depth normalization (middle), and monotonicity (right) for each normalization on dataset GSM3319047.

| GSM3360833 | |
|---|---|
| cells | 10,860 |
| genes | 60,623 |
| nonzero entries | 17,093,085 |
| density | 0.026 |
| total counts | 39,948,147 |
| mean counts / cell | 3,678 |
| mean counts / gene | 659 |
| min counts / cell | 178 |
| max counts / cell | 92,232 |
| overdispersion | 2.78 |

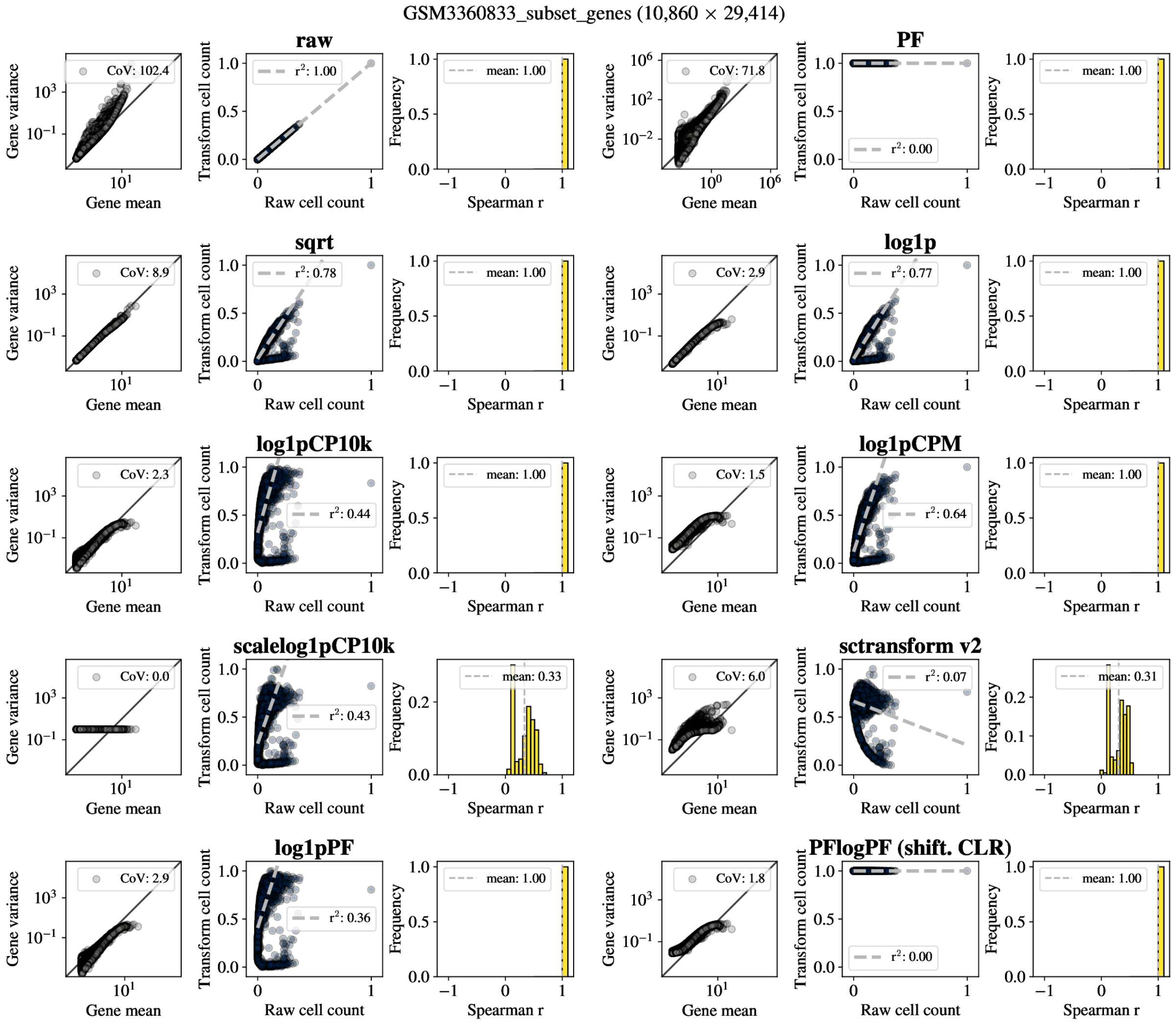


Supplementary Figure 1.206. **GSM3360833**. Per-method comparison of variance stabilization (left), depth normalization (middle), and monotonicity (right) for each normalization on dataset GSM3360833.

| GSM3360834 | |
|---|---|
| cells | 4,701 |
| genes | 60,623 |
| nonzero entries | 9,362,037 |
| density | 0.0329 |
| total counts | 20,880,440 |
| mean counts / cell | 4,442 |
| mean counts / gene | 344 |
| min counts / cell | 444 |
| max counts / cell | 24,271 |
| overdispersion | 0.284 |

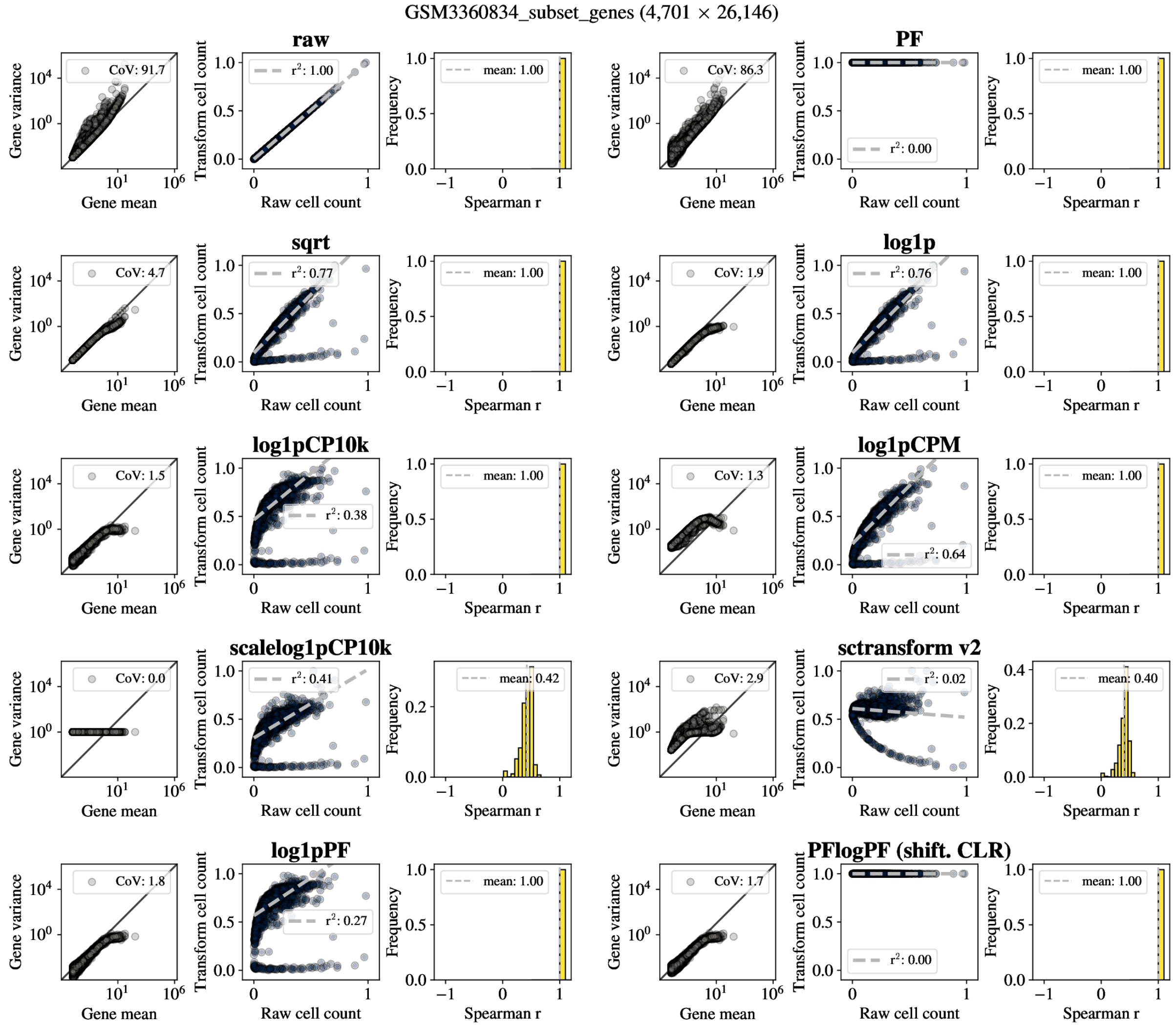


Supplementary Figure 1.207. **GSM3360834**. Per-method comparison of variance stabilization (left), depth normalization (middle), and monotonicity (right) for each normalization on dataset GSM3360834.

| **GSM3360835** | |
|---|---|
| cells | 8,620 |
| genes | 60,623 |
| nonzero entries | 9,936,579 |
| density | 0.019 |
| total counts | 21,668,079 |
| mean counts / cell | 2,514 |
| mean counts / gene | 357 |
| min counts / cell | 200 |
| max counts / cell | 27,531 |
| overdispersion | 11.5 |

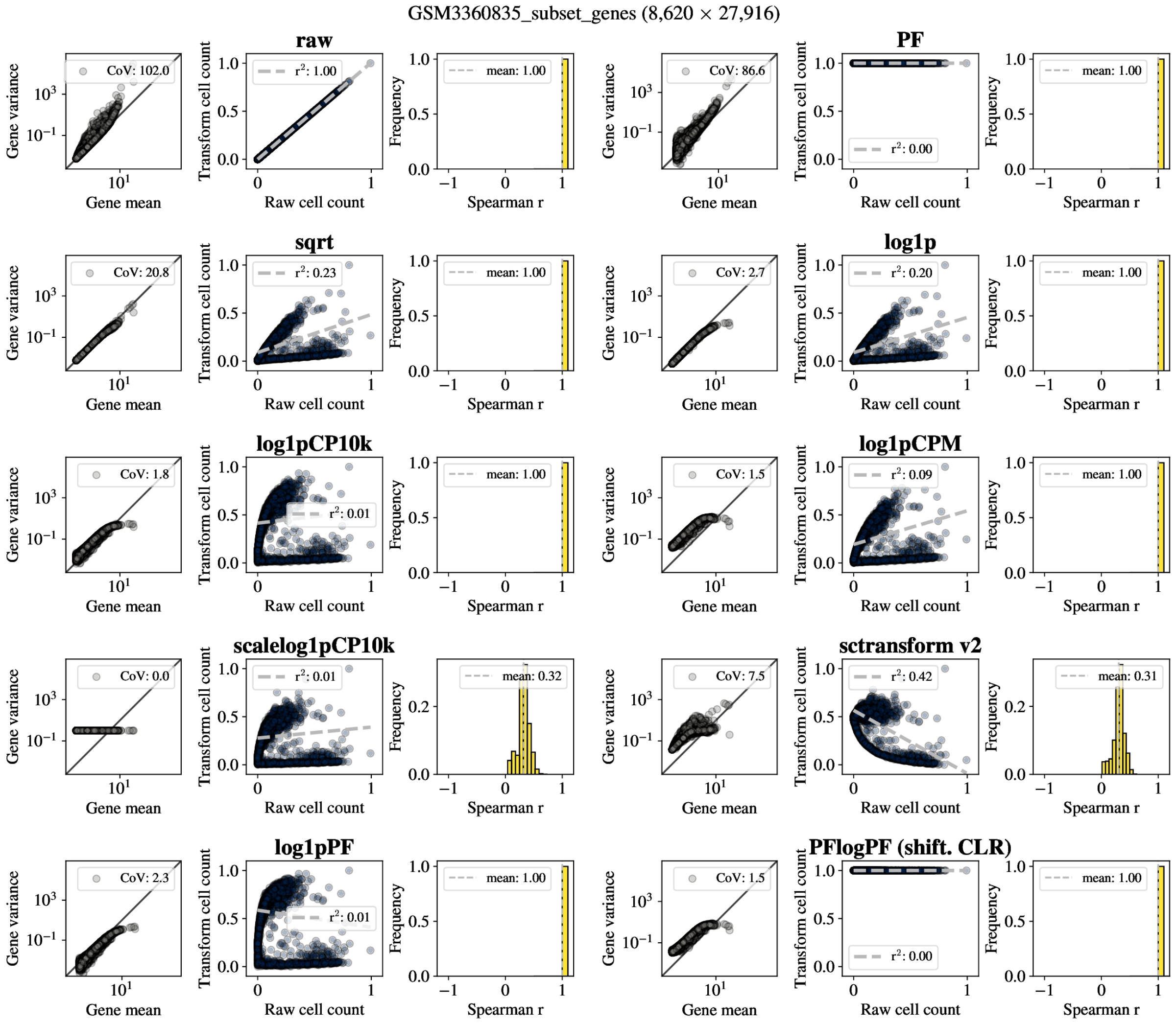


Supplementary Figure 1.208. **GSM3360835**. Per-method comparison of variance stabilization (left), depth normalization (middle), and monotonicity (right) for each normalization on dataset GSM3360835.

| GSM3360836 | |
|---|---|
| cells | 1,157 |
| genes | 60,623 |
| nonzero entries | 473,991 |
| density | 0.00676 |
| total counts | 1,202,273 |
| mean counts / cell | 1,039 |
| mean counts / gene | 20 |
| min counts / cell | 601 |
| max counts / cell | 4,657 |
| overdispersion | 0.291 |

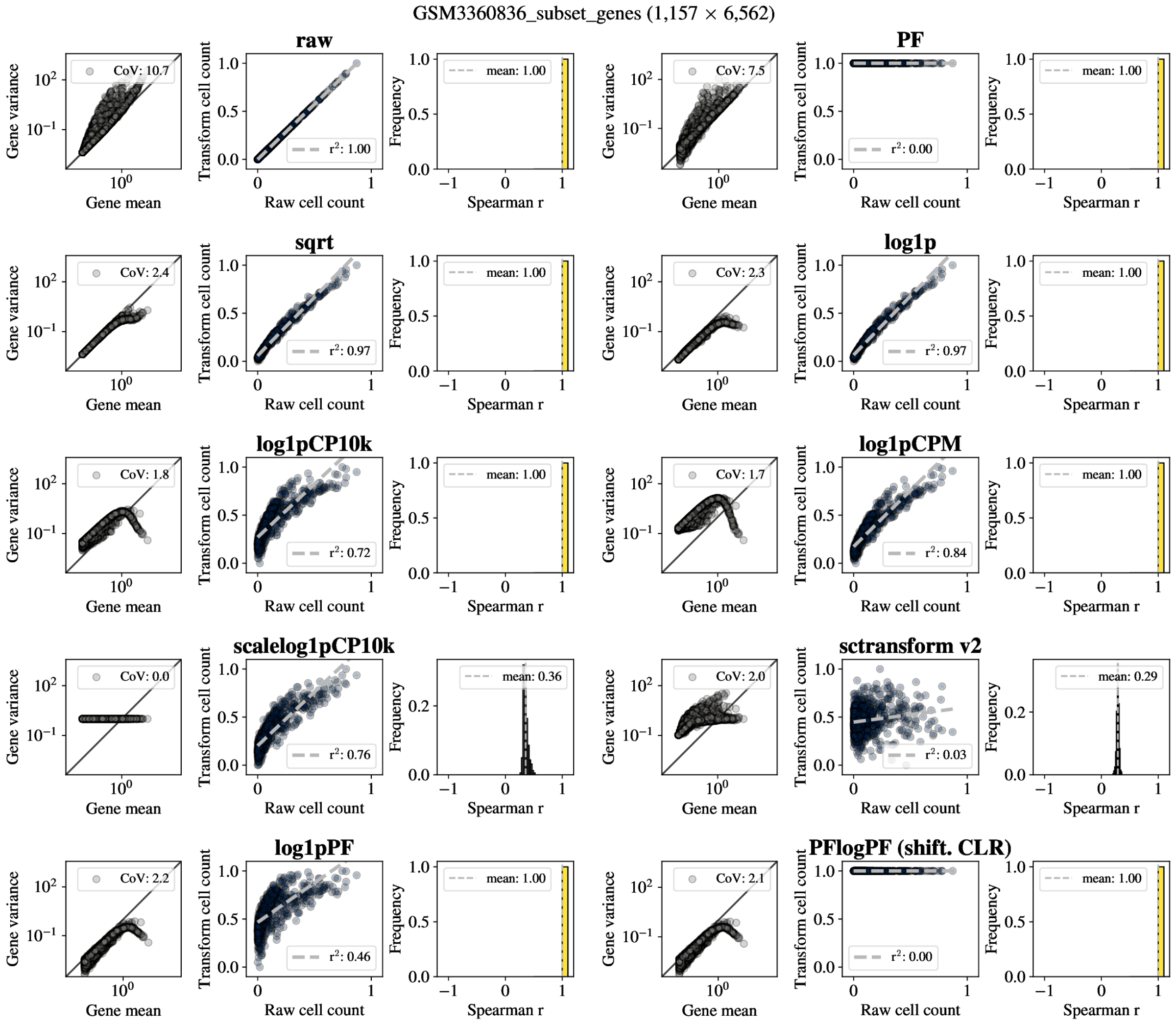


Supplementary Figure 1.209. **GSM3360836**. Per-method comparison of variance stabilization (left), depth normalization (middle), and monotonicity (right) for each normalization on dataset GSM3360836.

| **GSM3360837** | |
|---|---|
| cells | 12,947 |
| genes | 60,623 |
| nonzero entries | 13,222,986 |
| density | 0.0168 |
| total counts | 25,730,540 |
| mean counts / cell | 1,987 |
| mean counts / gene | 424 |
| min counts / cell | 164 |
| max counts / cell | 33,799 |
| overdispersion | 0.639 |

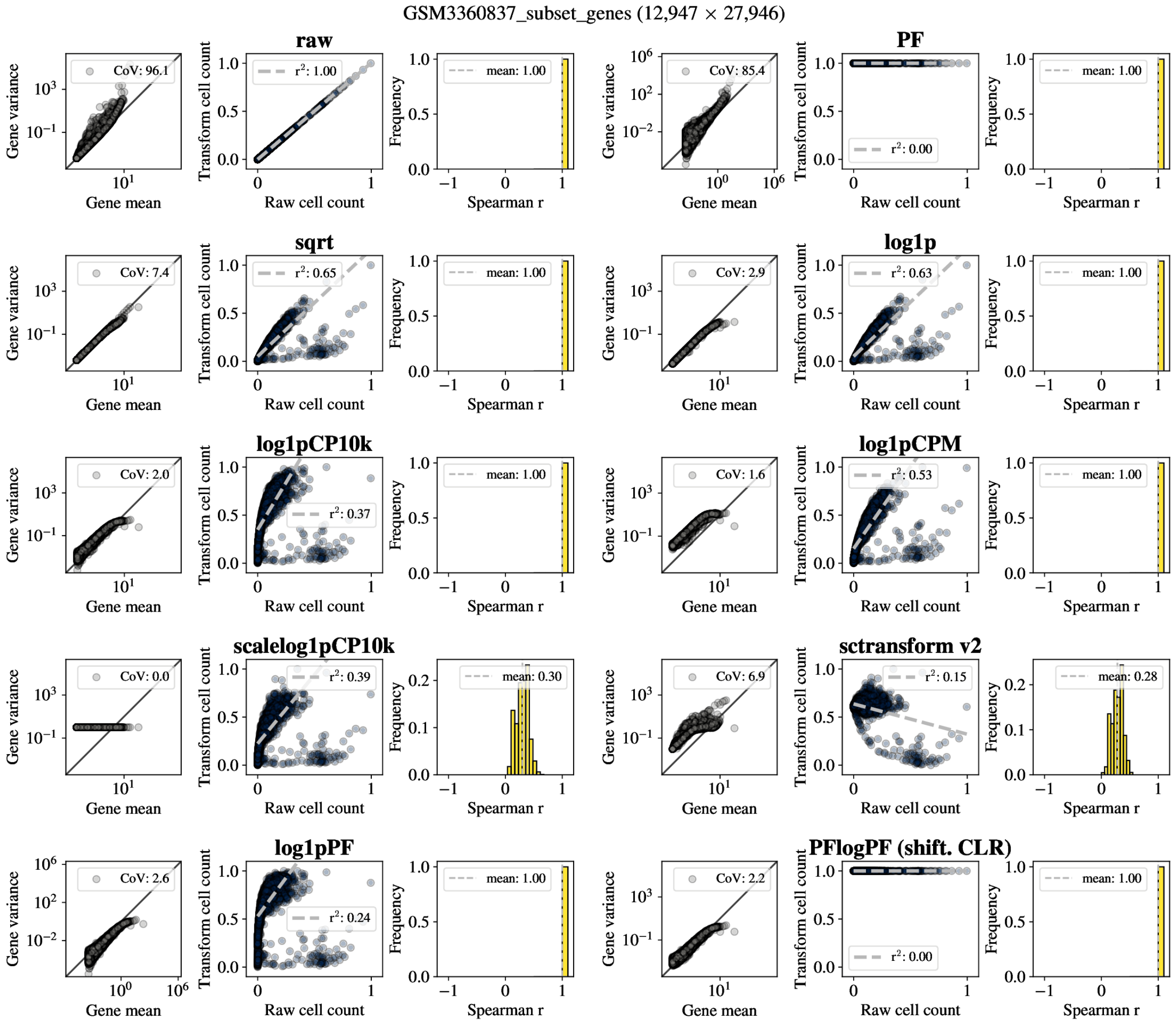


Supplementary Figure 1.210. **GSM3360837**. Per-method comparison of variance stabilization (left), depth normalization (middle), and monotonicity (right) for each normalization on dataset GSM3360837.

| GSM3360838 | |
|---|---|
| cells | 7,332 |
| genes | 60,623 |
| nonzero entries | 9,773,226 |
| density | 0.022 |
| total counts | 19,869,232 |
| mean counts / cell | 2,710 |
| mean counts / gene | 328 |
| min counts / cell | 127 |
| max counts / cell | 33,868 |
| overdispersion | 0.411 |

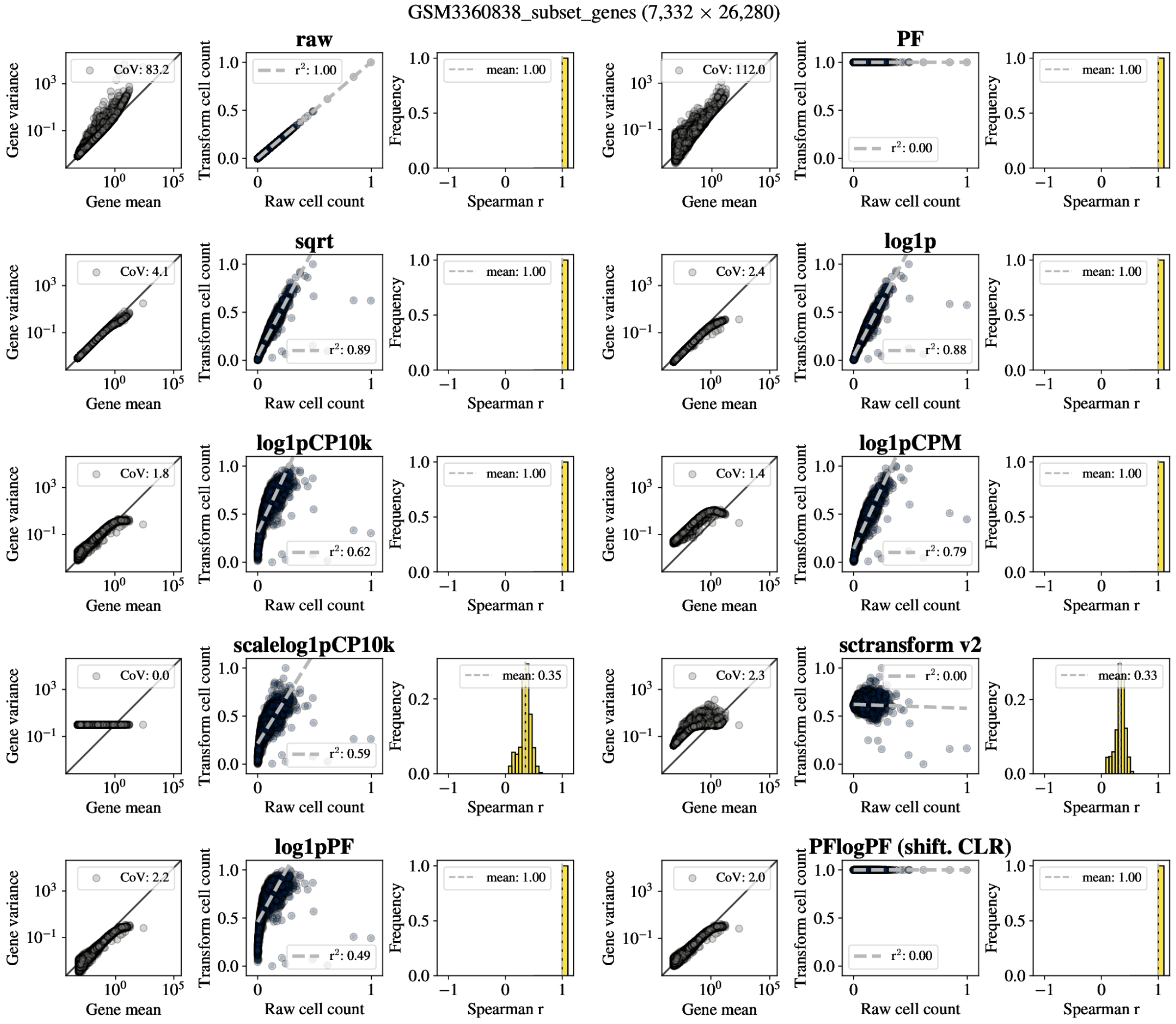


Supplementary Figure 1.211. **GSM3360838**. Per-method comparison of variance stabilization (left), depth normalization (middle), and monotonicity (right) for each normalization on dataset GSM3360838.

| GSM3375767 | |
|---|---|
| cells | 2,086 |
| genes | 60,623 |
| nonzero entries | 4,041,367 |
| density | 0.032 |
| total counts | 12,453,711 |
| mean counts / cell | 5,970 |
| mean counts / gene | 205 |
| min counts / cell | 1,532 |
| max counts / cell | 34,551 |
| overdispersion | 0.726 |

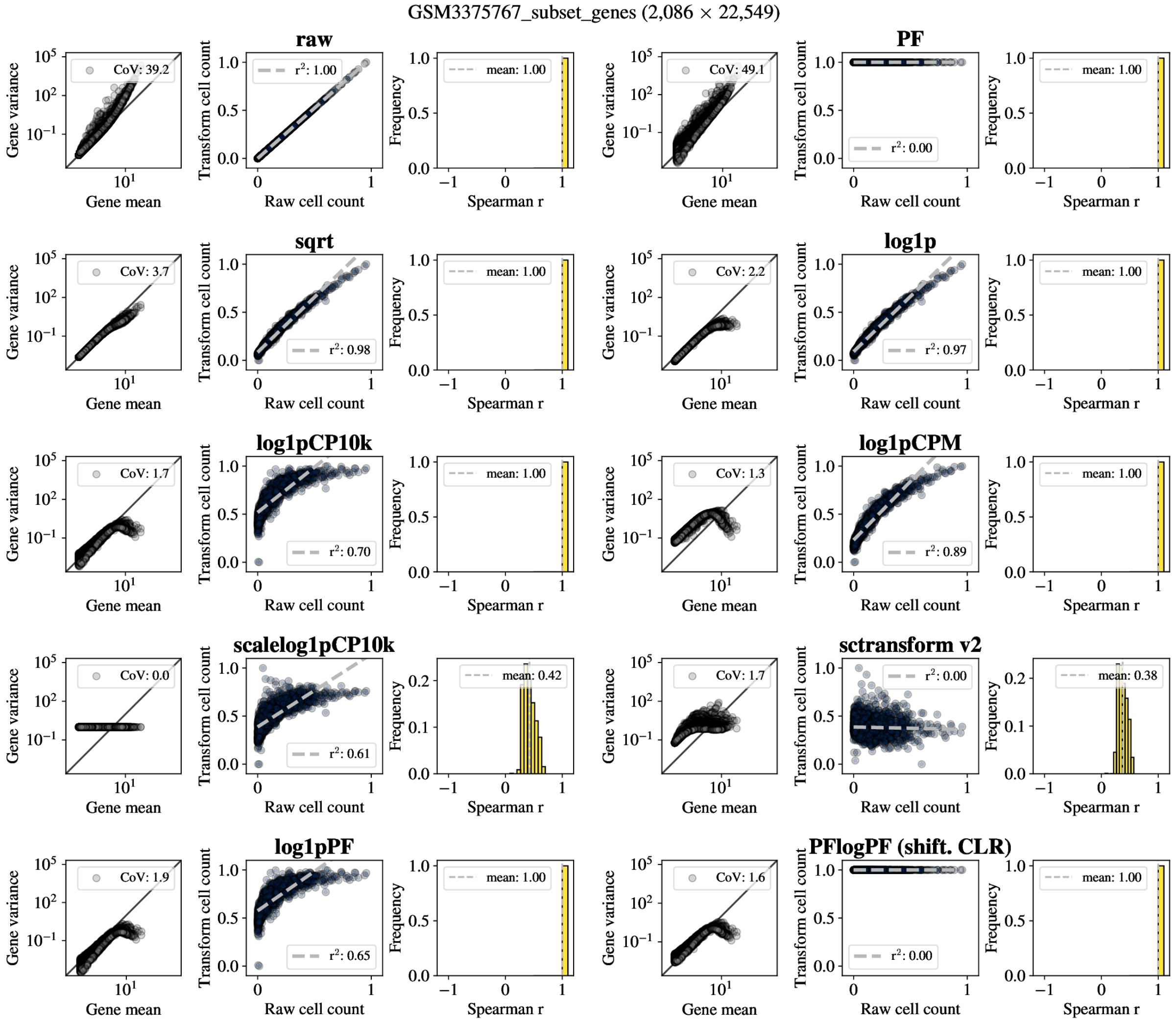


Supplementary Figure 1.212. **GSM3375767**. Per-method comparison of variance stabilization (left), depth normalization (middle), and monotonicity (right) for each normalization on dataset GSM3375767.

| **GSM3377671** | |
|---|---|
| cells | 2,965 |
| genes | 60,623 |
| nonzero entries | 10,879,648 |
| density | 0.0605 |
| total counts | 38,238,063 |
| mean counts / cell | 12,896 |
| mean counts / gene | 631 |
| min counts / cell | 3,997 |
| max counts / cell | 120,661 |
| overdispersion | 0.6 |

GSM3377671_subset_genes (2,965 × 29,131)

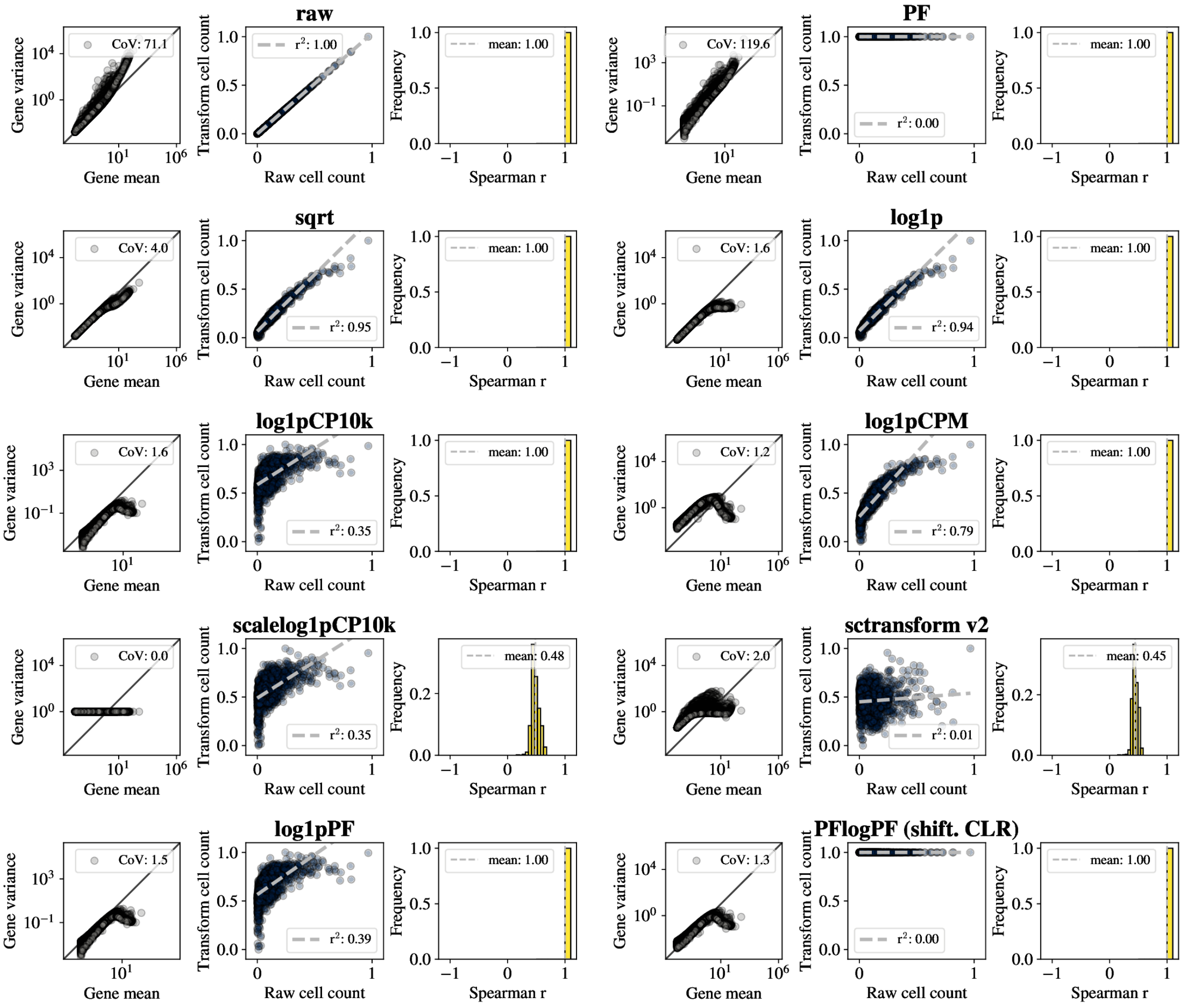


Supplementary Figure 1.213. **GSM3377671**. Per-method comparison of variance stabilization (left), depth normalization (middle), and monotonicity (right) for each normalization on dataset GSM3377671.

| GSM3377672 | |
|---|---|
| cells | 12,262 |
| genes | 60,623 |
| nonzero entries | 17,797,499 |
| density | 0.0239 |
| total counts | 57,080,943 |
| mean counts / cell | 4,655 |
| mean counts / gene | 942 |
| min counts / cell | 178 |
| max counts / cell | 142,110 |
| overdispersion | 2.86 |

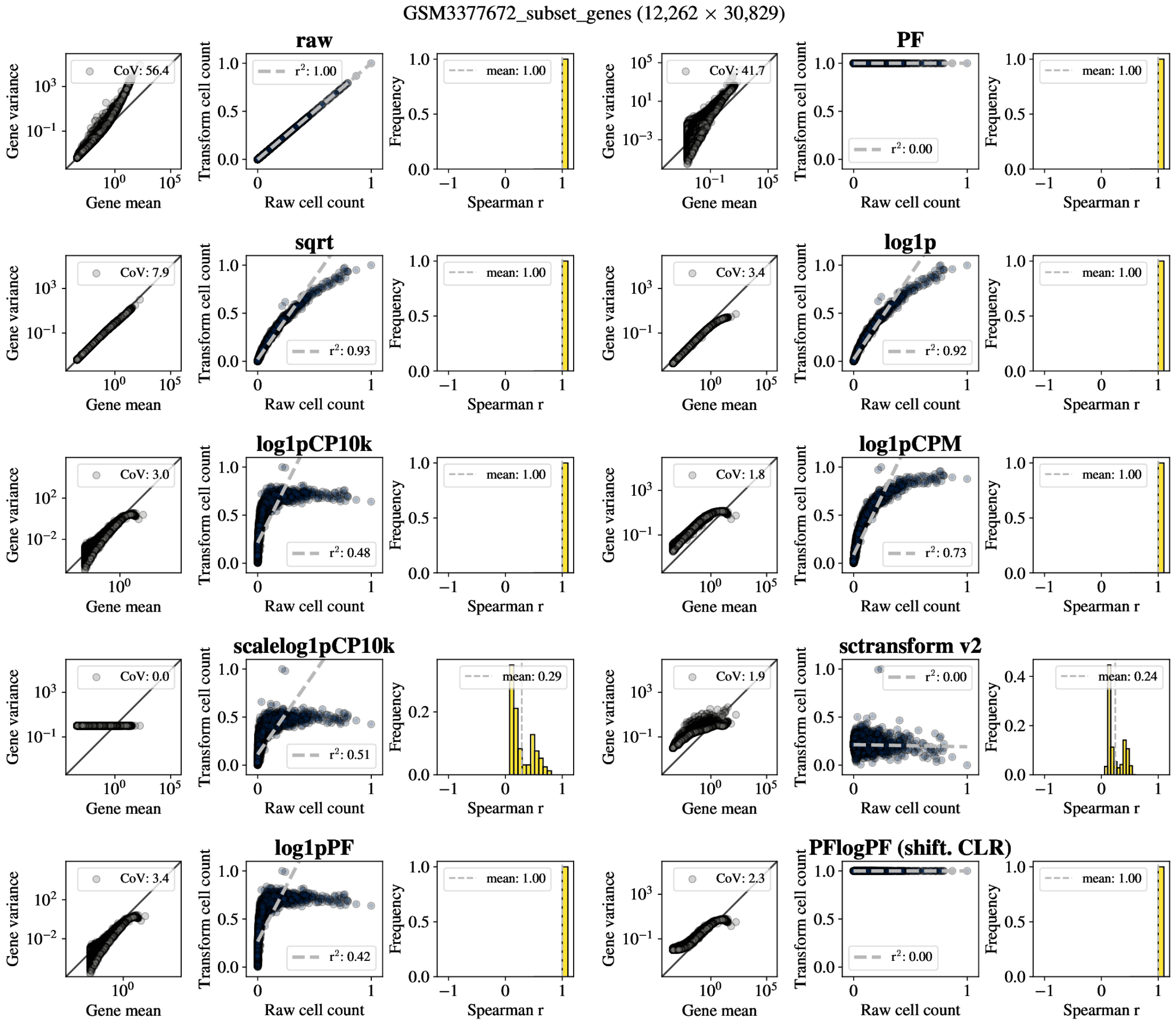


Supplementary Figure 1.214. **GSM3377672**. Per-method comparison of variance stabilization (left), depth normalization (middle), and monotonicity (right) for each normalization on dataset GSM3377672.

| **GSM3378054** | |
|---|---|
| cells | 2,733 |
| genes | 60,623 |
| nonzero entries | 8,307,144 |
| density | 0.0501 |
| total counts | 39,591,942 |
| mean counts / cell | 14,487 |
| mean counts / gene | 653 |
| min counts / cell | 281 |
| max counts / cell | 50,888 |
| overdispersion | 0.389 |

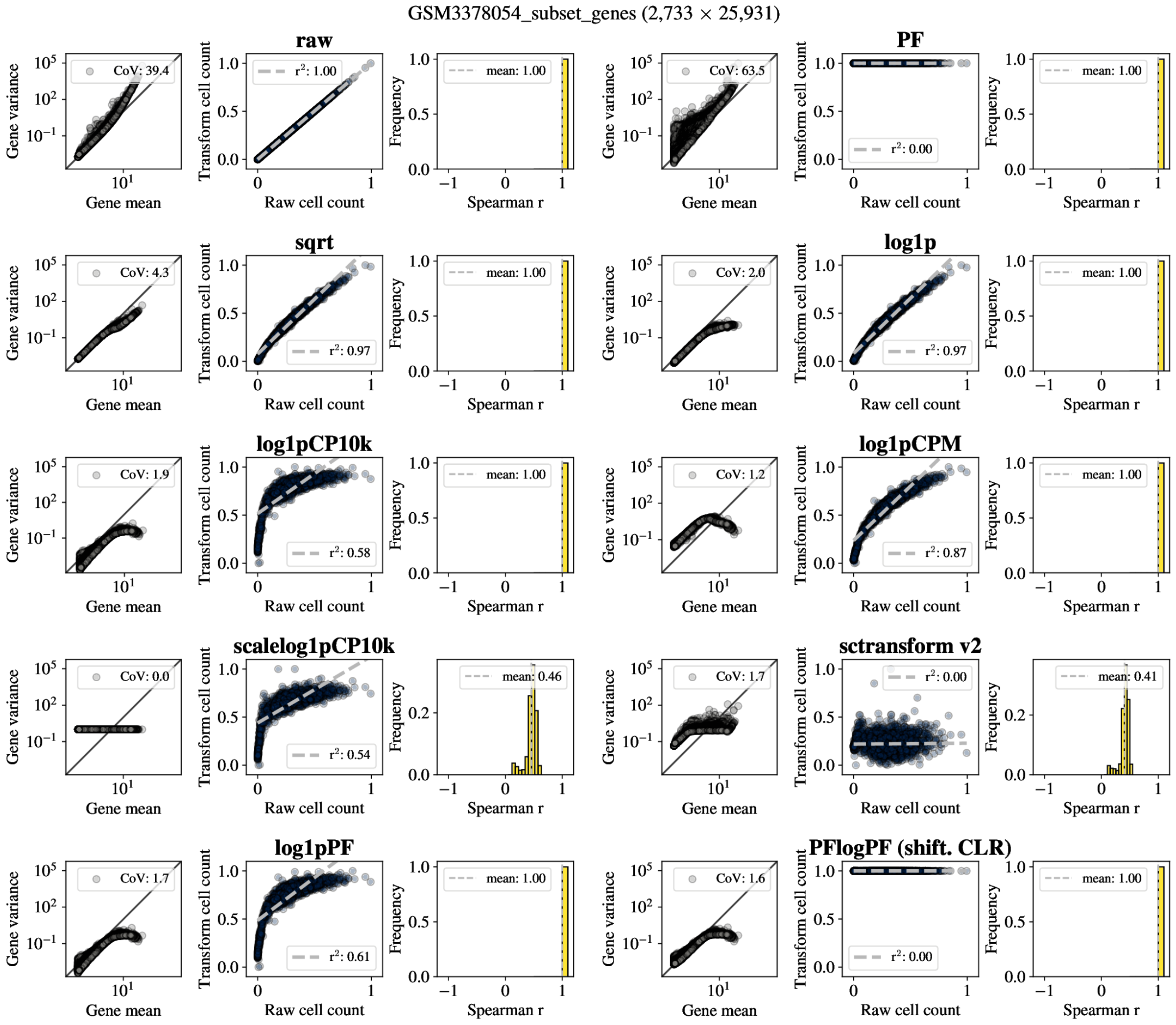


Supplementary Figure 1.215. **GSM3378054**. Per-method comparison of variance stabilization (left), depth normalization (middle), and monotonicity (right) for each normalization on dataset GSM3378054.

| GSM3378055 | |
|---|---|
| cells | 2,771 |
| genes | 60,623 |
| nonzero entries | 8,579,470 |
| density | 0.0511 |
| total counts | 41,126,009 |
| mean counts / cell | 14,842 |
| mean counts / gene | 678 |
| min counts / cell | 331 |
| max counts / cell | 68,877 |
| overdispersion | 0.337 |

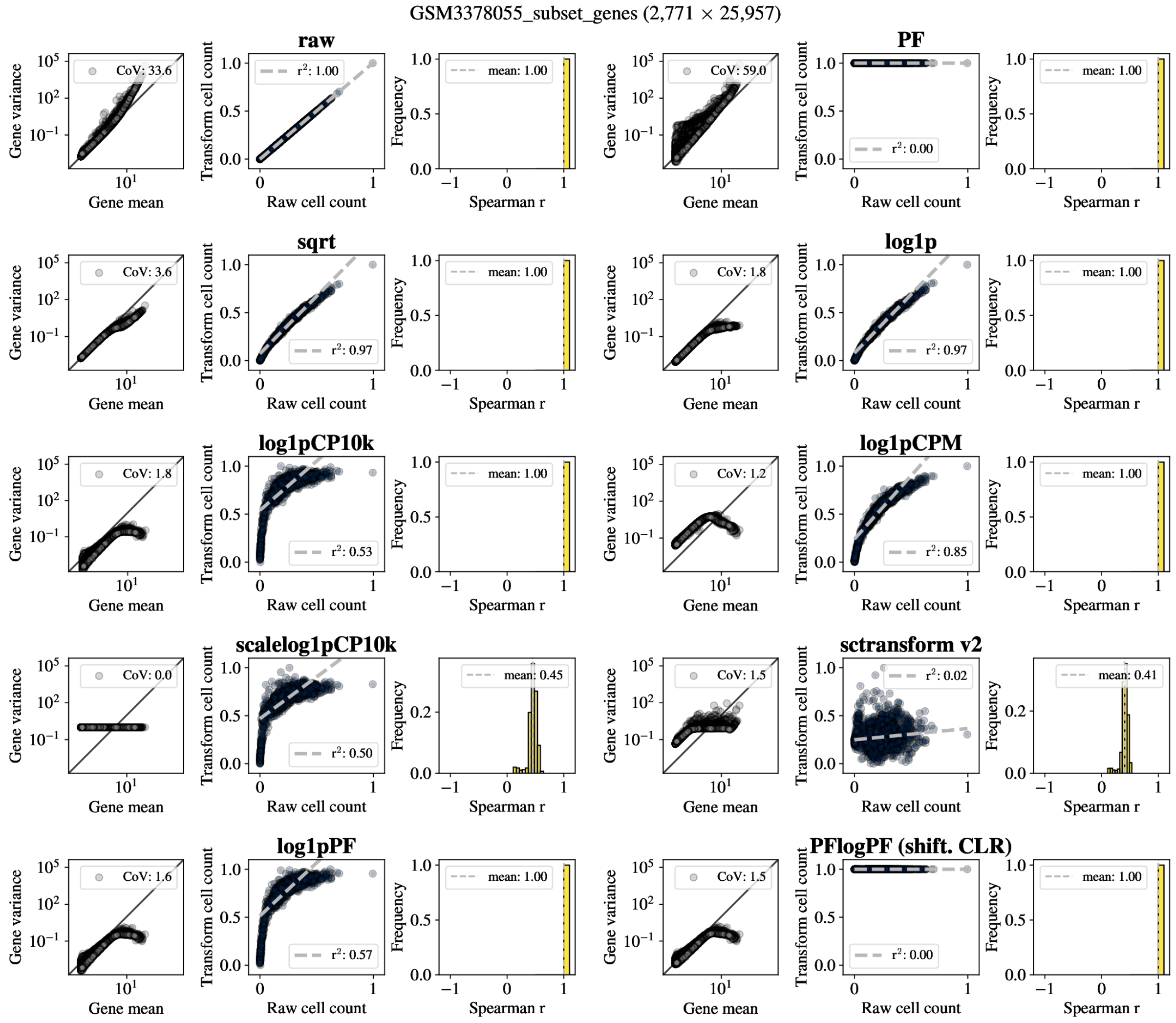


Supplementary Figure 1.216. **GSM3378055**. Per-method comparison of variance stabilization (left), depth normalization (middle), and monotonicity (right) for each normalization on dataset GSM3378055.

| GSM3378056 | |
|---|---|
| cells | 2,263 |
| genes | 60,623 |
| nonzero entries | 5,236,668 |
| density | 0.0382 |
| total counts | 22,451,092 |
| mean counts / cell | 9,921 |
| mean counts / gene | 370 |
| min counts / cell | 330 |
| max counts / cell | 35,601 |
| overdispersion | 0.369 |

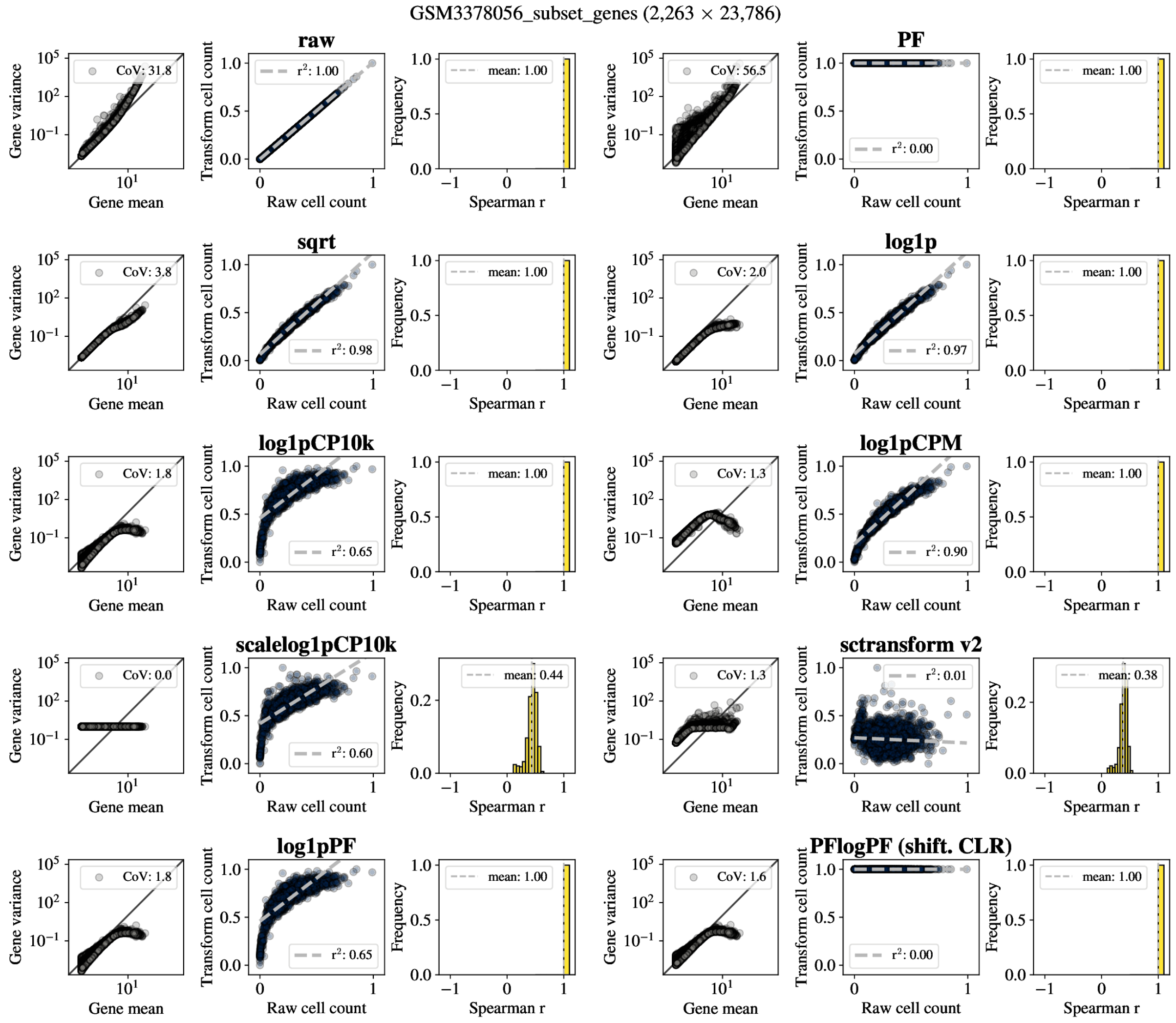


Supplementary Figure 1.217. **GSM3378056**. Per-method comparison of variance stabilization (left), depth normalization (middle), and monotonicity (right) for each normalization on dataset GSM3378056.

| GSM3378057 | |
|---|---|
| cells | 2,393 |
| genes | 60,623 |
| nonzero entries | 7,656,934 |
| density | 0.0528 |
| total counts | 37,208,587 |
| mean counts / cell | 15,549 |
| mean counts / gene | 614 |
| min counts / cell | 309 |
| max counts / cell | 43,953 |
| overdispersion | 0.262 |

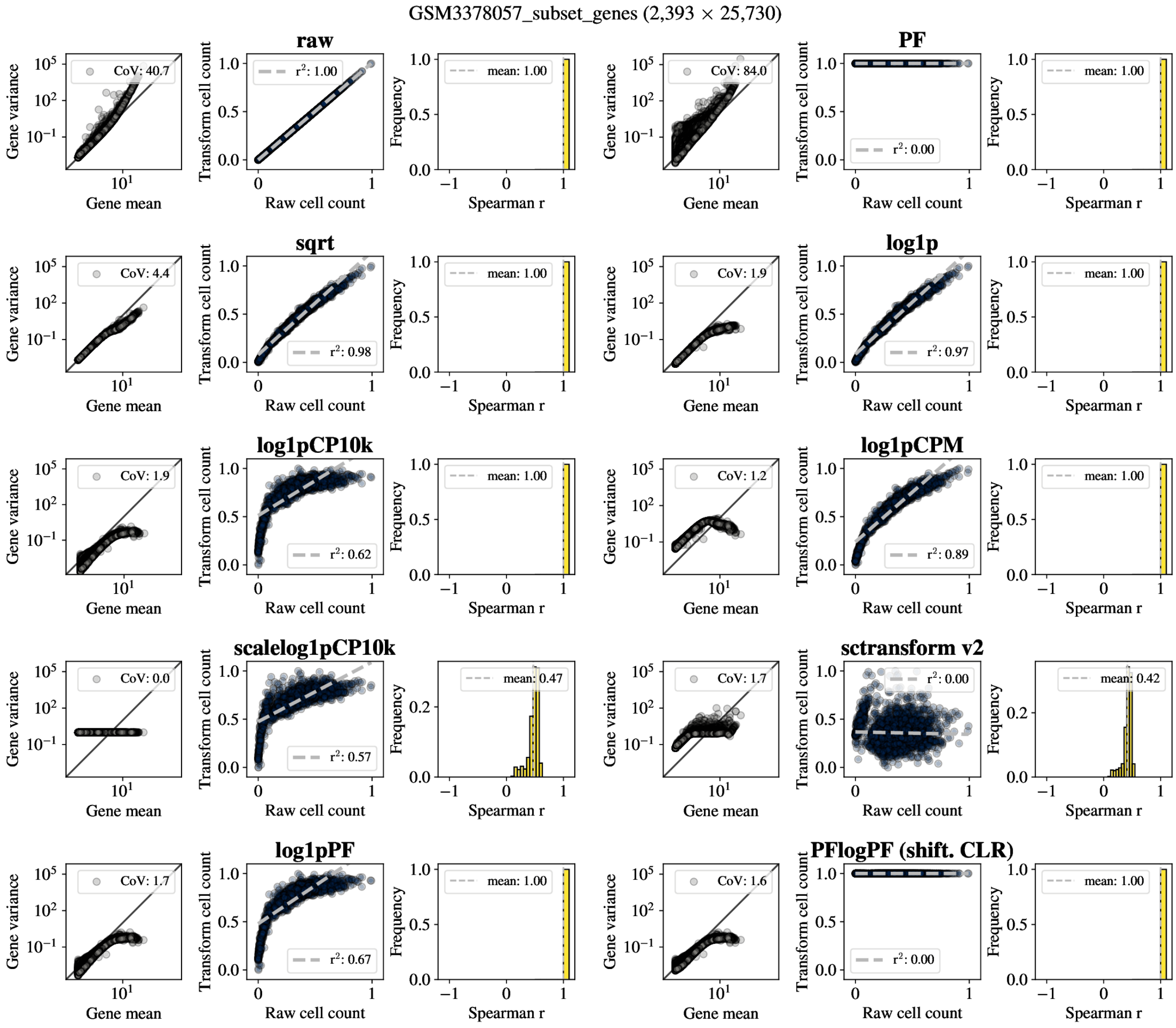


Supplementary Figure 1.218. **GSM3378057**. Per-method comparison of variance stabilization (left), depth normalization (middle), and monotonicity (right) for each normalization on dataset GSM3378057.

| GSM3378058 | |
|---|---|
| cells | 2,562 |
| genes | 60,623 |
| nonzero entries | 8,104,276 |
| density | 0.0522 |
| total counts | 37,454,584 |
| mean counts / cell | 14,619 |
| mean counts / gene | 618 |
| min counts / cell | 388 |
| max counts / cell | 49,553 |
| overdispersion | 0.25 |

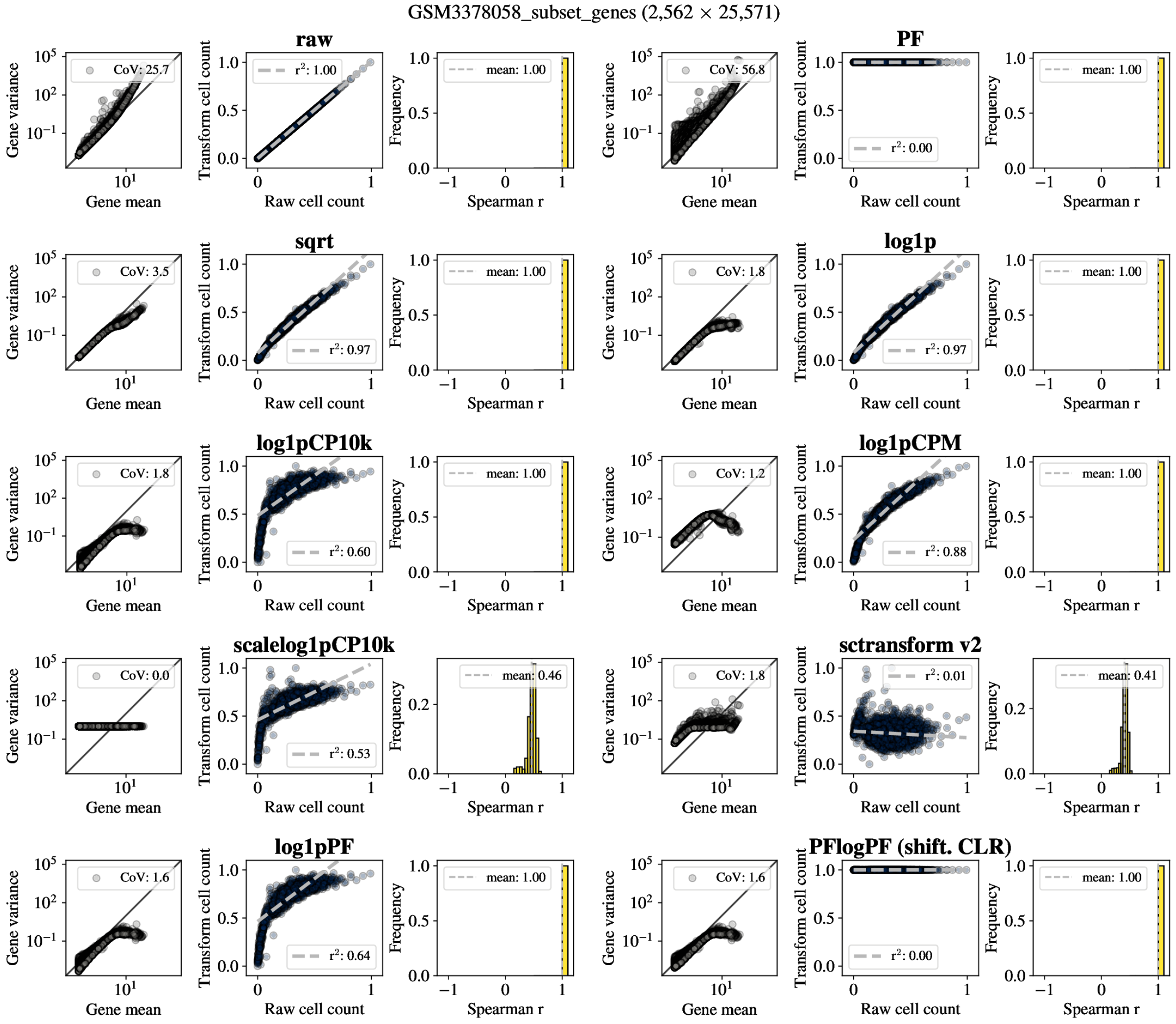


Supplementary Figure 1.219. **GSM3378058**. Per-method comparison of variance stabilization (left), depth normalization (middle), and monotonicity (right) for each normalization on dataset GSM3378058.

| GSM3378059 | |
|---|---|
| cells | 4,081 |
| genes | 60,623 |
| nonzero entries | 11,118,545 |
| density | 0.0449 |
| total counts | 48,306,867 |
| mean counts / cell | 11,837 |
| mean counts / gene | 797 |
| min counts / cell | 323 |
| max counts / cell | 44,225 |
| overdispersion | 0.404 |

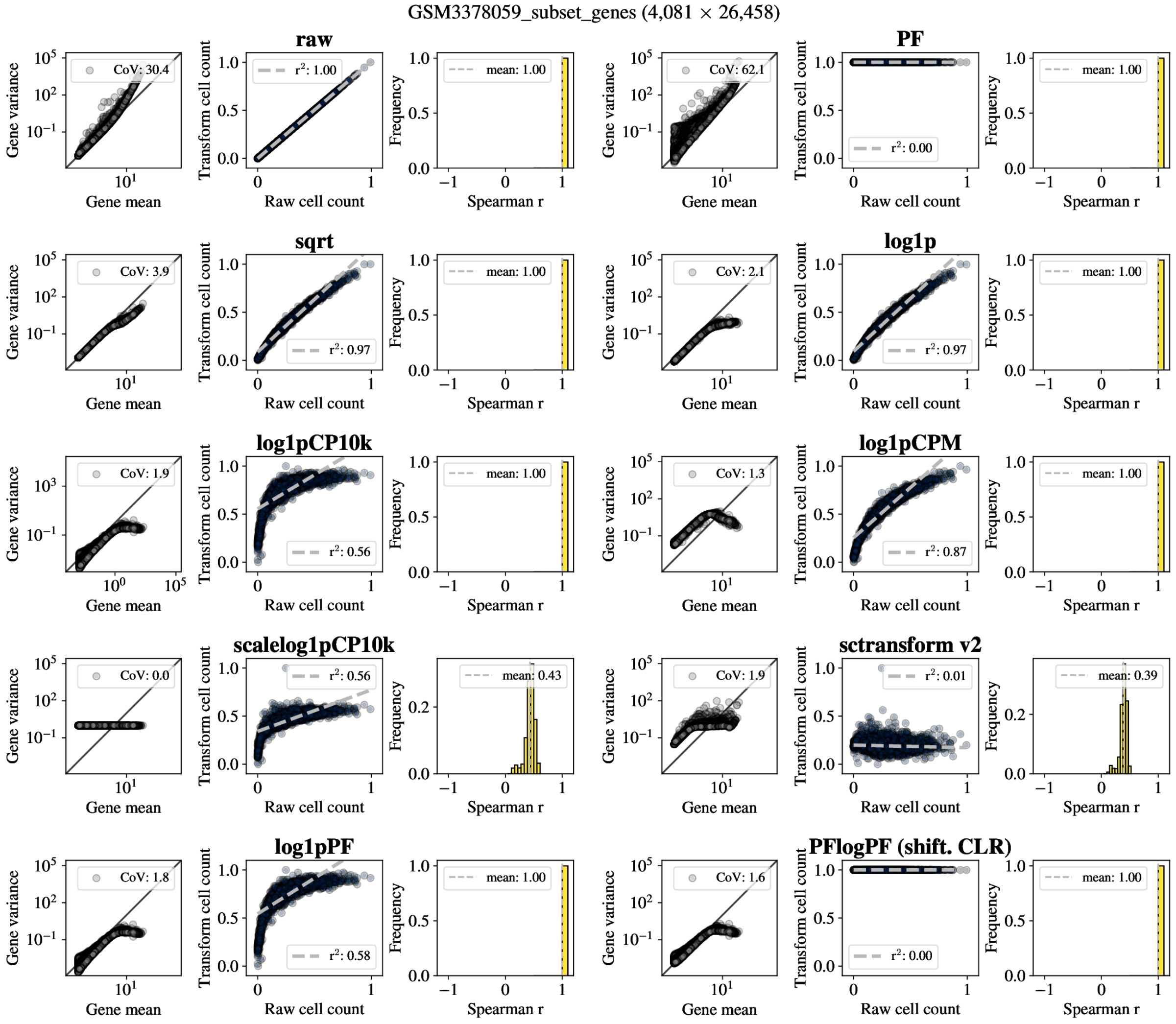


Supplementary Figure 1.220. **GSM3378059**. Per-method comparison of variance stabilization (left), depth normalization (middle), and monotonicity (right) for each normalization on dataset GSM3378059.

| GSM3396161 | |
|---|---|
| cells | 6,992 |
| genes | 60,623 |
| nonzero entries | 9,440,305 |
| density | 0.0223 |
| total counts | 36,326,609 |
| mean counts / cell | 5,195 |
| mean counts / gene | 599 |
| min counts / cell | 303 |
| max counts / cell | 72,269 |
| overdispersion | 9.16 |

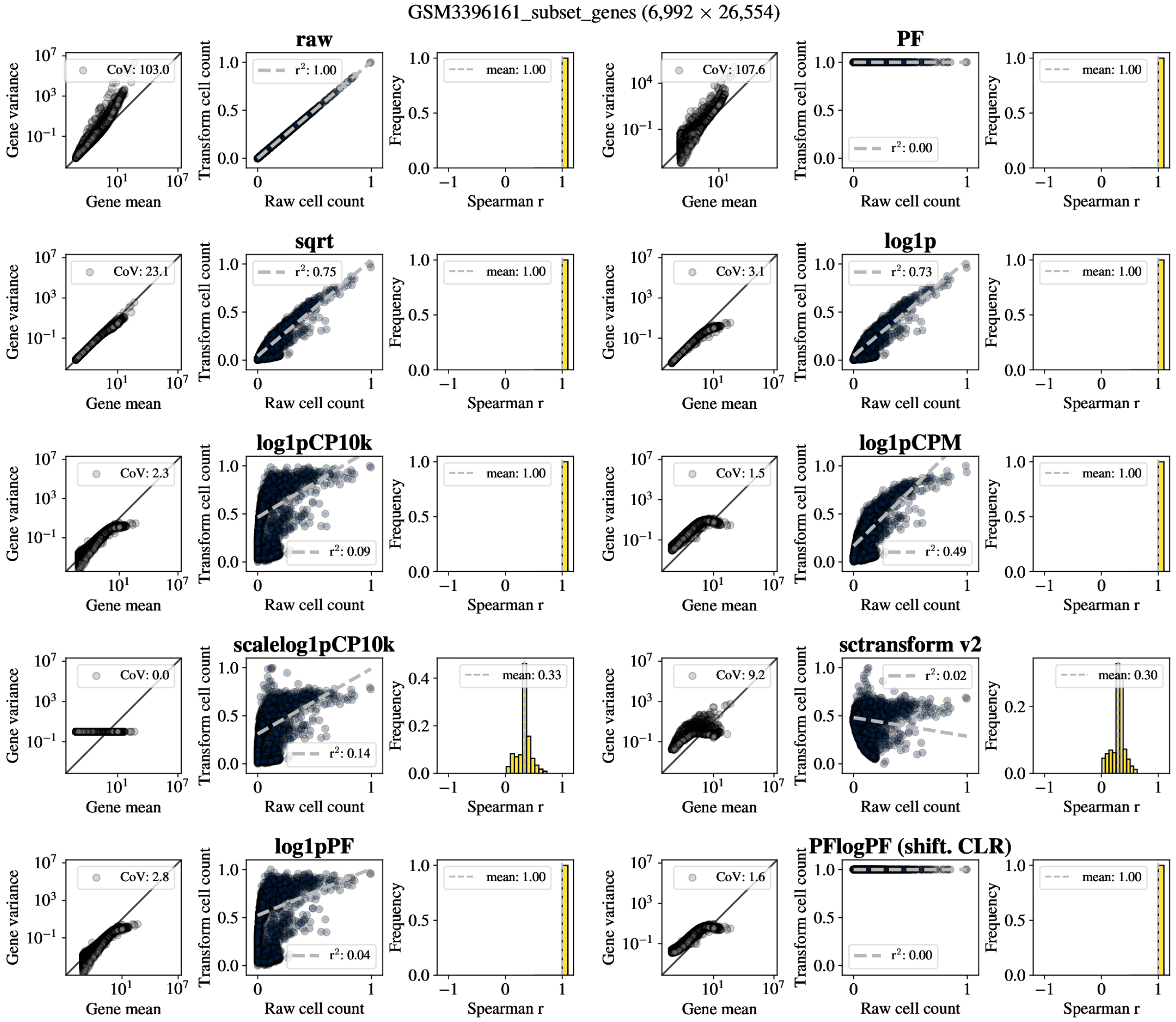


Supplementary Figure 1.221. **GSM3396161**. Per-method comparison of variance stabilization (left), depth normalization (middle), and monotonicity (right) for each normalization on dataset GSM3396161.

| GSM3396162 | |
|---|---|
| cells | 6,425 |
| genes | 60,623 |
| nonzero entries | 6,365,401 |
| density | 0.0163 |
| total counts | 22,497,091 |
| mean counts / cell | 3,501 |
| mean counts / gene | 371 |
| min counts / cell | 883 |
| max counts / cell | 35,518 |
| overdispersion | 8.35 |

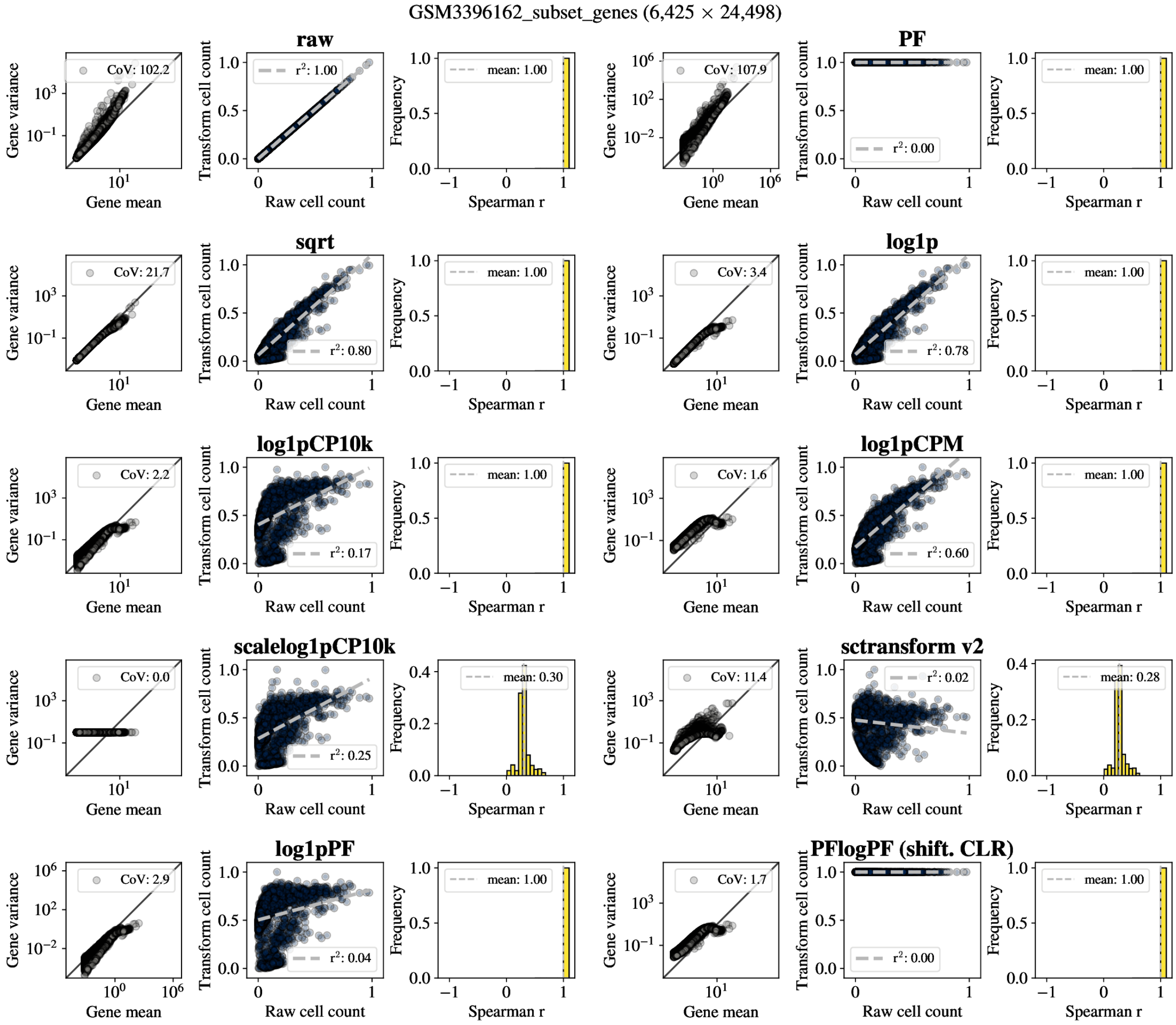


Supplementary Figure 1.222. **GSM3396162**. Per-method comparison of variance stabilization (left), depth normalization (middle), and monotonicity (right) for each normalization on dataset GSM3396162.

| GSM3396163 | |
|---|---|
| cells | 8,775 |
| genes | 60,623 |
| nonzero entries | 6,765,955 |
| density | 0.0127 |
| total counts | 26,810,913 |
| mean counts / cell | 3,055 |
| mean counts / gene | 442 |
| min counts / cell | 572 |
| max counts / cell | 37,925 |
| overdispersion | 3.81 |

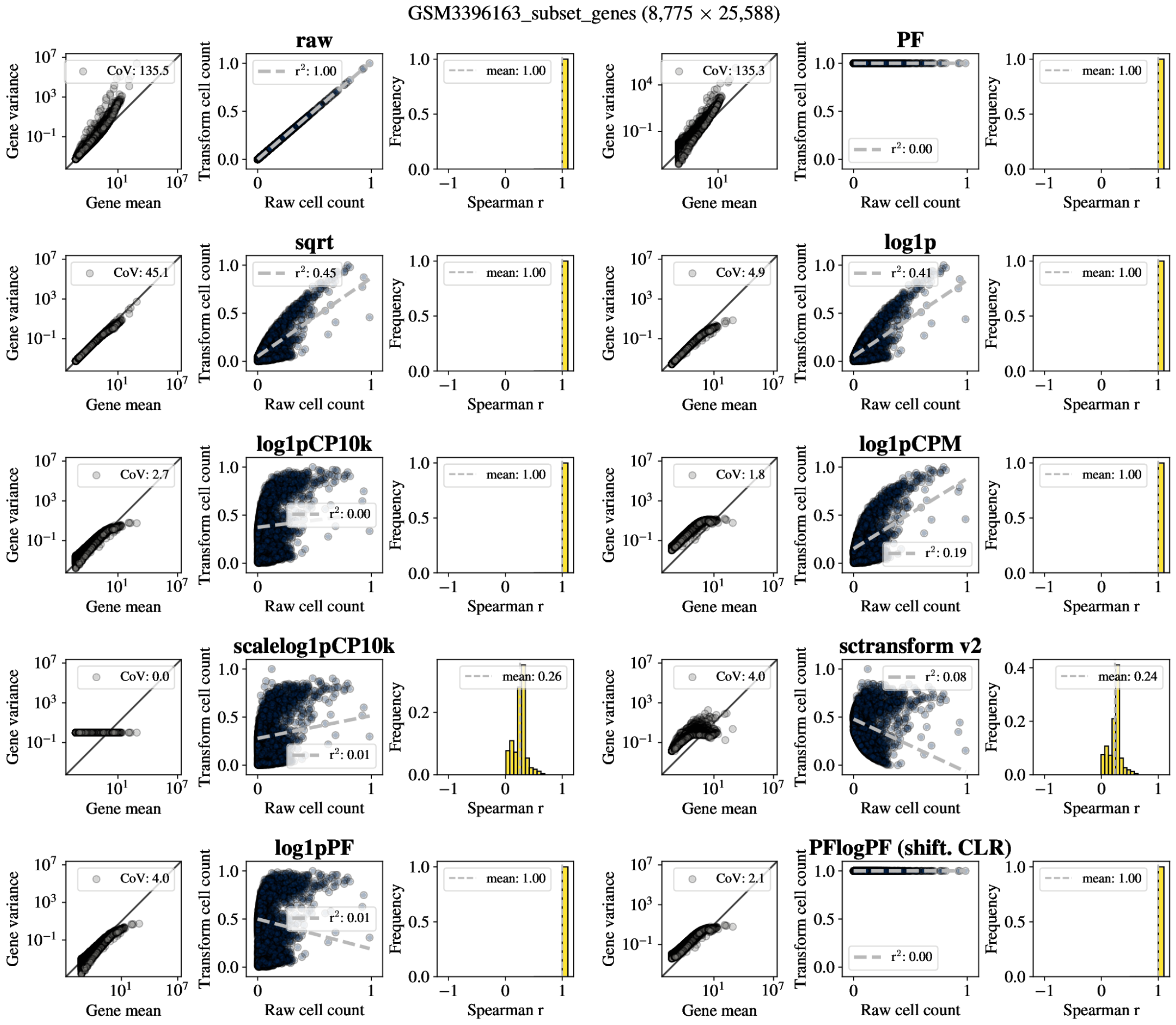


Supplementary Figure 1.223. **GSM3396163**. Per-method comparison of variance stabilization (left), depth normalization (middle), and monotonicity (right) for each normalization on dataset GSM3396163.

| GSM3396164 | |
|---|---|
| cells | 4,092 |
| genes | 60,623 |
| nonzero entries | 1,089,092 |
| density | 0.00439 |
| total counts | 3,918,255 |
| mean counts / cell | 958 |
| mean counts / gene | 65 |
| min counts / cell | 85 |
| max counts / cell | 23,866 |
| overdispersion | 12.6 |

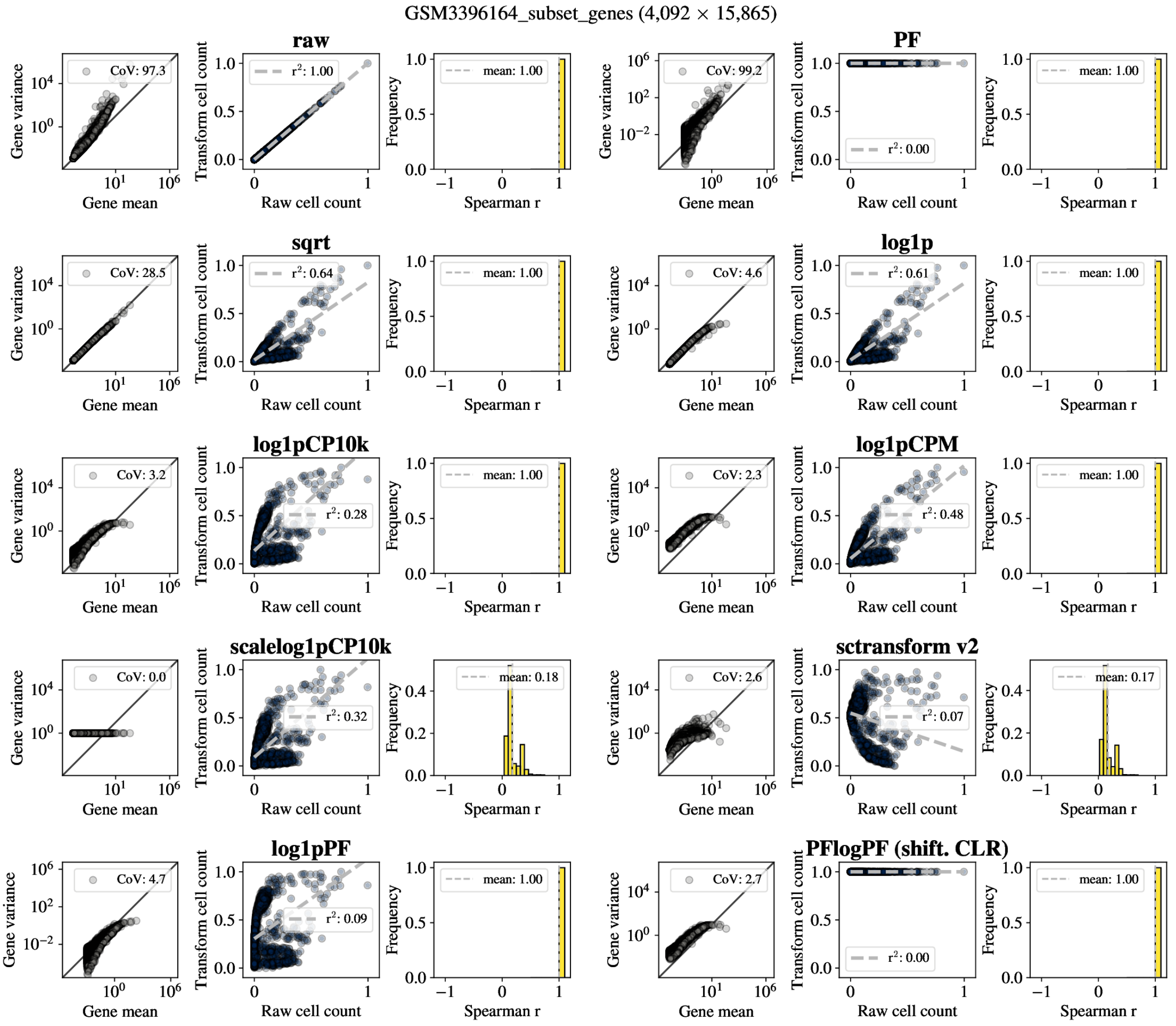


Supplementary Figure 1.224. **GSM3396164**. Per-method comparison of variance stabilization (left), depth normalization (middle), and monotonicity (right) for each normalization on dataset GSM3396164.

| GSM3396165 | |
|---|---|
| cells | 9,412 |
| genes | 60,623 |
| nonzero entries | 5,850,621 |
| density | 0.0103 |
| total counts | 22,899,579 |
| mean counts / cell | 2,433 |
| mean counts / gene | 378 |
| min counts / cell | 88 |
| max counts / cell | 40,691 |
| overdispersion | 4.97 |

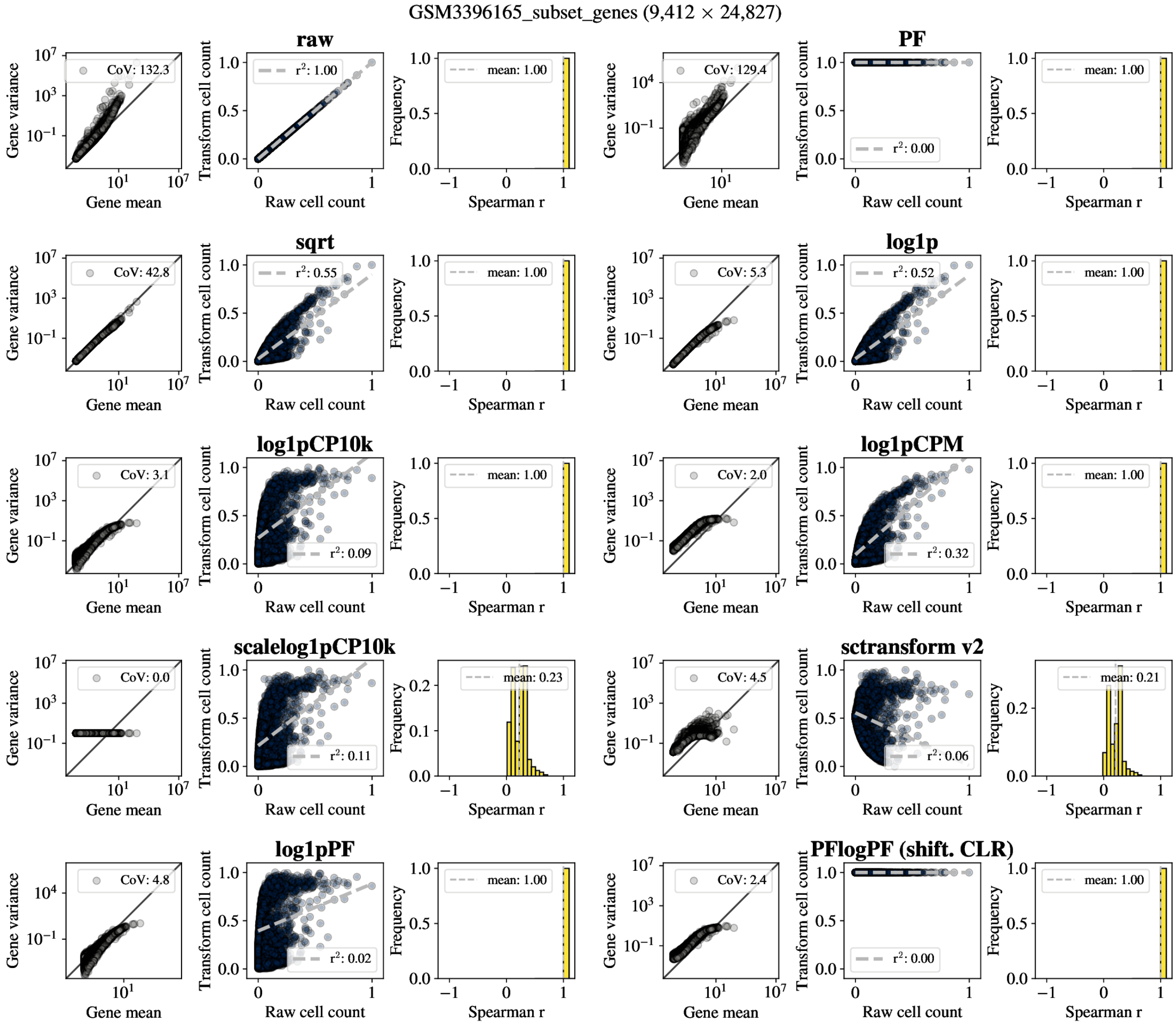


Supplementary Figure 1.225. **GSM3396165**. Per-method comparison of variance stabilization (left), depth normalization (middle), and monotonicity (right) for each normalization on dataset GSM3396165.

| GSM3396166 | |
|---|---|
| cells | 2,103 |
| genes | 60,623 |
| nonzero entries | 1,411,569 |
| density | 0.0111 |
| total counts | 12,636,725 |
| mean counts / cell | 6,009 |
| mean counts / gene | 208 |
| min counts / cell | 3,240 |
| max counts / cell | 40,091 |
| overdispersion | 0.302 |

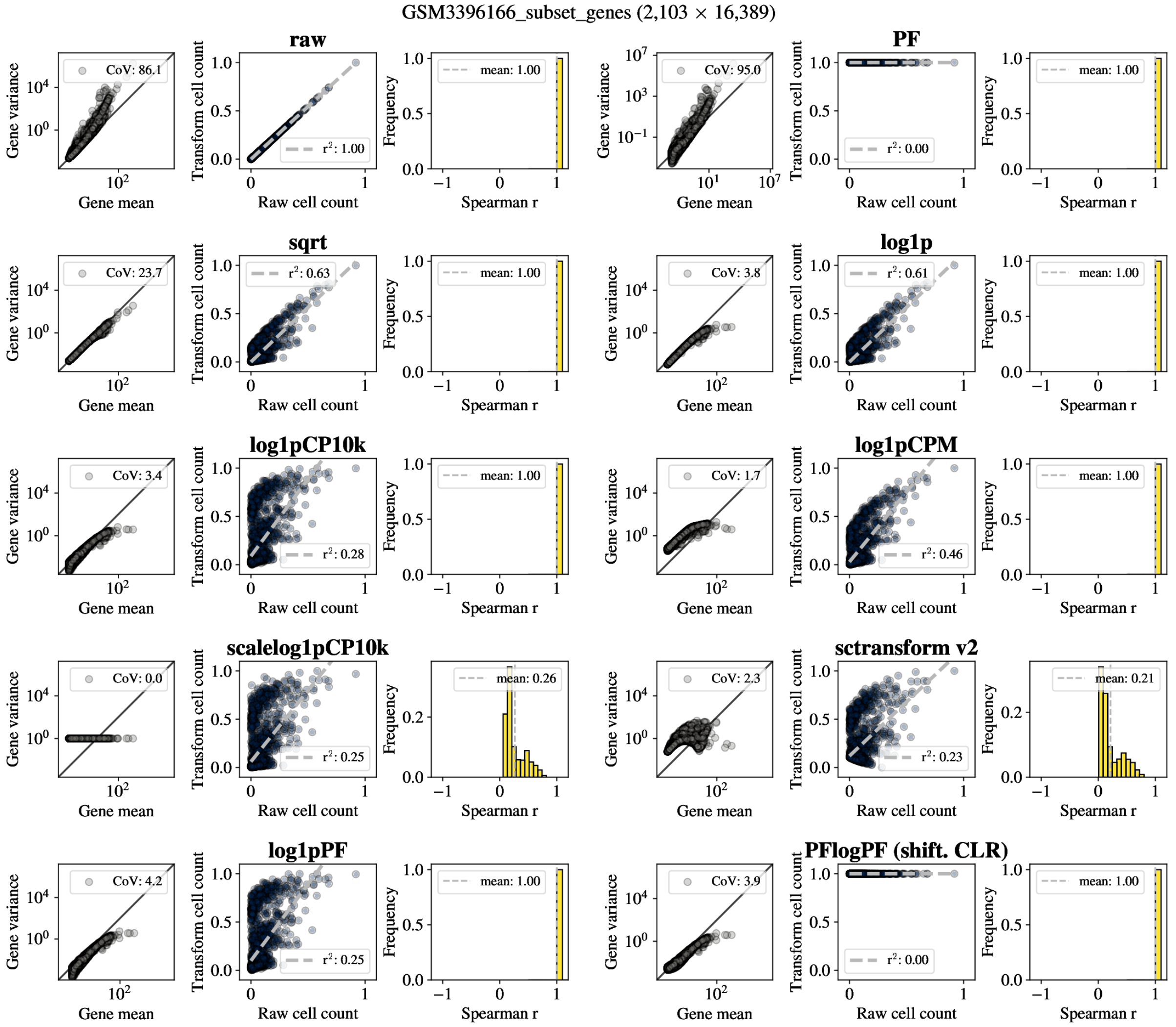


Supplementary Figure 1.226. **GSM3396166**. Per-method comparison of variance stabilization (left), depth normalization (middle), and monotonicity (right) for each normalization on dataset GSM3396166.

| GSM3396167 | |
|---|---|
| cells | 1,379 |
| genes | 60,623 |
| nonzero entries | 2,006,812 |
| density | 0.024 |
| total counts | 9,537,590 |
| mean counts / cell | 6,916 |
| mean counts / gene | 157 |
| min counts / cell | 3,533 |
| max counts / cell | 34,354 |
| overdispersion | 2.37 |

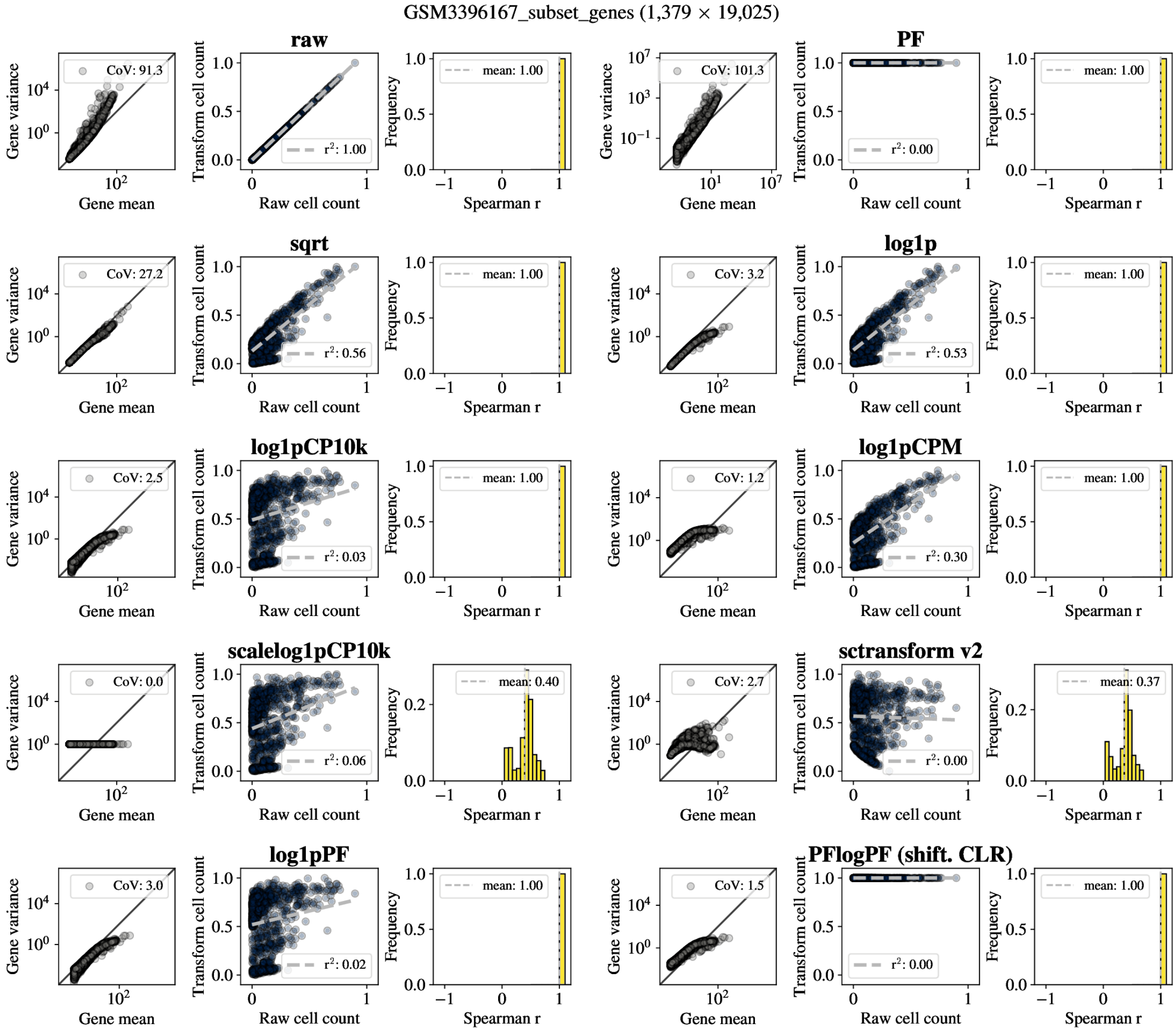


Supplementary Figure 1.227. **GSM3396167**. Per-method comparison of variance stabilization (left), depth normalization (middle), and monotonicity (right) for each normalization on dataset GSM3396167.

| GSM3396168 | |
|---|---|
| cells | 6,594 |
| genes | 60,623 |
| nonzero entries | 4,786,098 |
| density | 0.012 |
| total counts | 34,691,368 |
| mean counts / cell | 5,261 |
| mean counts / gene | 572 |
| min counts / cell | 143 |
| max counts / cell | 55,816 |
| overdispersion | 1.03 |

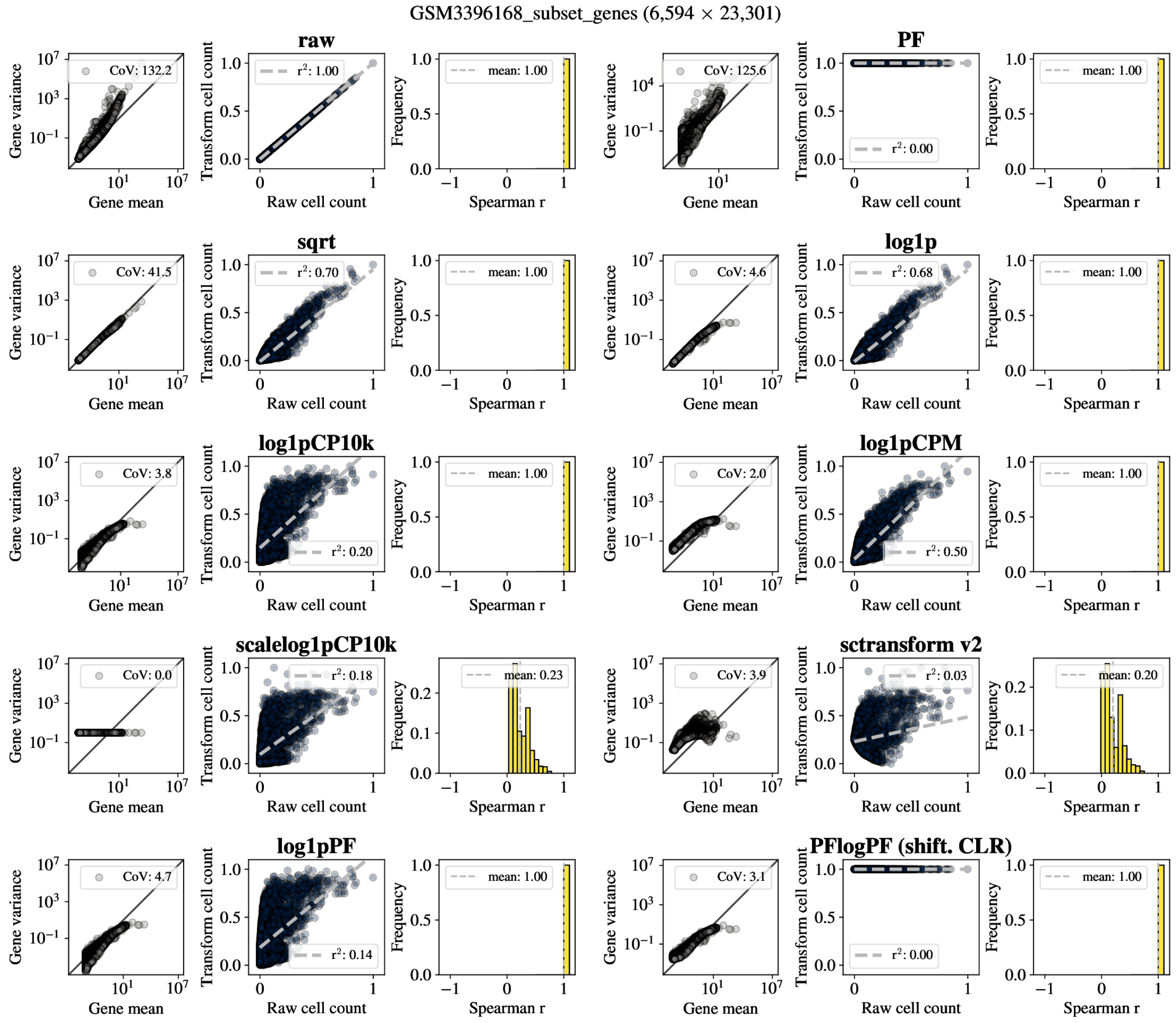


Supplementary Figure 1.228. **GSM3396168**. Per-method comparison of variance stabilization (left), depth normalization (middle), and monotonicity (right) for each normalization on dataset GSM3396168.

| GSM3396169 | |
|---|---|
| cells | 5,893 |
| genes | 60,623 |
| nonzero entries | 5,667,813 |
| density | 0.0159 |
| total counts | 24,820,029 |
| mean counts / cell | 4,212 |
| mean counts / gene | 409 |
| min counts / cell | 203 |
| max counts / cell | 54,819 |
| overdispersion | 4.59 |

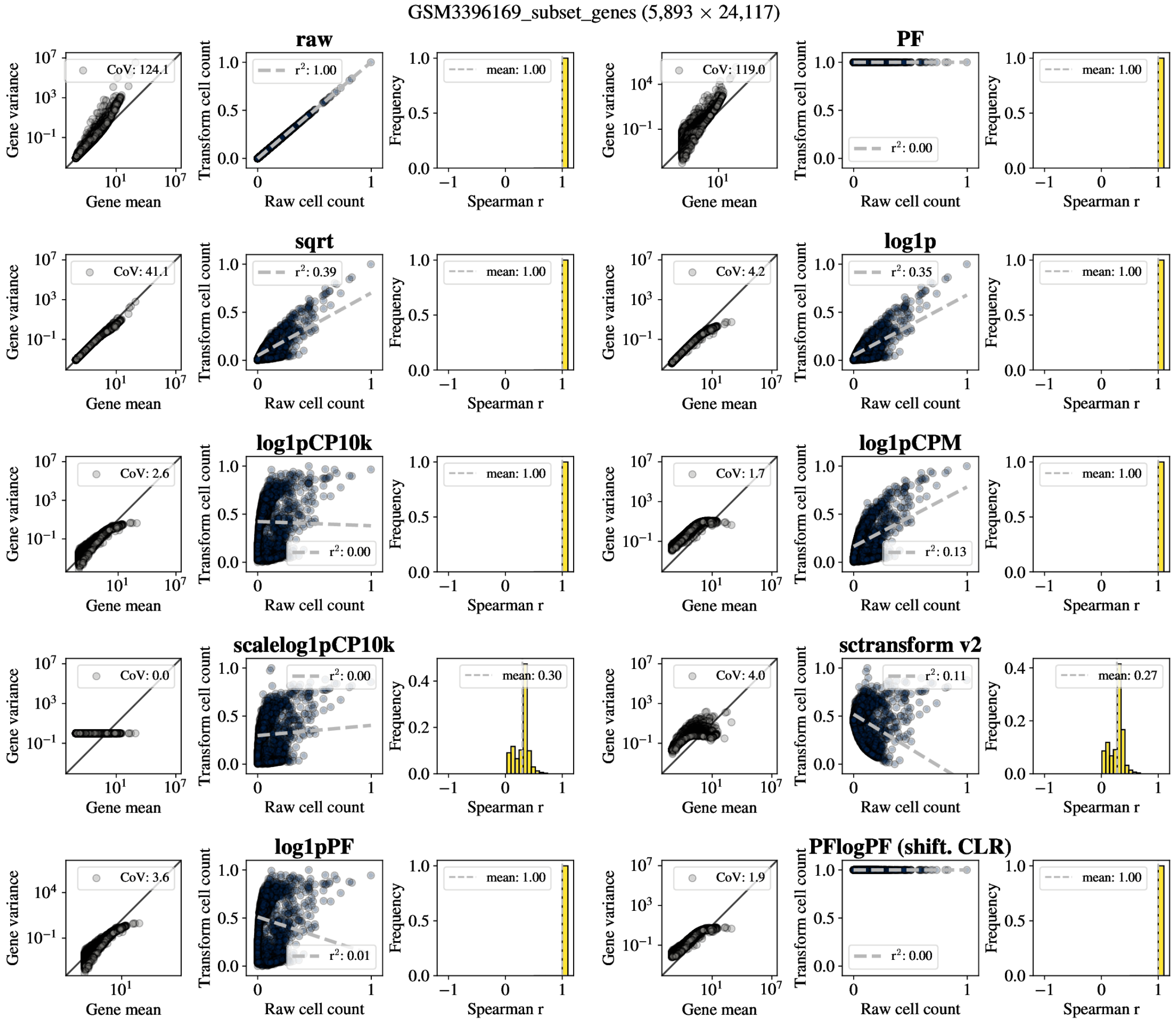


Supplementary Figure 1.229. **GSM3396169**. Per-method comparison of variance stabilization (left), depth normalization (middle), and monotonicity (right) for each normalization on dataset GSM3396169.

| GSM3396170 | |
|---|---|
| cells | 5,000 |
| genes | 60,623 |
| nonzero entries | 6,136,233 |
| density | 0.0202 |
| total counts | 25,872,455 |
| mean counts / cell | 5,174 |
| mean counts / gene | 427 |
| min counts / cell | 1,590 |
| max counts / cell | 63,436 |
| overdispersion | 4.61 |

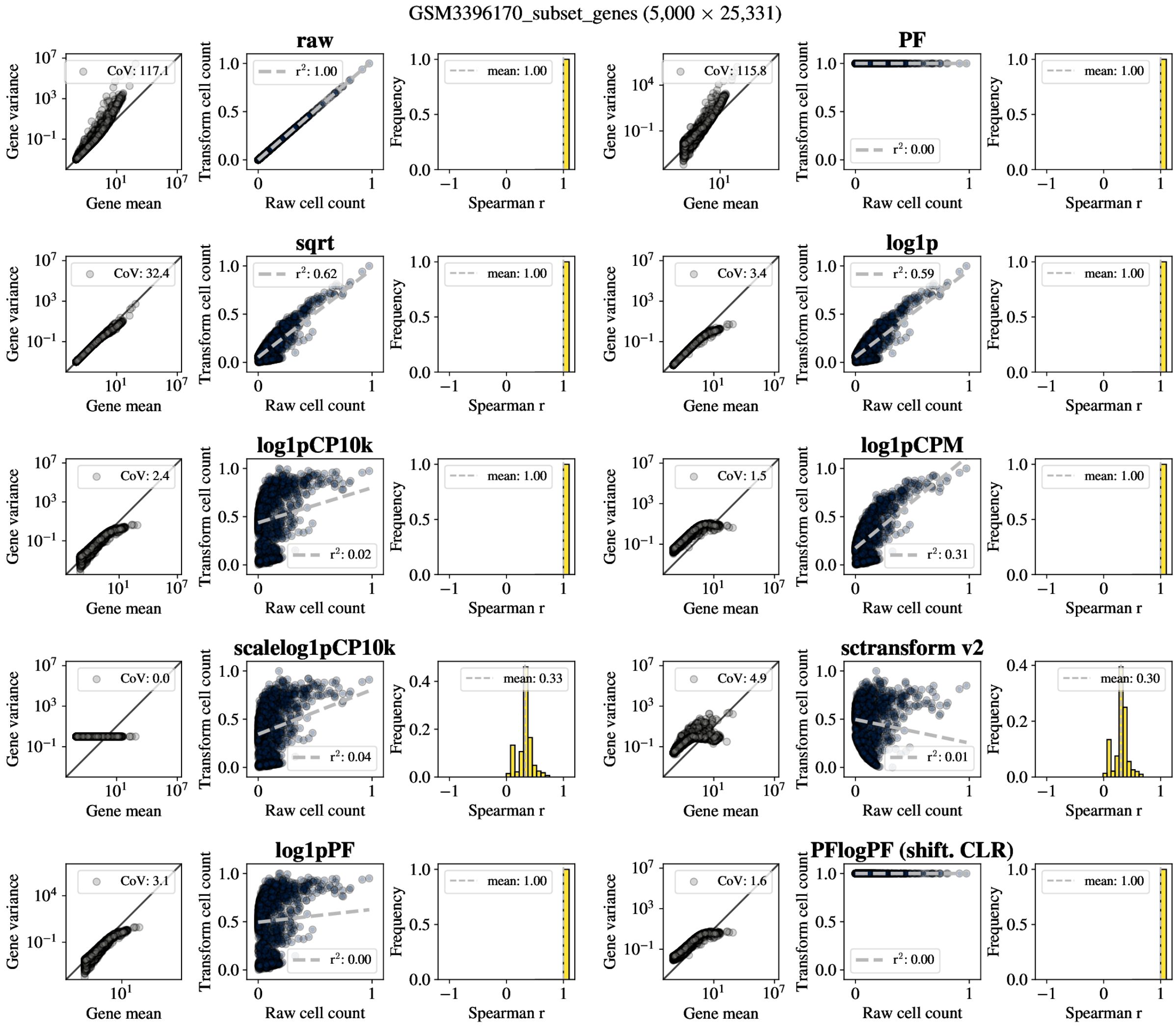


Supplementary Figure 1.230. **GSM3396170**. Per-method comparison of variance stabilization (left), depth normalization (middle), and monotonicity (right) for each normalization on dataset GSM3396170.

| GSM3396171 | |
|---|---|
| cells | 869 |
| genes | 60,623 |
| nonzero entries | 1,461,625 |
| density | 0.0277 |
| total counts | 7,287,420 |
| mean counts / cell | 8,386 |
| mean counts / gene | 120 |
| min counts / cell | 5,118 |
| max counts / cell | 51,417 |
| overdispersion | 2.15 |

GSM3396171_subset_genes (869 × 18,019)

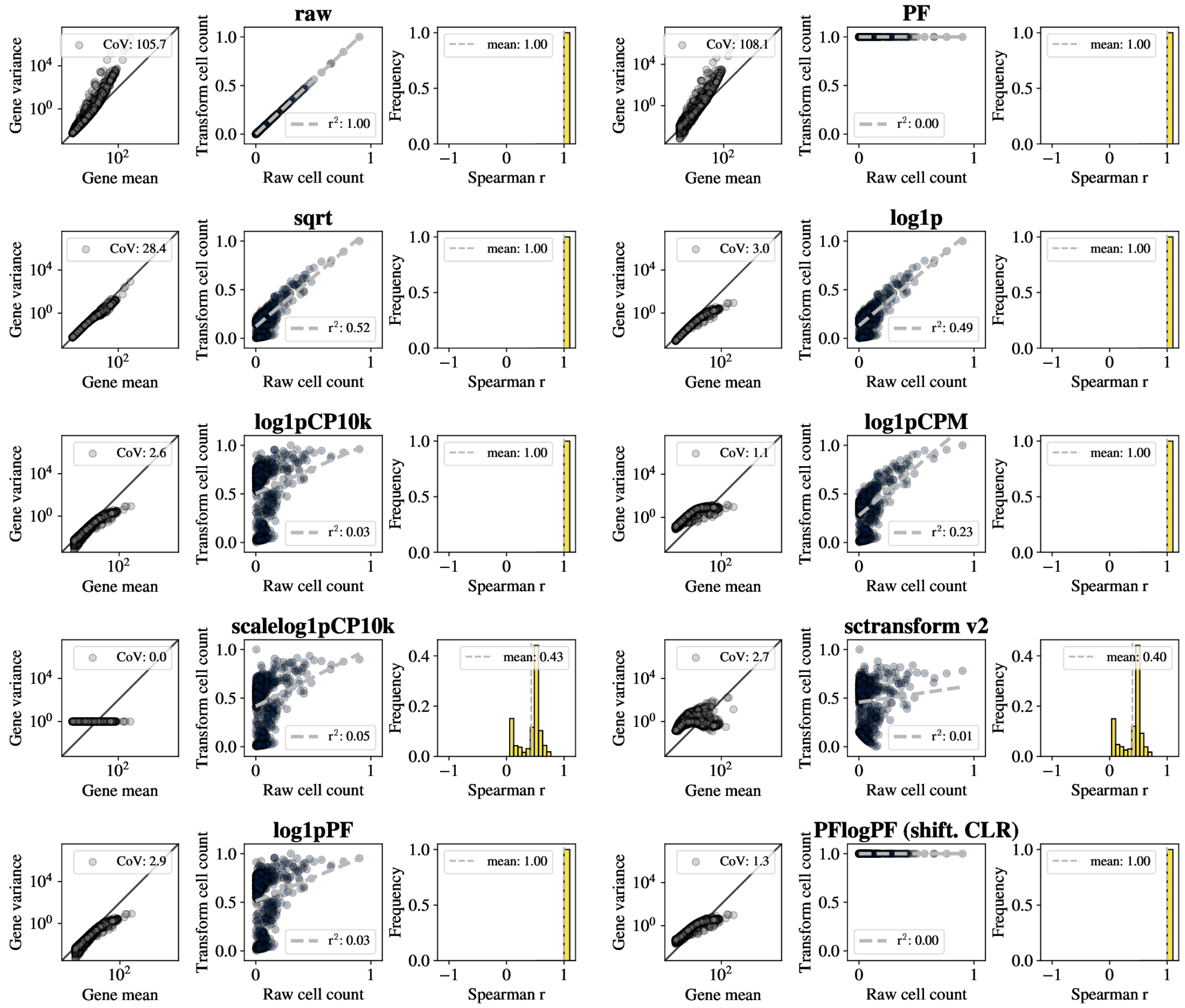


Supplementary Figure 1.231. **GSM3396171**. Per-method comparison of variance stabilization (left), depth normalization (middle), and monotonicity (right) for each normalization on dataset GSM3396171.

| GSM3396172 | |
|---|---|
| cells | 4,827 |
| genes | 60,623 |
| nonzero entries | 5,626,024 |
| density | 0.0192 |
| total counts | 18,463,603 |
| mean counts / cell | 3,825 |
| mean counts / gene | 305 |
| min counts / cell | 1,360 |
| max counts / cell | 42,624 |
| overdispersion | 7.78 |

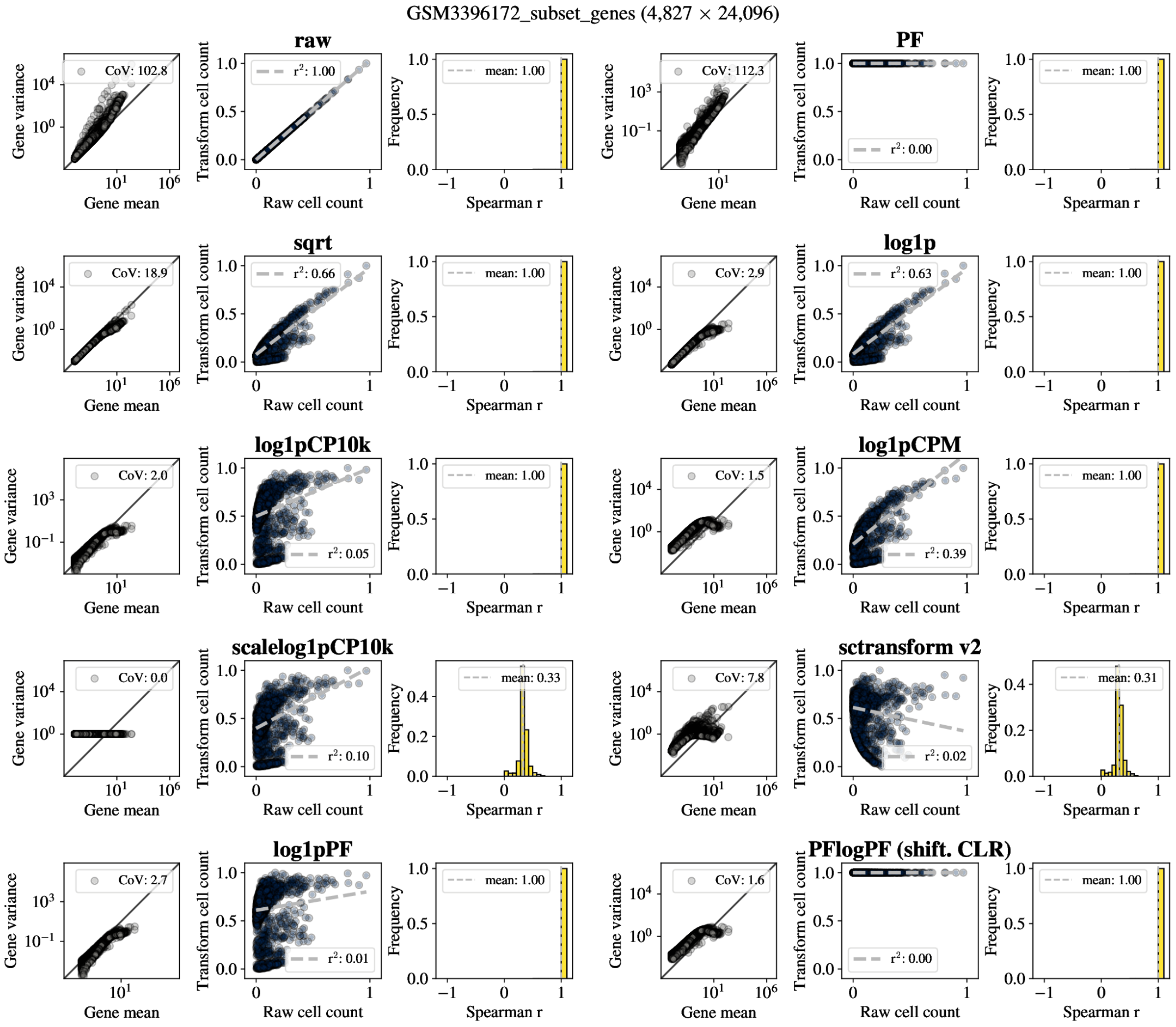


Supplementary Figure 1.232. **GSM3396172**. Per-method comparison of variance stabilization (left), depth normalization (middle), and monotonicity (right) for each normalization on dataset GSM3396172.

| GSM3396173 | |
|---|---|
| cells | 9,198 |
| genes | 60,623 |
| nonzero entries | 5,991,644 |
| density | 0.0107 |
| total counts | 29,223,580 |
| mean counts / cell | 3,177 |
| mean counts / gene | 482 |
| min counts / cell | 85 |
| max counts / cell | 48,765 |
| overdispersion | 4.25 |

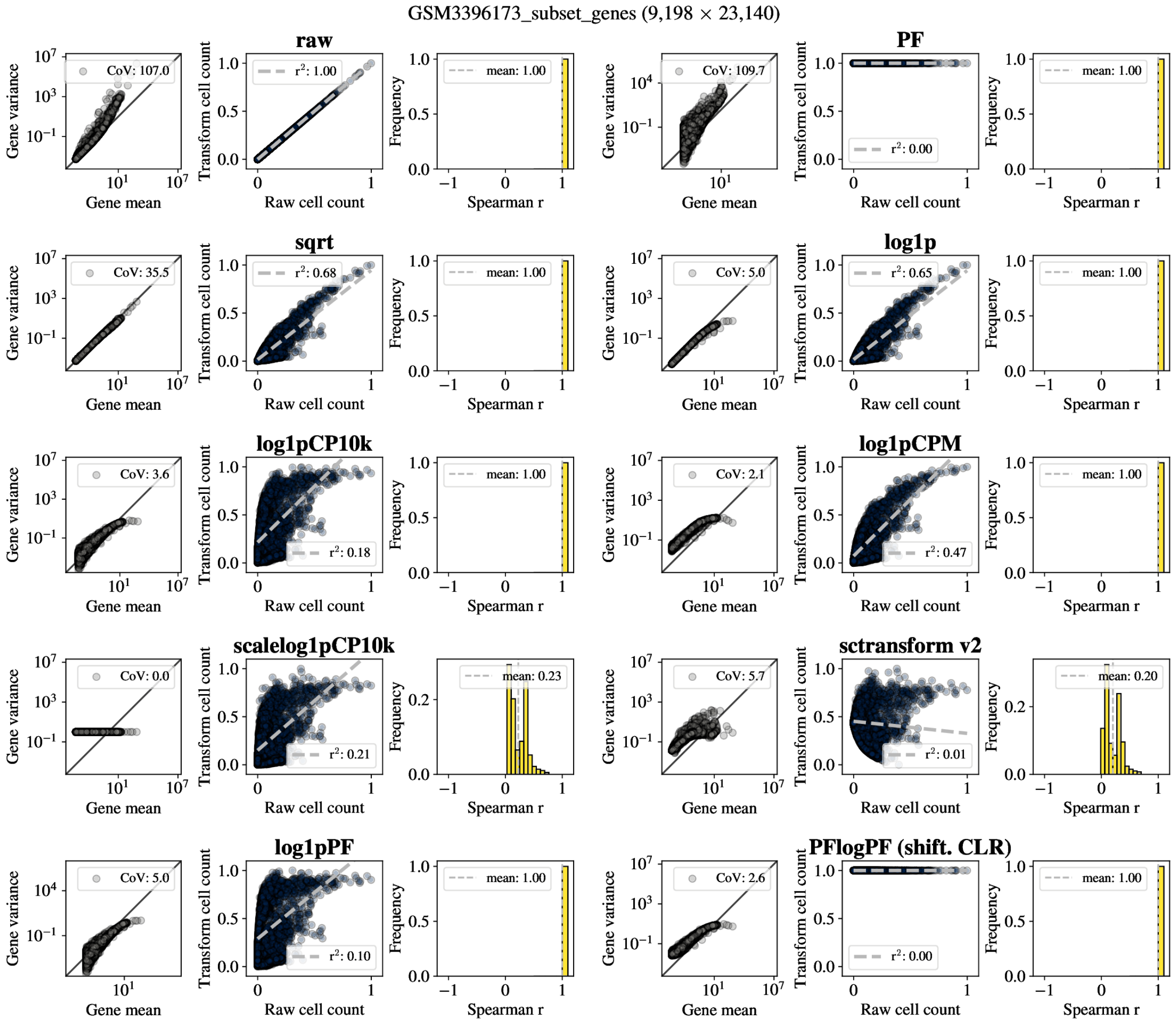


Supplementary Figure 1.233. **GSM3396173**. Per-method comparison of variance stabilization (left), depth normalization (middle), and monotonicity (right) for each normalization on dataset GSM3396173.

| GSM3396174 | |
|---|---|
| cells | 5,675 |
| genes | 60,623 |
| nonzero entries | 6,260,074 |
| density | 0.0182 |
| total counts | 20,541,259 |
| mean counts / cell | 3,620 |
| mean counts / gene | 339 |
| min counts / cell | 1,278 |
| max counts / cell | 45,440 |
| overdispersion | 11.2 |

GSM3396174_subset_genes (5,675 × 24,552)

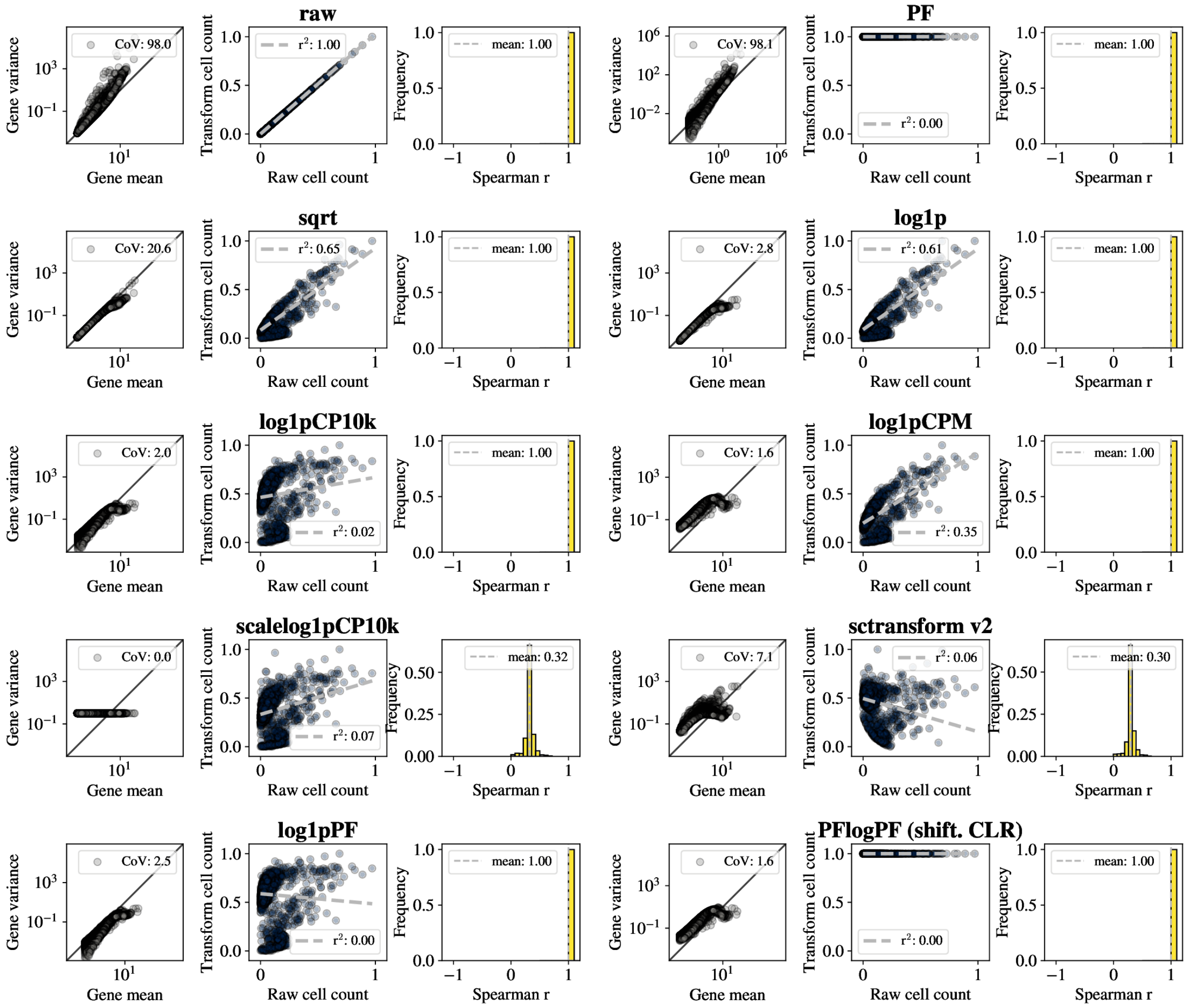


Supplementary Figure 1.234. **GSM3396174**. Per-method comparison of variance stabilization (left), depth normalization (middle), and monotonicity (right) for each normalization on dataset GSM3396174.

| GSM3396175 | |
|---|---|
| cells | 6,018 |
| genes | 60,623 |
| nonzero entries | 6,942,589 |
| density | 0.019 |
| total counts | 21,012,077 |
| mean counts / cell | 3,492 |
| mean counts / gene | 347 |
| min counts / cell | 1,315 |
| max counts / cell | 41,413 |
| overdispersion | 1.4 |

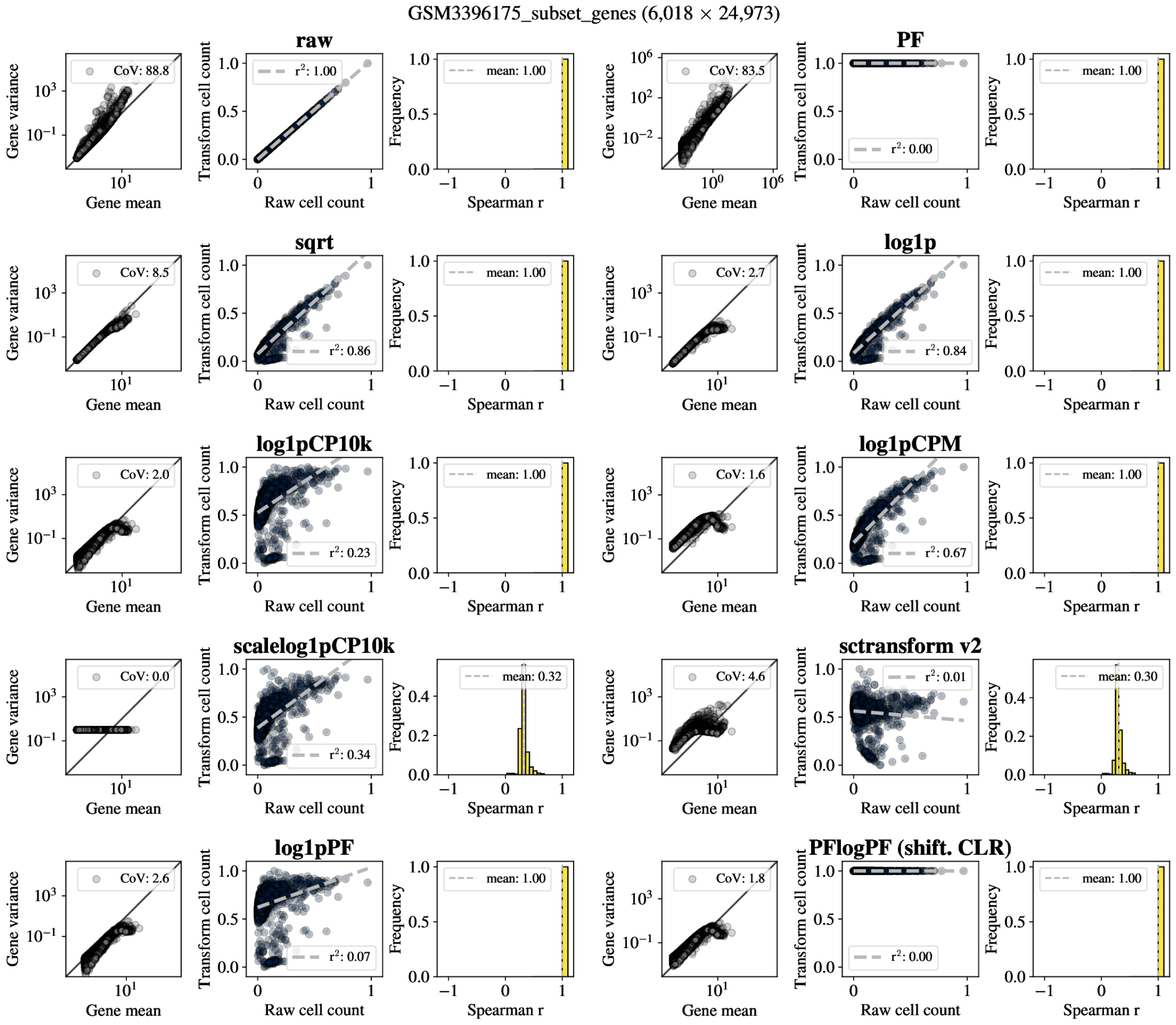


Supplementary Figure 1.235. **GSM3396175**. Per-method comparison of variance stabilization (left), depth normalization (middle), and monotonicity (right) for each normalization on dataset GSM3396175.

| GSM3396176 | |
|---|---|
| cells | 6,053 |
| genes | 60,623 |
| nonzero entries | 6,777,013 |
| density | 0.0185 |
| total counts | 34,225,718 |
| mean counts / cell | 5,654 |
| mean counts / gene | 565 |
| min counts / cell | 1,440 |
| max counts / cell | 53,890 |
| overdispersion | 2.99 |

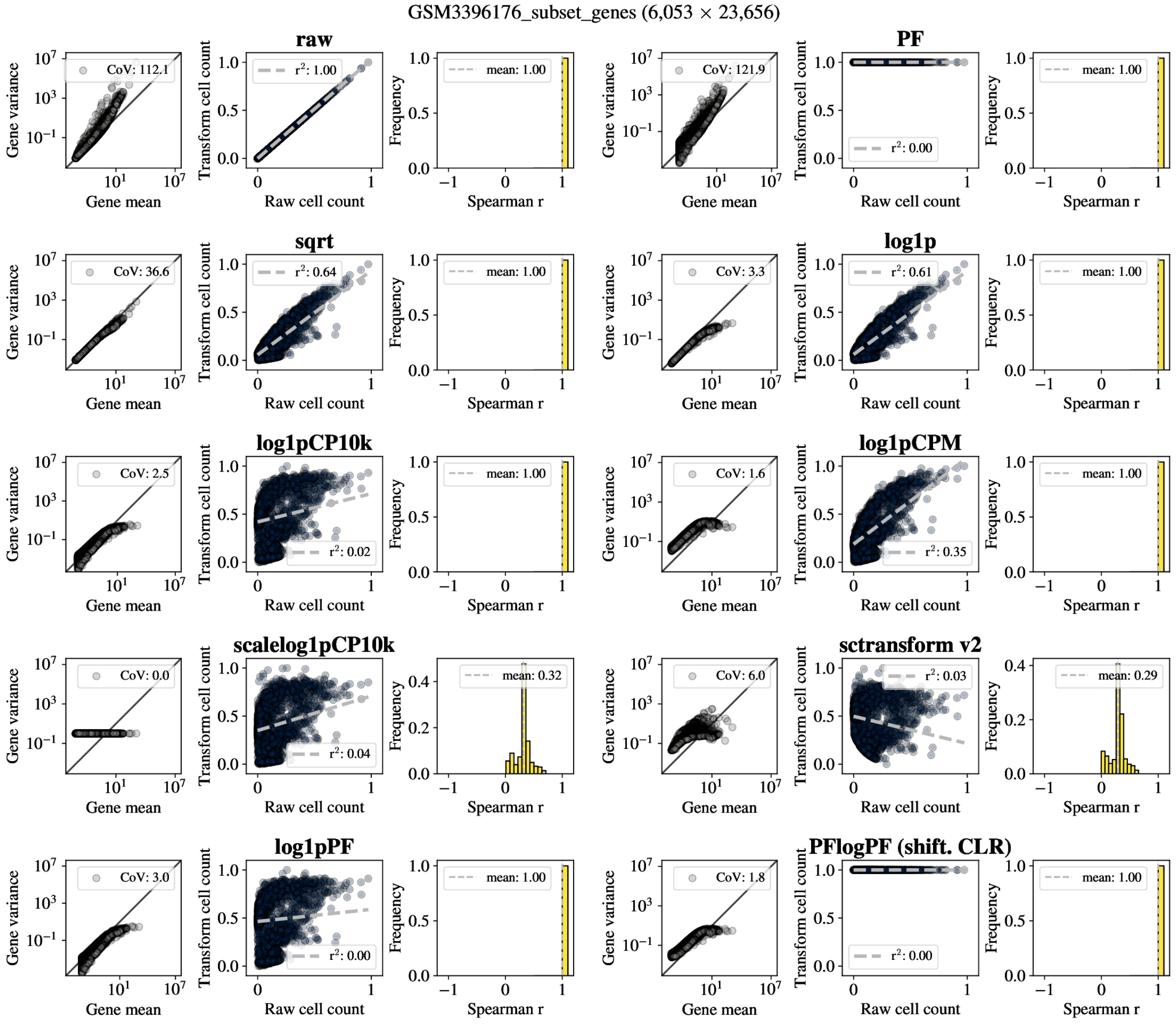


Supplementary Figure 1.236. **GSM3396176**. Per-method comparison of variance stabilization (left), depth normalization (middle), and monotonicity (right) for each normalization on dataset GSM3396176.

| GSM3396177 | |
|---|---|
| cells | 1,870 |
| genes | 60,623 |
| nonzero entries | 626,608 |
| density | 0.00553 |
| total counts | 3,721,878 |
| mean counts / cell | 1,990 |
| mean counts / gene | 61 |
| min counts / cell | 366 |
| max counts / cell | 14,047 |
| overdispersion | 1.8 |

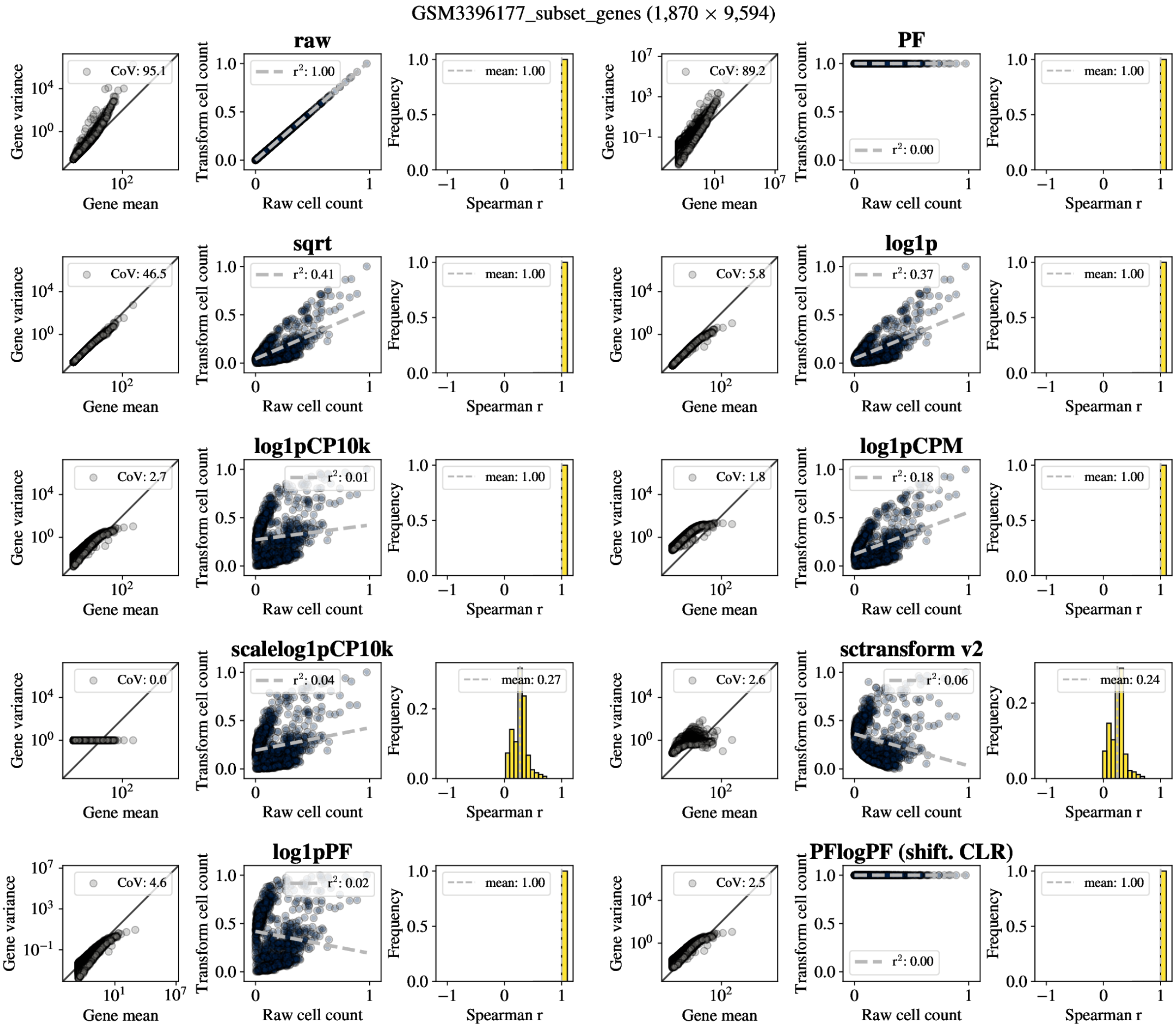


Supplementary Figure 1.237. **GSM3396177**. Per-method comparison of variance stabilization (left), depth normalization (middle), and monotonicity (right) for each normalization on dataset GSM3396177.

| GSM3396178 | |
|---|---|
| cells | 12,668 |
| genes | 60,623 |
| nonzero entries | 3,581,204 |
| density | 0.00466 |
| total counts | 14,037,831 |
| mean counts / cell | 1,108 |
| mean counts / gene | 232 |
| min counts / cell | 59 |
| max counts / cell | 32,567 |
| overdispersion | 9.14 |

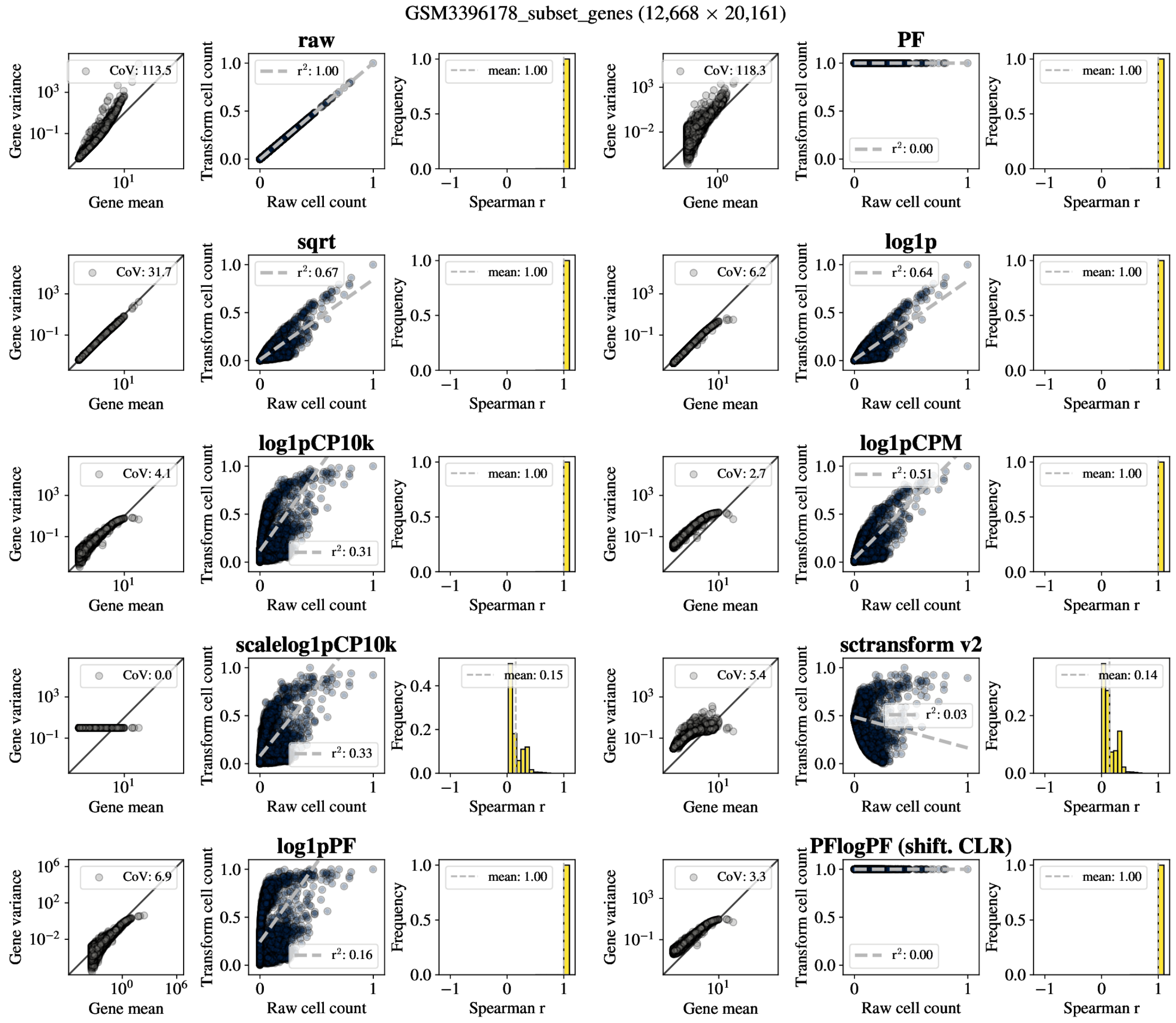


Supplementary Figure 1.238. **GSM3396178**. Per-method comparison of variance stabilization (left), depth normalization (middle), and monotonicity (right) for each normalization on dataset GSM3396178.

| GSM3396179 | |
|---|---|
| cells | 6,242 |
| genes | 60,623 |
| nonzero entries | 6,913,412 |
| density | 0.0183 |
| total counts | 21,807,503 |
| mean counts / cell | 3,494 |
| mean counts / gene | 360 |
| min counts / cell | 933 |
| max counts / cell | 41,050 |
| overdispersion | 6.7 |

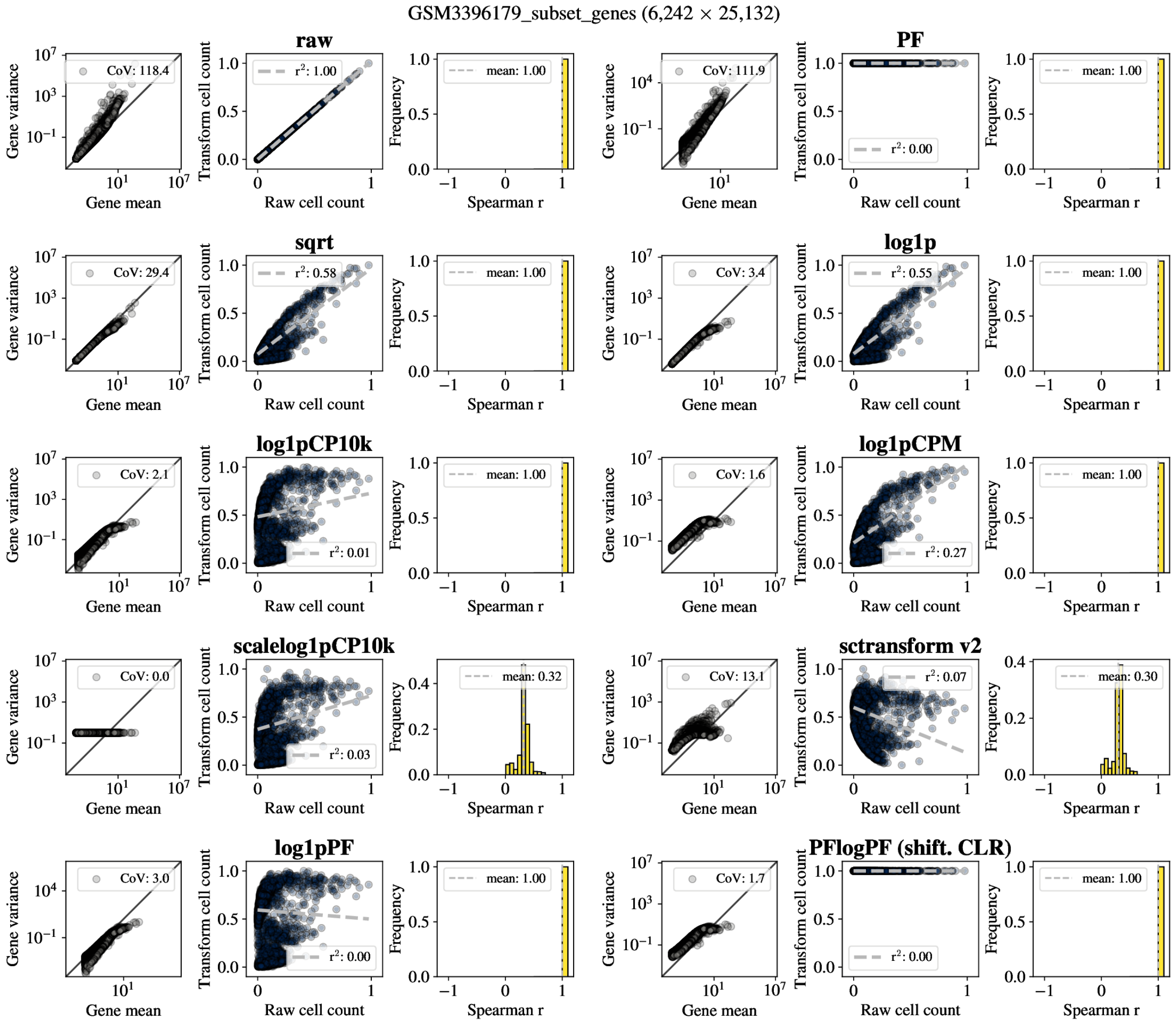


Supplementary Figure 1.239. **GSM3396179**. Per-method comparison of variance stabilization (left), depth normalization (middle), and monotonicity (right) for each normalization on dataset GSM3396179.

| **GSM3396180** | |
|---|---|
| cells | 1,126 |
| genes | 60,623 |
| nonzero entries | 853,954 |
| density | 0.0125 |
| total counts | 2,532,856 |
| mean counts / cell | 2,249 |
| mean counts / gene | 42 |
| min counts / cell | 522 |
| max counts / cell | 17,837 |
| overdispersion | 0.796 |

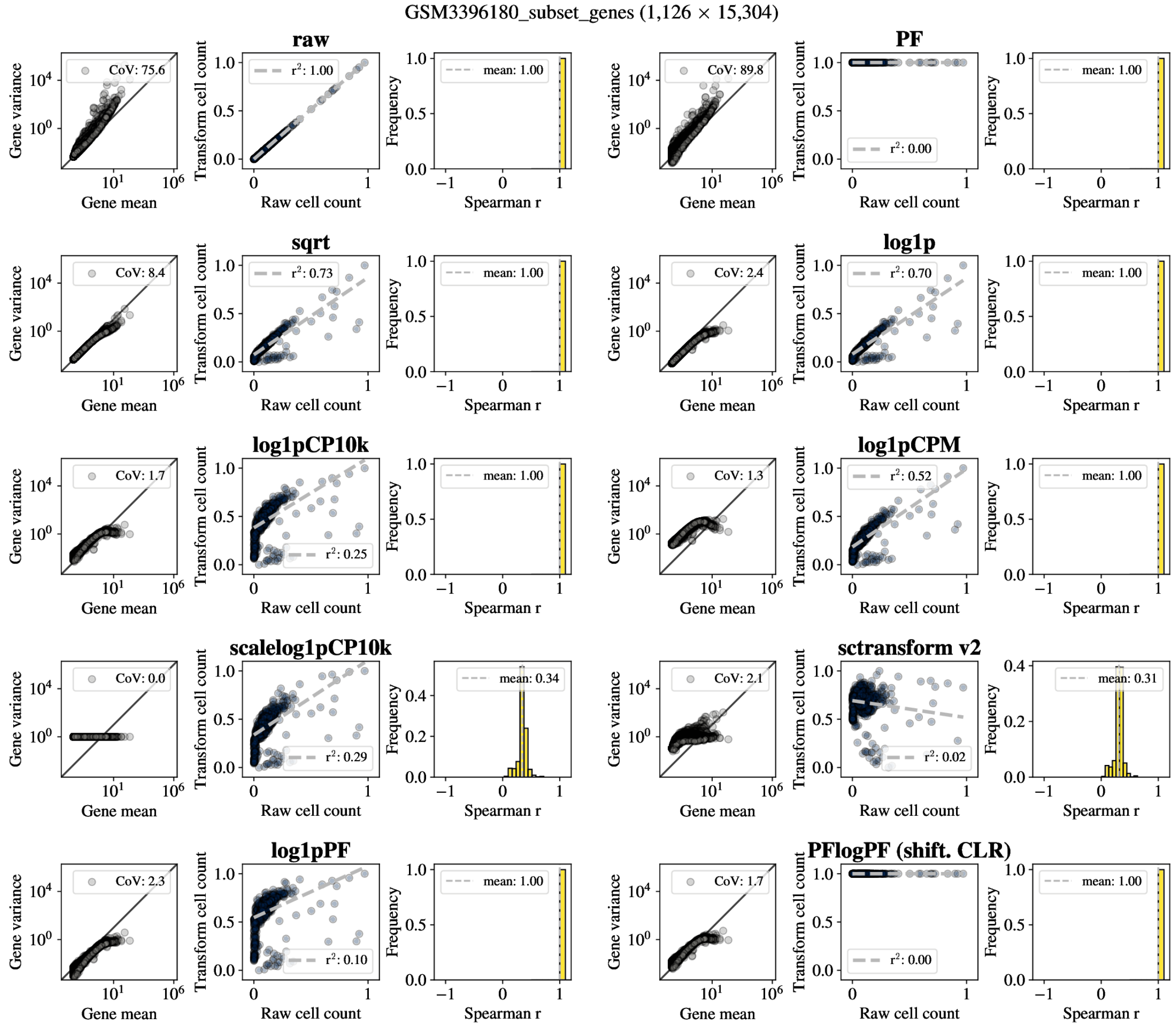


Supplementary Figure 1.240. **GSM3396180**. Per-method comparison of variance stabilization (left), depth normalization (middle), and monotonicity (right) for each normalization on dataset GSM3396180.

| GSM3396181 | |
|---|---|
| cells | 6,103 |
| genes | 60,623 |
| nonzero entries | 7,022,291 |
| density | 0.019 |
| total counts | 21,376,653 |
| mean counts / cell | 3,503 |
| mean counts / gene | 353 |
| min counts / cell | 918 |
| max counts / cell | 44,876 |
| overdispersion | 7.49 |

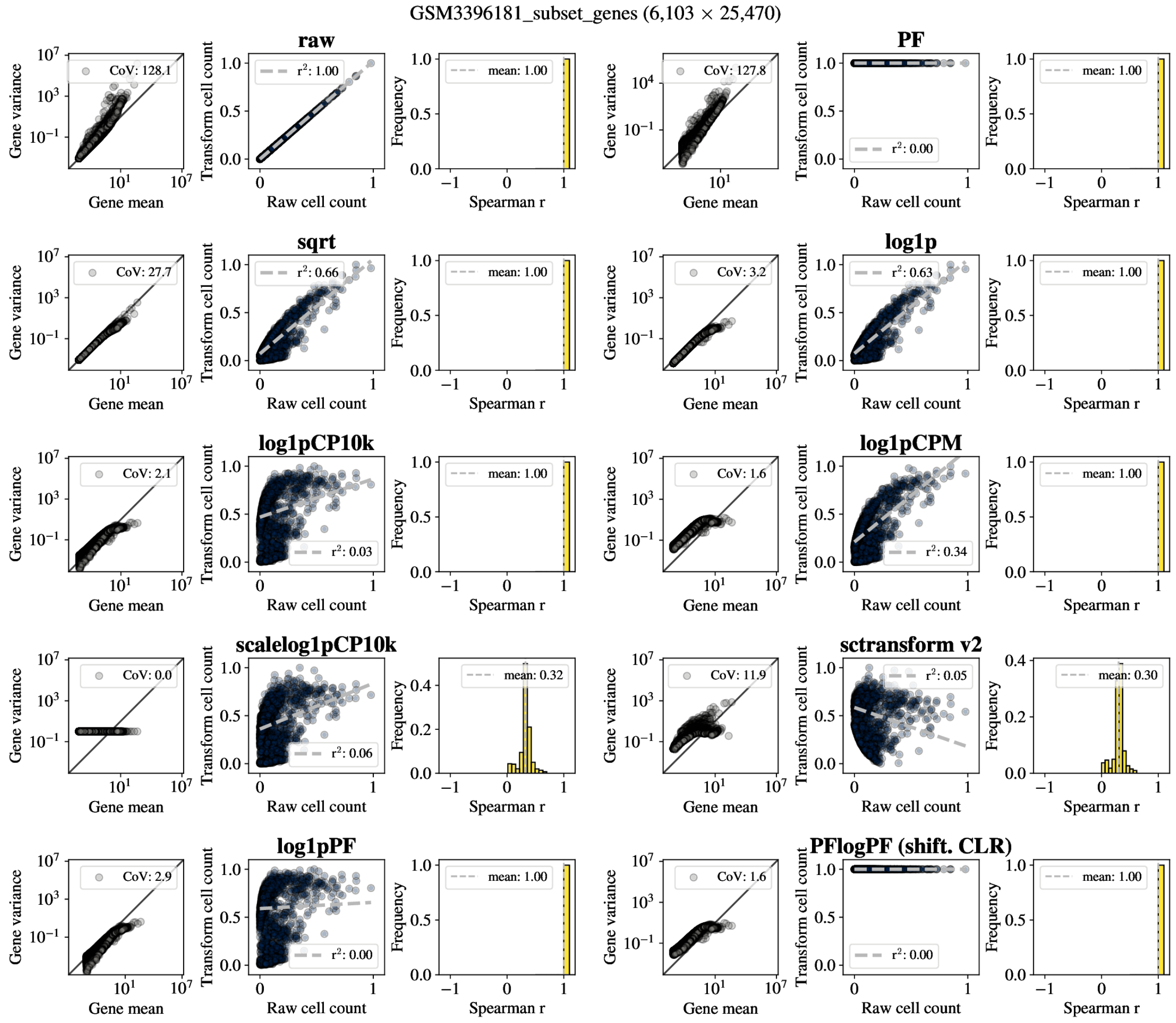


Supplementary Figure 1.241. **GSM3396181**. Per-method comparison of variance stabilization (left), depth normalization (middle), and monotonicity (right) for each normalization on dataset GSM3396181.

| GSM3396182 | |
|---|---|
| cells | 200 |
| genes | 60,623 |
| nonzero entries | 377,227 |
| density | 0.0311 |
| total counts | 1,517,361 |
| mean counts / cell | 7,587 |
| mean counts / gene | 25 |
| min counts / cell | 4,237 |
| max counts / cell | 27,446 |
| overdispersion | 2.42 |

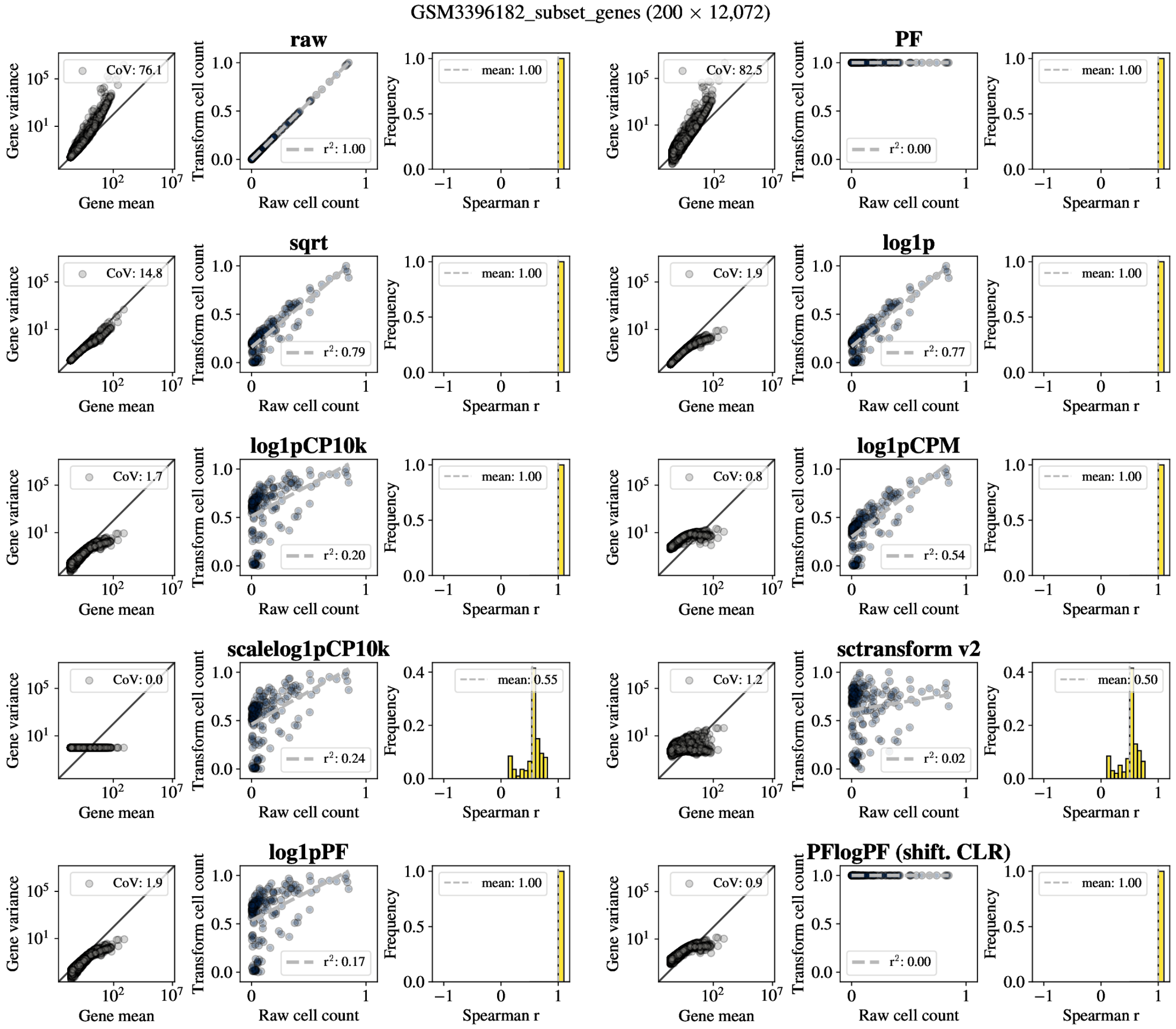


Supplementary Figure 1.242. **GSM3396182**. Per-method comparison of variance stabilization (left), depth normalization (middle), and monotonicity (right) for each normalization on dataset GSM3396182.

| GSM3396183 | |
|---|---|
| cells | 6,179 |
| genes | 60,623 |
| nonzero entries | 7,647,909 |
| density | 0.0204 |
| total counts | 28,098,441 |
| mean counts / cell | 4,547 |
| mean counts / gene | 463 |
| min counts / cell | 1,143 |
| max counts / cell | 47,653 |
| overdispersion | 6.64 |

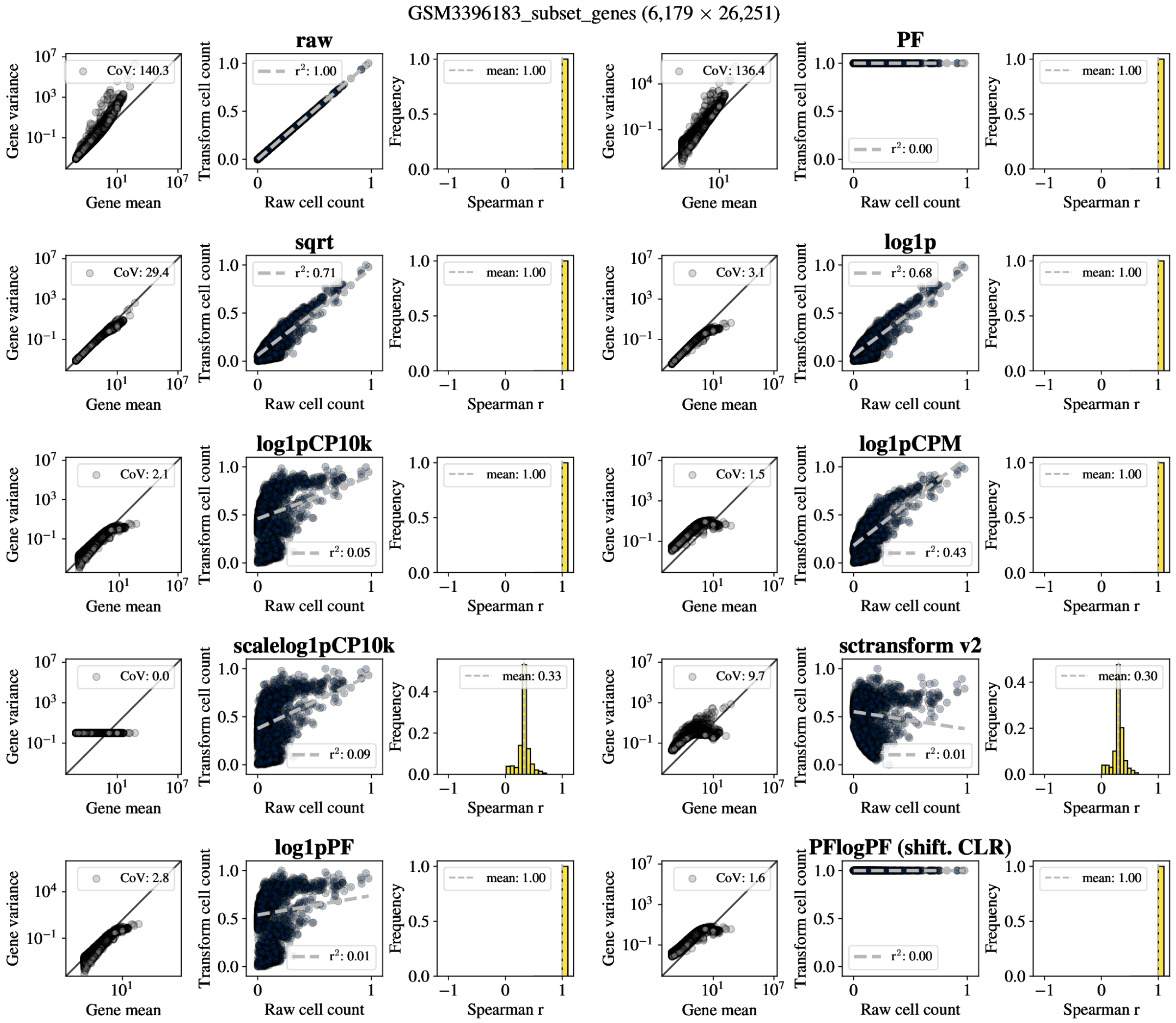


Supplementary Figure 1.243. **GSM3396183**. Per-method comparison of variance stabilization (left), depth normalization (middle), and monotonicity (right) for each normalization on dataset GSM3396183.

| GSM3396184 | |
|---|---|
| cells | 7,926 |
| genes | 60,623 |
| nonzero entries | 1,616,962 |
| density | 0.00337 |
| total counts | 8,119,713 |
| mean counts / cell | 1,024 |
| mean counts / gene | 134 |
| min counts / cell | 54 |
| max counts / cell | 17,646 |
| overdispersion | 11.2 |

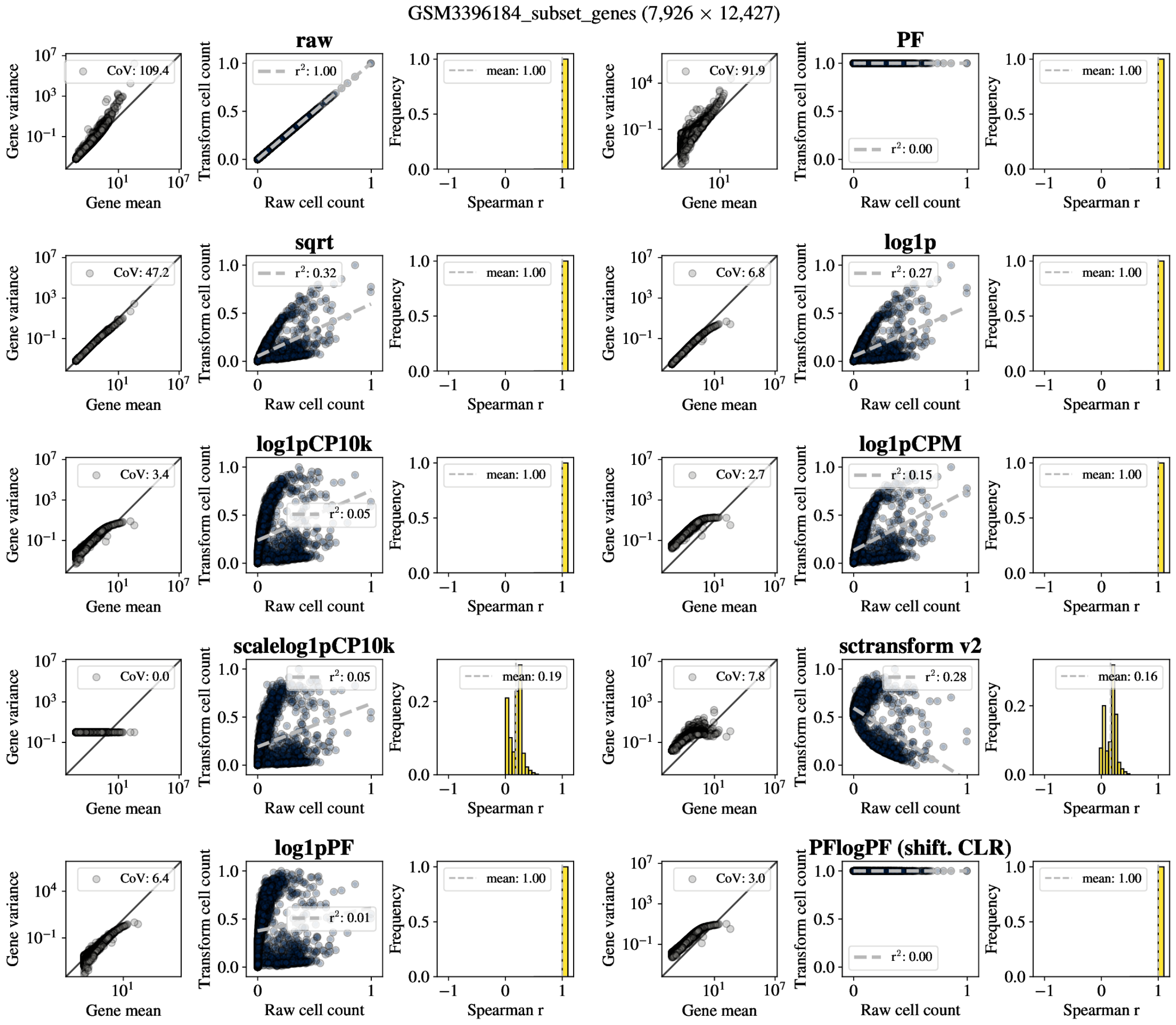


Supplementary Figure 1.244. **GSM3396184**. Per-method comparison of variance stabilization (left), depth normalization (middle), and monotonicity (right) for each normalization on dataset GSM3396184.

| GSM3396185 | |
|---|---|
| cells | 4,260 |
| genes | 60,623 |
| nonzero entries | 3,469,440 |
| density | 0.0134 |
| total counts | 10,903,099 |
| mean counts / cell | 2,559 |
| mean counts / gene | 180 |
| min counts / cell | 691 |
| max counts / cell | 36,979 |
| overdispersion | 7.36 |

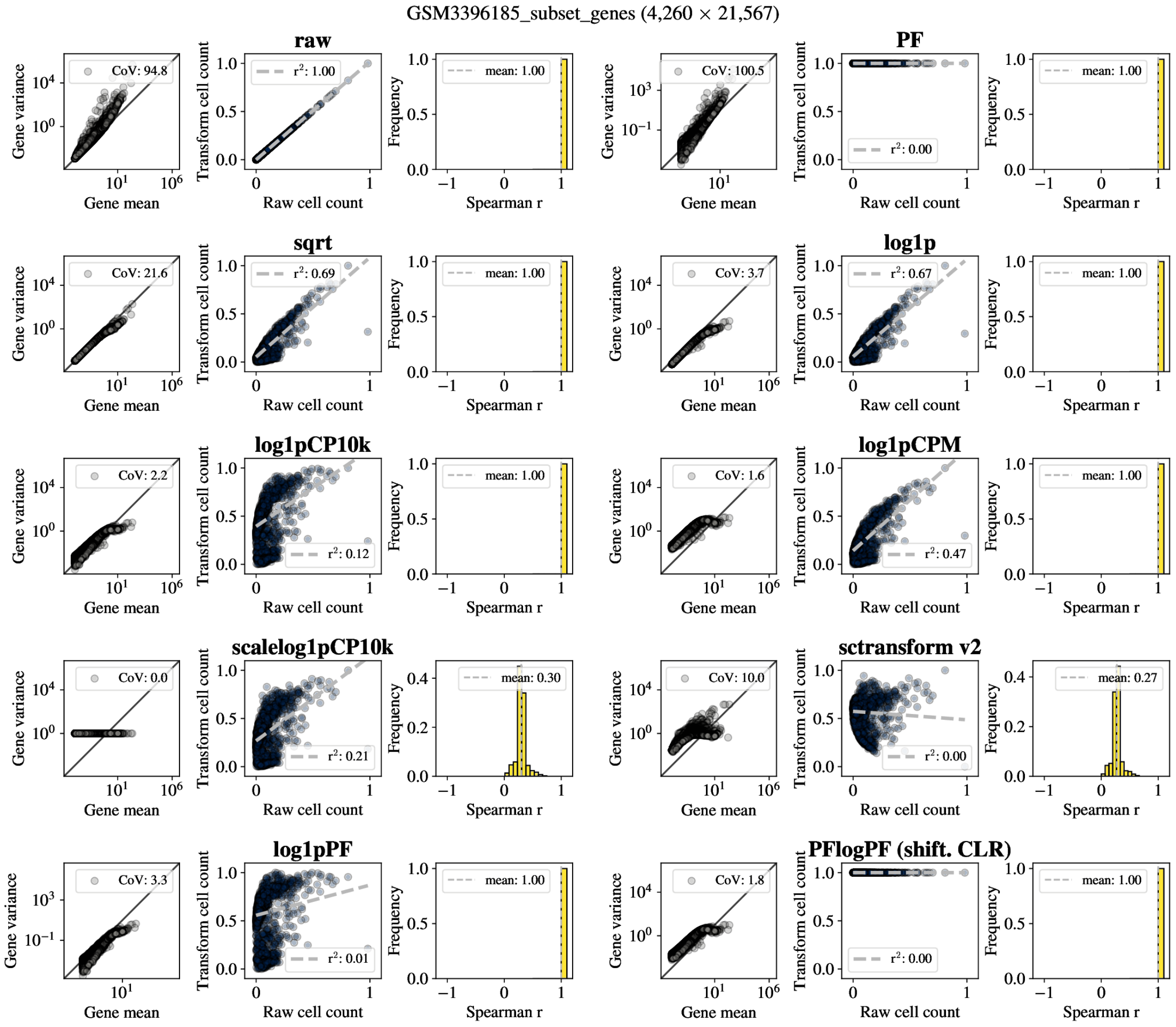


Supplementary Figure 1.245. **GSM3396185**. Per-method comparison of variance stabilization (left), depth normalization (middle), and monotonicity (right) for each normalization on dataset GSM3396185.

| GSM3402078 | |
|---|---|
| cells | 2,914 |
| genes | 60,623 |
| nonzero entries | 4,621,526 |
| density | 0.0262 |
| total counts | 13,586,560 |
| mean counts / cell | 4,663 |
| mean counts / gene | 224 |
| min counts / cell | 514 |
| max counts / cell | 60,558 |
| overdispersion | 1.22 |

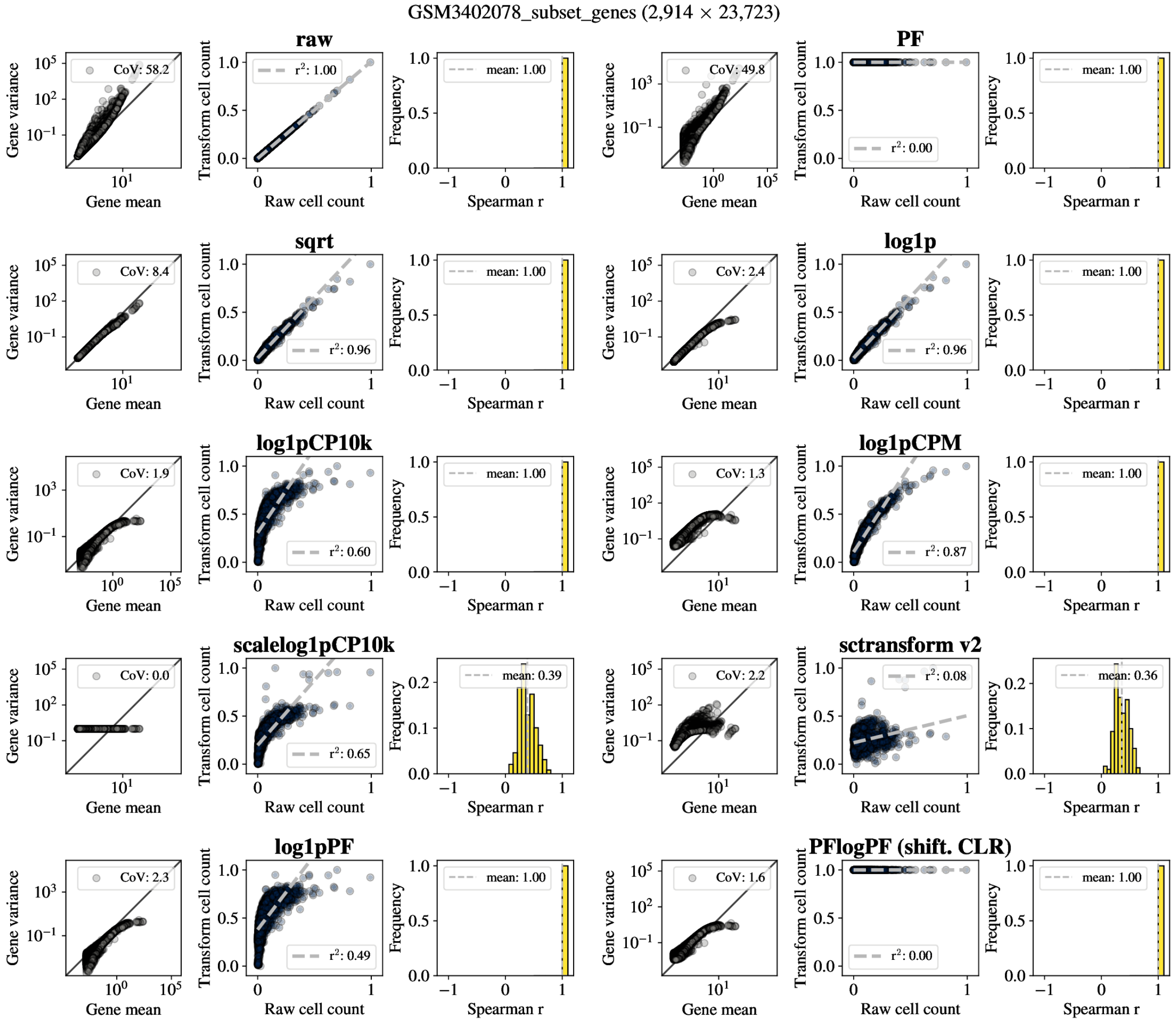


Supplementary Figure 1.246. **GSM3402078**. Per-method comparison of variance stabilization (left), depth normalization (middle), and monotonicity (right) for each normalization on dataset GSM3402078.

| GSM3402080 | |
|---|---|
| cells | 2,021 |
| genes | 60,623 |
| nonzero entries | 2,842,283 |
| density | 0.0232 |
| total counts | 8,501,463 |
| mean counts / cell | 4,207 |
| mean counts / gene | 140 |
| min counts / cell | 446 |
| max counts / cell | 43,048 |
| overdispersion | 2.14 |

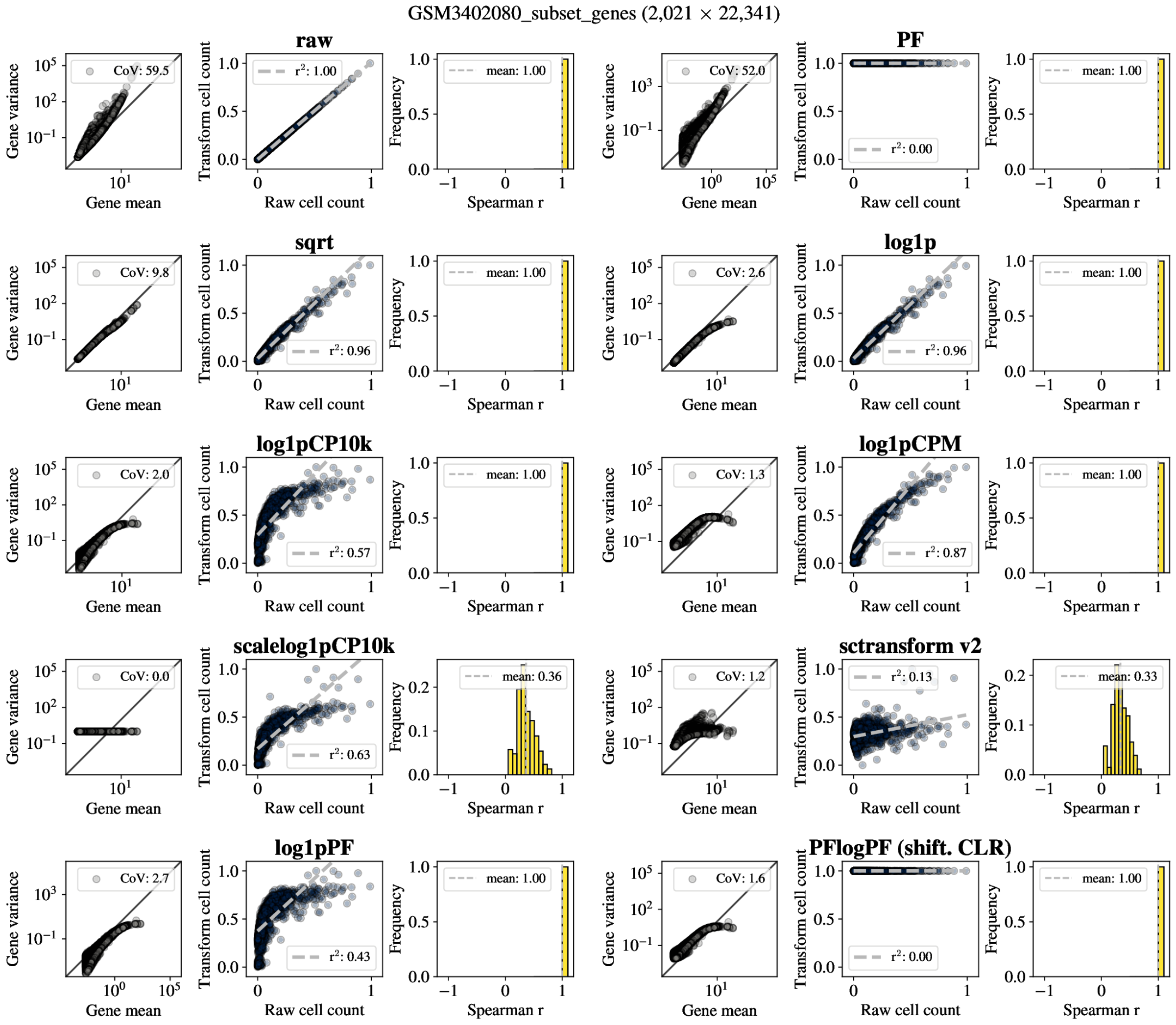


Supplementary Figure 1.247. **GSM3402080**. Per-method comparison of variance stabilization (left), depth normalization (middle), and monotonicity (right) for each normalization on dataset GSM3402080.

| GSM3439913 | |
|---|---|
| cells | 1,994 |
| genes | 60,623 |
| nonzero entries | 6,208,680 |
| density | 0.0514 |
| total counts | 26,243,300 |
| mean counts / cell | 13,161 |
| mean counts / gene | 433 |
| min counts / cell | 130 |
| max counts / cell | 57,736 |
| overdispersion | 0.982 |

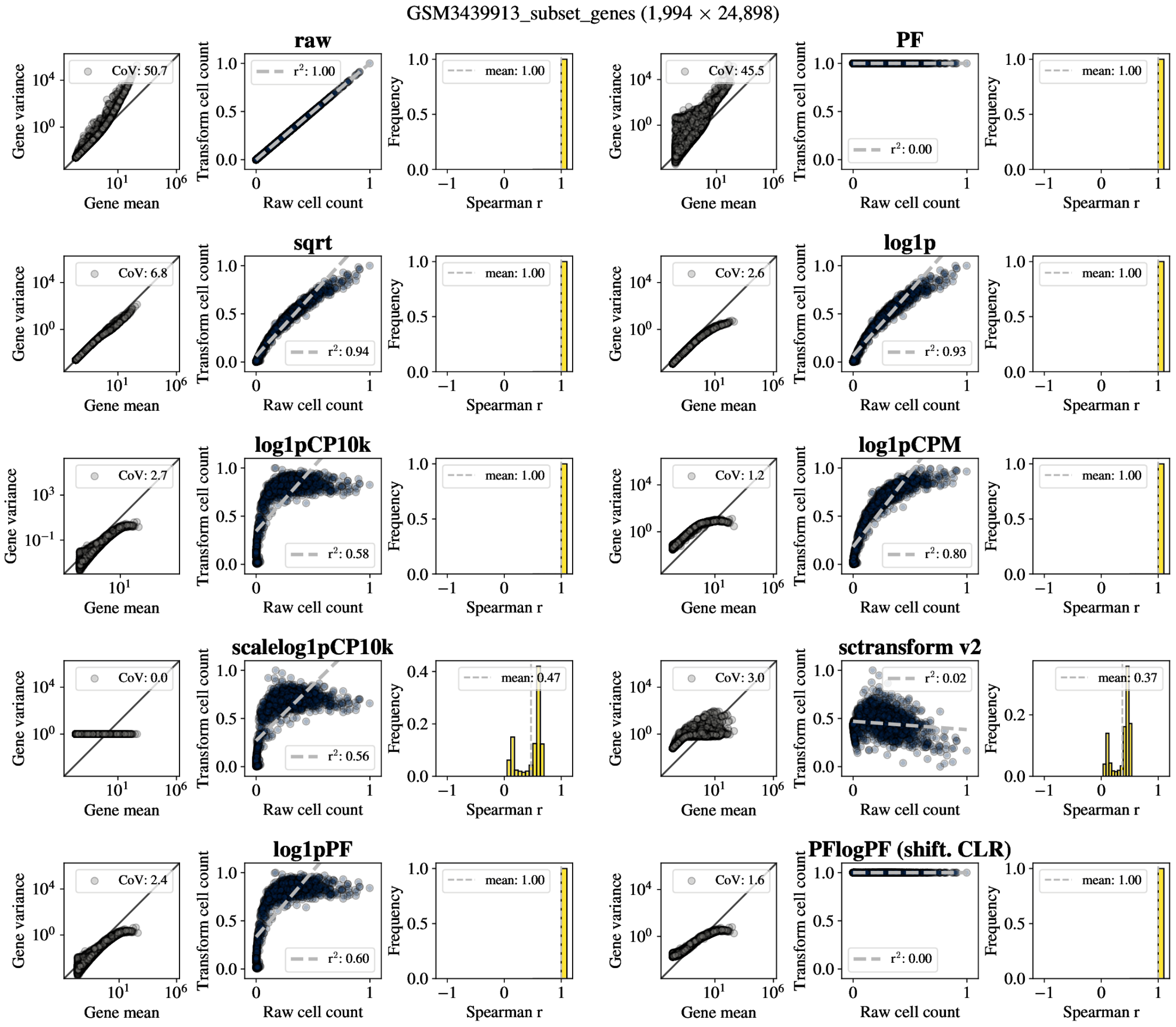


Supplementary Figure 1.248. **GSM3439913**. Per-method comparison of variance stabilization (left), depth normalization (middle), and monotonicity (right) for each normalization on dataset GSM3439913.

| GSM3439914 | |
|---|---|
| cells | 1,616 |
| genes | 60,623 |
| nonzero entries | 5,803,684 |
| density | 0.0592 |
| total counts | 21,130,668 |
| mean counts / cell | 13,076 |
| mean counts / gene | 349 |
| min counts / cell | 6,256 |
| max counts / cell | 43,409 |
| overdispersion | 0.452 |

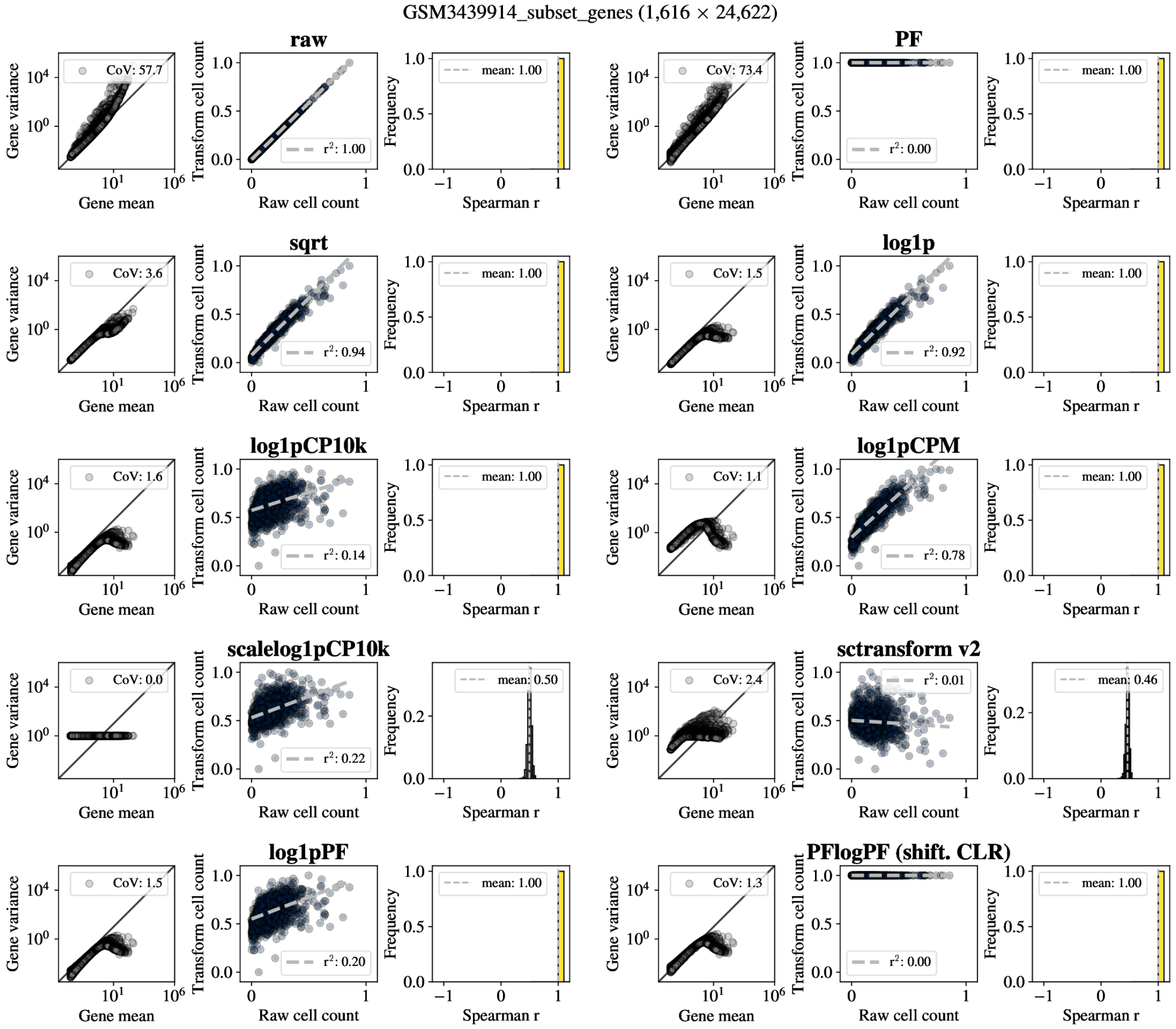


Supplementary Figure 1.249. **GSM3439914**. Per-method comparison of variance stabilization (left), depth normalization (middle), and monotonicity (right) for each normalization on dataset GSM3439914.

| GSM3439915 | |
|---|---|
| cells | 2,241 |
| genes | 60,623 |
| nonzero entries | 6,151,595 |
| density | 0.0453 |
| total counts | 36,306,920 |
| mean counts / cell | 16,201 |
| mean counts / gene | 599 |
| min counts / cell | 210 |
| max counts / cell | 80,366 |
| overdispersion | 1.14 |

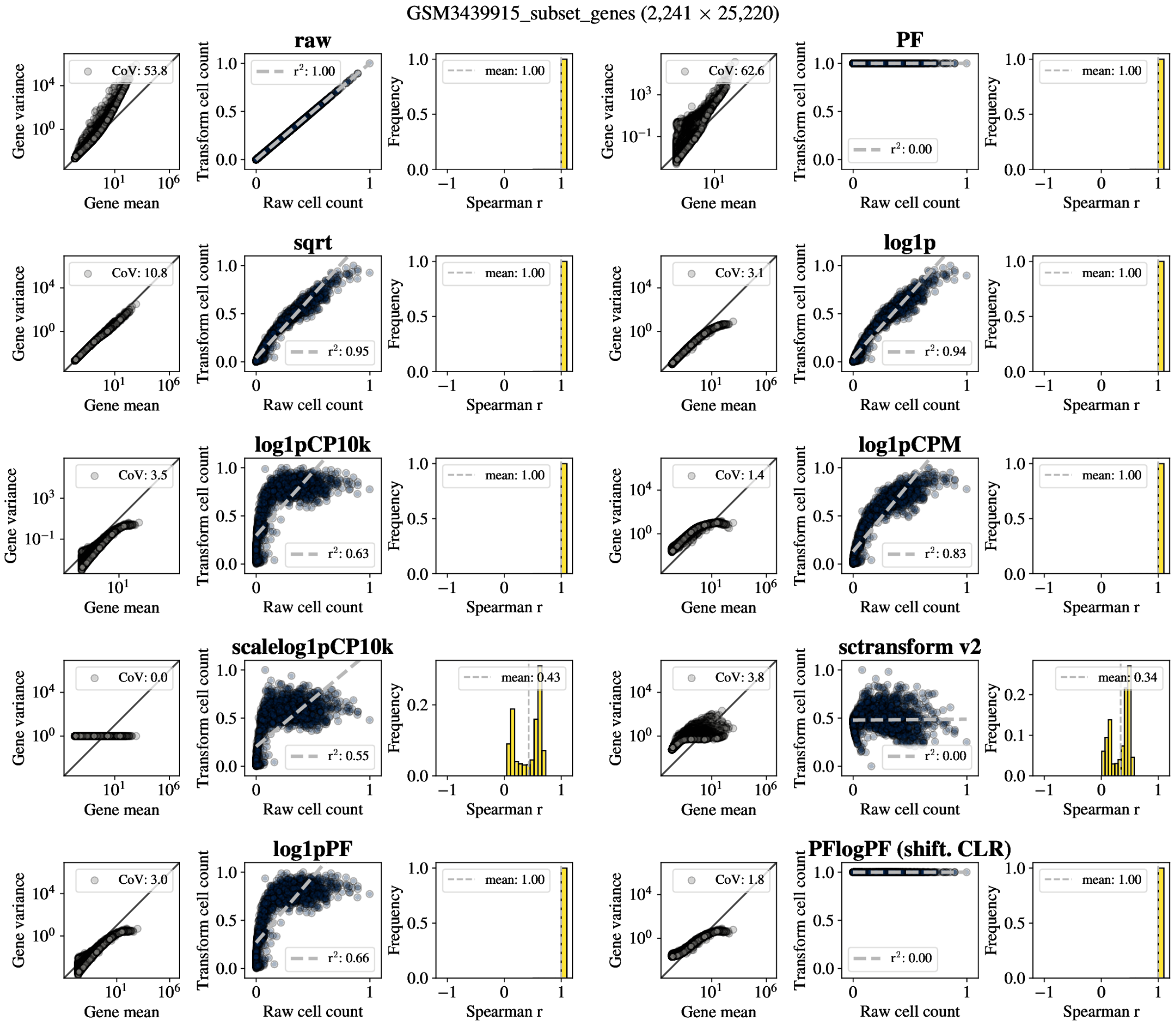


Supplementary Figure 1.250. **GSM3439915**. Per-method comparison of variance stabilization (left), depth normalization (middle), and monotonicity (right) for each normalization on dataset GSM3439915.

| GSM3439916 | |
|---|---|
| cells | 2,324 |
| genes | 60,623 |
| nonzero entries | 6,985,699 |
| density | 0.0496 |
| total counts | 38,650,994 |
| mean counts / cell | 16,631 |
| mean counts / gene | 638 |
| min counts / cell | 239 |
| max counts / cell | 80,241 |
| overdispersion | 1.43 |

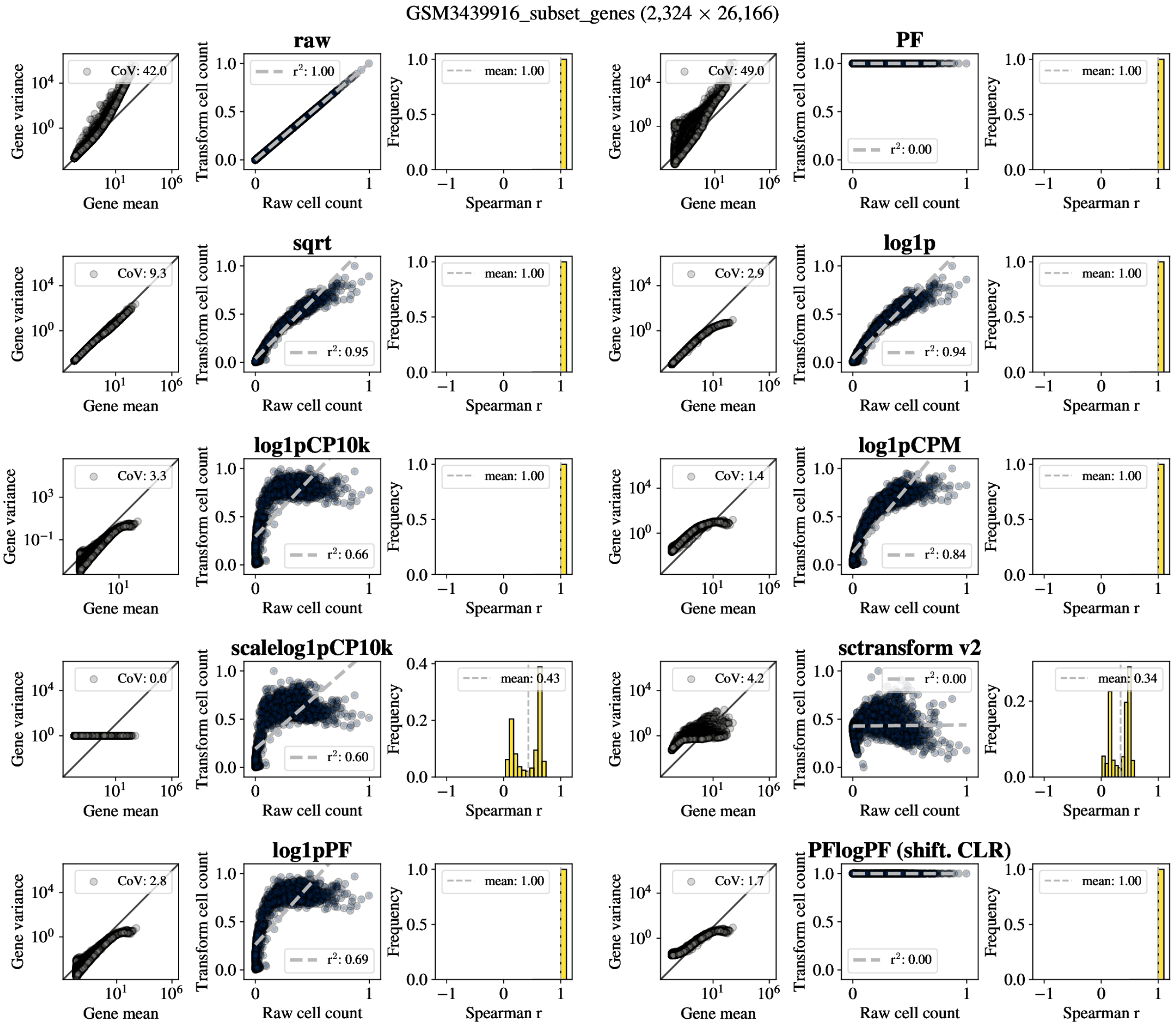


Supplementary Figure 1.251. **GSM3439916**. Per-method comparison of variance stabilization (left), depth normalization (middle), and monotonicity (right) for each normalization on dataset GSM3439916.

| GSM3439917 | |
|---|---|
| cells | 3,407 |
| genes | 60,623 |
| nonzero entries | 7,734,543 |
| density | 0.0374 |
| total counts | 43,500,723 |
| mean counts / cell | 12,768 |
| mean counts / gene | 718 |
| min counts / cell | 238 |
| max counts / cell | 109,555 |
| overdispersion | 4.54 |

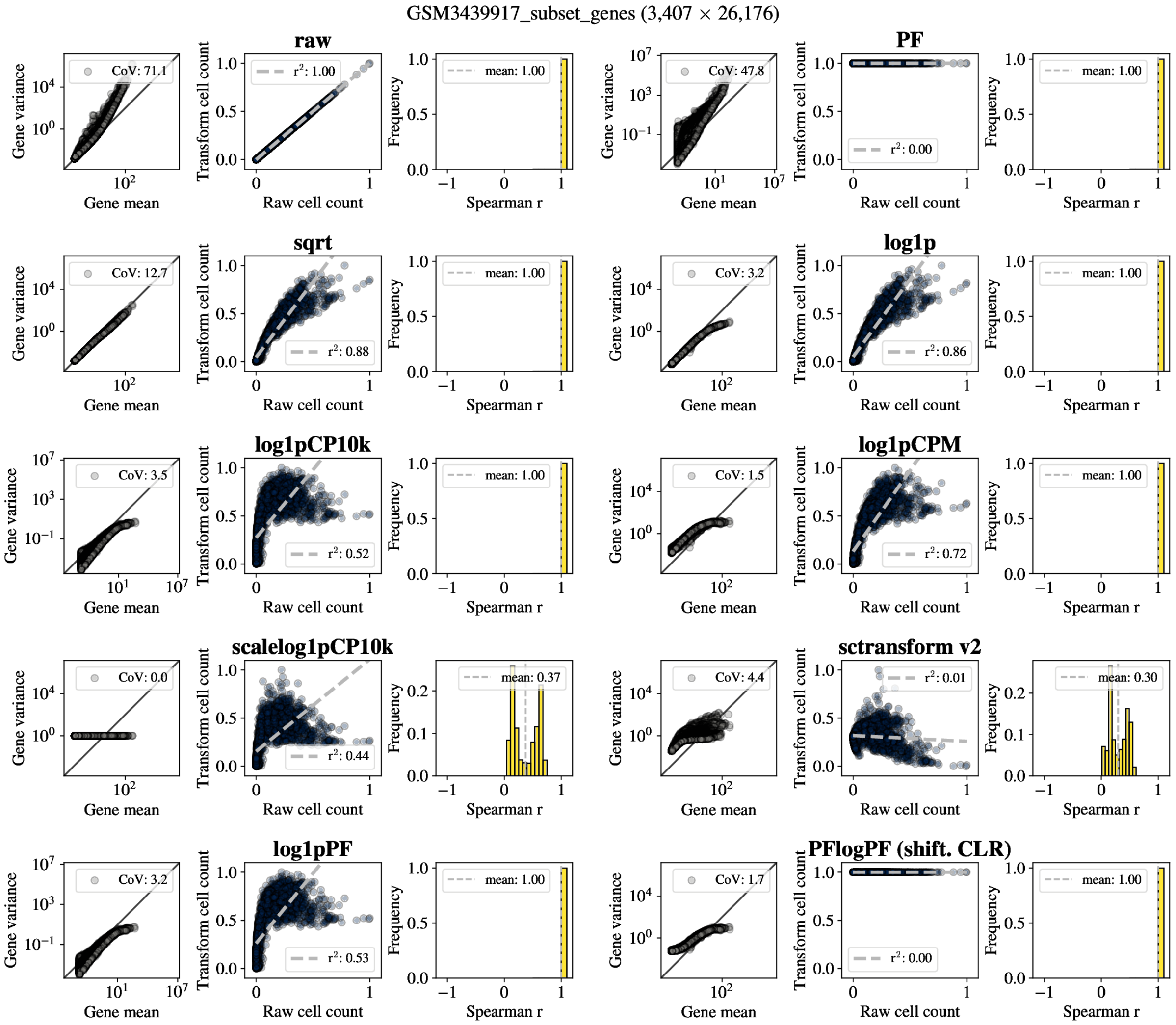


Supplementary Figure 1.252. **GSM3439917**. Per-method comparison of variance stabilization (left), depth normalization (middle), and monotonicity (right) for each normalization on dataset GSM3439917.

| GSM3439918 | |
|---|---|
| cells | 2,998 |
| genes | 60,623 |
| nonzero entries | 8,925,204 |
| density | 0.0491 |
| total counts | 43,929,711 |
| mean counts / cell | 14,653 |
| mean counts / gene | 725 |
| min counts / cell | 310 |
| max counts / cell | 89,030 |
| overdispersion | 1.12 |

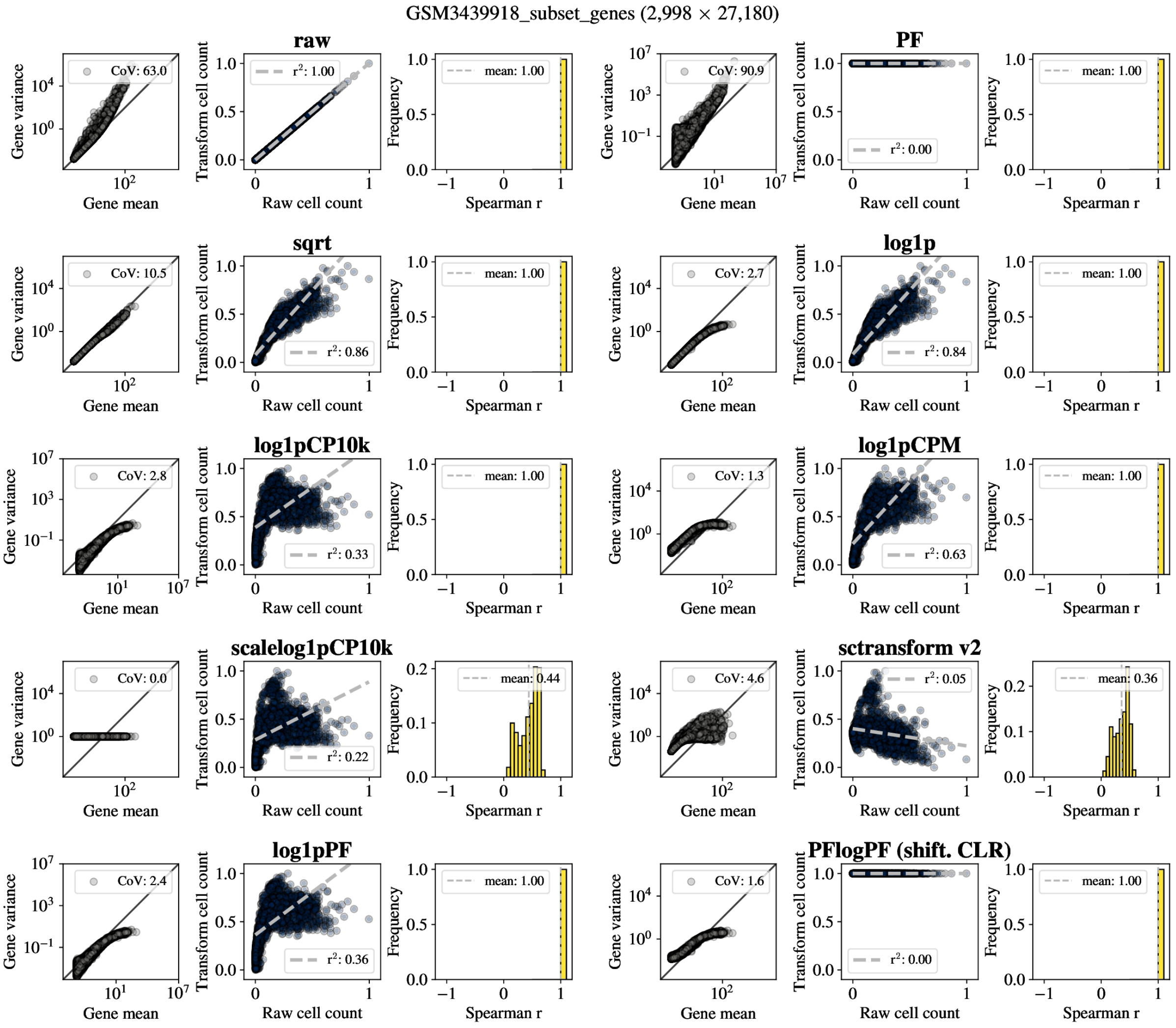


Supplementary Figure 1.253. **GSM3439918**. Per-method comparison of variance stabilization (left), depth normalization (middle), and monotonicity (right) for each normalization on dataset GSM3439918.

| GSM3439919 | |
|---|---|
| cells | 3,001 |
| genes | 60,623 |
| nonzero entries | 3,897,362 |
| density | 0.0214 |
| total counts | 17,126,960 |
| mean counts / cell | 5,707 |
| mean counts / gene | 283 |
| min counts / cell | 106 |
| max counts / cell | 127,170 |
| overdispersion | 4.56 |

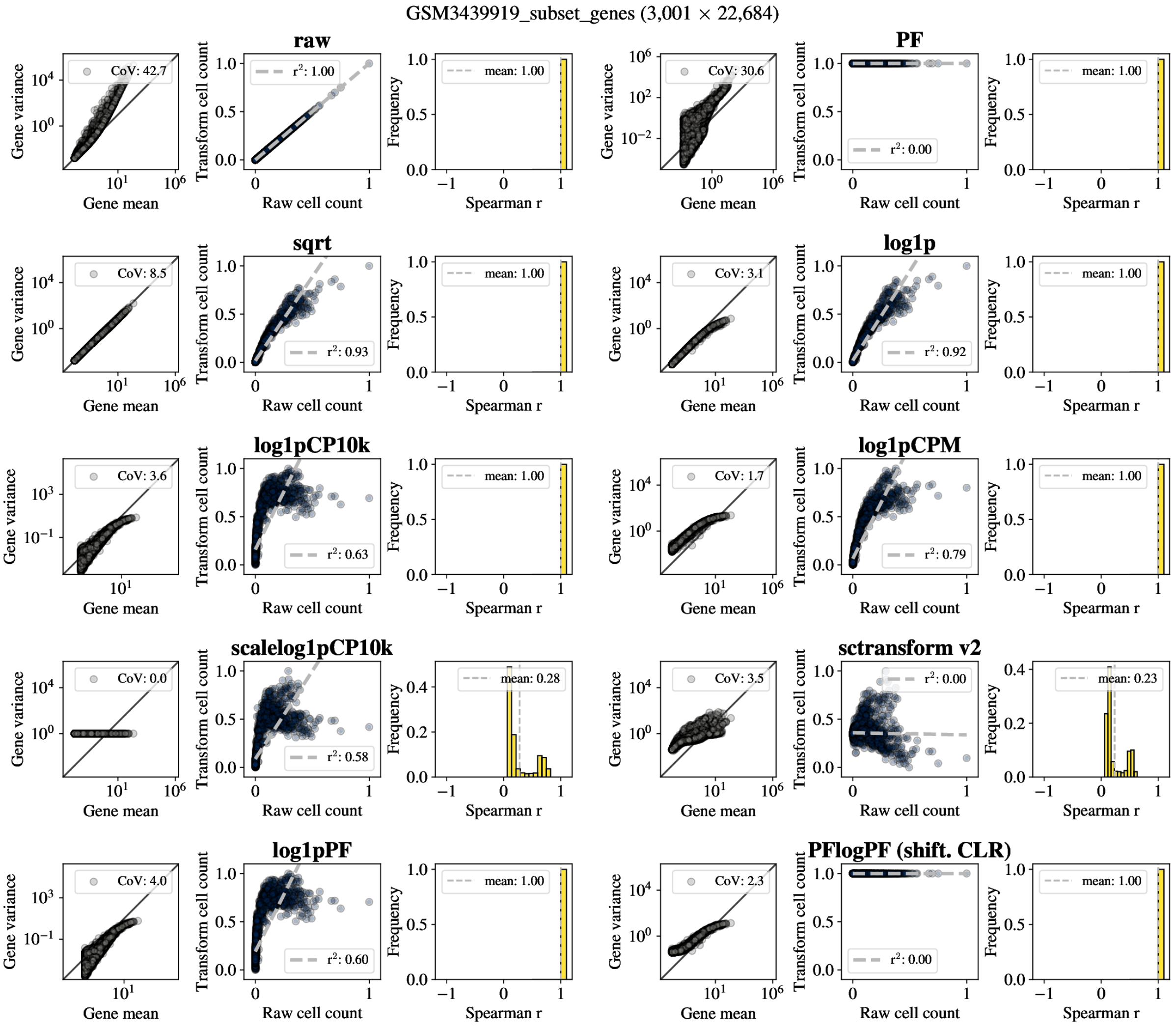


Supplementary Figure 1.254. **GSM3439919**. Per-method comparison of variance stabilization (left), depth normalization (middle), and monotonicity (right) for each normalization on dataset GSM3439919.

| GSM3439920 | |
|---|---|
| cells | 6,796 |
| genes | 60,623 |
| nonzero entries | 17,700,184 |
| density | 0.043 |
| total counts | 99,260,293 |
| mean counts / cell | 14,606 |
| mean counts / gene | 1,637 |
| min counts / cell | 399 |
| max counts / cell | 108,165 |
| overdispersion | 1.63 |

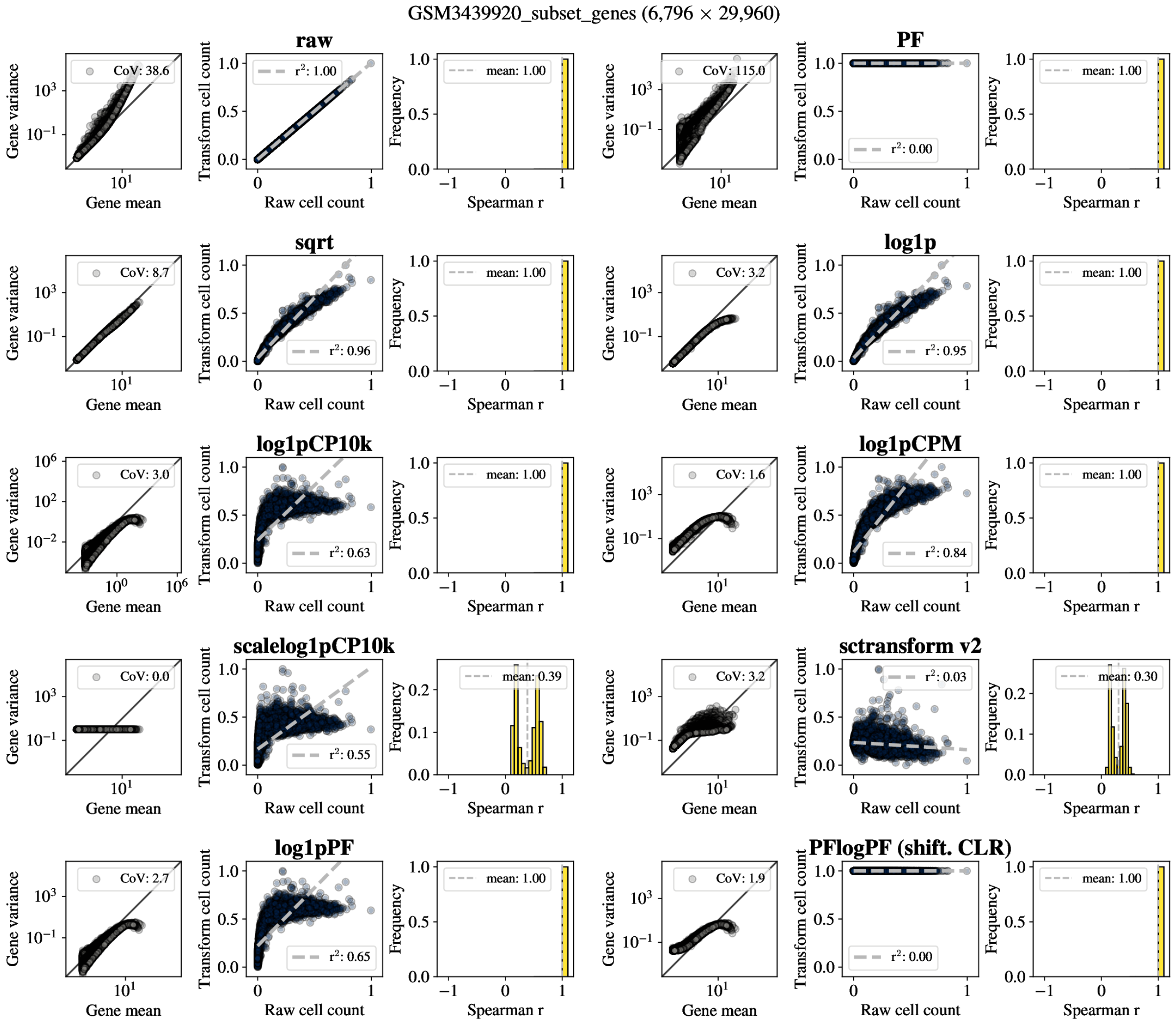


Supplementary Figure 1.255. **GSM3439920**. Per-method comparison of variance stabilization (left), depth normalization (middle), and monotonicity (right) for each normalization on dataset GSM3439920.

| GSM3439921 | |
|---|---|
| cells | 11,005 |
| genes | 60,623 |
| nonzero entries | 17,379,932 |
| density | 0.0261 |
| total counts | 73,858,828 |
| mean counts / cell | 6,711 |
| mean counts / gene | 1,218 |
| min counts / cell | 439 |
| max counts / cell | 92,157 |
| overdispersion | 2.88 |

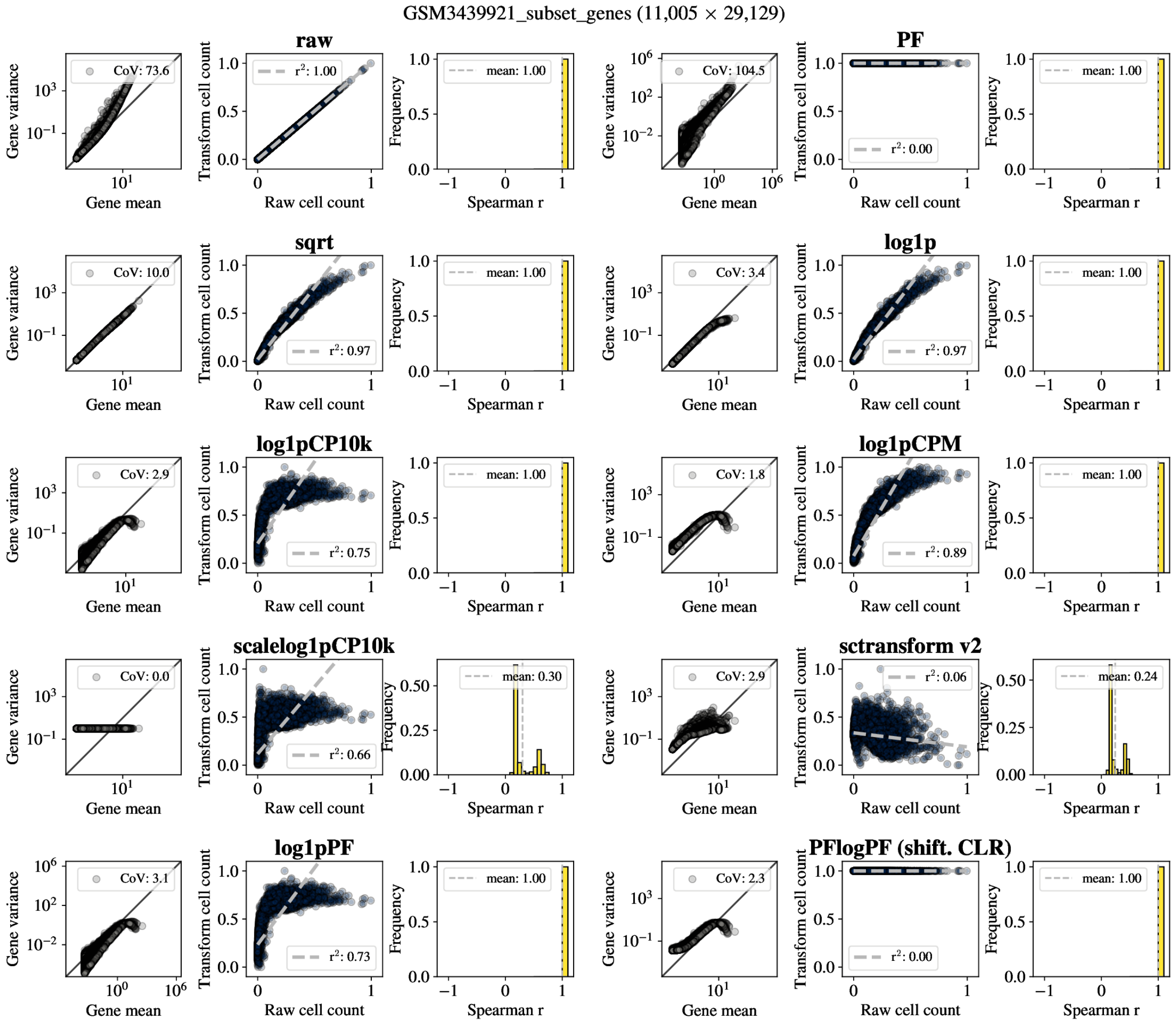


Supplementary Figure 1.256. **GSM3439921**. Per-method comparison of variance stabilization (left), depth normalization (middle), and monotonicity (right) for each normalization on dataset GSM3439921.

| GSM3439922 | |
|---|---|
| cells | 12,785 |
| genes | 60,623 |
| nonzero entries | 16,428,207 |
| density | 0.0212 |
| total counts | 66,119,644 |
| mean counts / cell | 5,172 |
| mean counts / gene | 1,091 |
| min counts / cell | 435 |
| max counts / cell | 55,526 |
| overdispersion | 2.93 |

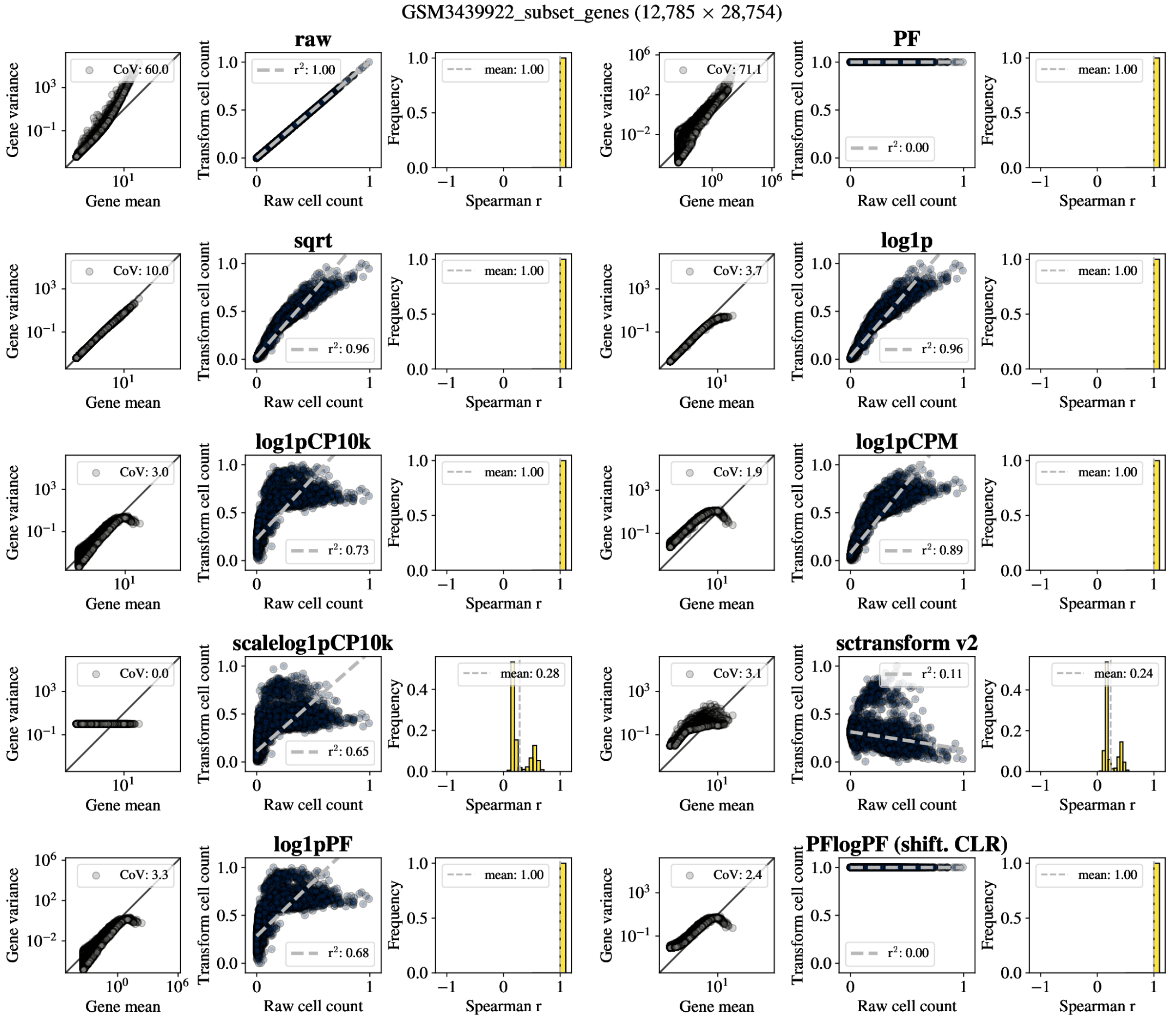


Supplementary Figure 1.257. **GSM3439922**. Per-method comparison of variance stabilization (left), depth normalization (middle), and monotonicity (right) for each normalization on dataset GSM3439922.

| GSM3439923 | |
|---|---|
| cells | 4,412 |
| genes | 60,623 |
| nonzero entries | 10,720,205 |
| density | 0.0401 |
| total counts | 38,961,578 |
| mean counts / cell | 8,831 |
| mean counts / gene | 643 |
| min counts / cell | 429 |
| max counts / cell | 62,524 |
| overdispersion | 1.99 |

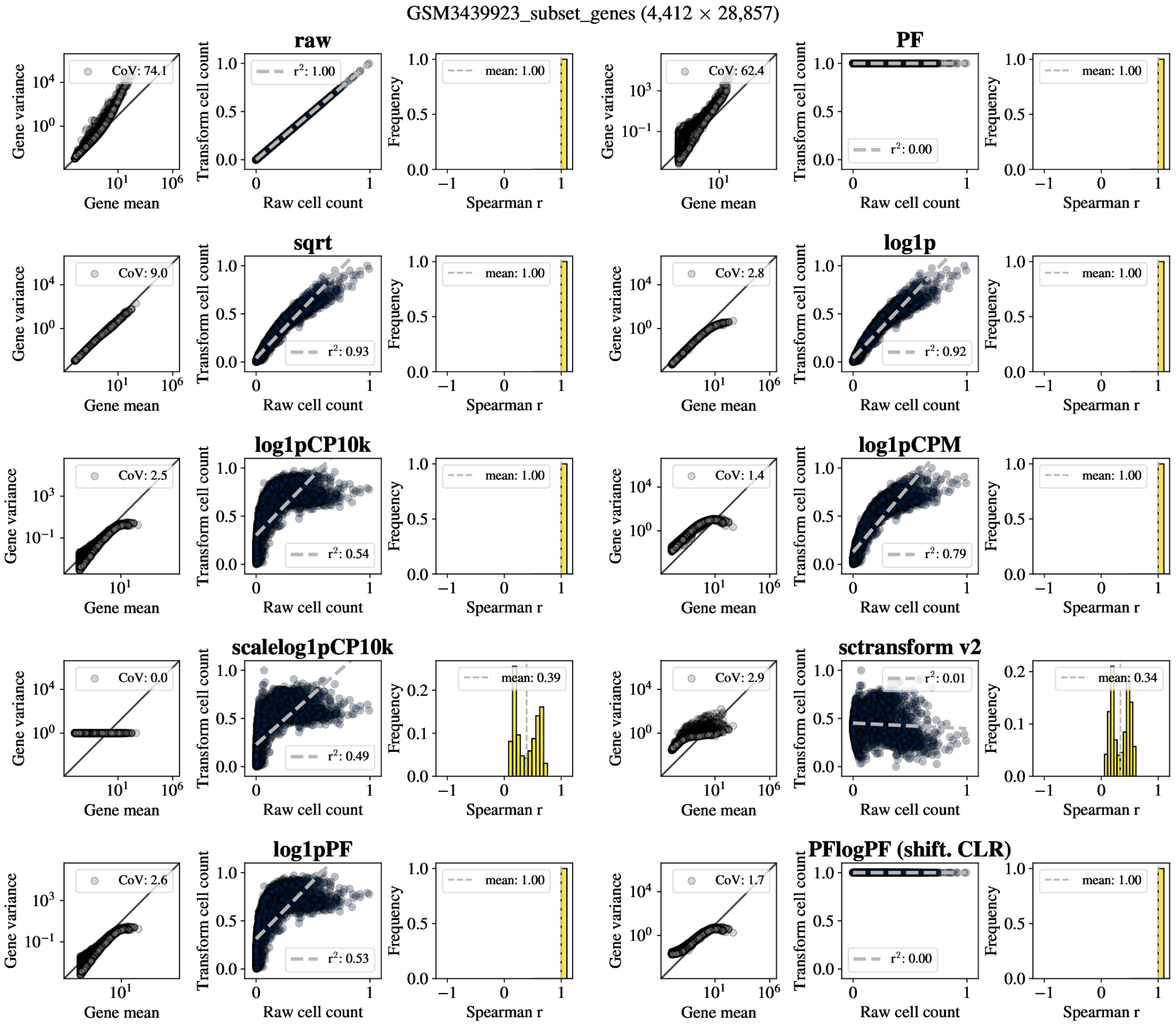


Supplementary Figure 1.258. **GSM3439923**. Per-method comparison of variance stabilization (left), depth normalization (middle), and monotonicity (right) for each normalization on dataset GSM3439923.

| GSM3439925 | |
|---|---|
| cells | 10,786 |
| genes | 60,623 |
| nonzero entries | 4,772,557 |
| density | 0.0073 |
| total counts | 10,691,172 |
| mean counts / cell | 991 |
| mean counts / gene | 176 |
| min counts / cell | 220 |
| max counts / cell | 11,310 |
| overdispersion | 1.42 |

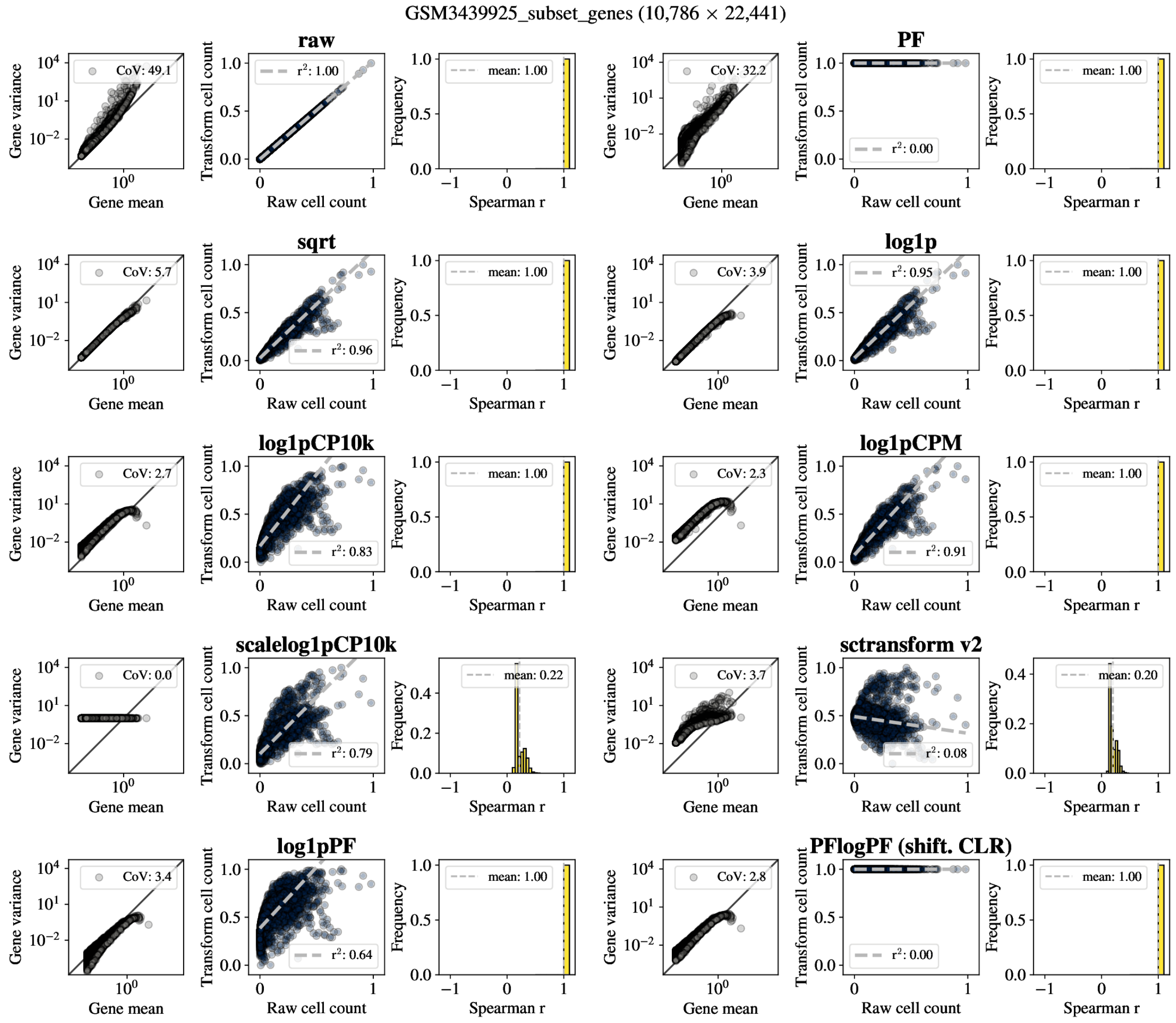


Supplementary Figure 1.259. **GSM3439925**. Per-method comparison of variance stabilization (left), depth normalization (middle), and monotonicity (right) for each normalization on dataset GSM3439925.

| GSM3439926 | |
|---|---|
| cells | 6,683 |
| genes | 60,623 |
| nonzero entries | 15,273,978 |
| density | 0.0377 |
| total counts | 48,937,222 |
| mean counts / cell | 7,323 |
| mean counts / gene | 807 |
| min counts / cell | 283 |
| max counts / cell | 44,836 |
| overdispersion | 1.34 |

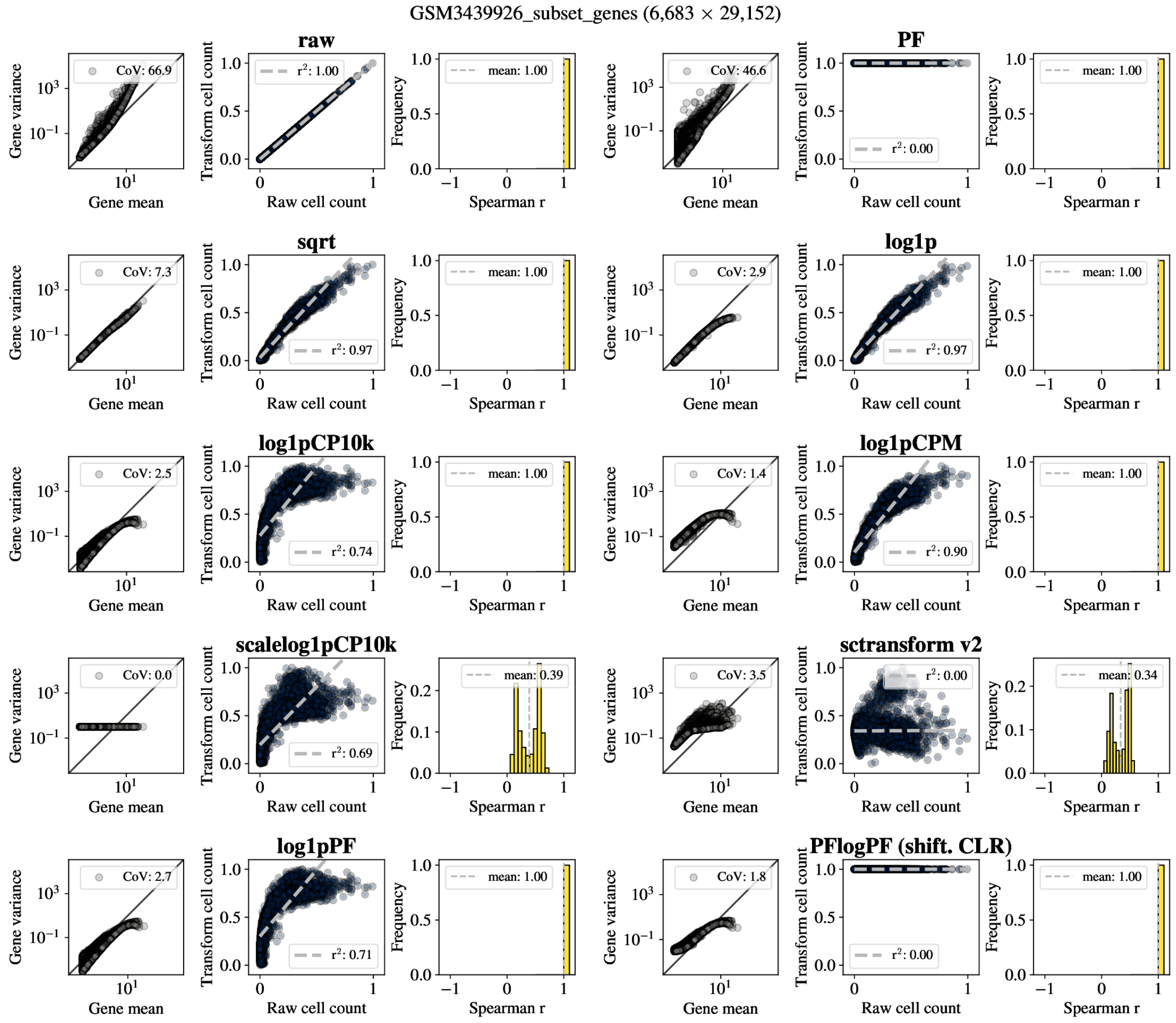


Supplementary Figure 1.260. **GSM3439926**. Per-method comparison of variance stabilization (left), depth normalization (middle), and monotonicity (right) for each normalization on dataset GSM3439926.

| GSM3439927 | |
|---|---|
| cells | 7,426 |
| genes | 60,623 |
| nonzero entries | 14,267,668 |
| density | 0.0317 |
| total counts | 44,031,418 |
| mean counts / cell | 5,929 |
| mean counts / gene | 726 |
| min counts / cell | 196 |
| max counts / cell | 55,863 |
| overdispersion | 9.07 |

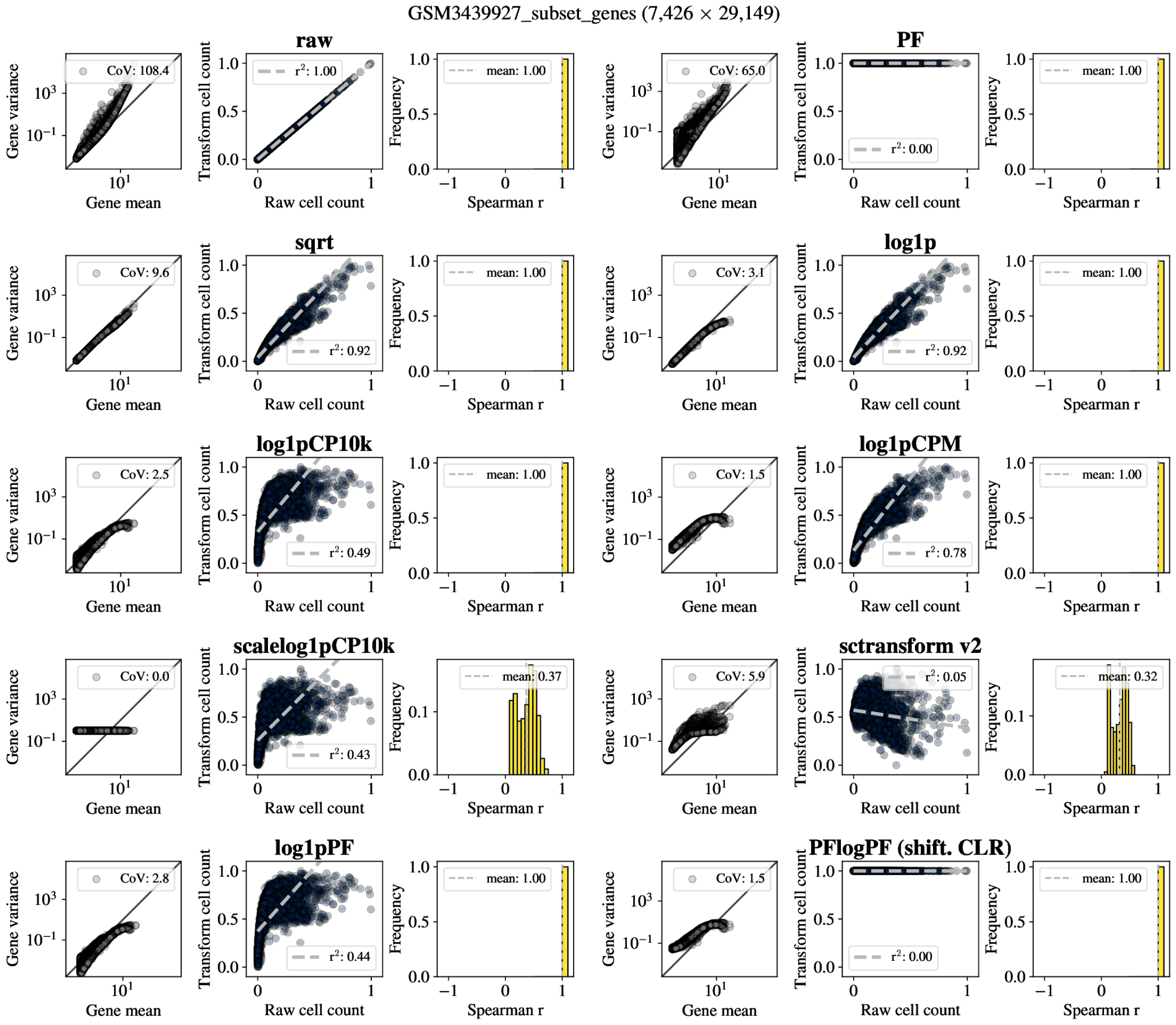


Supplementary Figure 1.261. **GSM3439927**. Per-method comparison of variance stabilization (left), depth normalization (middle), and monotonicity (right) for each normalization on dataset GSM3439927.

| GSM3439987 | |
|---|---|
| cells | 7,041 |
| genes | 60,623 |
| nonzero entries | 10,630,749 |
| density | 0.0249 |
| total counts | 39,751,114 |
| mean counts / cell | 5,646 |
| mean counts / gene | 656 |
| min counts / cell | 190 |
| max counts / cell | 51,351 |
| overdispersion | 2.32 |

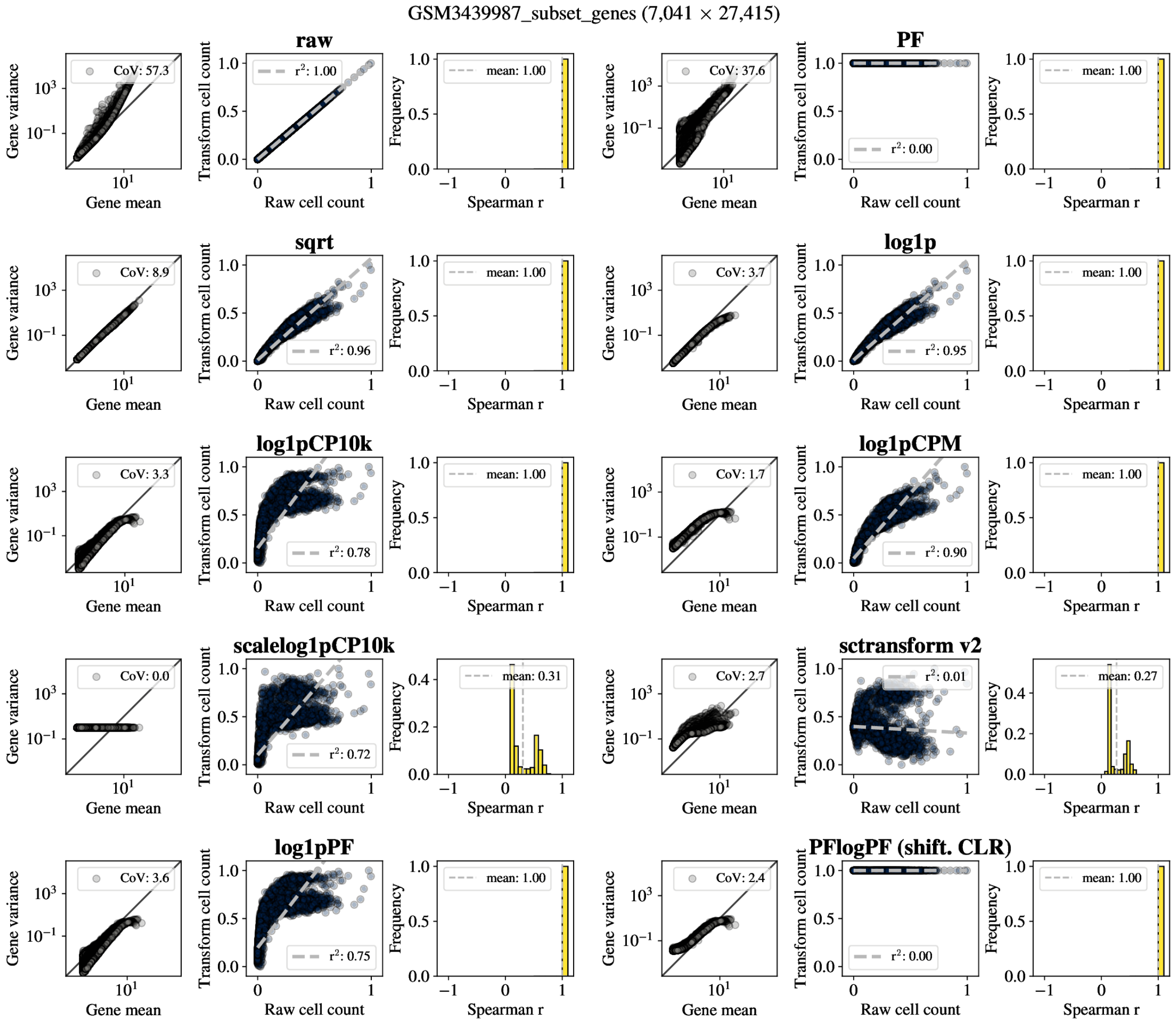


Supplementary Figure 1.262. **GSM3439987**. Per-method comparison of variance stabilization (left), depth normalization (middle), and monotonicity (right) for each normalization on dataset GSM3439987.

| **GSM3439988** | |
|---|---|
| cells | 2,518 |
| genes | 60,623 |
| nonzero entries | 5,785,054 |
| density | 0.0379 |
| total counts | 25,292,997 |
| mean counts / cell | 10,045 |
| mean counts / gene | 417 |
| min counts / cell | 197 |
| max counts / cell | 69,826 |
| overdispersion | 1.55 |

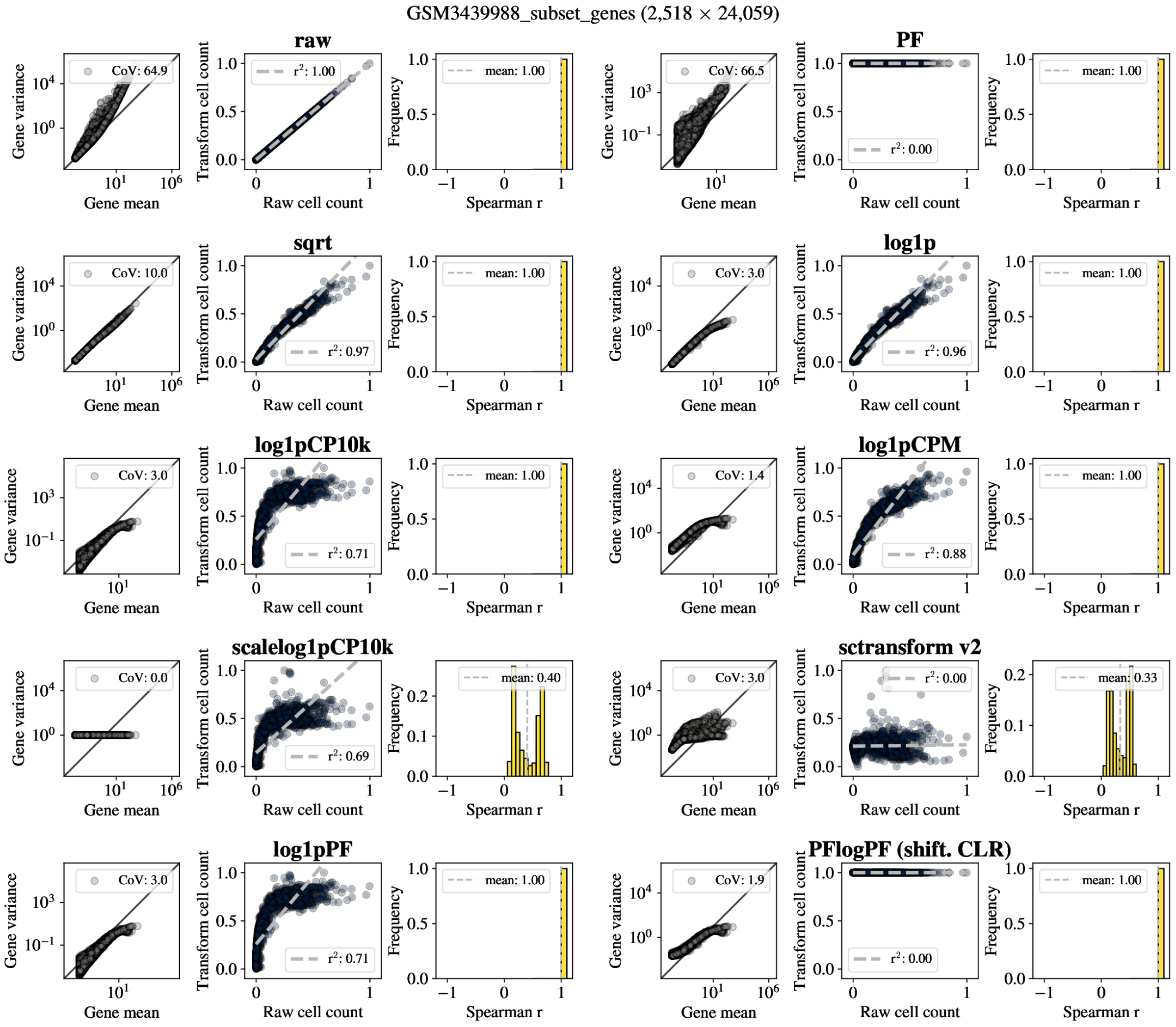


Supplementary Figure 1.263. **GSM3439988**. Per-method comparison of variance stabilization (left), depth normalization (middle), and monotonicity (right) for each normalization on dataset GSM3439988.

| GSM3439989 | |
|---|---|
| cells | 6,471 |
| genes | 60,623 |
| nonzero entries | 8,783,368 |
| density | 0.0224 |
| total counts | 38,244,404 |
| mean counts / cell | 5,910 |
| mean counts / gene | 631 |
| min counts / cell | 220 |
| max counts / cell | 96,766 |
| overdispersion | 3.76 |

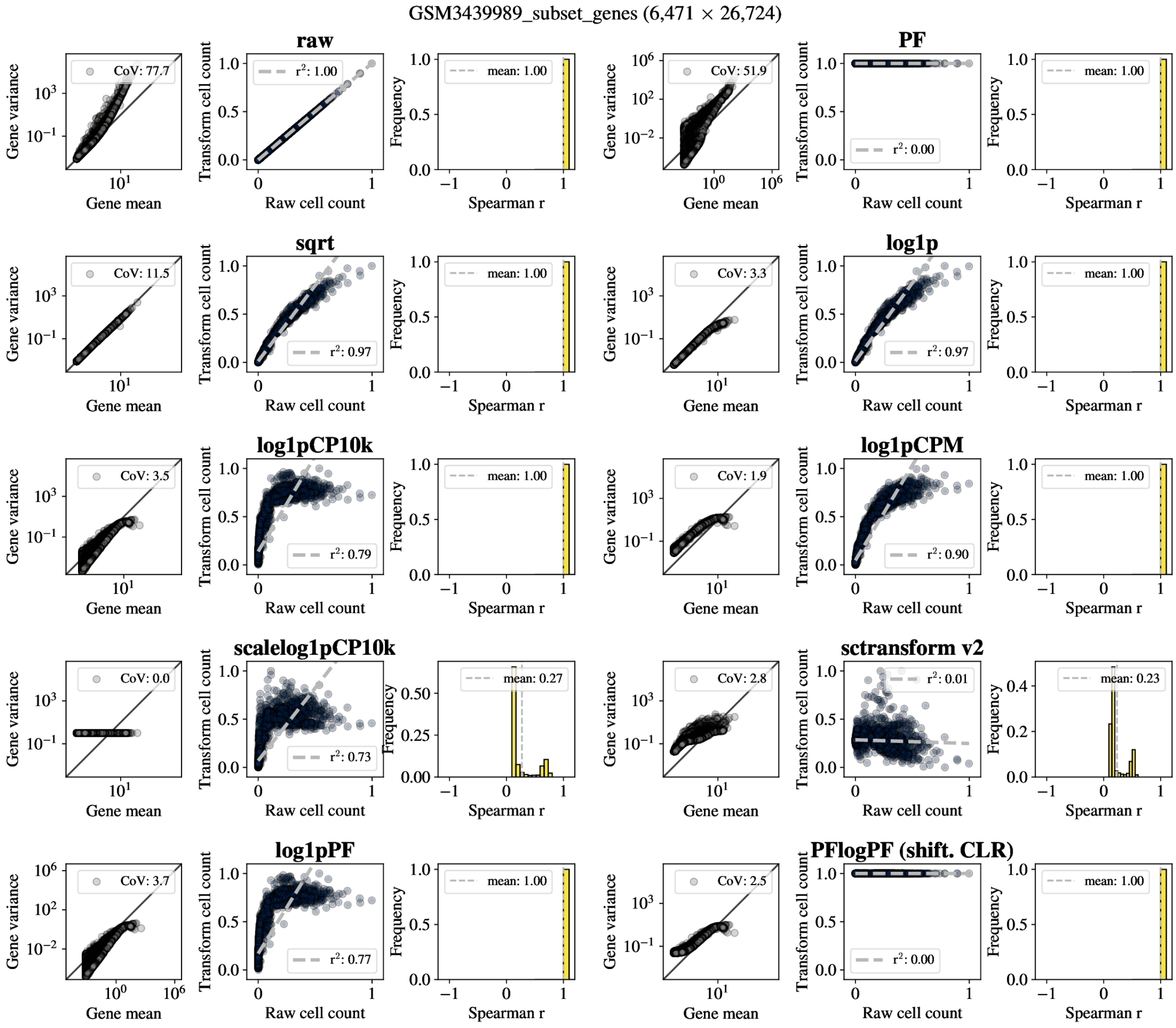


Supplementary Figure 1.264. **GSM3439989**. Per-method comparison of variance stabilization (left), depth normalization (middle), and monotonicity (right) for each normalization on dataset GSM3439989.

| GSM3509837 | |
|---|---|
| cells | 757 |
| genes | 60,623 |
| nonzero entries | 387,385 |
| density | 0.00844 |
| total counts | 1,046,881 |
| mean counts / cell | 1,383 |
| mean counts / gene | 17 |
| min counts / cell | 23 |
| max counts / cell | 23,376 |
| overdispersion | 9.87 |

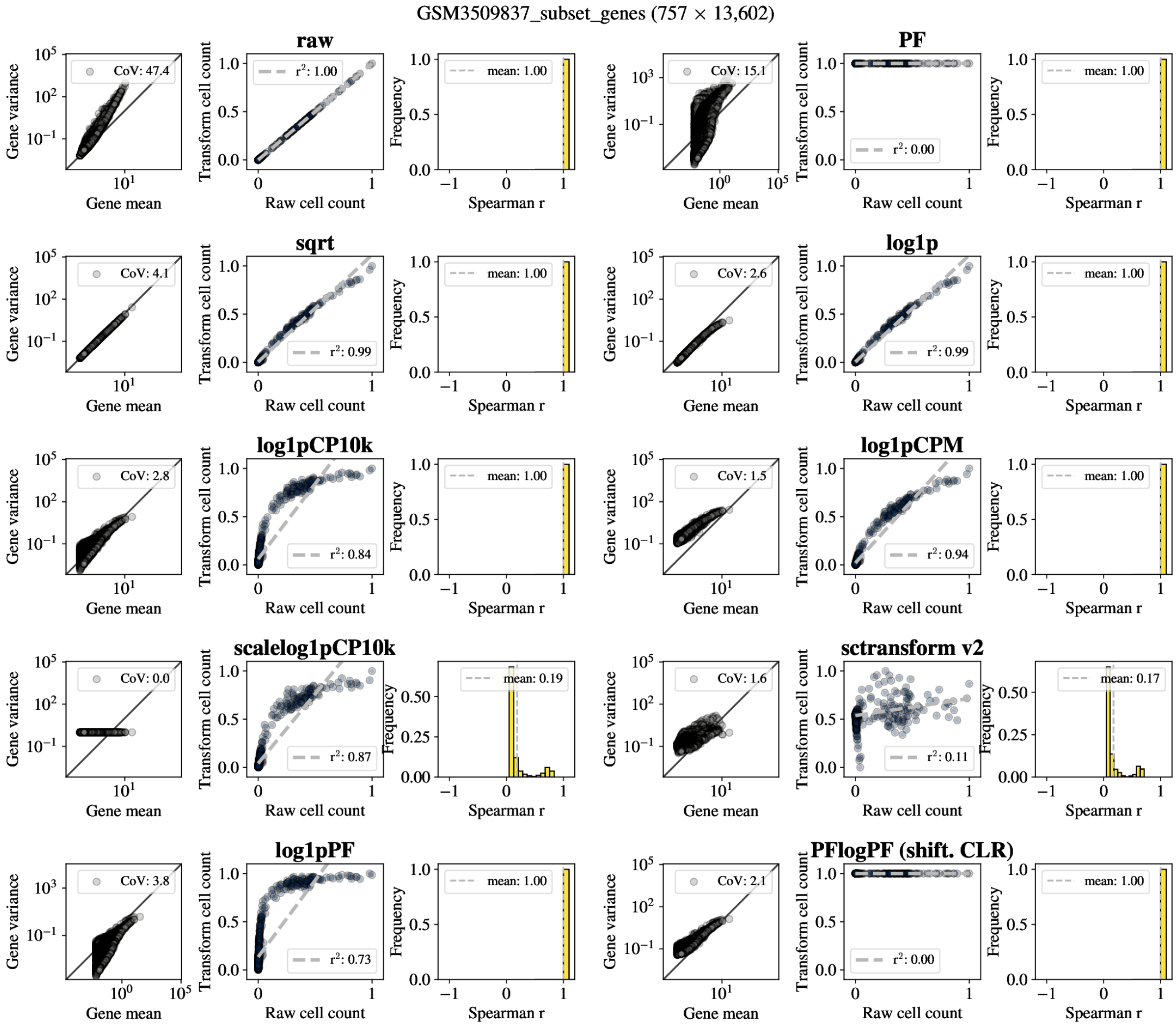


Supplementary Figure 1.265. **GSM3509837**. Per-method comparison of variance stabilization (left), depth normalization (middle), and monotonicity (right) for each normalization on dataset GSM3509837.

| GSM3509838 | |
|---|---|
| cells | 2,614 |
| genes | 60,623 |
| nonzero entries | 7,878,218 |
| density | 0.0497 |
| total counts | 18,111,725 |
| mean counts / cell | 6,929 |
| mean counts / gene | 299 |
| min counts / cell | 3,061 |
| max counts / cell | 26,985 |
| overdispersion | 0.309 |

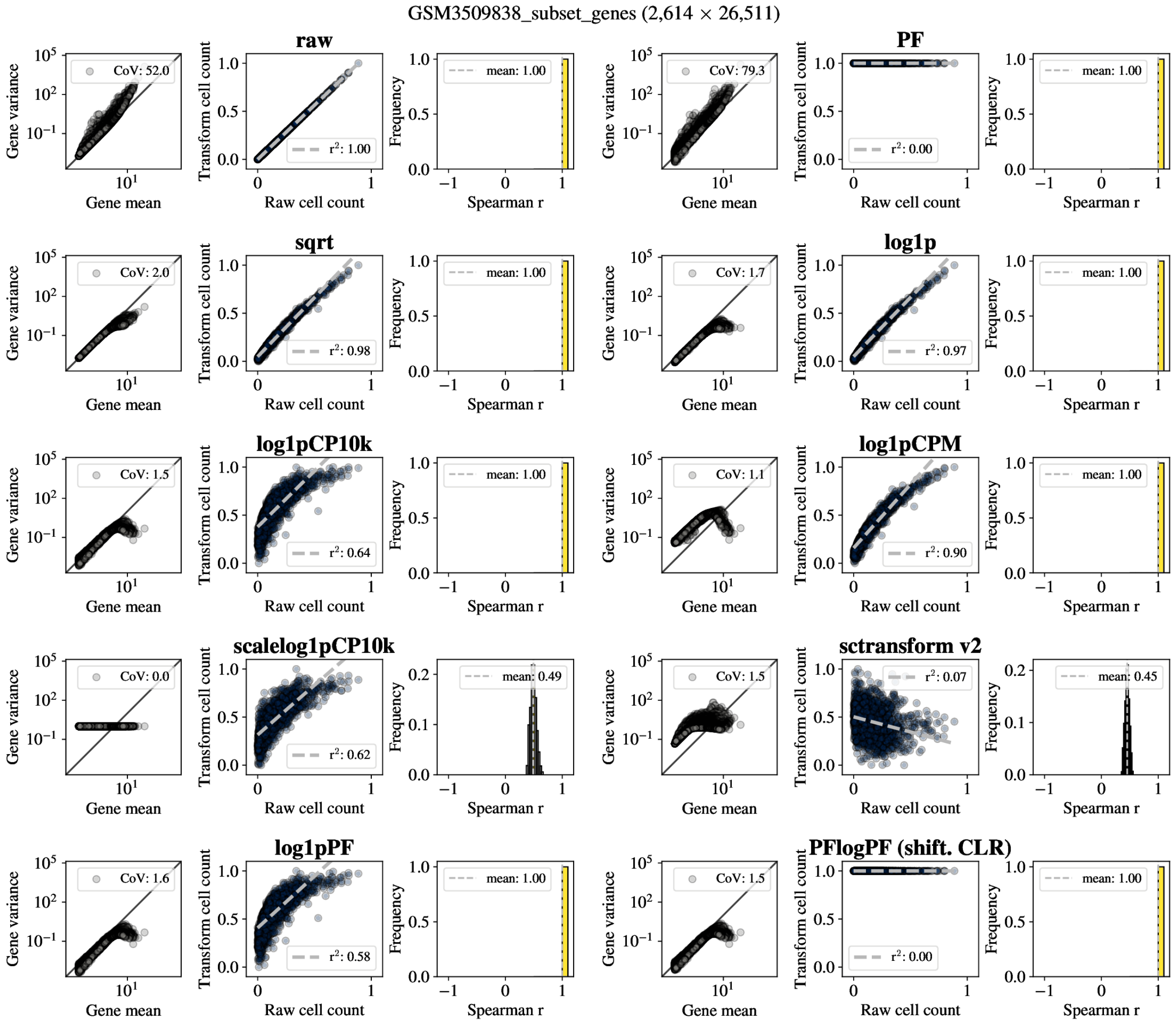


Supplementary Figure 1.266. **GSM3509838**. Per-method comparison of variance stabilization (left), depth normalization (middle), and monotonicity (right) for each normalization on dataset GSM3509838.

| GSM3509839 | |
|---|---|
| cells | 4,507 |
| genes | 60,623 |
| nonzero entries | 8,983,467 |
| density | 0.0329 |
| total counts | 18,573,781 |
| mean counts / cell | 4,121 |
| mean counts / gene | 306 |
| min counts / cell | 215 |
| max counts / cell | 29,452 |
| overdispersion | 0.479 |

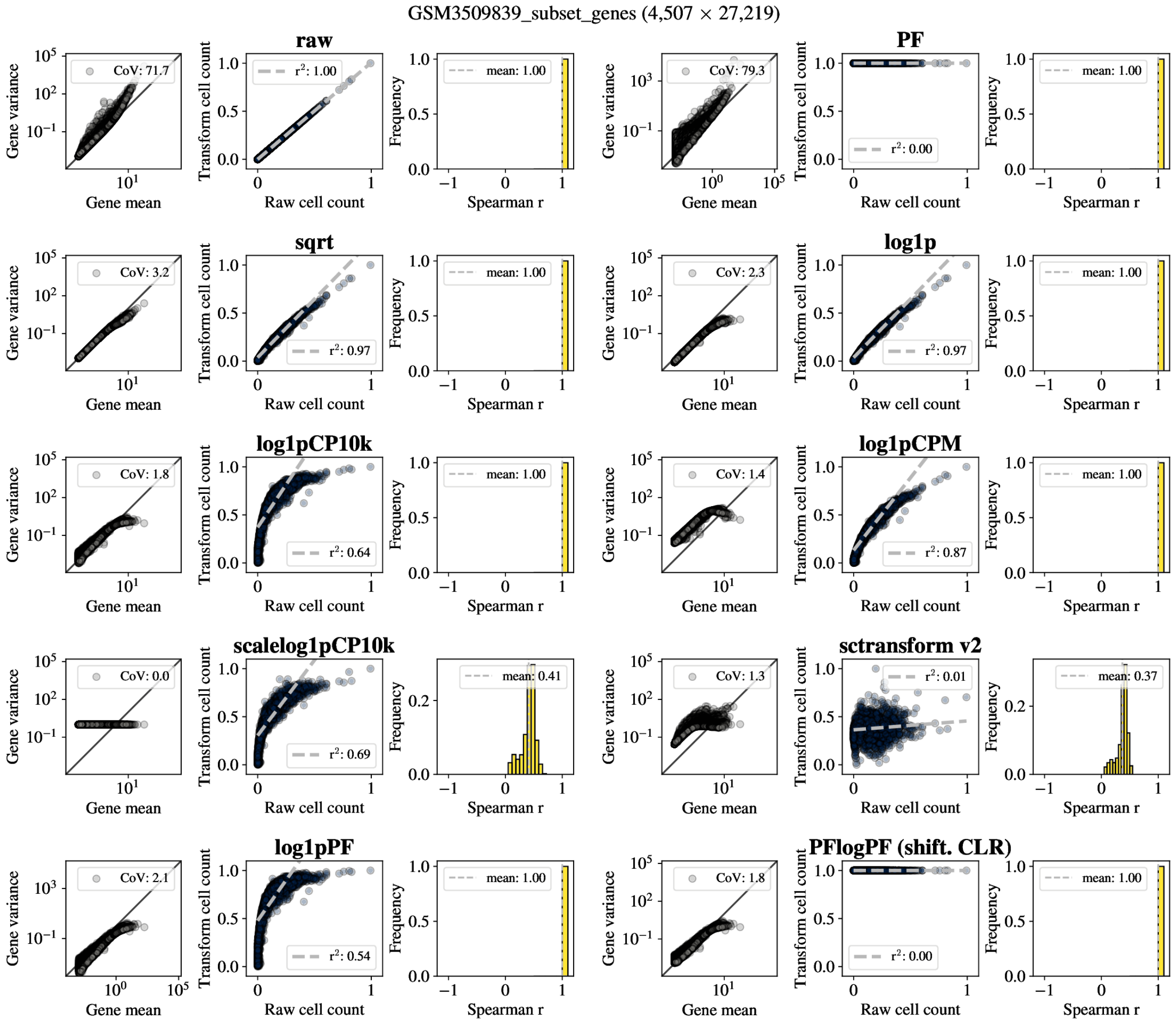


Supplementary Figure 1.267. **GSM3509839**. Per-method comparison of variance stabilization (left), depth normalization (middle), and monotonicity (right) for each normalization on dataset GSM3509839.

| GSM3509840 | |
|---|---|
| cells | 3,057 |
| genes | 60,623 |
| nonzero entries | 6,452,000 |
| density | 0.0348 |
| total counts | 14,327,642 |
| mean counts / cell | 4,687 |
| mean counts / gene | 236 |
| min counts / cell | 209 |
| max counts / cell | 24,173 |
| overdispersion | 0.435 |

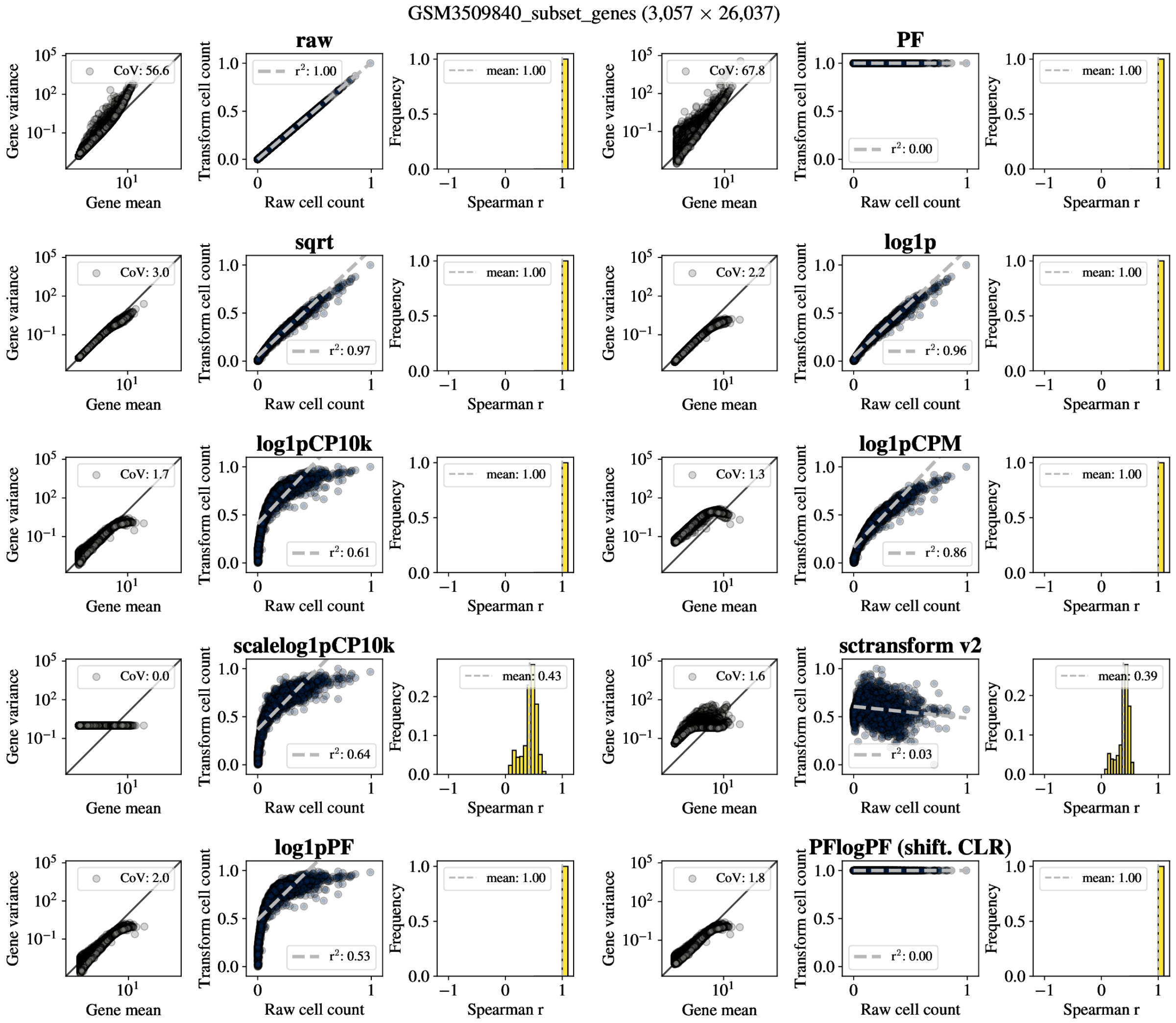


Supplementary Figure 1.268. **GSM3509840**. Per-method comparison of variance stabilization (left), depth normalization (middle), and monotonicity (right) for each normalization on dataset GSM3509840.

| GSM3516947 | |
|---|---|
| cells | 10,132 |
| genes | 60,623 |
| nonzero entries | 20,901,511 |
| density | 0.034 |
| total counts | 54,394,188 |
| mean counts / cell | 5,369 |
| mean counts / gene | 897 |
| min counts / cell | 1,653 |
| max counts / cell | 24,637 |
| overdispersion | 0.396 |

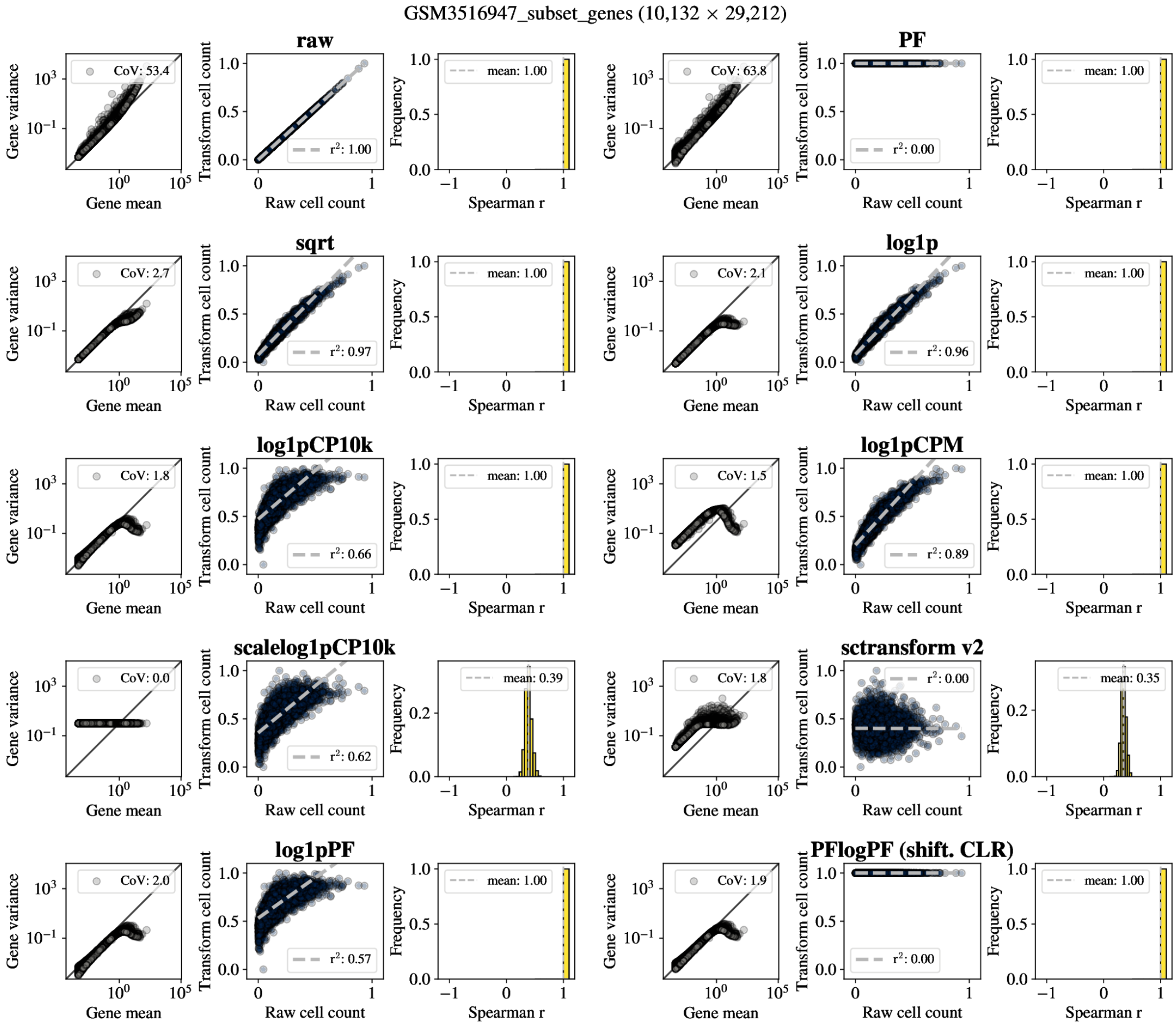


Supplementary Figure 1.269. **GSM3516947**. Per-method comparison of variance stabilization (left), depth normalization (middle), and monotonicity (right) for each normalization on dataset GSM3516947.

| GSM3516948 | |
|---|---|
| cells | 13,930 |
| genes | 60,623 |
| nonzero entries | 24,554,674 |
| density | 0.0291 |
| total counts | 78,793,238 |
| mean counts / cell | 5,656 |
| mean counts / gene | 1,300 |
| min counts / cell | 2,525 |
| max counts / cell | 31,285 |
| overdispersion | 0.432 |

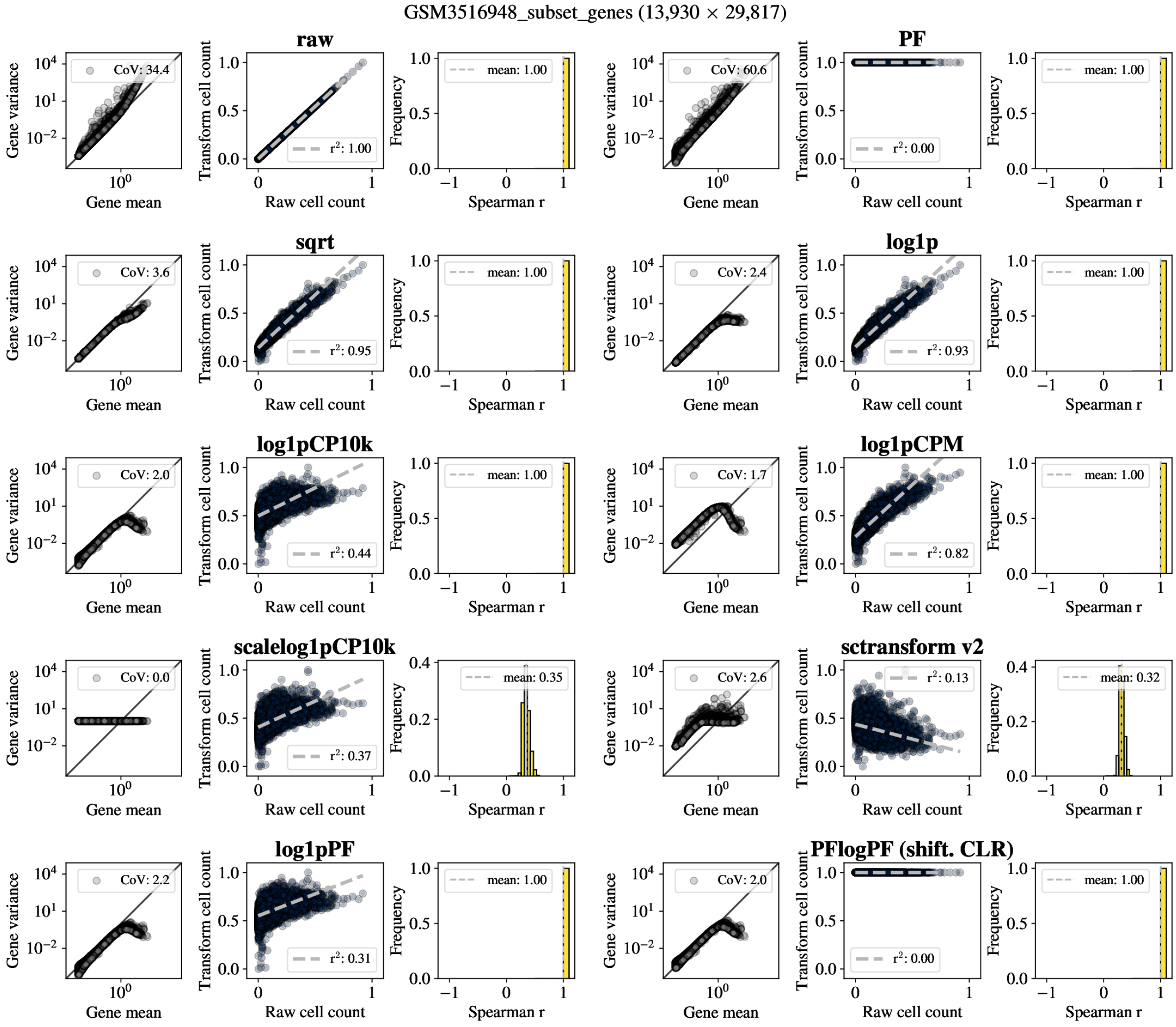


Supplementary Figure 1.270. **GSM3516948**. Per-method comparison of variance stabilization (left), depth normalization (middle), and monotonicity (right) for each normalization on dataset GSM3516948.

| GSM3526583 | |
|---|---|
| cells | 3,101 |
| genes | 60,623 |
| nonzero entries | 5,043,432 |
| density | 0.0268 |
| total counts | 11,078,451 |
| mean counts / cell | 3,573 |
| mean counts / gene | 183 |
| min counts / cell | 1,343 |
| max counts / cell | 29,985 |
| overdispersion | 0.627 |

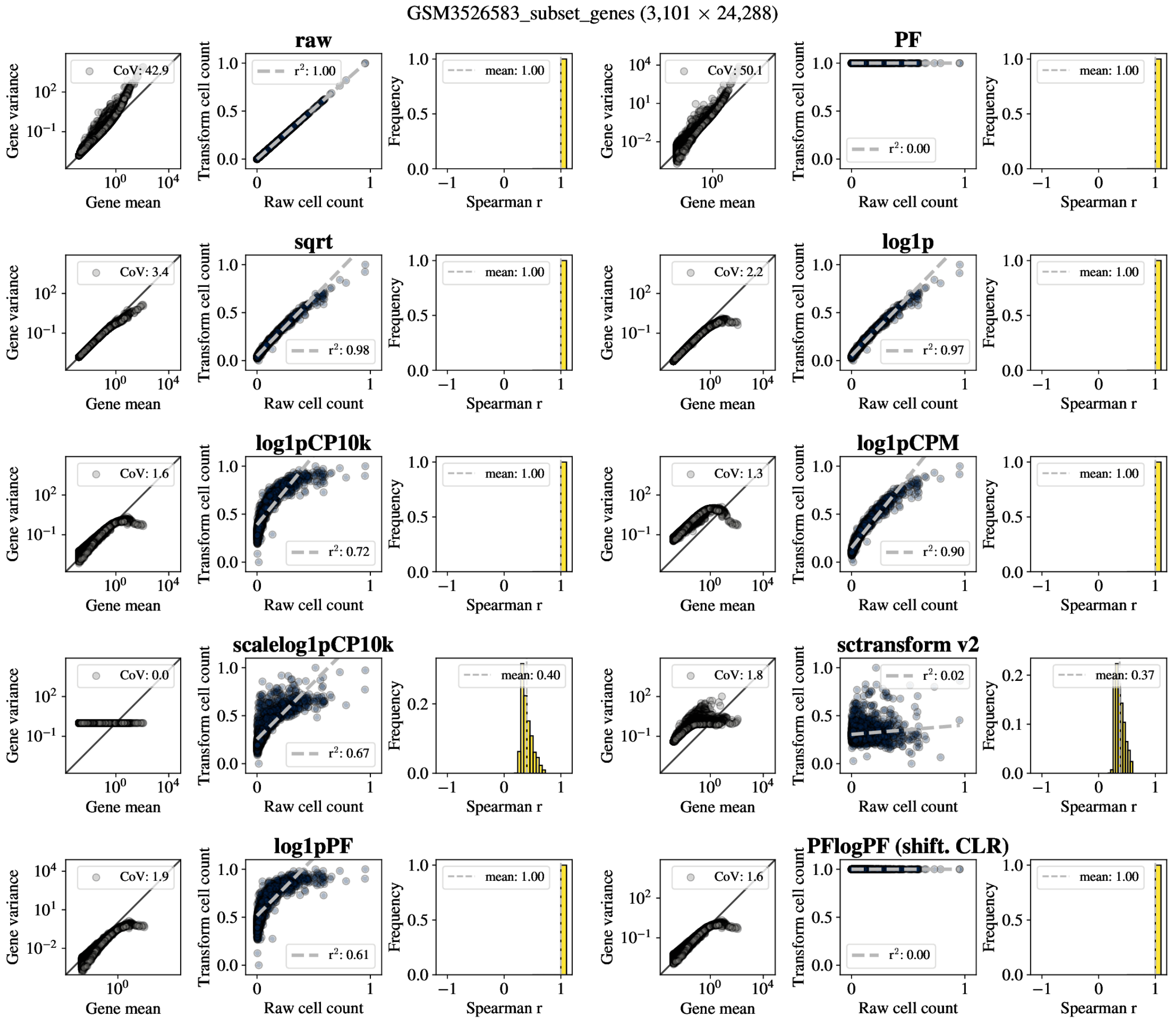


Supplementary Figure 1.271. **GSM3526583**. Per-method comparison of variance stabilization (left), depth normalization (middle), and monotonicity (right) for each normalization on dataset GSM3526583.

| GSM3526584 | |
|---|---|
| cells | 5,273 |
| genes | 60,623 |
| nonzero entries | 6,762,616 |
| density | 0.0212 |
| total counts | 14,937,647 |
| mean counts / cell | 2,833 |
| mean counts / gene | 246 |
| min counts / cell | 1,086 |
| max counts / cell | 45,801 |
| overdispersion | 0.373 |

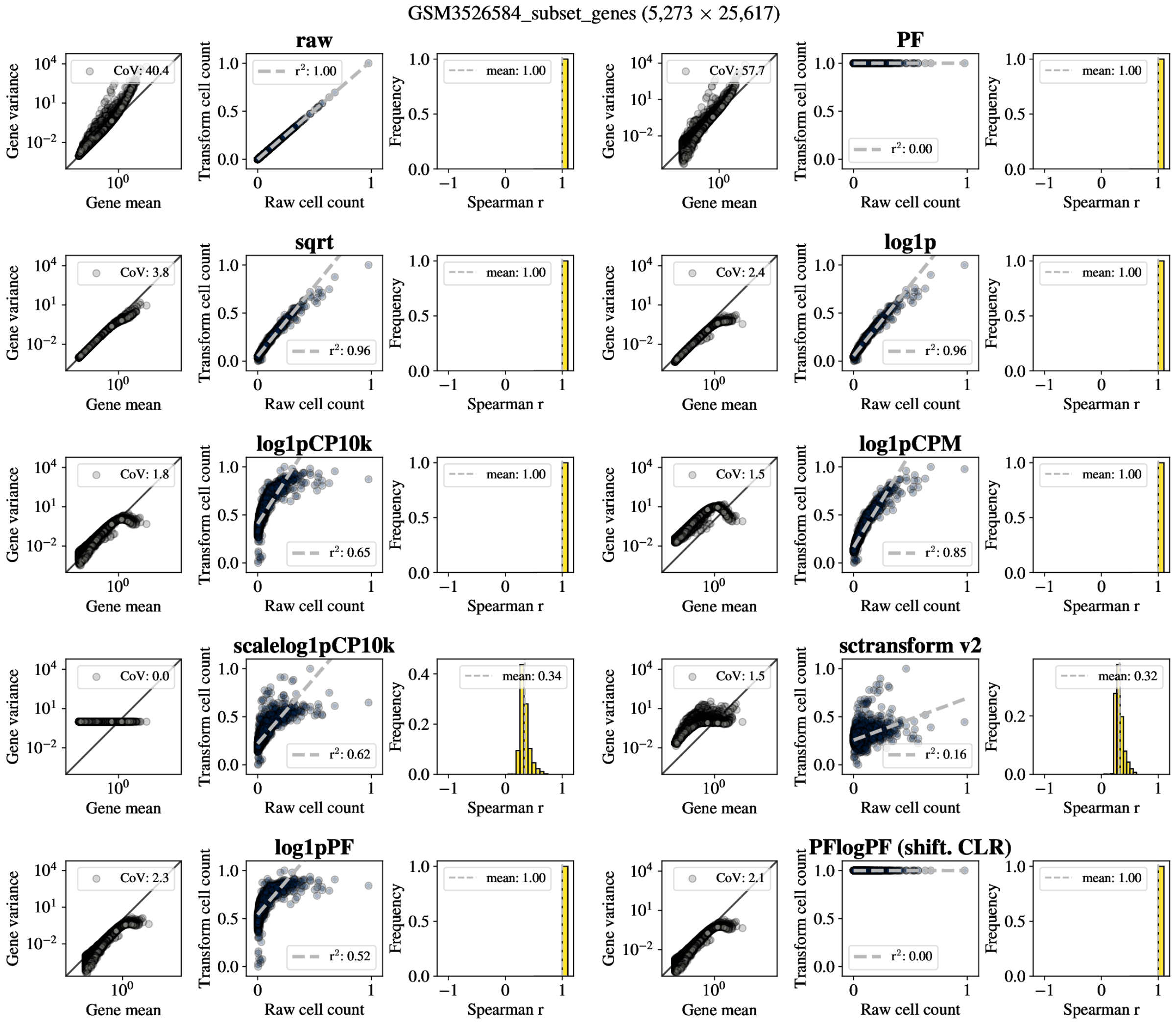


Supplementary Figure 1.272. **GSM3526584**. Per-method comparison of variance stabilization (left), depth normalization (middle), and monotonicity (right) for each normalization on dataset GSM3526584.

| GSM3526585 | |
|---|---|
| cells | 8,802 |
| genes | 60,623 |
| nonzero entries | 9,094,883 |
| density | 0.017 |
| total counts | 17,603,845 |
| mean counts / cell | 2,000 |
| mean counts / gene | 290 |
| min counts / cell | 365 |
| max counts / cell | 25,558 |
| overdispersion | 1.31 |

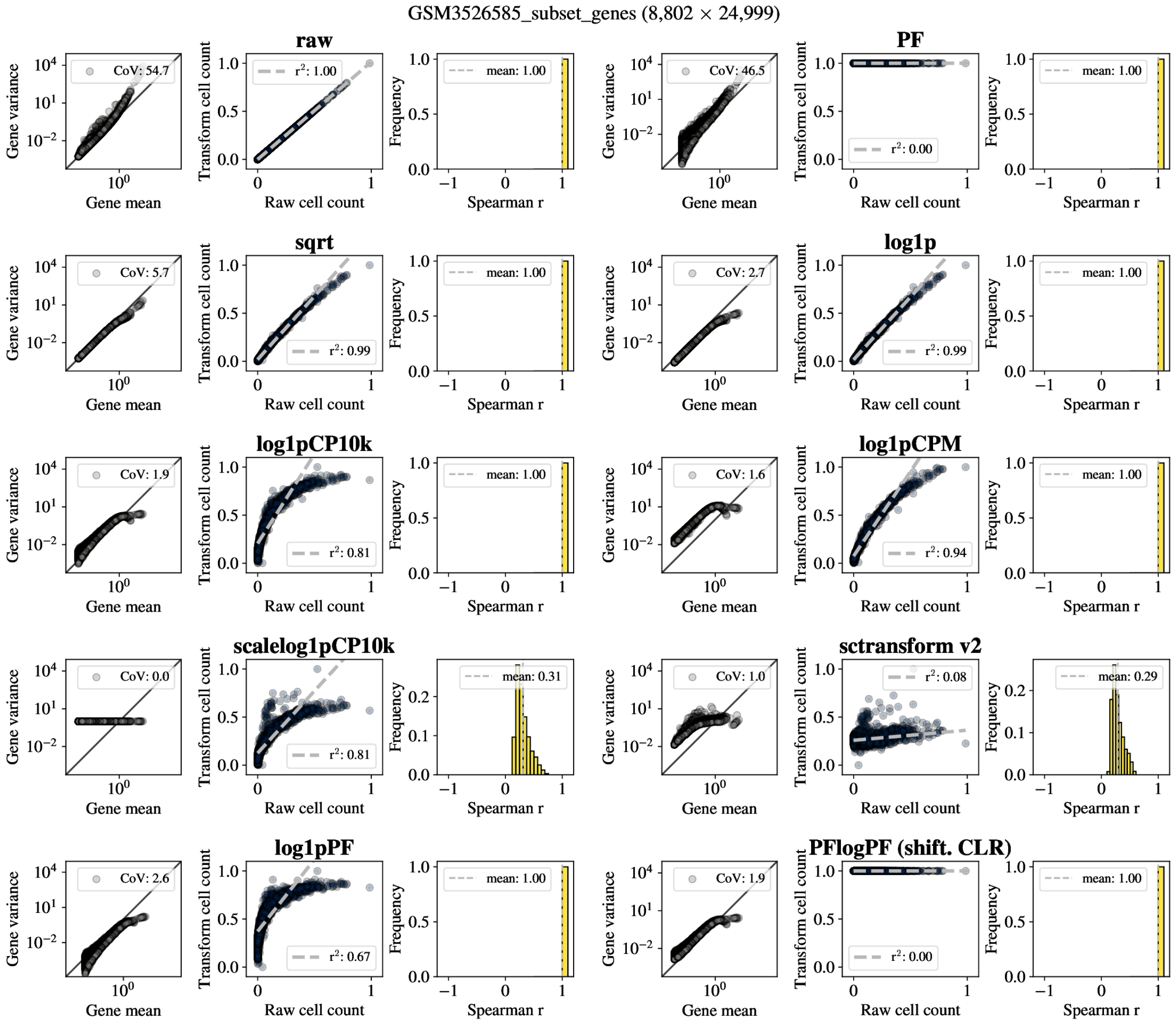


Supplementary Figure 1.273. **GSM3526585**. Per-method comparison of variance stabilization (left), depth normalization (middle), and monotonicity (right) for each normalization on dataset GSM3526585.

| GSM3526586 | |
|---|---|
| cells | 9,260 |
| genes | 60,623 |
| nonzero entries | 10,790,291 |
| density | 0.0192 |
| total counts | 21,766,351 |
| mean counts / cell | 2,351 |
| mean counts / gene | 359 |
| min counts / cell | 450 |
| max counts / cell | 21,596 |
| overdispersion | 2.15 |

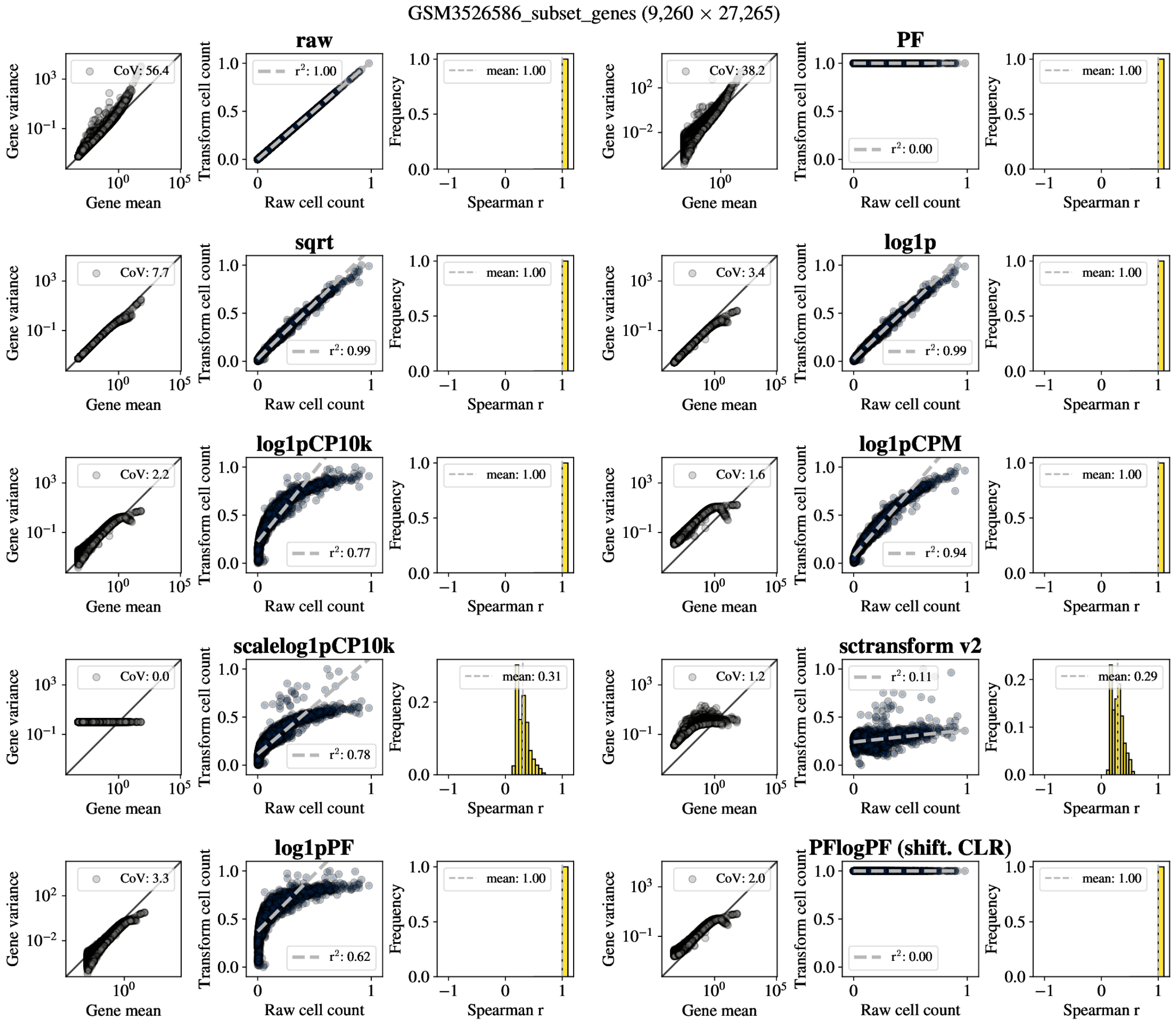


Supplementary Figure 1.274. **GSM3526586**. Per-method comparison of variance stabilization (left), depth normalization (middle), and monotonicity (right) for each normalization on dataset GSM3526586.

| GSM3526587 | |
|---|---|
| cells | 3,798 |
| genes | 60,623 |
| nonzero entries | 12,355,099 |
| density | 0.0537 |
| total counts | 29,013,098 |
| mean counts / cell | 7,639 |
| mean counts / gene | 479 |
| min counts / cell | 3,014 |
| max counts / cell | 48,738 |
| overdispersion | 3.33 |

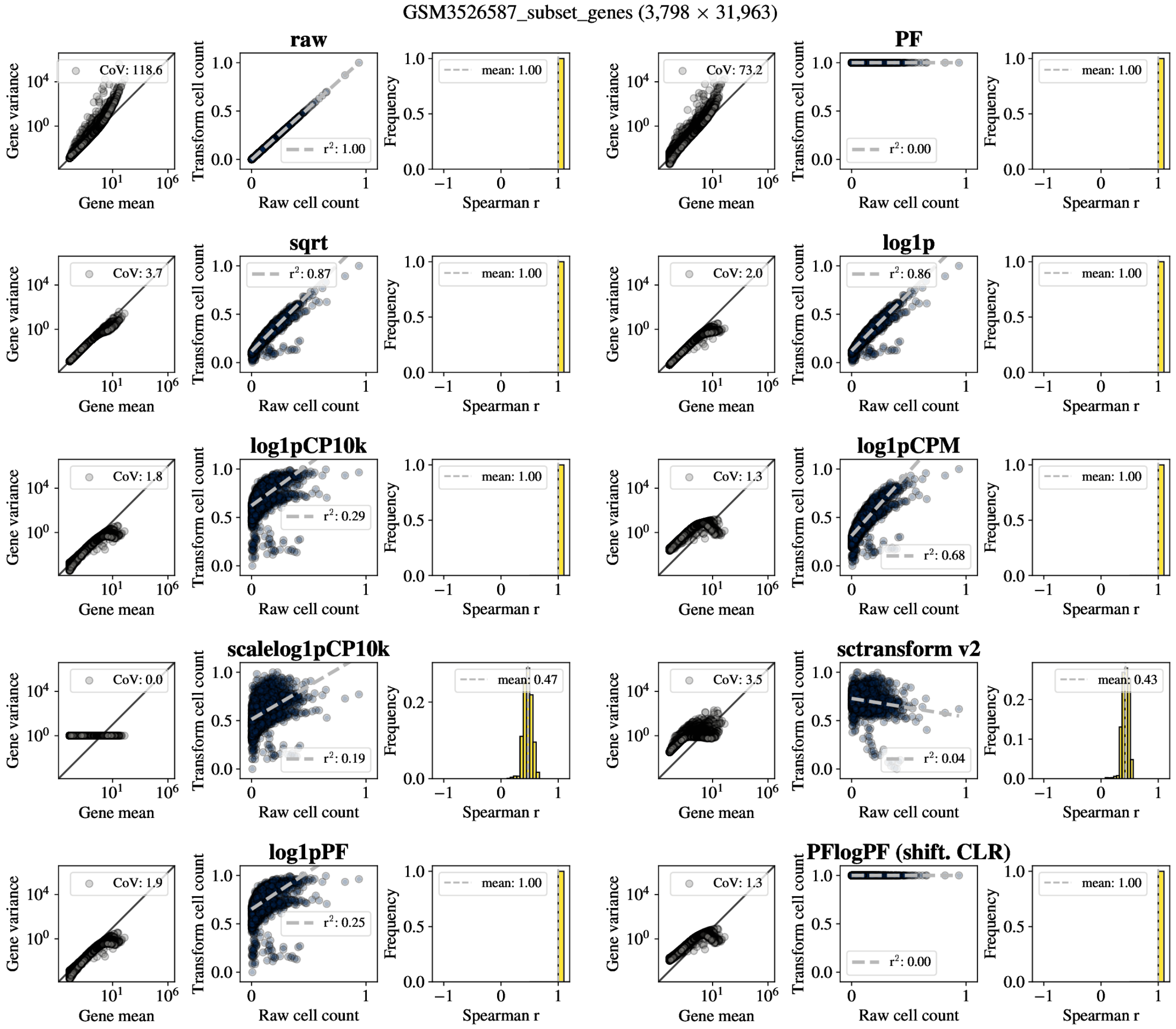


Supplementary Figure 1.275. **GSM3526587**. Per-method comparison of variance stabilization (left), depth normalization (middle), and monotonicity (right) for each normalization on dataset GSM3526587.

| GSM3526588 | |
|---|---|
| cells | 7,558 |
| genes | 60,623 |
| nonzero entries | 13,001,129 |
| density | 0.0284 |
| total counts | 29,408,714 |
| mean counts / cell | 3,891 |
| mean counts / gene | 485 |
| min counts / cell | 108 |
| max counts / cell | 32,336 |
| overdispersion | 1.49 |

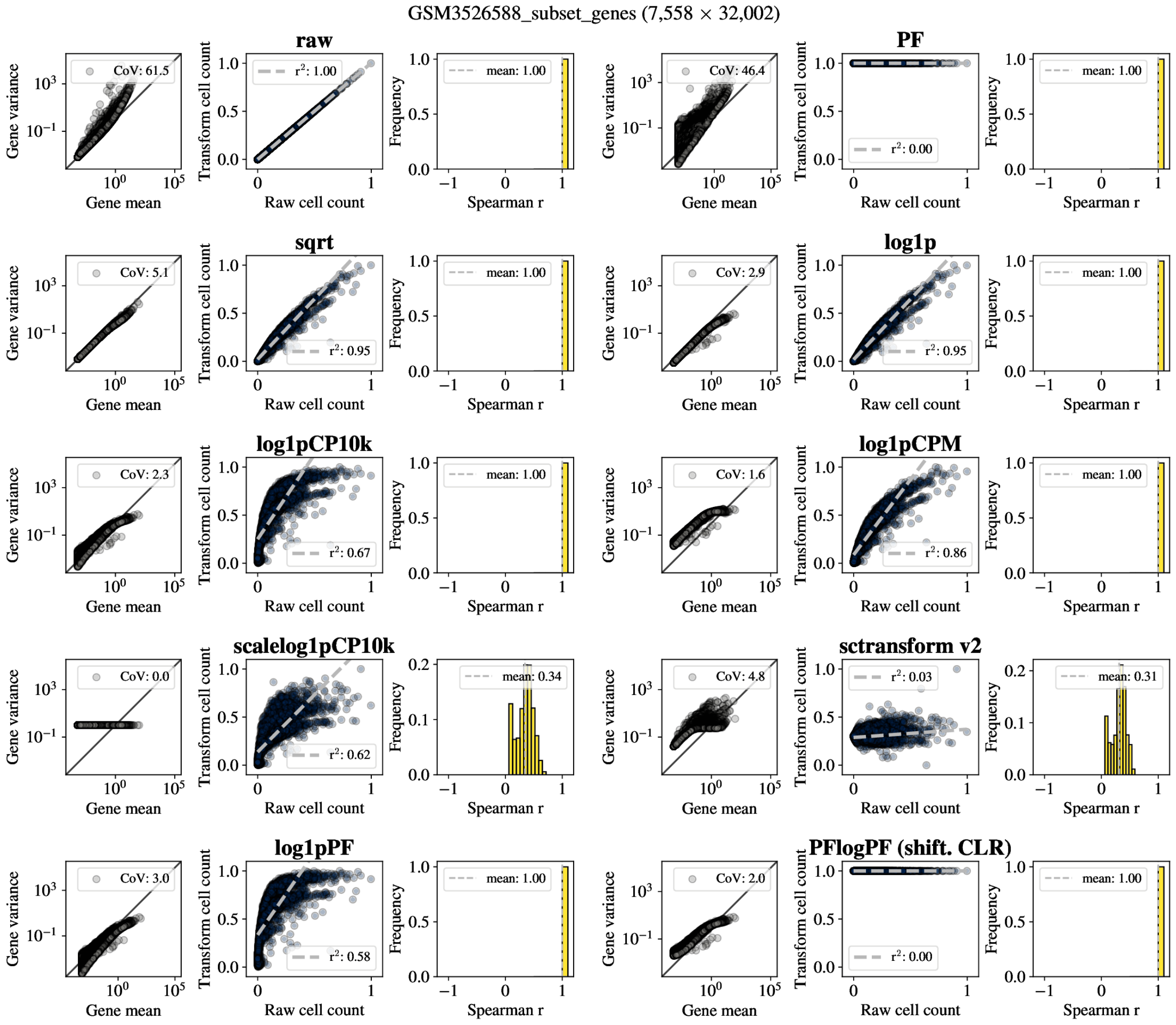


Supplementary Figure 1.276. **GSM3526588**. Per-method comparison of variance stabilization (left), depth normalization (middle), and monotonicity (right) for each normalization on dataset GSM3526588.

| GSM3526589 | |
|---|---|
| cells | 12,633 |
| genes | 60,623 |
| nonzero entries | 15,363,597 |
| density | 0.0201 |
| total counts | 33,925,554 |
| mean counts / cell | 2,685 |
| mean counts / gene | 560 |
| min counts / cell | 255 |
| max counts / cell | 50,190 |
| overdispersion | 2.4 |

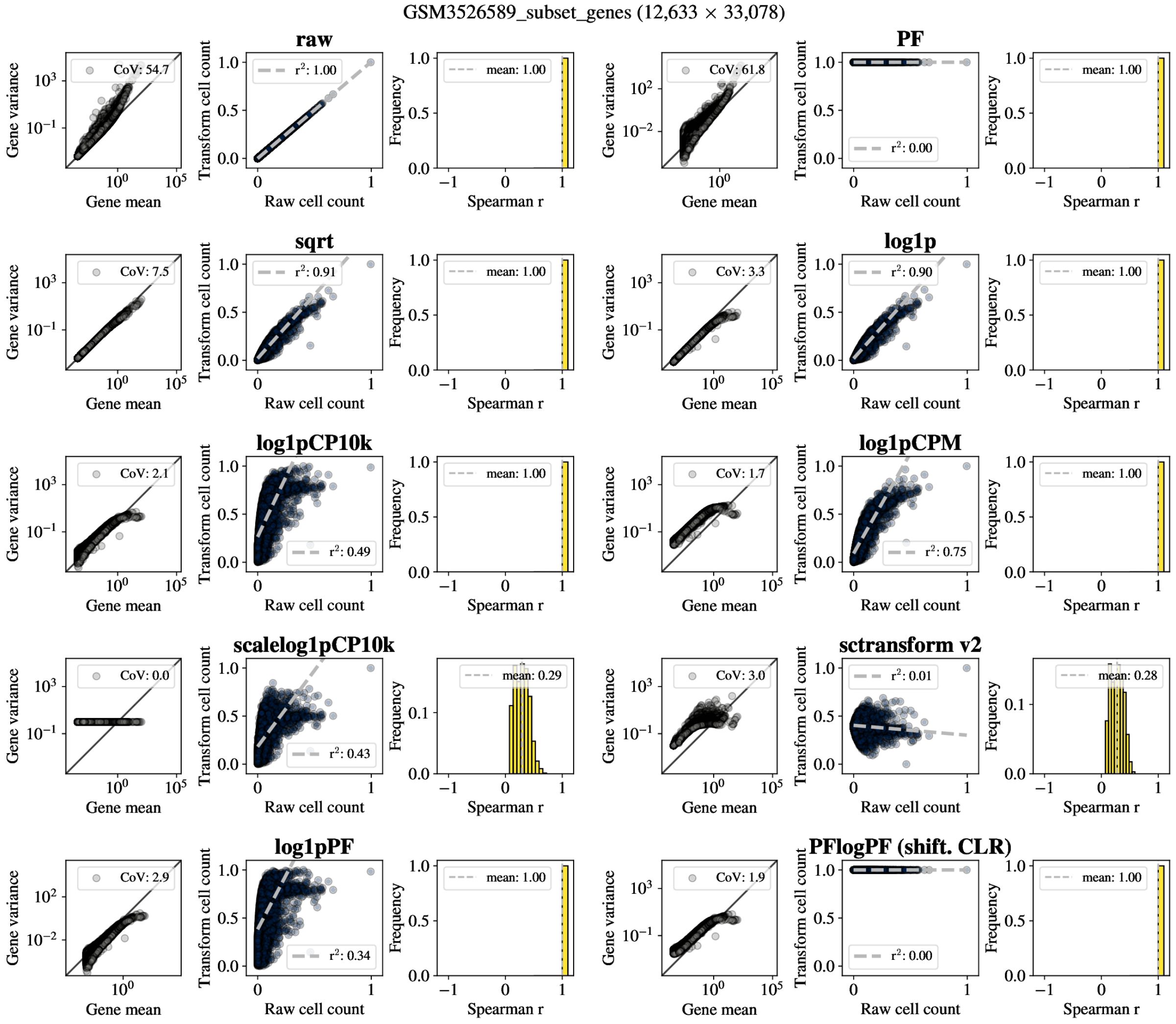


Supplementary Figure 1.277. **GSM3526589**. Per-method comparison of variance stabilization (left), depth normalization (middle), and monotonicity (right) for each normalization on dataset GSM3526589.

| GSM3526590 | |
|---|---|
| cells | 12,589 |
| genes | 60,623 |
| nonzero entries | 22,787,775 |
| density | 0.0299 |
| total counts | 50,990,995 |
| mean counts / cell | 4,050 |
| mean counts / gene | 841 |
| min counts / cell | 208 |
| max counts / cell | 33,860 |
| overdispersion | 1.6 |

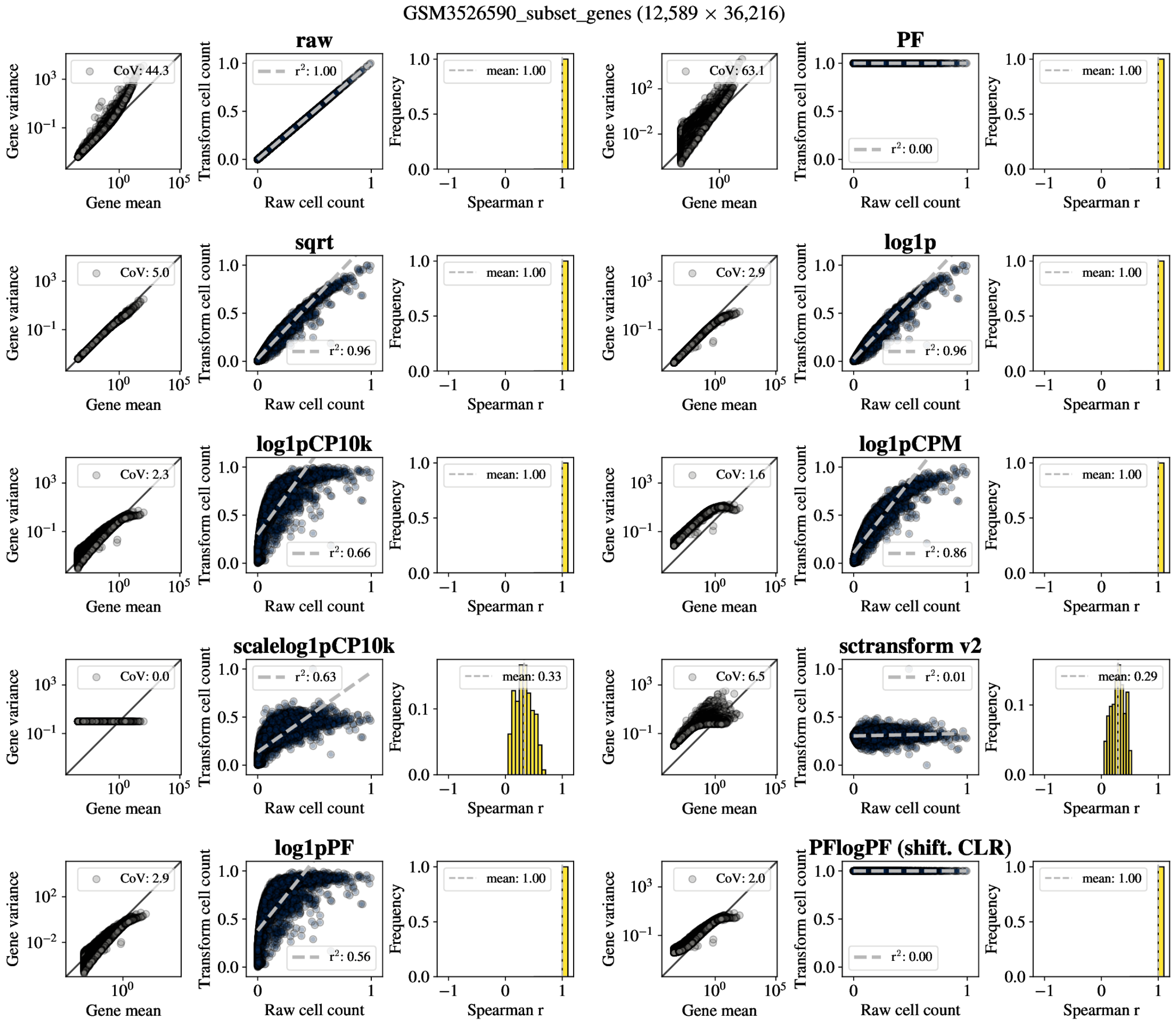


Supplementary Figure 1.278. **GSM3526590**. Per-method comparison of variance stabilization (left), depth normalization (middle), and monotonicity (right) for each normalization on dataset GSM3526590.

| GSM3529821 | |
|---|---|
| cells | 15,247 |
| genes | 60,623 |
| nonzero entries | 19,789,315 |
| density | 0.0214 |
| total counts | 76,621,808 |
| mean counts / cell | 5,025 |
| mean counts / gene | 1,264 |
| min counts / cell | 80 |
| max counts / cell | 39,154 |
| overdispersion | 1.82 |

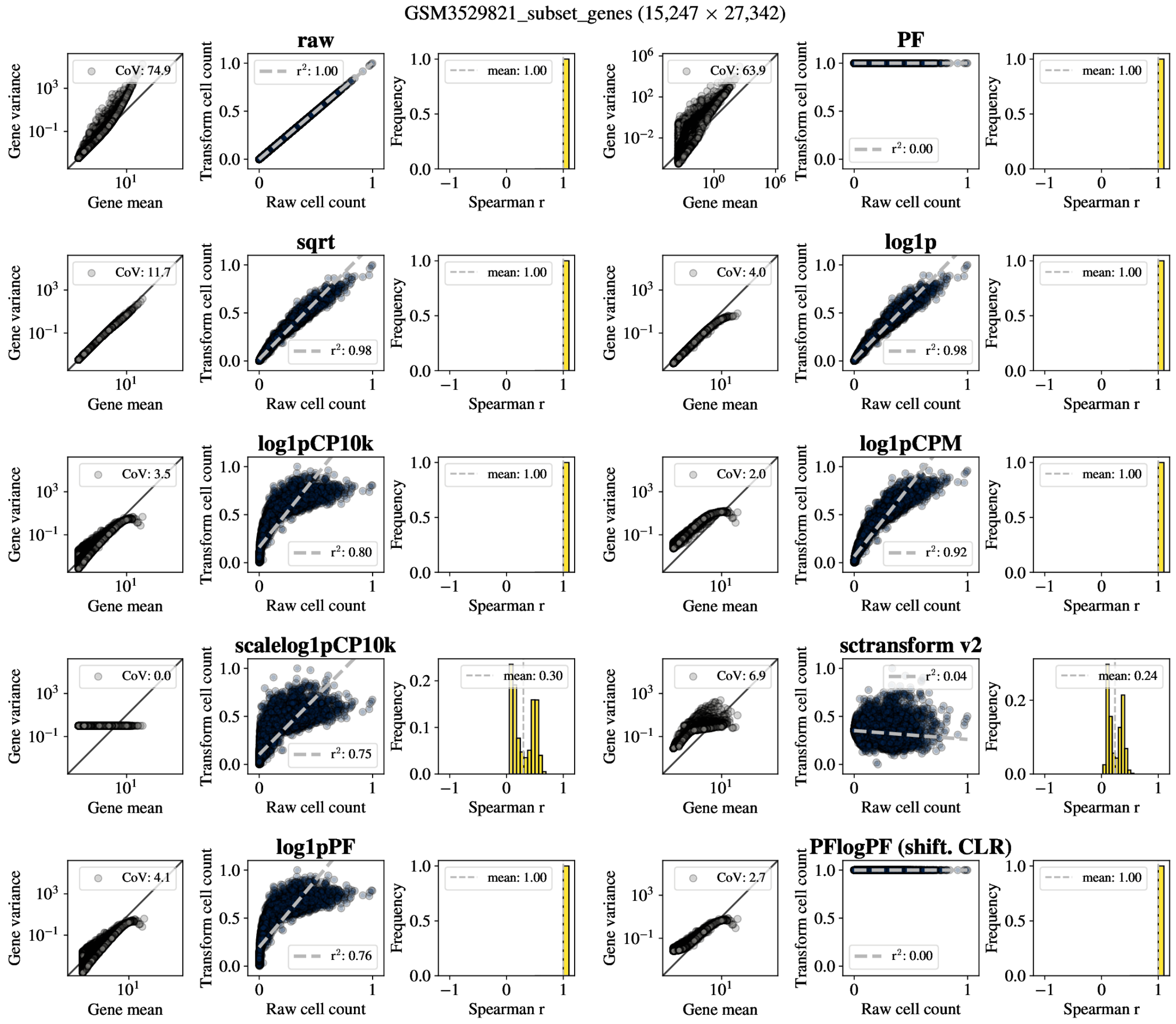


Supplementary Figure 1.279. **GSM3529821**. Per-method comparison of variance stabilization (left), depth normalization (middle), and monotonicity (right) for each normalization on dataset GSM3529821.

| GSM3535276 | |
|---|---|
| cells | 12,849 |
| genes | 60,623 |
| nonzero entries | 24,388,011 |
| density | 0.0313 |
| total counts | 63,027,759 |
| mean counts / cell | 4,905 |
| mean counts / gene | 1,040 |
| min counts / cell | 111 |
| max counts / cell | 81,906 |
| overdispersion | 0.997 |

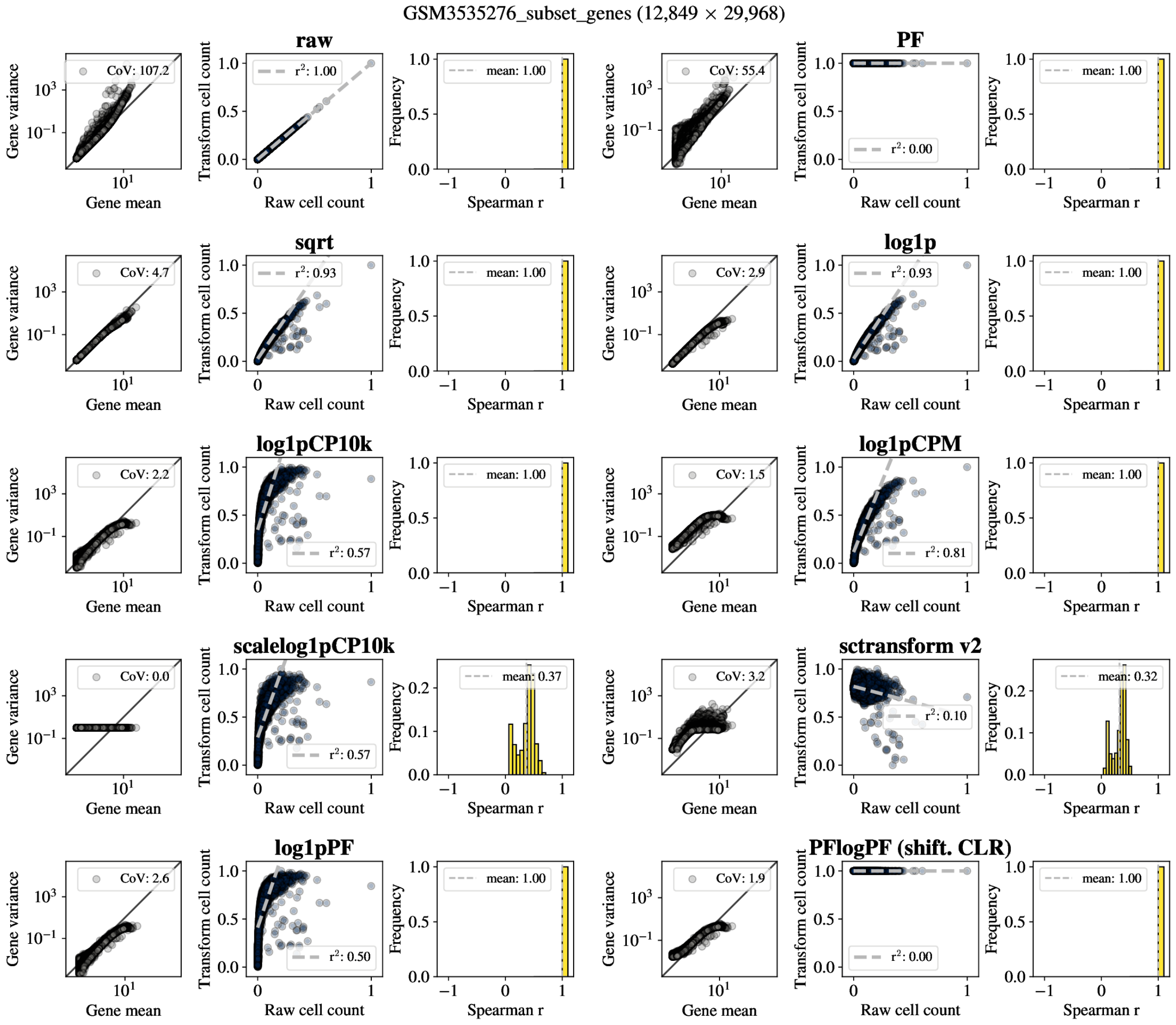


Supplementary Figure 1.280. **GSM3535276**. Per-method comparison of variance stabilization (left), depth normalization (middle), and monotonicity (right) for each normalization on dataset GSM3535276.

| GSM3535277 | |
|---|---|
| cells | 6,345 |
| genes | 60,623 |
| nonzero entries | 12,217,315 |
| density | 0.0318 |
| total counts | 33,306,101 |
| mean counts / cell | 5,249 |
| mean counts / gene | 549 |
| min counts / cell | 68 |
| max counts / cell | 71,174 |
| overdispersion | 1.12 |

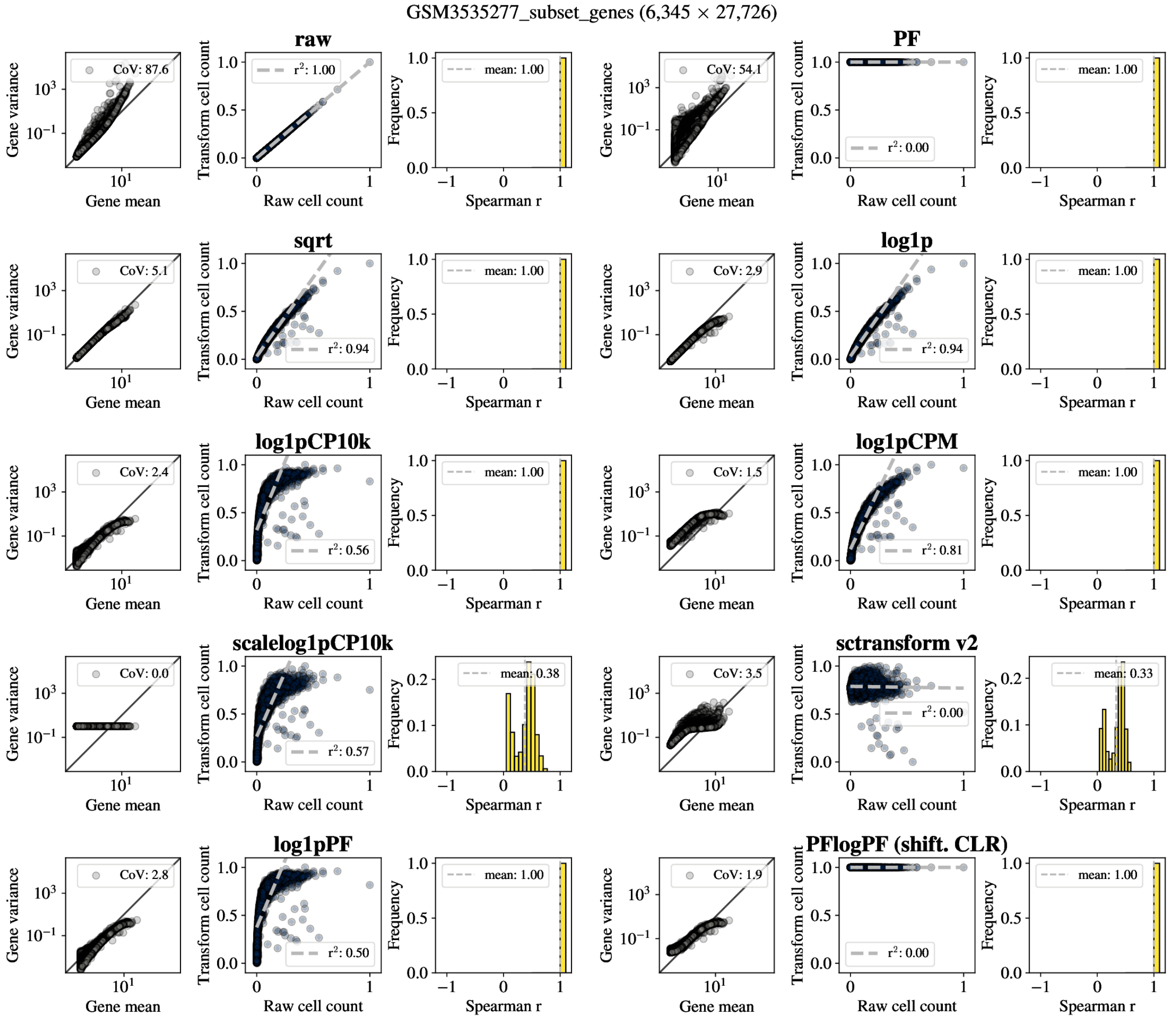


Supplementary Figure 1.281. **GSM3535277**. Per-method comparison of variance stabilization (left), depth normalization (middle), and monotonicity (right) for each normalization on dataset GSM3535277.

| GSM3535278 | |
|---|---|
| cells | 6,062 |
| genes | 60,623 |
| nonzero entries | 12,012,366 |
| density | 0.0327 |
| total counts | 29,451,076 |
| mean counts / cell | 4,858 |
| mean counts / gene | 486 |
| min counts / cell | 2,731 |
| max counts / cell | 37,918 |
| overdispersion | 0.411 |

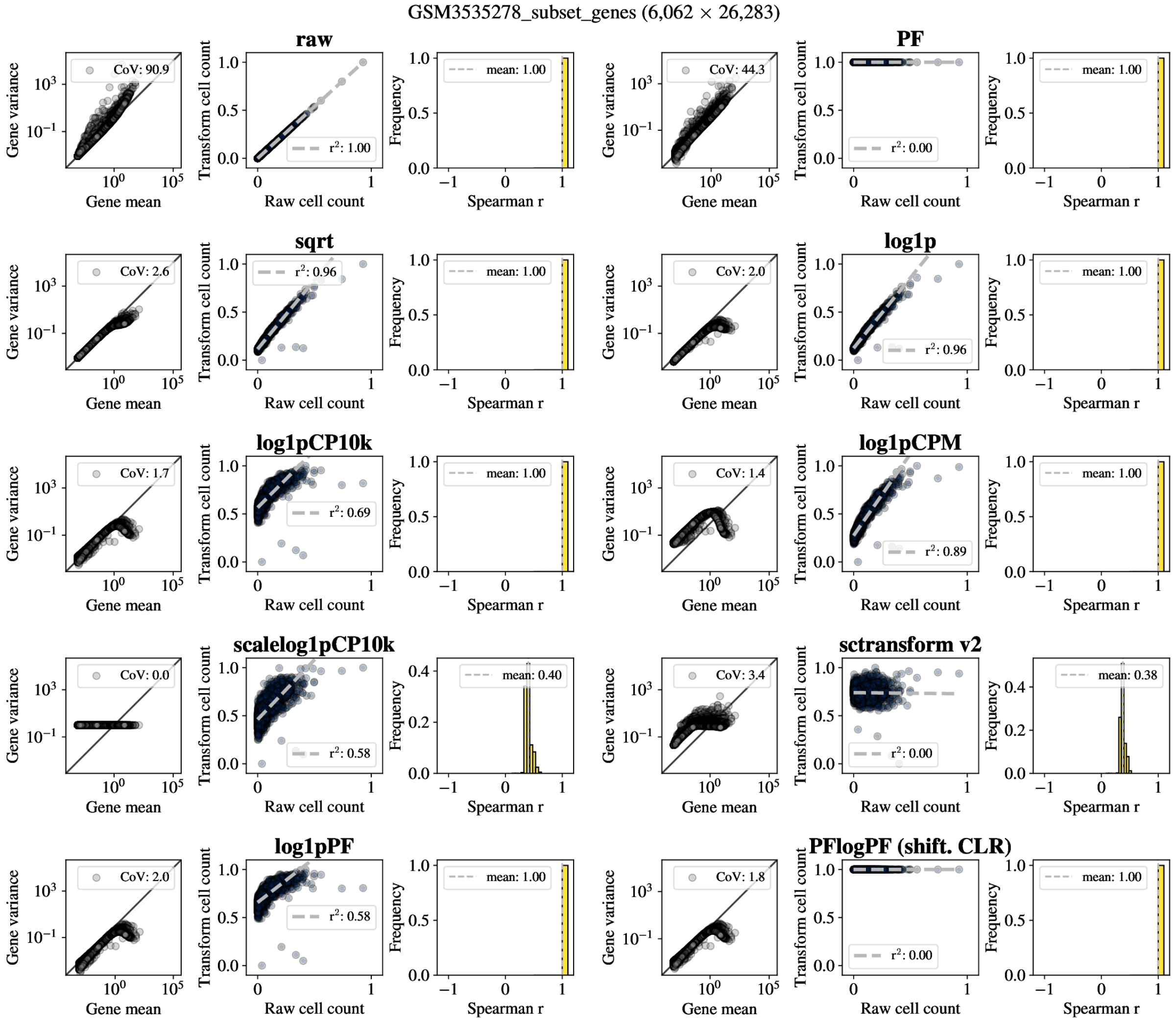


Supplementary Figure 1.282. **GSM3535278**. Per-method comparison of variance stabilization (left), depth normalization (middle), and monotonicity (right) for each normalization on dataset GSM3535278.

| GSM3535279 | |
|---|---|
| cells | 6,394 |
| genes | 60,623 |
| nonzero entries | 11,769,062 |
| density | 0.0304 |
| total counts | 32,926,259 |
| mean counts / cell | 5,150 |
| mean counts / gene | 543 |
| min counts / cell | 179 |
| max counts / cell | 38,309 |
| overdispersion | 1.03 |

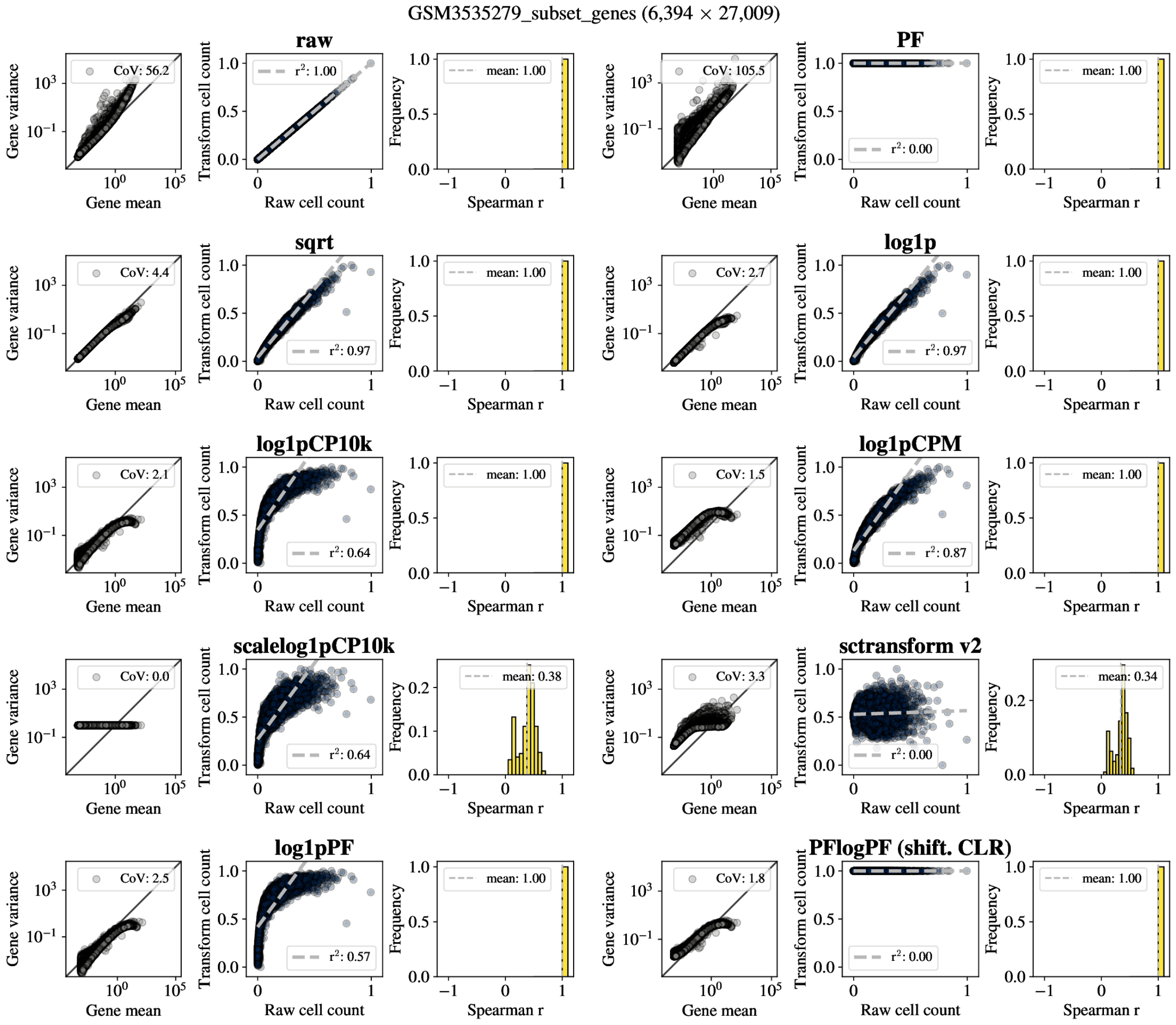


Supplementary Figure 1.283. **GSM3535279**. Per-method comparison of variance stabilization (left), depth normalization (middle), and monotonicity (right) for each normalization on dataset GSM3535279.

| GSM3535280 | |
|---|---|
| cells | 5,700 |
| genes | 60,623 |
| nonzero entries | 13,989,159 |
| density | 0.0405 |
| total counts | 38,429,239 |
| mean counts / cell | 6,742 |
| mean counts / gene | 634 |
| min counts / cell | 2,209 |
| max counts / cell | 40,376 |
| overdispersion | 0.573 |

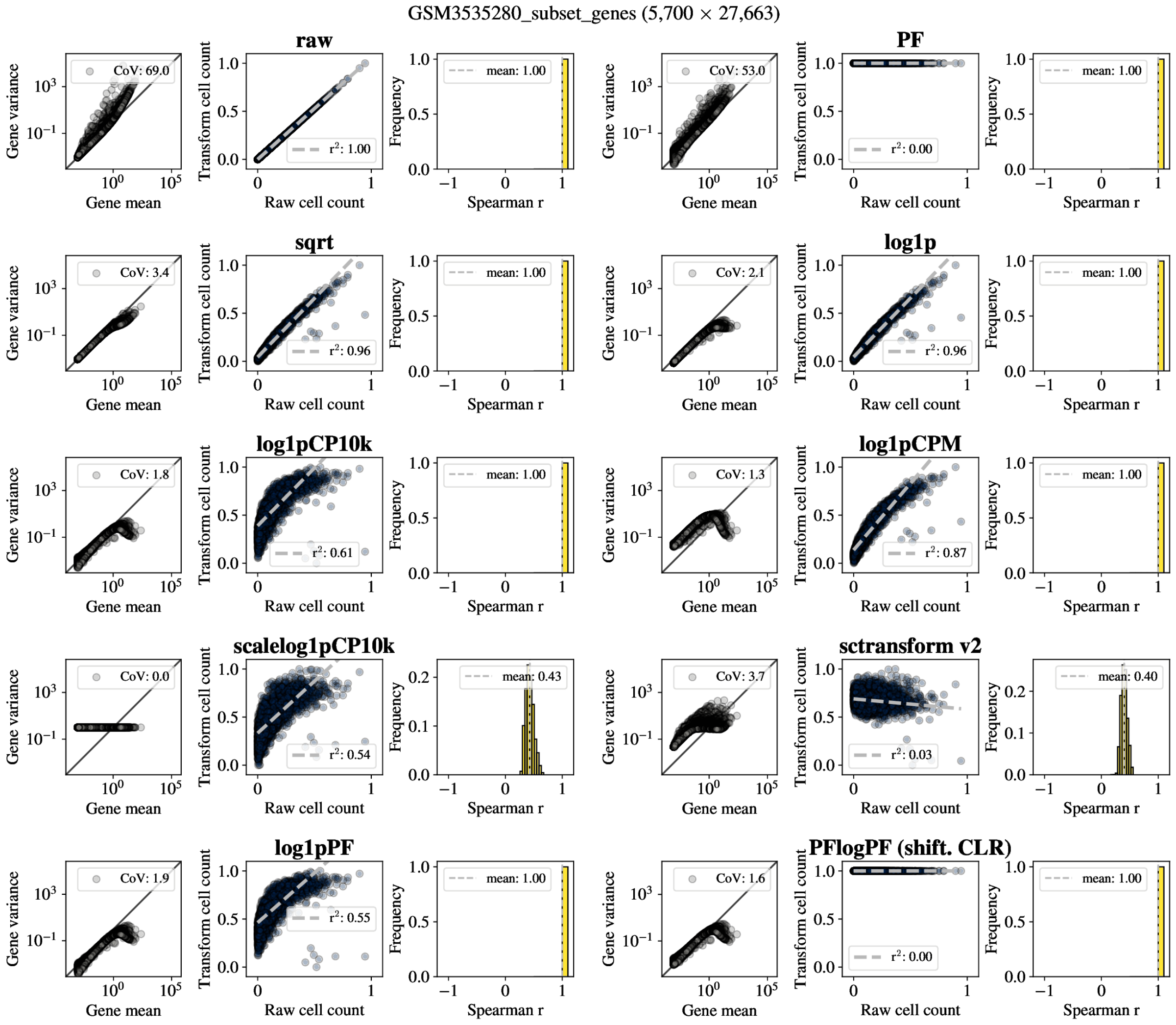


Supplementary Figure 1.284. **GSM3535280**. Per-method comparison of variance stabilization (left), depth normalization (middle), and monotonicity (right) for each normalization on dataset GSM3535280.

| GSM3535281 | |
|---|---|
| cells | 4,064 |
| genes | 60,623 |
| nonzero entries | 8,061,109 |
| density | 0.0327 |
| total counts | 19,063,463 |
| mean counts / cell | 4,691 |
| mean counts / gene | 314 |
| min counts / cell | 136 |
| max counts / cell | 23,103 |
| overdispersion | 0.473 |

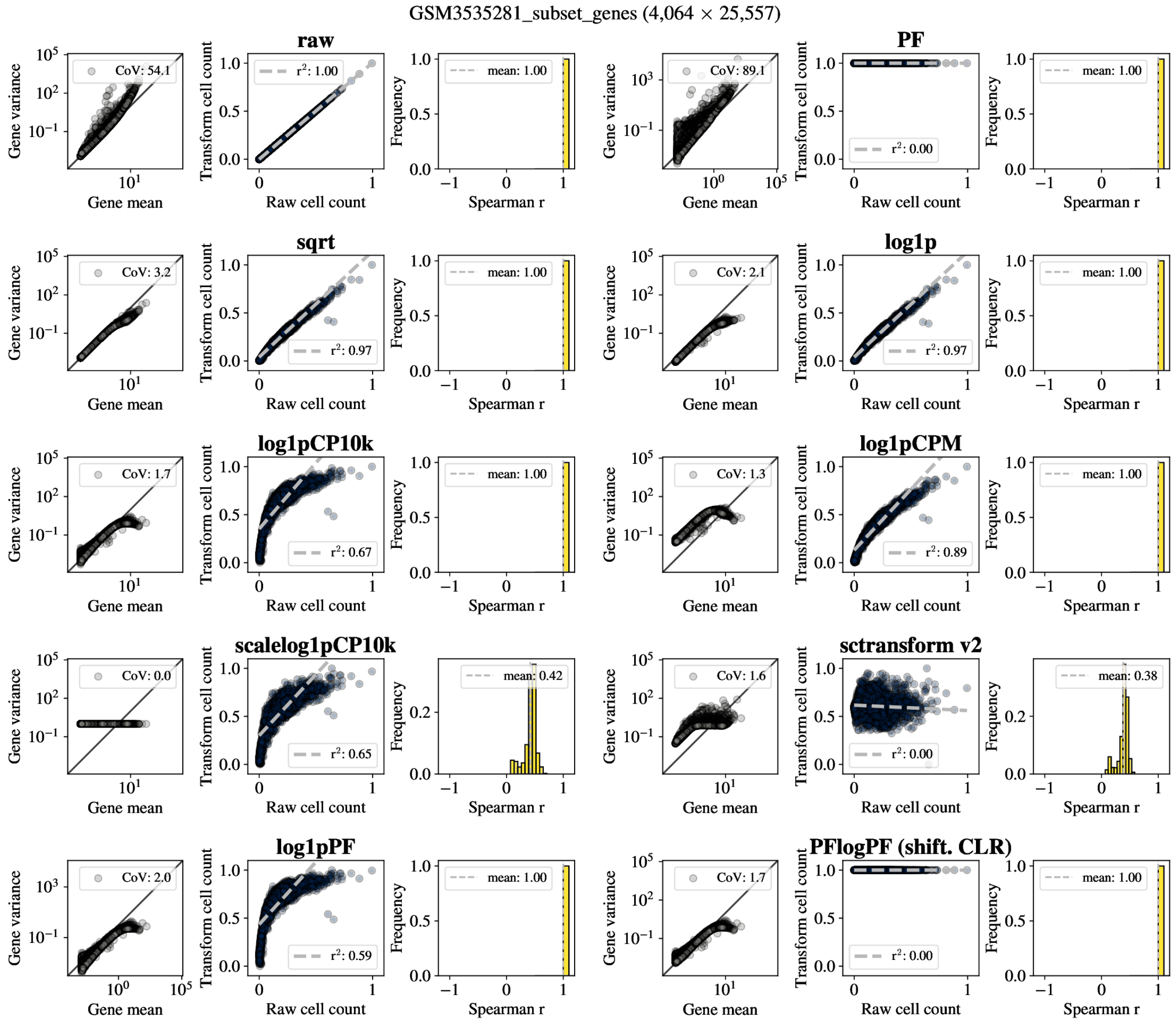


Supplementary Figure 1.285. **GSM3535281**. Per-method comparison of variance stabilization (left), depth normalization (middle), and monotonicity (right) for each normalization on dataset GSM3535281.

| GSM3557959 | |
|---|---|
| cells | 1,506 |
| genes | 60,623 |
| nonzero entries | 692,510 |
| density | 0.00759 |
| total counts | 1,969,885 |
| mean counts / cell | 1,308 |
| mean counts / gene | 32 |
| min counts / cell | 182 |
| max counts / cell | 16,941 |
| overdispersion | 0.959 |

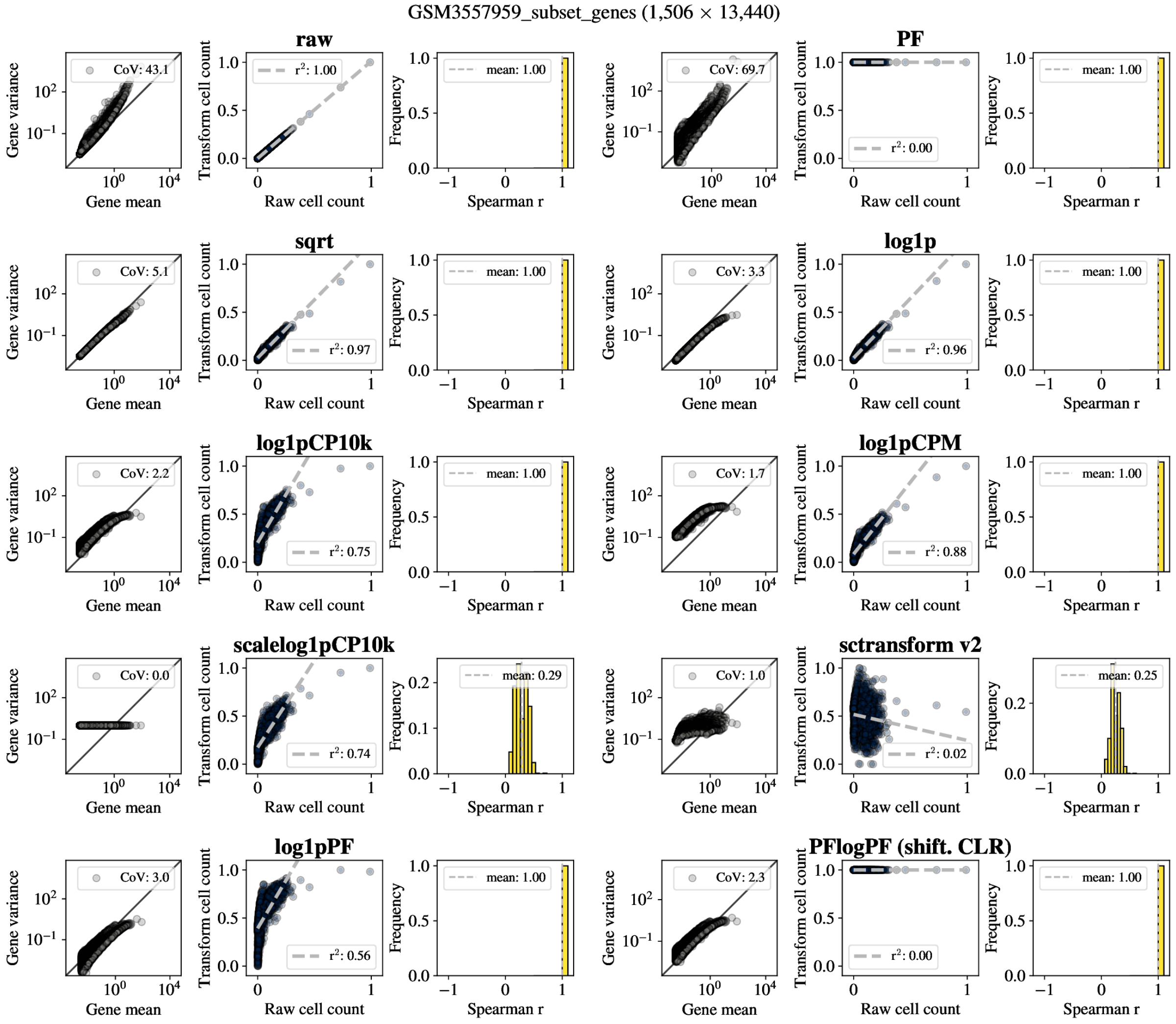


Supplementary Figure 1.286. **GSM3557959**. Per-method comparison of variance stabilization (left), depth normalization (middle), and monotonicity (right) for each normalization on dataset GSM3557959.

| GSM3557960 | |
|---|---|
| cells | 1,626 |
| genes | 60,623 |
| nonzero entries | 1,327,710 |
| density | 0.0135 |
| total counts | 3,628,822 |
| mean counts / cell | 2,232 |
| mean counts / gene | 60 |
| min counts / cell | 33 |
| max counts / cell | 16,159 |
| overdispersion | 2.36 |

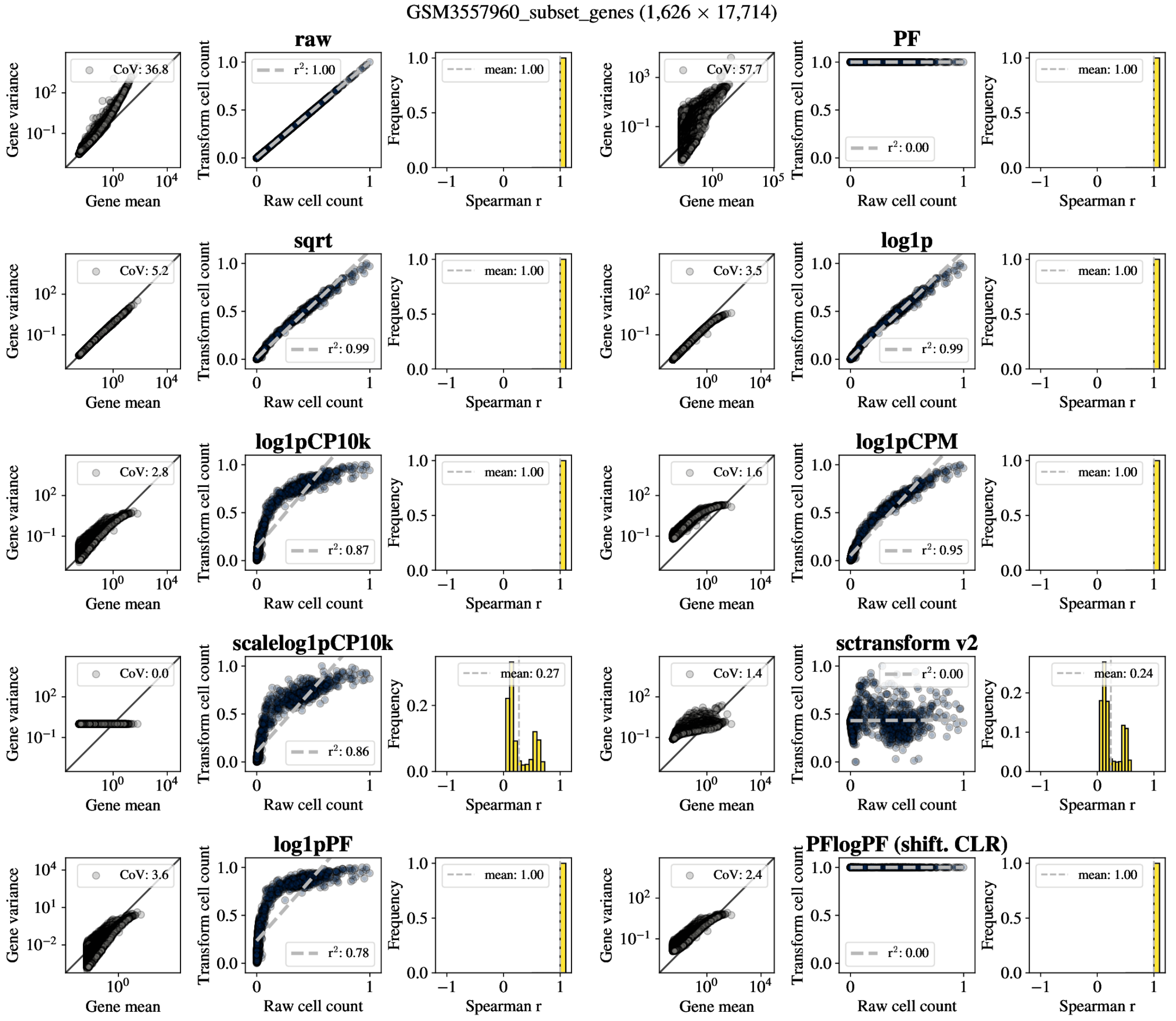


Supplementary Figure 1.287. **GSM3557960**. Per-method comparison of variance stabilization (left), depth normalization (middle), and monotonicity (right) for each normalization on dataset GSM3557960.

| GSM3557961 | |
|---|---|
| cells | 932 |
| genes | 60,623 |
| nonzero entries | 556,915 |
| density | 0.00986 |
| total counts | 1,232,498 |
| mean counts / cell | 1,322 |
| mean counts / gene | 20 |
| min counts / cell | 725 |
| max counts / cell | 5,213 |
| overdispersion | 0.311 |

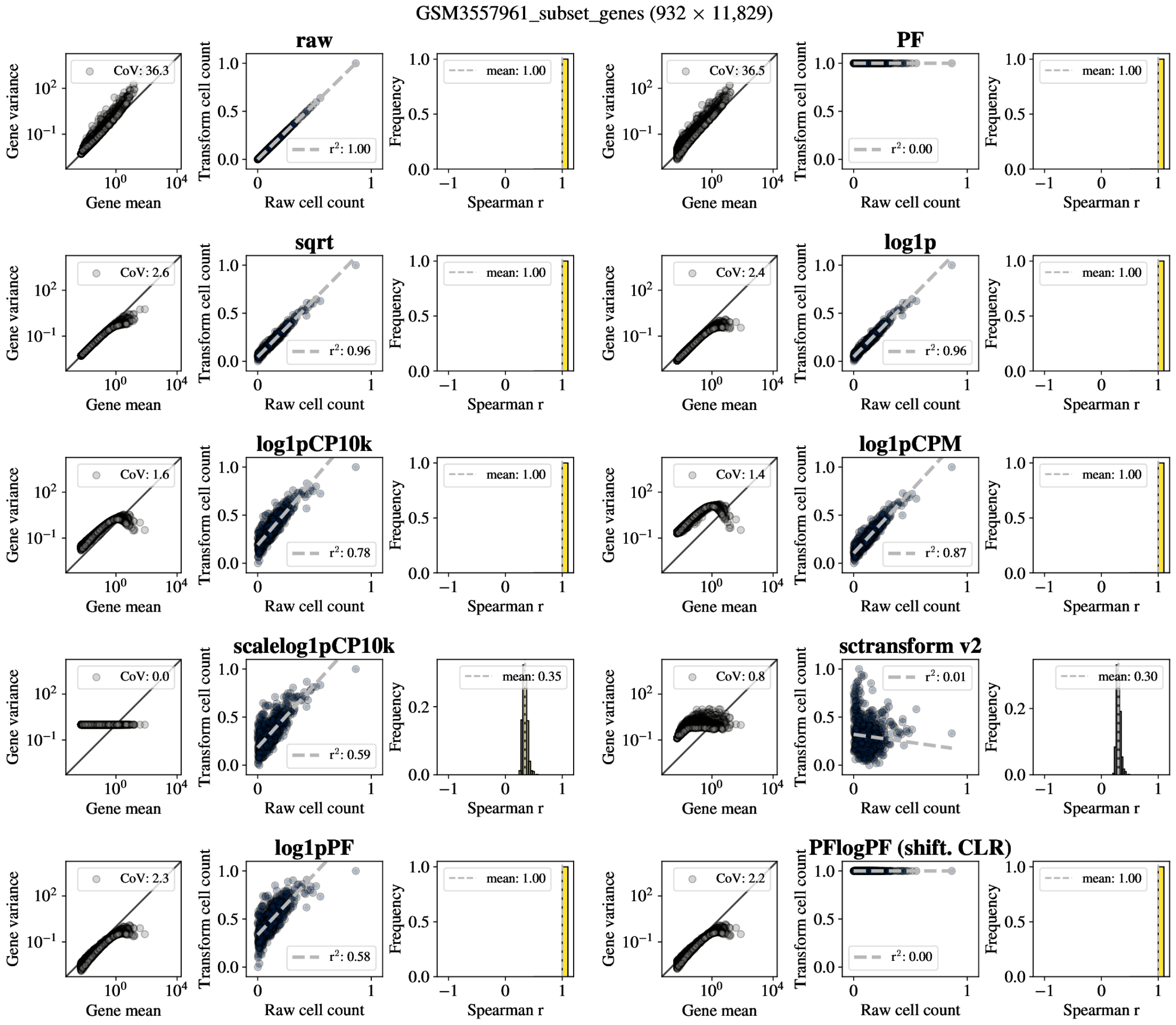


Supplementary Figure 1.288. **GSM3557961**. Per-method comparison of variance stabilization (left), depth normalization (middle), and monotonicity (right) for each normalization on dataset GSM3557961.

| GSM3557962 | |
|---|---|
| cells | 1,507 |
| genes | 60,623 |
| nonzero entries | 1,036,161 |
| density | 0.0113 |
| total counts | 2,373,910 |
| mean counts / cell | 1,575 |
| mean counts / gene | 39 |
| min counts / cell | 52 |
| max counts / cell | 16,644 |
| overdispersion | 0.733 |

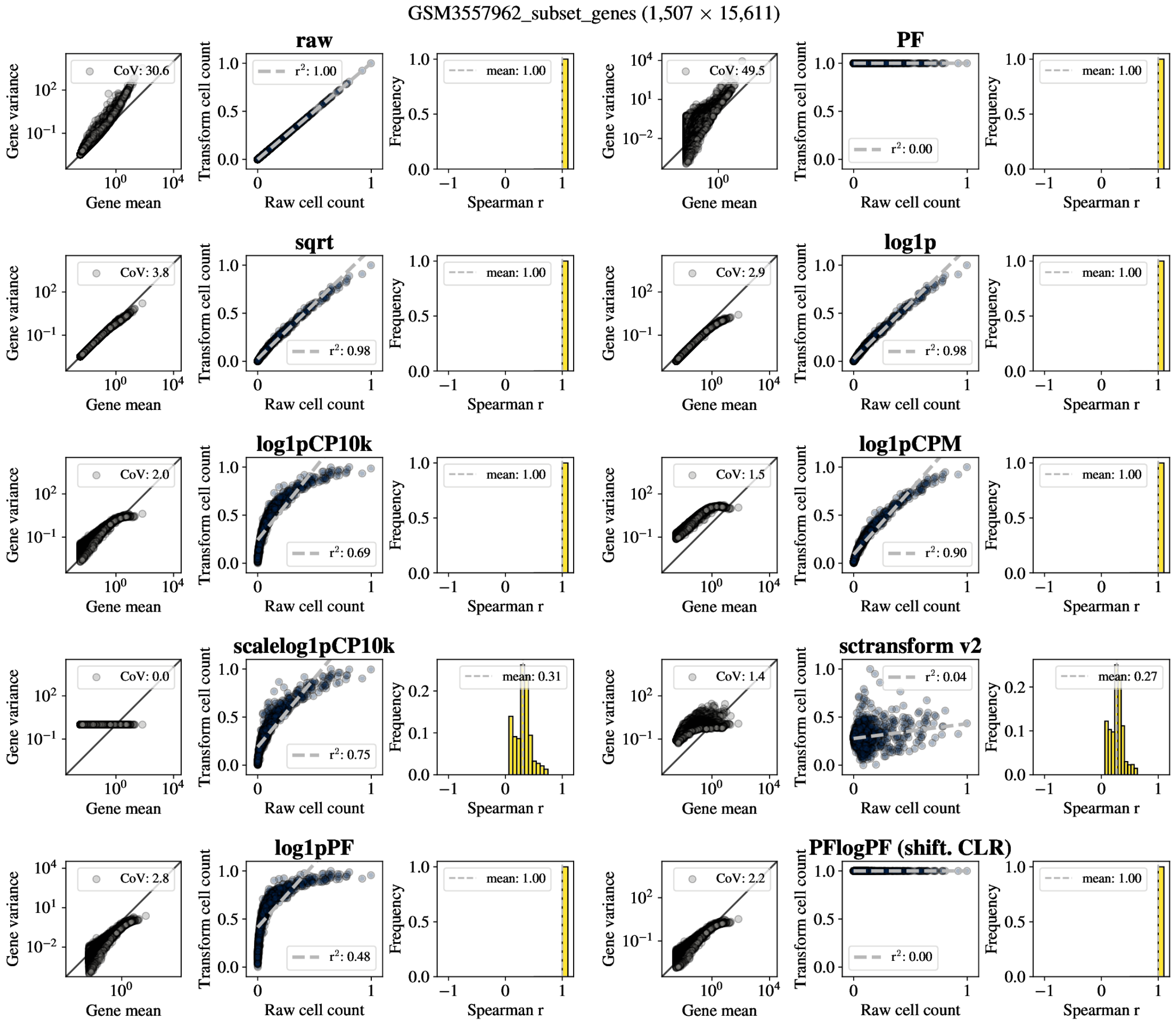


Supplementary Figure 1.289. **GSM3557962**. Per-method comparison of variance stabilization (left), depth normalization (middle), and monotonicity (right) for each normalization on dataset GSM3557962.

| GSM3557963 | |
|---|---|
| cells | 1,991 |
| genes | 60,623 |
| nonzero entries | 432,927 |
| density | 0.00359 |
| total counts | 706,672 |
| mean counts / cell | 355 |
| mean counts / gene | 12 |
| min counts / cell | 50 |
| max counts / cell | 3,605 |
| overdispersion | 2.92 |

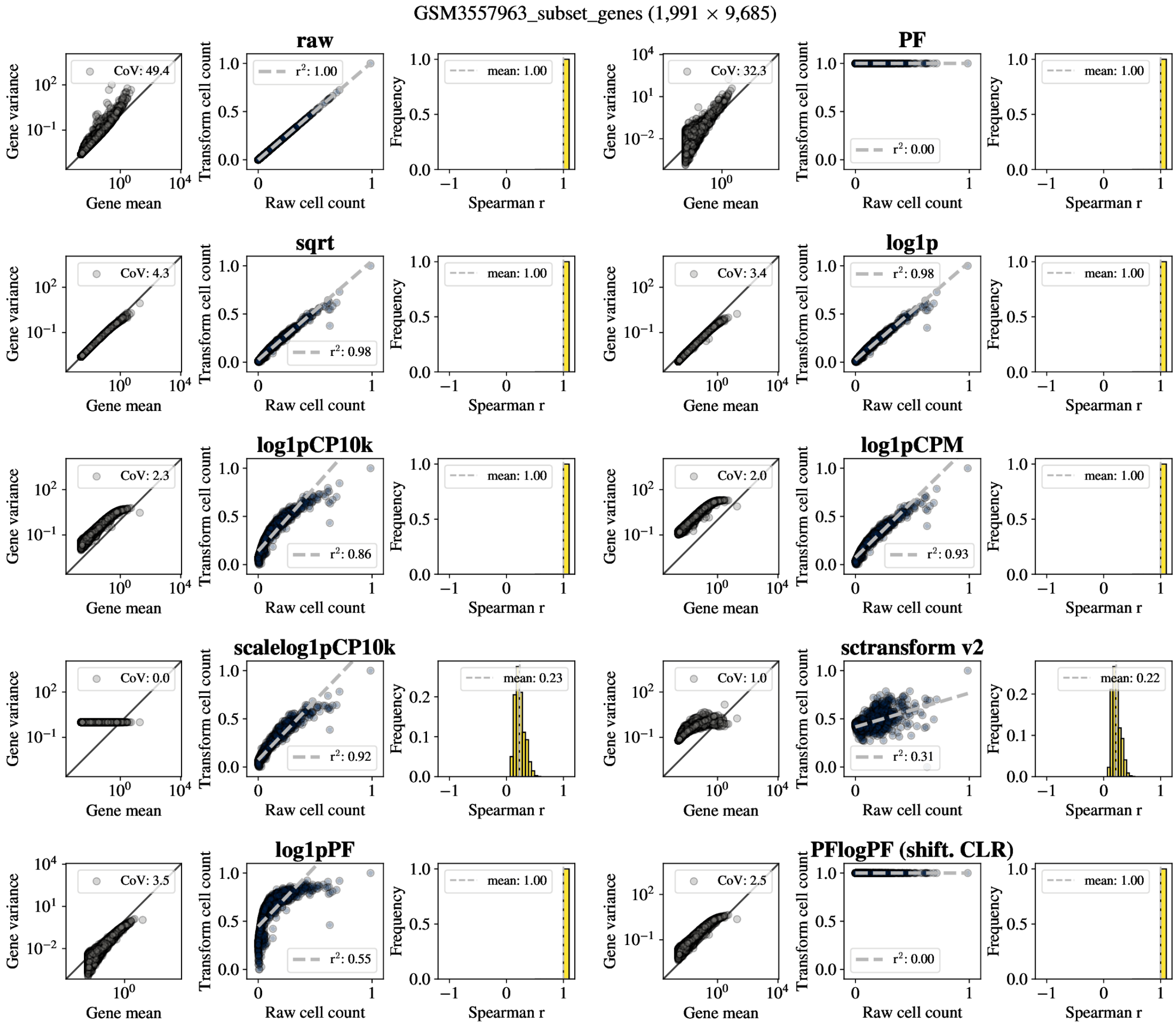


Supplementary Figure 1.290. **GSM3557963**. Per-method comparison of variance stabilization (left), depth normalization (middle), and monotonicity (right) for each normalization on dataset GSM3557963.

| GSM3557964 | |
|---|---|
| cells | 1,833 |
| genes | 60,623 |
| nonzero entries | 1,111,326 |
| density | 0.01 |
| total counts | 2,459,091 |
| mean counts / cell | 1,342 |
| mean counts / gene | 41 |
| min counts / cell | 125 |
| max counts / cell | 15,101 |
| overdispersion | 0.773 |

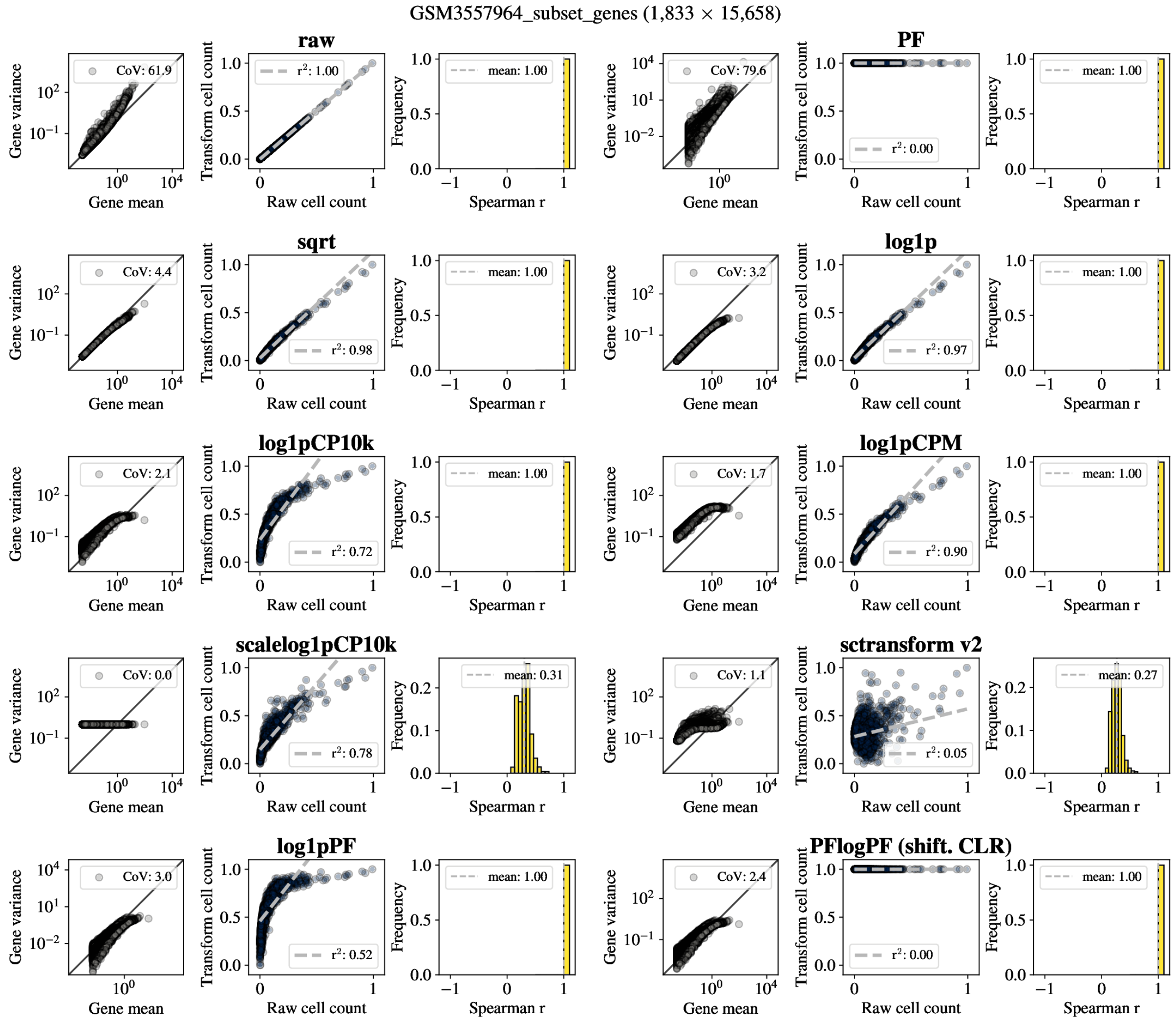


Supplementary Figure 1.291. **GSM3557964**. Per-method comparison of variance stabilization (left), depth normalization (middle), and monotonicity (right) for each normalization on dataset GSM3557964.

| GSM3557965 | |
|---|---|
| cells | 1,312 |
| genes | 60,623 |
| nonzero entries | 907,673 |
| density | 0.0114 |
| total counts | 2,339,215 |
| mean counts / cell | 1,783 |
| mean counts / gene | 39 |
| min counts / cell | 230 |
| max counts / cell | 14,743 |
| overdispersion | 0.681 |

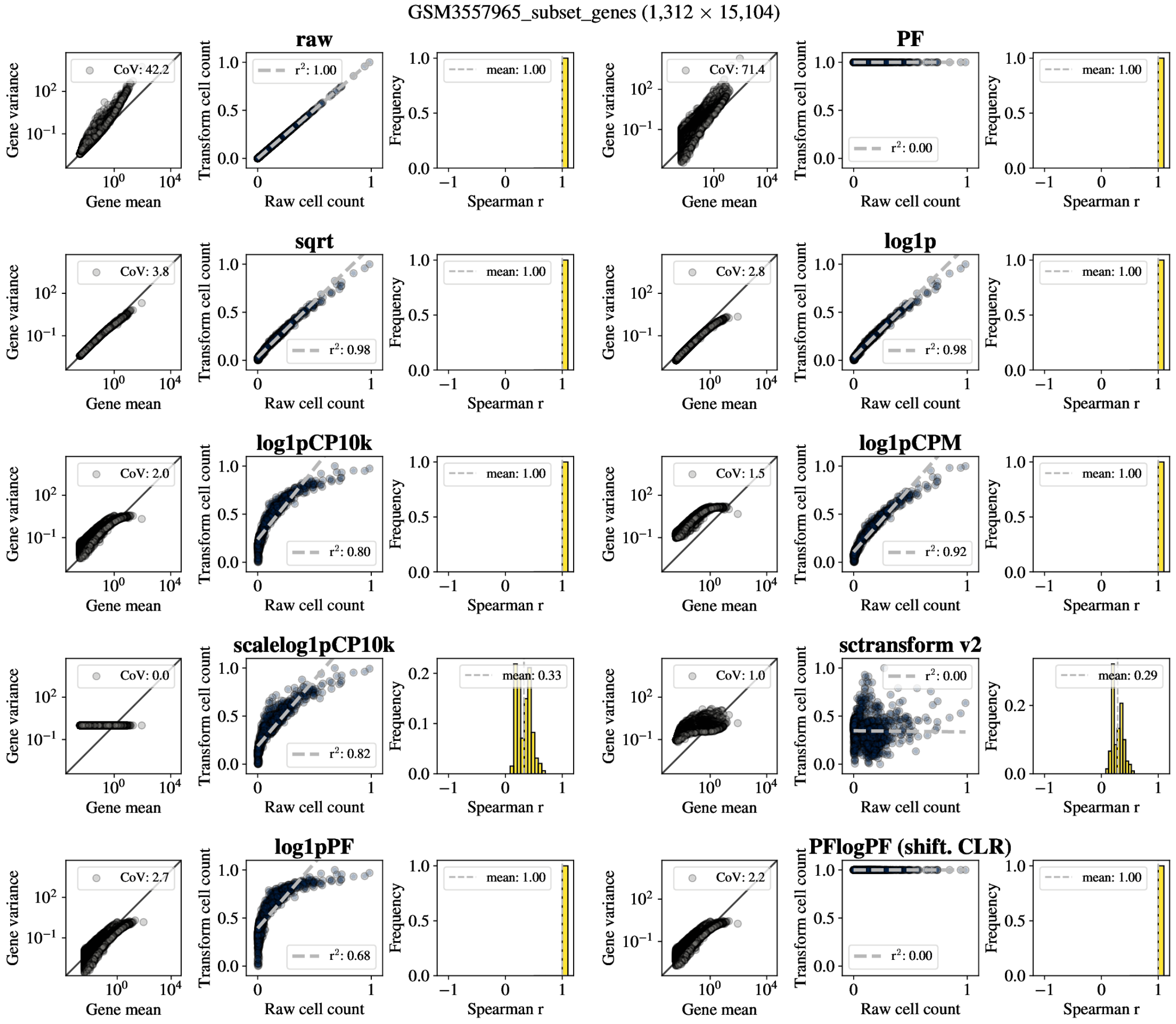


Supplementary Figure 1.292. **GSM3557965**. Per-method comparison of variance stabilization (left), depth normalization (middle), and monotonicity (right) for each normalization on dataset GSM3557965.

| GSM3557966 | |
|---|---|
| cells | 1,958 |
| genes | 60,623 |
| nonzero entries | 223,040 |
| density | 0.00188 |
| total counts | 392,834 |
| mean counts / cell | 201 |
| mean counts / gene | 6 |
| min counts / cell | 63 |
| max counts / cell | 7,191 |
| overdispersion | 10.5 |

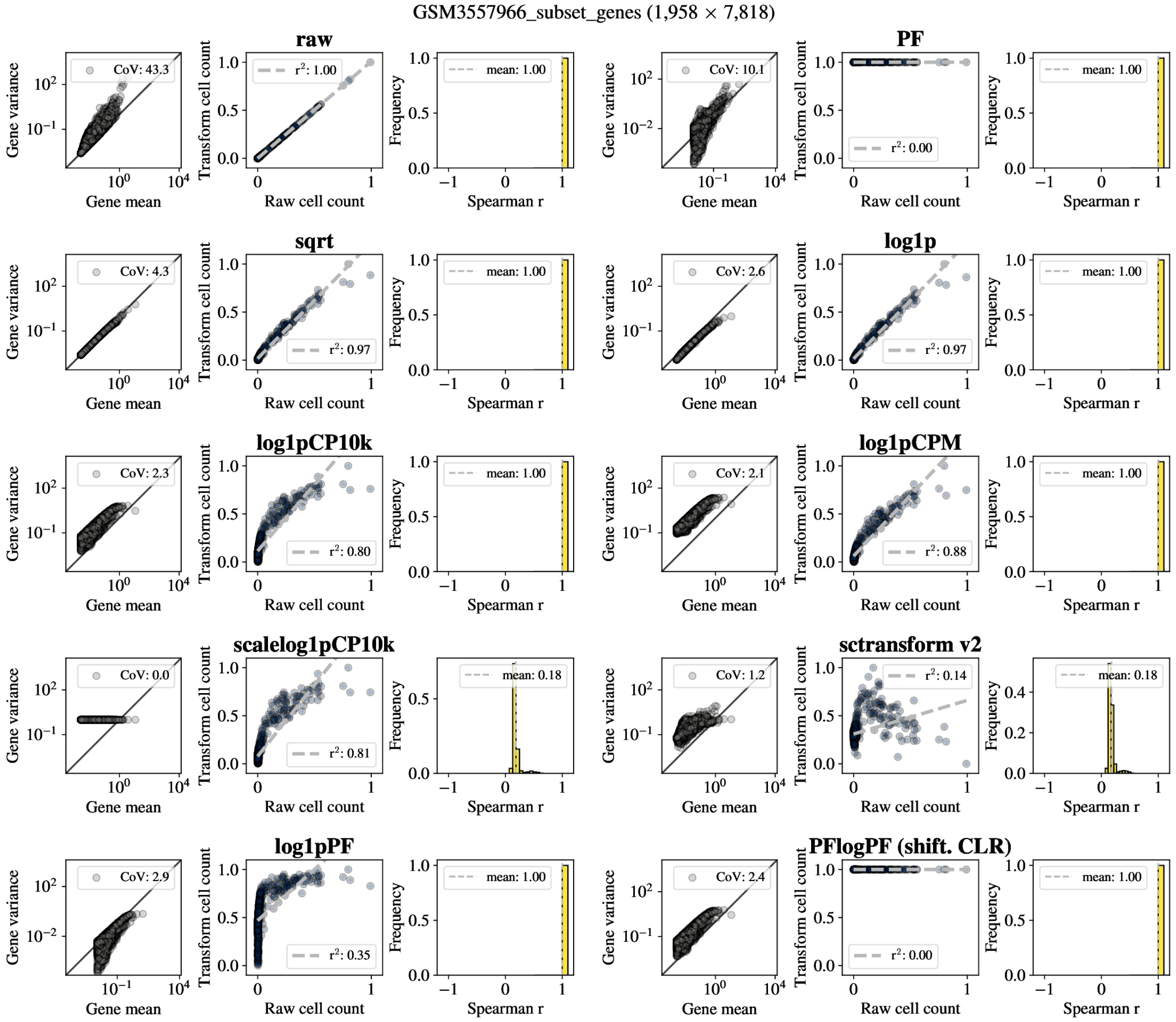


Supplementary Figure 1.293. **GSM3557966**. Per-method comparison of variance stabilization (left), depth normalization (middle), and monotonicity (right) for each normalization on dataset GSM3557966.

| GSM3557967 | |
|---|---|
| cells | 813 |
| genes | 60,623 |
| nonzero entries | 323,998 |
| density | 0.00657 |
| total counts | 738,135 |
| mean counts / cell | 908 |
| mean counts / gene | 12 |
| min counts / cell | 98 |
| max counts / cell | 20,207 |
| overdispersion | 2.01 |

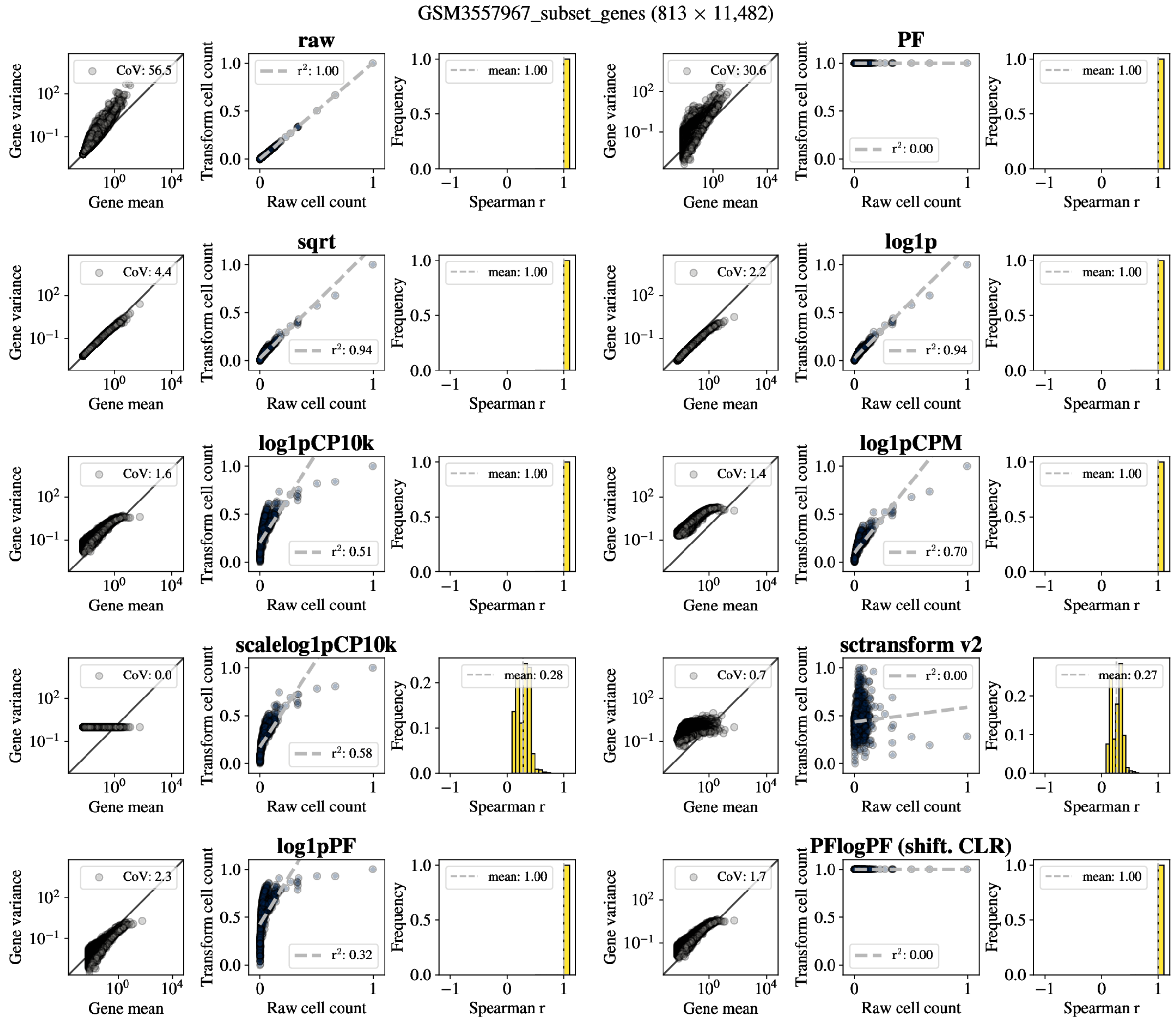


Supplementary Figure 1.294. **GSM3557967**. Per-method comparison of variance stabilization (left), depth normalization (middle), and monotonicity (right) for each normalization on dataset GSM3557967.

| GSM3557968 | |
|---|---|
| cells | 1,989 |
| genes | 60,623 |
| nonzero entries | 434,823 |
| density | 0.00361 |
| total counts | 800,639 |
| mean counts / cell | 403 |
| mean counts / gene | 13 |
| min counts / cell | 50 |
| max counts / cell | 4,823 |
| overdispersion | 3.09 |

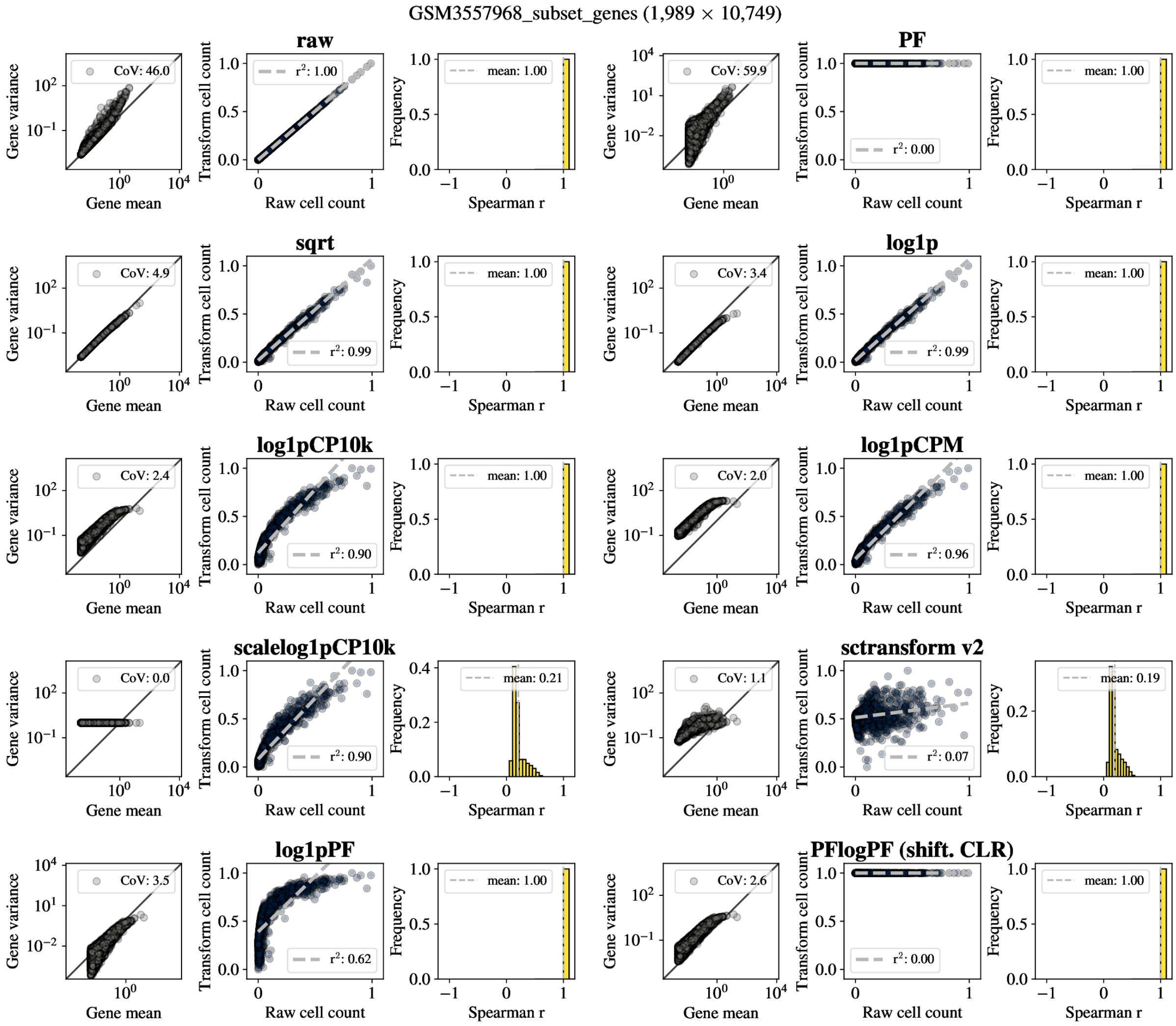


Supplementary Figure 1.295. **GSM3557968**. Per-method comparison of variance stabilization (left), depth normalization (middle), and monotonicity (right) for each normalization on dataset GSM3557968.

| GSM3557969 | |
|---|---|
| cells | 2,417 |
| genes | 60,623 |
| nonzero entries | 1,728,703 |
| density | 0.0118 |
| total counts | 3,682,996 |
| mean counts / cell | 1,524 |
| mean counts / gene | 61 |
| min counts / cell | 688 |
| max counts / cell | 8,161 |
| overdispersion | 0.182 |

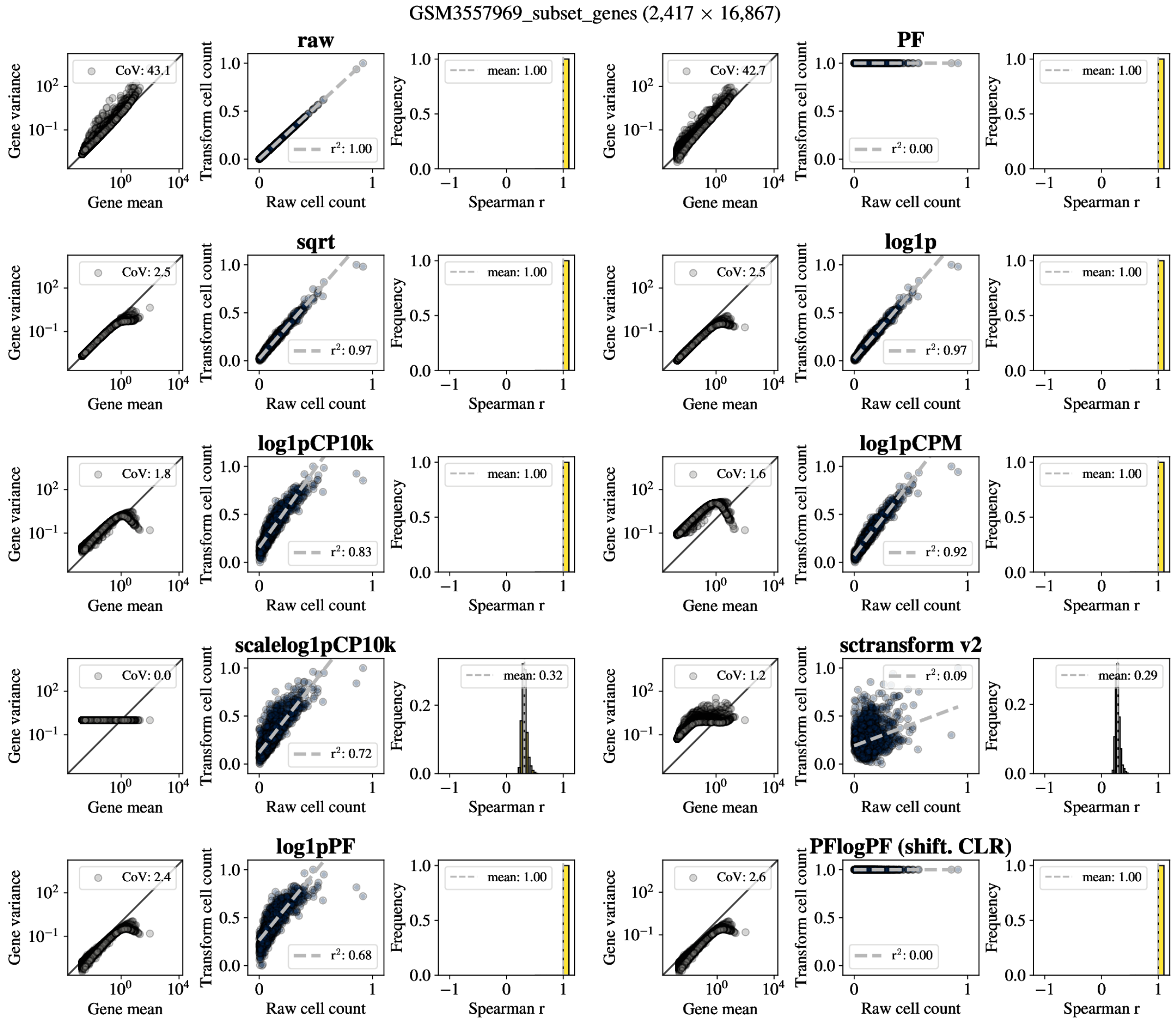


Supplementary Figure 1.296. **GSM3557969**. Per-method comparison of variance stabilization (left), depth normalization (middle), and monotonicity (right) for each normalization on dataset GSM3557969.

| GSM3557970 | |
|---|---|
| cells | 1,440 |
| genes | 60,623 |
| nonzero entries | 1,443,092 |
| density | 0.0165 |
| total counts | 3,412,473 |
| mean counts / cell | 2,370 |
| mean counts / gene | 56 |
| min counts / cell | 185 |
| max counts / cell | 12,908 |
| overdispersion | 0.886 |

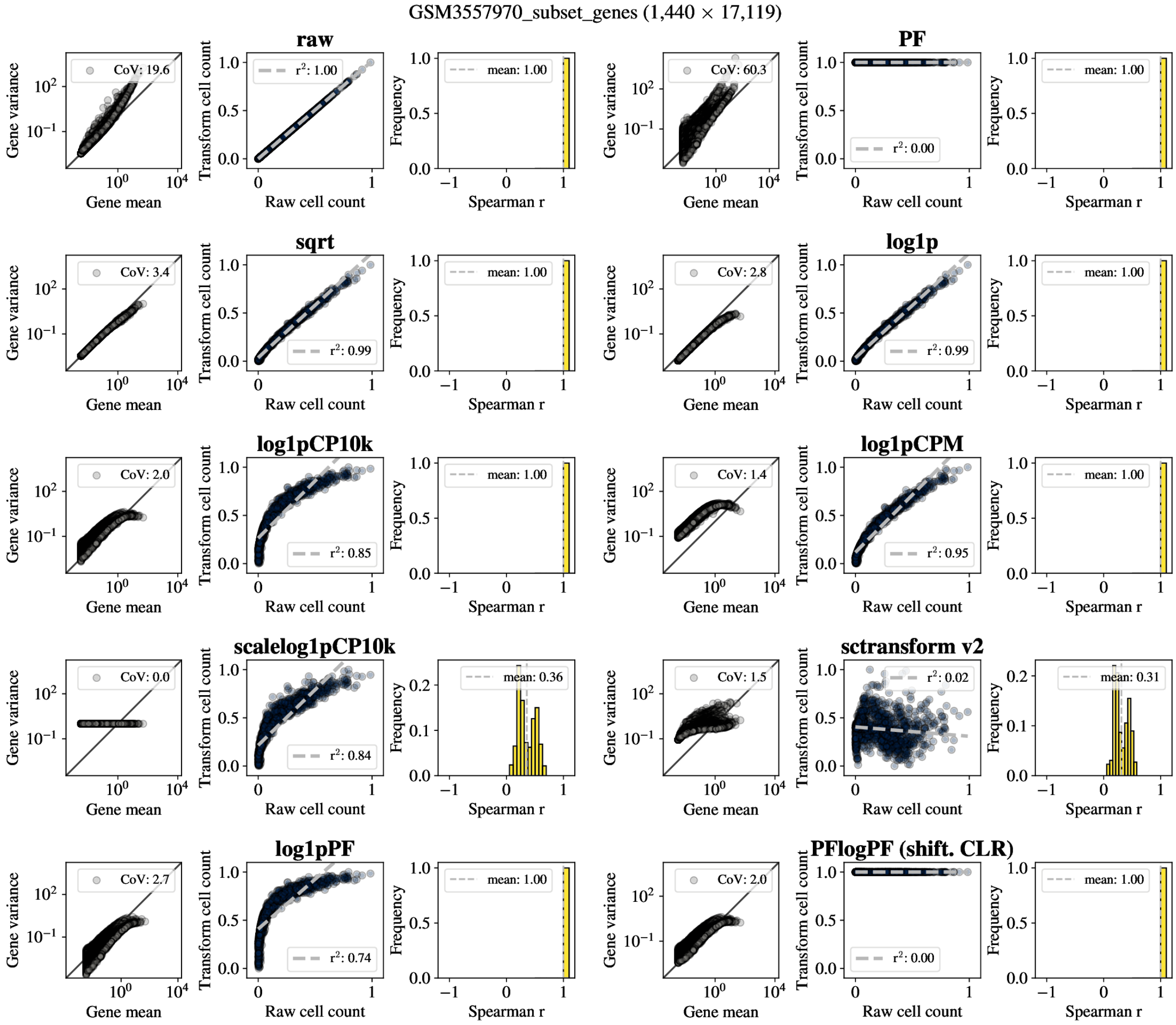


Supplementary Figure 1.297. **GSM3557970**. Per-method comparison of variance stabilization (left), depth normalization (middle), and monotonicity (right) for each normalization on dataset GSM3557970.

| GSM3557971 | |
|---|---|
| cells | 964 |
| genes | 60,623 |
| nonzero entries | 839,416 |
| density | 0.0144 |
| total counts | 1,955,745 |
| mean counts / cell | 2,029 |
| mean counts / gene | 32 |
| min counts / cell | 223 |
| max counts / cell | 11,502 |
| overdispersion | 0.798 |

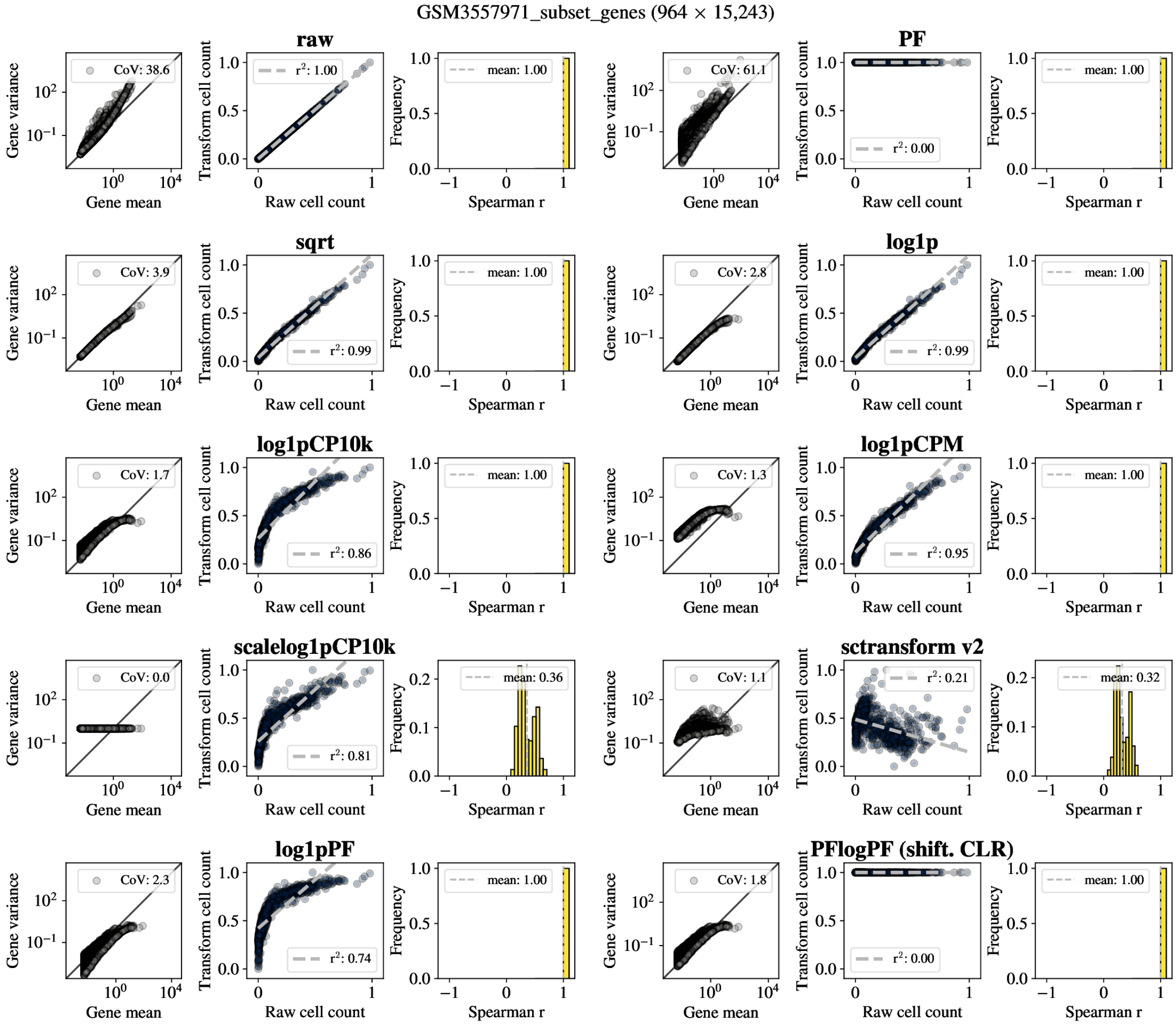


Supplementary Figure 1.298. **GSM3557971**. Per-method comparison of variance stabilization (left), depth normalization (middle), and monotonicity (right) for each normalization on dataset GSM3557971.

| GSM3557972 | |
|---|---|
| cells | 1,829 |
| genes | 60,623 |
| nonzero entries | 1,150,845 |
| density | 0.0104 |
| total counts | 2,385,133 |
| mean counts / cell | 1,304 |
| mean counts / gene | 39 |
| min counts / cell | 203 |
| max counts / cell | 21,862 |
| overdispersion | 0.775 |

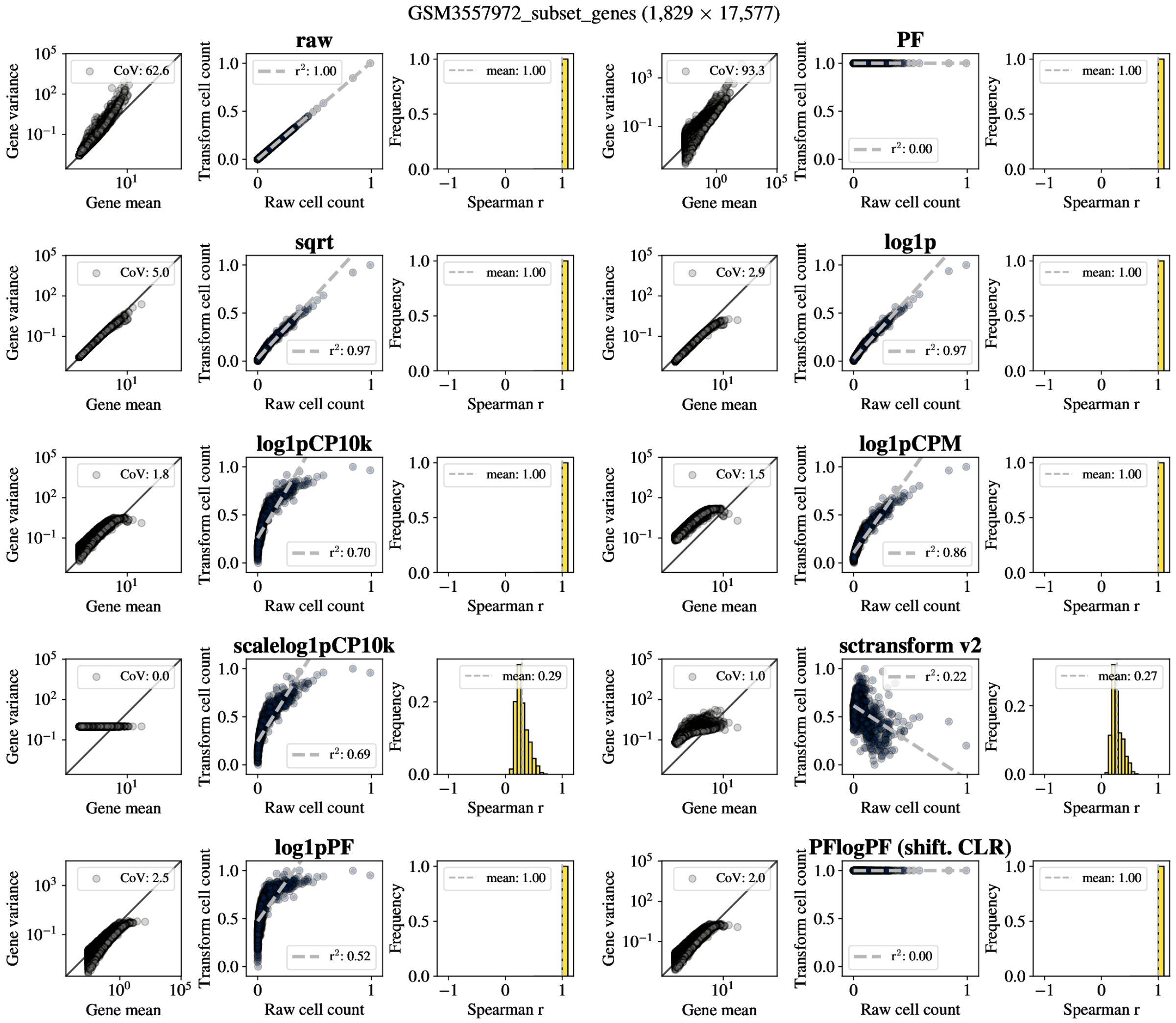


Supplementary Figure 1.299. **GSM3557972**. Per-method comparison of variance stabilization (left), depth normalization (middle), and monotonicity (right) for each normalization on dataset GSM3557972.

| GSM3557973 | |
|---|---|
| cells | 798 |
| genes | 60,623 |
| nonzero entries | 525,596 |
| density | 0.0109 |
| total counts | 1,564,269 |
| mean counts / cell | 1,960 |
| mean counts / gene | 26 |
| min counts / cell | 76 |
| max counts / cell | 16,827 |
| overdispersion | 2.56 |

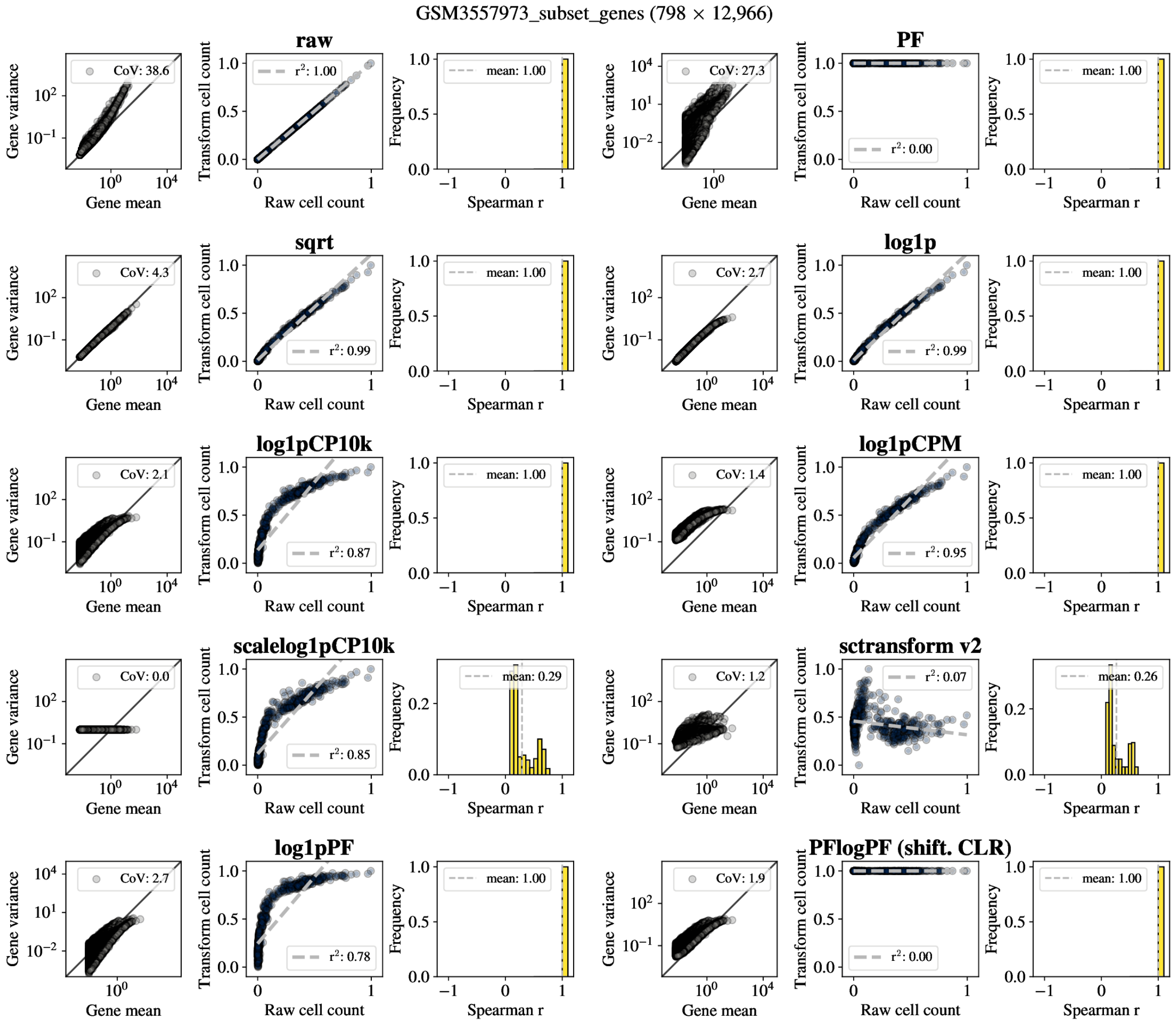


Supplementary Figure 1.300. **GSM3557973**. Per-method comparison of variance stabilization (left), depth normalization (middle), and monotonicity (right) for each normalization on dataset GSM3557973.

| GSM3558027 | |
|---|---|
| cells | 3,599 |
| genes | 60,623 |
| nonzero entries | 4,698,317 |
| density | 0.0215 |
| total counts | 17,382,771 |
| mean counts / cell | 4,830 |
| mean counts / gene | 287 |
| min counts / cell | 2,466 |
| max counts / cell | 27,529 |
| overdispersion | 0.771 |

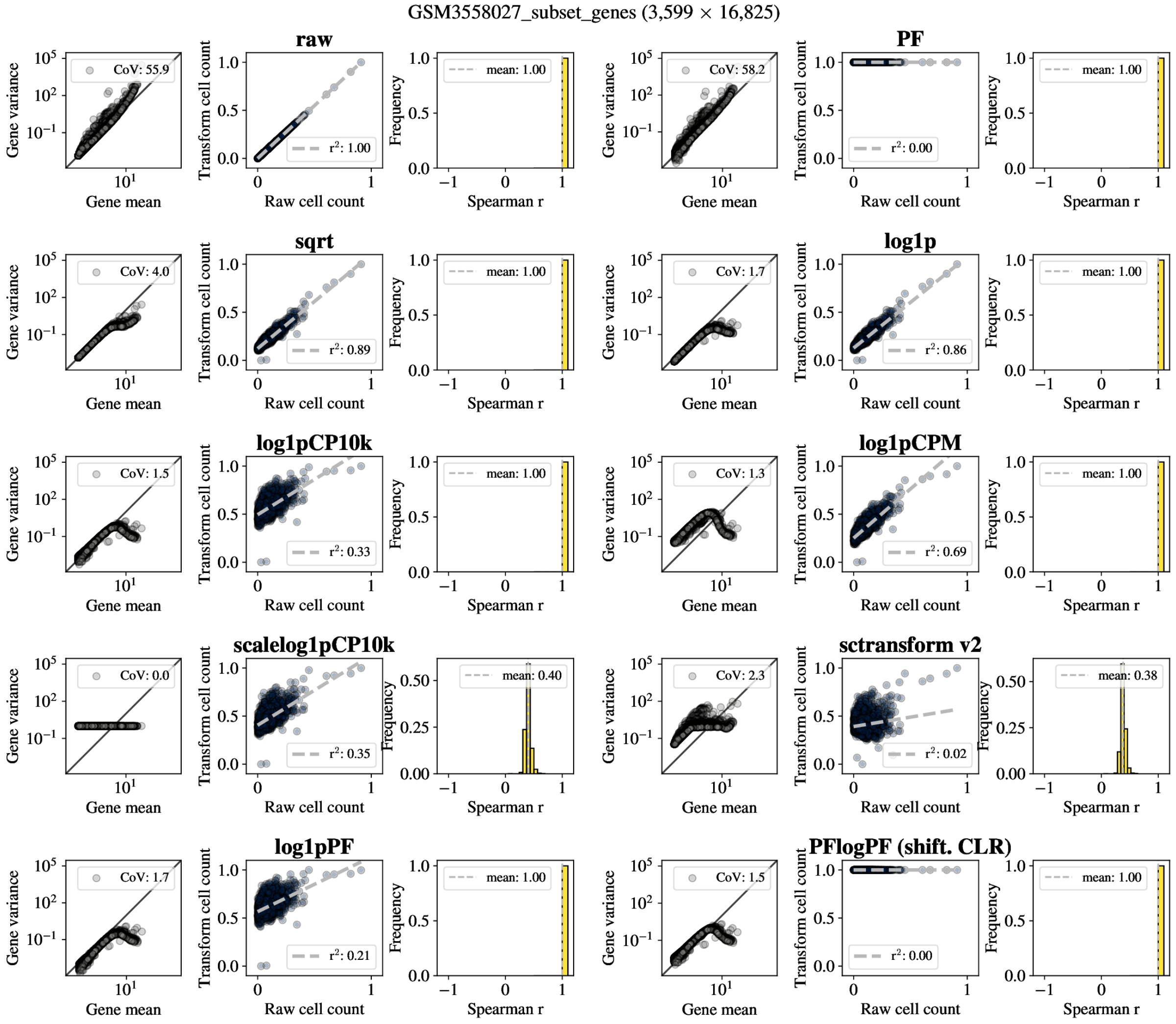


Supplementary Figure 1.301. **GSM3558027**. Per-method comparison of variance stabilization (left), depth normalization (middle), and monotonicity (right) for each normalization on dataset GSM3558027.

| GSM3576397 | |
|---|---|
| cells | 1,348 |
| genes | 60,623 |
| nonzero entries | 238,149 |
| density | 0.00291 |
| total counts | 572,860 |
| mean counts / cell | 425 |
| mean counts / gene | 9 |
| min counts / cell | 2 |
| max counts / cell | 2,548 |
| overdispersion | 2.66 |

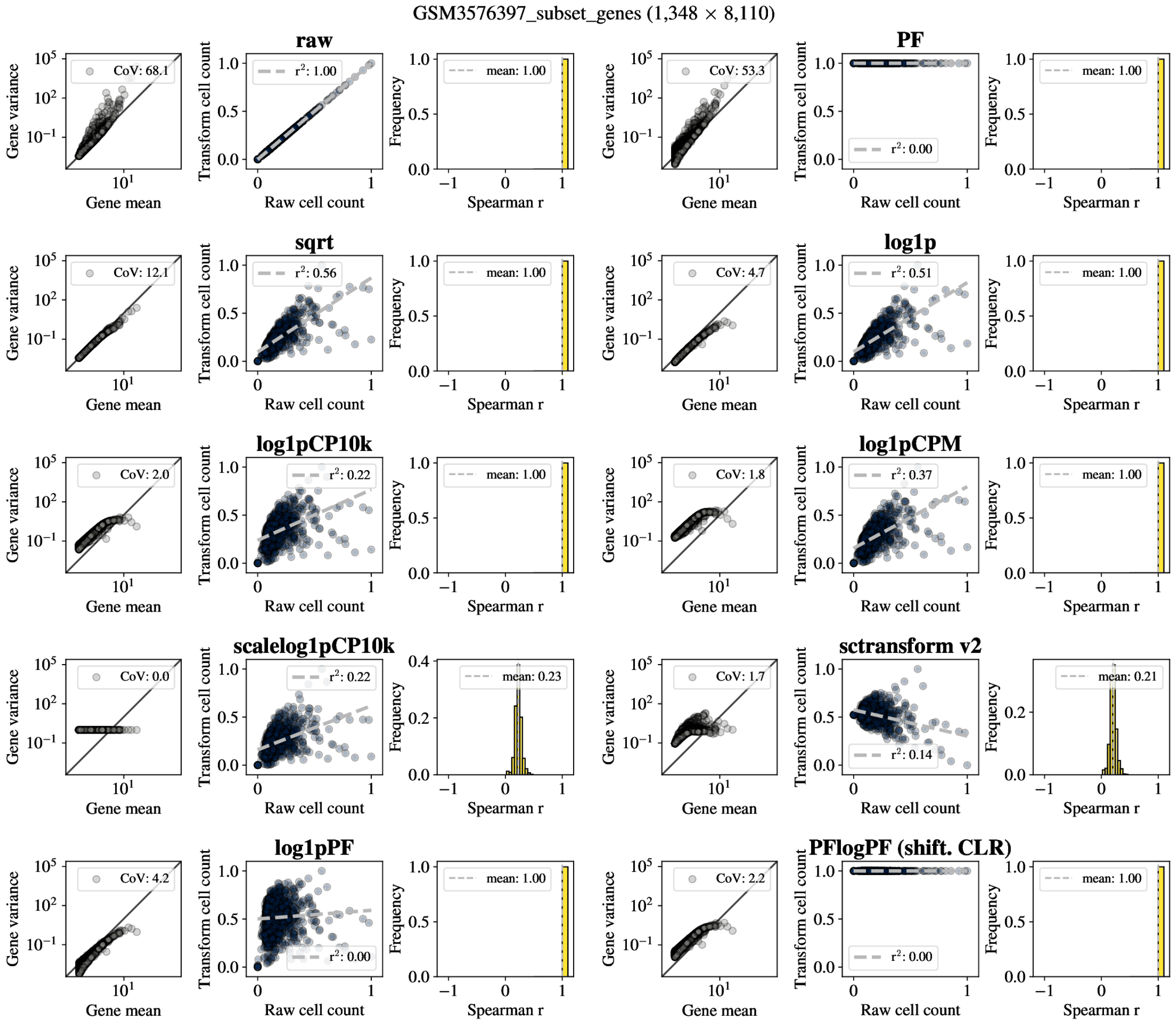


Supplementary Figure 1.302. **GSM3576397**. Per-method comparison of variance stabilization (left), depth normalization (middle), and monotonicity (right) for each normalization on dataset GSM3576397.

| GSM3576398 | |
|---|---|
| cells | 10,650 |
| genes | 60,623 |
| nonzero entries | 795,275 |
| density | 0.00123 |
| total counts | 1,844,999 |
| mean counts / cell | 173 |
| mean counts / gene | 30 |
| min counts / cell | 0 |
| max counts / cell | 10,839 |
| overdispersion | 3.3 |

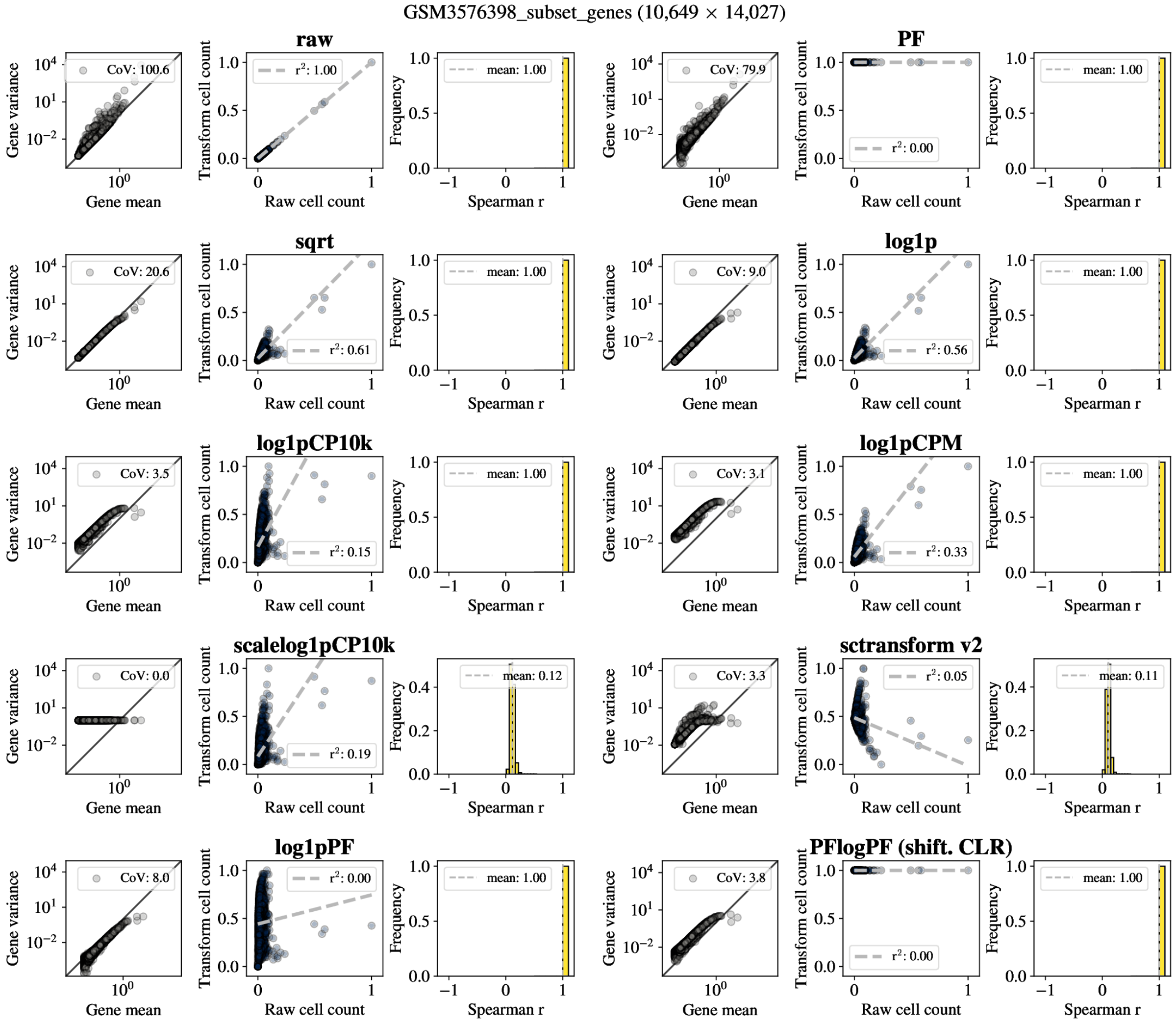


Supplementary Figure 1.303. **GSM3576398**. Per-method comparison of variance stabilization (left), depth normalization (middle), and monotonicity (right) for each normalization on dataset GSM3576398.

| GSM3576399 | |
|---|---|
| cells | 3,844 |
| genes | 60,623 |
| nonzero entries | 540,275 |
| density | 0.00232 |
| total counts | 1,151,372 |
| mean counts / cell | 300 |
| mean counts / gene | 19 |
| min counts / cell | 0 |
| max counts / cell | 3,340 |
| overdispersion | 3.21 |

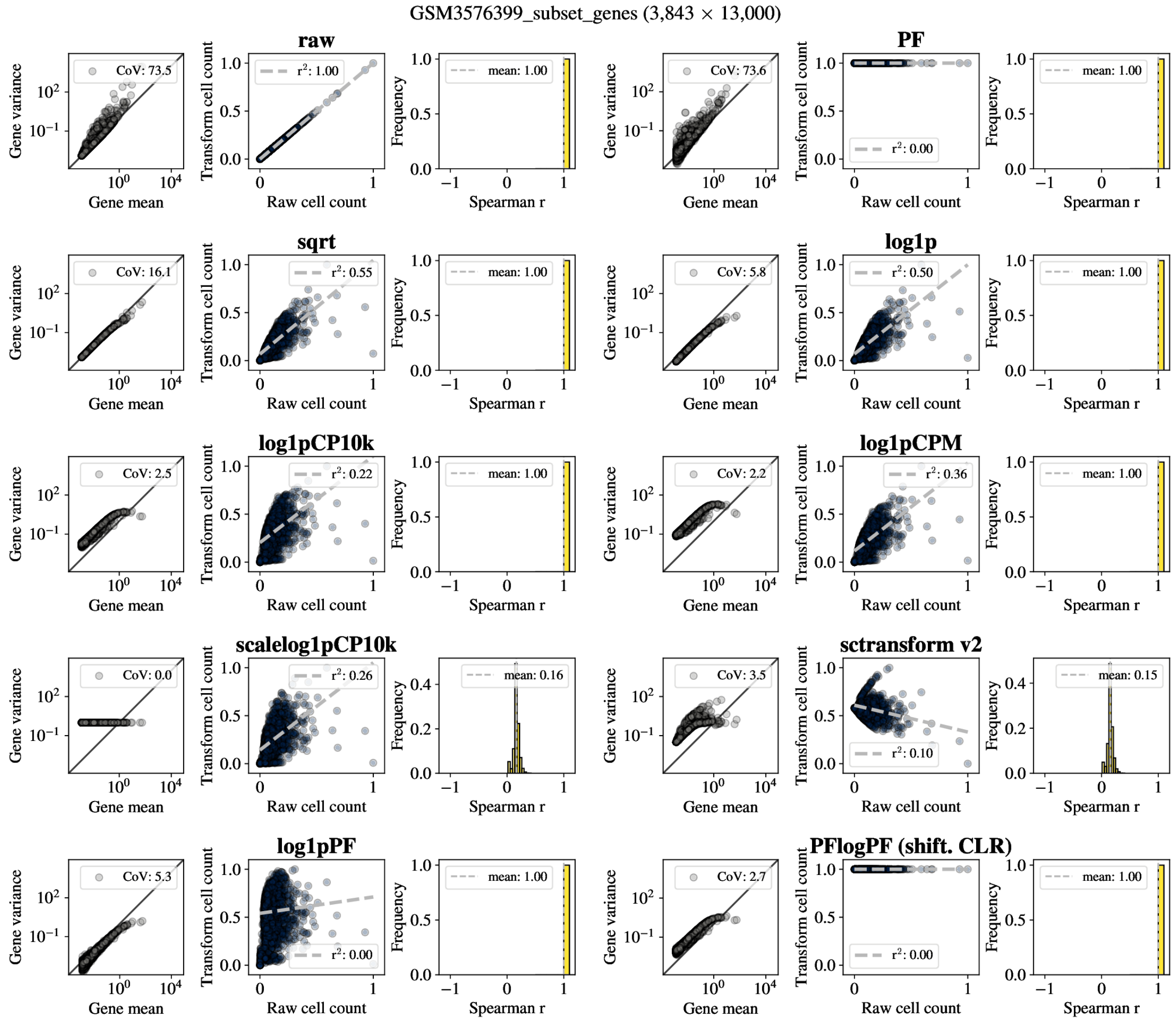


Supplementary Figure 1.304. **GSM3576399**. Per-method comparison of variance stabilization (left), depth normalization (middle), and monotonicity (right) for each normalization on dataset GSM3576399.

| GSM3576400 | |
|---|---|
| cells | 11,916 |
| genes | 60,623 |
| nonzero entries | 842,384 |
| density | 0.00117 |
| total counts | 2,274,454 |
| mean counts / cell | 191 |
| mean counts / gene | 38 |
| min counts / cell | 1 |
| max counts / cell | 6,427 |
| overdispersion | 1.5 |

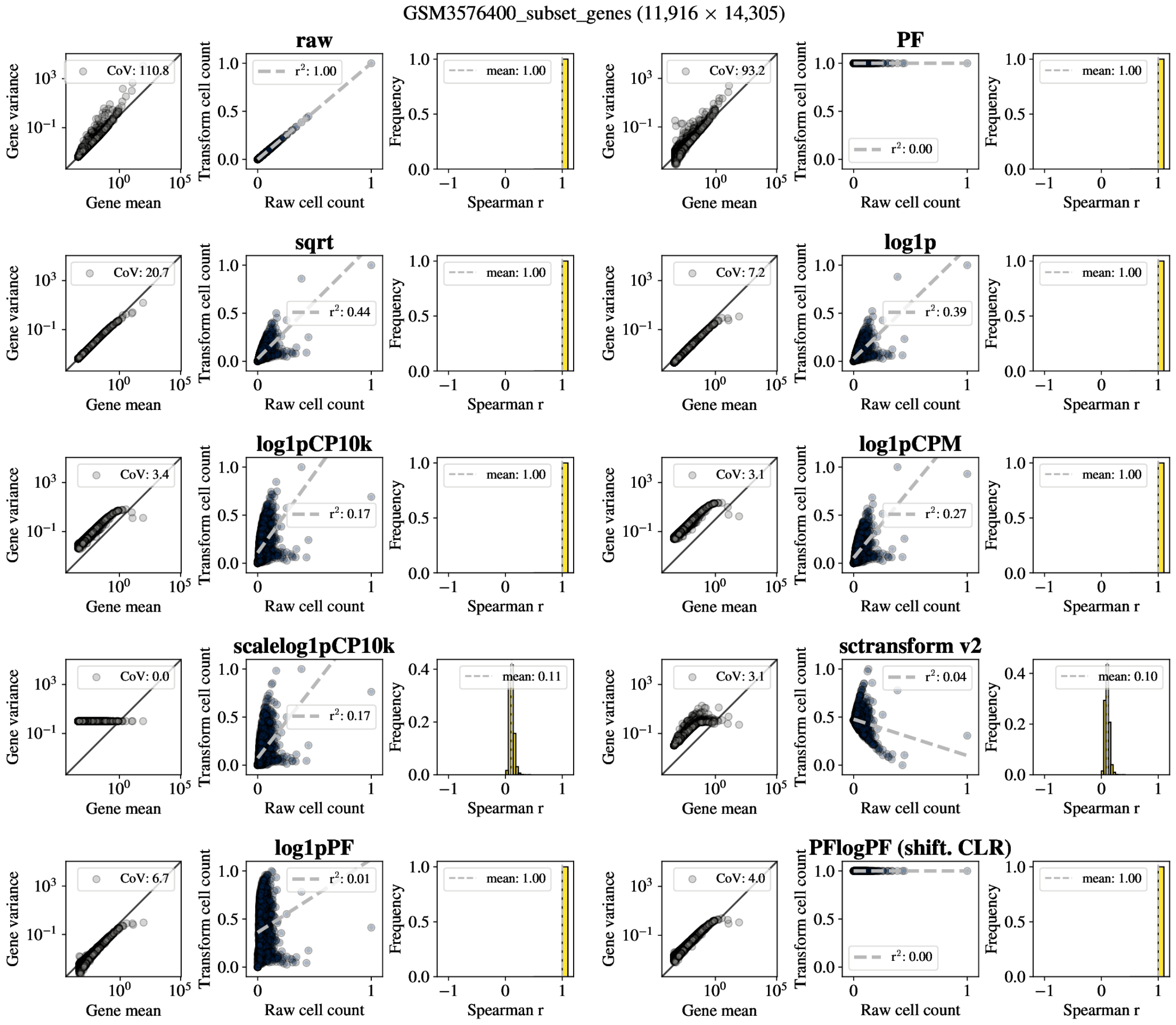


Supplementary Figure 1.305. **GSM3576400**. Per-method comparison of variance stabilization (left), depth normalization (middle), and monotonicity (right) for each normalization on dataset GSM3576400.

| GSM3576401 | |
|---|---|
| cells | 3,379 |
| genes | 60,623 |
| nonzero entries | 528,508 |
| density | 0.00258 |
| total counts | 1,201,013 |
| mean counts / cell | 355 |
| mean counts / gene | 20 |
| min counts / cell | 1 |
| max counts / cell | 6,127 |
| overdispersion | 3.47 |

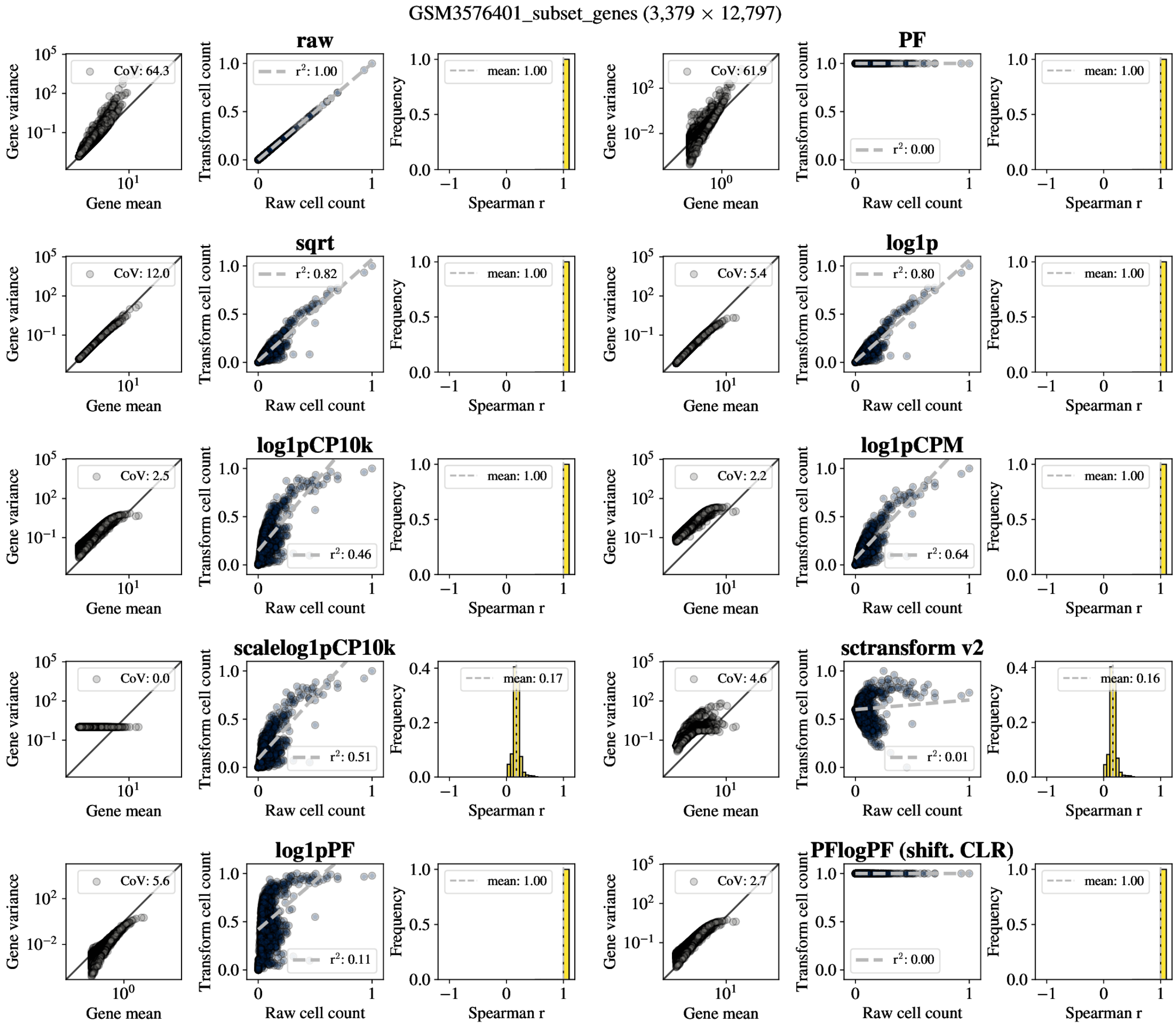


Supplementary Figure 1.306. **GSM3576401**. Per-method comparison of variance stabilization (left), depth normalization (middle), and monotonicity (right) for each normalization on dataset GSM3576401.

| GSM3576402 | |
|---|---|
| cells | 3,245 |
| genes | 60,623 |
| nonzero entries | 465,098 |
| density | 0.00236 |
| total counts | 844,791 |
| mean counts / cell | 260 |
| mean counts / gene | 14 |
| min counts / cell | 2 |
| max counts / cell | 1,793 |
| overdispersion | 2.94 |

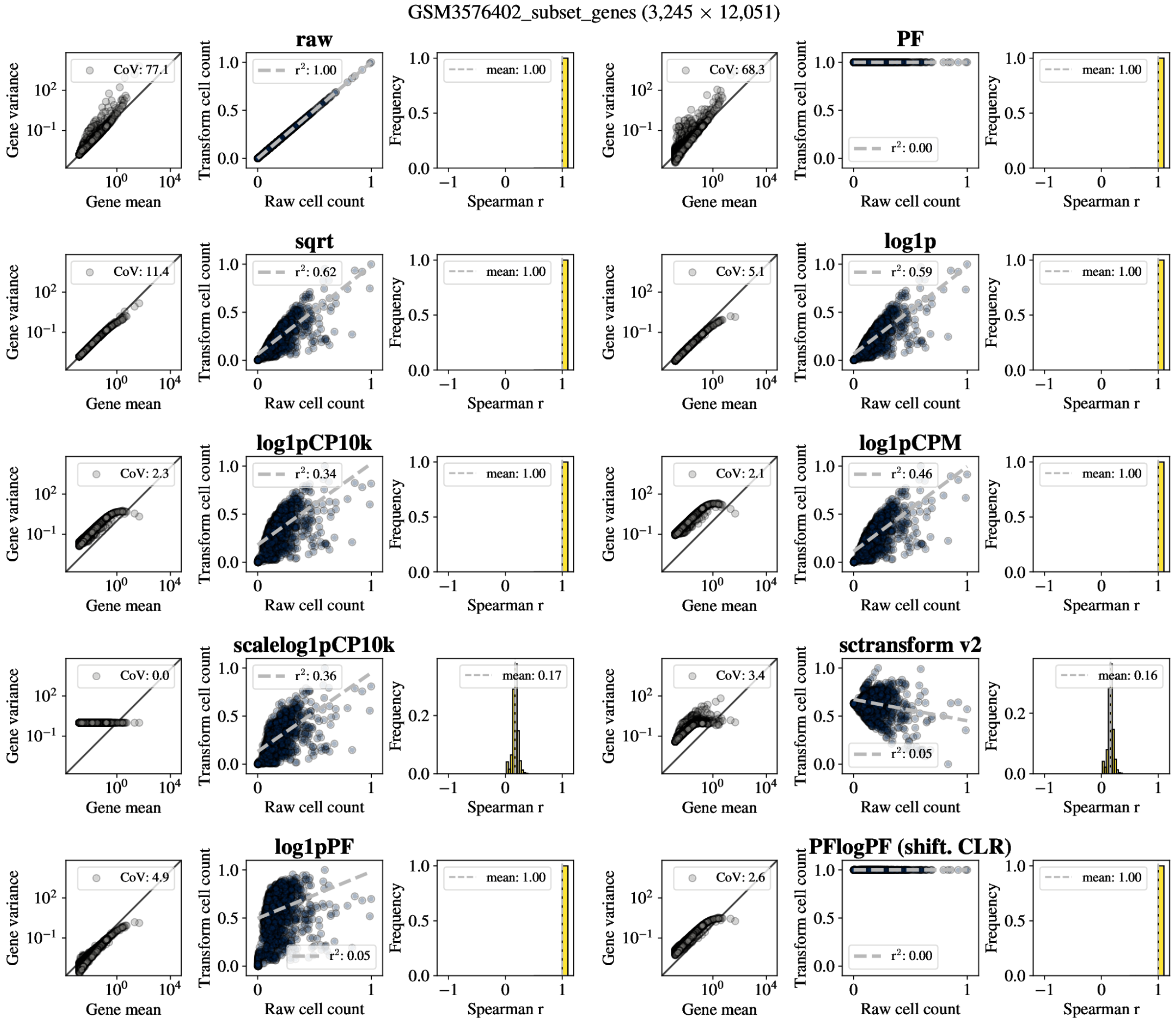


Supplementary Figure 1.307. **GSM3576402**. Per-method comparison of variance stabilization (left), depth normalization (middle), and monotonicity (right) for each normalization on dataset GSM3576402.

| GSM3576403 | |
|---|---|
| cells | 7,081 |
| genes | 60,623 |
| nonzero entries | 937,467 |
| density | 0.00218 |
| total counts | 1,576,767 |
| mean counts / cell | 223 |
| mean counts / gene | 26 |
| min counts / cell | 0 |
| max counts / cell | 2,244 |
| overdispersion | 4.77 |

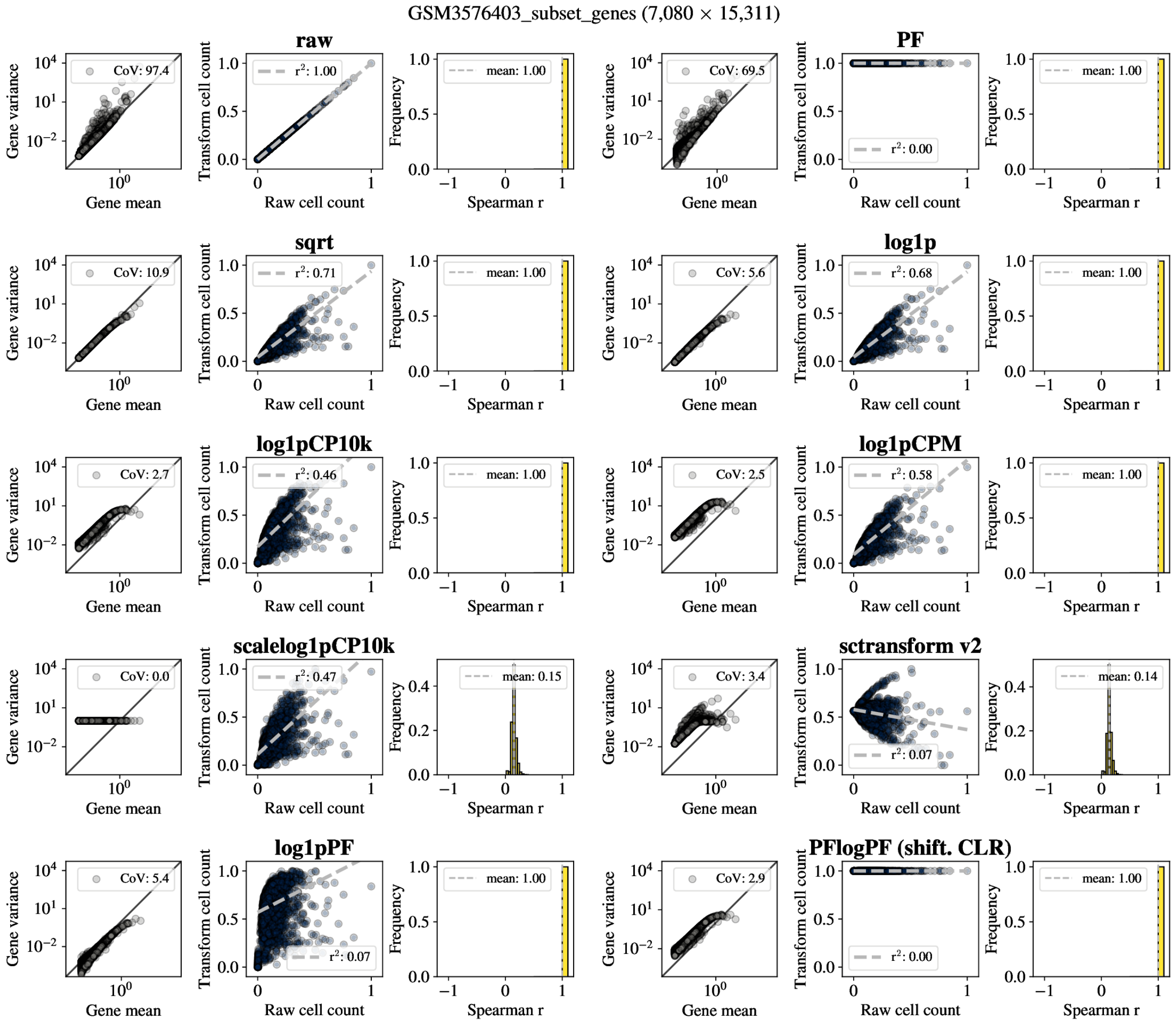


Supplementary Figure 1.308. **GSM3576403**. Per-method comparison of variance stabilization (left), depth normalization (middle), and monotonicity (right) for each normalization on dataset GSM3576403.

| GSM3576404 | |
|---|---|
| cells | 3,751 |
| genes | 60,623 |
| nonzero entries | 384,480 |
| density | 0.00169 |
| total counts | 727,225 |
| mean counts / cell | 194 |
| mean counts / gene | 12 |
| min counts / cell | 1 |
| max counts / cell | 1,665 |
| overdispersion | 3.53 |

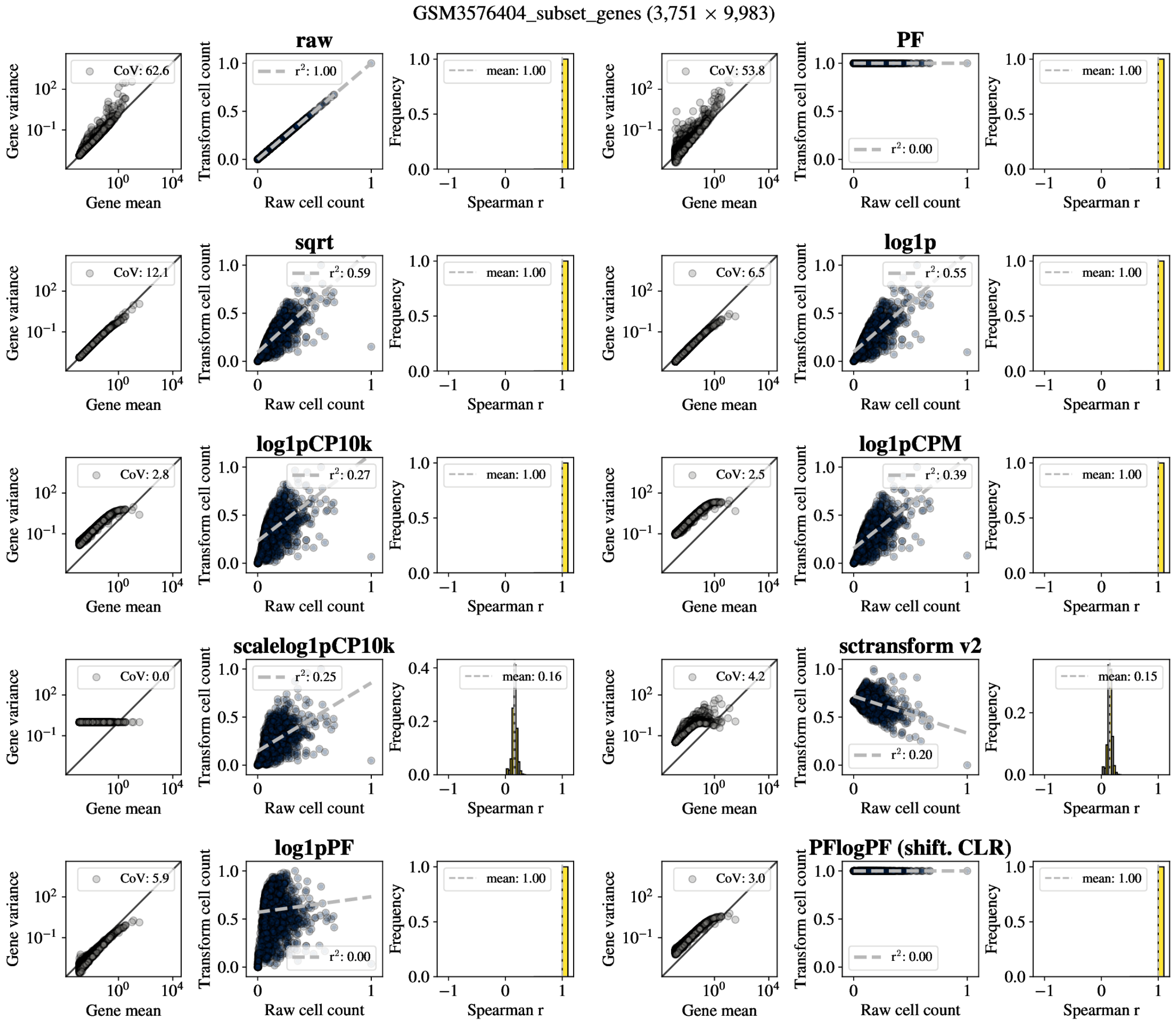


Supplementary Figure 1.309. **GSM3576404**. Per-method comparison of variance stabilization (left), depth normalization (middle), and monotonicity (right) for each normalization on dataset GSM3576404.

| GSM3576405 | |
|---|---|
| cells | 5,385 |
| genes | 60,623 |
| nonzero entries | 597,844 |
| density | 0.00183 |
| total counts | 1,078,216 |
| mean counts / cell | 200 |
| mean counts / gene | 18 |
| min counts / cell | 1 |
| max counts / cell | 2,059 |
| overdispersion | 3.75 |

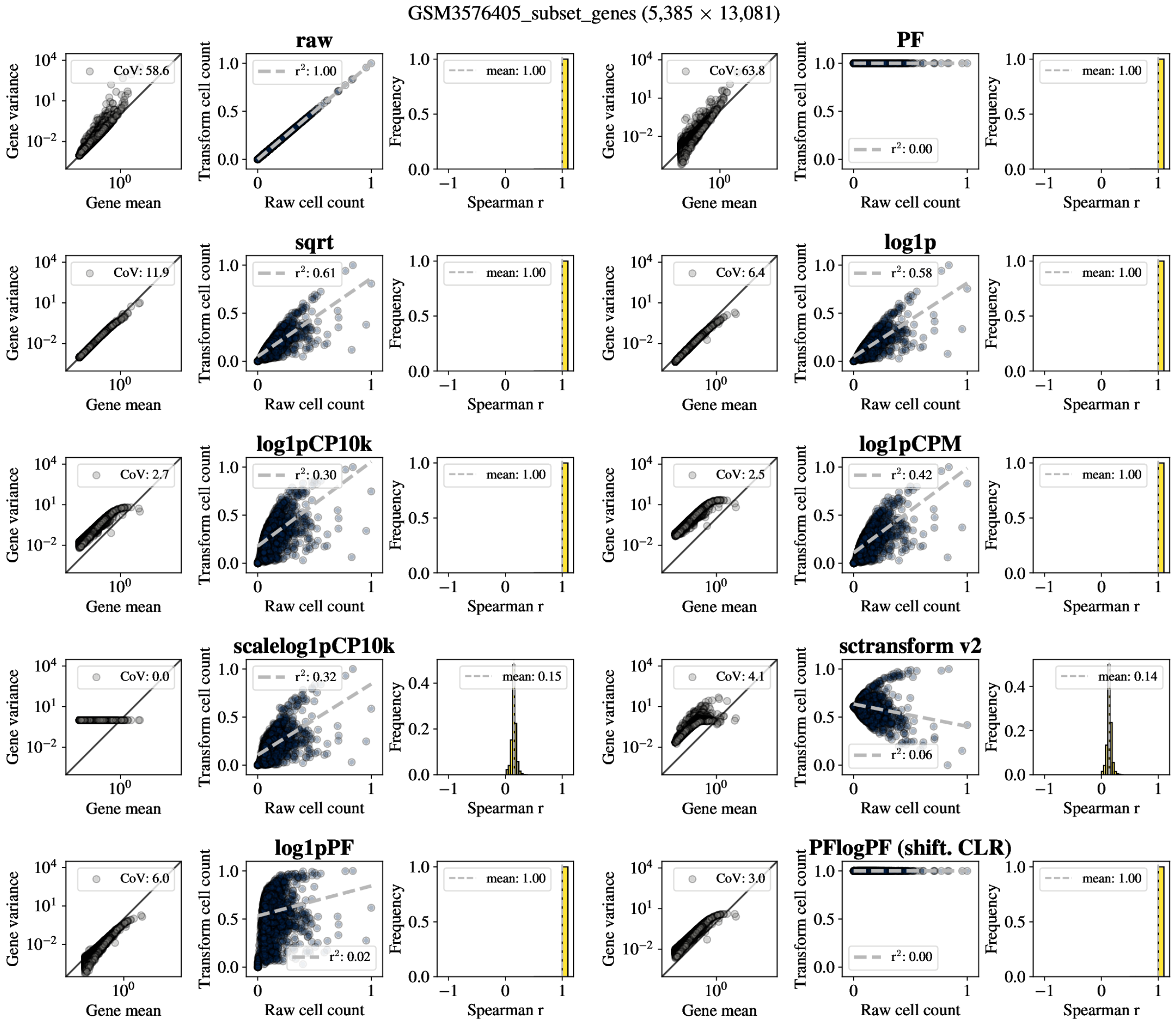


Supplementary Figure 1.310. **GSM3576405**. Per-method comparison of variance stabilization (left), depth normalization (middle), and monotonicity (right) for each normalization on dataset GSM3576405.

| GSM3576406 | |
|---|---|
| cells | 1,653 |
| genes | 60,623 |
| nonzero entries | 402,906 |
| density | 0.00402 |
| total counts | 735,020 |
| mean counts / cell | 445 |
| mean counts / gene | 12 |
| min counts / cell | 4 |
| max counts / cell | 3,743 |
| overdispersion | 2.6 |

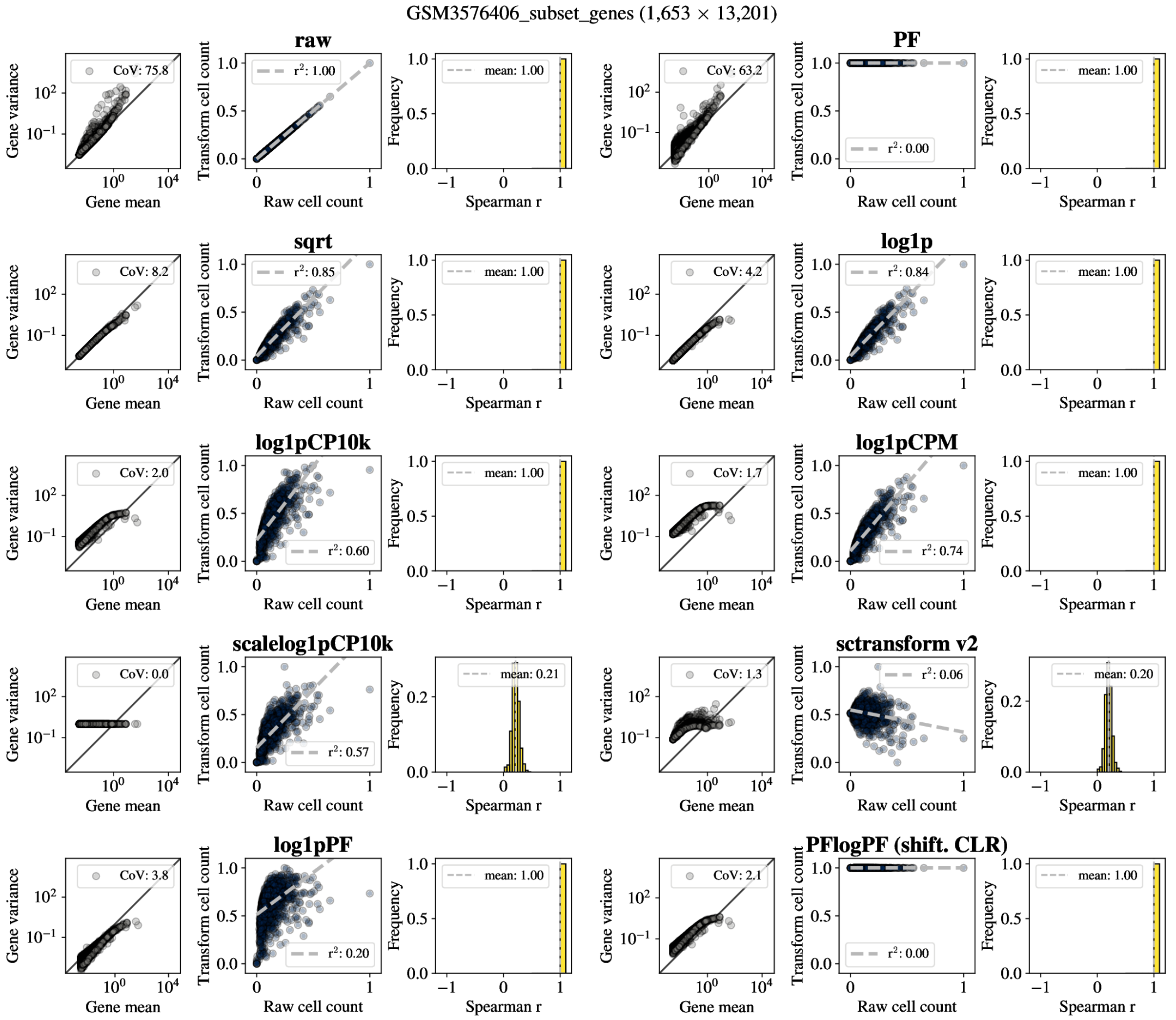


Supplementary Figure 1.311. **GSM3576406**. Per-method comparison of variance stabilization (left), depth normalization (middle), and monotonicity (right) for each normalization on dataset GSM3576406.

| **GSM3576407** | |
|---|---|
| cells | 1,341 |
| genes | 60,623 |
| nonzero entries | 301,569 |
| density | 0.00371 |
| total counts | 515,593 |
| mean counts / cell | 384 |
| mean counts / gene | 9 |
| min counts / cell | 2 |
| max counts / cell | 3,965 |
| overdispersion | 2.57 |

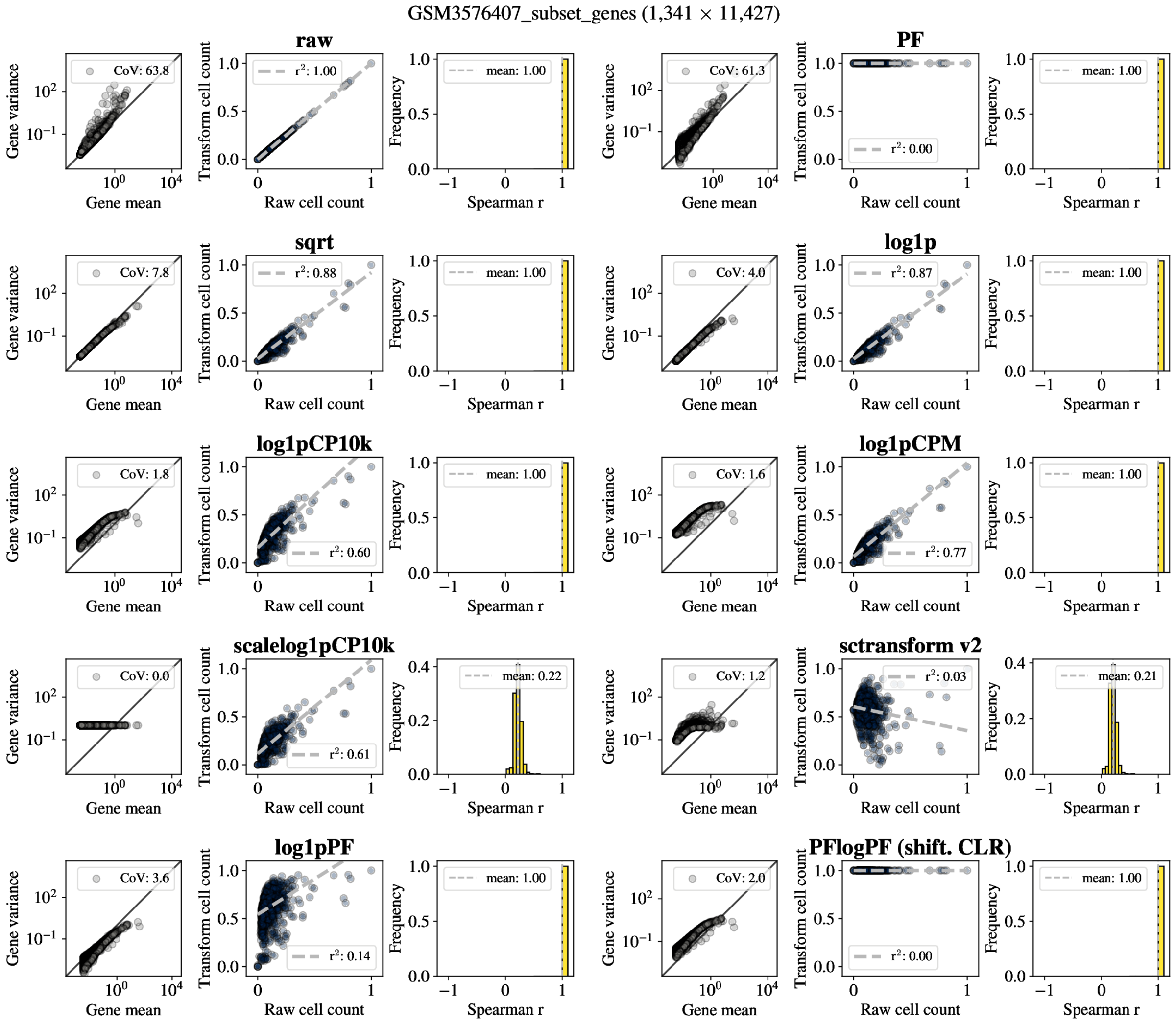


Supplementary Figure 1.312. **GSM3576407**. Per-method comparison of variance stabilization (left), depth normalization (middle), and monotonicity (right) for each normalization on dataset GSM3576407.

| GSM3576408 | |
|---|---|
| cells | 11,255 |
| genes | 60,623 |
| nonzero entries | 1,898,230 |
| density | 0.00278 |
| total counts | 3,130,806 |
| mean counts / cell | 278 |
| mean counts / gene | 52 |
| min counts / cell | 1 |
| max counts / cell | 6,689 |
| overdispersion | 2.99 |

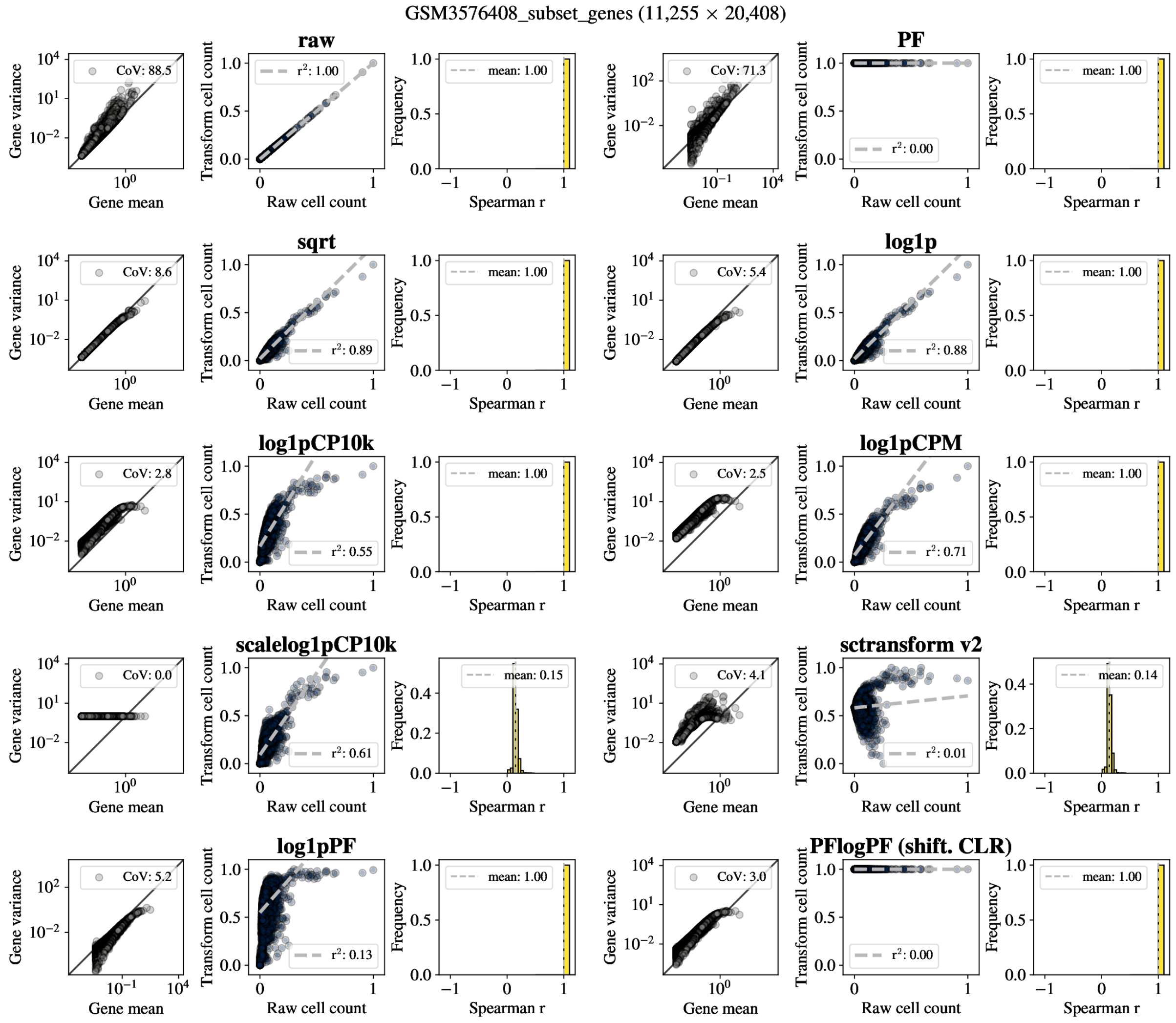


Supplementary Figure 1.313. **GSM3576408**. Per-method comparison of variance stabilization (left), depth normalization (middle), and monotonicity (right) for each normalization on dataset GSM3576408.

| GSM3576409 | |
|---|---|
| cells | 2,273 |
| genes | 60,623 |
| nonzero entries | 395,275 |
| density | 0.00287 |
| total counts | 797,099 |
| mean counts / cell | 351 |
| mean counts / gene | 13 |
| min counts / cell | 0 |
| max counts / cell | 2,998 |
| overdispersion | 4.37 |

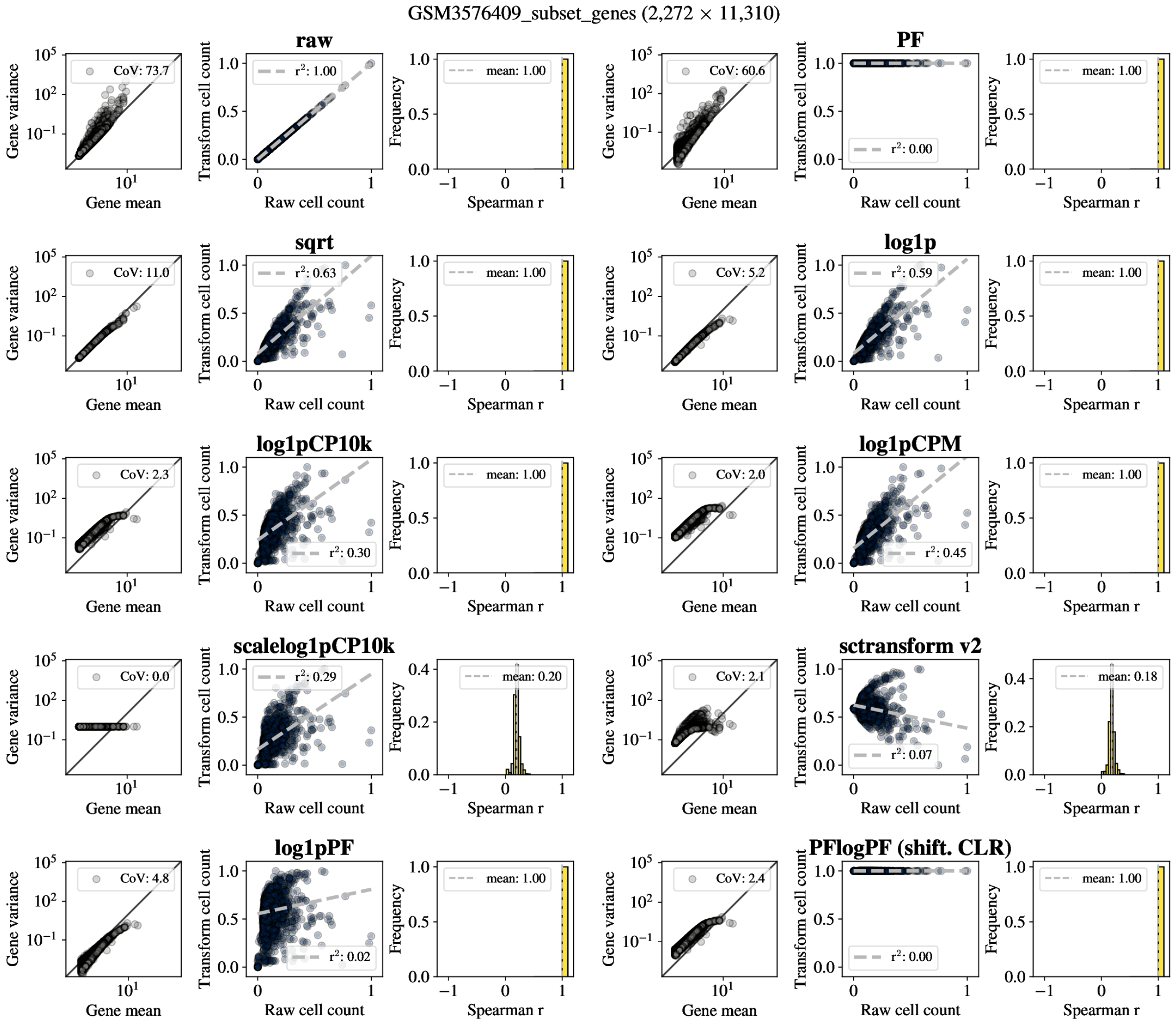


Supplementary Figure 1.314. **GSM3576409**. Per-method comparison of variance stabilization (left), depth normalization (middle), and monotonicity (right) for each normalization on dataset GSM3576409.

| GSM3576410 | |
|---|---|
| cells | 7,901 |
| genes | 60,623 |
| nonzero entries | 1,344,351 |
| density | 0.00281 |
| total counts | 3,108,393 |
| mean counts / cell | 393 |
| mean counts / gene | 51 |
| min counts / cell | 2 |
| max counts / cell | 5,268 |
| overdispersion | 2.46 |

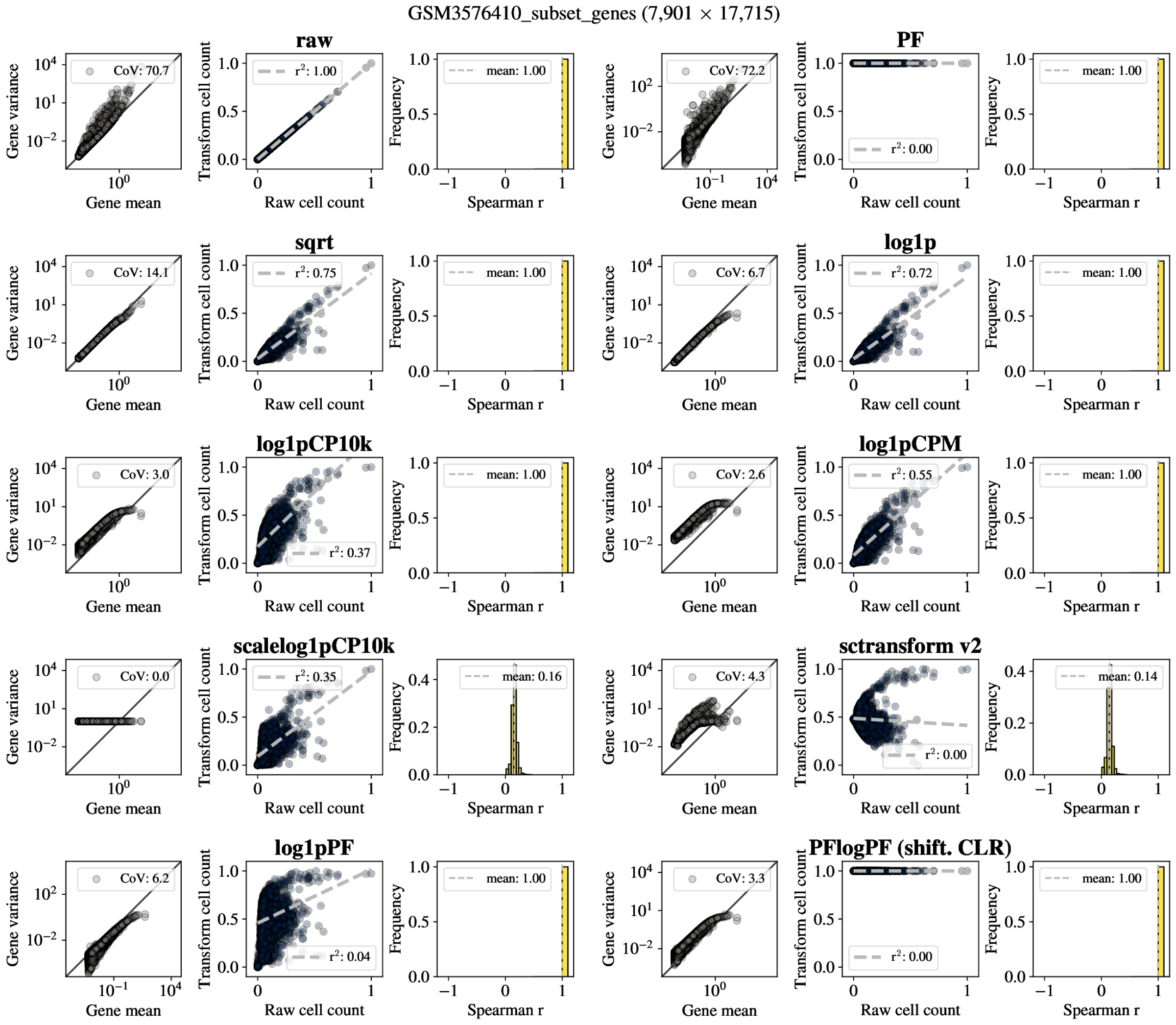


Supplementary Figure 1.315. **GSM3576410**. Per-method comparison of variance stabilization (left), depth normalization (middle), and monotonicity (right) for each normalization on dataset GSM3576410.

| GSM3576411 | |
|---|---|
| cells | 1,925 |
| genes | 60,623 |
| nonzero entries | 425,623 |
| density | 0.00365 |
| total counts | 726,863 |
| mean counts / cell | 378 |
| mean counts / gene | 12 |
| min counts / cell | 1 |
| max counts / cell | 2,326 |
| overdispersion | 2.44 |

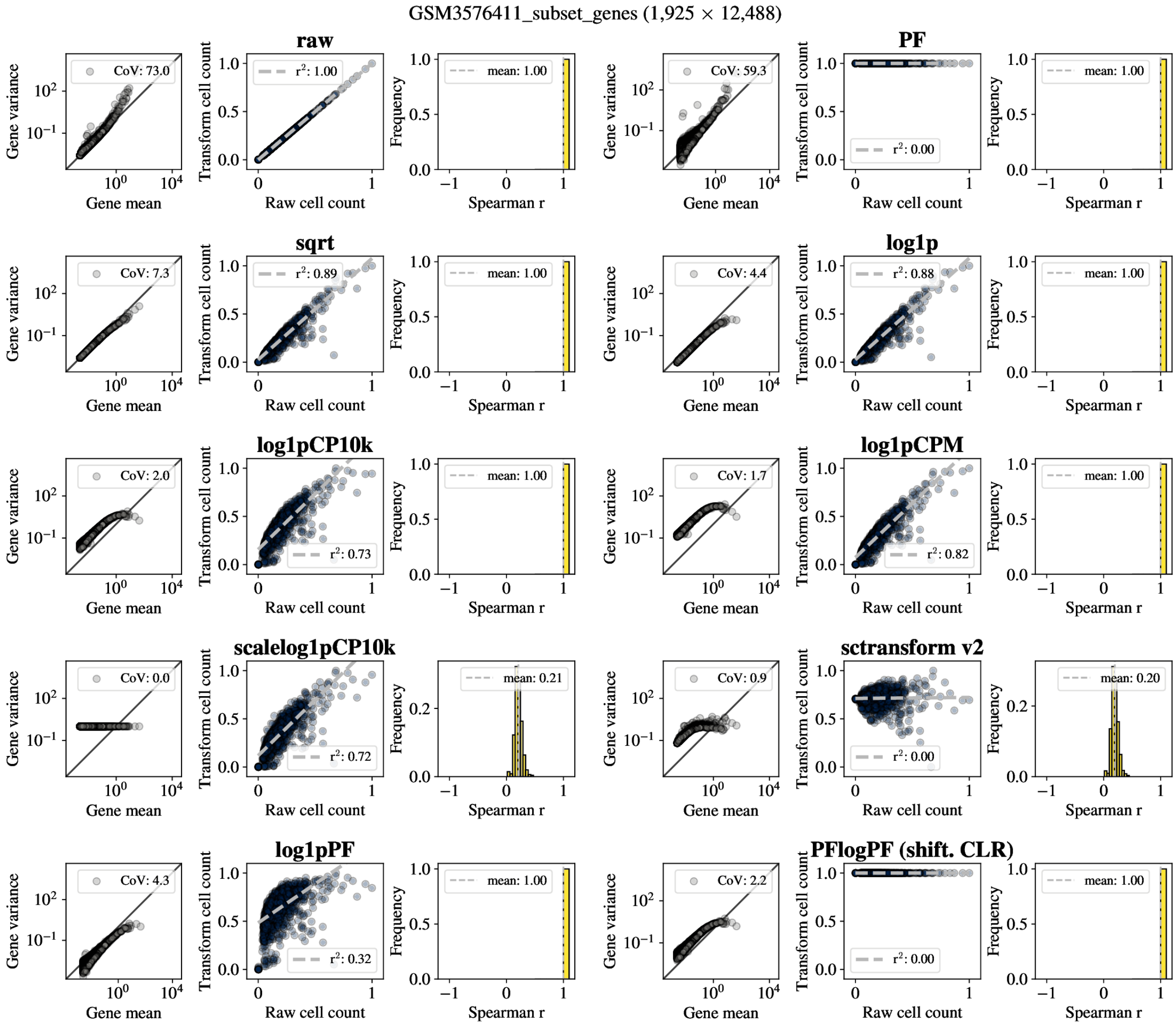


Supplementary Figure 1.316. **GSM3576411**. Per-method comparison of variance stabilization (left), depth normalization (middle), and monotonicity (right) for each normalization on dataset GSM3576411.

| GSM3576412 | |
|---|---|
| cells | 12,882 |
| genes | 60,623 |
| nonzero entries | 2,324,666 |
| density | 0.00298 |
| total counts | 4,780,214 |
| mean counts / cell | 371 |
| mean counts / gene | 79 |
| min counts / cell | 0 |
| max counts / cell | 2,922 |
| overdispersion | 2.62 |

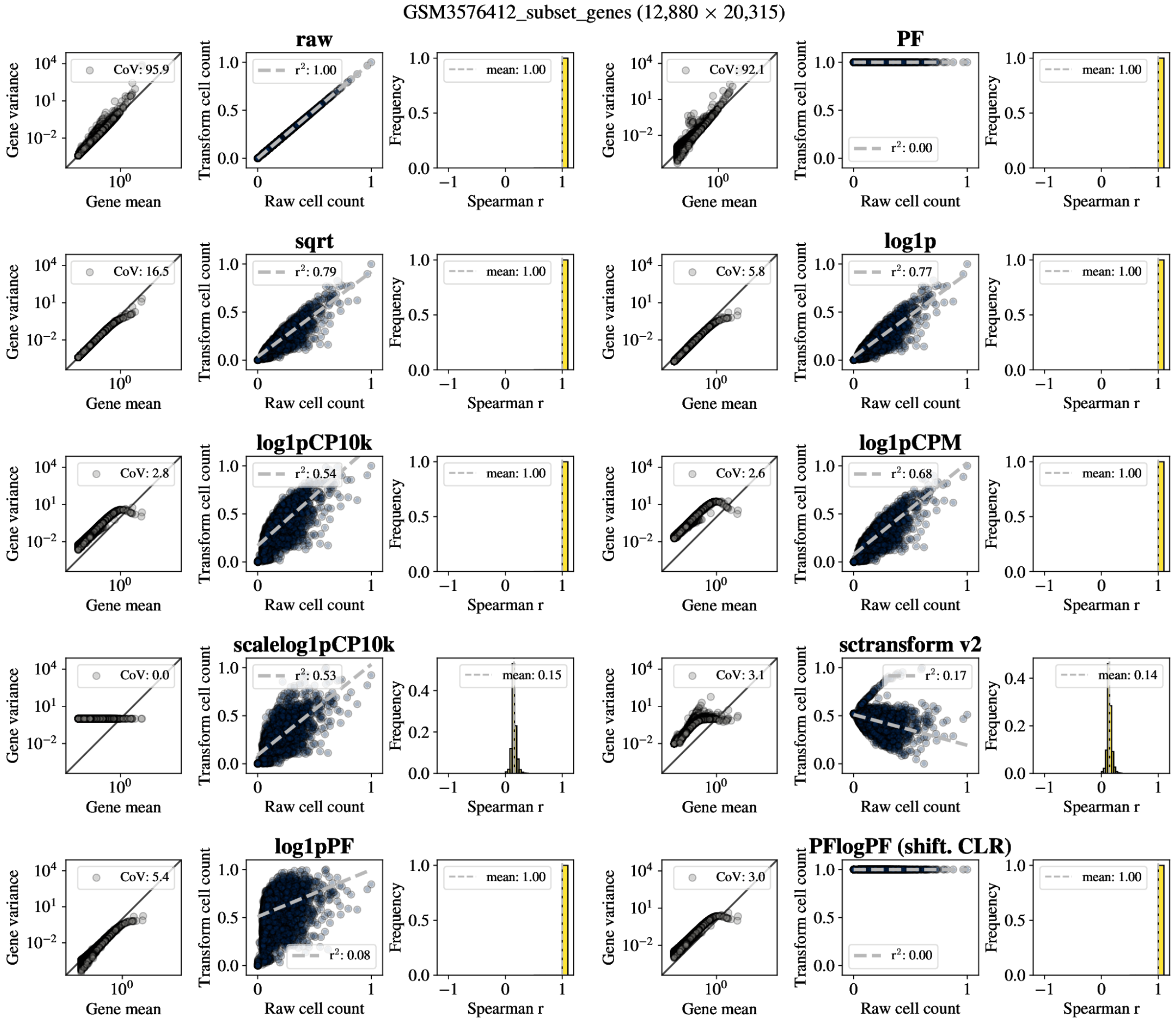


Supplementary Figure 1.317. **GSM3576412**. Per-method comparison of variance stabilization (left), depth normalization (middle), and monotonicity (right) for each normalization on dataset GSM3576412.

| GSM3576413 | |
|---|---|
| cells | 9,212 |
| genes | 60,623 |
| nonzero entries | 935,417 |
| density | 0.00167 |
| total counts | 1,581,212 |
| mean counts / cell | 172 |
| mean counts / gene | 26 |
| min counts / cell | 0 |
| max counts / cell | 1,810 |
| overdispersion | 2.95 |

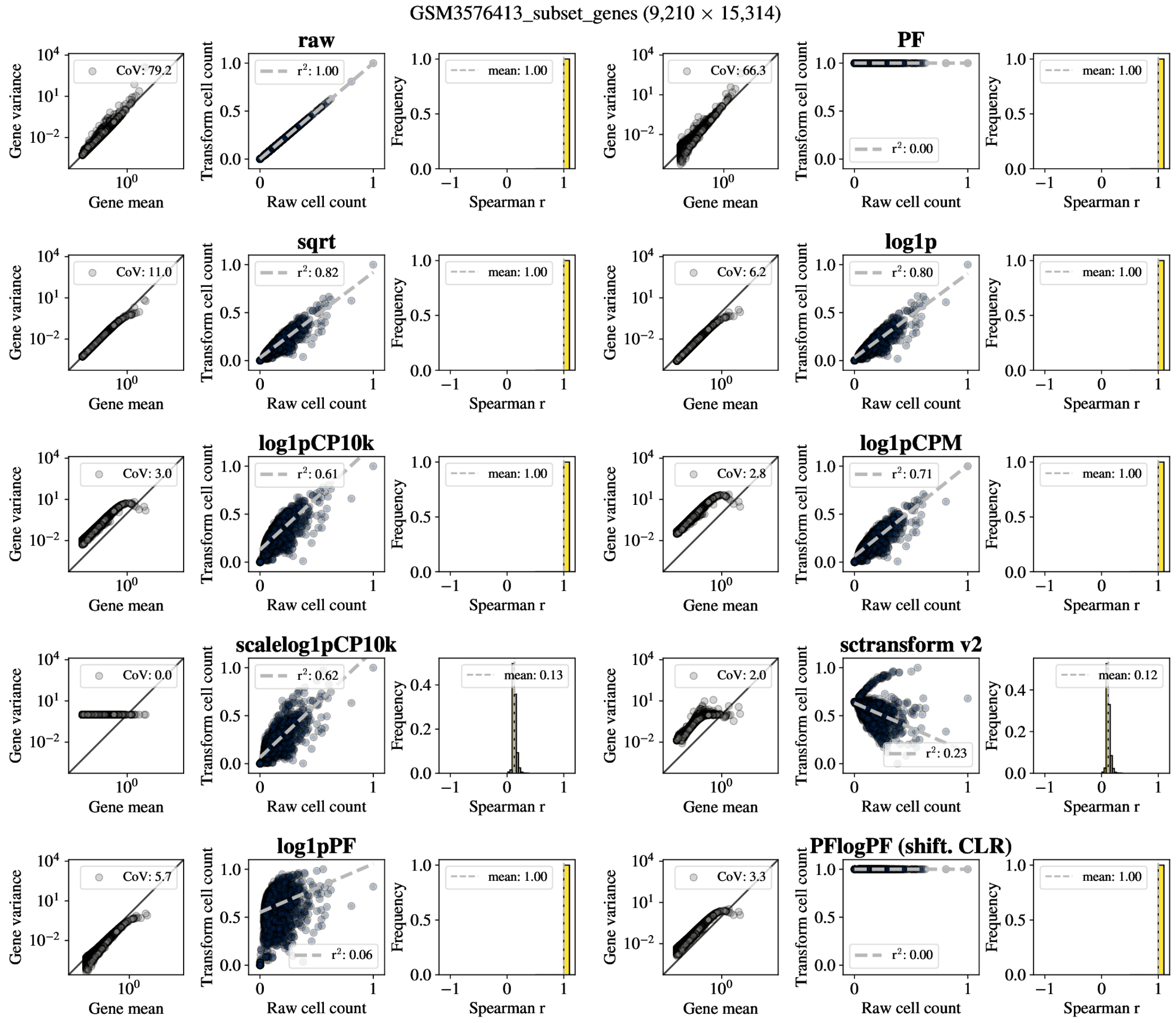


Supplementary Figure 1.318. **GSM3576413**. Per-method comparison of variance stabilization (left), depth normalization (middle), and monotonicity (right) for each normalization on dataset GSM3576413.

| GSM3576414 | |
|---|---|
| cells | 9,797 |
| genes | 60,623 |
| nonzero entries | 989,566 |
| density | 0.00167 |
| total counts | 1,942,685 |
| mean counts / cell | 198 |
| mean counts / gene | 32 |
| min counts / cell | 1 |
| max counts / cell | 4,404 |
| overdispersion | 2.33 |

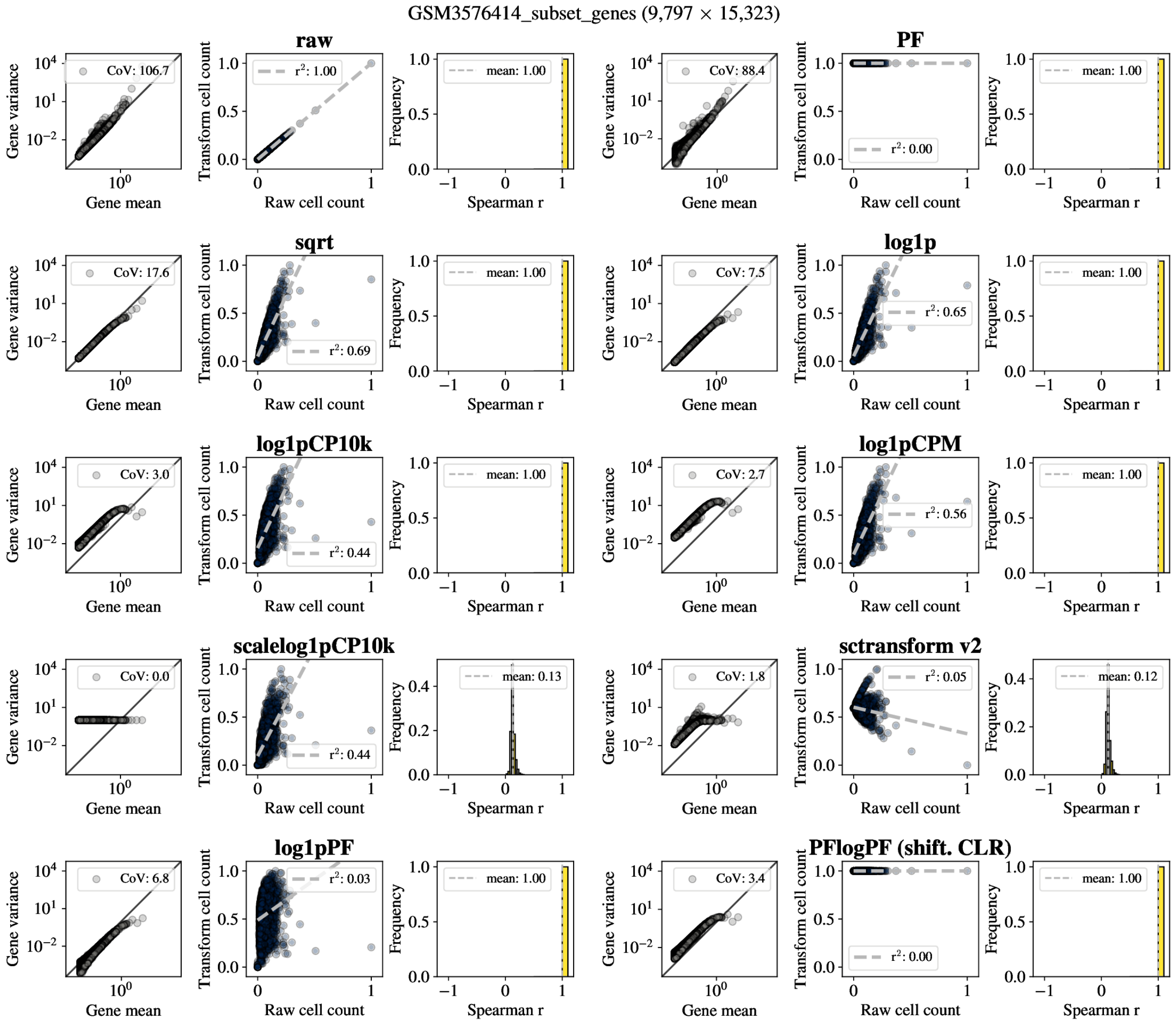


Supplementary Figure 1.319. **GSM3576414**. Per-method comparison of variance stabilization (left), depth normalization (middle), and monotonicity (right) for each normalization on dataset GSM3576414.

| GSM3576415 | |
|---|---|
| cells | 8,802 |
| genes | 60,623 |
| nonzero entries | 1,437,072 |
| density | 0.00269 |
| total counts | 2,318,037 |
| mean counts / cell | 263 |
| mean counts / gene | 38 |
| min counts / cell | 1 |
| max counts / cell | 3,488 |
| overdispersion | 3.18 |

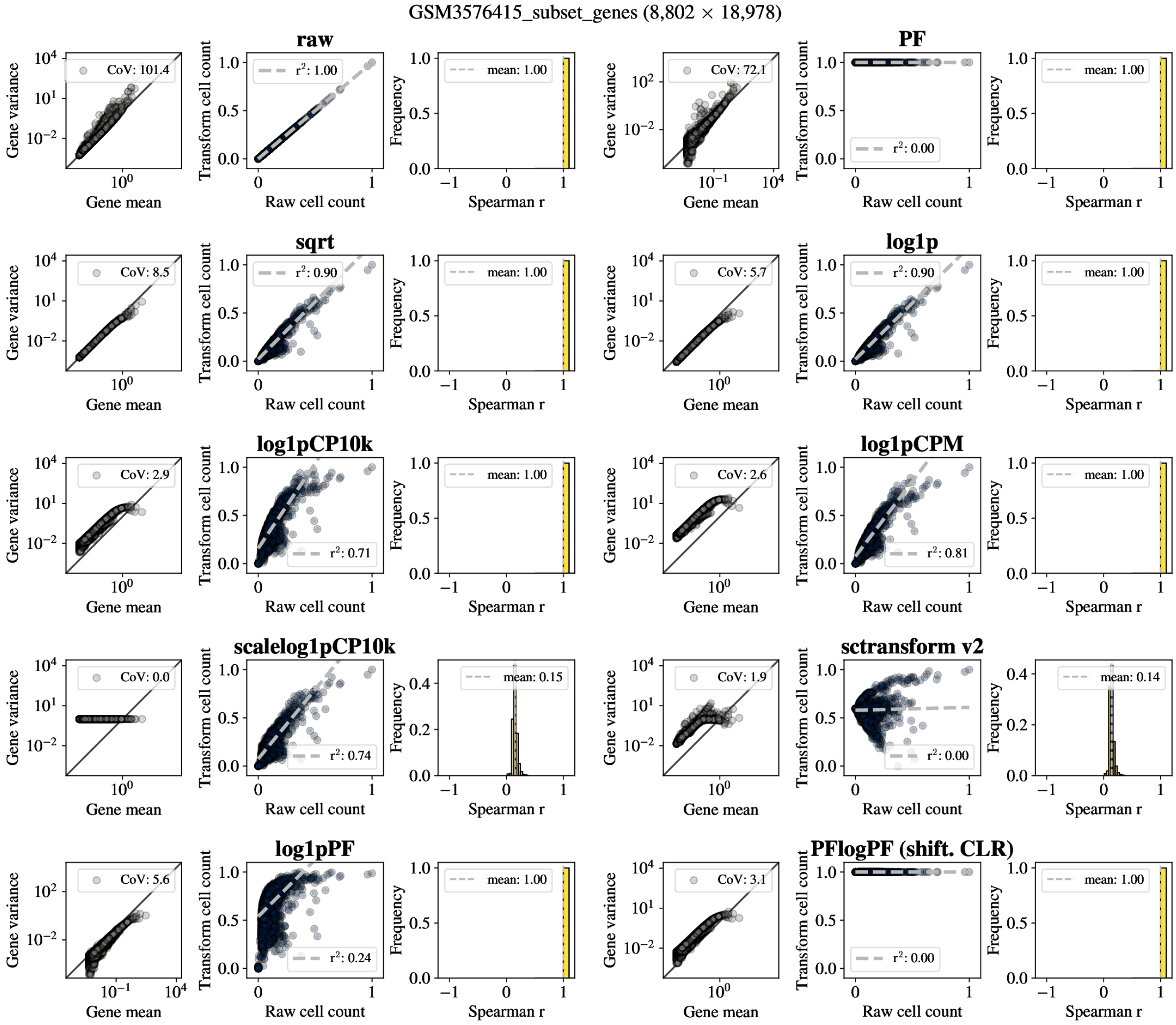


Supplementary Figure 1.320. **GSM3576415**. Per-method comparison of variance stabilization (left), depth normalization (middle), and monotonicity (right) for each normalization on dataset GSM3576415.

| GSM3576416 | |
|---|---|
| cells | 6,886 |
| genes | 60,623 |
| nonzero entries | 683,009 |
| density | 0.00164 |
| total counts | 1,105,585 |
| mean counts / cell | 161 |
| mean counts / gene | 18 |
| min counts / cell | 1 |
| max counts / cell | 3,249 |
| overdispersion | 1.88 |

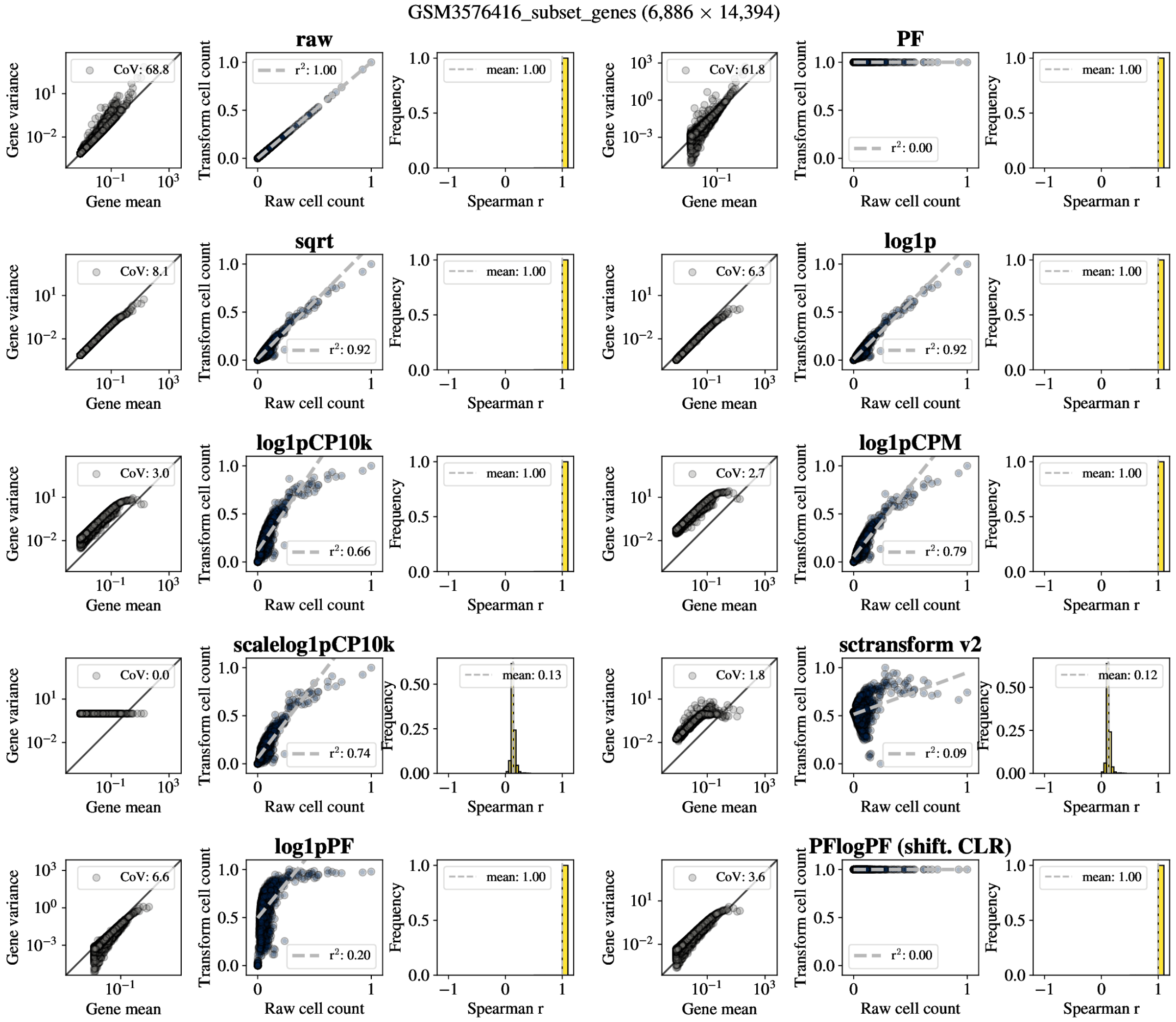


Supplementary Figure 1.321. **GSM3576416**. Per-method comparison of variance stabilization (left), depth normalization (middle), and monotonicity (right) for each normalization on dataset GSM3576416.

| GSM3576417 | |
|---|---|
| cells | 5,143 |
| genes | 60,623 |
| nonzero entries | 479,007 |
| density | 0.00154 |
| total counts | 744,338 |
| mean counts / cell | 145 |
| mean counts / gene | 12 |
| min counts / cell | 2 |
| max counts / cell | 1,134 |
| overdispersion | 2.91 |

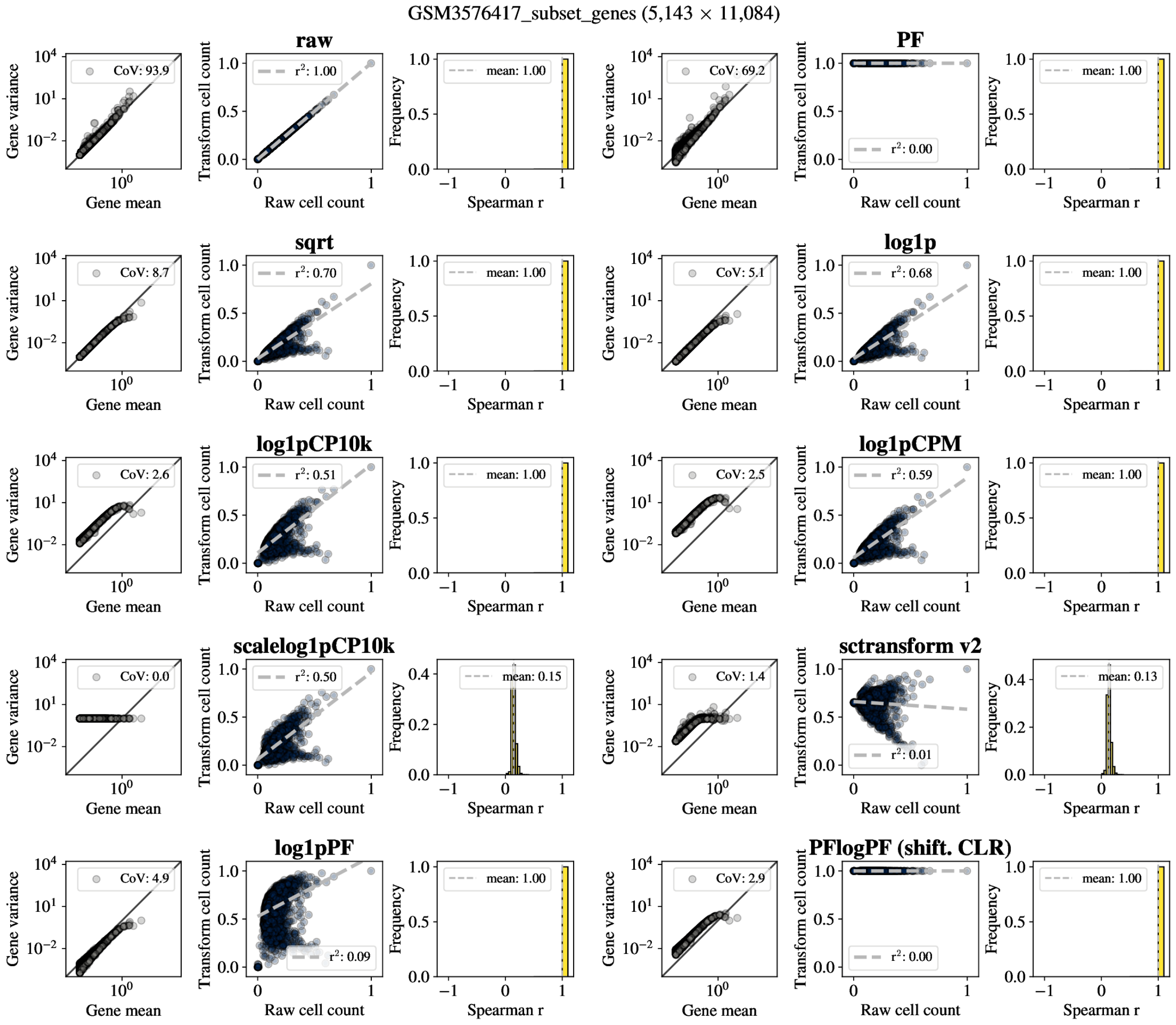


Supplementary Figure 1.322. **GSM3576417**. Per-method comparison of variance stabilization (left), depth normalization (middle), and monotonicity (right) for each normalization on dataset GSM3576417.

| GSM3576418 | |
|---|---|
| cells | 8,013 |
| genes | 60,623 |
| nonzero entries | 836,519 |
| density | 0.00172 |
| total counts | 1,188,333 |
| mean counts / cell | 148 |
| mean counts / gene | 20 |
| min counts / cell | 1 |
| max counts / cell | 1,174 |
| overdispersion | 3.65 |

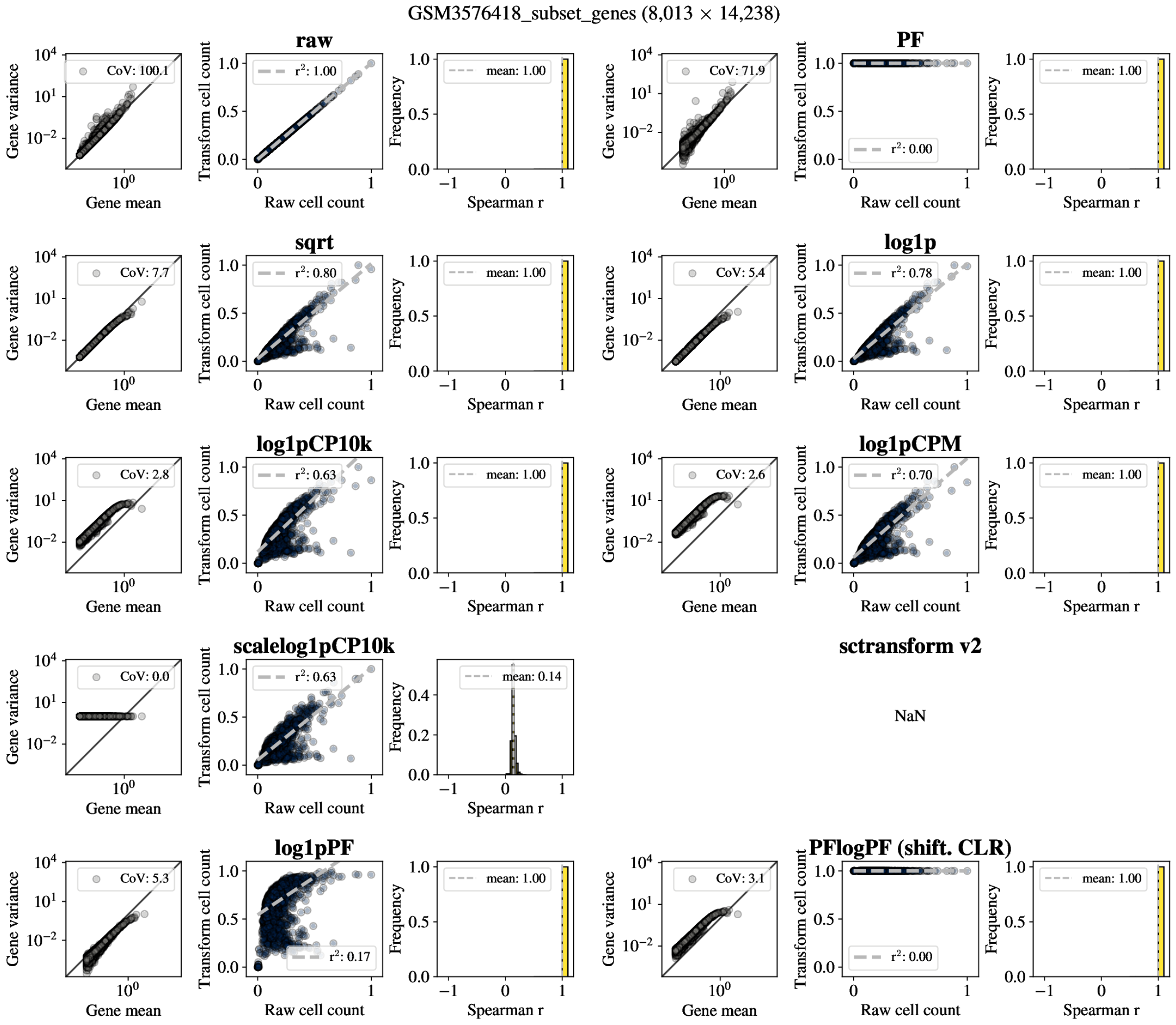


Supplementary Figure 1.323. **GSM3576418**. Per-method comparison of variance stabilization (left), depth normalization (middle), and monotonicity (right) for each normalization on dataset GSM3576418.

| GSM3576419 | |
|---|---|
| cells | 10,214 |
| genes | 60,623 |
| nonzero entries | 1,067,789 |
| density | 0.00172 |
| total counts | 1,494,107 |
| mean counts / cell | 146 |
| mean counts / gene | 25 |
| min counts / cell | 1 |
| max counts / cell | 1,517 |
| overdispersion | 4.35 |

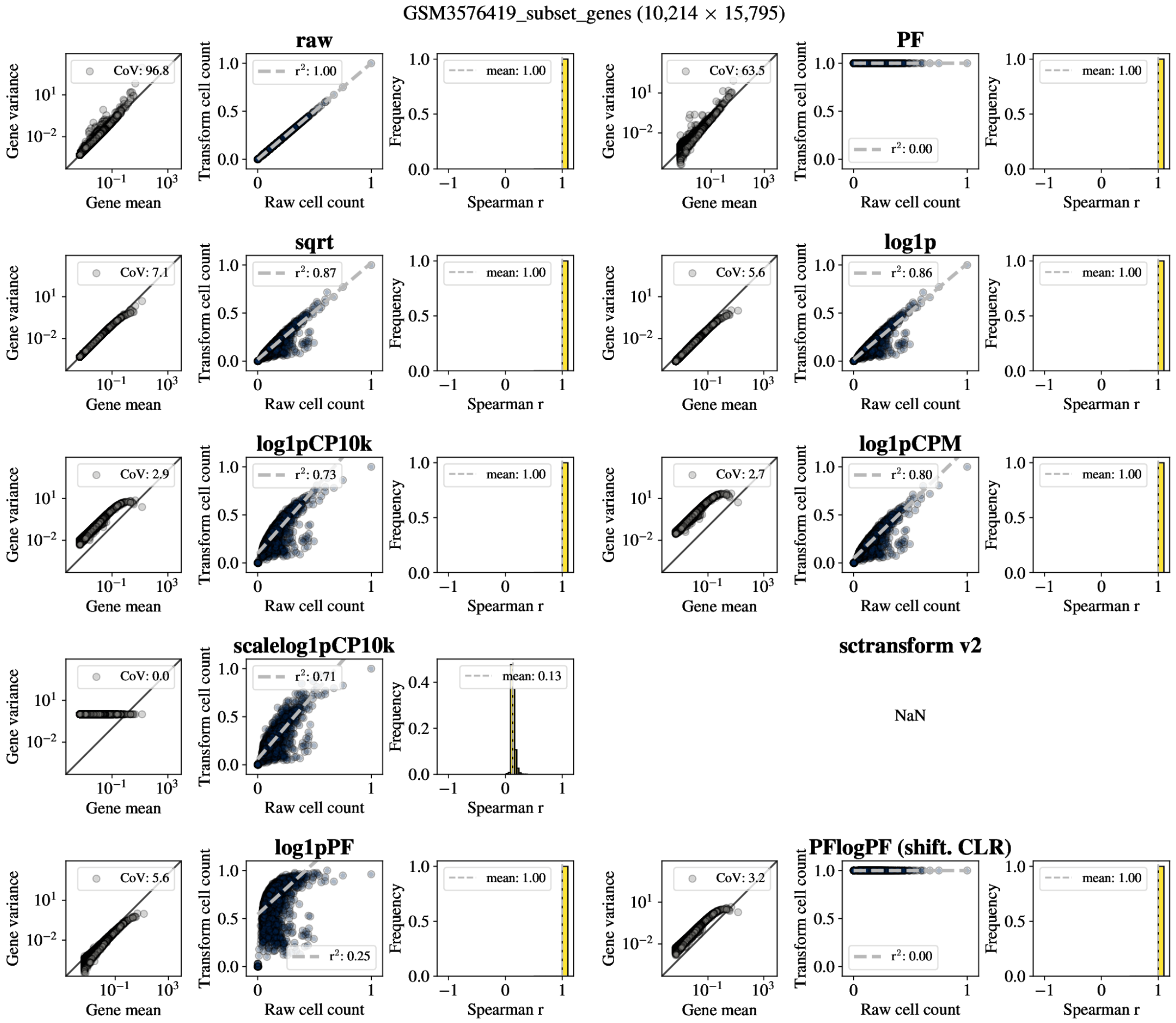


Supplementary Figure 1.324. **GSM3576419**. Per-method comparison of variance stabilization (left), depth normalization (middle), and monotonicity (right) for each normalization on dataset GSM3576419.

| GSM3576420 | |
|---|---|
| cells | 3,341 |
| genes | 60,623 |
| nonzero entries | 1,219,528 |
| density | 0.00602 |
| total counts | 2,003,823 |
| mean counts / cell | 600 |
| mean counts / gene | 33 |
| min counts / cell | 2 |
| max counts / cell | 5,778 |
| overdispersion | 2.83 |

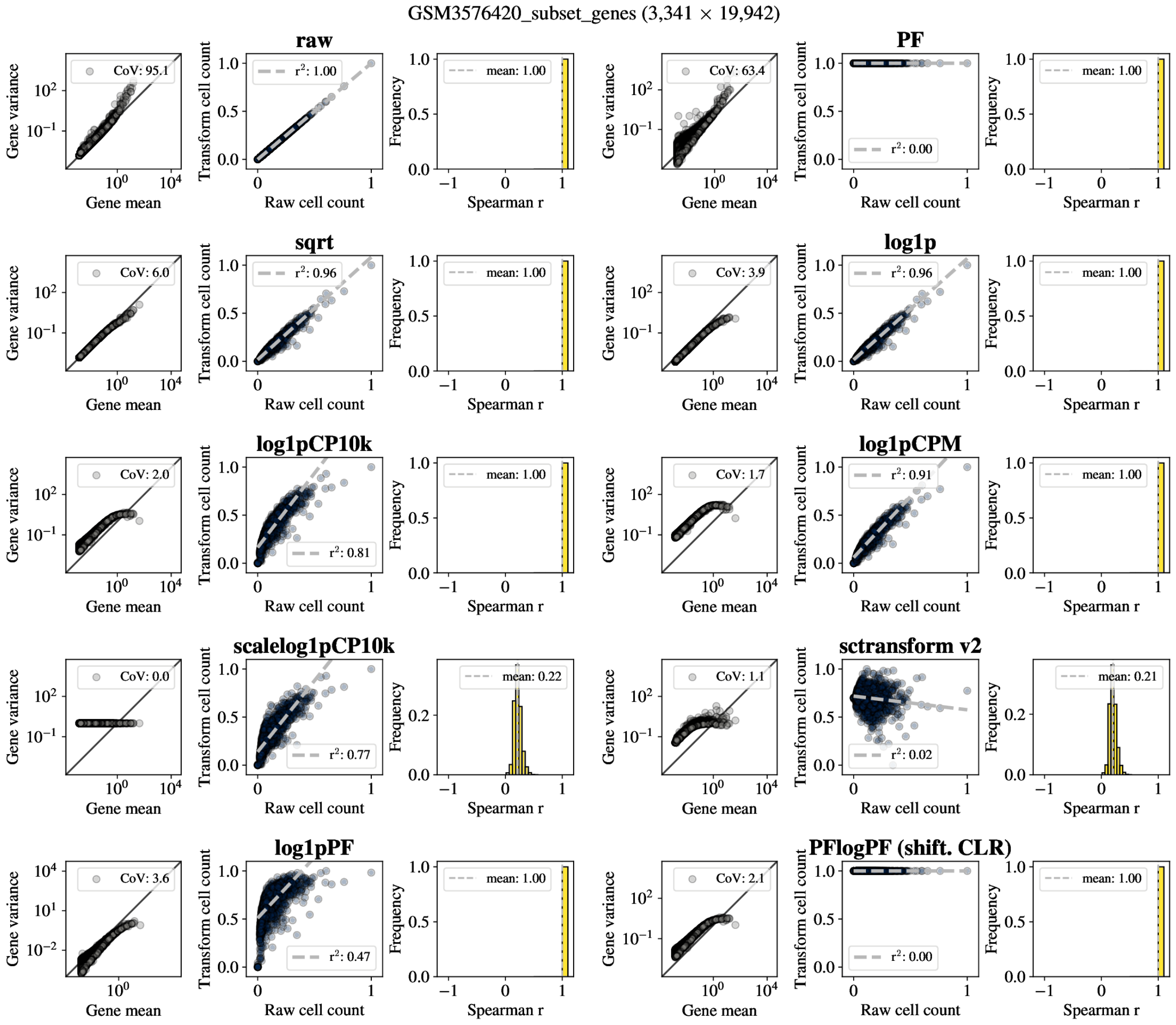


Supplementary Figure 1.325. **GSM3576420**. Per-method comparison of variance stabilization (left), depth normalization (middle), and monotonicity (right) for each normalization on dataset GSM3576420.

| GSM3576421 | |
|---|---|
| cells | 1,322 |
| genes | 60,623 |
| nonzero entries | 314,168 |
| density | 0.00392 |
| total counts | 482,911 |
| mean counts / cell | 365 |
| mean counts / gene | 8 |
| min counts / cell | 2 |
| max counts / cell | 2,305 |
| overdispersion | 2.28 |

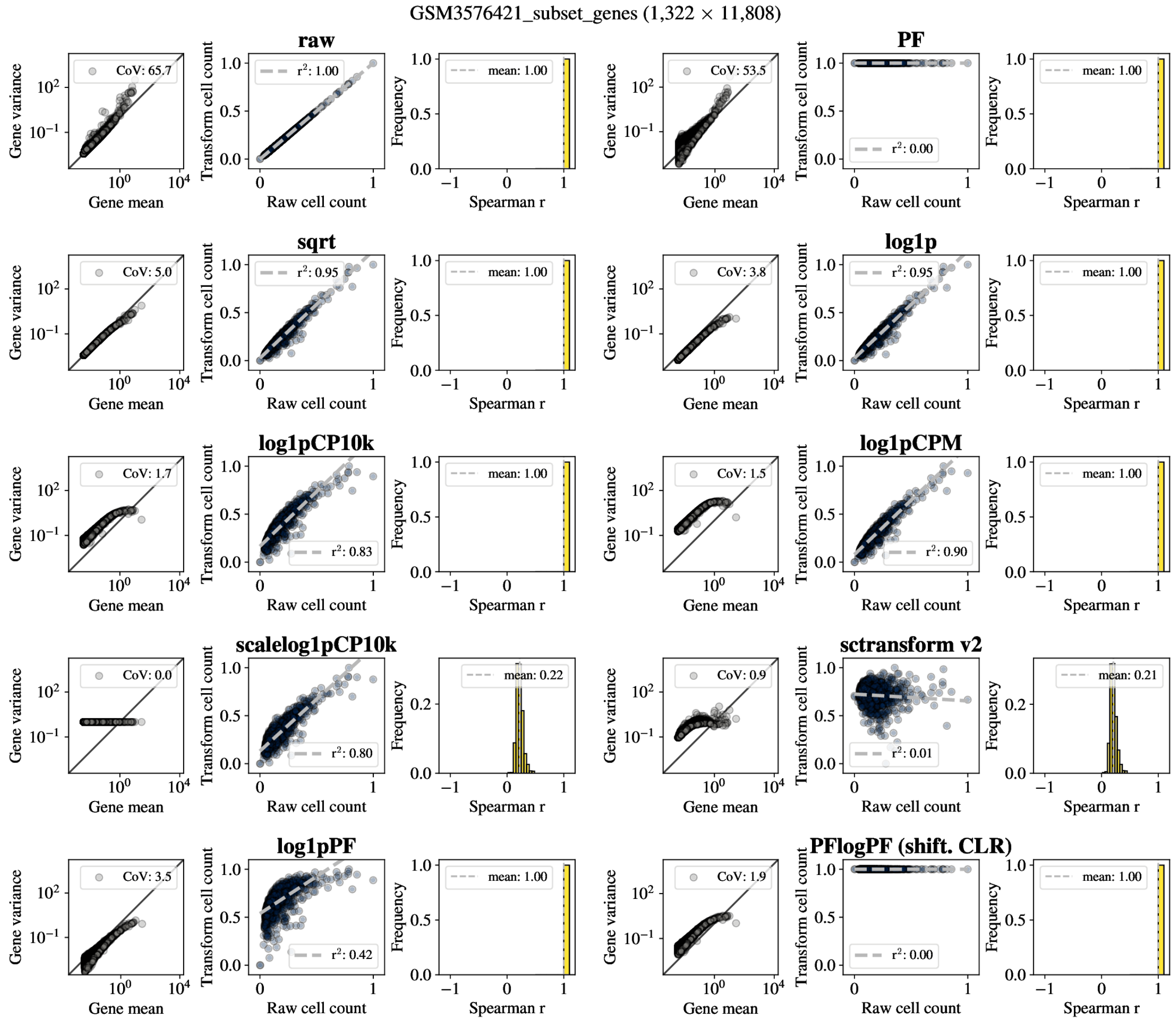


Supplementary Figure 1.326. **GSM3576421**. Per-method comparison of variance stabilization (left), depth normalization (middle), and monotonicity (right) for each normalization on dataset GSM3576421.

| GSM3576422 | |
|---|---|
| cells | 8,784 |
| genes | 60,623 |
| nonzero entries | 1,661,784 |
| density | 0.00312 |
| total counts | 2,628,095 |
| mean counts / cell | 299 |
| mean counts / gene | 43 |
| min counts / cell | 2 |
| max counts / cell | 6,294 |
| overdispersion | 2.52 |

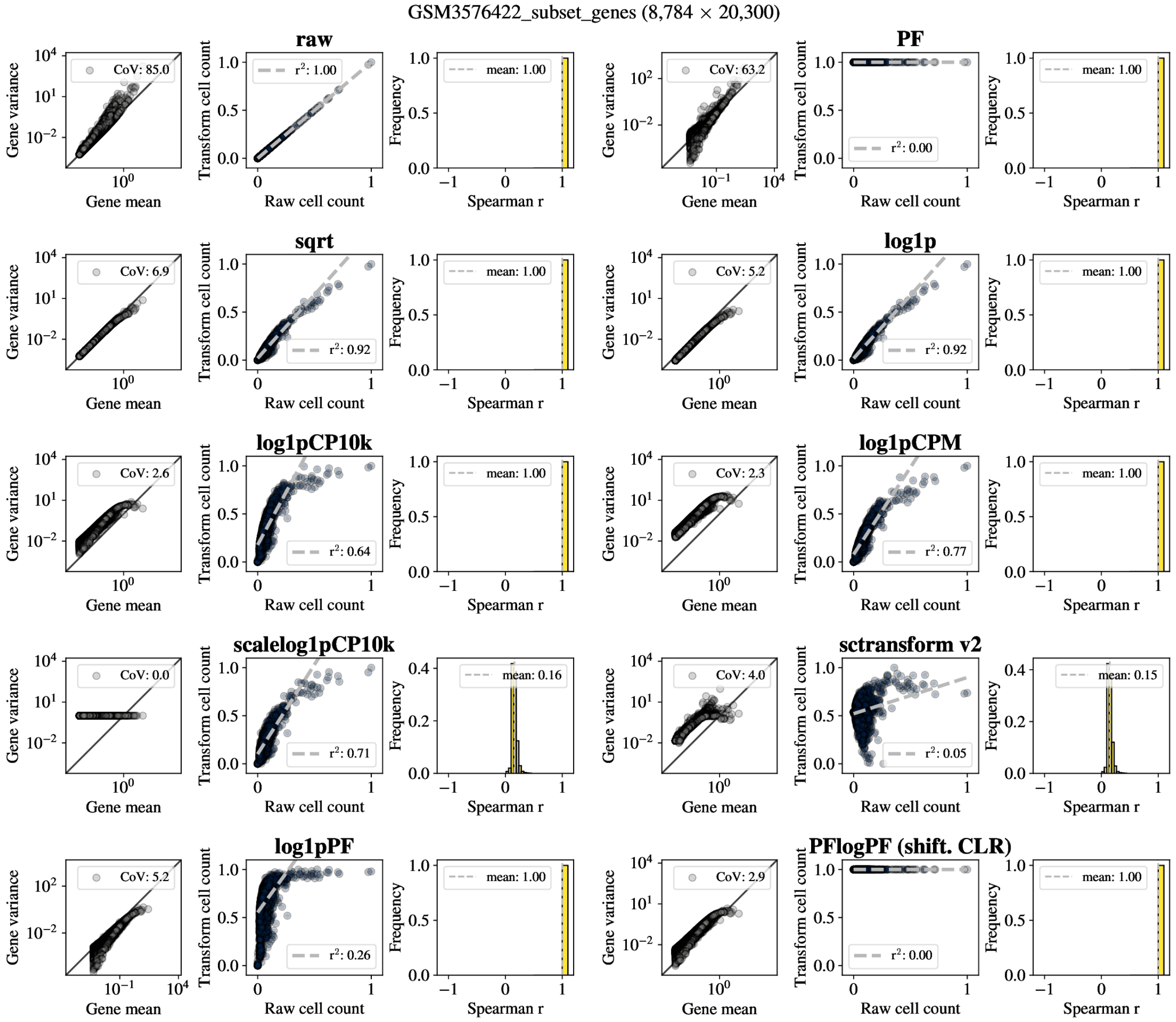


Supplementary Figure 1.327. **GSM3576422**. Per-method comparison of variance stabilization (left), depth normalization (middle), and monotonicity (right) for each normalization on dataset GSM3576422.

| GSM3576423 | |
|---|---|
| cells | 918 |
| genes | 60,623 |
| nonzero entries | 303,801 |
| density | 0.00546 |
| total counts | 559,583 |
| mean counts / cell | 610 |
| mean counts / gene | 9 |
| min counts / cell | 8 |
| max counts / cell | 5,328 |
| overdispersion | 3.3 |

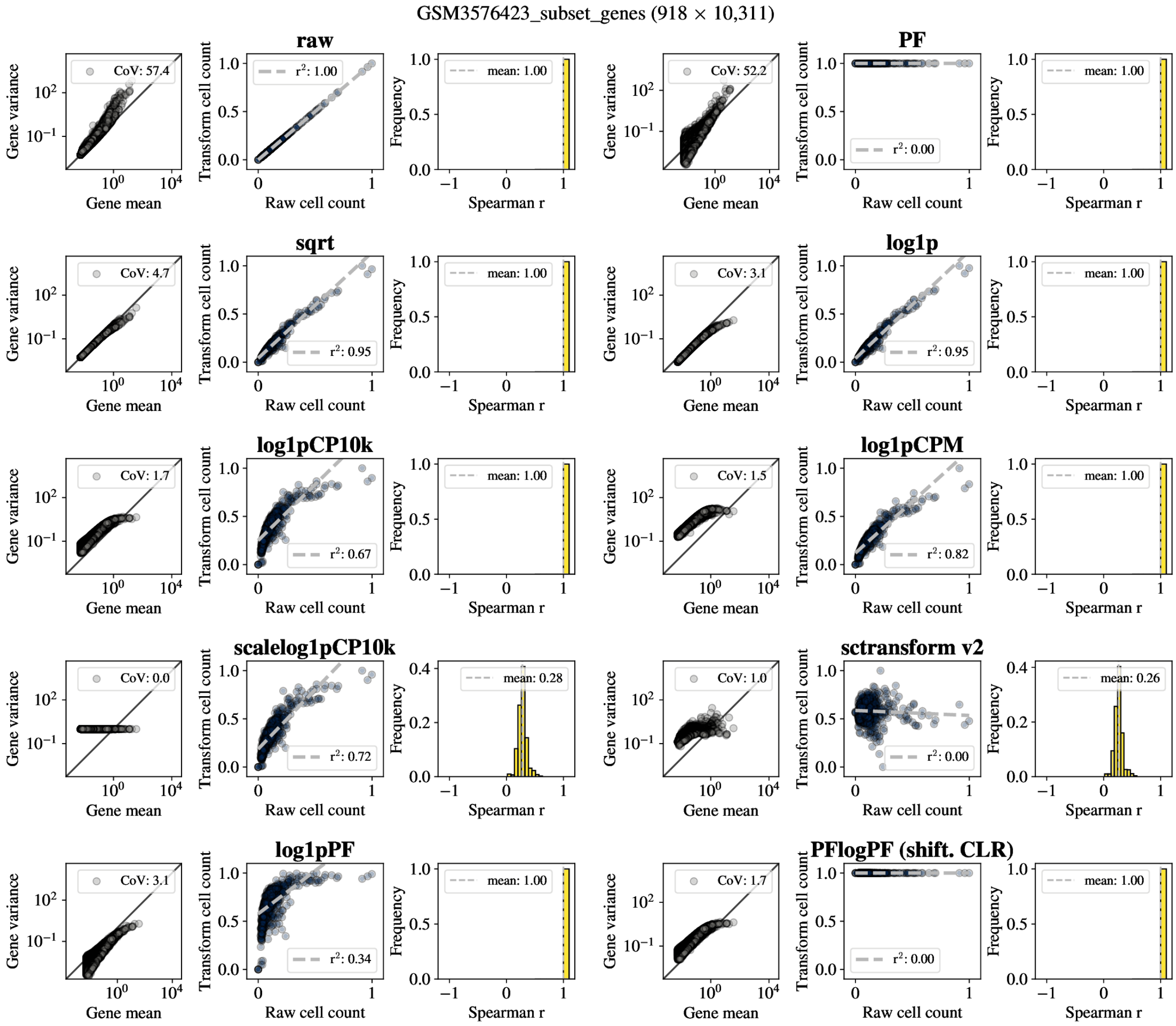


Supplementary Figure 1.328. **GSM3576423**. Per-method comparison of variance stabilization (left), depth normalization (middle), and monotonicity (right) for each normalization on dataset GSM3576423.

| GSM3576424 | |
|---|---|
| cells | 939 |
| genes | 60,623 |
| nonzero entries | 336,537 |
| density | 0.00591 |
| total counts | 614,581 |
| mean counts / cell | 655 |
| mean counts / gene | 10 |
| min counts / cell | 6 |
| max counts / cell | 2,879 |
| overdispersion | 4.14 |

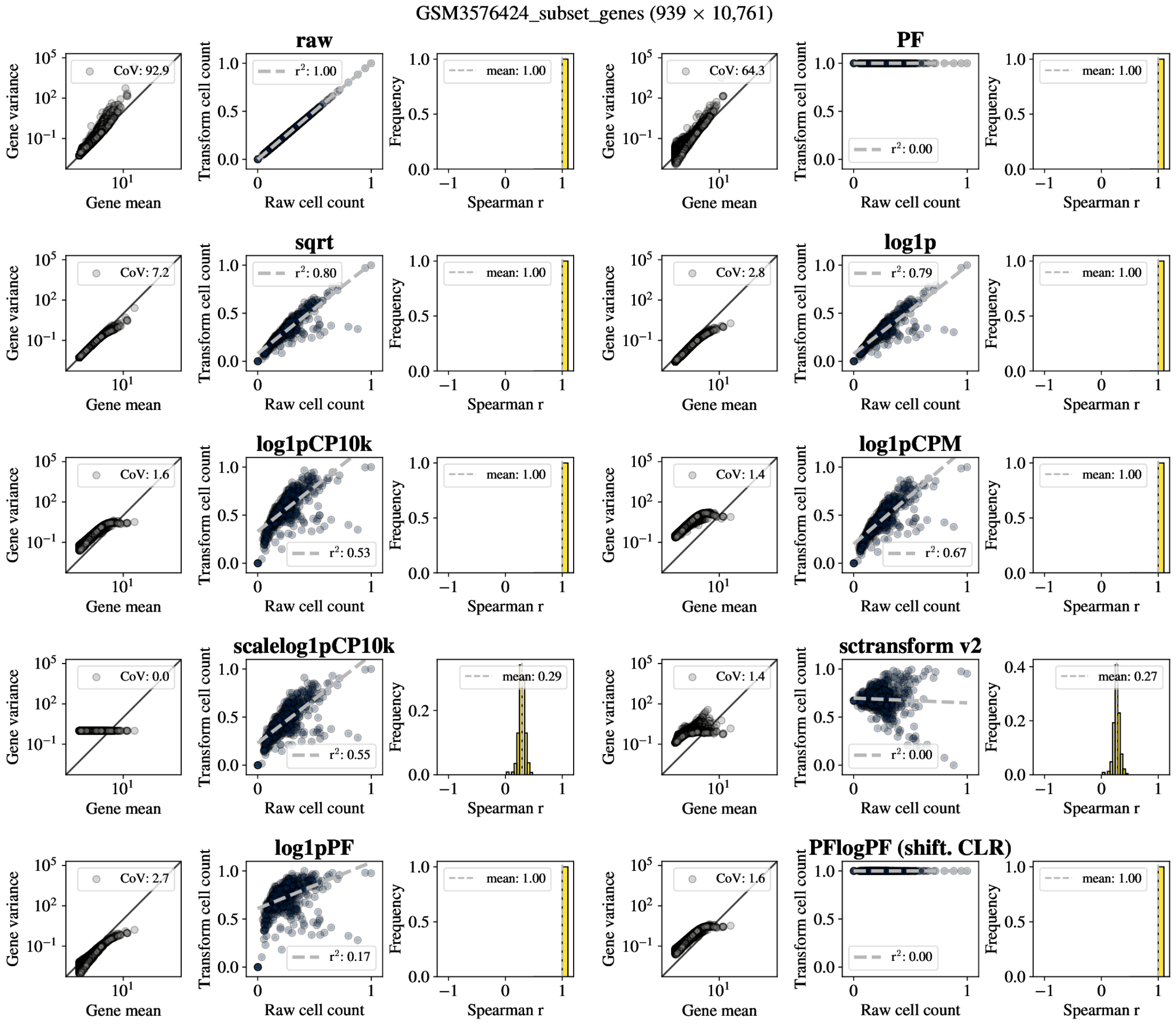


Supplementary Figure 1.329. **GSM3576424**. Per-method comparison of variance stabilization (left), depth normalization (middle), and monotonicity (right) for each normalization on dataset GSM3576424.

| GSM3576425 | |
|---|---|
| cells | 3,009 |
| genes | 60,623 |
| nonzero entries | 640,573 |
| density | 0.00351 |
| total counts | 977,569 |
| mean counts / cell | 325 |
| mean counts / gene | 16 |
| min counts / cell | 1 |
| max counts / cell | 2,090 |
| overdispersion | 2.73 |

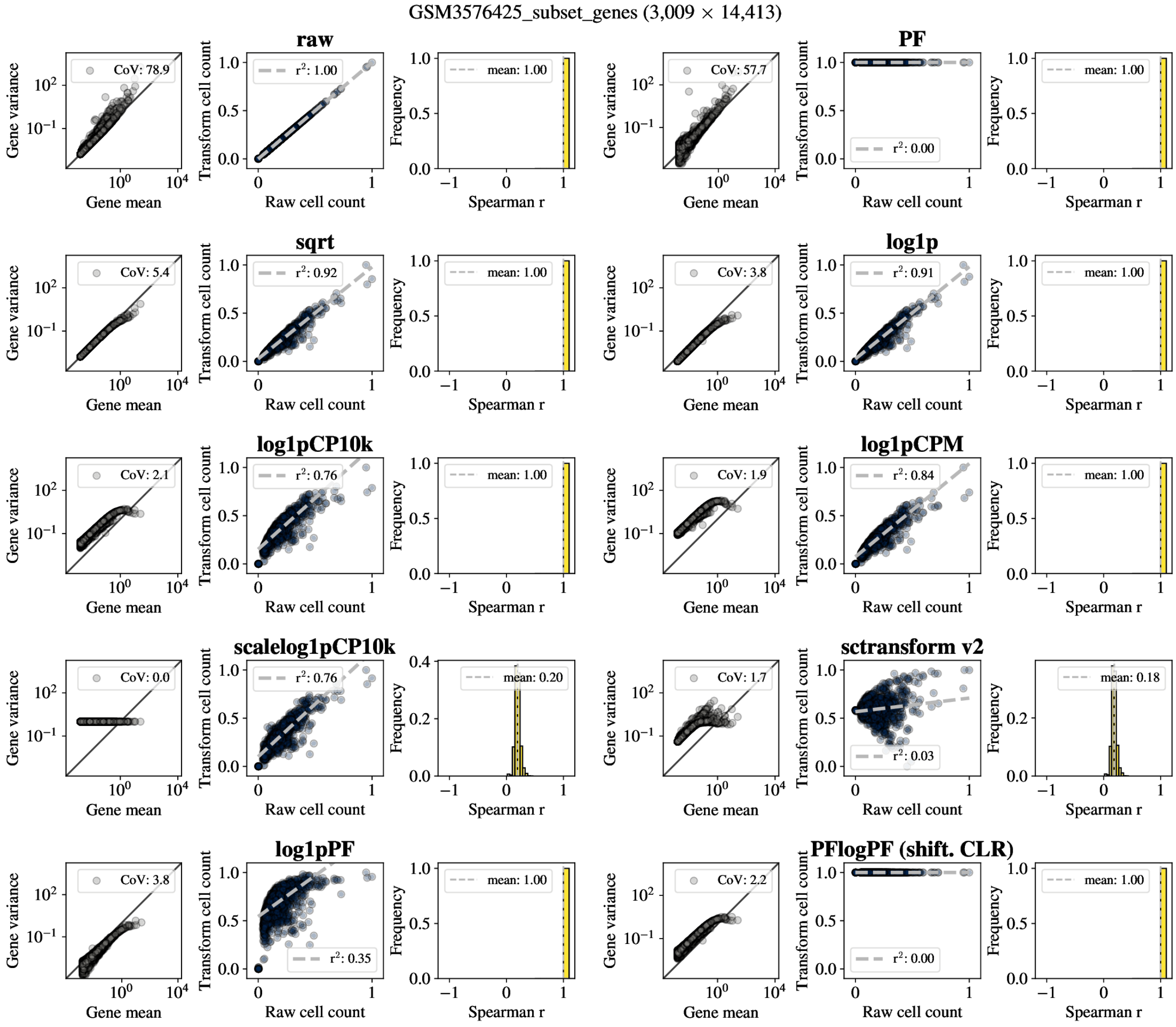


Supplementary Figure 1.330. **GSM3576425**. Per-method comparison of variance stabilization (left), depth normalization (middle), and monotonicity (right) for each normalization on dataset GSM3576425.

| GSM3576426 | |
|---|---|
| cells | 1,919 |
| genes | 60,623 |
| nonzero entries | 360,129 |
| density | 0.0031 |
| total counts | 589,554 |
| mean counts / cell | 307 |
| mean counts / gene | 10 |
| min counts / cell | 2 |
| max counts / cell | 2,978 |
| overdispersion | 4.29 |

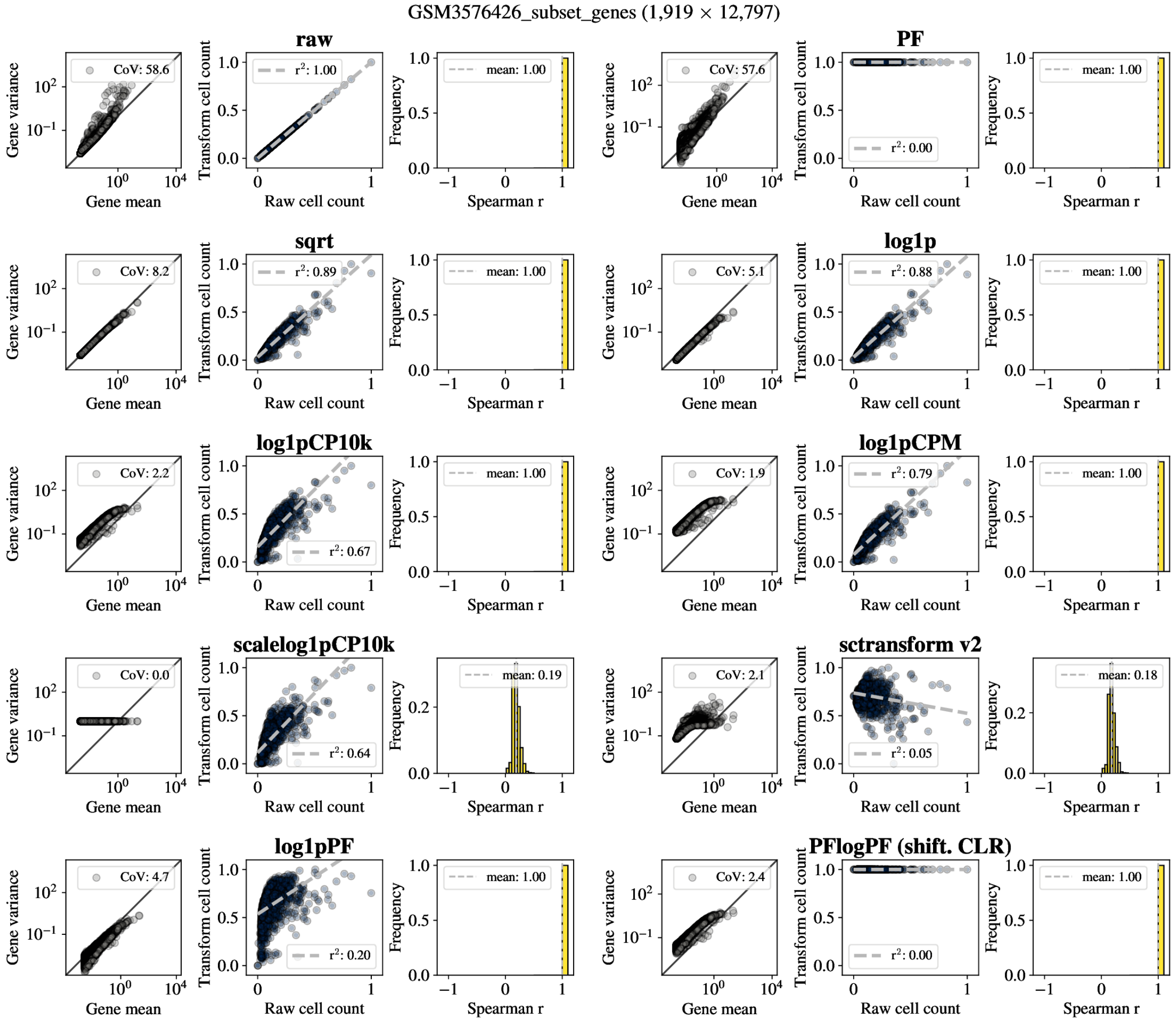


Supplementary Figure 1.331. **GSM3576426**. Per-method comparison of variance stabilization (left), depth normalization (middle), and monotonicity (right) for each normalization on dataset GSM3576426.

| **GSM3576427** | |
|---|---|
| cells | 1,479 |
| genes | 60,623 |
| nonzero entries | 414,396 |
| density | 0.00462 |
| total counts | 742,941 |
| mean counts / cell | 502 |
| mean counts / gene | 12 |
| min counts / cell | 7 |
| max counts / cell | 5,305 |
| overdispersion | 2.43 |

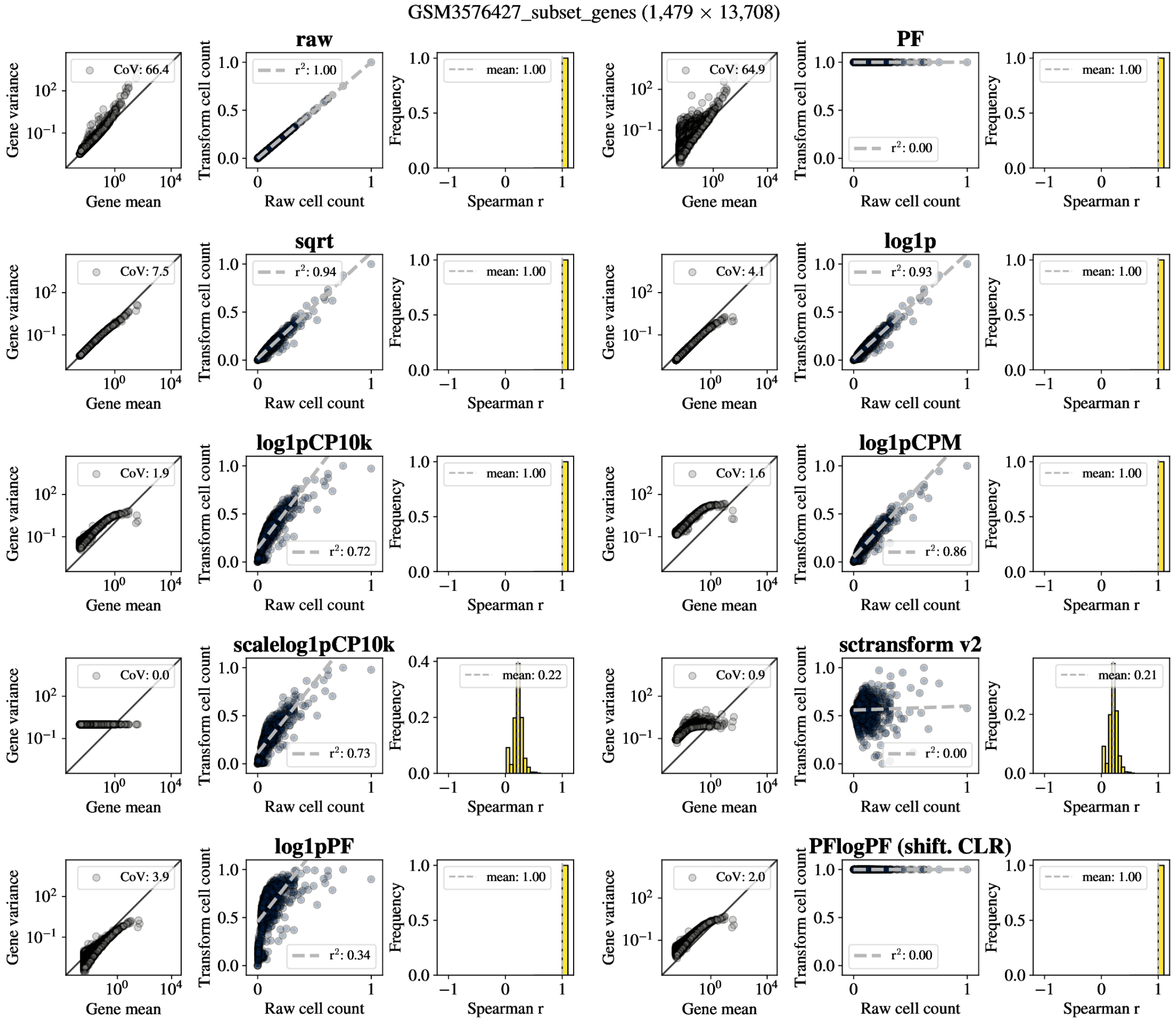


Supplementary Figure 1.332. **GSM3576427**. Per-method comparison of variance stabilization (left), depth normalization (middle), and monotonicity (right) for each normalization on dataset GSM3576427.

| GSM3576428 | |
|---|---|
| cells | 11,343 |
| genes | 60,623 |
| nonzero entries | 1,988,062 |
| density | 0.00289 |
| total counts | 3,114,455 |
| mean counts / cell | 275 |
| mean counts / gene | 51 |
| min counts / cell | 1 |
| max counts / cell | 5,417 |
| overdispersion | 2.55 |

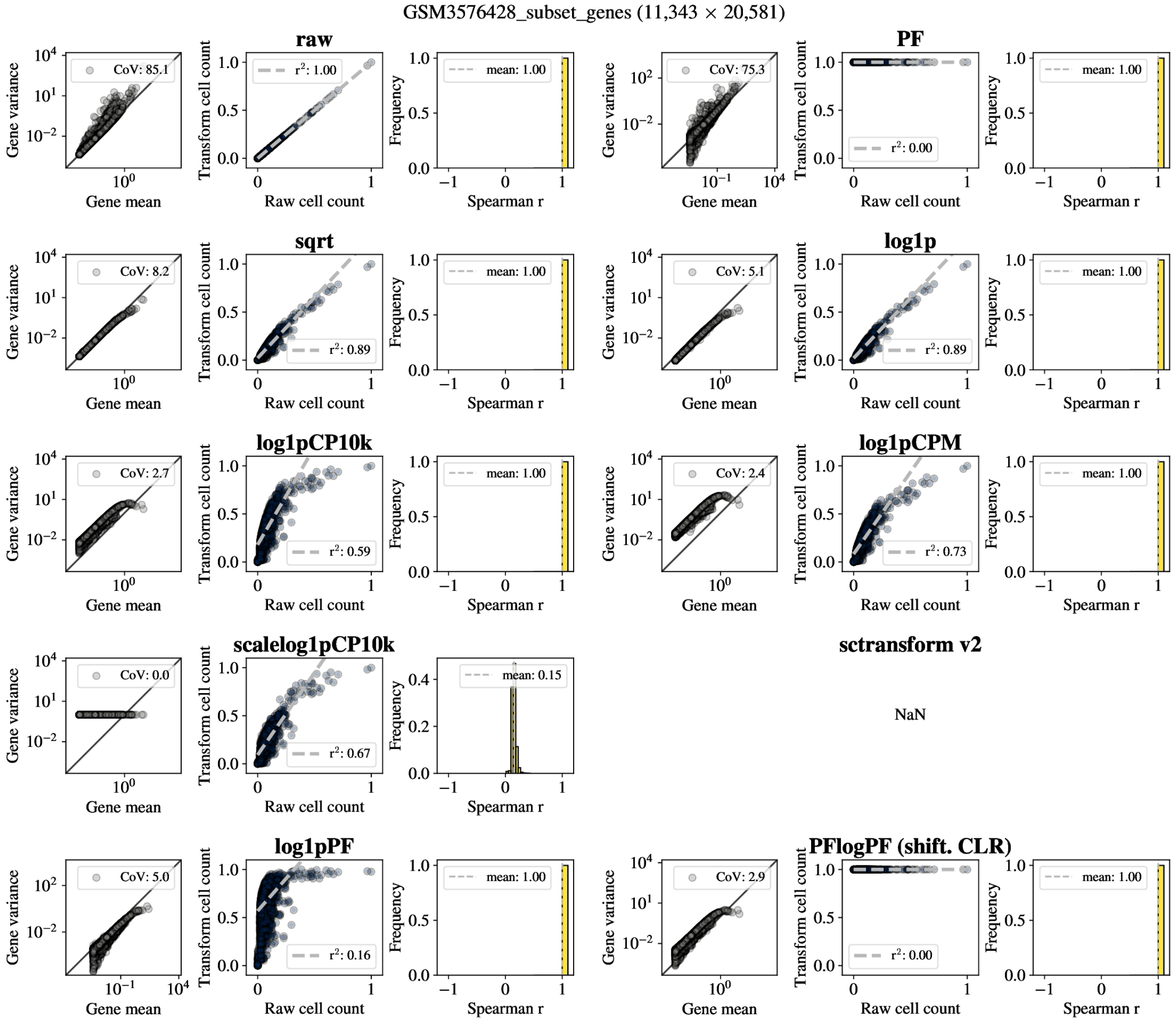


Supplementary Figure 1.333. **GSM3576428**. Per-method comparison of variance stabilization (left), depth normalization (middle), and monotonicity (right) for each normalization on dataset GSM3576428.

| GSM3576429 | |
|---|---|
| cells | 10,514 |
| genes | 60,623 |
| nonzero entries | 1,496,295 |
| density | 0.00235 |
| total counts | 2,318,164 |
| mean counts / cell | 220 |
| mean counts / gene | 38 |
| min counts / cell | 1 |
| max counts / cell | 3,965 |
| overdispersion | 3.05 |

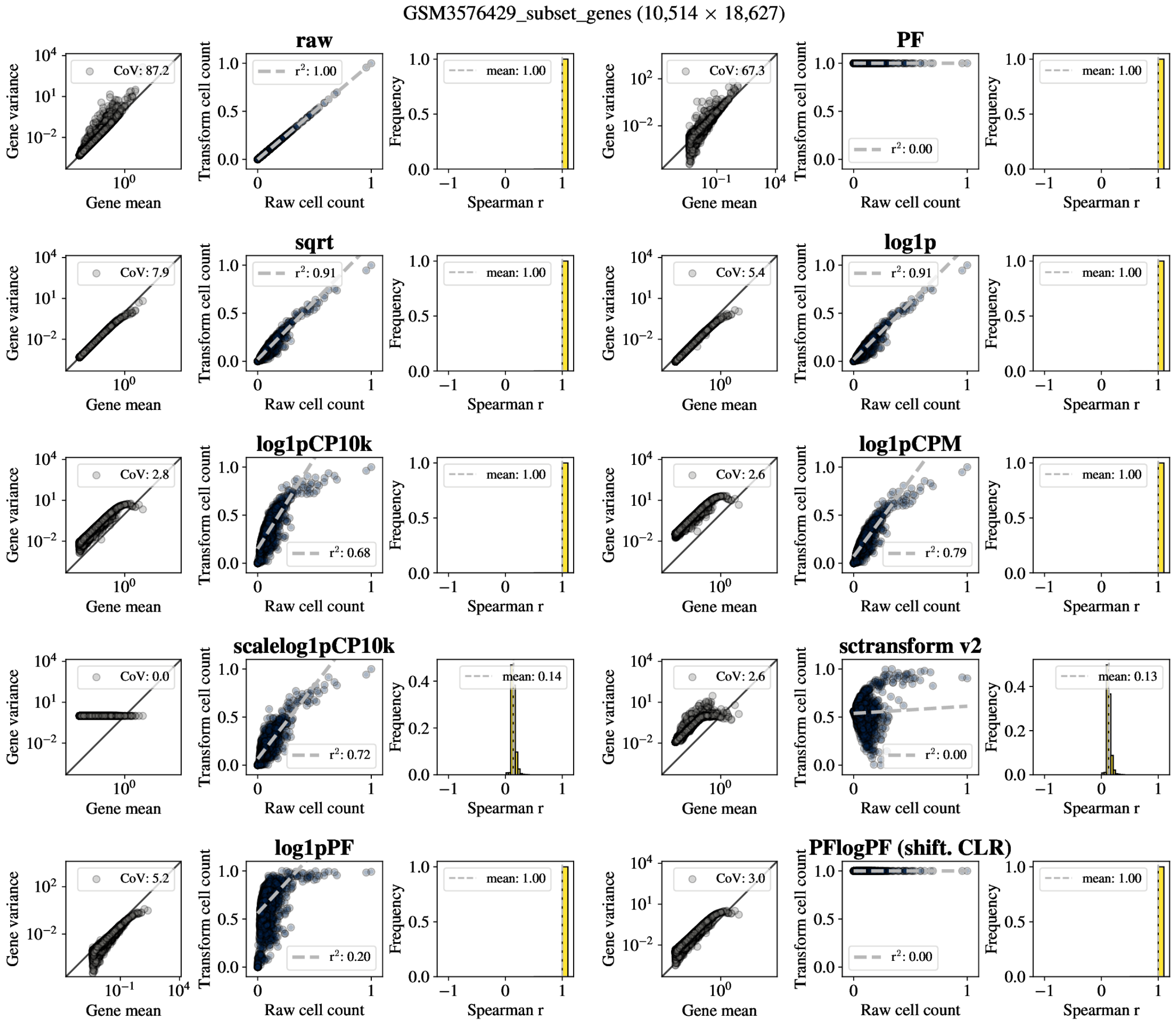


Supplementary Figure 1.334. **GSM3576429**. Per-method comparison of variance stabilization (left), depth normalization (middle), and monotonicity (right) for each normalization on dataset GSM3576429.

| **GSM3576430** | |
|---|---|
| cells | 8,977 |
| genes | 60,623 |
| nonzero entries | 1,335,170 |
| density | 0.00245 |
| total counts | 2,293,199 |
| mean counts / cell | 255 |
| mean counts / gene | 38 |
| min counts / cell | 0 |
| max counts / cell | 6,329 |
| overdispersion | 4.88 |

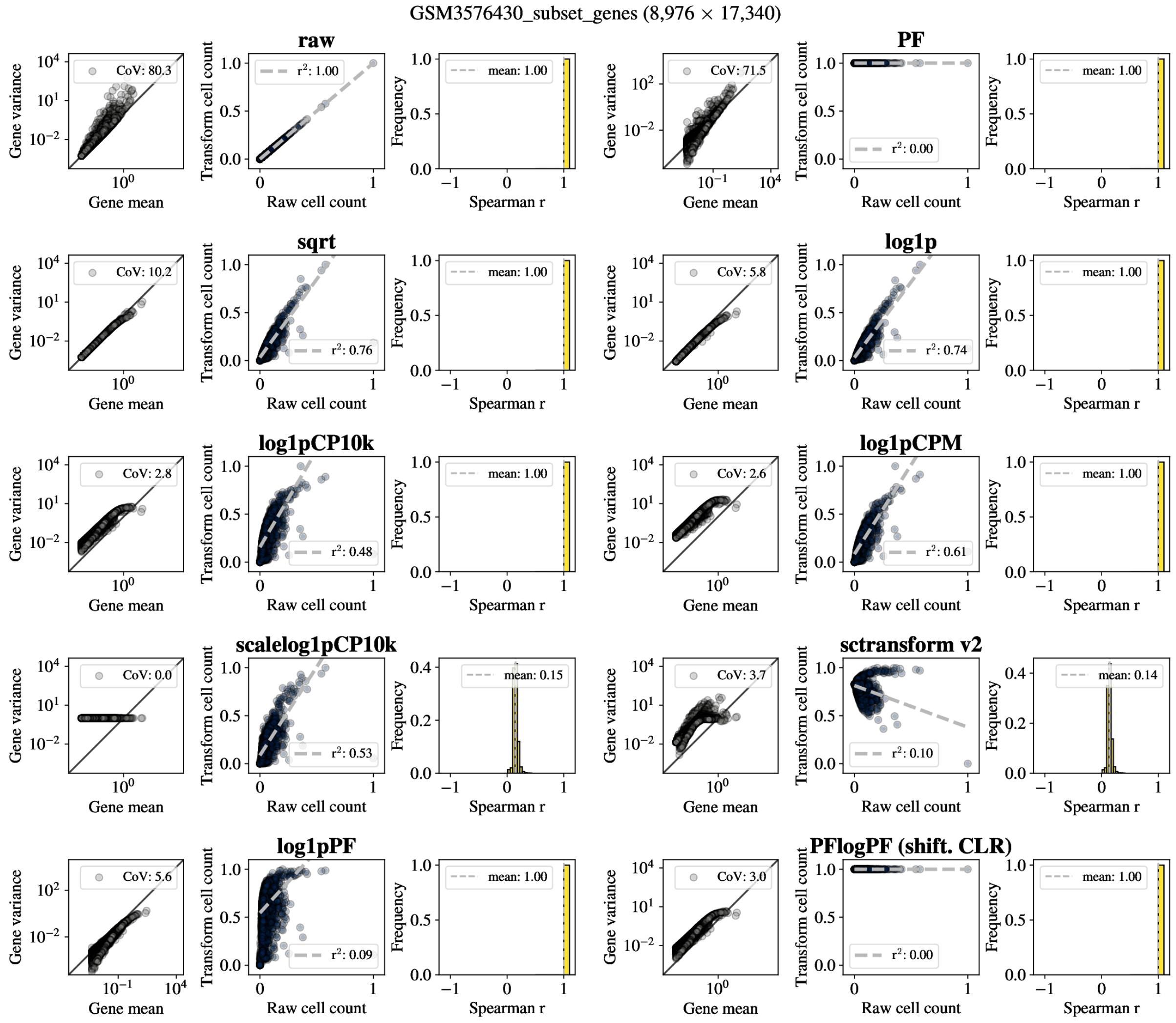


Supplementary Figure 1.335. **GSM3576430**. Per-method comparison of variance stabilization (left), depth normalization (middle), and monotonicity (right) for each normalization on dataset GSM3576430.

| GSM3587008 | |
|---|---|
| cells | 5,555 |
| genes | 60,623 |
| nonzero entries | 8,254,813 |
| density | 0.0245 |
| total counts | 32,971,581 |
| mean counts / cell | 5,935 |
| mean counts / gene | 544 |
| min counts / cell | 98 |
| max counts / cell | 69,833 |
| overdispersion | 1.65 |

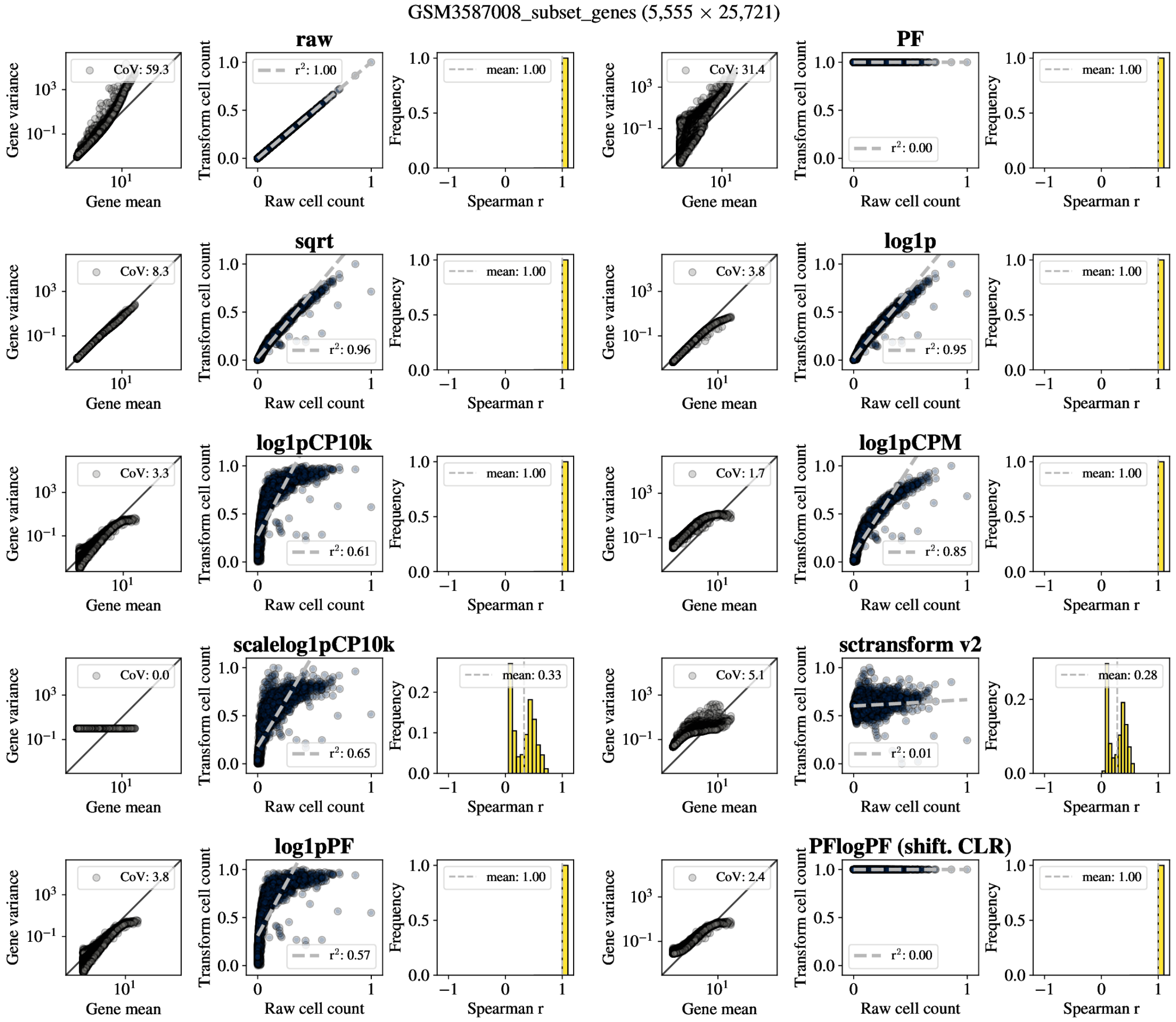


Supplementary Figure 1.336. **GSM3587008**. Per-method comparison of variance stabilization (left), depth normalization (middle), and monotonicity (right) for each normalization on dataset GSM3587008.

| GSM3587009 | |
|---|---|
| cells | 12,781 |
| genes | 60,623 |
| nonzero entries | 11,391,964 |
| density | 0.0147 |
| total counts | 38,703,617 |
| mean counts / cell | 3,028 |
| mean counts / gene | 638 |
| min counts / cell | 200 |
| max counts / cell | 97,989 |
| overdispersion | 2.2 |

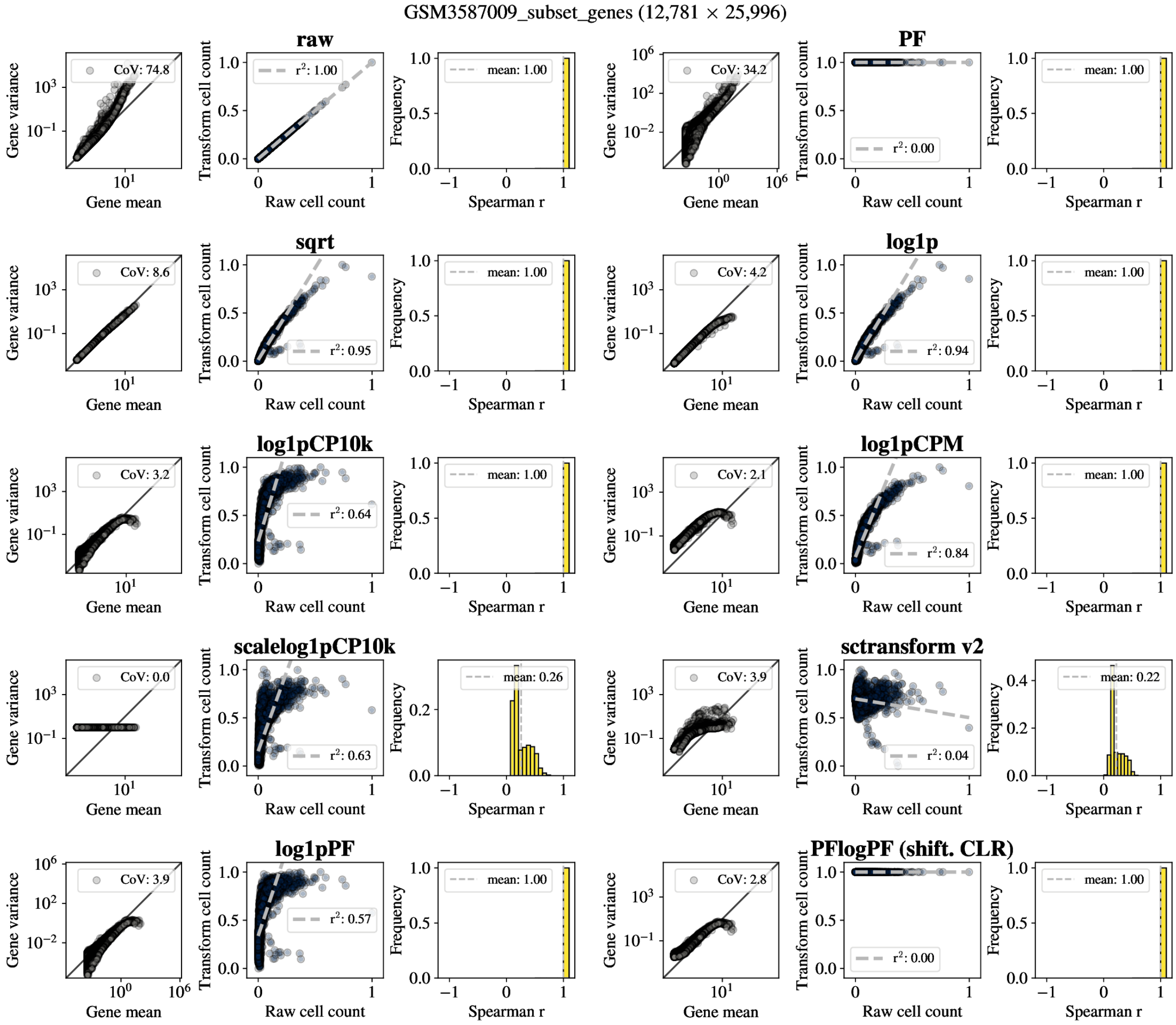


Supplementary Figure 1.337. **GSM3587009**. Per-method comparison of variance stabilization (left), depth normalization (middle), and monotonicity (right) for each normalization on dataset GSM3587009.

| **GSM3587010** | |
|---|---|
| cells | 5,604 |
| genes | 60,623 |
| nonzero entries | 7,292,195 |
| density | 0.0215 |
| total counts | 38,146,556 |
| mean counts / cell | 6,807 |
| mean counts / gene | 629 |
| min counts / cell | 75 |
| max counts / cell | 79,065 |
| overdispersion | 1.77 |

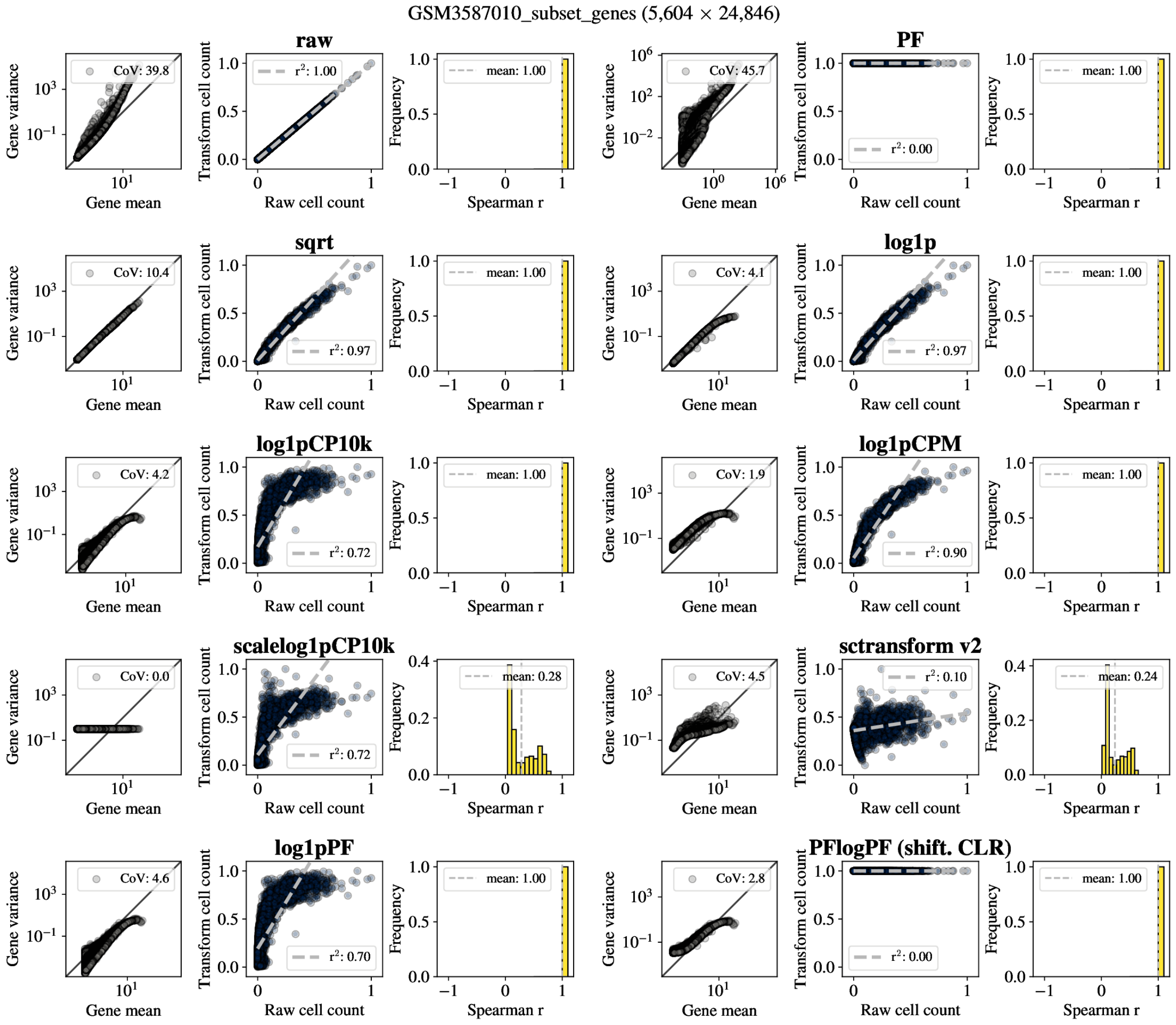


Supplementary Figure 1.338. **GSM3587010**. Per-method comparison of variance stabilization (left), depth normalization (middle), and monotonicity (right) for each normalization on dataset GSM3587010.

| GSM3587011 | |
|---|---|
| cells | 8,903 |
| genes | 60,623 |
| nonzero entries | 10,506,990 |
| density | 0.0195 |
| total counts | 46,189,682 |
| mean counts / cell | 5,188 |
| mean counts / gene | 762 |
| min counts / cell | 160 |
| max counts / cell | 90,069 |
| overdispersion | 1.73 |

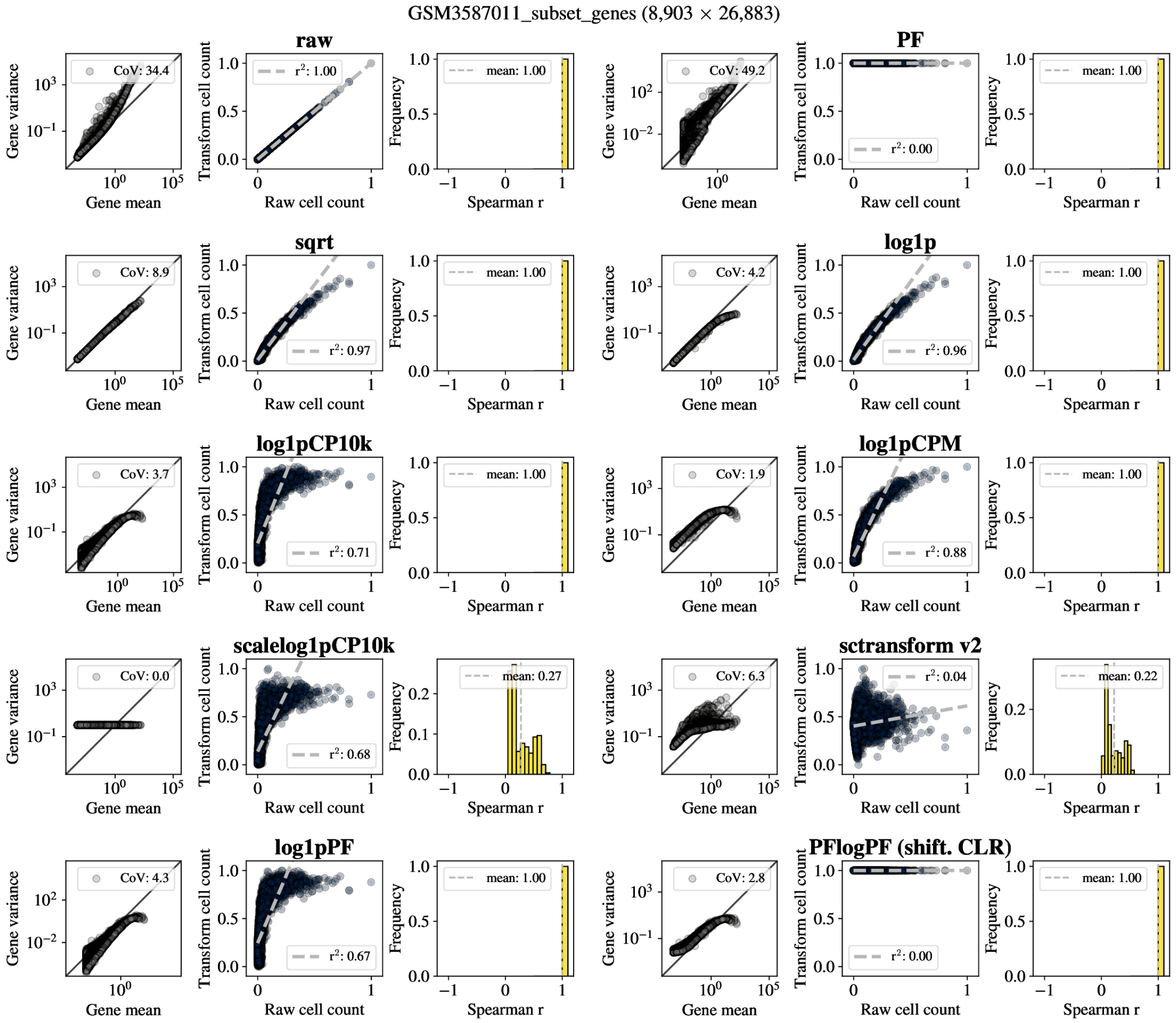


Supplementary Figure 1.339. **GSM3587011**. Per-method comparison of variance stabilization (left), depth normalization (middle), and monotonicity (right) for each normalization on dataset GSM3587011.

| GSM3587013 | |
|---|---|
| cells | 6,749 |
| genes | 60,623 |
| nonzero entries | 4,187,832 |
| density | 0.0102 |
| total counts | 22,496,143 |
| mean counts / cell | 3,333 |
| mean counts / gene | 371 |
| min counts / cell | 144 |
| max counts / cell | 94,552 |
| overdispersion | 0.925 |

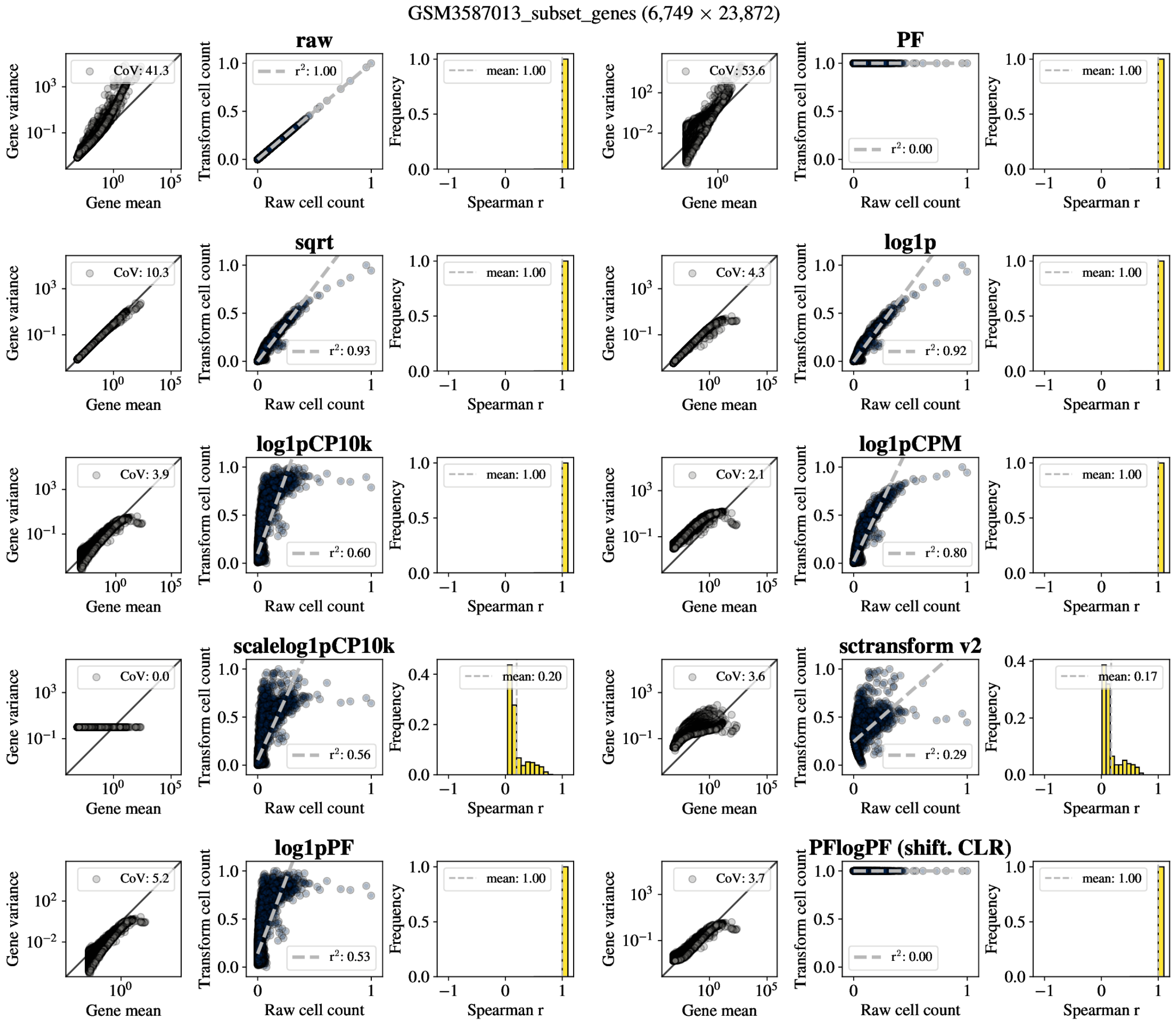


Supplementary Figure 1.340. **GSM3587013**. Per-method comparison of variance stabilization (left), depth normalization (middle), and monotonicity (right) for each normalization on dataset GSM3587013.

| GSM3665016 | |
|---|---|
| cells | 2,612 |
| genes | 60,623 |
| nonzero entries | 3,013,362 |
| density | 0.019 |
| total counts | 9,795,082 |
| mean counts / cell | 3,750 |
| mean counts / gene | 162 |
| min counts / cell | 51 |
| max counts / cell | 43,730 |
| overdispersion | 0.591 |

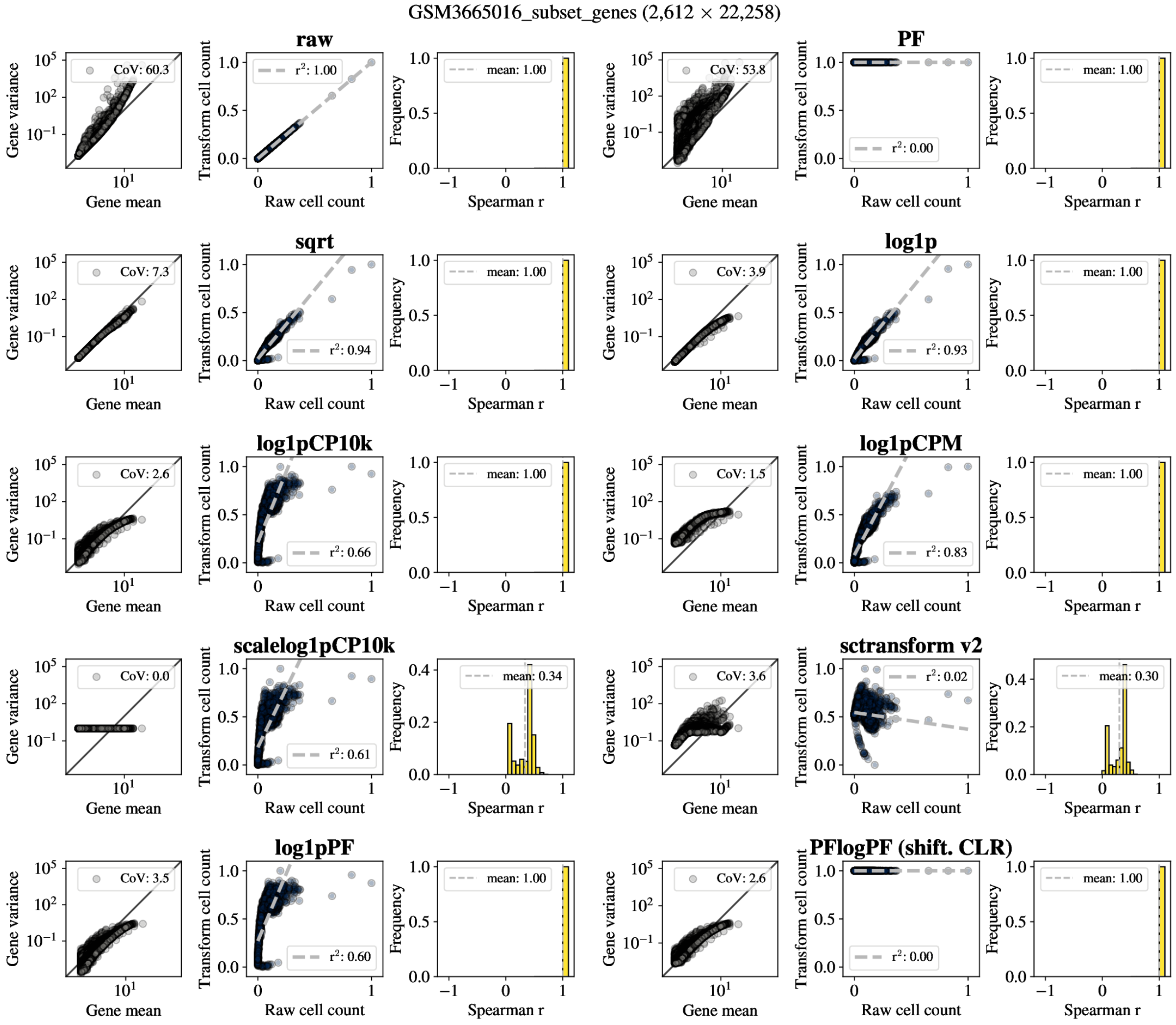


Supplementary Figure 1.341. **GSM3665016**. Per-method comparison of variance stabilization (left), depth normalization (middle), and monotonicity (right) for each normalization on dataset GSM3665016.

| GSM3665017 | |
|---|---|
| cells | 2,028 |
| genes | 60,623 |
| nonzero entries | 3,372,905 |
| density | 0.0274 |
| total counts | 11,361,476 |
| mean counts / cell | 5,602 |
| mean counts / gene | 187 |
| min counts / cell | 2,833 |
| max counts / cell | 18,056 |
| overdispersion | 0.175 |

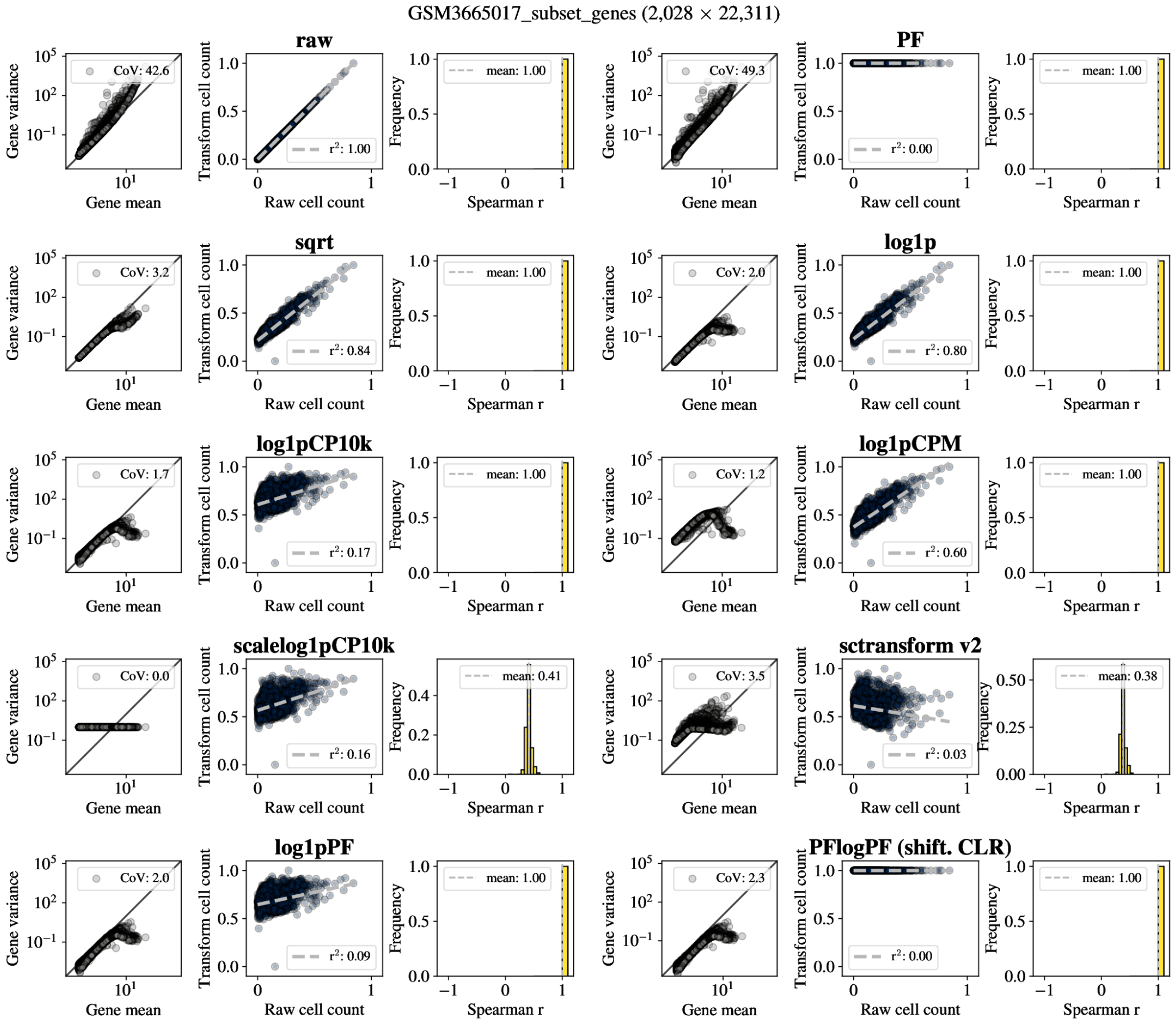


Supplementary Figure 1.342. **GSM3665017**. Per-method comparison of variance stabilization (left), depth normalization (middle), and monotonicity (right) for each normalization on dataset GSM3665017.

| GSM3665018 | |
|---|---|
| cells | 2,122 |
| genes | 60,623 |
| nonzero entries | 3,167,412 |
| density | 0.0246 |
| total counts | 9,466,877 |
| mean counts / cell | 4,461 |
| mean counts / gene | 156 |
| min counts / cell | 2,533 |
| max counts / cell | 43,670 |
| overdispersion | 0.115 |

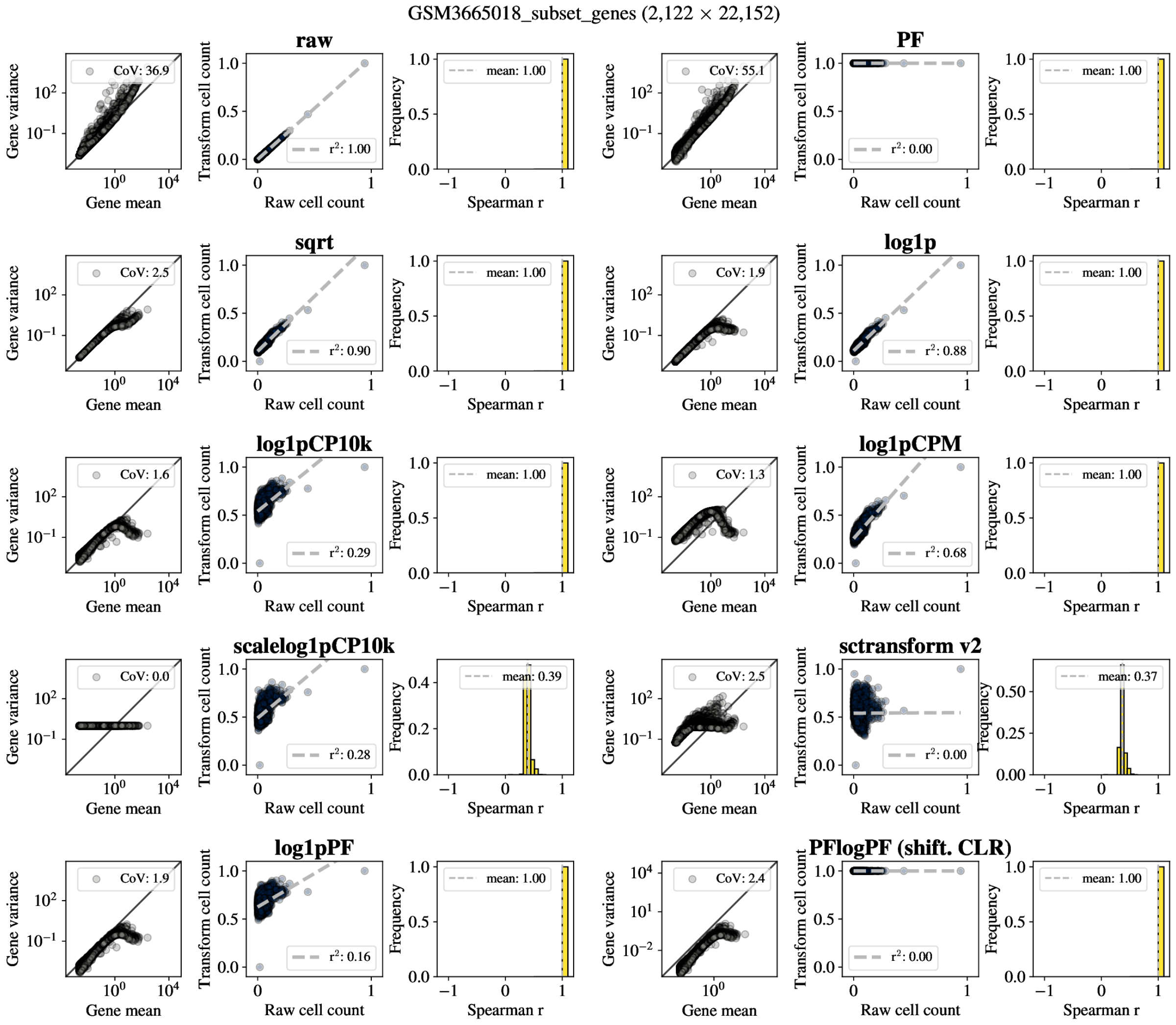


Supplementary Figure 1.343. **GSM3665018**. Per-method comparison of variance stabilization (left), depth normalization (middle), and monotonicity (right) for each normalization on dataset GSM3665018.

| GSM3665019 | |
|---|---|
| cells | 3,189 |
| genes | 60,623 |
| nonzero entries | 3,534,798 |
| density | 0.0183 |
| total counts | 9,842,002 |
| mean counts / cell | 3,086 |
| mean counts / gene | 162 |
| min counts / cell | 76 |
| max counts / cell | 24,847 |
| overdispersion | 0.444 |

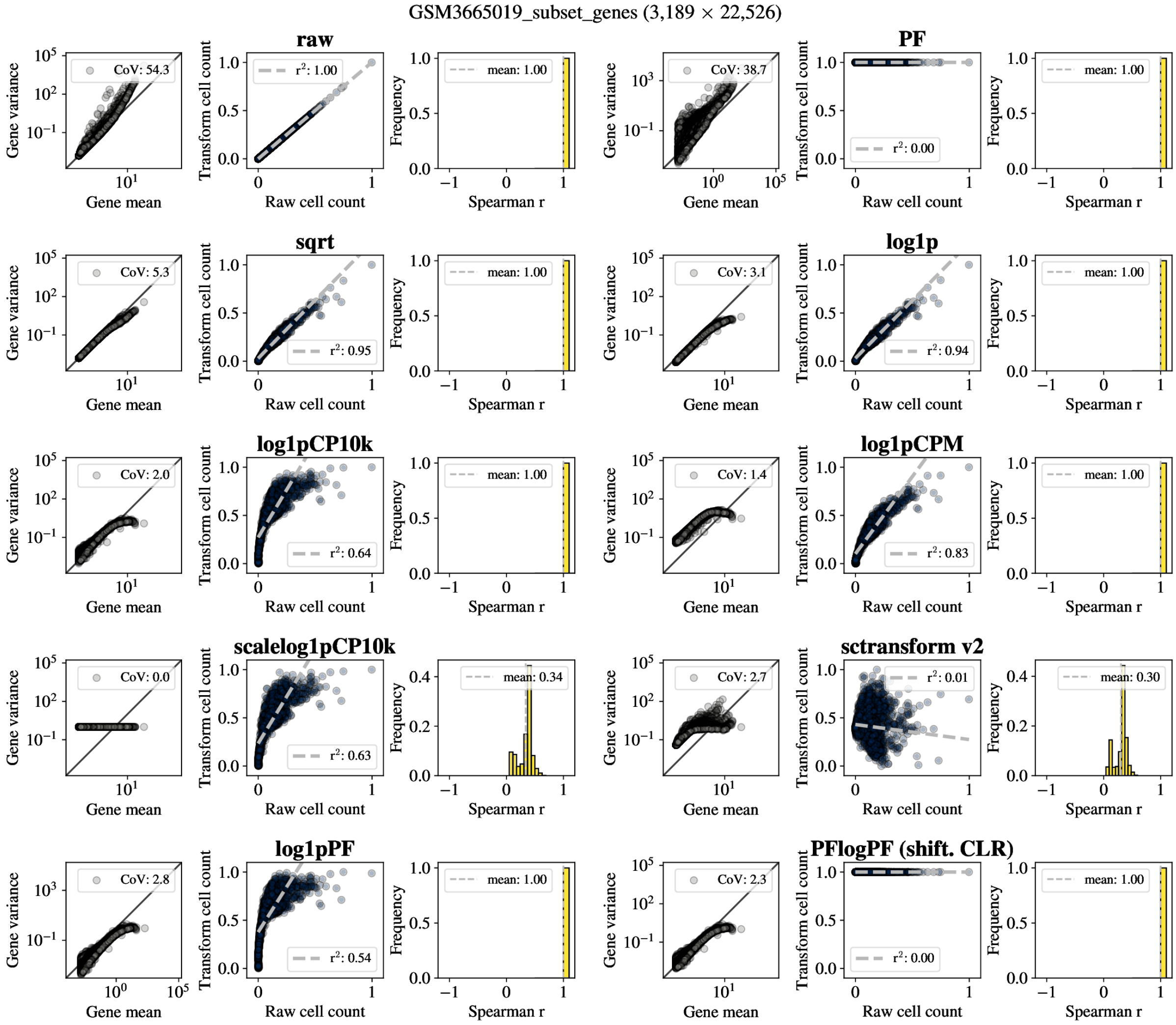


Supplementary Figure 1.344. **GSM3665019**. Per-method comparison of variance stabilization (left), depth normalization (middle), and monotonicity (right) for each normalization on dataset GSM3665019.

| GSM3667327 | |
|---|---|
| cells | 6,435 |
| genes | 60,623 |
| nonzero entries | 3,607,623 |
| density | 0.00925 |
| total counts | 12,024,531 |
| mean counts / cell | 1,869 |
| mean counts / gene | 198 |
| min counts / cell | 82 |
| max counts / cell | 144,797 |
| overdispersion | 6 |

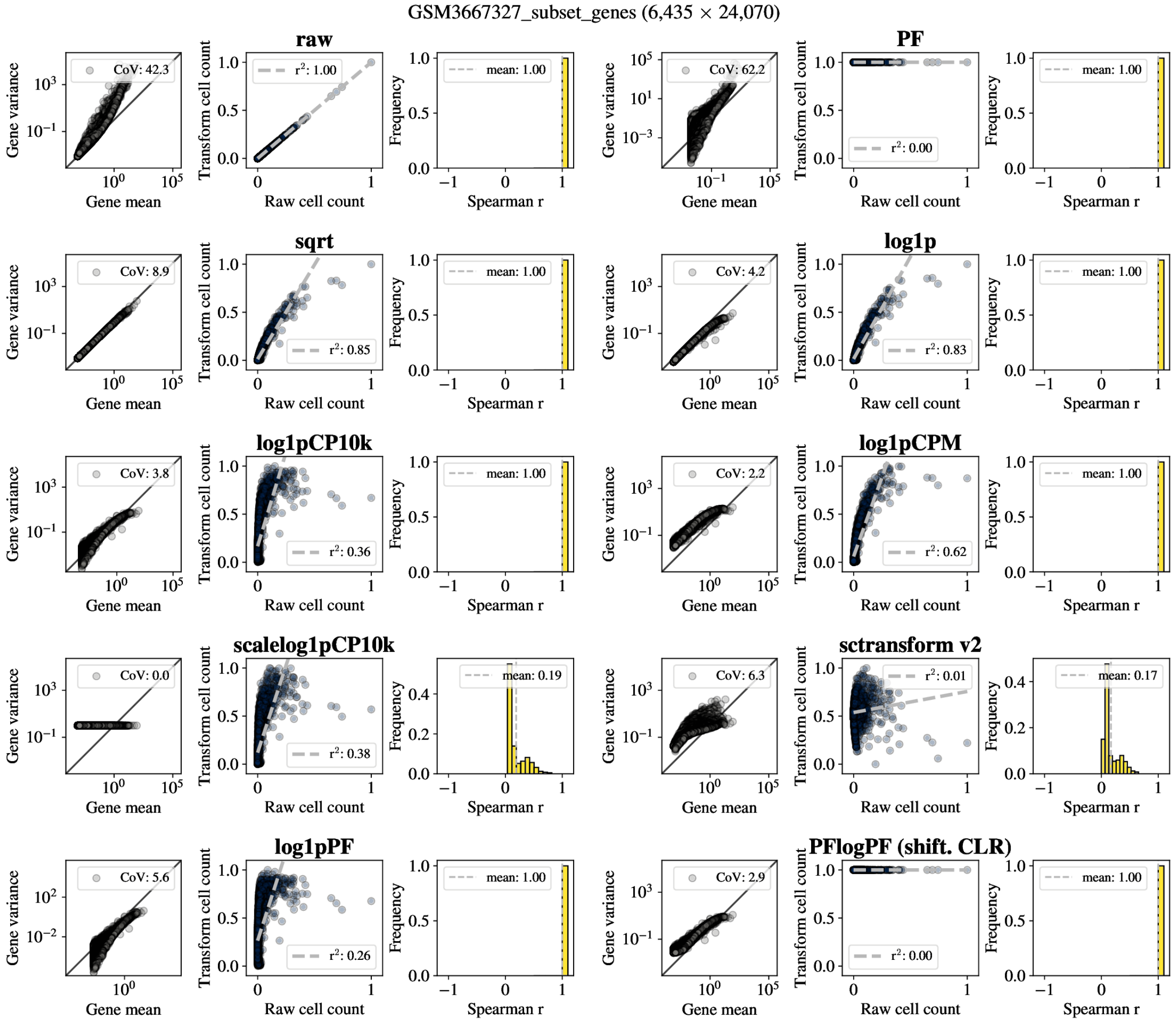


Supplementary Figure 1.345. **GSM3667327**. Per-method comparison of variance stabilization (left), depth normalization (middle), and monotonicity (right) for each normalization on dataset GSM3667327.

| GSM3667328 | |
|---|---|
| cells | 16,108 |
| genes | 60,623 |
| nonzero entries | 8,637,546 |
| density | 0.00885 |
| total counts | 33,042,035 |
| mean counts / cell | 2,051 |
| mean counts / gene | 545 |
| min counts / cell | 99 |
| max counts / cell | 127,000 |
| overdispersion | 4.69 |

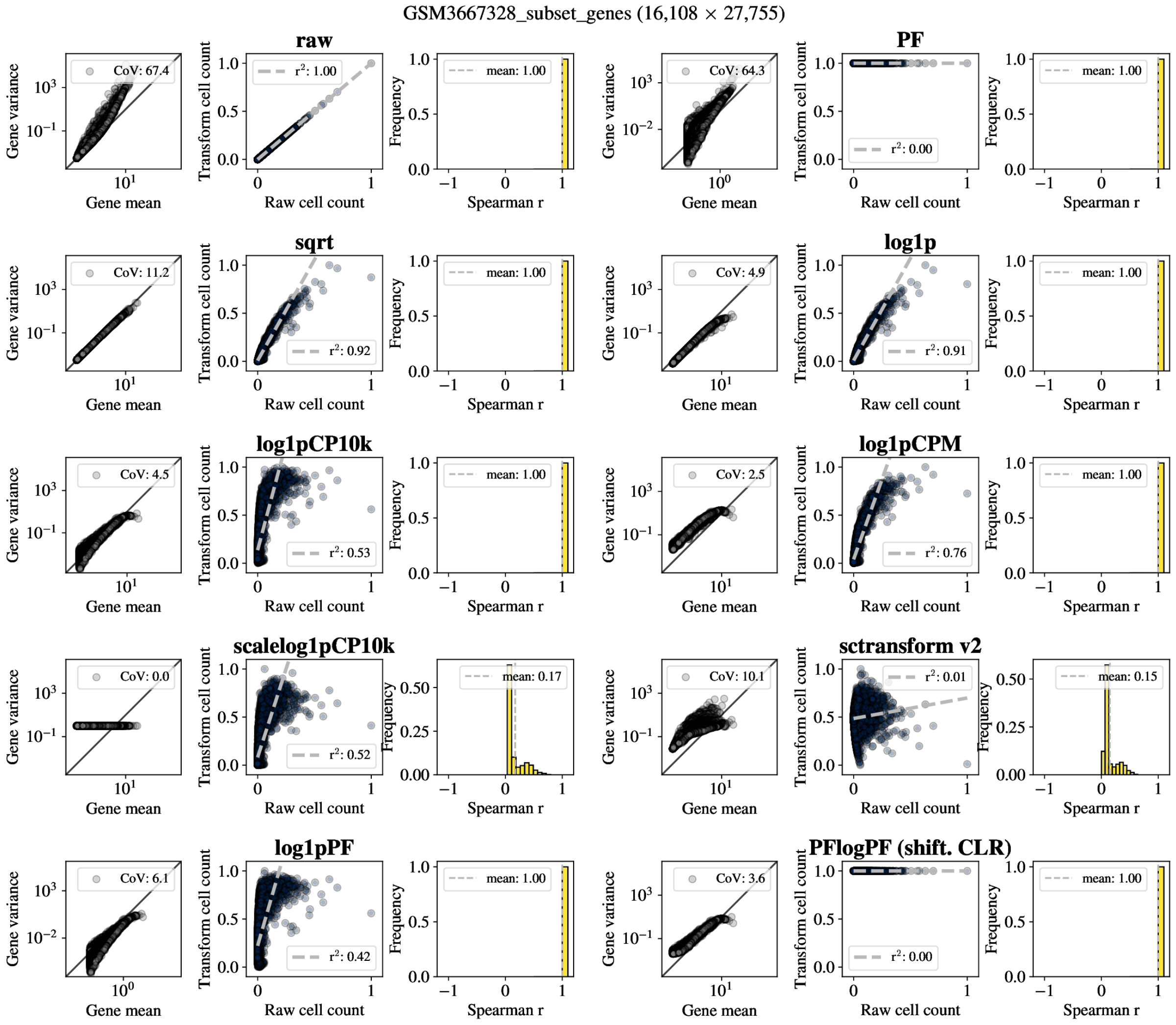


Supplementary Figure 1.346. **GSM3667328**. Per-method comparison of variance stabilization (left), depth normalization (middle), and monotonicity (right) for each normalization on dataset GSM3667328.

| GSM3667329 | |
|---|---|
| cells | 8,474 |
| genes | 60,623 |
| nonzero entries | 6,733,047 |
| density | 0.0131 |
| total counts | 29,551,920 |
| mean counts / cell | 3,487 |
| mean counts / gene | 487 |
| min counts / cell | 121 |
| max counts / cell | 108,478 |
| overdispersion | 5.45 |

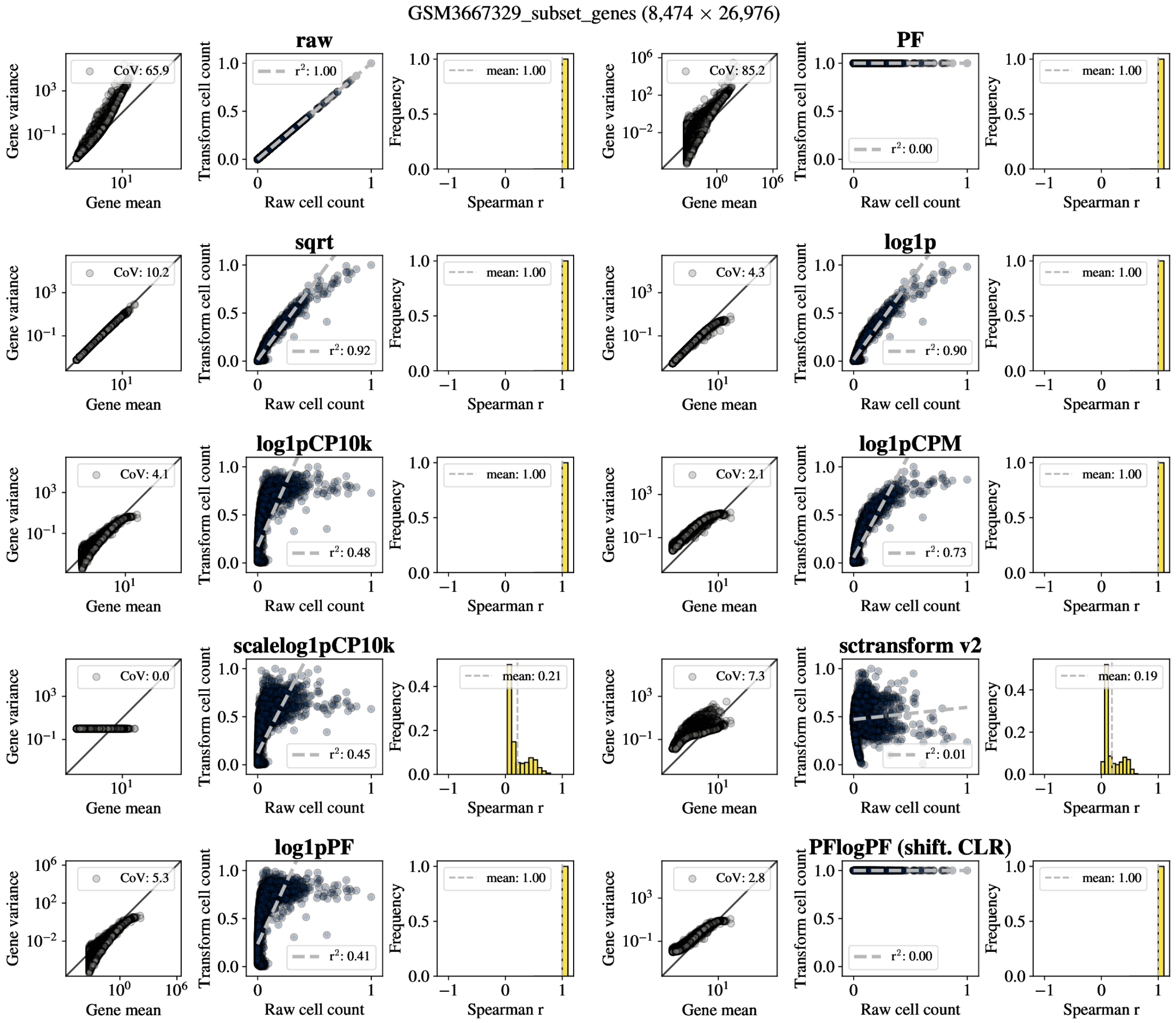


Supplementary Figure 1.347. **GSM3667329**. Per-method comparison of variance stabilization (left), depth normalization (middle), and monotonicity (right) for each normalization on dataset GSM3667329.

| GSM3711757 | |
|---|---|
| cells | 16,613 |
| genes | 60,623 |
| nonzero entries | 15,814,585 |
| density | 0.0157 |
| total counts | 41,236,247 |
| mean counts / cell | 2,482 |
| mean counts / gene | 680 |
| min counts / cell | 443 |
| max counts / cell | 95,446 |
| overdispersion | 0.589 |

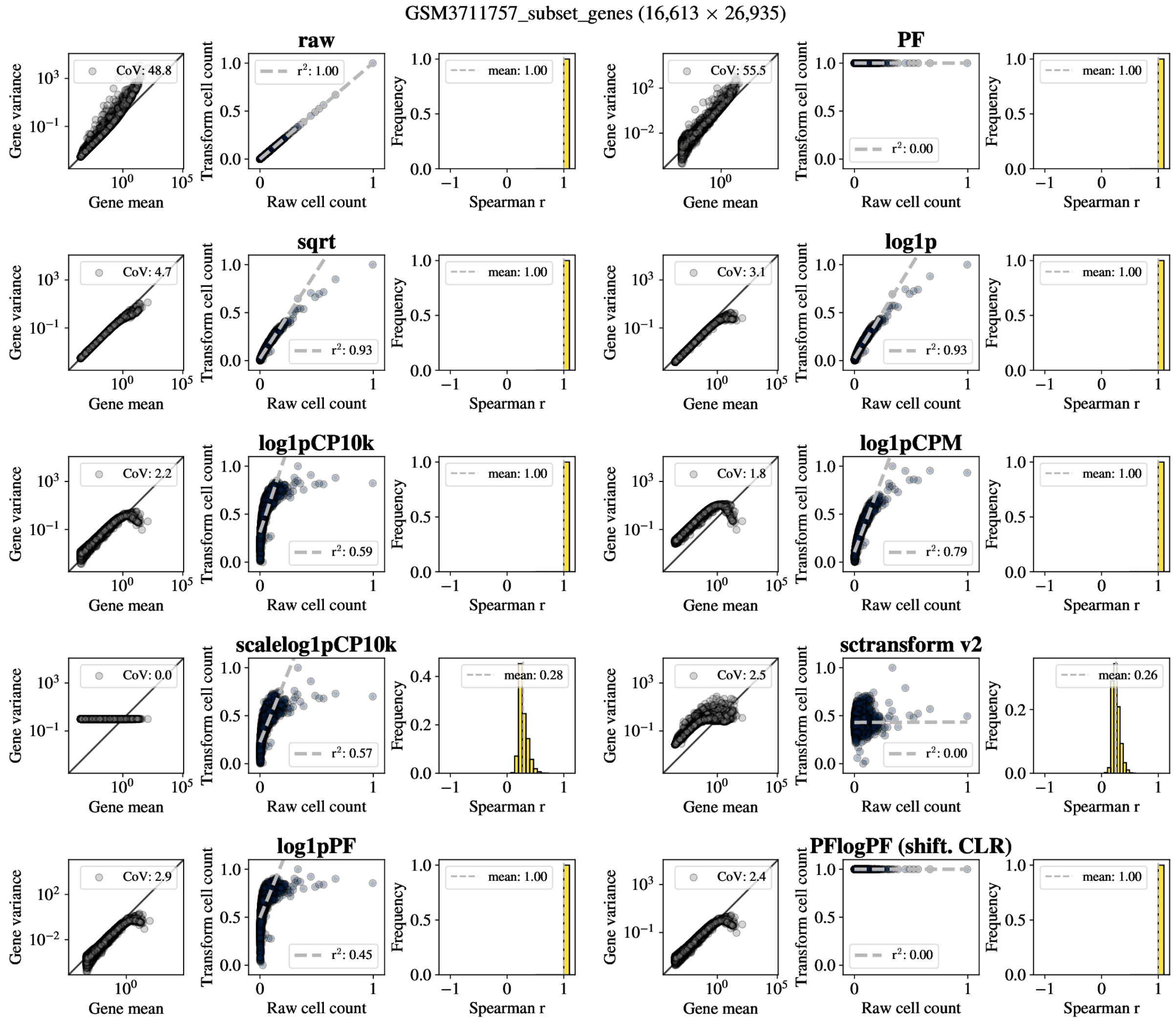


Supplementary Figure 1.348. **GSM3711757**. Per-method comparison of variance stabilization (left), depth normalization (middle), and monotonicity (right) for each normalization on dataset GSM3711757.

| GSM3711758 | |
|---|---|
| cells | 10,985 |
| genes | 60,623 |
| nonzero entries | 4,699,669 |
| density | 0.00706 |
| total counts | 12,881,429 |
| mean counts / cell | 1,173 |
| mean counts / gene | 212 |
| min counts / cell | 188 |
| max counts / cell | 28,784 |
| overdispersion | 2.52 |

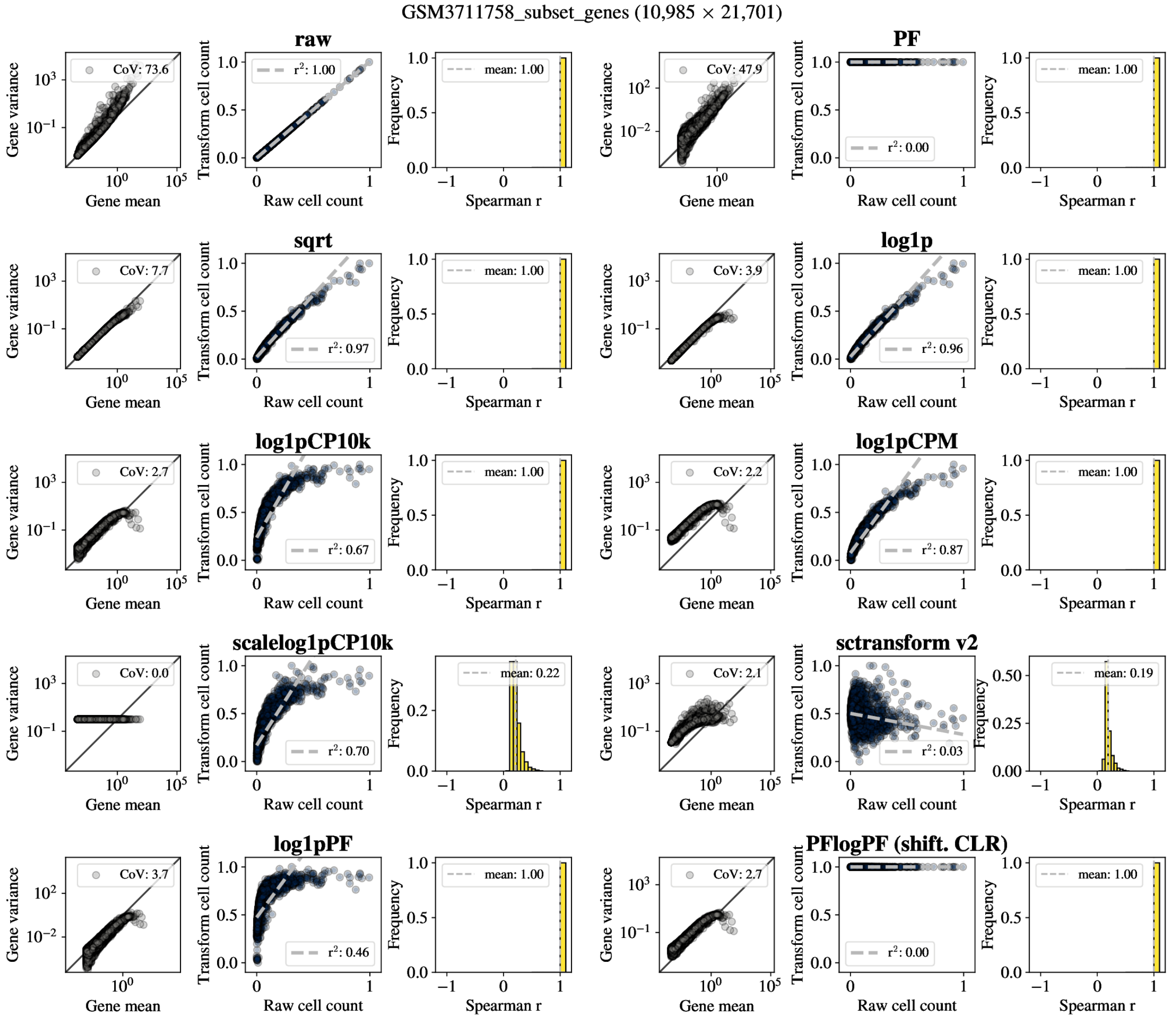


Supplementary Figure 1.349. **GSM3711758**. Per-method comparison of variance stabilization (left), depth normalization (middle), and monotonicity (right) for each normalization on dataset GSM3711758.

| GSM3711759 | |
|---|---|
| cells | 2,006 |
| genes | 60,623 |
| nonzero entries | 3,667,573 |
| density | 0.0302 |
| total counts | 14,076,770 |
| mean counts / cell | 7,017 |
| mean counts / gene | 232 |
| min counts / cell | 1,139 |
| max counts / cell | 140,197 |
| overdispersion | 5.44 |

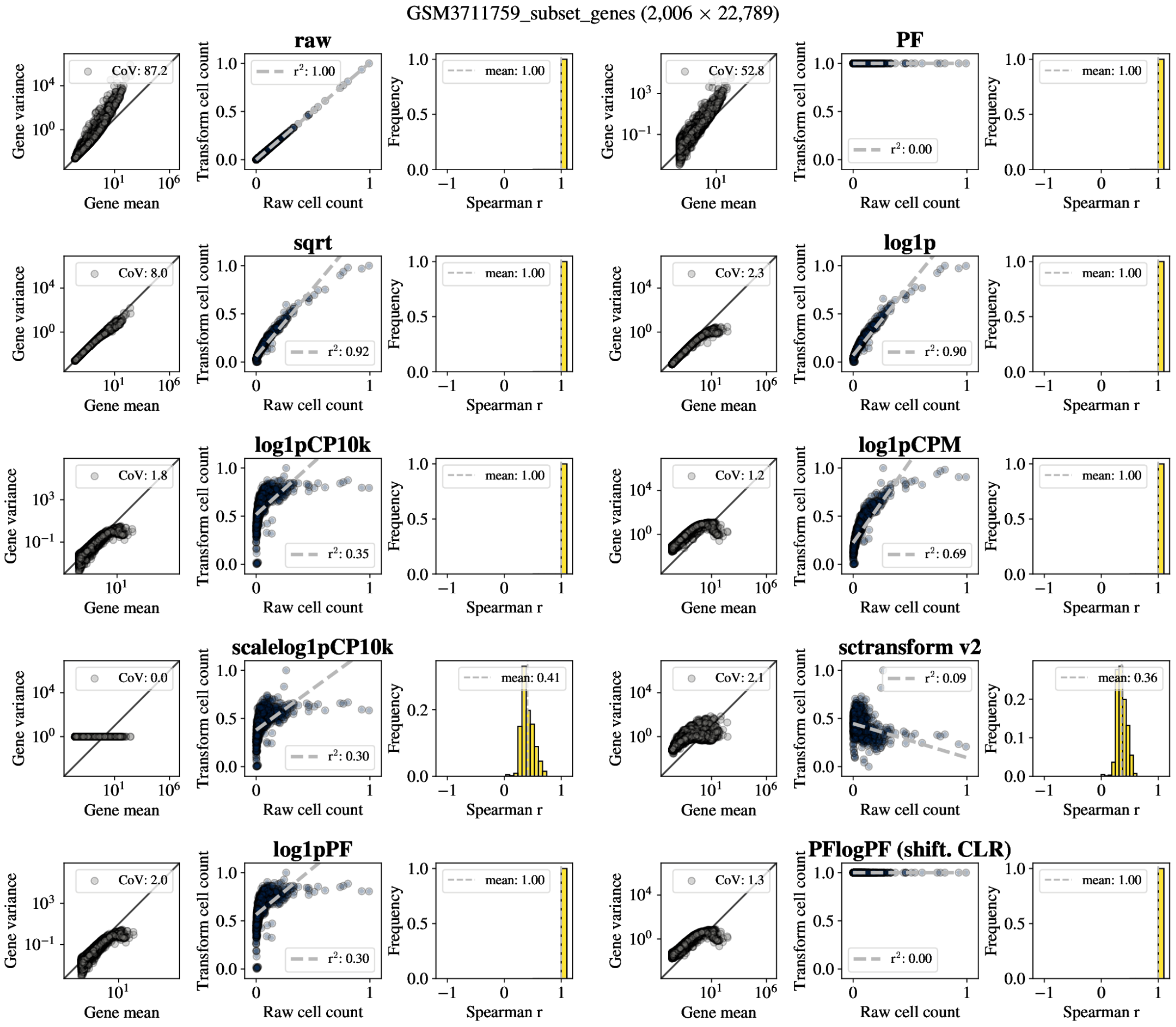


Supplementary Figure 1.350. **GSM3711759**. Per-method comparison of variance stabilization (left), depth normalization (middle), and monotonicity (right) for each normalization on dataset GSM3711759.

| GSM3711760 | |
|---|---|
| cells | 10,809 |
| genes | 60,623 |
| nonzero entries | 5,467,335 |
| density | 0.00834 |
| total counts | 12,835,327 |
| mean counts / cell | 1,187 |
| mean counts / gene | 212 |
| min counts / cell | 244 |
| max counts / cell | 29,334 |
| overdispersion | 1.14 |

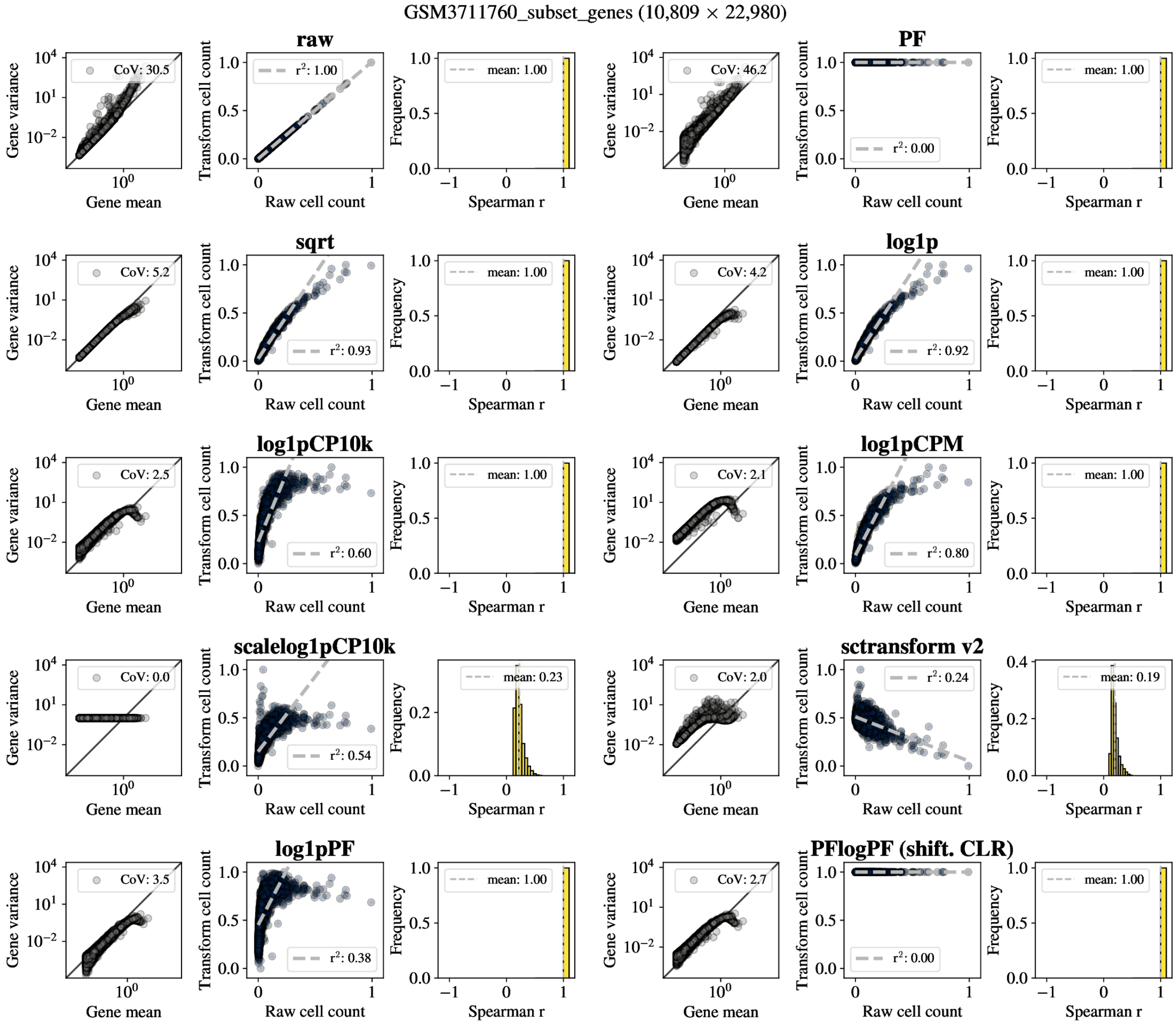


Supplementary Figure 1.351. **GSM3711760**. Per-method comparison of variance stabilization (left), depth normalization (middle), and monotonicity (right) for each normalization on dataset GSM3711760.

| GSM3711761 | |
|---|---|
| cells | 1,076 |
| genes | 60,623 |
| nonzero entries | 1,016,339 |
| density | 0.0156 |
| total counts | 3,454,448 |
| mean counts / cell | 3,210 |
| mean counts / gene | 57 |
| min counts / cell | 157 |
| max counts / cell | 72,238 |
| overdispersion | 2.33 |

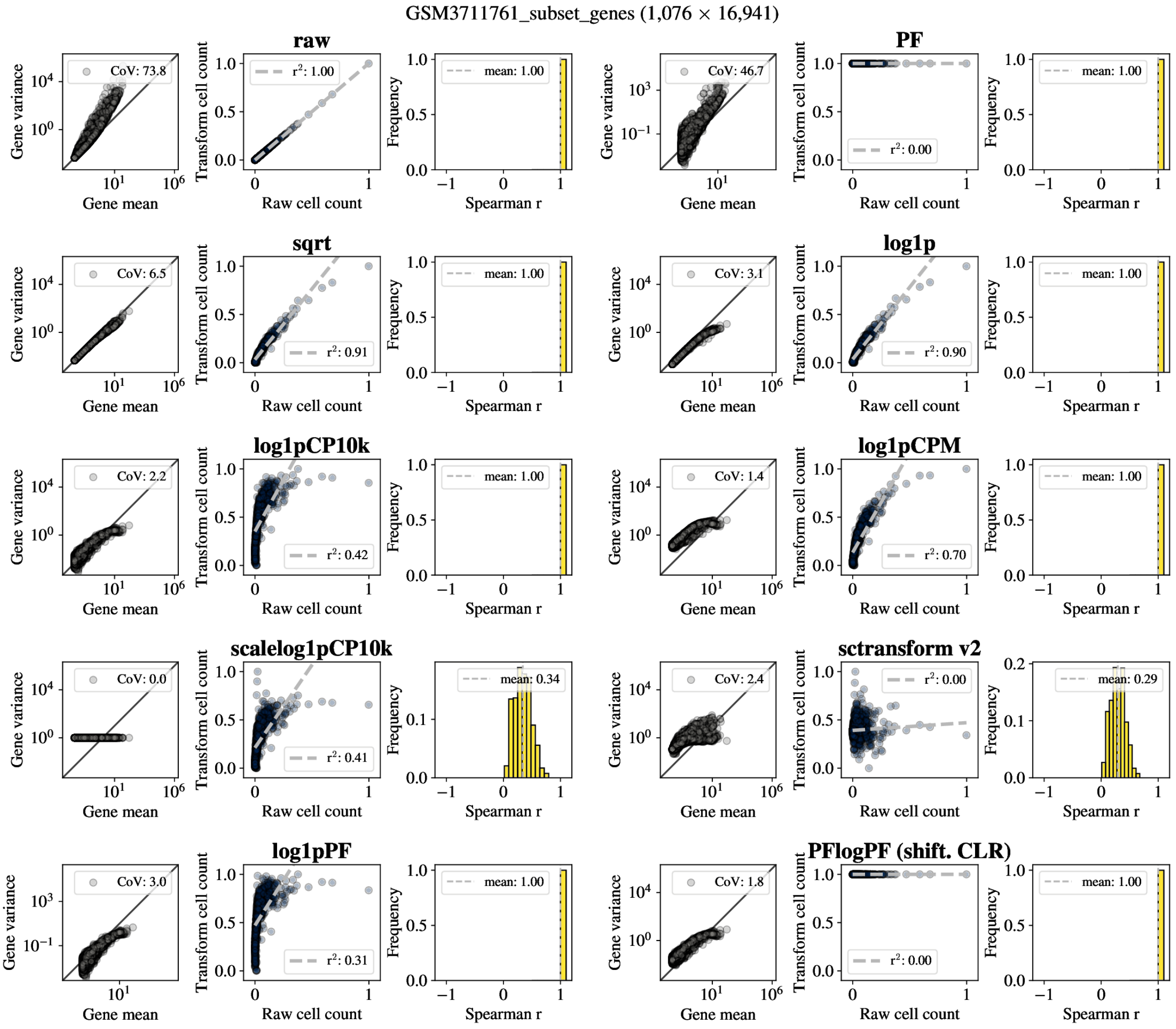


Supplementary Figure 1.352. **GSM3711761**. Per-method comparison of variance stabilization (left), depth normalization (middle), and monotonicity (right) for each normalization on dataset GSM3711761.

| GSM3711763 | |
|---|---|
| cells | 7,021 |
| genes | 60,623 |
| nonzero entries | 9,784,410 |
| density | 0.023 |
| total counts | 34,227,060 |
| mean counts / cell | 4,875 |
| mean counts / gene | 565 |
| min counts / cell | 823 |
| max counts / cell | 71,849 |
| overdispersion | 1.25 |

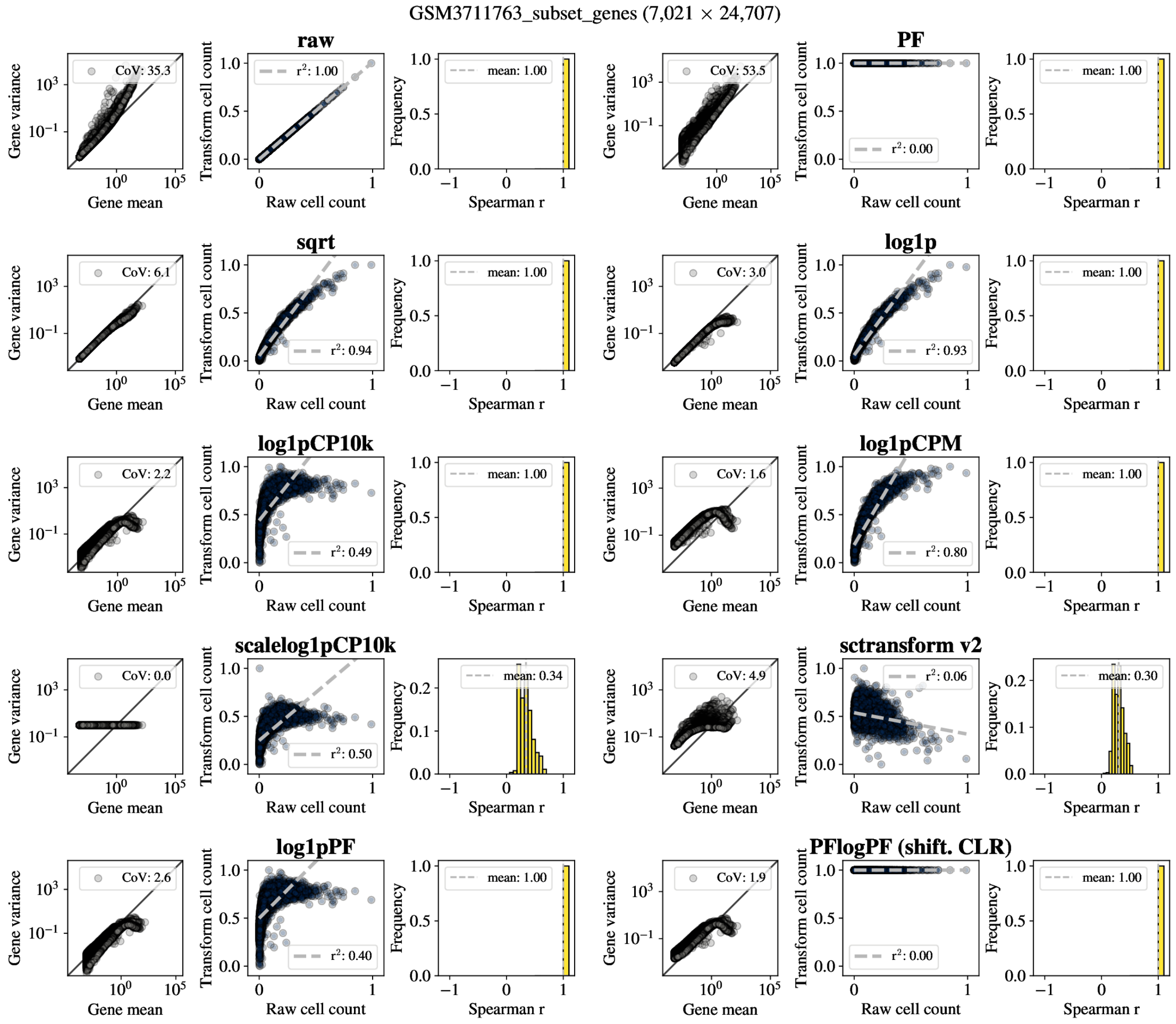


Supplementary Figure 1.353. **GSM3711763**. Per-method comparison of variance stabilization (left), depth normalization (middle), and monotonicity (right) for each normalization on dataset GSM3711763.

| GSM3711764 | |
|---|---|
| cells | 45,999 |
| genes | 60,623 |
| nonzero entries | 2,593,252 |
| density | 0.00093 |
| total counts | 3,231,785 |
| mean counts / cell | 70 |
| mean counts / gene | 53 |
| min counts / cell | 0 |
| max counts / cell | 13,954 |
| overdispersion | 3.18 |

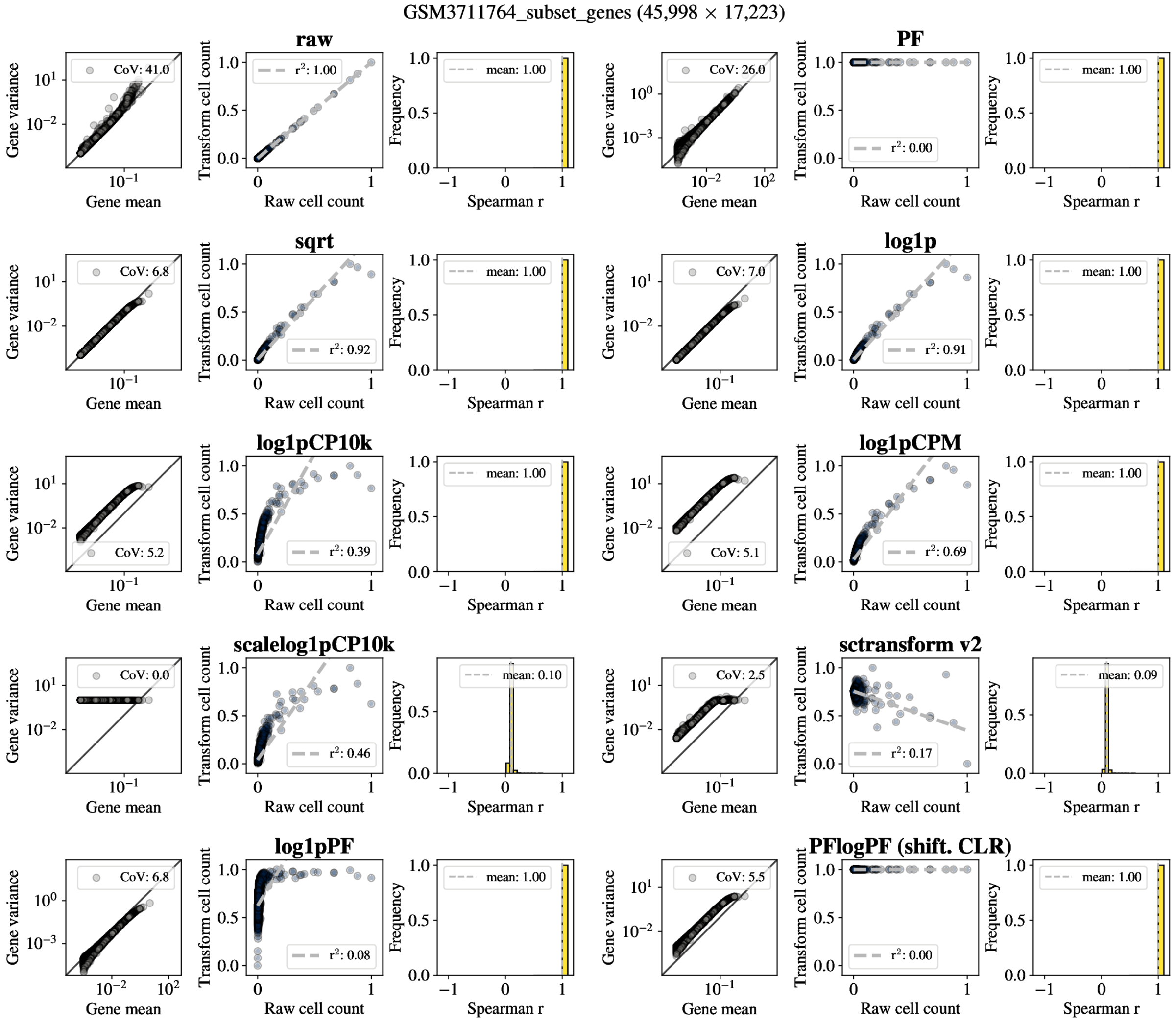


Supplementary Figure 1.354. **GSM3711764**. Per-method comparison of variance stabilization (left), depth normalization (middle), and monotonicity (right) for each normalization on dataset GSM3711764.

| GSM3711765 | |
|---|---|
| cells | 1,731 |
| genes | 60,623 |
| nonzero entries | 2,101,686 |
| density | 0.02 |
| total counts | 8,009,178 |
| mean counts / cell | 4,627 |
| mean counts / gene | 132 |
| min counts / cell | 275 |
| max counts / cell | 86,953 |
| overdispersion | 1.27 |

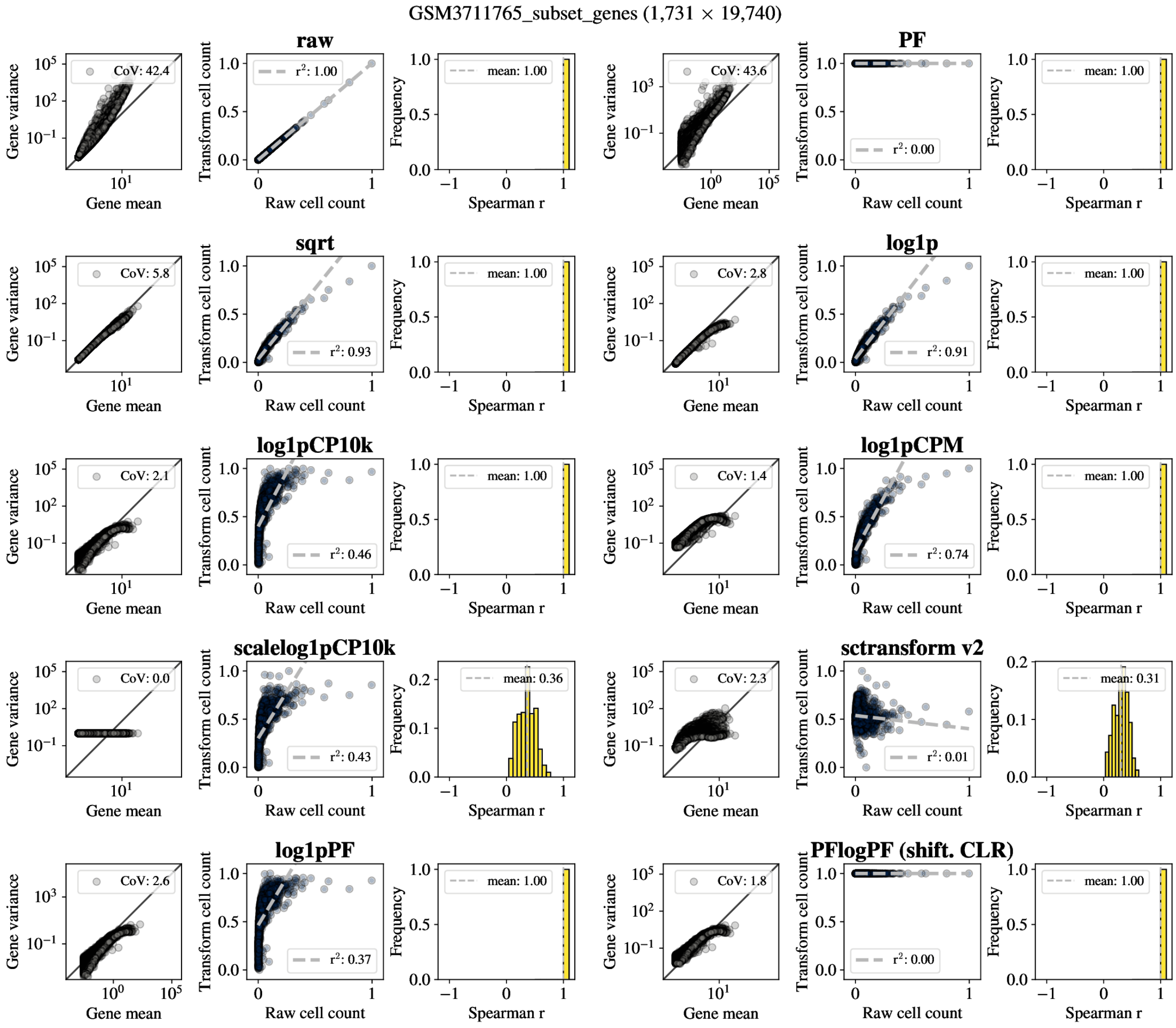


Supplementary Figure 1.355. **GSM3711765**. Per-method comparison of variance stabilization (left), depth normalization (middle), and monotonicity (right) for each normalization on dataset GSM3711765.

| GSM3711766 | |
|---|---|
| cells | 1,179 |
| genes | 60,623 |
| nonzero entries | 750,333 |
| density | 0.0105 |
| total counts | 1,920,999 |
| mean counts / cell | 1,629 |
| mean counts / gene | 32 |
| min counts / cell | 17 |
| max counts / cell | 37,824 |
| overdispersion | 11.8 |

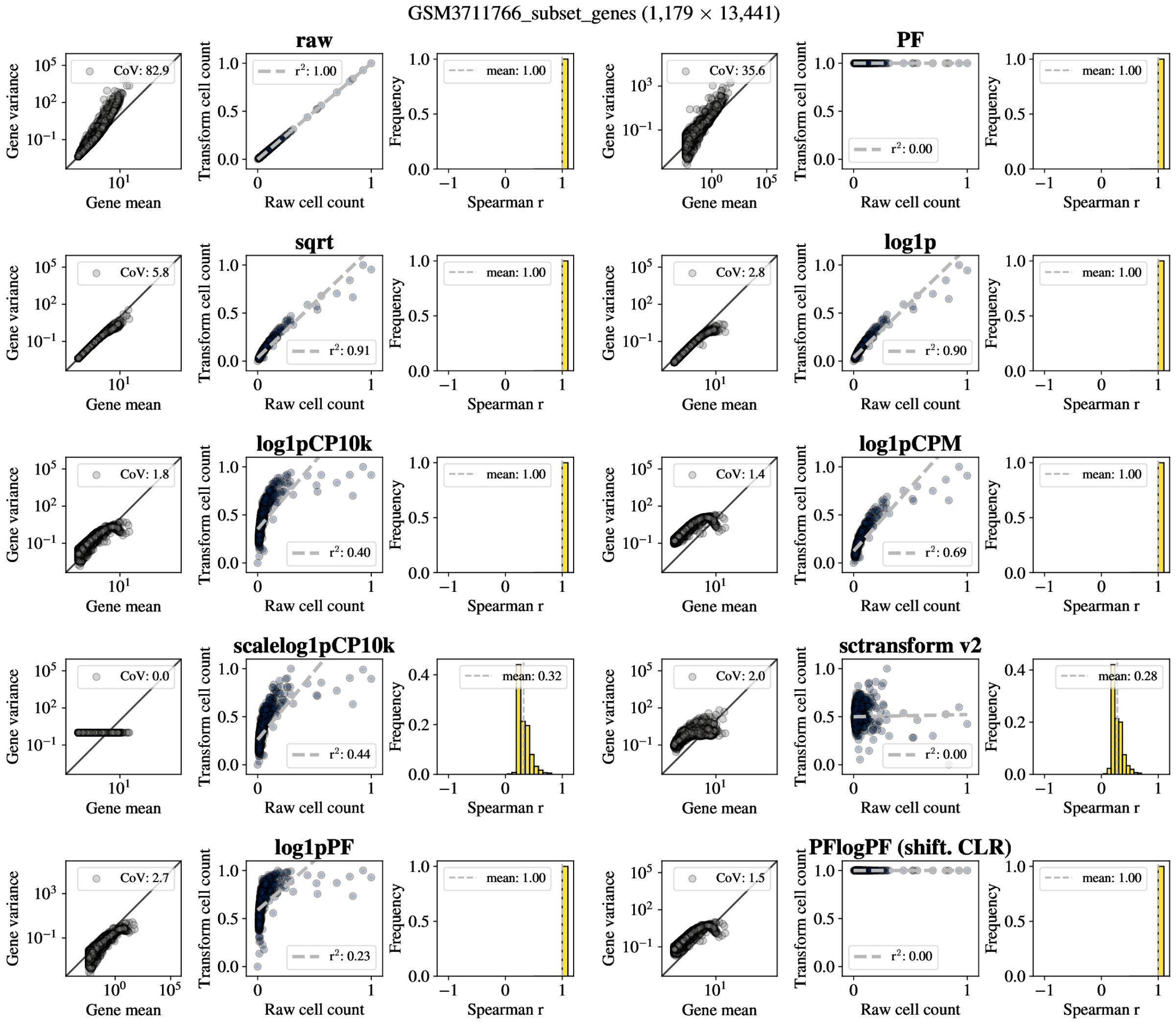


Supplementary Figure 1.356. **GSM3711766**. Per-method comparison of variance stabilization (left), depth normalization (middle), and monotonicity (right) for each normalization on dataset GSM3711766.

| GSM3711767 | |
|---|---|
| cells | 3,482 |
| genes | 60,623 |
| nonzero entries | 5,126,261 |
| density | 0.0243 |
| total counts | 19,836,914 |
| mean counts / cell | 5,697 |
| mean counts / gene | 327 |
| min counts / cell | 473 |
| max counts / cell | 99,294 |
| overdispersion | 2.02 |

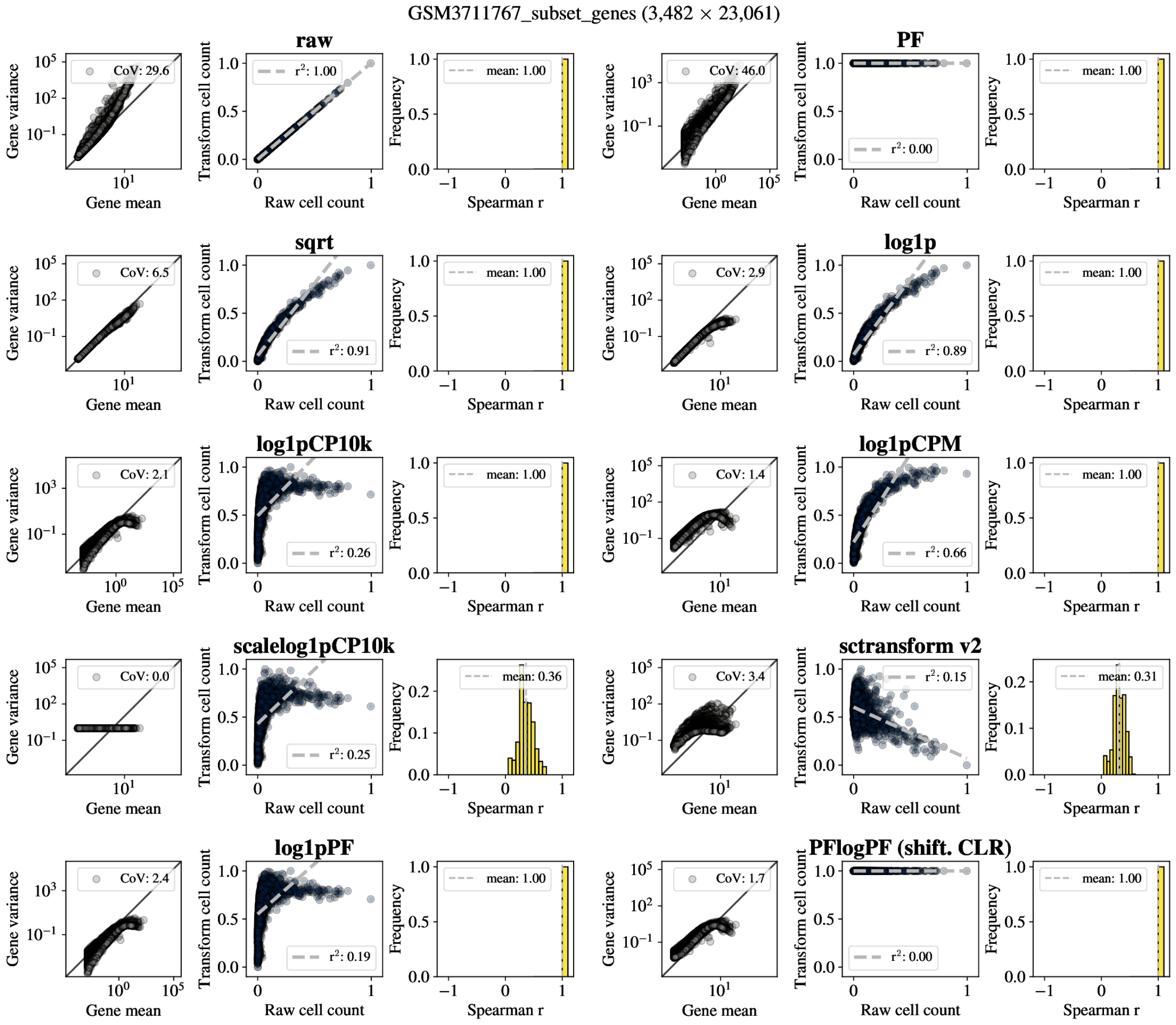


Supplementary Figure 1.357. **GSM3711767**. Per-method comparison of variance stabilization (left), depth normalization (middle), and monotonicity (right) for each normalization on dataset GSM3711767.

| **GSM3711768** | |
|---|---|
| cells | 5,203 |
| genes | 60,623 |
| nonzero entries | 3,162,212 |
| density | 0.01 |
| total counts | 7,338,829 |
| mean counts / cell | 1,410 |
| mean counts / gene | 121 |
| min counts / cell | 391 |
| max counts / cell | 17,454 |
| overdispersion | 1.22 |

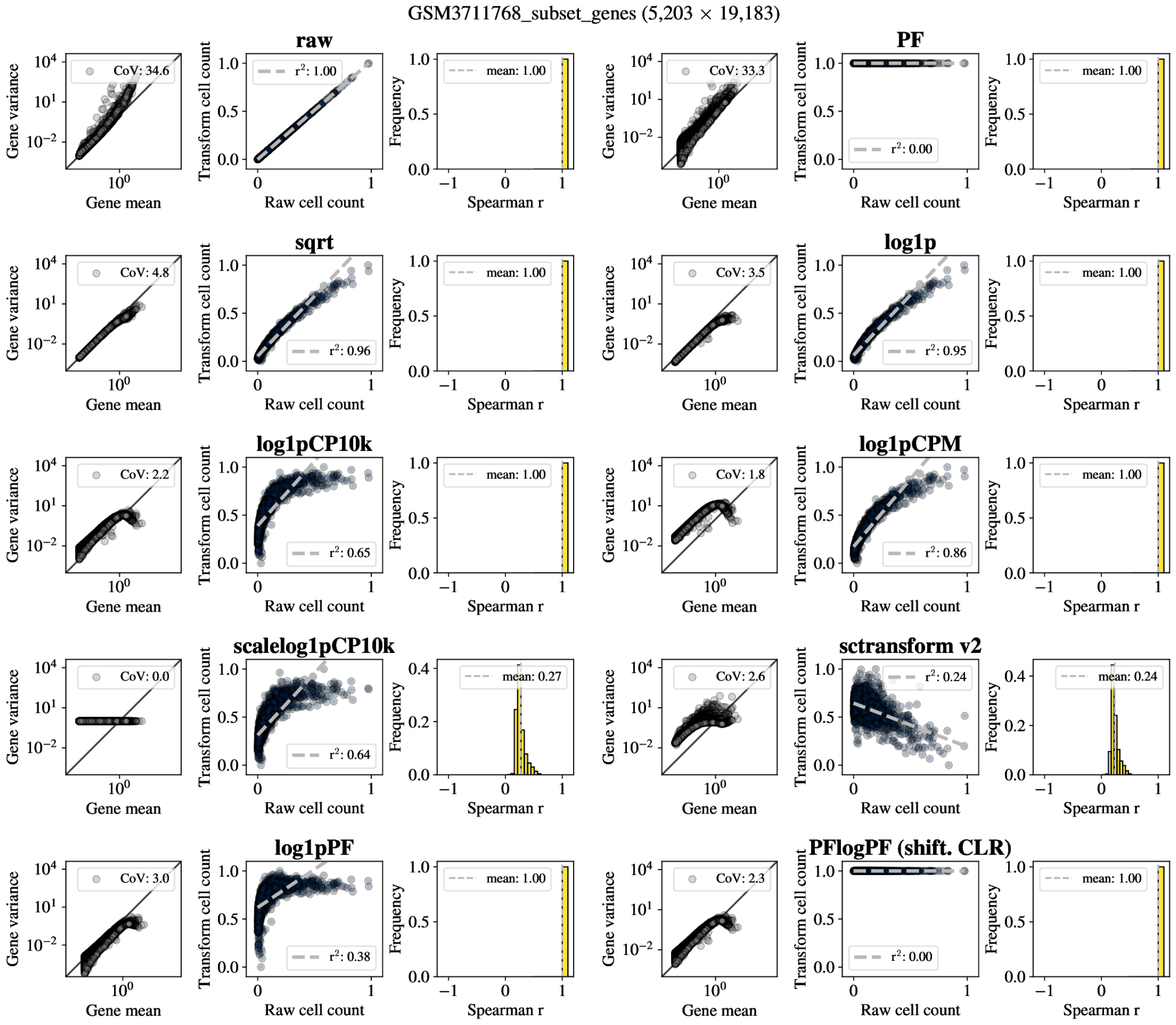


Supplementary Figure 1.358. **GSM3711768**. Per-method comparison of variance stabilization (left), depth normalization (middle), and monotonicity (right) for each normalization on dataset GSM3711768.

| GSM3711769 | |
|---|---|
| cells | 1,645 |
| genes | 60,623 |
| nonzero entries | 1,304,004 |
| density | 0.0131 |
| total counts | 3,415,131 |
| mean counts / cell | 2,076 |
| mean counts / gene | 56 |
| min counts / cell | 18 |
| max counts / cell | 41,759 |
| overdispersion | 8.68 |

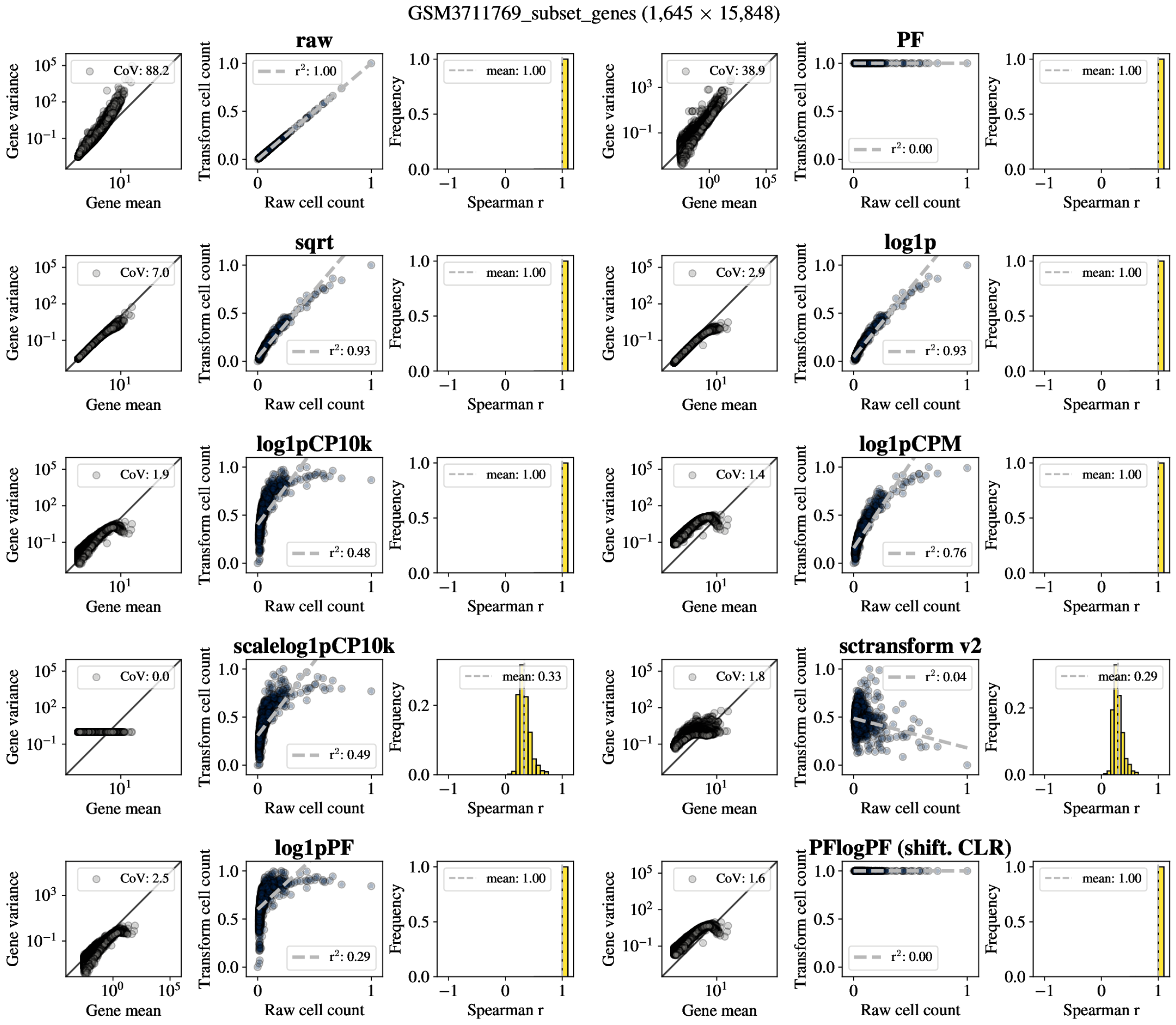


Supplementary Figure 1.359. **GSM3711769**. Per-method comparison of variance stabilization (left), depth normalization (middle), and monotonicity (right) for each normalization on dataset GSM3711769.

| **GSM3711770** | |
|---|---|
| cells | 408 |
| genes | 60,623 |
| nonzero entries | 324,243 |
| density | 0.0131 |
| total counts | 1,003,893 |
| mean counts / cell | 2,461 |
| mean counts / gene | 17 |
| min counts / cell | 237 |
| max counts / cell | 18,250 |
| overdispersion | 0.832 |

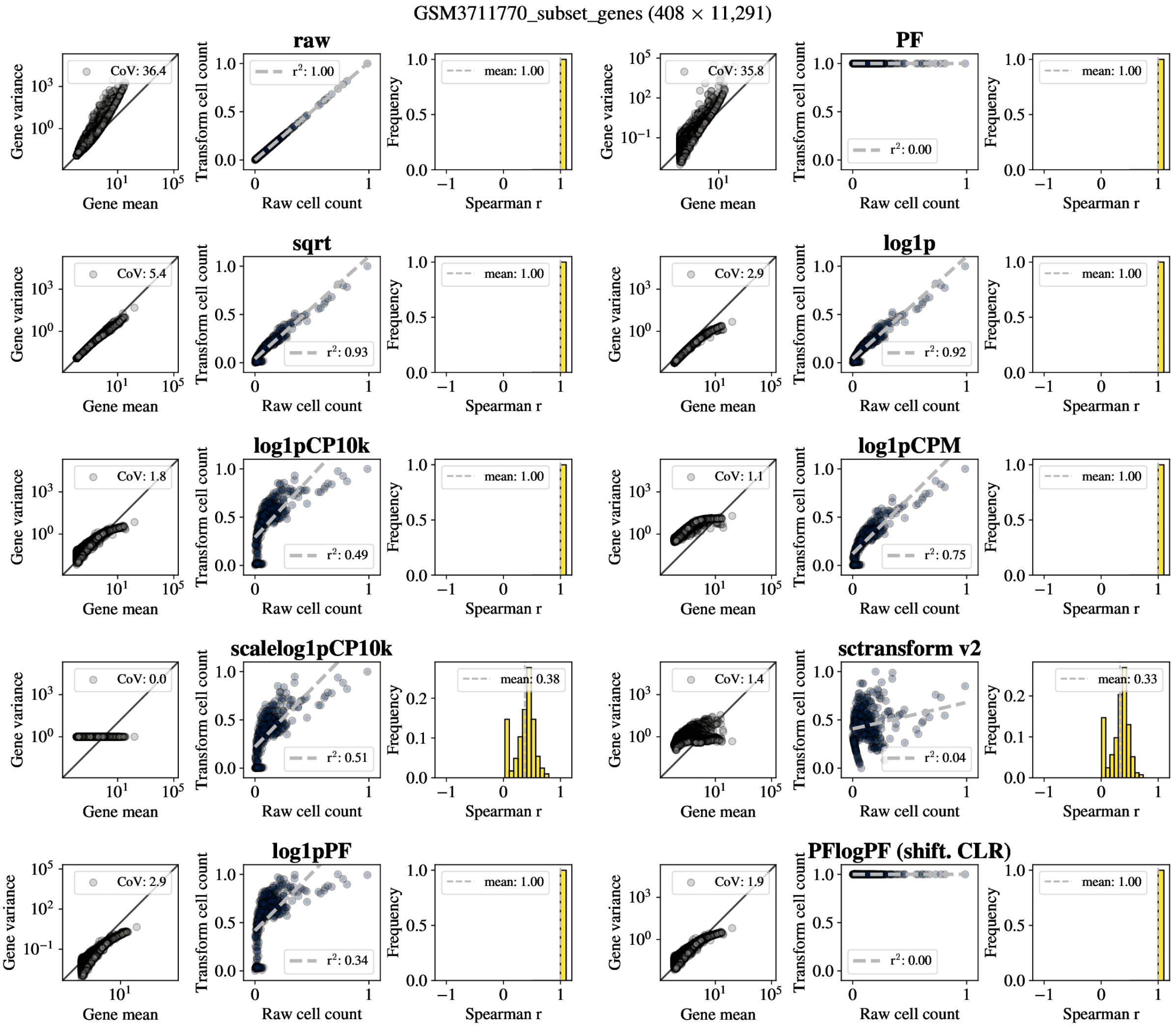


Supplementary Figure 1.360. **GSM3711770**. Per-method comparison of variance stabilization (left), depth normalization (middle), and monotonicity (right) for each normalization on dataset GSM3711770.

| GSM3711771 | |
|---|---|
| cells | 2,742 |
| genes | 60,623 |
| nonzero entries | 3,299,925 |
| density | 0.0199 |
| total counts | 12,278,278 |
| mean counts / cell | 4,478 |
| mean counts / gene | 203 |
| min counts / cell | 715 |
| max counts / cell | 57,529 |
| overdispersion | 2.17 |

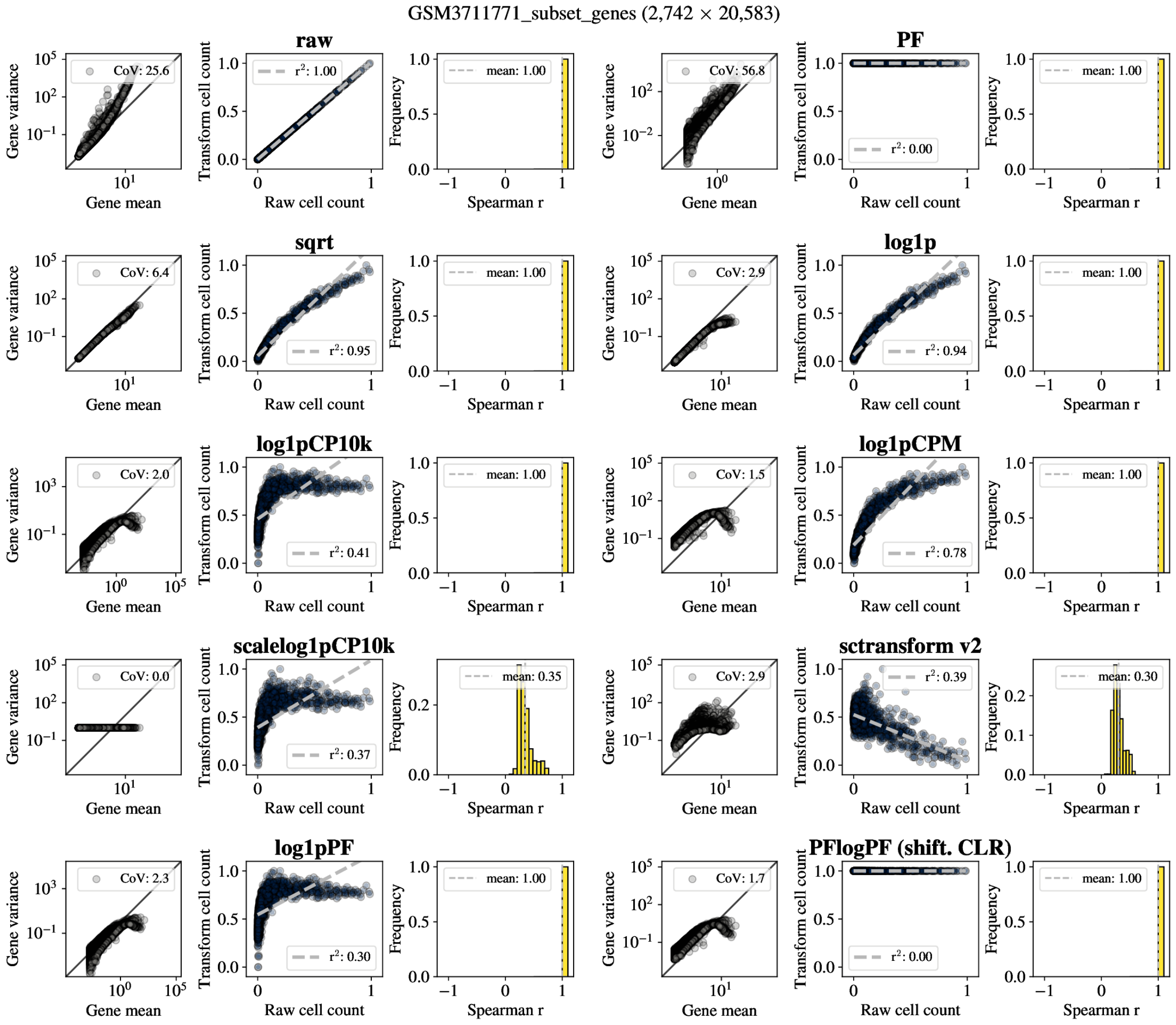


Supplementary Figure 1.361. **GSM3711771**. Per-method comparison of variance stabilization (left), depth normalization (middle), and monotonicity (right) for each normalization on dataset GSM3711771.

| GSM3711772 | |
|---|---|
| cells | 1,254 |
| genes | 60,623 |
| nonzero entries | 736,826 |
| density | 0.00969 |
| total counts | 2,673,882 |
| mean counts / cell | 2,132 |
| mean counts / gene | 44 |
| min counts / cell | 244 |
| max counts / cell | 23,471 |
| overdispersion | 1.98 |

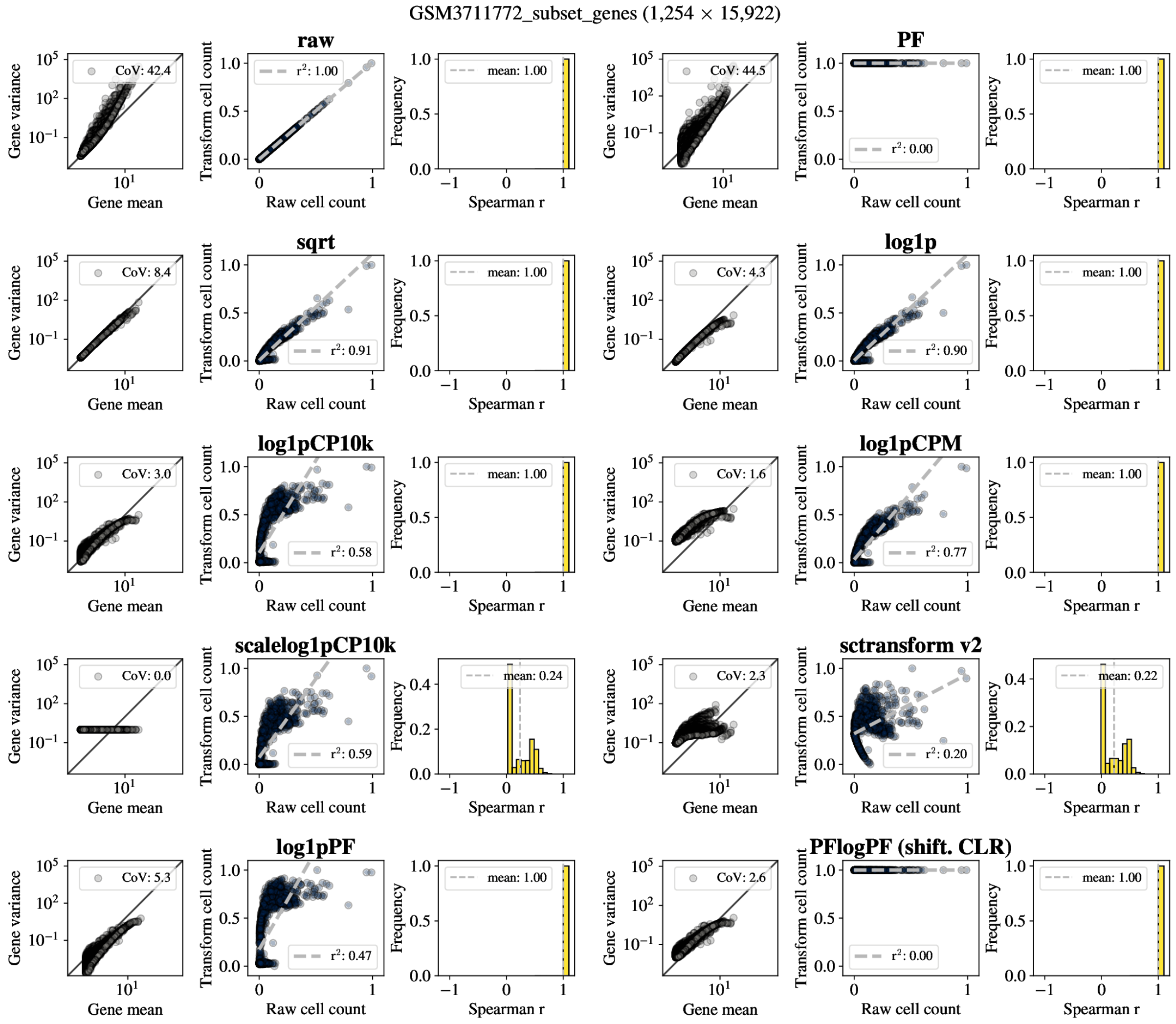


Supplementary Figure 1.362. **GSM3711772**. Per-method comparison of variance stabilization (left), depth normalization (middle), and monotonicity (right) for each normalization on dataset GSM3711772.

| GSM3711773 | |
|---|---|
| cells | 14,917 |
| genes | 60,623 |
| nonzero entries | 13,080,793 |
| density | 0.0145 |
| total counts | 40,200,974 |
| mean counts / cell | 2,695 |
| mean counts / gene | 663 |
| min counts / cell | 410 |
| max counts / cell | 61,557 |
| overdispersion | 1.51 |

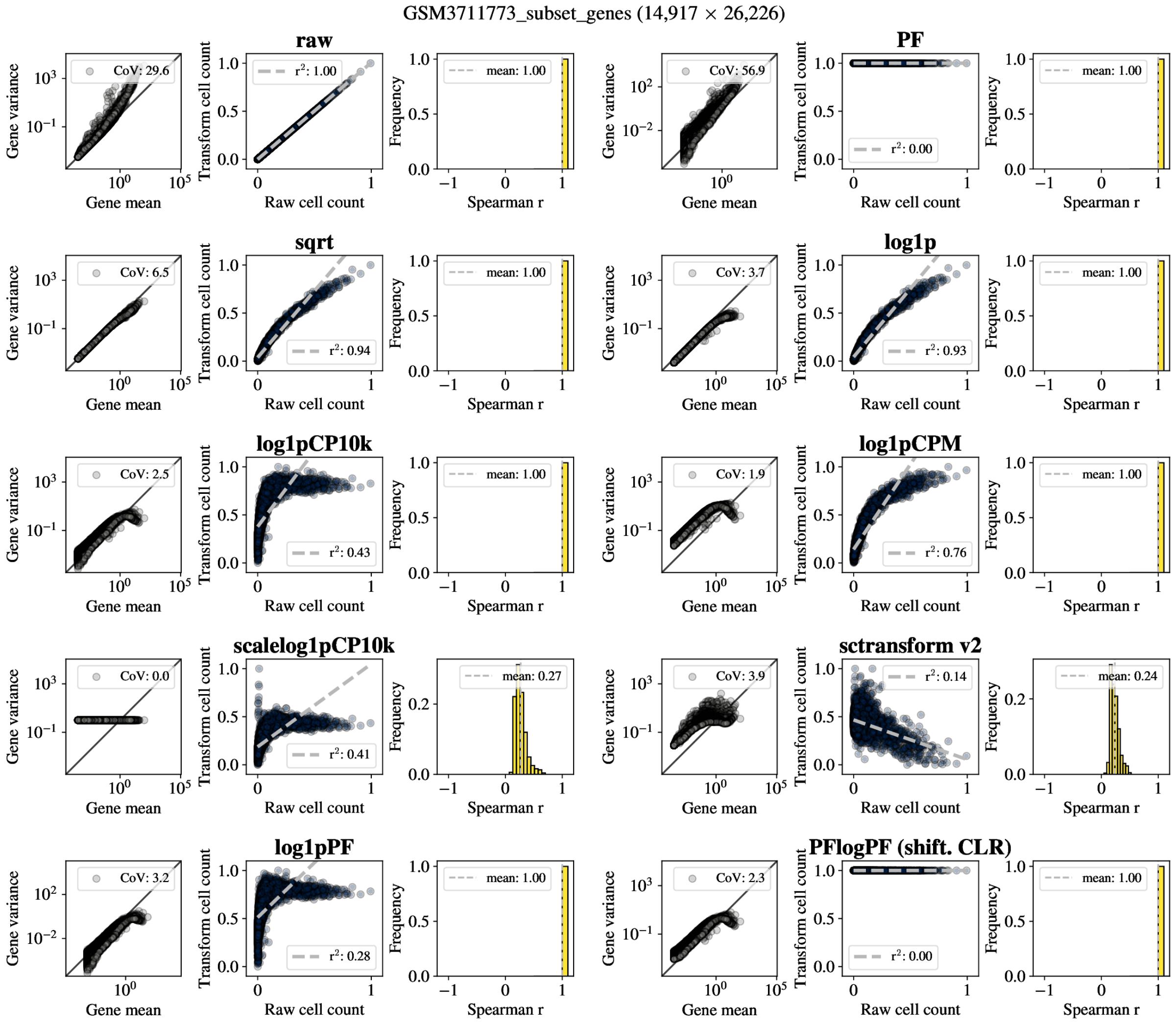


Supplementary Figure 1.363. **GSM3711773**. Per-method comparison of variance stabilization (left), depth normalization (middle), and monotonicity (right) for each normalization on dataset GSM3711773.

| GSM3711774 | |
|---|---|
| cells | 3,167 |
| genes | 60,623 |
| nonzero entries | 3,661,191 |
| density | 0.0191 |
| total counts | 11,770,596 |
| mean counts / cell | 3,717 |
| mean counts / gene | 194 |
| min counts / cell | 363 |
| max counts / cell | 55,248 |
| overdispersion | 0.317 |

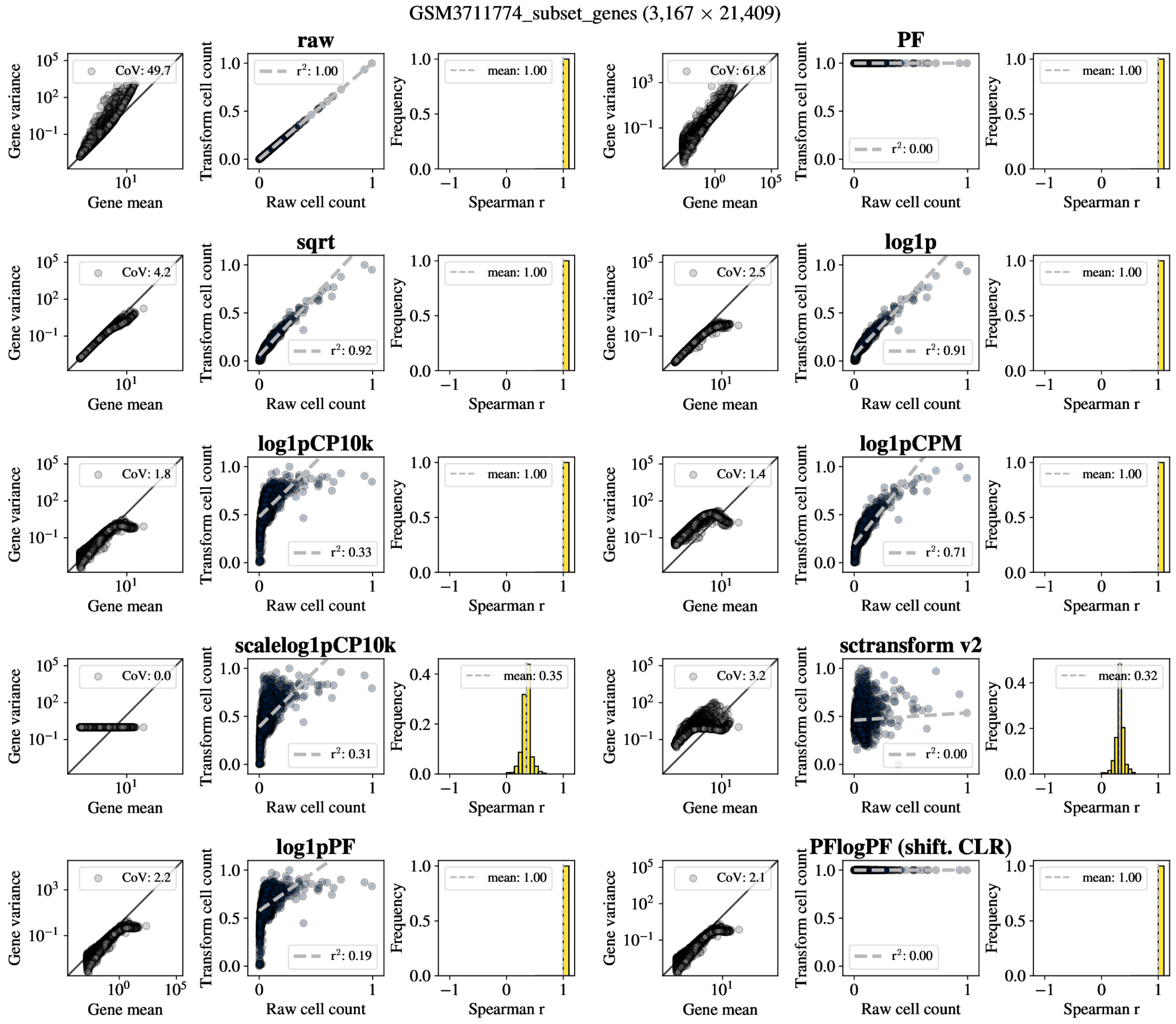


Supplementary Figure 1.364. **GSM3711774**. Per-method comparison of variance stabilization (left), depth normalization (middle), and monotonicity (right) for each normalization on dataset GSM3711774.

| GSM3711775 | |
|---|---|
| cells | 1,911 |
| genes | 60,623 |
| nonzero entries | 2,813,900 |
| density | 0.0243 |
| total counts | 12,025,327 |
| mean counts / cell | 6,293 |
| mean counts / gene | 198 |
| min counts / cell | 417 |
| max counts / cell | 129,404 |
| overdispersion | 2.71 |

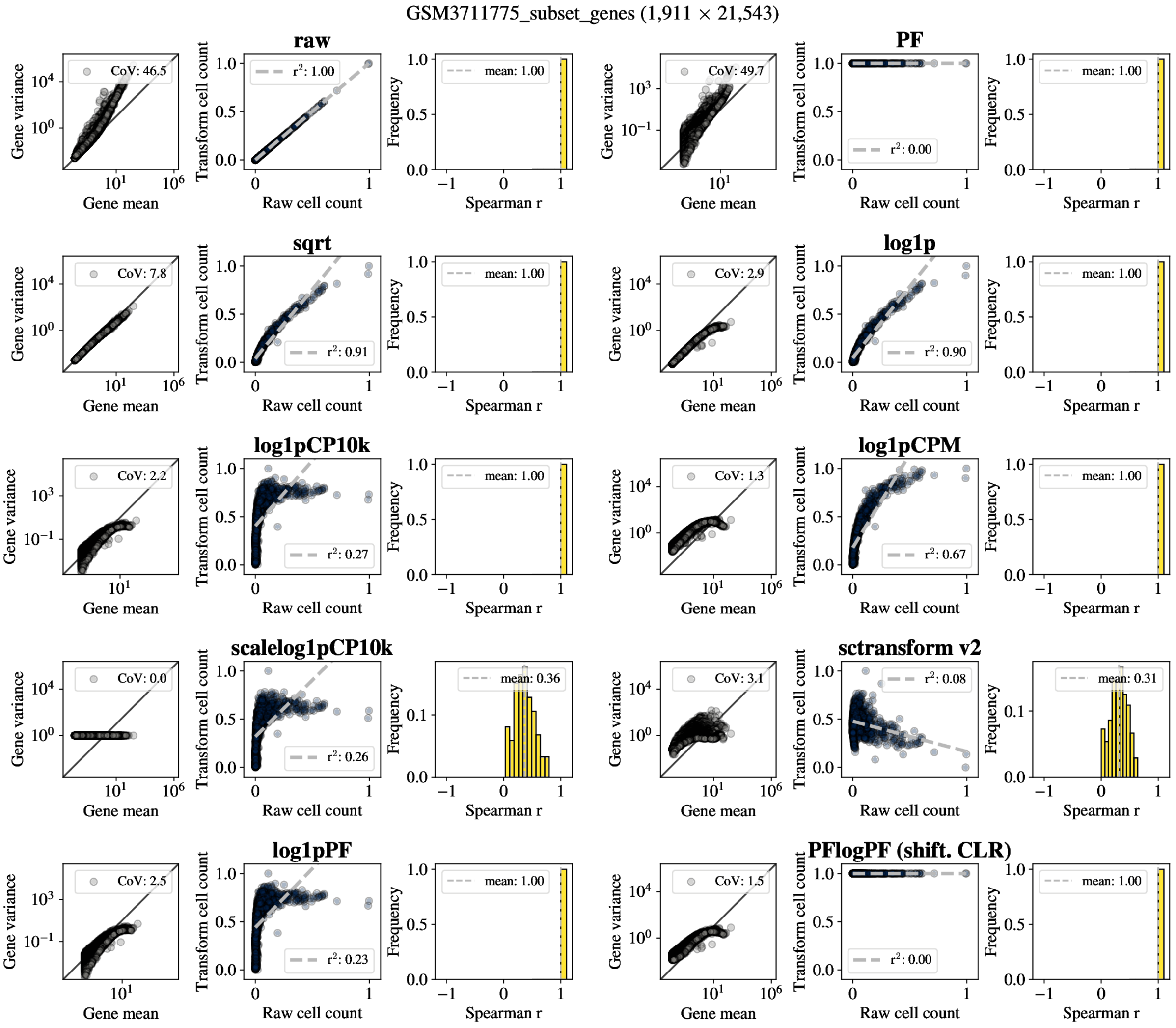


Supplementary Figure 1.365. **GSM3711775**. Per-method comparison of variance stabilization (left), depth normalization (middle), and monotonicity (right) for each normalization on dataset GSM3711775.

| GSM3711776 | |
|---|---|
| cells | 1,072 |
| genes | 60,623 |
| nonzero entries | 1,245,127 |
| density | 0.0192 |
| total counts | 3,878,944 |
| mean counts / cell | 3,618 |
| mean counts / gene | 64 |
| min counts / cell | 314 |
| max counts / cell | 24,425 |
| overdispersion | 0.783 |

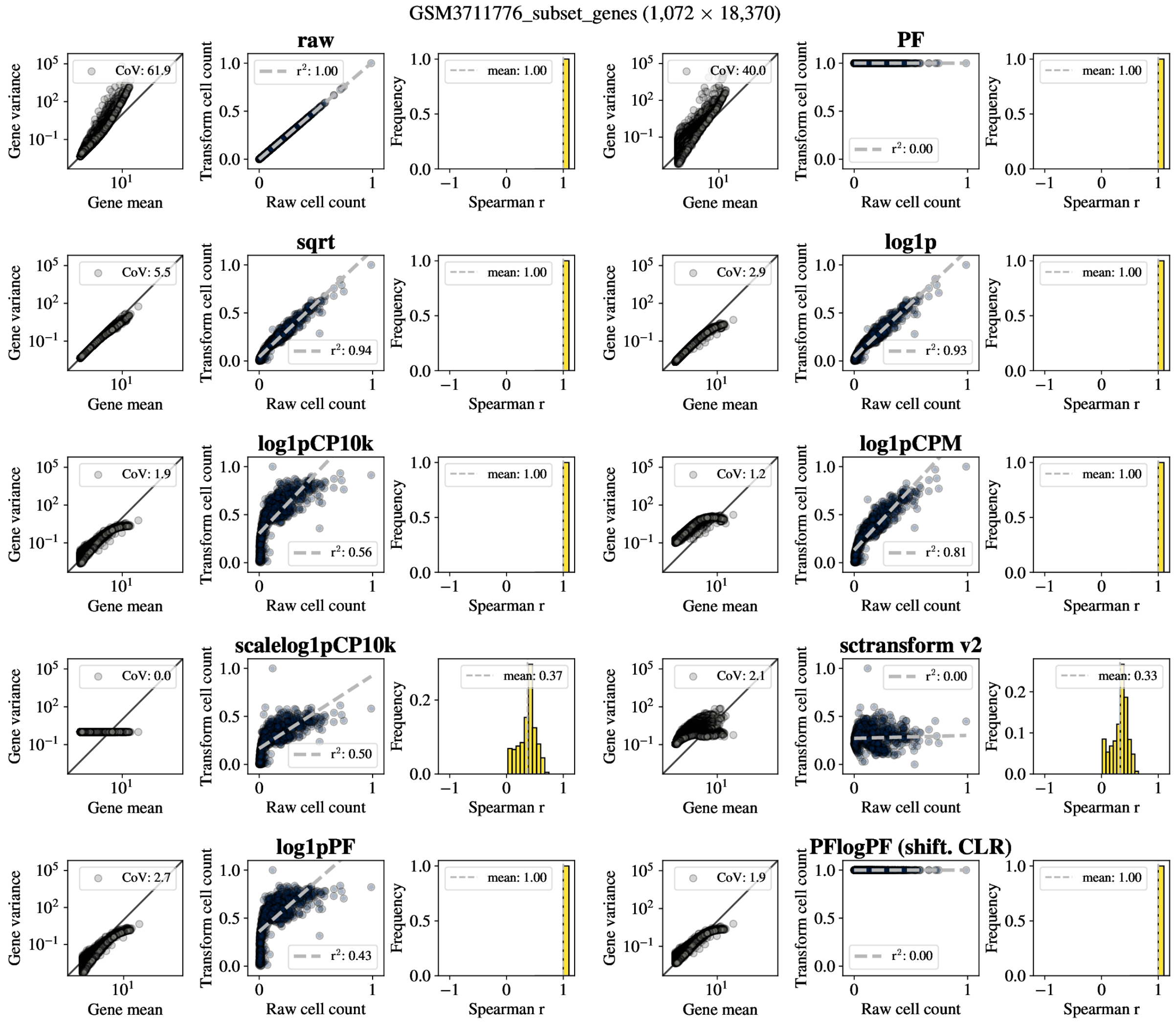


Supplementary Figure 1.366. **GSM3711776**. Per-method comparison of variance stabilization (left), depth normalization (middle), and monotonicity (right) for each normalization on dataset GSM3711776.

| GSM3711777 | |
|---|---|
| cells | 2,133 |
| genes | 60,623 |
| nonzero entries | 2,613,407 |
| density | 0.0202 |
| total counts | 9,000,877 |
| mean counts / cell | 4,220 |
| mean counts / gene | 148 |
| min counts / cell | 448 |
| max counts / cell | 49,284 |
| overdispersion | 0.997 |

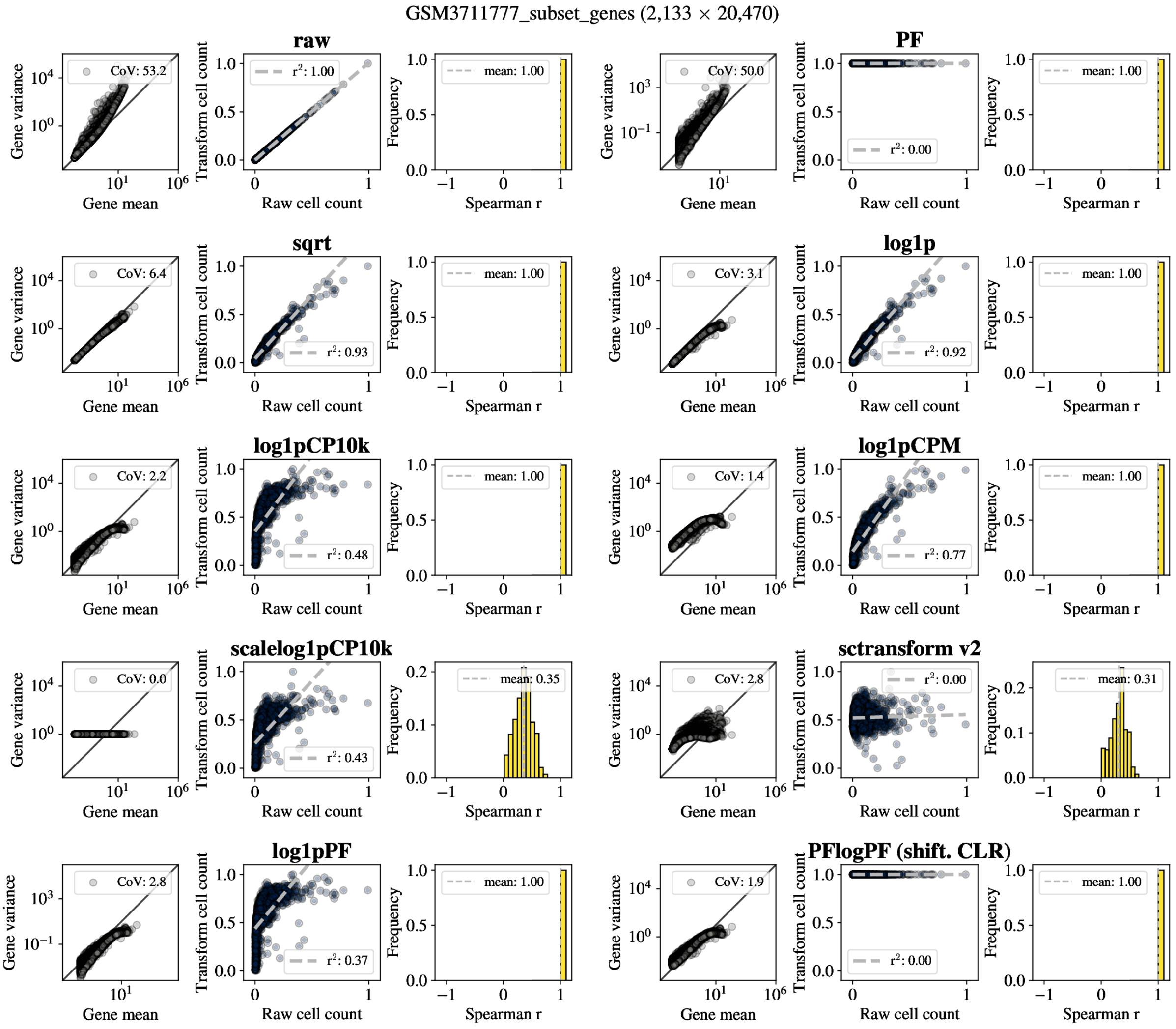


Supplementary Figure 1.367. **GSM3711777**. Per-method comparison of variance stabilization (left), depth normalization (middle), and monotonicity (right) for each normalization on dataset GSM3711777.

| GSM3711778 | |
|---|---|
| cells | 473 |
| genes | 60,623 |
| nonzero entries | 264,772 |
| density | 0.00923 |
| total counts | 685,404 |
| mean counts / cell | 1,449 |
| mean counts / gene | 11 |
| min counts / cell | 17 |
| max counts / cell | 15,002 |
| overdispersion | 2.04 |

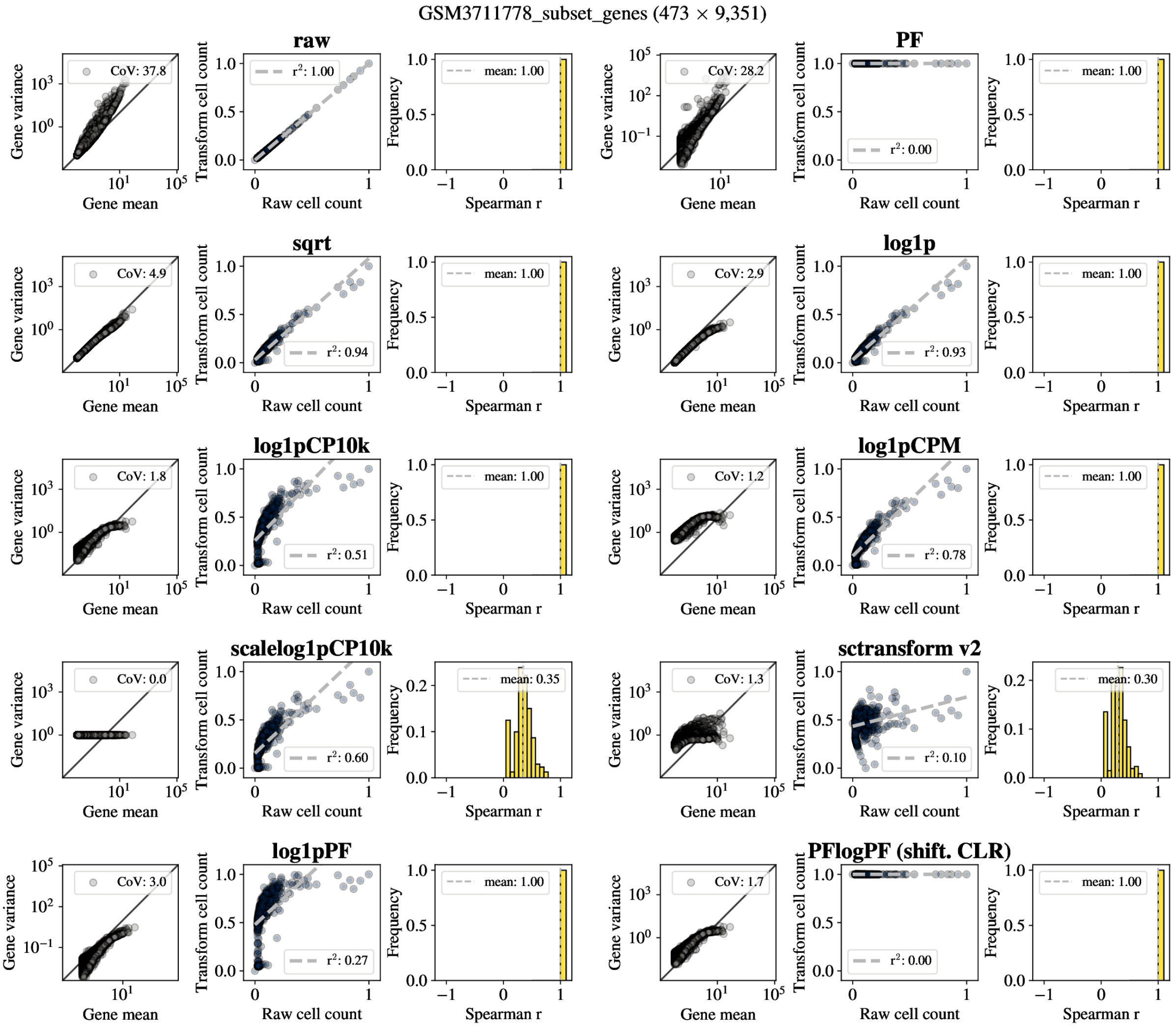


Supplementary Figure 1.368. **GSM3711778**. Per-method comparison of variance stabilization (left), depth normalization (middle), and monotonicity (right) for each normalization on dataset GSM3711778.

| **GSM3711779** | |
|---|---|
| cells | 4,077 |
| genes | 60,623 |
| nonzero entries | 2,853,269 |
| density | 0.0115 |
| total counts | 7,807,665 |
| mean counts / cell | 1,915 |
| mean counts / gene | 129 |
| min counts / cell | 441 |
| max counts / cell | 21,468 |
| overdispersion | 0.865 |

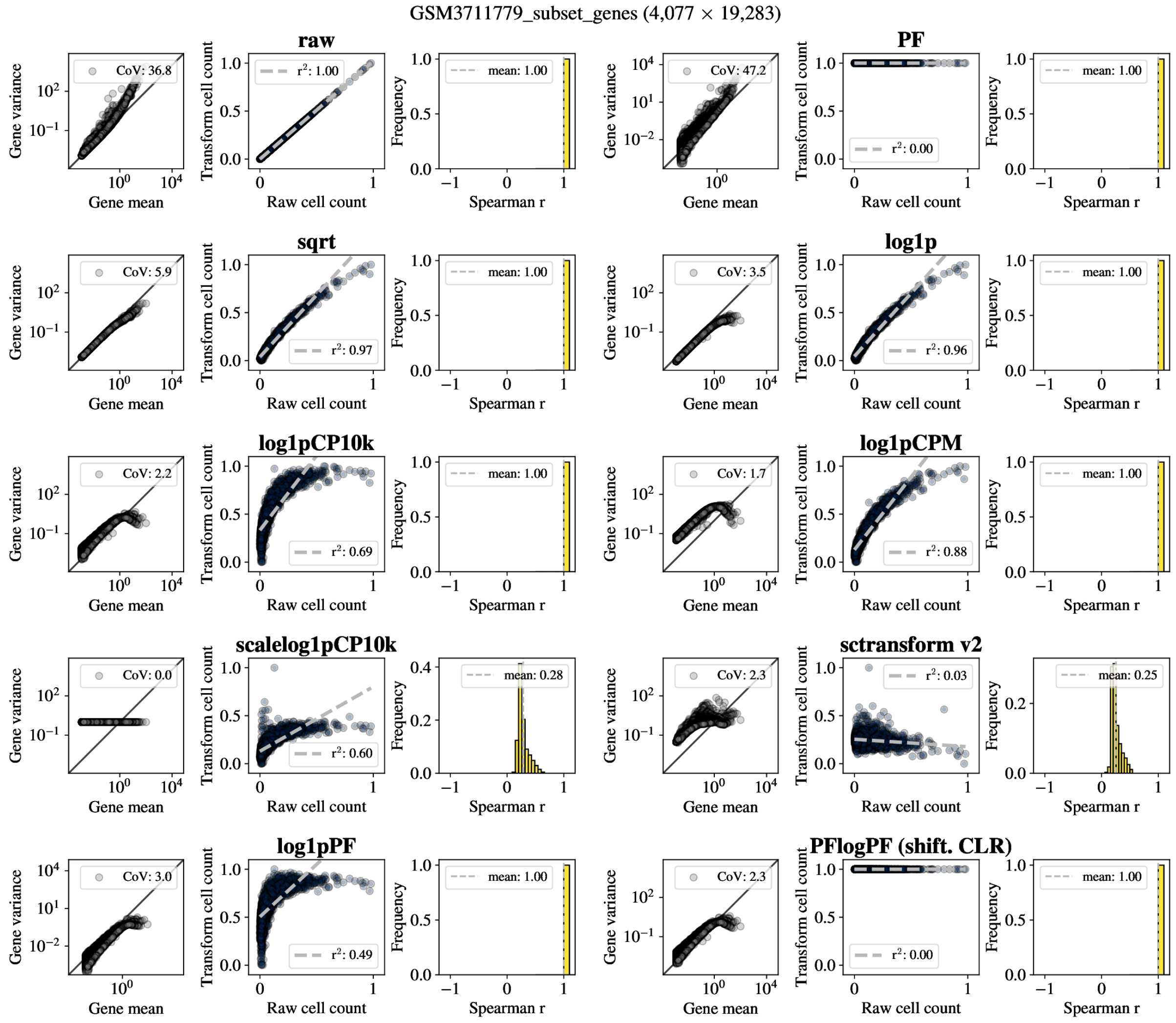


Supplementary Figure 1.369. **GSM3711779**. Per-method comparison of variance stabilization (left), depth normalization (middle), and monotonicity (right) for each normalization on dataset GSM3711779.

| GSM3711780 | |
|---|---|
| cells | 11,752 |
| genes | 60,623 |
| nonzero entries | 4,154,522 |
| density | 0.00583 |
| total counts | 8,687,076 |
| mean counts / cell | 739 |
| mean counts / gene | 143 |
| min counts / cell | 248 |
| max counts / cell | 117,065 |
| overdispersion | 6.79 |

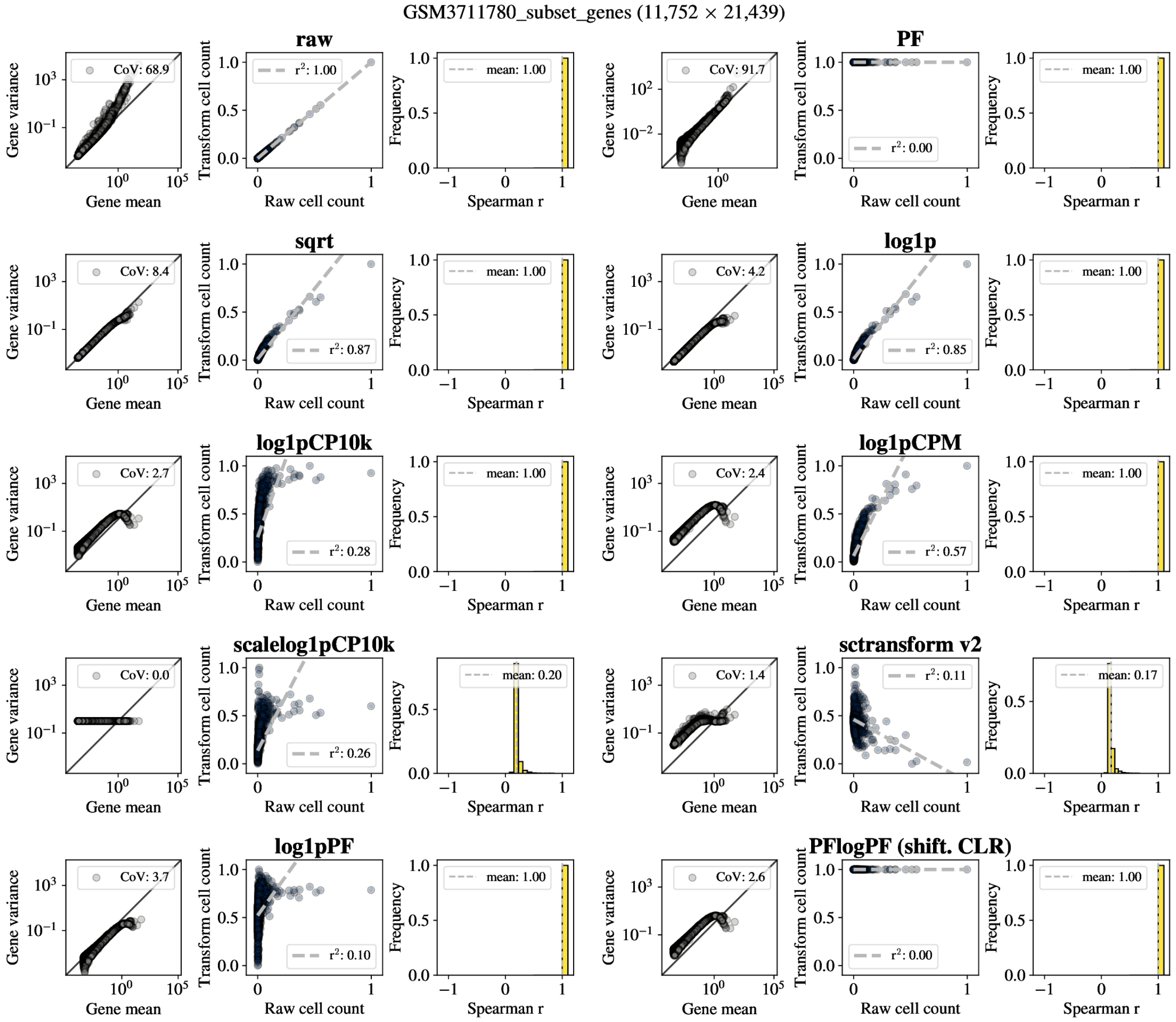


Supplementary Figure 1.370. **GSM3711780**. Per-method comparison of variance stabilization (left), depth normalization (middle), and monotonicity (right) for each normalization on dataset GSM3711780.

| GSM3711781 | |
|---|---|
| cells | 2,288 |
| genes | 60,623 |
| nonzero entries | 3,066,951 |
| density | 0.0221 |
| total counts | 10,928,512 |
| mean counts / cell | 4,776 |
| mean counts / gene | 180 |
| min counts / cell | 646 |
| max counts / cell | 69,074 |
| overdispersion | 9.08 |

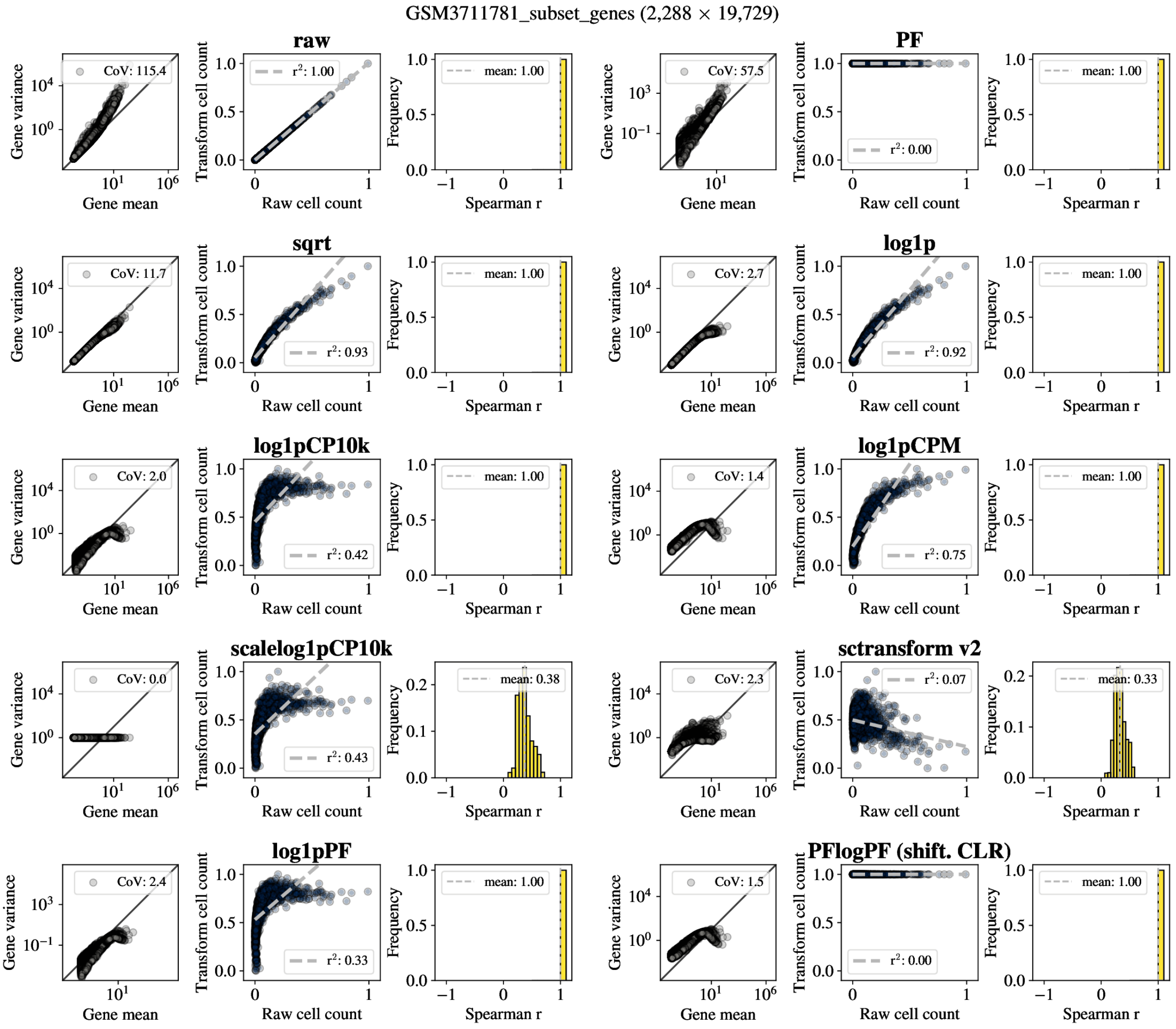


Supplementary Figure 1.371. **GSM3711781**. Per-method comparison of variance stabilization (left), depth normalization (middle), and monotonicity (right) for each normalization on dataset GSM3711781.

| GSM3711782 | |
|---|---|
| cells | 2,127 |
| genes | 60,623 |
| nonzero entries | 2,148,253 |
| density | 0.0167 |
| total counts | 6,508,164 |
| mean counts / cell | 3,060 |
| mean counts / gene | 107 |
| min counts / cell | 424 |
| max counts / cell | 43,268 |
| overdispersion | 0.941 |

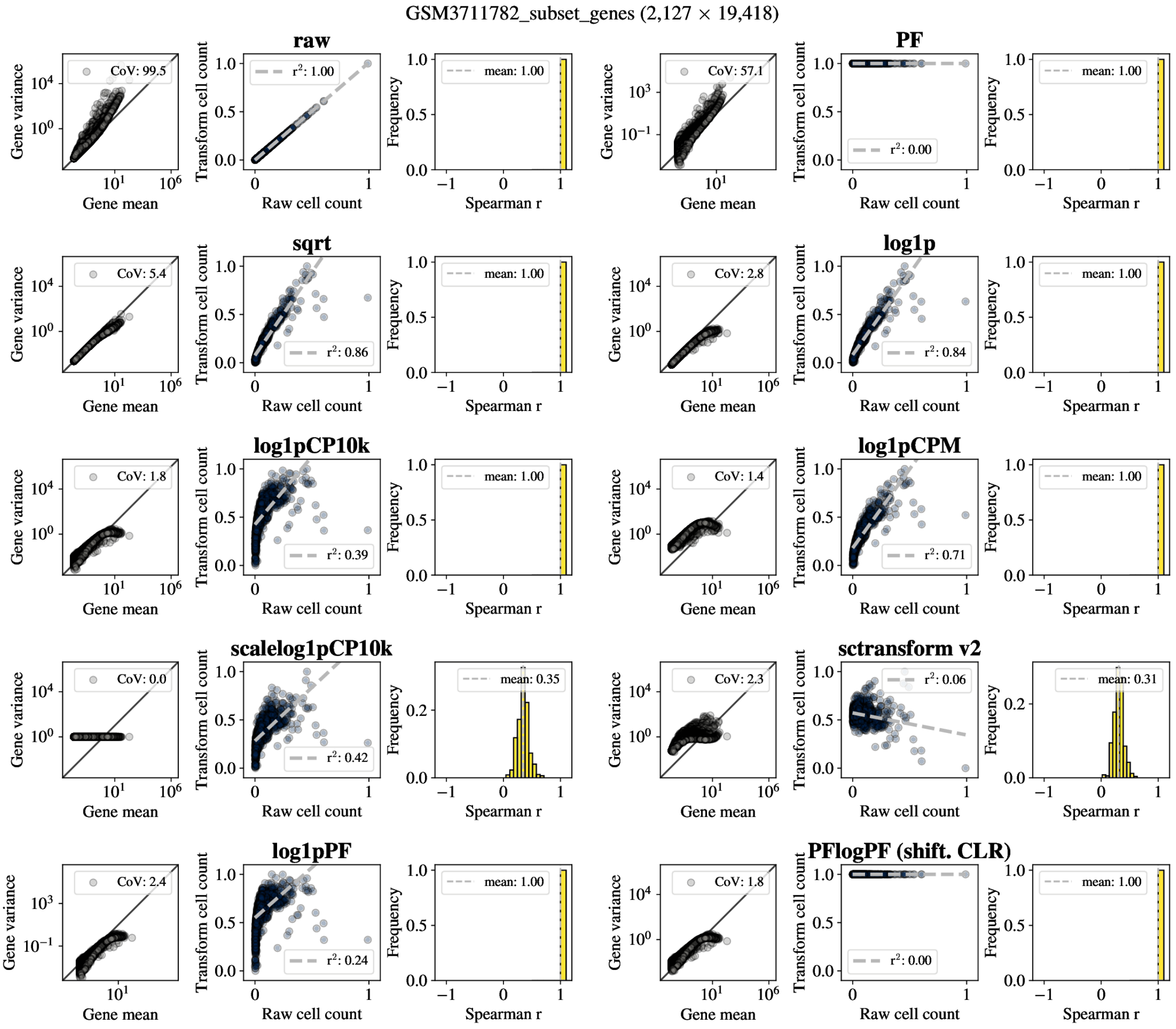


Supplementary Figure 1.372. **GSM3711782**. Per-method comparison of variance stabilization (left), depth normalization (middle), and monotonicity (right) for each normalization on dataset GSM3711782.

| GSM3711783 | |
|---|---|
| cells | 1,404 |
| genes | 60,623 |
| nonzero entries | 2,025,685 |
| density | 0.0238 |
| total counts | 6,029,952 |
| mean counts / cell | 4,295 |
| mean counts / gene | 99 |
| min counts / cell | 399 |
| max counts / cell | 35,659 |
| overdispersion | 0.544 |

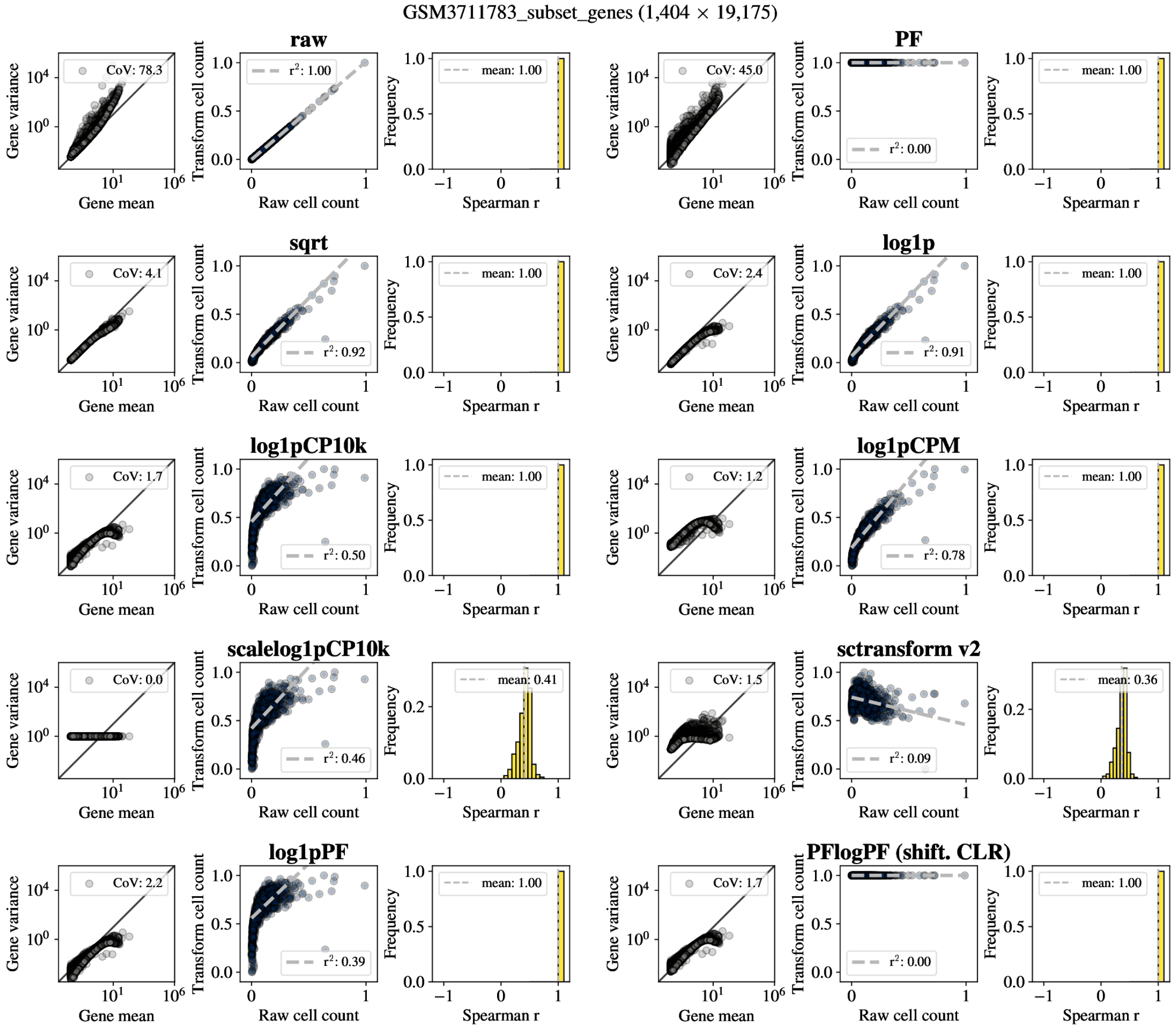


Supplementary Figure 1.373. **GSM3711783**. Per-method comparison of variance stabilization (left), depth normalization (middle), and monotonicity (right) for each normalization on dataset GSM3711783.

| GSM3711784 | |
|---|---|
| cells | 1,215 |
| genes | 60,623 |
| nonzero entries | 1,291,317 |
| density | 0.0175 |
| total counts | 4,221,785 |
| mean counts / cell | 3,475 |
| mean counts / gene | 70 |
| min counts / cell | 364 |
| max counts / cell | 58,505 |
| overdispersion | 0.528 |

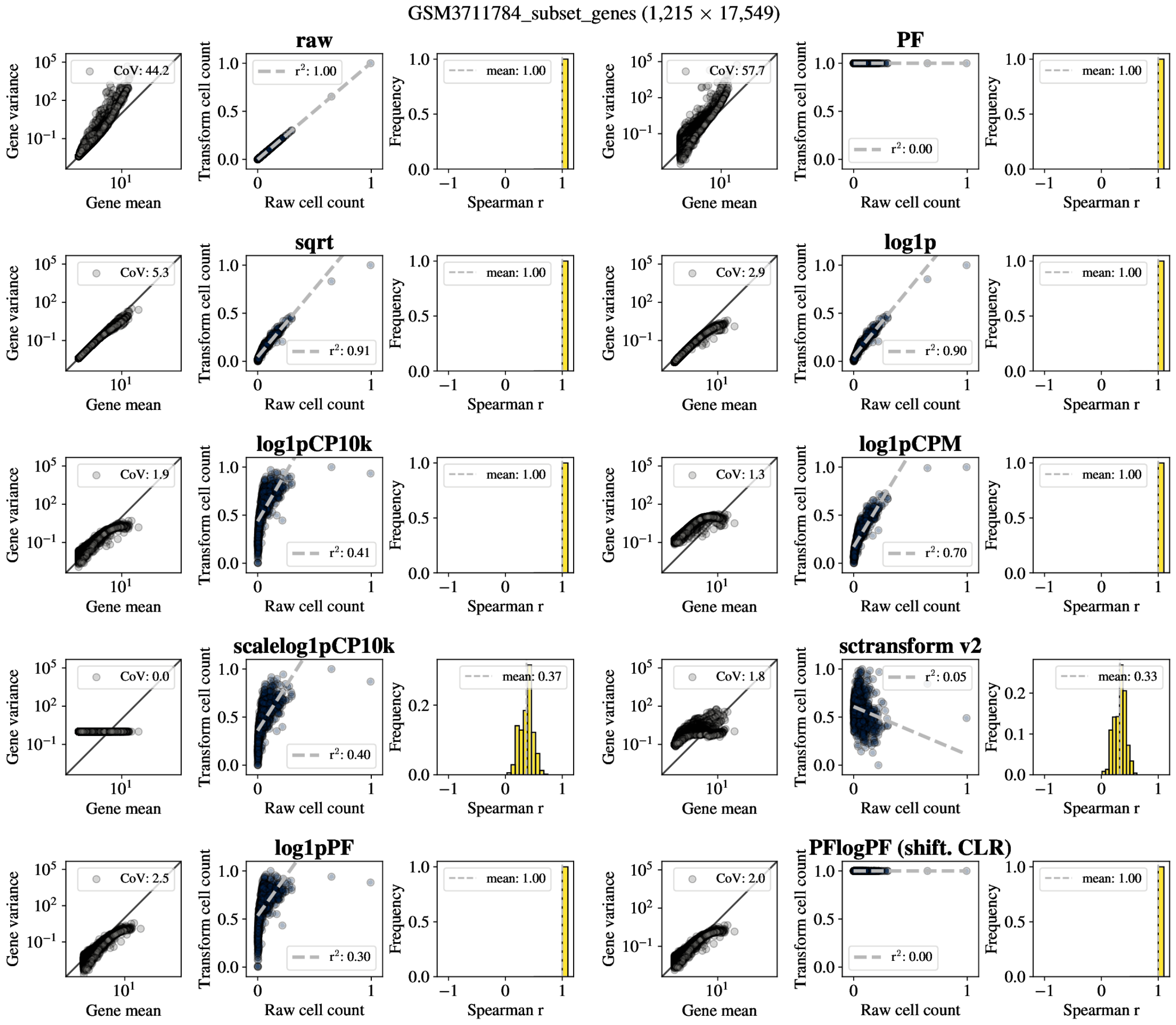


Supplementary Figure 1.374. **GSM3711784**. Per-method comparison of variance stabilization (left), depth normalization (middle), and monotonicity (right) for each normalization on dataset GSM3711784.

| GSM3711785 | |
|---|---|
| cells | 292 |
| genes | 60,623 |
| nonzero entries | 407,453 |
| density | 0.023 |
| total counts | 1,495,426 |
| mean counts / cell | 5,121 |
| mean counts / gene | 25 |
| min counts / cell | 399 |
| max counts / cell | 19,679 |
| overdispersion | 0.461 |

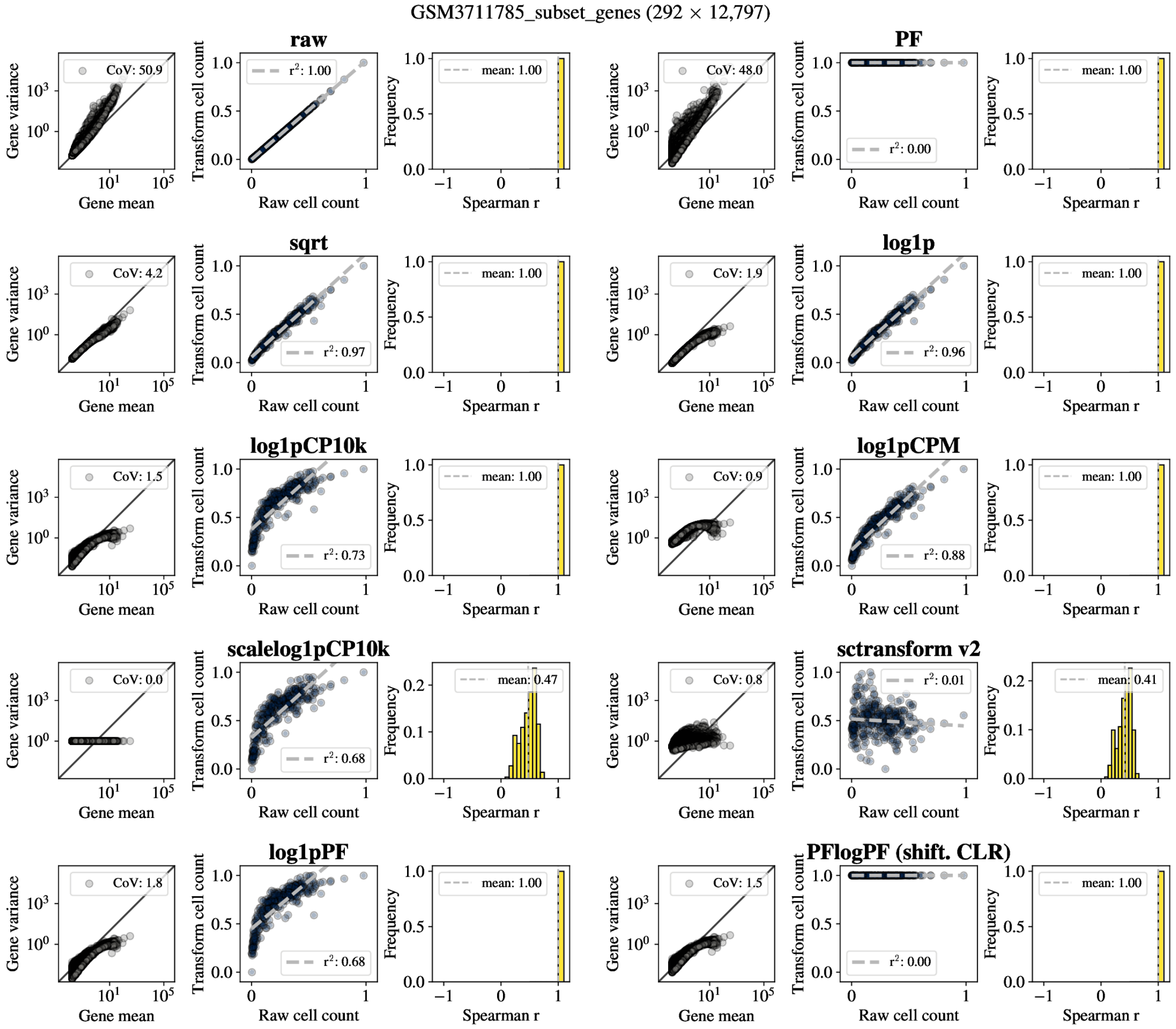


Supplementary Figure 1.375. **GSM3711785**. Per-method comparison of variance stabilization (left), depth normalization (middle), and monotonicity (right) for each normalization on dataset GSM3711785.

| GSM3711786 | |
|---|---|
| cells | 5,235 |
| genes | 60,623 |
| nonzero entries | 5,700,776 |
| density | 0.018 |
| total counts | 16,798,786 |
| mean counts / cell | 3,209 |
| mean counts / gene | 277 |
| min counts / cell | 408 |
| max counts / cell | 30,525 |
| overdispersion | 0.234 |

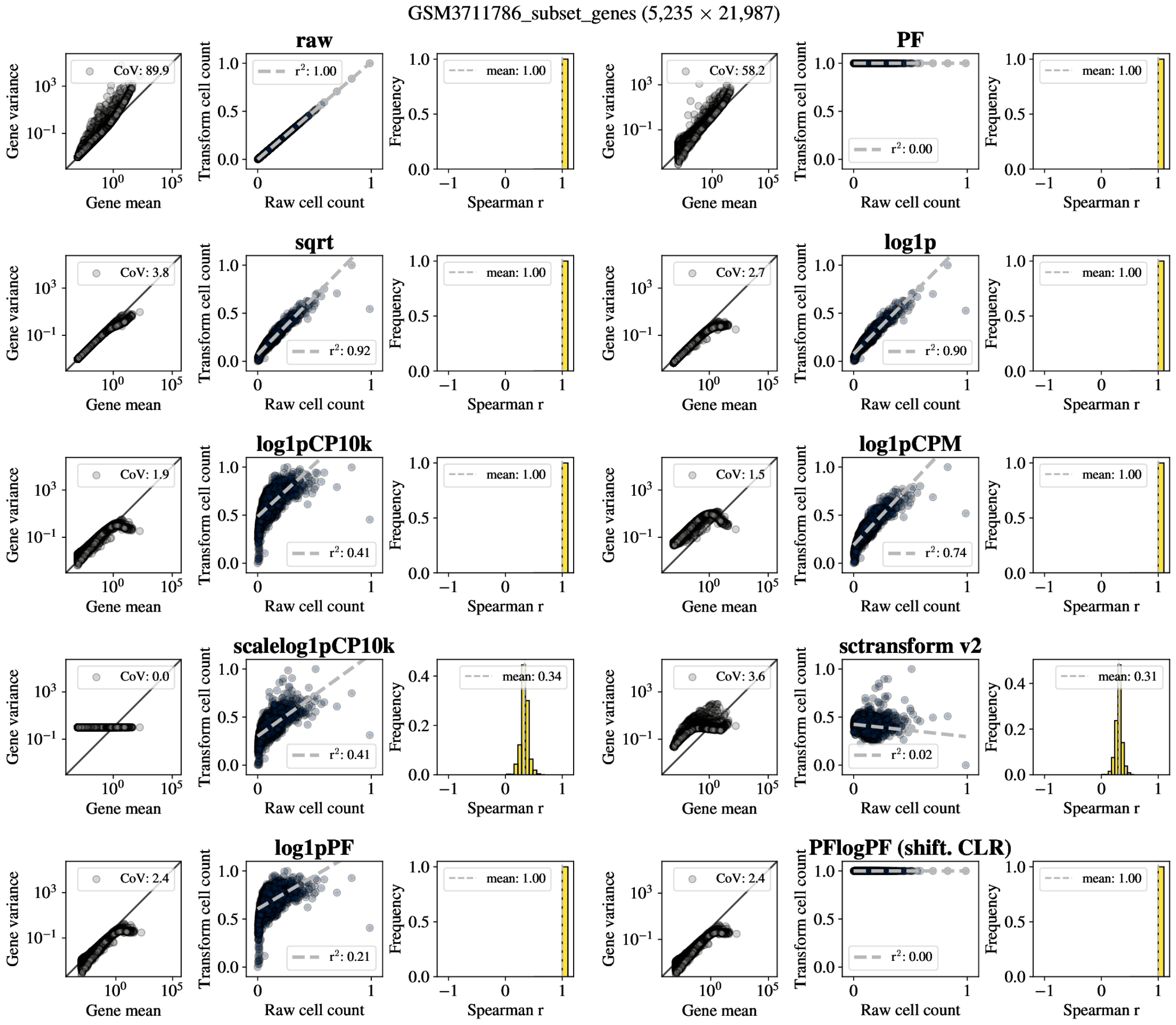


Supplementary Figure 1.376. **GSM3711786**. Per-method comparison of variance stabilization (left), depth normalization (middle), and monotonicity (right) for each normalization on dataset GSM3711786.

| GSM3711787 | |
|---|---|
| cells | 3,164 |
| genes | 60,623 |
| nonzero entries | 6,025,365 |
| density | 0.0314 |
| total counts | 19,396,340 |
| mean counts / cell | 6,130 |
| mean counts / gene | 320 |
| min counts / cell | 561 |
| max counts / cell | 32,767 |
| overdispersion | 0.266 |

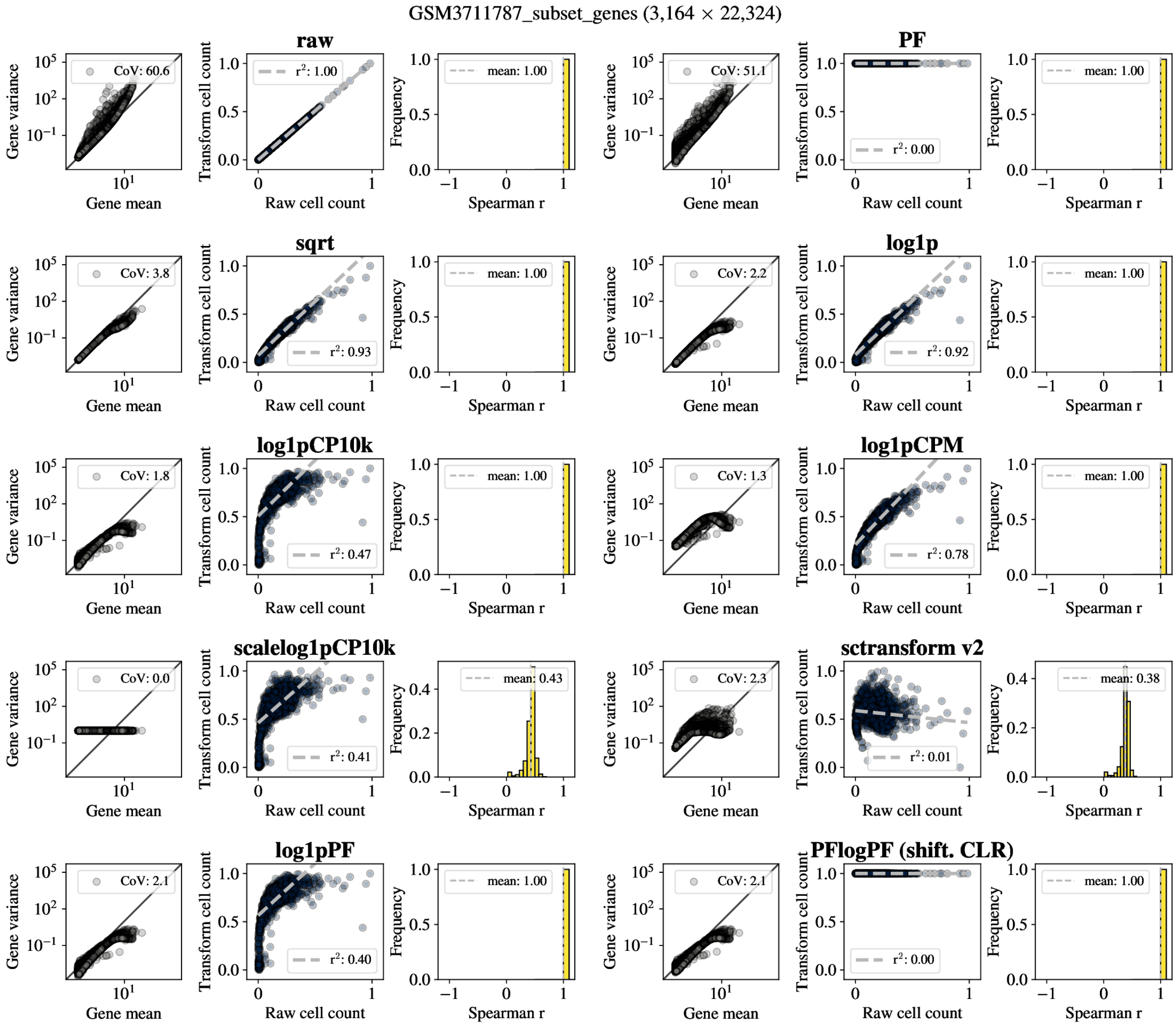


Supplementary Figure 1.377. **GSM3711787**. Per-method comparison of variance stabilization (left), depth normalization (middle), and monotonicity (right) for each normalization on dataset GSM3711787.

| GSM3717979 | |
|---|---|
| cells | 12,211 |
| genes | 60,623 |
| nonzero entries | 15,283,577 |
| density | 0.0206 |
| total counts | 37,123,283 |
| mean counts / cell | 3,040 |
| mean counts / gene | 612 |
| min counts / cell | 174 |
| max counts / cell | 27,128 |
| overdispersion | 0.412 |

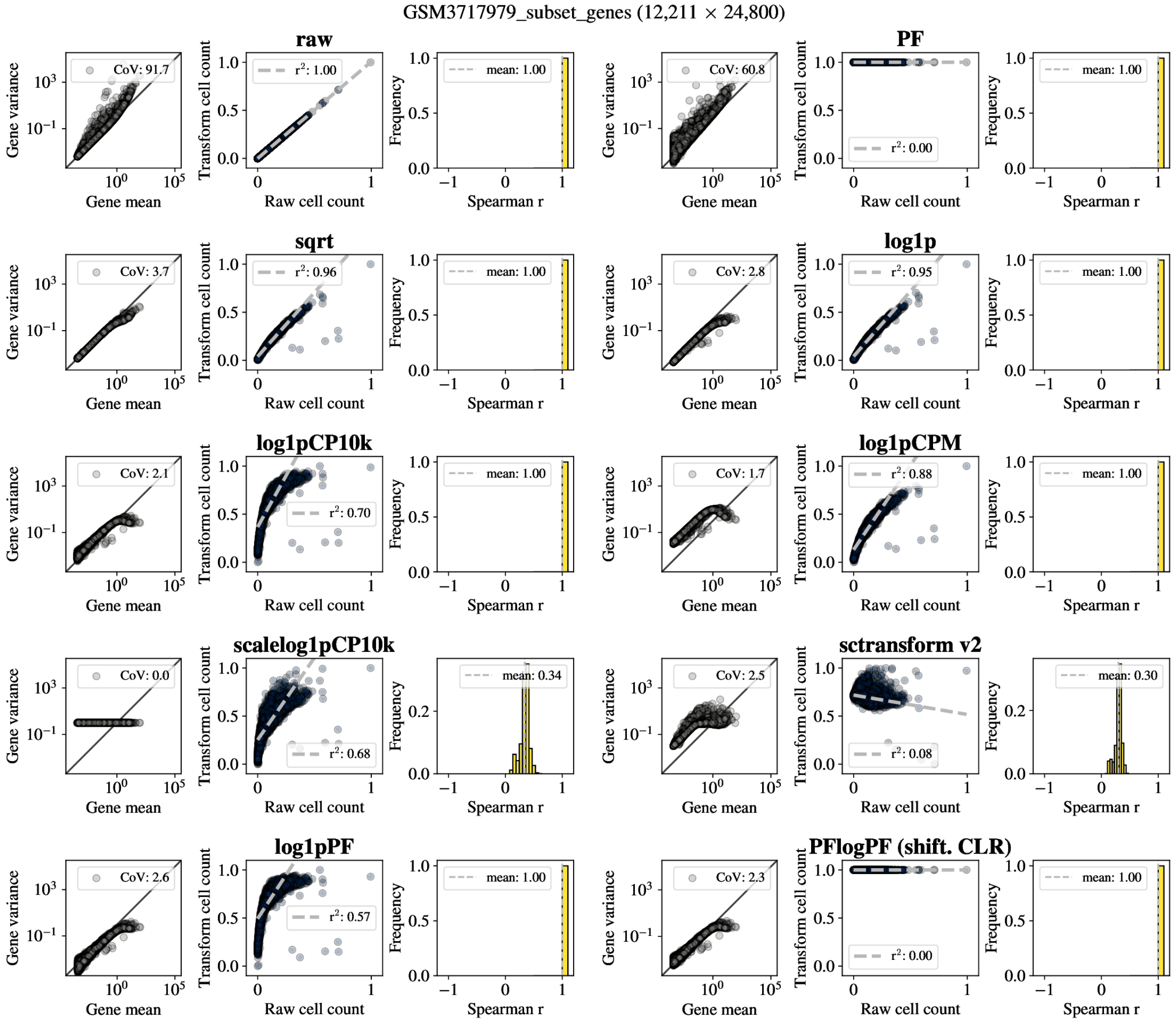


Supplementary Figure 1.378. **GSM3717979**. Per-method comparison of variance stabilization (left), depth normalization (middle), and monotonicity (right) for each normalization on dataset GSM3717979.

| GSM3723357 | |
|---|---|
| cells | 680 |
| genes | 60,623 |
| nonzero entries | 1,104,199 |
| density | 0.0268 |
| total counts | 4,566,432 |
| mean counts / cell | 6,715 |
| mean counts / gene | 75 |
| min counts / cell | 421 |
| max counts / cell | 101,114 |
| overdispersion | 1.99 |

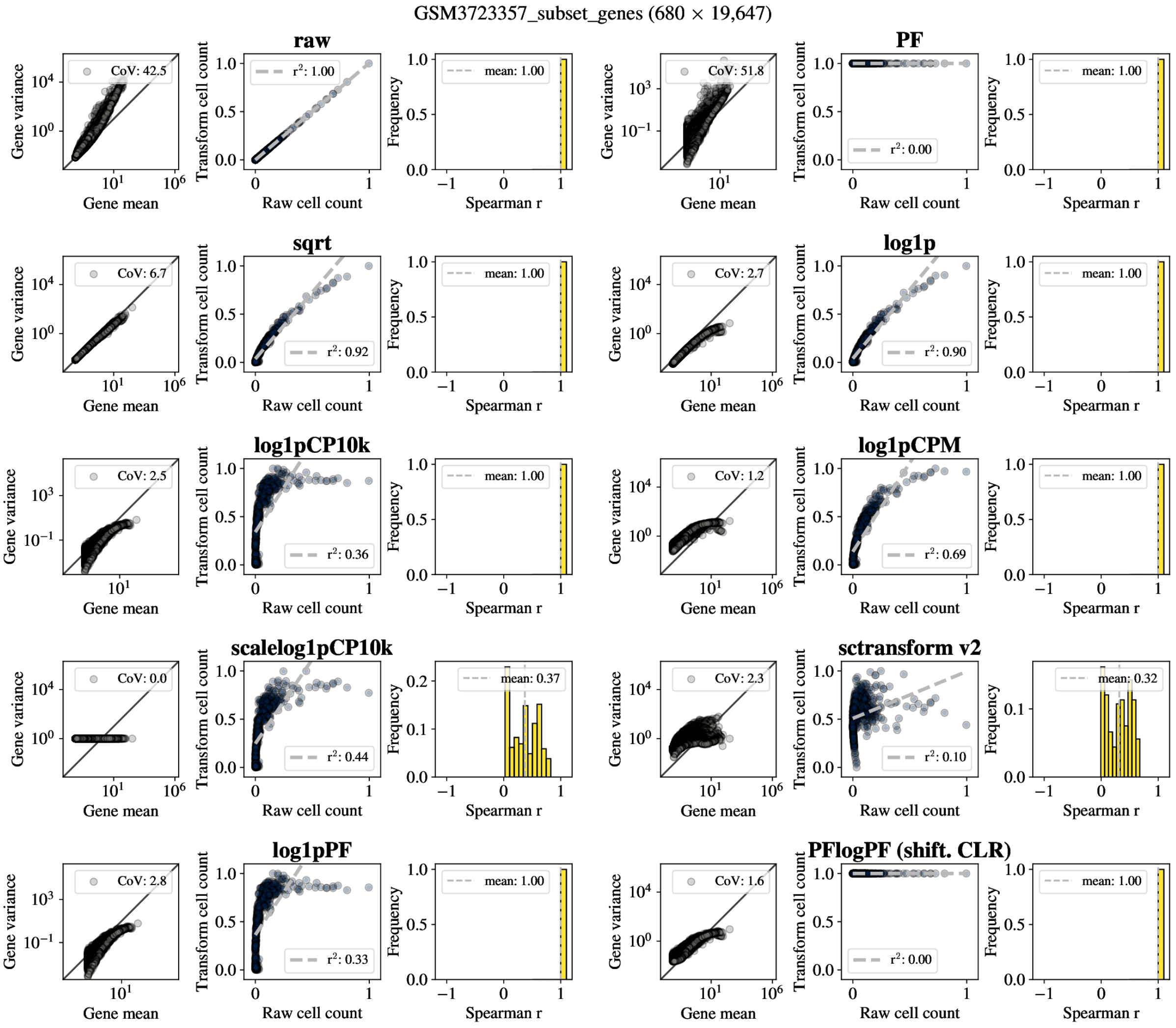


Supplementary Figure 1.379. **GSM3723357**. Per-method comparison of variance stabilization (left), depth normalization (middle), and monotonicity (right) for each normalization on dataset GSM3723357.

| GSM3723358 | |
|---|---|
| cells | 8,534 |
| genes | 60,623 |
| nonzero entries | 12,240,715 |
| density | 0.0237 |
| total counts | 41,029,001 |
| mean counts / cell | 4,808 |
| mean counts / gene | 677 |
| min counts / cell | 472 |
| max counts / cell | 63,619 |
| overdispersion | 1.06 |

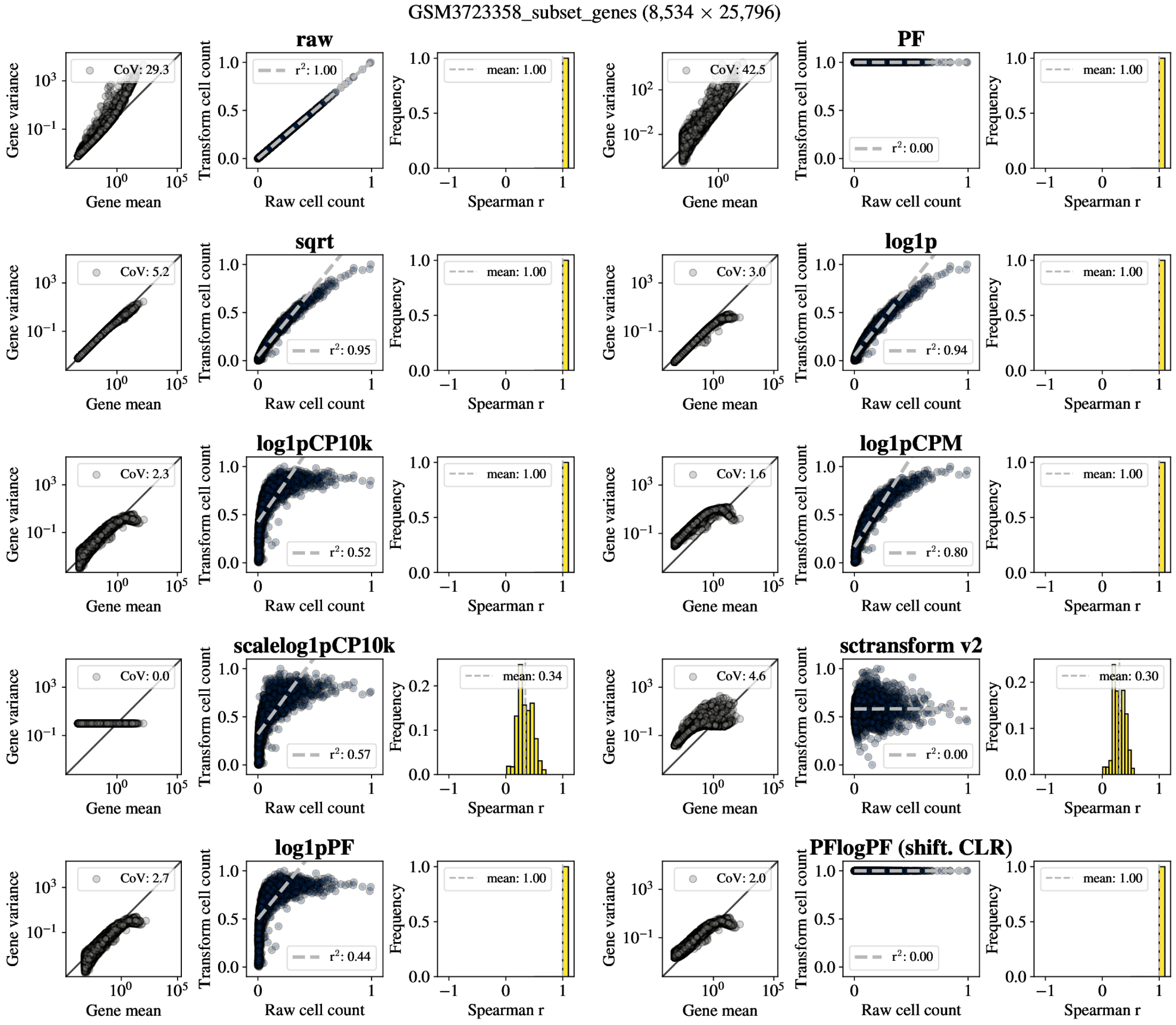


Supplementary Figure 1.380. **GSM3723358**. Per-method comparison of variance stabilization (left), depth normalization (middle), and monotonicity (right) for each normalization on dataset GSM3723358.

| GSM3723359 | |
|---|---|
| cells | 15,506 |
| genes | 60,623 |
| nonzero entries | 19,336,949 |
| density | 0.0206 |
| total counts | 58,260,087 |
| mean counts / cell | 3,757 |
| mean counts / gene | 961 |
| min counts / cell | 343 |
| max counts / cell | 35,617 |
| overdispersion | 0.54 |

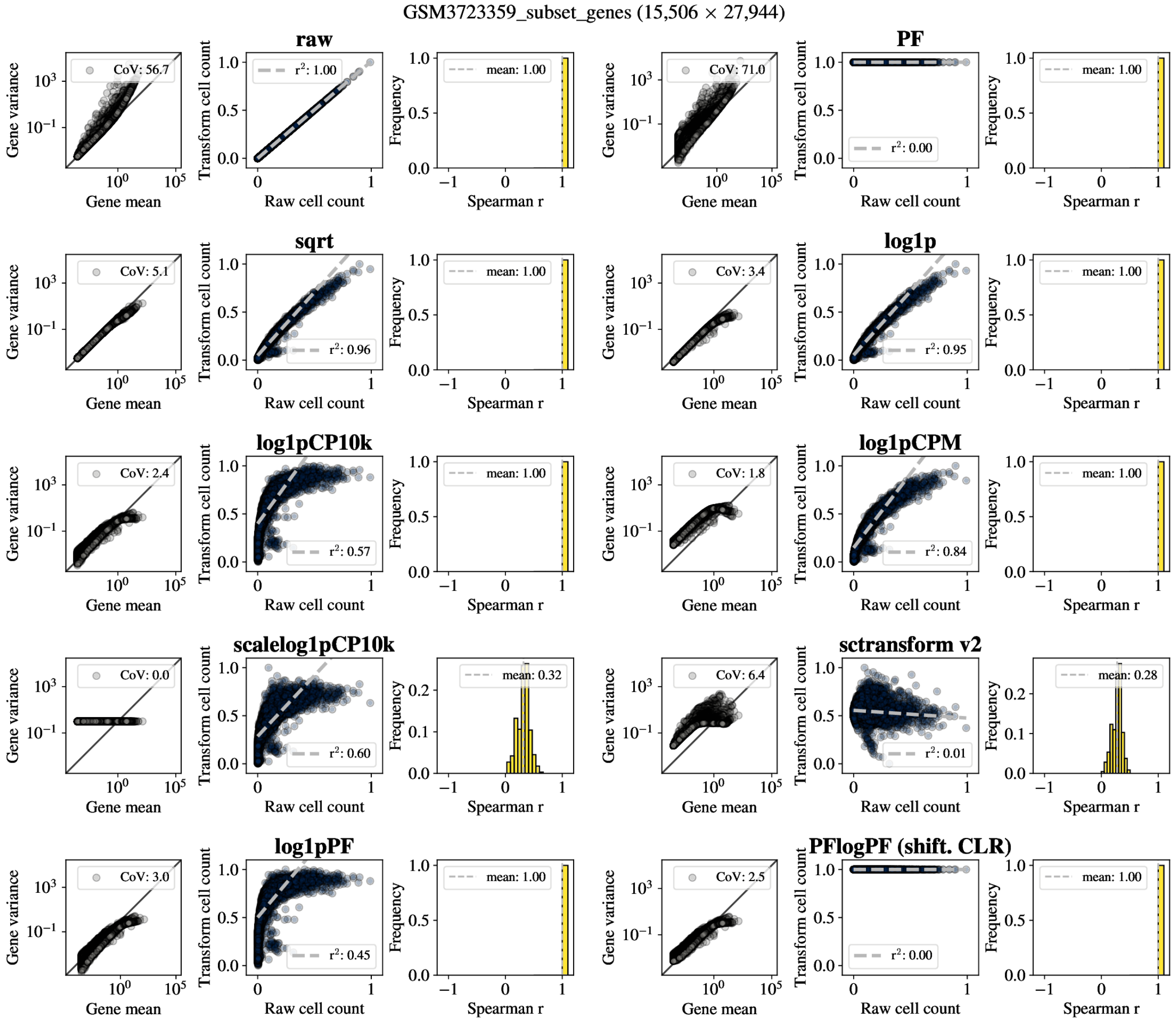


Supplementary Figure 1.381. **GSM3723359**. Per-method comparison of variance stabilization (left), depth normalization (middle), and monotonicity (right) for each normalization on dataset GSM3723359.

| GSM3731527 | |
|---|---|
| cells | 70,199 |
| genes | 60,623 |
| nonzero entries | 12,846,780 |
| density | 0.00302 |
| total counts | 42,846,890 |
| mean counts / cell | 610 |
| mean counts / gene | 707 |
| min counts / cell | 11 |
| max counts / cell | 50,478 |
| overdispersion | 19 |

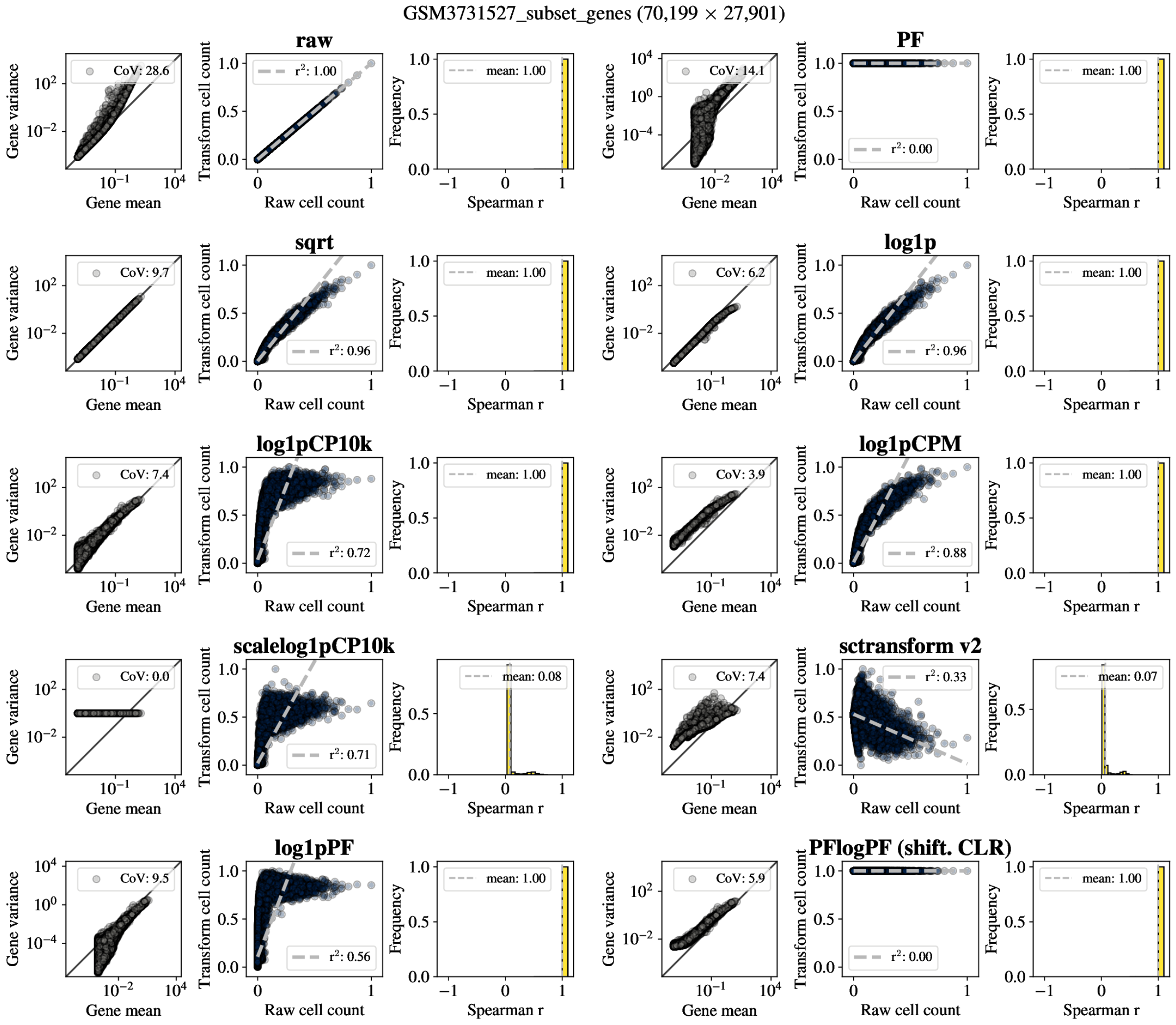


Supplementary Figure 1.382. **GSM3731527**. Per-method comparison of variance stabilization (left), depth normalization (middle), and monotonicity (right) for each normalization on dataset GSM3731527.

| GSM3732362 | |
|---|---|
| cells | 2,576 |
| genes | 60,623 |
| nonzero entries | 1,325,090 |
| density | 0.00849 |
| total counts | 3,110,323 |
| mean counts / cell | 1,207 |
| mean counts / gene | 51 |
| min counts / cell | 685 |
| max counts / cell | 3,753 |
| overdispersion | 0.122 |

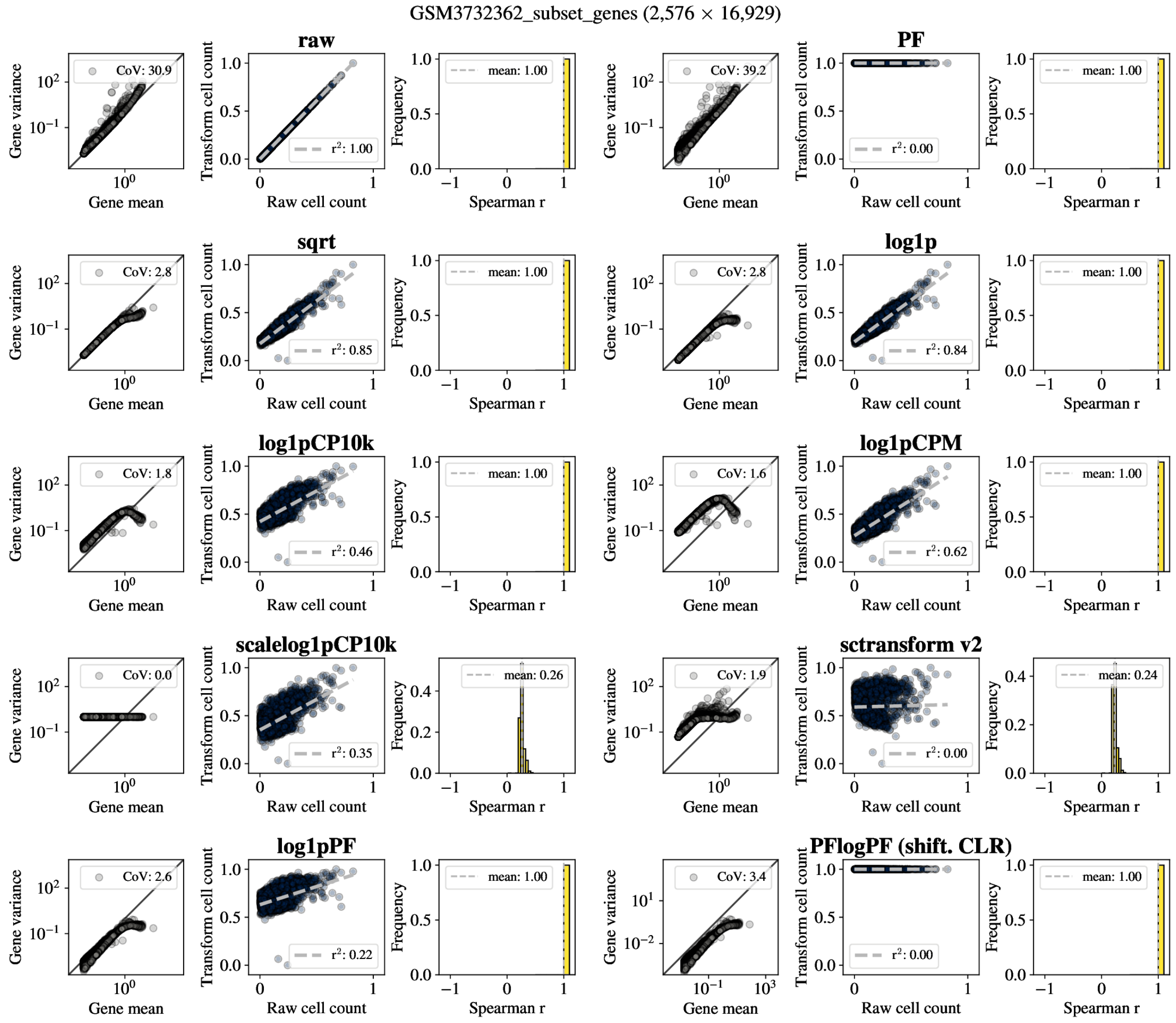


Supplementary Figure 1.383. **GSM3732362**. Per-method comparison of variance stabilization (left), depth normalization (middle), and monotonicity (right) for each normalization on dataset GSM3732362.

| GSM3732364 | |
|---|---|
| cells | 9,548 |
| genes | 60,623 |
| nonzero entries | 4,898,168 |
| density | 0.00846 |
| total counts | 12,267,639 |
| mean counts / cell | 1,285 |
| mean counts / gene | 202 |
| min counts / cell | 27 |
| max counts / cell | 16,830 |
| overdispersion | 1.73 |

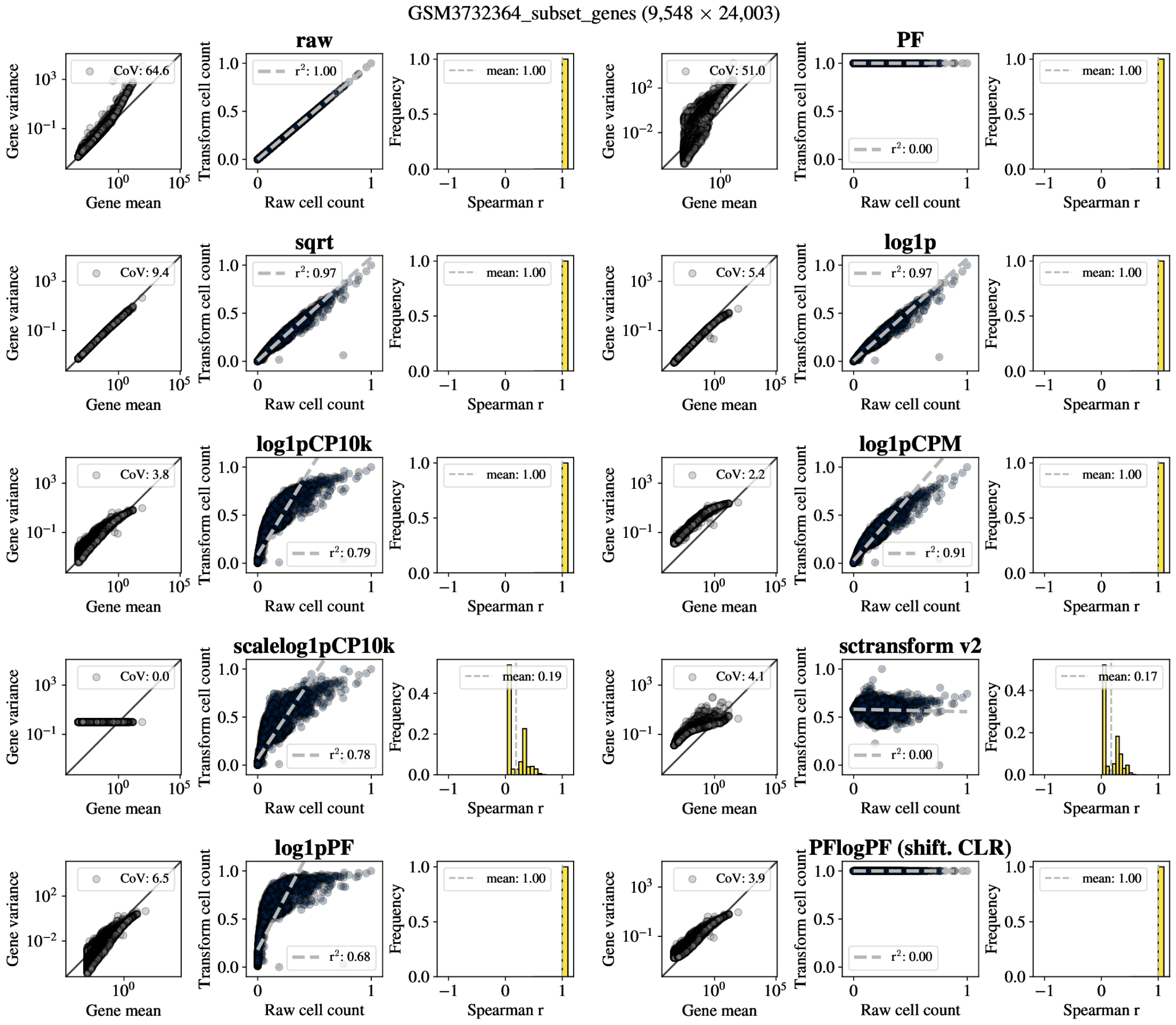


Supplementary Figure 1.384. **GSM3732364**. Per-method comparison of variance stabilization (left), depth normalization (middle), and monotonicity (right) for each normalization on dataset GSM3732364.

| GSM3732366 | |
|---|---|
| cells | 3,121 |
| genes | 60,623 |
| nonzero entries | 3,432,584 |
| density | 0.0181 |
| total counts | 10,238,609 |
| mean counts / cell | 3,281 |
| mean counts / gene | 169 |
| min counts / cell | 1,893 |
| max counts / cell | 16,287 |
| overdispersion | 0.146 |

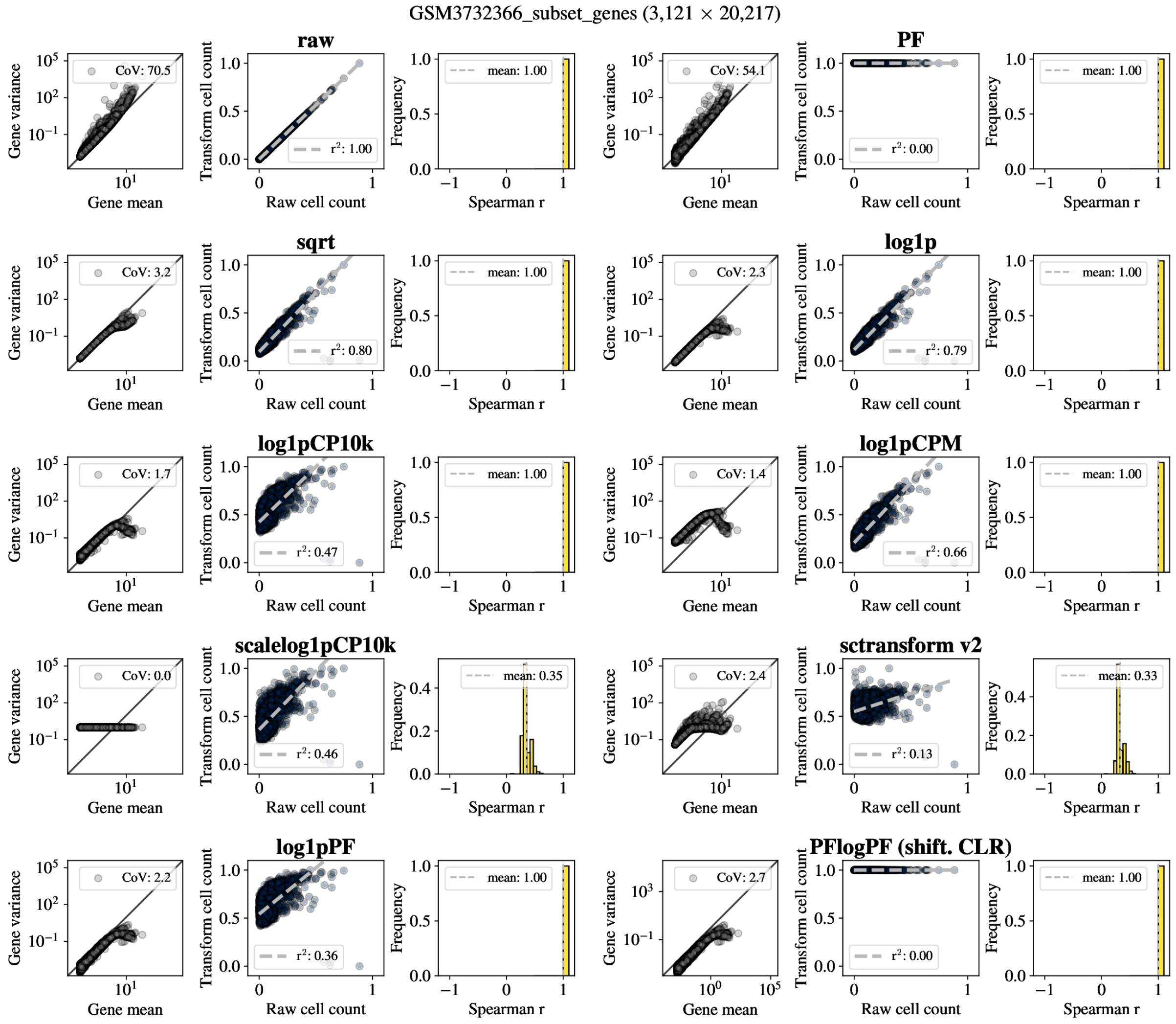


Supplementary Figure 1.385. **GSM3732366**. Per-method comparison of variance stabilization (left), depth normalization (middle), and monotonicity (right) for each normalization on dataset GSM3732366.

| GSM3732871 | |
|---|---|
| cells | 8,603 |
| genes | 60,623 |
| nonzero entries | 5,523,664 |
| density | 0.0106 |
| total counts | 12,273,900 |
| mean counts / cell | 1,427 |
| mean counts / gene | 202 |
| min counts / cell | 100 |
| max counts / cell | 38,512 |
| overdispersion | 3.49 |

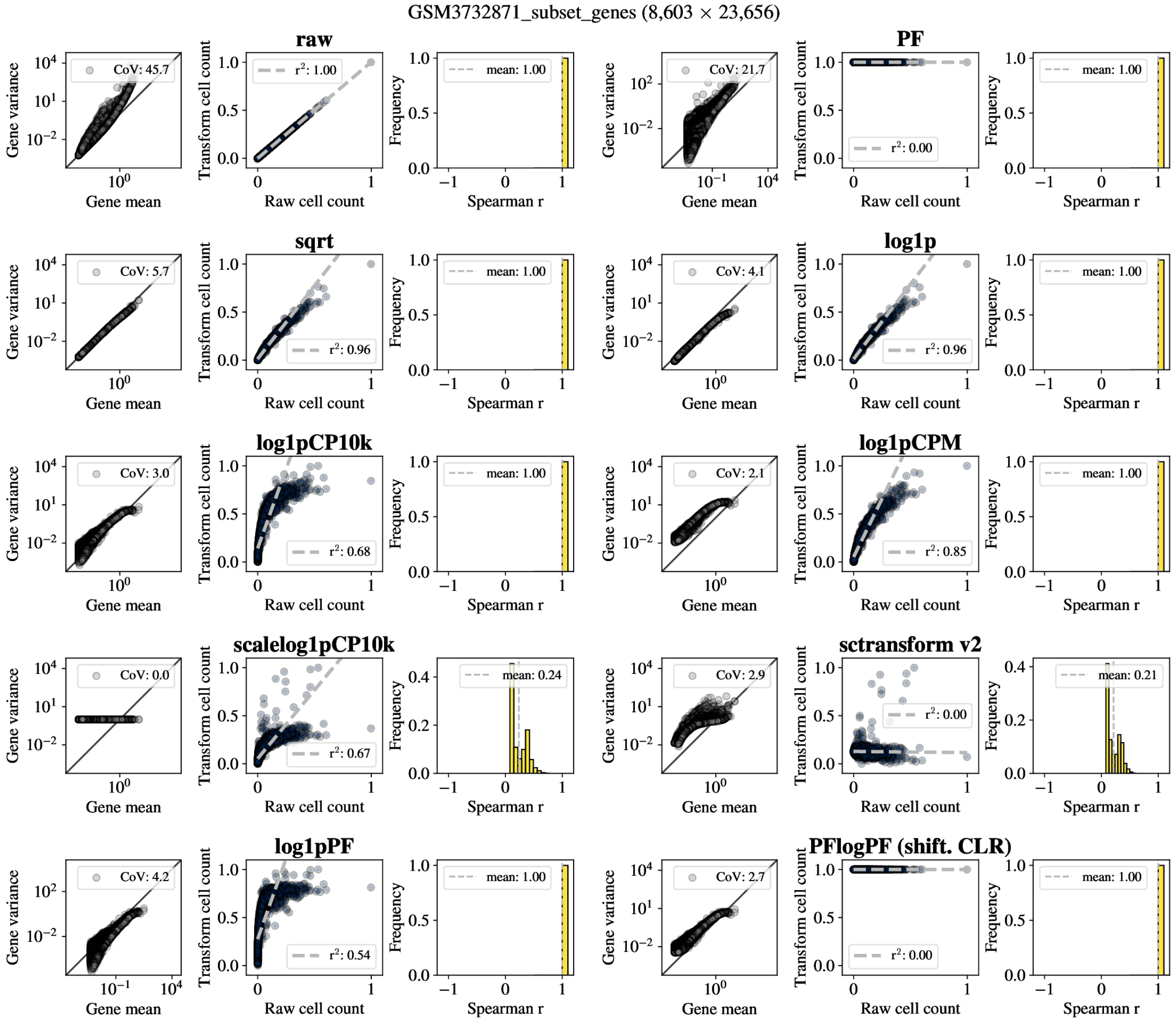


Supplementary Figure 1.386. **GSM3732871**. Per-method comparison of variance stabilization (left), depth normalization (middle), and monotonicity (right) for each normalization on dataset GSM3732871.

| GSM3734178 | |
|---|---|
| cells | 5,055 |
| genes | 60,623 |
| nonzero entries | 5,191,581 |
| density | 0.0169 |
| total counts | 15,549,073 |
| mean counts / cell | 3,076 |
| mean counts / gene | 256 |
| min counts / cell | 559 |
| max counts / cell | 18,879 |
| overdispersion | 0.206 |

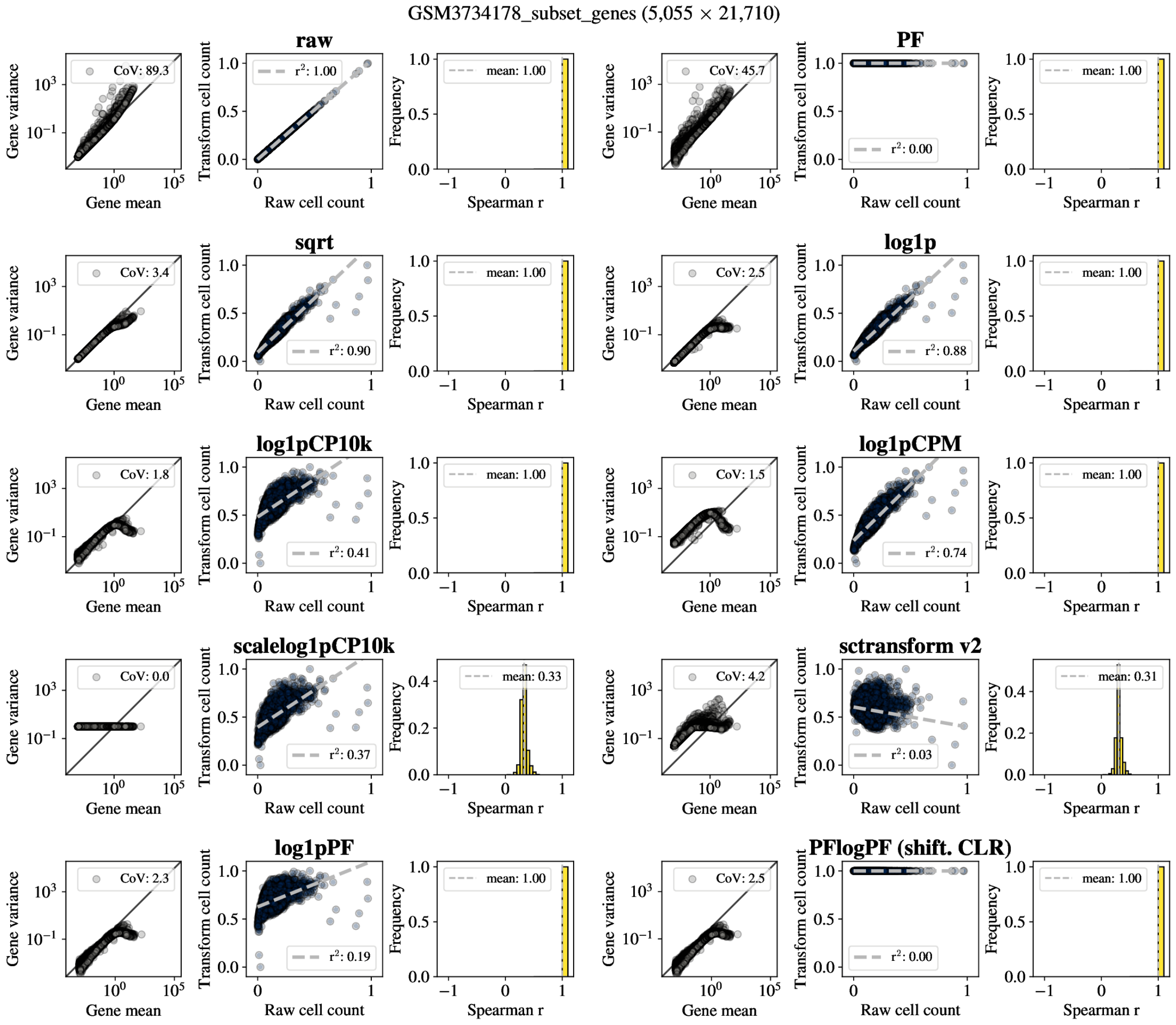


Supplementary Figure 1.387. **GSM3734178**. Per-method comparison of variance stabilization (left), depth normalization (middle), and monotonicity (right) for each normalization on dataset GSM3734178.

| GSM3734179 | |
|---|---|
| cells | 4,291 |
| genes | 60,623 |
| nonzero entries | 4,741,027 |
| density | 0.0182 |
| total counts | 13,318,935 |
| mean counts / cell | 3,104 |
| mean counts / gene | 220 |
| min counts / cell | 381 |
| max counts / cell | 33,643 |
| overdispersion | 0.216 |

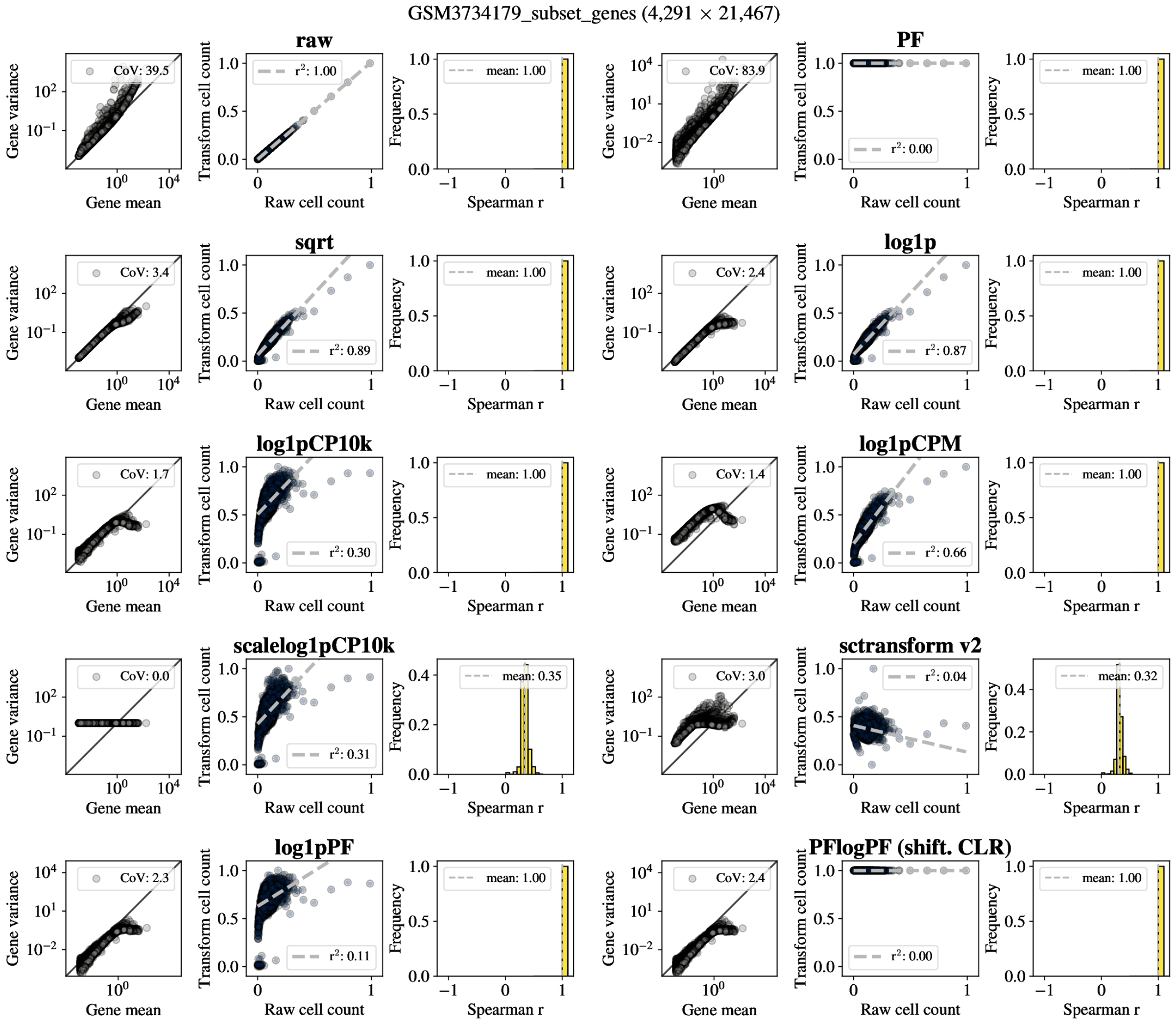


Supplementary Figure 1.388. **GSM3734179**. Per-method comparison of variance stabilization (left), depth normalization (middle), and monotonicity (right) for each normalization on dataset GSM3734179.

| GSM3734180 | |
|---|---|
| cells | 3,670 |
| genes | 60,623 |
| nonzero entries | 4,713,593 |
| density | 0.0212 |
| total counts | 13,575,006 |
| mean counts / cell | 3,699 |
| mean counts / gene | 224 |
| min counts / cell | 432 |
| max counts / cell | 20,265 |
| overdispersion | 0.259 |

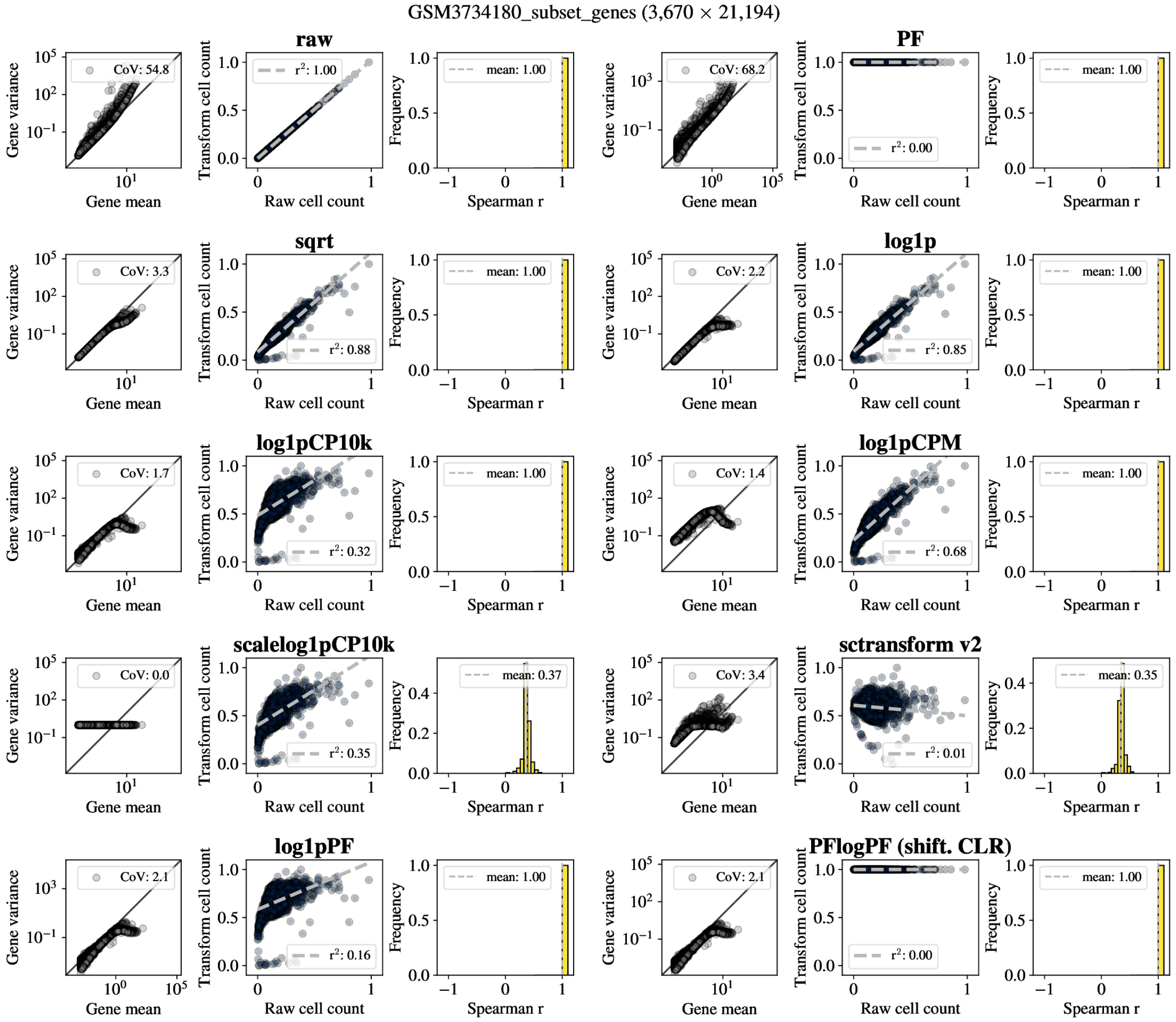


Supplementary Figure 1.389. **GSM3734180**. Per-method comparison of variance stabilization (left), depth normalization (middle), and monotonicity (right) for each normalization on dataset GSM3734180.

| GSM3734181 | |
|---|---|
| cells | 3,480 |
| genes | 60,623 |
| nonzero entries | 4,148,881 |
| density | 0.0197 |
| total counts | 12,153,044 |
| mean counts / cell | 3,492 |
| mean counts / gene | 200 |
| min counts / cell | 358 |
| max counts / cell | 23,241 |
| overdispersion | 0.201 |

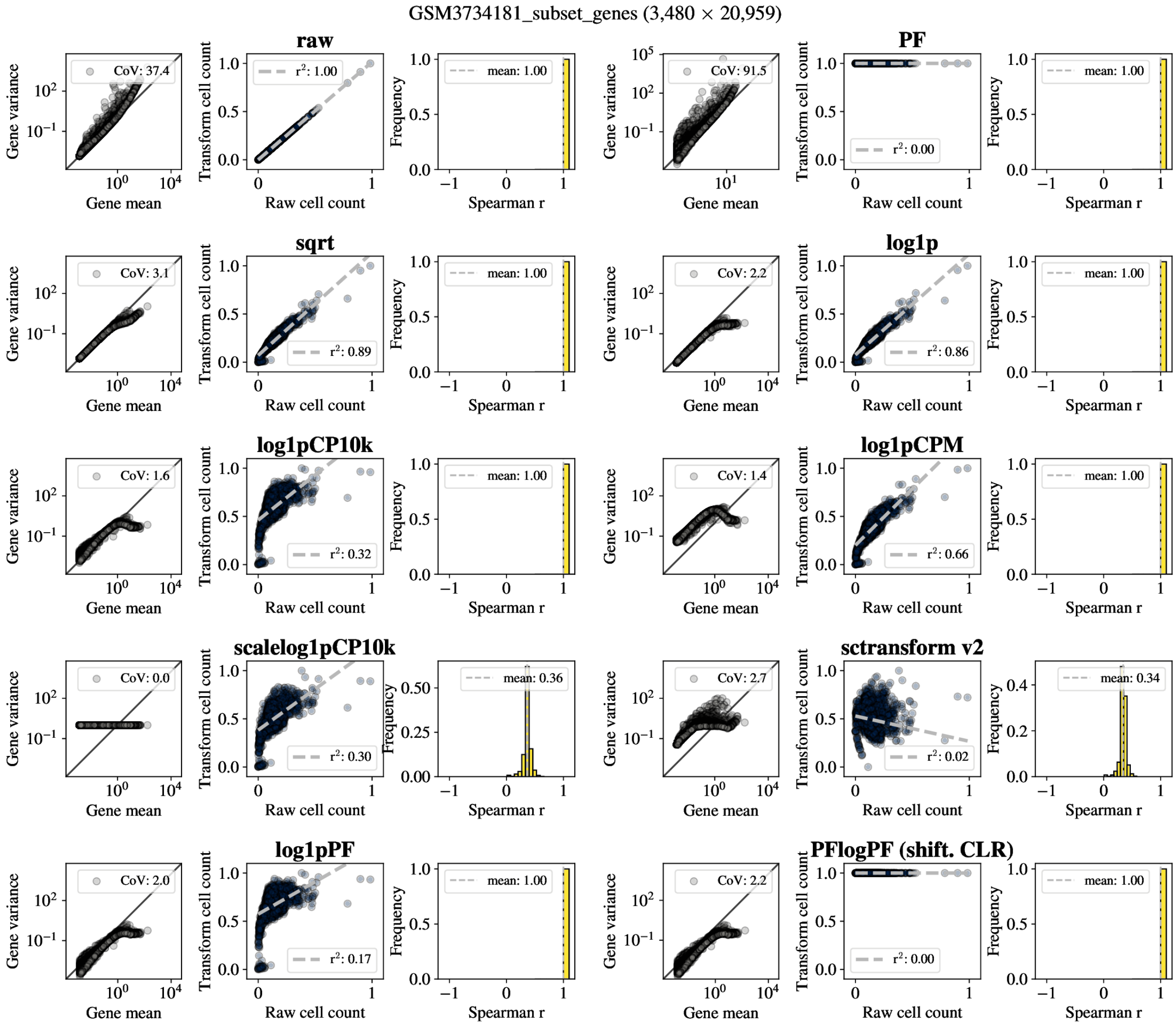


Supplementary Figure 1.390. **GSM3734181**. Per-method comparison of variance stabilization (left), depth normalization (middle), and monotonicity (right) for each normalization on dataset GSM3734181.

| GSM3734294 | |
|---|---|
| cells | 8,175 |
| genes | 60,623 |
| nonzero entries | 13,684,529 |
| density | 0.0276 |
| total counts | 37,412,424 |
| mean counts / cell | 4,576 |
| mean counts / gene | 617 |
| min counts / cell | 189 |
| max counts / cell | 44,954 |
| overdispersion | 0.98 |

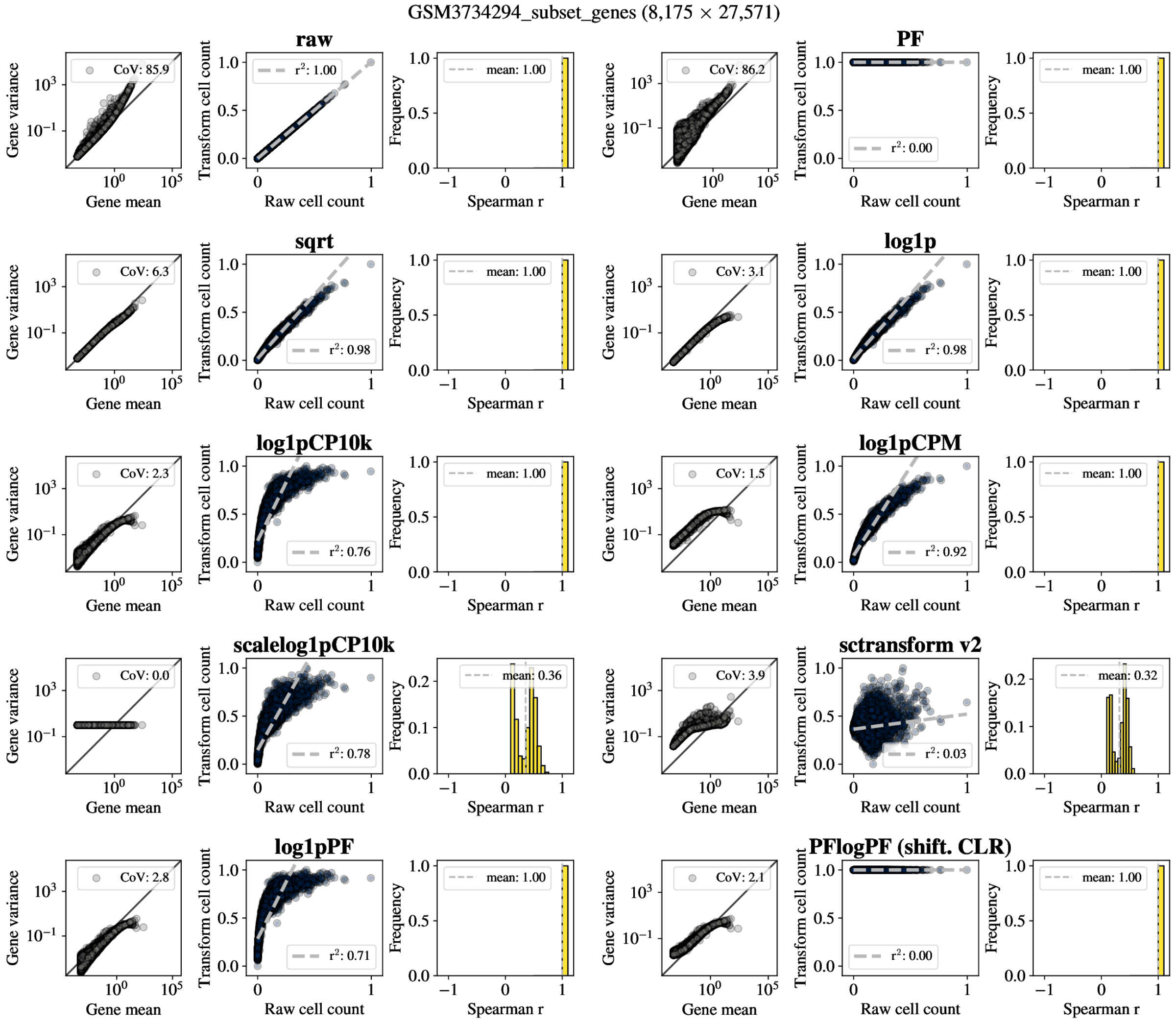


Supplementary Figure 1.391. **GSM3734294**. Per-method comparison of variance stabilization (left), depth normalization (middle), and monotonicity (right) for each normalization on dataset GSM3734294.

| GSM3734295 | |
|---|---|
| cells | 5,932 |
| genes | 60,623 |
| nonzero entries | 8,060,191 |
| density | 0.0224 |
| total counts | 24,938,999 |
| mean counts / cell | 4,204 |
| mean counts / gene | 411 |
| min counts / cell | 60 |
| max counts / cell | 49,846 |
| overdispersion | 1.44 |

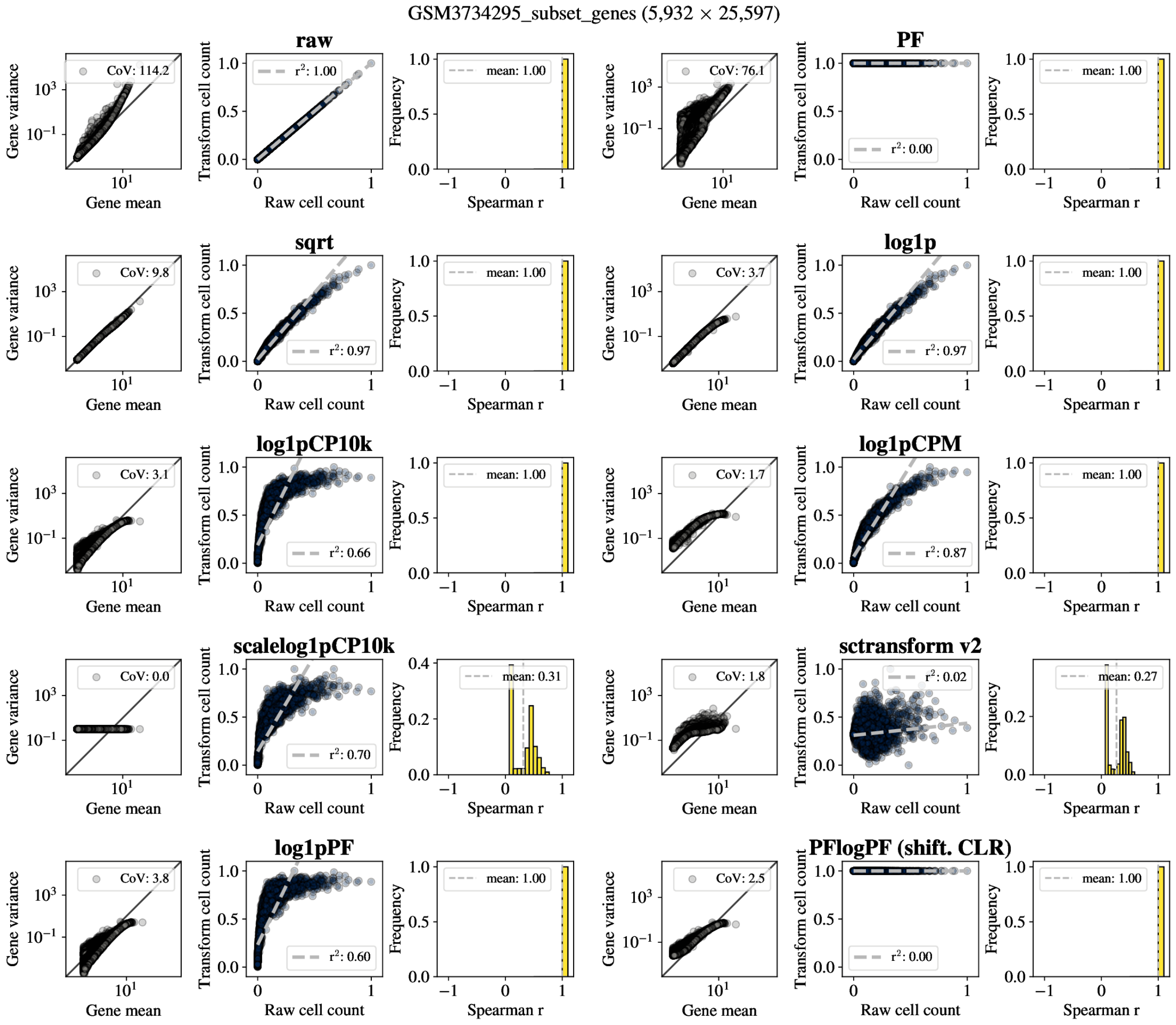


Supplementary Figure 1.392. **GSM3734295**. Per-method comparison of variance stabilization (left), depth normalization (middle), and monotonicity (right) for each normalization on dataset GSM3734295.

| GSM3734296 | |
|---|---|
| cells | 4,933 |
| genes | 60,623 |
| nonzero entries | 7,683,791 |
| density | 0.0257 |
| total counts | 17,954,801 |
| mean counts / cell | 3,640 |
| mean counts / gene | 296 |
| min counts / cell | 875 |
| max counts / cell | 20,459 |
| overdispersion | 0.466 |

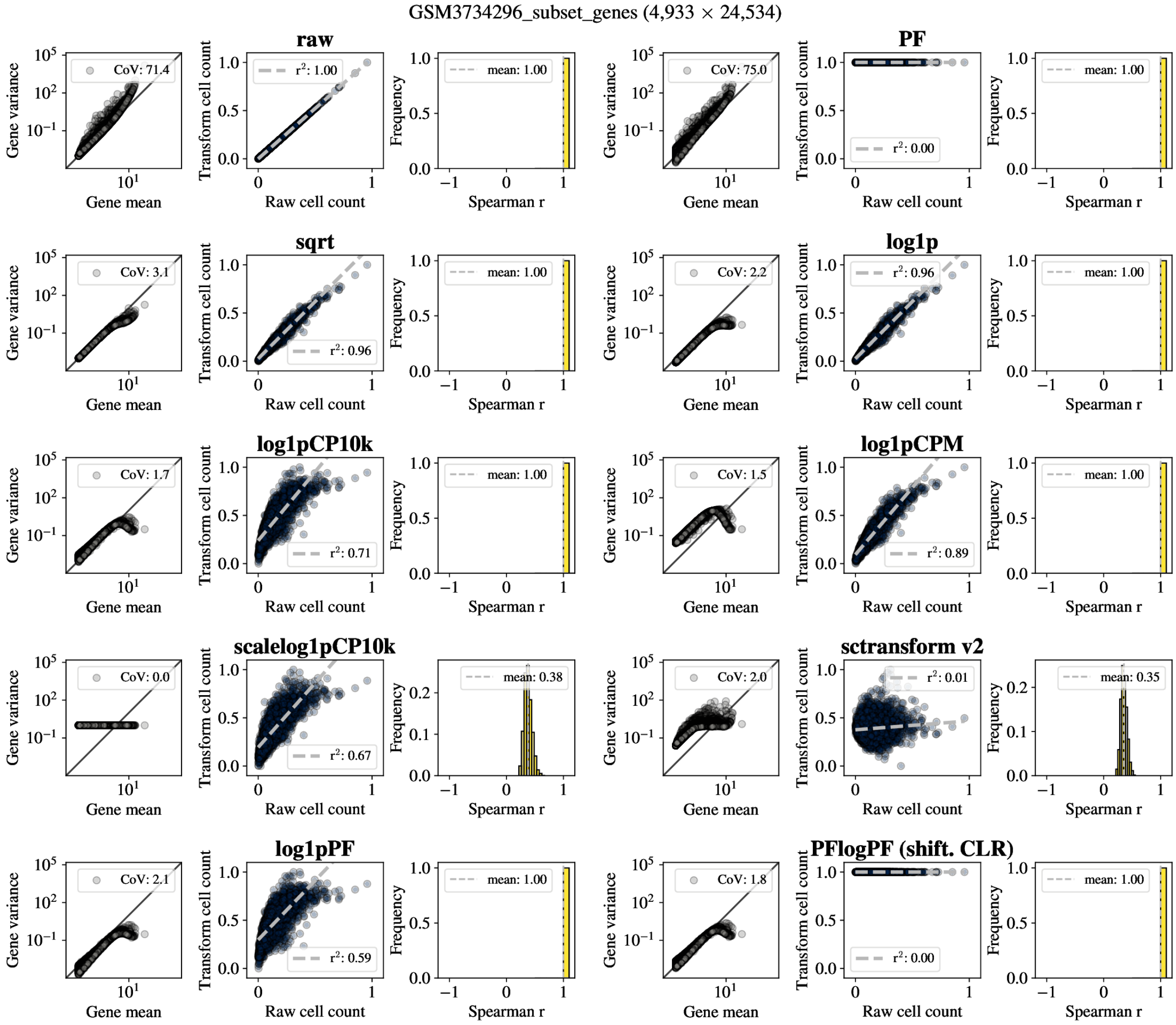


Supplementary Figure 1.393. **GSM3734296**. Per-method comparison of variance stabilization (left), depth normalization (middle), and monotonicity (right) for each normalization on dataset GSM3734296.

| GSM3734297 | |
|---|---|
| cells | 3,743 |
| genes | 60,623 |
| nonzero entries | 4,942,052 |
| density | 0.0218 |
| total counts | 11,485,083 |
| mean counts / cell | 3,068 |
| mean counts / gene | 189 |
| min counts / cell | 555 |
| max counts / cell | 19,831 |
| overdispersion | 0.492 |

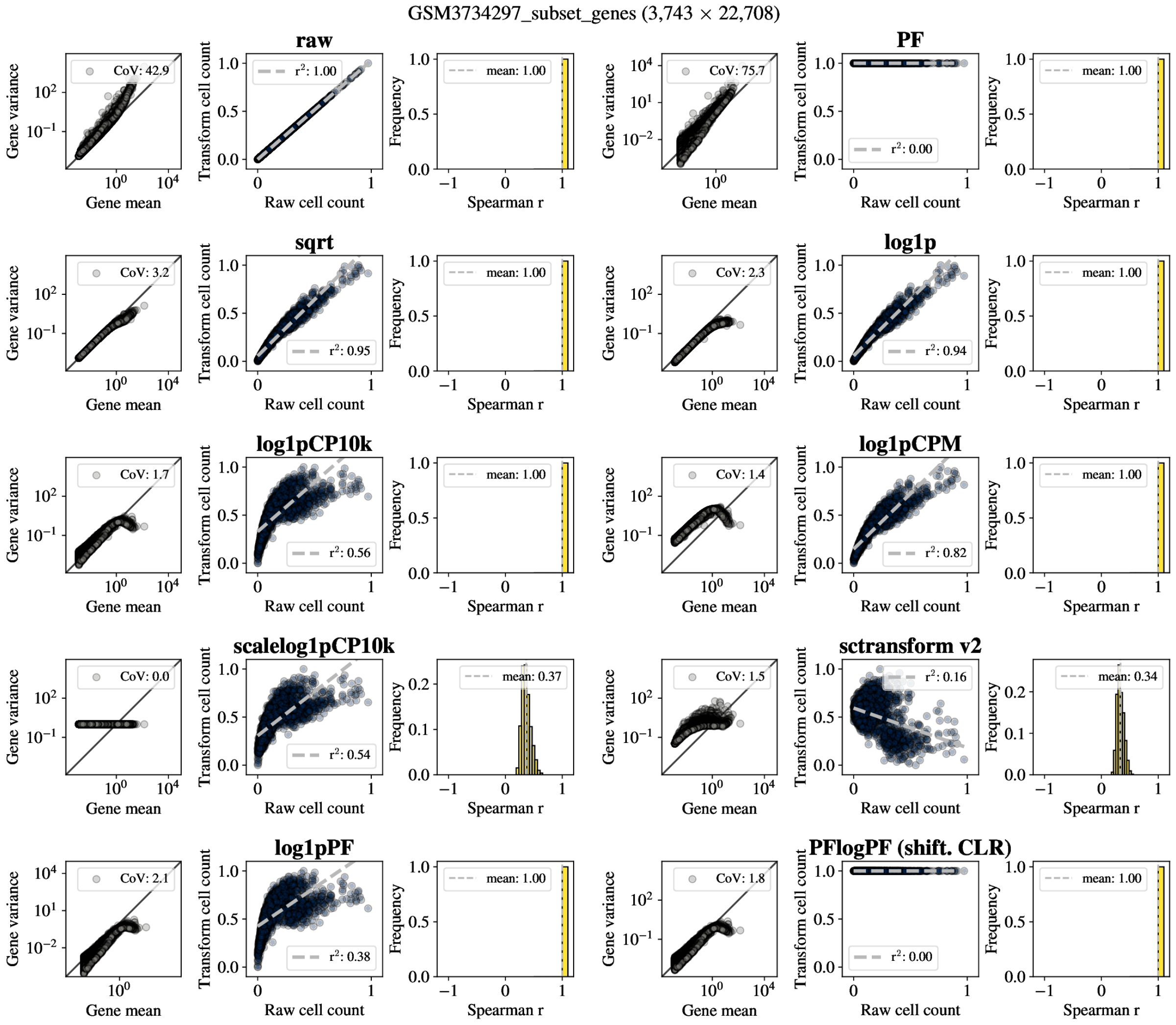


Supplementary Figure 1.394. **GSM3734297**. Per-method comparison of variance stabilization (left), depth normalization (middle), and monotonicity (right) for each normalization on dataset GSM3734297.

| GSM3735993 | |
|---|---|
| cells | 9,240 |
| genes | 60,623 |
| nonzero entries | 17,861,129 |
| density | 0.0319 |
| total counts | 70,899,230 |
| mean counts / cell | 7,673 |
| mean counts / gene | 1,170 |
| min counts / cell | 329 |
| max counts / cell | 50,112 |
| overdispersion | 1.77 |

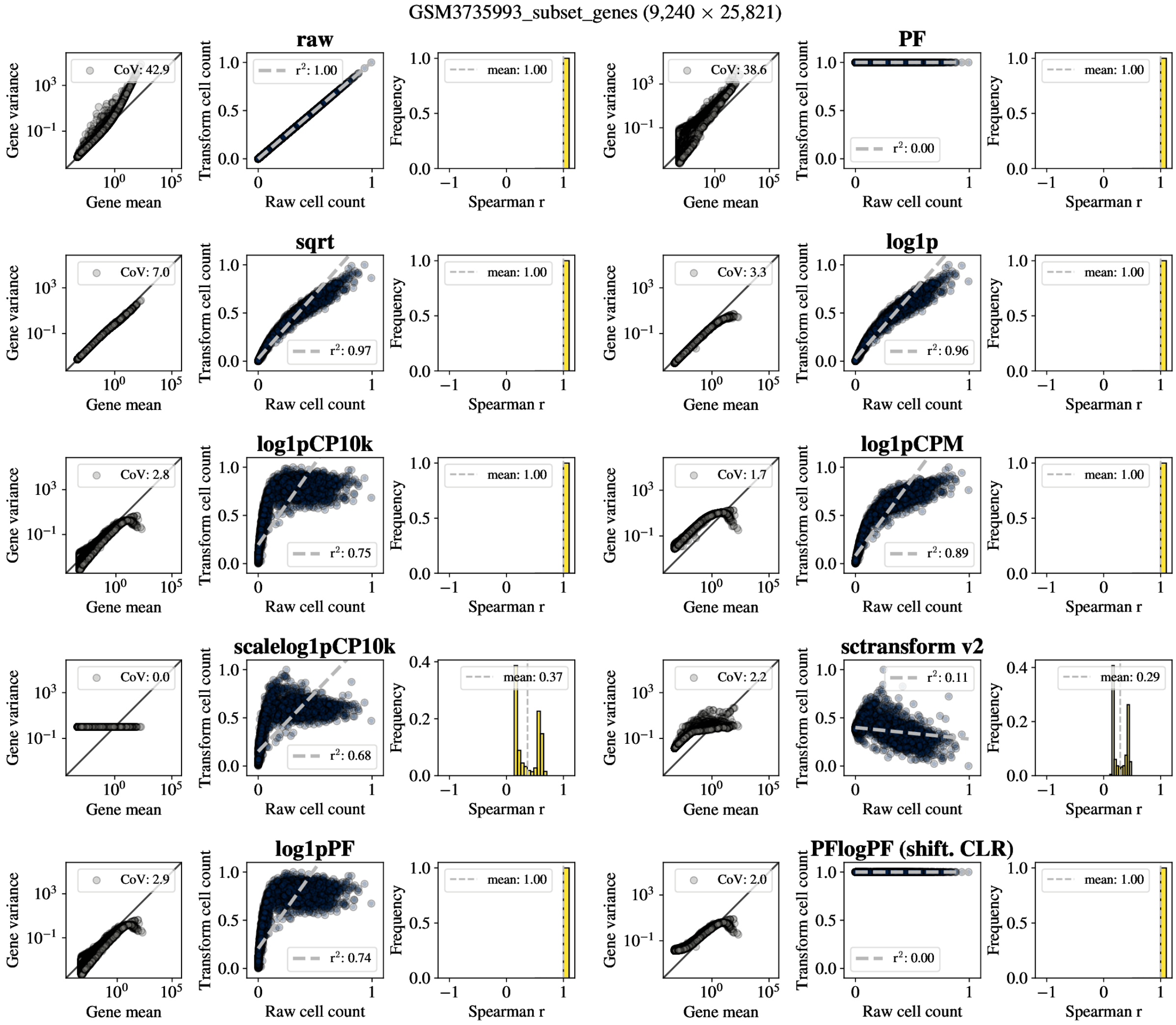


Supplementary Figure 1.395. **GSM3735993**. Per-method comparison of variance stabilization (left), depth normalization (middle), and monotonicity (right) for each normalization on dataset GSM3735993.

| GSM3735994 | |
|---|---|
| cells | 6,561 |
| genes | 60,623 |
| nonzero entries | 24,833,240 |
| density | 0.0624 |
| total counts | 133,262,750 |
| mean counts / cell | 20,311 |
| mean counts / gene | 2,198 |
| min counts / cell | 2,404 |
| max counts / cell | 73,862 |
| overdispersion | 0.374 |

GSM3735994_subset_genes (6,561 × 27,371)

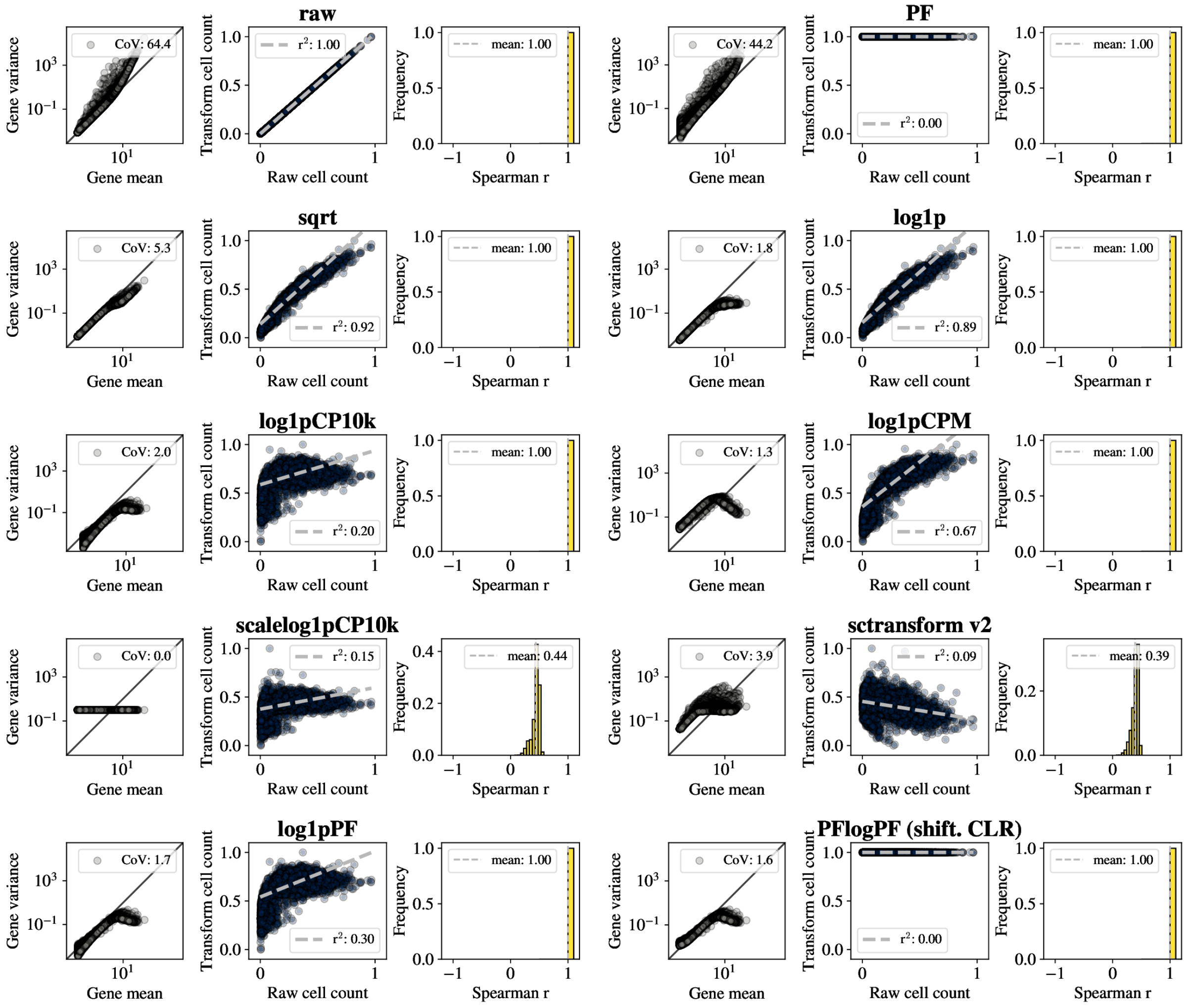


Supplementary Figure 1.396. **GSM3735994**. Per-method comparison of variance stabilization (left), depth normalization (middle), and monotonicity (right) for each normalization on dataset GSM3735994.

| GSM3738536 | |
|---|---|
| cells | 827 |
| genes | 60,623 |
| nonzero entries | 1,246,093 |
| density | 0.0249 |
| total counts | 3,510,175 |
| mean counts / cell | 4,244 |
| mean counts / gene | 58 |
| min counts / cell | 2,583 |
| max counts / cell | 24,127 |
| overdispersion | 0.101 |

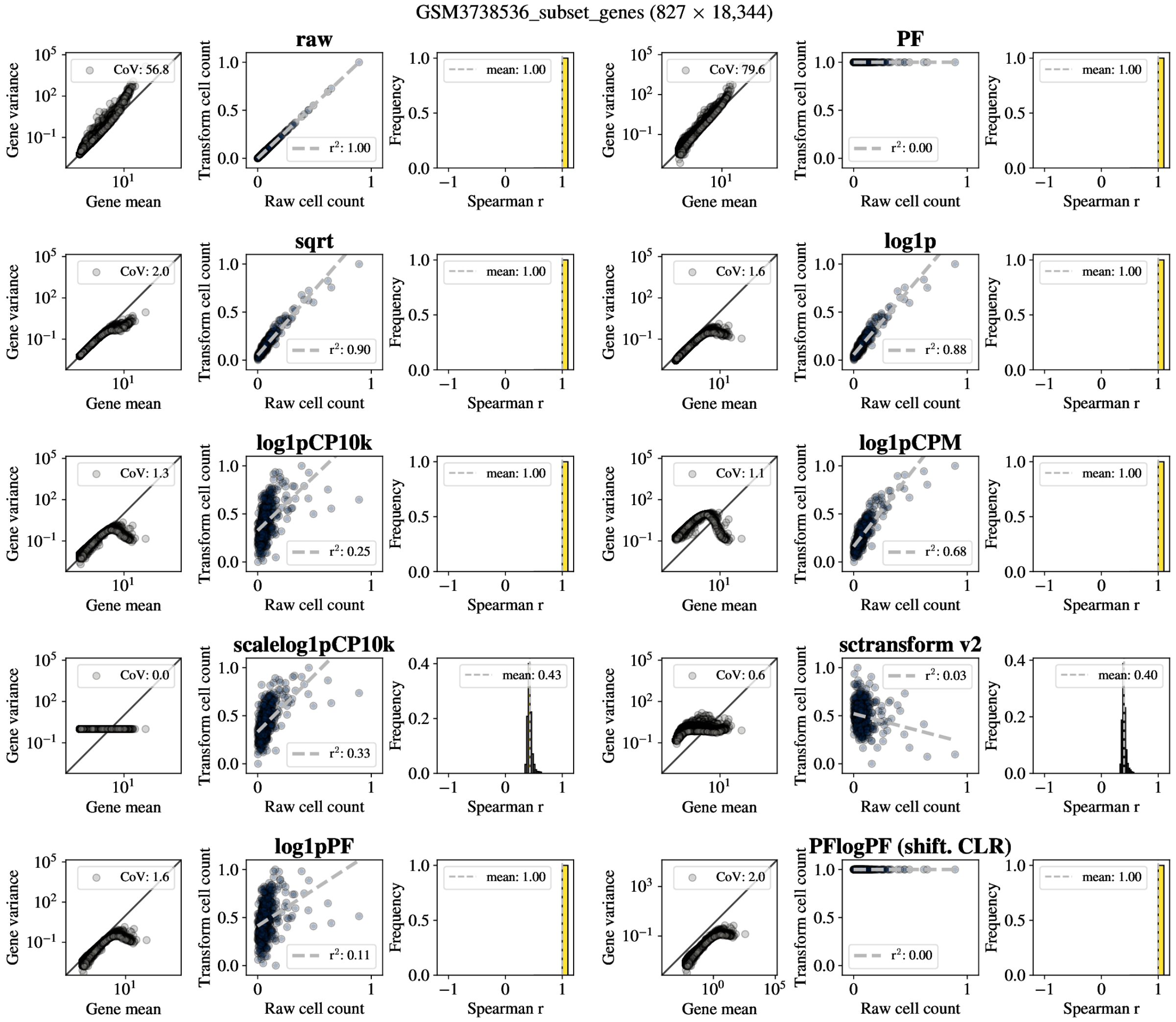


Supplementary Figure 1.397. **GSM3738536**. Per-method comparison of variance stabilization (left), depth normalization (middle), and monotonicity (right) for each normalization on dataset GSM3738536.

| GSM3738537 | |
|---|---|
| cells | 853 |
| genes | 60,623 |
| nonzero entries | 1,353,148 |
| density | 0.0262 |
| total counts | 3,726,034 |
| mean counts / cell | 4,368 |
| mean counts / gene | 61 |
| min counts / cell | 2,554 |
| max counts / cell | 20,713 |
| overdispersion | 0.123 |

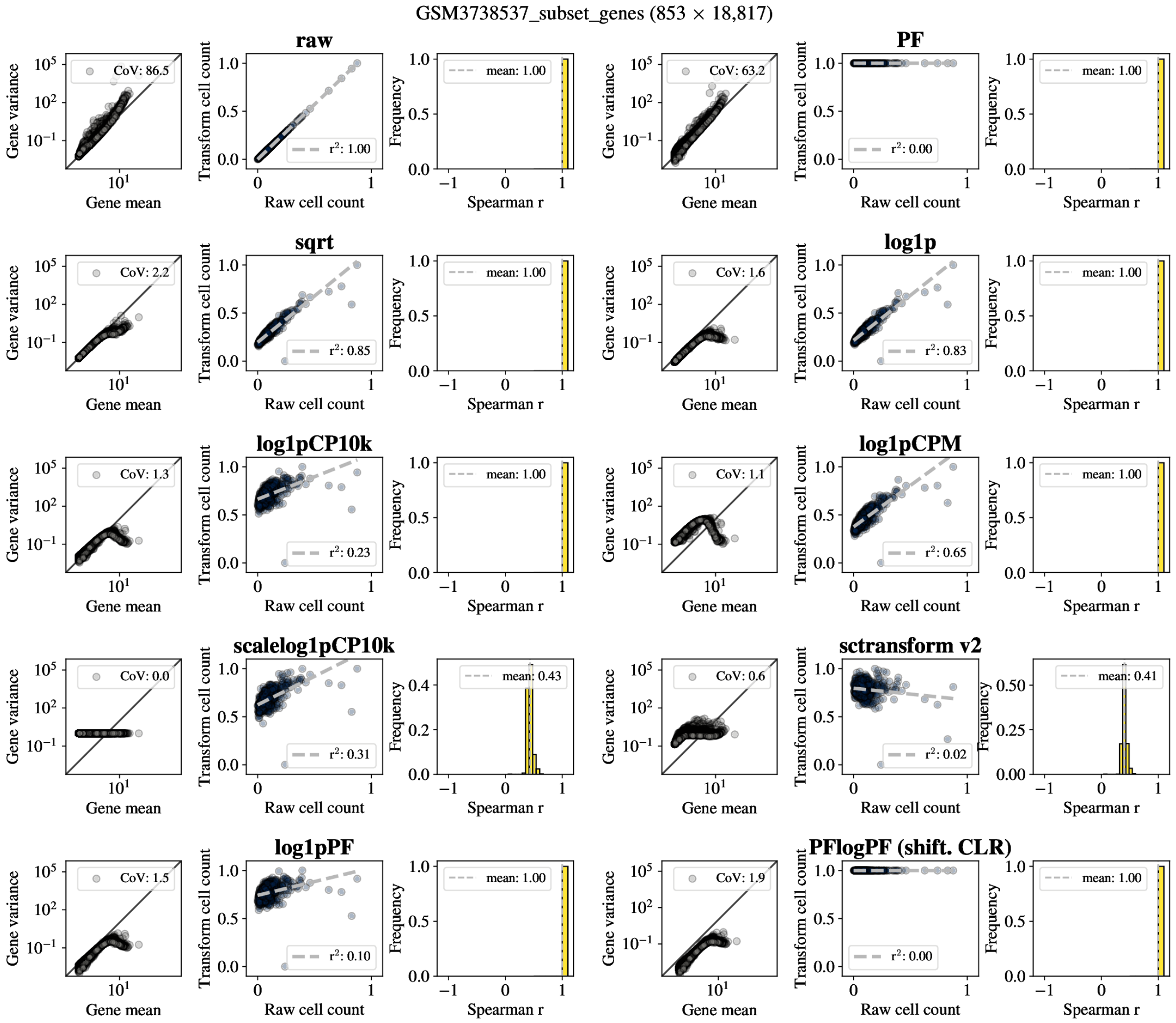


Supplementary Figure 1.398. **GSM3738537**. Per-method comparison of variance stabilization (left), depth normalization (middle), and monotonicity (right) for each normalization on dataset GSM3738537.

| GSM3738538 | |
|---|---|
| cells | 1,294 |
| genes | 60,623 |
| nonzero entries | 2,541,261 |
| density | 0.0324 |
| total counts | 6,873,291 |
| mean counts / cell | 5,312 |
| mean counts / gene | 113 |
| min counts / cell | 383 |
| max counts / cell | 40,703 |
| overdispersion | 0.103 |

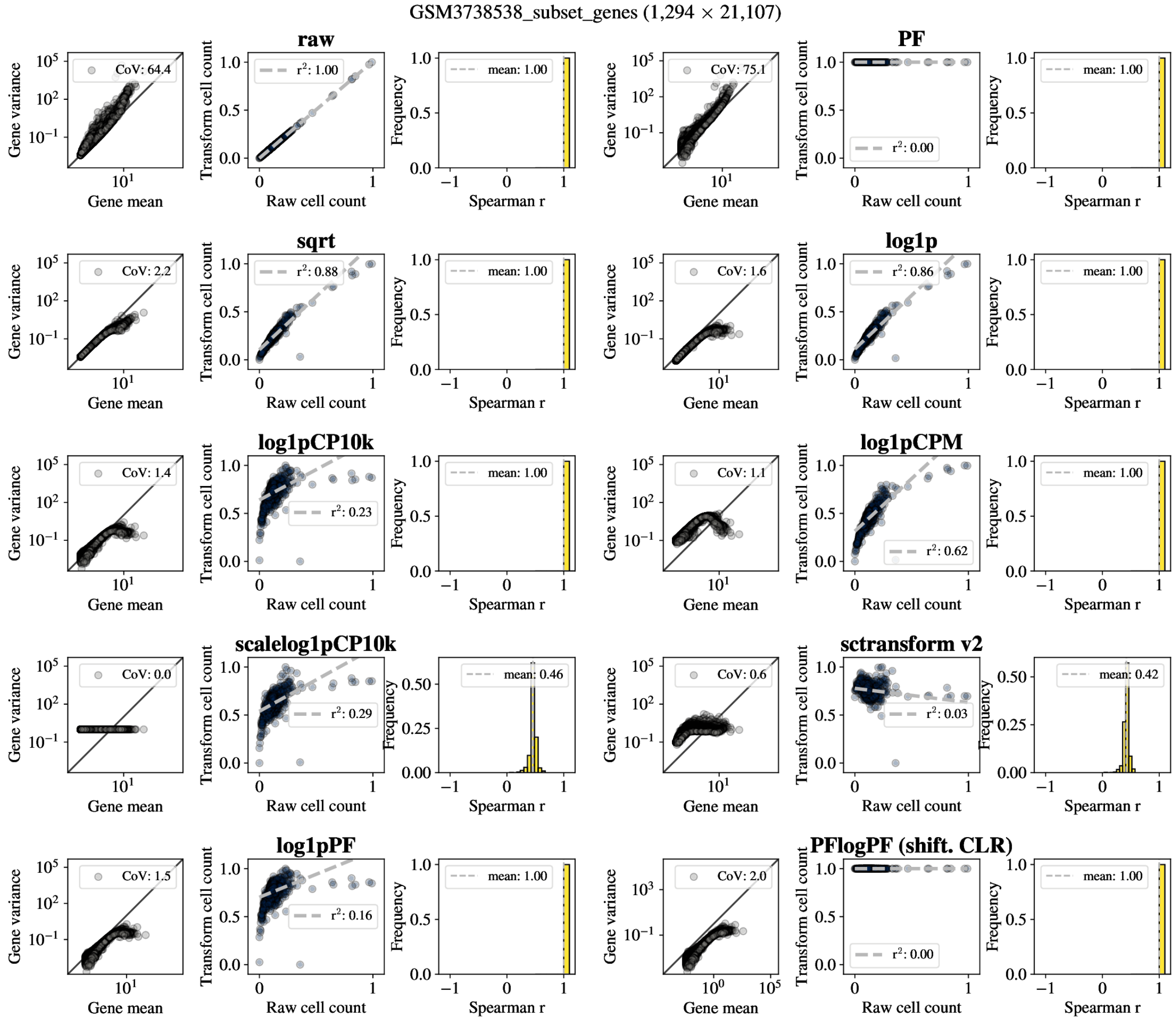


Supplementary Figure 1.399. **GSM3738538**. Per-method comparison of variance stabilization (left), depth normalization (middle), and monotonicity (right) for each normalization on dataset GSM3738538.

| GSM3738539 | |
|---|---|
| cells | 1,497 |
| genes | 60,623 |
| nonzero entries | 2,512,617 |
| density | 0.0277 |
| total counts | 7,037,645 |
| mean counts / cell | 4,701 |
| mean counts / gene | 116 |
| min counts / cell | 104 |
| max counts / cell | 65,629 |
| overdispersion | 0.208 |

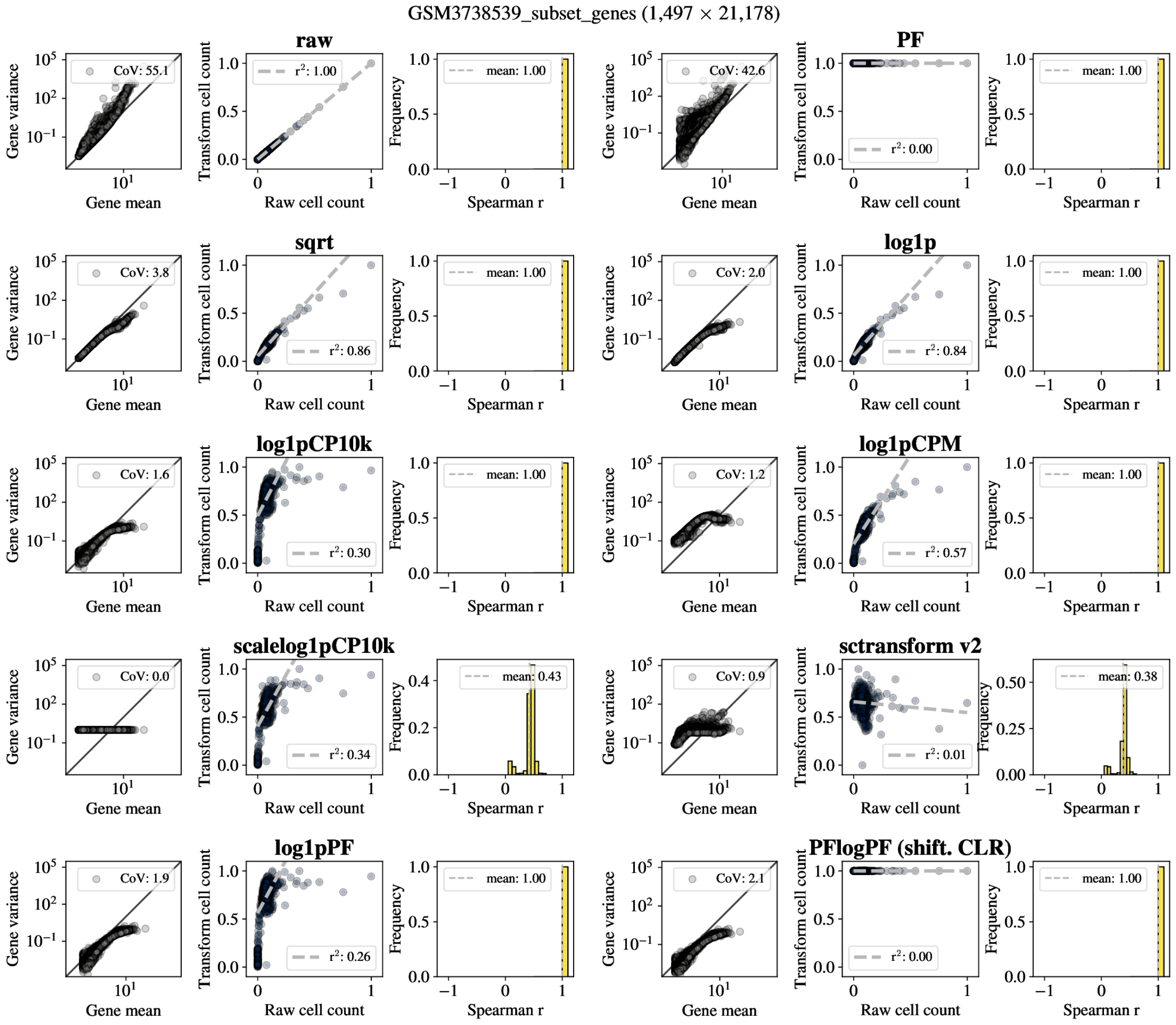


Supplementary Figure 1.400. **GSM3738539**. Per-method comparison of variance stabilization (left), depth normalization (middle), and monotonicity (right) for each normalization on dataset GSM3738539.

| GSM3738540 | |
|---|---|
| cells | 904 |
| genes | 60,623 |
| nonzero entries | 1,321,074 |
| density | 0.0241 |
| total counts | 3,364,794 |
| mean counts / cell | 3,722 |
| mean counts / gene | 56 |
| min counts / cell | 2,414 |
| max counts / cell | 13,159 |
| overdispersion | 0.0569 |

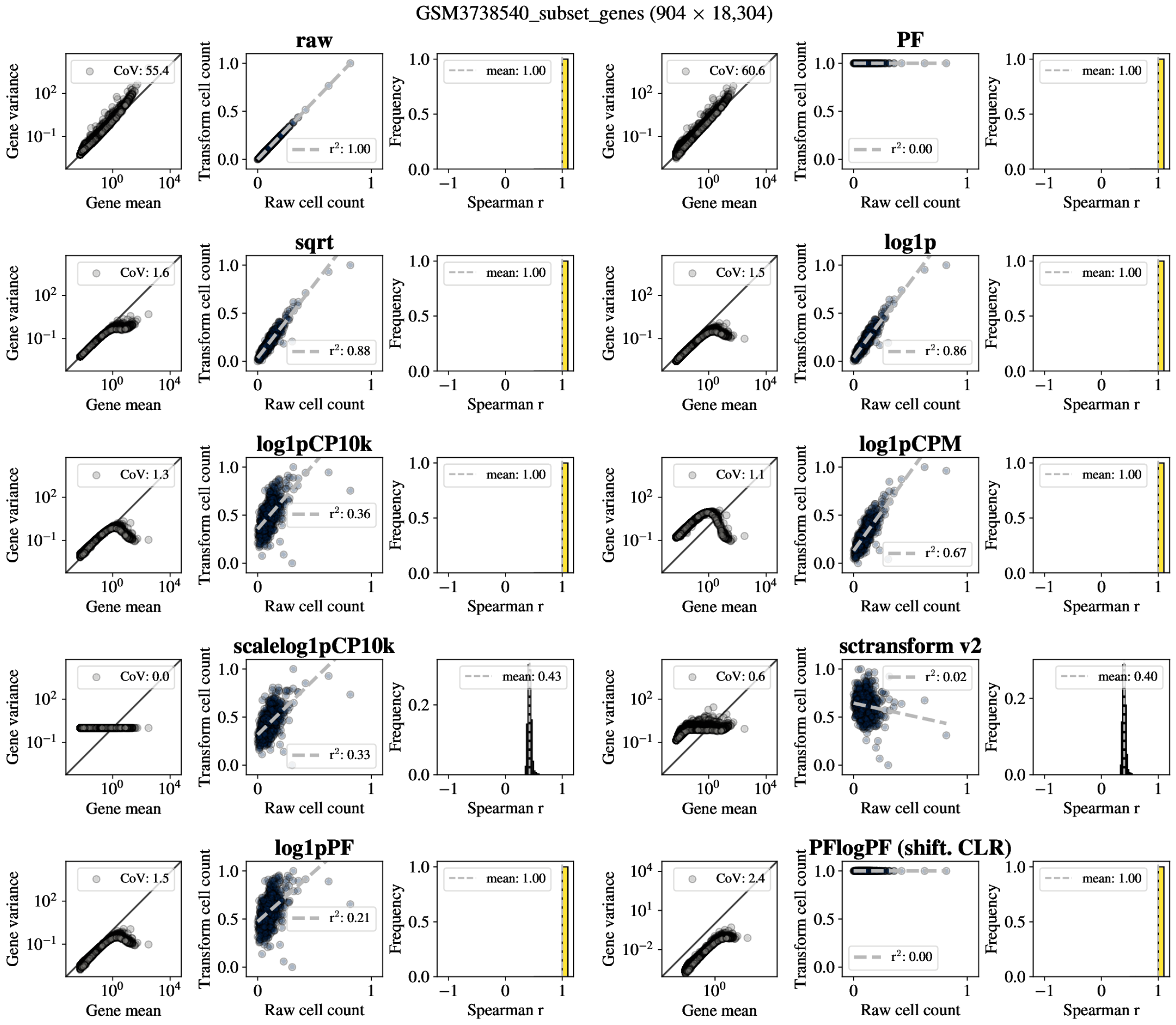


Supplementary Figure 1.401. **GSM3738540**. Per-method comparison of variance stabilization (left), depth normalization (middle), and monotonicity (right) for each normalization on dataset GSM3738540.

| GSM3738541 | |
|---|---|
| cells | 1,319 |
| genes | 60,623 |
| nonzero entries | 1,218,489 |
| density | 0.0152 |
| total counts | 3,208,217 |
| mean counts / cell | 2,432 |
| mean counts / gene | 53 |
| min counts / cell | 49 |
| max counts / cell | 36,099 |
| overdispersion | 0.807 |

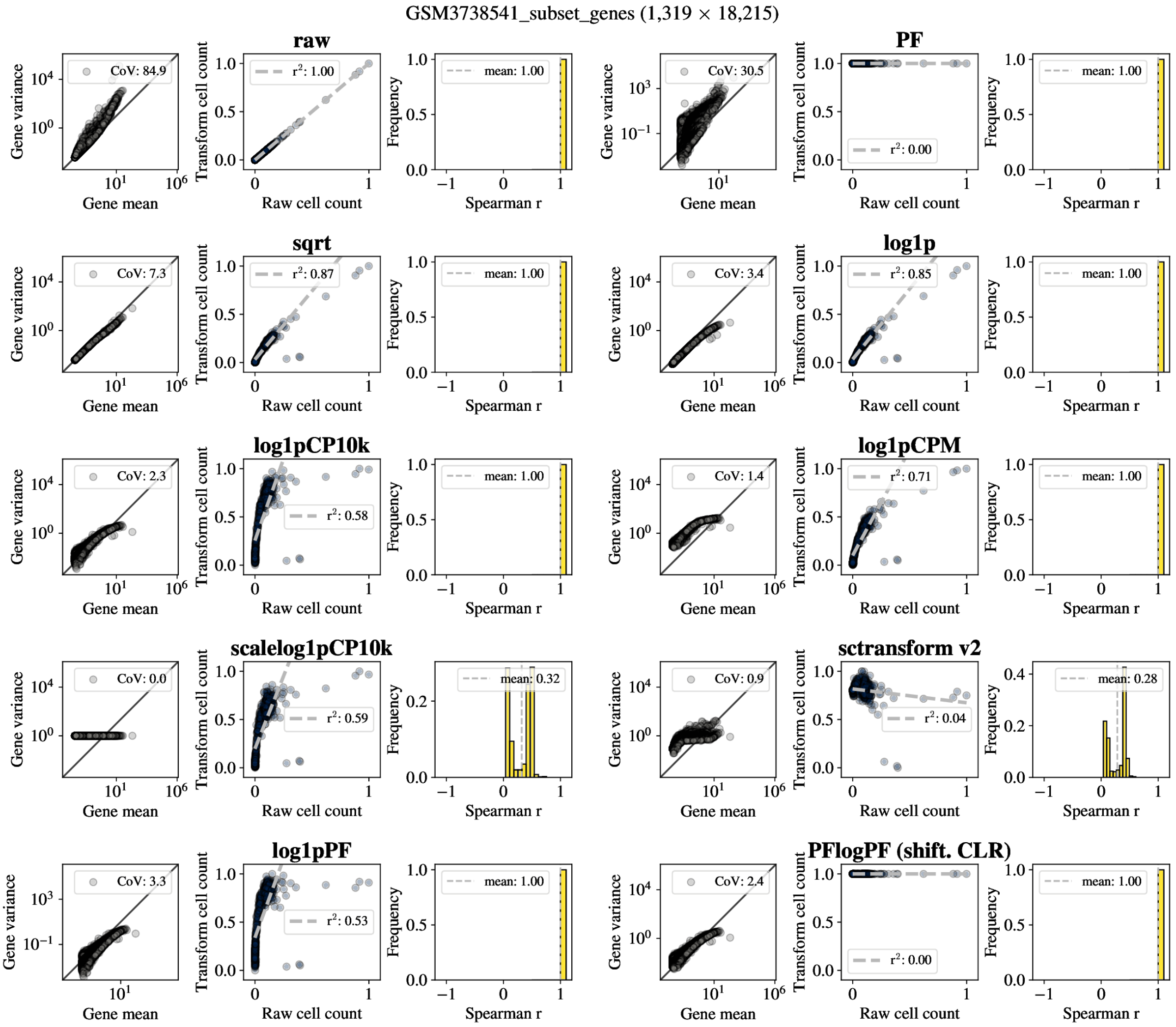


Supplementary Figure 1.402. **GSM3738541**. Per-method comparison of variance stabilization (left), depth normalization (middle), and monotonicity (right) for each normalization on dataset GSM3738541.

| GSM3738542 | |
|---|---|
| cells | 1,988 |
| genes | 60,623 |
| nonzero entries | 3,120,626 |
| density | 0.0259 |
| total counts | 7,742,927 |
| mean counts / cell | 3,895 |
| mean counts / gene | 128 |
| min counts / cell | 50 |
| max counts / cell | 15,263 |
| overdispersion | 0.302 |

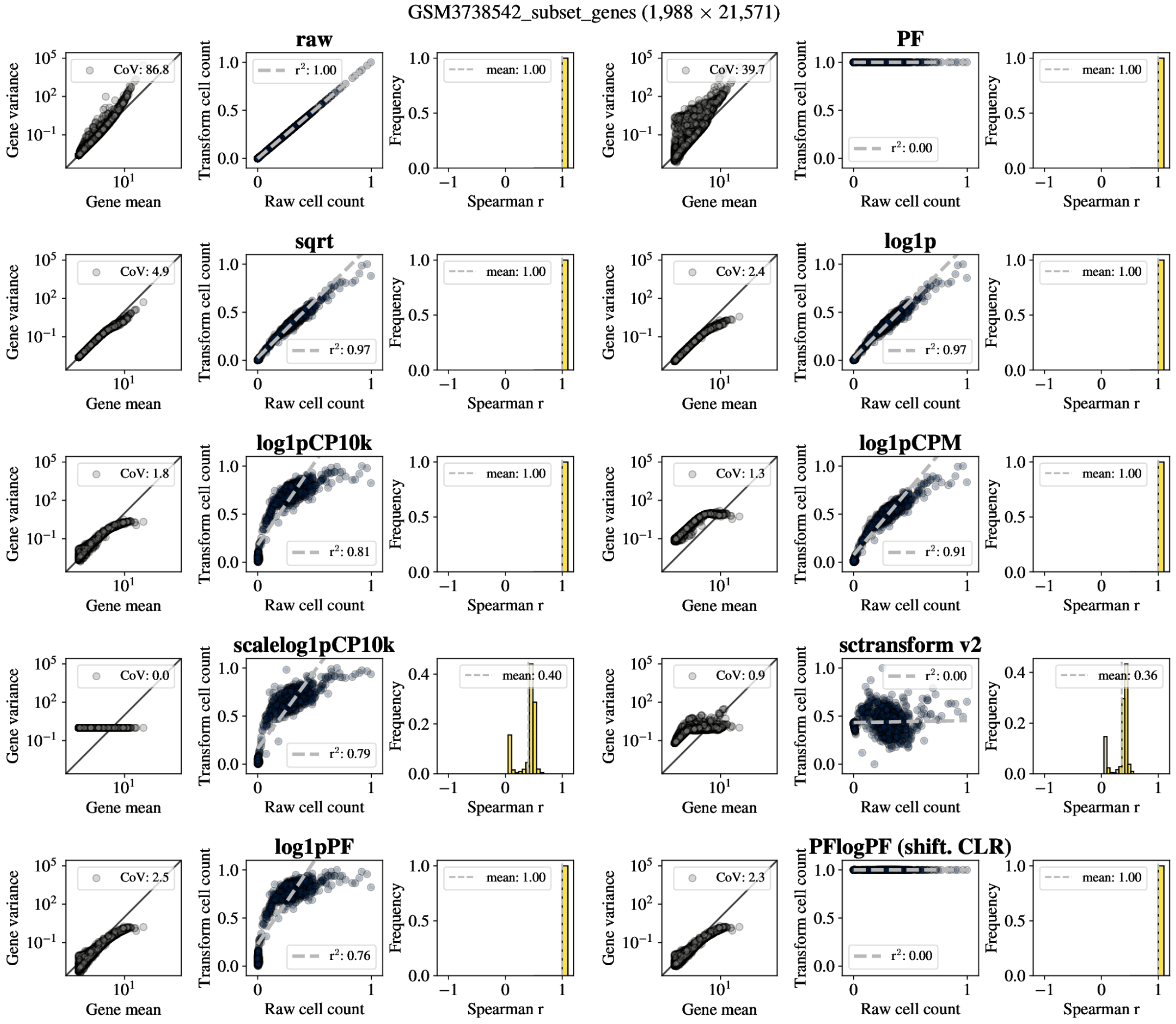


Supplementary Figure 1.403. **GSM3738542**. Per-method comparison of variance stabilization (left), depth normalization (middle), and monotonicity (right) for each normalization on dataset GSM3738542.

| **GSM3738543** | |
|---|---|
| cells | 3,719 |
| genes | 60,623 |
| nonzero entries | 2,826,297 |
| density | 0.0125 |
| total counts | 7,073,643 |
| mean counts / cell | 1,902 |
| mean counts / gene | 117 |
| min counts / cell | 29 |
| max counts / cell | 11,089 |
| overdispersion | 1.55 |

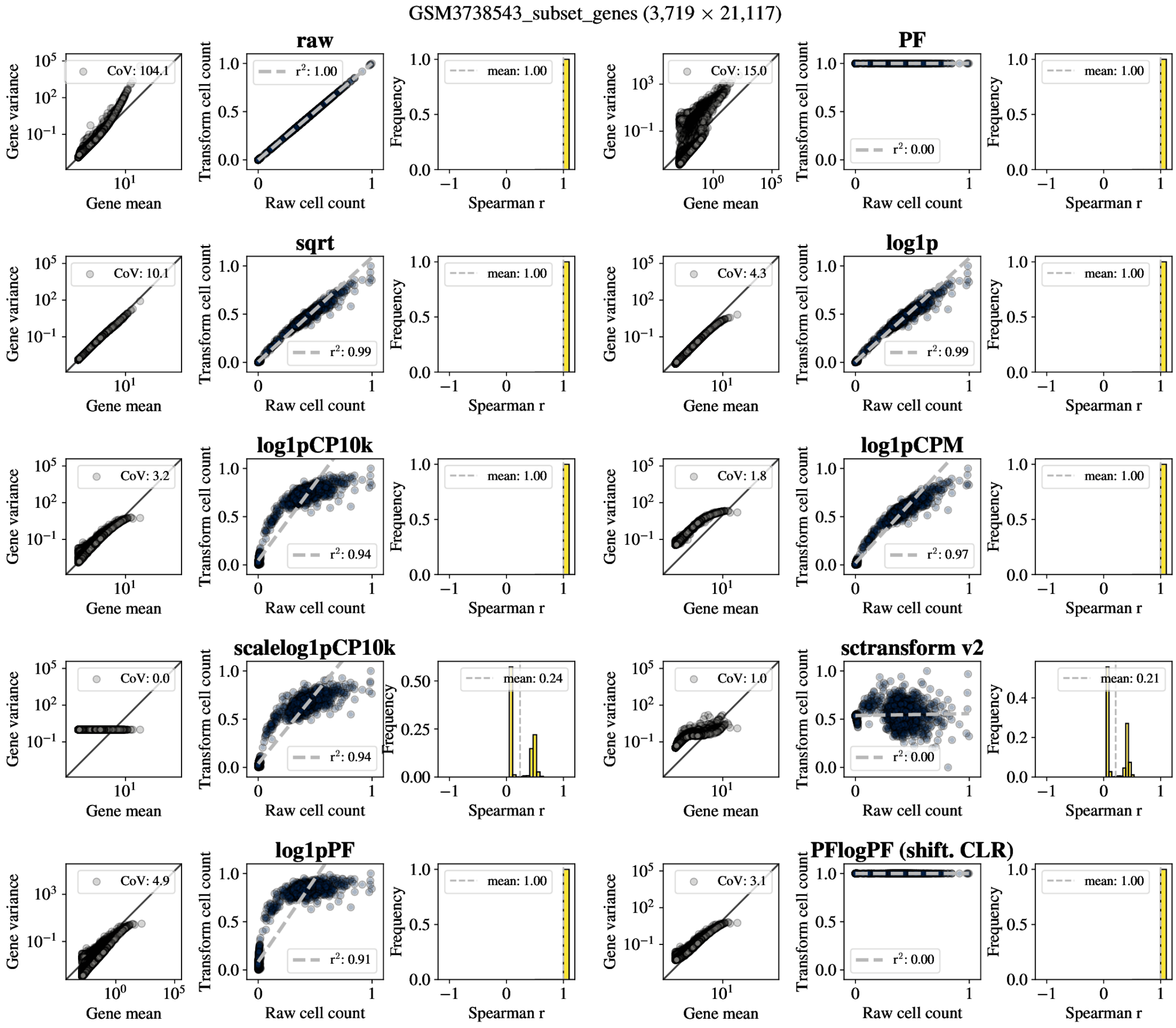


Supplementary Figure 1.404. **GSM3738543**. Per-method comparison of variance stabilization (left), depth normalization (middle), and monotonicity (right) for each normalization on dataset GSM3738543.

| GSM3739217 | |
|---|---|
| cells | 7,302 |
| genes | 60,623 |
| nonzero entries | 15,667,993 |
| density | 0.0354 |
| total counts | 51,101,771 |
| mean counts / cell | 6,998 |
| mean counts / gene | 843 |
| min counts / cell | 208 |
| max counts / cell | 39,833 |
| overdispersion | 0.648 |

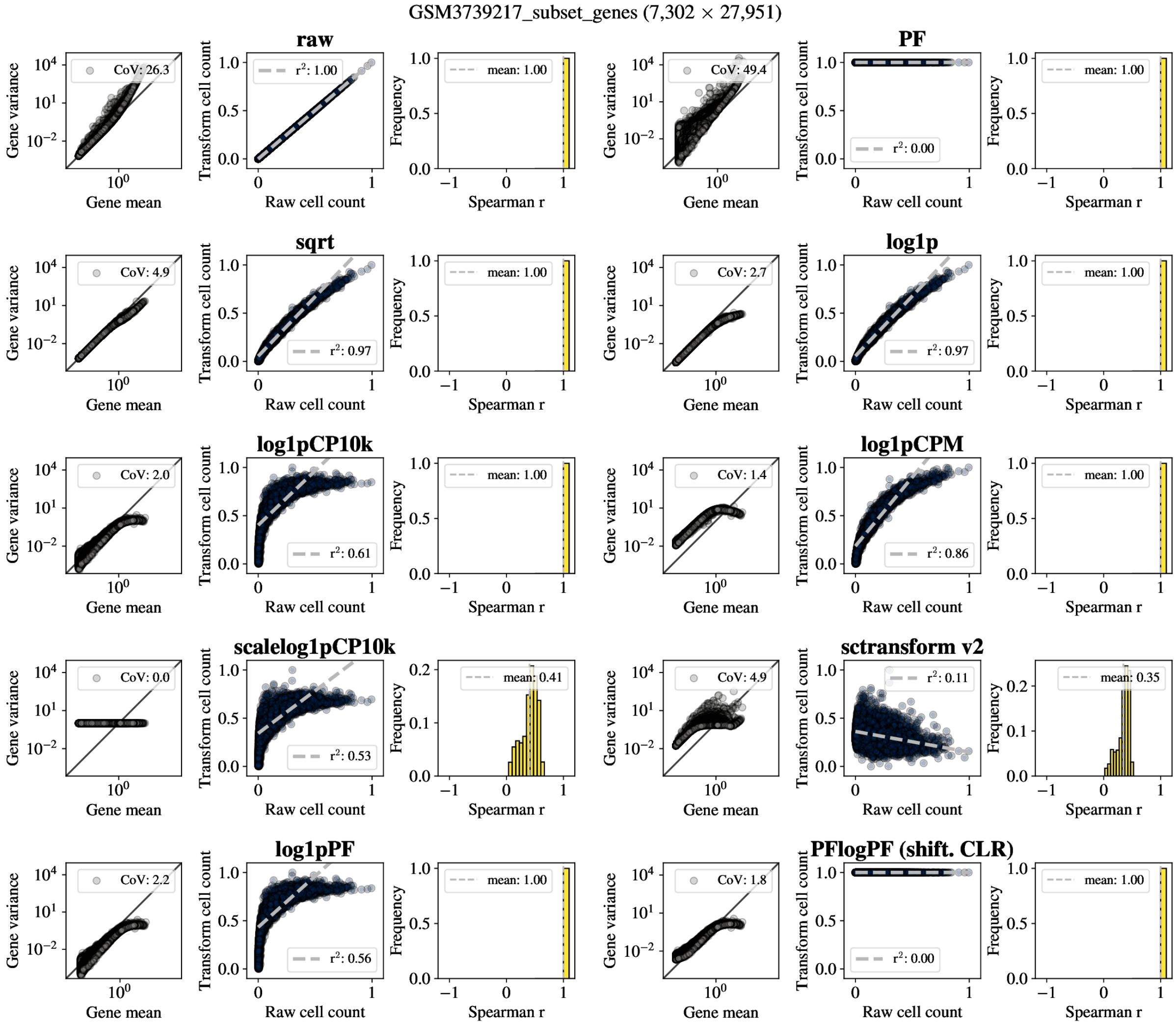


Supplementary Figure 1.405. **GSM3739217**. Per-method comparison of variance stabilization (left), depth normalization (middle), and monotonicity (right) for each normalization on dataset GSM3739217.

| GSM3739218 | |
|---|---|
| cells | 81 |
| genes | 60,623 |
| nonzero entries | 177,419 |
| density | 0.0361 |
| total counts | 581,038 |
| mean counts / cell | 7,173 |
| mean counts / gene | 10 |
| min counts / cell | 512 |
| max counts / cell | 32,125 |
| overdispersion | 2.71 |

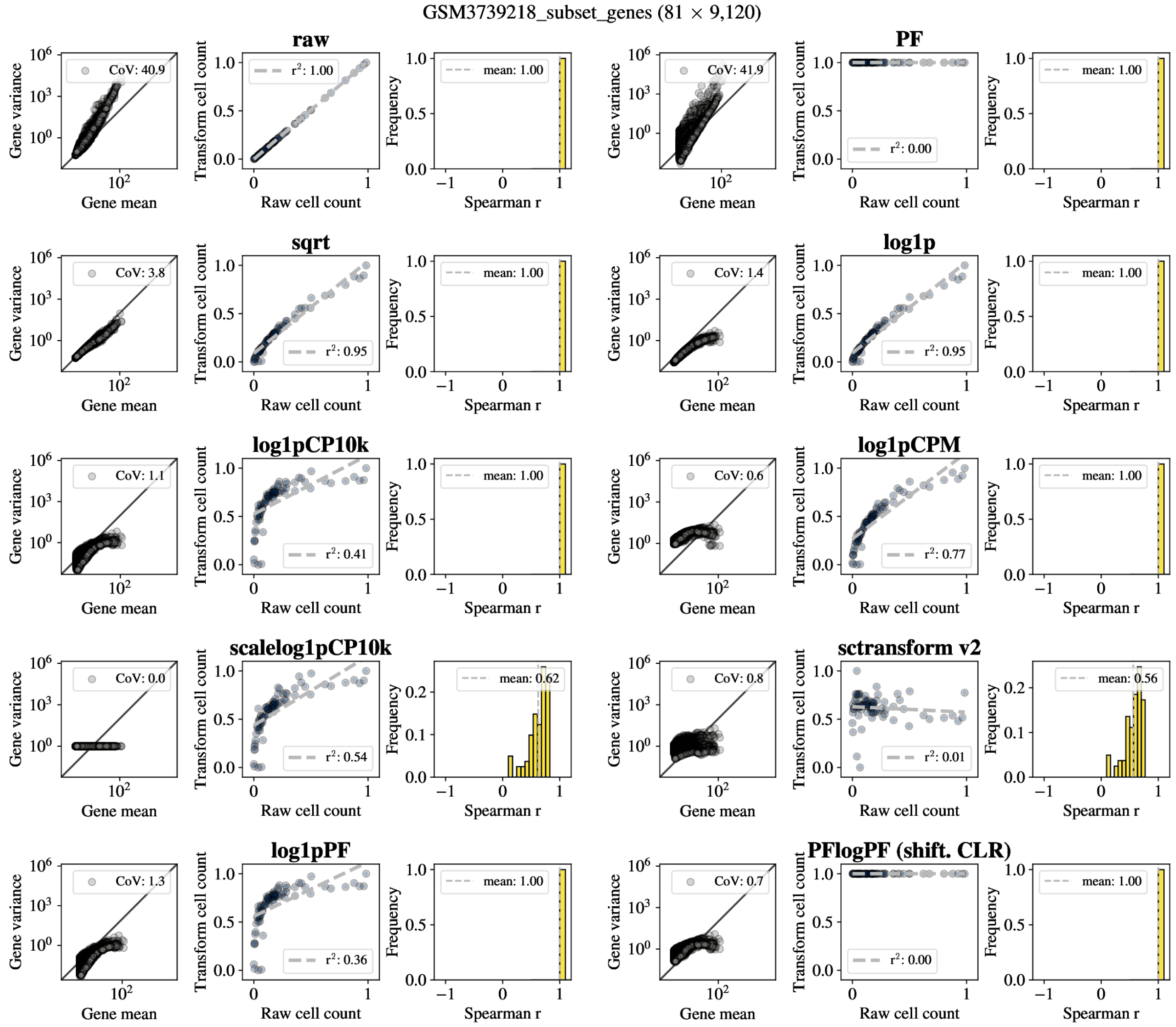


Supplementary Figure 1.406. **GSM3739218**. Per-method comparison of variance stabilization (left), depth normalization (middle), and monotonicity (right) for each normalization on dataset GSM3739218.

| GSM3739219 | |
|---|---|
| cells | 62 |
| genes | 60,623 |
| nonzero entries | 166,052 |
| density | 0.0442 |
| total counts | 513,769 |
| mean counts / cell | 8,287 |
| mean counts / gene | 8 |
| min counts / cell | 958 |
| max counts / cell | 23,340 |
| overdispersion | 0.443 |

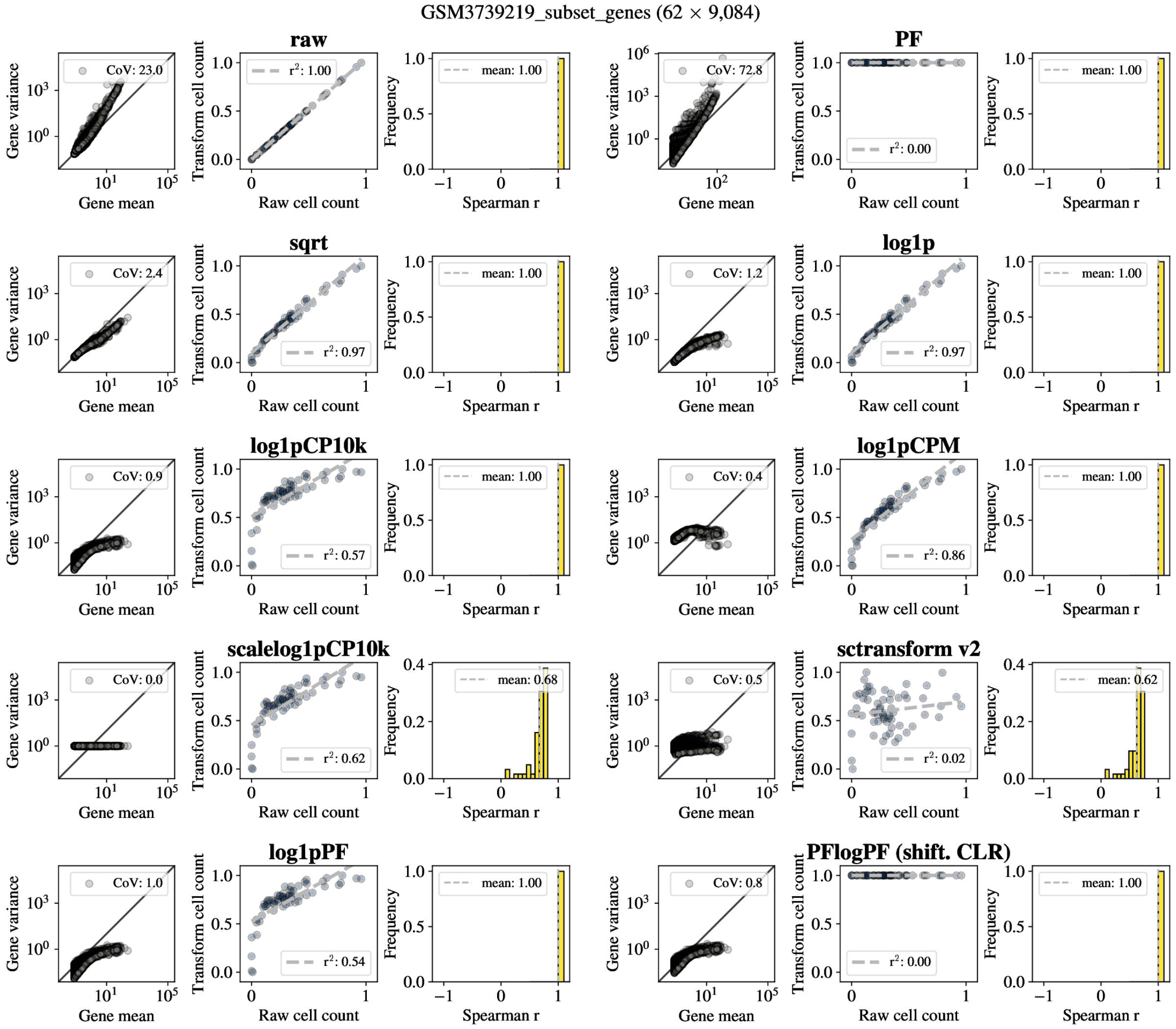


Supplementary Figure 1.407. **GSM3739219**. Per-method comparison of variance stabilization (left), depth normalization (middle), and monotonicity (right) for each normalization on dataset GSM3739219.

| GSM3739220 | |
|---|---|
| cells | 198 |
| genes | 60,623 |
| nonzero entries | 193,249 |
| density | 0.0161 |
| total counts | 581,112 |
| mean counts / cell | 2,935 |
| mean counts / gene | 10 |
| min counts / cell | 125 |
| max counts / cell | 20,472 |
| overdispersion | 2.29 |

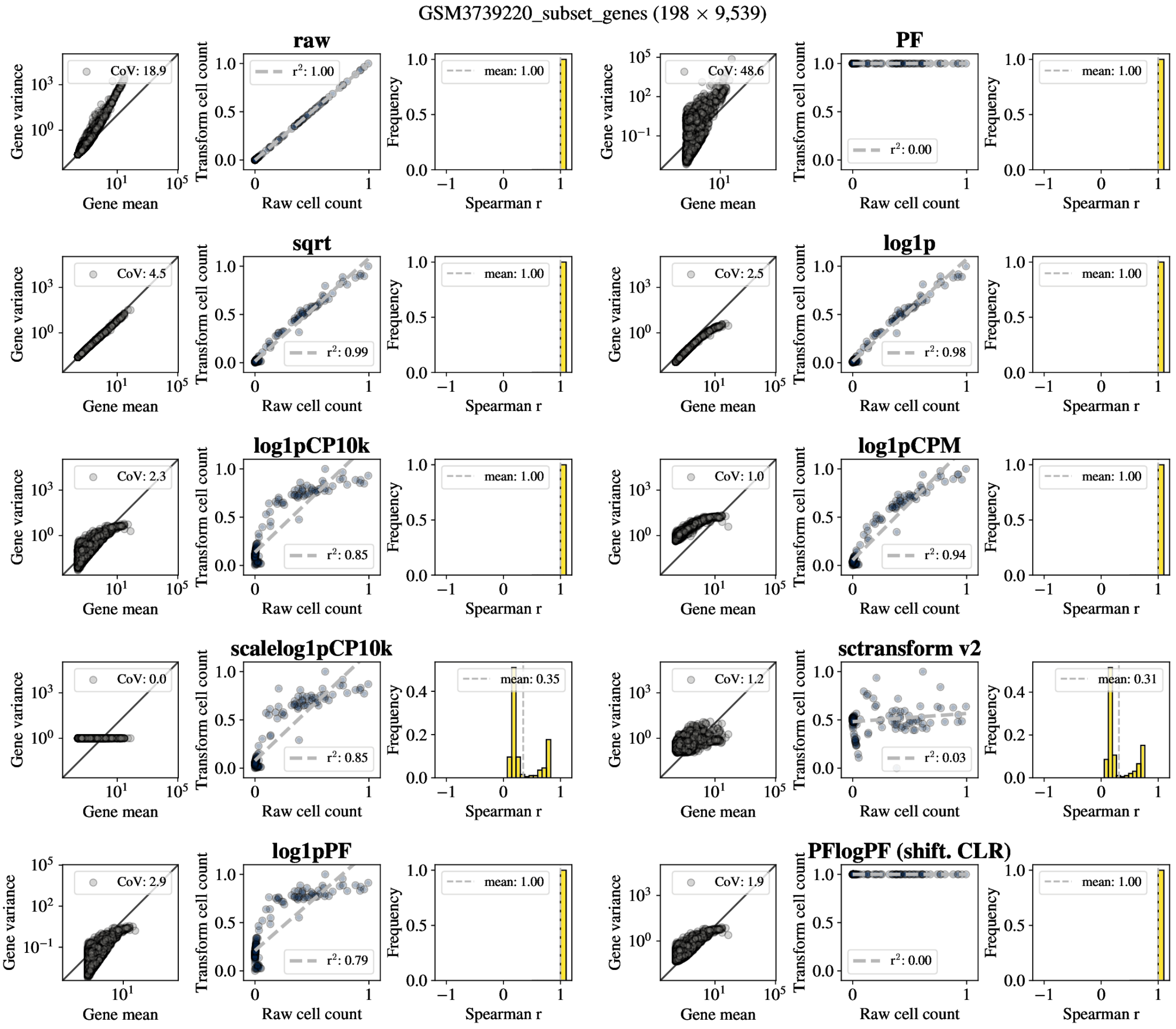


Supplementary Figure 1.408. **GSM3739220**. Per-method comparison of variance stabilization (left), depth normalization (middle), and monotonicity (right) for each normalization on dataset GSM3739220.

| GSM3745992 | |
|---|---|
| cells | 69 |
| genes | 60,623 |
| nonzero entries | 107,496 |
| density | 0.0257 |
| total counts | 690,850 |
| mean counts / cell | 10,012 |
| mean counts / gene | 11 |
| min counts / cell | 848 |
| max counts / cell | 67,217 |
| overdispersion | 1.08 |

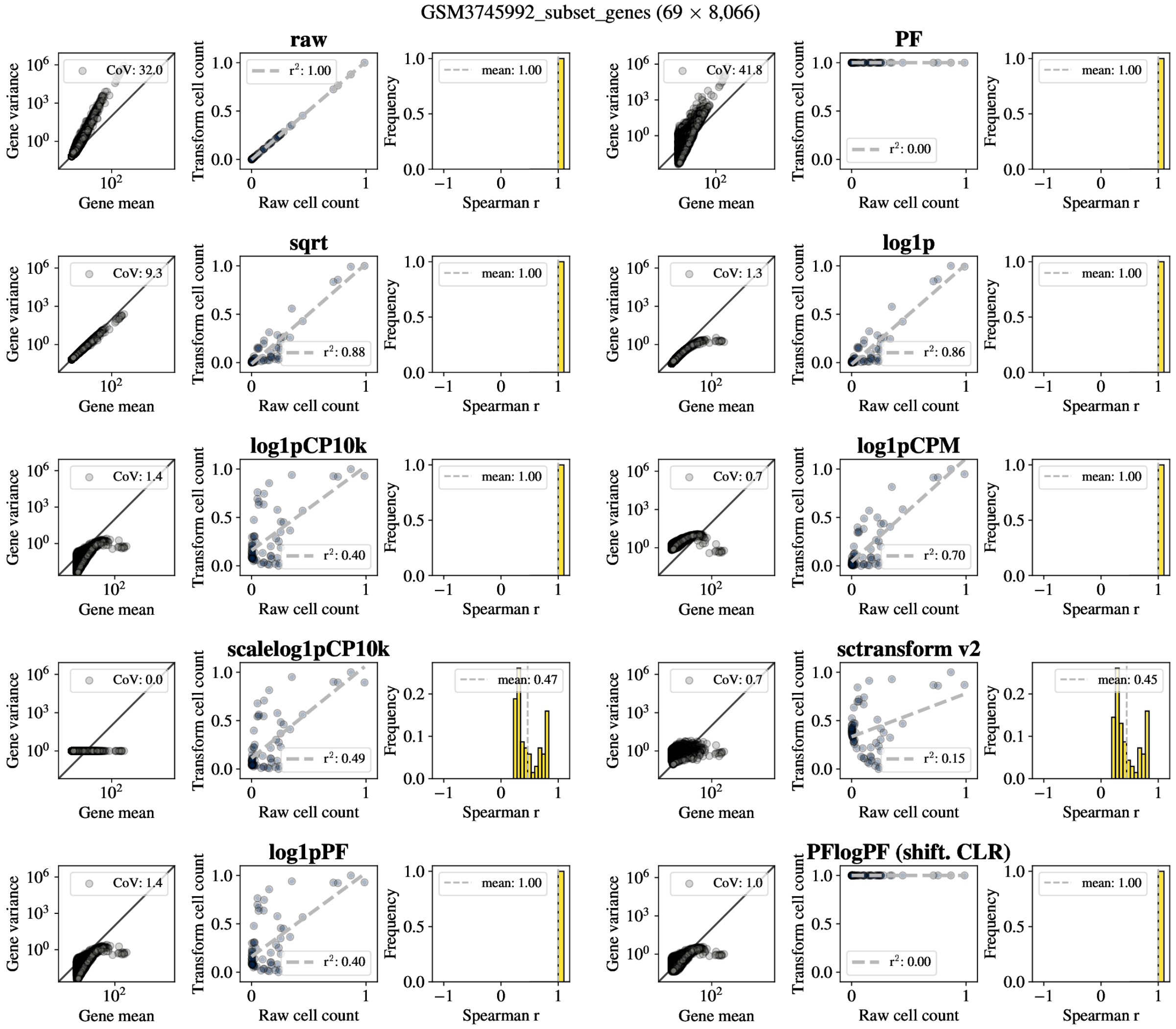


Supplementary Figure 1.409. **GSM3745992**. Per-method comparison of variance stabilization (left), depth normalization (middle), and monotonicity (right) for each normalization on dataset GSM3745992.

| GSM3745993 | |
|---|---|
| cells | 563 |
| genes | 60,623 |
| nonzero entries | 154,210 |
| density | 0.00452 |
| total counts | 707,731 |
| mean counts / cell | 1,257 |
| mean counts / gene | 12 |
| min counts / cell | 457 |
| max counts / cell | 87,451 |
| overdispersion | 5.94 |

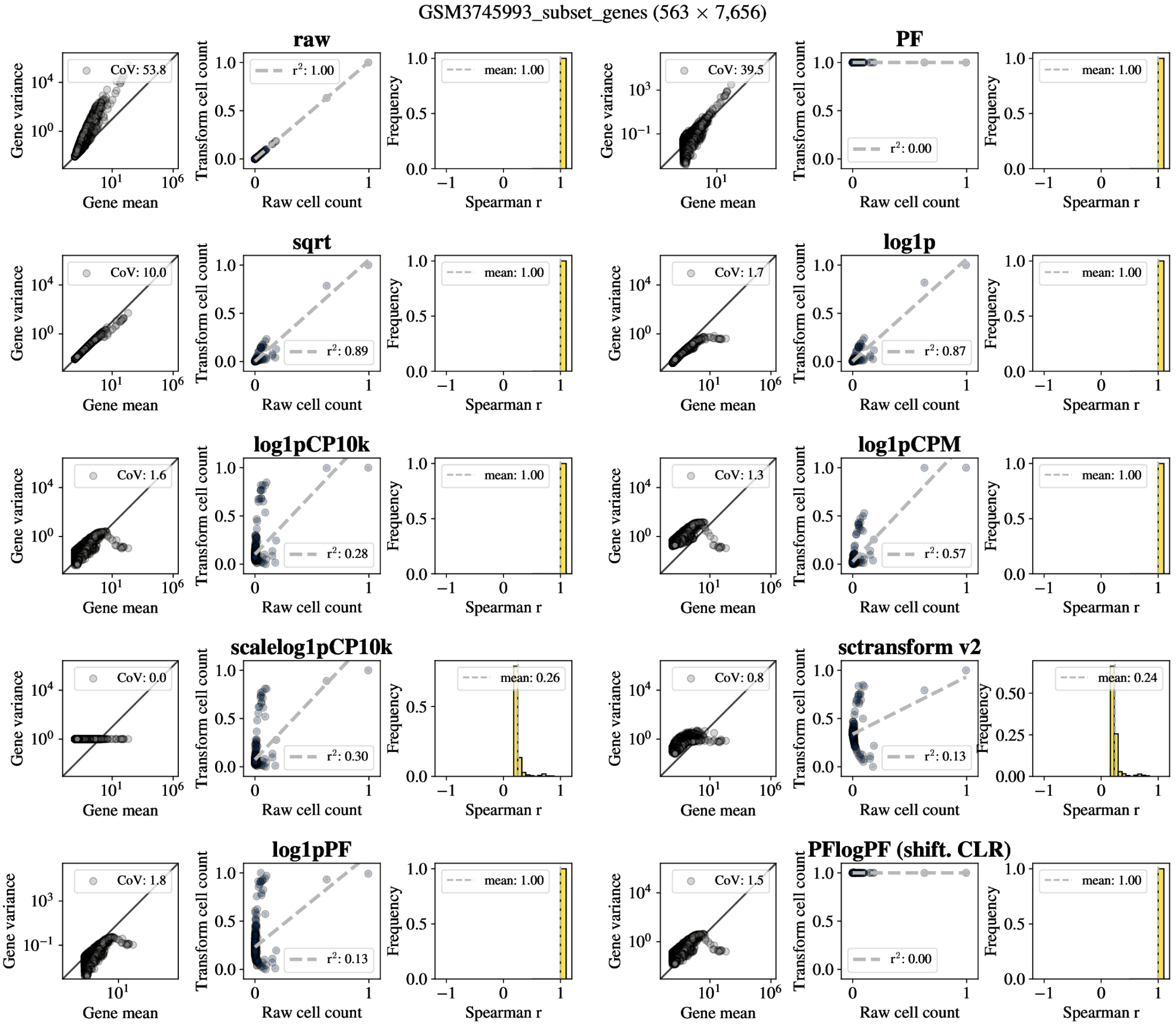


Supplementary Figure 1.410. **GSM3745993**. Per-method comparison of variance stabilization (left), depth normalization (middle), and monotonicity (right) for each normalization on dataset GSM3745993.

| GSM3745994 | |
|---|---|
| cells | 40 |
| genes | 60,623 |
| nonzero entries | 43,198 |
| density | 0.0178 |
| total counts | 538,306 |
| mean counts / cell | 13,458 |
| mean counts / gene | 9 |
| min counts / cell | 913 |
| max counts / cell | 76,211 |
| overdispersion | 1.25 |

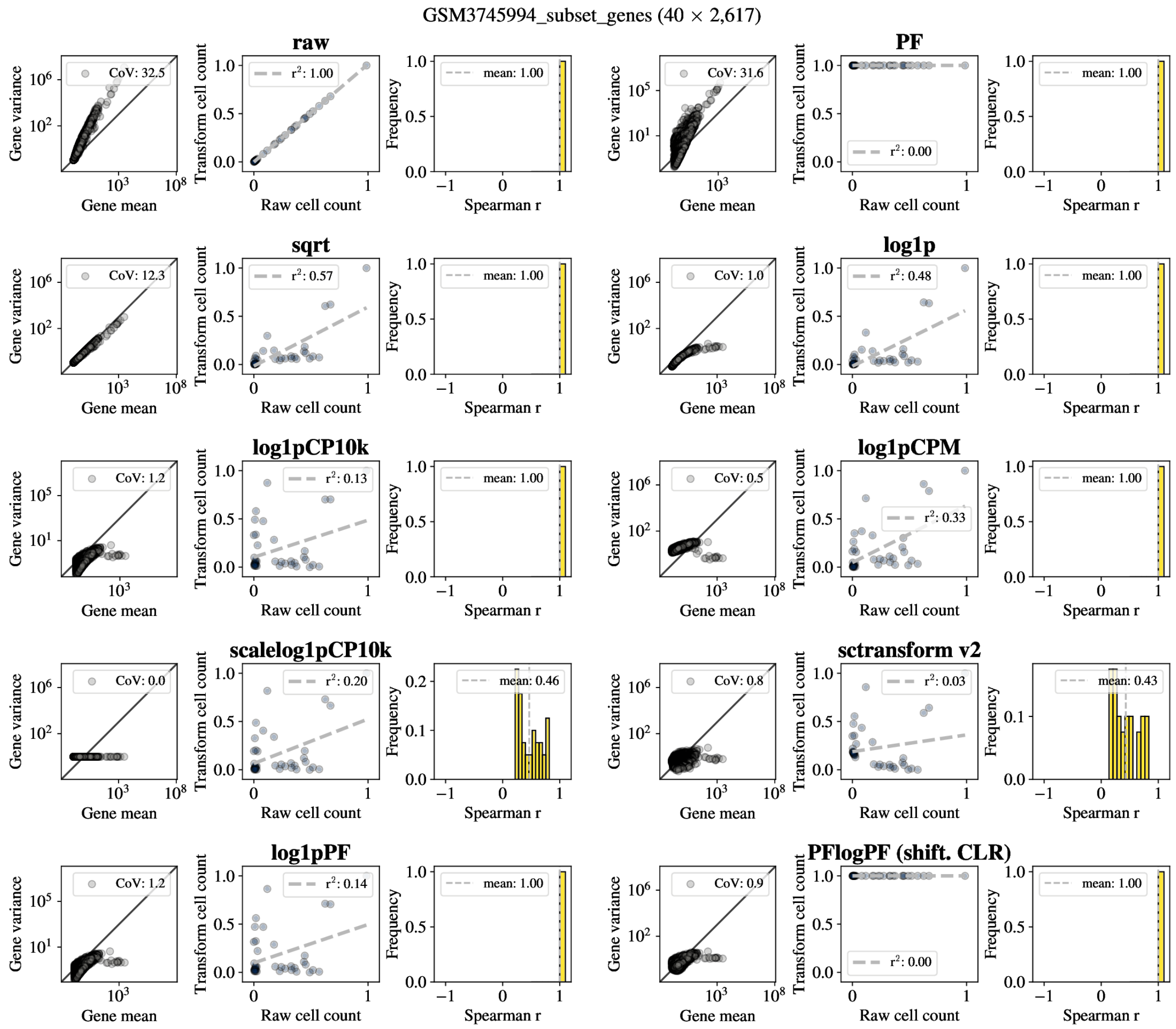


Supplementary Figure 1.411. **GSM3745994**. Per-method comparison of variance stabilization (left), depth normalization (middle), and monotonicity (right) for each normalization on dataset GSM3745994.

| GSM3745995 | |
|---|---|
| cells | 66 |
| genes | 60,623 |
| nonzero entries | 103,582 |
| density | 0.0259 |
| total counts | 321,591 |
| mean counts / cell | 4,873 |
| mean counts / gene | 5 |
| min counts / cell | 764 |
| max counts / cell | 22,024 |
| overdispersion | 1.69 |

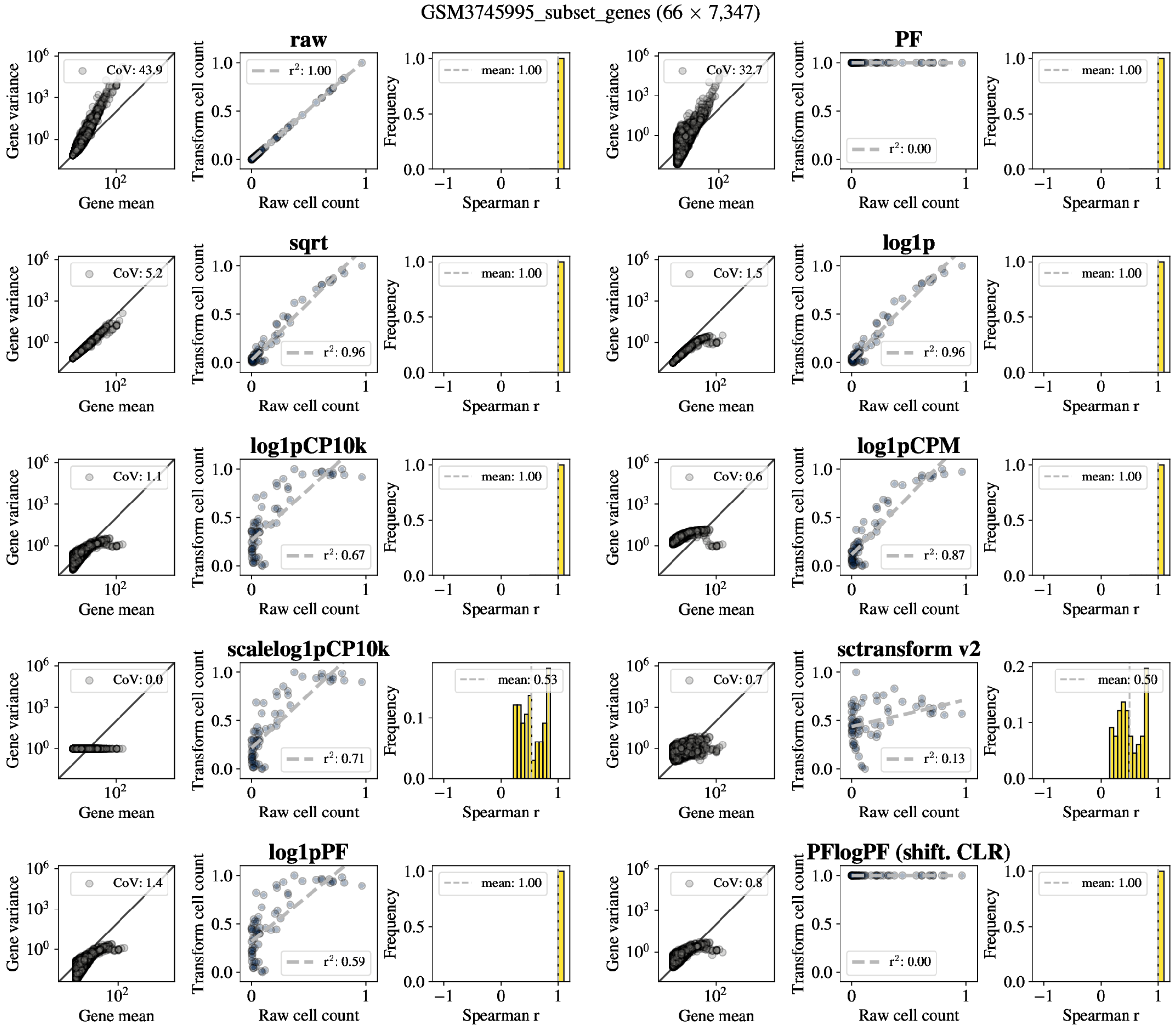


Supplementary Figure 1.412. **GSM3745995**. Per-method comparison of variance stabilization (left), depth normalization (middle), and monotonicity (right) for each normalization on dataset GSM3745995.

| GSM3745996 | |
|---|---|
| cells | 61 |
| genes | 60,623 |
| nonzero entries | 95,293 |
| density | 0.0258 |
| total counts | 336,318 |
| mean counts / cell | 5,513 |
| mean counts / gene | 6 |
| min counts / cell | 778 |
| max counts / cell | 22,793 |
| overdispersion | 1.22 |

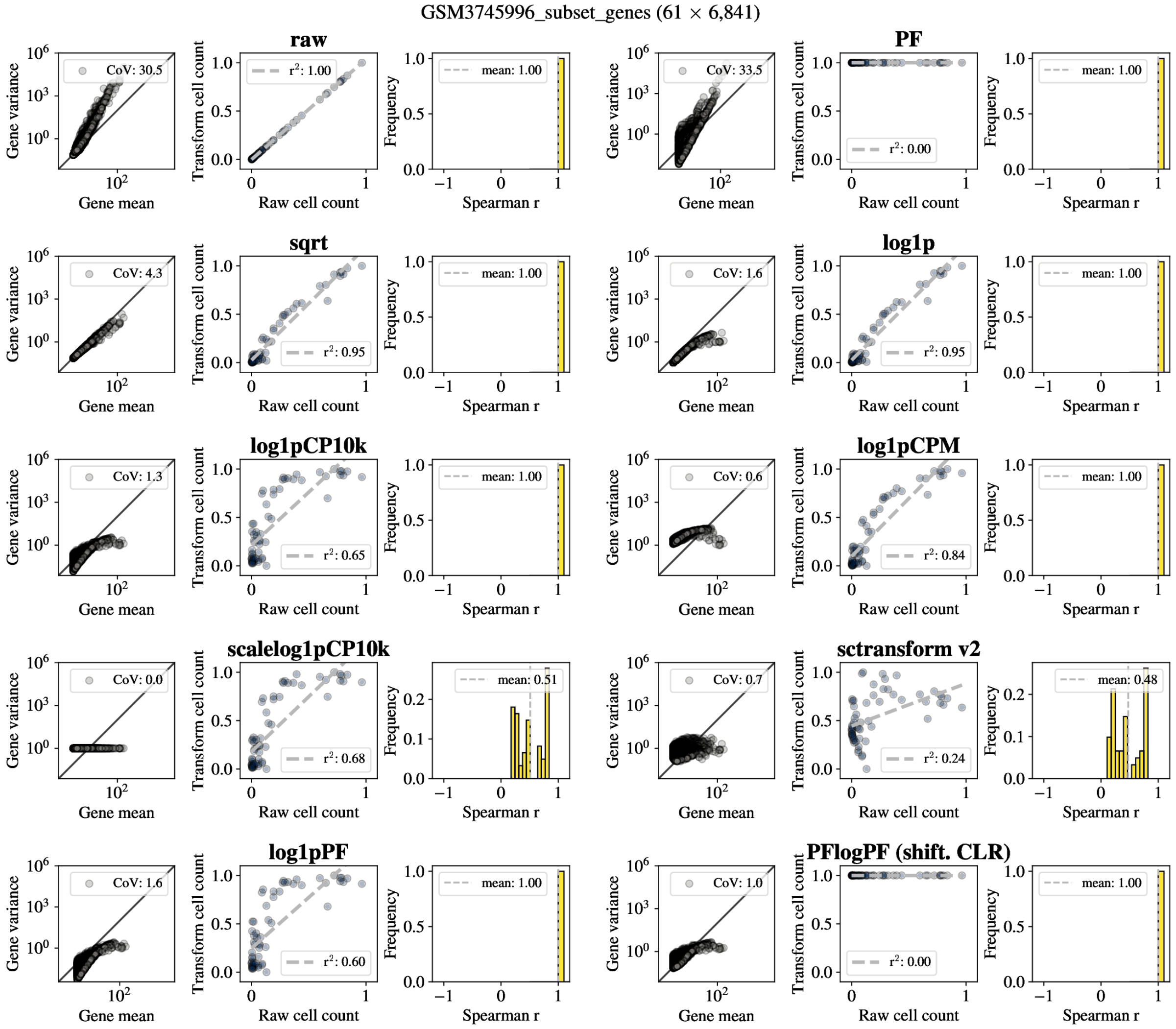


Supplementary Figure 1.413. **GSM3745996**. Per-method comparison of variance stabilization (left), depth normalization (middle), and monotonicity (right) for each normalization on dataset GSM3745996.

| GSM3745997 | |
|---|---|
| cells | 130 |
| genes | 60,623 |
| nonzero entries | 95,081 |
| density | 0.0121 |
| total counts | 292,598 |
| mean counts / cell | 2,251 |
| mean counts / gene | 5 |
| min counts / cell | 360 |
| max counts / cell | 20,870 |
| overdispersion | 1.41 |

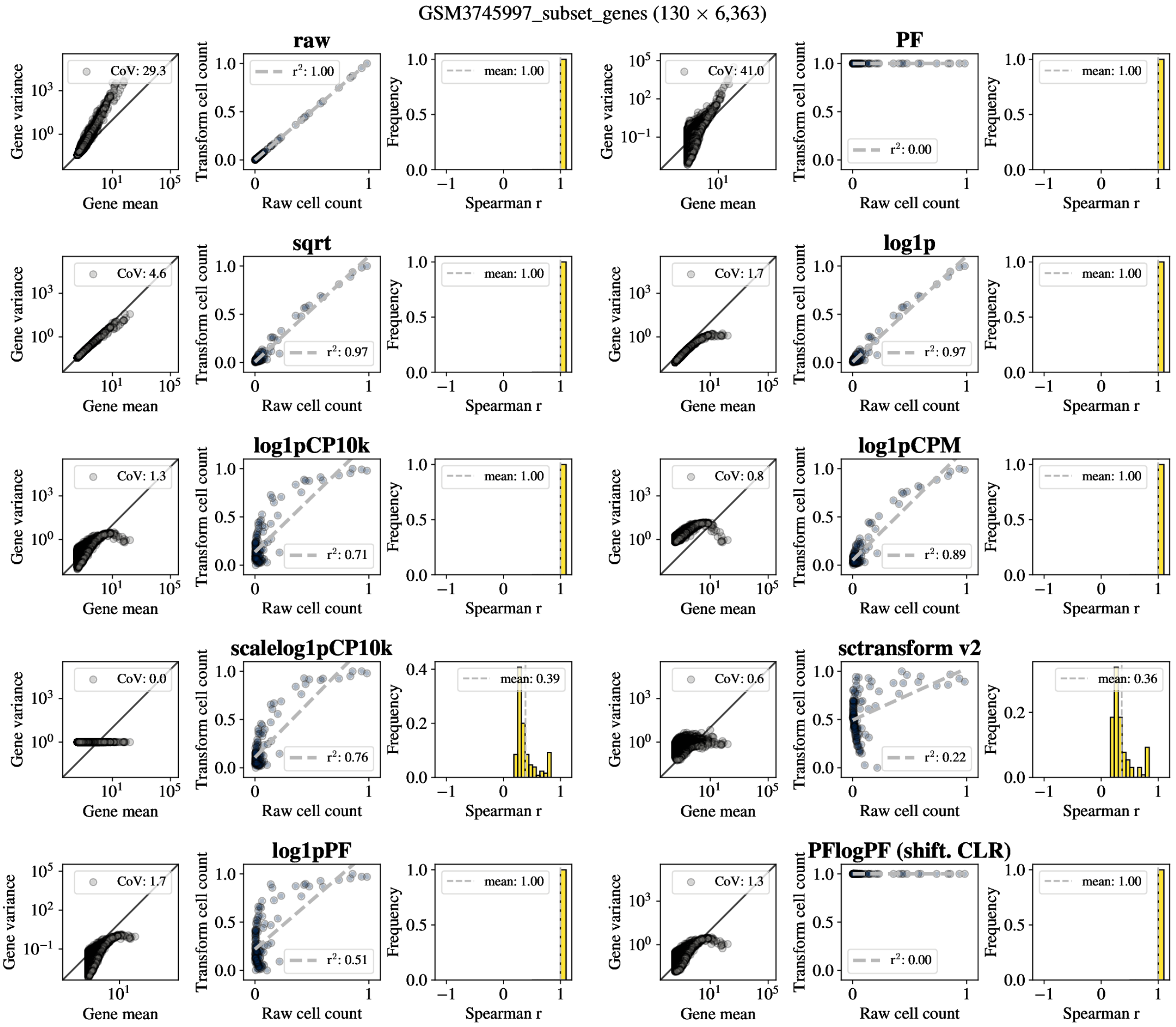


Supplementary Figure 1.414. **GSM3745997**. Per-method comparison of variance stabilization (left), depth normalization (middle), and monotonicity (right) for each normalization on dataset GSM3745997.

| GSM3755687 | |
|---|---|
| cells | 3,171 |
| genes | 60,623 |
| nonzero entries | 5,822,945 |
| density | 0.0303 |
| total counts | 19,207,762 |
| mean counts / cell | 6,057 |
| mean counts / gene | 317 |
| min counts / cell | 3,326 |
| max counts / cell | 94,177 |
| overdispersion | 0.86 |

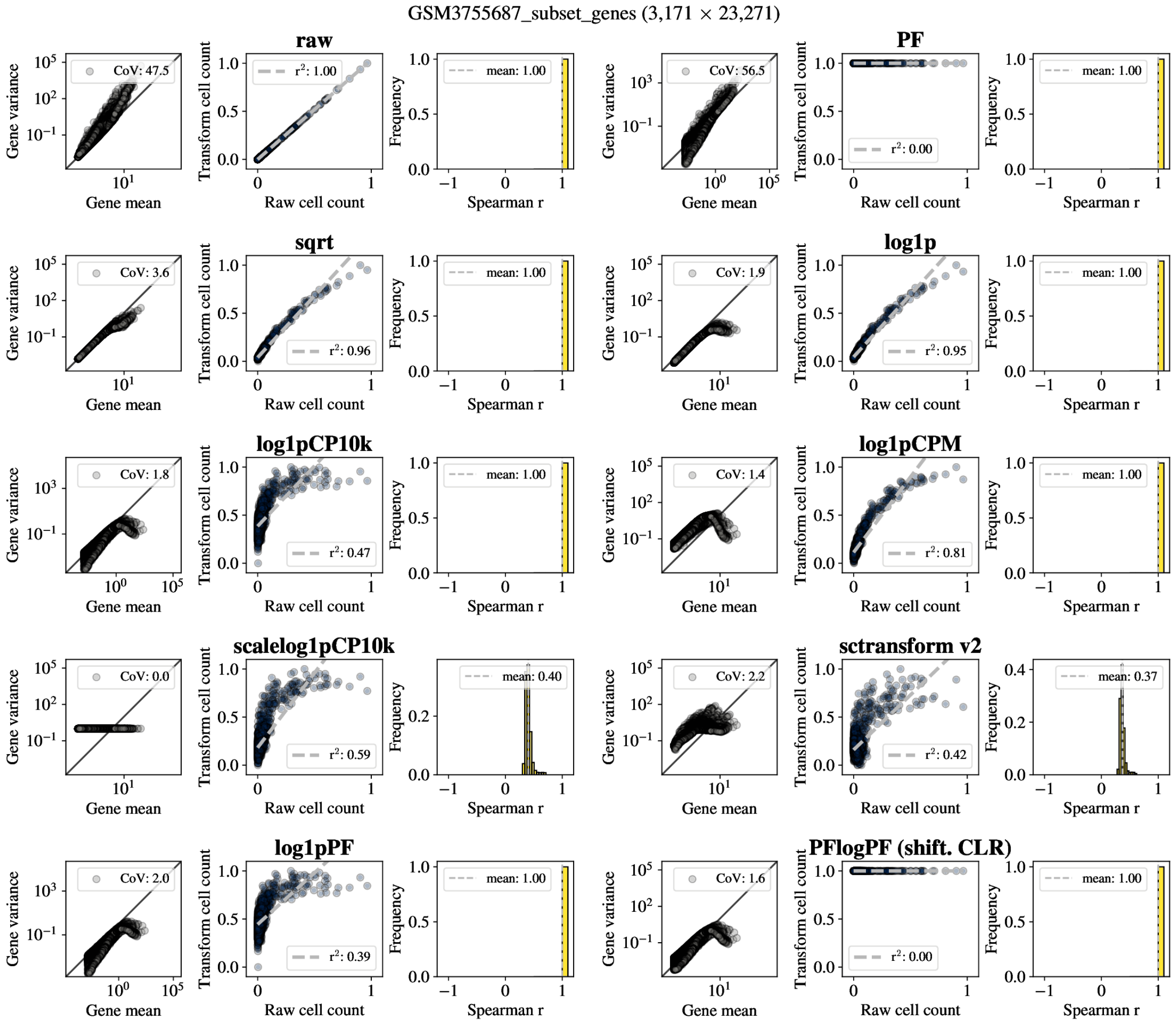


Supplementary Figure 1.415. **GSM3755687**. Per-method comparison of variance stabilization (left), depth normalization (middle), and monotonicity (right) for each normalization on dataset GSM3755687.

| GSM3755690 | |
|---|---|
| cells | 5,673 |
| genes | 60,623 |
| nonzero entries | 8,754,541 |
| density | 0.0255 |
| total counts | 27,911,943 |
| mean counts / cell | 4,920 |
| mean counts / gene | 460 |
| min counts / cell | 1,353 |
| max counts / cell | 63,362 |
| overdispersion | 0.955 |

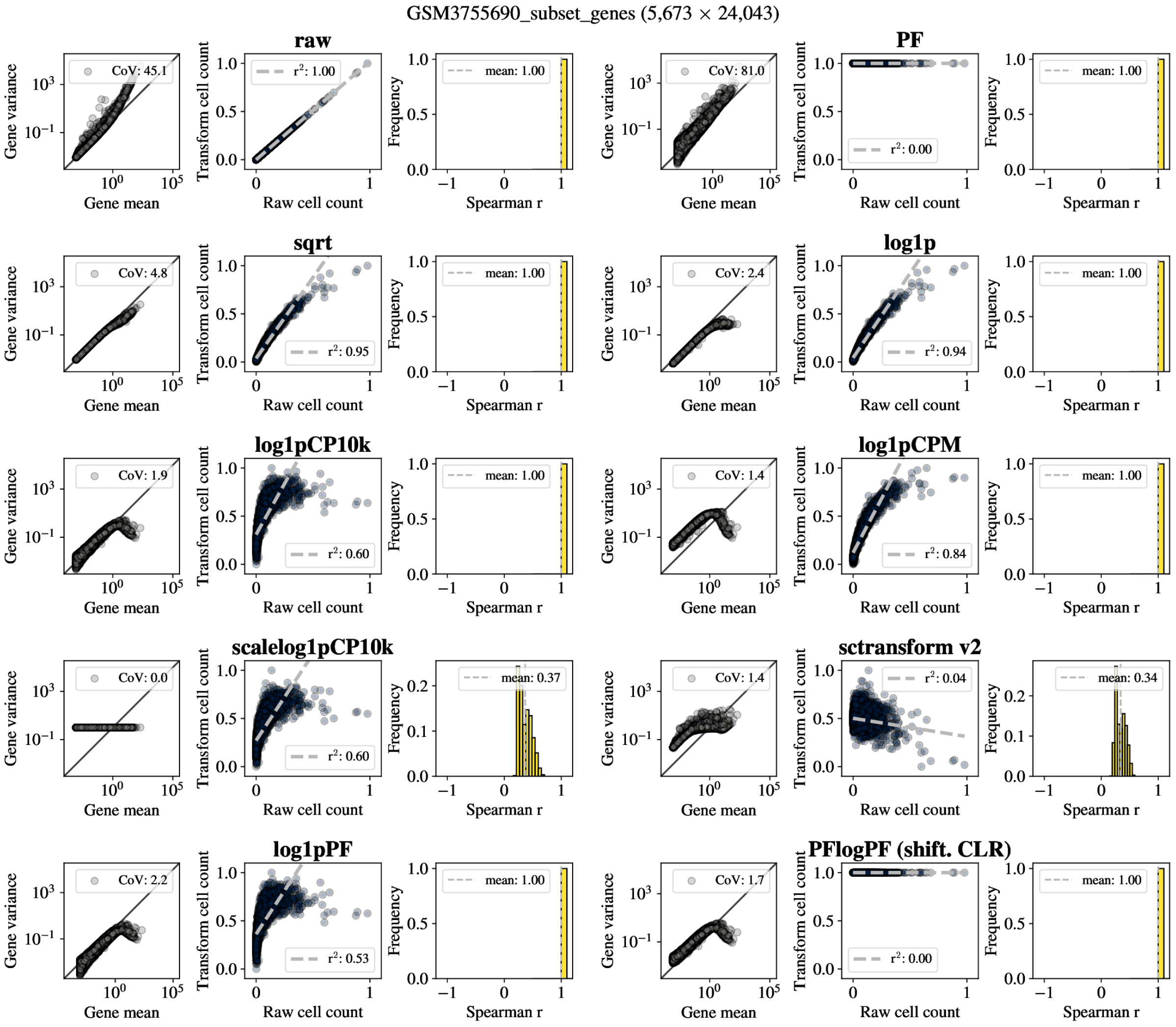


Supplementary Figure 1.416. **GSM3755690**. Per-method comparison of variance stabilization (left), depth normalization (middle), and monotonicity (right) for each normalization on dataset GSM3755690.

| GSM3755691 | |
|---|---|
| cells | 7,998 |
| genes | 60,623 |
| nonzero entries | 14,088,442 |
| density | 0.0291 |
| total counts | 42,396,566 |
| mean counts / cell | 5,301 |
| mean counts / gene | 699 |
| min counts / cell | 2,387 |
| max counts / cell | 56,356 |
| overdispersion | 0.475 |

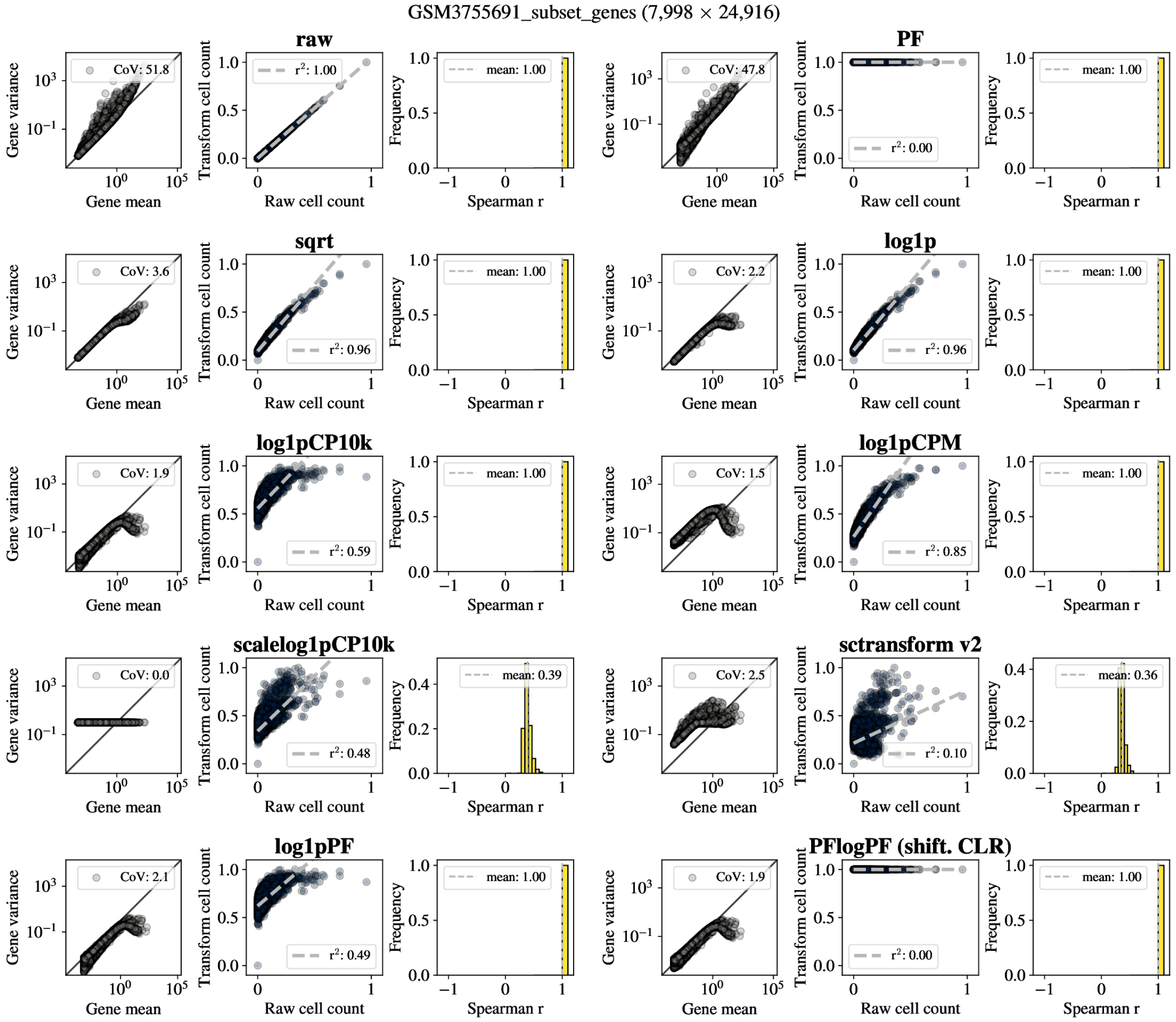


Supplementary Figure 1.417. **GSM3755691**. Per-method comparison of variance stabilization (left), depth normalization (middle), and monotonicity (right) for each normalization on dataset GSM3755691.

| GSM3755692 | |
|---|---|
| cells | 15,775 |
| genes | 60,623 |
| nonzero entries | 18,082,982 |
| density | 0.0189 |
| total counts | 48,082,162 |
| mean counts / cell | 3,048 |
| mean counts / gene | 793 |
| min counts / cell | 397 |
| max counts / cell | 53,134 |
| overdispersion | 0.737 |

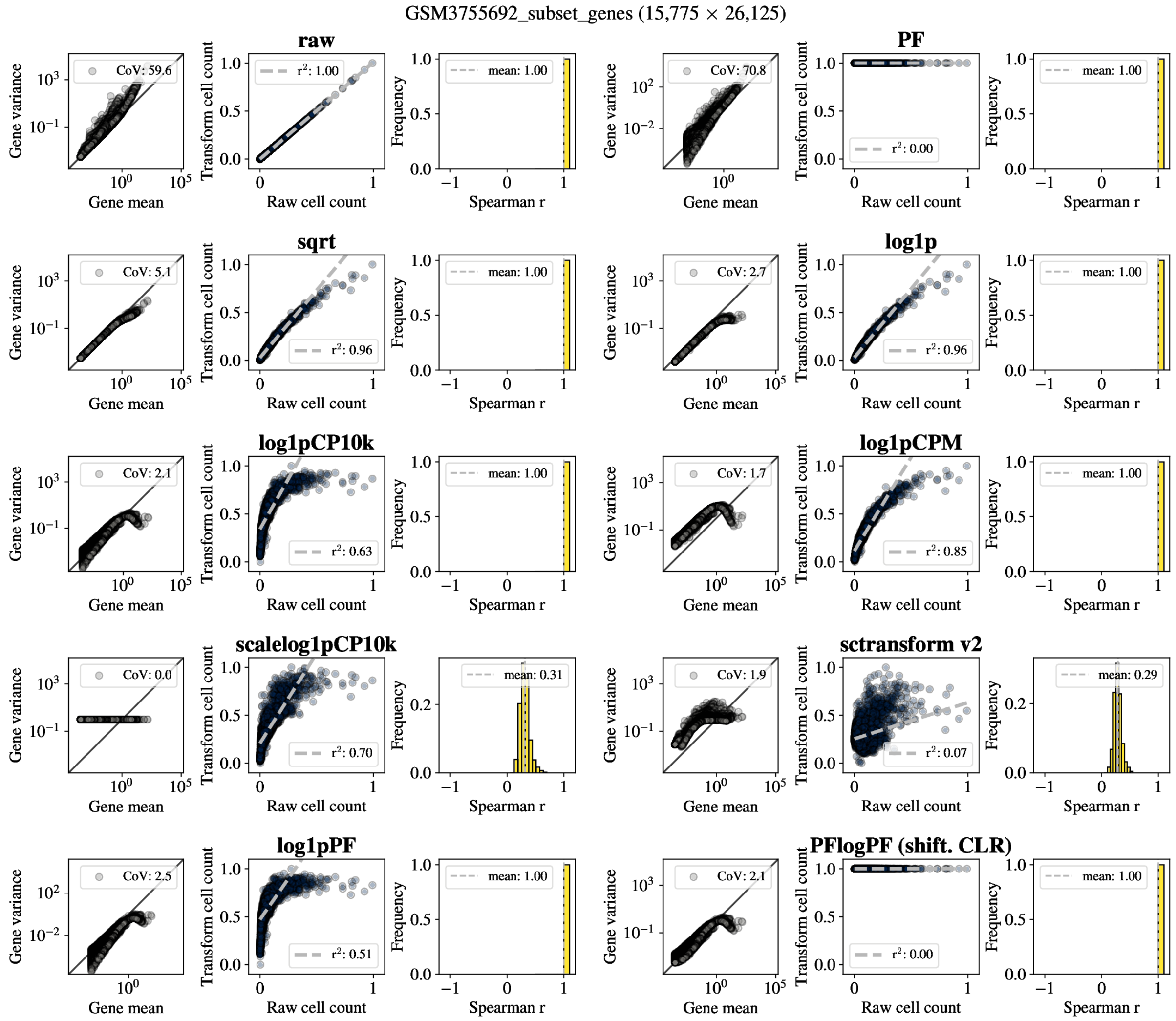


Supplementary Figure 1.418. **GSM3755692**. Per-method comparison of variance stabilization (left), depth normalization (middle), and monotonicity (right) for each normalization on dataset GSM3755692.

| GSM3755693 | |
|---|---|
| cells | 10,423 |
| genes | 60,623 |
| nonzero entries | 18,477,541 |
| density | 0.0292 |
| total counts | 54,368,745 |
| mean counts / cell | 5,216 |
| mean counts / gene | 897 |
| min counts / cell | 244 |
| max counts / cell | 32,789 |
| overdispersion | 0.678 |

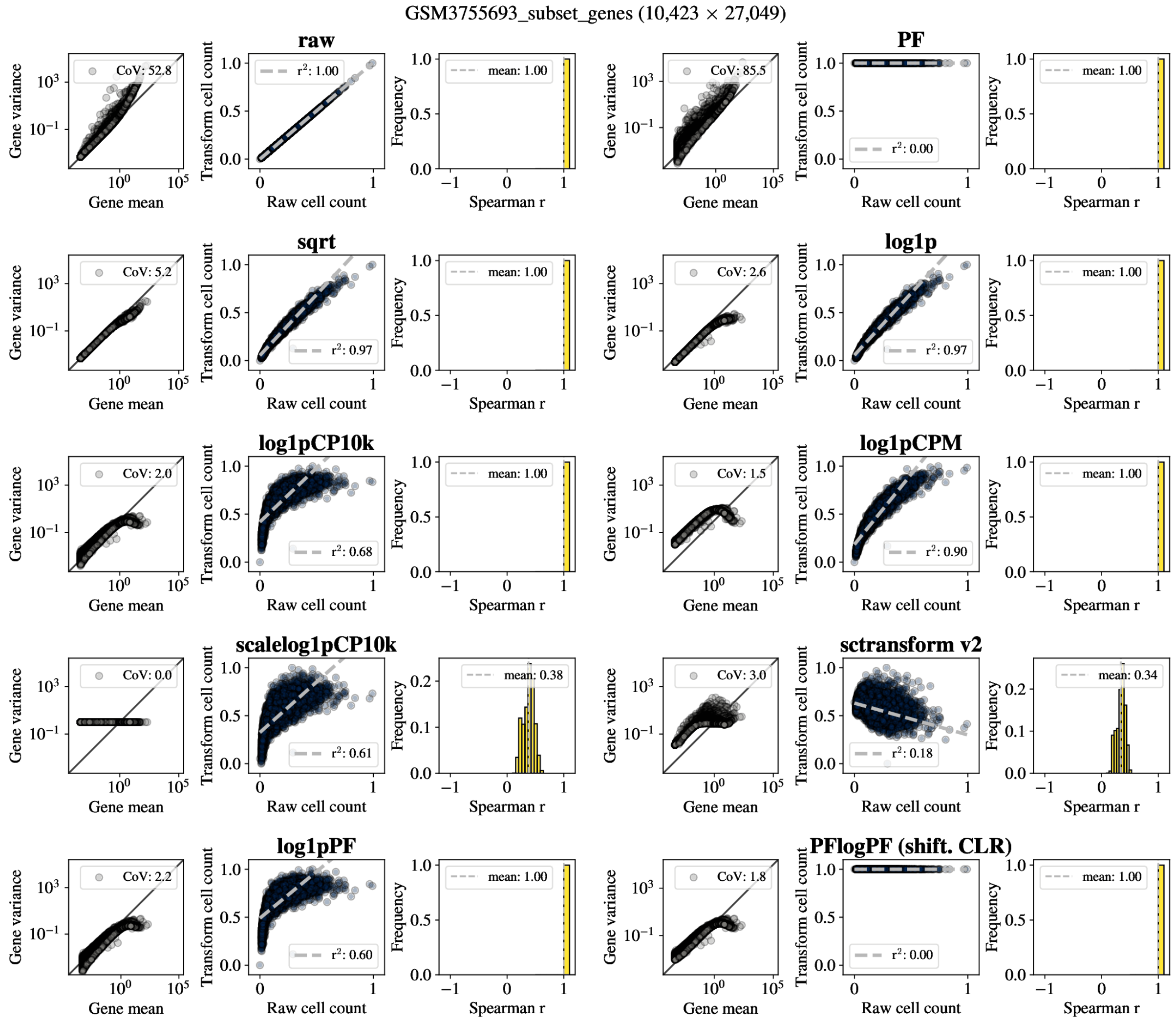


Supplementary Figure 1.419. **GSM3755693**. Per-method comparison of variance stabilization (left), depth normalization (middle), and monotonicity (right) for each normalization on dataset GSM3755693.

| GSM3755696 | |
|---|---|
| cells | 3,000 |
| genes | 60,623 |
| nonzero entries | 4,616,929 |
| density | 0.0254 |
| total counts | 14,970,752 |
| mean counts / cell | 4,990 |
| mean counts / gene | 247 |
| min counts / cell | 645 |
| max counts / cell | 117,660 |
| overdispersion | 2.6 |

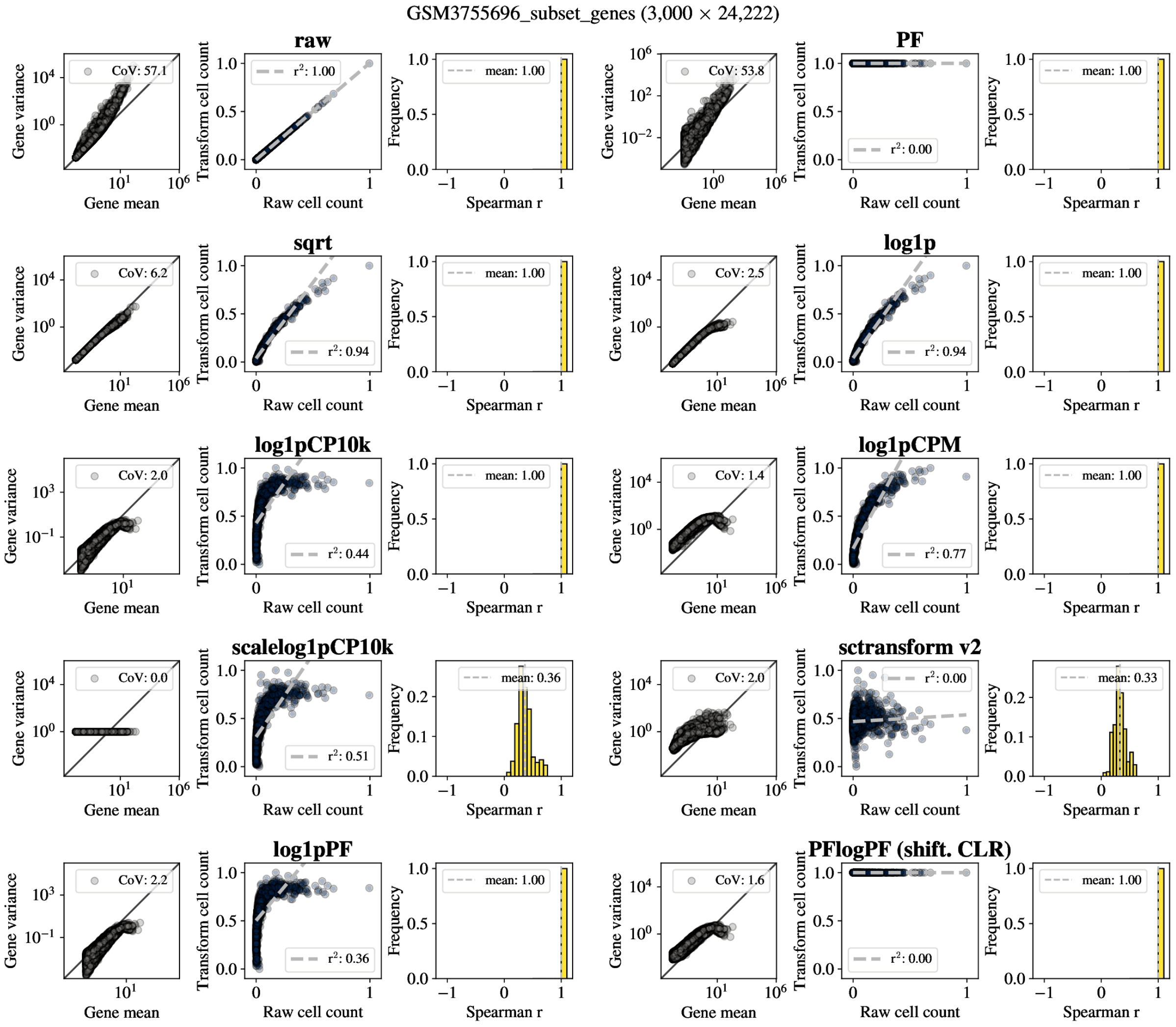


Supplementary Figure 1.420. **GSM3755696**. Per-method comparison of variance stabilization (left), depth normalization (middle), and monotonicity (right) for each normalization on dataset GSM3755696.

| GSM3755697 | |
|---|---|
| cells | 1,622 |
| genes | 60,623 |
| nonzero entries | 3,607,787 |
| density | 0.0367 |
| total counts | 12,768,281 |
| mean counts / cell | 7,872 |
| mean counts / gene | 211 |
| min counts / cell | 4,024 |
| max counts / cell | 93,788 |
| overdispersion | 0.61 |

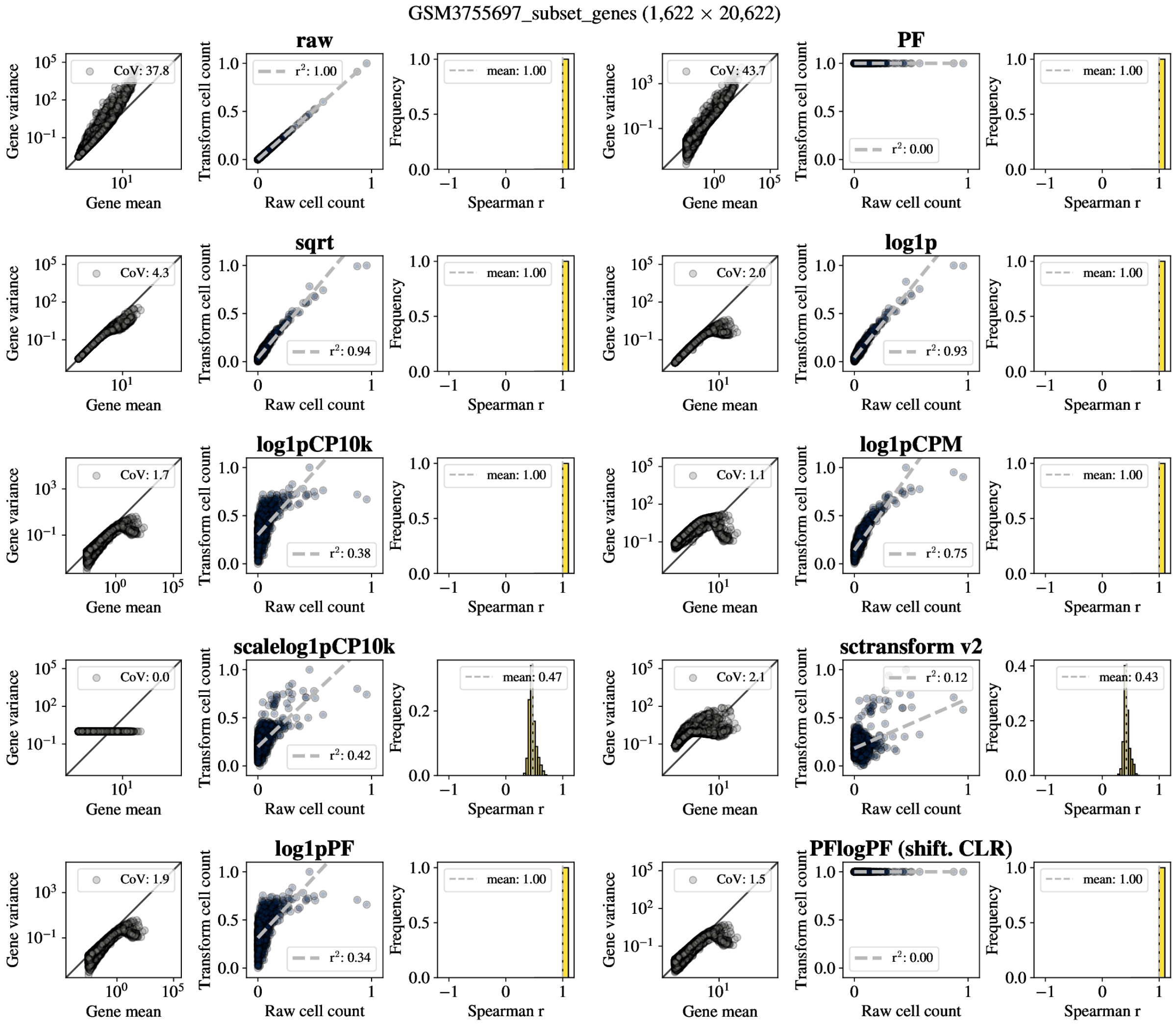


Supplementary Figure 1.421. **GSM3755697**. Per-method comparison of variance stabilization (left), depth normalization (middle), and monotonicity (right) for each normalization on dataset GSM3755697.

| GSM3755698 | |
|---|---|
| cells | 5,802 |
| genes | 60,623 |
| nonzero entries | 12,838,912 |
| density | 0.0365 |
| total counts | 54,612,525 |
| mean counts / cell | 9,413 |
| mean counts / gene | 901 |
| min counts / cell | 1,592 |
| max counts / cell | 109,990 |
| overdispersion | 1.76 |

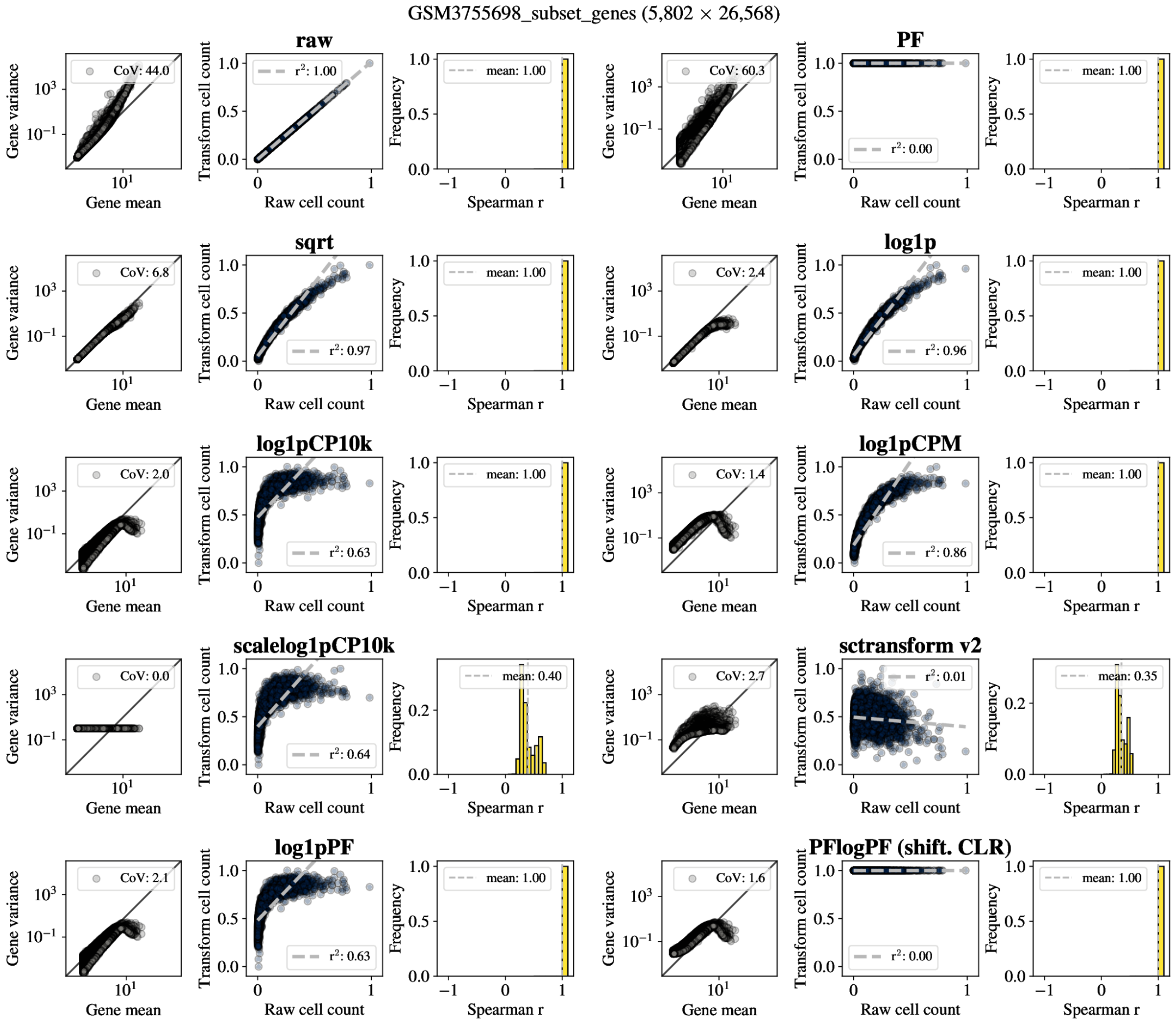


Supplementary Figure 1.422. **GSM3755698**. Per-method comparison of variance stabilization (left), depth normalization (middle), and monotonicity (right) for each normalization on dataset GSM3755698.

| **GSM3755699** | |
|---|---|
| cells | 12,723 |
| genes | 60,623 |
| nonzero entries | 15,944,087 |
| density | 0.0207 |
| total counts | 46,866,650 |
| mean counts / cell | 3,684 |
| mean counts / gene | 773 |
| min counts / cell | 119 |
| max counts / cell | 87,028 |
| overdispersion | 1.05 |

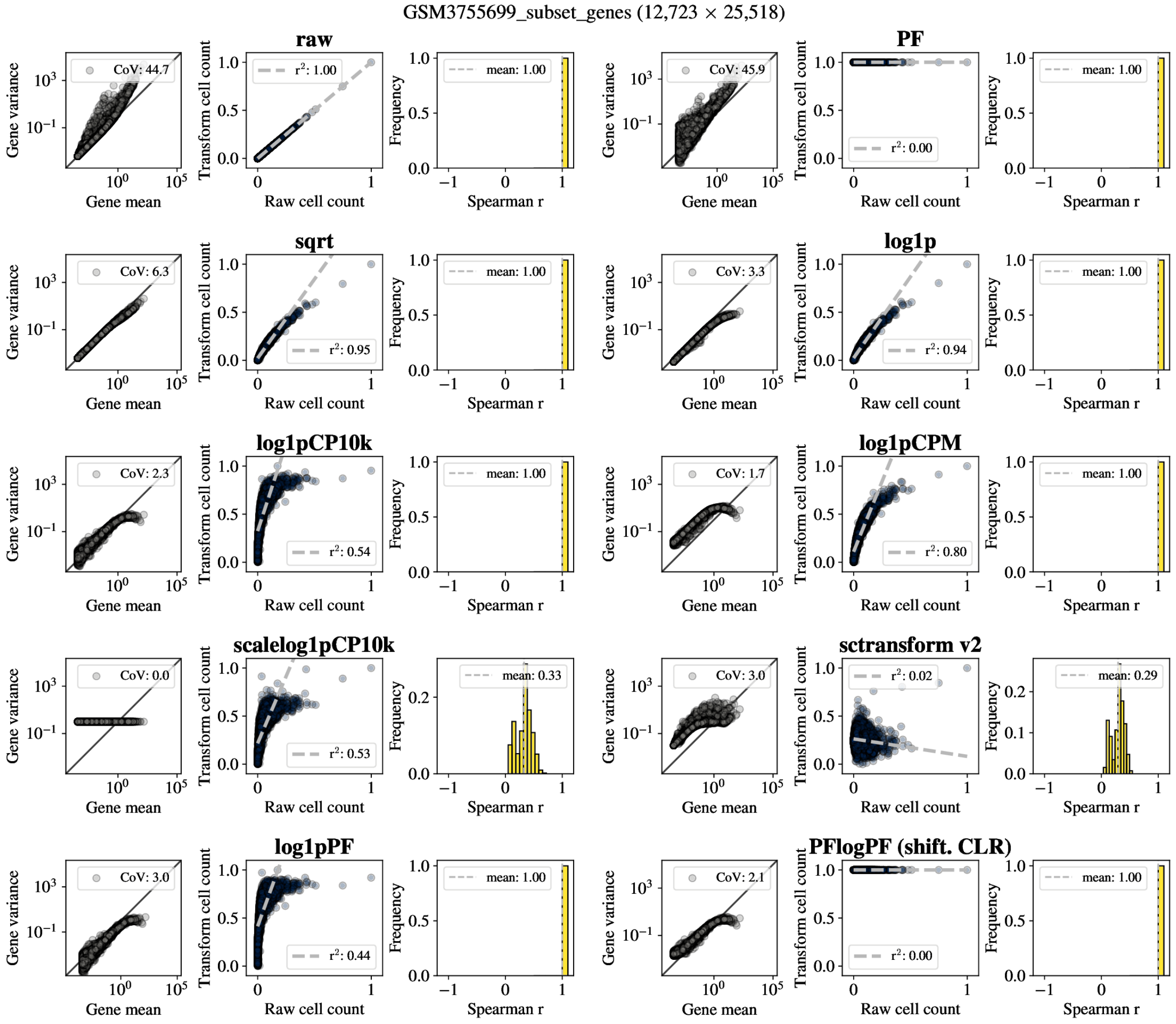


Supplementary Figure 1.423. **GSM3755699**. Per-method comparison of variance stabilization (left), depth normalization (middle), and monotonicity (right) for each normalization on dataset GSM3755699.

| GSM3758115 | |
|---|---|
| cells | 8,509 |
| genes | 60,623 |
| nonzero entries | 7,093,949 |
| density | 0.0138 |
| total counts | 18,390,873 |
| mean counts / cell | 2,161 |
| mean counts / gene | 303 |
| min counts / cell | 276 |
| max counts / cell | 84,412 |
| overdispersion | 1.62 |

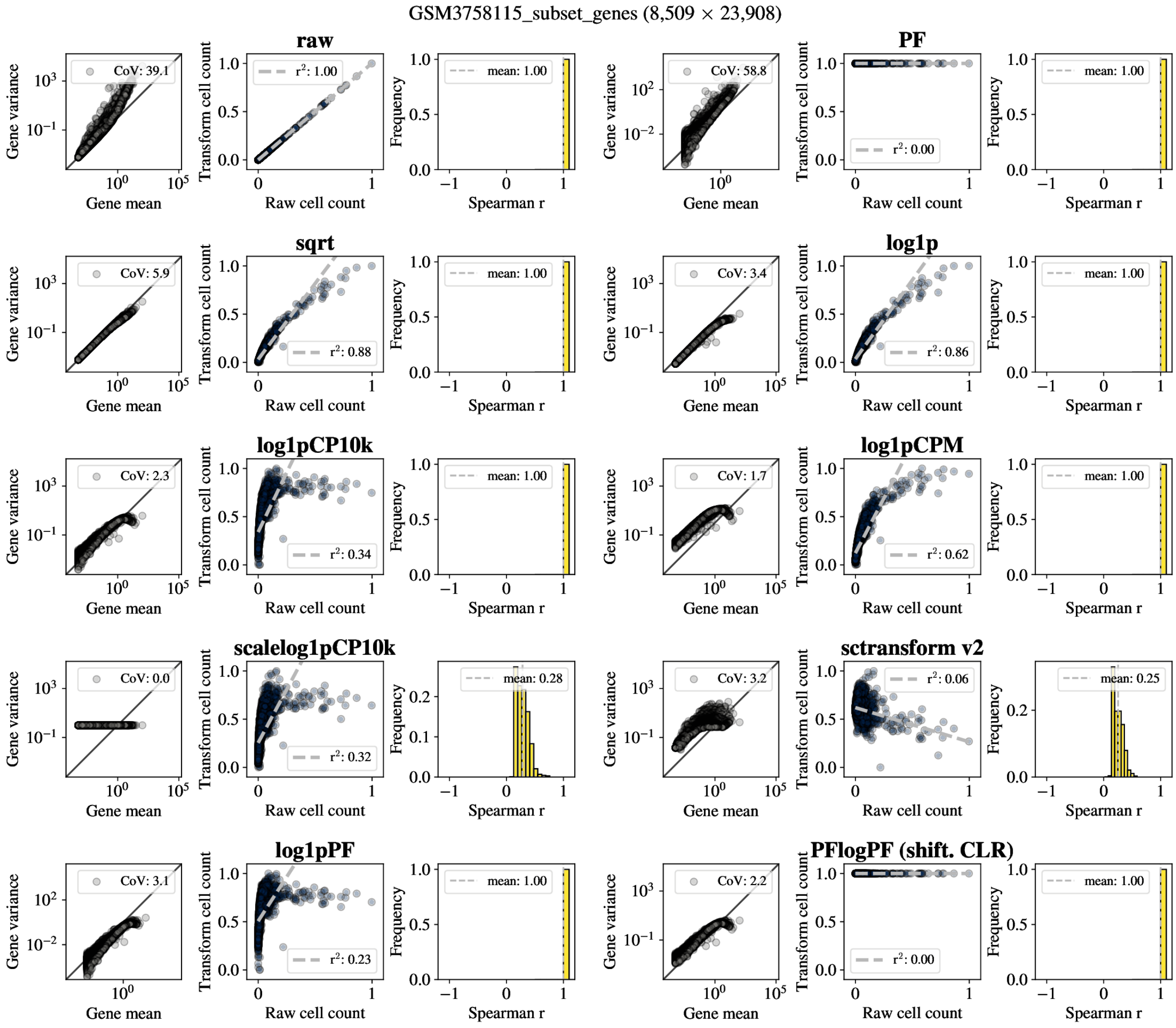


Supplementary Figure 1.424. **GSM3758115**. Per-method comparison of variance stabilization (left), depth normalization (middle), and monotonicity (right) for each normalization on dataset GSM3758115.

| GSM3758116 | |
|---|---|
| cells | 3,867 |
| genes | 60,623 |
| nonzero entries | 3,845,312 |
| density | 0.0164 |
| total counts | 11,700,400 |
| mean counts / cell | 3,026 |
| mean counts / gene | 193 |
| min counts / cell | 207 |
| max counts / cell | 57,966 |
| overdispersion | 5.14 |

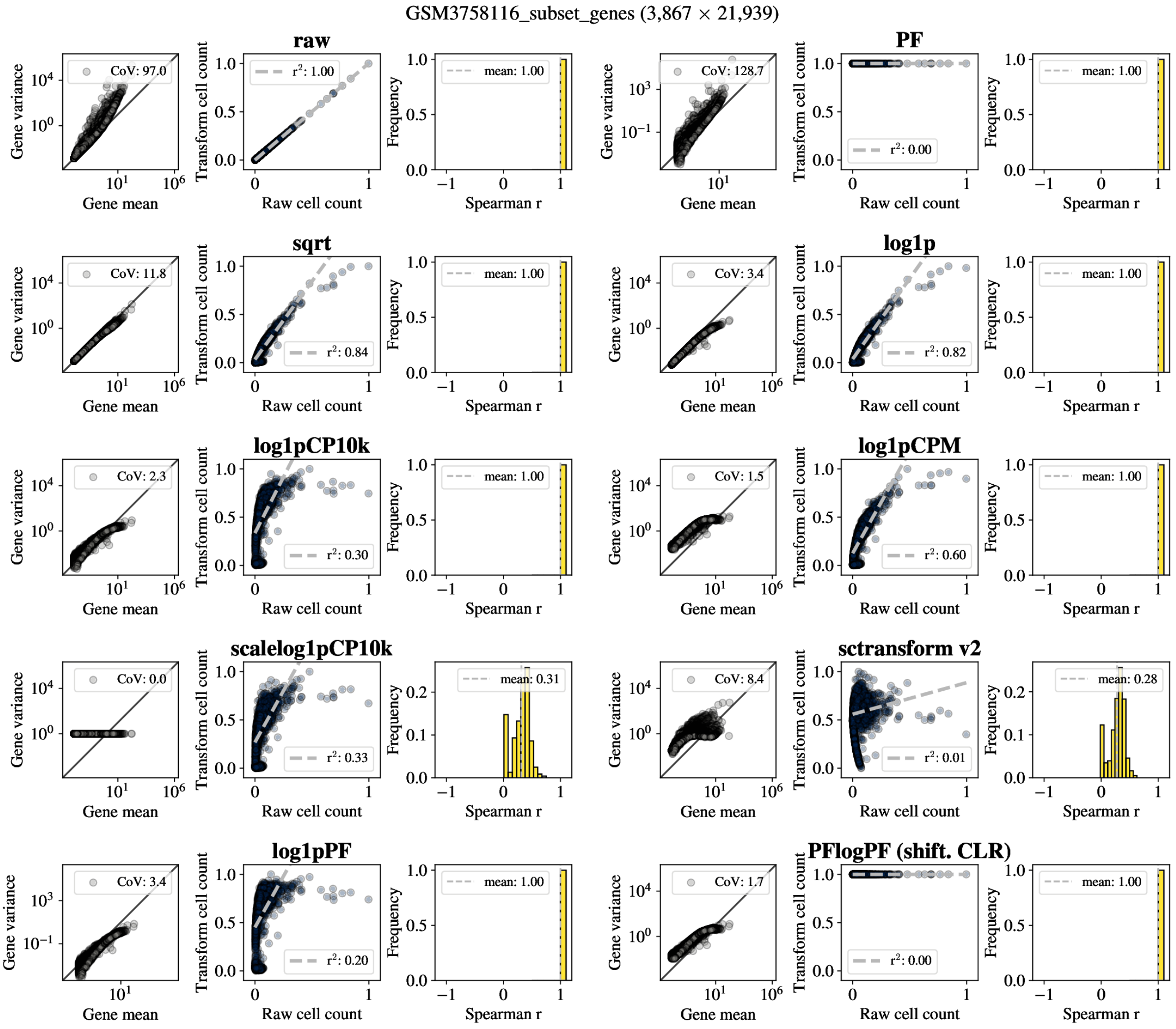


Supplementary Figure 1.425. **GSM3758116**. Per-method comparison of variance stabilization (left), depth normalization (middle), and monotonicity (right) for each normalization on dataset GSM3758116.

| GSM3758117 | |
|---|---|
| cells | 5,881 |
| genes | 60,623 |
| nonzero entries | 5,892,158 |
| density | 0.0165 |
| total counts | 14,965,411 |
| mean counts / cell | 2,545 |
| mean counts / gene | 247 |
| min counts / cell | 303 |
| max counts / cell | 42,381 |
| overdispersion | 0.994 |

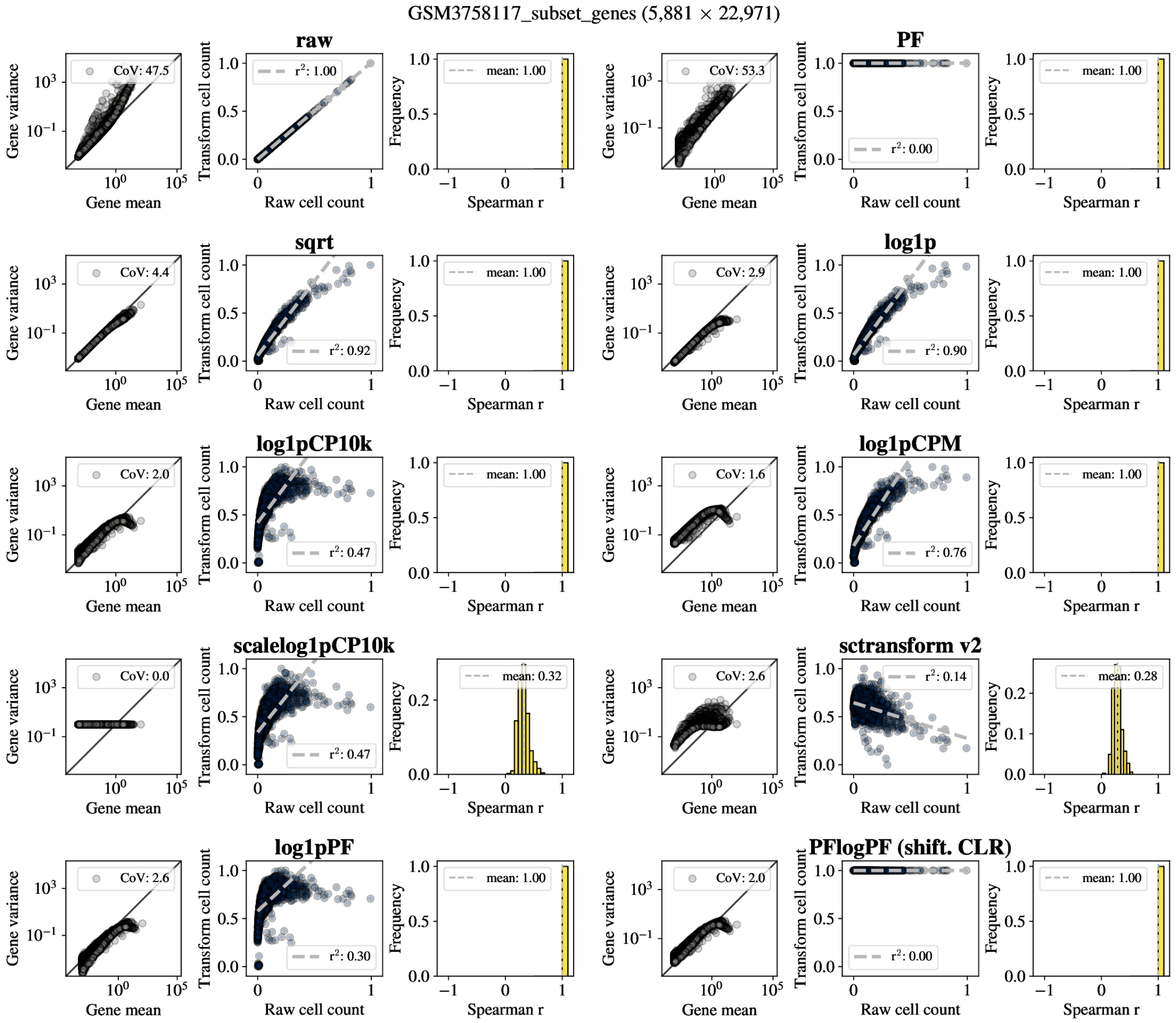


Supplementary Figure 1.426. **GSM3758117**. Per-method comparison of variance stabilization (left), depth normalization (middle), and monotonicity (right) for each normalization on dataset GSM3758117.

| GSM3758118 | |
|---|---|
| cells | 6,192 |
| genes | 60,623 |
| nonzero entries | 5,557,115 |
| density | 0.0148 |
| total counts | 14,131,299 |
| mean counts / cell | 2,282 |
| mean counts / gene | 233 |
| min counts / cell | 276 |
| max counts / cell | 47,419 |
| overdispersion | 0.912 |

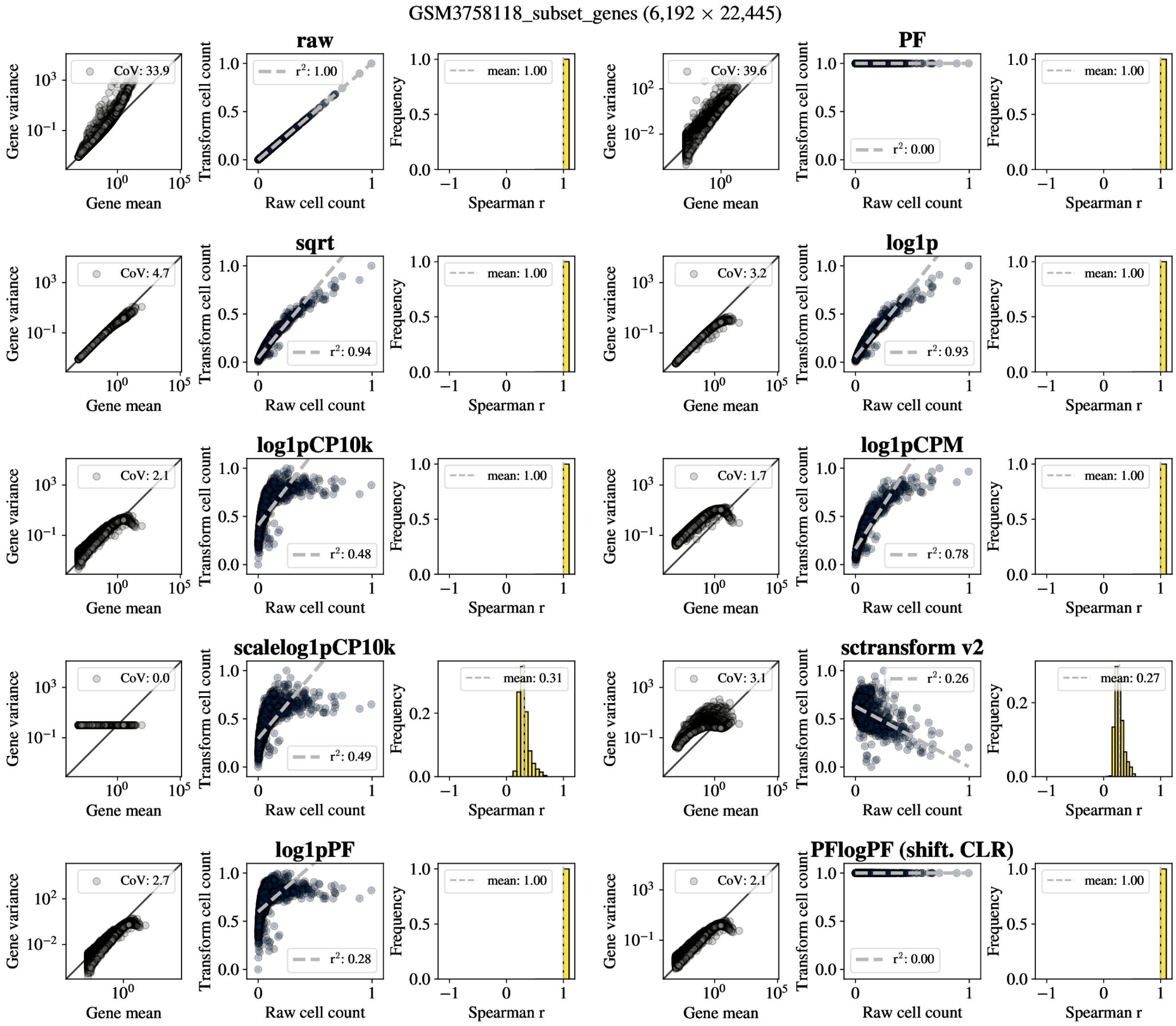


Supplementary Figure 1.427. **GSM3758118**. Per-method comparison of variance stabilization (left), depth normalization (middle), and monotonicity (right) for each normalization on dataset GSM3758118.

| GSM3758119 | |
| --- | --- |
| cells | 6,397 |
| genes | 60,623 |
| nonzero entries | 5,060,300 |
| density | 0.013 |
| total counts | 11,848,669 |
| mean counts / cell | 1,852 |
| mean counts / gene | 195 |
| min counts / cell | 295 |
| max counts / cell | 22,549 |
| overdispersion | 0.715 |

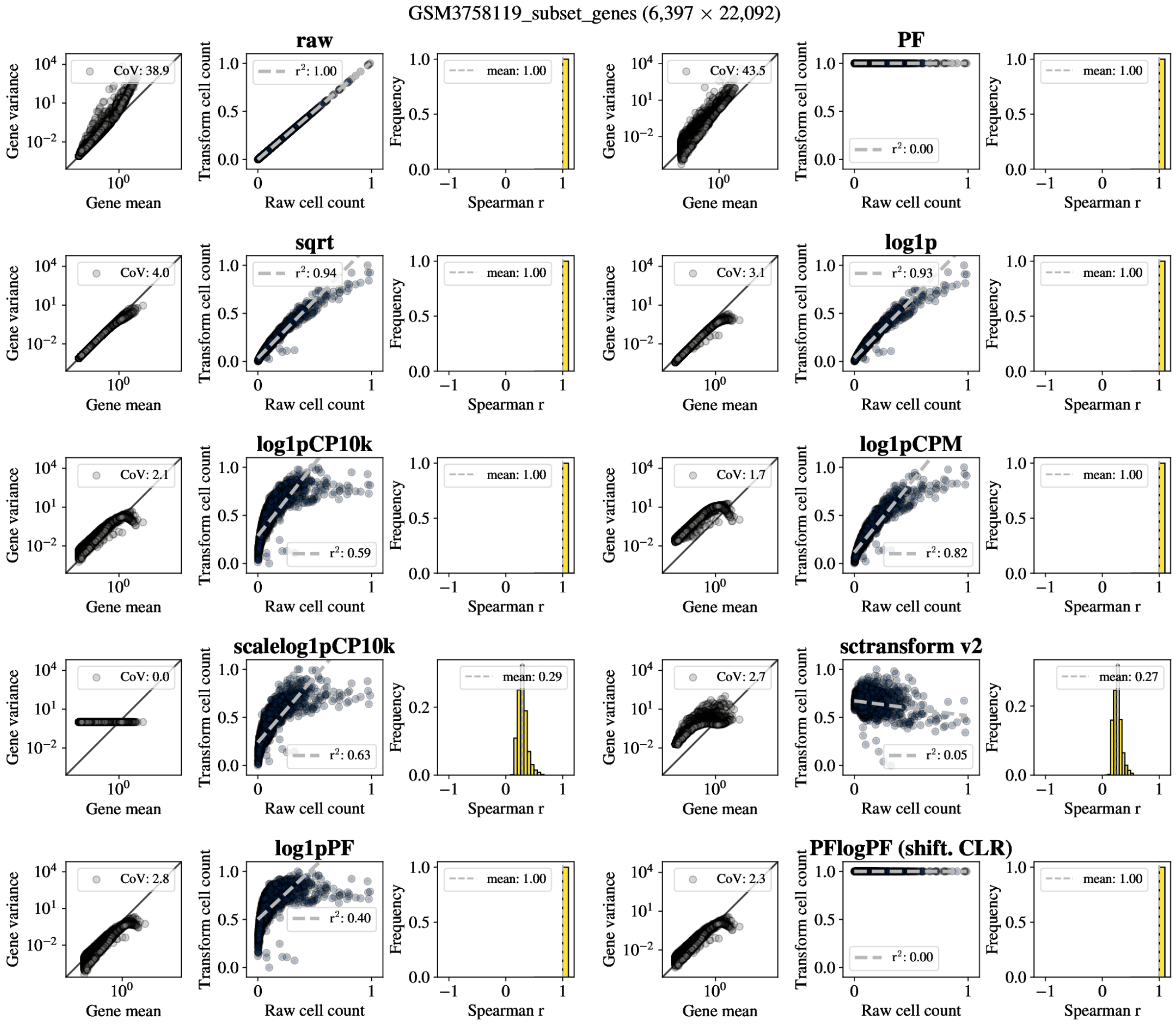


Supplementary Figure 1.428. **GSM3758119**. Per-method comparison of variance stabilization (left), depth normalization (middle), and monotonicity (right) for each normalization on dataset GSM3758119.

| GSM3770749 | |
|---|---|
| cells | 6,028 |
| genes | 60,623 |
| nonzero entries | 11,454,886 |
| density | 0.0313 |
| total counts | 23,206,413 |
| mean counts / cell | 3,850 |
| mean counts / gene | 383 |
| min counts / cell | 1,372 |
| max counts / cell | 48,028 |
| overdispersion | 0.367 |

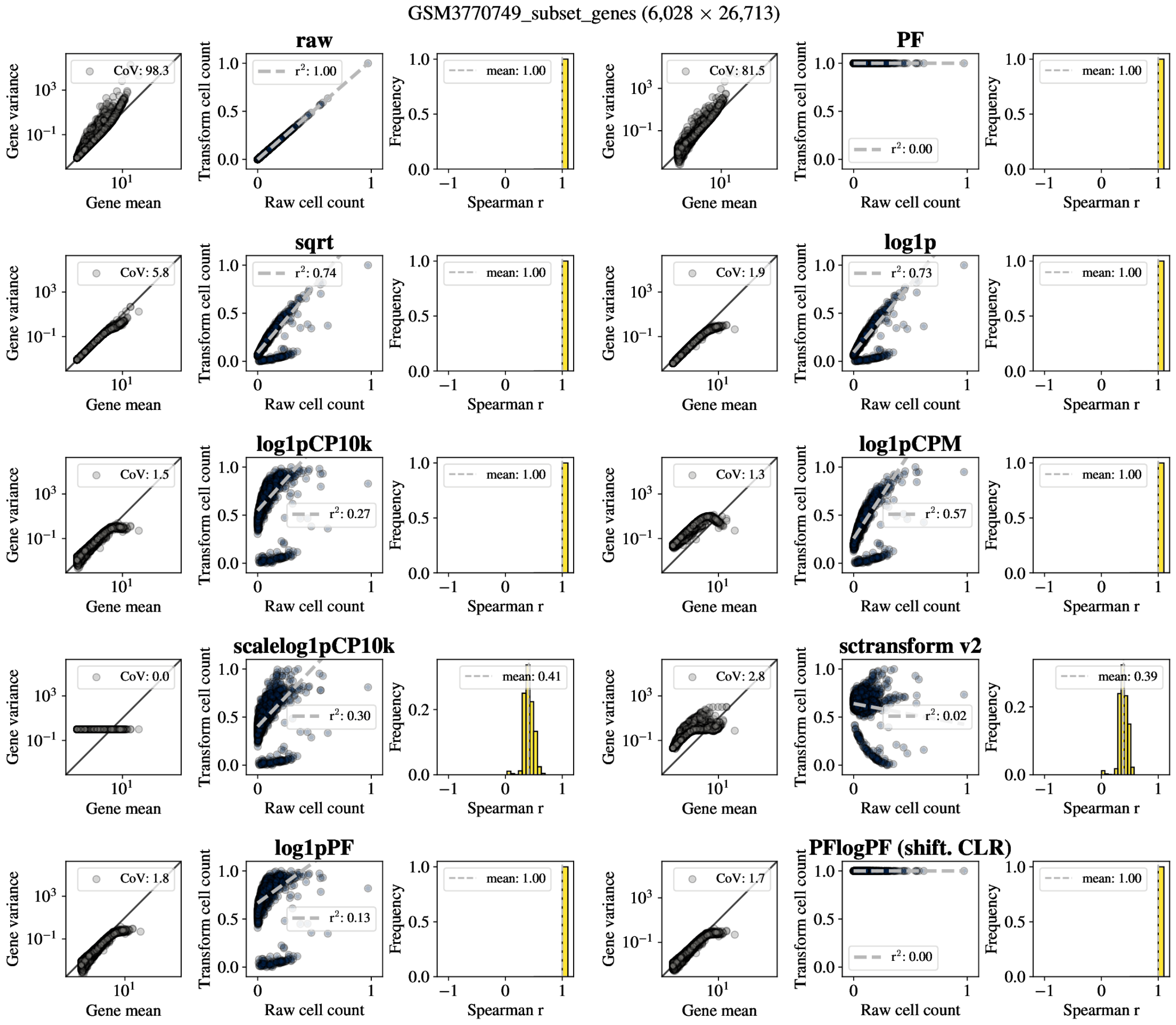


Supplementary Figure 1.429. **GSM3770749**. Per-method comparison of variance stabilization (left), depth normalization (middle), and monotonicity (right) for each normalization on dataset GSM3770749.

| GSM3770750 | |
|---|---|
| cells | 5,929 |
| genes | 60,623 |
| nonzero entries | 11,345,273 |
| density | 0.0316 |
| total counts | 23,266,612 |
| mean counts / cell | 3,924 |
| mean counts / gene | 384 |
| min counts / cell | 1,438 |
| max counts / cell | 35,319 |
| overdispersion | 0.371 |

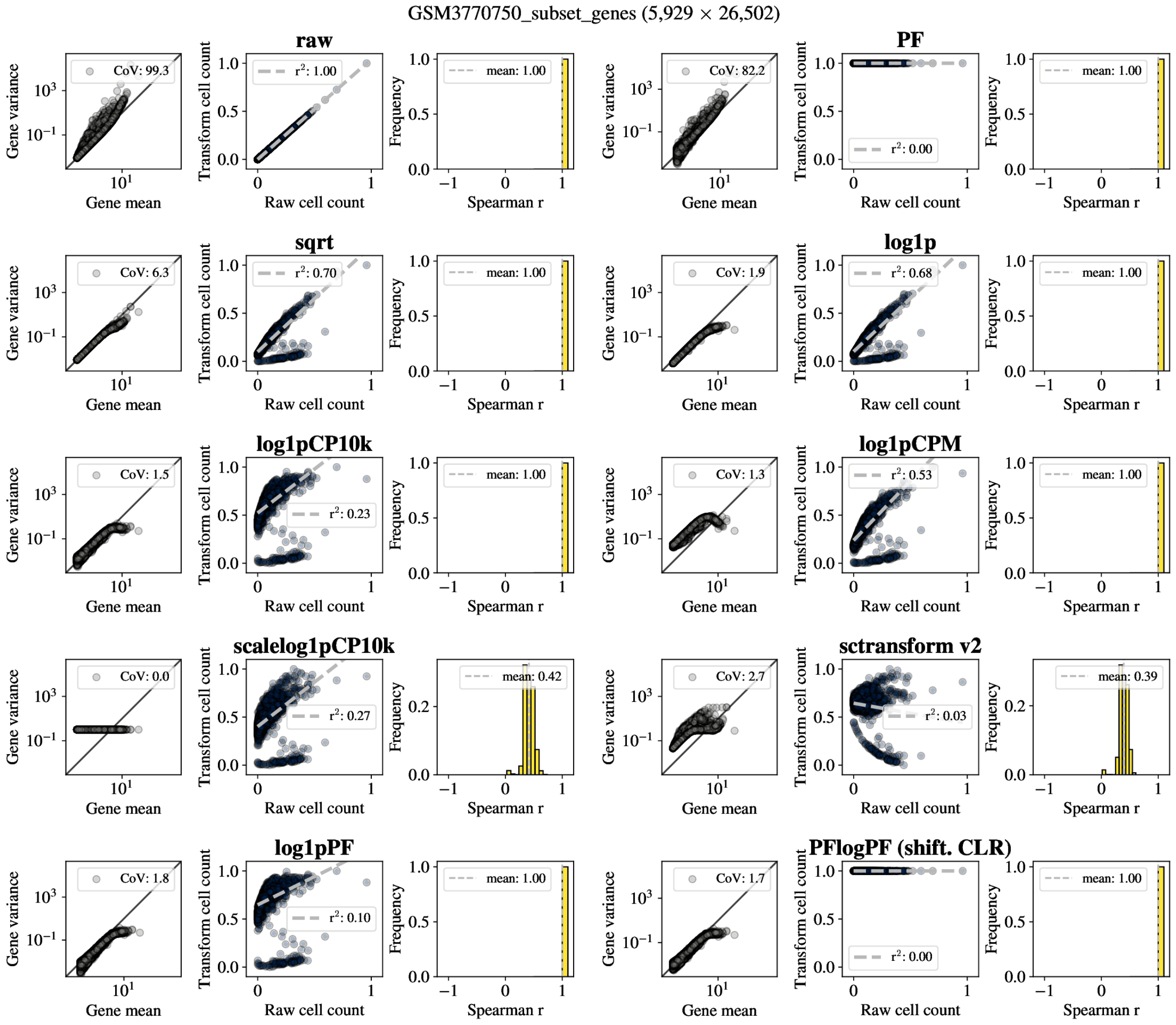


Supplementary Figure 1.430. **GSM3770750**. Per-method comparison of variance stabilization (left), depth normalization (middle), and monotonicity (right) for each normalization on dataset GSM3770750.

| GSM3773108 | |
|---|---|
| cells | 77 |
| genes | 60,623 |
| nonzero entries | 9,979 |
| density | 0.00214 |
| total counts | 16,238 |
| mean counts / cell | 211 |
| mean counts / gene | 0 |
| min counts / cell | 94 |
| max counts / cell | 405 |
| overdispersion | 0.488 |

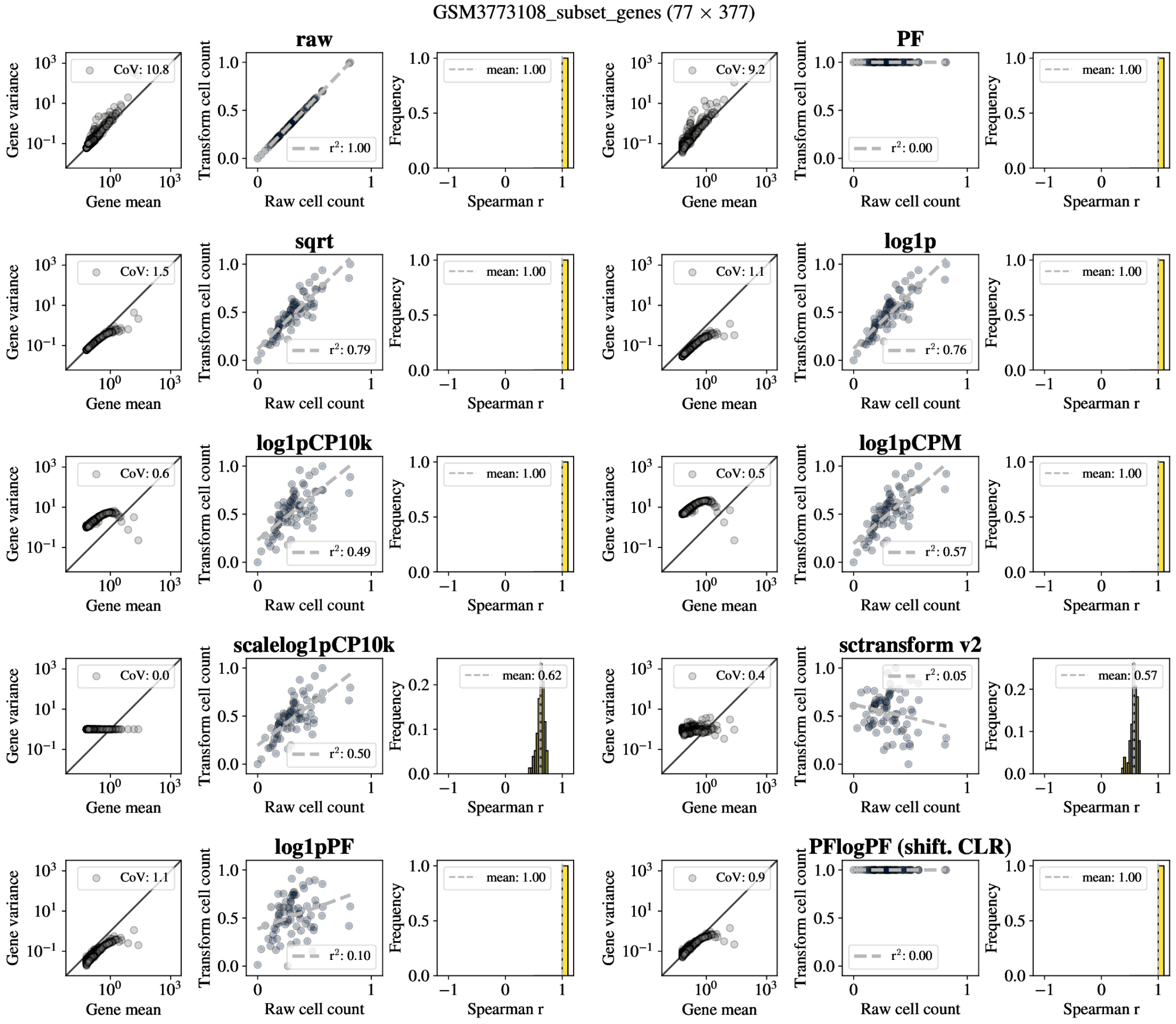


Supplementary Figure 1.431. **GSM3773108**. Per-method comparison of variance stabilization (left), depth normalization (middle), and monotonicity (right) for each normalization on dataset GSM3773108.

| GSM3773111 | |
|---|---|
| cells | 79 |
| genes | 60,623 |
| nonzero entries | 9,515 |
| density | 0.00199 |
| total counts | 16,441 |
| mean counts / cell | 208 |
| mean counts / gene | 0 |
| min counts / cell | 89 |
| max counts / cell | 430 |
| overdispersion | 0.566 |

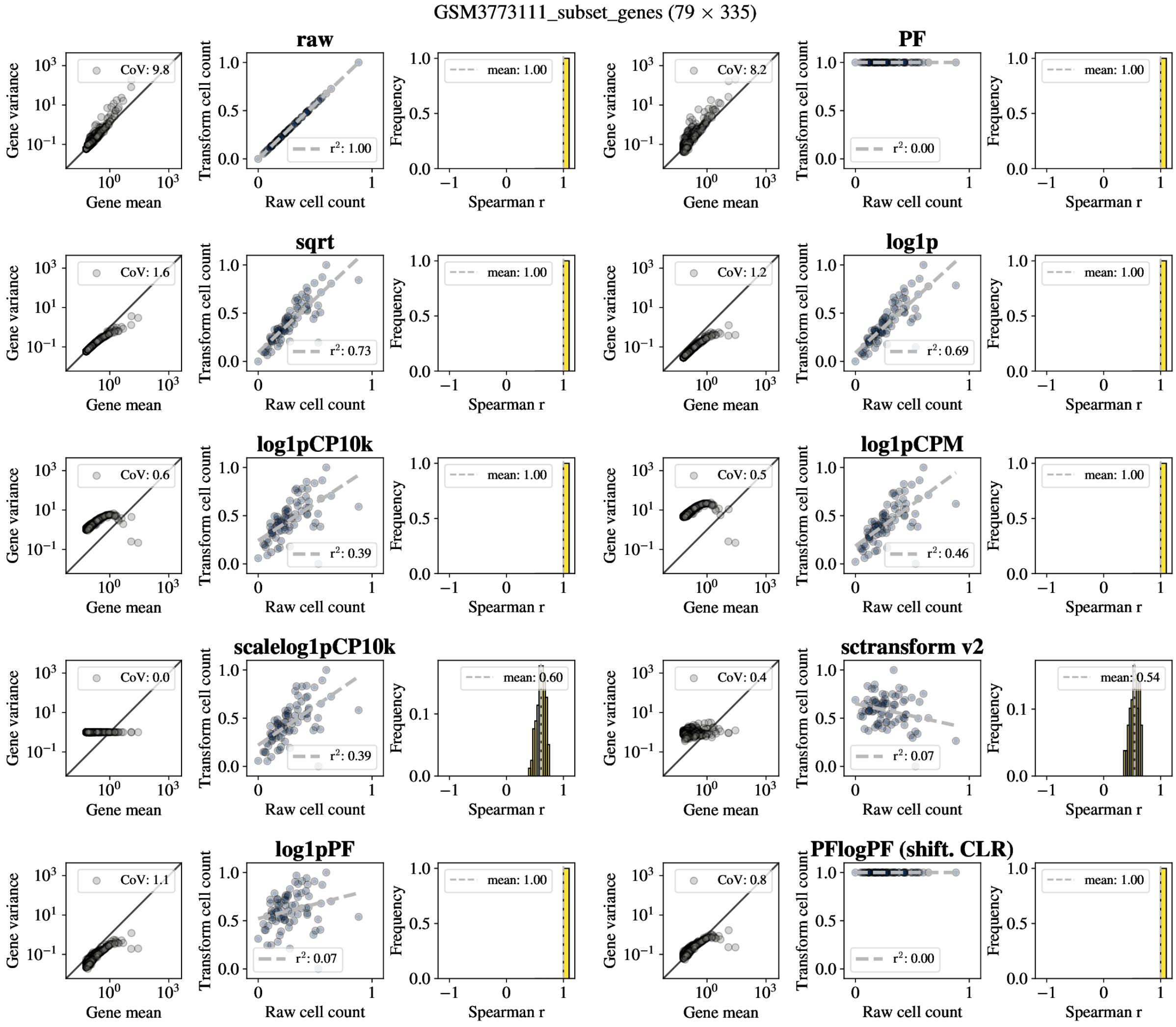


Supplementary Figure 1.432. **GSM3773111**. Per-method comparison of variance stabilization (left), depth normalization (middle), and monotonicity (right) for each normalization on dataset GSM3773111.

| GSM3773112 | |
|---|---|
| cells | 79 |
| genes | 60,623 |
| nonzero entries | 8,383 |
| density | 0.00175 |
| total counts | 15,586 |
| mean counts / cell | 197 |
| mean counts / gene | 0 |
| min counts / cell | 63 |
| max counts / cell | 456 |
| overdispersion | 0.727 |

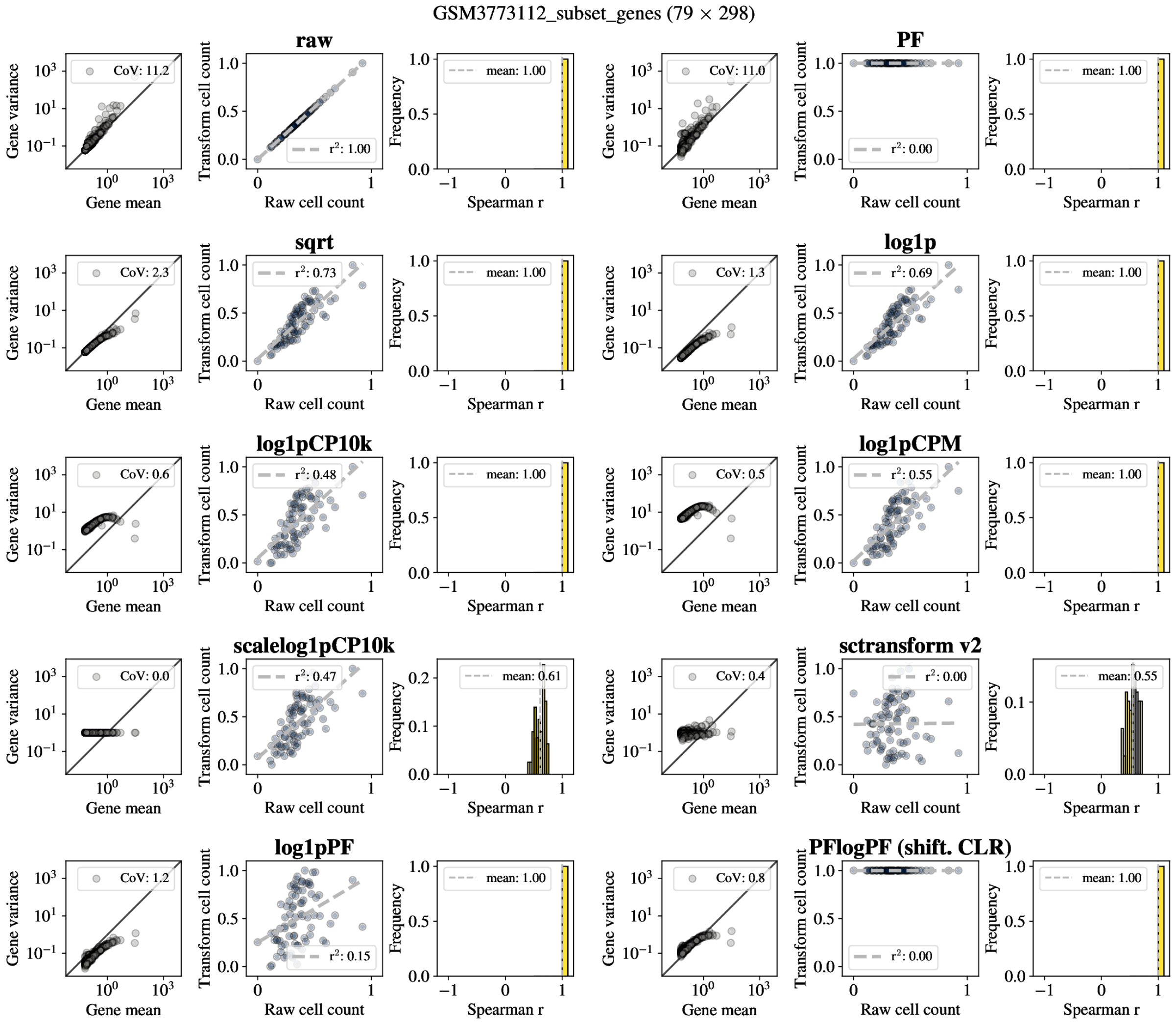


Supplementary Figure 1.433. **GSM3773112**. Per-method comparison of variance stabilization (left), depth normalization (middle), and monotonicity (right) for each normalization on dataset GSM3773112.

| GSM3773114 | |
|---|---|
| cells | 62 |
| genes | 60,623 |
| nonzero entries | 7,380 |
| density | 0.00196 |
| total counts | 14,698 |
| mean counts / cell | 237 |
| mean counts / gene | 0 |
| min counts / cell | 64 |
| max counts / cell | 915 |
| overdispersion | 0.607 |

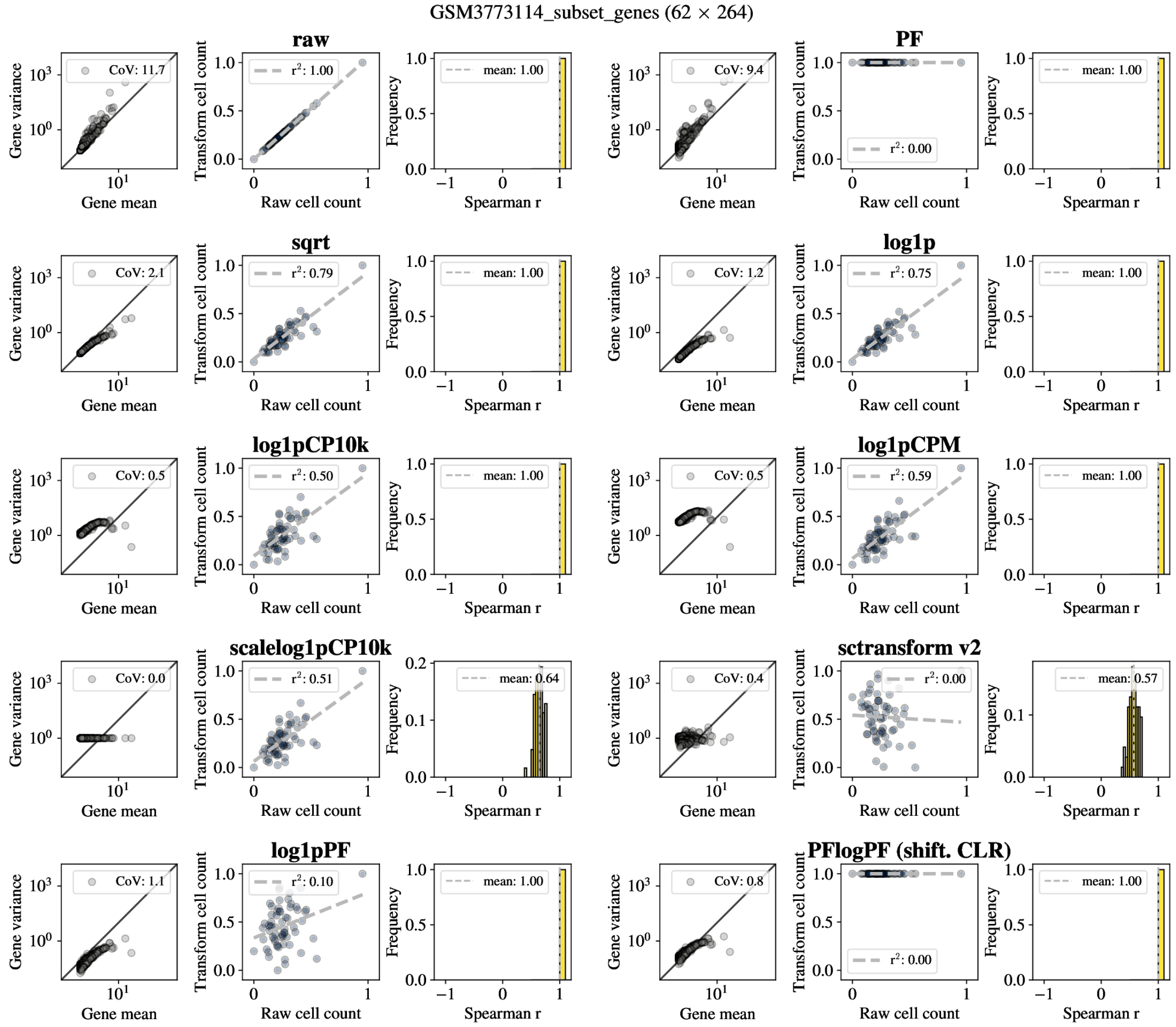


Supplementary Figure 1.434. **GSM3773114**. Per-method comparison of variance stabilization (left), depth normalization (middle), and monotonicity (right) for each normalization on dataset GSM3773114.

| GSM3773116 | |
|---|---|
| cells | 84 |
| genes | 60,623 |
| nonzero entries | 9,131 |
| density | 0.00179 |
| total counts | 17,635 |
| mean counts / cell | 210 |
| mean counts / gene | 0 |
| min counts / cell | 69 |
| max counts / cell | 449 |
| overdispersion | 0.319 |

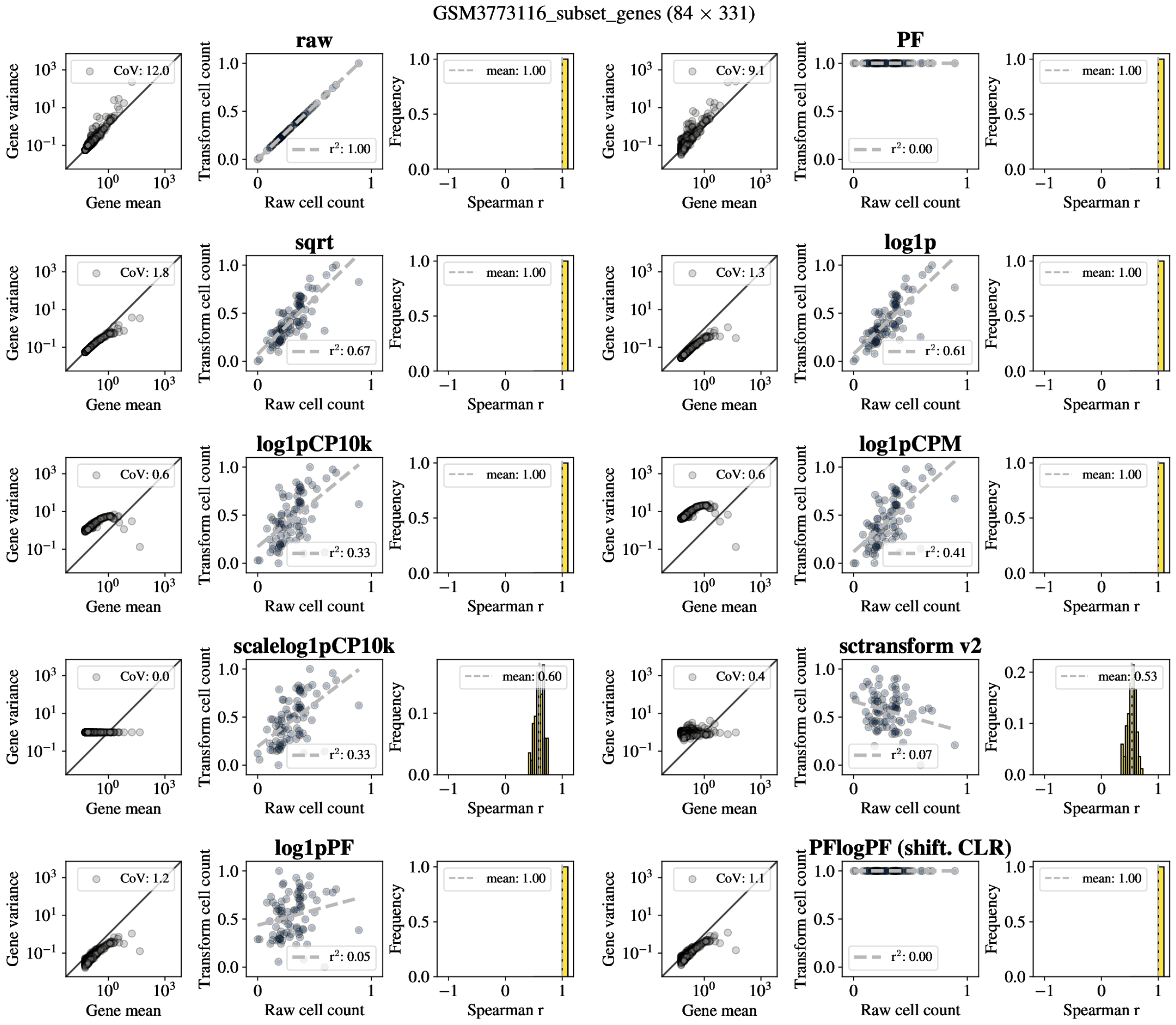


Supplementary Figure 1.435. **GSM3773116**. Per-method comparison of variance stabilization (left), depth normalization (middle), and monotonicity (right) for each normalization on dataset GSM3773116.

| GSM3773117 | |
|---|---|
| cells | 71 |
| genes | 60,623 |
| nonzero entries | 8,498 |
| density | 0.00197 |
| total counts | 16,080 |
| mean counts / cell | 226 |
| mean counts / gene | 0 |
| min counts / cell | 90 |
| max counts / cell | 666 |
| overdispersion | 0.628 |

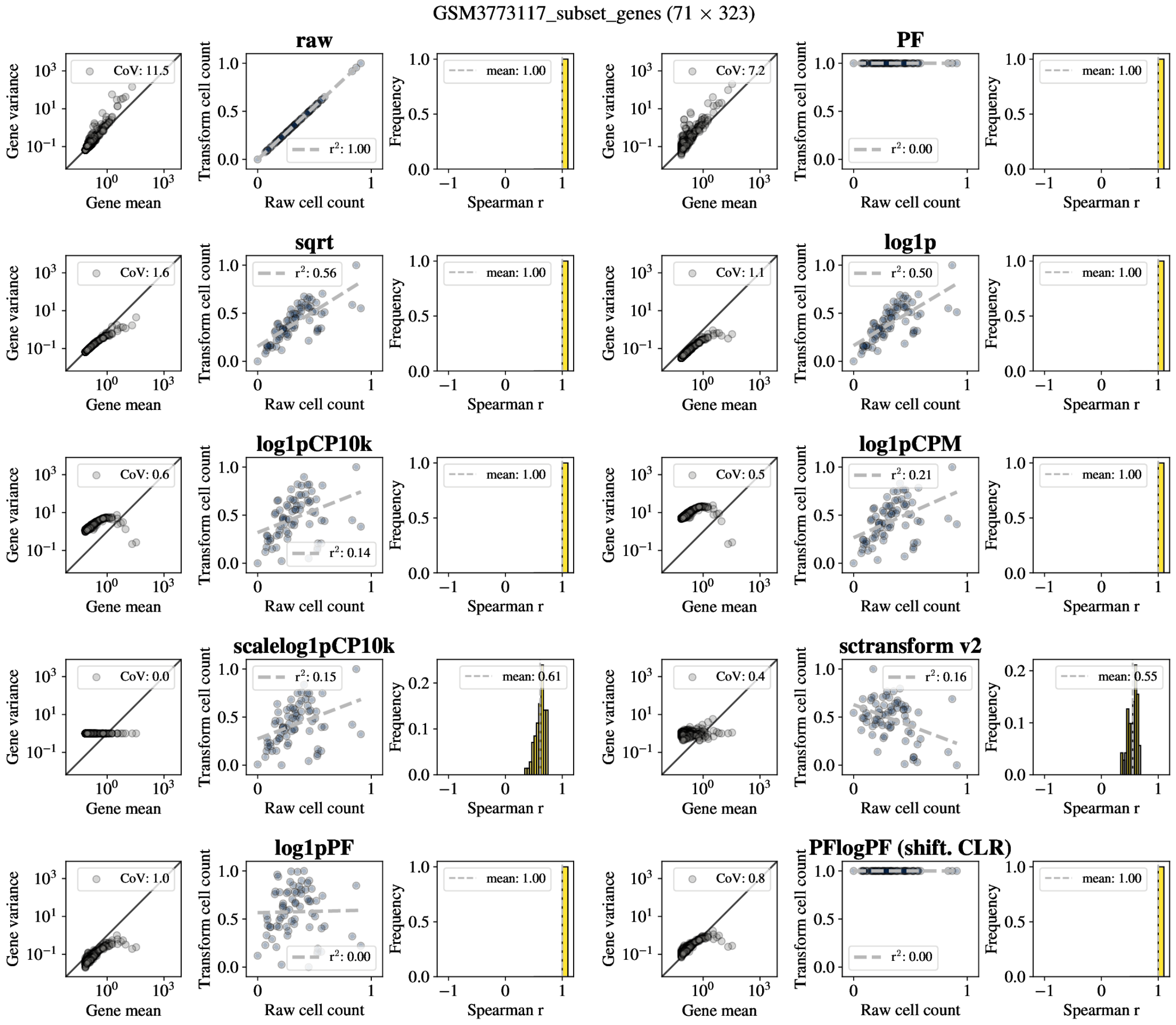


Supplementary Figure 1.436. **GSM3773117**. Per-method comparison of variance stabilization (left), depth normalization (middle), and monotonicity (right) for each normalization on dataset GSM3773117.

| GSM3773119 | |
|---|---|
| cells | 76 |
| genes | 60,623 |
| nonzero entries | 8,887 |
| density | 0.00193 |
| total counts | 15,458 |
| mean counts / cell | 203 |
| mean counts / gene | 0 |
| min counts / cell | 83 |
| max counts / cell | 540 |
| overdispersion | 0.519 |

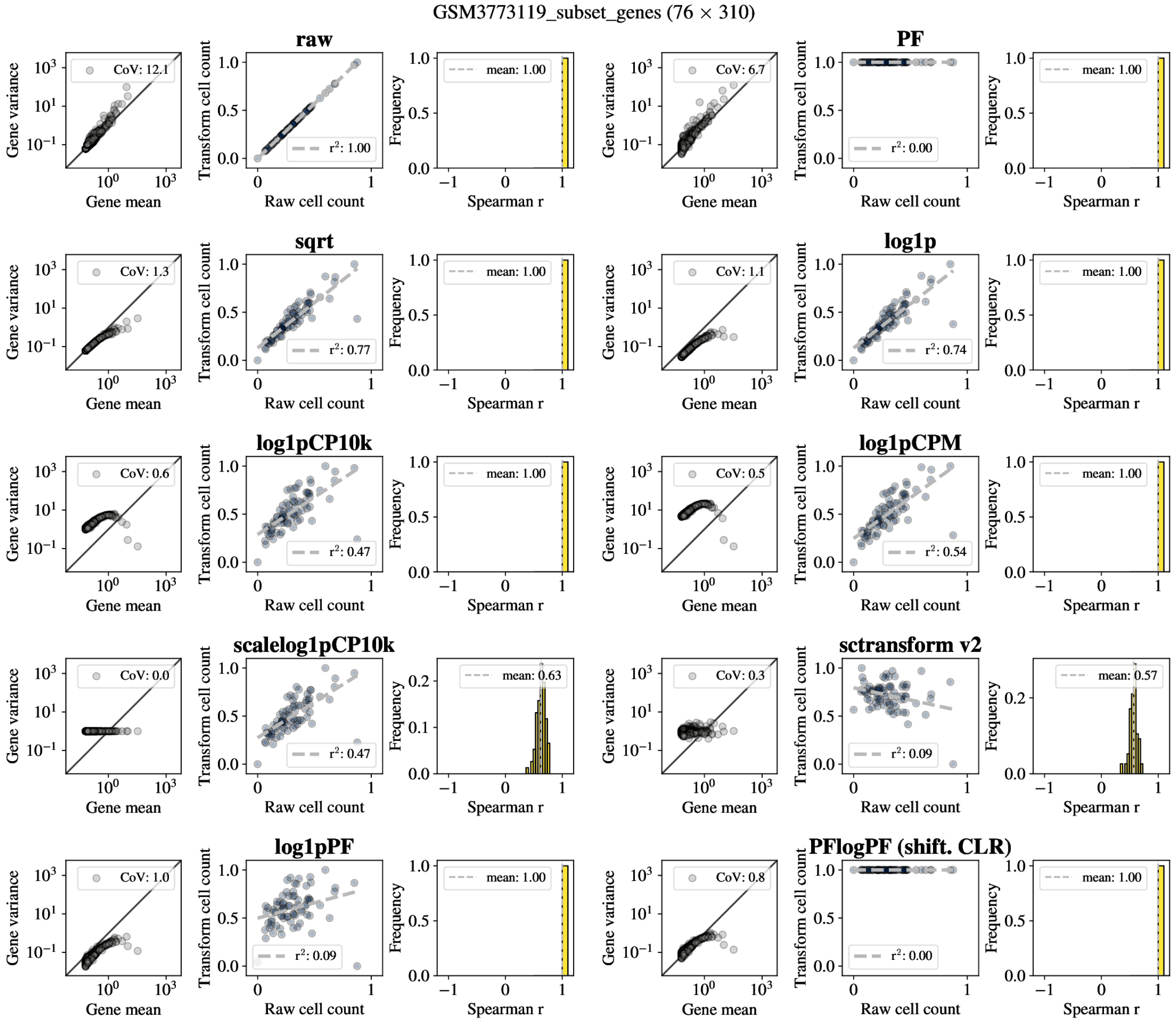


Supplementary Figure 1.437. **GSM3773119**. Per-method comparison of variance stabilization (left), depth normalization (middle), and monotonicity (right) for each normalization on dataset GSM3773119.

| GSM3773121 | |
|---|---|
| cells | 30 |
| genes | 60,623 |
| nonzero entries | 5,567 |
| density | 0.00306 |
| total counts | 10,589 |
| mean counts / cell | 353 |
| mean counts / gene | 0 |
| min counts / cell | 73 |
| max counts / cell | 740 |
| overdispersion | 0.429 |

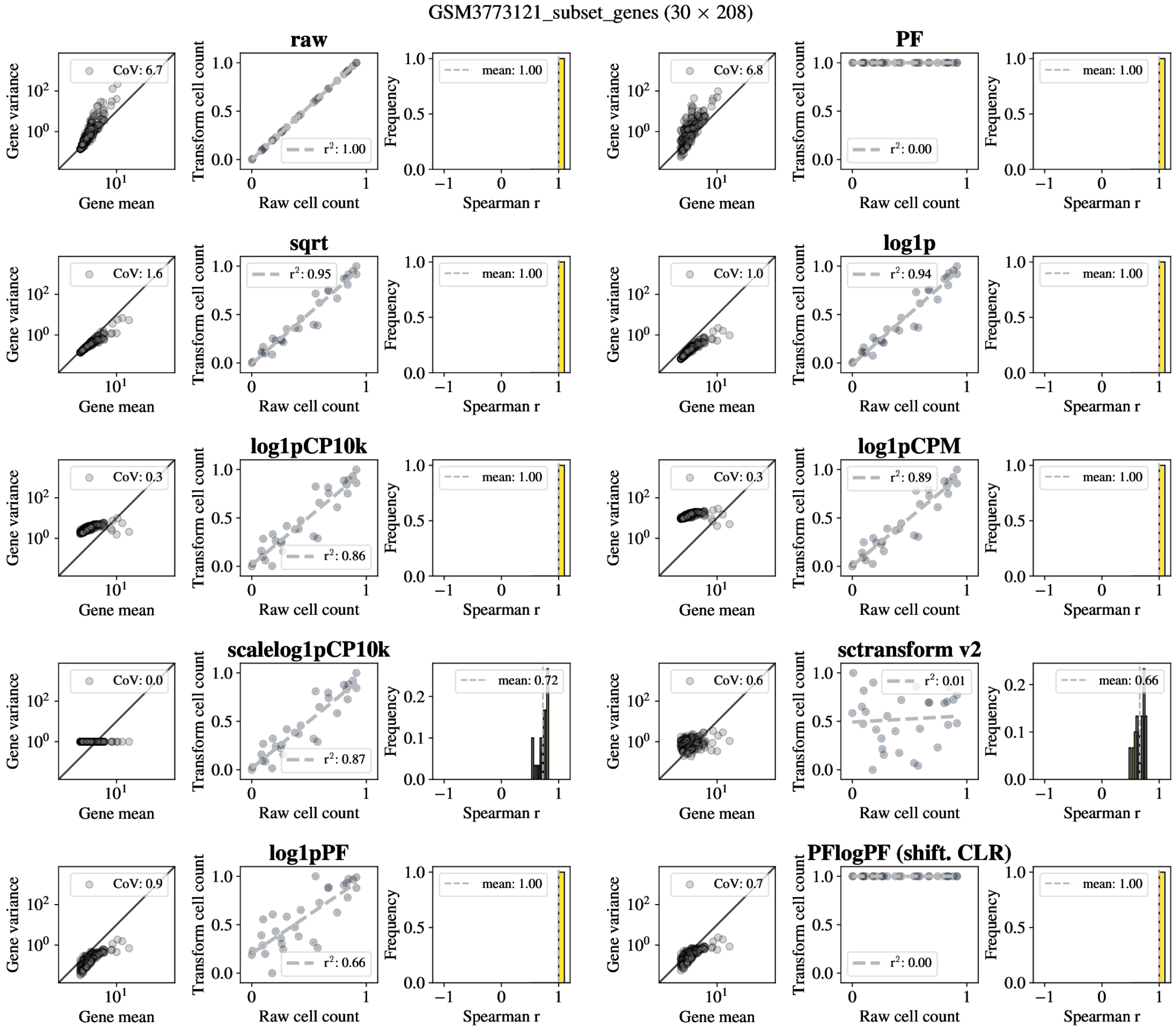


Supplementary Figure 1.438. **GSM3773121**. Per-method comparison of variance stabilization (left), depth normalization (middle), and monotonicity (right) for each normalization on dataset GSM3773121.

| **GSM3773124** | |
|---|---|
| cells | 66 |
| genes | 60,623 |
| nonzero entries | 8,733 |
| density | 0.00218 |
| total counts | 14,079 |
| mean counts / cell | 213 |
| mean counts / gene | 0 |
| min counts / cell | 67 |
| max counts / cell | 519 |
| overdispersion | 0.444 |

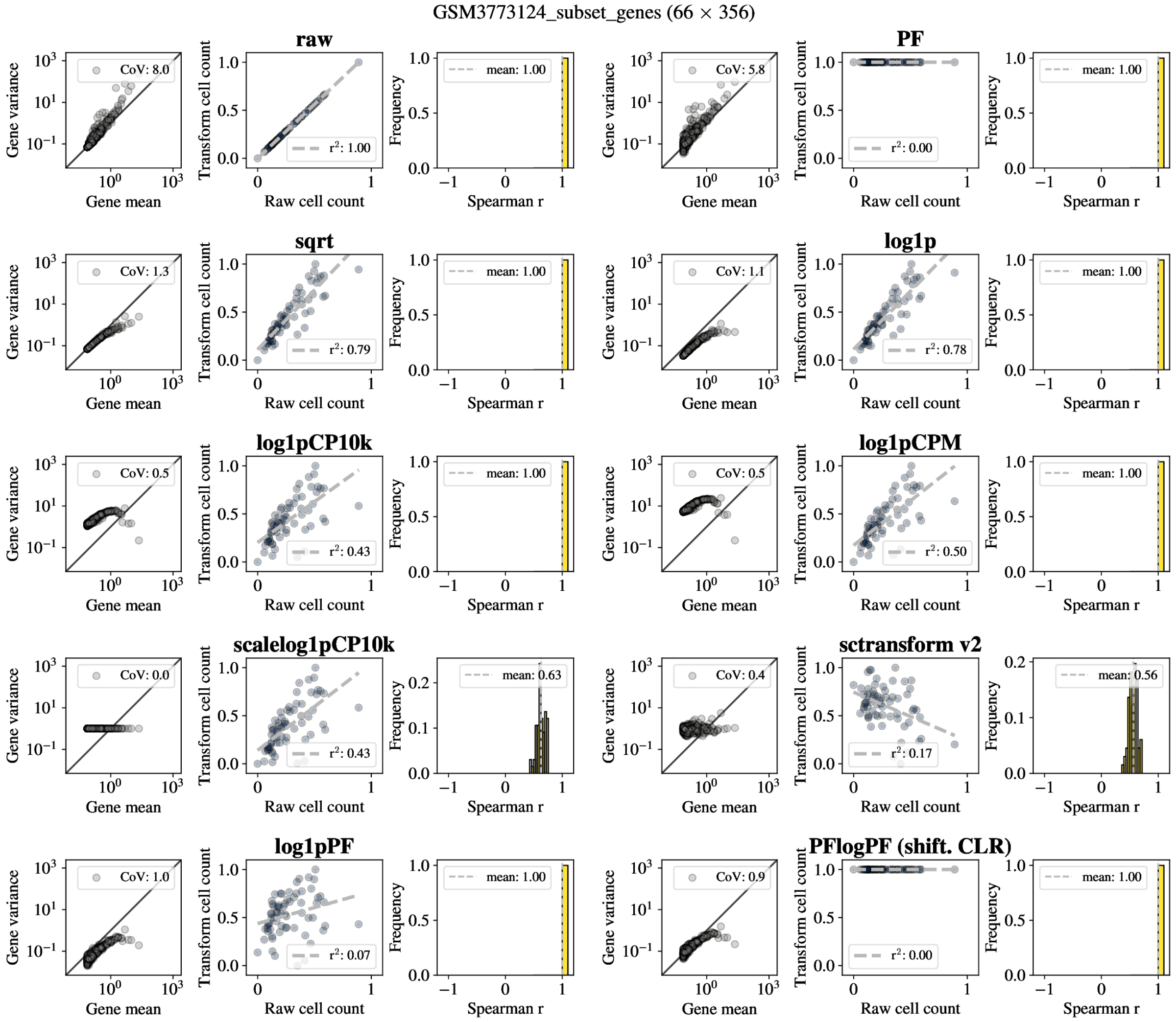


Supplementary Figure 1.439. **GSM3773124**. Per-method comparison of variance stabilization (left), depth normalization (middle), and monotonicity (right) for each normalization on dataset GSM3773124.

| GSM3773125 | |
|---|---|
| cells | 83 |
| genes | 60,623 |
| nonzero entries | 9,924 |
| density | 0.00197 |
| total counts | 17,423 |
| mean counts / cell | 210 |
| mean counts / gene | 0 |
| min counts / cell | 84 |
| max counts / cell | 581 |
| overdispersion | 0.649 |

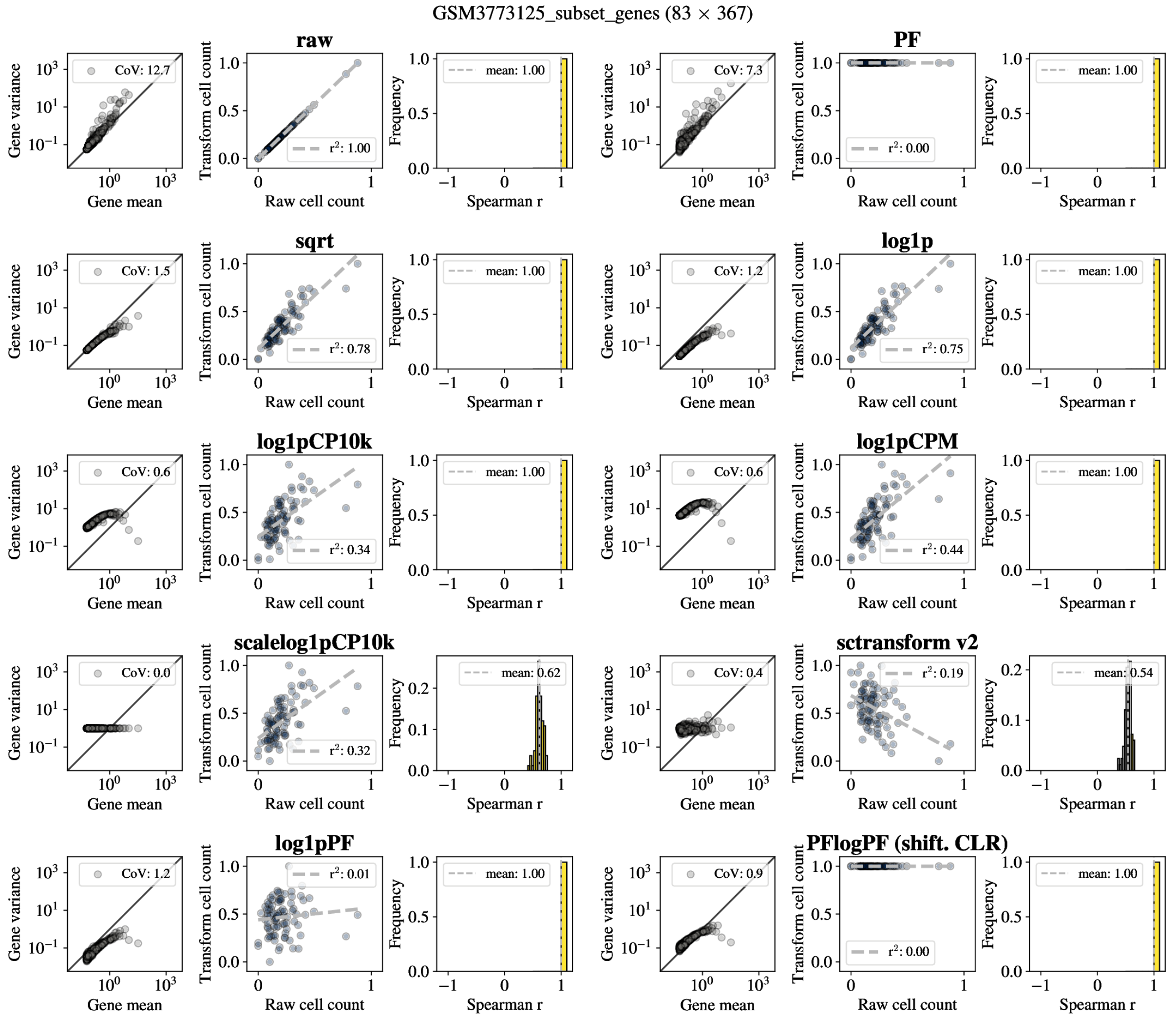


Supplementary Figure 1.440. **GSM3773125**. Per-method comparison of variance stabilization (left), depth normalization (middle), and monotonicity (right) for each normalization on dataset GSM3773125.

| GSM3954946 | |
|---|---|
| cells | 3,505 |
| genes | 60,623 |
| nonzero entries | 2,830,797 |
| density | 0.0133 |
| total counts | 8,492,308 |
| mean counts / cell | 2,423 |
| mean counts / gene | 140 |
| min counts / cell | 267 |
| max counts / cell | 20,891 |
| overdispersion | 2.11 |

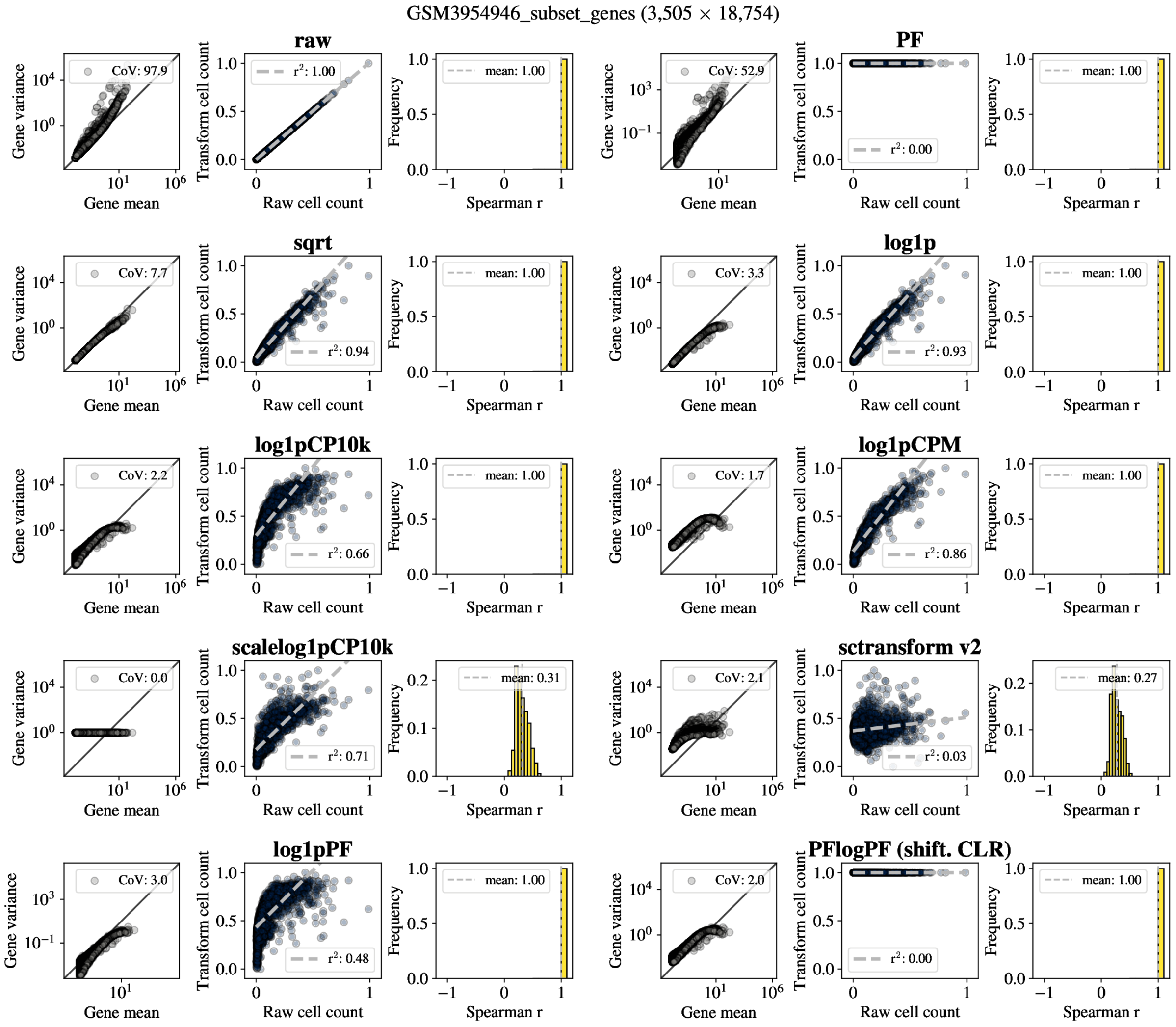


Supplementary Figure 1.441. **GSM3954946**. Per-method comparison of variance stabilization (left), depth normalization (middle), and monotonicity (right) for each normalization on dataset GSM3954946.

| GSM3954947 | |
|---|---|
| cells | 2,158 |
| genes | 60,623 |
| nonzero entries | 2,378,382 |
| density | 0.0182 |
| total counts | 9,021,247 |
| mean counts / cell | 4,180 |
| mean counts / gene | 149 |
| min counts / cell | 409 |
| max counts / cell | 41,414 |
| overdispersion | 31.9 |

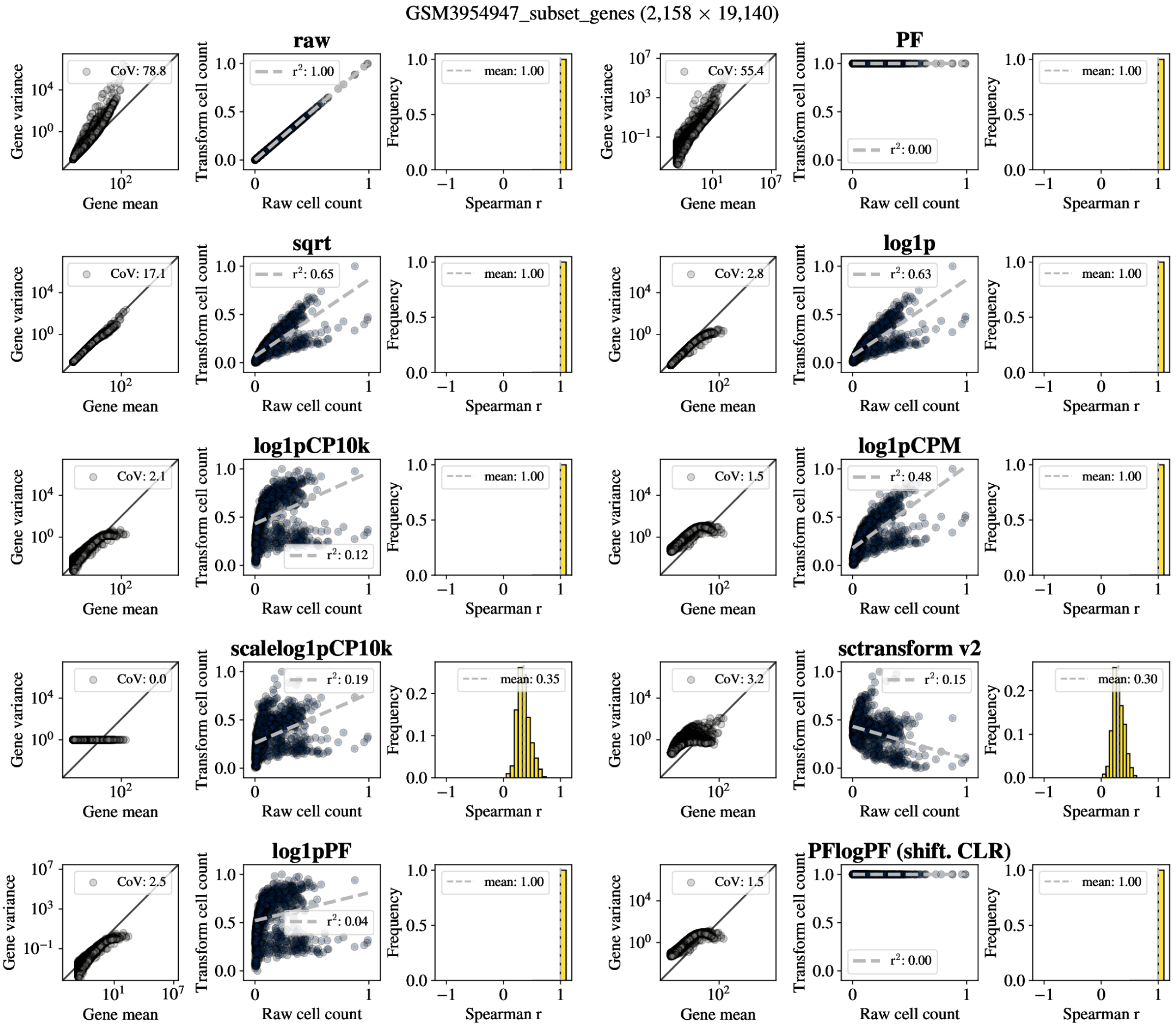


Supplementary Figure 1.442. **GSM3954947**. Per-method comparison of variance stabilization (left), depth normalization (middle), and monotonicity (right) for each normalization on dataset GSM3954947.

| GSM3954948 | |
|---|---|
| cells | 3,151 |
| genes | 60,623 |
| nonzero entries | 2,574,709 |
| density | 0.0135 |
| total counts | 9,128,694 |
| mean counts / cell | 2,897 |
| mean counts / gene | 151 |
| min counts / cell | 219 |
| max counts / cell | 43,978 |
| overdispersion | 29.7 |

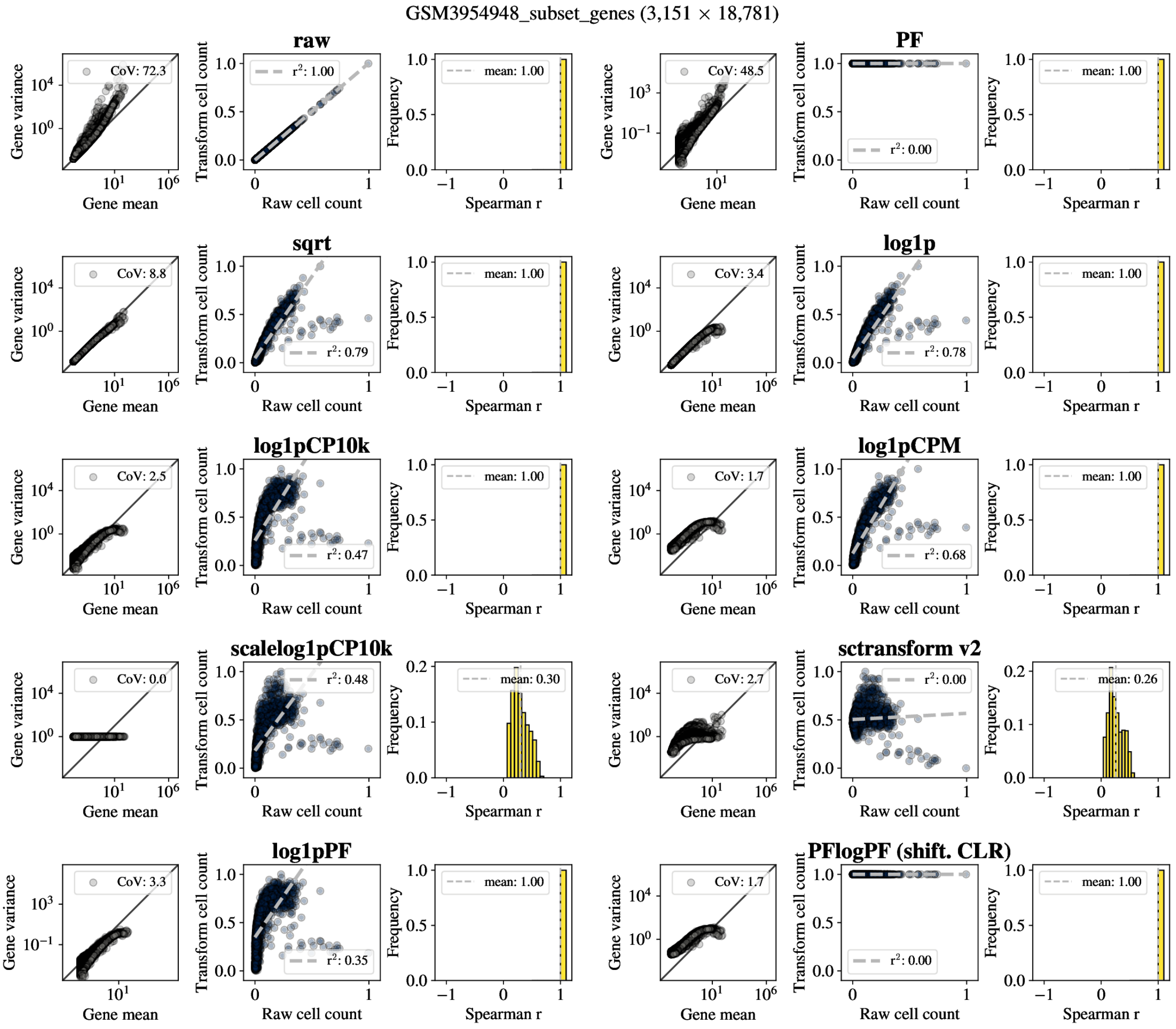


Supplementary Figure 1.443. **GSM3954948**. Per-method comparison of variance stabilization (left), depth normalization (middle), and monotonicity (right) for each normalization on dataset GSM3954948.

| GSM3954949 | |
|---|---|
| cells | 5,630 |
| genes | 60,623 |
| nonzero entries | 3,884,050 |
| density | 0.0114 |
| total counts | 11,387,402 |
| mean counts / cell | 2,023 |
| mean counts / gene | 188 |
| min counts / cell | 238 |
| max counts / cell | 20,108 |
| overdispersion | 19.6 |

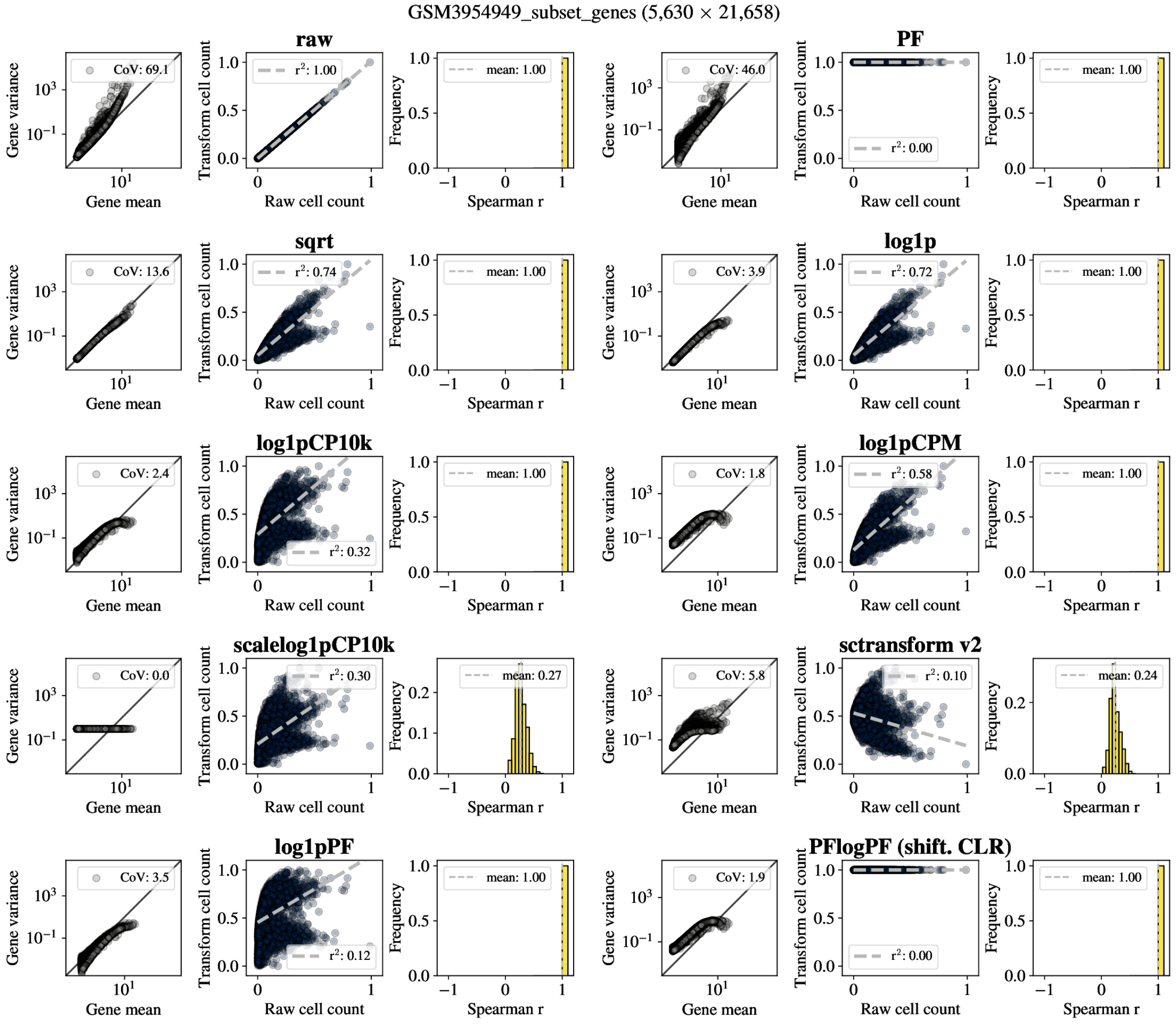


Supplementary Figure 1.444. **GSM3954949**. Per-method comparison of variance stabilization (left), depth normalization (middle), and monotonicity (right) for each normalization on dataset GSM3954949.

| GSM3954950 | |
|---|---|
| cells | 5,289 |
| genes | 60,623 |
| nonzero entries | 2,408,438 |
| density | 0.00751 |
| total counts | 5,582,697 |
| mean counts / cell | 1,056 |
| mean counts / gene | 92 |
| min counts / cell | 147 |
| max counts / cell | 8,670 |
| overdispersion | 24.3 |

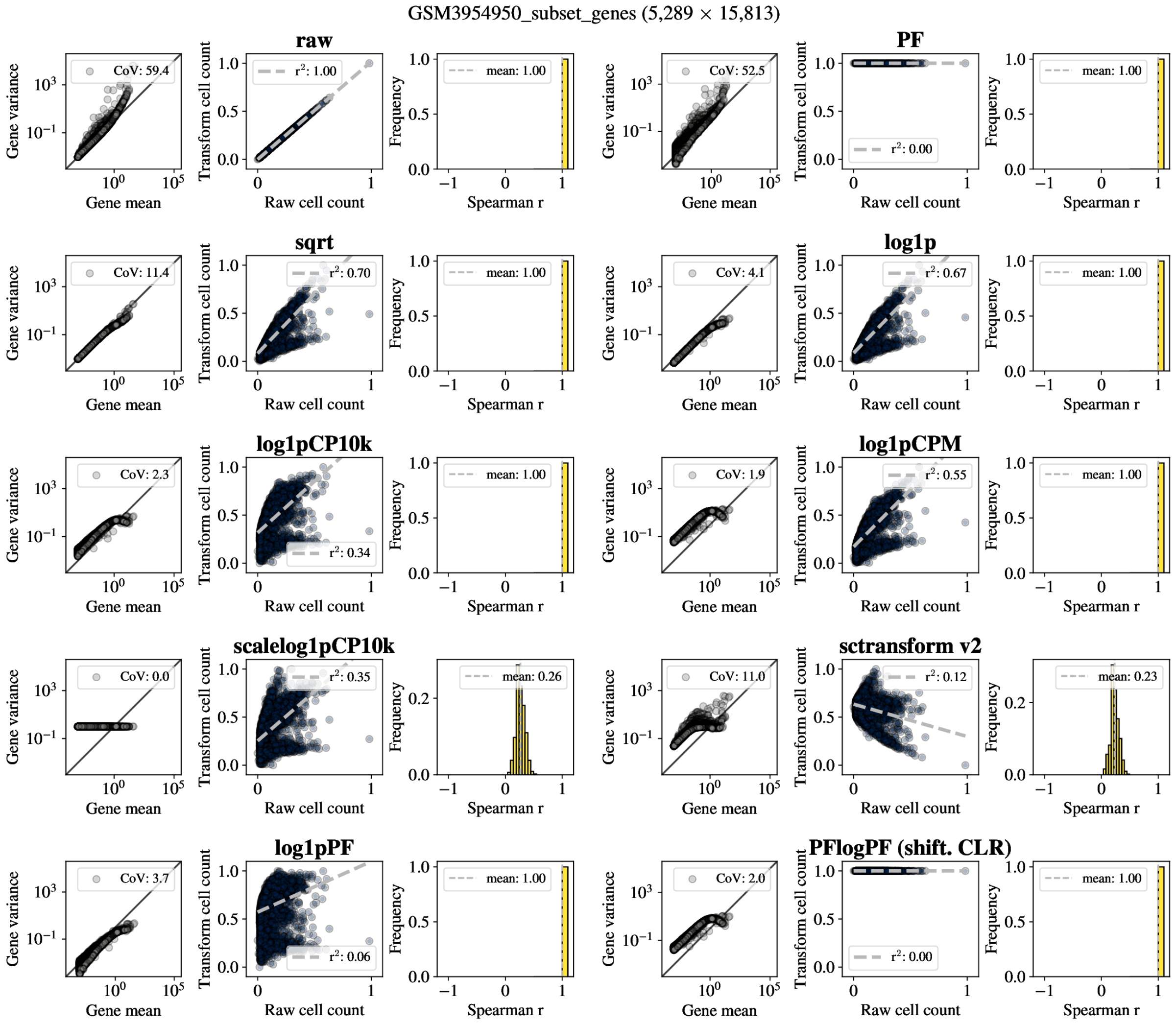


Supplementary Figure 1.445. **GSM3954950**. Per-method comparison of variance stabilization (left), depth normalization (middle), and monotonicity (right) for each normalization on dataset GSM3954950.

| GSM3954952 | |
|---|---|
| cells | 5,523 |
| genes | 60,623 |
| nonzero entries | 3,621,694 |
| density | 0.0108 |
| total counts | 13,440,152 |
| mean counts / cell | 2,433 |
| mean counts / gene | 222 |
| min counts / cell | 129 |
| max counts / cell | 44,048 |
| overdispersion | 7.21 |

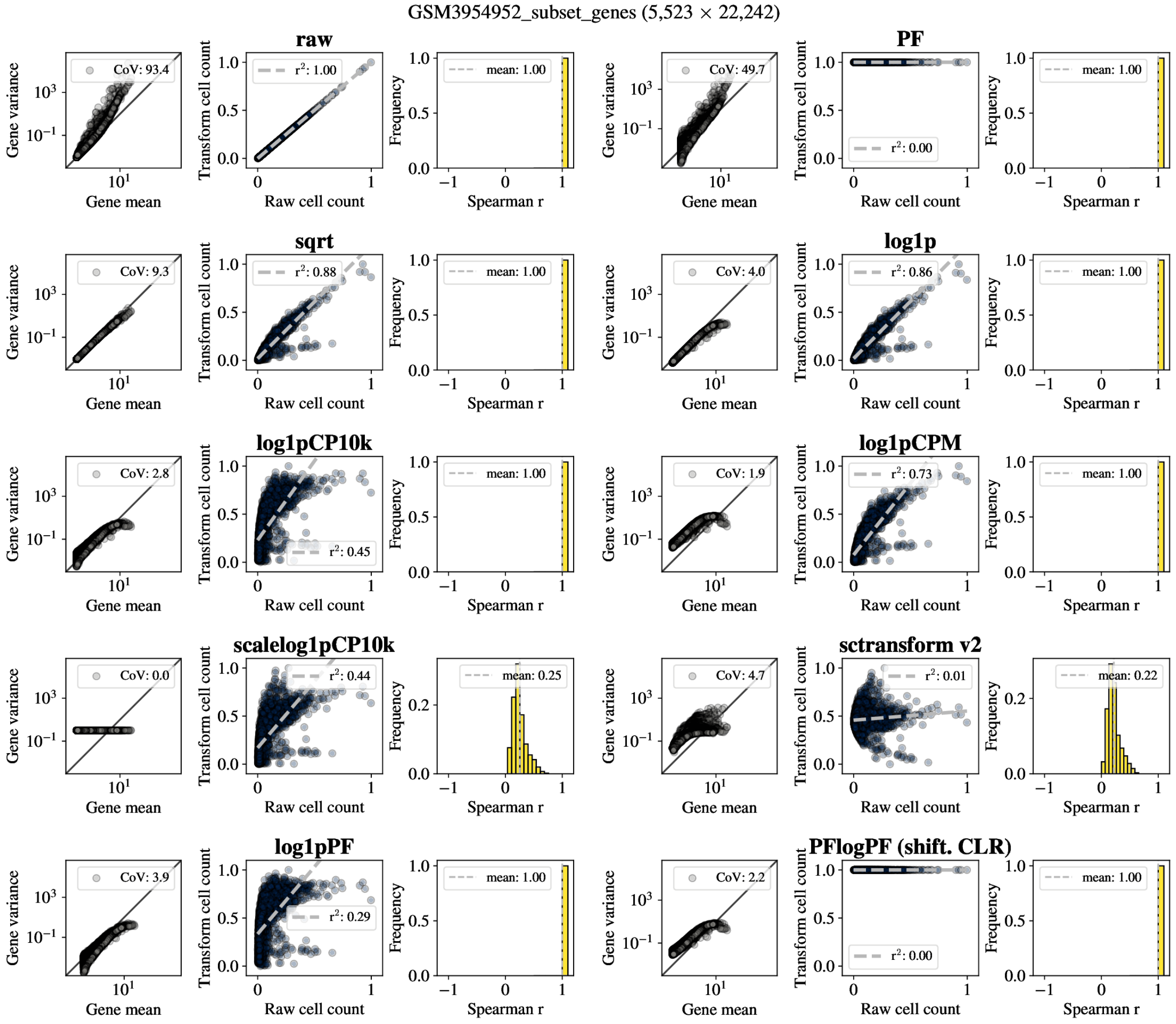


Supplementary Figure 1.446. **GSM3954952**. Per-method comparison of variance stabilization (left), depth normalization (middle), and monotonicity (right) for each normalization on dataset GSM3954952.

| GSM3954953 | |
|---|---|
| cells | 3,125 |
| genes | 60,623 |
| nonzero entries | 2,877,842 |
| density | 0.0152 |
| total counts | 10,442,863 |
| mean counts / cell | 3,342 |
| mean counts / gene | 172 |
| min counts / cell | 327 |
| max counts / cell | 48,174 |
| overdispersion | 34.8 |

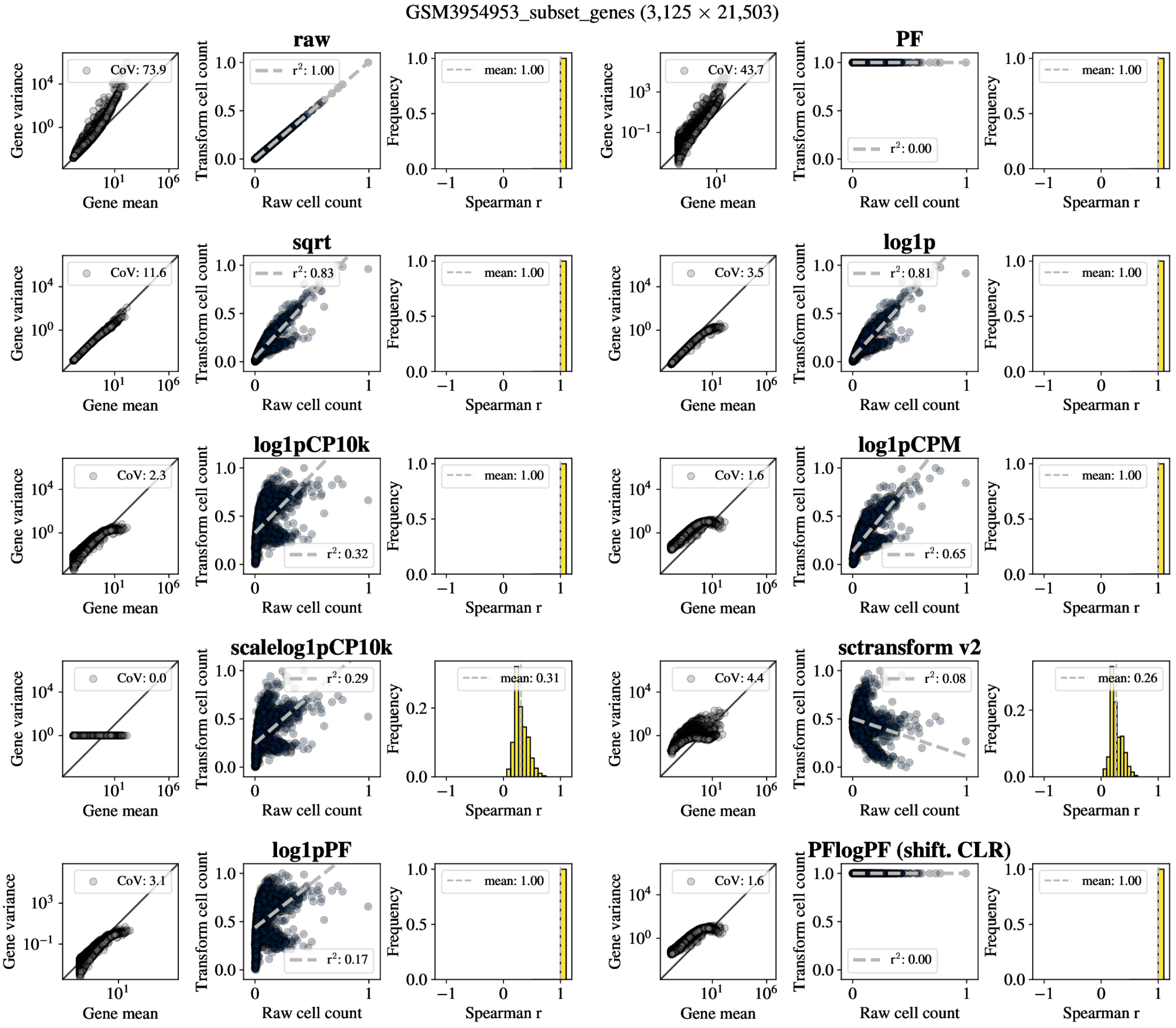


Supplementary Figure 1.447. **GSM3954953**. Per-method comparison of variance stabilization (left), depth normalization (middle), and monotonicity (right) for each normalization on dataset GSM3954953.

| GSM3954954 | |
|---|---|
| cells | 4,409 |
| genes | 60,623 |
| nonzero entries | 6,815,912 |
| density | 0.0255 |
| total counts | 34,801,745 |
| mean counts / cell | 7,893 |
| mean counts / gene | 574 |
| min counts / cell | 456 |
| max counts / cell | 54,050 |
| overdispersion | 21.6 |

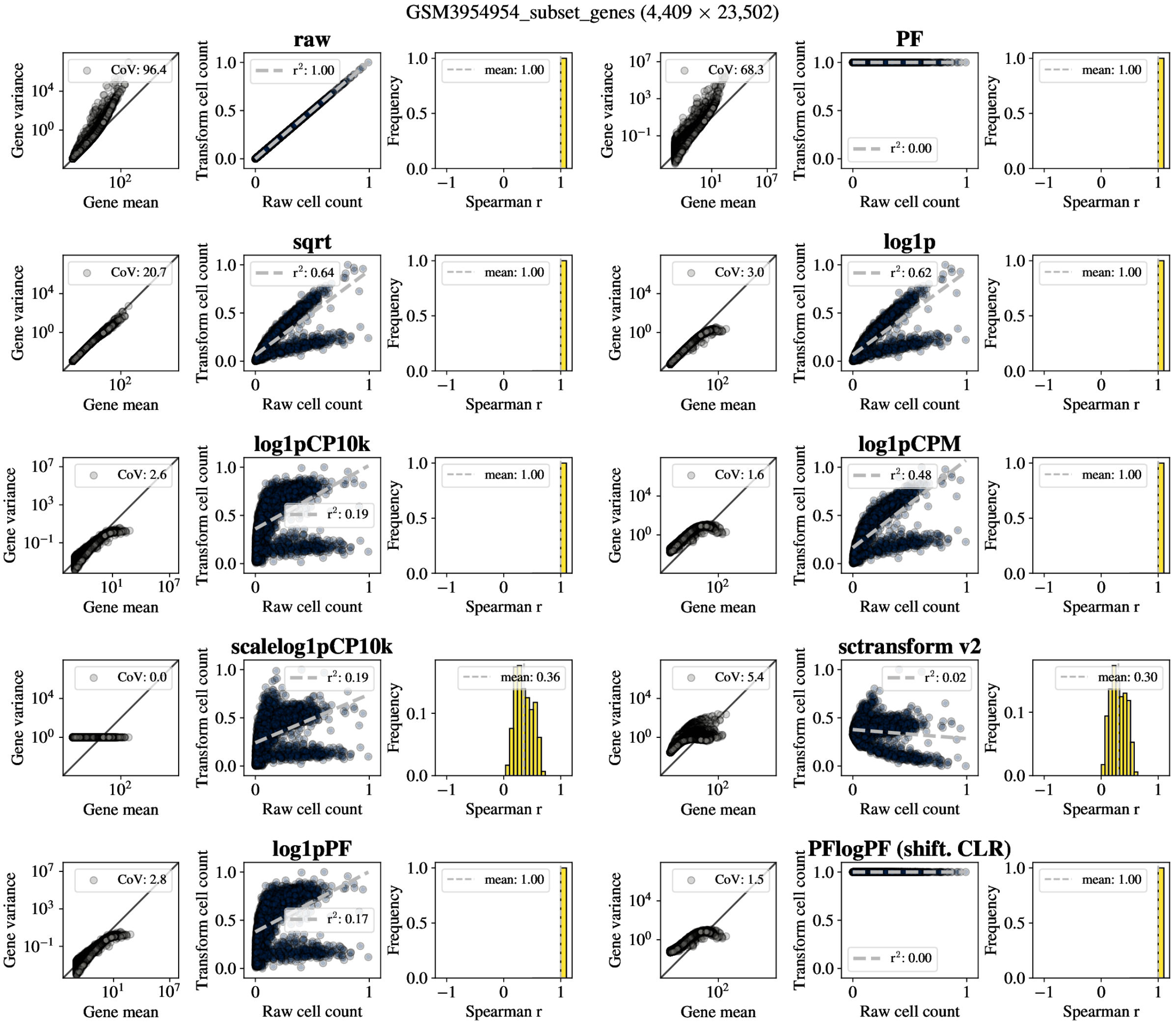


Supplementary Figure 1.448. **GSM3954954**. Per-method comparison of variance stabilization (left), depth normalization (middle), and monotonicity (right) for each normalization on dataset GSM3954954.

| GSM3954955 | |
|---|---|
| cells | 5,101 |
| genes | 60,623 |
| nonzero entries | 7,543,417 |
| density | 0.0244 |
| total counts | 36,038,080 |
| mean counts / cell | 7,065 |
| mean counts / gene | 594 |
| min counts / cell | 429 |
| max counts / cell | 66,298 |
| overdispersion | 11.1 |

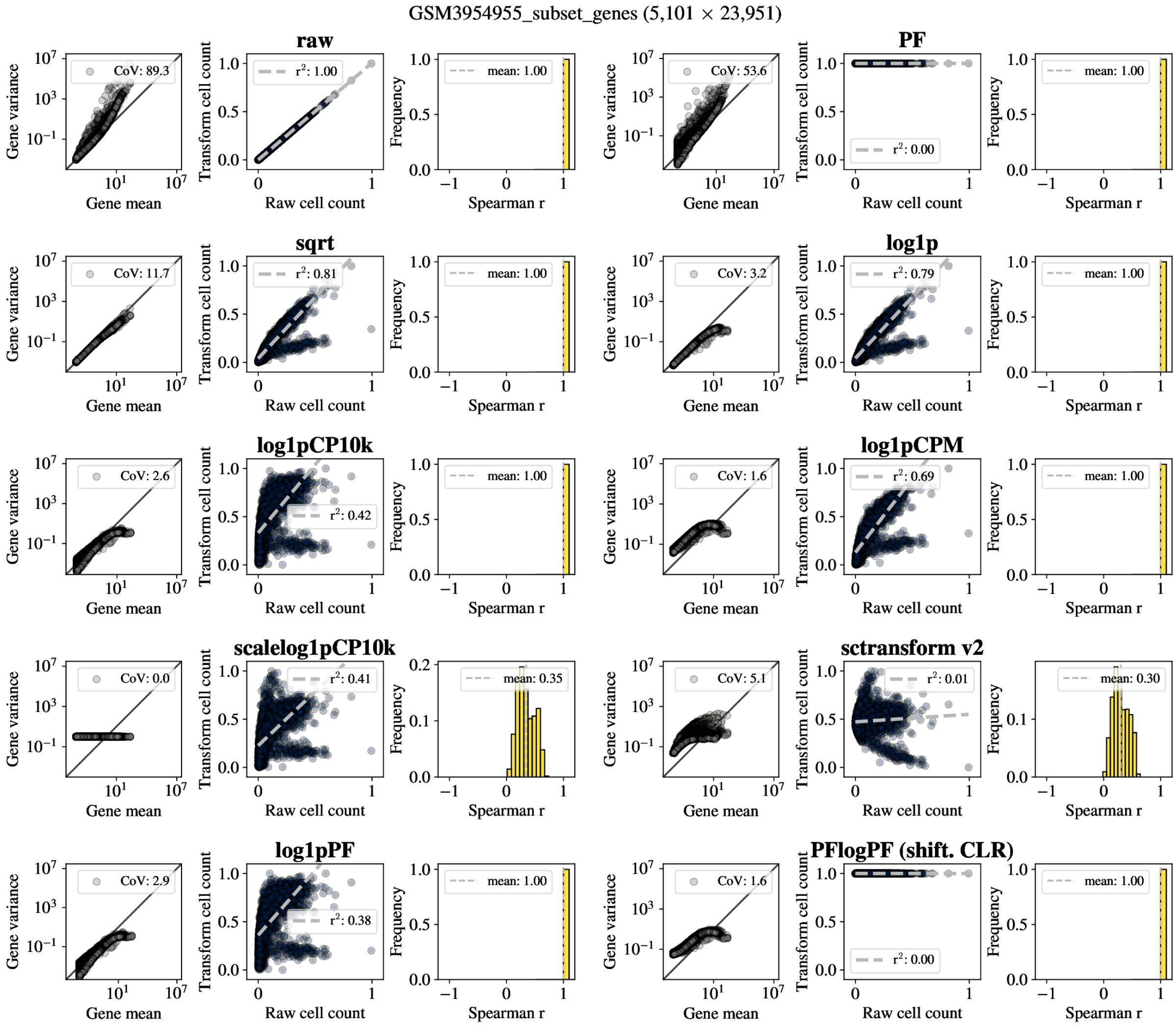


Supplementary Figure 1.449. **GSM3954955**. Per-method comparison of variance stabilization (left), depth normalization (middle), and monotonicity (right) for each normalization on dataset GSM3954955.

| GSM3954956 | |
|---|---|
| cells | 3,839 |
| genes | 60,623 |
| nonzero entries | 5,146,167 |
| density | 0.0221 |
| total counts | 20,396,285 |
| mean counts / cell | 5,313 |
| mean counts / gene | 336 |
| min counts / cell | 477 |
| max counts / cell | 52,931 |
| overdispersion | 1.04 |

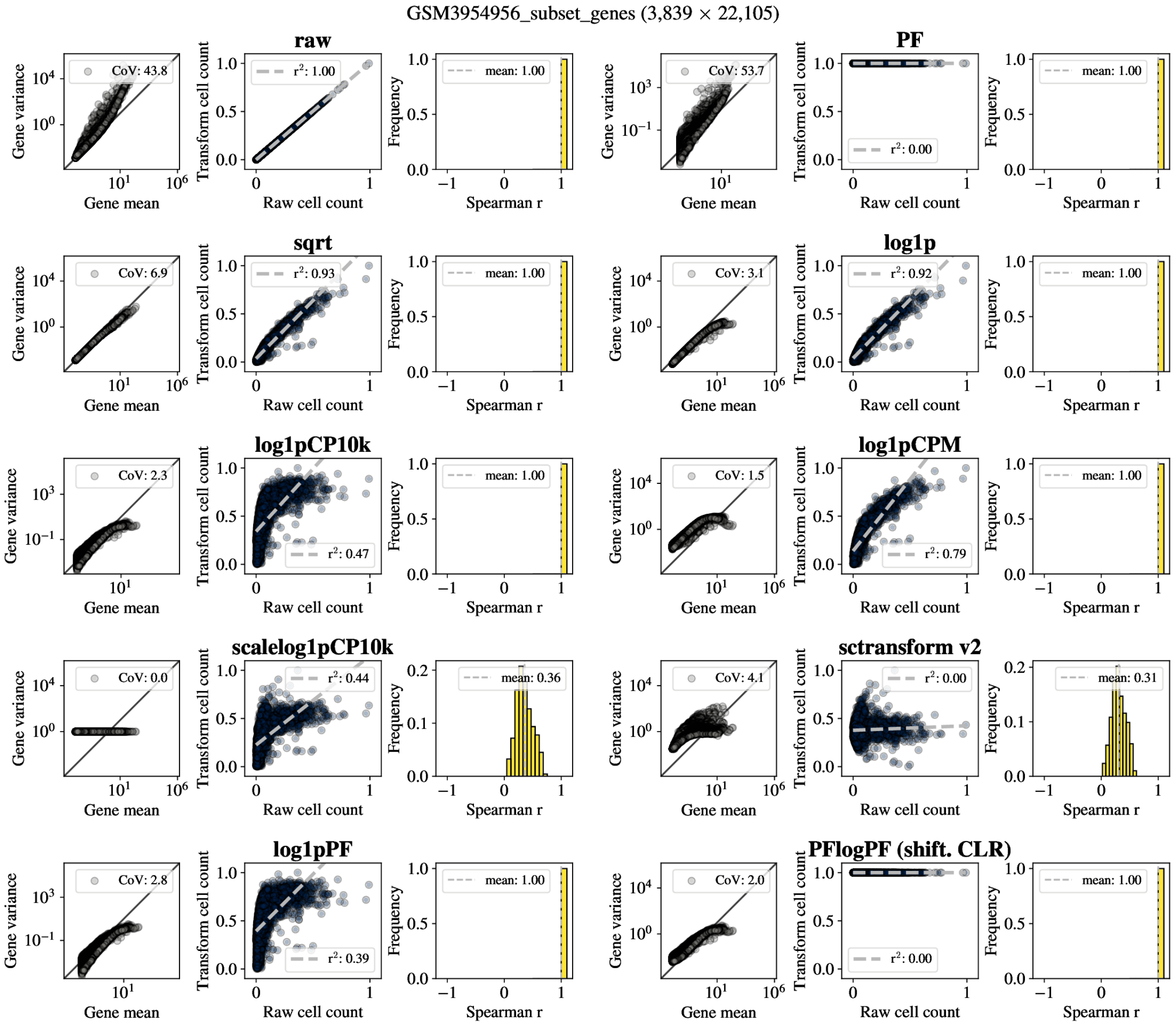


Supplementary Figure 1.450. **GSM3954956**. Per-method comparison of variance stabilization (left), depth normalization (middle), and monotonicity (right) for each normalization on dataset GSM3954956.

| GSM3954957 | |
|---|---|
| cells | 6,377 |
| genes | 60,623 |
| nonzero entries | 7,273,998 |
| density | 0.0188 |
| total counts | 30,868,699 |
| mean counts / cell | 4,841 |
| mean counts / gene | 509 |
| min counts / cell | 407 |
| max counts / cell | 51,225 |
| overdispersion | 6.46 |

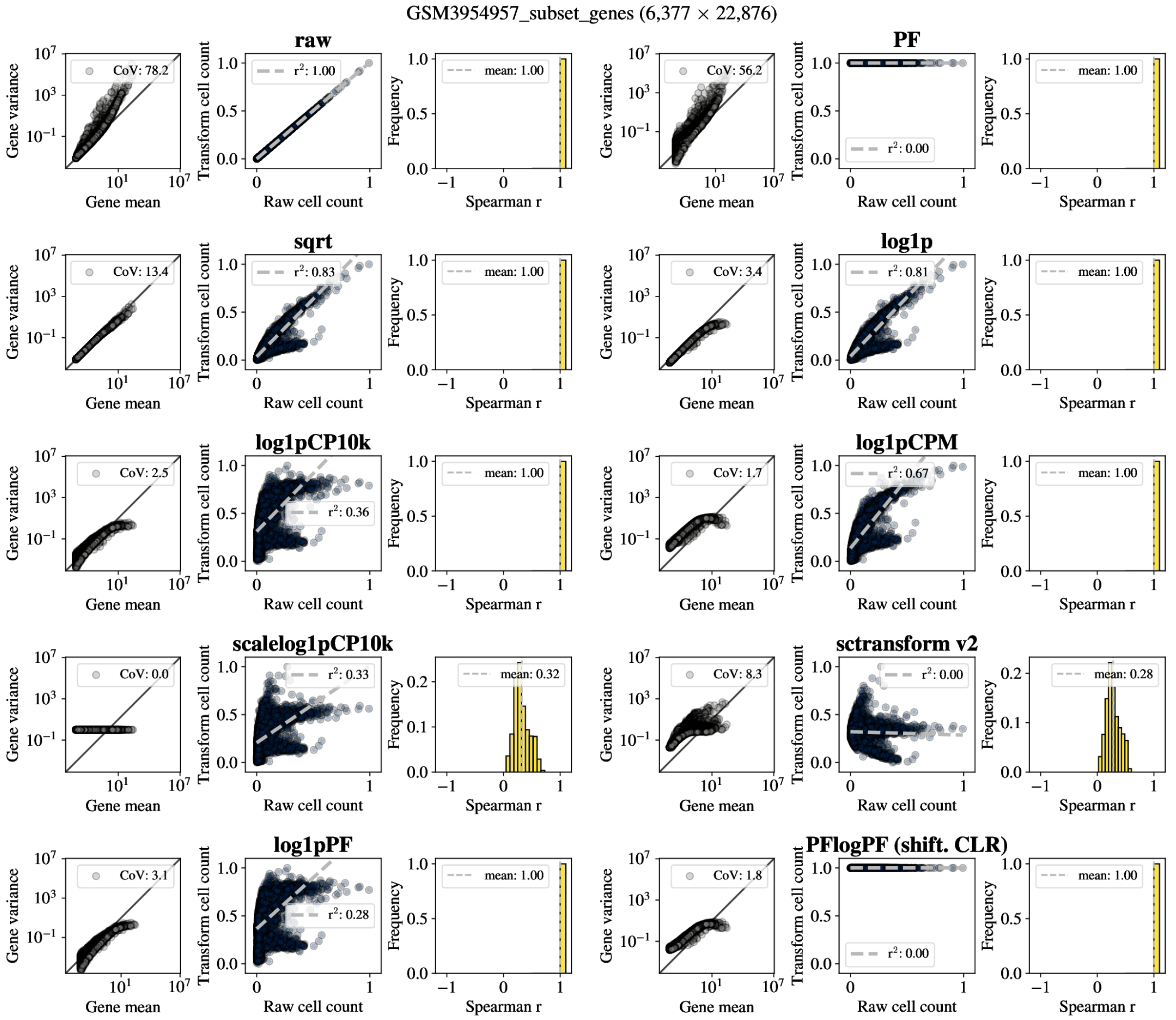


Supplementary Figure 1.451. **GSM3954957**. Per-method comparison of variance stabilization (left), depth normalization (middle), and monotonicity (right) for each normalization on dataset GSM3954957.

| GSM3954958 | |
|---|---|
| cells | 5,067 |
| genes | 60,623 |
| nonzero entries | 8,725,214 |
| density | 0.0284 |
| total counts | 40,995,713 |
| mean counts / cell | 8,091 |
| mean counts / gene | 676 |
| min counts / cell | 538 |
| max counts / cell | 91,971 |
| overdispersion | 1.31 |

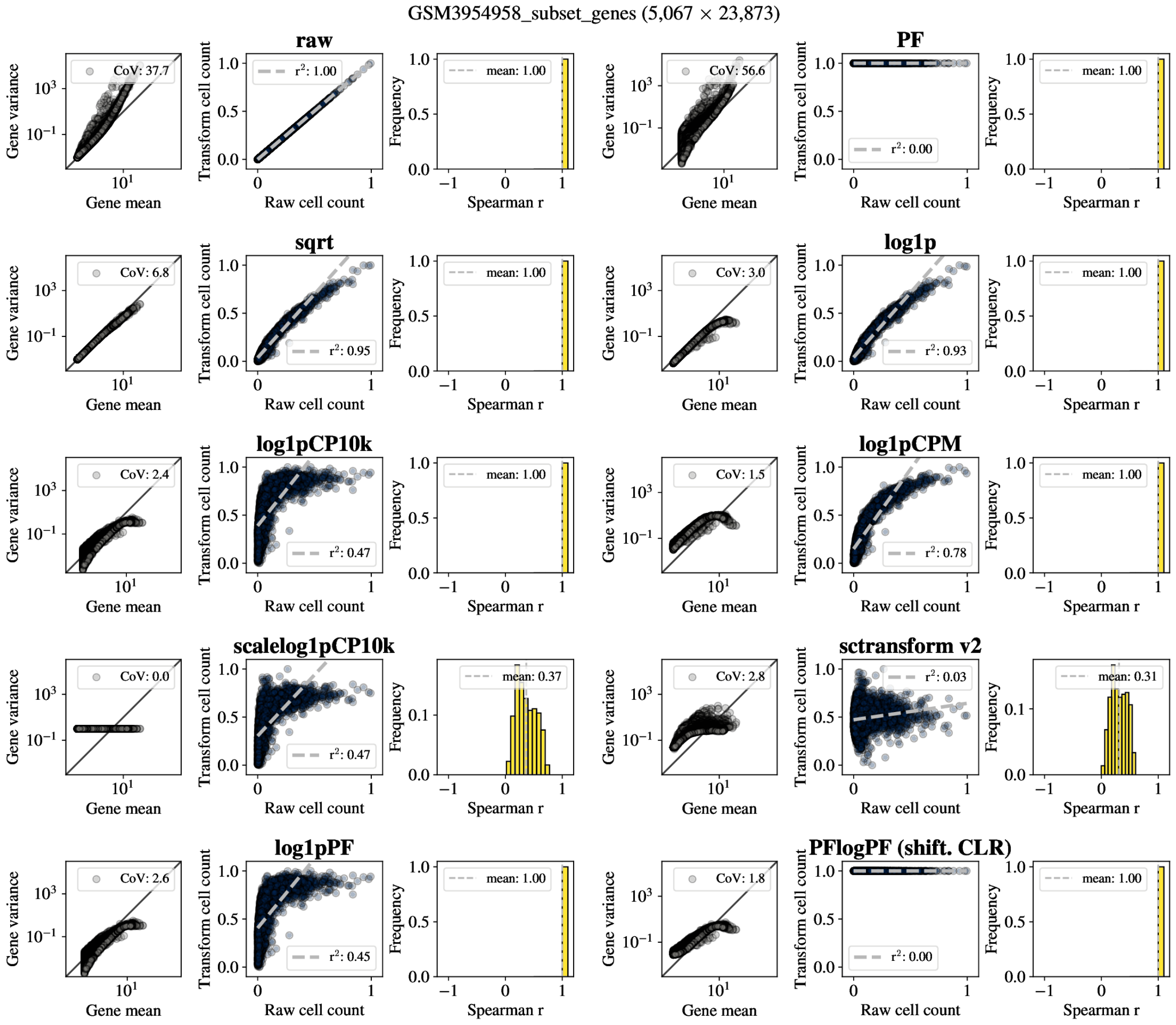


Supplementary Figure 1.452. **GSM3954958**. Per-method comparison of variance stabilization (left), depth normalization (middle), and monotonicity (right) for each normalization on dataset GSM3954958.

| GSM4037301 | |
|---|---|
| cells | 5,535 |
| genes | 60,623 |
| nonzero entries | 2,041,495 |
| density | 0.00608 |
| total counts | 5,081,863 |
| mean counts / cell | 918 |
| mean counts / gene | 84 |
| min counts / cell | 69 |
| max counts / cell | 9,871 |
| overdispersion | 3.77 |

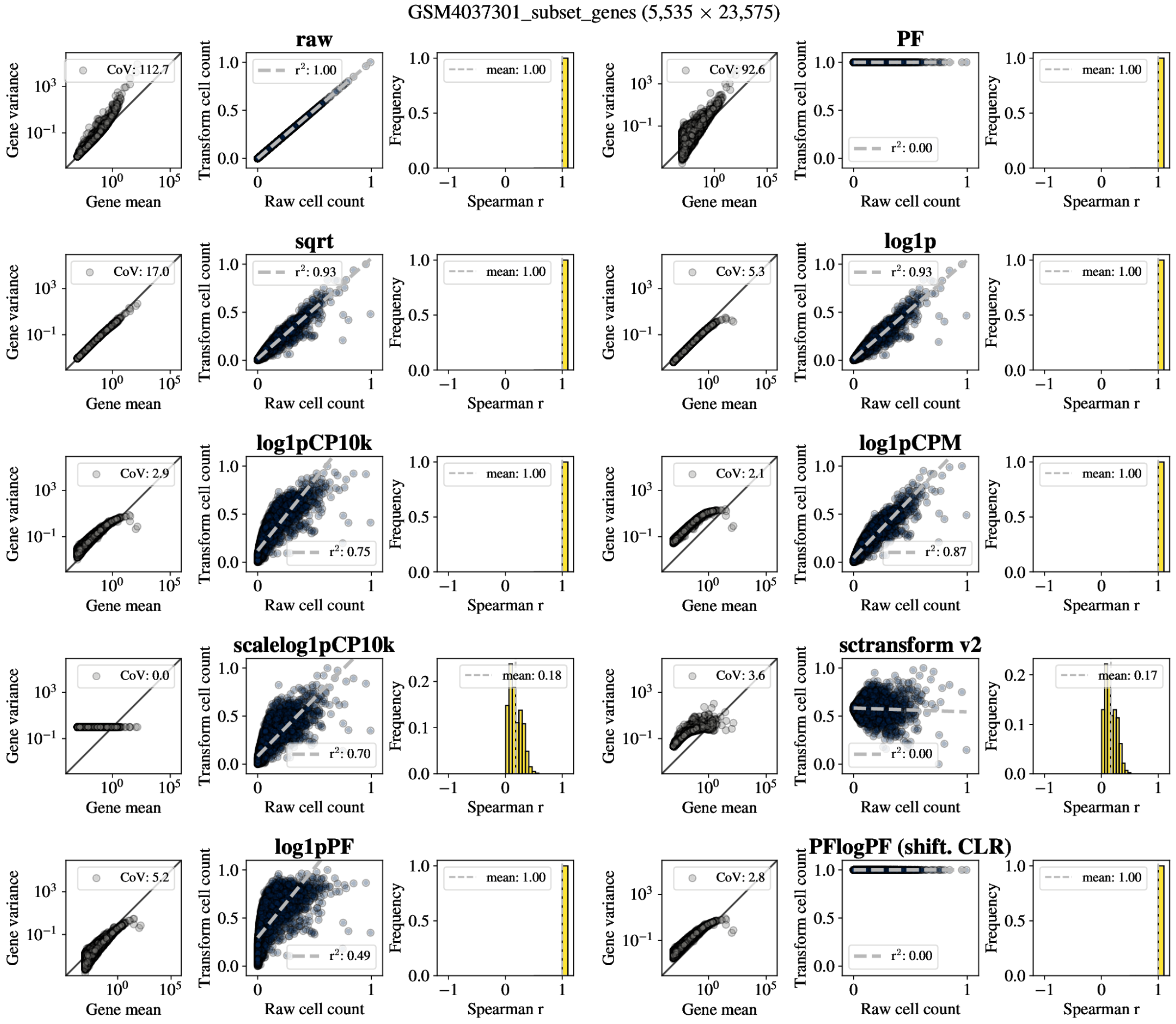


Supplementary Figure 1.453. **GSM4037301**. Per-method comparison of variance stabilization (left), depth normalization (middle), and monotonicity (right) for each normalization on dataset GSM4037301.

| GSM4037302 | |
|---|---|
| cells | 3,994 |
| genes | 60,623 |
| nonzero entries | 1,613,447 |
| density | 0.00666 |
| total counts | 3,312,677 |
| mean counts / cell | 829 |
| mean counts / gene | 55 |
| min counts / cell | 171 |
| max counts / cell | 7,257 |
| overdispersion | 1.82 |

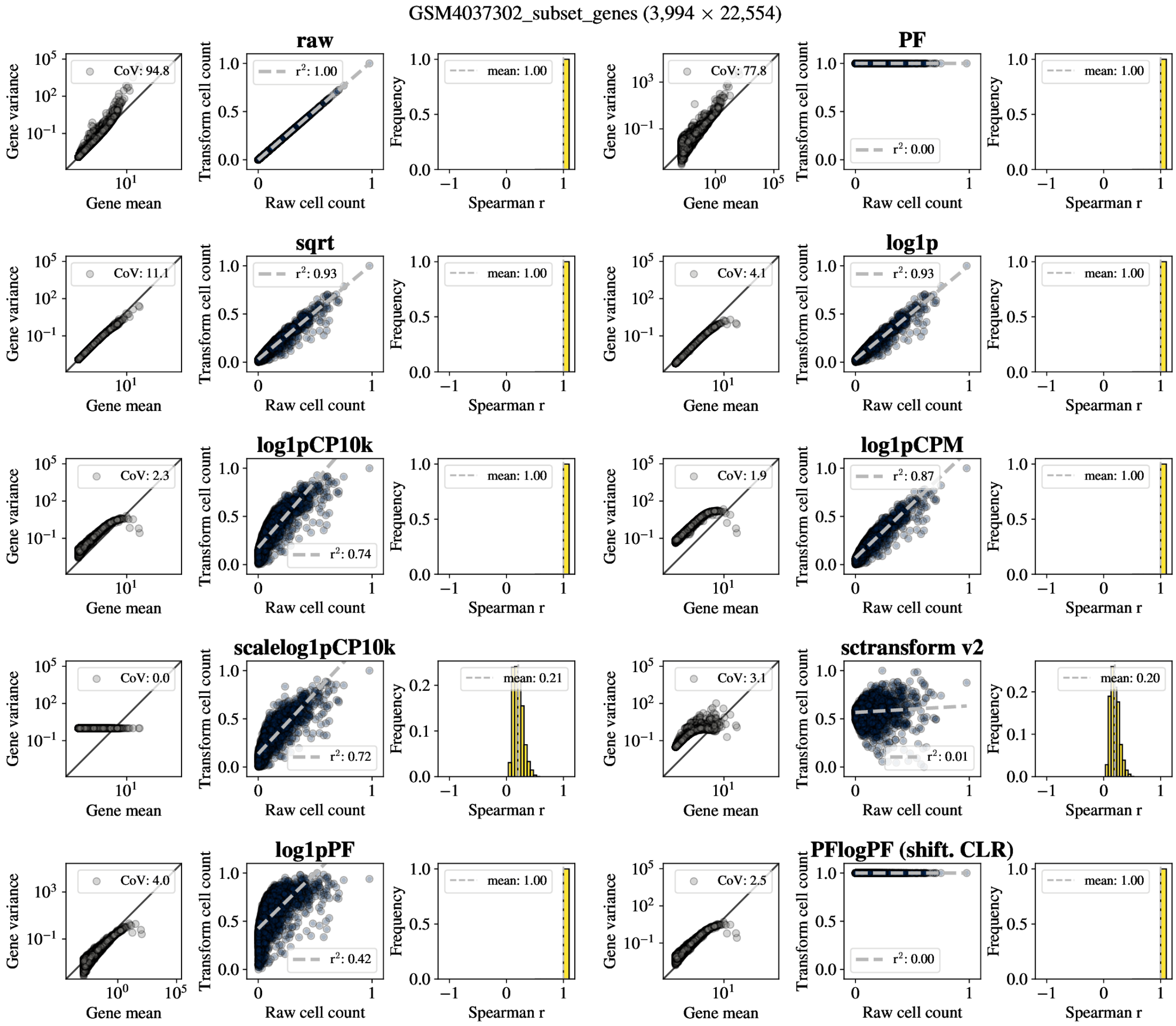


Supplementary Figure 1.454. **GSM4037302**. Per-method comparison of variance stabilization (left), depth normalization (middle), and monotonicity (right) for each normalization on dataset GSM4037302.

| GSM4037305 | |
|---|---|
| cells | 4,801 |
| genes | 60,623 |
| nonzero entries | 2,057,633 |
| density | 0.00707 |
| total counts | 4,774,351 |
| mean counts / cell | 994 |
| mean counts / gene | 79 |
| min counts / cell | 459 |
| max counts / cell | 4,653 |
| overdispersion | 1 |

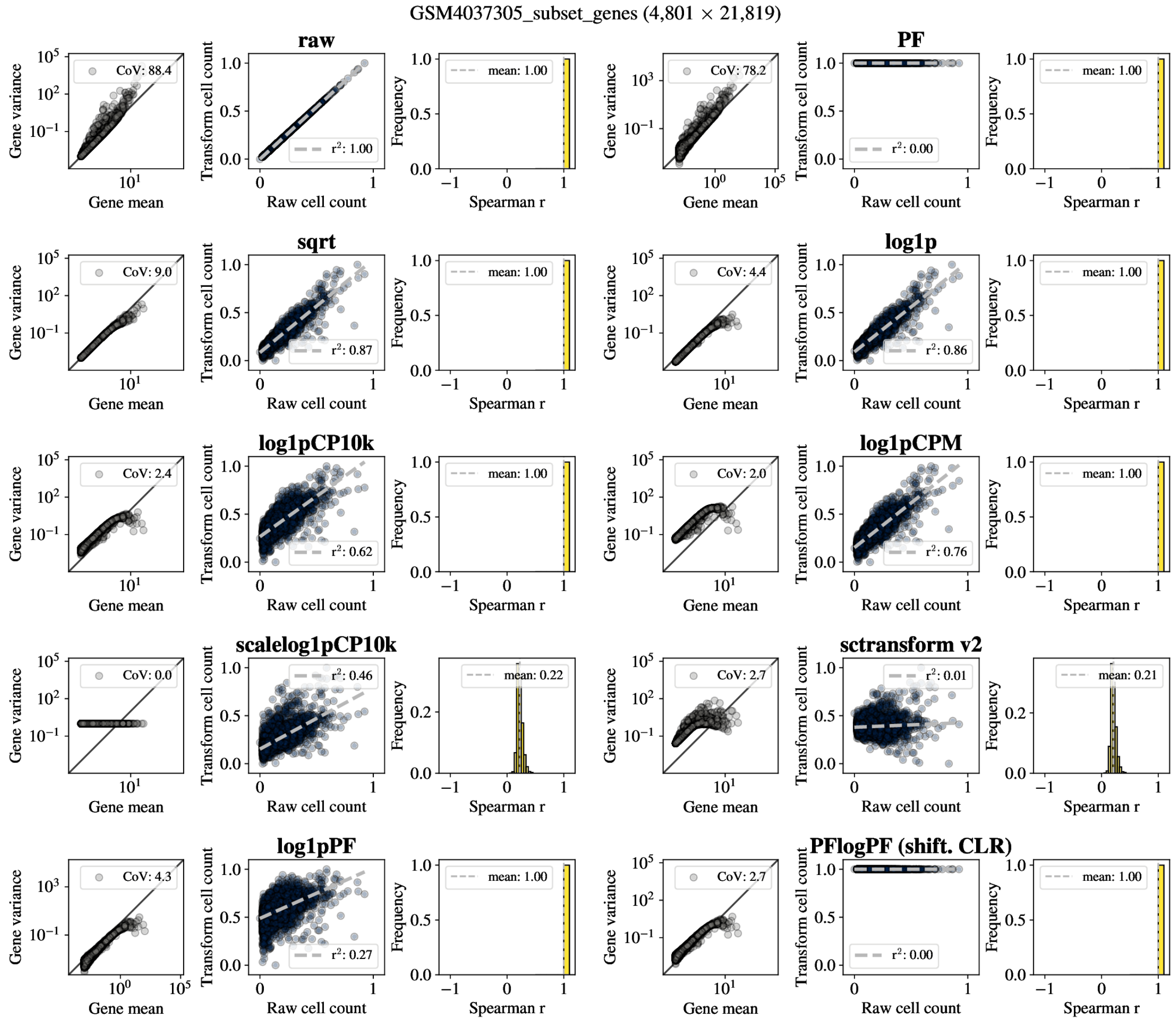


Supplementary Figure 1.455. **GSM4037305**. Per-method comparison of variance stabilization (left), depth normalization (middle), and monotonicity (right) for each normalization on dataset GSM4037305.

| GSM4037306 | |
|---|---|
| cells | 3,158 |
| genes | 60,623 |
| nonzero entries | 1,302,479 |
| density | 0.0068 |
| total counts | 2,804,110 |
| mean counts / cell | 888 |
| mean counts / gene | 46 |
| min counts / cell | 514 |
| max counts / cell | 3,739 |
| overdispersion | 1.71 |

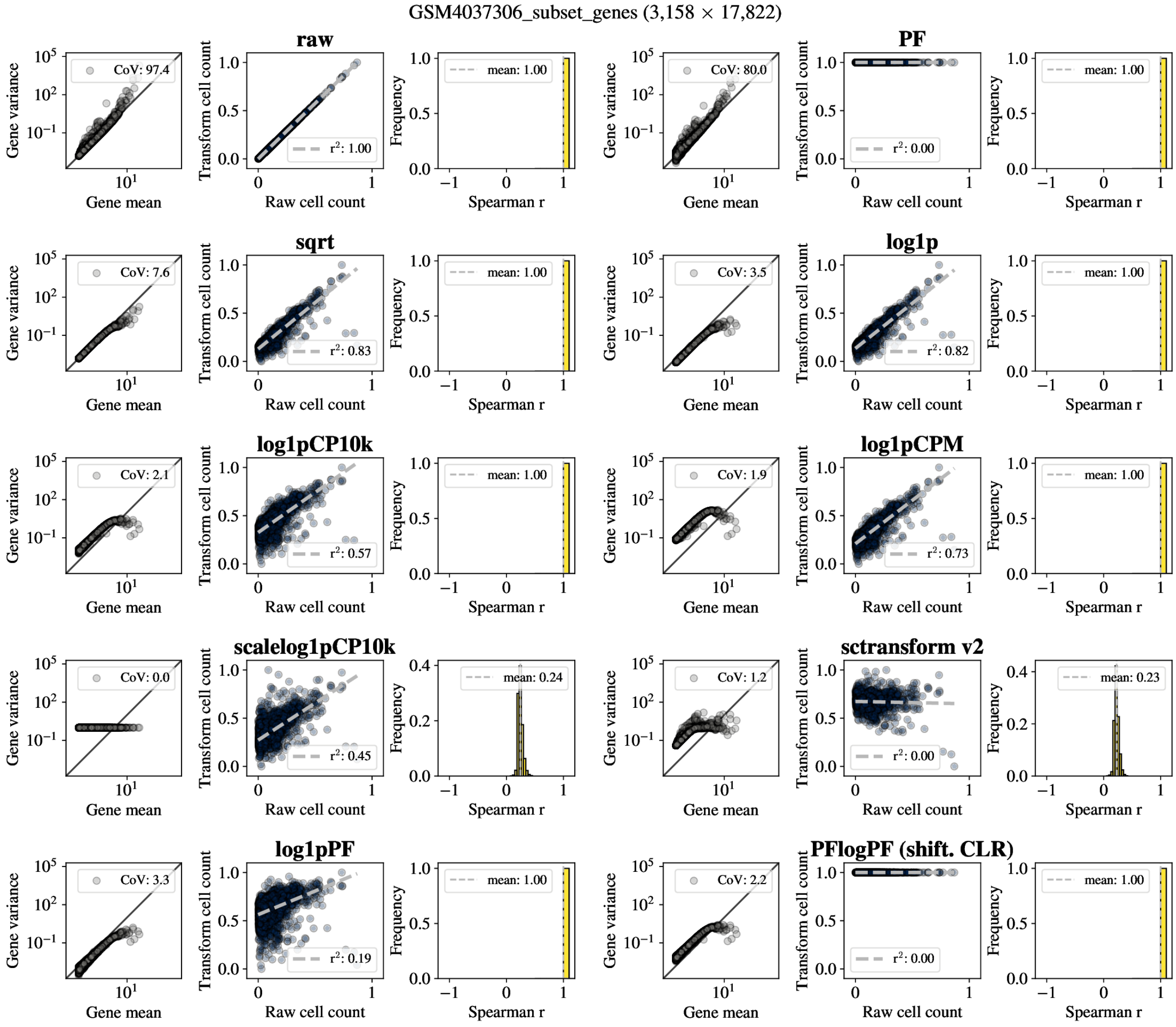


Supplementary Figure 1.456. **GSM4037306**. Per-method comparison of variance stabilization (left), depth normalization (middle), and monotonicity (right) for each normalization on dataset GSM4037306.

| GSM4037309 | |
|---|---|
| cells | 1,998 |
| genes | 60,623 |
| nonzero entries | 1,884,510 |
| density | 0.0156 |
| total counts | 4,103,093 |
| mean counts / cell | 2,054 |
| mean counts / gene | 68 |
| min counts / cell | 211 |
| max counts / cell | 20,570 |
| overdispersion | 3.08 |

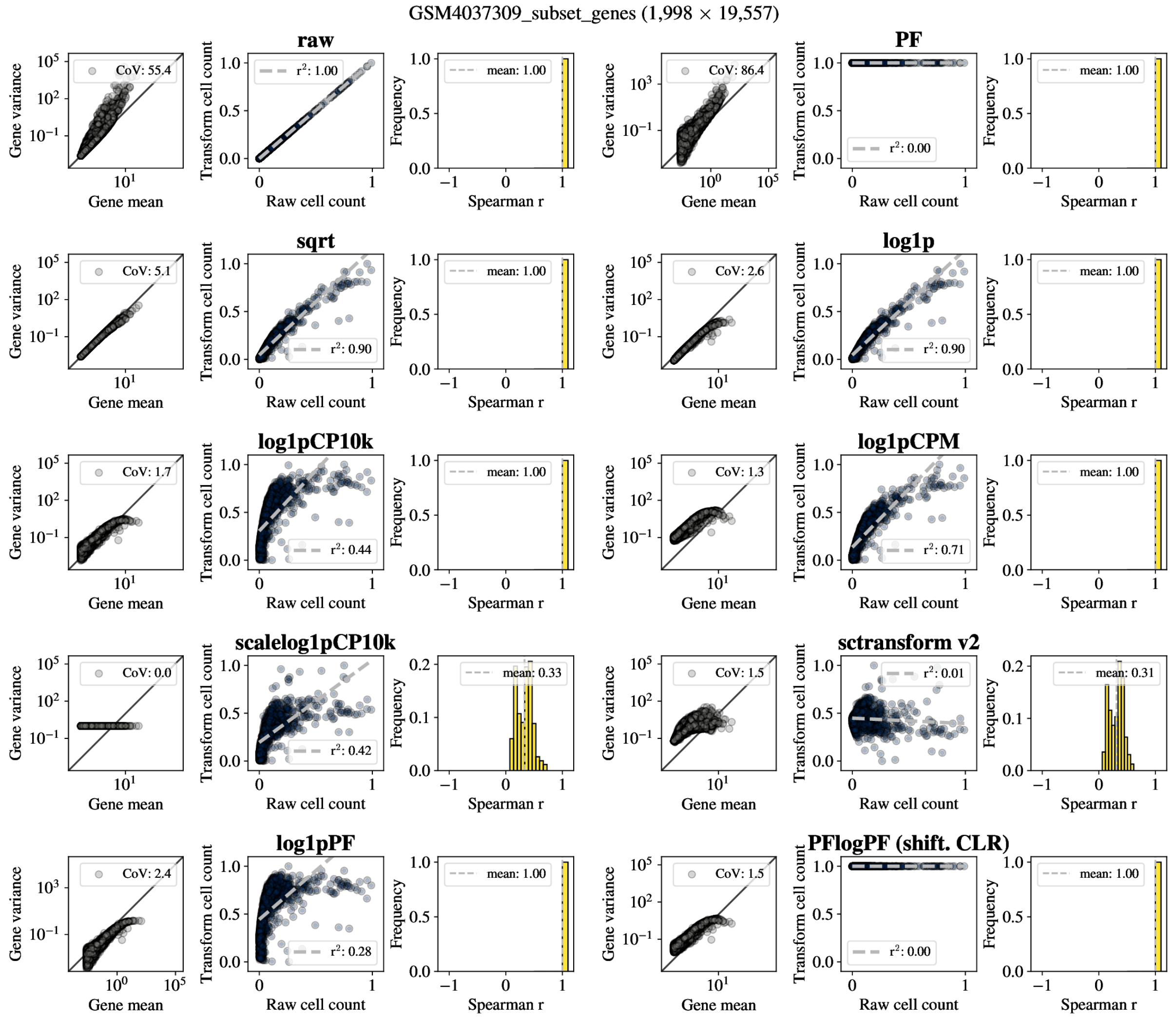


Supplementary Figure 1.457. **GSM4037309**. Per-method comparison of variance stabilization (left), depth normalization (middle), and monotonicity (right) for each normalization on dataset GSM4037309.

| GSM4037313 | |
|---|---|
| cells | 2,305 |
| genes | 60,623 |
| nonzero entries | 1,112,365 |
| density | 0.00796 |
| total counts | 2,229,582 |
| mean counts / cell | 967 |
| mean counts / gene | 37 |
| min counts / cell | 4 |
| max counts / cell | 4,645 |
| overdispersion | 1.7 |

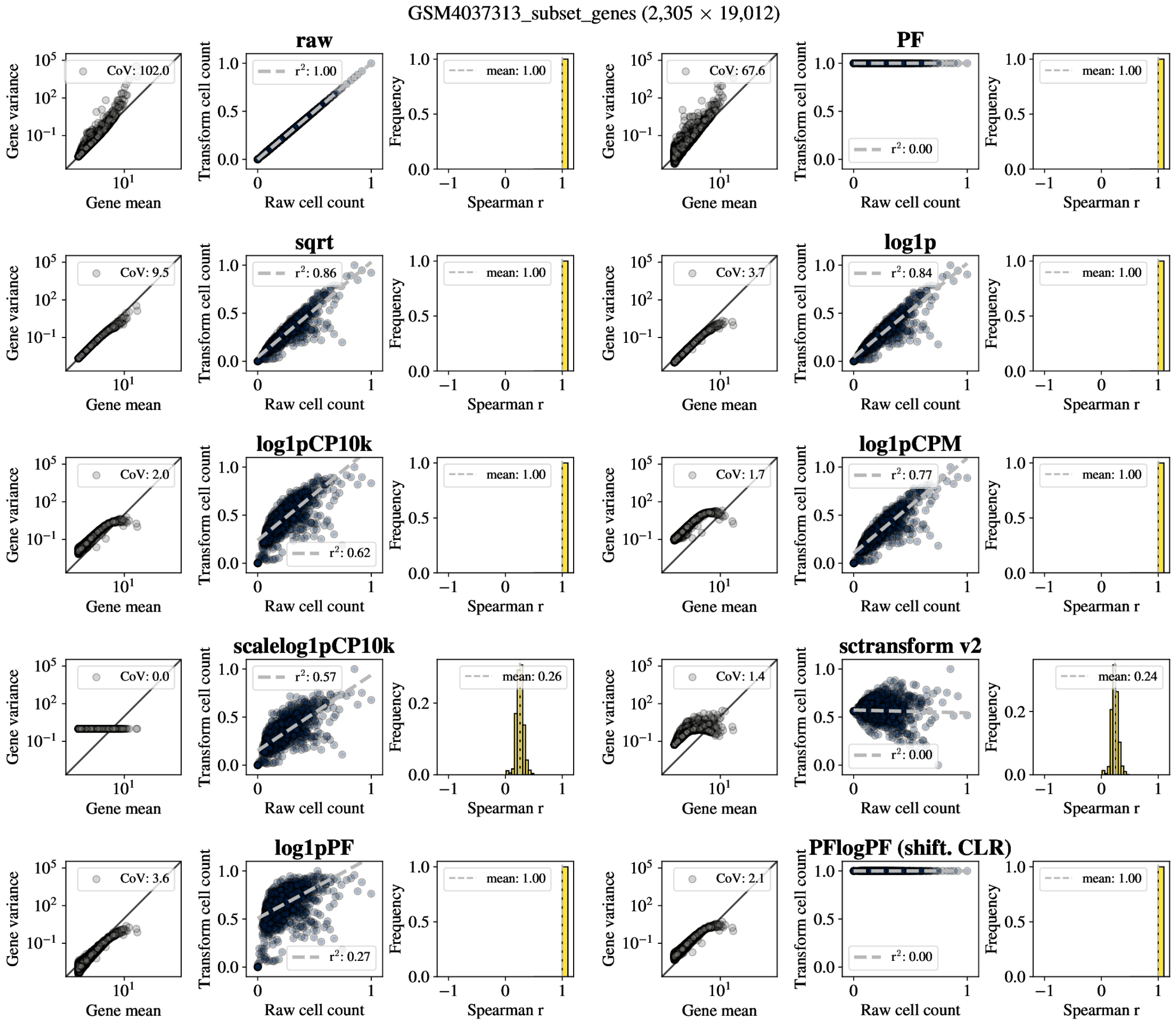


Supplementary Figure 1.458. **GSM4037313**. Per-method comparison of variance stabilization (left), depth normalization (middle), and monotonicity (right) for each normalization on dataset GSM4037313.

| GSM4037316 | |
|---|---|
| cells | 982 |
| genes | 60,623 |
| nonzero entries | 785,847 |
| density | 0.0132 |
| total counts | 2,051,378 |
| mean counts / cell | 2,089 |
| mean counts / gene | 34 |
| min counts / cell | 757 |
| max counts / cell | 12,082 |
| overdispersion | 2.31 |

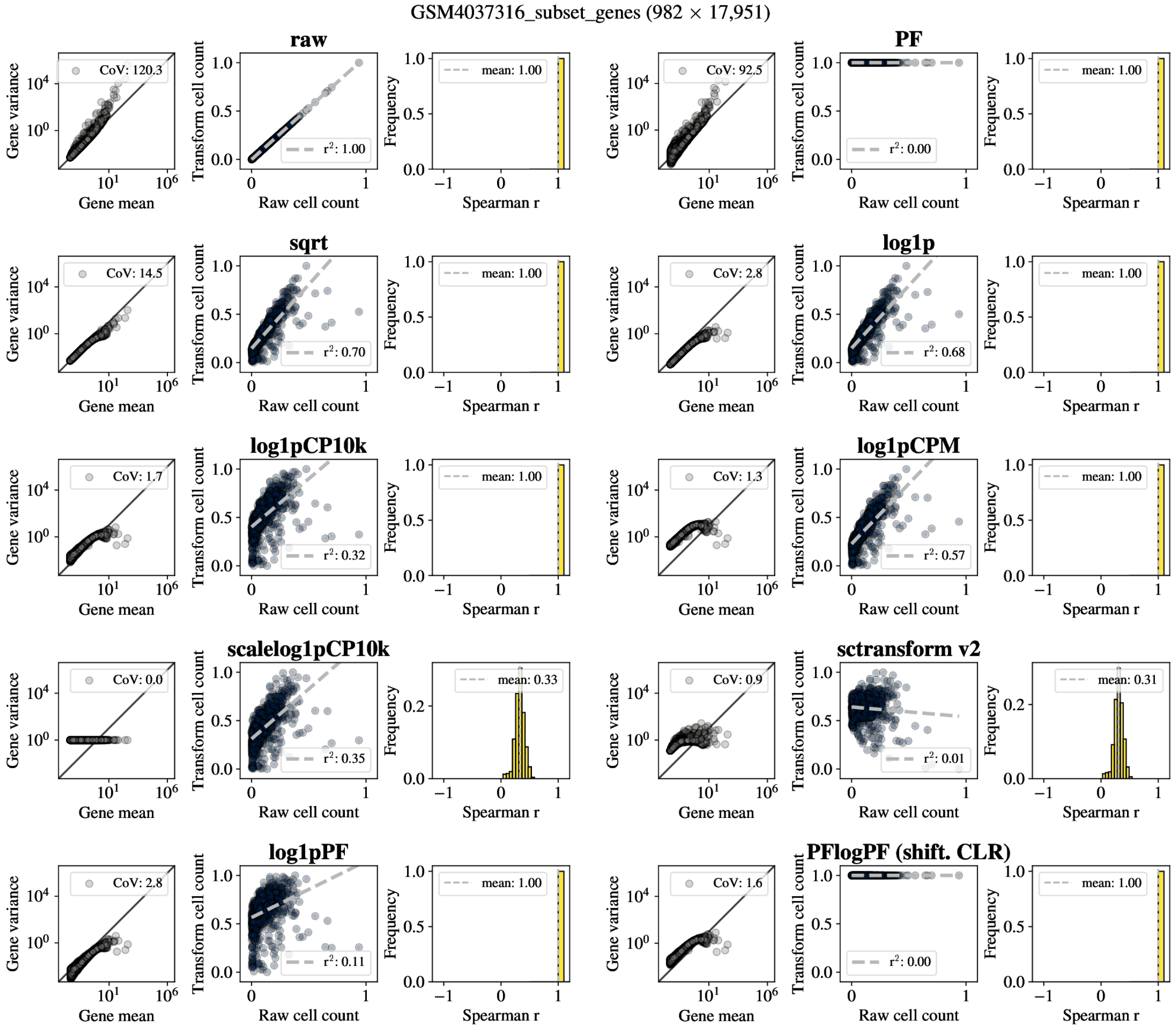


Supplementary Figure 1.459. **GSM4037316**. Per-method comparison of variance stabilization (left), depth normalization (middle), and monotonicity (right) for each normalization on dataset GSM4037316.

| GSM4037323 | |
|---|---|
| cells | 7,294 |
| genes | 60,623 |
| nonzero entries | 496,019 |
| density | 0.00112 |
| total counts | 810,767 |
| mean counts / cell | 111 |
| mean counts / gene | 13 |
| min counts / cell | 0 |
| max counts / cell | 740 |
| overdispersion | 1.13 |

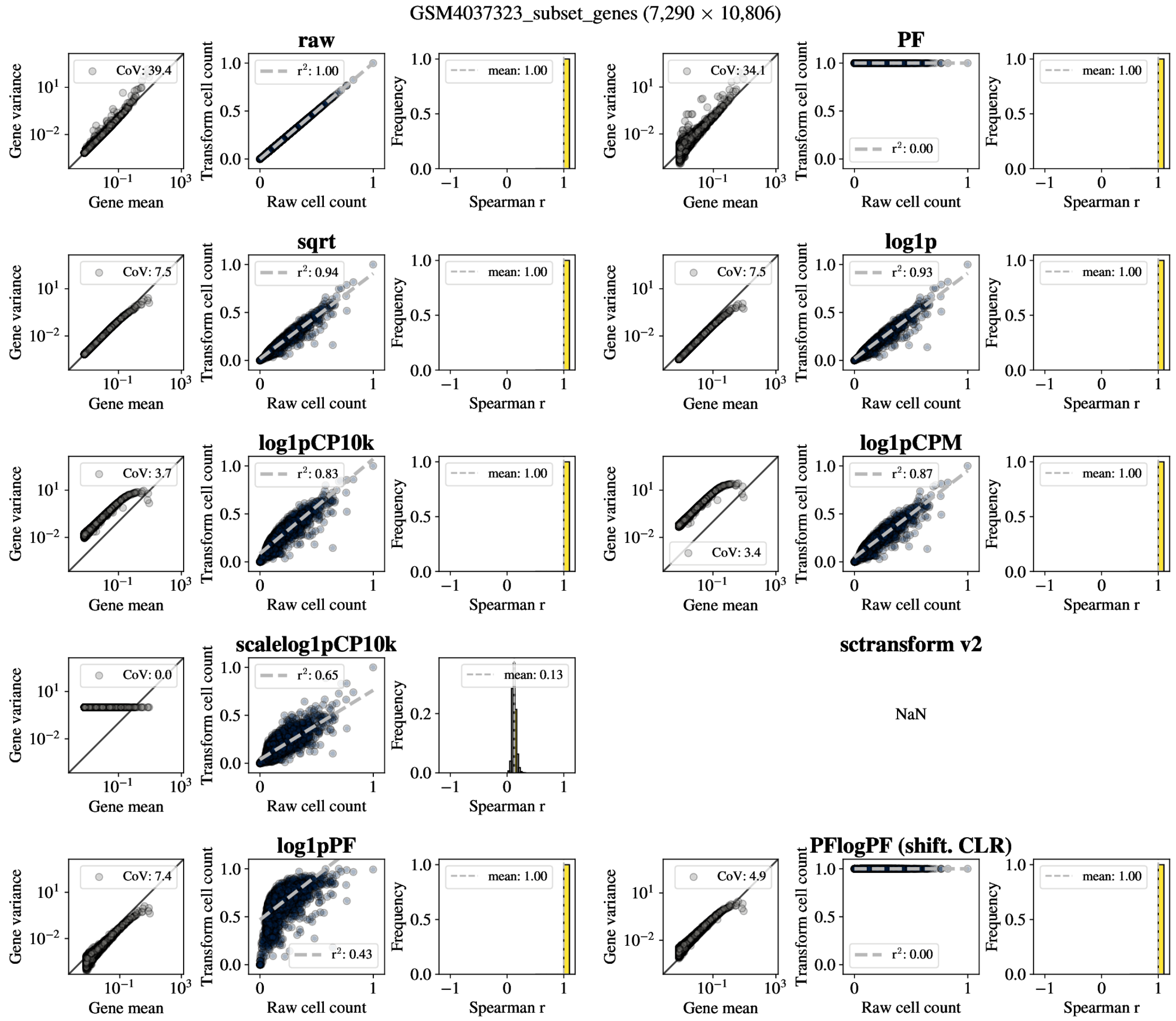


Supplementary Figure 1.460. **GSM4037323**. Per-method comparison of variance stabilization (left), depth normalization (middle), and monotonicity (right) for each normalization on dataset GSM4037323.

| GSM4037324 | |
|---|---|
| cells | 11,939 |
| genes | 60,623 |
| nonzero entries | 8,453,887 |
| density | 0.0117 |
| total counts | 26,006,509 |
| mean counts / cell | 2,178 |
| mean counts / gene | 429 |
| min counts / cell | 100 |
| max counts / cell | 32,746 |
| overdispersion | 10.1 |

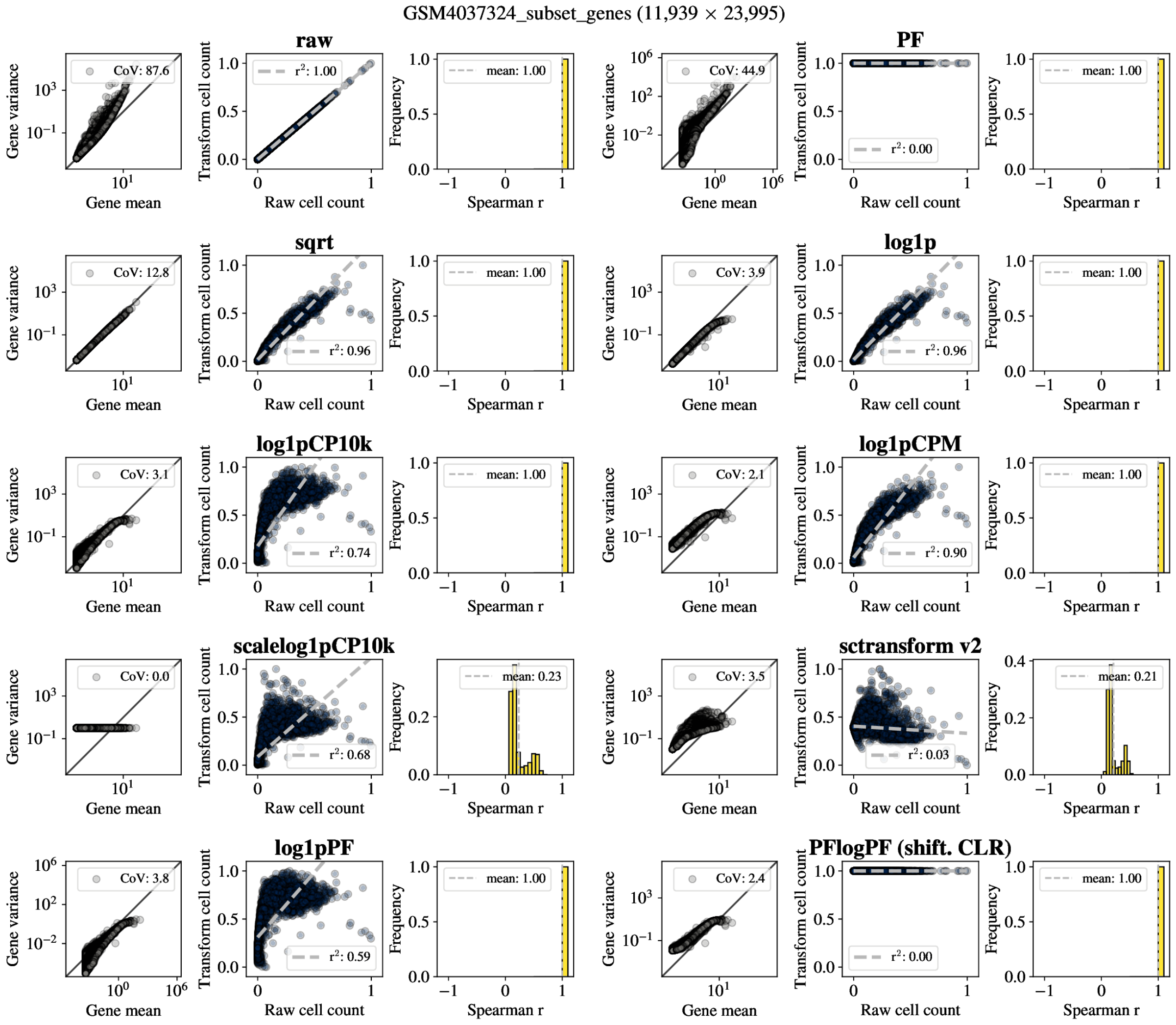


Supplementary Figure 1.461. **GSM4037324**. Per-method comparison of variance stabilization (left), depth normalization (middle), and monotonicity (right) for each normalization on dataset GSM4037324.

| GSM4037325 | |
|---|---|
| cells | 4,297 |
| genes | 60,623 |
| nonzero entries | 8,380,700 |
| density | 0.0322 |
| total counts | 23,281,884 |
| mean counts / cell | 5,418 |
| mean counts / gene | 384 |
| min counts / cell | 1,364 |
| max counts / cell | 52,044 |
| overdispersion | 1.56 |

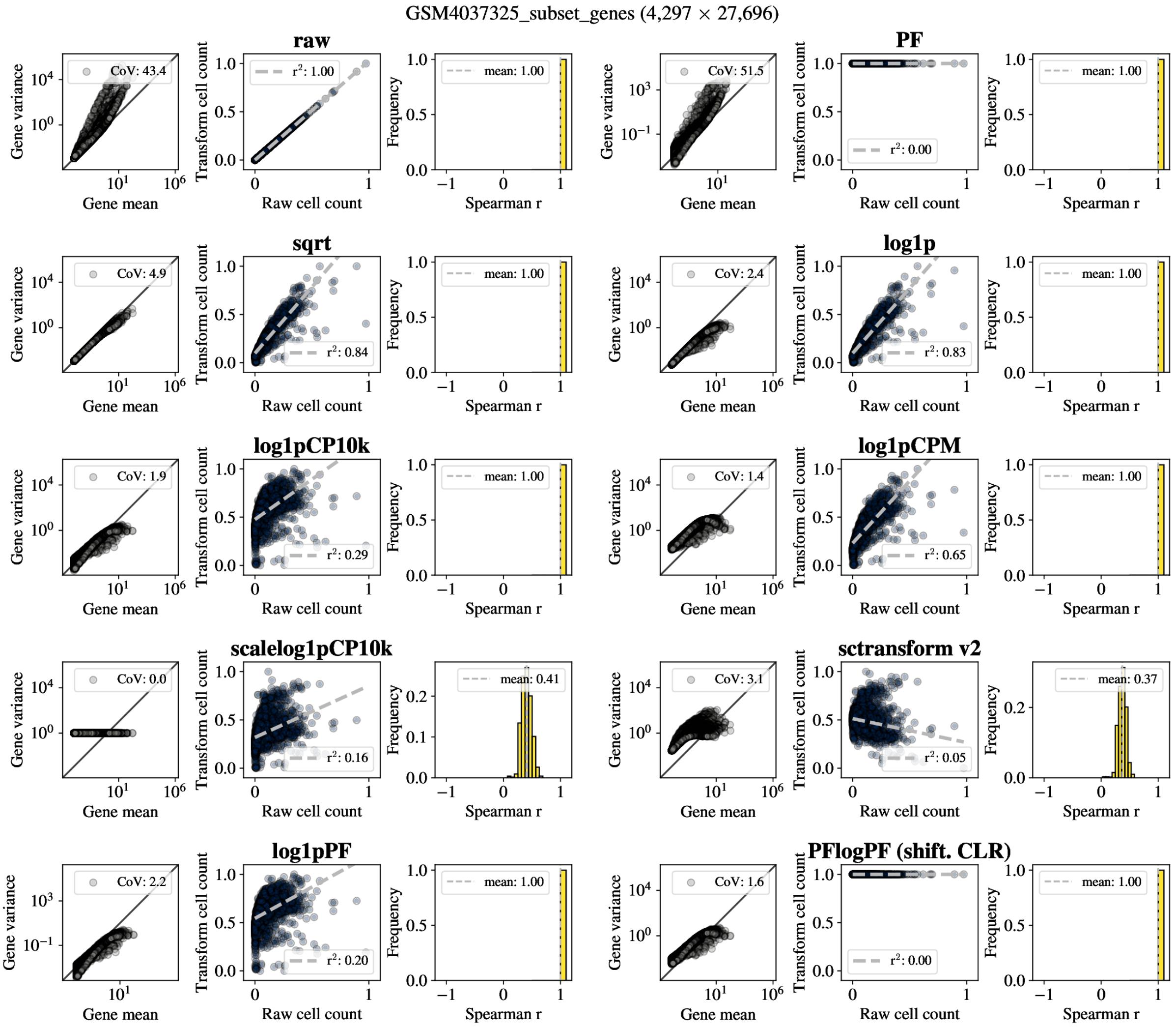


Supplementary Figure 1.462. **GSM4037325**. Per-method comparison of variance stabilization (left), depth normalization (middle), and monotonicity (right) for each normalization on dataset GSM4037325.

| GSM4037981 | |
|---|---|
| cells | 857 |
| genes | 60,623 |
| nonzero entries | 1,627,341 |
| density | 0.0313 |
| total counts | 6,873,973 |
| mean counts / cell | 8,021 |
| mean counts / gene | 113 |
| min counts / cell | 575 |
| max counts / cell | 184,498 |
| overdispersion | 2.93 |

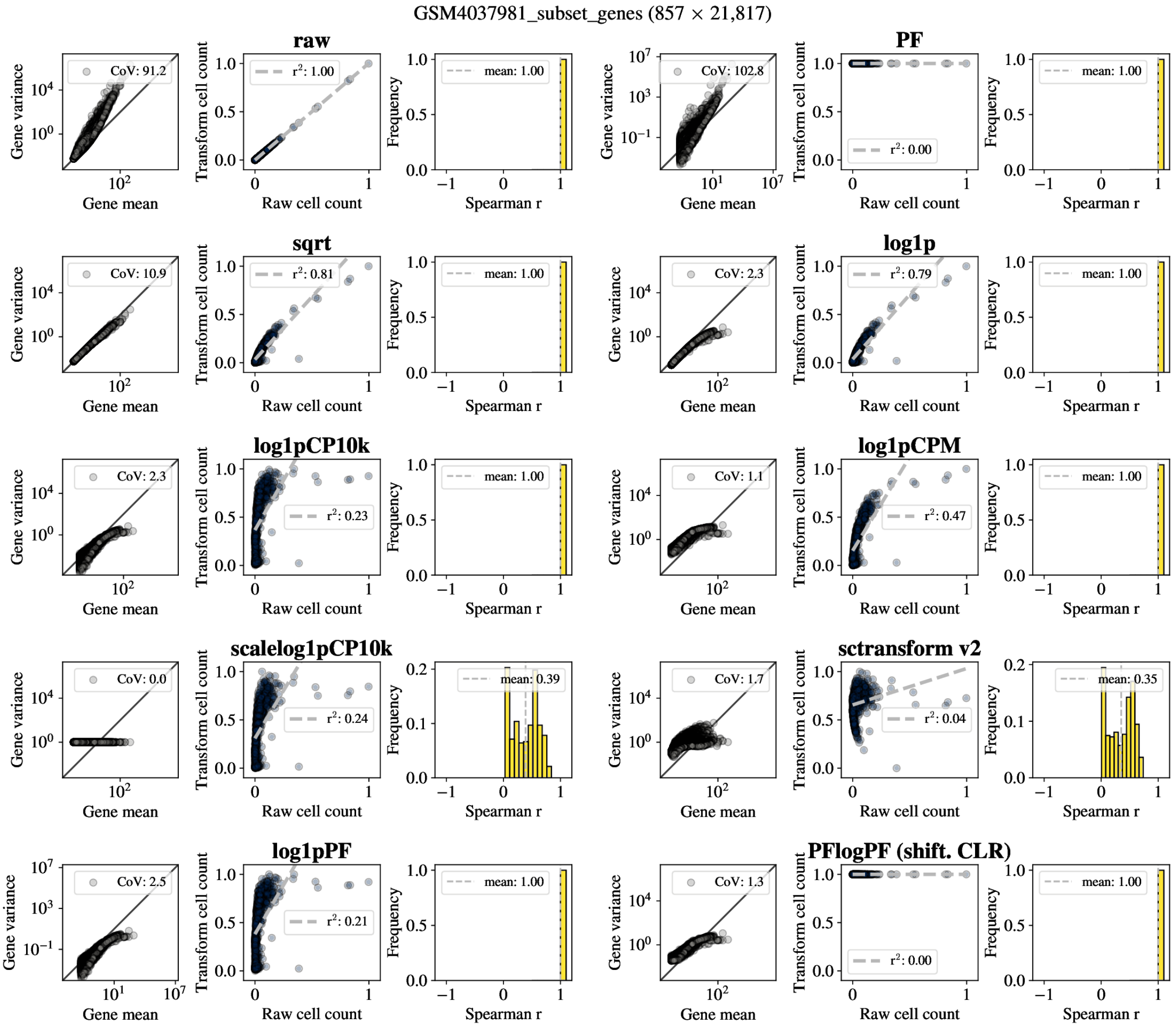


Supplementary Figure 1.463. **GSM4037981**. Per-method comparison of variance stabilization (left), depth normalization (middle), and monotonicity (right) for each normalization on dataset GSM4037981.

| GSM4037982 | |
|---|---|
| cells | 737 |
| genes | 60,623 |
| nonzero entries | 1,747,117 |
| density | 0.0391 |
| total counts | 8,247,485 |
| mean counts / cell | 11,191 |
| mean counts / gene | 136 |
| min counts / cell | 643 |
| max counts / cell | 199,360 |
| overdispersion | 4.41 |

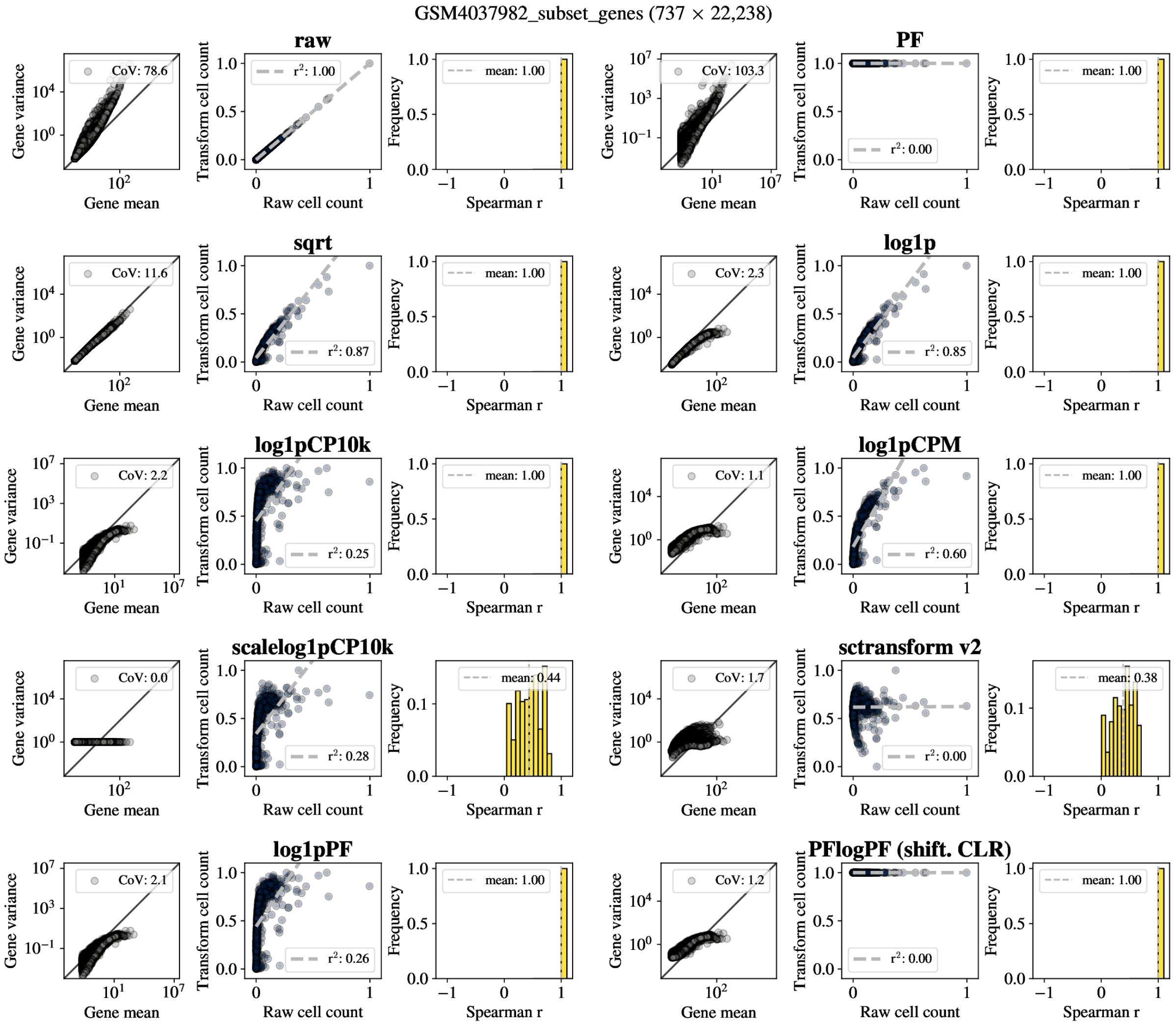


Supplementary Figure 1.464. **GSM4037982**. Per-method comparison of variance stabilization (left), depth normalization (middle), and monotonicity (right) for each normalization on dataset GSM4037982.

| GSM4037984 | |
|---|---|
| cells | 53 |
| genes | 60,623 |
| nonzero entries | 66,652 |
| density | 0.0207 |
| total counts | 264,831 |
| mean counts / cell | 4,997 |
| mean counts / gene | 4 |
| min counts / cell | 134 |
| max counts / cell | 27,703 |
| overdispersion | 3.09 |

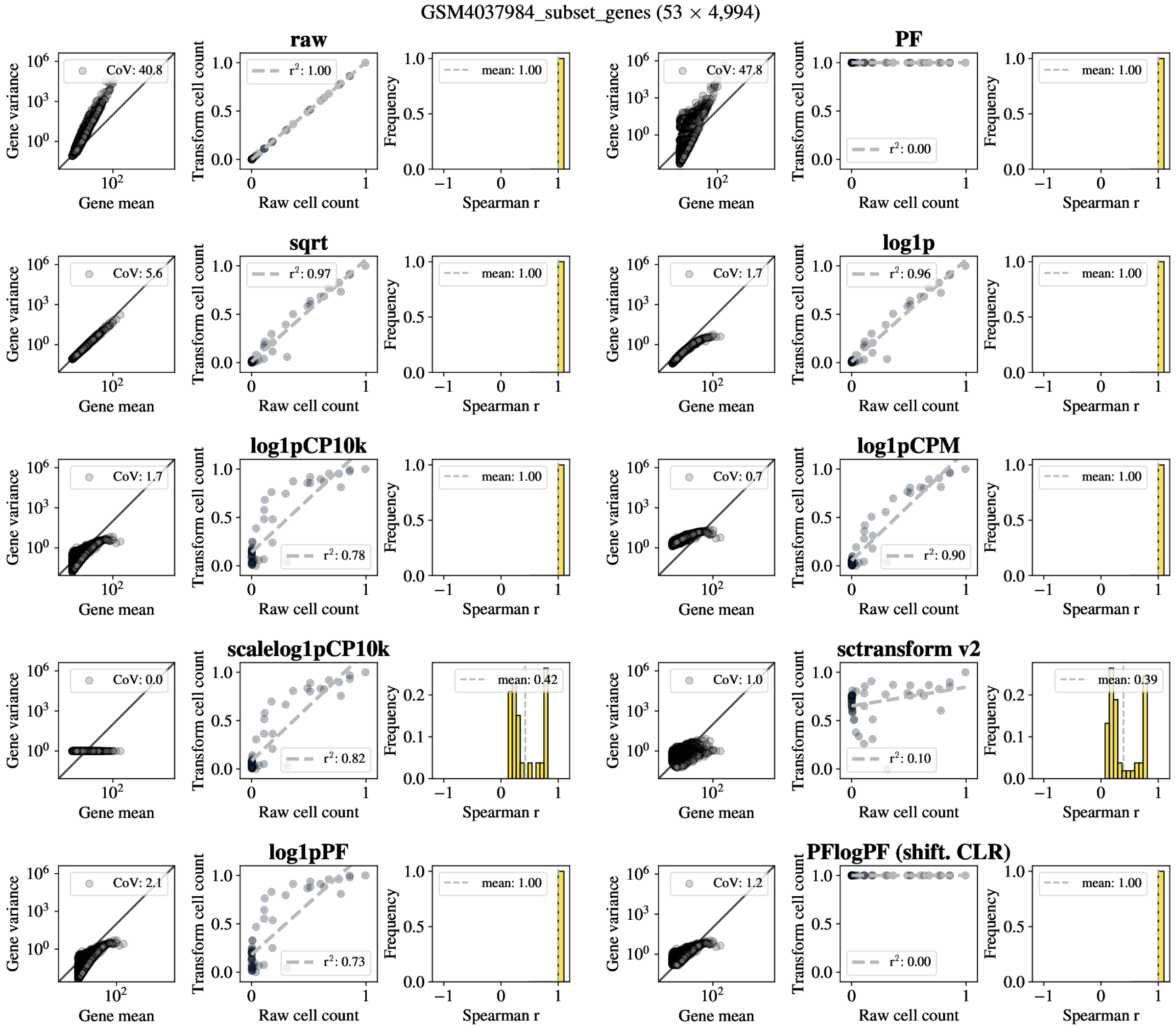


Supplementary Figure 1.465. **GSM4037984**. Per-method comparison of variance stabilization (left), depth normalization (middle), and monotonicity (right) for each normalization on dataset GSM4037984.

| GSM4037985 | |
|---|---|
| cells | 196 |
| genes | 60,623 |
| nonzero entries | 62,591 |
| density | 0.00527 |
| total counts | 223,359 |
| mean counts / cell | 1,140 |
| mean counts / gene | 4 |
| min counts / cell | 54 |
| max counts / cell | 30,784 |
| overdispersion | 20.8 |

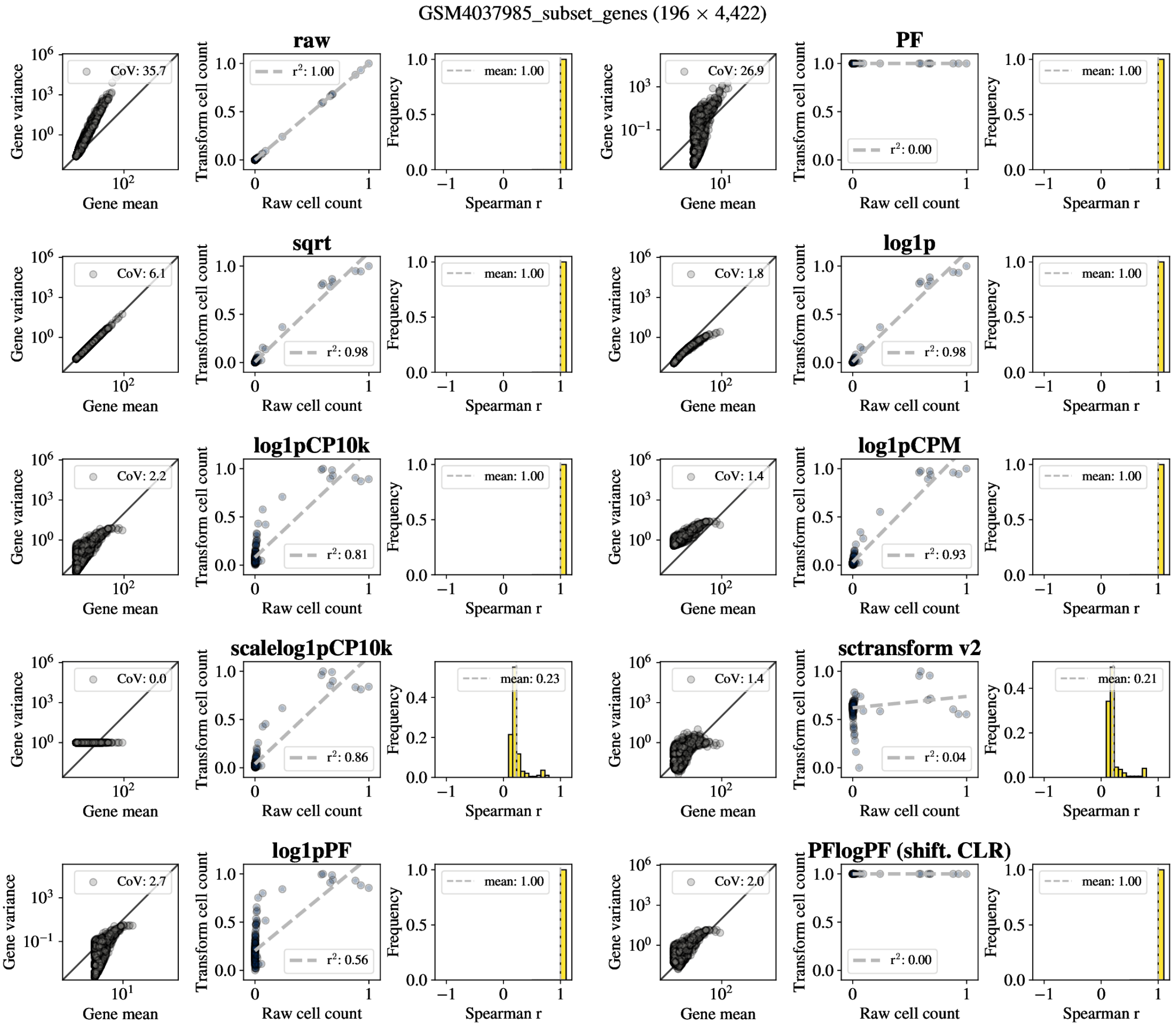


Supplementary Figure 1.466. **GSM4037985**. Per-method comparison of variance stabilization (left), depth normalization (middle), and monotonicity (right) for each normalization on dataset GSM4037985.

| GSM4037986 | |
|---|---|
| cells | 74 |
| genes | 60,623 |
| nonzero entries | 47,431 |
| density | 0.0106 |
| total counts | 181,135 |
| mean counts / cell | 2,448 |
| mean counts / gene | 3 |
| min counts / cell | 128 |
| max counts / cell | 40,795 |
| overdispersion | 10.8 |

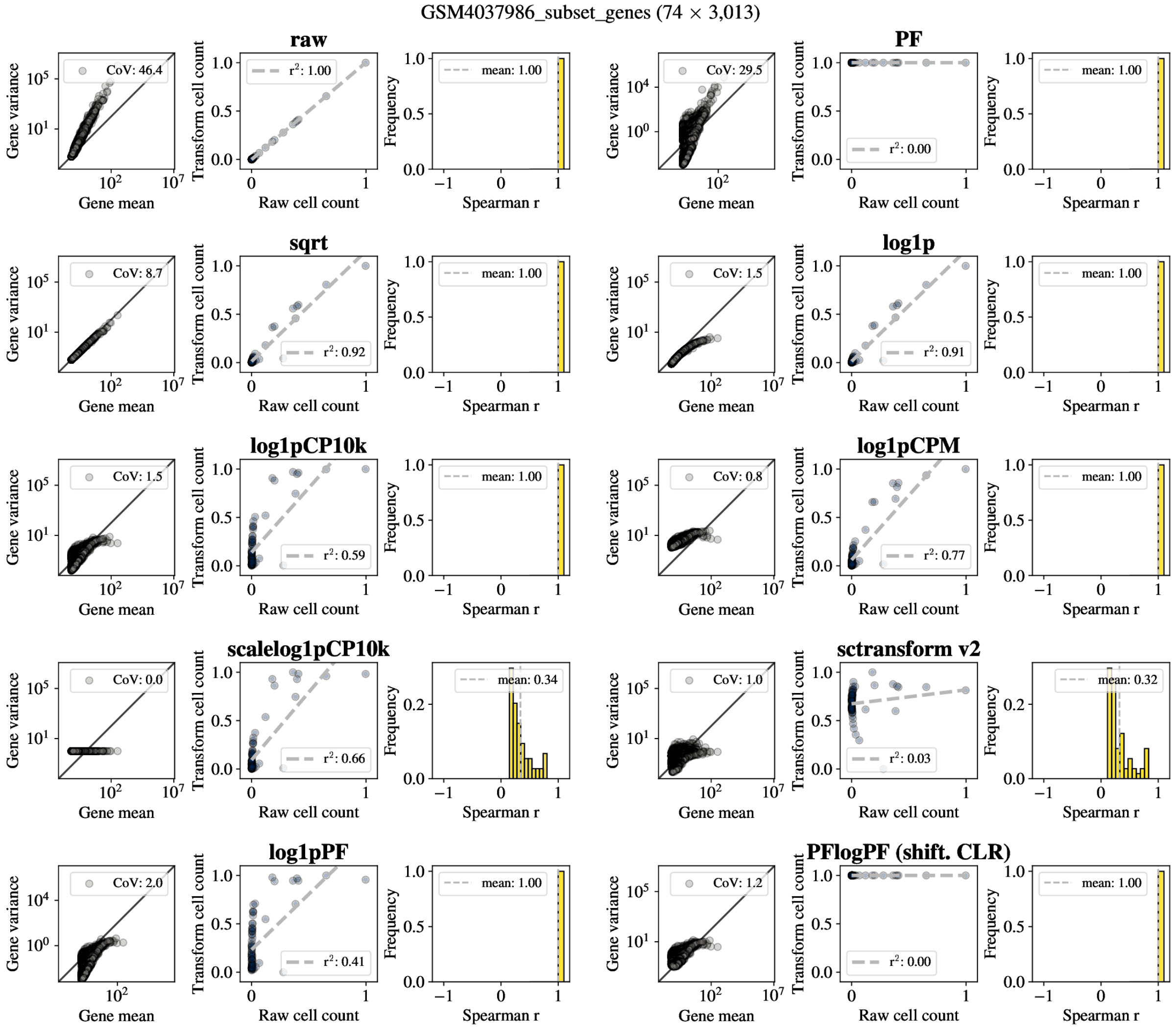


Supplementary Figure 1.467. **GSM4037986**. Per-method comparison of variance stabilization (left), depth normalization (middle), and monotonicity (right) for each normalization on dataset GSM4037986.

| GSM4037987 | |
|---|---|
| cells | 166 |
| genes | 60,623 |
| nonzero entries | 143,946 |
| density | 0.0143 |
| total counts | 603,931 |
| mean counts / cell | 3,638 |
| mean counts / gene | 10 |
| min counts / cell | 181 |
| max counts / cell | 74,025 |
| overdispersion | 5.14 |

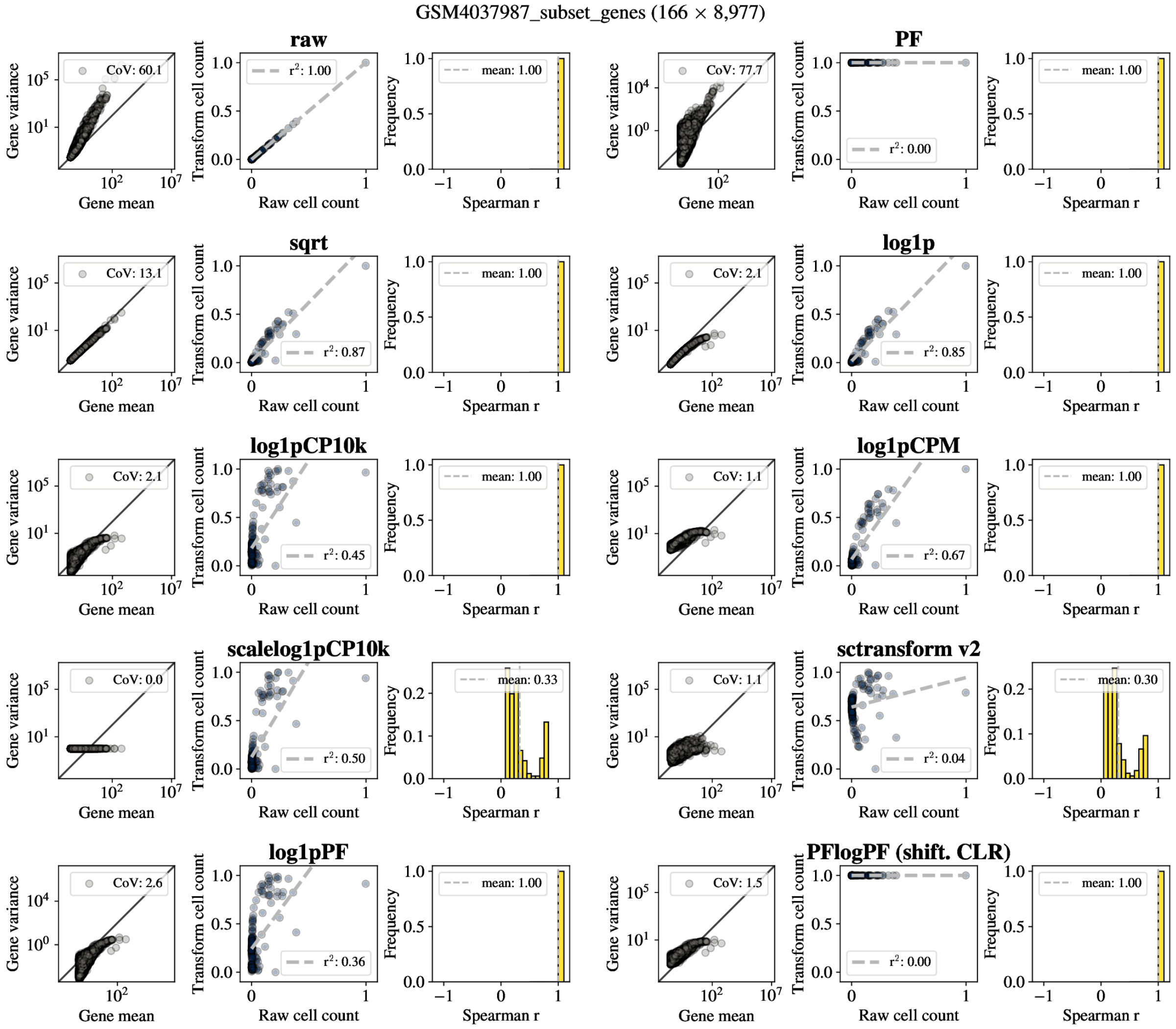


Supplementary Figure 1.468. **GSM4037987**. Per-method comparison of variance stabilization (left), depth normalization (middle), and monotonicity (right) for each normalization on dataset GSM4037987.

| GSM4037988 | |
|---|---|
| cells | 321 |
| genes | 60,623 |
| nonzero entries | 149,362 |
| density | 0.00768 |
| total counts | 503,843 |
| mean counts / cell | 1,570 |
| mean counts / gene | 8 |
| min counts / cell | 107 |
| max counts / cell | 62,060 |
| overdispersion | 11.5 |

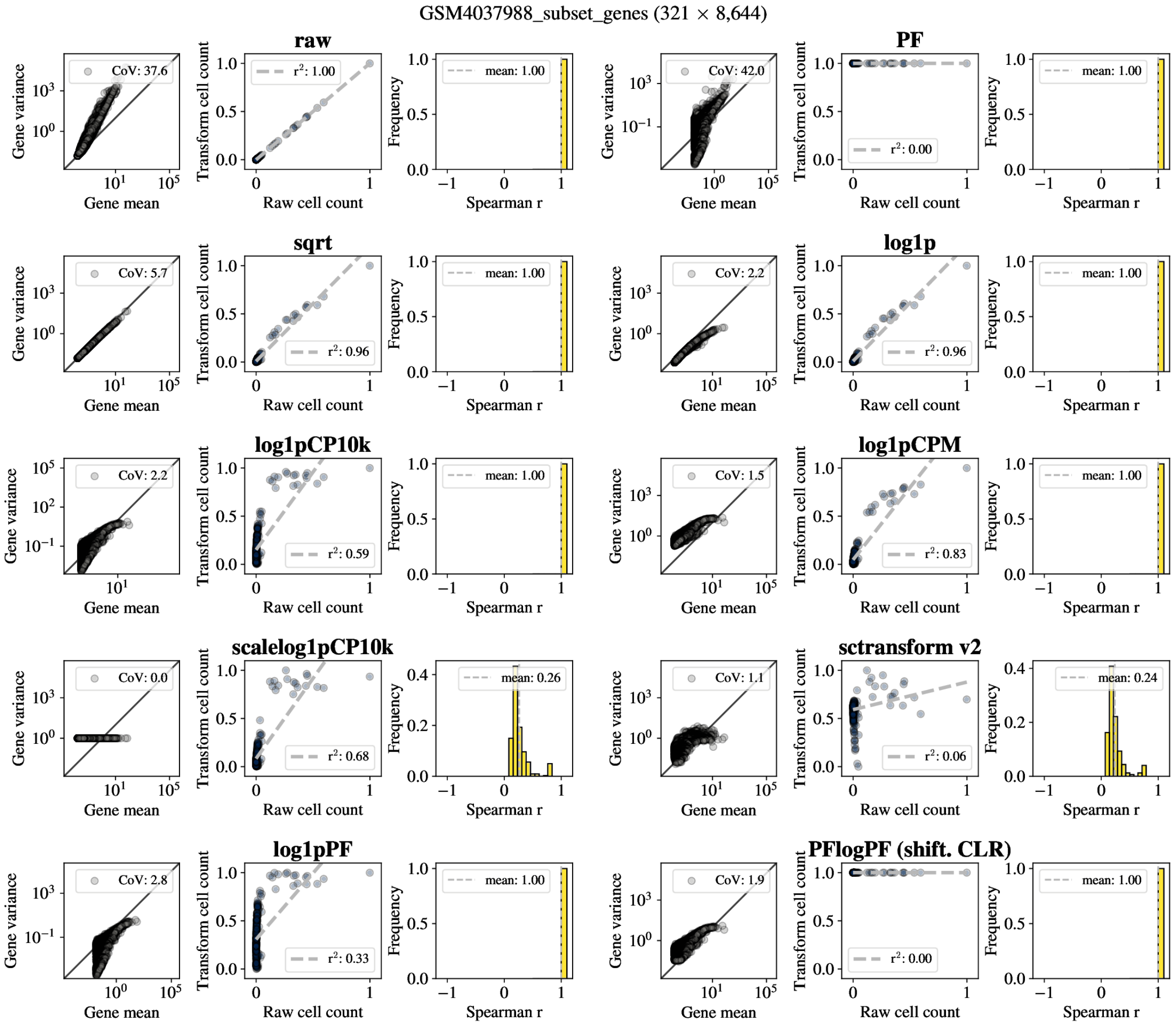


Supplementary Figure 1.469. **GSM4037988**. Per-method comparison of variance stabilization (left), depth normalization (middle), and monotonicity (right) for each normalization on dataset GSM4037988.

| **GSM4037989** | |
|---|---|
| cells | 1,914 |
| genes | 60,623 |
| nonzero entries | 124,626 |
| density | 0.00107 |
| total counts | 382,944 |
| mean counts / cell | 200 |
| mean counts / gene | 6 |
| min counts / cell | 5 |
| max counts / cell | 34,619 |
| overdispersion | 52.4 |

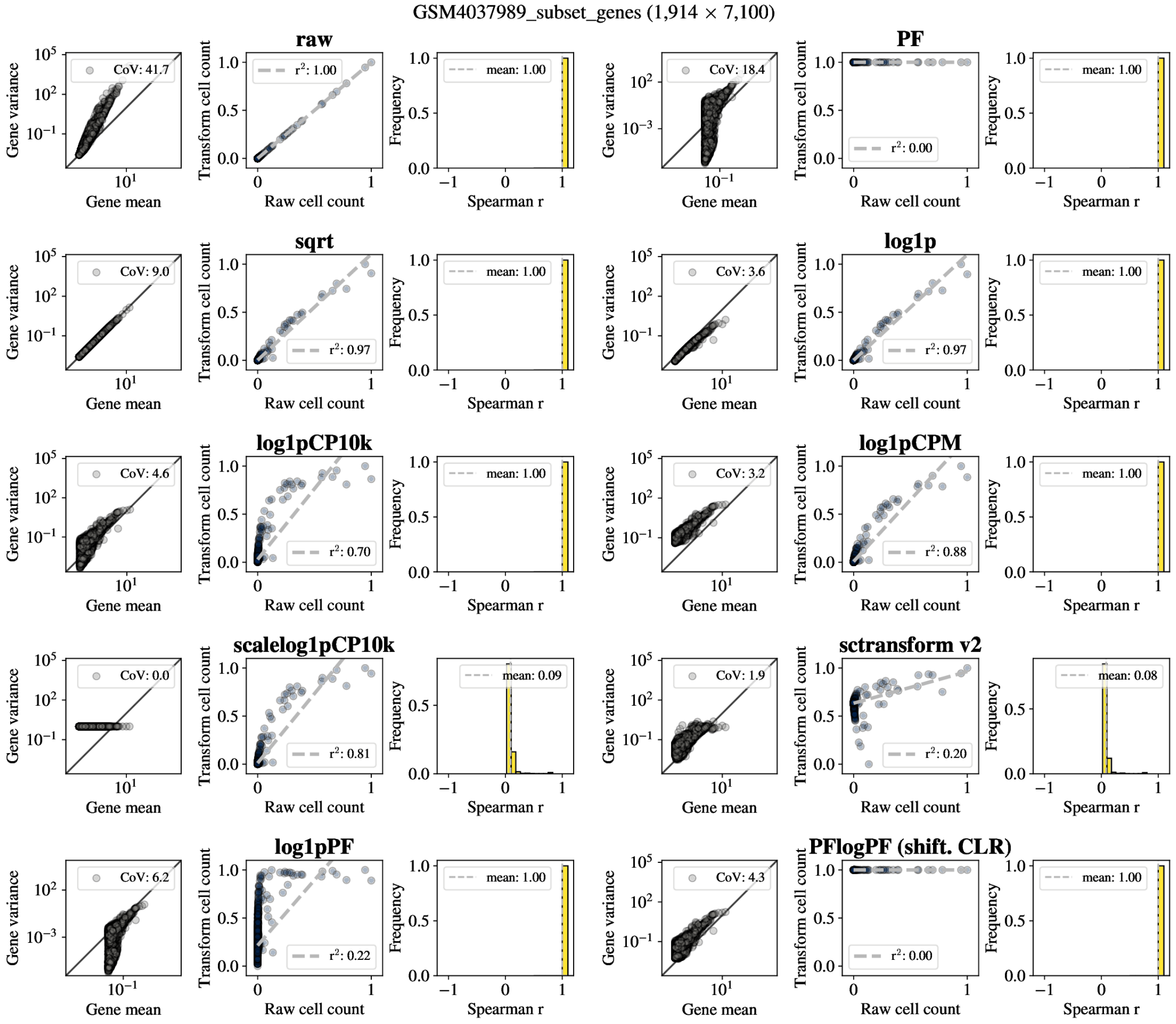


Supplementary Figure 1.470. **GSM4037989**. Per-method comparison of variance stabilization (left), depth normalization (middle), and monotonicity (right) for each normalization on dataset GSM4037989.

| GSM4037990 | |
|---|---|
| cells | 2,352 |
| genes | 60,623 |
| nonzero entries | 146,354 |
| density | 0.00103 |
| total counts | 515,994 |
| mean counts / cell | 219 |
| mean counts / gene | 9 |
| min counts / cell | 6 |
| max counts / cell | 105,700 |
| overdispersion | 134 |

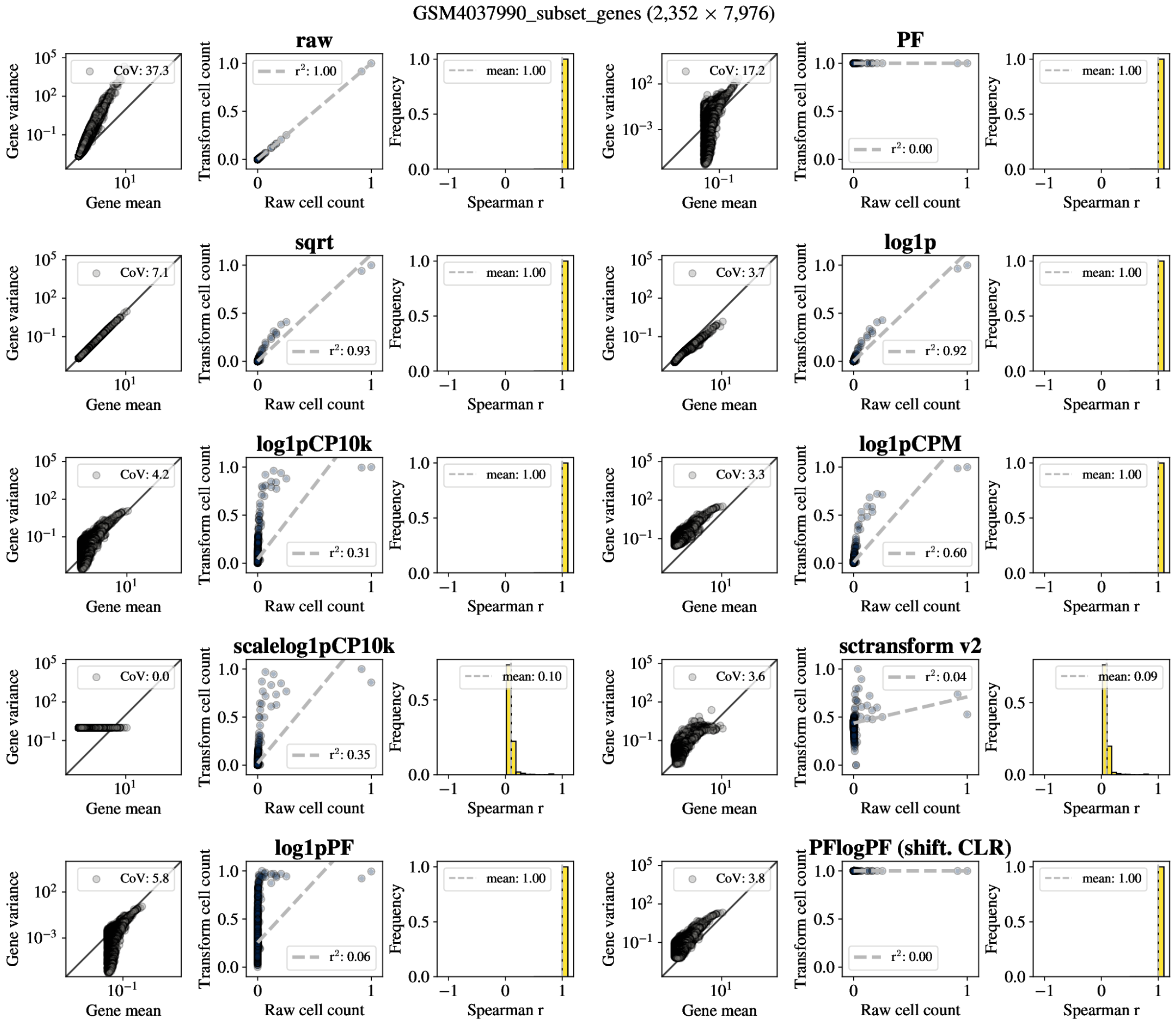


Supplementary Figure 1.471. **GSM4037990**. Per-method comparison of variance stabilization (left), depth normalization (middle), and monotonicity (right) for each normalization on dataset GSM4037990.

| GSM4037991 | |
|---|---|
| cells | 175 |
| genes | 60,623 |
| nonzero entries | 85,774 |
| density | 0.00809 |
| total counts | 380,478 |
| mean counts / cell | 2,174 |
| mean counts / gene | 6 |
| min counts / cell | 104 |
| max counts / cell | 76,962 |
| overdispersion | 6.79 |

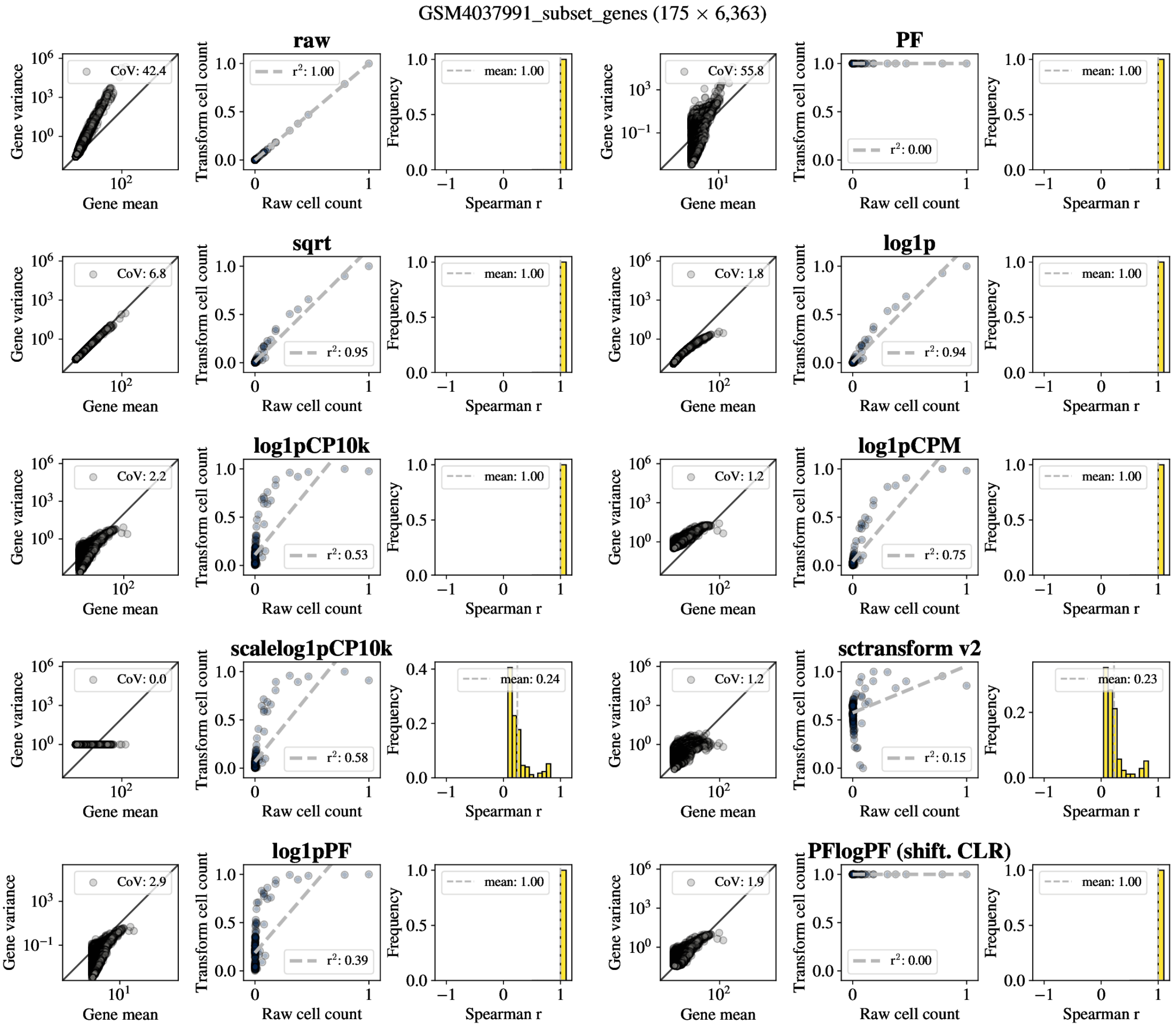


Supplementary Figure 1.472. **GSM4037991**. Per-method comparison of variance stabilization (left), depth normalization (middle), and monotonicity (right) for each normalization on dataset GSM4037991.

| GSM4037992 | |
|---|---|
| cells | 162 |
| genes | 60,623 |
| nonzero entries | 78,890 |
| density | 0.00803 |
| total counts | 278,851 |
| mean counts / cell | 1,721 |
| mean counts / gene | 5 |
| min counts / cell | 121 |
| max counts / cell | 49,220 |
| overdispersion | 5.9 |

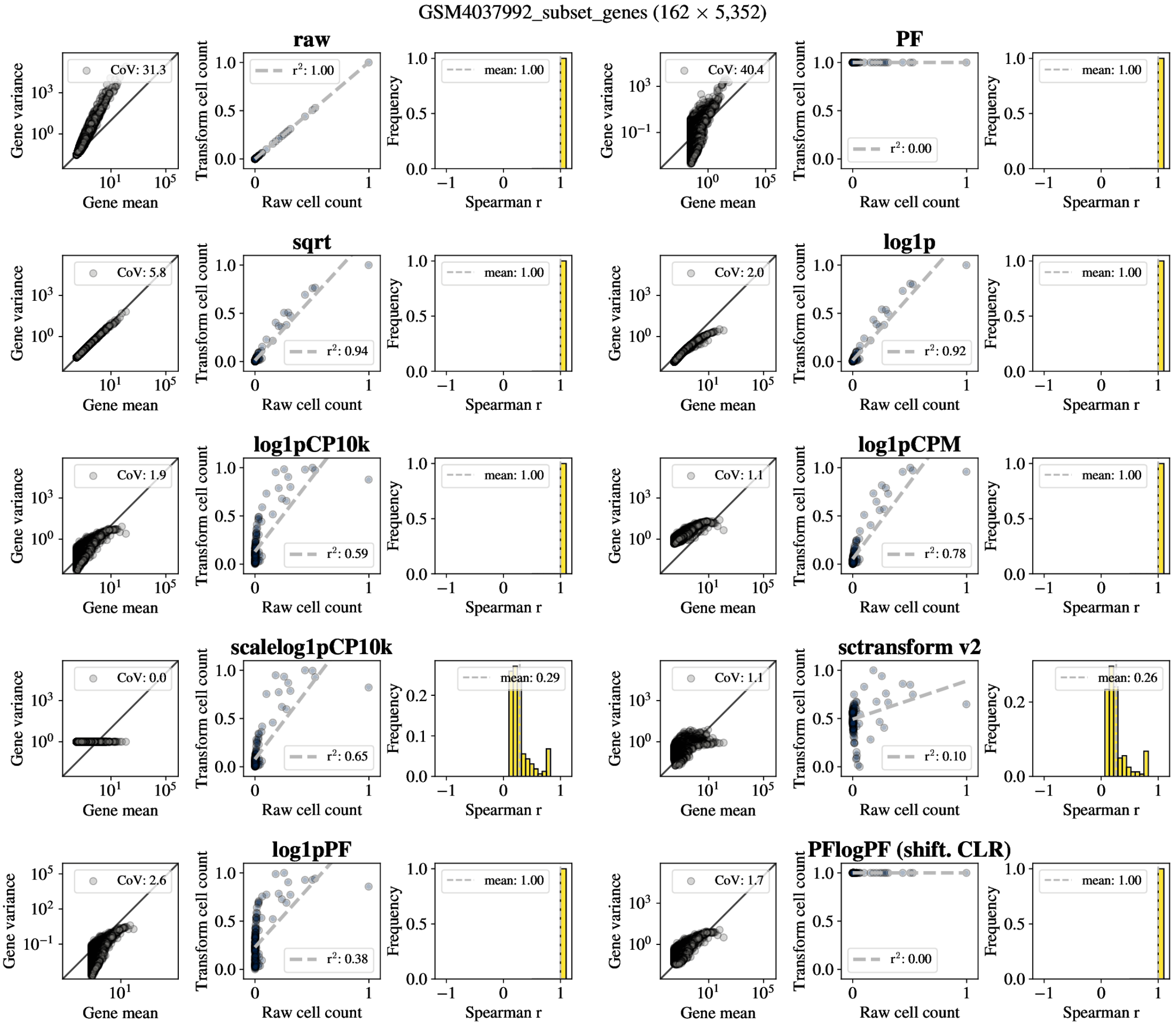


Supplementary Figure 1.473. **GSM4037992**. Per-method comparison of variance stabilization (left), depth normalization (middle), and monotonicity (right) for each normalization on dataset GSM4037992.

| GSM4037993 | |
|---|---|
| cells | 60 |
| genes | 60,623 |
| nonzero entries | 73,763 |
| density | 0.0203 |
| total counts | 266,106 |
| mean counts / cell | 4,435 |
| mean counts / gene | 4 |
| min counts / cell | 475 |
| max counts / cell | 32,222 |
| overdispersion | 1.19 |

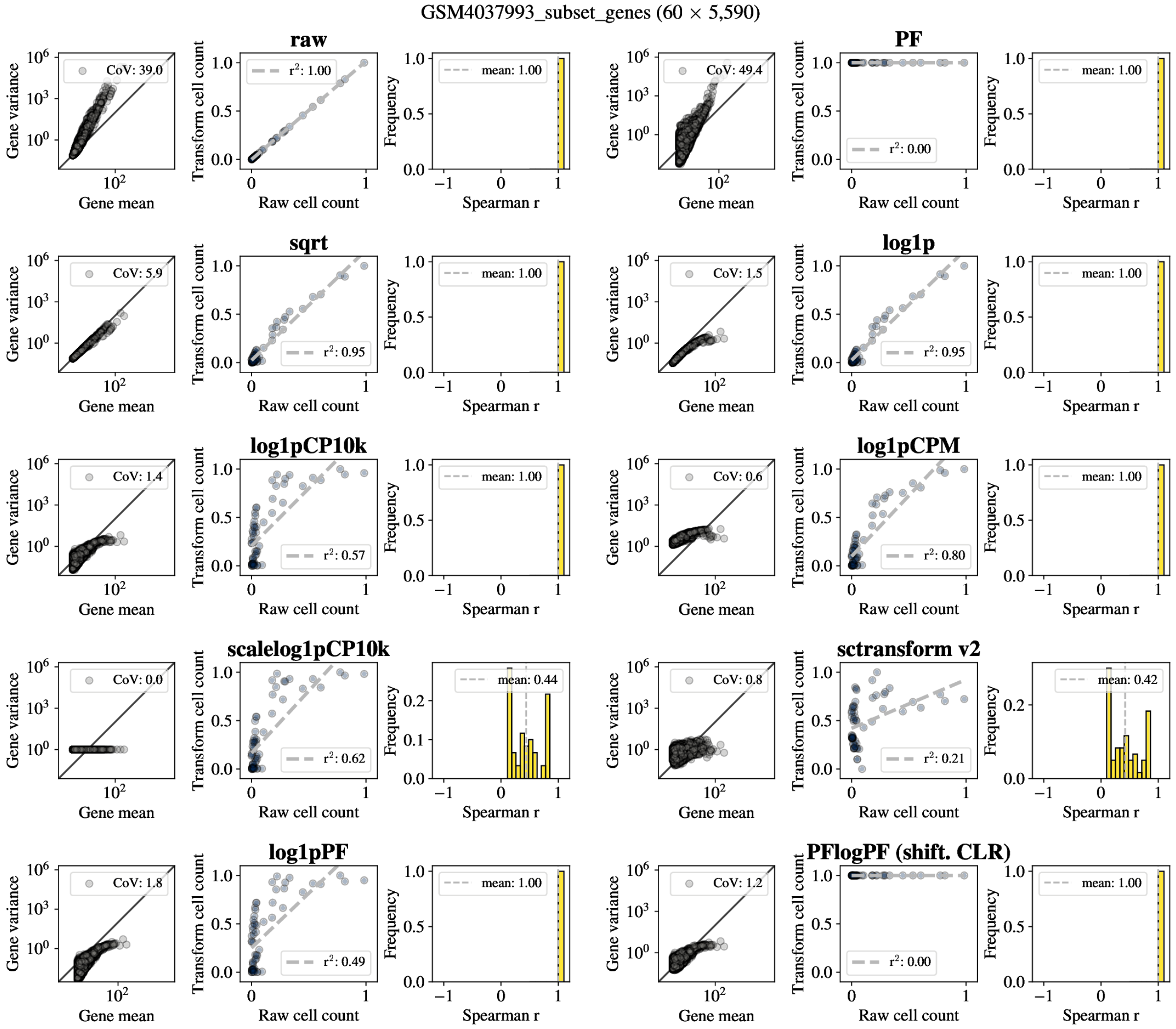


Supplementary Figure 1.474. **GSM4037993**. Per-method comparison of variance stabilization (left), depth normalization (middle), and monotonicity (right) for each normalization on dataset GSM4037993.

| GSM4037994 | |
|---|---|
| cells | 77 |
| genes | 60,623 |
| nonzero entries | 115,992 |
| density | 0.0248 |
| total counts | 436,335 |
| mean counts / cell | 5,667 |
| mean counts / gene | 7 |
| min counts / cell | 618 |
| max counts / cell | 43,607 |
| overdispersion | 4.24 |

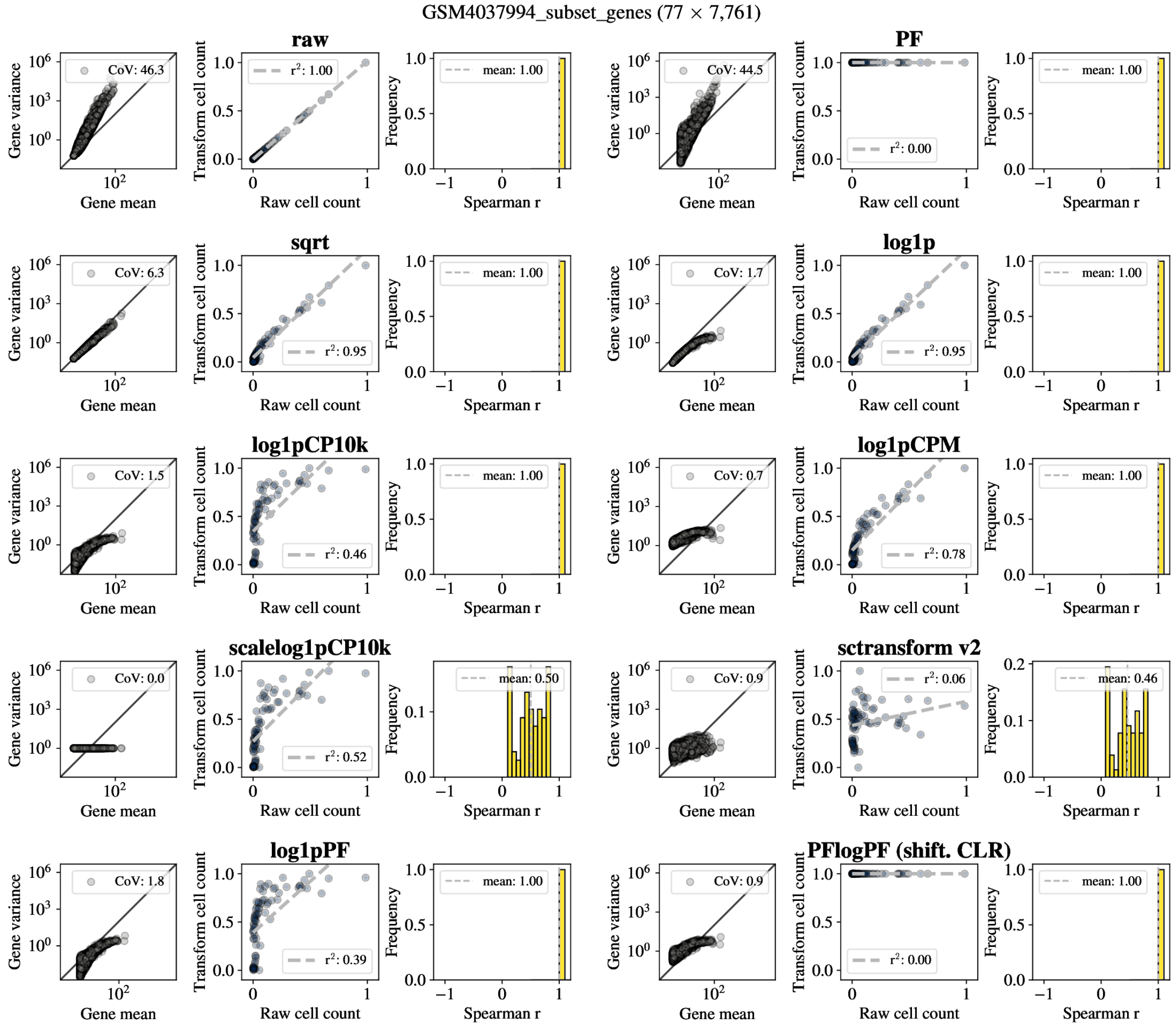


Supplementary Figure 1.475. **GSM4037994**. Per-method comparison of variance stabilization (left), depth normalization (middle), and monotonicity (right) for each normalization on dataset GSM4037994.

| GSM4080085 | |
|---|---|
| cells | 12,351 |
| genes | 60,623 |
| nonzero entries | 24,428,199 |
| density | 0.0326 |
| total counts | 107,412,094 |
| mean counts / cell | 8,697 |
| mean counts / gene | 1,772 |
| min counts / cell | 628 |
| max counts / cell | 87,051 |
| overdispersion | 2.59 |

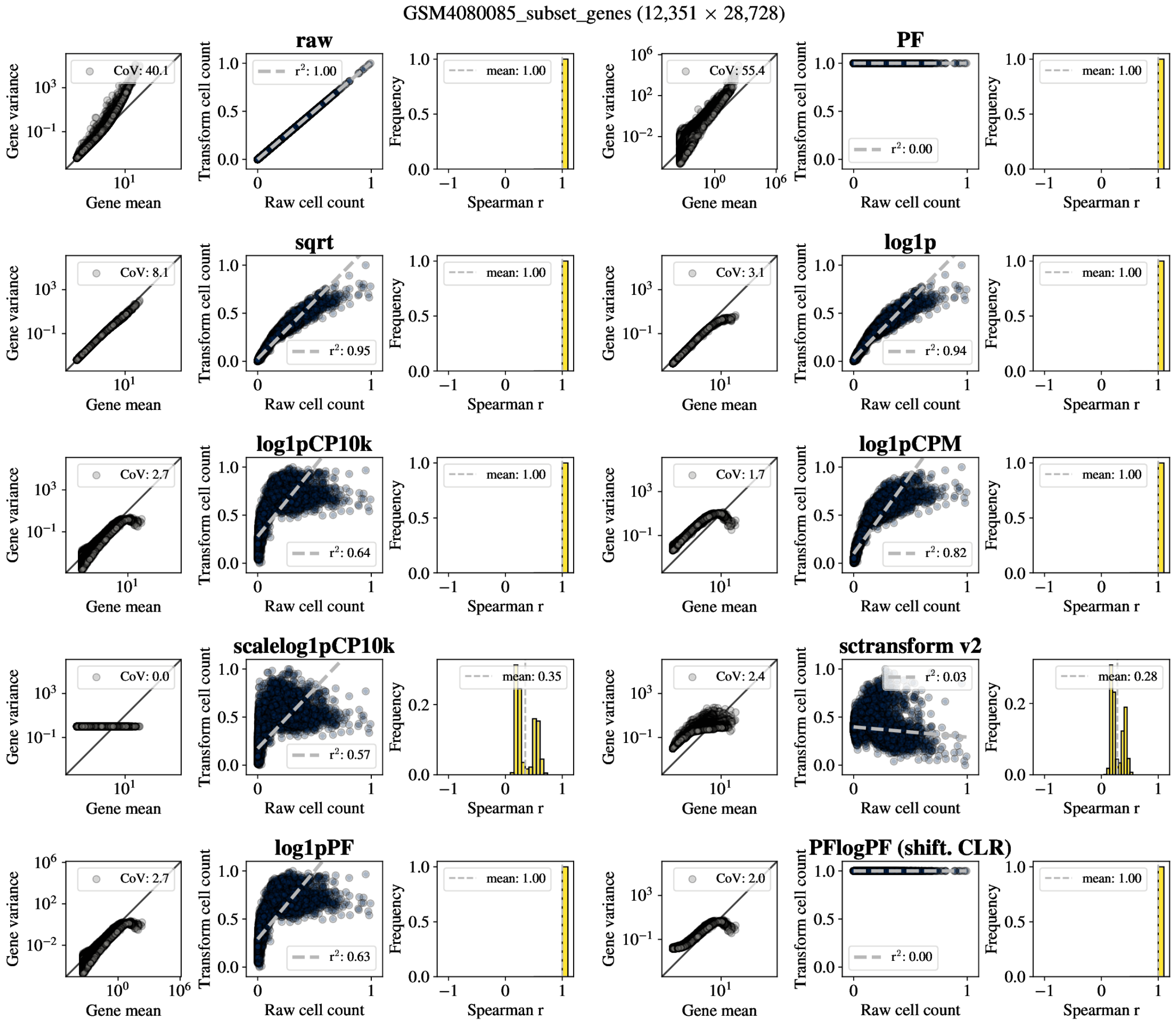


Supplementary Figure 1.476. **GSM4080085**. Per-method comparison of variance stabilization (left), depth normalization (middle), and monotonicity (right) for each normalization on dataset GSM4080085.

| GSM4080086 | |
|---|---|
| cells | 8,983 |
| genes | 60,623 |
| nonzero entries | 20,032,876 |
| density | 0.0368 |
| total counts | 77,514,201 |
| mean counts / cell | 8,629 |
| mean counts / gene | 1,279 |
| min counts / cell | 540 |
| max counts / cell | 65,714 |
| overdispersion | 1.72 |

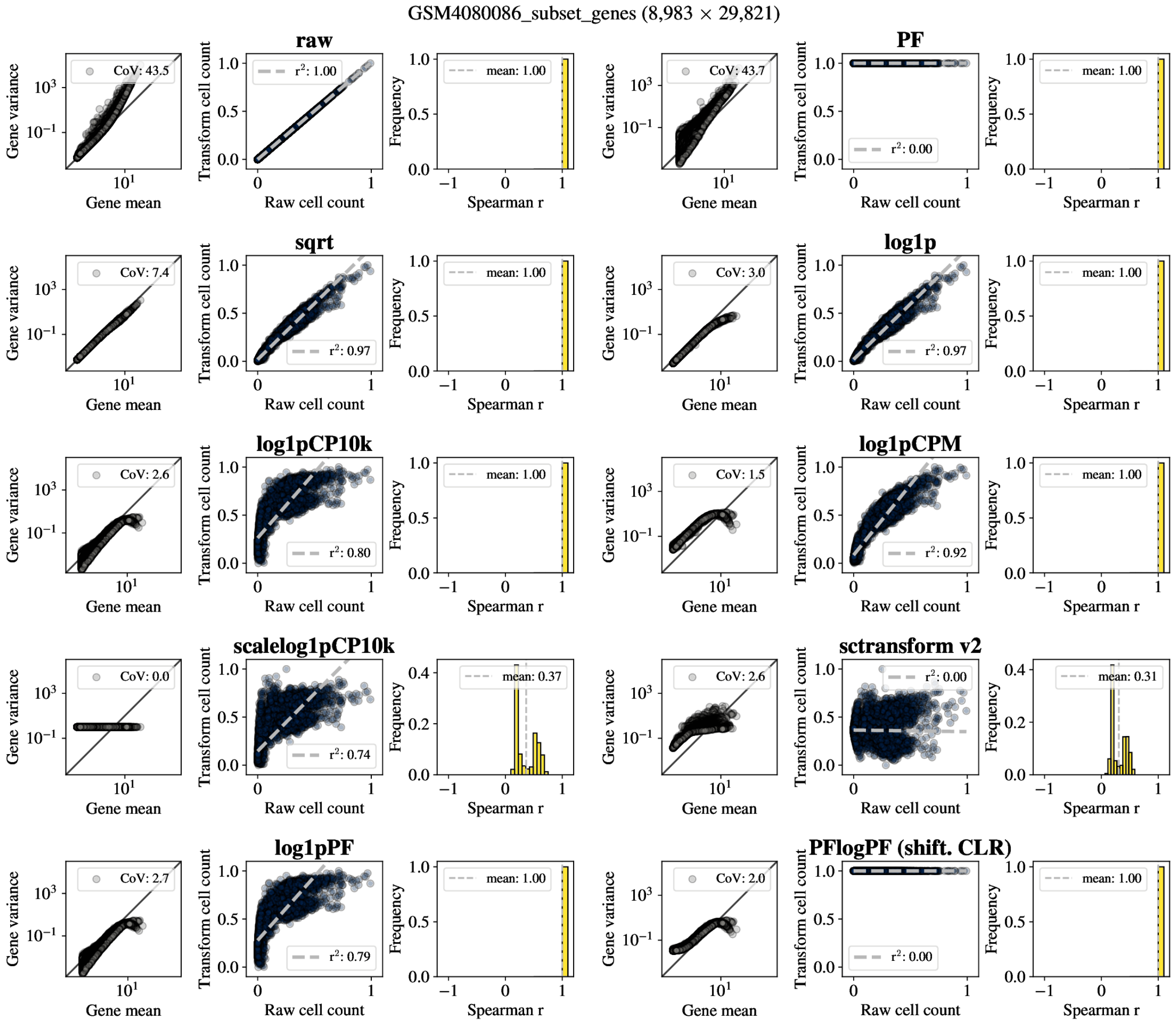


Supplementary Figure 1.477. **GSM4080086**. Per-method comparison of variance stabilization (left), depth normalization (middle), and monotonicity (right) for each normalization on dataset GSM4080086.

| GSM4080087 | |
| --- | --- |
| cells | 7,041 |
| genes | 60,623 |
| nonzero entries | 10,630,749 |
| density | 0.0249 |
| total counts | 39,751,114 |
| mean counts / cell | 5,646 |
| mean counts / gene | 656 |
| min counts / cell | 190 |
| max counts / cell | 51,351 |
| overdispersion | 2.32 |

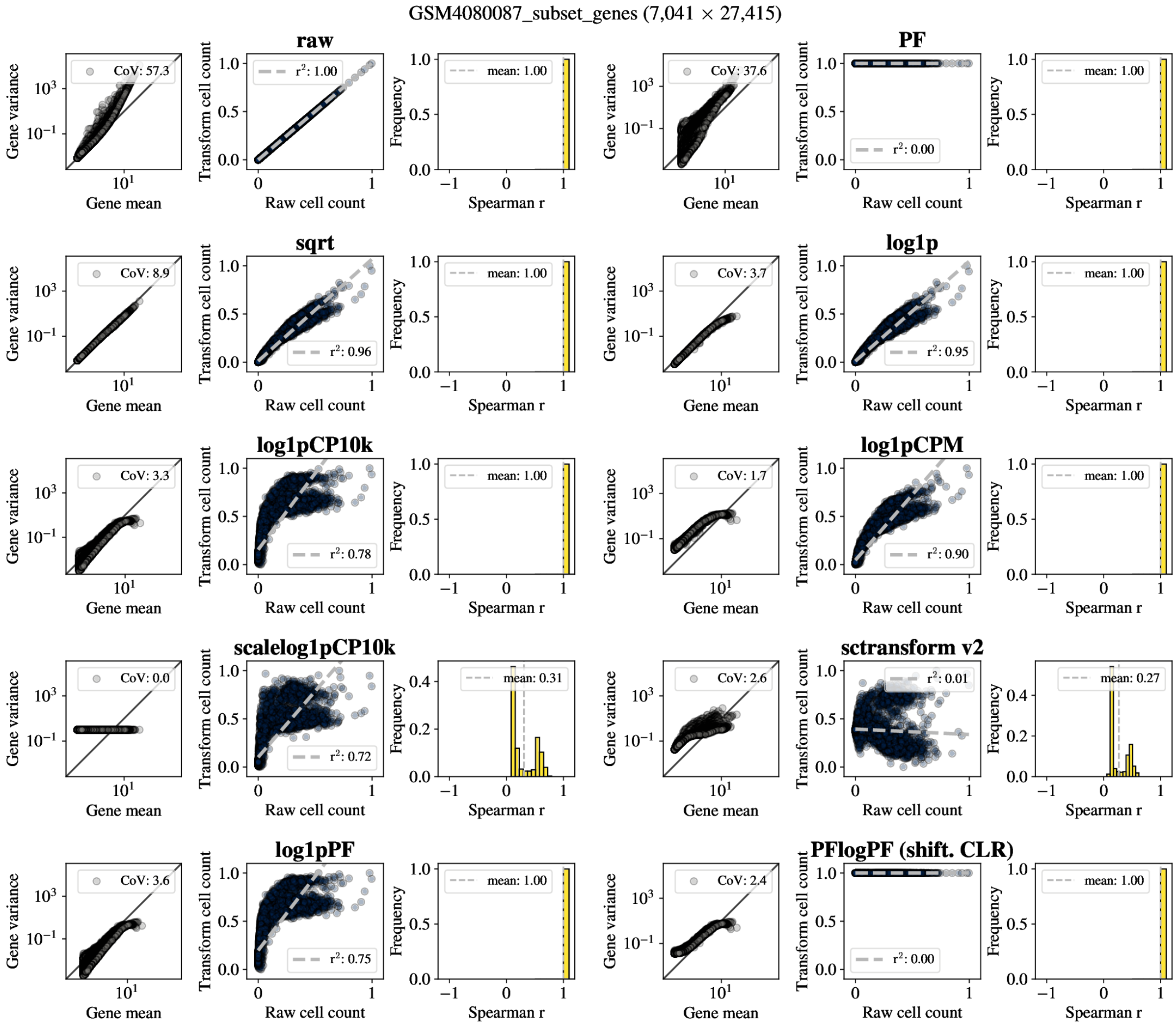


Supplementary Figure 1.478. **GSM4080087**. Per-method comparison of variance stabilization (left), depth normalization (middle), and monotonicity (right) for each normalization on dataset GSM4080087.

| GSM4080088 | |
| --- | --- |
| cells | 2,515 |
| genes | 60,623 |
| nonzero entries | 5,945,767 |
| density | 0.039 |
| total counts | 25,128,762 |
| mean counts / cell | 9,992 |
| mean counts / gene | 415 |
| min counts / cell | 195 |
| max counts / cell | 66,612 |
| overdispersion | 1.55 |

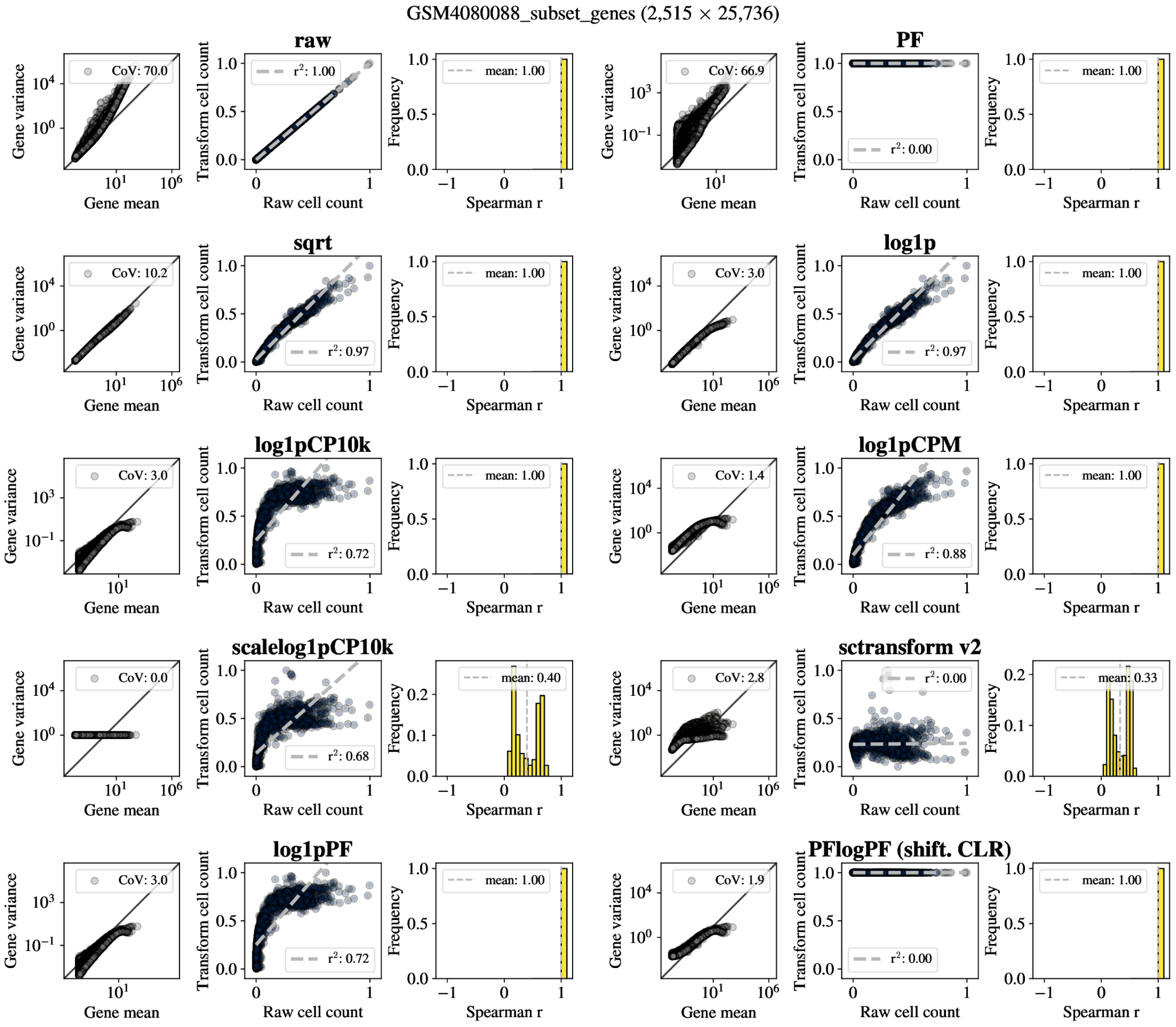


Supplementary Figure 1.479. **GSM4080088**. Per-method comparison of variance stabilization (left), depth normalization (middle), and monotonicity (right) for each normalization on dataset GSM4080088.

| GSM4080089 | |
|---|---|
| cells | 6,471 |
| genes | 60,623 |
| nonzero entries | 8,783,368 |
| density | 0.0224 |
| total counts | 38,244,404 |
| mean counts / cell | 5,910 |
| mean counts / gene | 631 |
| min counts / cell | 220 |
| max counts / cell | 96,766 |
| overdispersion | 3.76 |

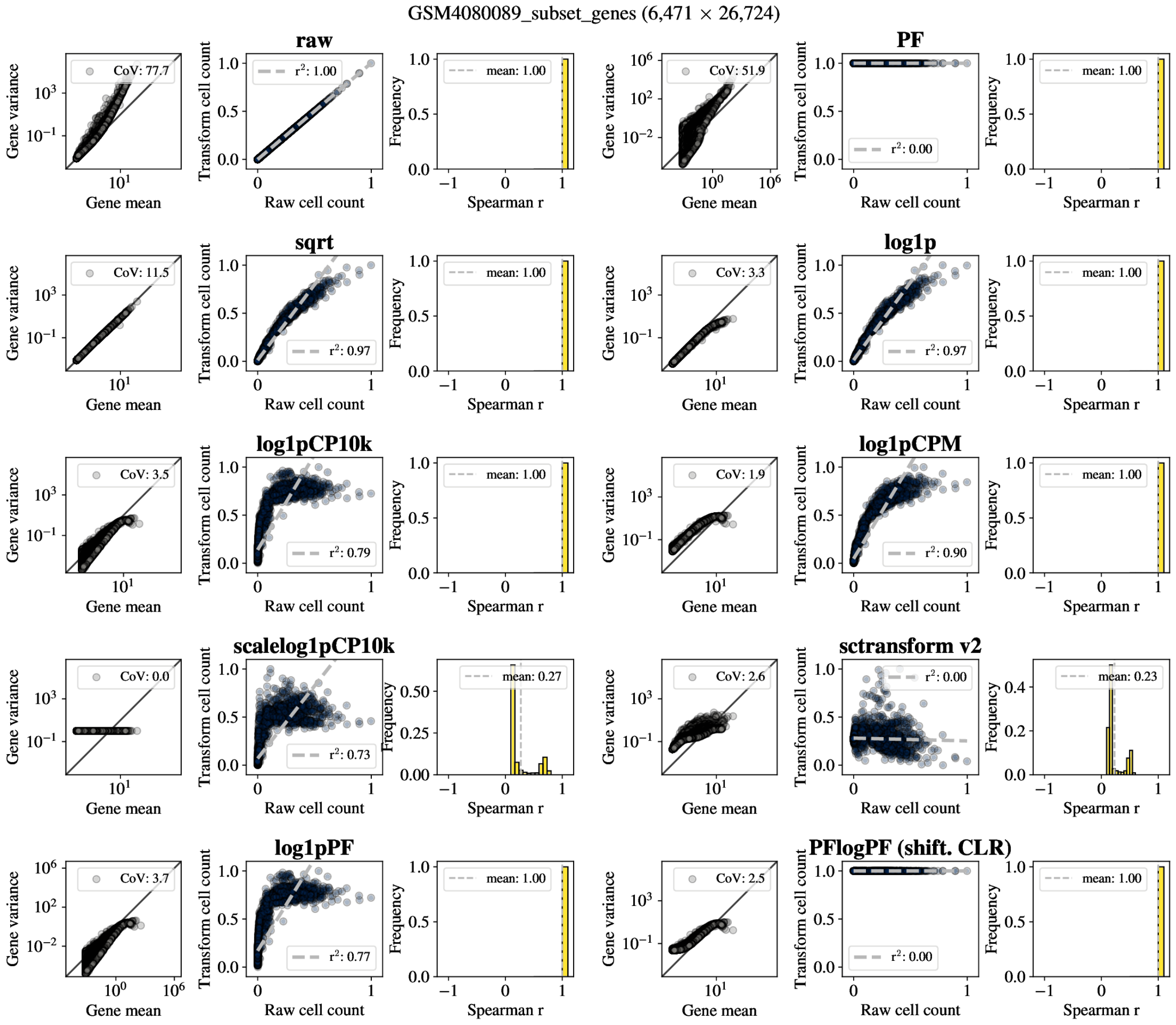


Supplementary Figure 1.480. **GSM4080089**. Per-method comparison of variance stabilization (left), depth normalization (middle), and monotonicity (right) for each normalization on dataset GSM4080089.

| GSM4080090 | |
|---|---|
| cells | 3,826 |
| genes | 60,623 |
| nonzero entries | 9,028,074 |
| density | 0.0389 |
| total counts | 39,963,575 |
| mean counts / cell | 10,445 |
| mean counts / gene | 659 |
| min counts / cell | 247 |
| max counts / cell | 104,611 |
| overdispersion | 1.64 |

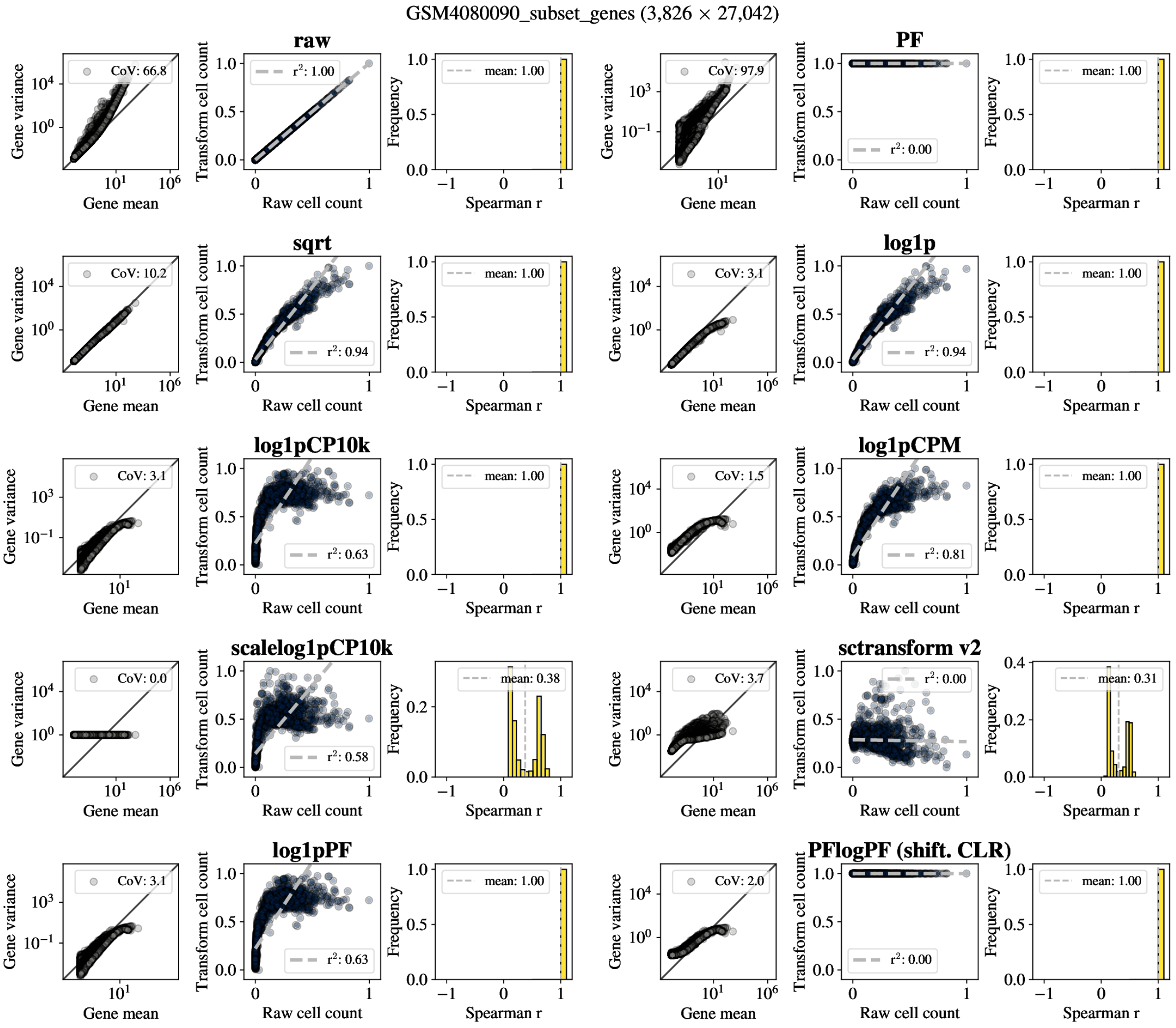


Supplementary Figure 1.481. **GSM4080090**. Per-method comparison of variance stabilization (left), depth normalization (middle), and monotonicity (right) for each normalization on dataset GSM4080090.

| GSM4142870 | |
|---|---|
| cells | 4,943 |
| genes | 60,623 |
| nonzero entries | 13,084,727 |
| density | 0.0437 |
| total counts | 49,610,986 |
| mean counts / cell | 10,037 |
| mean counts / gene | 818 |
| min counts / cell | 2,770 |
| max counts / cell | 52,374 |
| overdispersion | 0.697 |

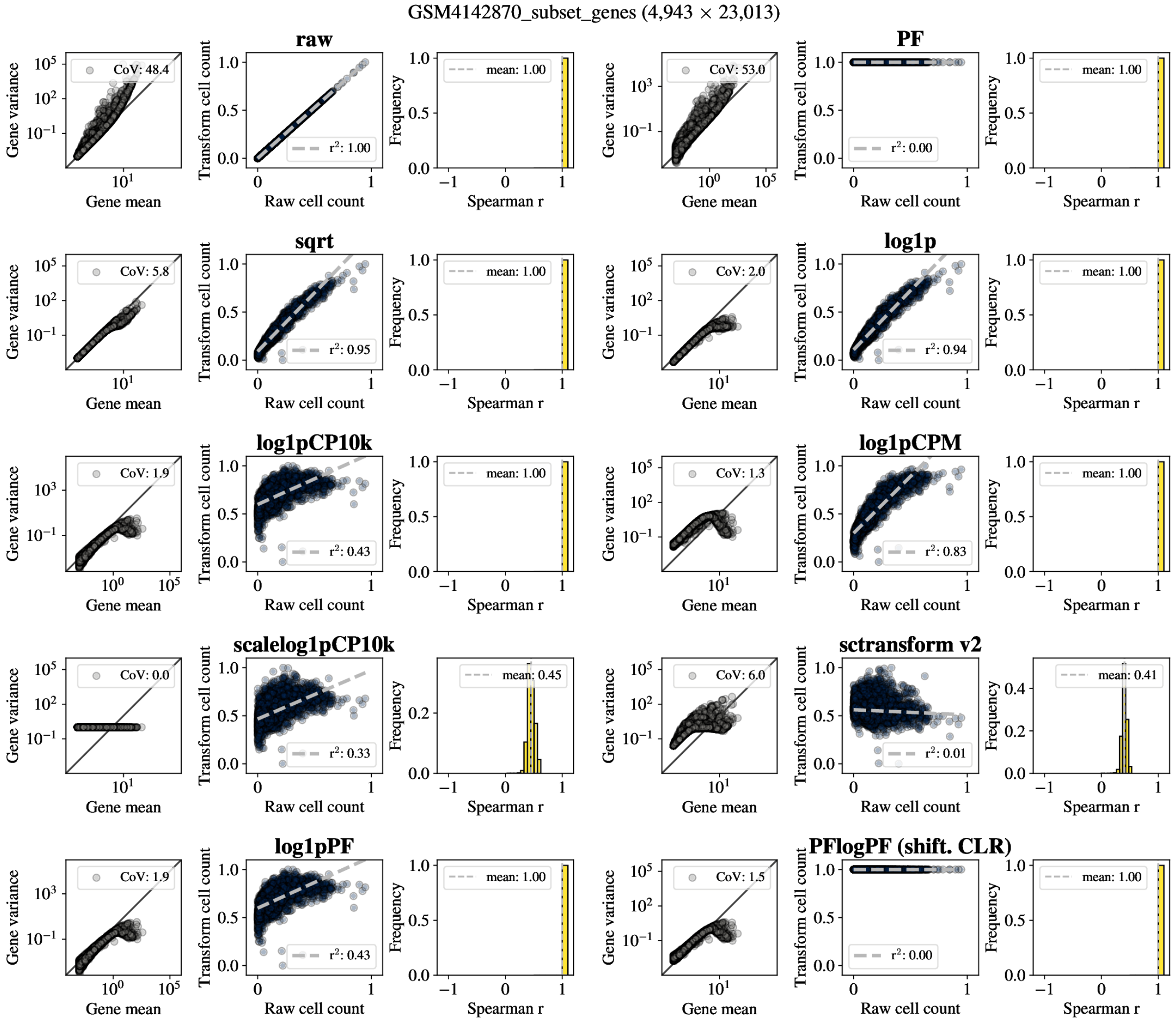


Supplementary Figure 1.482. **GSM4142870**. Per-method comparison of variance stabilization (left), depth normalization (middle), and monotonicity (right) for each normalization on dataset GSM4142870.

| GSM4142871 | |
|---|---|
| cells | 163 |
| genes | 60,623 |
| nonzero entries | 431,370 |
| density | 0.0437 |
| total counts | 1,636,711 |
| mean counts / cell | 10,041 |
| mean counts / gene | 27 |
| min counts / cell | 334 |
| max counts / cell | 83,241 |
| overdispersion | 3.09 |

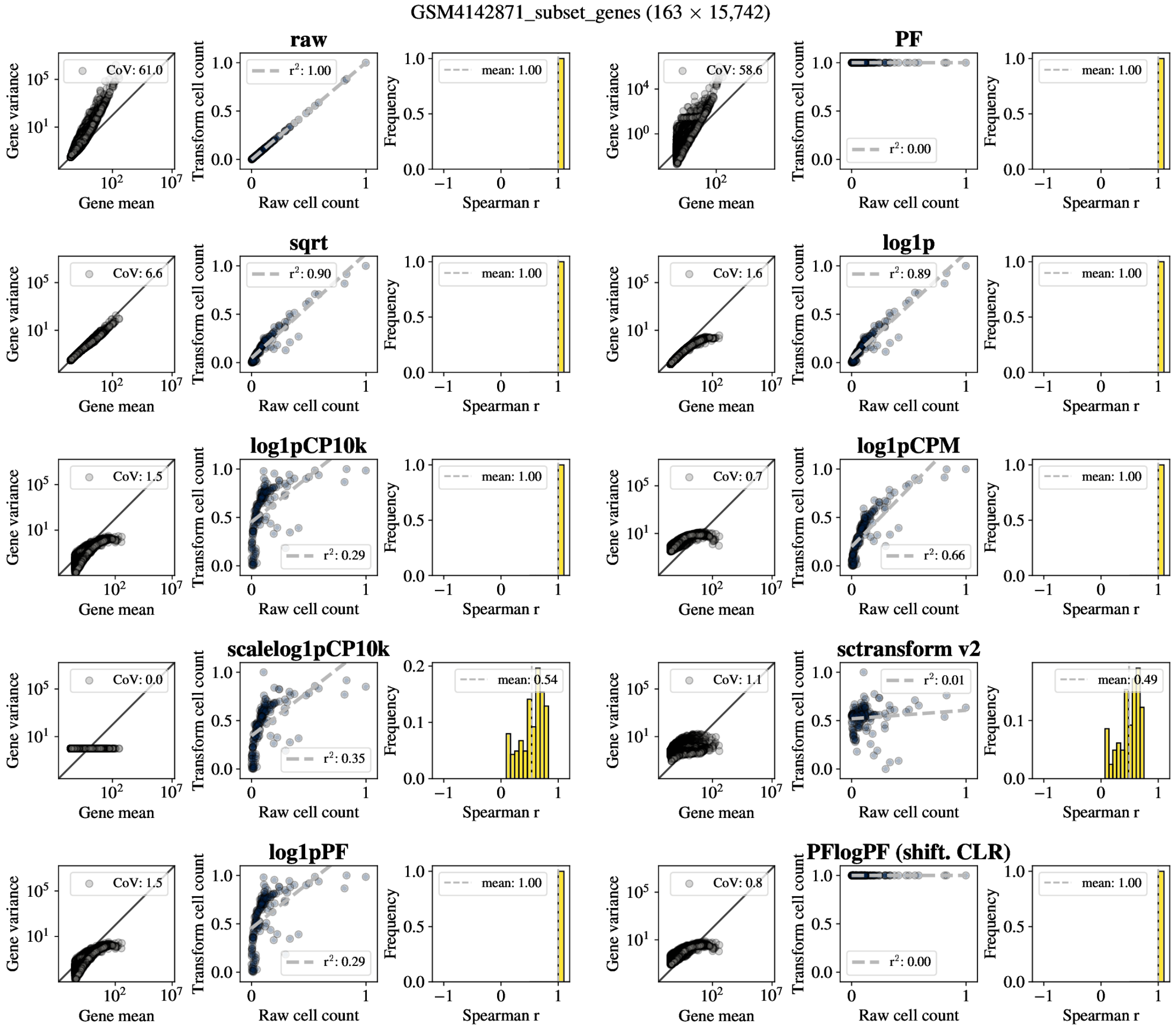


Supplementary Figure 1.483. **GSM4142871**. Per-method comparison of variance stabilization (left), depth normalization (middle), and monotonicity (right) for each normalization on dataset GSM4142871.

| GSM4142872 | |
|---|---|
| cells | 2,945 |
| genes | 60,623 |
| nonzero entries | 184,262 |
| density | 0.00103 |
| total counts | 419,111 |
| mean counts / cell | 142 |
| mean counts / gene | 7 |
| min counts / cell | 66 |
| max counts / cell | 990 |
| overdispersion | 0.89 |

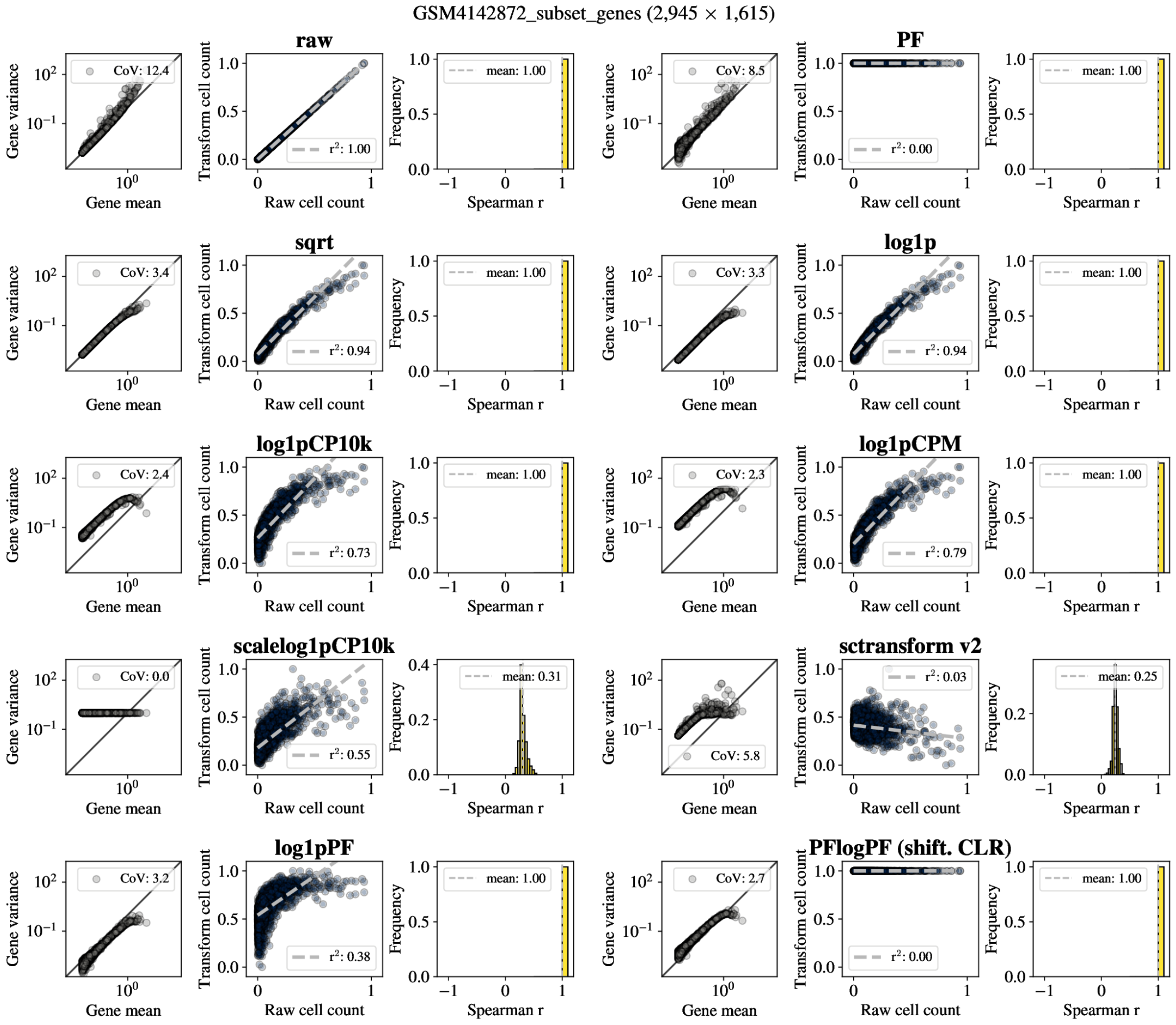


Supplementary Figure 1.484. **GSM4142872**. Per-method comparison of variance stabilization (left), depth normalization (middle), and monotonicity (right) for each normalization on dataset GSM4142872.

| GSM4142873 | |
|---|---|
| cells | 151 |
| genes | 60,623 |
| nonzero entries | 341,578 |
| density | 0.0373 |
| total counts | 1,405,673 |
| mean counts / cell | 9,309 |
| mean counts / gene | 23 |
| min counts / cell | 651 |
| max counts / cell | 68,143 |
| overdispersion | 11.5 |

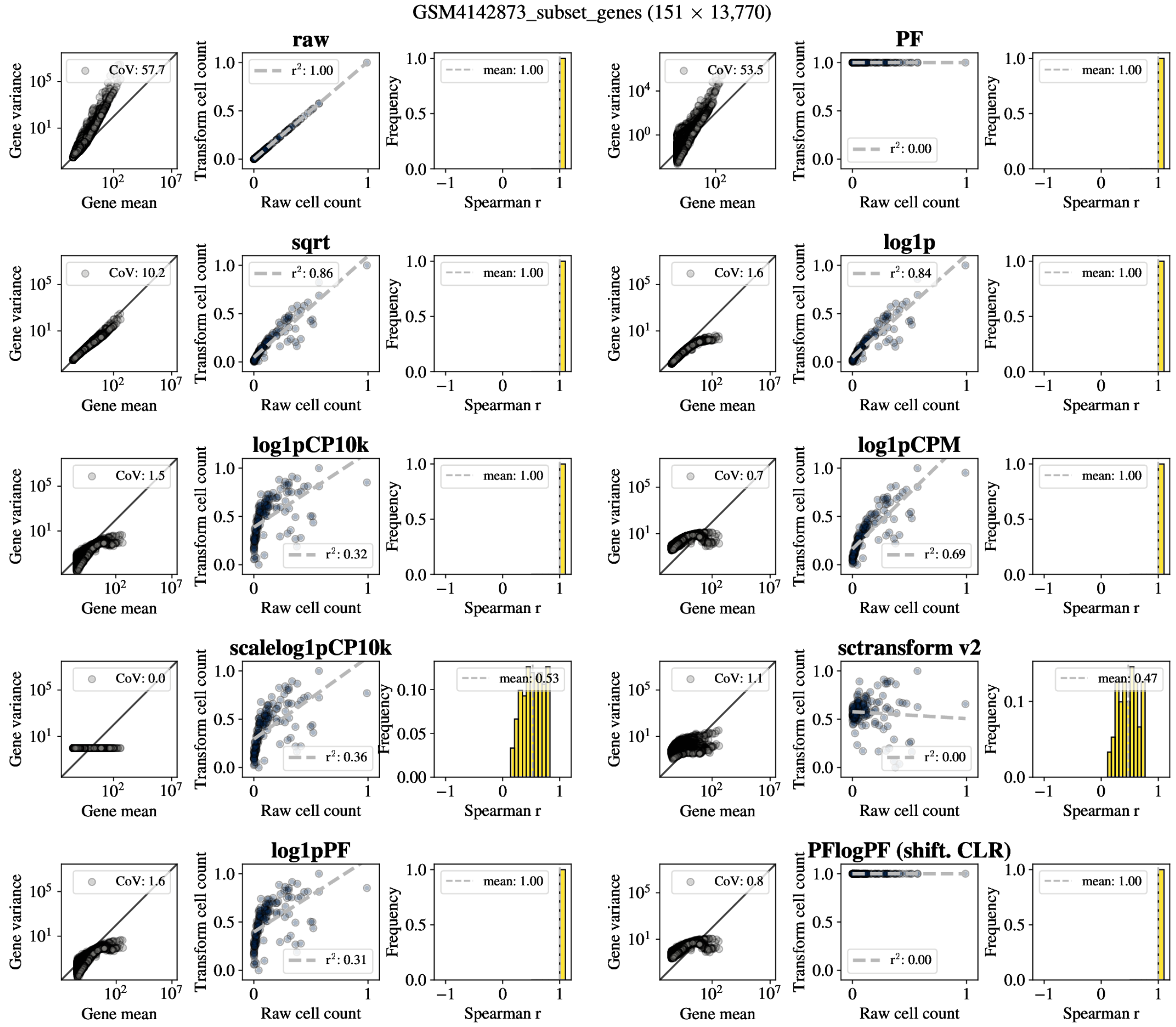


Supplementary Figure 1.485. **GSM4142873**. Per-method comparison of variance stabilization (left), depth normalization (middle), and monotonicity (right) for each normalization on dataset GSM4142873.

| GSM4145204 | |
|---|---|
| cells | 17,365 |
| genes | 60,623 |
| nonzero entries | 10,254,602 |
| density | 0.00974 |
| total counts | 27,484,511 |
| mean counts / cell | 1,583 |
| mean counts / gene | 453 |
| min counts / cell | 266 |
| max counts / cell | 28,573 |
| overdispersion | 0.988 |

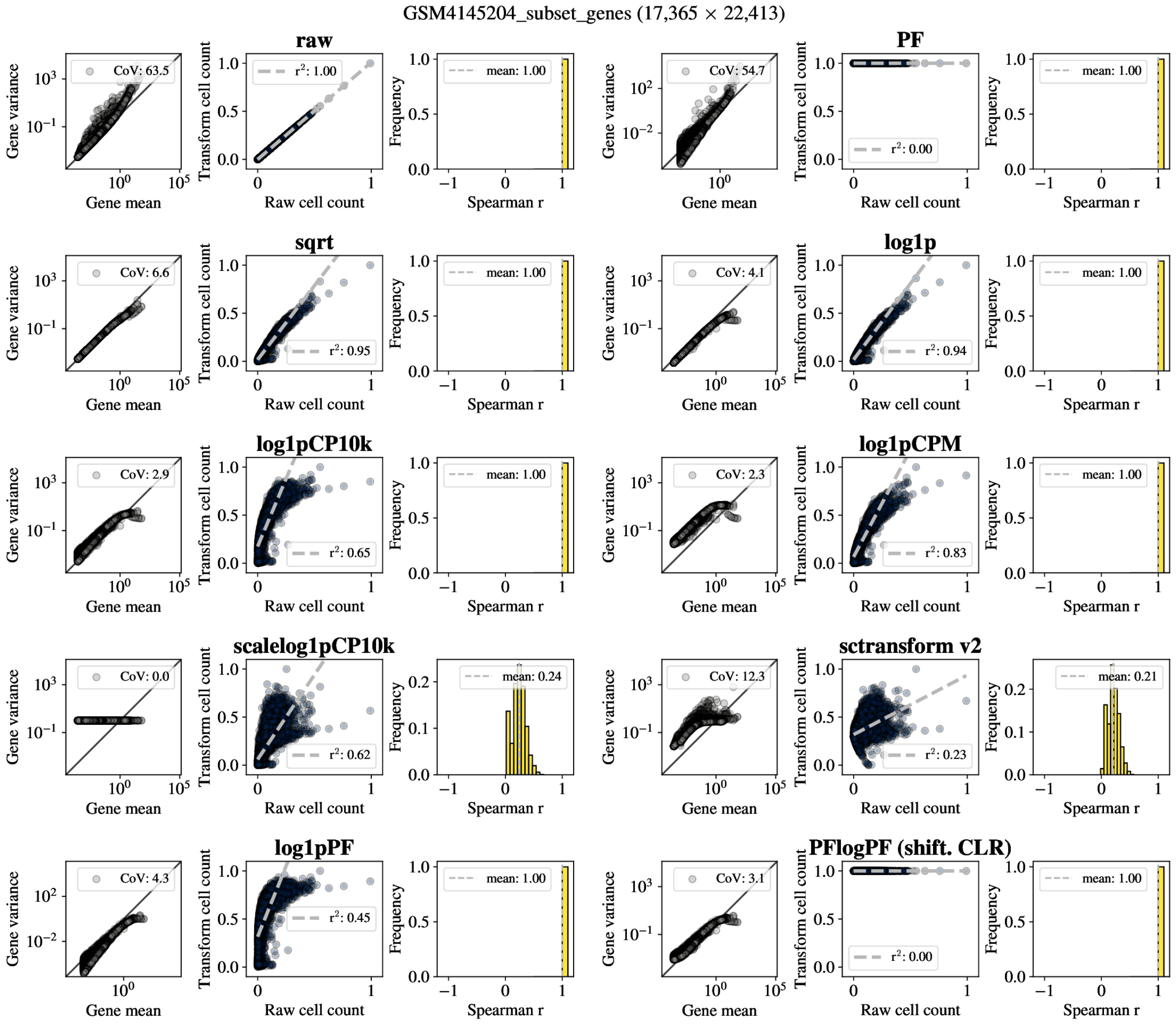


Supplementary Figure 1.486. **GSM4145204**. Per-method comparison of variance stabilization (left), depth normalization (middle), and monotonicity (right) for each normalization on dataset GSM4145204.

| GSM4272895 | |
|---|---|
| cells | 121 |
| genes | 60,623 |
| nonzero entries | 46,560 |
| density | 0.00635 |
| total counts | 118,691 |
| mean counts / cell | 981 |
| mean counts / gene | 2 |
| min counts / cell | 245 |
| max counts / cell | 16,551 |
| overdispersion | 1.52 |

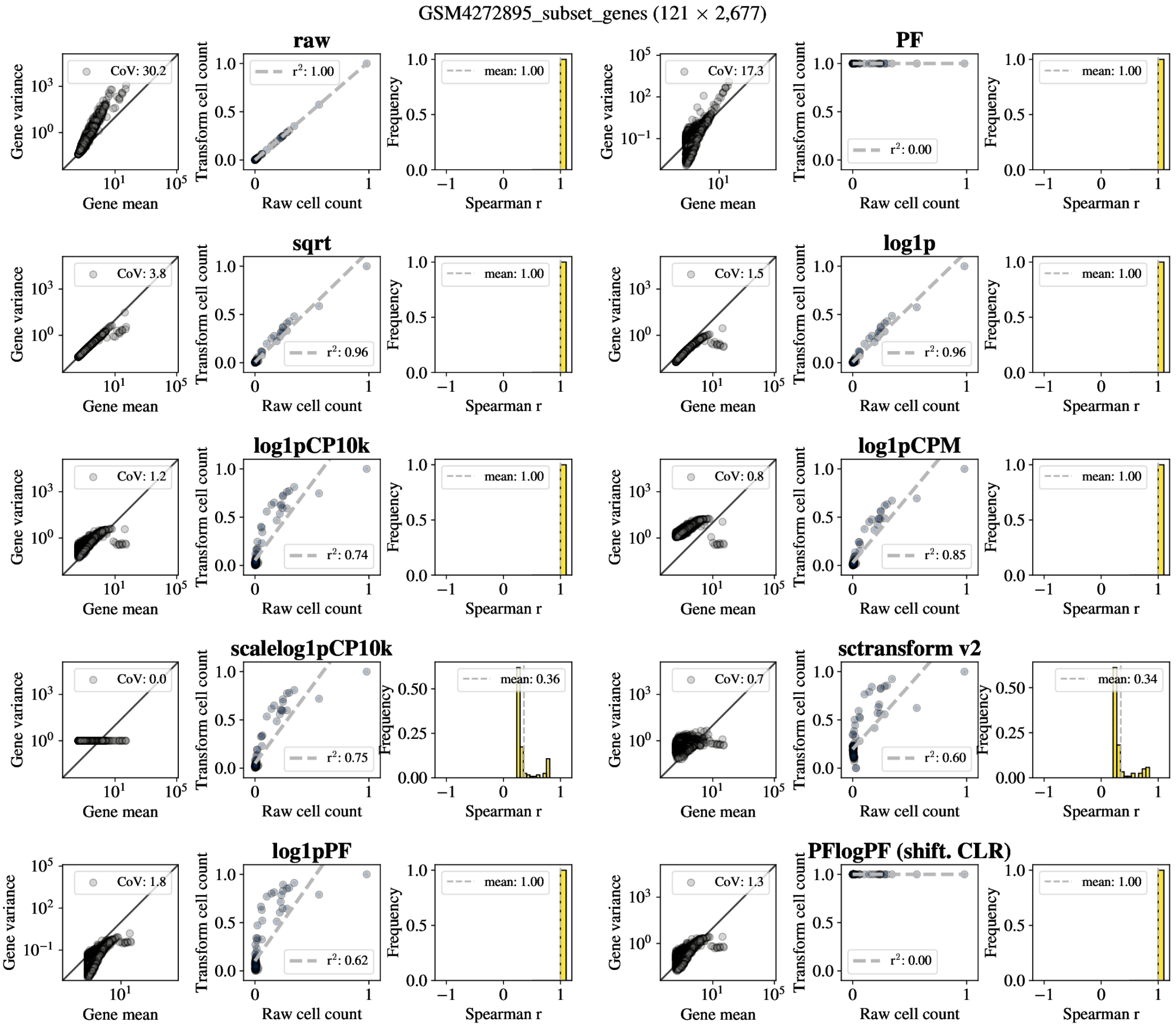


Supplementary Figure 1.487. **GSM4272895**. Per-method comparison of variance stabilization (left), depth normalization (middle), and monotonicity (right) for each normalization on dataset GSM4272895.

| GSM4272896 | |
|---|---|
| cells | 27 |
| genes | 60,623 |
| nonzero entries | 6,108 |
| density | 0.00373 |
| total counts | 21,099 |
| mean counts / cell | 781 |
| mean counts / gene | 0 |
| min counts / cell | 79 |
| max counts / cell | 7,945 |
| overdispersion | 8.52 |

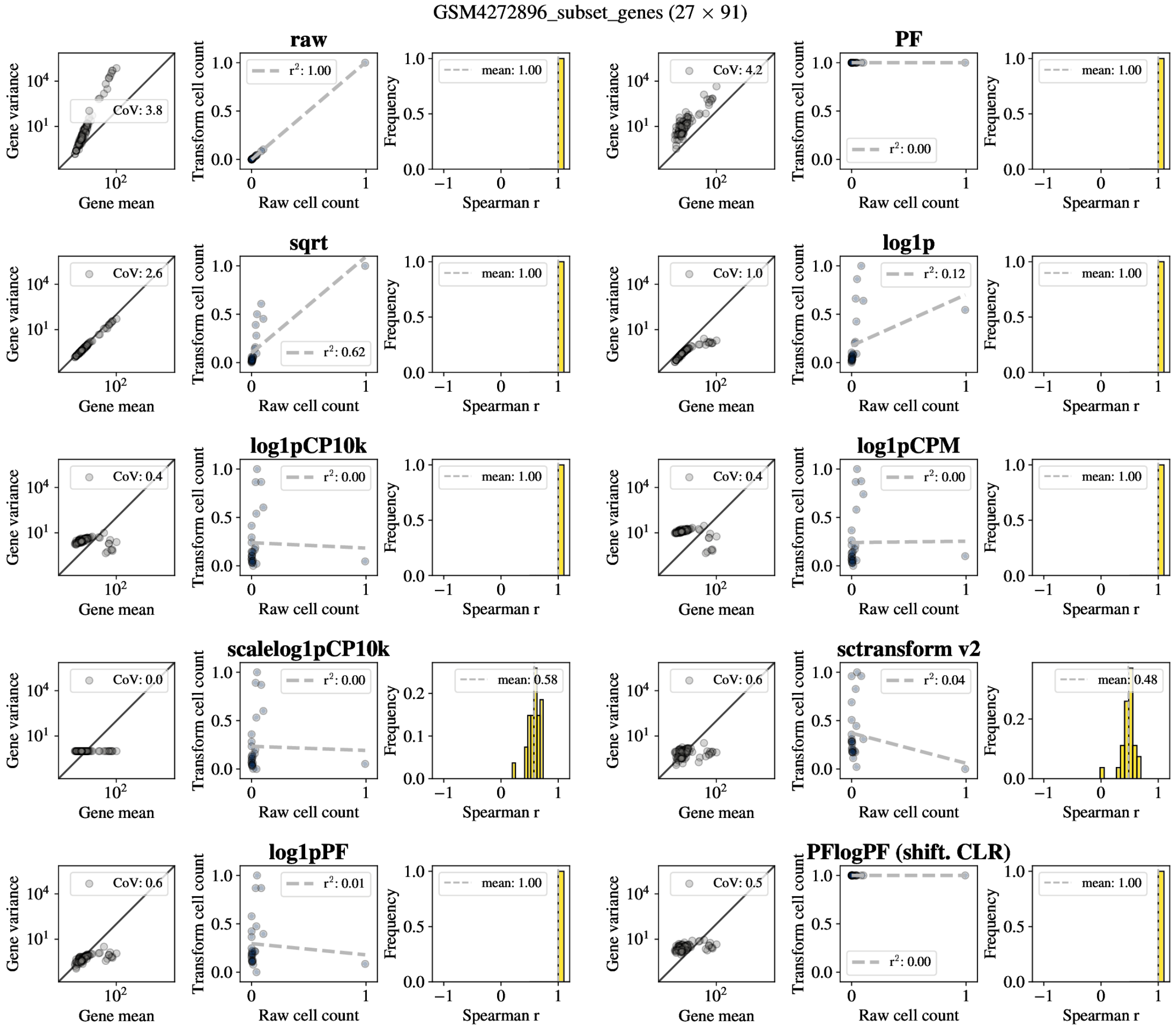


Supplementary Figure 1.488. **GSM4272896**. Per-method comparison of variance stabilization (left), depth normalization (middle), and monotonicity (right) for each normalization on dataset GSM4272896.

| GSM4272897 | |
|---|---|
| cells | 36 |
| genes | 60,623 |
| nonzero entries | 46,774 |
| density | 0.0214 |
| total counts | 157,942 |
| mean counts / cell | 4,387 |
| mean counts / gene | 3 |
| min counts / cell | 455 |
| max counts / cell | 22,548 |
| overdispersion | 5.78 |

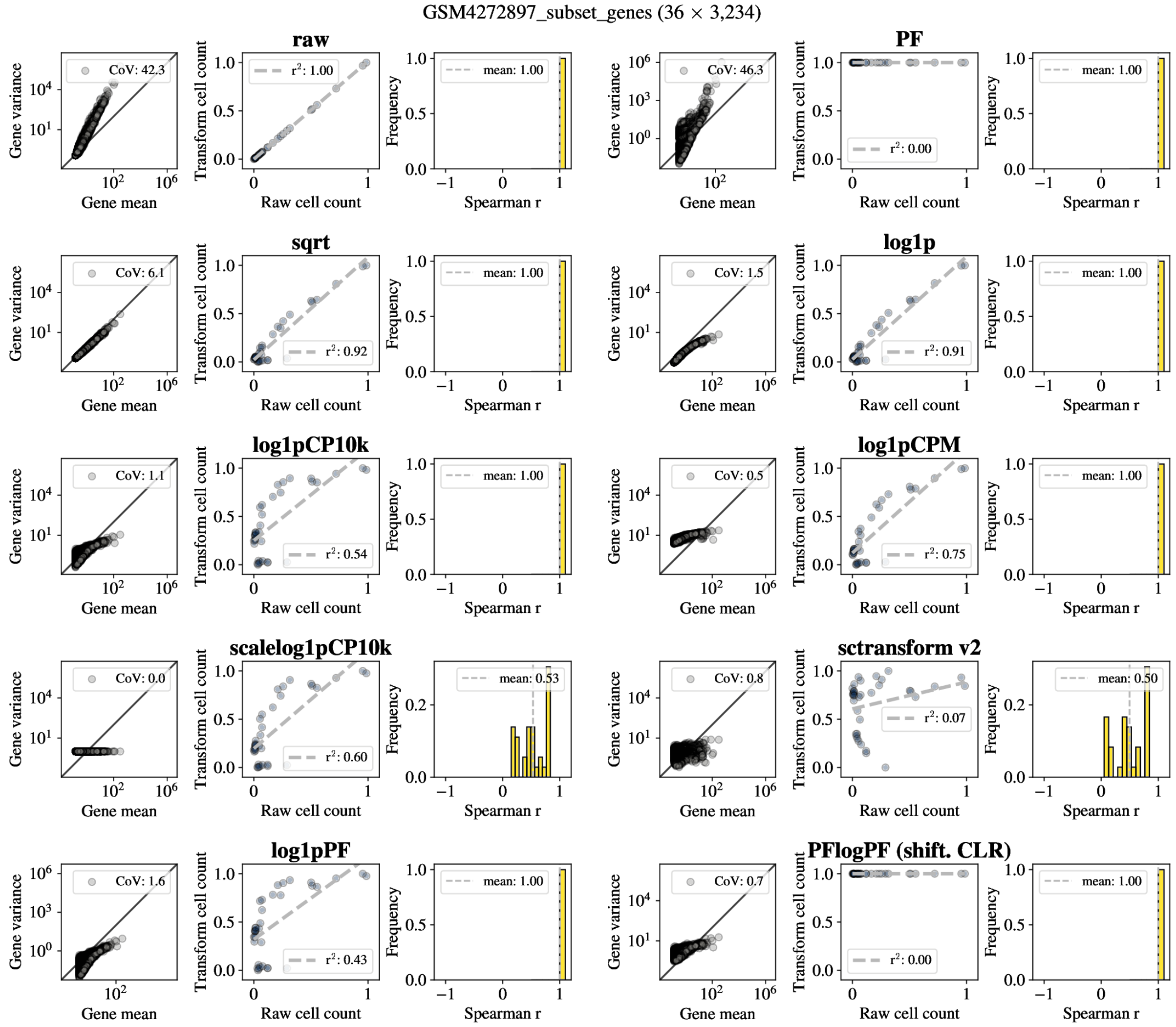


Supplementary Figure 1.489. **GSM4272897**. Per-method comparison of variance stabilization (left), depth normalization (middle), and monotonicity (right) for each normalization on dataset GSM4272897.

| GSM4272898 | |
|---|---|
| cells | 6,538 |
| genes | 60,623 |
| nonzero entries | 5,867,441 |
| density | 0.0148 |
| total counts | 13,427,407 |
| mean counts / cell | 2,054 |
| mean counts / gene | 221 |
| min counts / cell | 360 |
| max counts / cell | 28,776 |
| overdispersion | 0.775 |

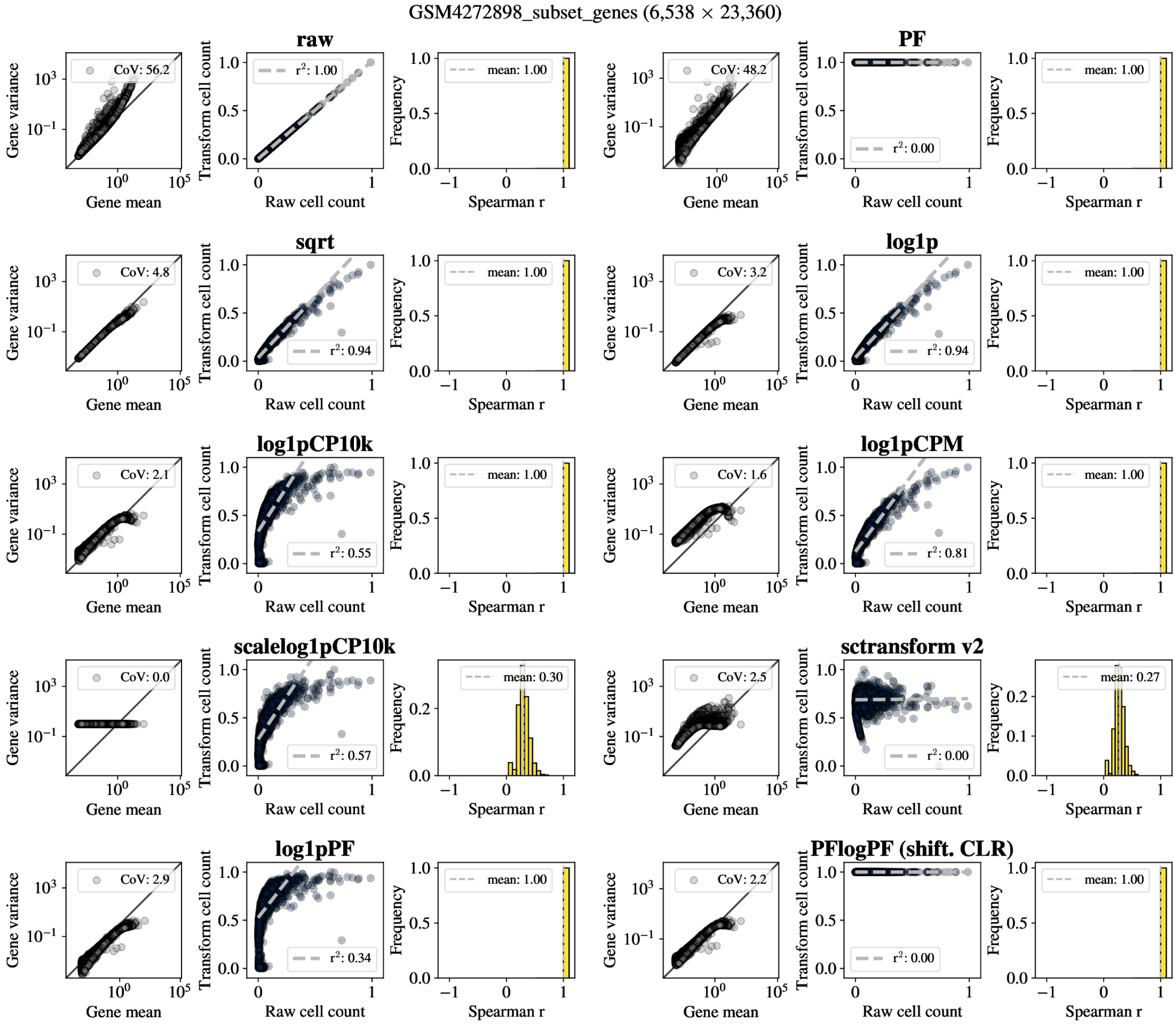


Supplementary Figure 1.490. **GSM4272898**. Per-method comparison of variance stabilization (left), depth normalization (middle), and monotonicity (right) for each normalization on dataset GSM4272898.

| GSM4272899 | |
|---|---|
| cells | 3,725 |
| genes | 60,623 |
| nonzero entries | 5,362,338 |
| density | 0.0237 |
| total counts | 18,042,317 |
| mean counts / cell | 4,844 |
| mean counts / gene | 298 |
| min counts / cell | 316 |
| max counts / cell | 61,529 |
| overdispersion | 0.993 |

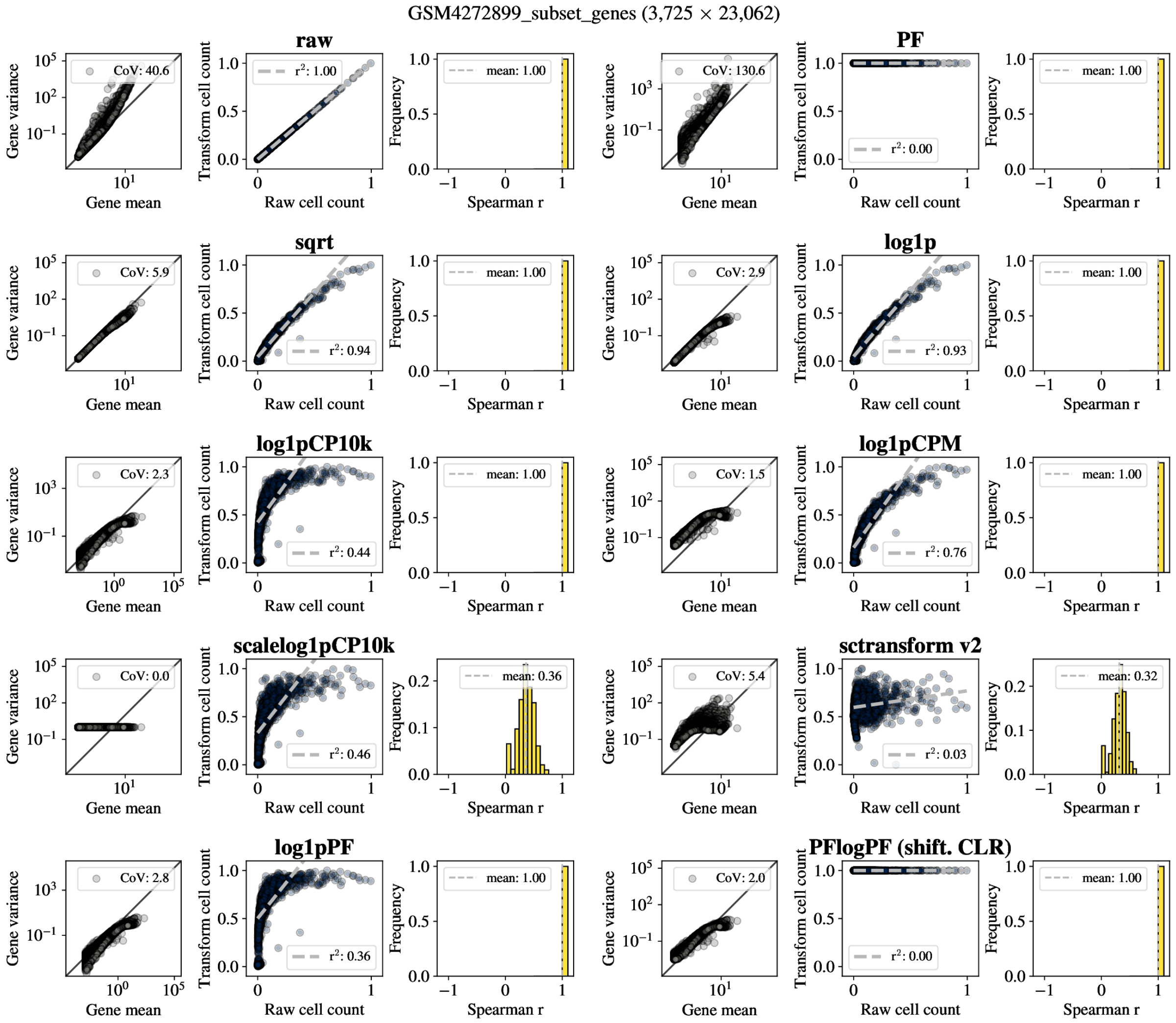


Supplementary Figure 1.491. **GSM4272899**. Per-method comparison of variance stabilization (left), depth normalization (middle), and monotonicity (right) for each normalization on dataset GSM4272899.

| GSM4272900 | |
|---|---|
| cells | 5,819 |
| genes | 60,623 |
| nonzero entries | 6,539,193 |
| density | 0.0185 |
| total counts | 17,484,232 |
| mean counts / cell | 3,005 |
| mean counts / gene | 288 |
| min counts / cell | 330 |
| max counts / cell | 57,730 |
| overdispersion | 0.856 |

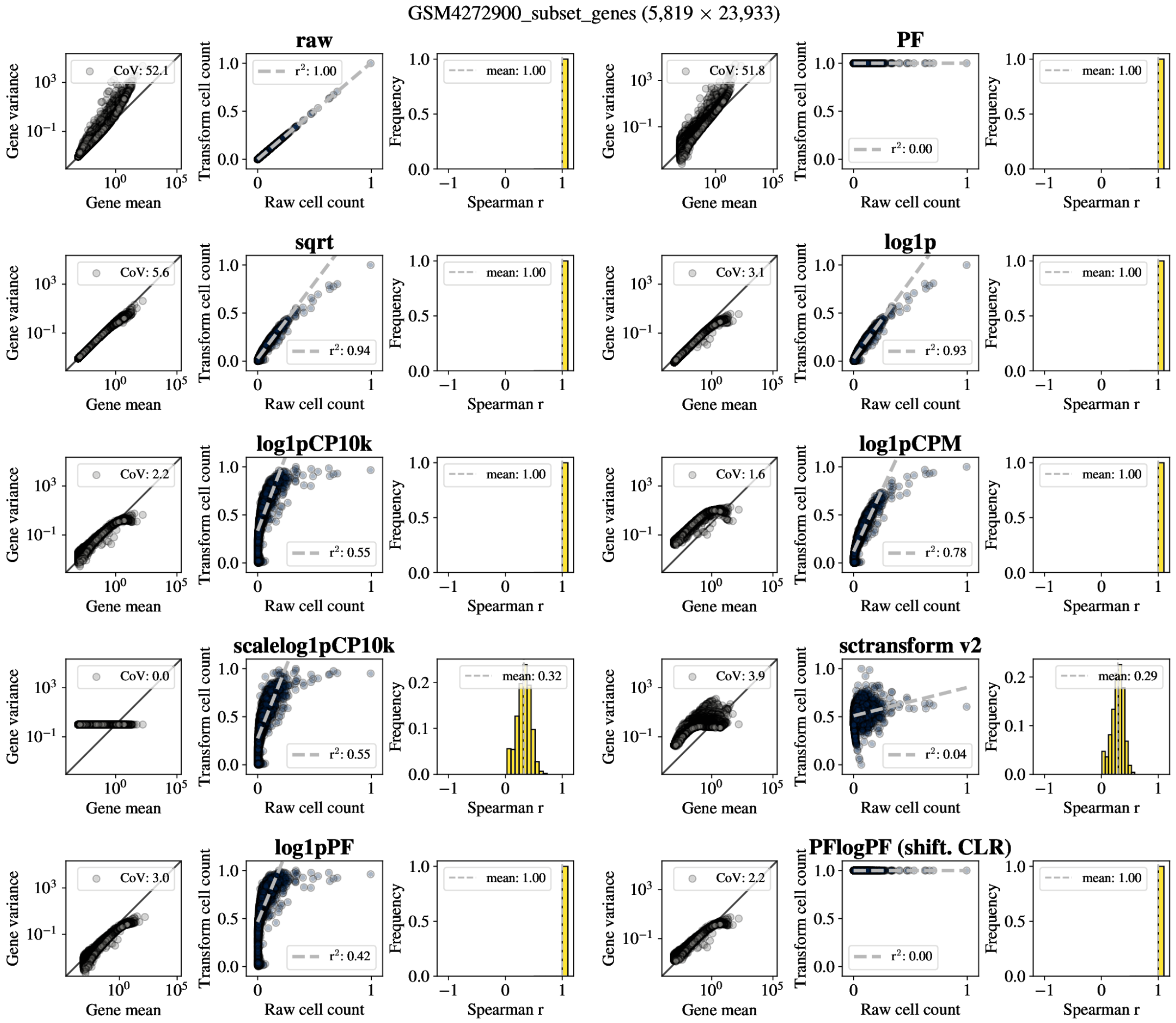


Supplementary Figure 1.492. **GSM4272900**. Per-method comparison of variance stabilization (left), depth normalization (middle), and monotonicity (right) for each normalization on dataset GSM4272900.

| GSM4272901 | |
|---|---|
| cells | 346 |
| genes | 60,623 |
| nonzero entries | 210,318 |
| density | 0.01 |
| total counts | 618,505 |
| mean counts / cell | 1,788 |
| mean counts / gene | 10 |
| min counts / cell | 597 |
| max counts / cell | 11,274 |
| overdispersion | 0.291 |

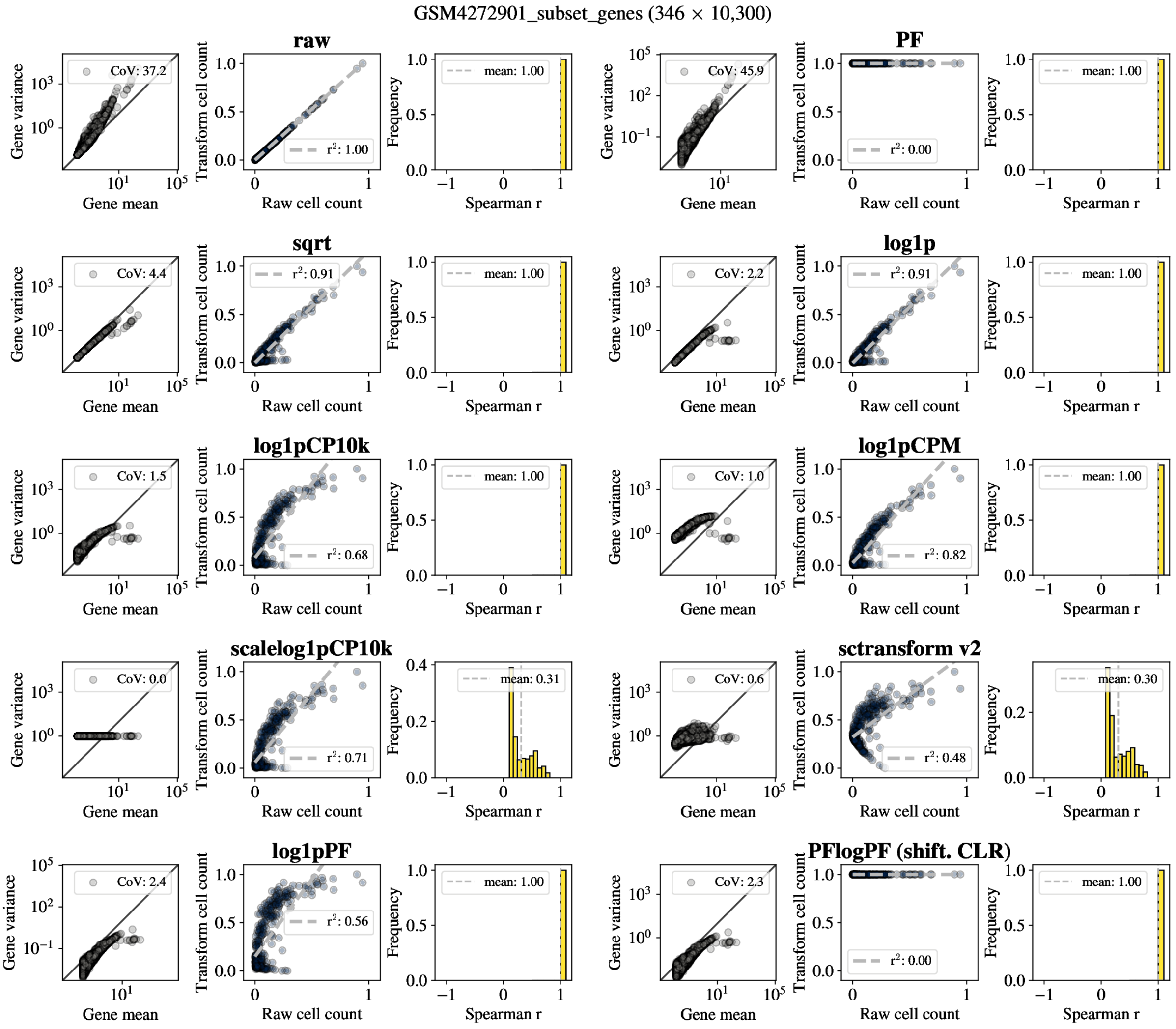


Supplementary Figure 1.493. **GSM4272901**. Per-method comparison of variance stabilization (left), depth normalization (middle), and monotonicity (right) for each normalization on dataset GSM4272901.

| GSM4272902 | |
|---|---|
| cells | 127 |
| genes | 60,623 |
| nonzero entries | 235,215 |
| density | 0.0306 |
| total counts | 552,225 |
| mean counts / cell | 4,348 |
| mean counts / gene | 9 |
| min counts / cell | 622 |
| max counts / cell | 21,924 |
| overdispersion | 0.579 |

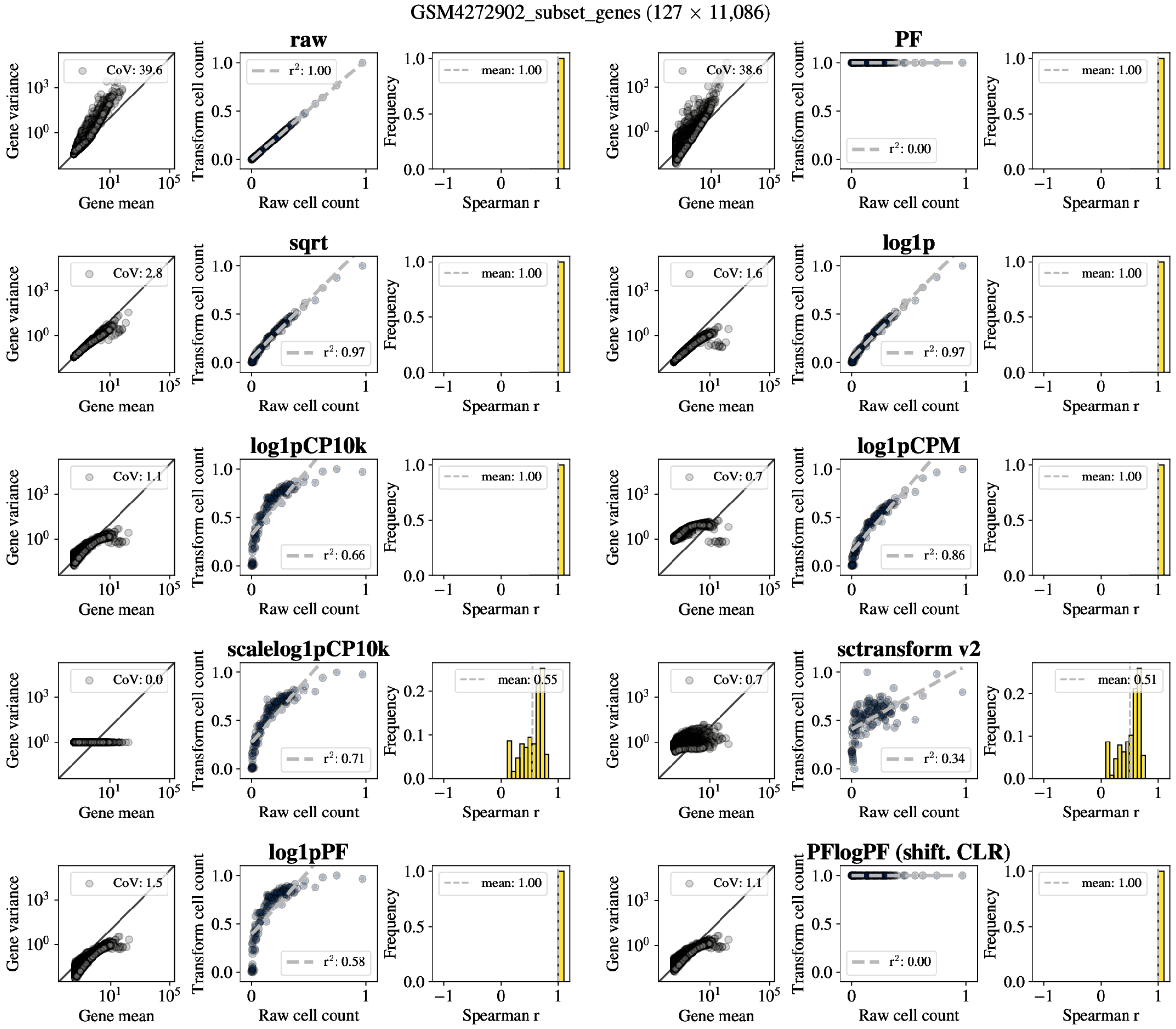


Supplementary Figure 1.494. **GSM4272902**. Per-method comparison of variance stabilization (left), depth normalization (middle), and monotonicity (right) for each normalization on dataset GSM4272902.

| GSM4339769 | |
|---|---|
| cells | 3,750 |
| genes | 60,623 |
| nonzero entries | 3,230,332 |
| density | 0.0142 |
| total counts | 9,904,750 |
| mean counts / cell | 2,641 |
| mean counts / gene | 163 |
| min counts / cell | 24 |
| max counts / cell | 18,811 |
| overdispersion | 2.38 |

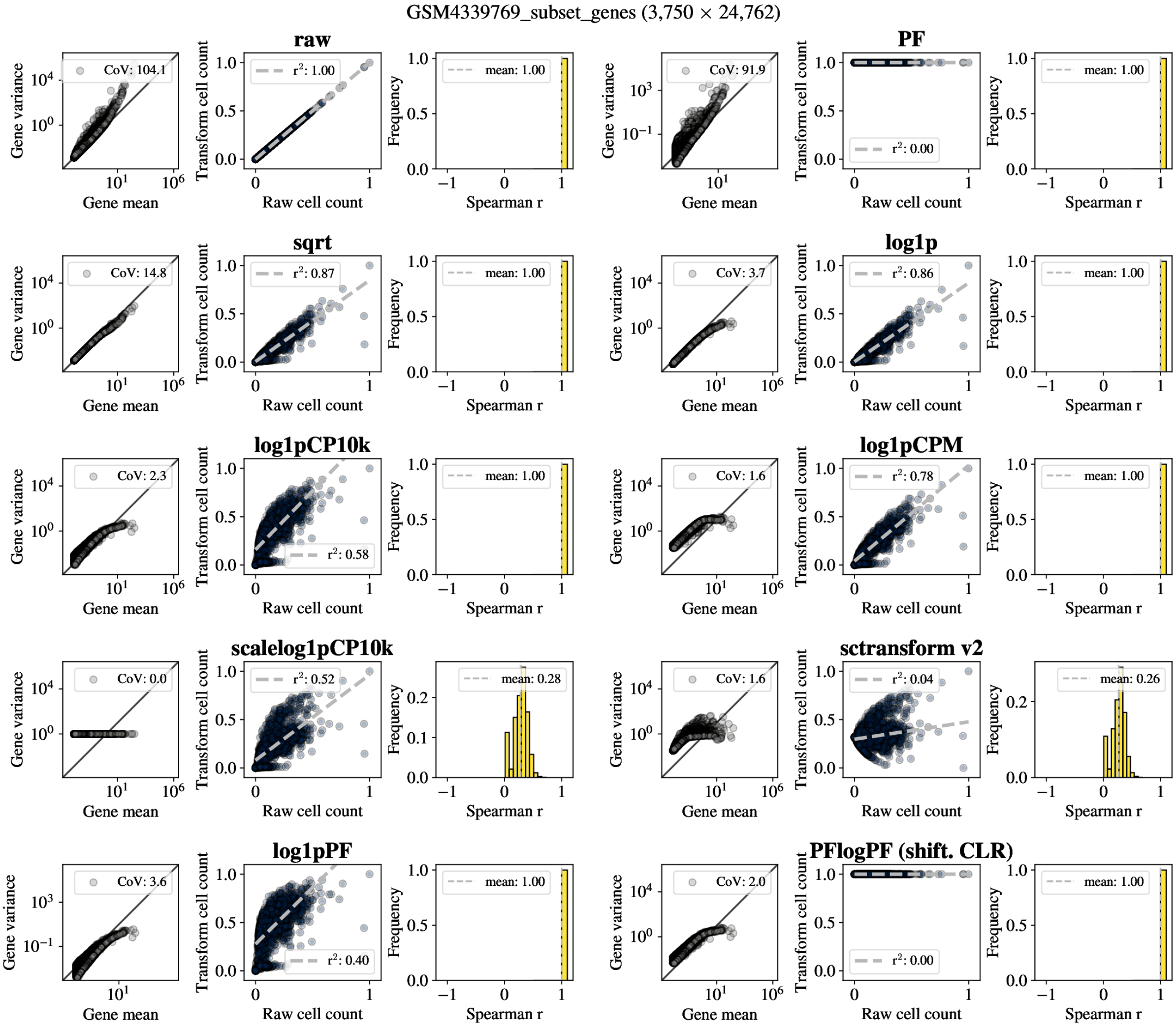


Supplementary Figure 1.495. **GSM4339769**. Per-method comparison of variance stabilization (left), depth normalization (middle), and monotonicity (right) for each normalization on dataset GSM4339769.

| GSM4463223 | |
|---|---|
| cells | 7,310 |
| genes | 60,623 |
| nonzero entries | 3,215,208 |
| density | 0.00726 |
| total counts | 9,600,715 |
| mean counts / cell | 1,313 |
| mean counts / gene | 158 |
| min counts / cell | 96 |
| max counts / cell | 9,511 |
| overdispersion | 1 |

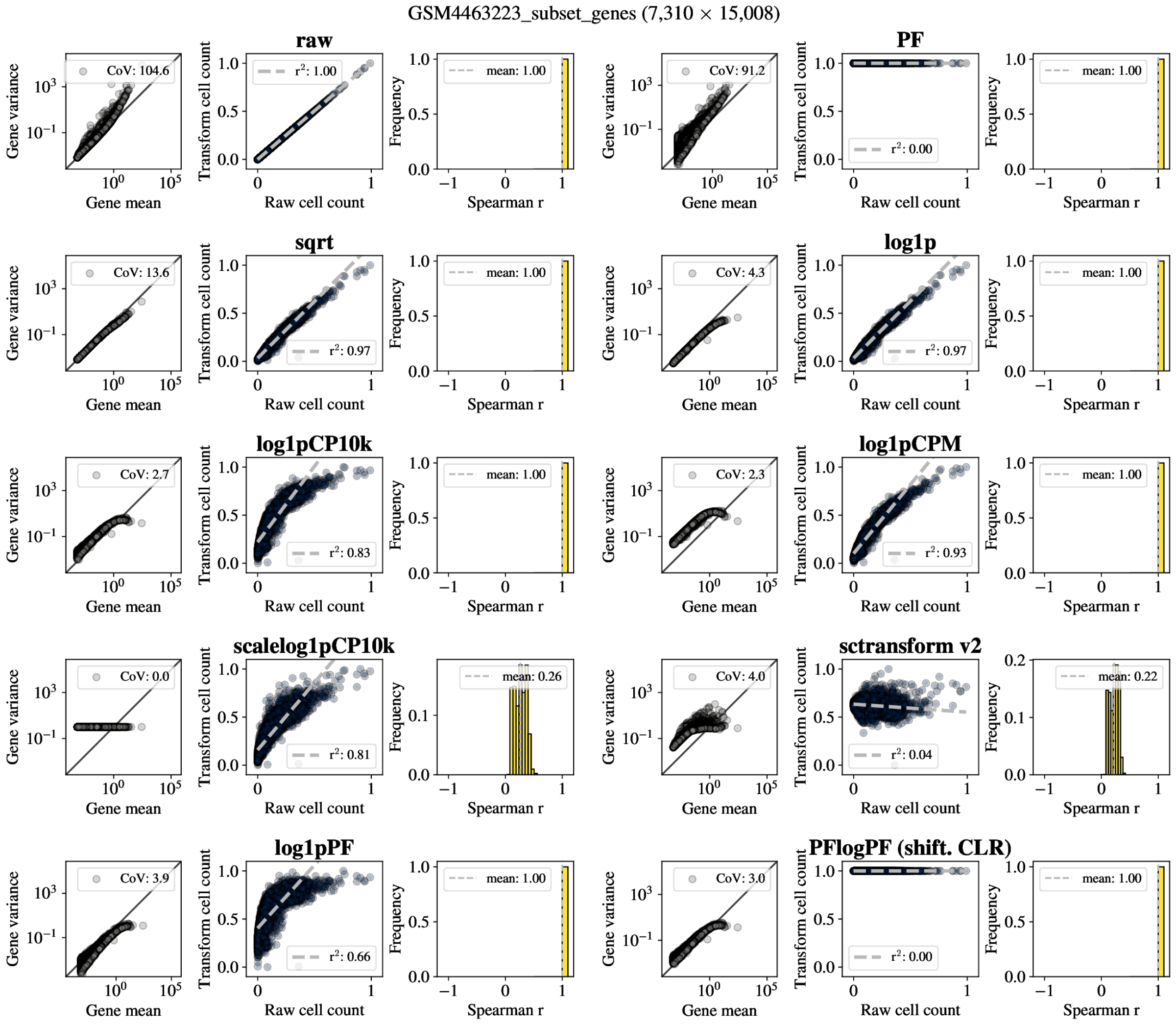


Supplementary Figure 1.496. **GSM4463223**. Per-method comparison of variance stabilization (left), depth normalization (middle), and monotonicity (right) for each normalization on dataset GSM4463223.

| GSM4463229 | |
|---|---|
| cells | 4,457 |
| genes | 60,623 |
| nonzero entries | 4,230,386 |
| density | 0.0157 |
| total counts | 13,272,672 |
| mean counts / cell | 2,978 |
| mean counts / gene | 219 |
| min counts / cell | 124 |
| max counts / cell | 72,563 |
| overdispersion | 3.82 |

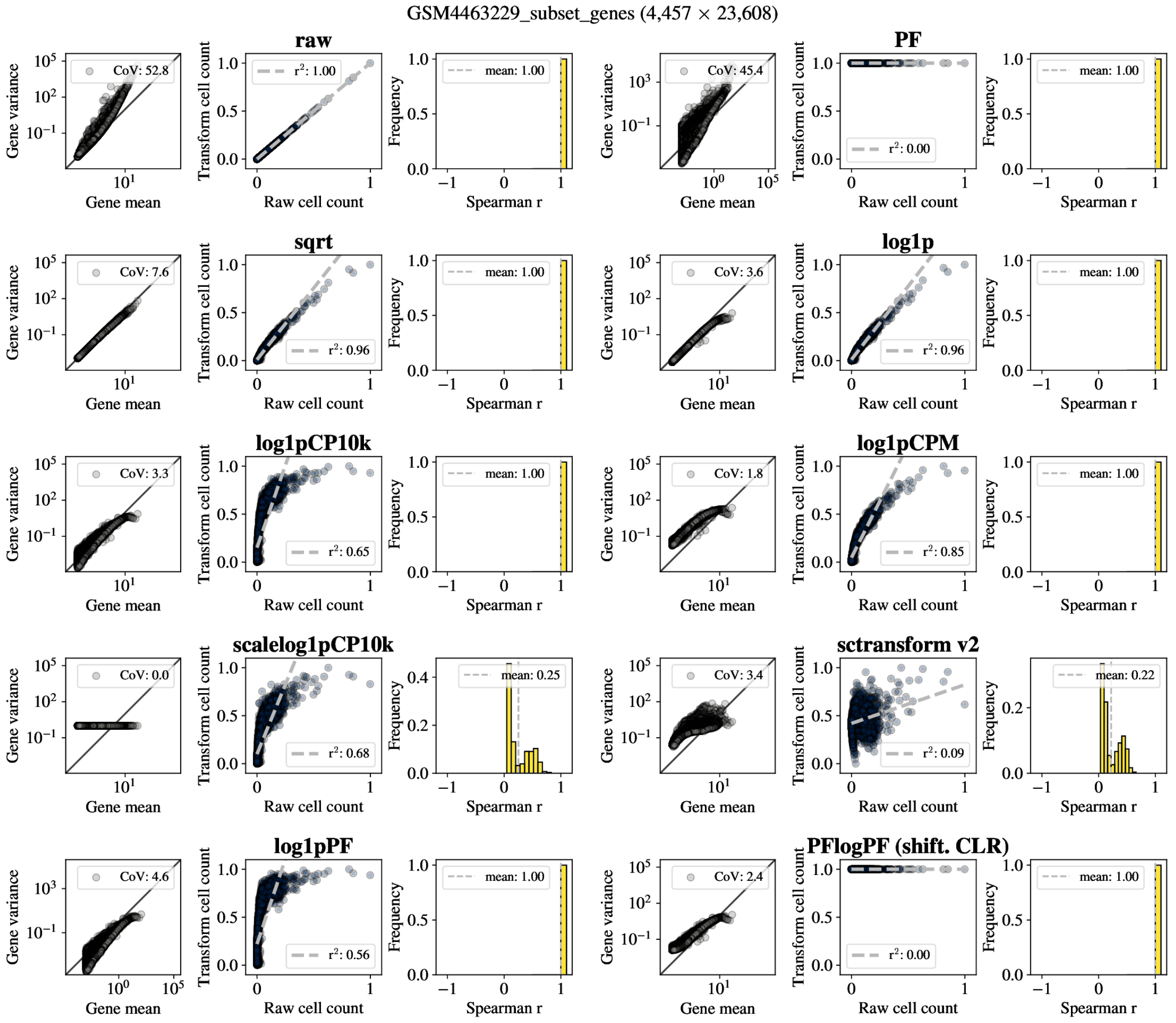


Supplementary Figure 1.497. **GSM4463229**. Per-method comparison of variance stabilization (left), depth normalization (middle), and monotonicity (right) for each normalization on dataset GSM4463229.

| GSM4463234 | |
|---|---|
| cells | 3,783 |
| genes | 60,623 |
| nonzero entries | 4,165,463 |
| density | 0.0182 |
| total counts | 13,104,032 |
| mean counts / cell | 3,464 |
| mean counts / gene | 216 |
| min counts / cell | 126 |
| max counts / cell | 68,097 |
| overdispersion | 3.46 |

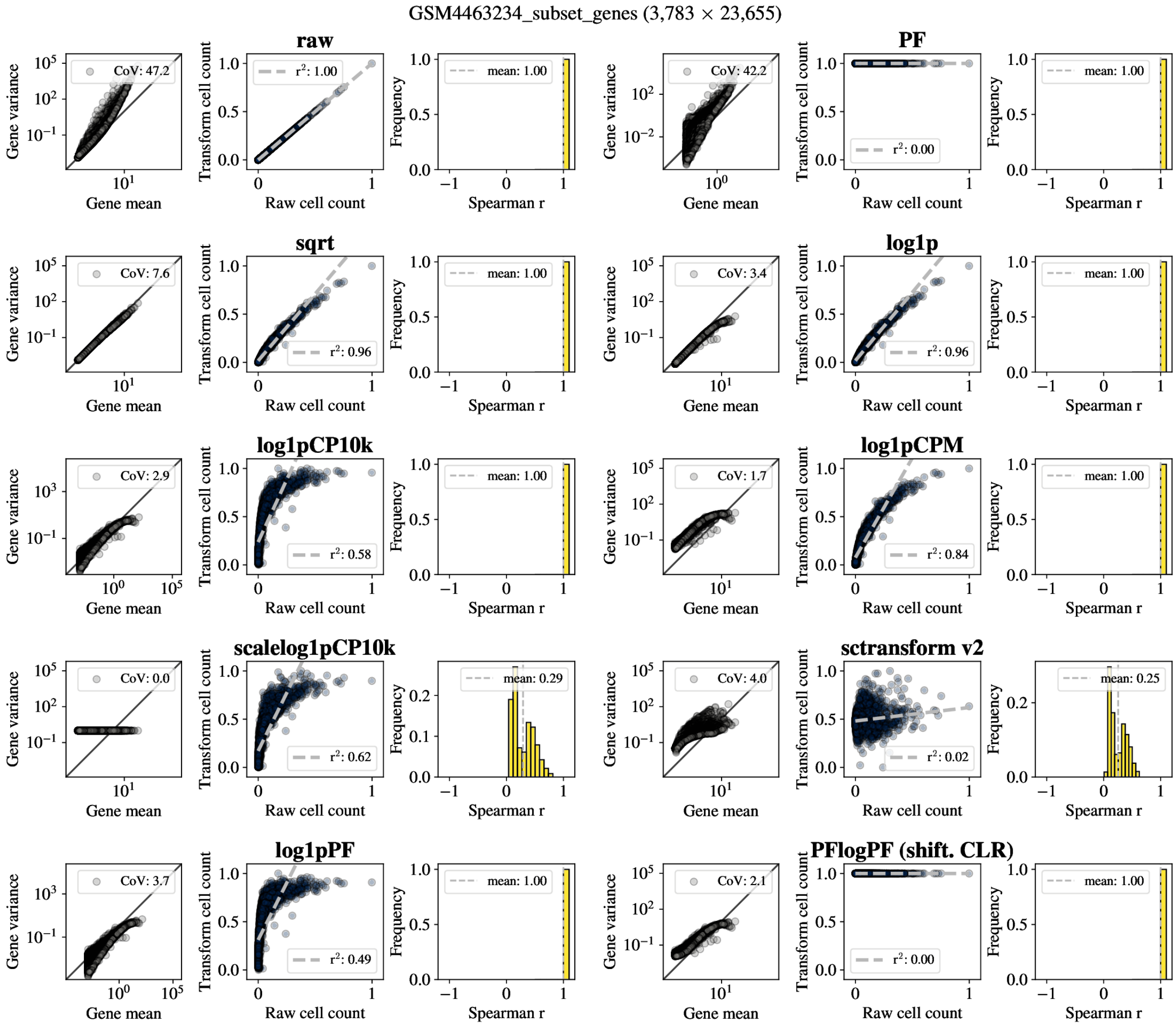


Supplementary Figure 1.498. **GSM4463234**. Per-method comparison of variance stabilization (left), depth normalization (middle), and monotonicity (right) for each normalization on dataset GSM4463234.

| **GSM4463239** | |
|---|---|
| cells | 5,312 |
| genes | 60,623 |
| nonzero entries | 3,751,982 |
| density | 0.0117 |
| total counts | 10,164,502 |
| mean counts / cell | 1,913 |
| mean counts / gene | 168 |
| min counts / cell | 350 |
| max counts / cell | 30,793 |
| overdispersion | 1.31 |

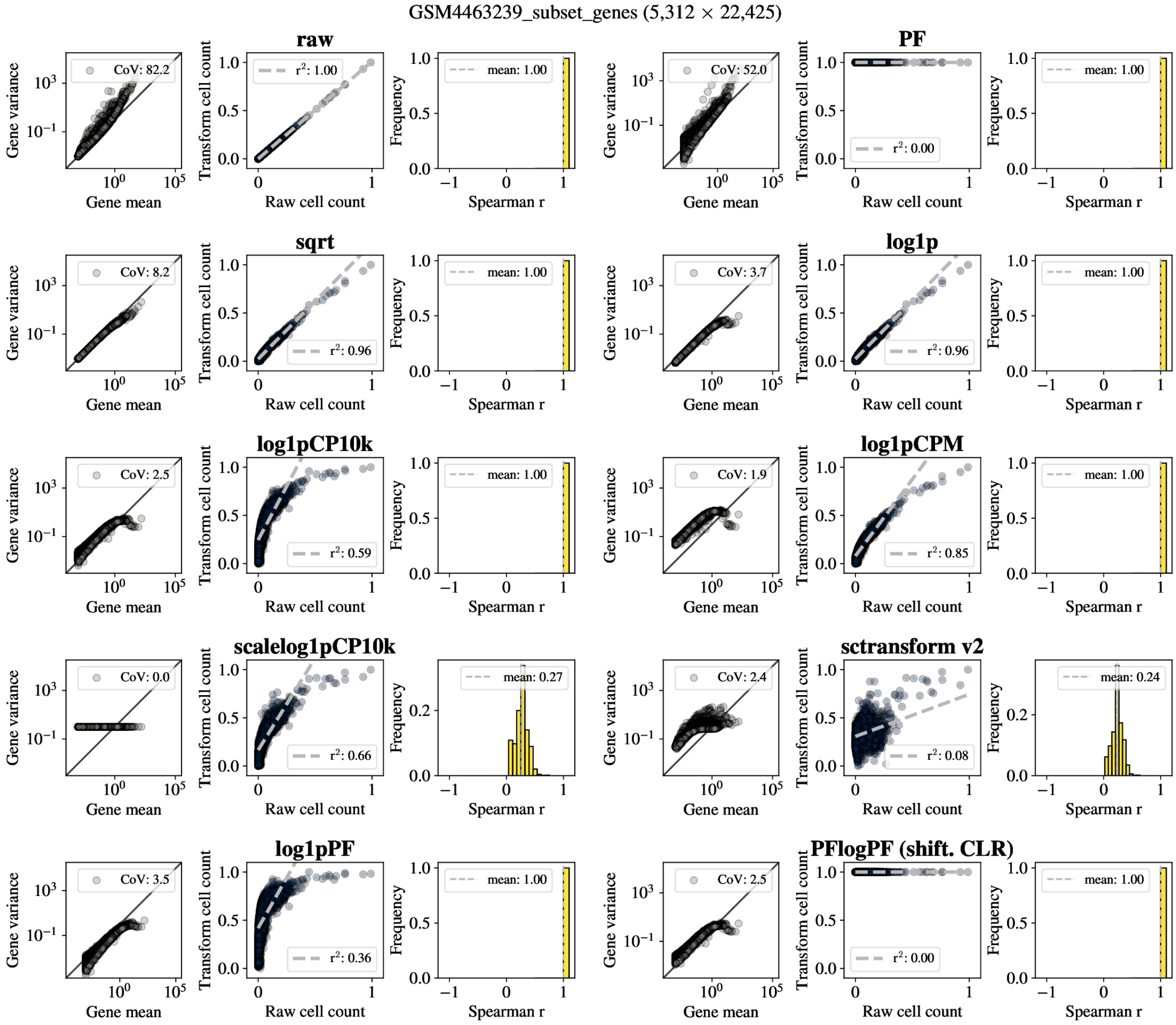


Supplementary Figure 1.499. **GSM4463239**. Per-method comparison of variance stabilization (left), depth normalization (middle), and monotonicity (right) for each normalization on dataset GSM4463239.

| GSM4463244 | |
|---|---|
| cells | 4,618 |
| genes | 60,623 |
| nonzero entries | 5,223,991 |
| density | 0.0187 |
| total counts | 17,104,043 |
| mean counts / cell | 3,704 |
| mean counts / gene | 282 |
| min counts / cell | 294 |
| max counts / cell | 55,007 |
| overdispersion | 2.62 |

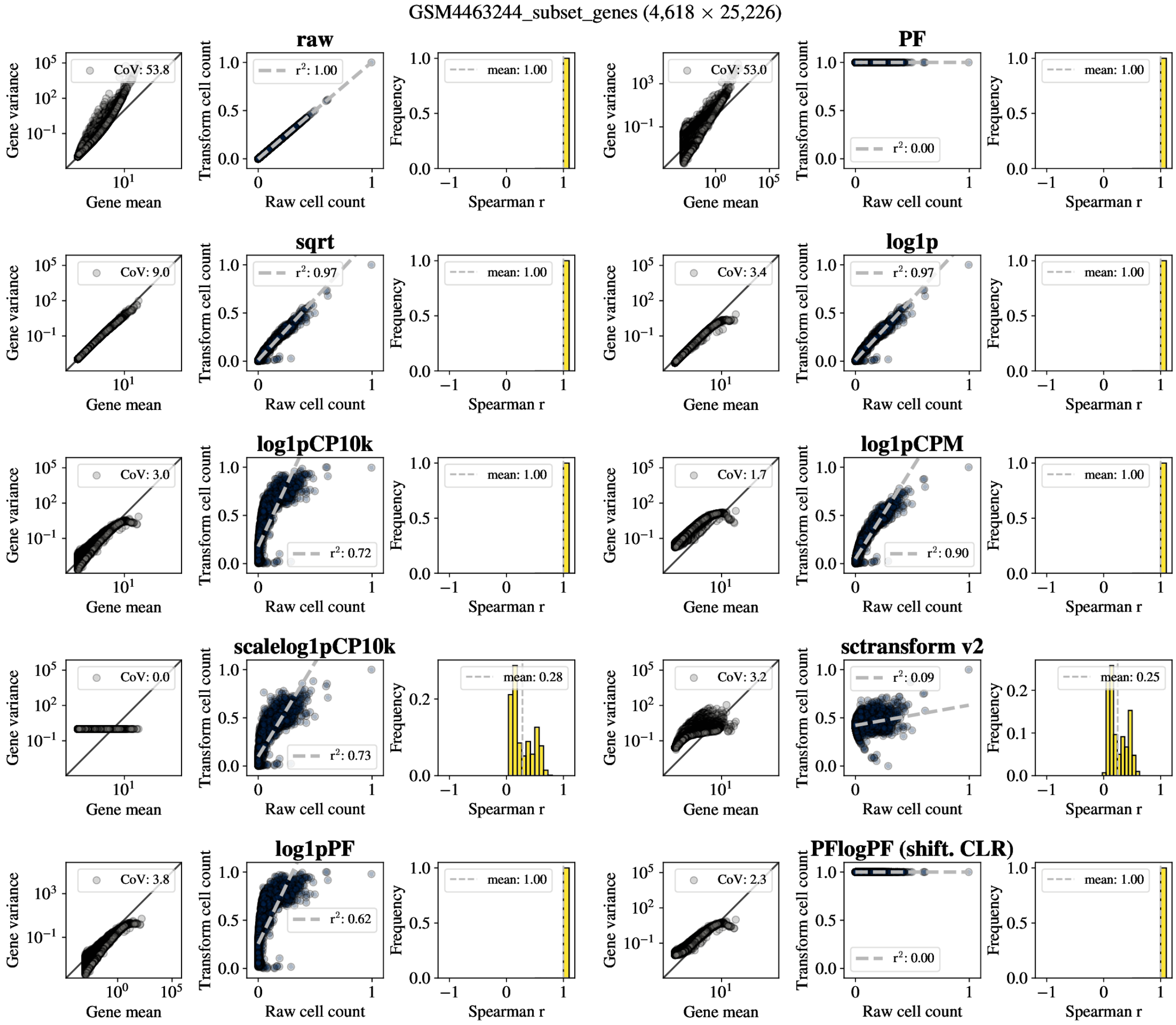


Supplementary Figure 1.500. **GSM4463244**. Per-method comparison of variance stabilization (left), depth normalization (middle), and monotonicity (right) for each normalization on dataset GSM4463244.

| GSM4463249 | |
|---|---|
| cells | 4,780 |
| genes | 60,623 |
| nonzero entries | 4,520,008 |
| density | 0.0156 |
| total counts | 10,226,099 |
| mean counts / cell | 2,139 |
| mean counts / gene | 169 |
| min counts / cell | 500 |
| max counts / cell | 37,697 |
| overdispersion | 1.29 |

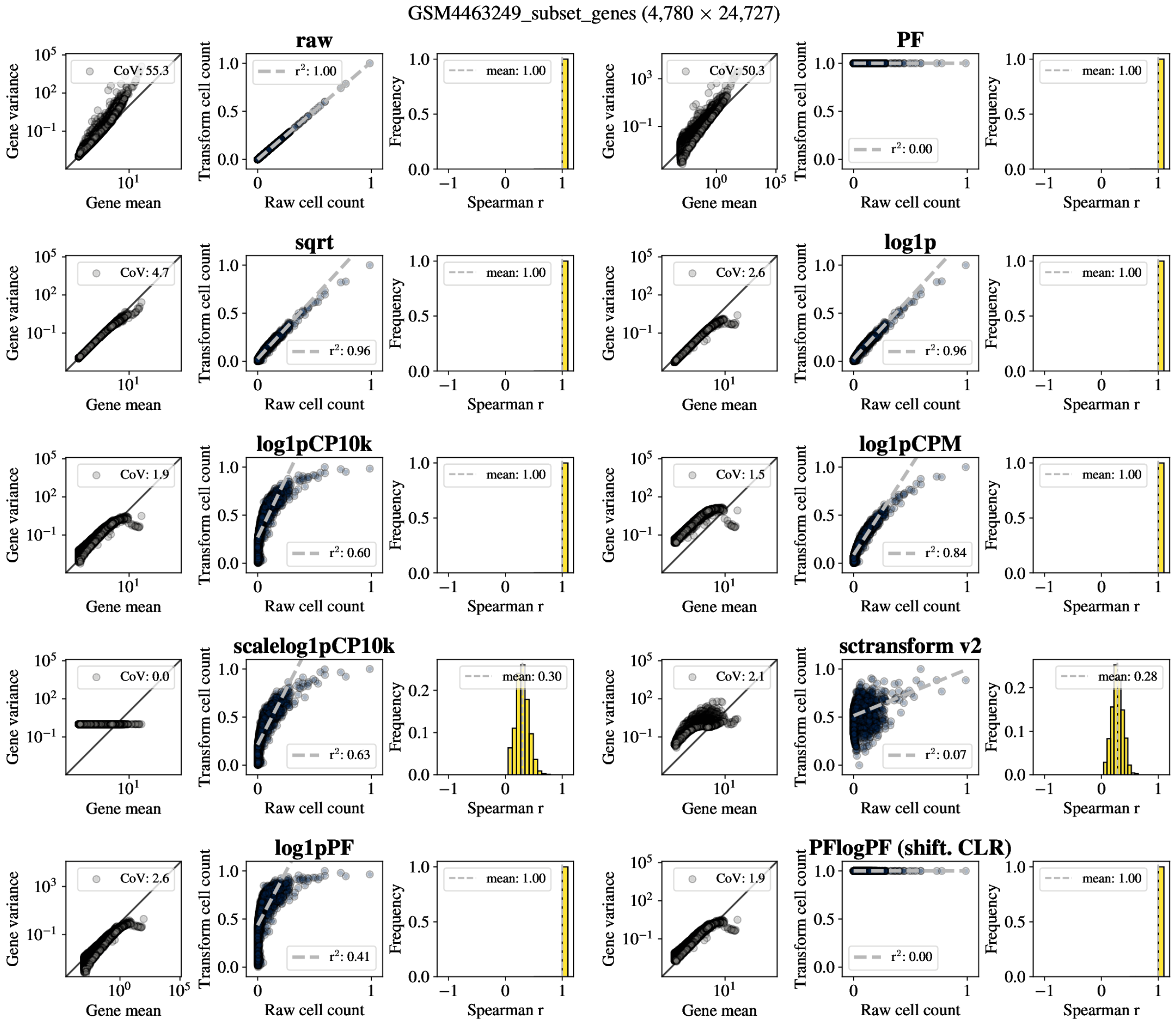

Supplementary Figure 1.501. **GSM4463249**. Per-method comparison of variance stabilization (left), depth normalization (middle), and monotonicity (right) for each normalization on dataset GSM4463249.

| GSM4463253 | |
|---|---|
| cells | 23,013 |
| genes | 60,623 |
| nonzero entries | 22,658,681 |
| density | 0.0162 |
| total counts | 87,984,942 |
| mean counts / cell | 3,823 |
| mean counts / gene | 1,451 |
| min counts / cell | 10 |
| max counts / cell | 47,239 |
| overdispersion | 0.542 |

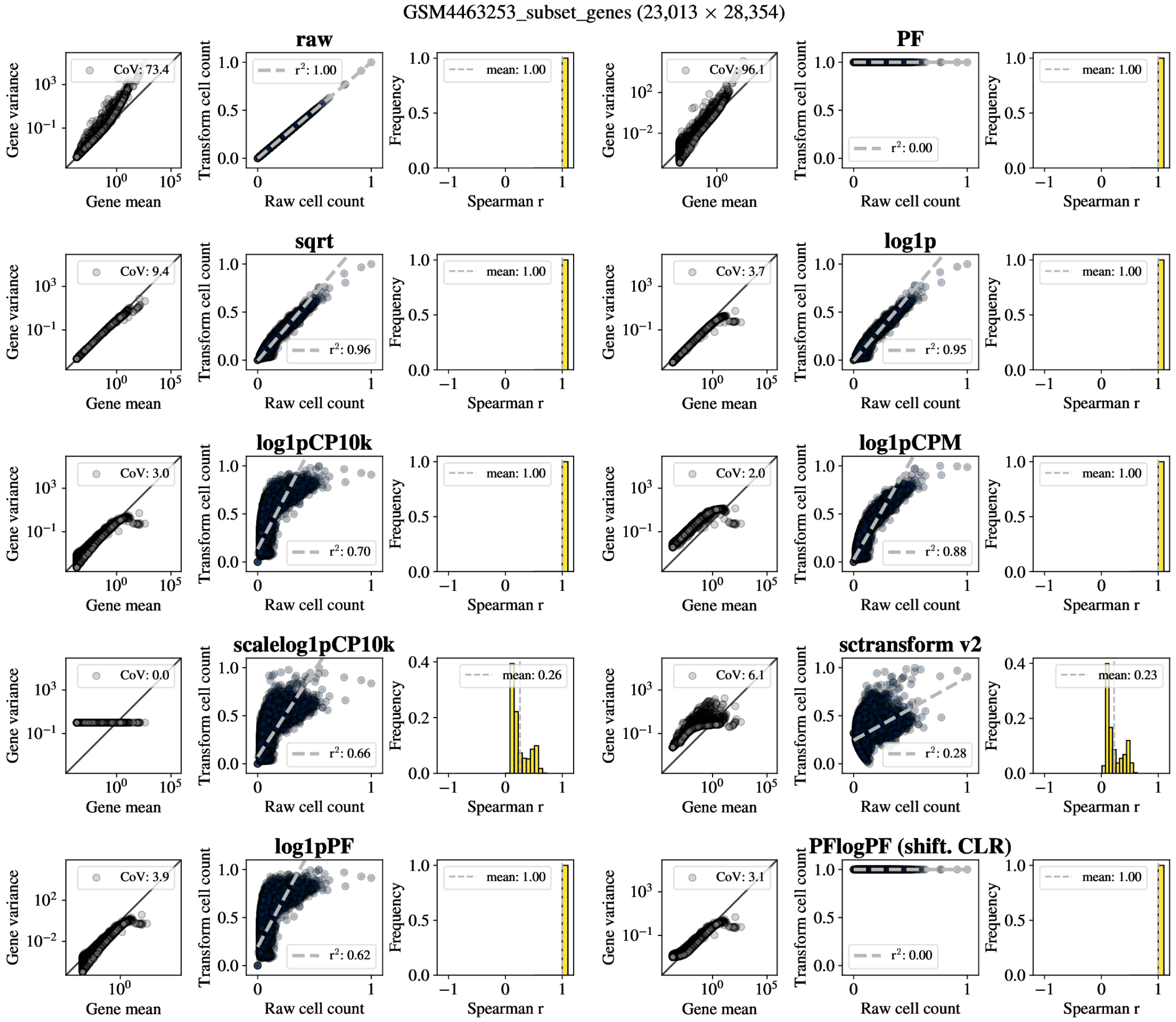


Supplementary Figure 1.502. **GSM4463253**. Per-method comparison of variance stabilization (left), depth normalization (middle), and monotonicity (right) for each normalization on dataset GSM4463253.

| GSM4475048 | |
|---|---|
| cells | 9,653 |
| genes | 60,623 |
| nonzero entries | 14,024,972 |
| density | 0.024 |
| total counts | 46,568,887 |
| mean counts / cell | 4,824 |
| mean counts / gene | 768 |
| min counts / cell | 480 |
| max counts / cell | 25,127 |
| overdispersion | 0.772 |

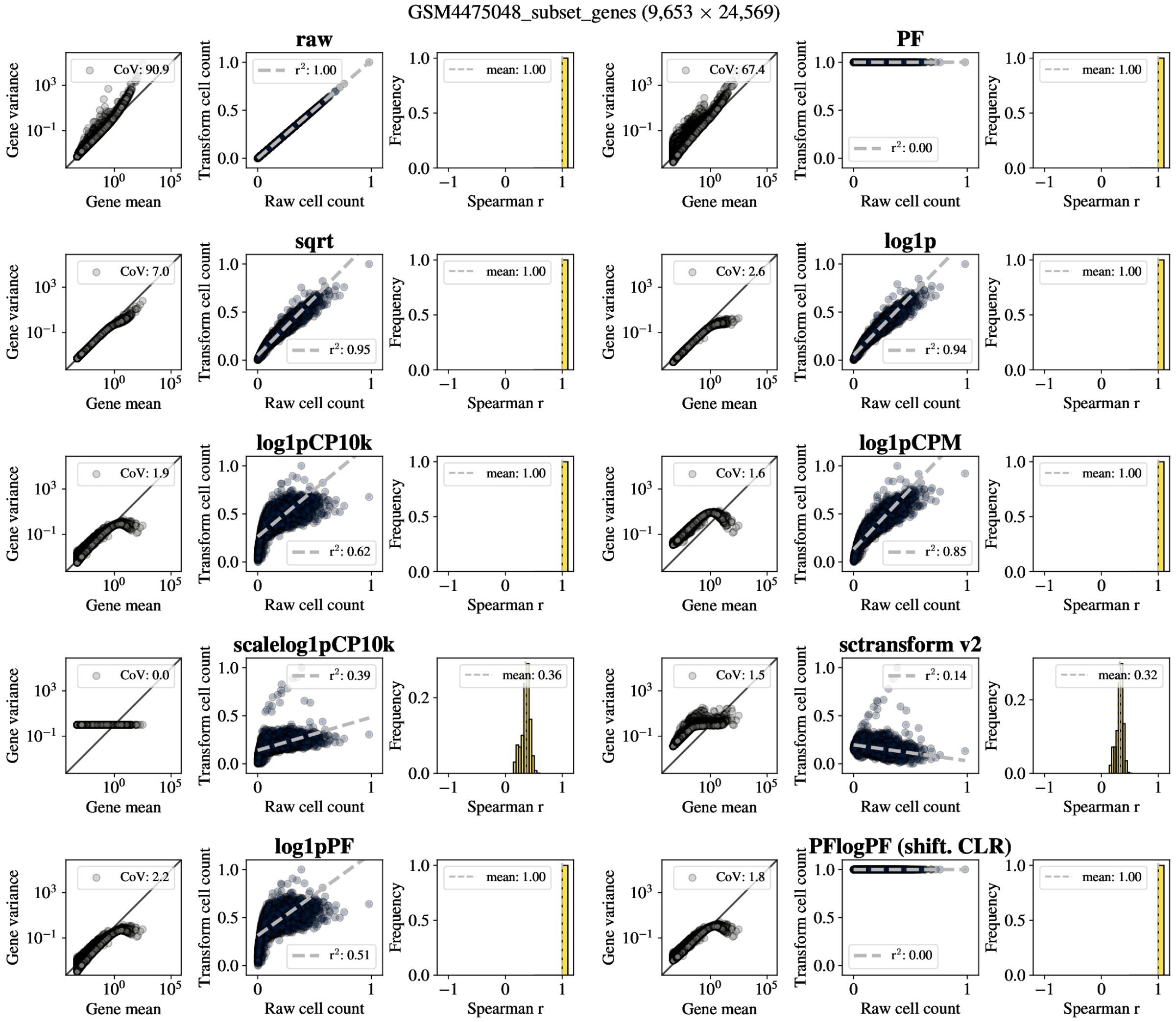


Supplementary Figure 1.503. **GSM4475048**. Per-method comparison of variance stabilization (left), depth normalization (middle), and monotonicity (right) for each normalization on dataset GSM4475048.

| GSM4475049 | |
|---|---|
| cells | 9,290 |
| genes | 60,623 |
| nonzero entries | 9,352,835 |
| density | 0.0166 |
| total counts | 27,259,082 |
| mean counts / cell | 2,934 |
| mean counts / gene | 450 |
| min counts / cell | 371 |
| max counts / cell | 16,889 |
| overdispersion | 0.753 |

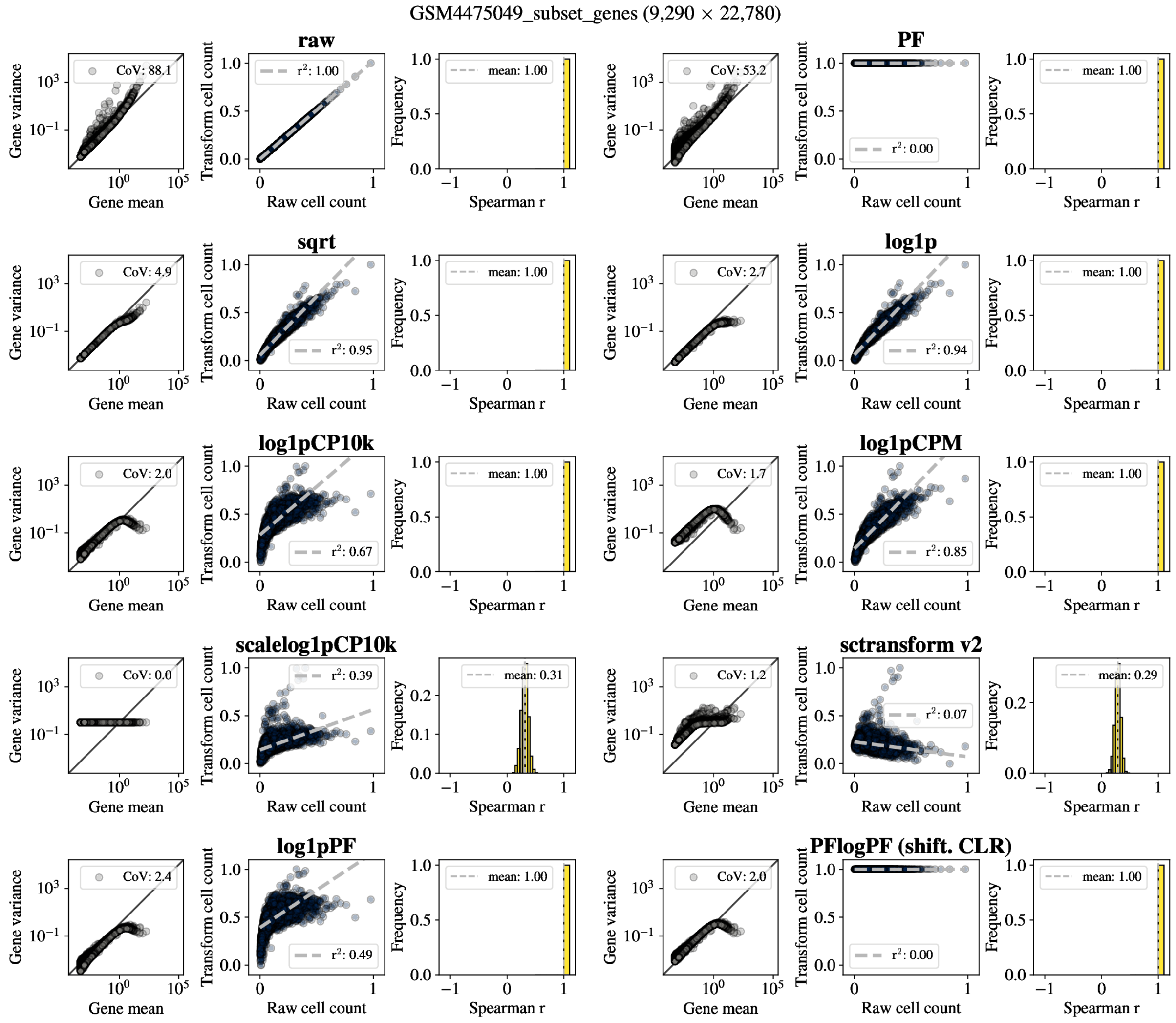


Supplementary Figure 1.504. **GSM4475049**. Per-method comparison of variance stabilization (left), depth normalization (middle), and monotonicity (right) for each normalization on dataset GSM4475049.

| GSM4475050 | |
|---|---|
| cells | 133 |
| genes | 60,623 |
| nonzero entries | 321,281 |
| density | 0.0398 |
| total counts | 1,361,885 |
| mean counts / cell | 10,240 |
| mean counts / gene | 22 |
| min counts / cell | 932 |
| max counts / cell | 34,417 |
| overdispersion | 1.74 |

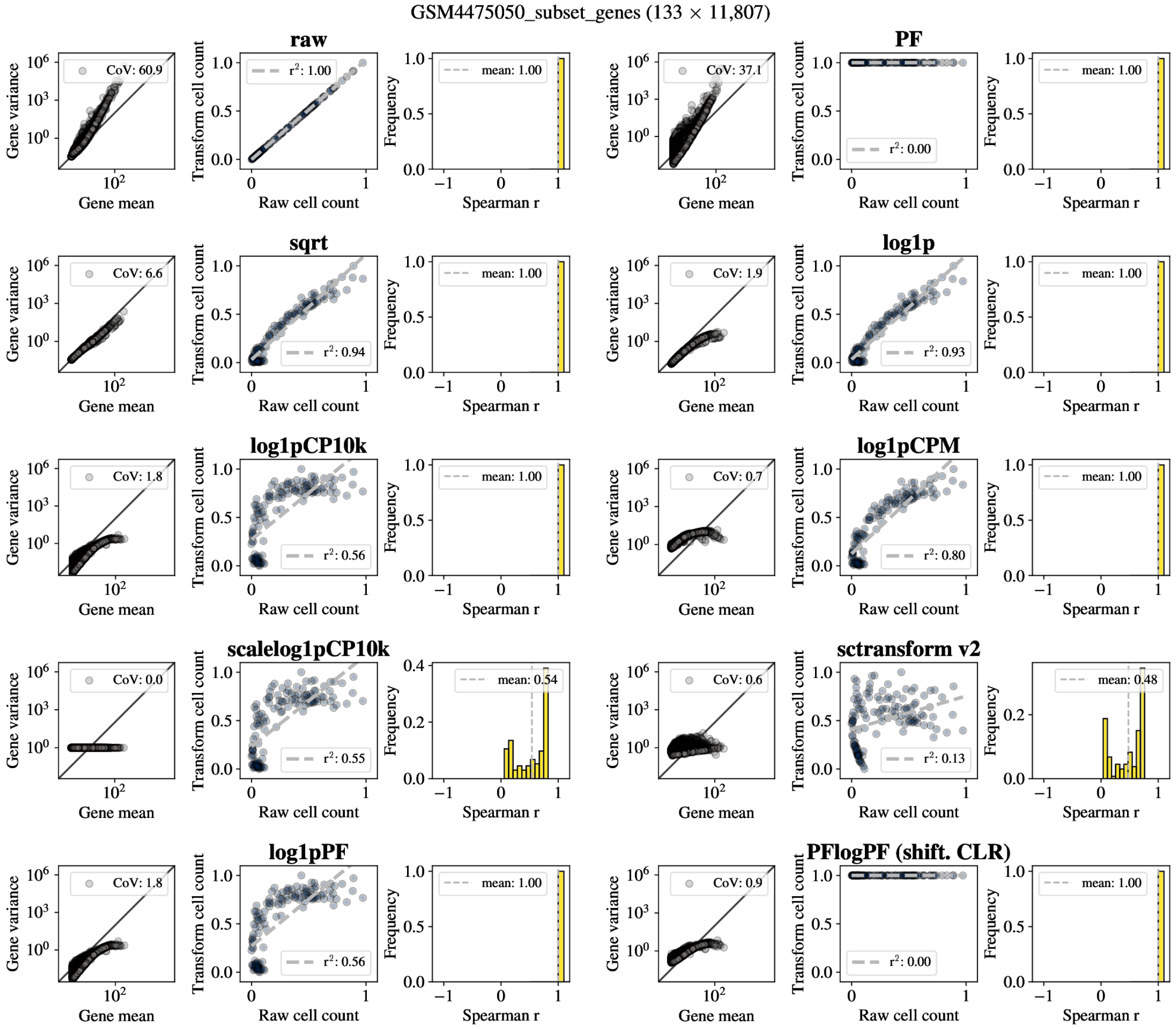


Supplementary Figure 1.505. **GSM4475050**. Per-method comparison of variance stabilization (left), depth normalization (middle), and monotonicity (right) for each normalization on dataset GSM4475050.

| GSM4475051 | |
|---|---|
| cells | 2,484 |
| genes | 60,623 |
| nonzero entries | 1,658,057 |
| density | 0.011 |
| total counts | 4,900,816 |
| mean counts / cell | 1,973 |
| mean counts / gene | 81 |
| min counts / cell | 26 |
| max counts / cell | 11,991 |
| overdispersion | 2.35 |

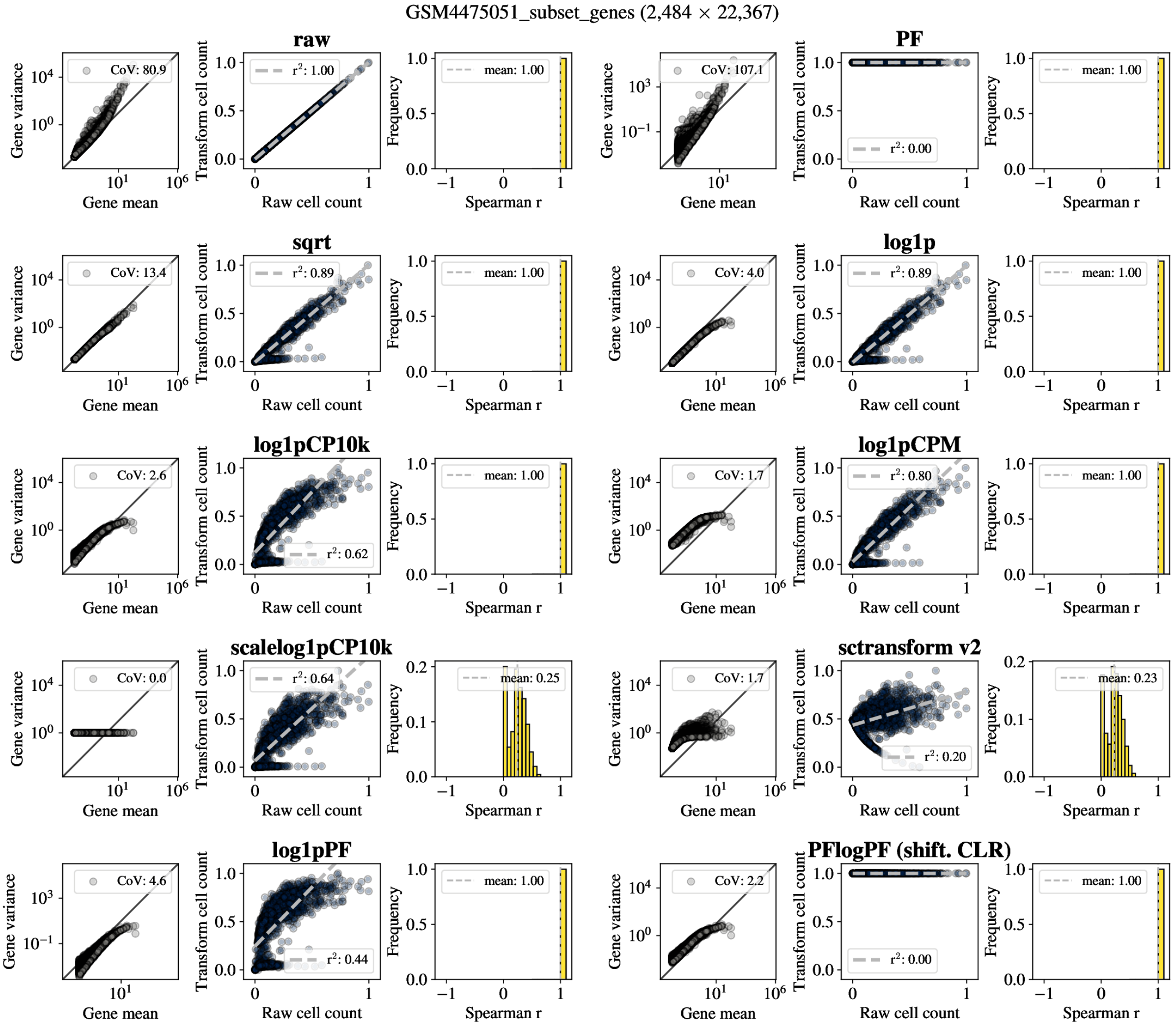


Supplementary Figure 1.506. **GSM4475051**. Per-method comparison of variance stabilization (left), depth normalization (middle), and monotonicity (right) for each normalization on dataset GSM4475051.

| GSM4475052 | |
|---|---|
| cells | 2,098 |
| genes | 60,623 |
| nonzero entries | 1,284,446 |
| density | 0.0101 |
| total counts | 3,518,188 |
| mean counts / cell | 1,677 |
| mean counts / gene | 58 |
| min counts / cell | 9 |
| max counts / cell | 14,652 |
| overdispersion | 1.36 |

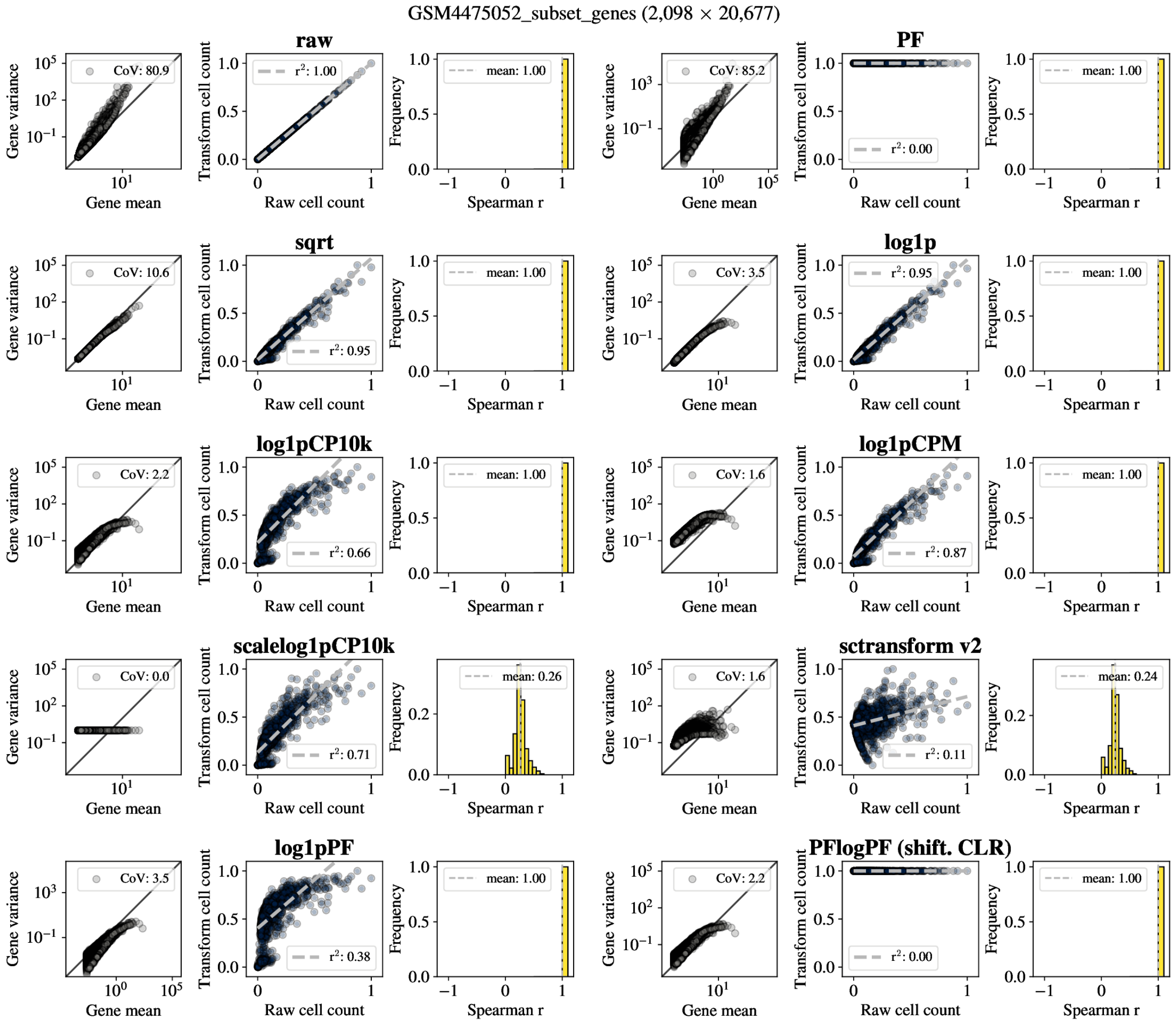


Supplementary Figure 1.507. **GSM4475052**. Per-method comparison of variance stabilization (left), depth normalization (middle), and monotonicity (right) for each normalization on dataset GSM4475052.

| GSM4475053 | |
|---|---|
| cells | 6,044 |
| genes | 60,623 |
| nonzero entries | 1,935,338 |
| density | 0.00528 |
| total counts | 5,192,362 |
| mean counts / cell | 859 |
| mean counts / gene | 86 |
| min counts / cell | 18 |
| max counts / cell | 12,922 |
| overdispersion | 1.22 |

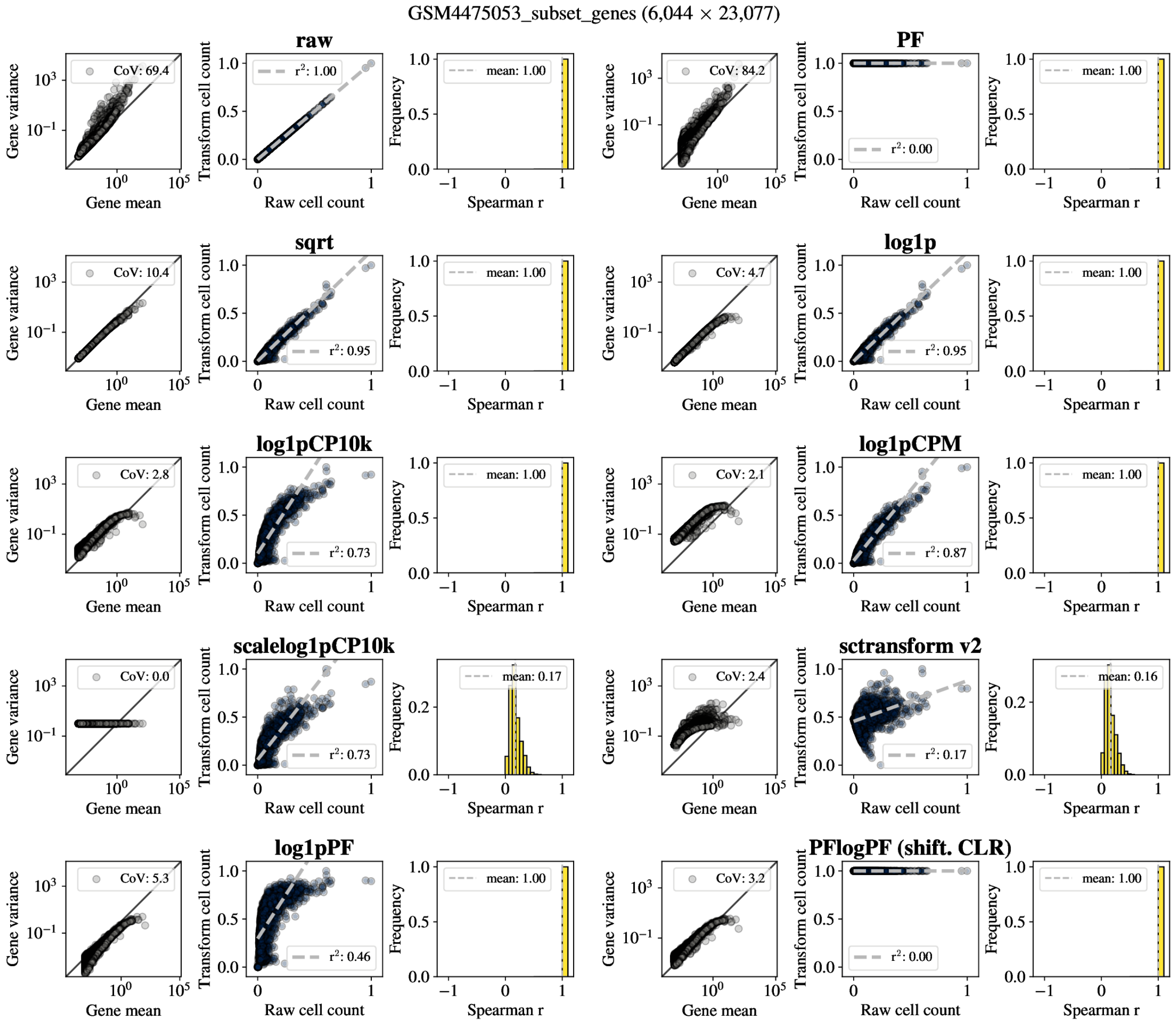


Supplementary Figure 1.508. **GSM4475053**. Per-method comparison of variance stabilization (left), depth normalization (middle), and monotonicity (right) for each normalization on dataset GSM4475053.

| GSM4769387 | |
|---|---|
| cells | 2,518 |
| genes | 60,623 |
| nonzero entries | 12,929,323 |
| density | 0.0847 |
| total counts | 72,485,575 |
| mean counts / cell | 28,787 |
| mean counts / gene | 1,196 |
| min counts / cell | 1,091 |
| max counts / cell | 159,454 |
| overdispersion | 1.63 |

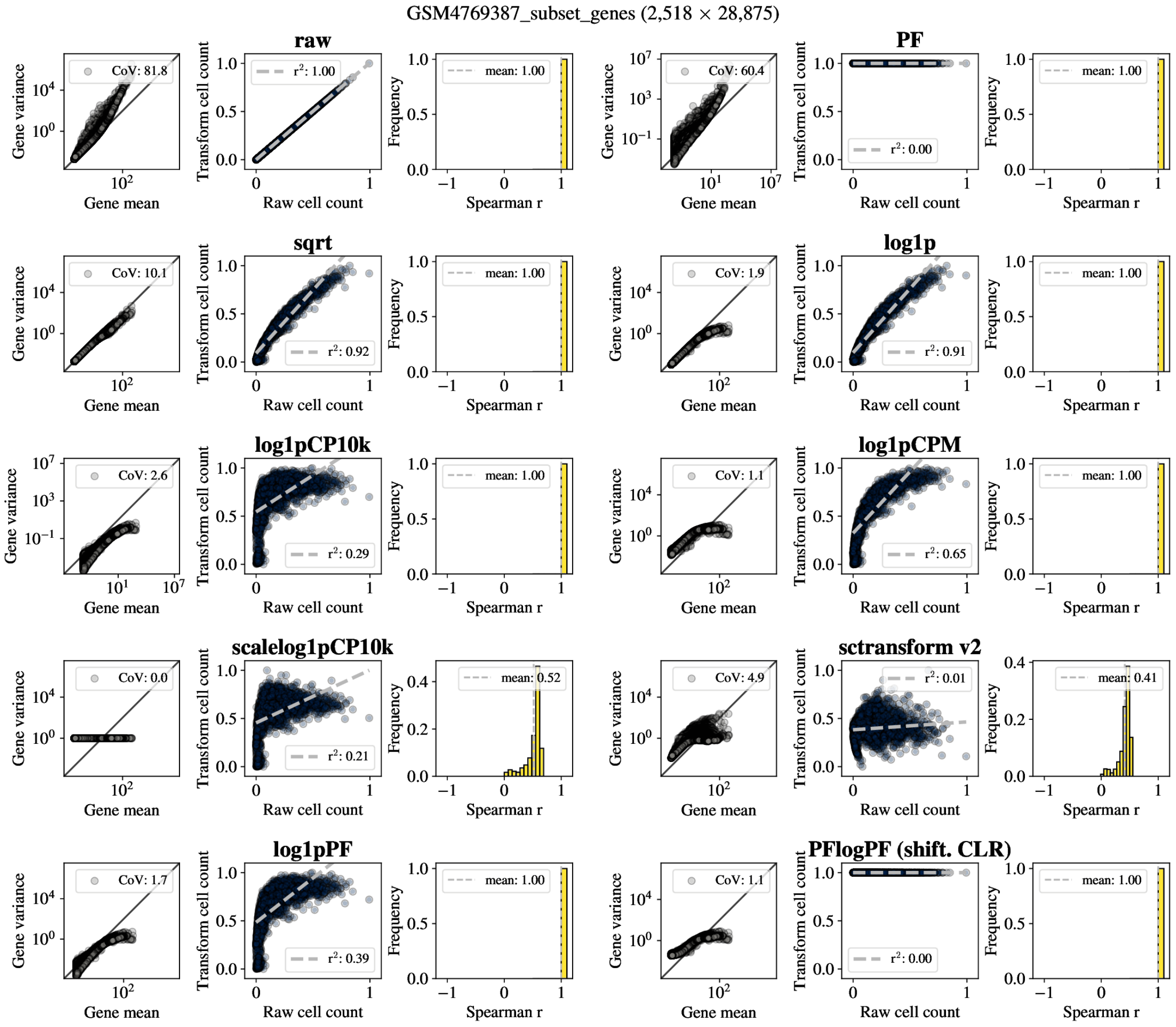


Supplementary Figure 1.509. **GSM4769387**. Per-method comparison of variance stabilization (left), depth normalization (middle), and monotonicity (right) for each normalization on dataset GSM4769387.

| GSM4769388 | |
|---|---|
| cells | 1,870 |
| genes | 60,623 |
| nonzero entries | 12,105,650 |
| density | 0.107 |
| total counts | 80,444,372 |
| mean counts / cell | 43,018 |
| mean counts / gene | 1,327 |
| min counts / cell | 1,111 |
| max counts / cell | 277,226 |
| overdispersion | 1.24 |

GSM4769388_subset_genes (1,870 × 29,083)

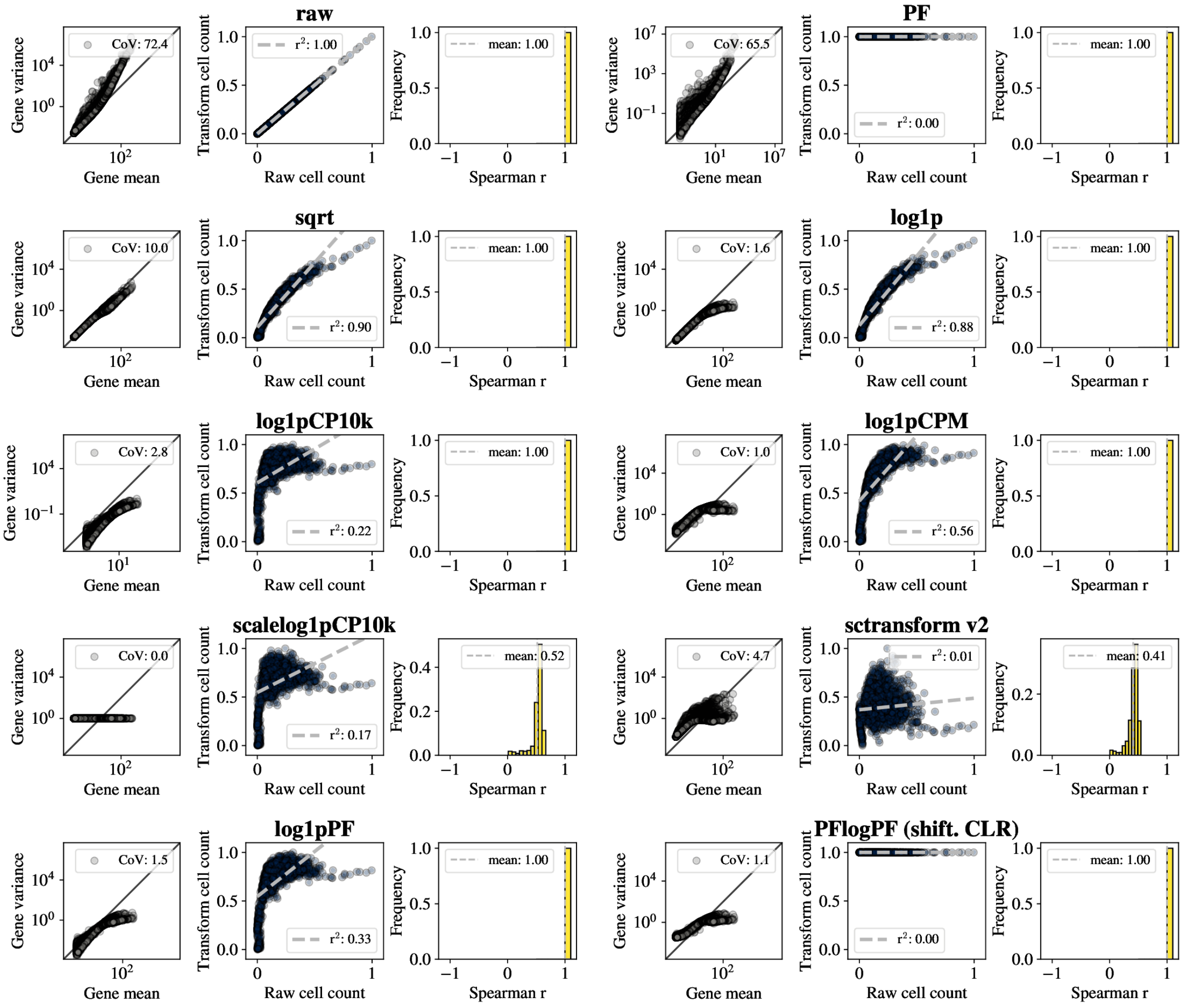


Supplementary Figure 1.510. **GSM4769388**. Per-method comparison of variance stabilization (left), depth normalization (middle), and monotonicity (right) for each normalization on dataset GSM4769388.

| GSM4769389 | |
|---|---|
| cells | 2,147 |
| genes | 60,623 |
| nonzero entries | 10,517,425 |
| density | 0.0808 |
| total counts | 58,151,317 |
| mean counts / cell | 27,085 |
| mean counts / gene | 959 |
| min counts / cell | 953 |
| max counts / cell | 148,808 |
| overdispersion | 0.555 |

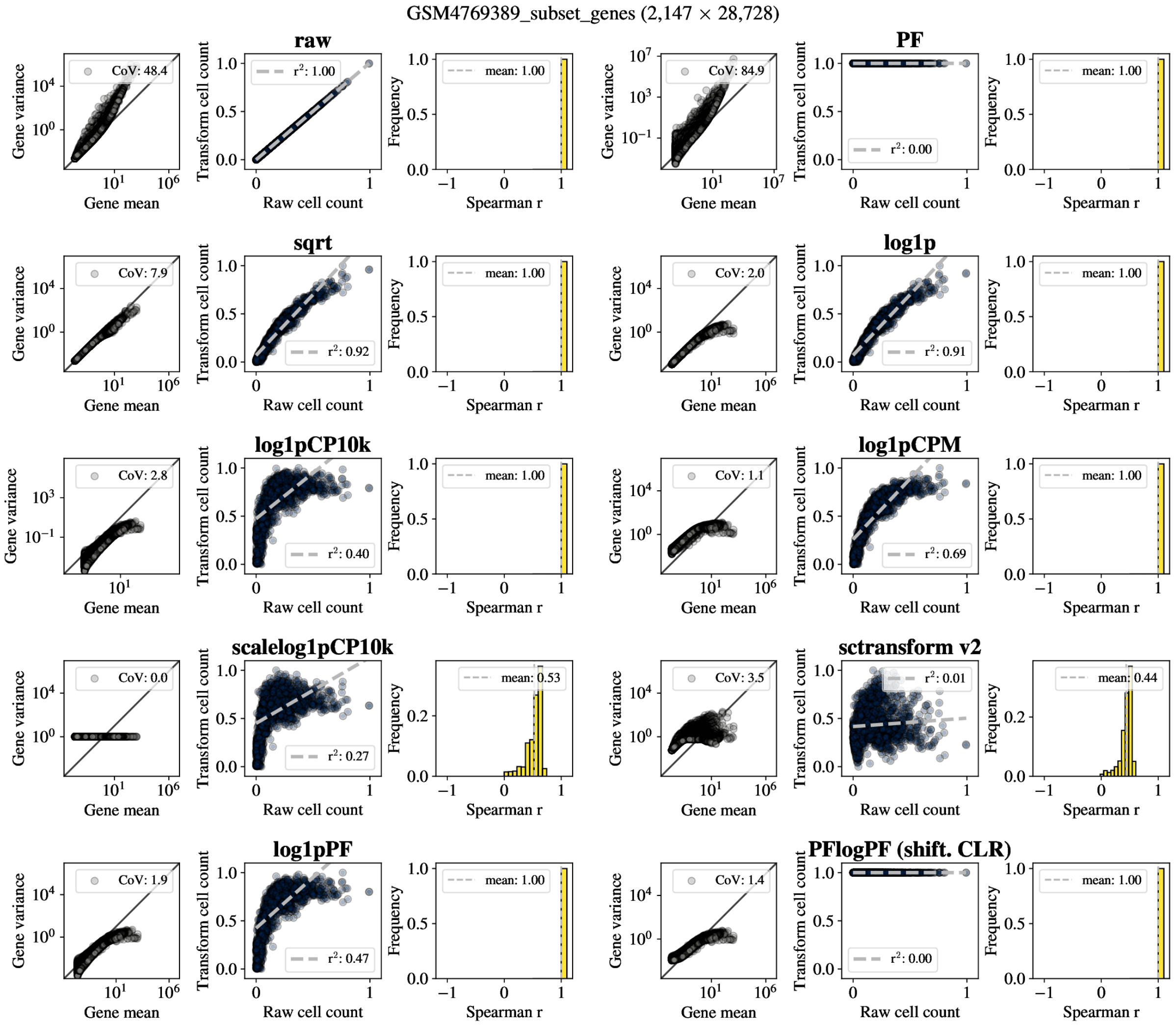


Supplementary Figure 1.511. **GSM4769389**. Per-method comparison of variance stabilization (left), depth normalization (middle), and monotonicity (right) for each normalization on dataset GSM4769389.

| GSM5051494 | |
|---|---|
| cells | 87 |
| genes | 60,623 |
| nonzero entries | 237,796 |
| density | 0.0451 |
| total counts | 717,296 |
| mean counts / cell | 8,245 |
| mean counts / gene | 12 |
| min counts / cell | 1,029 |
| max counts / cell | 70,175 |
| overdispersion | 10.4 |

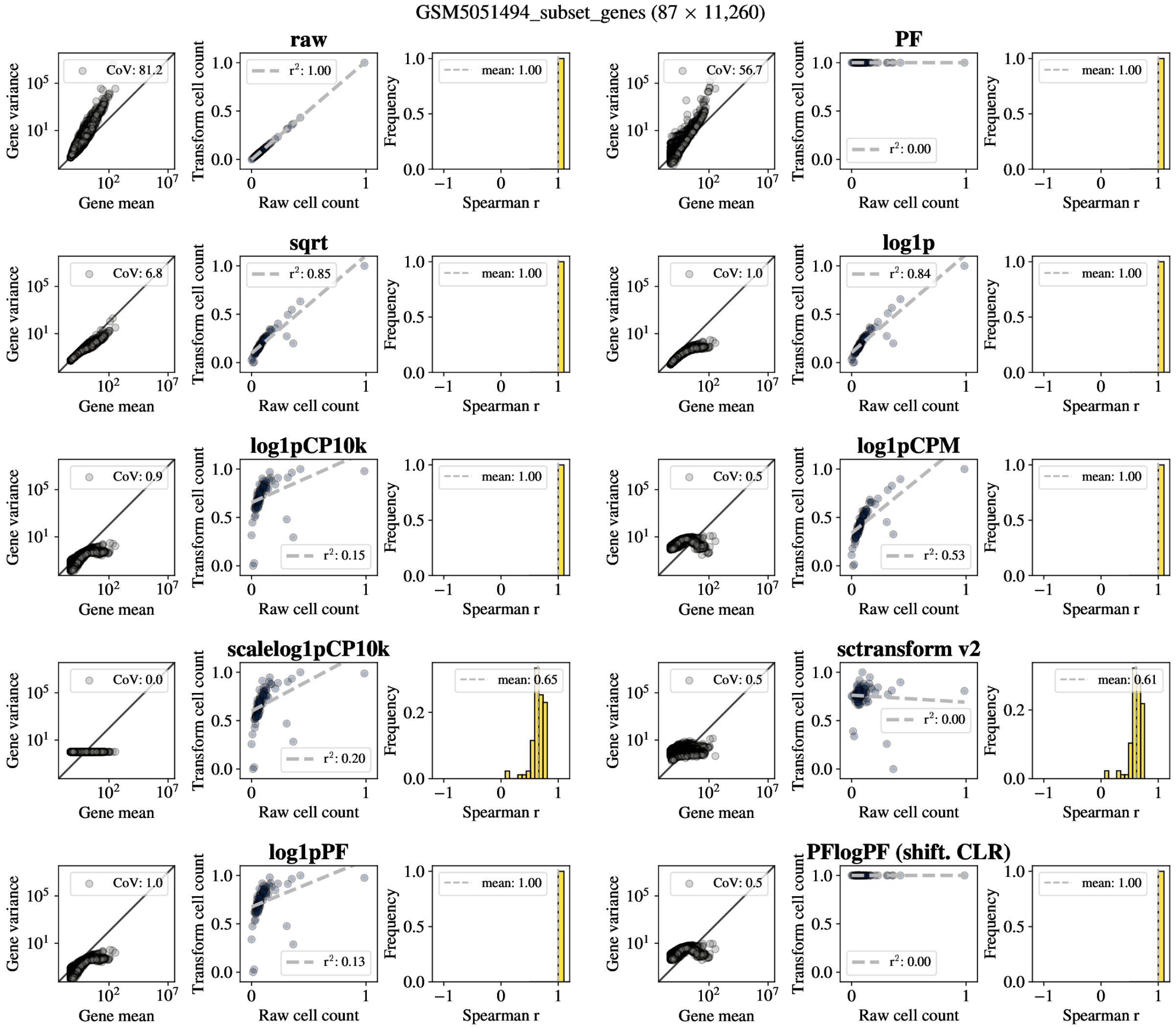


Supplementary Figure 1.512. **GSM5051494**. Per-method comparison of variance stabilization (left), depth normalization (middle), and monotonicity (right) for each normalization on dataset GSM5051494.

| GSM5051495 | |
|---|---|
| cells | 117 |
| genes | 60,623 |
| nonzero entries | 256,455 |
| density | 0.0362 |
| total counts | 615,314 |
| mean counts / cell | 5,259 |
| mean counts / gene | 10 |
| min counts / cell | 711 |
| max counts / cell | 17,483 |
| overdispersion | 0.536 |

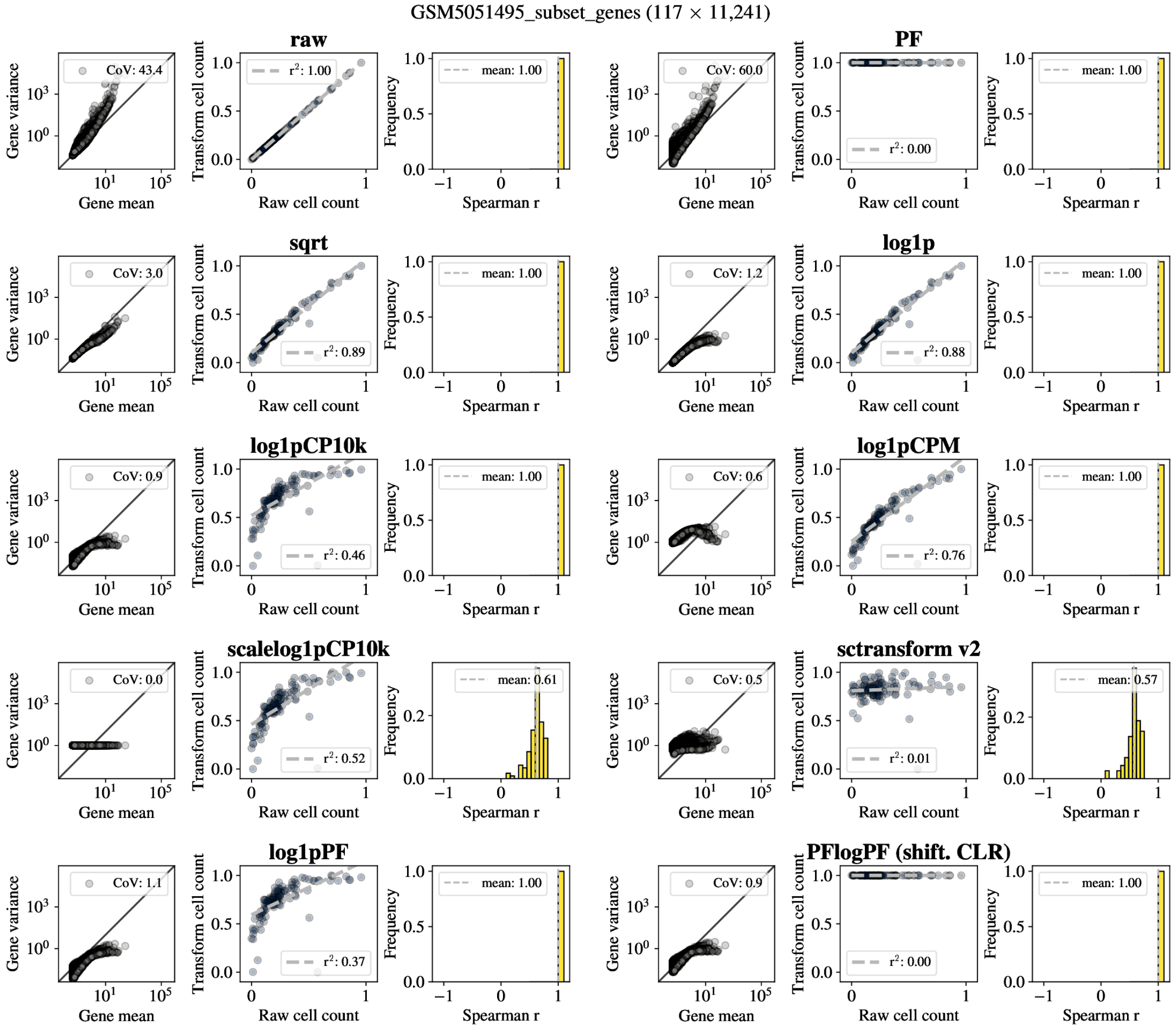


Supplementary Figure 1.513. **GSM5051495**. Per-method comparison of variance stabilization (left), depth normalization (middle), and monotonicity (right) for each normalization on dataset GSM5051495.

| GSM5051497 | |
|---|---|
| cells | 116 |
| genes | 60,623 |
| nonzero entries | 311,704 |
| density | 0.0443 |
| total counts | 823,839 |
| mean counts / cell | 7,102 |
| mean counts / gene | 14 |
| min counts / cell | 575 |
| max counts / cell | 31,354 |
| overdispersion | 1.21 |

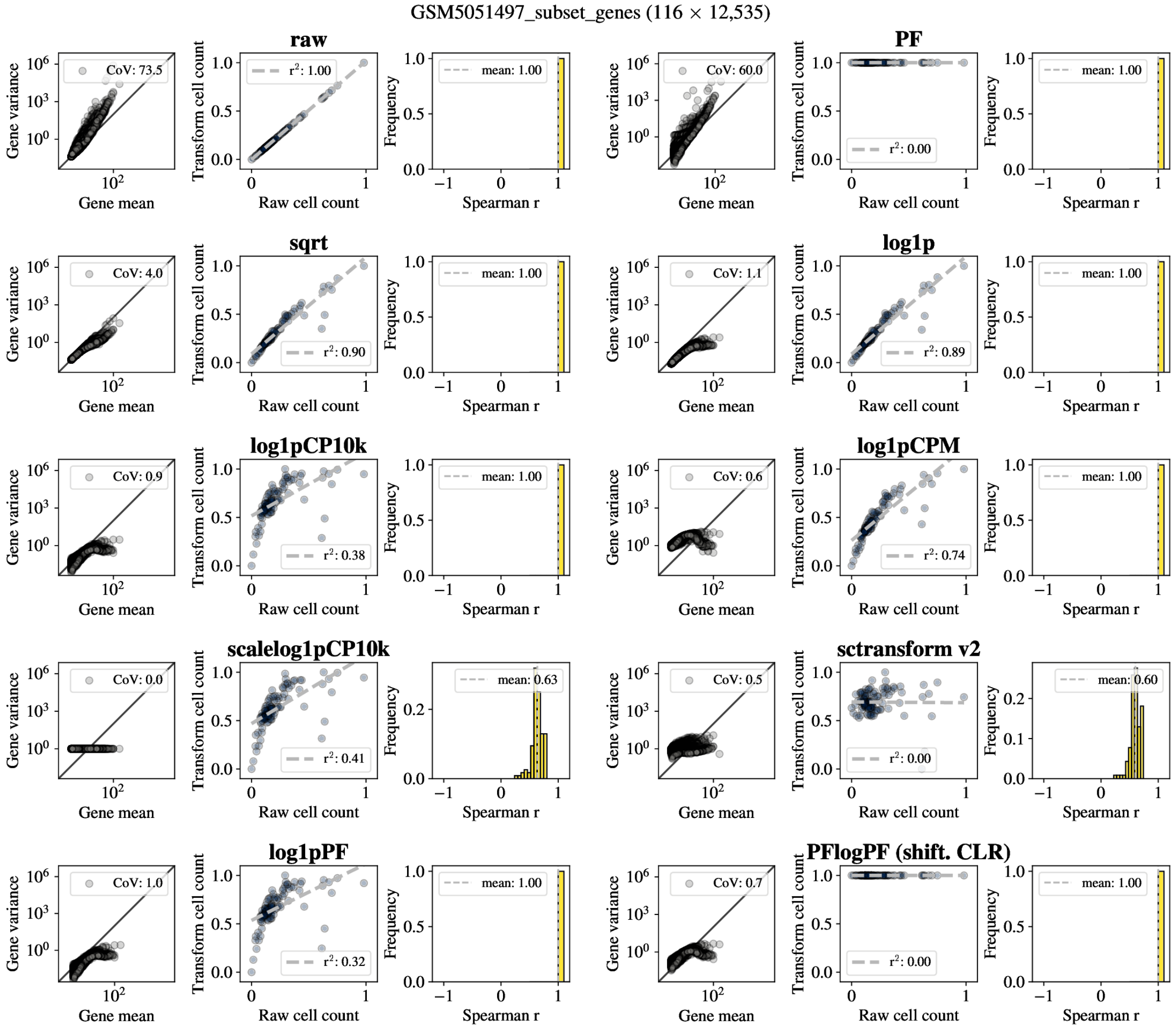


Supplementary Figure 1.514. **GSM5051497**. Per-method comparison of variance stabilization (left), depth normalization (middle), and monotonicity (right) for each normalization on dataset GSM5051497.

| GSM5051498 | |
|---|---|
| cells | 100 |
| genes | 60,623 |
| nonzero entries | 267,049 |
| density | 0.0441 |
| total counts | 758,250 |
| mean counts / cell | 7,582 |
| mean counts / gene | 13 |
| min counts / cell | 868 |
| max counts / cell | 42,098 |
| overdispersion | 21 |

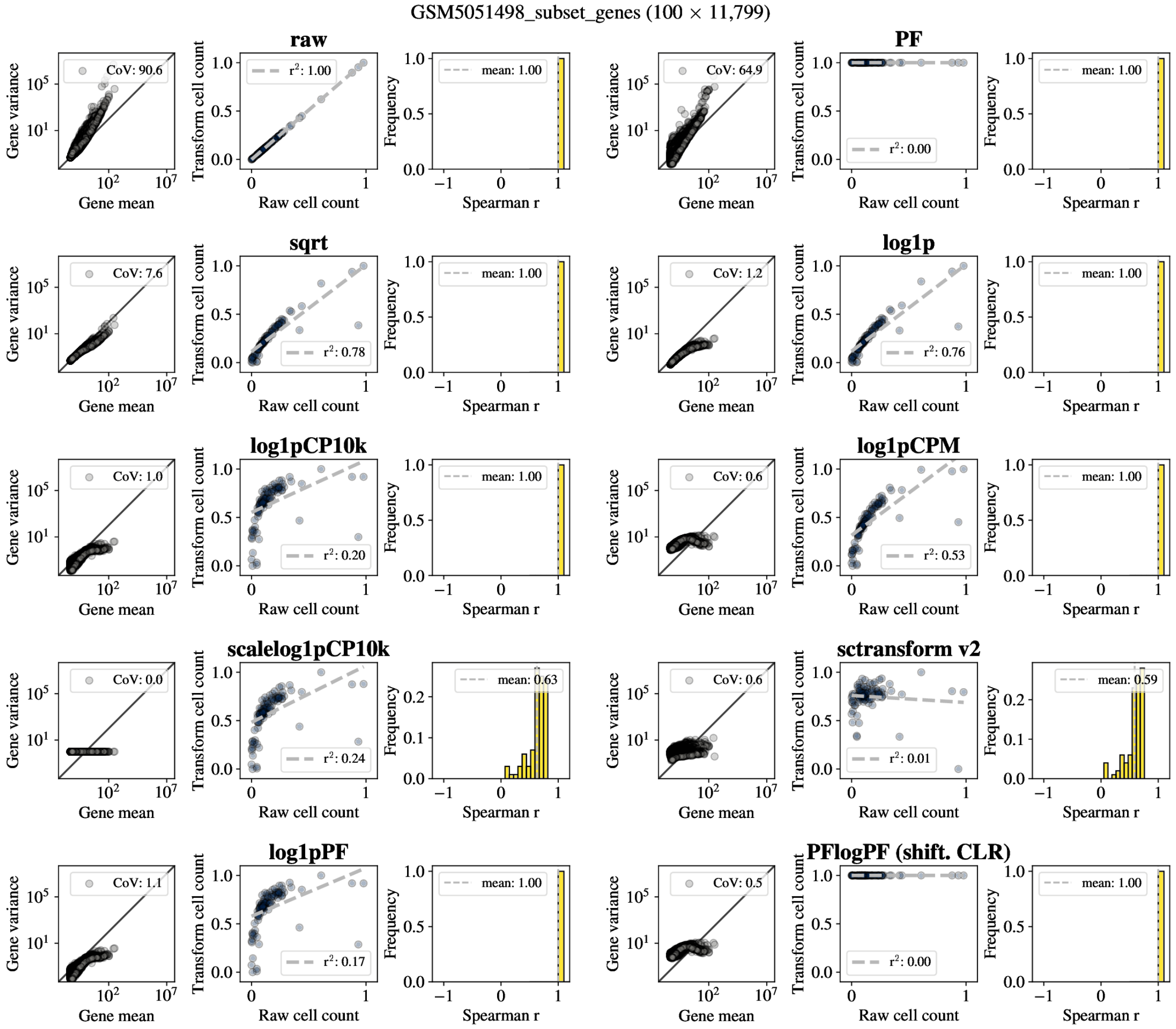


Supplementary Figure 1.515. **GSM5051498**. Per-method comparison of variance stabilization (left), depth normalization (middle), and monotonicity (right) for each normalization on dataset GSM5051498.

| **klein_2015** | |
|---|---|
| cells | 953 |
| genes | 25,435 |
| nonzero entries | 9,262,087 |
| density | 0.382 |
| total counts | 30,963,149 |
| mean counts / cell | 32,490 |
| mean counts / gene | 1,217 |
| min counts / cell | 12,721 |
| max counts / cell | 73,810 |
| overdispersion | 0.0428 |

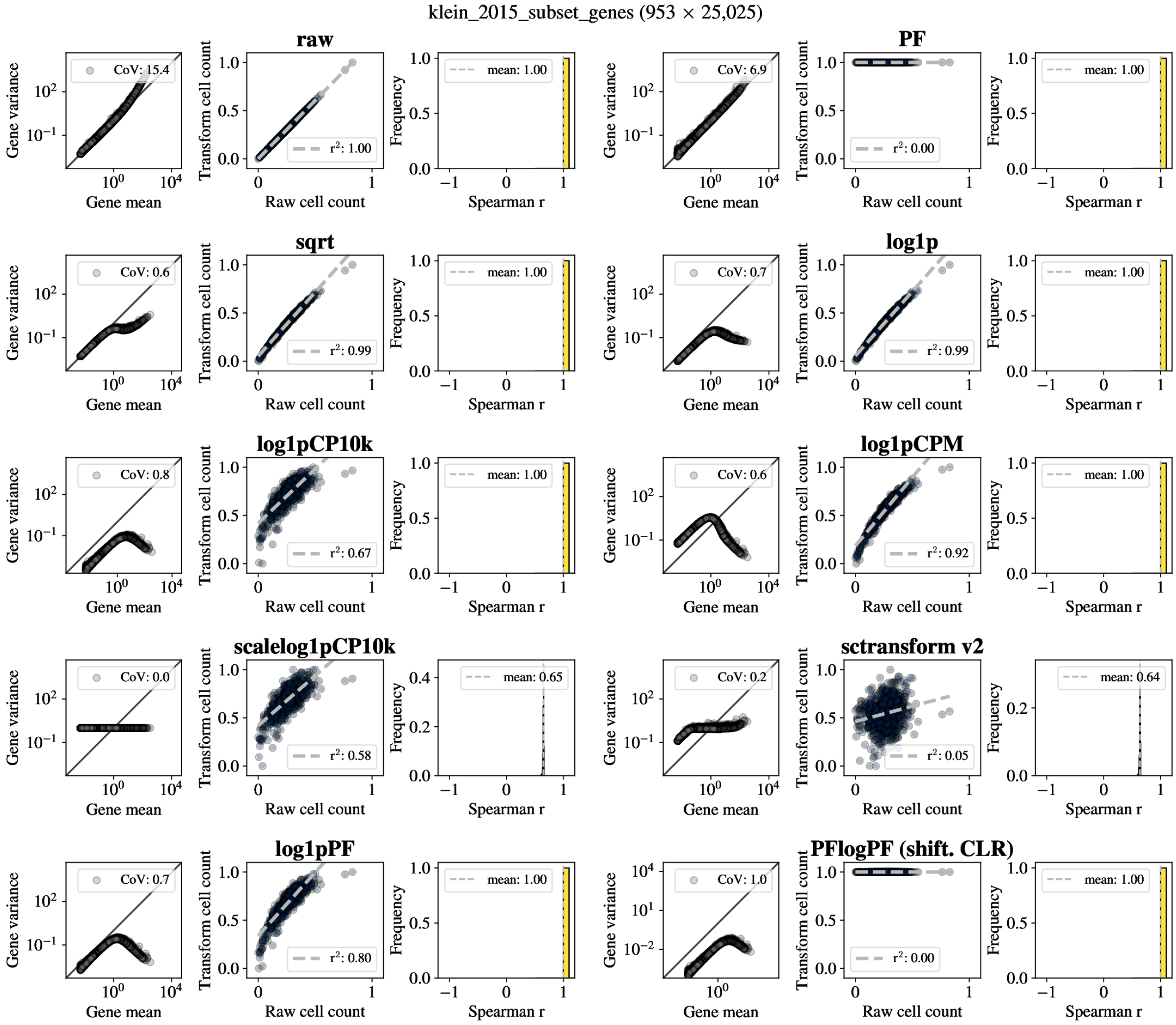


Supplementary Figure 1.516. **klein_2015**. Per-method comparison of variance stabilization (left), depth normalization (middle), and monotonicity (right) for each normalization on dataset klein_2015.

| macosko_2015 | |
|---|---|
| cells | 84 |
| genes | 80 |
| nonzero entries | 4,009 |
| density | 0.597 |
| total counts | 1,020,913 |
| mean counts / cell | 12,154 |
| mean counts / gene | 12,761 |
| min counts / cell | 4,337 |
| max counts / cell | 21,127 |
| overdispersion | 0.0481 |

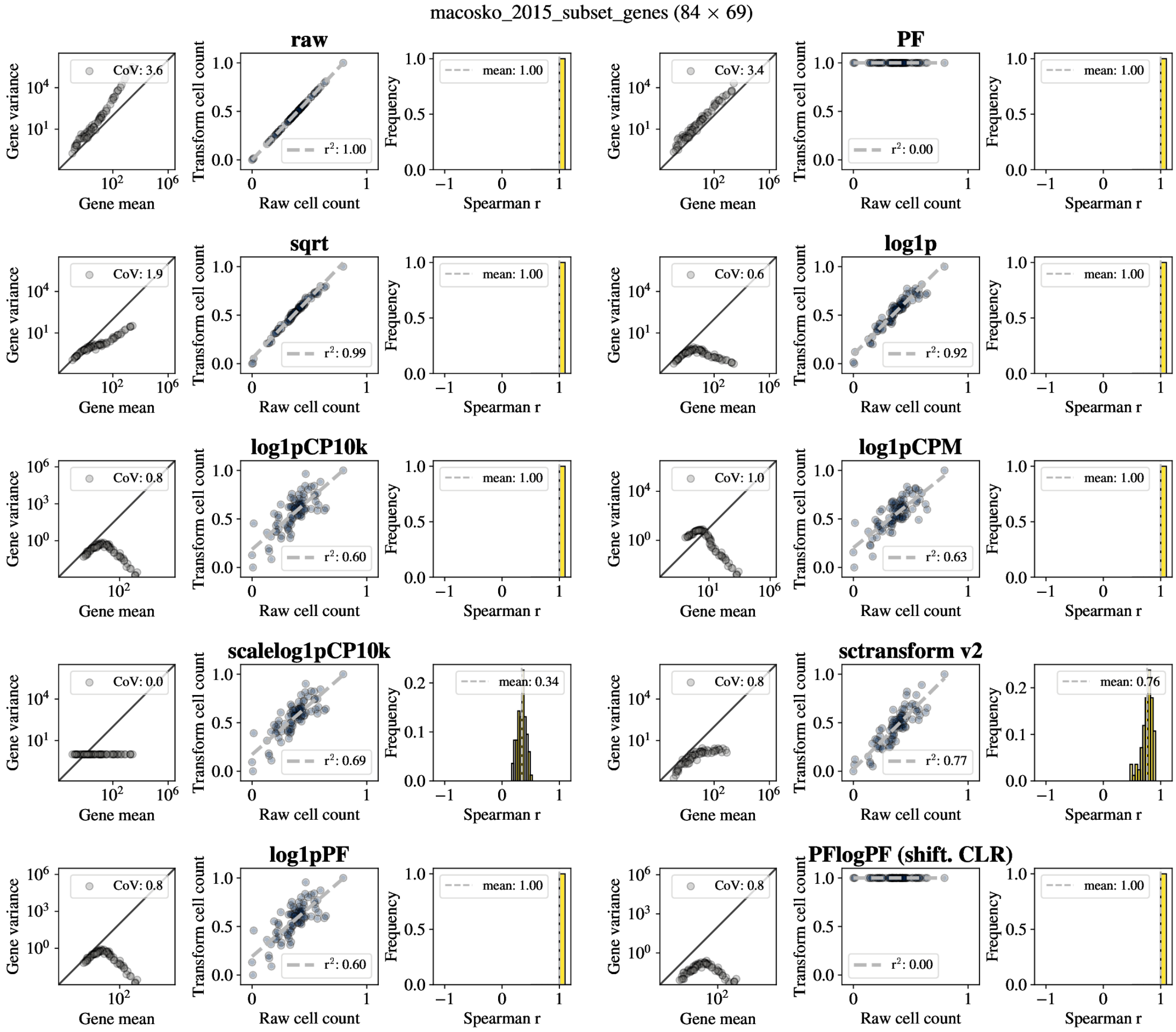


Supplementary Figure 1.517. **macosko_2015**. Per-method comparison of variance stabilization (left), depth normalization (middle), and monotonicity (right) for each normalization on dataset macosko_2015.

| padovan_merhar_2015 | |
|---|---|
| cells | 96 |
| genes | 25,590 |
| nonzero entries | 668,954 |
| density | 0.272 |
| total counts | 169,164,561 |
| mean counts / cell | 1,762,131 |
| mean counts / gene | 6,611 |
| min counts / cell | 259,281 |
| max counts / cell | 3,519,323 |
| overdispersion | 4.06 |

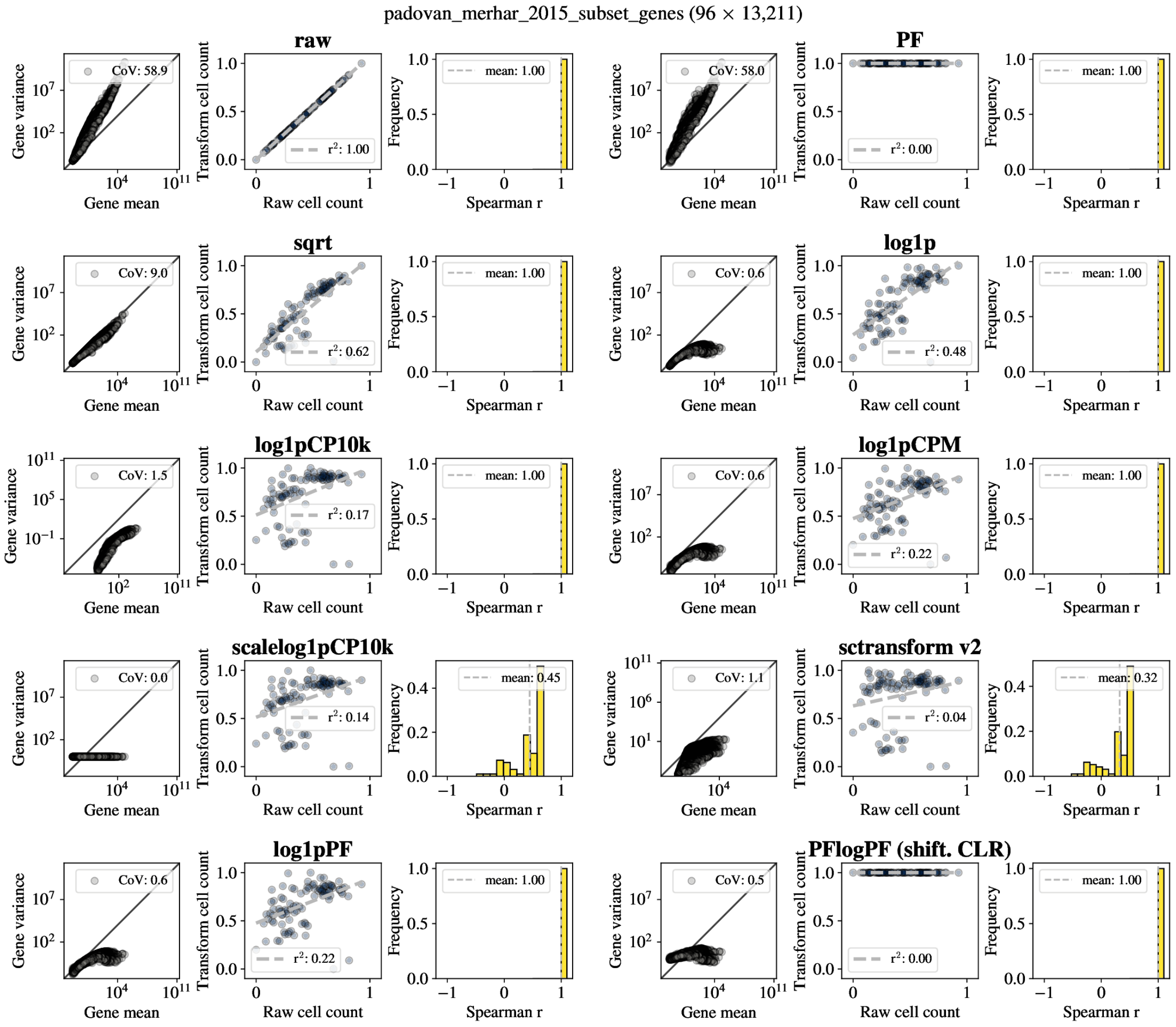


Supplementary Figure 1.518. **padovan_merhar_2015**. Per-method comparison of variance stabilization (left), depth normalization (middle), and monotonicity (right) for each normalization on dataset padovan_merhar_2015.

| pbmc3k | |
|---|---|
| cells | 2,700 |
| genes | 32,738 |
| nonzero entries | 2,286,884 |
| density | 0.0259 |
| total counts | 6,390,631 |
| mean counts / cell | 2,367 |
| mean counts / gene | 195 |
| min counts / cell | 548 |
| max counts / cell | 15,844 |
| overdispersion | 0.393 |

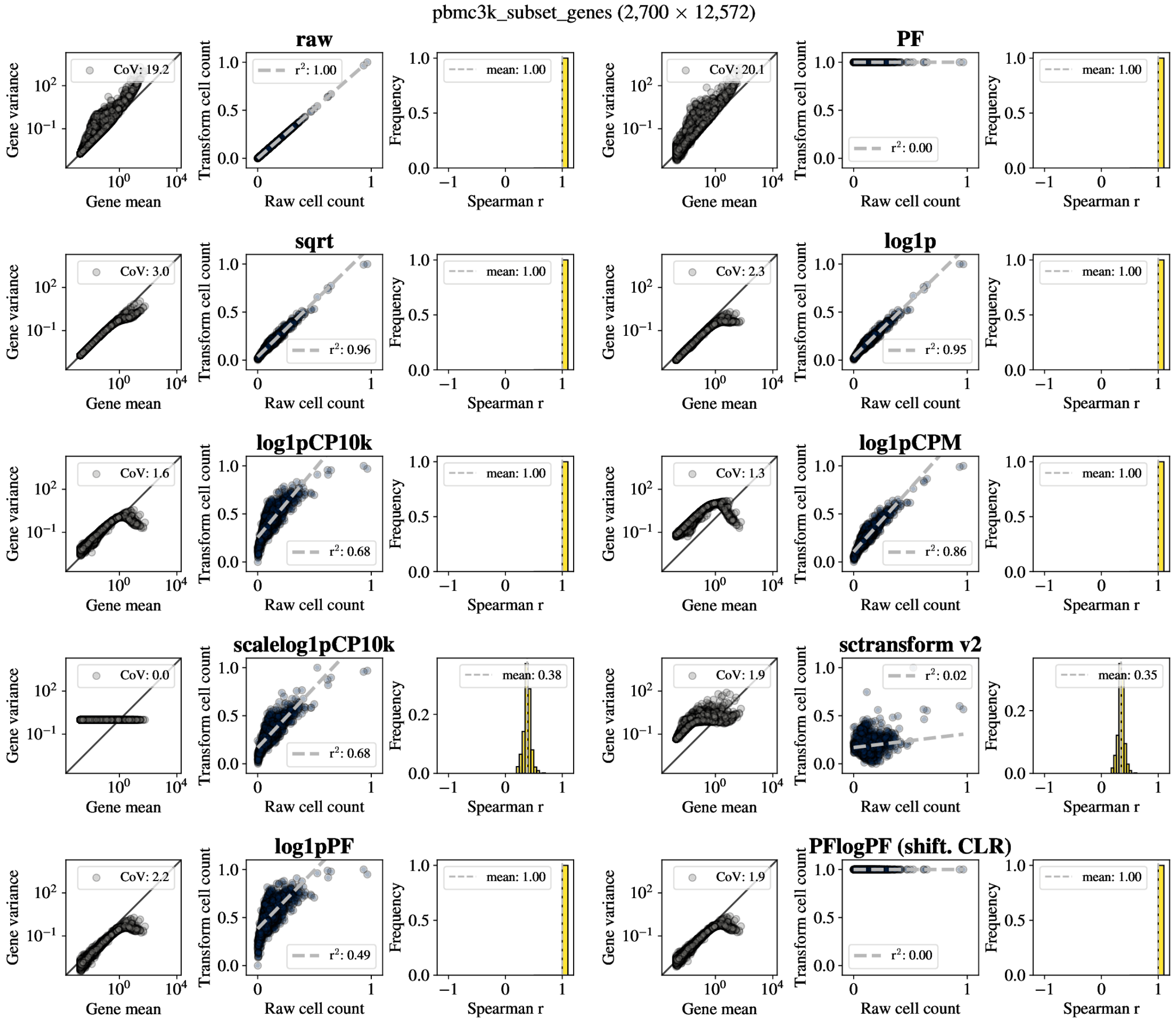


Supplementary Figure 1.519. **pbmc3k**. Per-method comparison of variance stabilization (left), depth normalization (middle), and monotonicity (right) for each normalization on dataset pbmc3k.

| SAMN14275543 | |
|---|---|
| cells | 6,814 |
| genes | 60,623 |
| nonzero entries | 2,555,781 |
| density | 0.00619 |
| total counts | 4,790,795 |
| mean counts / cell | 703 |
| mean counts / gene | 79 |
| min counts / cell | 1 |
| max counts / cell | 7,254 |
| overdispersion | 3.82 |

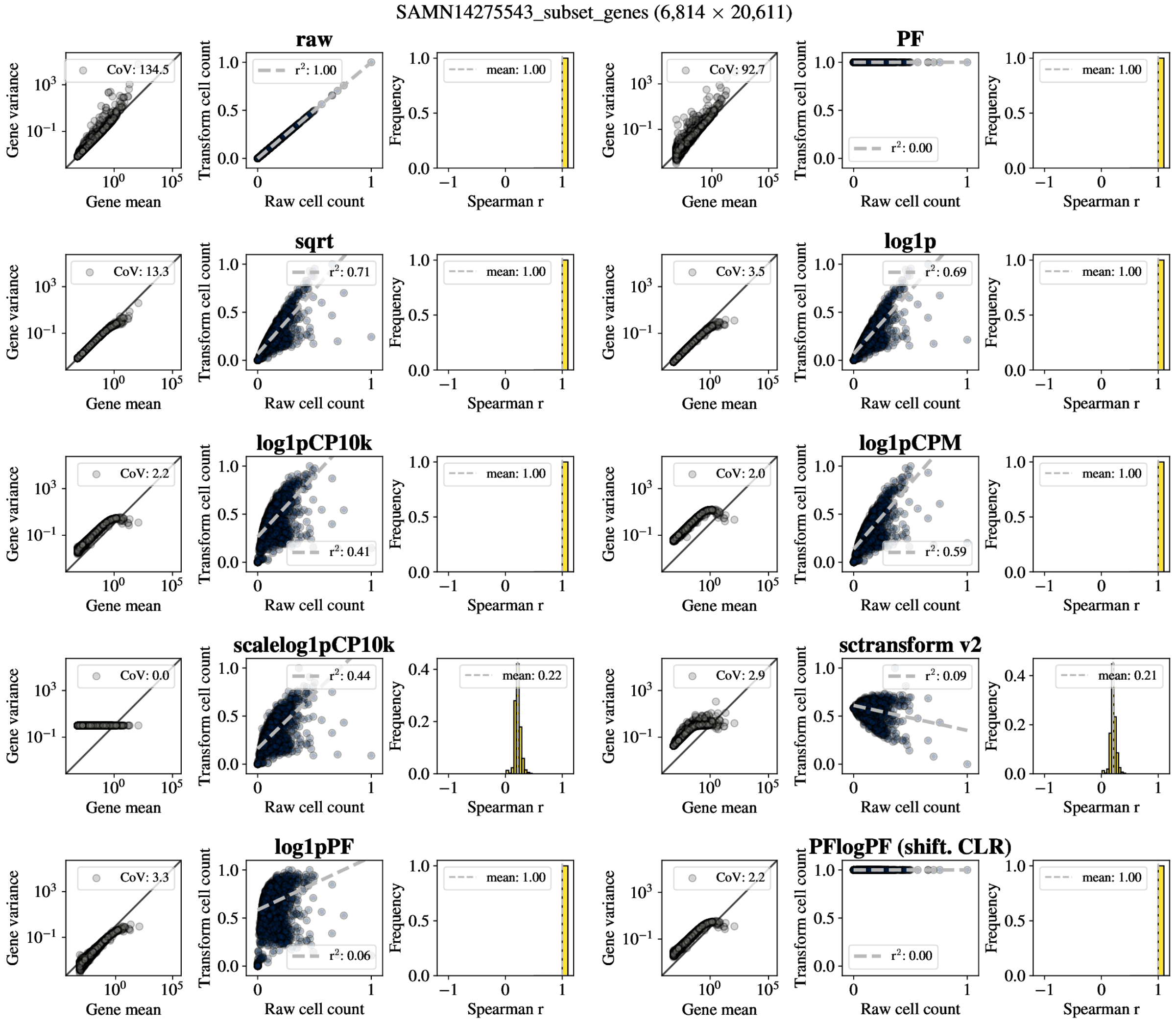


Supplementary Figure 1.520. **SAMN14275543**. Per-method comparison of variance stabilization (left), depth normalization (middle), and monotonicity (right) for each normalization on dataset SAMN14275543.

| SRX4733402 | |
|---|---|
| cells | 12,650 |
| genes | 60,623 |
| nonzero entries | 7,428,525 |
| density | 0.00969 |
| total counts | 16,630,190 |
| mean counts / cell | 1,315 |
| mean counts / gene | 274 |
| min counts / cell | 229 |
| max counts / cell | 58,149 |
| overdispersion | 3.05 |

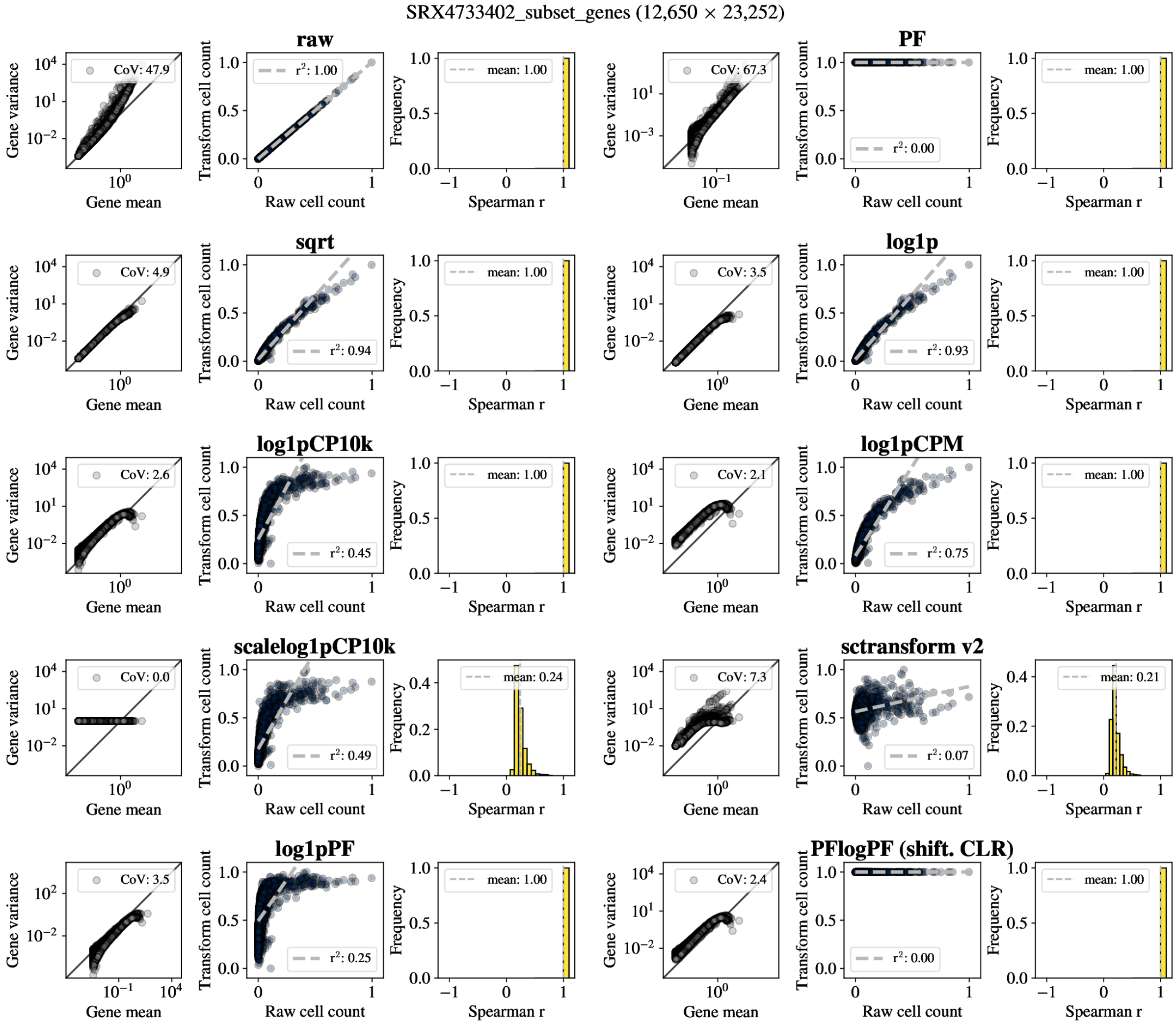


Supplementary Figure 1.521. **SRX4733402**. Per-method comparison of variance stabilization (left), depth normalization (middle), and monotonicity (right) for each normalization on dataset SRX4733402.

| SRX4733403 | |
|---|---|
| cells | 6,670 |
| genes | 60,623 |
| nonzero entries | 2,547,751 |
| density | 0.0063 |
| total counts | 4,444,233 |
| mean counts / cell | 666 |
| mean counts / gene | 73 |
| min counts / cell | 98 |
| max counts / cell | 7,999 |
| overdispersion | 1.17 |

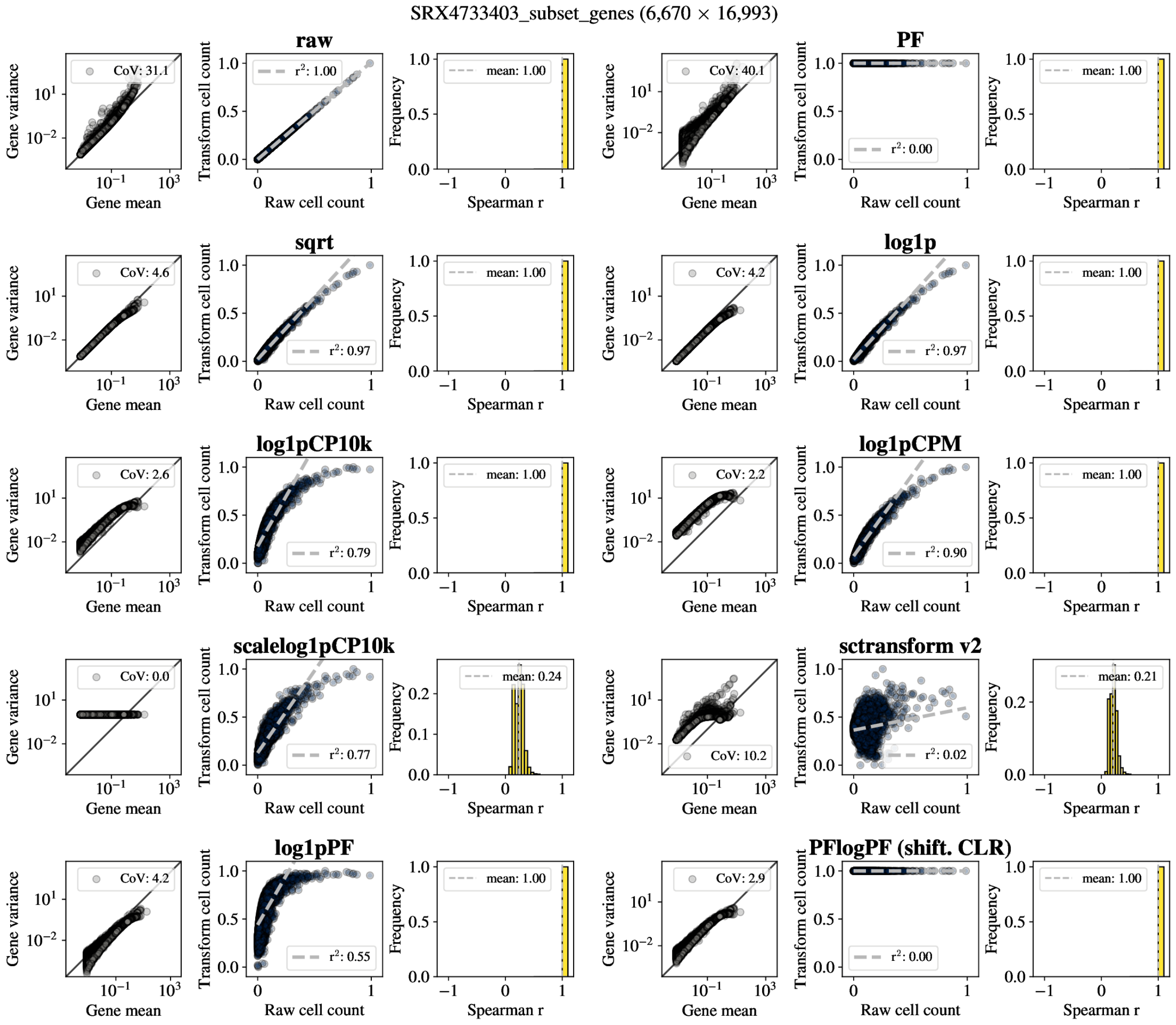


Supplementary Figure 1.522. **SRX4733403**. Per-method comparison of variance stabilization (left), depth normalization (middle), and monotonicity (right) for each normalization on dataset SRX4733403.

| SRX4733411 | |
|---|---|
| cells | 2,031 |
| genes | 60,623 |
| nonzero entries | 7,898,613 |
| density | 0.0642 |
| total counts | 52,043,097 |
| mean counts / cell | 25,624 |
| mean counts / gene | 858 |
| min counts / cell | 6,047 |
| max counts / cell | 97,107 |
| overdispersion | 0.553 |

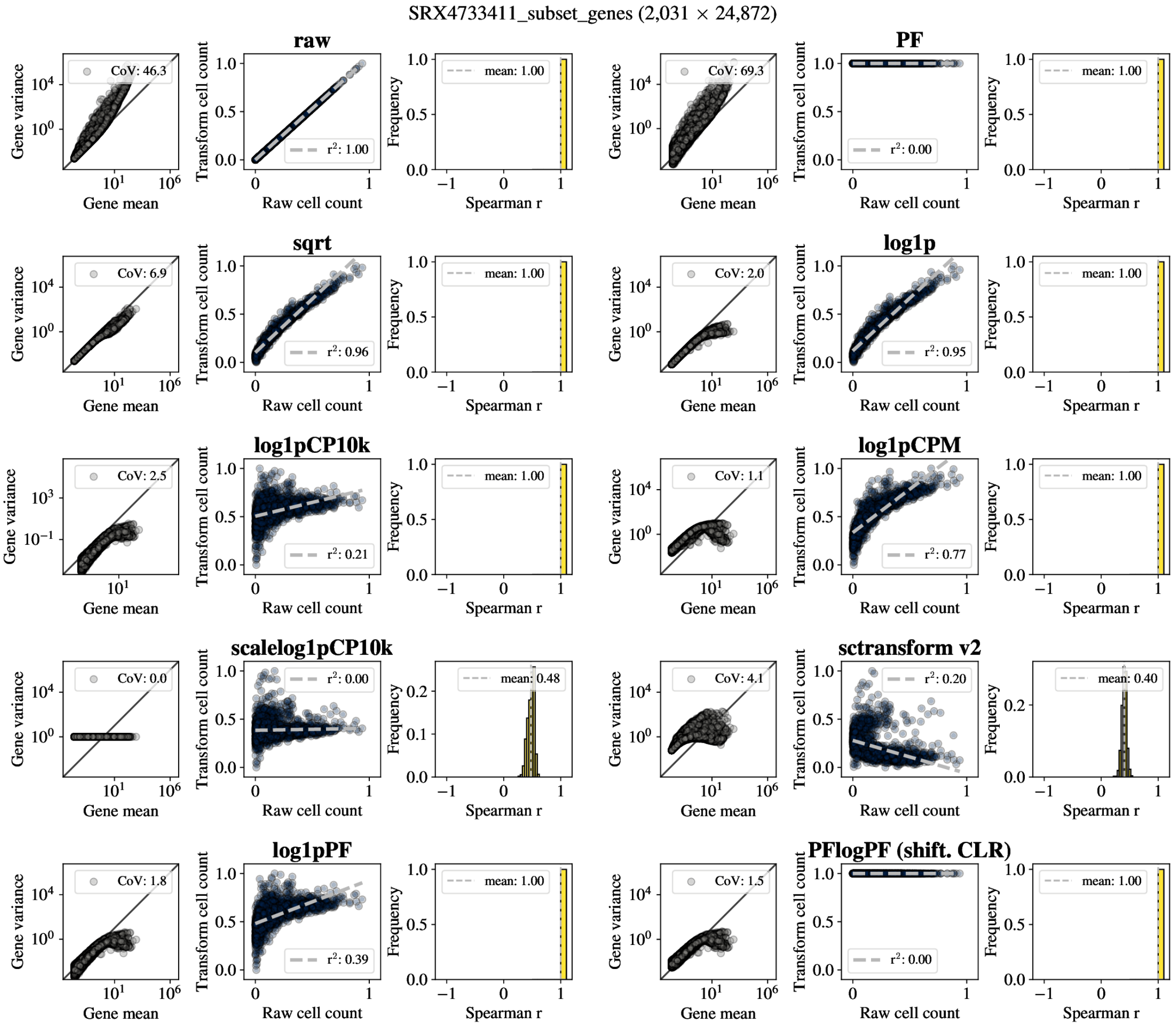


Supplementary Figure 1.523. **SRX4733411**. Per-method comparison of variance stabilization (left), depth normalization (middle), and monotonicity (right) for each normalization on dataset SRX4733411.

| svensson_2017_1 | |
|---|---|
| cells | 2,000 |
| genes | 24,116 |
| nonzero entries | 1,253,823 |
| density | 0.026 |
| total counts | 4,580,570 |
| mean counts / cell | 2,290 |
| mean counts / gene | 190 |
| min counts / cell | 476 |
| max counts / cell | 7,370 |
| overdispersion | 0.094 |

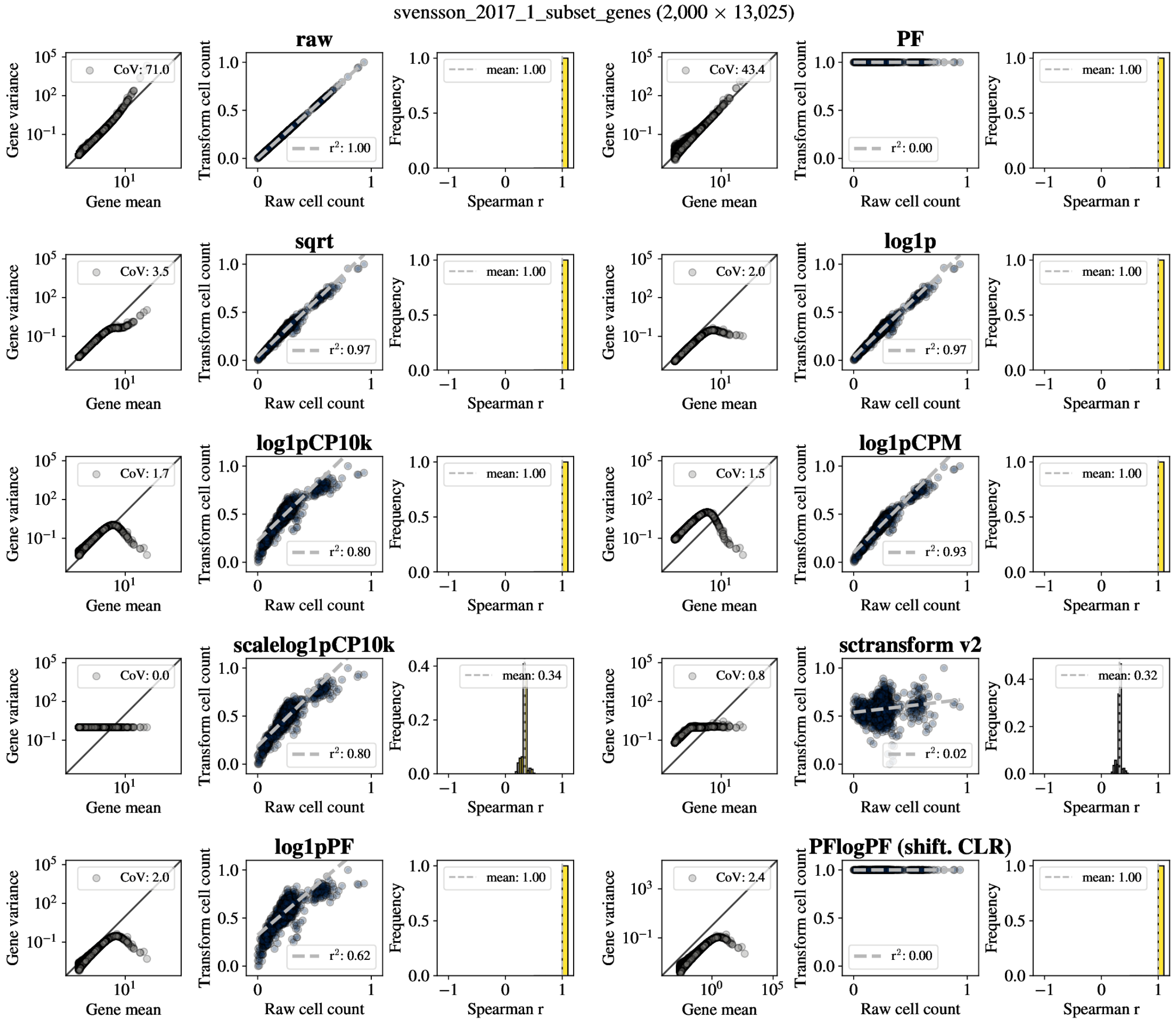


Supplementary Figure 1.524. **svensson_2017_1**. Per-method comparison of variance stabilization (left), depth normalization (middle), and monotonicity (right) for each normalization on dataset svensson_2017_1.

| svensson_2017_2 | |
|---|---|
| cells | 2,000 |
| genes | 24,116 |
| nonzero entries | 1,150,374 |
| density | 0.0239 |
| total counts | 4,474,561 |
| mean counts / cell | 2,237 |
| mean counts / gene | 186 |
| min counts / cell | 399 |
| max counts / cell | 26,254 |
| overdispersion | 0.369 |

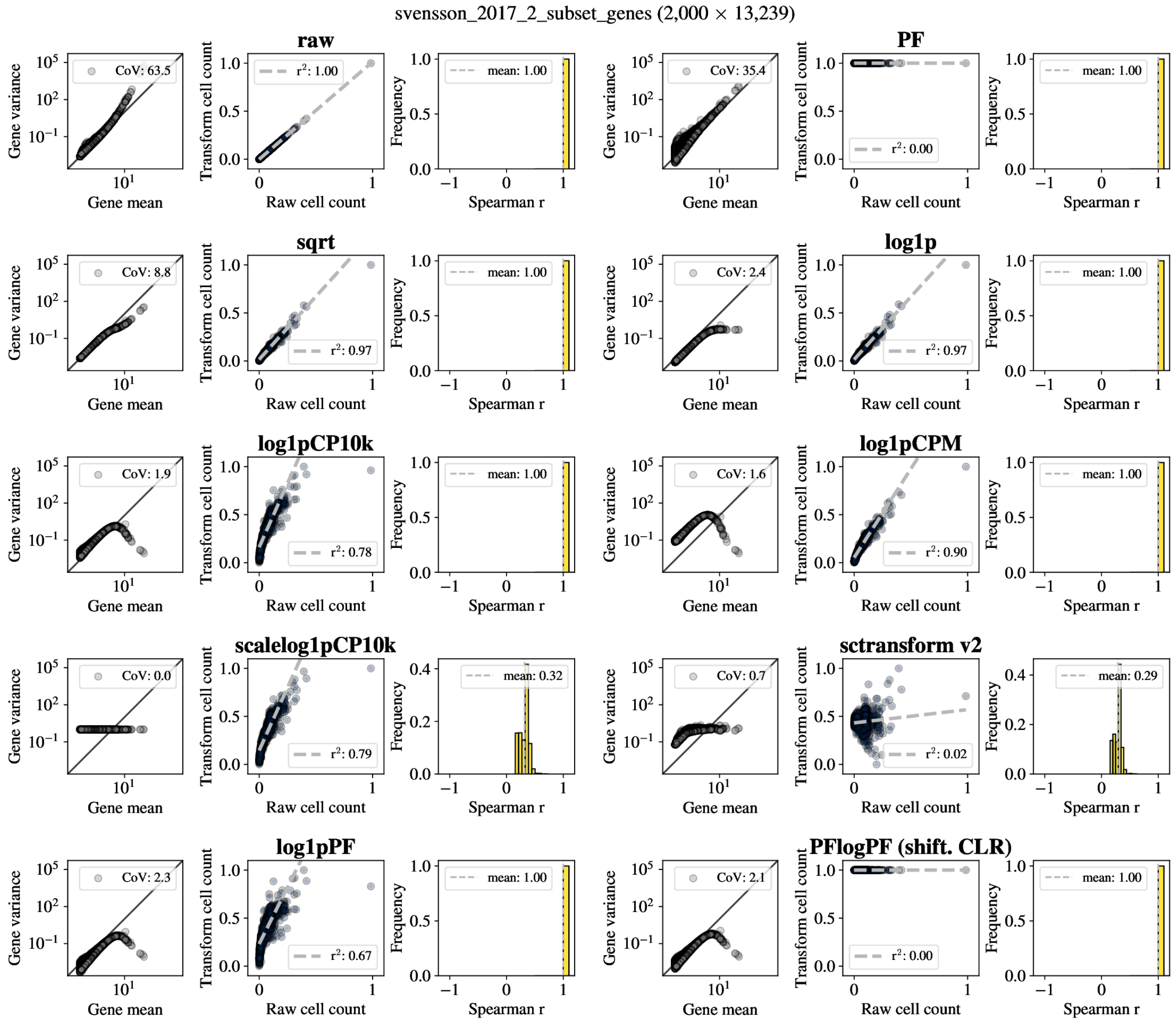


Supplementary Figure 1.525. **svensson_2017_2**. Per-method comparison of variance stabilization (left), depth normalization (middle), and monotonicity (right) for each normalization on dataset svensson_2017_2.

| zheng_2017 | |
|---|---|
| cells | 1,015 |
| genes | 92 |
| nonzero entries | 49,299 |
| density | 0.528 |
| total counts | 10,692,555 |
| mean counts / cell | 10,535 |
| mean counts / gene | 116,223 |
| min counts / cell | 1,913 |
| max counts / cell | 16,579 |
| overdispersion | 0.0416 |

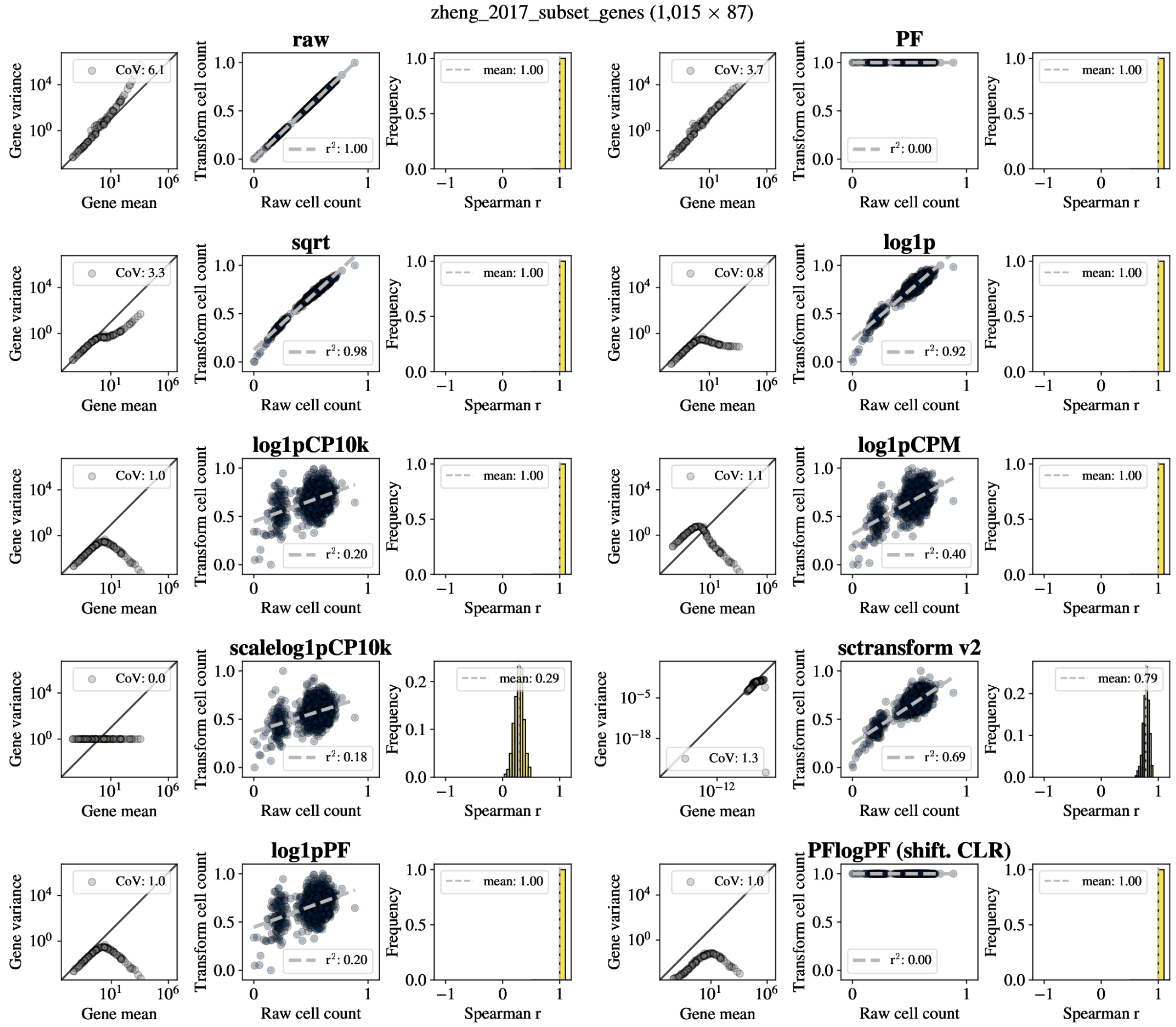


Supplementary Figure 1.526. **zheng_2017**. Per-method comparison of variance stabilization (left), depth normalization (middle), and monotonicity (right) for each normalization on dataset zheng_2017.
